# Choice of assembly software has a critical impact on virome characterisation

**DOI:** 10.1101/479105

**Authors:** Thomas D.S. Sutton, Adam G. Clooney, Feargal J. Ryan, R. Paul Ross, Colin Hill

## Abstract

**Background:** The viral component of microbial communities play a vital role in driving bacterial diversity, facilitating nutrient turnover and shaping community composition. Despite their importance, the vast majority of viral sequences are poorly annotated and share little or no homology to reference databases. As a result, investigation of the viral metagenome (virome) relies heavily on *de novo* assembly of short sequencing reads to recover compositional and functional information. Metagenomic assembly is particularly challenging for virome data, often resulting in fragmented assemblies and poor recovery of viral community members. Despite the essential role of assembly in virome analysis and difficulties posed by these data, current assembly comparisons have been limited to subsections of virome studies or bacterial datasets.

**Design:** This study presents the most comprehensive virome assembly comparison to date, featuring 16 metagenomic assembly approaches which have featured in human virome studies. Assemblers were assessed using four independent virome datasets, namely; simulated reads, two mock communities, viromes spiked with a known phage and human gut viromes.

**Results:** Assembly performance varied significantly across all test datasets, with SPAdes (meta) performing consistently well. Performance of MIRA and VICUNA varied, highlighting the importance of using a range of datasets when comparing assembly programs. It was also found that while some assemblers addressed the challenges of virome data better than others, all assemblers had limitations. Low read coverage and genomic repeats resulted in assemblies with poor genome recovery, high degrees of fragmentation and low accuracy contigs across all assemblers. These limitations must be considered when setting thresholds for downstream analysis and when drawing conclusions from virome data.

## Background

The rapid evolution of metagenomics and high throughput sequencing technologies has revolutionised the study of microbial communities, giving new insights into the role and identity of the uncultivated microbes which account for the majority of metagenomic sequences (Solden, Lloyd et al. 2016). However, the majority of microbial sequencing efforts have focused on the characterisation of prokaryotic microbes. Viral metagenomes (viromes) are dominated by novel sequences, often with up to 90% of sequences sharing little to no homology to reference databases (Aggarwala, Liang et al. 2017). Bacteriophage, the most abundant members of viral communities, play a key role in the shaping the composition of microbial communities and facilitate horizontal gene transfer (Paul 2008). Viromes have been shown to play a role in global geochemical cycles (Breitbart 2011) and have been studied in varied ecosystems including the ocean (Hurwitz and Sullivan 2013). Viromes of the human body are of particular interest, where they have been linked to disease status (Norman, Handley et al. 2015), maintaining human health (Manrique, Bolduc et al. 2016) and shaping the gut microbiome in early life (Lim, Zhou et al. 2015, McCann, Ryan et al. 2018). Due to the predominance of uncharacterised viral sequences “viral dark matter”; (Roux, Hallam et al. 2015), and the lack of a universal marker gene, virome studies rely on database independent analysis methods and depend heavily on *de novo* assembly to resolve viral genomes from metagenomic sequencing reads.

Metagenomic assemblers typically use de Bruijn graph (DBG) approaches to address the complexity and size of metagenomic datasets in an accurate and efficient manner. Microbial metagenomes pose significant challenges to DBG assembly when compared to single genome assemblies often complicating the DBG and leading to fragmentation and/or misassembly (Olson, Treangen et al. 2017). These challenges include uneven sequencing coverage of organisms within the metagenome, the presence of conserved regions across different species, repeat regions within genomes and the introduction of false *k-*mers by both closely related genomes at differing abundances and sequencing errors at high read coverage. This hampers the use of coverage statistics to resolve repeat regions between and within genomes (Olson, Treangen et al. 2017).

A wide array of metagenomic assembly programs have been employed, each addressing aspects of metagenomic challenges to varying degrees. However, many of these programs have been designed and optimised for bacterial metagenomes, which share many assembly challenges of viromes but to a lesser degree. Virome data is characterised by high proportions of repeat regions within viral genomes (Minot, Grunberg et al. 2012), hypervariable genomic regions associated with host interaction (Warwick-Dugdale, Solonenko et al. 2018) and high mutation rates which lead to increased metagenomic complexity and strain variation (Roux, Emerson et al. 2017). Low DNA yields also limit read coverage and often require a multiple displacement amplification (MDA) step which has been shown to preferentially amplify small single stranded DNA viruses (Kim and Bae 2011). Extremes in read coverage caused by MDA bias and dominant viral taxa such as crAssphage, which can make up large proportions of human gut viromes (Dutilh, Cassman et al. 2014), sequester sequencing resources and result in insufficient coverage of low abundance viruses. These challenges result in fragmented virome assemblies (García-López, Vázquez-Castellanos et al. 2015), limiting their use in downstream analysis. Despite benchmarks of bacterial metagenomes having highlighted failings and benefits of particular assembly programs, many poorly performing assemblers have featured in virome studies (Foulongne, Sauvage et al. 2012, Hannigan, Meisel et al. 2015, Guo, Hua et al. 2017).

Accurate comparison of metagenomic assemblers is complicated by the unknown composition of metagenomic datasets and the limited applicability of general assembly statistics such as N50 (Deng, Naccache et al. 2015, Vollmers, Wiegand et al. 2017). To address this, the accuracy and efficacy of metagenomic assembly programs is often evaluated using simulated datasets and mock communities of known composition. Although these simulated datasets are undergoing constant improvements (Sczyrba, Hofmann et al. 2017, Fritz, Hofmann et al. 2018), they have focused primarily on bacterial metagenomes and remain limited in their ability to accurately replicate the challenges of true metagenomes. While some virome-specific assembly benchmarks have been performed, many have been limited to a small number of assemblers, 454 data or subsections of virome studies and have exclusively used simulated data (Aguirre de Cárcer, Angly et al. 2014, Smits, Bodewes et al. 2014, Vázquez-Castellanos, García-López et al. 2014, García-López, Vázquez-Castellanos et al. 2015, Hesse, van Heusden et al. 2017, Roux, Emerson et al. 2017).

Here we expand upon previous studies and present a detailed investigation of assembly software for virome analysis which compares all those previously used in human virome studies to date, as well as other popular or more recently published assemblers (Table 1). We compare assembly efficacy and accuracy using simulated viromes, mock viral communities and human gut viromes spiked with a known exogenous bacteriophage. Furthermore we confirm these findings using human virome data from published datasets and assess computational parameters such as runtime and RAM usage. We also investigate in detail the impact of sequencing coverage and genomic repeats on assembly performance and highlight important considerations for future virome studies. Together these data comprise most comprehensive virome assembly benchmark to date.

**Table 1:**
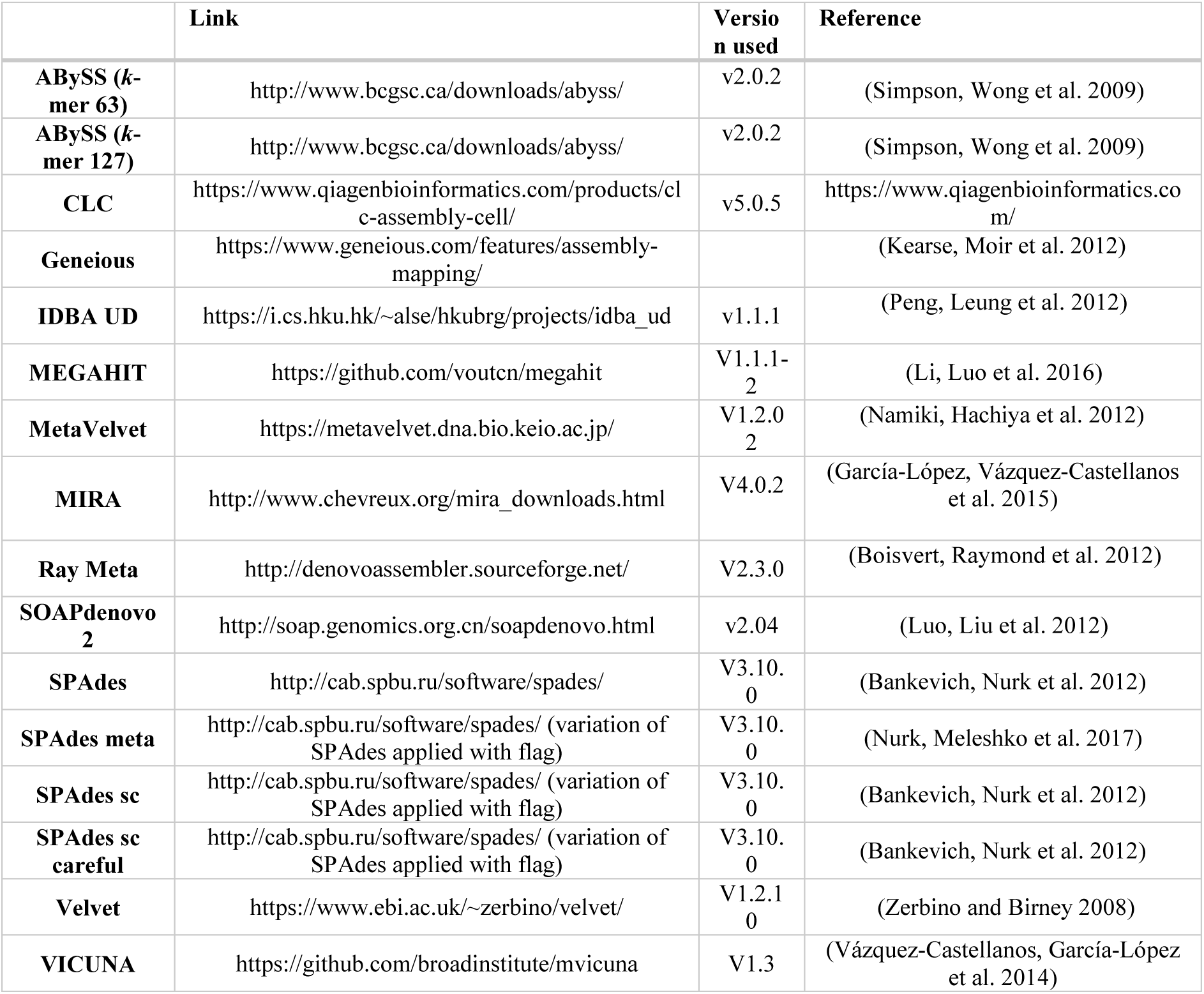
A list of assemblers used in this study

|  | Link | Version used | Reference |
| --- | --- | --- | --- |
| **ABYSS (k-mer 63)** | http://www.bcgsc.ca/downloads/abyss/ | v2.0.2 | (Simpson, Wong et al. 2009) |
| **ABYSS (k-mer 127)** | http://www.bcgsc.ca/downloads/abyss/ | v2.0.2 | (Simpson, Wong et al. 2009) |
| **CLC** | https://www.qiagenbioinformatics.com/products/clc-assembly-cell/ | v5.0.5 | https://www.qiagenbioinformatics.com/ |
| **Geneious** | https://www.geneious.com/features/assembly-mapping/ |  | (Kearse, Moir et al. 2012) |
| **IDBA UD** | https://i.cs.hku.hk/~alse/hkubrg/projects/idba_ud | v1.1.1 | (Peng, Leung et al. 2012) |
| **MEGAHIT** | https://github.com/voutcn/megahit | V1.1.1-2 | (Li, Luo et al. 2016) |
| **MetaVelvet** | https://metavelvet.dna.bio.keio.ac.jp/ | V1.2.0<br>2 | (Namiki, Hachiya et al. 2012) |
| **MIRA** | http://www.chevreux.org/mira_downloads.html | V4.0.2 | (García-López, Vázquez-Castellanos et al. 2015) |
| **Ray Meta** | http://denovoassembler.sourceforge.net/ | V2.3.0 | (Boisvert, Raymond et al. 2012) |
| **SOAPdenovo 2** | http://soap.genomics.org.cn/soapdenovo.html | v2.04 | (Luo, Liu et al. 2012) |
| **SPAdes** | http://cab.spbu.ru/software/spades/ | V3.10.<br>0 | (Bankevich, Nurk et al. 2012) |
| **SPAdes meta** | http://cab.spbu.ru/software/spades/ (variation of SPAdes applied with flag) | V3.10.<br>0 | (Nurk, Meleshko et al. 2017) |
| **SPAdes sc** | http://cab.spbu.ru/software/spades/ (variation of SPAdes applied with flag) | V3.10.<br>0 | (Bankevich, Nurk et al. 2012) |
| **SPAdes sc careful** | http://cab.spbu.ru/software/spades/ (variation of SPAdes applied with flag) | V3.10.<br>0 | (Bankevich, Nurk et al. 2012) |
| **Velvet** | https://www.ebi.ac.uk/~zerbino/velvet/ | V1.2.1<br>0 | (Zerbino and Birney 2008) |
| **VICUNA** | https://github.com/broadinstitute/mvicuna | V1.3 | (Vázquez-Castellanos, García-López et al. 2014) |

## Results

### Simulated virome dataset

The ability to accurately recover each of the 572 members of a published simulated community (Hesse, van Heusden et al. 2017) and the degree of fragmentation, was assessed by aligning the resulting contigs from each assembler to the reference genomes (Fig. 1). MetaVelvet was not included in this analysis as it failed to reach completion after seven days. Approximately half of the genomes in the community featured an average recovered genome fraction less than 75% and exhibited higher degrees of fragmentation (>10 contigs per genome on average) across all assemblers. For 87 of the 572 genomes there was an average recovered genome fraction of less than 20% across all assemblers (the low recovered genome fraction of VICUNA was excluded as an outlier). Of these genomes, 84 were present at low abundance (lowest 40% of all abundances normalised to genome length). The remaining three genomes were present at higher normalised abundances (50 – 80^th^ percentile) but featured the some of the highest proportions of genomic repeats (70^th^-90^th^ percentile).

**Figure 1:**
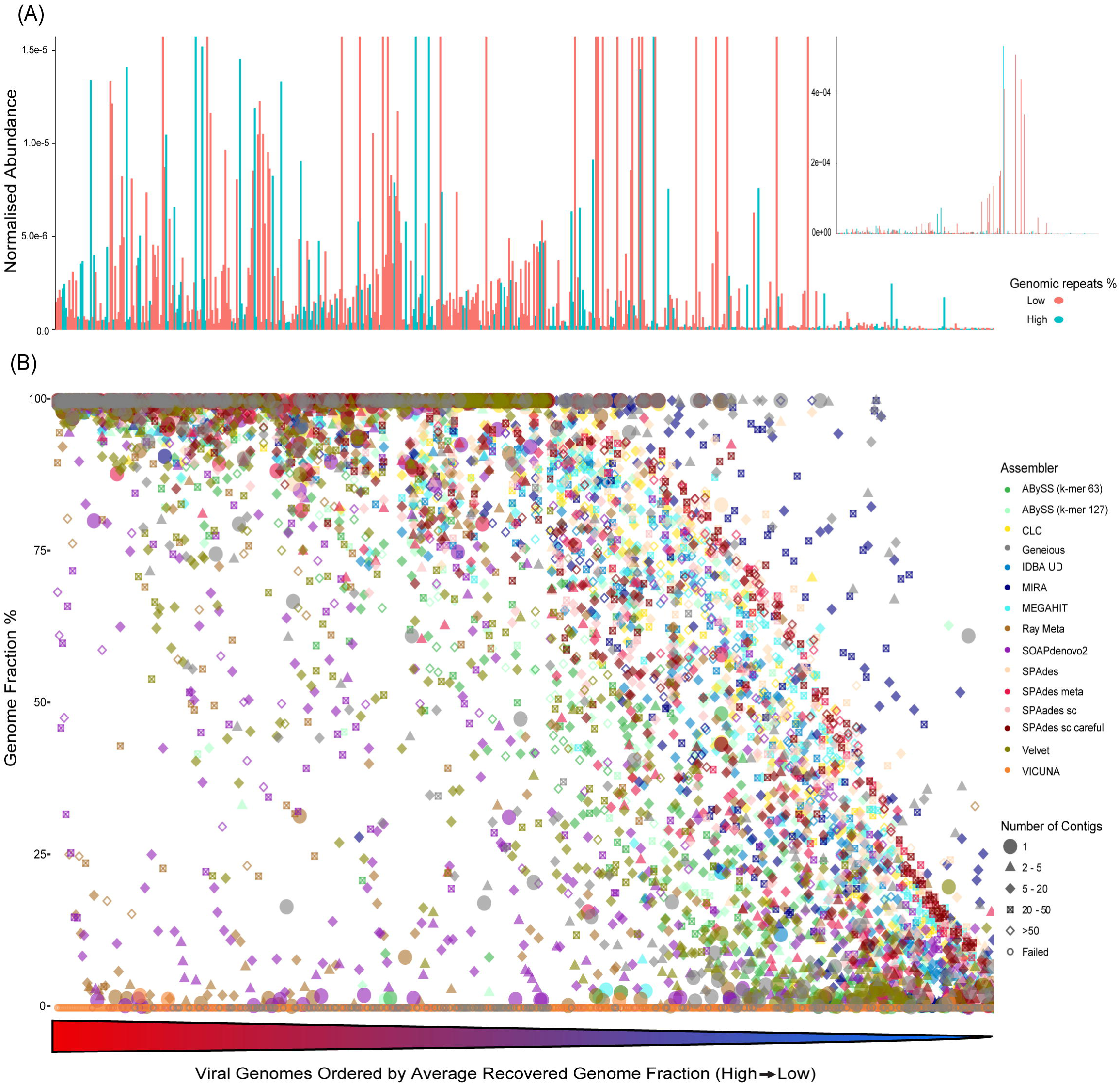
Relationship between percentage of each genome recovered (genome fraction), the number of contigs required for each combination of genome and assembler and the abundance and proportion of repeats for each genome. (A and B) Genomes are ordered by their average genome fraction across all assemblers from high to low along the x-axis. (A main) Relative abundance, normalized by genome length is plotted along y-axis with upper limit of 0.75% and colour of bars determined by proportion of repeat regions in each genome. Blue bars represent genomes with a high proportion of genomic repeats (4^th^ quartile of all genomes), red represents all other genomes below this quartile. (A insert) Expanded view of (A) without an upper limit of y value. (B) Percentage genome recovered is plotted along the y axis. Points are coloured by assembler with shape of the point is denoting number of contigs generated by each assembler for each genome.

Normalised genome abundance within the community had a strong positive correlation with recovered genome fraction across all assemblers (Supplementary Table 1, Additional file 5) and was verified using a linear model (Supplementary Table 2, Additional file 5), with the exception of SOAPdenovo2, which was negative. Normalised abundance also correlated negatively with the degree of fragmentation (number of contigs) across all assemblers except Velvet which was positively correlated and Geneious which was not significant (Supplementary Table1, Additional file 5). None of the genomes in the lower 30^th^ percentile of normalised abundance featured an average recovered genome fraction greater than 75%, further exemplifying the impact of low sequencing coverage. However high abundance did not consistently improve genome recovery and of the 172 genomes in the top 30% of normalised abundance, 20 featured an average genome fraction below 50%. The distance of the log transformed (due to extremes in values) normalised abundances from the mean was negatively correlated with recovered genome fraction across all assemblers (correlation coefficient: - 0.42, p-value < 2.2e^−16^). Of 171 genomes in the 40^th^ – 60^th^ percentile of normalised abundance 29 featured an average genome fraction below 50%. This indicates factors other than abundance may hamper genome recovery. MIRA and Geneious both recovered a greater fraction of low abundance genomes with fewer contigs than other assemblers. However, MIRA assemblies of 13 of the most abundant genomes in the community (highest 10%) exhibited the highest degree of fragmentation in the study, generating between 401 and 2983 contigs per genome.

The proportion of inverted repeats, palindromic repeats, tandem repeats and a total proportion of genomic repeats was calculated for each genome. The total percentage of repeat regions predicted in each genome was positively correlated with the degree of fragmentation observed in each assembly across all assemblers with the exception of Ray Meta (Supplementary Table 3, Additional file 5), and negatively correlated with recovered genome fraction across all assemblers except ABySS (*k*-mer 63/127), Geneious, and SOAPdenovo2. When this relationship between repeat regions and the recovered genome fraction was assessed using a linear model, correlations were significant for CLC, MIRA, Ray Meta, Velvet, and all parameters of SPAdes (Supplementary Table 2, Additional file 5). Both the proportion of repeat regions in a genome and the relative abundance of that genome contribute to the variation in recovered genome fraction, though each explain a separate aspect of this variation. No interaction was found between these two metrics.

VICUNA, Ray Meta, SOAPdenovo2, Geneious, ABySS (both *k*-mer sizes) and Velvet recovered under 50% of the total genome fraction (all genomes in the community). VICUNA produced just four contigs in total with high levels of mismatches (174 per 100kb on average) which could possibly linked to the format of the artificial reads as this was not observed in real sequencing data. The five assemblers which recovered the highest genome fraction overall were SPAdes (default), MEGAHIT, SPAdes (single cell), SPAdes (single cell + careful) and CLC. All assemblers achieving a minimum average genome fraction of 50% were subjected to a ranking system (Supplementary Table 4, Additional file 5). To compare both recovery and fragmentation assemblers were ordered from best to worst based on genome recovery and number of aligned contigs. The average rank resulted in Spades (default) performing best, recovering 72.2% overall genome sequences with 8230 contigs. The remaining top five assemblers of this combined rank were SPAdes (meta) 68.2% with 7419 contigs, SPAdes (single cell) 68.9% with 9506 contigs, CLC 68.6% with 9152 contigs and MEGAHIT 69.6% with 10083 contigs. The number of assemblies which recovered greater than 90% of the target genome in one single contig was compared (Fig 2). SPAdes (default) performed best, recovering 210, SPAdes (meta), SPAdes (single cell + careful), CLC, and SPAdes (single cell) each recovered 179, 168, 162 and 160 genomes respectively.

**Figure 2:**
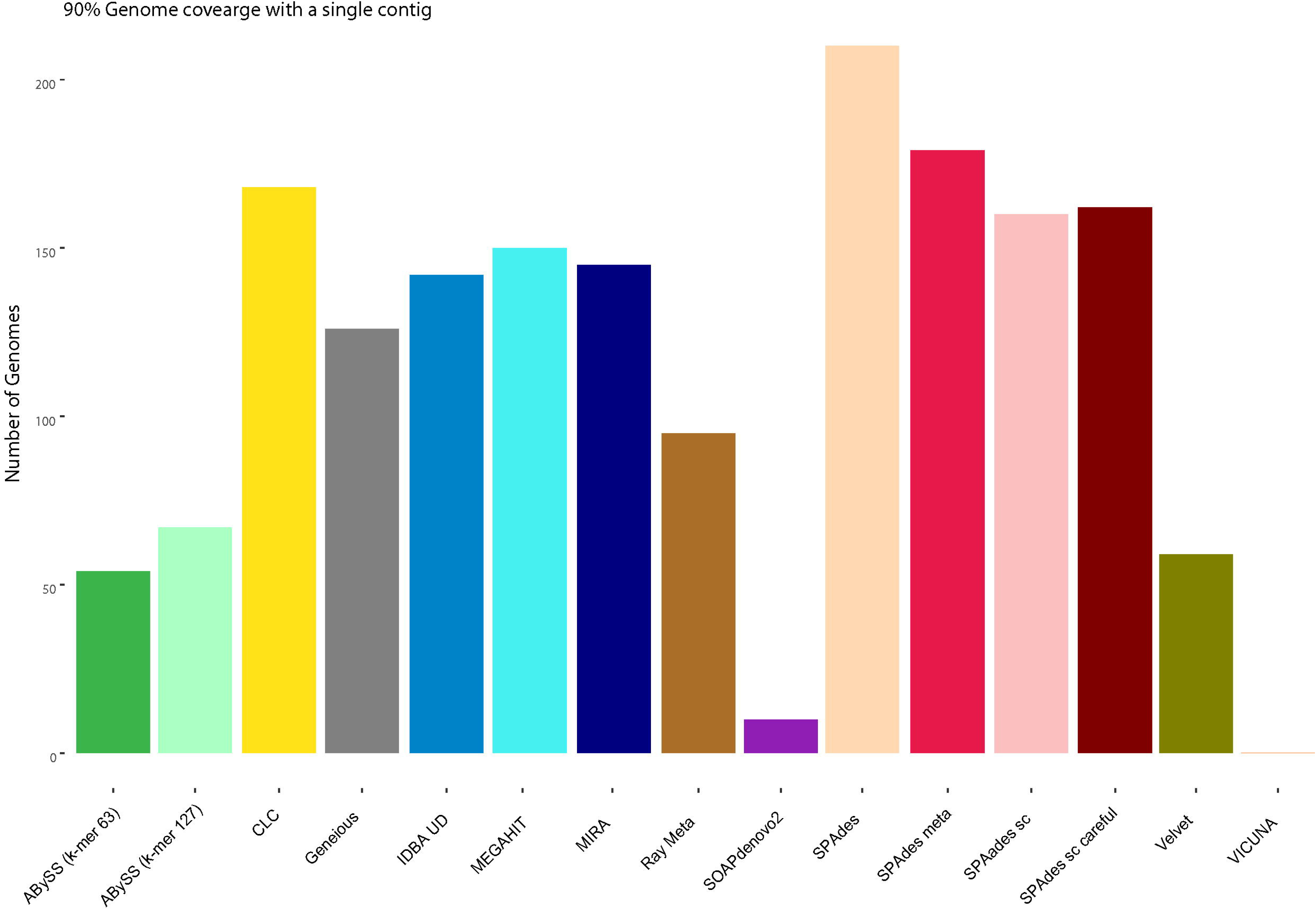
Number of contigs each assembler recovered to a minimum genome fraction of 90% in a single contig.

The accuracy of assemblies was assessed by calculating the average count of indels, mismatches, and misassemblies per 100kb across all genomes. These counts were normalised to the number of genomes each assembler recovered with a minimum genome fraction of 50%. These were ranked according to their performance in all three metrics (Supplementary Table 4, Additional file 5), with assemblies from Velvet having the lowest overall counts followed by ABySS, IDBA UD, MEGAHIT and Ray Meta. With the exception of Ray Meta and SOAPdenovo2, the number of mismatches per 100kb was negatively correlated with both genome abundance and recovered genome fraction across all assemblers (Supplementary Table 1, Additional file 5).

The rate of false positive (no alignment to reference genomes) and false negative (recovered genome fraction of 0%) contigs assembled allowed for the determination of sensitivity. A number of assemblers had a sensitivity greater than 97%, however each returned greater than 7,000 contigs, inferring a high degree of fragmentation (Table 2). MIRA assembled (partial or complete) 559 of the genomes with a false positive count of just four. However, this was achieved from more than 27,000 contigs. ABySS (both *k*-mer sizes), Geneious, Ray Meta and Velvet returned very few false positives but failed to detect many of the genomes present. SPAdes (meta) performed best with 558 of the 572 genomes detected and only five false positives resulting from 7419 contigs.

**Table 2:**
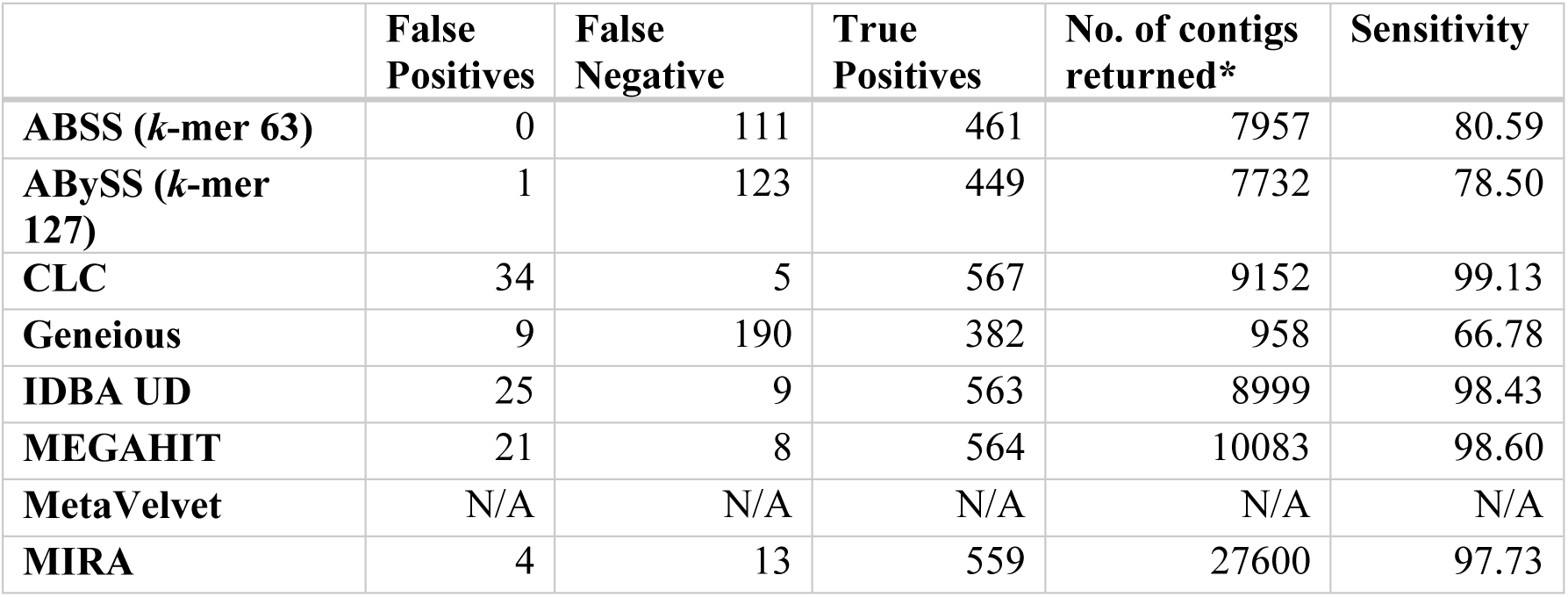
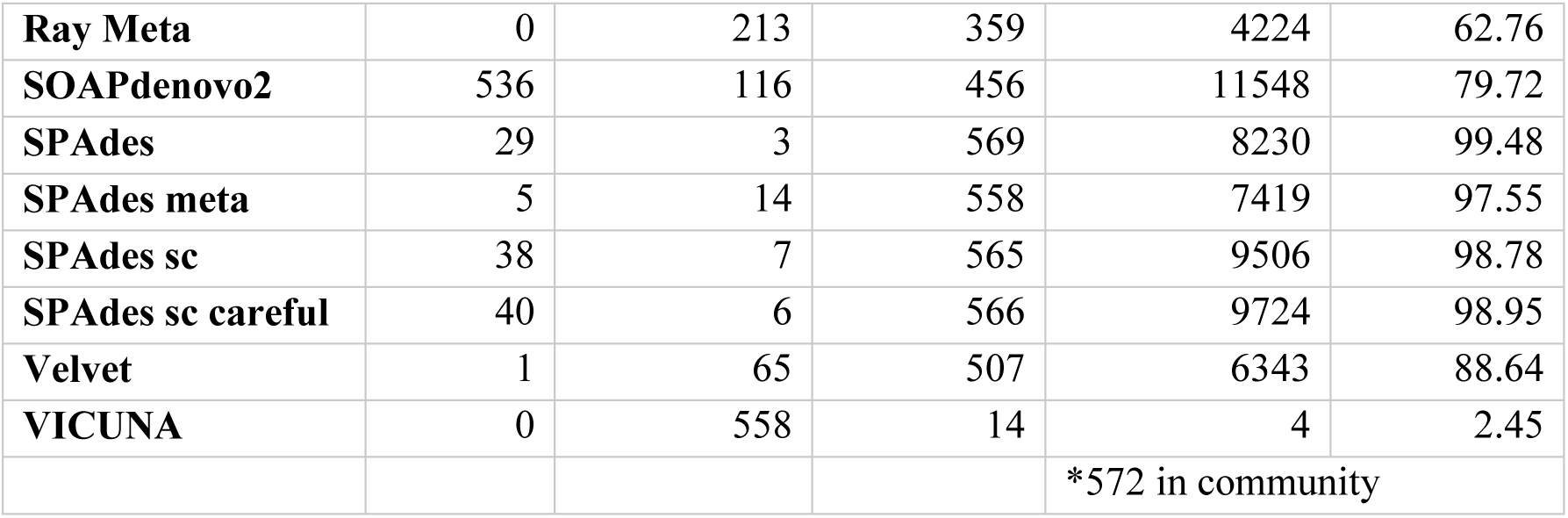
The number of false positive, false negative contigs generated by each assembler for the Simulated community, together with the sensitivity rates

### Mock community dataset

Two mock viral communities were used to investigate the impact of high and low abundance ssDNA viruses on assembly performance. Mock A (Table 3a) contained 12 viral genomes, 10 of which were at equal abundance (9.82% of the total community) and two ssDNA genomes (NC_001330 and NC_001422) at low abundance (0.92%). Analysis of this community showed that although some assemblers, namely CLC, Geneious, SPAdes (single cell) and VICUNA, detected all 12 genomes, this was at the expense of a large number of false positives (1143, 53, 1513 and 4969 respectively). Velvet and MetaVelvet generated no false positives, but failed to assemble three genomes, while ABySS (for both *k*-mers) generated a large number of false positives and failed to assemble four and six genomes, respectively. IDBA UD and Ray Meta outperformed the other assemblers with an equal number of contigs to genomes (12), followed by MEGAHIT, SPAdes (default) and SPAdes (meta) with 13, 14 and 14. Mock B (Table 3b) also contained 12 genomes but with a higher abundance of ssDNA genomes NC_001330 and NC_001422 (32.47%). VICUNA assemblies of Mock B improved upon those from Mock A as no false positives were generated, while the false positive rate in the MIRA assembler increased to 94 from none in Mock A. IDBA UD performed best followed by SPAdes (default), Ray Meta, MEGAHT and SPAdes (meta) based on sensitivity and number of contigs, while ABySS (both *k*-mer sizes) and SOAPdenovo2 had the lowest sensitivity. Despite being a relatively simple community consisting of 12 members, not all assemblers were able to recover all members (Supplementary Table 5-6, Additional file 5). A greater number of assemblers (six) failed to assemble all members of Mock B than Mock A (four). ABySS(*k*-mer 63), ABySS(*k*-mer 127), Velvet and MetaVelvet failed to assemble 6, 4, 3 and 3 genomes respectively, in Mock A and 6, 4, 1 and 1 genomes, respectively in Mock B. In addition, MIRA and SOAPdenovo2 failed to assemble 1 and 2 genomes respectively in Mock B.

**Table 3:**
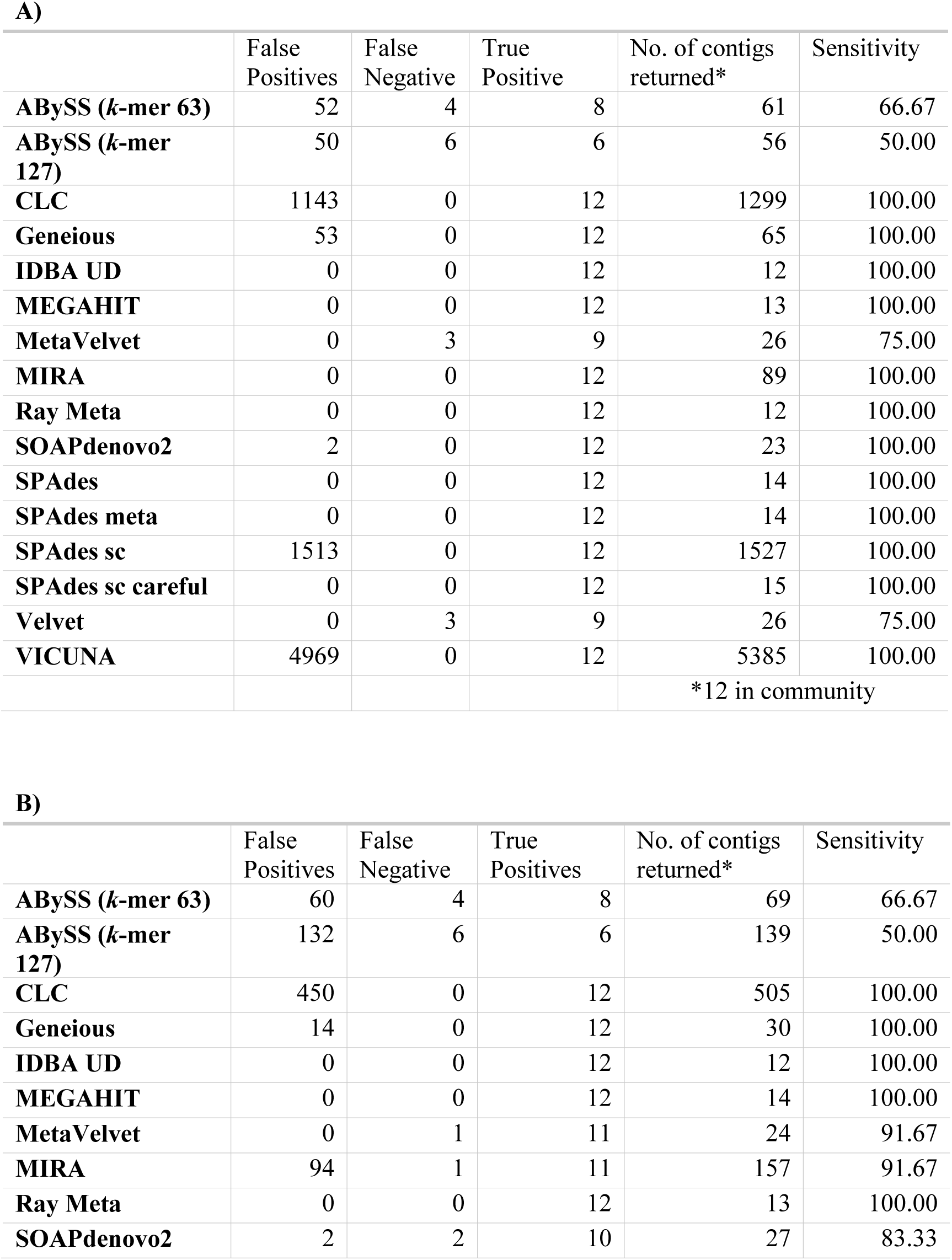
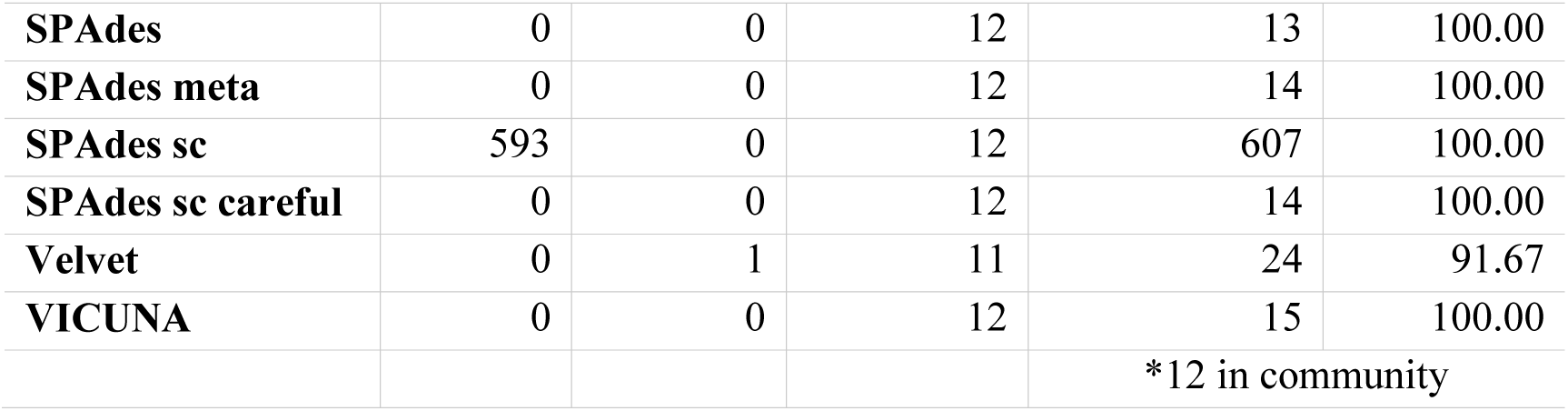
The number of false positive, false negative contigs generated by each assembler for a) Mock community A and b) Mock community B) along with the sensitivity rates for each

All but three VICUNA assemblies in Mock A exhibited a high level of fragmentation, generating 34.7 ± 35 (mean ± standard deviation) contigs per genome. Fragmentation was also seen in MIRA assemblies to a lesser degree with 7.4 ± 10 (mean ± standard deviation) contigs per genome on average. There was a high rate of fragmentation in CLC with one community member generating 144 contigs for genome KF302035. Average recovered genome fraction of 85.4 ± 6.4 % was skewed by ABySS (*k*-mer 63), ABySS (*k*-mer 127), Velvet, MetaVelvet, SOAPdenovo2, and VICUNA which recovered on average 49.5%, 66.6%, 73.8%, 73.8%, 29.7% and 76.6%, respectively. All other assemblers recovered over 99% of each genome in the community (Supplementary Figure 1, Additional file 6).

Closer inspection of the two ssDNA genomes present at lower relative abundance highlighted significant differences in the average number of indels across all assemblies of the NC_001330 and NC_001422 genomes versus other members of the community (p-value = 0.037). These genomes exhibited an average of 41.7 ± 18.5 and 9.4 ± 20.4 indels per 100kb, while all other genomes featured an average of 7.8 ± 18.9 indels per 100kb. The low abundant ssDNA genomes NC_001330 and NC_001422 also featured the highest average mismatches per 100kb at 148.7 ± 3 and 302.5 ± 10.7, respectively (Supplementary Figure 1, Additional file 6).

The degree of fragmentation observed by VICUNA and MIRA in Mock B was lower than in Mock A with a mean of 1.3 ± 0.89 and 5.3 ± 7.7 contigs per genome, respectively. CLC fragmented genome KF302035 in Mock B (44 contigs), but to a lesser degree than Mock A (144 contigs). MEGAHIT, which recovered at least 98% of all genomes in Mock A, also recovered over 98% of all genomes in Mock B except for the ssDNA genome NC_001422, of which 56.5% was recovered in two contigs. The majority of assemblies exhibited 147.9 ± 0 and 297 ± 1 mismatches per 100kb for NC_001330 and NC_001422 (high abundance ssDNA), respectively, identical values to those measured in Mock Velvet and MetaVelvet were exceptions with 184.2 and 860.2 for genome NC_001422 and NC_001330. A similar pattern of high values across a narrow range was also observed with the number of indels, with 49.3 to 32.9 present in all assemblies NC_001330. Genome NC_001422 featured 18.57 indels across all SPAdes assemblies (all parameters) and 860.2 across both Velvet and Metavelvet assemblies. All other assemblers which successfully recovered this genome did not feature any indels (Supplementary Figure 1, Additional file 6).

### Q33

Five assemblers failed to generate contigs which met alignment thresholds and were subsequently excluded from further analysis - namely ABySS (*k*-mer 63), ABySS (*k*-mer 127), SOAPdenovo2, Velvet and MetaVelvet. All remaining assemblers recovered over 90% of the spiked Q33 genome with the exception of MIRA (8.5%). Six assemblers recovered over 99% of the Q33 genome in a single contig - SPAdes (meta) 99.74%, MEGAHIT (99.6%), VICUNA (99.6%), Ray meta (99.6%), CLC (99.5%) and Geneious (99.1) (Fig. 3). However, only MEGAHIT assembled the Q33 genome with a contig equal in length to the genome itself. SPAdes (meta) and CLC generated assemblies shorter than the reference genome by 86 and 141 bases. VICUNA (723), Geneious (1765), and Ray Meta (9884) each generated assemblies longer than the reference genome. SPAdes (default) SPAdes (single cell), IDBA UD and SPAdes (single cell + careful) each assembled Q33 in 2, 3, 4, 5 and 5 contigs, respectively. Ray Meta and VICUNA assemblies had the lowest number of mismatches and indels per 100kb, however Ray Meta exhibited the highest rate of misassemblies (2 relocations, 1 inversion). All assemblers featured a minimum of one local misassembly with the exception of SPAdes (meta) did not feature any. The six best assemblies of the Q33 genome and the genome itself are syntenic (although occasionally on the reverse strand) and the start and end point were not conserved (Fig. 3).

**Figure 3:**
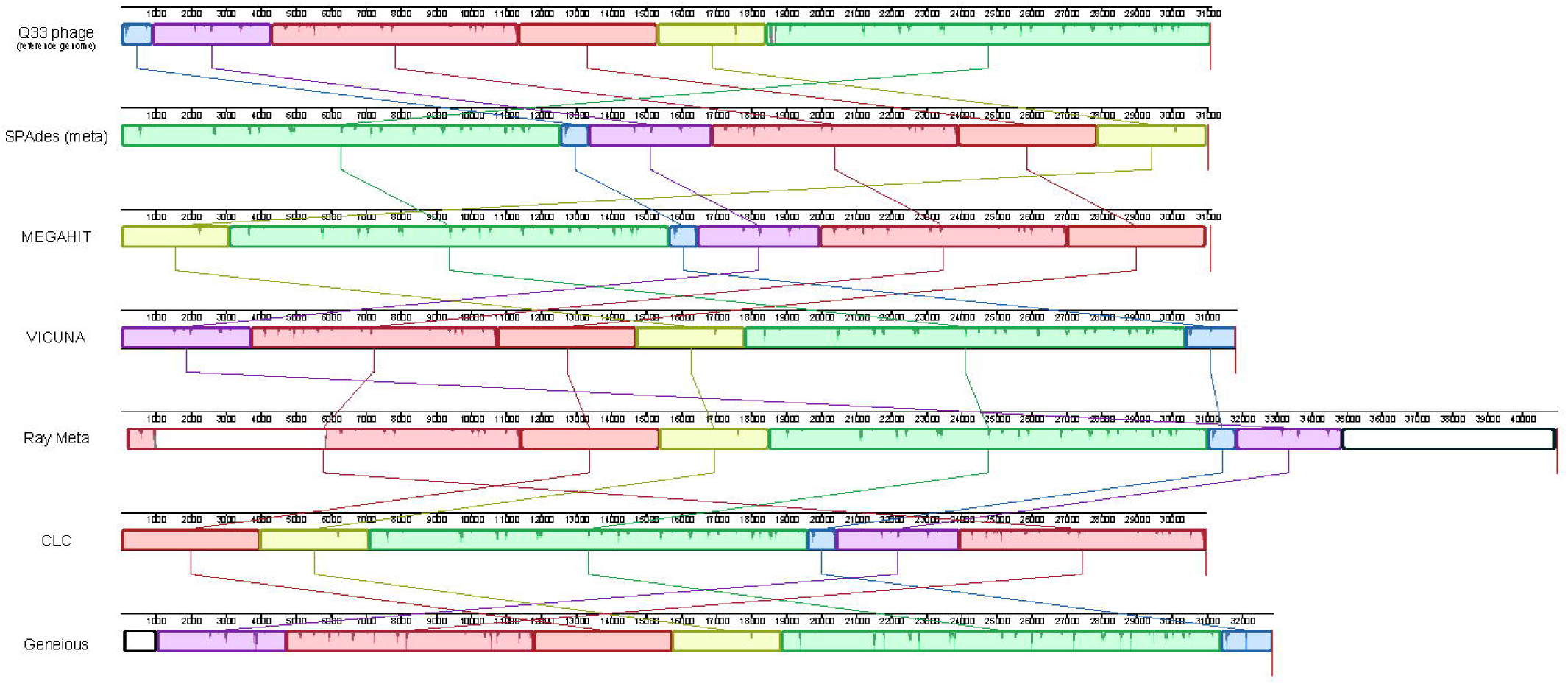
Mauve output of the Q33 reference genome (top) along with of the six assemblers which recovered >99% of the genome with a single contig. Assembly regions outside of locally collinear blocks which do not share homology to the reference genome are highlighted by a black outline. Reverse complement of assemblies in the opposite orientation to the reference were plotted for visualisation purposes (VICUNA, CLC, Geneious)

### Read depth analysis (Time and RAM)

Assemblers were compared for practicality by measuring the time to reach completion and maximum RAM usage via four published healthy human gut viromes (Manrique, Bolduc et al. 2016) and various sequencing depths. It must be noted that all assembly tasks were allocated five threads, however specifying the number of threads did not change the number of threads used by certain programs. MetaVelvet was not included in this analysis as it failed to reach completion after running for seven days. CLC and Geneious were performed on a desktop computer and therefore excluded from time and RAM analysis. Run time is dependent upon the number of reads and this is largely linear in scale with more reads leading to an increased assembly time (Fig. 4a). MIRA and Vicuna (Fig. 4a insert) were the slowest with MIRA taking over 15 times longer than the other software to assemble 3.5 million reads. SOAPdenovo2 had the shortest completion time followed by IBDA UD and Velvet. Most assemblers were consistent across samples (observed via error bars) with the exception of MIRA and Ray Meta. MIRA, Vicuna and Velvet (Fig. 4b insert) had the highest max RAM usage while the lowest was Ray Meta, IDBA UD and SPAdes (meta) (Fig. 4b). The majority of assemblers observed a linear scale pattern similar to that of run time.

**Figure 4:**
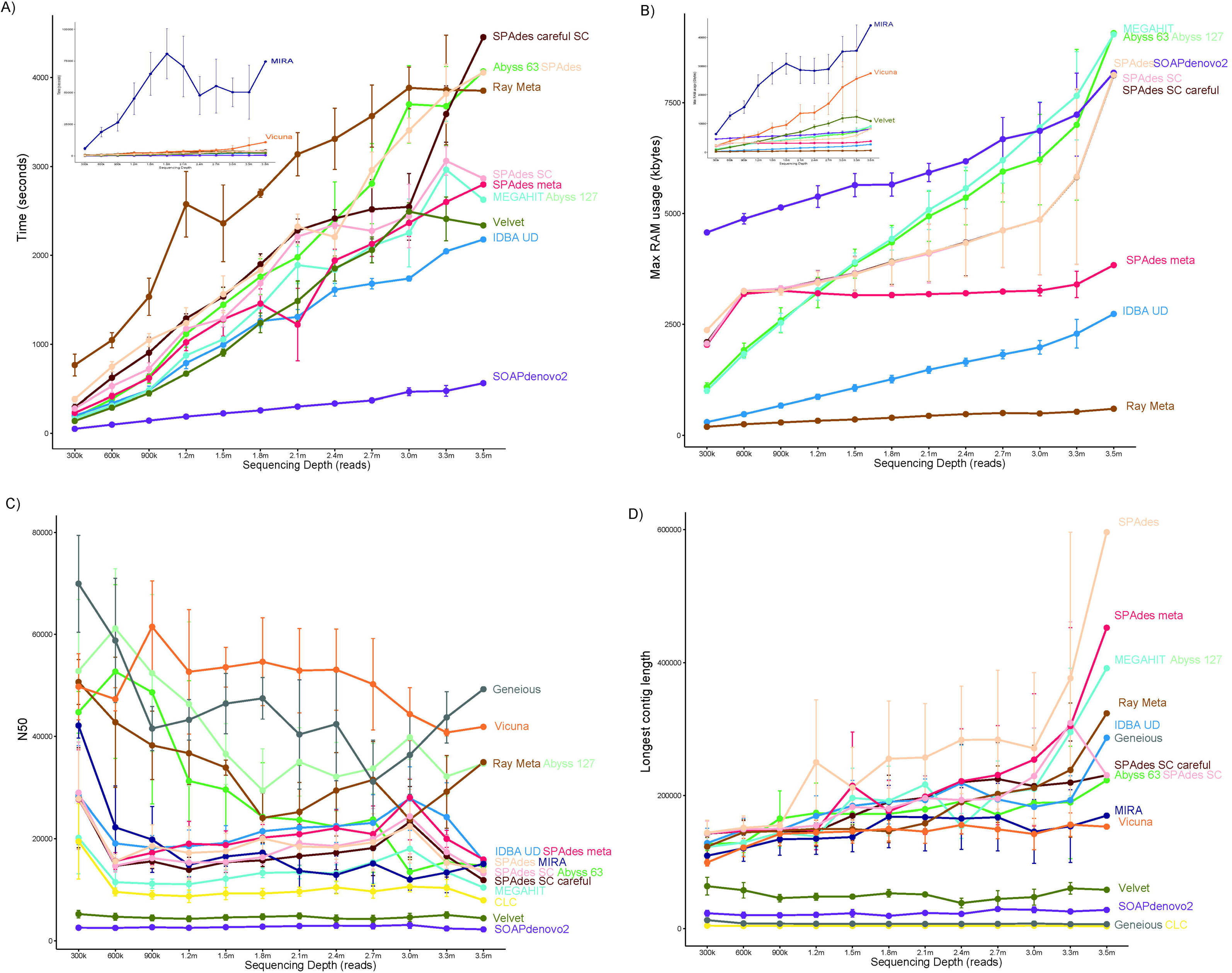
(A) Time, measured in seconds, for each assembly to reach completion successfully for each read subset, (B) the maximum RAM, measured in MB, used for each assembly for each read subset, (C) mean N50 length and (D) mean contig length for 4 samples for each assembly across the read subsets after filtering contigs less than 1000 bases. Points represent the mean time for the 4 samples while error bars are the standard error.

### Read depth analysis N50 and Longest contig length

For both the N50 (Fig. 4c) and the longest contig length (Fig. 4d), there was a large amount of variation between samples for the majority of assemblers. The longest contig length showed a large increase at the final sequencing depth. Particular assemblers, namely SPAdes (default), SPAdes (meta), MEGAHIT and ABySS (*k*-mer 127), produced longer contigs as the sequence depth was increased.

## Discussion

Many bacterial metagenomic assembly comparisons have highlighted that the choice of assembler has a significant impact on downstream analysis and the accuracy of the reconstructed metagenome (Mavromatis, Ivanova et al. 2007, Lindgreen, Adair et al. 2016, Greenwald, Klitgord et al. 2017, Vollmers, Wiegand et al. 2017). We have found this also to be true for viral metagenomes, where accurate and complete assembly are of particular importance given the lack of viral representation in reference databases. Virome studies depend heavily on the assembly step and possess many features which are challenging to successful assembly. In this study we compared the performance of those assemblers used to date in human viral metagenomics studies using datasets of known and unknown composition and varying complexity. These included a Q33 spiked virome, mock virome communities, a simulated virome and the “Healthy human gut phageome” (Manrique, Bolduc et al. 2016). Each dataset provided unique attributes allowing for comparison of assembly performance on a number of levels. The combination of artificial and real viromes used in this study allows for the comparison of various aspects of assembly performance across a range of datasets rather than depending on simulated viromes alone, as is commonly carried out in assembly comparisons (Mavromatis, Ivanova et al. 2007, Fritz, Hofmann et al. 2018).

The Simulated dataset featured 572 viral genomes at various relative abundances as published by Vázquez-Castellanos and colleagues (Vázquez-Castellanos, García-López et al. 2014). Fragmented assemblies of individual genomes within microbial communities hamper downstream analysis and limit the conclusions which can be drawn from metagenomic data such as taxonomic and functional profiles (Florea, Souvorov et al. 2011). Consequently, the percentage genome recovery and degree of fragmentation was assessed across each assembler, with SPAdes (default, meta and single cell) each performing well. VICUNA performed very poorly, recovering only four contigs with high numbers of mismatches and misassemblies, despite having performed well with other datasets and being designed to address challenges of heterogeneous viral populations (Yang, Charlebois et al. 2012). This failure may reflect the computational challenges relating to the format of the simulated reads, as benchmarks carried out within the VICUNA study itself only include actual sequencing reads (Yang, Charlebois et al. 2012). However, similar poor performance has been previously observed in virome assembly comparison using VICUNA and 454 reads (Vázquez-Castellanos, García-López et al. 2014). For those assemblers which could recover greater than 90% of the reference genome in a single contig, SPAdes (default) outperformed SPAdes (meta). This may be explained by a lack of strain variants in the dataset and the fact that SPAdes (meta) was optimised to combine strain variants of each species to form consensus sequences.

A subset of genomes were poorly recovered (<20% genome fraction) by nearly all assemblers. This observation indicates that there are challenging aspects of viral genomes and metagenomes which cannot be overcome with current assembly strategies. The strong positive correlations between the relative abundance and genome fraction suggest that a low abundance threshold applies to virome assembly, below which assemblies will consist of small fractions of the viral genome, and in most cases be highly fragmented. This detrimental impact of low coverage has been well established in previous assembly comparison studies (García-López, Vázquez-Castellanos et al. 2015, Roux, Emerson et al. 2017, Fritz, Hofmann et al. 2018). Highly abundant genomes also caused similar recovery and fragmentation issues across all assemblers, which is of particular importance due to the prevalence of extremely high abundance genomes in viral data (crAssphage, certain ssDNA viruses). As both abundance extremes are common in virome data, their impact must be considered when designing virome studies (i.e. sequencing depth). As relative abundance alone did not fully explain the variation in genome fraction recovered, the role of genomic repeats (a well-established assembly challenge (Acuña-Amador, Primot et al. 2018) was also investigated. However, genomic repeats could explain the variation in genome fraction recovered to a lesser degree than relative abundance, suggesting other factors contribute to poor genome recovery.

Compositional differences between final assemblies and viromes themselves must be taken into account when drawing conclusions about virome composition and setting parameters for downstream analysis. Challenges such as genomic content and strain variation are not currently addressed in human virome assembly strategies and impact the reconstruction of certain members of a virome. Hybrid sequencing, which uses both long and short reads to resolve genomic regions associated with poor assembly (Warwick-Dugdale, Solonenko et al. 2018) is a promising new technology which could address current virome assembly challenges. Extraction methods which may reduce the bias introduced by MDA steps include using Swift Biosciences 1S Plus kit (Roux, Solonenko et al. 2016) and/or increasing overall sequencing depth to improve coverage of lowly abundant viral genomes (in conjunction with an assembler which is less sensitive to high coverage sequences).

Performance of some assemblers in this study was hampered by high coverage datasets (namely overlap consensus assemblers). VICUNA assemblies exhibited the highest degree of fragmentation of all assemblers with Mock A, despite having resolved both high abundance ssDNA genomes of Mock B to a single contig. MIRA also exhibited a high degree of fragmentation with high abundance genomes in both simulated and mock datasets. However, MIRA was least affected by low abundance reads, recovering a greater genome fraction of low abundance genomes than other assemblers with fewer contigs. Assembly challenges of high coverage sequences in viromes may potentially be addressed by sub-setting reads similar to the assembly approach used by SLICEMBLER (Mirebrahim, Close et al. 2015).

Multi-assembler approaches such as the use of Geneious to generate consensus sequences from separate assemblers have been developed (Koren, Treangen et al. 2014, Schürch, Schipper et al. 2014, Deng, Naccache et al. 2015) but have not been included in human virome studies using short reads. MIRA assemblies of the Q33 genome and some low abundance genomes in the Simulated dataset were improved using Geneious, resolving greater genome fractions with fewer contigs (despite Geneious recovering a lower genome fraction of the Simulated dataset overall). It is possible that using these approaches could address issues facing each assembler, i.e. combine the assemblies of SPAdes (meta) which performs well across all 4 datasets but struggles to recover low abundant genomes, with MIRA assemblies which are less affected by low abundance but has difficulty resolving genomes of higher abundance. Comparison of multi-assembler approaches and combinations of various assemblers was not within the scope of this study, but should not be ruled out as a potential method of improving virome assembly in cases where composition could be assessed and obvious assembly challenges were known to be present.

Across all analysis methods in this study, SPAdes (meta) performed consistently well and would be our recommendation. It performed best in the Simulated data based on false positives, true positives and false negatives, best assembled the Q33 genome (recovery, fragmentation, misassemblies and genome size) and performed well with both mock communities in recovering all members accurately in one or two contigs. SPAdes (meta) RAM usage was low and did not increase to the same degree as other assemblers with increasing sequencing depth. This recommendation is in agreement with previous comparisons (Vollmers, Wiegand et al. 2017) which also suggested using SPAdes (meta) due to its ability to accurately assemble members of bacterial metagenomes. SPAdes (meta) is less able to accurately reconstruct micro-diversity as it generates a consensus assembly of “strain–contigs” in a metagenome, which means it is better equipped to address the high mutation rates observed in virome data (Nurk, Meleshko et al. 2017). This recommendation is also concurrent with a previous study (Roux, Emerson et al. 2017) which found IDBA UD, MEGAHIT and SPAdes (meta) to perform equally well when assessed using 14 simulated viromes. However, we found that SPAdes (meta) outperformed IDBA UD and MEGAHIT in the Q33 spiked dataset, RAM usage in relation to increasing sequencing depth, and in its ability to recover members of the Simulated virome in a single contig. This recommendation contradicts two previous assembly comparisons which found CLC (Hesse, van Heusden et al. 2017) and Velvet (White, Wang et al. 2017) to be best suited to virome data. However, SPAdes (meta) was not included in either study. Though SPAdes (meta) was out performed by MIRA in the assembly of low abundance genomes in the Simulated dataset, MIRA has limited application to large datasets. MEGAHIT also performed well across all datasets performing well in relation to recovery, fragmentation and accuracy, but encountered some recovery issues in mock datasets and minor accuracy issues with the Q33 genome.

The higher levels of accuracy (low mismatch indel and misassembly counts) of assemblers which performed poorly in other metrics namely (velvet and ABySS (*k*-mer 63), highlights the trade-off between accuracy and contiguity observed in previous assembly studies (Gritsenko, Nijkamp et al. 2012, Lin and Liao 2013). However, both IDBA-UD and MEGAHIT performed well for accuracy, genome recovery and fragmentation. These assemblers may be worth considering if strain level detail is of particular importance. The mock A and B datasets were used to assess the impact of amplification bias on assembly performance. All ssDNA assemblies featured an equal minimum number of mismatches across both Mock A and B. This may be caused by challenges in the genomes themselves hampering accurate assembly, but is more likely to reflect strain variation between genome sequence featured in the original publication and the genome of the phage used in the community itself.

The Q33 spiked virome consisted of pooled reads from three healthy human faecal samples, each of which having been spiked with 10^7^ PFU ml^−1^ of lactococcal phage Q33 prior to virome extraction. This allowed for assembly comparison of one abundant member of a challenging viral community. Despite the high relative abundance of the Q33 genome, only 6 assemblers could recover over 90% of the genome in a single contig, of these SPAdes (meta) and MEGAHIT reconstructed the Q33 genome accurately without the introduction foreign or chimeric DNA. It was also noted that the genome synteny was conserved across these six assemblies. This may reflect circularization of the linear Q33 genome during DNA extraction as the presence of cos sites has been previously predicted (Mahony, Martel et al. 2013).

The longest contigs of each assembler were only detected at the highest sequencing depths and varied across assemblers, which may indicate that high coverage is necessary to recover the largest viral genomes in a community. However, it is also possible that these long contigs may reflect misassemblies and duplication events caused by read errors at high sequencing depths which must be considered when analysing high coverage data. At almost all sequencing depths Geneious, Vicuna, Ray Meta and ABySS (k-mer 127) exhibited the highest N50 values, despite performing poorly in other metrics. This further highlights the limitation of using N50 alone as a metric of metagenomic assembly (Vollmers, Wiegand et al. 2017).

A further important consideration when performing any metagenomic assembly is practicality; size of dataset, computational resources, bioinformatic resources, and how much hands-on time is required from the end user. Both CLC and Geneious are available as a GUI (albeit requiring a licence fee) which widens their audience to researchers with limited command-line experience (CLC can also be run using the windows command line). However, this limits their practicality for large scale virome studies as they are limited to the computational power of desktop computers and are not suited to the assembly of large numbers of samples. Despite the limitations of computational power, CLC performed well in all datasets in terms of genome recovery and fragmentation. Of the freely available open source assemblers, MIRA and VICUNA are the least efficient in terms of RAM usage and assembly time, reflecting limitations of the overlap consensus approach to assembly. This limits their applicability to large virome datasets, and further increases the time required to carry out the Geneious assembly approach which requires the output of both assemblers. Despite the long runtime, VICUNA did not adhere to the number of cores specified, instead using all available cores. All other assemblers had a similar time requirements (with the exception of SOAPdenovo2 which performed poorly across all datasets). Of the assemblers which consistently performed well in terms of accuracy, genome fraction recovered and fragmentation, SPAdes (meta) was most efficient in terms of RAM usage, which did not increase to the same degree as other assemblers with increasing sequencing depth. MIRA stood out in terms of impracticality by generating by far the largest intermediate files of any assembler, requiring several gigabytes of storage space for intermediate files.

The combination of results from four datasets facilitates accurate comparison of assemblers as the limitations of each individual dataset vary. Application of Phi29 MDA to amplify virome DNA to sufficient quantities for sequencing can introduce significant bias and skew the original composition of the virome, making quantitative viromics difficult (Kim and Bae 2011, Roux, Solonenko et al. 2016). As a result, it is likely that true diversity of viral metagenomes is not being accurately captured using current virome extraction methods. However, as these procedures move away from steps known to introduce bias, greater diversity will be observed. In this sense, the level of complexity of the Q33 dataset, which pooled three independent human viromes, provides a useful testbed for metagenomic assemblers in future virome studies as extraction methods improve. Additionally, Q33 was not present in the viromes prior to spiking, assemblers were not challenged by the presence of native strain variations of Q33 genome. In this study, assemblers were compared without individual optimisation to the specific dataset. Feasibility dictates that, this “straight out of the box” approach to assembly is used by almost all metagenomic assembly comparisons. Additionally, as the true composition of metagenomes is unknown, any impact of parameter optimisation must be estimated from general assembly statistics such as N50 and longest contig which have been highlighted to be of limited usefulness (Aguirre de Cárcer, Angly et al. 2014, Vollmers, Wiegand et al. 2017).

## Conclusions

Of all assembly programs used in human virome studies, SPAdes (meta) addressed the challenges of virome data most effectively. However, all assemblers have limitations and are hampered by aspects of virome data. Low read coverage and high genomic repeats lead to assemblies with low recovered genome fraction and a higher degree of fragmentation, with the assemblies themselves being less accurate. This pattern was seen across all assemblers used in this study.

As assembler choice has significant implications for virome composition and the conclusions which can be drawn from a dataset, assemblers which performed poorly in this study (i.e. low genome recovery or accuracy and high degree of fragmentation) highlight a potential untapped resource in the sequence data of previously conducted virome studies. It is highly likely that many viral sequences were poorly assembled and reanalysis using a more effective assembler may yield new insights. Design of future virome studies should carefully consider the impact of sequencing depth, as extremes in read coverage will prevent the assembly and detection of viral genomes at both high and low abundance.

## Methods

Each assembler with the exception of Geneious and CLC was run as per manual with default parameters (unless stated) using a Lenovo x3650 M5 server with an intel Xeon processor E5-2690 v3 and 512Gb RAM. Geneious assembly approach mirrored that used in (Manrique, Bolduc et al. 2016) by generating consensus sequences from the assemblies of both MIRA and Vicuna. CLC and Geneious were run on a 64-bit windows 10 computer with an i5-4690 CPU and 16 GB of RAM.

### Data sources

Sequencing reads from mock communities A and B featured in (Roux, Solonenko et al. 2016), Simulated Virome dataset featured in (Hesse, van Heusden et al. 2017), reads used to compare the impact of sequencing depth on time and RAM usage featured in (Manrique, Bolduc et al. 2016) and human viromes spiked with 10^7^ PFU of Lactococcal phage Q33 (Mahony, Martel et al. 2013) and originated from (Shkoporov, Ryan et al. 2018).

### Read Pre-processing

Raw read quality was assessed with FastQC v0.11.5 and sequencing adapters were removed with cutadapt v1.9.1 (Martin 2011) for the mock, Spiked and healthy gut virome data sets. Trimming and filtering was carried out with Trimmomatic v0.36 (Bolger, Lohse et al. 2014) using parameters specific to each dataset. A sliding window size of 4 with a minimum Phred score of 30 and a minimum length of 60bp was used with reads from both mock communities. The leading 15bp and trailing 60bp were removed from “Healthy human gut phageome” reads and a sliding window of 4bp with a minimum phred score of 20 was applied. The leading 10bp and trailing 100bp were removed from the Q33 spiked virome reads and a sliding window size of 4bp with a minimum Phred score of 30. Filtered reads were through a minimum length filter of 60bp.

### Analysis methods

Quality filtered reads from the Q33 spiked dataset consisted of 3 individual viromes which were pooled and subsequently assembled. Contigs were aligned to the published Q33 using Blastn with an e-value cut-off of 1e^−20^. Top hit alignments to the Q33 genome with a minimum alignment length of 800 bases and which shared 95% identity were included in further analysis using QUAST (v. 4.4) (Gurevich, Saveliev et al. 2013) with “--unique mapping” flag. Further comparison and visualisation of Q33 assemblies was carried out using Mauve (v. 20150226, build 10) (Darling, Mau et al. 2010).

Alignment and comparison of assemblies from mock and simulated data sets to reference genomes was carried using MetaQUAST (v. 4.4) (Mikheenko, Saveliev et al. 2015) with “--unique mapping” flag. Correlations were carried out using Spearman method and plots were generated using the package ggplot2 (v 3.0.0) package in R (v.3.4.3). These correlations were validated using a linear model in R base library. For data which was not normally distributed, log transformation was carried out.

Reads from the “healthy human gut phageome” were analysed to compare the overall assembler efficiency and the impact of sequencing depth. Reads were randomly subset in pairs (both the forward and reverse read of a pair were retained) to different depths using an in-house python script. Samples were subset in increments of 300,000 reads to their respective maximum depth (2.7, 3.5, 3 and 3.3 million reads). The shell script *time* script, location */usr/bin/time*, was utilised to measure the maximum RAM and length of time for each assembly to reach completion. All assemblers were run using 5 threads where possible with the exception of CLC, Geneious, Ray Meta, Velvet and Vicuna. Ray Meta and Velvet were run with 10, 1 thread(s) respectively. Ray Meta failed to run with 5 while Velvet ran with 1 core despite 5 being allocated. Vicuna was also allocated 5 threads however used upwards of 20. MetaVelvet was run, but after 7 days had failed to reach completion and was therefore removed from the subsequent analysis of these metrics. Contig statistics and filtering (contigs greater than 1kb retained) were performed using the assembly-stats script from the Pathogen Informatics group at the Wellcome Sanger Institute (https://github.com/sanger-pathogens/assembly-stats).

## Supporting information

## Abbreviations/Glossary

The following terms; Genome fraction, N50, number of contigs, misassemblies, local misassemblies, are defined by QUAST (Mikheenko, Saveliev et al. 2015)

Genome fraction: “is the total number of aligned bases in the reference, divided by the genome size. A base in the reference genome is counted as aligned if there is at least one contig with at least one alignment to this base. Contigs from repeat regions may map to multiple places, and thus may be counted multiple times in this quantity.”
N50: “is the contig length such that using longer or equal length contigs produces half (50%) of the bases of the assembly. Usually there is no value that produces exactly 50%, so the technical definition is the maximum length x such that using contigs of length at least x accounts for at least 50% of the total assembly length.”
Number of contigs: “is the total number of contigs in the assembly that have size greater than or equal to 0 bp.”
Misassemblies: “is the number of positions in the assembled contigs where the left flanking sequence aligns over 1 kbp away from the right flanking sequence on the reference (relocation) or they overlap on more than 1 kbp (relocation) or flanking sequences align on different strands (inversion) or different chromosomes (translocation).”
Local misassemblies: “A local misassembly has two or more distinct alignments covering the breakpoint, the gap between left and right flanking sequences is less than 1 kbp and the left and right flanking sequences both are on the same strand of the same chromosome of the reference genome.”

## Data sources

Sequencing reads from mock communities A and B: http://datacommons.cyverse.org/browse/iplant/home/shared/iVirus/DNA_Viromes_library_comparison.

Simulated virome reads: https://figshare.com/articles/Simulated_virome_datasest_for_assembly_and_annotation_tests/5151163.

Reads used to compare the impact of sequencing depth on time and RAM usage from the NCBI SRA; http://www.ncbi.nlm.nih.gov/sra under the accession numbers SAMN04415496 to SAMN04415499

Human viromes spiked with 10^7^ PFU of Lactococcal phage Q33 phage http://www.ncbi.nlm.nih.gov/sra under the accession numbers SRX3240741, SRX3240716, SRX3240715

## Declarations

**Ethics approval and consent to participate**

Not applicable

## Consent for publication

Not applicable

## Authors’ contributions

TDSS, AGC, FJR, PR, and CH conceived and designed experiments. TDSS, AGC, FJR carried out bioinformatics analysis and drafted the manuscript. All authors approve and contributed to the final manuscript.

## Acknowledgements

We thank the authors of all datasets used in this study for the availability of their data.

## Additional files

Additional file 1. (html) Simulated Virome MetaQUAST output.

Additional file 2. (html) Mock virome A MetaQUAST output.

Additional file 3. (html) Mock virome B MetaQUAST output.

Additional file 4. (html) Q33 spiked virome QUAST output.

Additional file 5. (xls)

S. Table 1: Spearman correlation values from the relationships of indel, mismatch and misassembly counts, recovered genome fraction, abundance and total proportion of genomic repeats within the Simulated virome. *GF – Recovered genome fraction

Additional file 5. (xls)

Supplementary Table 2: Linear modelling correlation values comparing recovered genome fraction, total proportion of genomic repeats and abundance for the Simulated virome.

Additional file 5. (xls)

Supplementary Table 3: Spearman correlation values from the relationships of inverted, tandem, palindromic and total repeats, abundance and the number of contigs generated by each assembler for the Simulated virome.

Additional file 5. (xls)

Supplementary Table 4: (A) Ranking table comparing recovered genome fraction and contig numbers for assemblers which recovered at least 50% of the total genome fraction. (B) Ranking table of indel, mismatch and misassembly counts per 100kb, normalised to the number of genomes recovered to at least 50%.

Additional file 5. (xls)

Supplementary Table 5: Number of aligned and unaligned contigs generated by each assembler for Mock Community A.

Additional file 5. (xls)

Supplementary Table 6: Number of aligned and unaligned contigs generated by each assembler for Mock Community B.

Additional file 6. Supplementary Figure 1. Analysis of recovered genome fraction and indel/mismatch counts for Mock communities A and B. Triangles represent N/A values for mismatches and indels caused by assembly failures.

