## Supplementary material for "Choice of assembly software has a critical impact on virome characterisation"

### QUAST

**Quality Assessment Tool for Genome Assemblies**
by CAB

Loading...

Aligned to
""

Combined reference
Estimated reference size:
 bp
|
 bp
|
 references
|
 fragments
|
 % G+C
|
 chromosomes
  
 reads
|
 mapped
|
 properly paired
  
 genes
|
 operons

Unfortunately, JavaScript in your
browser is disabled or is not supported.
We need JavaScript to build report and plots.

Worst
Median
Best

Show heatmap

|  |
| --- |
| Contigs are ordered from largest (contig #1) to smallest.  Contigs are broken into nonoverlapping 100 bp windows. Plot shows numbers of windows for each GC percentage. |

[{"assembliesWithNs":["ABySS\_127","ABySS\_63","Ray\_Meta","SOAPdenovo2","SPAdes","SPAdes\_meta","SPAdes\_sc","SPAdes\_sc\_careful","Velvet","Geneious"],"minContig":500,"report":[["Genome statistics",[{"values":["49.166","45.196","68.601","66.146","69.665","65.873","25.520","38.068","72.273","68.138","68.854","68.645","47.738","0.004","41.262"],"quality":"More is better","isMain":true,"metricName":"Genome fraction (%)"},{"values":["1.025","1.010","1.008","1.006","1.010","2.694","1.029","1.034","1.011","1.005","1.005","1.006","1.002","1.082","1.048"],"quality":"Less is better","isMain":true,"metricName":"Duplication ratio"},{"values":[228547,212531,245567,244933,257479,257331,134892,134904,260428,260153,257672,257672,206050,562,434153],"quality":"More is better","isMain":true,"metricName":"Largest alignment"},{"values":[23674881,21425692,32442333,31244550,33018109,83297496,12328551,17984472,34285840,32128868,32454082,32393887,22451711,2023,20277960],"quality":"More is better","isMain":true,"metricName":"Total aligned length"},{"values":[9253,7426,16476,15678,14625,4305,13868,1876,29967,29468,14329,14453,11310,491,37621],"quality":"More is better","isMain":false,"metricName":"NA50"},{"values":[2263,2002,2863,2596,2447,2202,1996,679,3734,4000,2501,2392,3073,457,15111],"quality":"More is better","isMain":false,"metricName":"NA75"},{"values":[417,451,357,336,358,3580,169,1861,260,253,352,346,412,3,143],"quality":"Less is better","isMain":false,"metricName":"LA50"},{"values":[1793,1938,1688,1812,2007,10604,911,6849,1268,1105,1957,1988,1429,4,346],"quality":"Less is better","isMain":false,"metricName":"LA75"}]],["Misassemblies",[{"values":[47,24,117,106,69,361,21,33,165,95,133,129,21,0,471],"quality":"Less is better","isMain":true,"metricName":"# misassemblies"},{"values":[17,6,26,21,14,104,14,6,49,20,29,30,8,0,87],"quality":"Less is better","isMain":false,"metricName":" # relocations"},{"values":[0,0,0,0,0,0,0,0,0,0,0,0,0,0,0],"quality":"Less is better","isMain":false,"metricName":" # translocations"},{"values":[1,0,3,1,2,6,0,1,3,1,3,3,1,0,4],"quality":"Less is better","isMain":false,"metricName":" # inversions"},{"values":[29,18,88,84,53,251,7,26,113,74,101,96,12,0,380],"quality":"Less is better","isMain":false,"metricName":" # interspecies translocations"},{"values":[37,22,113,106,67,273,19,31,151,73,128,124,20,0,243],"quality":"Less is better","isMain":false,"metricName":"# misassembled contigs"},{"values":[1185158,483198,442144,439028,368452,5922378,838732,91438,1666997,1236945,1450687,1436341,55036,0,9289166],"quality":"Less is better","isMain":true,"metricName":"Misassembled contigs length"},{"values":[2,0,11,13,7,9,2,201,22,14,14,16,0,0,39],"quality":"Less is better","isMain":false,"metricName":"# possibly misassembled contigs"},{"values":[4,0,12,13,7,15,4,408,24,23,15,16,0,0,69],"quality":"Less is better","isMain":false,"metricName":" # possible misassemblies"},{"values":[37,13,34,123,166,280,67,208,231,98,135,128,34,0,212],"quality":"Less is better","isMain":false,"metricName":"# local misassemblies"},{"values":[0,0,3,1,0,0,0,542,0,0,0,0,0,0,0],"quality":"Less is better","isMain":false,"metricName":"# unaligned mis. contigs"}]],["Unaligned",[{"values":[0,1,34,25,21,4,0,536,29,5,38,40,1,0,9],"quality":"Less is better","isMain":false,"metricName":"# fully unaligned contigs"},{"values":[0,825,32945,21695,18765,5250,0,590096,34376,4567,43665,45195,806,0,23824],"quality":"Less is better","isMain":false,"metricName":"Fully unaligned length"},{"values":[2,1,19,22,12,11,3,925,42,28,49,52,1,0,42],"quality":"Less is better","isMain":false,"metricName":"# partially unaligned contigs"},{"values":[1304,1537,18315,17620,11111,10076,40039,1881407,48334,34530,44069,48589,736,0,73178],"quality":"Less is better","isMain":false,"metricName":"Partially unaligned length"}]],["Mismatches",[{"values":[6759,5747,25043,19644,24709,37700,3440,6572,33197,36945,30302,24586,5382,19,33734],"quality":"Less is better","isMain":false,"metricName":"# mismatches"},{"values":[303,185,968,846,928,1806,242,5257,1329,1487,1291,1127,208,6,2731],"quality":"Less is better","isMain":false,"metricName":"# indels"},{"values":[2295,1159,3206,4734,4518,6687,3291,250676,6173,12931,14888,13986,2642,6,9240],"quality":"Less is better","isMain":false,"metricName":"Indels length"},{"values":["29.27","27.07","77.72","63.23","75.52","121.85","28.70","36.76","97.80","115.45","93.70","76.26","24.00","939.20","174.07"],"quality":"Less is better","isMain":true,"metricName":"# mismatches per 100 kbp"},{"values":["1.31","0.87","3.00","2.72","2.84","5.84","2.02","29.40","3.92","4.65","3.99","3.50","0.93","296.59","14.09"],"quality":"Less is better","isMain":true,"metricName":"# indels per 100 kbp"},{"values":[225,144,876,680,785,1587,154,45,1115,1145,909,776,125,6,2447],"quality":"Less is better","isMain":false,"metricName":" # indels (<= 5 bp)"},{"values":[78,41,92,166,143,219,88,5212,214,342,382,351,83,0,284],"quality":"Less is better","isMain":false,"metricName":" # indels (> 5 bp)"},{"values":[2087,10174,70,0,0,31371,38470,1202313,16835,19343,37748,37921,561,0,58221],"quality":"Less is better","isMain":false,"metricName":"# N's"},{"values":["8.81","47.47","0.22","0.00","0.00","37.63","310.92","5734.93","48.93","60.09","115.85","116.57","2.50","0.00","285.24"],"quality":"Less is better","isMain":true,"metricName":"# N's per 100 kbp"}]],["Statistics without reference",[{"values":[7957,7732,9152,8999,10083,27600,4224,11548,8230,7419,9506,9724,6343,4,958],"quality":"Equal","isMain":true,"metricName":"# contigs"},{"values":[370491,365479,13713,28327,133775,28198,33995,51732,14948,12769,18136,24689,37706,58,958],"quality":"Equal","isMain":false,"metricName":"# contigs (>= 0 bp)"},{"values":[3819,3722,4786,4927,5079,19809,1984,5026,4475,4126,5189,5164,3617,0,954],"quality":"Equal","isMain":false,"metricName":"# contigs (>= 1000 bp)"},{"values":[795,723,1003,902,942,2706,351,686,973,900,939,918,895,0,566],"quality":"Equal","isMain":false,"metricName":"# contigs (>= 5000 bp)"},{"values":[373,313,545,467,485,870,194,236,506,520,472,461,466,0,360],"quality":"Equal","isMain":false,"metricName":"# contigs (>= 10000 bp)"},{"values":[149,129,270,244,250,311,111,64,279,291,231,234,160,0,229],"quality":"Equal","isMain":false,"metricName":"# contigs (>= 25000 bp)"},{"values":[46,35,86,81,88,107,22,8,111,115,84,86,34,0,107],"quality":"Equal","isMain":false,"metricName":"# contigs (>= 50000 bp)"},{"values":[254220,212531,245567,244933,257479,257331,168755,143833,260428,260153,257672,257672,206050,575,493731],"quality":"More is better","isMain":true,"metricName":"Largest contig"},{"values":[23681087,21432785,32520957,31301066,33063967,83362307,12372995,20964737,34403483,32191509,32583190,32531234,22456974,2189,20411466],"quality":"More is better","isMain":true,"metricName":"Total length"},{"values":[93104463,59887345,34080536,35290817,65789049,83623020,17261783,29532819,36698861,34069084,35239336,36159364,27134710,15629,20411466],"quality":"More is better","isMain":false,"metricName":"Total length (>= 0 bp)"},{"values":[20800608,18620360,29438093,28401161,29537783,77880567,10823994,16446893,31728463,29803343,29481300,29259287,20558915,0,20407831],"quality":"More is better","isMain":true,"metricName":"Total length (>= 1000 bp)"},{"values":[14587695,12424905,21868488,20293423,21202095,38464194,7670699,8166787,24731787,23426530,21004194,20839174,14756571,0,19256390],"quality":"More is better","isMain":false,"metricName":"Total length (>= 5000 bp)"},{"values":[11616775,9592773,18720616,17300896,18060662,26066031,6604341,5081779,21481760,20792976,17792033,17713386,11805214,0,17812757],"quality":"More is better","isMain":true,"metricName":"Total length (>= 10000 bp)"},{"values":[8108365,6795598,14580506,13894703,14553640,18199348,5281051,2486355,18006271,17361350,14192549,14305769,7061500,0,15726390],"quality":"More is better","isMain":false,"metricName":"Total length (>= 25000 bp)"},{"values":[4335830,3342681,7723720,7755862,8368074,10612165,1923336,594195,11519040,10848548,8526821,8602205,2580830,0,11104104],"quality":"More is better","isMain":true,"metricName":"Total length (>= 50000 bp)"},{"values":[9466,7528,16694,15712,14657,4365,14676,3018,31536,31099,14816,14847,11310,553,56961],"quality":"More is better","isMain":false,"metricName":"N50"},{"values":[2272,2012,2912,2634,2463,2215,2050,1125,3906,4156,2570,2460,3081,548,28523],"quality":"More is better","isMain":false,"metricName":"N75"},{"values":[397,444,354,334,356,3396,160,1291,251,246,345,340,412,2,91],"quality":"Less is better","isMain":false,"metricName":"L50"},{"values":[1752,1917,1664,1794,1985,10356,873,4343,1215,1057,1902,1935,1426,3,214],"quality":"Less is better","isMain":false,"metricName":"L75"}]],["Predicted genes",[]],["Similarity statistics",[{"values":[34,33,51,54,57,21,11,4,55,50,51,50,36,0,14],"quality":"Equal","isMain":false,"metricName":"# similar correct contigs"},{"values":[0,0,0,0,0,0,0,0,0,0,0,0,0,0,0],"quality":"Equal","isMain":false,"metricName":"# similar misassembled blocks"}]],["Reference statistics",[{"values":[46967092,46967092,46967092,46967092,46967092,46967092,46967092,46967092,46967092,46967092,46967092,46967092,46967092,46967092,46967092],"quality":"Equal","isMain":false,"metricName":"Reference length"},{"values":[572,572,572,572,572,572,572,572,572,572,572,572,572,572,572],"quality":"Equal","isMain":false,"metricName":"Reference fragments"}]]],"referenceName":"combined\_reference","date":"18 July 2018, Wednesday, 16:04:27","order":[0,1,2,3,4,5,6,7,8,9,10,11,12,13,14],"assembliesNames":["ABySS\_127","ABySS\_63","CLC","IDBA\_UD","MEGAHIT","MIRA","Ray\_Meta","SOAPdenovo2","SPAdes","SPAdes\_meta","SPAdes\_sc","SPAdes\_sc\_careful","Velvet","VICUNA","Geneious"]},{"assembliesWithNs":null,"minContig":500,"report":[["Genome statistics",[{"values":["98.731","97.587","98.149","98.343","98.558","99.118","80.547","66.126","98.892","98.500","98.558","98.549","96.347","99.194"],"quality":"More is better","isMain":true,"metricName":"Genome fraction (%)"},{"values":["1.000","1.000","1.000","1.000","1.001","1.000","1.023","1.062","1.000","1.000","1.001","1.001","1.000","1.000"],"quality":"Less is better","isMain":true,"metricName":"Duplication ratio"},{"values":[116413,124904,82194,116359,116438,129456,4525,7835,129161,125482,125772,125772,70038,129555],"quality":"More is better","isMain":true,"metricName":"Largest alignment"},{"values":[128978,127500,128163,128444,128849,129456,107580,87507,129161,128655,128761,128765,125867,129555],"quality":"More is better","isMain":true,"metricName":"Total aligned length"},{"values":[116413,124904,82194,116359,116438,129456,1135,4490,129161,127023,127297,127297,70038,129608],"quality":"More is better","isMain":false,"metricName":"NG50"},{"values":[116413,124904,34031,116359,116438,129456,672,897,129161,127023,127297,127297,45092,129608],"quality":"More is better","isMain":false,"metricName":"NG75"},{"values":[116413,124904,82194,116359,116438,129456,1266,3154,129161,125482,125772,125772,70038,129555],"quality":"More is better","isMain":false,"metricName":"NA50"},{"values":[116413,124904,34031,116359,116438,129456,955,897,129161,125482,125772,125772,45092,129555],"quality":"More is better","isMain":false,"metricName":"NA75"},{"values":[116413,124904,82194,116359,116438,129456,1135,1889,129161,125482,125772,125772,70038,129555],"quality":"More is better","isMain":true,"metricName":"NGA50"},{"values":[116413,124904,34031,116359,116438,129456,672,null,129161,125482,125772,125772,45092,129555],"quality":"More is better","isMain":false,"metricName":"NGA75"},{"values":[1,1,1,1,1,1,38,11,1,1,1,1,1,1],"quality":"Less is better","isMain":false,"metricName":"LG50"},{"values":[1,1,2,1,1,1,74,29,1,1,1,1,2,1],"quality":"Less is better","isMain":false,"metricName":"LG75"},{"values":[1,1,1,1,1,1,28,10,1,1,1,1,1,1],"quality":"Less is better","isMain":false,"metricName":"LA50"},{"values":[1,1,2,1,1,1,52,26,1,1,1,1,2,1],"quality":"Less is better","isMain":false,"metricName":"LA75"},{"values":[1,1,1,1,1,1,38,15,1,1,1,1,1,1],"quality":"Less is better","isMain":true,"metricName":"LGA50"},{"values":[1,1,2,1,1,1,74,null,1,1,1,1,2,1],"quality":"Less is better","isMain":false,"metricName":"LGA75"}]],["Misassemblies",[{"values":[0,0,0,0,0,0,0,0,0,1,1,1,0,0],"quality":"Less is better","isMain":true,"metricName":"# misassemblies"},{"values":[0,0,0,0,0,0,0,0,0,1,1,1,0,0],"quality":"Less is better","isMain":false,"metricName":" # relocations"},{"values":[0,0,0,0,0,0,0,0,0,0,0,0,0,0],"quality":"Less is better","isMain":false,"metricName":" # translocations"},{"values":[0,0,0,0,0,0,0,0,0,0,0,0,0,0],"quality":"Less is better","isMain":false,"metricName":" # inversions"},{"values":[0,0,0,0,0,0,0,0,0,1,1,1,0,0],"quality":"Less is better","isMain":false,"metricName":"# misassembled contigs"},{"values":[0,0,0,0,0,0,0,0,0,127023,127297,127297,0,0],"quality":"Less is better","isMain":true,"metricName":"Misassembled contigs length"},{"values":[0,0,1,0,0,0,0,2,0,1,1,1,0,0],"quality":"Less is better","isMain":false,"metricName":"# local misassemblies"},{"values":[0,0,0,0,0,0,0,0,0,0,0,0,0,0],"quality":"Less is better","isMain":false,"metricName":"# unaligned mis. contigs"}]],["Unaligned",[{"values":[0,0,0,0,0,0,0,0,0,0,0,0,0,0],"quality":"Less is better","isMain":false,"metricName":"# fully unaligned contigs"},{"values":[0,0,0,0,0,0,0,0,0,0,0,0,0,0],"quality":"Less is better","isMain":false,"metricName":"Fully unaligned length"},{"values":[0,0,0,0,0,0,0,6,0,0,0,0,0,0],"quality":"Less is better","isMain":false,"metricName":"# partially unaligned contigs"},{"values":[0,0,0,0,0,0,0,13499,0,0,0,0,0,0],"quality":"Less is better","isMain":false,"metricName":"Partially unaligned length"}]],["Mismatches",[{"values":[0,0,20,0,0,0,0,2,0,15,16,3,0,0],"quality":"Less is better","isMain":false,"metricName":"# mismatches"},{"values":[1,0,11,2,1,1,1,66,1,4,4,2,0,3],"quality":"Less is better","isMain":false,"metricName":"# indels"},{"values":[29,0,39,50,29,29,18,3117,29,41,27,50,0,50],"quality":"Less is better","isMain":false,"metricName":"Indels length"},{"values":["0.00","0.00","15.60","0.00","0.00","0.00","0.00","2.32","0.00","11.66","12.43","2.33","0.00","0.00"],"quality":"Less is better","isMain":true,"metricName":"# mismatches per 100 kbp"},{"values":["0.78","0.00","8.58","1.56","0.78","0.77","0.95","76.42","0.77","3.11","3.11","1.55","0.00","2.32"],"quality":"Less is better","isMain":true,"metricName":"# indels per 100 kbp"},{"values":[0,0,8,0,0,0,0,0,0,2,3,0,0,1],"quality":"Less is better","isMain":false,"metricName":" # indels (<= 5 bp)"},{"values":[1,0,3,2,1,1,1,66,1,2,1,2,0,2],"quality":"Less is better","isMain":false,"metricName":" # indels (> 5 bp)"},{"values":[0,0,0,0,0,0,0,9025,0,0,10,0,0,2],"quality":"Less is better","isMain":false,"metricName":"# N's"},{"values":["0.00","0.00","0.00","0.00","0.00","0.00","0.00","8574.82","0.00","0.00","7.76","0.00","0.00","1.54"],"quality":"Less is better","isMain":true,"metricName":"# N's per 100 kbp"}]],["Statistics without reference",[{"values":[3,4,5,4,4,1,90,39,1,2,2,2,5,1],"quality":"Equal","isMain":true,"metricName":"# contigs"},{"values":[3,2,5,3,3,1,47,25,1,2,2,2,5,1],"quality":"Equal","isMain":false,"metricName":"# contigs (>= 1000 bp)"},{"values":[2,1,3,2,2,1,0,7,1,1,1,1,3,1],"quality":"Equal","isMain":false,"metricName":"# contigs (>= 5000 bp)"},{"values":[1,1,2,1,1,1,0,0,1,1,1,1,2,1],"quality":"Equal","isMain":false,"metricName":"# contigs (>= 10000 bp)"},{"values":[1,1,2,1,1,1,0,0,1,1,1,1,2,1],"quality":"Equal","isMain":false,"metricName":"# contigs (>= 25000 bp)"},{"values":[1,1,1,1,1,1,0,0,1,1,1,1,1,1],"quality":"Equal","isMain":false,"metricName":"# contigs (>= 50000 bp)"},{"values":[116413,124904,82194,116359,116438,129456,4525,9504,129161,127023,127297,127297,70038,129608],"quality":"More is better","isMain":true,"metricName":"Largest contig"},{"values":[128978,127500,128163,128444,128849,129456,107628,105250,129161,128655,128791,128791,125867,129608],"quality":"More is better","isMain":true,"metricName":"Total length"},{"values":[128978,126431,128163,127848,128227,129456,75860,94813,129161,128655,128791,128791,125867,129608],"quality":"More is better","isMain":true,"metricName":"Total length (>= 1000 bp)"},{"values":[125955,124904,124529,125847,125063,129456,0,50268,129161,127023,127297,127297,123395,129608],"quality":"More is better","isMain":false,"metricName":"Total length (>= 5000 bp)"},{"values":[116413,124904,116225,116359,116438,129456,0,0,129161,127023,127297,127297,115130,129608],"quality":"More is better","isMain":true,"metricName":"Total length (>= 10000 bp)"},{"values":[116413,124904,116225,116359,116438,129456,0,0,129161,127023,127297,127297,115130,129608],"quality":"More is better","isMain":false,"metricName":"Total length (>= 25000 bp)"},{"values":[116413,124904,82194,116359,116438,129456,0,0,129161,127023,127297,127297,70038,129608],"quality":"More is better","isMain":true,"metricName":"Total length (>= 50000 bp)"},{"values":[116413,124904,82194,116359,116438,129456,1266,4691,129161,127023,127297,127297,70038,129608],"quality":"More is better","isMain":false,"metricName":"N50"},{"values":[116413,124904,34031,116359,116438,129456,955,2300,129161,127023,127297,127297,45092,129608],"quality":"More is better","isMain":false,"metricName":"N75"},{"values":[1,1,1,1,1,1,28,8,1,1,1,1,1,1],"quality":"Less is better","isMain":false,"metricName":"L50"},{"values":[1,1,2,1,1,1,52,15,1,1,1,1,2,1],"quality":"Less is better","isMain":false,"metricName":"L75"},{"values":["45.77","45.64","45.66","45.73","45.76","45.88","45.77","45.48","45.80","45.72","45.74","45.74","45.44","45.90"],"quality":"Equal","isMain":false,"metricName":"GC (%)"}]],["Predicted genes",[]],["Similarity statistics",[{"values":[0,0,0,0,0,0,0,0,0,0,0,0,0,0],"quality":"Equal","isMain":false,"metricName":"# similar correct contigs"},{"values":[0,0,0,0,0,0,0,0,0,0,0,0,0,0],"quality":"Equal","isMain":false,"metricName":"# similar misassembled blocks"}]],["Reference statistics",[{"values":[130608,130608,130608,130608,130608,130608,130608,130608,130608,130608,130608,130608,130608,130608],"quality":"Equal","isMain":false,"metricName":"Reference length"},{"values":[1,1,1,1,1,1,1,1,1,1,1,1,1,1],"quality":"Equal","isMain":false,"metricName":"Reference fragments"},{"values":["46.17","46.17","46.17","46.17","46.17","46.17","46.17","46.17","46.17","46.17","46.17","46.17","46.17","46.17"],"quality":"Equal","isMain":false,"metricName":"Reference GC (%)"}]]],"referenceName":"gi\_10140926\_ref\_NC","date":"18 July 2018, Wednesday, 16:04:50","order":[0,1,2,3,4,5,6,7,8,9,10,11,12,13],"assembliesNames":["ABySS\_127","ABySS\_63","CLC","IDBA\_UD","MEGAHIT","MIRA","Ray\_Meta","SOAPdenovo2","SPAdes","SPAdes\_meta","SPAdes\_sc","SPAdes\_sc\_careful","Velvet","Geneious"]},{"assembliesWithNs":null,"minContig":500,"report":[["Genome statistics",[{"values":["69.825","56.488","88.359","72.029","88.359","100.000","1.169","25.144","88.070","77.745","57.447","57.758","22.093","100.000"],"quality":"More is better","isMain":true,"metricName":"Genome fraction (%)"},{"values":["1.027","1.019","1.000","1.069","1.043","1.488","1.000","1.456","1.013","1.006","1.005","1.009","1.000","1.488"],"quality":"Less is better","isMain":true,"metricName":"Duplication ratio"},{"values":[7566,2623,11497,4437,6258,32865,519,1993,12908,10422,2774,2774,2035,38902],"quality":"More is better","isMain":true,"metricName":"Largest alignment"},{"values":[31823,25547,39224,34186,40887,66062,519,11160,39598,34675,25616,25864,9806,66035],"quality":"More is better","isMain":true,"metricName":"Total aligned length"},{"values":[2002,644,7271,1497,3370,44529,null,590,6082,4710,912,912,null,45094],"quality":"More is better","isMain":false,"metricName":"NG50"},{"values":[null,null,4396,794,1776,44529,null,null,4384,889,null,null,null,45094],"quality":"More is better","isMain":false,"metricName":"NG75"},{"values":[4618,1168,7271,1497,3370,27133,519,null,7568,9749,1289,1883,777,27133],"quality":"More is better","isMain":false,"metricName":"NA50"},{"values":[1746,928,5387,828,1776,null,519,null,5112,4099,1198,1198,627,null],"quality":"More is better","isMain":false,"metricName":"NA75"},{"values":[2002,644,7271,1237,3370,32865,null,null,6082,4657,912,912,null,38902],"quality":"More is better","isMain":true,"metricName":"NGA50"},{"values":[null,null,4396,588,1748,27133,null,null,4384,889,null,null,null,38902],"quality":"More is better","isMain":false,"metricName":"NGA75"},{"values":[5,18,3,7,5,1,null,17,3,3,14,13,null,1],"quality":"Less is better","isMain":false,"metricName":"LG50"},{"values":[null,null,5,17,9,1,null,null,5,7,null,null,null,1],"quality":"Less is better","isMain":false,"metricName":"LG75"},{"values":[3,8,3,7,5,2,1,null,2,2,6,6,5,2],"quality":"Less is better","isMain":false,"metricName":"LA50"},{"values":[6,14,4,16,9,null,1,null,4,4,11,10,8,null],"quality":"Less is better","isMain":false,"metricName":"LA75"},{"values":[5,18,3,10,5,1,null,null,3,3,14,13,null,1],"quality":"Less is better","isMain":true,"metricName":"LGA50"},{"values":[null,null,5,23,10,2,null,null,5,7,null,null,null,1],"quality":"Less is better","isMain":false,"metricName":"LGA75"}]],["Misassemblies",[{"values":[0,0,0,0,0,0,0,0,0,0,0,0,0,0],"quality":"Less is better","isMain":true,"metricName":"# misassemblies"},{"values":[0,0,0,0,0,0,0,0,0,0,0,0,0,0],"quality":"Less is better","isMain":false,"metricName":" # relocations"},{"values":[0,0,0,0,0,0,0,0,0,0,0,0,0,0],"quality":"Less is better","isMain":false,"metricName":" # translocations"},{"values":[0,0,0,0,0,0,0,0,0,0,0,0,0,0],"quality":"Less is better","isMain":false,"metricName":" # inversions"},{"values":[0,0,0,0,0,0,0,0,0,0,0,0,0,0],"quality":"Less is better","isMain":false,"metricName":"# misassembled contigs"},{"values":[0,0,0,0,0,0,0,0,0,0,0,0,0,0],"quality":"Less is better","isMain":true,"metricName":"Misassembled contigs length"},{"values":[0,0,0,0,0,4,0,0,0,0,0,0,0,4],"quality":"Less is better","isMain":false,"metricName":"# local misassemblies"},{"values":[0,0,0,0,0,0,0,8,0,0,0,0,0,0],"quality":"Less is better","isMain":false,"metricName":"# unaligned mis. contigs"}]],["Unaligned",[{"values":[0,0,0,0,0,0,0,0,0,0,0,0,0,0],"quality":"Less is better","isMain":false,"metricName":"# fully unaligned contigs"},{"values":[0,0,0,0,0,0,0,0,0,0,0,0,0,0],"quality":"Less is better","isMain":false,"metricName":"Fully unaligned length"},{"values":[0,0,0,1,1,2,0,7,0,0,0,0,0,2],"quality":"Less is better","isMain":false,"metricName":"# partially unaligned contigs"},{"values":[0,0,0,3839,1338,23588,0,8149,0,0,0,0,0,23588],"quality":"Less is better","isMain":false,"metricName":"Partially unaligned length"}]],["Mismatches",[{"values":[1,24,2,101,30,637,0,2,0,270,1,8,3,579],"quality":"Less is better","isMain":false,"metricName":"# mismatches"},{"values":[0,0,0,2,0,18,0,11,0,10,0,0,0,12],"quality":"Less is better","isMain":false,"metricName":"# indels"},{"values":[0,0,0,3,0,22,0,527,0,11,0,0,0,14],"quality":"Less is better","isMain":false,"metricName":"Indels length"},{"values":["3.23","95.72","5.10","315.92","76.50","1435.17","0.00","17.92","0.00","782.45","3.92","31.21","30.59","1304.49"],"quality":"Less is better","isMain":true,"metricName":"# mismatches per 100 kbp"},{"values":["0.00","0.00","0.00","6.26","0.00","40.55","0.00","98.57","0.00","28.98","0.00","0.00","0.00","27.04"],"quality":"Less is better","isMain":true,"metricName":"# indels per 100 kbp"},{"values":[0,0,0,2,0,18,0,0,0,10,0,0,0,12],"quality":"Less is better","isMain":false,"metricName":" # indels (<= 5 bp)"},{"values":[0,0,0,0,0,0,0,11,0,0,0,0,0,0],"quality":"Less is better","isMain":false,"metricName":" # indels (> 5 bp)"},{"values":[0,0,0,0,0,2,0,8685,0,0,0,0,0,2],"quality":"Less is better","isMain":false,"metricName":"# N's"},{"values":["0.00","0.00","0.00","0.00","0.00","2.23","0.00","35594.26","0.00","0.00","0.00","0.00","0.00","2.23"],"quality":"Less is better","isMain":true,"metricName":"# N's per 100 kbp"}]],["Statistics without reference",[{"values":[14,23,8,24,16,3,1,21,9,9,19,18,12,2],"quality":"Equal","isMain":true,"metricName":"# contigs"},{"values":[9,12,7,13,12,3,0,8,6,6,12,11,2,2],"quality":"Equal","isMain":false,"metricName":"# contigs (>= 1000 bp)"},{"values":[2,0,4,1,2,3,0,0,4,2,0,0,0,2],"quality":"Equal","isMain":false,"metricName":"# contigs (>= 5000 bp)"},{"values":[0,0,1,0,0,2,0,0,1,1,0,0,0,2],"quality":"Equal","isMain":false,"metricName":"# contigs (>= 10000 bp)"},{"values":[0,0,0,0,0,2,0,0,0,0,0,0,0,2],"quality":"Equal","isMain":false,"metricName":"# contigs (>= 25000 bp)"},{"values":[0,0,0,0,0,0,0,0,0,0,0,0,0,0],"quality":"Equal","isMain":false,"metricName":"# contigs (>= 50000 bp)"},{"values":[7566,2623,11497,8276,7596,44529,519,4789,12908,10422,2774,2774,2035,45094],"quality":"More is better","isMain":true,"metricName":"Largest contig"},{"values":[31823,25547,39224,38025,42225,89650,519,24400,39598,34728,25616,25864,9806,89623],"quality":"More is better","isMain":true,"metricName":"Total length"},{"values":[28509,17796,38584,30288,38914,89650,0,15218,37467,32681,20749,20708,3038,89623],"quality":"More is better","isMain":true,"metricName":"Total length (>= 1000 bp)"},{"values":[13362,0,31512,8276,13066,89650,0,0,31670,20171,0,0,0,89623],"quality":"More is better","isMain":false,"metricName":"Total length (>= 5000 bp)"},{"values":[0,0,11497,0,0,83586,0,0,12908,10422,0,0,0,89623],"quality":"More is better","isMain":true,"metricName":"Total length (>= 10000 bp)"},{"values":[0,0,0,0,0,83586,0,0,0,0,0,0,0,89623],"quality":"More is better","isMain":false,"metricName":"Total length (>= 25000 bp)"},{"values":[0,0,0,0,0,0,0,0,0,0,0,0,0,0],"quality":"More is better","isMain":true,"metricName":"Total length (>= 50000 bp)"},{"values":[4618,1168,7271,1948,4301,39057,519,1215,7568,9749,1289,1883,777,45094],"quality":"More is better","isMain":false,"metricName":"N50"},{"values":[1746,928,5387,1173,1927,39057,519,852,5112,4099,1198,1198,627,44529],"quality":"More is better","isMain":false,"metricName":"N75"},{"values":[3,8,3,5,4,2,1,6,2,2,6,6,5,1],"quality":"Less is better","isMain":false,"metricName":"L50"},{"values":[6,14,4,12,8,2,1,12,4,4,11,10,8,2],"quality":"Less is better","isMain":false,"metricName":"L75"},{"values":["45.07","44.85","45.20","44.97","45.06","45.15","44.89","45.87","45.16","45.15","44.92","45.40","44.43","45.14"],"quality":"Equal","isMain":false,"metricName":"GC (%)"}]],["Predicted genes",[]],["Similarity statistics",[{"values":[0,0,0,0,0,0,0,0,0,0,0,0,0,0],"quality":"Equal","isMain":false,"metricName":"# similar correct contigs"},{"values":[0,0,0,0,0,0,0,0,0,0,0,0,0,0],"quality":"Equal","isMain":false,"metricName":"# similar misassembled blocks"}]],["Reference statistics",[{"values":[44385,44385,44385,44385,44385,44385,44385,44385,44385,44385,44385,44385,44385,44385],"quality":"Equal","isMain":false,"metricName":"Reference length"},{"values":[1,1,1,1,1,1,1,1,1,1,1,1,1,1],"quality":"Equal","isMain":false,"metricName":"Reference fragments"},{"values":["45.25","45.25","45.25","45.25","45.25","45.25","45.25","45.25","45.25","45.25","45.25","45.25","45.25","45.25"],"quality":"Equal","isMain":false,"metricName":"Reference GC (%)"}]]],"referenceName":"gi\_108861984\_ref\_NC","date":"18 July 2018, Wednesday, 16:05:01","order":[0,1,2,3,4,5,6,7,8,9,10,11,12,13],"assembliesNames":["ABySS\_127","ABySS\_63","CLC","IDBA\_UD","MEGAHIT","MIRA","Ray\_Meta","SOAPdenovo2","SPAdes","SPAdes\_meta","SPAdes\_sc","SPAdes\_sc\_careful","Velvet","Geneious"]},{"assembliesWithNs":null,"minContig":500,"report":[["Genome statistics",[{"values":["93.904","87.920","97.369","98.333","98.267","98.005","96.698","98.605","99.012","98.246","98.246","97.905","46.200"],"quality":"More is better","isMain":true,"metricName":"Genome fraction (%)"},{"values":["1.003","1.002","1.009","1.001","1.000","1.004","1.015","1.000","1.001","1.000","1.000","1.001","1.000"],"quality":"Less is better","isMain":true,"metricName":"Duplication ratio"},{"values":[9098,5767,47123,42239,69688,22384,41759,69798,65942,69809,69798,33833,20359],"quality":"More is better","isMain":true,"metricName":"Largest alignment"},{"values":[98596,92272,102862,103054,102907,103026,101854,103262,103792,102903,102892,102591,48376],"quality":"More is better","isMain":true,"metricName":"Total aligned length"},{"values":[3036,1954,22975,20500,69688,10632,7921,69798,65942,69809,69798,11670,null],"quality":"More is better","isMain":false,"metricName":"NG50"},{"values":[1378,965,11459,18965,19007,8379,5138,33464,33839,33094,33094,6800,null],"quality":"More is better","isMain":false,"metricName":"NG75"},{"values":[3173,2331,22975,20500,69688,10632,7680,69798,65942,69809,69798,11670,16549],"quality":"More is better","isMain":false,"metricName":"NA50"},{"values":[1735,1308,11459,18965,19007,8379,5023,33464,33839,33094,33094,6800,16549],"quality":"More is better","isMain":false,"metricName":"NA75"},{"values":[3036,1954,22975,20500,69688,10632,7680,69798,65942,69809,69798,11670,null],"quality":"More is better","isMain":true,"metricName":"NGA50"},{"values":[1378,965,11459,18965,19007,8379,5023,33464,33839,33094,33094,6800,null],"quality":"More is better","isMain":false,"metricName":"NGA75"},{"values":[11,15,2,2,1,3,3,1,1,1,1,3,null],"quality":"Less is better","isMain":false,"metricName":"LG50"},{"values":[24,34,3,3,2,6,7,2,2,2,2,5,null],"quality":"Less is better","isMain":false,"metricName":"LG75"},{"values":[10,12,2,2,1,3,3,1,1,1,1,3,2],"quality":"Less is better","isMain":false,"metricName":"LA50"},{"values":[21,26,3,3,2,6,7,2,2,2,2,5,2],"quality":"Less is better","isMain":false,"metricName":"LA75"},{"values":[11,15,2,2,1,3,3,1,1,1,1,3,null],"quality":"Less is better","isMain":true,"metricName":"LGA50"},{"values":[24,34,3,3,2,6,7,2,2,2,2,5,null],"quality":"Less is better","isMain":false,"metricName":"LGA75"}]],["Misassemblies",[{"values":[0,0,0,0,0,0,0,0,0,0,0,0,0],"quality":"Less is better","isMain":true,"metricName":"# misassemblies"},{"values":[0,0,0,0,0,0,0,0,0,0,0,0,0],"quality":"Less is better","isMain":false,"metricName":" # relocations"},{"values":[0,0,0,0,0,0,0,0,0,0,0,0,0],"quality":"Less is better","isMain":false,"metricName":" # translocations"},{"values":[0,0,0,0,0,0,0,0,0,0,0,0,0],"quality":"Less is better","isMain":false,"metricName":" # inversions"},{"values":[0,0,0,0,0,0,0,0,0,0,0,0,0],"quality":"Less is better","isMain":false,"metricName":"# misassembled contigs"},{"values":[0,0,0,0,0,0,0,0,0,0,0,0,0],"quality":"Less is better","isMain":true,"metricName":"Misassembled contigs length"},{"values":[0,0,0,0,0,0,0,0,0,0,0,0,0],"quality":"Less is better","isMain":false,"metricName":"# local misassemblies"},{"values":[0,0,0,0,0,0,0,0,0,0,0,0,0],"quality":"Less is better","isMain":false,"metricName":"# unaligned mis. contigs"}]],["Unaligned",[{"values":[0,0,0,0,0,0,0,0,0,0,0,0,0],"quality":"Less is better","isMain":false,"metricName":"# fully unaligned contigs"},{"values":[0,0,0,0,0,0,0,0,0,0,0,0,0],"quality":"Less is better","isMain":false,"metricName":"Fully unaligned length"},{"values":[0,0,0,0,0,0,1,0,0,0,0,0,0],"quality":"Less is better","isMain":false,"metricName":"# partially unaligned contigs"},{"values":[0,0,0,0,0,0,616,0,0,0,0,0,0],"quality":"Less is better","isMain":false,"metricName":"Partially unaligned length"}]],["Mismatches",[{"values":[9,1,12,8,9,5,10,17,20,13,7,11,4],"quality":"Less is better","isMain":false,"metricName":"# mismatches"},{"values":[0,0,3,1,0,0,19,0,5,3,2,0,0],"quality":"Less is better","isMain":false,"metricName":"# indels"},{"values":[0,0,29,42,0,0,888,0,141,119,108,0,0],"quality":"Less is better","isMain":false,"metricName":"Indels length"},{"values":["9.15","1.09","11.77","7.77","8.75","4.87","9.88","16.47","19.29","12.64","6.80","10.73","8.27"],"quality":"Less is better","isMain":true,"metricName":"# mismatches per 100 kbp"},{"values":["0.00","0.00","2.94","0.97","0.00","0.00","18.77","0.00","4.82","2.92","1.94","0.00","0.00"],"quality":"Less is better","isMain":true,"metricName":"# indels per 100 kbp"},{"values":[0,0,2,0,0,0,0,0,1,0,0,0,0],"quality":"Less is better","isMain":false,"metricName":" # indels (<= 5 bp)"},{"values":[0,0,1,1,0,0,19,0,4,3,2,0,0],"quality":"Less is better","isMain":false,"metricName":" # indels (> 5 bp)"},{"values":[0,0,0,0,0,27,1673,0,0,10,0,0,14],"quality":"Less is better","isMain":false,"metricName":"# N's"},{"values":["0.00","0.00","0.00","0.00","0.00","26.21","1618.16","0.00","0.00","9.72","0.00","0.00","28.94"],"quality":"Less is better","isMain":true,"metricName":"# N's per 100 kbp"}]],["Statistics without reference",[{"values":[44,52,7,6,3,12,16,2,3,2,2,12,3],"quality":"Equal","isMain":true,"metricName":"# contigs"},{"values":[33,33,7,6,3,12,15,2,3,2,2,12,3],"quality":"Equal","isMain":false,"metricName":"# contigs (>= 1000 bp)"},{"values":[3,2,5,4,3,8,7,2,2,2,2,6,3],"quality":"Equal","isMain":false,"metricName":"# contigs (>= 5000 bp)"},{"values":[0,0,3,4,3,4,1,2,2,2,2,4,3],"quality":"Equal","isMain":false,"metricName":"# contigs (>= 10000 bp)"},{"values":[0,0,1,1,1,0,1,2,2,2,2,1,0],"quality":"Equal","isMain":false,"metricName":"# contigs (>= 25000 bp)"},{"values":[0,0,0,0,1,0,0,1,1,1,1,0,0],"quality":"Equal","isMain":false,"metricName":"# contigs (>= 50000 bp)"},{"values":[9098,5767,47123,42239,69688,22384,41951,69798,65942,69809,69798,33833,20359],"quality":"More is better","isMain":true,"metricName":"Largest contig"},{"values":[98596,92272,102862,103054,102907,103026,103389,103262,103792,102903,102892,102591,48376],"quality":"More is better","isMain":true,"metricName":"Total length"},{"values":[89999,77813,102862,103054,102907,103026,102797,103262,103792,102903,102892,102591,48376],"quality":"More is better","isMain":true,"metricName":"Total length (>= 1000 bp)"},{"values":[23025,11371,97613,96046,102907,94179,81888,103262,99781,102903,102892,84688,48376],"quality":"More is better","isMain":false,"metricName":"Total length (>= 5000 bp)"},{"values":[0,0,81557,96046,102907,63987,41951,103262,99781,102903,102892,72822,48376],"quality":"More is better","isMain":true,"metricName":"Total length (>= 10000 bp)"},{"values":[0,0,47123,42239,69688,0,41951,103262,99781,102903,102892,33833,0],"quality":"More is better","isMain":false,"metricName":"Total length (>= 25000 bp)"},{"values":[0,0,0,0,69688,0,0,69798,65942,69809,69798,0,0],"quality":"More is better","isMain":true,"metricName":"Total length (>= 50000 bp)"},{"values":[3173,2331,22975,20500,69688,10632,7921,69798,65942,69809,69798,11670,16549],"quality":"More is better","isMain":false,"metricName":"N50"},{"values":[1735,1308,11459,18965,19007,8379,5138,33464,33839,33094,33094,6800,16549],"quality":"More is better","isMain":false,"metricName":"N75"},{"values":[10,12,2,2,1,3,3,1,1,1,1,3,2],"quality":"Less is better","isMain":false,"metricName":"L50"},{"values":[21,26,3,3,2,6,7,2,2,2,2,5,2],"quality":"Less is better","isMain":false,"metricName":"L75"},{"values":["32.64","32.86","32.85","32.85","32.86","32.67","32.91","32.82","32.78","32.87","32.87","32.89","33.20"],"quality":"Equal","isMain":false,"metricName":"GC (%)"}]],["Predicted genes",[]],["Similarity statistics",[{"values":[0,0,0,0,0,0,0,0,0,0,0,0,0],"quality":"Equal","isMain":false,"metricName":"# similar correct contigs"},{"values":[0,0,0,0,0,0,0,0,0,0,0,0,0],"quality":"Equal","isMain":false,"metricName":"# similar misassembled blocks"}]],["Reference statistics",[{"values":[104710,104710,104710,104710,104710,104710,104710,104710,104710,104710,104710,104710,104710],"quality":"Equal","isMain":false,"metricName":"Reference length"},{"values":[1,1,1,1,1,1,1,1,1,1,1,1,1],"quality":"Equal","isMain":false,"metricName":"Reference fragments"},{"values":["32.67","32.67","32.67","32.67","32.67","32.67","32.67","32.67","32.67","32.67","32.67","32.67","32.67"],"quality":"Equal","isMain":false,"metricName":"Reference GC (%)"}]]],"referenceName":"gi\_109255272\_ref\_NC","date":"18 July 2018, Wednesday, 16:05:11","order":[0,1,2,3,4,5,6,7,8,9,10,11,12],"assembliesNames":["ABySS\_127","ABySS\_63","CLC","IDBA\_UD","MEGAHIT","MIRA","SOAPdenovo2","SPAdes","SPAdes\_meta","SPAdes\_sc","SPAdes\_sc\_careful","Velvet","Geneious"]},{"assembliesWithNs":null,"minContig":500,"report":[["Genome statistics",[{"values":["95.332","90.004","90.241","97.586","95.361","98.803","0.285","79.773","99.101","98.347","89.821","89.788","82.239","81.986"],"quality":"More is better","isMain":true,"metricName":"Genome fraction (%)"},{"values":["1.004","1.006","1.008","1.003","1.002","1.032","1.000","1.044","1.005","1.002","1.003","1.003","1.001","1.010"],"quality":"Less is better","isMain":true,"metricName":"Duplication ratio"},{"values":[17617,7345,22772,43111,23802,25140,544,18347,78360,74238,22035,22035,18370,37817],"quality":"More is better","isMain":true,"metricName":"Largest alignment"},{"values":[182964,173011,172599,187067,182667,194553,544,153228,190222,188242,171925,171923,157252,157960],"quality":"More is better","isMain":true,"metricName":"Total aligned length"},{"values":[4022,2874,15405,28370,15064,24294,null,7491,181334,51932,14477,14475,7558,30458],"quality":"More is better","isMain":false,"metricName":"NG50"},{"values":[2476,1504,6419,21396,9343,8751,null,2851,181334,25133,8053,8053,2097,8751],"quality":"More is better","isMain":false,"metricName":"NG75"},{"values":[4248,3133,15405,28370,15064,12922,544,8460,72104,51932,15084,15084,9724,23441],"quality":"More is better","isMain":false,"metricName":"NA50"},{"values":[2852,1758,7579,21396,10356,6777,544,3904,72104,25133,9674,9674,4368,10989],"quality":"More is better","isMain":false,"metricName":"NA75"},{"values":[4022,2874,10833,28370,15064,12922,null,6223,72104,51932,14218,14216,7558,12922],"quality":"More is better","isMain":true,"metricName":"NGA50"},{"values":[2476,1504,5312,21396,9343,7474,null,1919,72104,25133,8053,8053,2097,6153],"quality":"More is better","isMain":false,"metricName":"NGA75"},{"values":[14,20,5,3,5,3,null,8,1,2,6,6,8,3],"quality":"Less is better","isMain":false,"metricName":"LG50"},{"values":[28,43,11,5,10,7,null,19,1,3,10,10,20,5],"quality":"Less is better","isMain":false,"metricName":"LG75"},{"values":[13,17,5,3,5,5,1,7,2,2,5,5,6,3],"quality":"Less is better","isMain":false,"metricName":"LA50"},{"values":[26,35,10,5,9,11,1,14,2,3,9,9,12,6],"quality":"Less is better","isMain":false,"metricName":"LA75"},{"values":[14,20,6,3,5,5,null,9,2,2,6,6,8,4],"quality":"Less is better","isMain":true,"metricName":"LGA50"},{"values":[28,43,12,5,10,10,null,22,2,3,10,10,20,9],"quality":"Less is better","isMain":false,"metricName":"LGA75"}]],["Misassemblies",[{"values":[0,0,3,0,0,5,0,0,2,0,0,0,0,5],"quality":"Less is better","isMain":true,"metricName":"# misassemblies"},{"values":[0,0,3,0,0,5,0,0,2,0,0,0,0,5],"quality":"Less is better","isMain":false,"metricName":" # relocations"},{"values":[0,0,0,0,0,0,0,0,0,0,0,0,0,0],"quality":"Less is better","isMain":false,"metricName":" # translocations"},{"values":[0,0,0,0,0,0,0,0,0,0,0,0,0,0],"quality":"Less is better","isMain":false,"metricName":" # inversions"},{"values":[0,0,3,0,0,2,0,0,1,0,0,0,0,2],"quality":"Less is better","isMain":false,"metricName":"# misassembled contigs"},{"values":[0,0,40812,0,0,63908,0,0,181334,0,0,0,0,95370],"quality":"Less is better","isMain":true,"metricName":"Misassembled contigs length"},{"values":[0,0,0,0,1,2,0,2,3,0,0,0,0,2],"quality":"Less is better","isMain":false,"metricName":"# local misassemblies"},{"values":[0,0,0,0,0,0,0,1,0,0,0,0,0,0],"quality":"Less is better","isMain":false,"metricName":"# unaligned mis. contigs"}]],["Unaligned",[{"values":[0,0,0,0,0,0,0,0,0,0,0,0,0,0],"quality":"Less is better","isMain":false,"metricName":"# fully unaligned contigs"},{"values":[0,0,0,0,0,0,0,0,0,0,0,0,0,0],"quality":"Less is better","isMain":false,"metricName":"Fully unaligned length"},{"values":[0,0,0,0,0,0,0,1,1,0,0,0,0,0],"quality":"Less is better","isMain":false,"metricName":"# partially unaligned contigs"},{"values":[0,0,0,0,0,0,0,2325,903,0,0,0,0,0],"quality":"Less is better","isMain":false,"metricName":"Partially unaligned length"}]],["Mismatches",[{"values":[14,5,42,4,9,84,0,4,29,14,13,7,12,78],"quality":"Less is better","isMain":false,"metricName":"# mismatches"},{"values":[2,0,13,2,0,33,0,50,3,0,6,6,2,34],"quality":"Less is better","isMain":false,"metricName":"# indels"},{"values":[46,0,98,46,0,128,0,2439,3,0,164,166,16,91],"quality":"Less is better","isMain":false,"metricName":"Indels length"},{"values":["7.68","2.91","24.35","2.14","4.94","44.49","0.00","2.62","15.31","7.45","7.57","4.08","7.64","49.78"],"quality":"Less is better","isMain":true,"metricName":"# mismatches per 100 kbp"},{"values":["1.10","0.00","7.54","1.07","0.00","17.48","0.00","32.80","1.58","0.00","3.50","3.50","1.27","21.70"],"quality":"Less is better","isMain":true,"metricName":"# indels per 100 kbp"},{"values":[0,0,11,0,0,29,0,0,3,0,2,2,1,31],"quality":"Less is better","isMain":false,"metricName":" # indels (<= 5 bp)"},{"values":[2,0,2,2,0,4,0,50,0,0,4,4,1,3],"quality":"Less is better","isMain":false,"metricName":" # indels (> 5 bp)"},{"values":[0,0,0,0,0,47,0,7534,1003,0,10,0,0,44],"quality":"Less is better","isMain":false,"metricName":"# N's"},{"values":["0.00","0.00","0.00","0.00","0.00","24.12","0.00","4665.62","524.51","0.00","5.81","0.00","0.00","27.82"],"quality":"Less is better","isMain":true,"metricName":"# N's per 100 kbp"}]],["Statistics without reference",[{"values":[58,77,24,16,26,21,1,31,4,7,27,27,31,7],"quality":"Equal","isMain":true,"metricName":"# contigs"},{"values":[47,52,20,10,17,16,0,27,3,5,16,16,26,7],"quality":"Equal","isMain":false,"metricName":"# contigs (>= 1000 bp)"},{"values":[10,10,12,7,12,13,0,10,2,5,10,10,10,7],"quality":"Equal","isMain":false,"metricName":"# contigs (>= 5000 bp)"},{"values":[1,0,6,7,9,5,0,5,1,5,8,8,5,4],"quality":"Equal","isMain":false,"metricName":"# contigs (>= 10000 bp)"},{"values":[0,0,1,3,0,2,0,0,1,4,0,0,0,3],"quality":"Equal","isMain":false,"metricName":"# contigs (>= 25000 bp)"},{"values":[0,0,0,0,0,1,0,0,1,2,0,0,0,1],"quality":"Equal","isMain":false,"metricName":"# contigs (>= 50000 bp)"},{"values":[17617,7345,27071,43111,23802,51400,544,18991,181334,74238,22035,22035,18370,51400],"quality":"More is better","isMain":true,"metricName":"Largest contig"},{"values":[182964,173011,173748,187067,182667,194841,544,161479,191225,188242,172184,172182,157252,158182],"quality":"More is better","isMain":true,"metricName":"Total length"},{"values":[175252,155664,170859,182584,176379,191047,0,158624,190268,186795,163819,163817,153621,158182],"quality":"More is better","isMain":true,"metricName":"Total length (>= 1000 bp)"},{"values":[81386,61031,154930,174508,165958,185331,0,108371,189110,186795,145179,145177,109872,158182],"quality":"More is better","isMain":false,"metricName":"Total length (>= 5000 bp)"},{"values":[17617,0,109726,174508,143114,129854,0,71400,181334,186795,127452,127450,73222,136817],"quality":"More is better","isMain":true,"metricName":"Total length (>= 10000 bp)"},{"values":[0,0,27071,100641,0,76540,0,0,181334,176356,0,0,0,125828],"quality":"More is better","isMain":false,"metricName":"Total length (>= 25000 bp)"},{"values":[0,0,0,0,0,51400,0,0,181334,126170,0,0,0,51400],"quality":"More is better","isMain":true,"metricName":"Total length (>= 50000 bp)"},{"values":[4248,3133,15405,28370,15064,24294,544,8460,181334,51932,15084,15084,9724,43970],"quality":"More is better","isMain":false,"metricName":"N50"},{"values":[2852,1758,7946,21396,10356,8751,544,4641,181334,25133,9674,9674,4368,30458],"quality":"More is better","isMain":false,"metricName":"N75"},{"values":[13,17,5,3,5,3,1,7,1,2,5,5,6,2],"quality":"Less is better","isMain":false,"metricName":"L50"},{"values":[26,35,9,5,9,7,1,13,1,3,9,9,12,3],"quality":"Less is better","isMain":false,"metricName":"L75"},{"values":["47.99","48.19","48.41","48.01","48.14","47.79","49.82","48.98","47.92","47.97","48.47","48.47","48.89","47.88"],"quality":"Equal","isMain":false,"metricName":"GC (%)"}]],["Predicted genes",[]],["Similarity statistics",[{"values":[0,0,0,0,0,0,0,0,0,0,0,0,0,0],"quality":"Equal","isMain":false,"metricName":"# similar correct contigs"},{"values":[0,0,0,0,0,0,0,0,0,0,0,0,0,0],"quality":"Equal","isMain":false,"metricName":"# similar misassembled blocks"}]],["Reference statistics",[{"values":[191100,191100,191100,191100,191100,191100,191100,191100,191100,191100,191100,191100,191100,191100],"quality":"Equal","isMain":false,"metricName":"Reference length"},{"values":[1,1,1,1,1,1,1,1,1,1,1,1,1,1],"quality":"Equal","isMain":false,"metricName":"Reference fragments"},{"values":["47.89","47.89","47.89","47.89","47.89","47.89","47.89","47.89","47.89","47.89","47.89","47.89","47.89","47.89"],"quality":"Equal","isMain":false,"metricName":"Reference GC (%)"}]]],"referenceName":"gi\_109287878\_ref\_NC","date":"18 July 2018, Wednesday, 16:05:23","order":[0,1,2,3,4,5,6,7,8,9,10,11,12,13],"assembliesNames":["ABySS\_127","ABySS\_63","CLC","IDBA\_UD","MEGAHIT","MIRA","Ray\_Meta","SOAPdenovo2","SPAdes","SPAdes\_meta","SPAdes\_sc","SPAdes\_sc\_careful","Velvet","Geneious"]},{"assembliesWithNs":null,"minContig":500,"report":[["Genome statistics",[{"values":["93.486","85.497","100.000","100.000","100.000","98.786","98.841","100.000","100.000","100.000","100.000","99.968","80.258"],"quality":"More is better","isMain":true,"metricName":"Genome fraction (%)"},{"values":["1.002","1.003","1.001","1.003","1.004","1.036","1.016","1.004","1.002","1.002","1.002","1.000","1.000"],"quality":"Less is better","isMain":true,"metricName":"Duplication ratio"},{"values":[8101,4507,34543,34624,34666,16683,15551,34652,34580,34580,34580,34514,16683],"quality":"More is better","isMain":true,"metricName":"Largest alignment"},{"values":[32329,29599,34543,34624,34666,35328,34199,34652,34580,34580,34580,34514,27709],"quality":"More is better","isMain":true,"metricName":"Total aligned length"},{"values":[1796,1451,34543,34624,34666,11026,9199,34652,34580,34580,34580,34514,11026],"quality":"More is better","isMain":false,"metricName":"NG50"},{"values":[1367,817,34543,34624,34666,11026,4330,34652,34580,34580,34580,34514,11026],"quality":"More is better","isMain":false,"metricName":"NG75"},{"values":[1989,1674,34543,34624,34666,11026,9199,34652,34580,34580,34580,34514,16683],"quality":"More is better","isMain":false,"metricName":"NA50"},{"values":[1469,1317,34543,34624,34666,11026,4237,34652,34580,34580,34580,34514,11026],"quality":"More is better","isMain":false,"metricName":"NA75"},{"values":[1796,1451,34543,34624,34666,11026,9199,34652,34580,34580,34580,34514,11026],"quality":"More is better","isMain":true,"metricName":"NGA50"},{"values":[1367,817,34543,34624,34666,11026,4237,34652,34580,34580,34580,34514,11026],"quality":"More is better","isMain":false,"metricName":"NGA75"},{"values":[6,8,1,1,1,2,2,1,1,1,1,1,2],"quality":"Less is better","isMain":false,"metricName":"LG50"},{"values":[11,15,1,1,1,2,3,1,1,1,1,1,2],"quality":"Less is better","isMain":false,"metricName":"LG75"},{"values":[5,6,1,1,1,2,2,1,1,1,1,1,1],"quality":"Less is better","isMain":false,"metricName":"LA50"},{"values":[10,11,1,1,1,2,3,1,1,1,1,1,2],"quality":"Less is better","isMain":false,"metricName":"LA75"},{"values":[6,8,1,1,1,2,2,1,1,1,1,1,2],"quality":"Less is better","isMain":true,"metricName":"LGA50"},{"values":[11,15,1,1,1,2,3,1,1,1,1,1,2],"quality":"Less is better","isMain":false,"metricName":"LGA75"}]],["Misassemblies",[{"values":[0,0,0,0,0,0,0,0,0,0,0,0,0],"quality":"Less is better","isMain":true,"metricName":"# misassemblies"},{"values":[0,0,0,0,0,0,0,0,0,0,0,0,0],"quality":"Less is better","isMain":false,"metricName":" # relocations"},{"values":[0,0,0,0,0,0,0,0,0,0,0,0,0],"quality":"Less is better","isMain":false,"metricName":" # translocations"},{"values":[0,0,0,0,0,0,0,0,0,0,0,0,0],"quality":"Less is better","isMain":false,"metricName":" # inversions"},{"values":[0,0,0,0,0,0,0,0,0,0,0,0,0],"quality":"Less is better","isMain":false,"metricName":"# misassembled contigs"},{"values":[0,0,0,0,0,0,0,0,0,0,0,0,0],"quality":"Less is better","isMain":true,"metricName":"Misassembled contigs length"},{"values":[0,0,0,1,1,0,0,1,0,0,0,0,0],"quality":"Less is better","isMain":false,"metricName":"# local misassemblies"},{"values":[0,0,0,0,0,0,0,0,0,0,0,0,0],"quality":"Less is better","isMain":false,"metricName":"# unaligned mis. contigs"}]],["Unaligned",[{"values":[0,0,0,0,0,0,0,0,0,0,0,0,0],"quality":"Less is better","isMain":false,"metricName":"# fully unaligned contigs"},{"values":[0,0,0,0,0,0,0,0,0,0,0,0,0],"quality":"Less is better","isMain":false,"metricName":"Fully unaligned length"},{"values":[0,0,0,0,0,0,0,0,0,0,0,0,0],"quality":"Less is better","isMain":false,"metricName":"# partially unaligned contigs"},{"values":[0,0,0,0,0,0,0,0,0,0,0,0,0],"quality":"Less is better","isMain":false,"metricName":"Partially unaligned length"}]],["Mismatches",[{"values":[3,1,3,0,0,0,1,9,5,2,0,1,0],"quality":"Less is better","isMain":false,"metricName":"# mismatches"},{"values":[0,0,1,0,0,0,5,0,1,1,1,0,0],"quality":"Less is better","isMain":false,"metricName":"# indels"},{"values":[0,0,18,0,0,0,227,0,55,55,55,0,0],"quality":"Less is better","isMain":false,"metricName":"Indels length"},{"values":["9.29","3.39","8.69","0.00","0.00","0.00","2.93","26.07","14.48","5.79","0.00","2.90","0.00"],"quality":"Less is better","isMain":true,"metricName":"# mismatches per 100 kbp"},{"values":["0.00","0.00","2.90","0.00","0.00","0.00","14.65","0.00","2.90","2.90","2.90","0.00","0.00"],"quality":"Less is better","isMain":true,"metricName":"# indels per 100 kbp"},{"values":[0,0,0,0,0,0,0,0,0,0,0,0,0],"quality":"Less is better","isMain":false,"metricName":" # indels (<= 5 bp)"},{"values":[0,0,1,0,0,0,5,0,1,1,1,0,0],"quality":"Less is better","isMain":false,"metricName":" # indels (> 5 bp)"},{"values":[0,0,0,0,0,6,526,0,0,0,0,0,3],"quality":"Less is better","isMain":false,"metricName":"# N's"},{"values":["0.00","0.00","0.00","0.00","0.00","16.98","1516.64","0.00","0.00","0.00","0.00","0.00","10.83"],"quality":"Less is better","isMain":true,"metricName":"# N's per 100 kbp"}]],["Statistics without reference",[{"values":[17,20,1,1,1,5,5,1,1,1,1,1,2],"quality":"Equal","isMain":true,"metricName":"# contigs"},{"values":[13,12,1,1,1,3,5,1,1,1,1,1,2],"quality":"Equal","isMain":false,"metricName":"# contigs (>= 1000 bp)"},{"values":[1,0,1,1,1,3,2,1,1,1,1,1,2],"quality":"Equal","isMain":false,"metricName":"# contigs (>= 5000 bp)"},{"values":[0,0,1,1,1,2,1,1,1,1,1,1,2],"quality":"Equal","isMain":false,"metricName":"# contigs (>= 10000 bp)"},{"values":[0,0,1,1,1,0,0,1,1,1,1,1,0],"quality":"Equal","isMain":false,"metricName":"# contigs (>= 25000 bp)"},{"values":[0,0,0,0,0,0,0,0,0,0,0,0,0],"quality":"Equal","isMain":false,"metricName":"# contigs (>= 50000 bp)"},{"values":[8101,4507,34543,34624,34666,16683,15941,34652,34580,34580,34580,34514,16683],"quality":"More is better","isMain":true,"metricName":"Largest contig"},{"values":[32329,29599,34543,34624,34666,35328,34682,34652,34580,34580,34580,34514,27709],"quality":"More is better","isMain":true,"metricName":"Total length"},{"values":[29114,23589,34543,34624,34666,34106,34682,34652,34580,34580,34580,34514,27709],"quality":"More is better","isMain":true,"metricName":"Total length (>= 1000 bp)"},{"values":[8101,0,34543,34624,34666,34106,25140,34652,34580,34580,34580,34514,27709],"quality":"More is better","isMain":false,"metricName":"Total length (>= 5000 bp)"},{"values":[0,0,34543,34624,34666,27709,15941,34652,34580,34580,34580,34514,27709],"quality":"More is better","isMain":true,"metricName":"Total length (>= 10000 bp)"},{"values":[0,0,34543,34624,34666,0,0,34652,34580,34580,34580,34514,0],"quality":"More is better","isMain":false,"metricName":"Total length (>= 25000 bp)"},{"values":[0,0,0,0,0,0,0,0,0,0,0,0,0],"quality":"More is better","isMain":true,"metricName":"Total length (>= 50000 bp)"},{"values":[1989,1674,34543,34624,34666,11026,9199,34652,34580,34580,34580,34514,16683],"quality":"More is better","isMain":false,"metricName":"N50"},{"values":[1469,1317,34543,34624,34666,11026,4330,34652,34580,34580,34580,34514,11026],"quality":"More is better","isMain":false,"metricName":"N75"},{"values":[5,6,1,1,1,2,2,1,1,1,1,1,1],"quality":"Less is better","isMain":false,"metricName":"L50"},{"values":[10,11,1,1,1,2,3,1,1,1,1,1,2],"quality":"Less is better","isMain":false,"metricName":"L75"},{"values":["41.43","41.57","41.58","41.56","41.54","41.73","41.59","41.52","41.56","41.57","41.57","41.57","41.81"],"quality":"Equal","isMain":false,"metricName":"GC (%)"}]],["Predicted genes",[]],["Similarity statistics",[{"values":[0,0,0,0,0,0,0,0,0,0,0,0,0],"quality":"Equal","isMain":false,"metricName":"# similar correct contigs"},{"values":[0,0,0,0,0,0,0,0,0,0,0,0,0],"quality":"Equal","isMain":false,"metricName":"# similar misassembled blocks"}]],["Reference statistics",[{"values":[34525,34525,34525,34525,34525,34525,34525,34525,34525,34525,34525,34525,34525],"quality":"Equal","isMain":false,"metricName":"Reference length"},{"values":[1,1,1,1,1,1,1,1,1,1,1,1,1],"quality":"Equal","isMain":false,"metricName":"Reference fragments"},{"values":["41.57","41.57","41.57","41.57","41.57","41.57","41.57","41.57","41.57","41.57","41.57","41.57","41.57"],"quality":"Equal","isMain":false,"metricName":"Reference GC (%)"}]]],"referenceName":"gi\_109289936\_ref\_NC","date":"18 July 2018, Wednesday, 16:05:33","order":[0,1,2,3,4,5,6,7,8,9,10,11,12],"assembliesNames":["ABySS\_127","ABySS\_63","CLC","IDBA\_UD","MEGAHIT","MIRA","SOAPdenovo2","SPAdes","SPAdes\_meta","SPAdes\_sc","SPAdes\_sc\_careful","Velvet","Geneious"]},{"assembliesWithNs":null,"minContig":500,"report":[["Genome statistics",[{"values":["67.039","59.392","92.833","94.784","94.567","83.171","89.604","97.501","96.220","96.797","96.622","67.313","21.707"],"quality":"More is better","isMain":true,"metricName":"Genome fraction (%)"},{"values":["1.001","1.001","1.016","1.002","1.000","1.027","1.017","1.001","1.003","1.003","1.002","1.000","1.000"],"quality":"Less is better","isMain":true,"metricName":"Duplication ratio"},{"values":[3135,3305,9377,13715,13879,6185,7925,13879,13879,13879,13879,12739,7220],"quality":"More is better","isMain":true,"metricName":"Largest alignment"},{"values":[108374,95975,152006,153284,152721,137876,145498,157611,155833,156736,156400,108719,35056],"quality":"More is better","isMain":true,"metricName":"Total aligned length"},{"values":[821,692,4084,7434,5625,2473,2239,6493,5667,6305,6305,2688,null],"quality":"More is better","isMain":false,"metricName":"NG50"},{"values":[null,null,1810,4013,3337,1100,1197,4013,3176,3739,3403,null,null],"quality":"More is better","isMain":false,"metricName":"NG75"},{"values":[1238,1067,4482,5058,5056,2608,2293,5843,5667,5233,5233,4049,3480],"quality":"More is better","isMain":false,"metricName":"NA50"},{"values":[821,761,2081,2400,3166,1510,1418,3176,3461,2402,2402,2322,1708],"quality":"More is better","isMain":false,"metricName":"NA75"},{"values":[821,692,4084,5711,5056,2335,2132,5843,5667,5667,5667,2405,null],"quality":"More is better","isMain":true,"metricName":"NGA50"},{"values":[null,null,1670,2880,3166,1088,1187,3583,3176,2880,2880,null,null],"quality":"More is better","isMain":false,"metricName":"NGA75"},{"values":[54,69,14,7,9,21,21,8,9,9,9,16,null],"quality":"Less is better","isMain":false,"metricName":"LG50"},{"values":[null,null,29,15,18,46,46,15,18,17,17,null,null],"quality":"Less is better","isMain":false,"metricName":"LG75"},{"values":[26,29,13,10,10,18,18,9,9,10,10,9,5],"quality":"Less is better","isMain":false,"metricName":"LA50"},{"values":[55,57,26,21,20,35,38,19,17,21,21,19,9],"quality":"Less is better","isMain":false,"metricName":"LA75"},{"values":[54,69,14,9,10,22,21,9,9,9,9,17,null],"quality":"Less is better","isMain":true,"metricName":"LGA50"},{"values":[null,null,30,19,20,48,46,17,18,19,19,null,null],"quality":"Less is better","isMain":false,"metricName":"LGA75"}]],["Misassemblies",[{"values":[0,0,0,0,0,0,0,0,0,0,0,0,0],"quality":"Less is better","isMain":true,"metricName":"# misassemblies"},{"values":[0,0,0,0,0,0,0,0,0,0,0,0,0],"quality":"Less is better","isMain":false,"metricName":" # relocations"},{"values":[0,0,0,0,0,0,0,0,0,0,0,0,0],"quality":"Less is better","isMain":false,"metricName":" # translocations"},{"values":[0,0,0,0,0,0,0,0,0,0,0,0,0],"quality":"Less is better","isMain":false,"metricName":" # inversions"},{"values":[0,0,0,0,0,0,0,0,0,0,0,0,0],"quality":"Less is better","isMain":false,"metricName":"# misassembled contigs"},{"values":[0,0,0,0,0,0,0,0,0,0,0,0,0],"quality":"Less is better","isMain":true,"metricName":"Misassembled contigs length"},{"values":[0,0,0,0,0,0,0,0,0,0,0,0,0],"quality":"Less is better","isMain":false,"metricName":"# local misassemblies"},{"values":[0,0,0,0,0,0,0,0,0,0,0,0,0],"quality":"Less is better","isMain":false,"metricName":"# unaligned mis. contigs"}]],["Unaligned",[{"values":[0,0,0,0,0,0,0,0,0,0,0,0,0],"quality":"Less is better","isMain":false,"metricName":"# fully unaligned contigs"},{"values":[0,0,0,0,0,0,0,0,0,0,0,0,0],"quality":"Less is better","isMain":false,"metricName":"Fully unaligned length"},{"values":[0,0,0,2,1,1,0,2,0,1,1,2,3],"quality":"Less is better","isMain":false,"metricName":"# partially unaligned contigs"},{"values":[0,0,0,15607,6147,2697,0,8329,0,9833,9833,3391,10451],"quality":"Less is better","isMain":false,"metricName":"Partially unaligned length"}]],["Mismatches",[{"values":[15,6,99,57,50,24,47,102,133,77,69,57,31],"quality":"Less is better","isMain":false,"metricName":"# mismatches"},{"values":[0,0,1,1,1,1,22,1,0,0,0,1,9],"quality":"Less is better","isMain":false,"metricName":"# indels"},{"values":[0,0,1,1,1,1,1030,1,0,0,0,1,11],"quality":"Less is better","isMain":false,"metricName":"Indels length"},{"values":["13.86","6.26","66.04","37.24","32.74","17.87","32.48","64.79","85.60","49.26","44.22","52.44","88.44"],"quality":"Less is better","isMain":true,"metricName":"# mismatches per 100 kbp"},{"values":["0.00","0.00","0.67","0.65","0.65","0.74","15.21","0.64","0.00","0.00","0.00","0.92","25.68"],"quality":"Less is better","isMain":true,"metricName":"# indels per 100 kbp"},{"values":[0,0,1,1,1,1,0,1,0,0,0,1,9],"quality":"Less is better","isMain":false,"metricName":" # indels (<= 5 bp)"},{"values":[0,0,0,0,0,0,22,0,0,0,0,0,0],"quality":"Less is better","isMain":false,"metricName":" # indels (> 5 bp)"},{"values":[0,0,0,0,0,92,2148,0,0,0,112,0,335],"quality":"Less is better","isMain":false,"metricName":"# N's"},{"values":["0.00","0.00","0.00","0.00","0.00","65.45","1460.13","0.00","0.00","0.00","67.37","0.00","736.15"],"quality":"Less is better","isMain":true,"metricName":"# N's per 100 kbp"}]],["Statistics without reference",[{"values":[94,94,61,38,37,70,75,35,38,40,40,36,9],"quality":"Equal","isMain":true,"metricName":"# contigs"},{"values":[37,31,41,35,35,49,55,34,36,38,38,32,9],"quality":"Equal","isMain":false,"metricName":"# contigs (>= 1000 bp)"},{"values":[0,0,8,12,10,4,5,11,9,11,11,7,3],"quality":"Equal","isMain":false,"metricName":"# contigs (>= 5000 bp)"},{"values":[0,0,0,4,4,0,0,4,4,5,5,1,0],"quality":"Equal","isMain":false,"metricName":"# contigs (>= 10000 bp)"},{"values":[0,0,0,0,0,0,0,0,0,0,0,0,0],"quality":"Equal","isMain":false,"metricName":"# contigs (>= 25000 bp)"},{"values":[0,0,0,0,0,0,0,0,0,0,0,0,0],"quality":"Equal","isMain":false,"metricName":"# contigs (>= 50000 bp)"},{"values":[3135,3305,9377,19021,13879,6185,7925,19179,13879,13879,13879,12739,9897],"quality":"More is better","isMain":true,"metricName":"Largest contig"},{"values":[108374,95975,152234,168894,158868,140573,147110,165943,155845,166572,166236,112113,45507],"quality":"More is better","isMain":true,"metricName":"Total length"},{"values":[66032,50066,137226,166642,157383,125021,132066,165147,154527,165086,164750,108984,45507],"quality":"More is better","isMain":true,"metricName":"Total length (>= 1000 bp)"},{"values":[0,0,54484,110104,89022,23758,31857,103787,83553,97833,97833,47954,25851],"quality":"More is better","isMain":false,"metricName":"Total length (>= 5000 bp)"},{"values":[0,0,0,56031,49672,0,0,56716,49367,58414,58414,12739,0],"quality":"More is better","isMain":true,"metricName":"Total length (>= 10000 bp)"},{"values":[0,0,0,0,0,0,0,0,0,0,0,0,0],"quality":"More is better","isMain":false,"metricName":"Total length (>= 25000 bp)"},{"values":[0,0,0,0,0,0,0,0,0,0,0,0,0],"quality":"More is better","isMain":true,"metricName":"Total length (>= 50000 bp)"},{"values":[1238,1067,4482,6344,5625,2683,2293,6493,5667,6305,6305,4843,6107],"quality":"More is better","isMain":false,"metricName":"N50"},{"values":[821,761,2081,3586,3337,1517,1443,3795,3461,3176,3176,2365,4104],"quality":"More is better","isMain":false,"metricName":"N75"},{"values":[26,29,13,8,9,17,18,8,9,9,9,9,3],"quality":"Less is better","isMain":false,"metricName":"L50"},{"values":[55,57,26,16,18,34,37,16,17,18,18,18,5],"quality":"Less is better","isMain":false,"metricName":"L75"},{"values":["40.95","41.00","40.92","41.28","41.15","41.17","40.92","41.20","40.94","41.15","41.14","41.21","41.92"],"quality":"Equal","isMain":false,"metricName":"GC (%)"}]],["Predicted genes",[]],["Similarity statistics",[{"values":[0,0,0,1,1,0,0,1,1,1,1,1,0],"quality":"Equal","isMain":false,"metricName":"# similar correct contigs"},{"values":[0,0,0,0,0,0,0,0,0,0,0,0,0],"quality":"Equal","isMain":false,"metricName":"# similar misassembled blocks"}]],["Reference statistics",[{"values":[161475,161475,161475,161475,161475,161475,161475,161475,161475,161475,161475,161475,161475],"quality":"Equal","isMain":false,"metricName":"Reference length"},{"values":[1,1,1,1,1,1,1,1,1,1,1,1,1],"quality":"Equal","isMain":false,"metricName":"Reference fragments"},{"values":["41.04","41.04","41.04","41.04","41.04","41.04","41.04","41.04","41.04","41.04","41.04","41.04","41.04"],"quality":"Equal","isMain":false,"metricName":"Reference GC (%)"}]]],"referenceName":"gi\_109289986\_ref\_NC","date":"18 July 2018, Wednesday, 16:05:44","order":[0,1,2,3,4,5,6,7,8,9,10,11,12],"assembliesNames":["ABySS\_127","ABySS\_63","CLC","IDBA\_UD","MEGAHIT","MIRA","SOAPdenovo2","SPAdes","SPAdes\_meta","SPAdes\_sc","SPAdes\_sc\_careful","Velvet","Geneious"]},{"assembliesWithNs":null,"minContig":500,"report":[["Genome statistics",[{"values":["9.421","6.474","0.708","4.198","4.792","23.560","2.879","2.405","12.168","2.192","9.473","9.548","4.243","5.123"],"quality":"More is better","isMain":true,"metricName":"Genome fraction (%)"},{"values":["1.000","1.000","1.000","1.000","1.000","1.015","1.000","1.000","1.000","0.999","1.008","1.008","1.000","1.000"],"quality":"Less is better","isMain":true,"metricName":"Duplication ratio"},{"values":[1094,955,527,1069,1003,1938,865,877,5225,882,4314,4314,891,3816],"quality":"More is better","isMain":true,"metricName":"Largest alignment"},{"values":[7017,4822,527,3126,3568,17808,2144,1791,9063,1632,7110,7166,3160,3816],"quality":"More is better","isMain":true,"metricName":"Total aligned length"},{"values":[908,911,527,911,995,1272,665,177,5225,882,4314,4314,794,3816],"quality":"More is better","isMain":false,"metricName":"NA50"},{"values":[638,837,527,593,988,1053,614,null,967,750,823,823,773,3816],"quality":"More is better","isMain":false,"metricName":"NA75"},{"values":[null,null,null,null,null,null,null,null,null,null,null,null,null,null],"quality":"More is better","isMain":true,"metricName":"NGA50"},{"values":[4,3,1,2,2,6,2,3,1,1,1,1,2,1],"quality":"Less is better","isMain":false,"metricName":"LA50"},{"values":[7,4,1,3,3,10,3,null,3,2,3,3,3,1],"quality":"Less is better","isMain":false,"metricName":"LA75"}]],["Misassemblies",[{"values":[0,0,0,0,0,0,0,0,0,0,0,0,0,0],"quality":"Less is better","isMain":true,"metricName":"# misassemblies"},{"values":[0,0,0,0,0,0,0,0,0,0,0,0,0,0],"quality":"Less is better","isMain":false,"metricName":" # relocations"},{"values":[0,0,0,0,0,0,0,0,0,0,0,0,0,0],"quality":"Less is better","isMain":false,"metricName":" # translocations"},{"values":[0,0,0,0,0,0,0,0,0,0,0,0,0,0],"quality":"Less is better","isMain":false,"metricName":" # inversions"},{"values":[0,0,0,0,0,0,0,0,0,0,0,0,0,0],"quality":"Less is better","isMain":false,"metricName":"# misassembled contigs"},{"values":[0,0,0,0,0,0,0,0,0,0,0,0,0,0],"quality":"Less is better","isMain":true,"metricName":"Misassembled contigs length"},{"values":[0,0,0,0,0,0,0,0,0,0,0,0,0,1],"quality":"Less is better","isMain":false,"metricName":"# local misassemblies"},{"values":[0,0,0,0,0,0,0,0,0,0,0,0,0,0],"quality":"Less is better","isMain":false,"metricName":"# unaligned mis. contigs"}]],["Unaligned",[{"values":[0,0,0,0,0,0,0,0,0,0,0,0,0,0],"quality":"Less is better","isMain":false,"metricName":"# fully unaligned contigs"},{"values":[0,0,0,0,0,0,0,0,0,0,0,0,0,0],"quality":"Less is better","isMain":false,"metricName":"Fully unaligned length"},{"values":[0,0,0,0,0,0,0,1,1,0,0,0,0,0],"quality":"Less is better","isMain":false,"metricName":"# partially unaligned contigs"},{"values":[0,0,0,0,0,0,0,1754,593,0,0,0,0,0],"quality":"Less is better","isMain":false,"metricName":"Partially unaligned length"}]],["Mismatches",[{"values":[0,3,2,5,7,39,1,1,24,6,30,30,5,29],"quality":"Less is better","isMain":false,"metricName":"# mismatches"},{"values":[0,0,0,1,1,3,0,0,0,1,3,3,0,6],"quality":"Less is better","isMain":false,"metricName":"# indels"},{"values":[0,0,0,1,1,18,0,0,0,1,3,3,0,8],"quality":"Less is better","isMain":false,"metricName":"Indels length"},{"values":["0.00","62.21","379.51","159.90","196.13","222.25","46.64","55.83","264.81","367.42","425.17","421.82","158.23","759.96"],"quality":"Less is better","isMain":true,"metricName":"# mismatches per 100 kbp"},{"values":["0.00","0.00","0.00","31.98","28.02","17.10","0.00","0.00","0.00","61.24","42.52","42.18","0.00","157.23"],"quality":"Less is better","isMain":true,"metricName":"# indels per 100 kbp"},{"values":[0,0,0,1,1,2,0,0,0,1,3,3,0,6],"quality":"Less is better","isMain":false,"metricName":" # indels (<= 5 bp)"},{"values":[0,0,0,0,0,1,0,0,0,0,0,0,0,0],"quality":"Less is better","isMain":false,"metricName":" # indels (> 5 bp)"},{"values":[0,0,0,0,0,12,0,448,593,0,0,0,0,71],"quality":"Less is better","isMain":false,"metricName":"# N's"},{"values":["0.00","0.00","0.00","0.00","0.00","67.39","0.00","12637.52","6141.26","0.00","0.00","0.00","0.00","1860.59"],"quality":"Less is better","isMain":true,"metricName":"# N's per 100 kbp"}]],["Statistics without reference",[{"values":[9,6,1,4,4,15,3,3,6,2,5,5,4,1],"quality":"Equal","isMain":true,"metricName":"# contigs"},{"values":[2,0,0,1,1,11,0,1,2,0,1,1,0,1],"quality":"Equal","isMain":false,"metricName":"# contigs (>= 1000 bp)"},{"values":[0,0,0,0,0,0,0,0,1,0,0,0,0,0],"quality":"Equal","isMain":false,"metricName":"# contigs (>= 5000 bp)"},{"values":[0,0,0,0,0,0,0,0,0,0,0,0,0,0],"quality":"Equal","isMain":false,"metricName":"# contigs (>= 10000 bp)"},{"values":[0,0,0,0,0,0,0,0,0,0,0,0,0,0],"quality":"Equal","isMain":false,"metricName":"# contigs (>= 25000 bp)"},{"values":[0,0,0,0,0,0,0,0,0,0,0,0,0,0],"quality":"Equal","isMain":false,"metricName":"# contigs (>= 50000 bp)"},{"values":[1094,955,527,1069,1003,1938,865,1931,5818,882,4319,4319,891,3816],"quality":"More is better","isMain":true,"metricName":"Largest contig"},{"values":[7017,4822,527,3126,3568,17808,2144,3545,9656,1632,7115,7171,3160,3816],"quality":"More is better","isMain":true,"metricName":"Total length"},{"values":[2094,0,0,1069,1003,14672,0,1931,6952,0,4319,4319,0,3816],"quality":"More is better","isMain":true,"metricName":"Total length (>= 1000 bp)"},{"values":[0,0,0,0,0,0,0,0,5818,0,0,0,0,0],"quality":"More is better","isMain":false,"metricName":"Total length (>= 5000 bp)"},{"values":[0,0,0,0,0,0,0,0,0,0,0,0,0,0],"quality":"More is better","isMain":true,"metricName":"Total length (>= 10000 bp)"},{"values":[0,0,0,0,0,0,0,0,0,0,0,0,0,0],"quality":"More is better","isMain":false,"metricName":"Total length (>= 25000 bp)"},{"values":[0,0,0,0,0,0,0,0,0,0,0,0,0,0],"quality":"More is better","isMain":true,"metricName":"Total length (>= 50000 bp)"},{"values":[908,911,527,911,995,1272,665,1931,5818,882,4319,4319,794,3816],"quality":"More is better","isMain":false,"metricName":"N50"},{"values":[638,837,527,593,988,1053,614,877,967,750,823,823,773,3816],"quality":"More is better","isMain":false,"metricName":"N75"},{"values":[4,3,1,2,2,6,2,1,1,1,1,1,2,1],"quality":"Less is better","isMain":false,"metricName":"L50"},{"values":[7,4,1,3,3,10,3,2,3,2,3,3,3,1],"quality":"Less is better","isMain":false,"metricName":"L75"},{"values":["63.18","63.36","70.40","63.15","63.09","63.65","65.44","58.22","63.41","63.79","63.61","63.65","61.11","63.68"],"quality":"Equal","isMain":false,"metricName":"GC (%)"}]],["Predicted genes",[]],["Similarity statistics",[{"values":[0,0,0,0,0,0,0,0,0,0,0,0,0,0],"quality":"Equal","isMain":false,"metricName":"# similar correct contigs"},{"values":[0,0,0,0,0,0,0,0,0,0,0,0,0,0],"quality":"Equal","isMain":false,"metricName":"# similar misassembled blocks"}]],["Reference statistics",[{"values":[74483,74483,74483,74483,74483,74483,74483,74483,74483,74483,74483,74483,74483,74483],"quality":"Equal","isMain":false,"metricName":"Reference length"},{"values":[1,1,1,1,1,1,1,1,1,1,1,1,1,1],"quality":"Equal","isMain":false,"metricName":"Reference fragments"},{"values":["62.94","62.94","62.94","62.94","62.94","62.94","62.94","62.94","62.94","62.94","62.94","62.94","62.94","62.94"],"quality":"Equal","isMain":false,"metricName":"Reference GC (%)"}]]],"referenceName":"gi\_109302753\_ref\_NC","date":"18 July 2018, Wednesday, 16:05:55","order":[0,1,2,3,4,5,6,7,8,9,10,11,12,13],"assembliesNames":["ABySS\_127","ABySS\_63","CLC","IDBA\_UD","MEGAHIT","MIRA","Ray\_Meta","SOAPdenovo2","SPAdes","SPAdes\_meta","SPAdes\_sc","SPAdes\_sc\_careful","Velvet","Geneious"]},{"assembliesWithNs":null,"minContig":500,"report":[["Genome statistics",[{"values":["24.314","3.805","21.030","31.193","29.216","67.785","13.298","25.985","17.178","19.720","21.677","11.577","26.381"],"quality":"More is better","isMain":true,"metricName":"Genome fraction (%)"},{"values":["1.005","1.000","1.011","1.011","1.013","1.132","1.135","1.000","1.031","1.005","1.004","1.004","1.001"],"quality":"Less is better","isMain":true,"metricName":"Duplication ratio"},{"values":[1288,954,1277,1429,1772,4100,1200,3101,4049,1221,1194,1484,5259],"quality":"More is better","isMain":true,"metricName":"Largest alignment"},{"values":[13888,2163,12019,17931,16674,42684,7604,14770,9805,11211,12324,6582,15009],"quality":"More is better","isMain":true,"metricName":"Total aligned length"},{"values":[null,null,null,null,null,1703,null,null,null,null,null,null,null],"quality":"More is better","isMain":false,"metricName":"NG50"},{"values":[null,null,null,null,null,1192,null,null,null,null,null,null,null],"quality":"More is better","isMain":false,"metricName":"NG75"},{"values":[726,639,711,912,813,1410,760,1641,566,970,970,1019,5172],"quality":"More is better","isMain":false,"metricName":"NA50"},{"values":[646,570,609,633,591,1018,608,755,null,732,620,719,2058],"quality":"More is better","isMain":false,"metricName":"NA75"},{"values":[null,null,null,null,null,1357,null,null,null,null,null,null,null],"quality":"More is better","isMain":true,"metricName":"NGA50"},{"values":[null,null,null,null,null,339,null,null,null,null,null,null,null],"quality":"More is better","isMain":false,"metricName":"NGA75"},{"values":[null,null,null,null,null,11,null,null,null,null,null,null,null],"quality":"Less is better","isMain":false,"metricName":"LG50"},{"values":[null,null,null,null,null,21,null,null,null,null,null,null,null],"quality":"Less is better","isMain":false,"metricName":"LG75"},{"values":[8,2,7,8,8,13,5,4,6,6,6,3,2],"quality":"Less is better","isMain":false,"metricName":"LA50"},{"values":[13,3,11,14,14,24,9,8,null,9,10,5,4],"quality":"Less is better","isMain":false,"metricName":"LA75"},{"values":[null,null,null,null,null,15,null,null,null,null,null,null,null],"quality":"Less is better","isMain":true,"metricName":"LGA50"},{"values":[null,null,null,null,null,29,null,null,null,null,null,null,null],"quality":"Less is better","isMain":false,"metricName":"LGA75"}]],["Misassemblies",[{"values":[0,0,0,0,0,0,0,0,0,0,0,0,0],"quality":"Less is better","isMain":true,"metricName":"# misassemblies"},{"values":[0,0,0,0,0,0,0,0,0,0,0,0,0],"quality":"Less is better","isMain":false,"metricName":" # relocations"},{"values":[0,0,0,0,0,0,0,0,0,0,0,0,0],"quality":"Less is better","isMain":false,"metricName":" # translocations"},{"values":[0,0,0,0,0,0,0,0,0,0,0,0,0],"quality":"Less is better","isMain":false,"metricName":" # inversions"},{"values":[0,0,0,0,0,0,0,0,0,0,0,0,0],"quality":"Less is better","isMain":false,"metricName":"# misassembled contigs"},{"values":[0,0,0,0,0,0,0,0,0,0,0,0,0],"quality":"Less is better","isMain":true,"metricName":"Misassembled contigs length"},{"values":[0,0,0,0,0,1,0,1,0,0,0,0,2],"quality":"Less is better","isMain":false,"metricName":"# local misassemblies"},{"values":[0,0,0,0,0,0,0,0,1,0,0,0,0],"quality":"Less is better","isMain":false,"metricName":"# unaligned mis. contigs"}]],["Unaligned",[{"values":[0,0,0,0,0,0,0,0,0,0,0,0,0],"quality":"Less is better","isMain":false,"metricName":"# fully unaligned contigs"},{"values":[0,0,0,0,0,0,0,0,0,0,0,0,0],"quality":"Less is better","isMain":false,"metricName":"Fully unaligned length"},{"values":[0,0,0,0,1,4,1,1,1,0,0,0,3],"quality":"Less is better","isMain":false,"metricName":"# partially unaligned contigs"},{"values":[0,0,0,0,937,7488,576,1613,7876,0,0,0,3896],"quality":"Less is better","isMain":false,"metricName":"Partially unaligned length"}]],["Mismatches",[{"values":[9,0,34,8,54,323,0,82,83,12,13,4,159],"quality":"Less is better","isMain":false,"metricName":"# mismatches"},{"values":[0,0,0,0,2,2,1,10,4,0,0,0,10],"quality":"Less is better","isMain":false,"metricName":"# indels"},{"values":[0,0,0,0,2,2,44,17,10,0,0,0,13],"quality":"Less is better","isMain":false,"metricName":"Indels length"},{"values":["65.11","0.00","284.38","45.11","325.11","838.16","0.00","555.07","849.89","107.04","105.49","60.77","1060.14"],"quality":"Less is better","isMain":true,"metricName":"# mismatches per 100 kbp"},{"values":["0.00","0.00","0.00","0.00","12.04","5.19","13.23","67.69","40.96","0.00","0.00","0.00","66.68"],"quality":"Less is better","isMain":true,"metricName":"# indels per 100 kbp"},{"values":[0,0,0,0,2,2,0,9,4,0,0,0,10],"quality":"Less is better","isMain":false,"metricName":" # indels (<= 5 bp)"},{"values":[0,0,0,0,0,0,1,1,0,0,0,0,0],"quality":"Less is better","isMain":false,"metricName":" # indels (> 5 bp)"},{"values":[0,0,0,0,0,65,886,0,0,0,0,0,372],"quality":"Less is better","isMain":false,"metricName":"# N's"},{"values":["0.00","0.00","0.00","0.00","0.00","127.21","9674.60","0.00","0.00","0.00","0.00","0.00","1967.73"],"quality":"Less is better","isMain":true,"metricName":"# N's per 100 kbp"}]],["Statistics without reference",[{"values":[19,3,16,21,20,29,10,12,6,13,15,7,4],"quality":"Equal","isMain":true,"metricName":"# contigs"},{"values":[3,0,3,7,8,27,2,6,4,5,5,3,4],"quality":"Equal","isMain":false,"metricName":"# contigs (>= 1000 bp)"},{"values":[0,0,0,0,0,0,0,0,1,0,0,0,2],"quality":"Equal","isMain":false,"metricName":"# contigs (>= 5000 bp)"},{"values":[0,0,0,0,0,0,0,0,1,0,0,0,0],"quality":"Equal","isMain":false,"metricName":"# contigs (>= 10000 bp)"},{"values":[0,0,0,0,0,0,0,0,0,0,0,0,0],"quality":"Equal","isMain":false,"metricName":"# contigs (>= 25000 bp)"},{"values":[0,0,0,0,0,0,0,0,0,0,0,0,0],"quality":"Equal","isMain":false,"metricName":"# contigs (>= 50000 bp)"},{"values":[1288,954,1298,1429,1772,4100,1515,4714,10052,1275,1248,1484,6718],"quality":"More is better","isMain":true,"metricName":"Largest contig"},{"values":[13888,2163,12091,17931,17765,51097,9158,16383,17940,11265,12378,6611,18905],"quality":"More is better","isMain":true,"metricName":"Total length"},{"values":[3373,0,3474,8285,10058,49374,2737,12417,16491,5599,5572,3701,18905],"quality":"More is better","isMain":true,"metricName":"Total length (>= 1000 bp)"},{"values":[0,0,0,0,0,0,0,0,10052,0,0,0,12699],"quality":"More is better","isMain":false,"metricName":"Total length (>= 5000 bp)"},{"values":[0,0,0,0,0,0,0,0,10052,0,0,0,0],"quality":"More is better","isMain":true,"metricName":"Total length (>= 10000 bp)"},{"values":[0,0,0,0,0,0,0,0,0,0,0,0,0],"quality":"More is better","isMain":false,"metricName":"Total length (>= 25000 bp)"},{"values":[0,0,0,0,0,0,0,0,0,0,0,0,0],"quality":"More is better","isMain":true,"metricName":"Total length (>= 50000 bp)"},{"values":[726,639,765,912,1064,1834,935,1671,10052,970,970,1019,5981],"quality":"More is better","isMain":false,"metricName":"N50"},{"values":[646,570,609,633,623,1357,760,1054,4049,732,620,719,4148],"quality":"More is better","isMain":false,"metricName":"N75"},{"values":[8,2,6,8,7,10,4,3,1,6,6,3,2],"quality":"Less is better","isMain":false,"metricName":"L50"},{"values":[13,3,11,14,13,18,7,6,2,9,10,5,3],"quality":"Less is better","isMain":false,"metricName":"L75"},{"values":["60.60","61.40","61.20","61.01","61.18","61.63","61.00","61.16","61.32","61.43","61.43","61.90","61.84"],"quality":"Equal","isMain":false,"metricName":"GC (%)"}]],["Predicted genes",[]],["Similarity statistics",[{"values":[0,0,0,0,0,0,0,0,0,0,0,0,0],"quality":"Equal","isMain":false,"metricName":"# similar correct contigs"},{"values":[0,0,0,0,0,0,0,0,0,0,0,0,0],"quality":"Equal","isMain":false,"metricName":"# similar misassembled blocks"}]],["Reference statistics",[{"values":[56852,56852,56852,56852,56852,56852,56852,56852,56852,56852,56852,56852,56852],"quality":"Equal","isMain":false,"metricName":"Reference length"},{"values":[1,1,1,1,1,1,1,1,1,1,1,1,1],"quality":"Equal","isMain":false,"metricName":"Reference fragments"},{"values":["61.53","61.53","61.53","61.53","61.53","61.53","61.53","61.53","61.53","61.53","61.53","61.53","61.53"],"quality":"Equal","isMain":false,"metricName":"Reference GC (%)"}]]],"referenceName":"gi\_109392186\_ref\_NC","date":"18 July 2018, Wednesday, 16:06:05","order":[0,1,2,3,4,5,6,7,8,9,10,11,12],"assembliesNames":["ABySS\_127","ABySS\_63","CLC","IDBA\_UD","MEGAHIT","MIRA","SOAPdenovo2","SPAdes","SPAdes\_meta","SPAdes\_sc","SPAdes\_sc\_careful","Velvet","Geneious"]},{"assembliesWithNs":null,"minContig":500,"report":[["Genome statistics",[{"values":["100.000","93.298","99.604","99.438","95.280","100.000","39.475","11.171","100.000","22.218","88.652","91.882","22.212","100.000"],"quality":"More is better","isMain":true,"metricName":"Genome fraction (%)"},{"values":["1.181","1.012","1.015","1.011","1.011","1.020","1.023","1.196","1.002","1.007","1.001","1.000","1.003","1.004"],"quality":"Less is better","isMain":true,"metricName":"Duplication ratio"},{"values":[67655,9532,12657,17344,11959,70925,1609,566,70781,11333,30166,30684,2168,70925],"quality":"More is better","isMain":true,"metricName":"Largest alignment"},{"values":[83474,66736,71408,70996,68049,72090,28529,7893,70781,15710,62638,64918,15736,70925],"quality":"More is better","isMain":true,"metricName":"Total aligned length"},{"values":[83474,4532,11699,8740,8013,70925,null,null,70781,null,11212,30059,null,70925],"quality":"More is better","isMain":false,"metricName":"NG50"},{"values":[83474,1778,9347,4792,2023,70925,null,null,70781,null,3606,30059,null,70925],"quality":"More is better","isMain":false,"metricName":"NG75"},{"values":[67655,4920,11699,8740,8013,70925,781,null,70781,11333,11212,30059,758,70925],"quality":"More is better","isMain":false,"metricName":"NA50"},{"values":[67655,1782,9347,4792,2061,70925,640,null,70781,2735,6122,30059,593,70925],"quality":"More is better","isMain":false,"metricName":"NA75"},{"values":[67655,4532,11699,8740,8013,70925,null,null,70781,null,11212,30059,null,70925],"quality":"More is better","isMain":true,"metricName":"NGA50"},{"values":[67655,1778,9347,4792,2023,70925,null,null,70781,null,3606,30059,null,70925],"quality":"More is better","isMain":false,"metricName":"NGA75"},{"values":[1,6,3,3,4,1,null,null,1,null,2,2,null,1],"quality":"Less is better","isMain":false,"metricName":"LG50"},{"values":[1,13,5,7,11,1,null,null,1,null,5,2,null,1],"quality":"Less is better","isMain":false,"metricName":"LG75"},{"values":[1,5,3,3,4,1,12,null,1,1,2,2,8,1],"quality":"Less is better","isMain":false,"metricName":"LA50"},{"values":[1,12,5,7,10,1,22,null,1,2,3,2,14,1],"quality":"Less is better","isMain":false,"metricName":"LA75"},{"values":[1,6,3,3,4,1,null,null,1,null,2,2,null,1],"quality":"Less is better","isMain":true,"metricName":"LGA50"},{"values":[1,13,5,7,11,1,null,null,1,null,5,2,null,1],"quality":"Less is better","isMain":false,"metricName":"LGA75"}]],["Misassemblies",[{"values":[1,0,0,0,0,0,0,0,0,0,0,0,0,0],"quality":"Less is better","isMain":true,"metricName":"# misassemblies"},{"values":[1,0,0,0,0,0,0,0,0,0,0,0,0,0],"quality":"Less is better","isMain":false,"metricName":" # relocations"},{"values":[0,0,0,0,0,0,0,0,0,0,0,0,0,0],"quality":"Less is better","isMain":false,"metricName":" # translocations"},{"values":[0,0,0,0,0,0,0,0,0,0,0,0,0,0],"quality":"Less is better","isMain":false,"metricName":" # inversions"},{"values":[1,0,0,0,0,0,0,0,0,0,0,0,0,0],"quality":"Less is better","isMain":false,"metricName":"# misassembled contigs"},{"values":[83474,0,0,0,0,0,0,0,0,0,0,0,0,0],"quality":"Less is better","isMain":true,"metricName":"Misassembled contigs length"},{"values":[0,0,0,0,0,1,0,0,1,1,0,0,0,1],"quality":"Less is better","isMain":false,"metricName":"# local misassemblies"},{"values":[0,0,0,0,0,0,0,8,0,0,0,0,0,0],"quality":"Less is better","isMain":false,"metricName":"# unaligned mis. contigs"}]],["Unaligned",[{"values":[0,0,0,0,0,0,0,0,0,0,0,0,0,0],"quality":"Less is better","isMain":false,"metricName":"# fully unaligned contigs"},{"values":[0,0,0,0,0,0,0,0,0,0,0,0,0,0],"quality":"Less is better","isMain":false,"metricName":"Fully unaligned length"},{"values":[0,0,0,0,0,0,0,10,0,1,0,0,0,0],"quality":"Less is better","isMain":false,"metricName":"# partially unaligned contigs"},{"values":[0,0,0,0,0,0,0,13384,0,2236,0,0,0,0],"quality":"Less is better","isMain":false,"metricName":"Partially unaligned length"}]],["Mismatches",[{"values":[0,400,140,0,14,3,1,3,21,113,216,0,0,0],"quality":"Less is better","isMain":false,"metricName":"# mismatches"},{"values":[0,10,3,0,0,0,0,4,0,15,10,0,0,0],"quality":"Less is better","isMain":false,"metricName":"# indels"},{"values":[0,14,3,0,0,0,0,193,0,23,14,0,0,0],"quality":"Less is better","isMain":false,"metricName":"Indels length"},{"values":["0.00","606.81","198.94","0.00","20.80","4.25","3.59","38.01","29.72","719.84","344.85","0.00","0.00","0.00"],"quality":"Less is better","isMain":true,"metricName":"# mismatches per 100 kbp"},{"values":["0.00","15.17","4.26","0.00","0.00","0.00","0.00","50.68","0.00","95.55","15.97","0.00","0.00","0.00"],"quality":"Less is better","isMain":true,"metricName":"# indels per 100 kbp"},{"values":[0,10,3,0,0,0,0,0,0,15,10,0,0,0],"quality":"Less is better","isMain":false,"metricName":" # indels (<= 5 bp)"},{"values":[0,0,0,0,0,0,0,4,0,0,0,0,0,0],"quality":"Less is better","isMain":false,"metricName":" # indels (> 5 bp)"},{"values":[0,5,0,0,0,0,0,7800,0,104,0,0,0,3],"quality":"Less is better","isMain":false,"metricName":"# N's"},{"values":["0.00","7.49","0.00","0.00","0.00","0.00","0.00","34170.06","0.00","576.18","0.00","0.00","0.00","4.23"],"quality":"Less is better","isMain":true,"metricName":"# N's per 100 kbp"}]],["Statistics without reference",[{"values":[1,27,11,13,23,3,33,18,1,3,9,3,21,1],"quality":"Equal","isMain":true,"metricName":"# contigs"},{"values":[1,18,11,12,18,1,10,10,1,3,9,3,1,1],"quality":"Equal","isMain":false,"metricName":"# contigs (>= 1000 bp)"},{"values":[1,4,6,5,4,1,0,0,1,1,3,2,0,1],"quality":"Equal","isMain":false,"metricName":"# contigs (>= 5000 bp)"},{"values":[1,0,3,2,1,1,0,0,1,1,2,2,0,1],"quality":"Equal","isMain":false,"metricName":"# contigs (>= 10000 bp)"},{"values":[1,0,0,0,0,1,0,0,1,0,1,2,0,1],"quality":"Equal","isMain":false,"metricName":"# contigs (>= 25000 bp)"},{"values":[1,0,0,0,0,1,0,0,1,0,0,0,0,1],"quality":"Equal","isMain":false,"metricName":"# contigs (>= 50000 bp)"},{"values":[83474,9532,12657,17344,11959,70925,1609,3173,70781,13569,30166,30684,2168,70925],"quality":"More is better","isMain":true,"metricName":"Largest contig"},{"values":[83474,66736,71408,70996,68049,72090,28536,22827,70781,18050,62672,64918,15736,70925],"quality":"More is better","isMain":true,"metricName":"Total length"},{"values":[83474,60127,71408,70146,64515,70925,13275,18385,70781,18050,62672,64918,2168,70925],"quality":"More is better","isMain":true,"metricName":"Total length (>= 1000 bp)"},{"values":[83474,28451,61992,47156,36176,70925,0,0,70781,13569,47508,60743,0,70925],"quality":"More is better","isMain":false,"metricName":"Total length (>= 5000 bp)"},{"values":[83474,0,36905,27887,11959,70925,0,0,70781,13569,41378,60743,0,70925],"quality":"More is better","isMain":true,"metricName":"Total length (>= 10000 bp)"},{"values":[83474,0,0,0,0,70925,0,0,70781,0,30166,60743,0,70925],"quality":"More is better","isMain":false,"metricName":"Total length (>= 25000 bp)"},{"values":[83474,0,0,0,0,70925,0,0,70781,0,0,0,0,70925],"quality":"More is better","isMain":true,"metricName":"Total length (>= 50000 bp)"},{"values":[83474,4920,11699,8740,8013,70925,781,1601,70781,13569,11212,30059,758,70925],"quality":"More is better","isMain":false,"metricName":"N50"},{"values":[83474,1782,9347,4792,2061,70925,640,1405,70781,13569,6130,30059,593,70925],"quality":"More is better","isMain":false,"metricName":"N75"},{"values":[1,5,3,3,4,1,12,6,1,1,2,2,8,1],"quality":"Less is better","isMain":false,"metricName":"L50"},{"values":[1,12,5,7,10,1,22,9,1,1,3,2,14,1],"quality":"Less is better","isMain":false,"metricName":"L75"},{"values":["69.06","69.05","69.09","69.10","69.10","69.10","69.12","69.90","69.10","69.56","69.22","69.13","69.27","69.09"],"quality":"Equal","isMain":false,"metricName":"GC (%)"}]],["Predicted genes",[]],["Similarity statistics",[{"values":[0,0,0,0,0,0,0,0,0,0,0,0,0,0],"quality":"Equal","isMain":false,"metricName":"# similar correct contigs"},{"values":[0,0,0,0,0,0,0,0,0,0,0,0,0,0],"quality":"Equal","isMain":false,"metricName":"# similar misassembled blocks"}]],["Reference statistics",[{"values":[70654,70654,70654,70654,70654,70654,70654,70654,70654,70654,70654,70654,70654,70654],"quality":"Equal","isMain":false,"metricName":"Reference length"},{"values":[1,1,1,1,1,1,1,1,1,1,1,1,1,1],"quality":"Equal","isMain":false,"metricName":"Reference fragments"},{"values":["69.11","69.11","69.11","69.11","69.11","69.11","69.11","69.11","69.11","69.11","69.11","69.11","69.11","69.11"],"quality":"Equal","isMain":false,"metricName":"Reference GC (%)"}]]],"referenceName":"gi\_109392353\_ref\_NC","date":"18 July 2018, Wednesday, 16:06:16","order":[0,1,2,3,4,5,6,7,8,9,10,11,12,13],"assembliesNames":["ABySS\_127","ABySS\_63","CLC","IDBA\_UD","MEGAHIT","MIRA","Ray\_Meta","SOAPdenovo2","SPAdes","SPAdes\_meta","SPAdes\_sc","SPAdes\_sc\_careful","Velvet","Geneious"]},{"assembliesWithNs":null,"minContig":500,"report":[["Genome statistics",[{"values":["98.286","100.000","100.000","99.871","100.000","94.503","7.334","100.000","99.318","99.303","98.611","63.216","100.000"],"quality":"More is better","isMain":true,"metricName":"Genome fraction (%)"},{"values":["1.000","1.011","1.006","1.000","1.023","1.058","1.000","1.002","1.005","1.002","1.000","1.008","1.006"],"quality":"Less is better","isMain":true,"metricName":"Duplication ratio"},{"values":[39636,29441,47448,51980,52342,16781,801,52174,51745,40665,40860,3212,52362],"quality":"More is better","isMain":true,"metricName":"Largest alignment"},{"values":[51155,52636,52344,51980,53242,51553,3817,52174,51745,51794,51324,33139,52362],"quality":"More is better","isMain":true,"metricName":"Total aligned length"},{"values":[39636,29441,47448,51980,52342,8017,null,52174,51942,40665,40860,772,52364],"quality":"More is better","isMain":false,"metricName":"NG50"},{"values":[39636,12158,47448,51980,52342,3644,null,52174,51942,40665,40860,null,52364],"quality":"More is better","isMain":false,"metricName":"NG75"},{"values":[39636,29441,47448,51980,52342,8017,627,52174,51745,40665,40860,1107,52362],"quality":"More is better","isMain":false,"metricName":"NA50"},{"values":[39636,12158,47448,51980,52342,3165,591,52174,51745,40665,40860,776,52362],"quality":"More is better","isMain":false,"metricName":"NA75"},{"values":[39636,29441,47448,51980,52342,8017,null,52174,51745,40665,40860,772,52362],"quality":"More is better","isMain":true,"metricName":"NGA50"},{"values":[39636,12158,47448,51980,52342,3165,null,52174,51745,40665,40860,null,52362],"quality":"More is better","isMain":false,"metricName":"NGA75"},{"values":[1,1,1,1,1,3,null,1,1,1,1,20,1],"quality":"Less is better","isMain":false,"metricName":"LG50"},{"values":[1,2,1,1,1,5,null,1,1,1,1,null,1],"quality":"Less is better","isMain":false,"metricName":"LG75"},{"values":[1,1,1,1,1,3,3,1,1,1,1,10,1],"quality":"Less is better","isMain":false,"metricName":"LA50"},{"values":[1,2,1,1,1,5,5,1,1,1,1,19,1],"quality":"Less is better","isMain":false,"metricName":"LA75"},{"values":[1,1,1,1,1,3,null,1,1,1,1,20,1],"quality":"Less is better","isMain":true,"metricName":"LGA50"},{"values":[1,2,1,1,1,5,null,1,1,1,1,null,1],"quality":"Less is better","isMain":false,"metricName":"LGA75"}]],["Misassemblies",[{"values":[0,0,0,0,0,0,0,0,0,0,0,0,0],"quality":"Less is better","isMain":true,"metricName":"# misassemblies"},{"values":[0,0,0,0,0,0,0,0,0,0,0,0,0],"quality":"Less is better","isMain":false,"metricName":" # relocations"},{"values":[0,0,0,0,0,0,0,0,0,0,0,0,0],"quality":"Less is better","isMain":false,"metricName":" # translocations"},{"values":[0,0,0,0,0,0,0,0,0,0,0,0,0],"quality":"Less is better","isMain":false,"metricName":" # inversions"},{"values":[0,0,0,0,0,0,0,0,0,0,0,0,0],"quality":"Less is better","isMain":false,"metricName":"# misassembled contigs"},{"values":[0,0,0,0,0,0,0,0,0,0,0,0,0],"quality":"Less is better","isMain":true,"metricName":"Misassembled contigs length"},{"values":[0,0,0,0,1,0,0,1,1,0,0,0,1],"quality":"Less is better","isMain":false,"metricName":"# local misassemblies"},{"values":[0,0,0,0,0,0,0,0,0,0,0,0,0],"quality":"Less is better","isMain":false,"metricName":"# unaligned mis. contigs"}]],["Unaligned",[{"values":[0,0,0,0,0,0,0,0,0,0,0,0,0],"quality":"Less is better","isMain":false,"metricName":"# fully unaligned contigs"},{"values":[0,0,0,0,0,0,0,0,0,0,0,0,0],"quality":"Less is better","isMain":false,"metricName":"Fully unaligned length"},{"values":[0,0,0,0,0,0,0,0,0,0,0,0,0],"quality":"Less is better","isMain":false,"metricName":"# partially unaligned contigs"},{"values":[0,0,0,0,0,0,0,0,0,0,0,0,0],"quality":"Less is better","isMain":false,"metricName":"Partially unaligned length"}]],["Mismatches",[{"values":[31,0,2,1,1,4,0,0,190,31,0,1,0],"quality":"Less is better","isMain":false,"metricName":"# mismatches"},{"values":[0,0,0,0,0,0,0,0,9,0,0,0,1],"quality":"Less is better","isMain":false,"metricName":"# indels"},{"values":[0,0,0,0,0,0,0,0,10,0,0,0,1],"quality":"Less is better","isMain":false,"metricName":"Indels length"},{"values":["60.60","0.00","3.84","1.92","1.92","8.13","0.00","0.00","367.56","59.98","0.00","3.04","0.00"],"quality":"Less is better","isMain":true,"metricName":"# mismatches per 100 kbp"},{"values":["0.00","0.00","0.00","0.00","0.00","0.00","0.00","0.00","17.41","0.00","0.00","0.00","1.92"],"quality":"Less is better","isMain":true,"metricName":"# indels per 100 kbp"},{"values":[0,0,0,0,0,0,0,0,9,0,0,0,1],"quality":"Less is better","isMain":false,"metricName":" # indels (<= 5 bp)"},{"values":[0,0,0,0,0,0,0,0,0,0,0,0,0],"quality":"Less is better","isMain":false,"metricName":" # indels (> 5 bp)"},{"values":[0,0,0,0,2,0,0,0,0,0,0,0,446],"quality":"Less is better","isMain":false,"metricName":"# N's"},{"values":["0.00","0.00","0.00","0.00","3.76","0.00","0.00","0.00","0.00","0.00","0.00","0.00","851.73"],"quality":"Less is better","isMain":true,"metricName":"# N's per 100 kbp"}]],["Statistics without reference",[{"values":[3,4,3,1,2,11,6,1,1,3,2,31,1],"quality":"Equal","isMain":true,"metricName":"# contigs"},{"values":[3,3,2,1,1,10,0,1,1,2,2,12,1],"quality":"Equal","isMain":false,"metricName":"# contigs (>= 1000 bp)"},{"values":[2,3,1,1,1,3,0,1,1,2,2,0,1],"quality":"Equal","isMain":false,"metricName":"# contigs (>= 5000 bp)"},{"values":[2,3,1,1,1,1,0,1,1,2,2,0,1],"quality":"Equal","isMain":false,"metricName":"# contigs (>= 10000 bp)"},{"values":[1,1,1,1,1,0,0,1,1,1,1,0,1],"quality":"Equal","isMain":false,"metricName":"# contigs (>= 25000 bp)"},{"values":[0,0,0,1,1,0,0,1,1,0,0,0,1],"quality":"Equal","isMain":false,"metricName":"# contigs (>= 50000 bp)"},{"values":[39636,29441,47448,51980,52342,16781,801,52174,51942,40665,40860,3212,52364],"quality":"More is better","isMain":true,"metricName":"Largest contig"},{"values":[51155,52636,52344,51980,53242,52032,3817,52174,51942,51794,51324,33169,52364],"quality":"More is better","isMain":true,"metricName":"Total length"},{"values":[51155,52113,51696,51980,52342,51145,0,52174,51942,51129,51324,19333,52364],"quality":"More is better","isMain":true,"metricName":"Total length (>= 1000 bp)"},{"values":[49816,52113,47448,51980,52342,34023,0,52174,51942,51129,51324,0,52364],"quality":"More is better","isMain":false,"metricName":"Total length (>= 5000 bp)"},{"values":[49816,52113,47448,51980,52342,16781,0,52174,51942,51129,51324,0,52364],"quality":"More is better","isMain":true,"metricName":"Total length (>= 10000 bp)"},{"values":[39636,29441,47448,51980,52342,0,0,52174,51942,40665,40860,0,52364],"quality":"More is better","isMain":false,"metricName":"Total length (>= 25000 bp)"},{"values":[0,0,0,51980,52342,0,0,52174,51942,0,0,0,52364],"quality":"More is better","isMain":true,"metricName":"Total length (>= 50000 bp)"},{"values":[39636,29441,47448,51980,52342,8017,627,52174,51942,40665,40860,1107,52364],"quality":"More is better","isMain":false,"metricName":"N50"},{"values":[39636,12158,47448,51980,52342,3644,591,52174,51942,40665,40860,776,52364],"quality":"More is better","isMain":false,"metricName":"N75"},{"values":[1,1,1,1,1,3,3,1,1,1,1,10,1],"quality":"Less is better","isMain":false,"metricName":"L50"},{"values":[1,2,1,1,1,5,5,1,1,1,1,19,1],"quality":"Less is better","isMain":false,"metricName":"L75"},{"values":["62.88","62.84","62.88","62.88","62.85","62.78","63.82","62.88","62.92","62.89","62.88","62.80","62.86"],"quality":"Equal","isMain":false,"metricName":"GC (%)"}]],["Predicted genes",[]],["Similarity statistics",[{"values":[0,0,0,0,0,0,0,0,0,0,0,0,0],"quality":"Equal","isMain":false,"metricName":"# similar correct contigs"},{"values":[0,0,0,0,0,0,0,0,0,0,0,0,0],"quality":"Equal","isMain":false,"metricName":"# similar misassembled blocks"}]],["Reference statistics",[{"values":[52047,52047,52047,52047,52047,52047,52047,52047,52047,52047,52047,52047,52047],"quality":"Equal","isMain":false,"metricName":"Reference length"},{"values":[1,1,1,1,1,1,1,1,1,1,1,1,1],"quality":"Equal","isMain":false,"metricName":"Reference fragments"},{"values":["62.88","62.88","62.88","62.88","62.88","62.88","62.88","62.88","62.88","62.88","62.88","62.88","62.88"],"quality":"Equal","isMain":false,"metricName":"Reference GC (%)"}]]],"referenceName":"gi\_109392453\_ref\_NC","date":"18 July 2018, Wednesday, 16:06:26","order":[0,1,2,3,4,5,6,7,8,9,10,11,12],"assembliesNames":["ABySS\_63","CLC","IDBA\_UD","MEGAHIT","MIRA","Ray\_Meta","SOAPdenovo2","SPAdes","SPAdes\_meta","SPAdes\_sc","SPAdes\_sc\_careful","Velvet","Geneious"]},{"assembliesWithNs":null,"minContig":500,"report":[["Genome statistics",[{"values":["11.624","4.173","9.822","8.069","14.425","32.209","3.612","4.730","11.724","2.331","12.500","5.592","4.491","9.779"],"quality":"More is better","isMain":true,"metricName":"Genome fraction (%)"},{"values":["1.014","1.006","1.002","1.016","1.043","1.039","1.001","1.112","1.021","1.000","1.037","1.000","1.000","1.000"],"quality":"Less is better","isMain":true,"metricName":"Duplication ratio"},{"values":[1468,948,1944,3369,2668,6575,1052,1012,2778,868,10440,1012,3030,6738],"quality":"More is better","isMain":true,"metricName":"Largest alignment"},{"values":[18125,6454,15124,12607,23126,51466,5554,7273,18408,3584,19495,8598,6906,15036],"quality":"More is better","isMain":true,"metricName":"Total aligned length"},{"values":[798,722,1254,1127,1012,1908,986,543,1353,738,10440,738,824,5846],"quality":"More is better","isMain":false,"metricName":"NA50"},{"values":[612,625,820,672,641,1106,879,387,774,600,639,631,606,5846],"quality":"More is better","isMain":false,"metricName":"NA75"},{"values":[null,null,null,null,null,null,null,null,null,null,null,null,null,null],"quality":"More is better","isMain":true,"metricName":"NGA50"},{"values":[9,4,5,4,7,8,3,7,6,3,1,5,2,2],"quality":"Less is better","isMain":false,"metricName":"LA50"},{"values":[15,7,9,7,13,17,5,11,10,4,7,9,5,2],"quality":"Less is better","isMain":false,"metricName":"LA75"}]],["Misassemblies",[{"values":[0,0,0,0,0,0,0,0,0,0,0,0,0,0],"quality":"Less is better","isMain":true,"metricName":"# misassemblies"},{"values":[0,0,0,0,0,0,0,0,0,0,0,0,0,0],"quality":"Less is better","isMain":false,"metricName":" # relocations"},{"values":[0,0,0,0,0,0,0,0,0,0,0,0,0,0],"quality":"Less is better","isMain":false,"metricName":" # translocations"},{"values":[0,0,0,0,0,0,0,0,0,0,0,0,0,0],"quality":"Less is better","isMain":false,"metricName":" # inversions"},{"values":[0,0,0,0,0,0,0,0,0,0,0,0,0,0],"quality":"Less is better","isMain":false,"metricName":"# misassembled contigs"},{"values":[0,0,0,0,0,0,0,0,0,0,0,0,0,0],"quality":"Less is better","isMain":true,"metricName":"Misassembled contigs length"},{"values":[0,0,0,0,0,0,0,0,0,0,0,0,0,0],"quality":"Less is better","isMain":false,"metricName":"# local misassemblies"},{"values":[0,0,0,0,0,0,0,0,0,0,0,0,0,0],"quality":"Less is better","isMain":false,"metricName":"# unaligned mis. contigs"}]],["Unaligned",[{"values":[0,0,0,0,0,0,0,0,0,0,0,0,0,0],"quality":"Less is better","isMain":false,"metricName":"# fully unaligned contigs"},{"values":[0,0,0,0,0,0,0,0,0,0,0,0,0,0],"quality":"Less is better","isMain":false,"metricName":"Fully unaligned length"},{"values":[0,0,0,0,0,0,0,1,0,0,0,0,0,0],"quality":"Less is better","isMain":false,"metricName":"# partially unaligned contigs"},{"values":[0,0,0,0,0,0,0,654,0,0,0,0,0,0],"quality":"Less is better","isMain":false,"metricName":"Partially unaligned length"}]],["Mismatches",[{"values":[4,1,40,27,63,97,28,0,20,11,95,1,7,52],"quality":"Less is better","isMain":false,"metricName":"# mismatches"},{"values":[0,0,1,1,19,5,0,2,0,0,6,0,0,4],"quality":"Less is better","isMain":false,"metricName":"# indels"},{"values":[0,0,1,1,60,7,0,102,0,0,6,0,0,7],"quality":"Less is better","isMain":false,"metricName":"Indels length"},{"values":["22.38","15.59","264.85","217.60","284.03","195.85","504.14","0.00","110.94","306.92","494.28","11.63","101.36","345.81"],"quality":"Less is better","isMain":true,"metricName":"# mismatches per 100 kbp"},{"values":["0.00","0.00","6.62","8.06","85.66","10.10","0.00","27.50","0.00","0.00","31.22","0.00","0.00","26.60"],"quality":"Less is better","isMain":true,"metricName":"# indels per 100 kbp"},{"values":[0,0,1,1,17,5,0,0,0,0,6,0,0,4],"quality":"Less is better","isMain":false,"metricName":" # indels (<= 5 bp)"},{"values":[0,0,0,0,2,0,0,2,0,0,0,0,0,0],"quality":"Less is better","isMain":false,"metricName":" # indels (> 5 bp)"},{"values":[0,0,0,0,0,38,0,831,0,0,435,0,0,71],"quality":"Less is better","isMain":false,"metricName":"# N's"},{"values":["0.00","0.00","0.00","0.00","0.00","73.83","0.00","9504.75","0.00","0.00","2182.64","0.00","0.00","472.20"],"quality":"Less is better","isMain":true,"metricName":"# N's per 100 kbp"}]],["Statistics without reference",[{"values":[23,9,14,12,23,33,6,14,17,5,14,12,7,3],"quality":"Equal","isMain":true,"metricName":"# contigs"},{"values":[4,0,8,4,8,22,2,1,8,0,2,1,1,3],"quality":"Equal","isMain":false,"metricName":"# contigs (>= 1000 bp)"},{"values":[0,0,0,0,0,1,0,0,0,0,1,0,0,2],"quality":"Equal","isMain":false,"metricName":"# contigs (>= 5000 bp)"},{"values":[0,0,0,0,0,0,0,0,0,0,1,0,0,0],"quality":"Equal","isMain":false,"metricName":"# contigs (>= 10000 bp)"},{"values":[0,0,0,0,0,0,0,0,0,0,0,0,0,0],"quality":"Equal","isMain":false,"metricName":"# contigs (>= 25000 bp)"},{"values":[0,0,0,0,0,0,0,0,0,0,0,0,0,0],"quality":"Equal","isMain":false,"metricName":"# contigs (>= 50000 bp)"},{"values":[1468,948,1946,3369,2668,6575,1058,1012,2778,868,10875,1012,3030,6738],"quality":"More is better","isMain":true,"metricName":"Largest contig"},{"values":[18125,6454,15126,12607,23126,51467,5560,8743,18408,3584,19930,8598,6906,15036],"quality":"More is better","isMain":true,"metricName":"Total length"},{"values":[5220,0,11346,7259,13523,43904,2093,1012,12758,0,11887,1012,3030,15036],"quality":"More is better","isMain":true,"metricName":"Total length (>= 1000 bp)"},{"values":[0,0,0,0,0,6575,0,0,0,0,10875,0,0,12584],"quality":"More is better","isMain":false,"metricName":"Total length (>= 5000 bp)"},{"values":[0,0,0,0,0,0,0,0,0,0,10875,0,0,0],"quality":"More is better","isMain":true,"metricName":"Total length (>= 10000 bp)"},{"values":[0,0,0,0,0,0,0,0,0,0,0,0,0,0],"quality":"More is better","isMain":false,"metricName":"Total length (>= 25000 bp)"},{"values":[0,0,0,0,0,0,0,0,0,0,0,0,0,0],"quality":"More is better","isMain":true,"metricName":"Total length (>= 50000 bp)"},{"values":[798,722,1254,1127,1012,1908,986,558,1353,738,10875,738,824,5846],"quality":"More is better","isMain":false,"metricName":"N50"},{"values":[612,625,1012,672,641,1106,879,540,774,600,738,631,606,5846],"quality":"More is better","isMain":false,"metricName":"N75"},{"values":[9,4,5,4,7,8,3,6,6,3,1,5,2,2],"quality":"Less is better","isMain":false,"metricName":"L50"},{"values":[15,7,8,7,13,17,5,10,10,4,6,9,5,2],"quality":"Less is better","isMain":false,"metricName":"L75"},{"values":["64.88","63.99","64.86","64.67","65.21","64.80","64.68","64.83","64.97","62.64","64.57","64.48","65.90","66.52"],"quality":"Equal","isMain":false,"metricName":"GC (%)"}]],["Predicted genes",[]],["Similarity statistics",[{"values":[0,0,0,0,0,0,0,0,0,0,0,0,0,0],"quality":"Equal","isMain":false,"metricName":"# similar correct contigs"},{"values":[0,0,0,0,0,0,0,0,0,0,0,0,0,0],"quality":"Equal","isMain":false,"metricName":"# similar misassembled blocks"}]],["Reference statistics",[{"values":[153766,153766,153766,153766,153766,153766,153766,153766,153766,153766,153766,153766,153766,153766],"quality":"Equal","isMain":false,"metricName":"Reference length"},{"values":[1,1,1,1,1,1,1,1,1,1,1,1,1,1],"quality":"Equal","isMain":false,"metricName":"Reference fragments"},{"values":["64.74","64.74","64.74","64.74","64.74","64.74","64.74","64.74","64.74","64.74","64.74","64.74","64.74","64.74"],"quality":"Equal","isMain":false,"metricName":"Reference GC (%)"}]]],"referenceName":"gi\_109393216\_ref\_NC","date":"18 July 2018, Wednesday, 16:06:37","order":[0,1,2,3,4,5,6,7,8,9,10,11,12,13],"assembliesNames":["ABySS\_127","ABySS\_63","CLC","IDBA\_UD","MEGAHIT","MIRA","Ray\_Meta","SOAPdenovo2","SPAdes","SPAdes\_meta","SPAdes\_sc","SPAdes\_sc\_careful","Velvet","Geneious"]},{"assembliesWithNs":null,"minContig":500,"report":[["Genome statistics",[{"values":["27.478","20.725","88.287","80.430","80.009","47.841","83.747","87.876","88.590","88.603","88.603","17.384","10.510"],"quality":"More is better","isMain":true,"metricName":"Genome fraction (%)"},{"values":["1.001","1.000","1.001","1.000","1.000","1.008","1.003","1.001","1.001","1.001","1.001","1.002","1.000"],"quality":"Less is better","isMain":true,"metricName":"Duplication ratio"},{"values":[1743,1220,6874,5107,6705,3855,4851,6859,7004,6859,6859,5008,5155],"quality":"More is better","isMain":true,"metricName":"Largest alignment"},{"values":[21554,16257,69354,63108,62760,37825,65789,68989,69549,69559,69559,13636,8244],"quality":"More is better","isMain":true,"metricName":"Total aligned length"},{"values":[null,null,1844,1649,1655,null,1530,1993,1993,1993,1993,null,null],"quality":"More is better","isMain":false,"metricName":"NG50"},{"values":[null,null,926,717,658,null,690,1308,1153,1308,1308,null,null],"quality":"More is better","isMain":false,"metricName":"NG75"},{"values":[849,733,2057,1844,1861,1358,1727,2063,2063,2063,2063,2385,5155],"quality":"More is better","isMain":false,"metricName":"NA50"},{"values":[770,629,1474,1404,1381,974,1012,1602,1602,1602,1602,957,3089],"quality":"More is better","isMain":false,"metricName":"NA75"},{"values":[null,null,1844,1649,1655,null,1530,1993,1993,1993,1993,null,null],"quality":"More is better","isMain":true,"metricName":"NGA50"},{"values":[null,null,926,717,658,null,686,1308,1153,1308,1308,null,null],"quality":"More is better","isMain":false,"metricName":"NGA75"},{"values":[null,null,13,17,15,null,16,13,13,13,13,null,null],"quality":"Less is better","isMain":false,"metricName":"LG50"},{"values":[null,null,26,34,33,null,35,25,25,25,25,null,null],"quality":"Less is better","isMain":false,"metricName":"LG75"},{"values":[9,9,11,13,11,10,13,11,11,11,11,2,1],"quality":"Less is better","isMain":false,"metricName":"LA50"},{"values":[16,15,21,22,21,18,24,20,20,20,20,5,2],"quality":"Less is better","isMain":false,"metricName":"LA75"},{"values":[null,null,13,17,15,null,16,13,13,13,13,null,null],"quality":"Less is better","isMain":true,"metricName":"LGA50"},{"values":[null,null,26,34,33,null,36,25,25,25,25,null,null],"quality":"Less is better","isMain":false,"metricName":"LGA75"}]],["Misassemblies",[{"values":[0,0,0,0,0,0,0,0,0,0,0,0,0],"quality":"Less is better","isMain":true,"metricName":"# misassemblies"},{"values":[0,0,0,0,0,0,0,0,0,0,0,0,0],"quality":"Less is better","isMain":false,"metricName":" # relocations"},{"values":[0,0,0,0,0,0,0,0,0,0,0,0,0],"quality":"Less is better","isMain":false,"metricName":" # translocations"},{"values":[0,0,0,0,0,0,0,0,0,0,0,0,0],"quality":"Less is better","isMain":false,"metricName":" # inversions"},{"values":[0,0,0,0,0,0,0,0,0,0,0,0,0],"quality":"Less is better","isMain":false,"metricName":"# misassembled contigs"},{"values":[0,0,0,0,0,0,0,0,0,0,0,0,0],"quality":"Less is better","isMain":true,"metricName":"Misassembled contigs length"},{"values":[0,0,0,0,1,0,0,1,0,1,1,0,0],"quality":"Less is better","isMain":false,"metricName":"# local misassemblies"},{"values":[0,0,0,0,0,0,0,0,0,0,0,0,0],"quality":"Less is better","isMain":false,"metricName":"# unaligned mis. contigs"}]],["Unaligned",[{"values":[0,0,0,0,0,0,0,0,0,0,0,0,0],"quality":"Less is better","isMain":false,"metricName":"# fully unaligned contigs"},{"values":[0,0,0,0,0,0,0,0,0,0,0,0,0],"quality":"Less is better","isMain":false,"metricName":"Fully unaligned length"},{"values":[0,0,0,0,0,0,0,0,0,0,0,0,0],"quality":"Less is better","isMain":false,"metricName":"# partially unaligned contigs"},{"values":[0,0,0,0,0,0,0,0,0,0,0,0,0],"quality":"Less is better","isMain":false,"metricName":"Partially unaligned length"}]],["Mismatches",[{"values":[2,0,55,33,30,10,42,73,60,52,35,3,0],"quality":"Less is better","isMain":false,"metricName":"# mismatches"},{"values":[0,0,0,0,0,0,1,0,0,0,0,0,0],"quality":"Less is better","isMain":false,"metricName":"# indels"},{"values":[0,0,0,0,0,0,57,0,0,0,0,0,0],"quality":"Less is better","isMain":false,"metricName":"Indels length"},{"values":["9.28","0.00","79.42","52.31","47.80","26.65","63.93","105.90","86.34","74.82","50.36","22.00","0.00"],"quality":"Less is better","isMain":true,"metricName":"# mismatches per 100 kbp"},{"values":["0.00","0.00","0.00","0.00","0.00","0.00","1.52","0.00","0.00","0.00","0.00","0.00","0.00"],"quality":"Less is better","isMain":true,"metricName":"# indels per 100 kbp"},{"values":[0,0,0,0,0,0,0,0,0,0,0,0,0],"quality":"Less is better","isMain":false,"metricName":" # indels (<= 5 bp)"},{"values":[0,0,0,0,0,0,1,0,0,0,0,0,0],"quality":"Less is better","isMain":false,"metricName":" # indels (> 5 bp)"},{"values":[0,0,0,0,0,42,107,0,0,0,0,0,4],"quality":"Less is better","isMain":false,"metricName":"# N's"},{"values":["0.00","0.00","0.00","0.00","0.00","111.04","162.32","0.00","0.00","0.00","0.00","0.00","48.52"],"quality":"Less is better","isMain":true,"metricName":"# N's per 100 kbp"}]],["Statistics without reference",[{"values":[24,22,40,40,39,30,47,37,38,38,38,9,2],"quality":"Equal","isMain":true,"metricName":"# contigs"},{"values":[7,3,25,27,26,17,24,26,26,26,26,4,2],"quality":"Equal","isMain":false,"metricName":"# contigs (>= 1000 bp)"},{"values":[0,0,2,1,1,0,0,2,2,2,2,1,1],"quality":"Equal","isMain":false,"metricName":"# contigs (>= 5000 bp)"},{"values":[0,0,0,0,0,0,0,0,0,0,0,0,0],"quality":"Equal","isMain":false,"metricName":"# contigs (>= 10000 bp)"},{"values":[0,0,0,0,0,0,0,0,0,0,0,0,0],"quality":"Equal","isMain":false,"metricName":"# contigs (>= 25000 bp)"},{"values":[0,0,0,0,0,0,0,0,0,0,0,0,0],"quality":"Equal","isMain":false,"metricName":"# contigs (>= 50000 bp)"},{"values":[1743,1220,6874,5107,6705,3855,4874,6859,7004,6859,6859,5038,5155],"quality":"More is better","isMain":true,"metricName":"Largest contig"},{"values":[21583,16257,69354,63108,62760,37825,65919,68989,69549,69559,69559,13666,8244],"quality":"More is better","isMain":true,"metricName":"Total length"},{"values":[9302,3325,58209,53401,54050,27884,49856,60622,60612,60622,60622,10101,8244],"quality":"More is better","isMain":true,"metricName":"Total length (>= 1000 bp)"},{"values":[0,0,12150,5107,6705,0,0,12135,12280,12135,12135,5038,5155],"quality":"More is better","isMain":false,"metricName":"Total length (>= 5000 bp)"},{"values":[0,0,0,0,0,0,0,0,0,0,0,0,0],"quality":"More is better","isMain":true,"metricName":"Total length (>= 10000 bp)"},{"values":[0,0,0,0,0,0,0,0,0,0,0,0,0],"quality":"More is better","isMain":false,"metricName":"Total length (>= 25000 bp)"},{"values":[0,0,0,0,0,0,0,0,0,0,0,0,0],"quality":"More is better","isMain":true,"metricName":"Total length (>= 50000 bp)"},{"values":[849,733,2057,1844,1861,1358,1727,2063,2063,2063,2063,2385,5155],"quality":"More is better","isMain":false,"metricName":"N50"},{"values":[770,629,1474,1404,1381,974,1012,1602,1602,1602,1602,957,3089],"quality":"More is better","isMain":false,"metricName":"N75"},{"values":[9,9,11,13,11,10,13,11,11,11,11,2,1],"quality":"Less is better","isMain":false,"metricName":"L50"},{"values":[16,15,21,22,21,18,24,20,20,20,20,5,2],"quality":"Less is better","isMain":false,"metricName":"L75"},{"values":["57.65","57.53","56.82","56.81","56.80","57.20","56.88","56.81","56.81","56.82","56.82","58.02","56.98"],"quality":"Equal","isMain":false,"metricName":"GC (%)"}]],["Predicted genes",[]],["Similarity statistics",[{"values":[0,0,0,0,0,0,0,0,0,0,0,0,0],"quality":"Equal","isMain":false,"metricName":"# similar correct contigs"},{"values":[0,0,0,0,0,0,0,0,0,0,0,0,0],"quality":"Equal","isMain":false,"metricName":"# similar misassembled blocks"}]],["Reference statistics",[{"values":[78441,78441,78441,78441,78441,78441,78441,78441,78441,78441,78441,78441,78441],"quality":"Equal","isMain":false,"metricName":"Reference length"},{"values":[1,1,1,1,1,1,1,1,1,1,1,1,1],"quality":"Equal","isMain":false,"metricName":"Reference fragments"},{"values":["56.85","56.85","56.85","56.85","56.85","56.85","56.85","56.85","56.85","56.85","56.85","56.85","56.85"],"quality":"Equal","isMain":false,"metricName":"Reference GC (%)"}]]],"referenceName":"gi\_109521437\_ref\_NC","date":"18 July 2018, Wednesday, 16:06:48","order":[0,1,2,3,4,5,6,7,8,9,10,11,12],"assembliesNames":["ABySS\_127","ABySS\_63","CLC","IDBA\_UD","MEGAHIT","MIRA","SOAPdenovo2","SPAdes","SPAdes\_meta","SPAdes\_sc","SPAdes\_sc\_careful","Velvet","Geneious"]},{"assembliesWithNs":null,"minContig":500,"report":[["Genome statistics",[{"values":["99.695","100.000","99.068","99.824","99.836","28.120","100.000","6.311","99.881","100.000","99.876","99.881","93.499"],"quality":"More is better","isMain":true,"metricName":"Genome fraction (%)"},{"values":["1.008","1.007","1.007","1.009","1.011","1.002","1.132","1.000","1.041","1.001","1.008","1.011","1.004"],"quality":"Less is better","isMain":true,"metricName":"Duplication ratio"},{"values":[35003,35983,25891,32718,31553,2849,48398,712,39183,67243,48701,49006,6864],"quality":"More is better","isMain":true,"metricName":"Largest alignment"},{"values":[67536,67660,66992,67664,67834,18930,76051,4240,68886,67243,67230,67438,63014],"quality":"More is better","isMain":true,"metricName":"Total aligned length"},{"values":[35003,35983,10470,18305,18683,null,48398,null,40166,67243,49021,49326,2692],"quality":"More is better","isMain":false,"metricName":"NG50"},{"values":[32533,31677,8190,18305,9134,null,27653,null,23100,67243,18651,9910,1245],"quality":"More is better","isMain":false,"metricName":"NG75"},{"values":[35003,35983,10470,18305,18683,1276,48398,593,39183,67243,48701,49006,2692],"quality":"More is better","isMain":false,"metricName":"NA50"},{"values":[32533,31677,8190,18305,9134,920,27653,571,22331,67243,18529,9791,1695],"quality":"More is better","isMain":false,"metricName":"NA75"},{"values":[35003,35983,10470,18305,18683,null,48398,null,39183,67243,48701,49006,2692],"quality":"More is better","isMain":true,"metricName":"NGA50"},{"values":[32533,31677,8190,18305,9134,null,27653,null,22331,67243,18529,9791,1245],"quality":"More is better","isMain":false,"metricName":"NGA75"},{"values":[1,1,2,2,2,null,1,null,1,1,1,1,7],"quality":"Less is better","isMain":false,"metricName":"LG50"},{"values":[2,2,4,2,3,null,2,null,2,1,2,2,17],"quality":"Less is better","isMain":false,"metricName":"LG75"},{"values":[1,1,2,2,2,5,1,4,1,1,1,1,7],"quality":"Less is better","isMain":false,"metricName":"LA50"},{"values":[2,2,4,2,3,10,2,5,2,1,2,2,14],"quality":"Less is better","isMain":false,"metricName":"LA75"},{"values":[1,1,2,2,2,null,1,null,1,1,1,1,7],"quality":"Less is better","isMain":true,"metricName":"LGA50"},{"values":[2,2,4,2,3,null,2,null,2,1,2,2,17],"quality":"Less is better","isMain":false,"metricName":"LGA75"}]],["Misassemblies",[{"values":[0,0,0,0,0,0,0,0,0,0,0,0,0],"quality":"Less is better","isMain":true,"metricName":"# misassemblies"},{"values":[0,0,0,0,0,0,0,0,0,0,0,0,0],"quality":"Less is better","isMain":false,"metricName":" # relocations"},{"values":[0,0,0,0,0,0,0,0,0,0,0,0,0],"quality":"Less is better","isMain":false,"metricName":" # translocations"},{"values":[0,0,0,0,0,0,0,0,0,0,0,0,0],"quality":"Less is better","isMain":false,"metricName":" # inversions"},{"values":[0,0,0,0,0,0,0,0,0,0,0,0,0],"quality":"Less is better","isMain":false,"metricName":"# misassembled contigs"},{"values":[0,0,0,0,0,0,0,0,0,0,0,0,0],"quality":"Less is better","isMain":true,"metricName":"Misassembled contigs length"},{"values":[0,0,0,0,0,0,0,0,0,0,1,1,0],"quality":"Less is better","isMain":false,"metricName":"# local misassemblies"},{"values":[0,0,0,0,0,0,0,0,0,0,0,0,0],"quality":"Less is better","isMain":false,"metricName":"# unaligned mis. contigs"}]],["Unaligned",[{"values":[0,0,0,0,0,0,0,0,0,0,0,0,0],"quality":"Less is better","isMain":false,"metricName":"# fully unaligned contigs"},{"values":[0,0,0,0,0,0,0,0,0,0,0,0,0],"quality":"Less is better","isMain":false,"metricName":"Fully unaligned length"},{"values":[0,0,0,0,0,0,0,0,1,0,0,0,0],"quality":"Less is better","isMain":false,"metricName":"# partially unaligned contigs"},{"values":[0,0,0,0,0,0,0,0,769,0,0,0,0],"quality":"Less is better","isMain":false,"metricName":"Partially unaligned length"}]],["Mismatches",[{"values":[0,0,10,0,14,12,3,0,0,0,3,0,4],"quality":"Less is better","isMain":false,"metricName":"# mismatches"},{"values":[1,1,0,0,4,0,0,0,1,1,3,1,0],"quality":"Less is better","isMain":false,"metricName":"# indels"},{"values":[9,9,0,0,51,0,0,0,42,55,57,42,0],"quality":"Less is better","isMain":false,"metricName":"Indels length"},{"values":["0.00","0.00","15.02","0.00","20.87","63.52","4.47","0.00","0.00","0.00","4.47","0.00","6.37"],"quality":"Less is better","isMain":true,"metricName":"# mismatches per 100 kbp"},{"values":["1.49","1.49","0.00","0.00","5.96","0.00","0.00","0.00","1.49","1.49","4.47","1.49","0.00"],"quality":"Less is better","isMain":true,"metricName":"# indels per 100 kbp"},{"values":[0,0,0,0,1,0,0,0,0,0,0,0,0],"quality":"Less is better","isMain":false,"metricName":" # indels (<= 5 bp)"},{"values":[1,1,0,0,3,0,0,0,1,1,3,1,0],"quality":"Less is better","isMain":false,"metricName":" # indels (> 5 bp)"},{"values":[0,5,0,0,0,1,0,0,1752,0,122,118,0],"quality":"Less is better","isMain":false,"metricName":"# N's"},{"values":["0.00","7.39","0.00","0.00","0.00","5.28","0.00","0.00","2480.25","0.00","180.28","173.84","0.00"],"quality":"Less is better","isMain":true,"metricName":"# N's per 100 kbp"}]],["Statistics without reference",[{"values":[2,2,10,8,6,17,2,7,10,1,2,6,29],"quality":"Equal","isMain":true,"metricName":"# contigs"},{"values":[2,2,7,8,5,6,2,0,3,1,2,4,23],"quality":"Equal","isMain":false,"metricName":"# contigs (>= 1000 bp)"},{"values":[2,2,5,3,4,0,2,0,2,1,2,2,3],"quality":"Equal","isMain":false,"metricName":"# contigs (>= 5000 bp)"},{"values":[2,2,2,2,2,0,2,0,2,1,2,1,0],"quality":"Equal","isMain":false,"metricName":"# contigs (>= 10000 bp)"},{"values":[2,2,1,1,1,0,2,0,1,1,1,1,0],"quality":"Equal","isMain":false,"metricName":"# contigs (>= 25000 bp)"},{"values":[0,0,0,0,0,0,0,0,0,1,0,0,0],"quality":"Equal","isMain":false,"metricName":"# contigs (>= 50000 bp)"},{"values":[35003,35983,25891,32718,31553,2849,48398,712,40166,67243,49021,49326,6864],"quality":"More is better","isMain":true,"metricName":"Largest contig"},{"values":[67536,67660,67043,67664,67834,18930,76051,4240,70638,67243,67672,67877,63061],"quality":"More is better","isMain":true,"metricName":"Total length"},{"values":[67536,67660,65189,67664,66905,10613,76051,0,66122,67243,67672,66025,58395],"quality":"More is better","isMain":true,"metricName":"Total length (>= 1000 bp)"},{"values":[67536,67660,58138,59493,64495,0,76051,0,63266,67243,67672,59236,18236],"quality":"More is better","isMain":false,"metricName":"Total length (>= 5000 bp)"},{"values":[67536,67660,36361,51023,50236,0,76051,0,63266,67243,67672,49326,0],"quality":"More is better","isMain":true,"metricName":"Total length (>= 10000 bp)"},{"values":[67536,67660,25891,32718,31553,0,76051,0,40166,67243,49021,49326,0],"quality":"More is better","isMain":false,"metricName":"Total length (>= 25000 bp)"},{"values":[0,0,0,0,0,0,0,0,0,67243,0,0,0],"quality":"More is better","isMain":true,"metricName":"Total length (>= 50000 bp)"},{"values":[35003,35983,10470,18305,18683,1276,48398,593,40166,67243,49021,49326,2692],"quality":"More is better","isMain":false,"metricName":"N50"},{"values":[32533,31677,8190,18305,9134,920,27653,571,23100,67243,18651,9910,1695],"quality":"More is better","isMain":false,"metricName":"N75"},{"values":[1,1,2,2,2,5,1,4,1,1,1,1,7],"quality":"Less is better","isMain":false,"metricName":"L50"},{"values":[2,2,4,2,3,10,2,5,2,1,2,2,14],"quality":"Less is better","isMain":false,"metricName":"L75"},{"values":["69.04","68.96","69.00","68.95","68.98","68.84","69.06","69.15","68.93","68.97","68.97","68.98","68.91"],"quality":"Equal","isMain":false,"metricName":"GC (%)"}]],["Predicted genes",[]],["Similarity statistics",[{"values":[0,0,0,0,0,0,0,0,0,0,0,0,0],"quality":"Equal","isMain":false,"metricName":"# similar correct contigs"},{"values":[0,0,0,0,0,0,0,0,0,0,0,0,0],"quality":"Equal","isMain":false,"metricName":"# similar misassembled blocks"}]],["Reference statistics",[{"values":[67188,67188,67188,67188,67188,67188,67188,67188,67188,67188,67188,67188,67188],"quality":"Equal","isMain":false,"metricName":"Reference length"},{"values":[1,1,1,1,1,1,1,1,1,1,1,1,1],"quality":"Equal","isMain":false,"metricName":"Reference fragments"},{"values":["68.96","68.96","68.96","68.96","68.96","68.96","68.96","68.96","68.96","68.96","68.96","68.96","68.96"],"quality":"Equal","isMain":false,"metricName":"Reference GC (%)"}]]],"referenceName":"gi\_109521758\_ref\_NC","date":"18 July 2018, Wednesday, 16:06:58","order":[0,1,2,3,4,5,6,7,8,9,10,11,12],"assembliesNames":["ABySS\_127","ABySS\_63","CLC","IDBA\_UD","MEGAHIT","MIRA","Ray\_Meta","SOAPdenovo2","SPAdes","SPAdes\_meta","SPAdes\_sc","SPAdes\_sc\_careful","Velvet"]},{"assembliesWithNs":null,"minContig":500,"report":[["Genome statistics",[{"values":["16.290","1.932","3.105","10.582","8.926","71.316","3.112","12.549","6.046","3.462","11.836","0.882","4.056"],"quality":"More is better","isMain":true,"metricName":"Genome fraction (%)"},{"values":["1.000","1.000","1.000","1.000","1.010","1.100","1.000","1.025","1.000","1.000","1.064","1.000","1.000"],"quality":"Less is better","isMain":true,"metricName":"Duplication ratio"},{"values":[1102,673,1041,1898,1759,4134,1042,1926,1853,1854,5780,609,2801],"quality":"More is better","isMain":true,"metricName":"Largest alignment"},{"values":[11250,1334,2144,7308,6164,54155,2149,8832,4175,2391,8696,609,2801],"quality":"More is better","isMain":true,"metricName":"Total aligned length"},{"values":[null,null,null,null,null,1515,null,null,null,null,null,null,null],"quality":"More is better","isMain":false,"metricName":"NG50"},{"values":[null,null,null,null,null,925,null,null,null,null,null,null,null],"quality":"More is better","isMain":false,"metricName":"NG75"},{"values":[686,673,593,636,858,1838,595,672,657,1854,5780,609,2801],"quality":"More is better","isMain":false,"metricName":"NA50"},{"values":[594,661,593,545,619,1394,595,584,551,1854,1854,609,2801],"quality":"More is better","isMain":false,"metricName":"NA75"},{"values":[null,null,null,null,null,1515,null,null,null,null,null,null,null],"quality":"More is better","isMain":true,"metricName":"NGA50"},{"values":[null,null,null,null,null,925,null,null,null,null,null,null,null],"quality":"More is better","isMain":false,"metricName":"NGA75"},{"values":[null,null,null,null,null,16,null,null,null,null,null,null,null],"quality":"Less is better","isMain":false,"metricName":"LG50"},{"values":[null,null,null,null,null,30,null,null,null,null,null,null,null],"quality":"Less is better","isMain":false,"metricName":"LG75"},{"values":[7,1,2,4,3,11,2,4,2,1,1,1,1],"quality":"Less is better","isMain":false,"metricName":"LA50"},{"values":[11,2,2,7,5,20,2,8,4,1,2,1,1],"quality":"Less is better","isMain":false,"metricName":"LA75"},{"values":[null,null,null,null,null,16,null,null,null,null,null,null,null],"quality":"Less is better","isMain":true,"metricName":"LGA50"},{"values":[null,null,null,null,null,30,null,null,null,null,null,null,null],"quality":"Less is better","isMain":false,"metricName":"LGA75"}]],["Misassemblies",[{"values":[0,0,0,0,0,0,0,0,0,0,0,0,0],"quality":"Less is better","isMain":true,"metricName":"# misassemblies"},{"values":[0,0,0,0,0,0,0,0,0,0,0,0,0],"quality":"Less is better","isMain":false,"metricName":" # relocations"},{"values":[0,0,0,0,0,0,0,0,0,0,0,0,0],"quality":"Less is better","isMain":false,"metricName":" # translocations"},{"values":[0,0,0,0,0,0,0,0,0,0,0,0,0],"quality":"Less is better","isMain":false,"metricName":" # inversions"},{"values":[0,0,0,0,0,0,0,0,0,0,0,0,0],"quality":"Less is better","isMain":false,"metricName":"# misassembled contigs"},{"values":[0,0,0,0,0,0,0,0,0,0,0,0,0],"quality":"Less is better","isMain":true,"metricName":"Misassembled contigs length"},{"values":[0,0,0,0,0,0,0,0,0,0,0,0,0],"quality":"Less is better","isMain":false,"metricName":"# local misassemblies"},{"values":[0,0,0,0,0,0,0,0,0,0,0,0,0],"quality":"Less is better","isMain":false,"metricName":"# unaligned mis. contigs"}]],["Unaligned",[{"values":[0,0,0,0,0,0,0,0,0,0,0,0,0],"quality":"Less is better","isMain":false,"metricName":"# fully unaligned contigs"},{"values":[0,0,0,0,0,0,0,0,0,0,0,0,0],"quality":"Less is better","isMain":false,"metricName":"Fully unaligned length"},{"values":[0,0,0,0,0,0,0,0,0,0,0,0,0],"quality":"Less is better","isMain":false,"metricName":"# partially unaligned contigs"},{"values":[0,0,0,0,0,0,0,0,0,0,0,0,0],"quality":"Less is better","isMain":false,"metricName":"Partially unaligned length"}]],["Mismatches",[{"values":[5,0,0,0,13,154,0,11,3,0,91,0,7],"quality":"Less is better","isMain":false,"metricName":"# mismatches"},{"values":[0,0,0,0,0,5,0,0,0,0,1,0,0],"quality":"Less is better","isMain":false,"metricName":"# indels"},{"values":[0,0,0,0,0,6,0,0,0,0,3,0,0],"quality":"Less is better","isMain":false,"metricName":"Indels length"},{"values":["44.44","0.00","0.00","0.00","210.90","312.69","0.00","126.93","71.86","0.00","1113.29","0.00","249.91"],"quality":"Less is better","isMain":true,"metricName":"# mismatches per 100 kbp"},{"values":["0.00","0.00","0.00","0.00","0.00","10.15","0.00","0.00","0.00","0.00","12.23","0.00","0.00"],"quality":"Less is better","isMain":true,"metricName":"# indels per 100 kbp"},{"values":[0,0,0,0,0,5,0,0,0,0,1,0,0],"quality":"Less is better","isMain":false,"metricName":" # indels (<= 5 bp)"},{"values":[0,0,0,0,0,0,0,0,0,0,0,0,0],"quality":"Less is better","isMain":false,"metricName":" # indels (> 5 bp)"},{"values":[0,0,0,0,0,66,0,0,0,0,0,0,21],"quality":"Less is better","isMain":false,"metricName":"# N's"},{"values":["0.00","0.00","0.00","0.00","0.00","121.86","0.00","0.00","0.00","0.00","0.00","0.00","749.73"],"quality":"Less is better","isMain":true,"metricName":"# N's per 100 kbp"}]],["Statistics without reference",[{"values":[16,2,3,10,7,33,3,11,5,2,4,1,1],"quality":"Equal","isMain":true,"metricName":"# contigs"},{"values":[1,0,1,1,1,28,1,2,1,1,2,0,1],"quality":"Equal","isMain":false,"metricName":"# contigs (>= 1000 bp)"},{"values":[0,0,0,0,0,0,0,0,0,0,1,0,0],"quality":"Equal","isMain":false,"metricName":"# contigs (>= 5000 bp)"},{"values":[0,0,0,0,0,0,0,0,0,0,0,0,0],"quality":"Equal","isMain":false,"metricName":"# contigs (>= 10000 bp)"},{"values":[0,0,0,0,0,0,0,0,0,0,0,0,0],"quality":"Equal","isMain":false,"metricName":"# contigs (>= 25000 bp)"},{"values":[0,0,0,0,0,0,0,0,0,0,0,0,0],"quality":"Equal","isMain":false,"metricName":"# contigs (>= 50000 bp)"},{"values":[1102,673,1041,1898,1759,4134,1042,1926,1853,1854,5780,609,2801],"quality":"More is better","isMain":true,"metricName":"Largest contig"},{"values":[11250,1334,2144,7308,6225,54160,2149,8883,4175,2391,8696,609,2801],"quality":"More is better","isMain":true,"metricName":"Total length"},{"values":[1102,0,1041,1898,1759,50084,1042,3371,1853,1854,7634,0,2801],"quality":"More is better","isMain":true,"metricName":"Total length (>= 1000 bp)"},{"values":[0,0,0,0,0,0,0,0,0,0,5780,0,0],"quality":"More is better","isMain":false,"metricName":"Total length (>= 5000 bp)"},{"values":[0,0,0,0,0,0,0,0,0,0,0,0,0],"quality":"More is better","isMain":true,"metricName":"Total length (>= 10000 bp)"},{"values":[0,0,0,0,0,0,0,0,0,0,0,0,0],"quality":"More is better","isMain":false,"metricName":"Total length (>= 25000 bp)"},{"values":[0,0,0,0,0,0,0,0,0,0,0,0,0],"quality":"More is better","isMain":true,"metricName":"Total length (>= 50000 bp)"},{"values":[686,673,593,636,858,1838,595,672,657,1854,5780,609,2801],"quality":"More is better","isMain":false,"metricName":"N50"},{"values":[594,661,593,545,668,1394,595,596,551,1854,1854,609,2801],"quality":"More is better","isMain":false,"metricName":"N75"},{"values":[7,1,2,4,3,11,2,4,2,1,1,1,1],"quality":"Less is better","isMain":false,"metricName":"L50"},{"values":[11,2,2,7,5,20,2,7,4,1,2,1,1],"quality":"Less is better","isMain":false,"metricName":"L75"},{"values":["66.62","63.94","63.71","66.80","65.67","67.30","63.75","67.43","65.89","66.16","66.89","59.28","69.82"],"quality":"Equal","isMain":false,"metricName":"GC (%)"}]],["Predicted genes",[]],["Similarity statistics",[{"values":[0,0,0,0,0,0,0,0,0,0,0,0,0],"quality":"Equal","isMain":false,"metricName":"# similar correct contigs"},{"values":[0,0,0,0,0,0,0,0,0,0,0,0,0],"quality":"Equal","isMain":false,"metricName":"# similar misassembled blocks"}]],["Reference statistics",[{"values":[69059,69059,69059,69059,69059,69059,69059,69059,69059,69059,69059,69059,69059],"quality":"Equal","isMain":false,"metricName":"Reference length"},{"values":[1,1,1,1,1,1,1,1,1,1,1,1,1],"quality":"Equal","isMain":false,"metricName":"Reference fragments"},{"values":["67.34","67.34","67.34","67.34","67.34","67.34","67.34","67.34","67.34","67.34","67.34","67.34","67.34"],"quality":"Equal","isMain":false,"metricName":"Reference GC (%)"}]]],"referenceName":"gi\_109521840\_ref\_NC","date":"18 July 2018, Wednesday, 16:07:09","order":[0,1,2,3,4,5,6,7,8,9,10,11,12],"assembliesNames":["ABySS\_127","ABySS\_63","CLC","IDBA\_UD","MEGAHIT","MIRA","SOAPdenovo2","SPAdes","SPAdes\_meta","SPAdes\_sc","SPAdes\_sc\_careful","Velvet","Geneious"]},{"assembliesWithNs":null,"minContig":500,"report":[["Genome statistics",[{"values":["1.789","0.991","0.849","0.729","4.039","0.980","0.980"],"quality":"More is better","isMain":true,"metricName":"Genome fraction (%)"},{"values":["1.000","1.000","1.000","1.511","1.045","1.000","1.000"],"quality":"Less is better","isMain":true,"metricName":"Duplication ratio"},{"values":[1159,642,550,472,617,635,635],"quality":"More is better","isMain":true,"metricName":"Largest alignment"},{"values":[1159,642,550,472,2734,635,635],"quality":"More is better","isMain":true,"metricName":"Total aligned length"},{"values":[1159,642,550,472,523,635,635],"quality":"More is better","isMain":false,"metricName":"NA50"},{"values":[1159,642,550,null,520,635,635],"quality":"More is better","isMain":false,"metricName":"NA75"},{"values":[null,null,null,null,null,null,null],"quality":"More is better","isMain":true,"metricName":"NGA50"},{"values":[1,1,1,1,3,1,1],"quality":"Less is better","isMain":false,"metricName":"LA50"},{"values":[1,1,1,null,4,1,1],"quality":"Less is better","isMain":false,"metricName":"LA75"}]],["Misassemblies",[{"values":[0,0,0,0,0,0,0],"quality":"Less is better","isMain":true,"metricName":"# misassemblies"},{"values":[0,0,0,0,0,0,0],"quality":"Less is better","isMain":false,"metricName":" # relocations"},{"values":[0,0,0,0,0,0,0],"quality":"Less is better","isMain":false,"metricName":" # translocations"},{"values":[0,0,0,0,0,0,0],"quality":"Less is better","isMain":false,"metricName":" # inversions"},{"values":[0,0,0,0,0,0,0],"quality":"Less is better","isMain":false,"metricName":"# misassembled contigs"},{"values":[0,0,0,0,0,0,0],"quality":"Less is better","isMain":true,"metricName":"Misassembled contigs length"},{"values":[0,0,0,0,0,0,0],"quality":"Less is better","isMain":false,"metricName":"# local misassemblies"},{"values":[0,0,0,0,0,0,0],"quality":"Less is better","isMain":false,"metricName":"# unaligned mis. contigs"}]],["Unaligned",[{"values":[0,0,0,0,0,0,0],"quality":"Less is better","isMain":false,"metricName":"# fully unaligned contigs"},{"values":[0,0,0,0,0,0,0],"quality":"Less is better","isMain":false,"metricName":"Fully unaligned length"},{"values":[0,0,0,0,0,0,0],"quality":"Less is better","isMain":false,"metricName":"# partially unaligned contigs"},{"values":[0,0,0,0,0,0,0],"quality":"Less is better","isMain":false,"metricName":"Partially unaligned length"}]],["Mismatches",[{"values":[3,6,1,11,4,1,1],"quality":"Less is better","isMain":false,"metricName":"# mismatches"},{"values":[0,0,0,0,0,0,0],"quality":"Less is better","isMain":false,"metricName":"# indels"},{"values":[0,0,0,0,0,0,0],"quality":"Less is better","isMain":false,"metricName":"Indels length"},{"values":["258.84","934.58","181.82","2330.51","152.85","157.48","157.48"],"quality":"Less is better","isMain":true,"metricName":"# mismatches per 100 kbp"},{"values":["0.00","0.00","0.00","0.00","0.00","0.00","0.00"],"quality":"Less is better","isMain":true,"metricName":"# indels per 100 kbp"},{"values":[0,0,0,0,0,0,0],"quality":"Less is better","isMain":false,"metricName":" # indels (<= 5 bp)"},{"values":[0,0,0,0,0,0,0],"quality":"Less is better","isMain":false,"metricName":" # indels (> 5 bp)"},{"values":[0,0,0,47,0,0,0],"quality":"Less is better","isMain":false,"metricName":"# N's"},{"values":["0.00","0.00","0.00","6591.87","0.00","0.00","0.00"],"quality":"Less is better","isMain":true,"metricName":"# N's per 100 kbp"}]],["Statistics without reference",[{"values":[1,1,1,1,5,1,1],"quality":"Equal","isMain":true,"metricName":"# contigs"},{"values":[1,0,0,0,0,0,0],"quality":"Equal","isMain":false,"metricName":"# contigs (>= 1000 bp)"},{"values":[0,0,0,0,0,0,0],"quality":"Equal","isMain":false,"metricName":"# contigs (>= 5000 bp)"},{"values":[0,0,0,0,0,0,0],"quality":"Equal","isMain":false,"metricName":"# contigs (>= 10000 bp)"},{"values":[0,0,0,0,0,0,0],"quality":"Equal","isMain":false,"metricName":"# contigs (>= 25000 bp)"},{"values":[0,0,0,0,0,0,0],"quality":"Equal","isMain":false,"metricName":"# contigs (>= 50000 bp)"},{"values":[1159,642,550,713,617,635,635],"quality":"More is better","isMain":true,"metricName":"Largest contig"},{"values":[1159,642,550,713,2734,635,635],"quality":"More is better","isMain":true,"metricName":"Total length"},{"values":[1159,0,0,0,0,0,0],"quality":"More is better","isMain":true,"metricName":"Total length (>= 1000 bp)"},{"values":[0,0,0,0,0,0,0],"quality":"More is better","isMain":false,"metricName":"Total length (>= 5000 bp)"},{"values":[0,0,0,0,0,0,0],"quality":"More is better","isMain":true,"metricName":"Total length (>= 10000 bp)"},{"values":[0,0,0,0,0,0,0],"quality":"More is better","isMain":false,"metricName":"Total length (>= 25000 bp)"},{"values":[0,0,0,0,0,0,0],"quality":"More is better","isMain":true,"metricName":"Total length (>= 50000 bp)"},{"values":[1159,642,550,713,523,635,635],"quality":"More is better","isMain":false,"metricName":"N50"},{"values":[1159,642,550,713,520,635,635],"quality":"More is better","isMain":false,"metricName":"N75"},{"values":[1,1,1,1,3,1,1],"quality":"Less is better","isMain":false,"metricName":"L50"},{"values":[1,1,1,1,4,1,1],"quality":"Less is better","isMain":false,"metricName":"L75"},{"values":["57.81","58.88","63.45","60.36","58.19","57.32","57.32"],"quality":"Equal","isMain":false,"metricName":"GC (%)"}]],["Predicted genes",[]],["Similarity statistics",[{"values":[0,0,0,0,0,0,0],"quality":"Equal","isMain":false,"metricName":"# similar correct contigs"},{"values":[0,0,0,0,0,0,0],"quality":"Equal","isMain":false,"metricName":"# similar misassembled blocks"}]],["Reference statistics",[{"values":[64787,64787,64787,64787,64787,64787,64787],"quality":"Equal","isMain":false,"metricName":"Reference length"},{"values":[1,1,1,1,1,1,1],"quality":"Equal","isMain":false,"metricName":"Reference fragments"},{"values":["59.66","59.66","59.66","59.66","59.66","59.66","59.66"],"quality":"Equal","isMain":false,"metricName":"Reference GC (%)"}]]],"referenceName":"gi\_109522567\_ref\_NC","date":"18 July 2018, Wednesday, 16:07:16","order":[0,1,2,3,4,5,6],"assembliesNames":["ABySS\_127","CLC","MEGAHIT","SOAPdenovo2","SPAdes","SPAdes\_sc","SPAdes\_sc\_careful"]},{"assembliesWithNs":null,"minContig":500,"report":[["Genome statistics",[{"values":["13.854","7.201","13.373","0.919","9.660","14.914","14.775","14.593","14.782"],"quality":"More is better","isMain":true,"metricName":"Genome fraction (%)"},{"values":["1.000","1.000","1.000","1.000","1.000","1.000","1.000","1.000","1.000"],"quality":"Less is better","isMain":true,"metricName":"Duplication ratio"},{"values":[1968,1678,1624,1170,1380,1968,1968,1968,1968],"quality":"More is better","isMain":true,"metricName":"Largest alignment"},{"values":[30597,15905,29535,2030,21336,32939,32631,32230,32647],"quality":"More is better","isMain":true,"metricName":"Total aligned length"},{"values":[850,852,771,1170,718,874,850,850,850],"quality":"More is better","isMain":false,"metricName":"NA50"},{"values":[738,557,628,860,580,742,742,738,738],"quality":"More is better","isMain":false,"metricName":"NA75"},{"values":[null,null,null,null,null,null,null,null,null],"quality":"More is better","isMain":true,"metricName":"NGA50"},{"values":[14,8,16,1,11,15,15,15,15],"quality":"Less is better","isMain":false,"metricName":"LA50"},{"values":[24,15,26,2,20,25,25,25,25],"quality":"Less is better","isMain":false,"metricName":"LA75"}]],["Misassemblies",[{"values":[0,0,0,0,0,0,0,0,0],"quality":"Less is better","isMain":true,"metricName":"# misassemblies"},{"values":[0,0,0,0,0,0,0,0,0],"quality":"Less is better","isMain":false,"metricName":" # relocations"},{"values":[0,0,0,0,0,0,0,0,0],"quality":"Less is better","isMain":false,"metricName":" # translocations"},{"values":[0,0,0,0,0,0,0,0,0],"quality":"Less is better","isMain":false,"metricName":" # inversions"},{"values":[0,0,0,0,0,0,0,0,0],"quality":"Less is better","isMain":false,"metricName":"# misassembled contigs"},{"values":[0,0,0,0,0,0,0,0,0],"quality":"Less is better","isMain":true,"metricName":"Misassembled contigs length"},{"values":[0,0,1,0,0,0,0,0,0],"quality":"Less is better","isMain":false,"metricName":"# local misassemblies"},{"values":[0,0,0,0,0,0,0,0,0],"quality":"Less is better","isMain":false,"metricName":"# unaligned mis. contigs"}]],["Unaligned",[{"values":[0,0,0,0,0,0,0,0,0],"quality":"Less is better","isMain":false,"metricName":"# fully unaligned contigs"},{"values":[0,0,0,0,0,0,0,0,0],"quality":"Less is better","isMain":false,"metricName":"Fully unaligned length"},{"values":[0,1,0,0,0,0,0,0,0],"quality":"Less is better","isMain":false,"metricName":"# partially unaligned contigs"},{"values":[0,1456,0,0,0,0,0,0,0],"quality":"Less is better","isMain":false,"metricName":"Partially unaligned length"}]],["Mismatches",[{"values":[50,26,41,0,21,53,48,45,44],"quality":"Less is better","isMain":false,"metricName":"# mismatches"},{"values":[0,0,1,0,0,0,0,0,0],"quality":"Less is better","isMain":false,"metricName":"# indels"},{"values":[0,0,78,0,0,0,0,0,0],"quality":"Less is better","isMain":false,"metricName":"Indels length"},{"values":["163.41","163.47","138.82","0.00","98.43","160.90","147.10","139.62","134.78"],"quality":"Less is better","isMain":true,"metricName":"# mismatches per 100 kbp"},{"values":["0.00","0.00","3.39","0.00","0.00","0.00","0.00","0.00","0.00"],"quality":"Less is better","isMain":true,"metricName":"# indels per 100 kbp"},{"values":[0,0,0,0,0,0,0,0,0],"quality":"Less is better","isMain":false,"metricName":" # indels (<= 5 bp)"},{"values":[0,0,1,0,0,0,0,0,0],"quality":"Less is better","isMain":false,"metricName":" # indels (> 5 bp)"},{"values":[0,0,0,2,0,0,0,0,0],"quality":"Less is better","isMain":false,"metricName":"# N's"},{"values":["0.00","0.00","0.00","98.52","0.00","0.00","0.00","0.00","0.00"],"quality":"Less is better","isMain":true,"metricName":"# N's per 100 kbp"}]],["Statistics without reference",[{"values":[36,20,39,2,29,38,38,38,38],"quality":"Equal","isMain":true,"metricName":"# contigs"},{"values":[9,6,4,1,3,11,10,10,10],"quality":"Equal","isMain":false,"metricName":"# contigs (>= 1000 bp)"},{"values":[0,0,0,0,0,0,0,0,0],"quality":"Equal","isMain":false,"metricName":"# contigs (>= 5000 bp)"},{"values":[0,0,0,0,0,0,0,0,0],"quality":"Equal","isMain":false,"metricName":"# contigs (>= 10000 bp)"},{"values":[0,0,0,0,0,0,0,0,0],"quality":"Equal","isMain":false,"metricName":"# contigs (>= 25000 bp)"},{"values":[0,0,0,0,0,0,0,0,0],"quality":"Equal","isMain":false,"metricName":"# contigs (>= 50000 bp)"},{"values":[1968,1716,1624,1170,1380,1968,1968,1968,1968],"quality":"More is better","isMain":true,"metricName":"Largest contig"},{"values":[30597,17361,29535,2030,21336,32939,32632,32230,32647],"quality":"More is better","isMain":true,"metricName":"Total length"},{"values":[11335,7884,5108,1170,3713,13489,12464,12464,12464],"quality":"More is better","isMain":true,"metricName":"Total length (>= 1000 bp)"},{"values":[0,0,0,0,0,0,0,0,0],"quality":"More is better","isMain":false,"metricName":"Total length (>= 5000 bp)"},{"values":[0,0,0,0,0,0,0,0,0],"quality":"More is better","isMain":true,"metricName":"Total length (>= 10000 bp)"},{"values":[0,0,0,0,0,0,0,0,0],"quality":"More is better","isMain":false,"metricName":"Total length (>= 25000 bp)"},{"values":[0,0,0,0,0,0,0,0,0],"quality":"More is better","isMain":true,"metricName":"Total length (>= 50000 bp)"},{"values":[850,909,771,1170,718,874,850,850,850],"quality":"More is better","isMain":false,"metricName":"N50"},{"values":[738,695,628,860,580,742,742,738,738],"quality":"More is better","isMain":false,"metricName":"N75"},{"values":[14,7,16,1,11,15,15,15,15],"quality":"Less is better","isMain":false,"metricName":"L50"},{"values":[24,13,26,2,20,25,25,25,25],"quality":"Less is better","isMain":false,"metricName":"L75"},{"values":["54.71","53.37","55.68","62.82","54.49","55.19","55.15","54.96","54.98"],"quality":"Equal","isMain":false,"metricName":"GC (%)"}]],["Predicted genes",[]],["Similarity statistics",[{"values":[0,0,0,0,0,0,0,0,0],"quality":"Equal","isMain":false,"metricName":"# similar correct contigs"},{"values":[0,0,0,0,0,0,0,0,0],"quality":"Equal","isMain":false,"metricName":"# similar misassembled blocks"}]],["Reference statistics",[{"values":[220859,220859,220859,220859,220859,220859,220859,220859,220859],"quality":"Equal","isMain":false,"metricName":"Reference length"},{"values":[1,1,1,1,1,1,1,1,1],"quality":"Equal","isMain":false,"metricName":"Reference fragments"},{"values":["54.55","54.55","54.55","54.55","54.55","54.55","54.55","54.55","54.55"],"quality":"Equal","isMain":false,"metricName":"Reference GC (%)"}]]],"referenceName":"gi\_109638461\_ref\_NC","date":"18 July 2018, Wednesday, 16:07:24","order":[0,1,2,3,4,5,6,7,8],"assembliesNames":["CLC","IDBA\_UD","MEGAHIT","MIRA","SOAPdenovo2","SPAdes","SPAdes\_meta","SPAdes\_sc","SPAdes\_sc\_careful"]},{"assembliesWithNs":null,"minContig":500,"report":[["Genome statistics",[{"values":["2.560","0.255","49.530","34.524","43.893","7.275","38.526","51.300","51.329","51.561","51.562","0.258"],"quality":"More is better","isMain":true,"metricName":"Genome fraction (%)"},{"values":["1.010","1.000","1.002","1.003","1.001","1.000","1.000","1.003","1.004","1.003","1.003","1.000"],"quality":"Less is better","isMain":true,"metricName":"Duplication ratio"},{"values":[825,590,3177,2826,2775,1645,3177,3177,3177,3177,3177,598],"quality":"More is better","isMain":true,"metricName":"Largest alignment"},{"values":[5933,590,114863,80035,101767,16863,89323,118950,119461,119563,119561,598],"quality":"More is better","isMain":true,"metricName":"Total aligned length"},{"values":[null,null,null,null,null,null,null,541,535,547,547,null],"quality":"More is better","isMain":false,"metricName":"NG50"},{"values":[530,590,934,924,828,1067,853,939,938,941,941,null],"quality":"More is better","isMain":false,"metricName":"NA50"},{"values":[522,590,799,681,646,756,652,799,801,801,801,null],"quality":"More is better","isMain":false,"metricName":"NA75"},{"values":[null,null,null,null,null,null,null,535,535,535,535,null],"quality":"More is better","isMain":true,"metricName":"NGA50"},{"values":[null,null,null,null,null,null,null,117,118,117,117,null],"quality":"Less is better","isMain":false,"metricName":"LG50"},{"values":[5,1,44,33,45,7,37,46,45,44,44,null],"quality":"Less is better","isMain":false,"metricName":"LA50"},{"values":[8,1,77,59,80,12,67,80,79,78,78,null],"quality":"Less is better","isMain":false,"metricName":"LA75"},{"values":[null,null,null,null,null,null,null,119,118,118,118,null],"quality":"Less is better","isMain":true,"metricName":"LGA50"}]],["Misassemblies",[{"values":[0,0,0,1,0,0,0,0,0,0,0,0],"quality":"Less is better","isMain":true,"metricName":"# misassemblies"},{"values":[0,0,0,1,0,0,0,0,0,0,0,0],"quality":"Less is better","isMain":false,"metricName":" # relocations"},{"values":[0,0,0,0,0,0,0,0,0,0,0,0],"quality":"Less is better","isMain":false,"metricName":" # translocations"},{"values":[0,0,0,0,0,0,0,0,0,0,0,0],"quality":"Less is better","isMain":false,"metricName":" # inversions"},{"values":[0,0,0,1,0,0,0,0,0,0,0,0],"quality":"Less is better","isMain":false,"metricName":"# misassembled contigs"},{"values":[0,0,0,1415,0,0,0,0,0,0,0,0],"quality":"Less is better","isMain":true,"metricName":"Misassembled contigs length"},{"values":[0,0,0,0,1,0,0,0,4,3,3,0],"quality":"Less is better","isMain":false,"metricName":"# local misassemblies"},{"values":[0,0,0,0,0,0,0,0,0,0,0,0],"quality":"Less is better","isMain":false,"metricName":"# unaligned mis. contigs"}]],["Unaligned",[{"values":[0,0,0,0,0,0,0,0,0,0,0,0],"quality":"Less is better","isMain":false,"metricName":"# fully unaligned contigs"},{"values":[0,0,0,0,0,0,0,0,0,0,0,0],"quality":"Less is better","isMain":false,"metricName":"Fully unaligned length"},{"values":[0,0,0,1,0,0,0,1,1,0,0,1],"quality":"Less is better","isMain":false,"metricName":"# partially unaligned contigs"},{"values":[0,0,0,790,0,0,0,790,548,0,0,43951],"quality":"Less is better","isMain":false,"metricName":"Partially unaligned length"}]],["Mismatches",[{"values":[2,0,176,91,132,5,98,212,202,184,151,0],"quality":"Less is better","isMain":false,"metricName":"# mismatches"},{"values":[1,0,2,6,9,0,1,1,6,5,3,0],"quality":"Less is better","isMain":false,"metricName":"# indels"},{"values":[1,0,2,6,68,0,42,1,13,12,10,0],"quality":"Less is better","isMain":false,"metricName":"Indels length"},{"values":["33.70","0.00","153.30","113.71","129.74","29.65","109.74","178.28","169.78","153.95","126.34","0.00"],"quality":"Less is better","isMain":true,"metricName":"# mismatches per 100 kbp"},{"values":["16.85","0.00","1.74","7.50","8.85","0.00","1.12","0.84","5.04","4.18","2.51","0.00"],"quality":"Less is better","isMain":true,"metricName":"# indels per 100 kbp"},{"values":[1,0,2,6,7,0,0,1,5,4,2,0],"quality":"Less is better","isMain":false,"metricName":" # indels (<= 5 bp)"},{"values":[0,0,0,0,2,0,1,0,1,1,1,0],"quality":"Less is better","isMain":false,"metricName":" # indels (> 5 bp)"},{"values":[0,0,0,0,0,37,0,0,0,0,0,4],"quality":"Less is better","isMain":false,"metricName":"# N's"},{"values":["0.00","0.00","0.00","0.00","0.00","219.42","0.00","0.00","0.00","0.00","0.00","8.98"],"quality":"Less is better","isMain":true,"metricName":"# N's per 100 kbp"}]],["Statistics without reference",[{"values":[10,1,121,90,123,18,105,124,125,124,124,1],"quality":"Equal","isMain":true,"metricName":"# contigs"},{"values":[0,0,37,27,26,8,23,42,38,40,40,1],"quality":"Equal","isMain":false,"metricName":"# contigs (>= 1000 bp)"},{"values":[0,0,0,0,0,0,0,0,0,0,0,1],"quality":"Equal","isMain":false,"metricName":"# contigs (>= 5000 bp)"},{"values":[0,0,0,0,0,0,0,0,0,0,0,1],"quality":"Equal","isMain":false,"metricName":"# contigs (>= 10000 bp)"},{"values":[0,0,0,0,0,0,0,0,0,0,0,1],"quality":"Equal","isMain":false,"metricName":"# contigs (>= 25000 bp)"},{"values":[0,0,0,0,0,0,0,0,0,0,0,0],"quality":"Equal","isMain":false,"metricName":"# contigs (>= 50000 bp)"},{"values":[858,590,3177,2826,2775,1645,3177,3177,3177,3177,3177,44549],"quality":"More is better","isMain":true,"metricName":"Largest contig"},{"values":[5993,590,114989,81029,101799,16863,89323,120079,120009,119935,119933,44549],"quality":"More is better","isMain":true,"metricName":"Total length"},{"values":[0,0,51532,35703,34326,9739,32370,57969,54132,57016,57014,44549],"quality":"More is better","isMain":true,"metricName":"Total length (>= 1000 bp)"},{"values":[0,0,0,0,0,0,0,0,0,0,0,44549],"quality":"More is better","isMain":false,"metricName":"Total length (>= 5000 bp)"},{"values":[0,0,0,0,0,0,0,0,0,0,0,44549],"quality":"More is better","isMain":true,"metricName":"Total length (>= 10000 bp)"},{"values":[0,0,0,0,0,0,0,0,0,0,0,44549],"quality":"More is better","isMain":false,"metricName":"Total length (>= 25000 bp)"},{"values":[0,0,0,0,0,0,0,0,0,0,0,0],"quality":"More is better","isMain":true,"metricName":"Total length (>= 50000 bp)"},{"values":[530,590,938,952,828,1067,853,958,938,941,941,44549],"quality":"More is better","isMain":false,"metricName":"N50"},{"values":[522,590,799,707,646,756,652,805,801,801,801,44549],"quality":"More is better","isMain":false,"metricName":"N75"},{"values":[5,1,44,32,45,7,37,45,45,44,44,1],"quality":"Less is better","isMain":false,"metricName":"L50"},{"values":[8,1,77,58,80,12,67,79,79,78,78,1],"quality":"Less is better","isMain":false,"metricName":"L75"},{"values":["54.68","51.53","53.66","53.33","53.33","53.43","53.83","53.15","53.24","53.49","53.49","35.38"],"quality":"Equal","isMain":false,"metricName":"GC (%)"}]],["Predicted genes",[]],["Similarity statistics",[{"values":[0,0,0,0,0,0,0,0,0,0,0,0],"quality":"Equal","isMain":false,"metricName":"# similar correct contigs"},{"values":[0,0,0,0,0,0,0,0,0,0,0,0],"quality":"Equal","isMain":false,"metricName":"# similar misassembled blocks"}]],["Reference statistics",[{"values":[231801,231801,231801,231801,231801,231801,231801,231801,231801,231801,231801,231801],"quality":"Equal","isMain":false,"metricName":"Reference length"},{"values":[1,1,1,1,1,1,1,1,1,1,1,1],"quality":"Equal","isMain":false,"metricName":"Reference fragments"},{"values":["52.77","52.77","52.77","52.77","52.77","52.77","52.77","52.77","52.77","52.77","52.77","52.77"],"quality":"Equal","isMain":false,"metricName":"Reference GC (%)"}]]],"referenceName":"gi\_109638594\_ref\_NC","date":"18 July 2018, Wednesday, 16:07:34","order":[0,1,2,3,4,5,6,7,8,9,10,11],"assembliesNames":["ABySS\_127","ABySS\_63","CLC","IDBA\_UD","MEGAHIT","MIRA","SOAPdenovo2","SPAdes","SPAdes\_meta","SPAdes\_sc","SPAdes\_sc\_careful","Geneious"]},{"assembliesWithNs":null,"minContig":500,"report":[["Genome statistics",[{"values":["66.791","55.896","93.114","96.526","96.493","85.887","77.221","97.544","97.354","97.196","96.995","80.795","35.025"],"quality":"More is better","isMain":true,"metricName":"Genome fraction (%)"},{"values":["1.000","1.000","1.029","1.005","1.004","1.035","1.030","1.005","1.004","1.010","1.009","1.000","1.004"],"quality":"Less is better","isMain":true,"metricName":"Duplication ratio"},{"values":[3908,3017,10597,15484,11748,6222,6428,16545,16537,16432,16160,12870,7623],"quality":"More is better","isMain":true,"metricName":"Largest alignment"},{"values":[88176,73802,125501,127709,127843,116992,102921,128985,128563,129198,128950,106670,46200],"quality":"More is better","isMain":true,"metricName":"Total aligned length"},{"values":[820,586,4009,6137,6068,2468,1724,8667,6712,6668,6668,4218,null],"quality":"More is better","isMain":false,"metricName":"NG50"},{"values":[null,null,2707,3358,4046,1397,645,5113,4638,4638,4638,1364,null],"quality":"More is better","isMain":false,"metricName":"NG75"},{"values":[1144,960,4099,6137,6068,2617,1885,8667,6481,6665,6665,4432,4347],"quality":"More is better","isMain":false,"metricName":"NA50"},{"values":[820,728,2707,3354,4369,1736,1080,6121,4638,4638,4120,2434,3992],"quality":"More is better","isMain":false,"metricName":"NA75"},{"values":[820,586,4009,6137,6068,2468,1665,8667,6481,6665,6665,4218,null],"quality":"More is better","isMain":true,"metricName":"NGA50"},{"values":[null,null,2581,3354,4046,1294,568,4638,4046,4120,4120,1364,null],"quality":"More is better","isMain":false,"metricName":"NGA75"},{"values":[45,63,12,8,8,19,24,6,6,7,7,11,null],"quality":"Less is better","isMain":false,"metricName":"LG50"},{"values":[null,null,22,15,15,35,53,11,11,12,12,26,null],"quality":"Less is better","isMain":false,"metricName":"LG75"},{"values":[22,24,11,8,8,16,18,6,6,7,7,8,4],"quality":"Less is better","isMain":false,"metricName":"LA50"},{"values":[45,47,21,15,14,29,36,10,11,12,13,16,7],"quality":"Less is better","isMain":false,"metricName":"LA75"},{"values":[45,63,12,8,8,19,25,6,6,7,7,11,null],"quality":"Less is better","isMain":true,"metricName":"LGA50"},{"values":[null,null,22,15,15,36,60,11,12,13,13,26,null],"quality":"Less is better","isMain":false,"metricName":"LGA75"}]],["Misassemblies",[{"values":[0,0,0,0,0,0,0,0,0,0,0,0,0],"quality":"Less is better","isMain":true,"metricName":"# misassemblies"},{"values":[0,0,0,0,0,0,0,0,0,0,0,0,0],"quality":"Less is better","isMain":false,"metricName":" # relocations"},{"values":[0,0,0,0,0,0,0,0,0,0,0,0,0],"quality":"Less is better","isMain":false,"metricName":" # translocations"},{"values":[0,0,0,0,0,0,0,0,0,0,0,0,0],"quality":"Less is better","isMain":false,"metricName":" # inversions"},{"values":[0,0,0,0,0,0,0,0,0,0,0,0,0],"quality":"Less is better","isMain":false,"metricName":"# misassembled contigs"},{"values":[0,0,0,0,0,0,0,0,0,0,0,0,0],"quality":"Less is better","isMain":true,"metricName":"Misassembled contigs length"},{"values":[0,0,0,0,0,1,0,1,1,1,1,0,1],"quality":"Less is better","isMain":false,"metricName":"# local misassemblies"},{"values":[0,0,0,0,0,0,0,0,0,0,0,0,0],"quality":"Less is better","isMain":false,"metricName":"# unaligned mis. contigs"}]],["Unaligned",[{"values":[0,0,0,0,0,0,0,0,0,0,0,0,0],"quality":"Less is better","isMain":false,"metricName":"# fully unaligned contigs"},{"values":[0,0,0,0,0,0,0,0,0,0,0,0,0],"quality":"Less is better","isMain":false,"metricName":"Fully unaligned length"},{"values":[0,0,0,1,1,1,2,1,0,2,4,0,0],"quality":"Less is better","isMain":false,"metricName":"# partially unaligned contigs"},{"values":[0,0,0,741,672,1518,2009,537,0,1278,3400,0,0],"quality":"Less is better","isMain":false,"metricName":"Partially unaligned length"}]],["Mismatches",[{"values":[12,4,226,92,120,56,26,204,58,324,301,16,4],"quality":"Less is better","isMain":false,"metricName":"# mismatches"},{"values":[2,0,19,3,4,2,28,3,7,16,15,0,1],"quality":"Less is better","isMain":false,"metricName":"# indels"},{"values":[5,0,43,97,119,51,1312,68,39,152,145,0,31],"quality":"Less is better","isMain":false,"metricName":"Indels length"},{"values":["13.61","5.42","183.88","72.21","94.22","49.40","25.51","158.44","45.14","252.54","235.10","15.00","8.65"],"quality":"Less is better","isMain":true,"metricName":"# mismatches per 100 kbp"},{"values":["2.27","0.00","15.46","2.35","3.14","1.76","27.47","2.33","5.45","12.47","11.72","0.00","2.16"],"quality":"Less is better","isMain":true,"metricName":"# indels per 100 kbp"},{"values":[2,0,17,1,2,0,0,2,6,12,12,0,0],"quality":"Less is better","isMain":false,"metricName":" # indels (<= 5 bp)"},{"values":[0,0,2,2,2,2,28,1,1,4,3,0,1],"quality":"Less is better","isMain":false,"metricName":" # indels (> 5 bp)"},{"values":[0,0,0,0,0,127,3179,10,124,10,0,0,68],"quality":"Less is better","isMain":false,"metricName":"# N's"},{"values":["0.00","0.00","0.00","0.00","0.00","106.90","2970.31","7.69","96.09","7.64","0.00","0.00","146.55"],"quality":"Less is better","isMain":true,"metricName":"# N's per 100 kbp"}]],["Statistics without reference",[{"values":[78,77,46,31,28,56,67,22,24,26,26,35,11],"quality":"Equal","isMain":true,"metricName":"# contigs"},{"values":[31,23,31,28,26,42,42,19,22,23,23,29,11],"quality":"Equal","isMain":false,"metricName":"# contigs (>= 1000 bp)"},{"values":[0,0,7,10,10,2,2,11,10,11,11,5,3],"quality":"Equal","isMain":false,"metricName":"# contigs (>= 5000 bp)"},{"values":[0,0,1,2,2,0,0,3,3,3,3,1,0],"quality":"Equal","isMain":false,"metricName":"# contigs (>= 10000 bp)"},{"values":[0,0,0,0,0,0,0,0,0,0,0,0,0],"quality":"Equal","isMain":false,"metricName":"# contigs (>= 25000 bp)"},{"values":[0,0,0,0,0,0,0,0,0,0,0,0,0],"quality":"Equal","isMain":false,"metricName":"# contigs (>= 50000 bp)"},{"values":[3908,3017,10597,15484,11748,6222,6428,16545,16777,16432,17146,12870,7623],"quality":"More is better","isMain":true,"metricName":"Largest contig"},{"values":[88176,73802,126430,128816,128518,118805,107026,129996,129039,130914,132641,106673,46400],"quality":"More is better","isMain":true,"metricName":"Total length"},{"values":[53823,36498,116092,127042,127402,108235,90572,128152,127755,128708,130435,102599,46400],"quality":"More is better","isMain":true,"metricName":"Total length (>= 1000 bp)"},{"values":[0,0,47993,79327,80678,11540,11770,103260,94407,95222,95936,39674,19127],"quality":"More is better","isMain":false,"metricName":"Total length (>= 5000 bp)"},{"values":[0,0,10597,26070,23208,0,0,44778,45004,38510,39224,12870,0],"quality":"More is better","isMain":true,"metricName":"Total length (>= 10000 bp)"},{"values":[0,0,0,0,0,0,0,0,0,0,0,0,0],"quality":"More is better","isMain":false,"metricName":"Total length (>= 25000 bp)"},{"values":[0,0,0,0,0,0,0,0,0,0,0,0,0],"quality":"More is better","isMain":true,"metricName":"Total length (>= 50000 bp)"},{"values":[1144,960,4099,6137,6068,2621,2049,8667,6712,6668,6668,4432,4347],"quality":"More is better","isMain":false,"metricName":"N50"},{"values":[820,728,2872,3358,4369,1852,1303,6121,4638,4638,4638,2434,3992],"quality":"More is better","isMain":false,"metricName":"N75"},{"values":[22,24,11,8,8,16,17,6,6,7,7,8,4],"quality":"Less is better","isMain":false,"metricName":"L50"},{"values":[45,47,20,15,14,29,34,10,11,12,12,16,7],"quality":"Less is better","isMain":false,"metricName":"L75"},{"values":["54.56","54.76","54.82","54.93","55.19","54.93","54.64","55.02","55.11","54.92","54.88","54.78","54.61"],"quality":"Equal","isMain":false,"metricName":"GC (%)"}]],["Predicted genes",[]],["Similarity statistics",[{"values":[0,0,0,0,0,0,0,0,0,0,0,0,0],"quality":"Equal","isMain":false,"metricName":"# similar correct contigs"},{"values":[0,0,0,0,0,0,0,0,0,0,0,0,0],"quality":"Equal","isMain":false,"metricName":"# similar misassembled blocks"}]],["Reference statistics",[{"values":[131995,131995,131995,131995,131995,131995,131995,131995,131995,131995,131995,131995,131995],"quality":"Equal","isMain":false,"metricName":"Reference length"},{"values":[1,1,1,1,1,1,1,1,1,1,1,1,1],"quality":"Equal","isMain":false,"metricName":"Reference fragments"},{"values":["55.13","55.13","55.13","55.13","55.13","55.13","55.13","55.13","55.13","55.13","55.13","55.13","55.13"],"quality":"Equal","isMain":false,"metricName":"Reference GC (%)"}]]],"referenceName":"gi\_11038431\_ref\_NC","date":"18 July 2018, Wednesday, 16:07:45","order":[0,1,2,3,4,5,6,7,8,9,10,11,12],"assembliesNames":["ABySS\_127","ABySS\_63","CLC","IDBA\_UD","MEGAHIT","MIRA","SOAPdenovo2","SPAdes","SPAdes\_meta","SPAdes\_sc","SPAdes\_sc\_careful","Velvet","Geneious"]},{"assembliesWithNs":null,"minContig":500,"report":[["Genome statistics",[{"values":["18.477","11.254","71.537","62.884","66.211","31.571","66.185","76.773","74.965","75.681","76.727","7.885","7.102"],"quality":"More is better","isMain":true,"metricName":"Genome fraction (%)"},{"values":["1.000","1.000","1.007","1.009","1.000","1.032","1.005","1.003","1.004","1.009","1.005","1.000","1.063"],"quality":"Less is better","isMain":true,"metricName":"Duplication ratio"},{"values":[1466,1682,3647,3406,4353,1783,4182,4737,3647,4737,4737,3161,2884],"quality":"More is better","isMain":true,"metricName":"Largest alignment"},{"values":[18662,11366,72644,63534,66889,32705,67066,77574,76039,76467,77519,7964,7471],"quality":"More is better","isMain":true,"metricName":"Total aligned length"},{"values":[null,null,1093,973,788,null,806,1144,1144,1400,1400,null,null],"quality":"More is better","isMain":false,"metricName":"NG50"},{"values":[null,null,null,null,null,null,null,528,501,652,667,null,null],"quality":"More is better","isMain":false,"metricName":"NG75"},{"values":[631,841,1631,1548,1597,1163,1235,1665,1641,1660,1660,1543,null],"quality":"More is better","isMain":false,"metricName":"NA50"},{"values":[565,696,963,935,783,860,806,935,973,918,852,942,null],"quality":"More is better","isMain":false,"metricName":"NA75"},{"values":[null,null,1093,851,783,null,806,1144,1144,1144,1144,null,null],"quality":"More is better","isMain":true,"metricName":"NGA50"},{"values":[null,null,null,null,null,null,null,528,501,501,529,null,null],"quality":"More is better","isMain":false,"metricName":"NGA75"},{"values":[null,null,26,28,30,null,33,23,25,21,21,null,null],"quality":"Less is better","isMain":false,"metricName":"LG50"},{"values":[null,null,null,null,null,null,null,55,57,49,49,null,null],"quality":"Less is better","isMain":false,"metricName":"LG75"},{"values":[10,6,15,15,14,12,16,15,16,16,16,2,null],"quality":"Less is better","isMain":false,"metricName":"LA50"},{"values":[18,9,30,28,31,20,33,31,31,32,33,4,null],"quality":"Less is better","isMain":false,"metricName":"LA75"},{"values":[null,null,26,30,31,null,33,23,25,23,23,null,null],"quality":"Less is better","isMain":true,"metricName":"LGA50"},{"values":[null,null,null,null,null,null,null,55,57,55,54,null,null],"quality":"Less is better","isMain":false,"metricName":"LGA75"}]],["Misassemblies",[{"values":[0,0,0,0,0,1,0,0,0,0,0,0,0],"quality":"Less is better","isMain":true,"metricName":"# misassemblies"},{"values":[0,0,0,0,0,1,0,0,0,0,0,0,0],"quality":"Less is better","isMain":false,"metricName":" # relocations"},{"values":[0,0,0,0,0,0,0,0,0,0,0,0,0],"quality":"Less is better","isMain":false,"metricName":" # translocations"},{"values":[0,0,0,0,0,0,0,0,0,0,0,0,0],"quality":"Less is better","isMain":false,"metricName":" # inversions"},{"values":[0,0,0,0,0,1,0,0,0,0,0,0,0],"quality":"Less is better","isMain":false,"metricName":"# misassembled contigs"},{"values":[0,0,0,0,0,1183,0,0,0,0,0,0,0],"quality":"Less is better","isMain":true,"metricName":"Misassembled contigs length"},{"values":[0,0,0,0,0,1,0,0,0,0,0,0,1],"quality":"Less is better","isMain":false,"metricName":"# local misassemblies"},{"values":[0,0,0,0,0,0,0,0,0,0,0,0,0],"quality":"Less is better","isMain":false,"metricName":"# unaligned mis. contigs"}]],["Unaligned",[{"values":[0,0,0,0,0,0,0,0,0,0,0,0,0],"quality":"Less is better","isMain":false,"metricName":"# fully unaligned contigs"},{"values":[0,0,0,0,0,0,0,0,0,0,0,0,0],"quality":"Less is better","isMain":false,"metricName":"Fully unaligned length"},{"values":[0,0,0,1,1,0,0,0,0,1,1,0,1],"quality":"Less is better","isMain":false,"metricName":"# partially unaligned contigs"},{"values":[0,0,0,1669,513,0,0,0,0,2600,2600,0,9044],"quality":"Less is better","isMain":false,"metricName":"Partially unaligned length"}]],["Mismatches",[{"values":[5,1,79,67,45,51,38,70,64,66,71,1,30],"quality":"Less is better","isMain":false,"metricName":"# mismatches"},{"values":[0,0,2,1,0,8,2,0,0,2,5,0,0],"quality":"Less is better","isMain":false,"metricName":"# indels"},{"values":[0,0,2,1,0,10,100,0,0,3,12,0,0],"quality":"Less is better","isMain":false,"metricName":"Indels length"},{"values":["26.79","8.80","109.34","105.49","67.29","159.94","56.85","90.28","84.53","86.35","91.62","12.56","418.24"],"quality":"Less is better","isMain":true,"metricName":"# mismatches per 100 kbp"},{"values":["0.00","0.00","2.77","1.57","0.00","25.09","2.99","0.00","0.00","2.62","6.45","0.00","0.00"],"quality":"Less is better","isMain":true,"metricName":"# indels per 100 kbp"},{"values":[0,0,2,1,0,8,0,0,0,2,4,0,0],"quality":"Less is better","isMain":false,"metricName":" # indels (<= 5 bp)"},{"values":[0,0,0,0,0,0,2,0,0,0,1,0,0],"quality":"Less is better","isMain":false,"metricName":" # indels (> 5 bp)"},{"values":[0,0,0,0,0,74,184,0,0,0,0,0,107],"quality":"Less is better","isMain":false,"metricName":"# N's"},{"values":["0.00","0.00","0.00","0.00","0.00","224.77","273.80","0.00","0.00","0.00","0.00","0.00","641.83"],"quality":"Less is better","isMain":true,"metricName":"# N's per 100 kbp"}]],["Statistics without reference",[{"values":[26,14,56,49,56,30,59,58,57,56,57,6,4],"quality":"Equal","isMain":true,"metricName":"# contigs"},{"values":[4,3,27,25,21,17,23,28,30,30,30,3,4],"quality":"Equal","isMain":false,"metricName":"# contigs (>= 1000 bp)"},{"values":[0,0,0,0,0,0,0,0,0,0,0,0,1],"quality":"Equal","isMain":false,"metricName":"# contigs (>= 5000 bp)"},{"values":[0,0,0,0,0,0,0,0,0,0,0,0,0],"quality":"Equal","isMain":false,"metricName":"# contigs (>= 10000 bp)"},{"values":[0,0,0,0,0,0,0,0,0,0,0,0,0],"quality":"Equal","isMain":false,"metricName":"# contigs (>= 25000 bp)"},{"values":[0,0,0,0,0,0,0,0,0,0,0,0,0],"quality":"Equal","isMain":false,"metricName":"# contigs (>= 50000 bp)"},{"values":[1466,1682,3647,3406,4353,1849,4182,4737,3647,4737,4737,3161,9947],"quality":"More is better","isMain":true,"metricName":"Largest contig"},{"values":[18662,11366,72742,65722,67402,32922,67202,77786,76039,79702,80464,7964,16671],"quality":"More is better","isMain":true,"metricName":"Total length"},{"values":[5110,3818,52090,47964,43016,23219,41832,56321,56436,60954,60955,5797,16671],"quality":"More is better","isMain":true,"metricName":"Total length (>= 1000 bp)"},{"values":[0,0,0,0,0,0,0,0,0,0,0,0,9947],"quality":"More is better","isMain":false,"metricName":"Total length (>= 5000 bp)"},{"values":[0,0,0,0,0,0,0,0,0,0,0,0,0],"quality":"More is better","isMain":true,"metricName":"Total length (>= 10000 bp)"},{"values":[0,0,0,0,0,0,0,0,0,0,0,0,0],"quality":"More is better","isMain":false,"metricName":"Total length (>= 25000 bp)"},{"values":[0,0,0,0,0,0,0,0,0,0,0,0,0],"quality":"More is better","isMain":true,"metricName":"Total length (>= 50000 bp)"},{"values":[631,841,1631,1726,1597,1195,1235,1665,1641,1764,1764,1543,9947],"quality":"More is better","isMain":false,"metricName":"N50"},{"values":[565,696,963,974,788,908,806,935,973,1019,1007,942,2884],"quality":"More is better","isMain":false,"metricName":"N75"},{"values":[10,6,15,14,14,11,16,15,16,15,15,2,1],"quality":"Less is better","isMain":false,"metricName":"L50"},{"values":[18,9,30,27,30,19,33,31,31,29,30,4,2],"quality":"Less is better","isMain":false,"metricName":"L75"},{"values":["42.35","40.36","41.90","41.59","41.68","40.40","41.91","41.56","41.87","41.45","41.39","42.09","40.33"],"quality":"Equal","isMain":false,"metricName":"GC (%)"}]],["Predicted genes",[]],["Similarity statistics",[{"values":[0,0,0,0,0,0,0,0,0,0,0,0,0],"quality":"Equal","isMain":false,"metricName":"# similar correct contigs"},{"values":[0,0,0,0,0,0,0,0,0,0,0,0,0],"quality":"Equal","isMain":false,"metricName":"# similar misassembled blocks"}]],["Reference statistics",[{"values":[100999,100999,100999,100999,100999,100999,100999,100999,100999,100999,100999,100999,100999],"quality":"Equal","isMain":false,"metricName":"Reference length"},{"values":[1,1,1,1,1,1,1,1,1,1,1,1,1],"quality":"Equal","isMain":false,"metricName":"Reference fragments"},{"values":["40.69","40.69","40.69","40.69","40.69","40.69","40.69","40.69","40.69","40.69","40.69","40.69","40.69"],"quality":"Equal","isMain":false,"metricName":"Reference GC (%)"}]]],"referenceName":"gi\_11068003\_ref\_NC","date":"18 July 2018, Wednesday, 16:07:56","order":[0,1,2,3,4,5,6,7,8,9,10,11,12],"assembliesNames":["ABySS\_127","ABySS\_63","CLC","IDBA\_UD","MEGAHIT","MIRA","SOAPdenovo2","SPAdes","SPAdes\_meta","SPAdes\_sc","SPAdes\_sc\_careful","Velvet","Geneious"]},{"assembliesWithNs":null,"minContig":500,"report":[["Genome statistics",[{"values":["99.957","100.000","100.000","100.000","81.416","72.484","100.000","100.000","100.000","100.000","100.000"],"quality":"More is better","isMain":true,"metricName":"Genome fraction (%)"},{"values":["1.000","1.002","1.003","1.009","1.026","1.001","1.003","1.001","1.001","1.001","1.001"],"quality":"Less is better","isMain":true,"metricName":"Duplication ratio"},{"values":[43991,44109,44151,44402,12878,2436,44137,44065,44065,44065,39780],"quality":"More is better","isMain":true,"metricName":"Largest alignment"},{"values":[43991,44109,44151,44402,36747,31921,44137,44065,44065,44065,44061],"quality":"More is better","isMain":true,"metricName":"Total aligned length"},{"values":[44010,44109,44151,44402,7432,750,44137,44065,44065,44065,39780],"quality":"More is better","isMain":false,"metricName":"NG50"},{"values":[44010,44109,44151,44402,4051,null,44137,44065,44065,44065,39780],"quality":"More is better","isMain":false,"metricName":"NG75"},{"values":[43991,44109,44151,44402,9120,1219,44137,44065,44065,44065,39780],"quality":"More is better","isMain":false,"metricName":"NA50"},{"values":[43991,44109,44151,44402,7432,706,44137,44065,44065,44065,39780],"quality":"More is better","isMain":false,"metricName":"NA75"},{"values":[43991,44109,44151,44402,7432,750,44137,44065,44065,44065,39780],"quality":"More is better","isMain":true,"metricName":"NGA50"},{"values":[43991,44109,44151,44402,4051,null,44137,44065,44065,44065,39780],"quality":"More is better","isMain":false,"metricName":"NGA75"},{"values":[1,1,1,1,3,18,1,1,1,1,1],"quality":"Less is better","isMain":false,"metricName":"LG50"},{"values":[1,1,1,1,4,null,1,1,1,1,1],"quality":"Less is better","isMain":false,"metricName":"LG75"},{"values":[1,1,1,1,2,11,1,1,1,1,1],"quality":"Less is better","isMain":false,"metricName":"LA50"},{"values":[1,1,1,1,3,21,1,1,1,1,1],"quality":"Less is better","isMain":false,"metricName":"LA75"},{"values":[1,1,1,1,3,18,1,1,1,1,1],"quality":"Less is better","isMain":true,"metricName":"LGA50"},{"values":[1,1,1,1,4,null,1,1,1,1,1],"quality":"Less is better","isMain":false,"metricName":"LGA75"}]],["Misassemblies",[{"values":[0,0,0,0,0,0,0,0,0,0,0],"quality":"Less is better","isMain":true,"metricName":"# misassemblies"},{"values":[0,0,0,0,0,0,0,0,0,0,0],"quality":"Less is better","isMain":false,"metricName":" # relocations"},{"values":[0,0,0,0,0,0,0,0,0,0,0],"quality":"Less is better","isMain":false,"metricName":" # translocations"},{"values":[0,0,0,0,0,0,0,0,0,0,0],"quality":"Less is better","isMain":false,"metricName":" # inversions"},{"values":[0,0,0,0,0,0,0,0,0,0,0],"quality":"Less is better","isMain":false,"metricName":"# misassembled contigs"},{"values":[0,0,0,0,0,0,0,0,0,0,0],"quality":"Less is better","isMain":true,"metricName":"Misassembled contigs length"},{"values":[0,1,1,1,0,0,1,0,0,0,0],"quality":"Less is better","isMain":false,"metricName":"# local misassemblies"},{"values":[0,0,0,0,0,0,0,0,0,0,0],"quality":"Less is better","isMain":false,"metricName":"# unaligned mis. contigs"}]],["Unaligned",[{"values":[0,0,0,0,0,0,0,0,0,0,0],"quality":"Less is better","isMain":false,"metricName":"# fully unaligned contigs"},{"values":[0,0,0,0,0,0,0,0,0,0,0],"quality":"Less is better","isMain":false,"metricName":"Fully unaligned length"},{"values":[0,0,0,0,0,0,0,0,0,0,0],"quality":"Less is better","isMain":false,"metricName":"# partially unaligned contigs"},{"values":[0,0,0,0,0,0,0,0,0,0,0],"quality":"Less is better","isMain":false,"metricName":"Partially unaligned length"}]],["Mismatches",[{"values":[0,0,0,0,0,2,0,0,0,0,0],"quality":"Less is better","isMain":false,"metricName":"# mismatches"},{"values":[0,0,0,0,0,0,0,1,1,1,0],"quality":"Less is better","isMain":false,"metricName":"# indels"},{"values":[0,0,0,0,0,0,0,55,55,55,0],"quality":"Less is better","isMain":false,"metricName":"Indels length"},{"values":["0.00","0.00","0.00","0.00","0.00","6.27","0.00","0.00","0.00","0.00","0.00"],"quality":"Less is better","isMain":true,"metricName":"# mismatches per 100 kbp"},{"values":["0.00","0.00","0.00","0.00","0.00","0.00","0.00","2.27","2.27","2.27","0.00"],"quality":"Less is better","isMain":true,"metricName":"# indels per 100 kbp"},{"values":[0,0,0,0,0,0,0,0,0,0,0],"quality":"Less is better","isMain":false,"metricName":" # indels (<= 5 bp)"},{"values":[0,0,0,0,0,0,0,1,1,1,0],"quality":"Less is better","isMain":false,"metricName":" # indels (> 5 bp)"},{"values":[0,0,0,0,0,0,0,0,0,0,0],"quality":"Less is better","isMain":false,"metricName":"# N's"},{"values":["0.00","0.00","0.00","0.00","0.00","0.00","0.00","0.00","0.00","0.00","0.00"],"quality":"Less is better","isMain":true,"metricName":"# N's per 100 kbp"}]],["Statistics without reference",[{"values":[1,1,1,1,5,33,1,1,1,1,2],"quality":"Equal","isMain":true,"metricName":"# contigs"},{"values":[1,1,1,1,5,12,1,1,1,1,2],"quality":"Equal","isMain":false,"metricName":"# contigs (>= 1000 bp)"},{"values":[1,1,1,1,3,0,1,1,1,1,1],"quality":"Equal","isMain":false,"metricName":"# contigs (>= 5000 bp)"},{"values":[1,1,1,1,1,0,1,1,1,1,1],"quality":"Equal","isMain":false,"metricName":"# contigs (>= 10000 bp)"},{"values":[1,1,1,1,0,0,1,1,1,1,1],"quality":"Equal","isMain":false,"metricName":"# contigs (>= 25000 bp)"},{"values":[0,0,0,0,0,0,0,0,0,0,0],"quality":"Equal","isMain":false,"metricName":"# contigs (>= 50000 bp)"},{"values":[44010,44109,44151,44402,12878,2436,44137,44065,44065,44065,39780],"quality":"More is better","isMain":true,"metricName":"Largest contig"},{"values":[44010,44109,44151,44402,36747,31921,44137,44065,44065,44065,44061],"quality":"More is better","isMain":true,"metricName":"Total length"},{"values":[44010,44109,44151,44402,36747,17212,44137,44065,44065,44065,44061],"quality":"More is better","isMain":true,"metricName":"Total length (>= 1000 bp)"},{"values":[44010,44109,44151,44402,29430,0,44137,44065,44065,44065,39780],"quality":"More is better","isMain":false,"metricName":"Total length (>= 5000 bp)"},{"values":[44010,44109,44151,44402,12878,0,44137,44065,44065,44065,39780],"quality":"More is better","isMain":true,"metricName":"Total length (>= 10000 bp)"},{"values":[44010,44109,44151,44402,0,0,44137,44065,44065,44065,39780],"quality":"More is better","isMain":false,"metricName":"Total length (>= 25000 bp)"},{"values":[0,0,0,0,0,0,0,0,0,0,0],"quality":"More is better","isMain":true,"metricName":"Total length (>= 50000 bp)"},{"values":[44010,44109,44151,44402,9120,1219,44137,44065,44065,44065,39780],"quality":"More is better","isMain":false,"metricName":"N50"},{"values":[44010,44109,44151,44402,7432,706,44137,44065,44065,44065,39780],"quality":"More is better","isMain":false,"metricName":"N75"},{"values":[1,1,1,1,2,11,1,1,1,1,1],"quality":"Less is better","isMain":false,"metricName":"L50"},{"values":[1,1,1,1,3,21,1,1,1,1,1],"quality":"Less is better","isMain":false,"metricName":"L75"},{"values":["43.73","43.72","43.71","43.68","43.79","43.86","43.73","43.74","43.74","43.74","43.71"],"quality":"Equal","isMain":false,"metricName":"GC (%)"}]],["Predicted genes",[]],["Similarity statistics",[{"values":[0,0,0,0,0,0,0,0,0,0,0],"quality":"Equal","isMain":false,"metricName":"# similar correct contigs"},{"values":[0,0,0,0,0,0,0,0,0,0,0],"quality":"Equal","isMain":false,"metricName":"# similar misassembled blocks"}]],["Reference statistics",[{"values":[44010,44010,44010,44010,44010,44010,44010,44010,44010,44010,44010],"quality":"Equal","isMain":false,"metricName":"Reference length"},{"values":[1,1,1,1,1,1,1,1,1,1,1],"quality":"Equal","isMain":false,"metricName":"Reference fragments"},{"values":["43.72","43.72","43.72","43.72","43.72","43.72","43.72","43.72","43.72","43.72","43.72"],"quality":"Equal","isMain":false,"metricName":"Reference GC (%)"}]]],"referenceName":"gi\_111146921\_ref\_NC","date":"18 July 2018, Wednesday, 16:08:06","order":[0,1,2,3,4,5,6,7,8,9,10],"assembliesNames":["CLC","IDBA\_UD","MEGAHIT","MIRA","Ray\_Meta","SOAPdenovo2","SPAdes","SPAdes\_meta","SPAdes\_sc","SPAdes\_sc\_careful","Velvet"]},{"assembliesWithNs":null,"minContig":500,"report":[["Genome statistics",[{"values":["46.803","38.990","74.793","89.047","88.046","79.467","40.442","84.962","88.080","86.513","78.610","54.854","36.528"],"quality":"More is better","isMain":true,"metricName":"Genome fraction (%)"},{"values":["1.004","1.005","1.042","1.008","1.035","1.045","1.062","1.005","1.012","1.014","1.014","1.002","1.004"],"quality":"Less is better","isMain":true,"metricName":"Duplication ratio"},{"values":[3429,3422,5855,10014,7581,5869,2180,15821,14710,15405,11808,5276,10963],"quality":"More is better","isMain":true,"metricName":"Largest alignment"},{"values":[92950,77553,154157,177222,179942,164423,80652,168612,175822,172514,156664,108592,72256],"quality":"More is better","isMain":true,"metricName":"Total aligned length"},{"values":[null,null,2231,4397,2616,2356,null,5495,4862,3194,2048,654,null],"quality":"More is better","isMain":false,"metricName":"NG50"},{"values":[null,null,1013,2113,1515,992,null,2570,2290,1547,692,null,null],"quality":"More is better","isMain":false,"metricName":"NG75"},{"values":[971,828,1846,4541,2768,2581,811,6932,6933,3467,2571,1656,5660],"quality":"More is better","isMain":false,"metricName":"NA50"},{"values":[702,677,700,2321,1811,1725,611,3467,2576,2048,1498,983,3986],"quality":"More is better","isMain":false,"metricName":"NA75"},{"values":[null,null,1777,4397,2505,2356,null,5487,4844,3188,2048,651,null],"quality":"More is better","isMain":true,"metricName":"NGA50"},{"values":[null,null,578,1996,1515,992,null,2392,2057,1173,668,null,null],"quality":"More is better","isMain":false,"metricName":"NGA75"},{"values":[null,null,21,15,23,30,null,11,11,17,26,65,null],"quality":"Less is better","isMain":false,"metricName":"LG50"},{"values":[null,null,51,33,47,61,null,24,27,37,68,null,null],"quality":"Less is better","isMain":false,"metricName":"LG75"},{"values":[33,32,31,13,20,23,39,9,9,15,17,23,5],"quality":"Less is better","isMain":false,"metricName":"LA50"},{"values":[61,58,69,27,40,42,73,18,21,31,37,44,8],"quality":"Less is better","isMain":false,"metricName":"LA75"},{"values":[null,null,35,15,23,30,null,11,11,18,26,66,null],"quality":"Less is better","isMain":true,"metricName":"LGA50"},{"values":[null,null,89,34,47,61,null,25,27,41,70,null,null],"quality":"Less is better","isMain":false,"metricName":"LGA75"}]],["Misassemblies",[{"values":[0,0,0,0,0,0,0,0,0,0,0,0,0],"quality":"Less is better","isMain":true,"metricName":"# misassemblies"},{"values":[0,0,0,0,0,0,0,0,0,0,0,0,0],"quality":"Less is better","isMain":false,"metricName":" # relocations"},{"values":[0,0,0,0,0,0,0,0,0,0,0,0,0],"quality":"Less is better","isMain":false,"metricName":" # translocations"},{"values":[0,0,0,0,0,0,0,0,0,0,0,0,0],"quality":"Less is better","isMain":false,"metricName":" # inversions"},{"values":[0,0,0,0,0,0,0,0,0,0,0,0,0],"quality":"Less is better","isMain":false,"metricName":"# misassembled contigs"},{"values":[0,0,0,0,0,0,0,0,0,0,0,0,0],"quality":"Less is better","isMain":true,"metricName":"Misassembled contigs length"},{"values":[0,0,1,1,1,0,0,1,1,2,1,1,0],"quality":"Less is better","isMain":false,"metricName":"# local misassemblies"},{"values":[0,0,1,0,0,0,1,0,0,0,0,0,0],"quality":"Less is better","isMain":false,"metricName":"# unaligned mis. contigs"}]],["Unaligned",[{"values":[0,0,0,0,0,0,0,0,0,0,0,0,0],"quality":"Less is better","isMain":false,"metricName":"# fully unaligned contigs"},{"values":[0,0,0,0,0,0,0,0,0,0,0,0,0],"quality":"Less is better","isMain":false,"metricName":"Fully unaligned length"},{"values":[0,0,2,1,1,0,5,2,1,2,0,0,0],"quality":"Less is better","isMain":false,"metricName":"# partially unaligned contigs"},{"values":[0,0,26563,861,786,0,6824,1847,1061,3193,0,0,0],"quality":"Less is better","isMain":false,"metricName":"Partially unaligned length"}]],["Mismatches",[{"values":[58,43,570,424,539,284,127,541,588,538,447,153,162],"quality":"Less is better","isMain":false,"metricName":"# mismatches"},{"values":[3,0,87,42,59,44,36,54,64,61,47,5,27],"quality":"Less is better","isMain":false,"metricName":"# indels"},{"values":[8,0,315,124,187,139,1665,137,194,202,155,80,183],"quality":"Less is better","isMain":false,"metricName":"Indels length"},{"values":["62.57","55.69","384.80","240.42","309.10","180.45","158.56","321.51","337.07","314.00","287.11","140.83","223.93"],"quality":"Less is better","isMain":true,"metricName":"# mismatches per 100 kbp"},{"values":["3.24","0.00","58.73","23.82","33.83","27.96","44.95","32.09","36.69","35.60","30.19","4.60","37.32"],"quality":"Less is better","isMain":true,"metricName":"# indels per 100 kbp"},{"values":[3,0,78,38,54,38,1,51,58,54,42,3,21],"quality":"Less is better","isMain":false,"metricName":" # indels (<= 5 bp)"},{"values":[0,0,9,4,5,6,35,3,6,7,5,2,6],"quality":"Less is better","isMain":false,"metricName":" # indels (> 5 bp)"},{"values":[0,0,0,0,0,514,5683,0,422,1371,730,40,421],"quality":"Less is better","isMain":false,"metricName":"# N's"},{"values":["0.00","0.00","0.00","0.00","0.00","312.48","6182.21","0.00","237.73","774.97","462.39","36.74","579.90"],"quality":"Less is better","isMain":true,"metricName":"# N's per 100 kbp"}]],["Statistics without reference",[{"values":[99,91,98,59,80,83,98,37,48,71,83,82,16],"quality":"Equal","isMain":true,"metricName":"# contigs"},{"values":[32,20,51,49,60,59,28,36,41,48,49,44,16],"quality":"Equal","isMain":false,"metricName":"# contigs (>= 1000 bp)"},{"values":[0,0,5,11,6,2,0,12,10,8,6,1,7],"quality":"Equal","isMain":false,"metricName":"# contigs (>= 5000 bp)"},{"values":[0,0,1,1,0,0,0,4,4,1,1,0,1],"quality":"Equal","isMain":false,"metricName":"# contigs (>= 10000 bp)"},{"values":[0,0,1,0,0,0,0,0,0,0,0,0,0],"quality":"Equal","isMain":false,"metricName":"# contigs (>= 25000 bp)"},{"values":[0,0,0,0,0,0,0,0,0,0,0,0,0],"quality":"Equal","isMain":false,"metricName":"# contigs (>= 50000 bp)"},{"values":[3429,3422,30159,10014,7581,5869,3228,15821,14710,15405,11808,5276,10963],"quality":"More is better","isMain":true,"metricName":"Largest contig"},{"values":[93031,77587,180960,178713,181248,164488,91925,170962,177514,176909,157876,108859,72599],"quality":"More is better","isMain":true,"metricName":"Total length"},{"values":[46126,28653,148981,170826,166023,146727,42302,170346,172102,161281,133587,82361,72599],"quality":"More is better","isMain":true,"metricName":"Total length (>= 1000 bp)"},{"values":[0,0,52425,82427,39575,11050,0,107808,96297,66553,43343,5276,51070],"quality":"More is better","isMain":false,"metricName":"Total length (>= 5000 bp)"},{"values":[0,0,30159,10014,0,0,0,48840,49362,15405,11808,0,10963],"quality":"More is better","isMain":true,"metricName":"Total length (>= 10000 bp)"},{"values":[0,0,30159,0,0,0,0,0,0,0,0,0,0],"quality":"More is better","isMain":false,"metricName":"Total length (>= 25000 bp)"},{"values":[0,0,0,0,0,0,0,0,0,0,0,0,0],"quality":"More is better","isMain":true,"metricName":"Total length (>= 50000 bp)"},{"values":[971,828,2584,4541,2845,2581,914,6932,6933,3602,2724,1656,5660],"quality":"More is better","isMain":false,"metricName":"N50"},{"values":[702,689,1471,2321,1869,1725,708,3470,2739,2127,1498,1014,3986],"quality":"More is better","isMain":false,"metricName":"N75"},{"values":[33,32,17,13,20,23,32,9,9,14,17,23,5],"quality":"Less is better","isMain":false,"metricName":"L50"},{"values":[61,58,41,27,39,42,61,18,21,29,37,44,8],"quality":"Less is better","isMain":false,"metricName":"L75"},{"values":["33.31","33.49","32.42","33.16","33.29","33.22","33.18","33.27","33.31","33.20","33.25","33.15","33.29"],"quality":"Equal","isMain":false,"metricName":"GC (%)"}]],["Predicted genes",[]],["Similarity statistics",[{"values":[0,0,0,0,0,0,0,0,0,0,0,0,0],"quality":"Equal","isMain":false,"metricName":"# similar correct contigs"},{"values":[0,0,0,0,0,0,0,0,0,0,0,0,0],"quality":"Equal","isMain":false,"metricName":"# similar misassembled blocks"}]],["Reference statistics",[{"values":[198050,198050,198050,198050,198050,198050,198050,198050,198050,198050,198050,198050,198050],"quality":"Equal","isMain":false,"metricName":"Reference length"},{"values":[1,1,1,1,1,1,1,1,1,1,1,1,1],"quality":"Equal","isMain":false,"metricName":"Reference fragments"},{"values":["33.26","33.26","33.26","33.26","33.26","33.26","33.26","33.26","33.26","33.26","33.26","33.26","33.26"],"quality":"Equal","isMain":false,"metricName":"Reference GC (%)"}]]],"referenceName":"gi\_113195177\_ref\_NC","date":"18 July 2018, Wednesday, 16:08:18","order":[0,1,2,3,4,5,6,7,8,9,10,11,12],"assembliesNames":["ABySS\_127","ABySS\_63","CLC","IDBA\_UD","MEGAHIT","MIRA","SOAPdenovo2","SPAdes","SPAdes\_meta","SPAdes\_sc","SPAdes\_sc\_careful","Velvet","Geneious"]},{"assembliesWithNs":null,"minContig":500,"report":[["Genome statistics",[{"values":["18.954","12.201","76.896","71.552","76.463","41.590","69.176","83.007","83.825","83.656","83.788","9.004","1.112"],"quality":"More is better","isMain":true,"metricName":"Genome fraction (%)"},{"values":["1.001","1.001","1.018","1.004","1.001","1.005","1.004","1.004","1.001","1.003","1.004","1.000","1.000"],"quality":"Less is better","isMain":true,"metricName":"Duplication ratio"},{"values":[1309,1117,5124,4865,4799,2919,5061,5124,5124,5124,5124,3008,1868],"quality":"More is better","isMain":true,"metricName":"Largest alignment"},{"values":[31872,20518,130706,120379,128559,70248,116455,139525,141015,140601,140923,15130,1868],"quality":"More is better","isMain":true,"metricName":"Total aligned length"},{"values":[null,null,1259,1321,1360,null,994,1831,1653,1787,1882,null,null],"quality":"More is better","isMain":false,"metricName":"NG50"},{"values":[null,null,667,566,554,null,null,905,832,891,905,null,null],"quality":"More is better","isMain":false,"metricName":"NG75"},{"values":[713,743,1460,1699,1799,1199,1409,1914,1983,1945,1983,1667,1868],"quality":"More is better","isMain":false,"metricName":"NA50"},{"values":[587,631,868,1039,1031,794,891,1113,1255,1210,1210,1072,1868],"quality":"More is better","isMain":false,"metricName":"NA75"},{"values":[null,null,1121,1231,1289,null,966,1653,1653,1667,1667,null,null],"quality":"More is better","isMain":true,"metricName":"NGA50"},{"values":[null,null,572,null,554,null,null,828,832,832,854,null,null],"quality":"More is better","isMain":false,"metricName":"NGA75"},{"values":[null,null,34,34,37,null,47,29,31,30,30,null,null],"quality":"Less is better","isMain":false,"metricName":"LG50"},{"values":[null,null,81,80,87,null,null,61,67,63,61,null,null],"quality":"Less is better","isMain":false,"metricName":"LG75"},{"values":[18,12,25,25,25,22,26,25,24,25,25,4,1],"quality":"Less is better","isMain":false,"metricName":"LA50"},{"values":[30,19,56,49,48,40,51,50,47,49,49,7,1],"quality":"Less is better","isMain":false,"metricName":"LA75"},{"values":[null,null,38,39,38,null,48,32,31,31,31,null,null],"quality":"Less is better","isMain":true,"metricName":"LGA50"},{"values":[null,null,90,null,87,null,null,68,67,67,66,null,null],"quality":"Less is better","isMain":false,"metricName":"LGA75"}]],["Misassemblies",[{"values":[0,0,0,0,1,0,0,0,0,0,0,0,0],"quality":"Less is better","isMain":true,"metricName":"# misassemblies"},{"values":[0,0,0,0,1,0,0,0,0,0,0,0,0],"quality":"Less is better","isMain":false,"metricName":" # relocations"},{"values":[0,0,0,0,0,0,0,0,0,0,0,0,0],"quality":"Less is better","isMain":false,"metricName":" # translocations"},{"values":[0,0,0,0,0,0,0,0,0,0,0,0,0],"quality":"Less is better","isMain":false,"metricName":" # inversions"},{"values":[0,0,0,0,1,0,0,0,0,0,0,0,0],"quality":"Less is better","isMain":false,"metricName":"# misassembled contigs"},{"values":[0,0,0,0,1648,0,0,0,0,0,0,0,0],"quality":"Less is better","isMain":true,"metricName":"Misassembled contigs length"},{"values":[0,0,0,0,0,0,0,0,0,0,0,0,0],"quality":"Less is better","isMain":false,"metricName":"# local misassemblies"},{"values":[0,0,0,0,0,0,0,0,0,0,0,0,0],"quality":"Less is better","isMain":false,"metricName":"# unaligned mis. contigs"}]],["Unaligned",[{"values":[0,0,0,0,0,0,0,0,0,0,0,0,0],"quality":"Less is better","isMain":false,"metricName":"# fully unaligned contigs"},{"values":[0,0,0,0,0,0,0,0,0,0,0,0,0],"quality":"Less is better","isMain":false,"metricName":"Fully unaligned length"},{"values":[0,0,3,2,0,0,0,3,0,3,4,0,0],"quality":"Less is better","isMain":false,"metricName":"# partially unaligned contigs"},{"values":[0,0,4426,6381,0,0,0,5228,0,3361,4612,0,0],"quality":"Less is better","isMain":false,"metricName":"Partially unaligned length"}]],["Mismatches",[{"values":[5,3,180,198,96,21,92,180,134,146,165,4,0],"quality":"Less is better","isMain":false,"metricName":"# mismatches"},{"values":[0,0,2,2,0,0,4,2,0,2,4,0,0],"quality":"Less is better","isMain":false,"metricName":"# indels"},{"values":[0,0,2,2,0,0,183,3,0,3,5,0,0],"quality":"Less is better","isMain":false,"metricName":"Indels length"},{"values":["15.70","14.63","139.30","164.67","74.71","30.05","79.14","129.05","95.13","103.86","117.19","26.44","0.00"],"quality":"Less is better","isMain":true,"metricName":"# mismatches per 100 kbp"},{"values":["0.00","0.00","1.55","1.66","0.00","0.00","3.44","1.43","0.00","1.42","2.84","0.00","0.00"],"quality":"Less is better","isMain":true,"metricName":"# indels per 100 kbp"},{"values":[0,0,2,2,0,0,0,2,0,2,4,0,0],"quality":"Less is better","isMain":false,"metricName":" # indels (<= 5 bp)"},{"values":[0,0,0,0,0,0,4,0,0,0,0,0,0],"quality":"Less is better","isMain":false,"metricName":" # indels (> 5 bp)"},{"values":[0,0,0,0,0,94,383,0,0,0,0,0,3],"quality":"Less is better","isMain":false,"metricName":"# N's"},{"values":["0.00","0.00","0.00","0.00","0.00","133.81","328.16","0.00","0.00","0.00","0.00","0.00","160.60"],"quality":"Less is better","isMain":true,"metricName":"# N's per 100 kbp"}]],["Statistics without reference",[{"values":[44,28,98,81,91,67,93,87,88,87,87,12,1],"quality":"Equal","isMain":true,"metricName":"# contigs"},{"values":[7,2,47,51,49,27,46,56,56,57,57,7,1],"quality":"Equal","isMain":false,"metricName":"# contigs (>= 1000 bp)"},{"values":[0,0,3,1,0,0,1,3,2,3,3,0,0],"quality":"Equal","isMain":false,"metricName":"# contigs (>= 5000 bp)"},{"values":[0,0,0,0,0,0,0,0,0,0,0,0,0],"quality":"Equal","isMain":false,"metricName":"# contigs (>= 10000 bp)"},{"values":[0,0,0,0,0,0,0,0,0,0,0,0,0],"quality":"Equal","isMain":false,"metricName":"# contigs (>= 25000 bp)"},{"values":[0,0,0,0,0,0,0,0,0,0,0,0,0],"quality":"Equal","isMain":false,"metricName":"# contigs (>= 50000 bp)"},{"values":[1309,1117,5225,8204,4799,2919,5061,5225,5124,5225,5225,3008,1868],"quality":"More is better","isMain":true,"metricName":"Largest contig"},{"values":[31872,20518,135921,127072,128559,70248,116713,145300,141015,144344,145917,15130,1868],"quality":"More is better","isMain":true,"metricName":"Total length"},{"values":[8011,2146,98622,103951,98042,41693,83162,121415,116728,121219,122792,12140,1868],"quality":"More is better","isMain":true,"metricName":"Total length (>= 1000 bp)"},{"values":[0,0,15410,8204,0,0,5061,15410,10185,15410,15410,0,0],"quality":"More is better","isMain":false,"metricName":"Total length (>= 5000 bp)"},{"values":[0,0,0,0,0,0,0,0,0,0,0,0,0],"quality":"More is better","isMain":true,"metricName":"Total length (>= 10000 bp)"},{"values":[0,0,0,0,0,0,0,0,0,0,0,0,0],"quality":"More is better","isMain":false,"metricName":"Total length (>= 25000 bp)"},{"values":[0,0,0,0,0,0,0,0,0,0,0,0,0],"quality":"More is better","isMain":true,"metricName":"Total length (>= 50000 bp)"},{"values":[713,743,1653,1891,1799,1199,1409,2025,1983,2007,2025,1667,1868],"quality":"More is better","isMain":false,"metricName":"N50"},{"values":[587,631,928,1135,1031,794,891,1259,1255,1311,1311,1072,1868],"quality":"More is better","isMain":false,"metricName":"N75"},{"values":[18,12,23,21,25,22,26,23,24,24,24,4,1],"quality":"Less is better","isMain":false,"metricName":"L50"},{"values":[30,19,51,43,48,40,51,46,47,46,46,7,1],"quality":"Less is better","isMain":false,"metricName":"L75"},{"values":["48.14","47.70","48.49","48.76","48.57","48.58","48.42","48.62","48.52","48.45","48.51","48.29","45.36"],"quality":"Equal","isMain":false,"metricName":"GC (%)"}]],["Predicted genes",[]],["Similarity statistics",[{"values":[0,0,0,0,0,0,0,0,0,0,0,0,0],"quality":"Equal","isMain":false,"metricName":"# similar correct contigs"},{"values":[0,0,0,0,0,0,0,0,0,0,0,0,0],"quality":"Equal","isMain":false,"metricName":"# similar misassembled blocks"}]],["Reference statistics",[{"values":[168041,168041,168041,168041,168041,168041,168041,168041,168041,168041,168041,168041,168041],"quality":"Equal","isMain":false,"metricName":"Reference length"},{"values":[1,1,1,1,1,1,1,1,1,1,1,1,1],"quality":"Equal","isMain":false,"metricName":"Reference fragments"},{"values":["48.57","48.57","48.57","48.57","48.57","48.57","48.57","48.57","48.57","48.57","48.57","48.57","48.57"],"quality":"Equal","isMain":false,"metricName":"Reference GC (%)"}]]],"referenceName":"gi\_114679849\_ref\_NC","date":"18 July 2018, Wednesday, 16:08:30","order":[0,1,2,3,4,5,6,7,8,9,10,11,12],"assembliesNames":["ABySS\_127","ABySS\_63","CLC","IDBA\_UD","MEGAHIT","MIRA","SOAPdenovo2","SPAdes","SPAdes\_meta","SPAdes\_sc","SPAdes\_sc\_careful","Velvet","Geneious"]},{"assembliesWithNs":null,"minContig":500,"report":[["Genome statistics",[{"values":["20.269","14.013","19.970","2.629","7.303","22.118","18.030","21.348","21.014"],"quality":"More is better","isMain":true,"metricName":"Genome fraction (%)"},{"values":["1.083","1.008","1.002","1.000","1.001","1.020","1.019","1.031","1.031"],"quality":"Less is better","isMain":true,"metricName":"Duplication ratio"},{"values":[1629,1552,1516,1008,1497,1687,1687,2247,2247],"quality":"More is better","isMain":true,"metricName":"Largest alignment"},{"values":[28952,18857,26874,3534,9826,29739,24248,28693,28243],"quality":"More is better","isMain":true,"metricName":"Total aligned length"},{"values":[1041,844,1050,987,888,1041,860,1126,1123],"quality":"More is better","isMain":false,"metricName":"NA50"},{"values":[687,621,728,810,630,809,654,781,781],"quality":"More is better","isMain":false,"metricName":"NA75"},{"values":[null,null,null,null,null,null,null,null,null],"quality":"More is better","isMain":true,"metricName":"NGA50"},{"values":[12,9,11,2,5,12,12,11,11],"quality":"Less is better","isMain":false,"metricName":"LA50"},{"values":[21,16,19,3,8,21,19,19,19],"quality":"Less is better","isMain":false,"metricName":"LA75"}]],["Misassemblies",[{"values":[1,0,0,0,0,0,0,0,0],"quality":"Less is better","isMain":true,"metricName":"# misassemblies"},{"values":[1,0,0,0,0,0,0,0,0],"quality":"Less is better","isMain":false,"metricName":" # relocations"},{"values":[0,0,0,0,0,0,0,0,0],"quality":"Less is better","isMain":false,"metricName":" # translocations"},{"values":[0,0,0,0,0,0,0,0,0],"quality":"Less is better","isMain":false,"metricName":" # inversions"},{"values":[1,0,0,0,0,0,0,0,0],"quality":"Less is better","isMain":false,"metricName":"# misassembled contigs"},{"values":[723,0,0,0,0,0,0,0,0],"quality":"Less is better","isMain":true,"metricName":"Misassembled contigs length"},{"values":[0,0,0,0,0,1,0,1,1],"quality":"Less is better","isMain":false,"metricName":"# local misassemblies"},{"values":[0,0,0,0,0,0,0,0,0],"quality":"Less is better","isMain":false,"metricName":"# unaligned mis. contigs"}]],["Unaligned",[{"values":[0,0,0,0,0,0,0,0,0],"quality":"Less is better","isMain":false,"metricName":"# fully unaligned contigs"},{"values":[0,0,0,0,0,0,0,0,0],"quality":"Less is better","isMain":false,"metricName":"Fully unaligned length"},{"values":[1,1,0,0,0,1,0,1,1],"quality":"Less is better","isMain":false,"metricName":"# partially unaligned contigs"},{"values":[698,620,0,0,0,755,0,1565,1565],"quality":"Less is better","isMain":false,"metricName":"Partially unaligned length"}]],["Mismatches",[{"values":[354,189,343,12,114,422,184,388,372],"quality":"Less is better","isMain":false,"metricName":"# mismatches"},{"values":[9,8,19,0,2,19,5,19,19],"quality":"Less is better","isMain":false,"metricName":"# indels"},{"values":[31,31,47,0,9,62,18,57,57],"quality":"Less is better","isMain":false,"metricName":"Indels length"},{"values":["1299.32","1003.40","1277.80","339.56","1161.25","1419.39","759.20","1352.10","1317.00"],"quality":"Less is better","isMain":true,"metricName":"# mismatches per 100 kbp"},{"values":["33.03","42.47","70.78","0.00","20.37","63.91","20.63","66.21","67.27"],"quality":"Less is better","isMain":true,"metricName":"# indels per 100 kbp"},{"values":[7,6,17,0,1,16,3,16,16],"quality":"Less is better","isMain":false,"metricName":" # indels (<= 5 bp)"},{"values":[2,2,2,0,1,3,2,3,3],"quality":"Less is better","isMain":false,"metricName":" # indels (> 5 bp)"},{"values":[0,0,0,28,0,0,0,0,0],"quality":"Less is better","isMain":false,"metricName":"# N's"},{"values":["0.00","0.00","0.00","792.30","0.00","0.00","0.00","0.00","0.00"],"quality":"Less is better","isMain":true,"metricName":"# N's per 100 kbp"}]],["Statistics without reference",[{"values":[32,23,30,4,12,30,29,27,27],"quality":"Equal","isMain":true,"metricName":"# contigs"},{"values":[12,6,12,1,2,14,6,17,16],"quality":"Equal","isMain":false,"metricName":"# contigs (>= 1000 bp)"},{"values":[0,0,0,0,0,0,0,0,0],"quality":"Equal","isMain":false,"metricName":"# contigs (>= 5000 bp)"},{"values":[0,0,0,0,0,0,0,0,0],"quality":"Equal","isMain":false,"metricName":"# contigs (>= 10000 bp)"},{"values":[0,0,0,0,0,0,0,0,0],"quality":"Equal","isMain":false,"metricName":"# contigs (>= 25000 bp)"},{"values":[0,0,0,0,0,0,0,0,0],"quality":"Equal","isMain":false,"metricName":"# contigs (>= 50000 bp)"},{"values":[1629,1552,1516,1008,1497,1846,1687,2247,2247],"quality":"More is better","isMain":true,"metricName":"Largest contig"},{"values":[30194,19604,26884,3534,9826,31080,24688,31143,30693],"quality":"More is better","isMain":true,"metricName":"Total length"},{"values":[15571,7353,14634,1008,2806,18990,7611,23762,22460],"quality":"More is better","isMain":true,"metricName":"Total length (>= 1000 bp)"},{"values":[0,0,0,0,0,0,0,0,0],"quality":"More is better","isMain":false,"metricName":"Total length (>= 5000 bp)"},{"values":[0,0,0,0,0,0,0,0,0],"quality":"More is better","isMain":true,"metricName":"Total length (>= 10000 bp)"},{"values":[0,0,0,0,0,0,0,0,0],"quality":"More is better","isMain":false,"metricName":"Total length (>= 25000 bp)"},{"values":[0,0,0,0,0,0,0,0,0],"quality":"More is better","isMain":true,"metricName":"Total length (>= 50000 bp)"},{"values":[1041,963,1053,987,888,1123,866,1271,1264],"quality":"More is better","isMain":false,"metricName":"N50"},{"values":[809,651,728,810,630,866,716,1001,945],"quality":"More is better","isMain":false,"metricName":"N75"},{"values":[12,9,11,2,5,11,12,10,10],"quality":"Less is better","isMain":false,"metricName":"L50"},{"values":[20,15,19,3,8,19,19,17,17],"quality":"Less is better","isMain":false,"metricName":"L75"},{"values":["40.03","40.42","40.15","42.16","39.87","40.18","39.75","40.73","40.69"],"quality":"Equal","isMain":false,"metricName":"GC (%)"}]],["Predicted genes",[]],["Similarity statistics",[{"values":[0,0,0,0,0,0,0,0,0],"quality":"Equal","isMain":false,"metricName":"# similar correct contigs"},{"values":[0,0,0,0,0,0,0,0,0],"quality":"Equal","isMain":false,"metricName":"# similar misassembled blocks"}]],["Reference statistics",[{"values":[134417,134417,134417,134417,134417,134417,134417,134417,134417],"quality":"Equal","isMain":false,"metricName":"Reference length"},{"values":[1,1,1,1,1,1,1,1,1],"quality":"Equal","isMain":false,"metricName":"Reference fragments"},{"values":["40.69","40.69","40.69","40.69","40.69","40.69","40.69","40.69","40.69"],"quality":"Equal","isMain":false,"metricName":"Reference GC (%)"}]]],"referenceName":"gi\_114680053\_ref\_NC","date":"18 July 2018, Wednesday, 16:08:38","order":[0,1,2,3,4,5,6,7,8],"assembliesNames":["CLC","IDBA\_UD","MEGAHIT","MIRA","SOAPdenovo2","SPAdes","SPAdes\_meta","SPAdes\_sc","SPAdes\_sc\_careful"]},{"assembliesWithNs":null,"minContig":500,"report":[["Genome statistics",[{"values":["9.538","3.947","11.464","6.660","12.166","12.933","12.401","12.401"],"quality":"More is better","isMain":true,"metricName":"Genome fraction (%)"},{"values":["1.000","1.048","1.002","1.000","1.000","1.004","1.009","1.009"],"quality":"Less is better","isMain":true,"metricName":"Duplication ratio"},{"values":[1291,1147,1084,1291,1291,1897,1291,1291],"quality":"More is better","isMain":true,"metricName":"Largest alignment"},{"values":[14967,6202,18014,10451,19091,20345,19482,19482],"quality":"More is better","isMain":true,"metricName":"Total aligned length"},{"values":[751,536,694,682,751,751,751,751],"quality":"More is better","isMain":false,"metricName":"NA50"},{"values":[548,null,611,560,634,667,582,582],"quality":"More is better","isMain":false,"metricName":"NA75"},{"values":[null,null,null,null,null,null,null,null],"quality":"More is better","isMain":true,"metricName":"NGA50"},{"values":[9,7,12,7,12,11,12,12],"quality":"Less is better","isMain":false,"metricName":"LA50"},{"values":[15,null,19,11,19,18,20,20],"quality":"Less is better","isMain":false,"metricName":"LA75"}]],["Misassemblies",[{"values":[1,0,0,0,0,0,0,0],"quality":"Less is better","isMain":true,"metricName":"# misassemblies"},{"values":[1,0,0,0,0,0,0,0],"quality":"Less is better","isMain":false,"metricName":" # relocations"},{"values":[0,0,0,0,0,0,0,0],"quality":"Less is better","isMain":false,"metricName":" # translocations"},{"values":[0,0,0,0,0,0,0,0],"quality":"Less is better","isMain":false,"metricName":" # inversions"},{"values":[1,0,0,0,0,0,0,0],"quality":"Less is better","isMain":false,"metricName":"# misassembled contigs"},{"values":[504,0,0,0,0,0,0,0],"quality":"Less is better","isMain":true,"metricName":"Misassembled contigs length"},{"values":[0,0,1,0,0,1,0,0],"quality":"Less is better","isMain":false,"metricName":"# local misassemblies"},{"values":[0,0,0,0,0,0,0,0],"quality":"Less is better","isMain":false,"metricName":"# unaligned mis. contigs"}]],["Unaligned",[{"values":[0,0,0,0,0,0,0,0],"quality":"Less is better","isMain":false,"metricName":"# fully unaligned contigs"},{"values":[0,0,0,0,0,0,0,0],"quality":"Less is better","isMain":false,"metricName":"Fully unaligned length"},{"values":[1,2,1,0,2,0,2,2],"quality":"Less is better","isMain":false,"metricName":"# partially unaligned contigs"},{"values":[900,2092,747,0,1757,0,1757,1757],"quality":"Less is better","isMain":false,"metricName":"Partially unaligned length"}]],["Mismatches",[{"values":[51,12,40,19,53,47,66,61],"quality":"Less is better","isMain":false,"metricName":"# mismatches"},{"values":[5,1,1,0,2,5,1,1],"quality":"Less is better","isMain":false,"metricName":"# indels"},{"values":[6,8,2,0,2,29,22,22],"quality":"Less is better","isMain":false,"metricName":"Indels length"},{"values":["340.75","193.74","222.35","181.80","277.62","231.58","339.16","313.46"],"quality":"Less is better","isMain":true,"metricName":"# mismatches per 100 kbp"},{"values":["33.41","16.14","5.56","0.00","10.48","24.64","5.14","5.14"],"quality":"Less is better","isMain":true,"metricName":"# indels per 100 kbp"},{"values":[5,0,1,0,2,4,0,0],"quality":"Less is better","isMain":false,"metricName":" # indels (<= 5 bp)"},{"values":[0,1,0,0,0,1,1,1],"quality":"Less is better","isMain":false,"metricName":" # indels (> 5 bp)"},{"values":[0,0,0,0,0,0,0,0],"quality":"Less is better","isMain":false,"metricName":"# N's"},{"values":["0.00","0.00","0.00","0.00","0.00","0.00","0.00","0.00"],"quality":"Less is better","isMain":true,"metricName":"# N's per 100 kbp"}]],["Statistics without reference",[{"values":[20,10,26,15,26,26,27,27],"quality":"Equal","isMain":true,"metricName":"# contigs"},{"values":[3,3,3,1,5,3,4,4],"quality":"Equal","isMain":false,"metricName":"# contigs (>= 1000 bp)"},{"values":[0,0,0,0,0,0,0,0],"quality":"Equal","isMain":false,"metricName":"# contigs (>= 5000 bp)"},{"values":[0,0,0,0,0,0,0,0],"quality":"Equal","isMain":false,"metricName":"# contigs (>= 10000 bp)"},{"values":[0,0,0,0,0,0,0,0],"quality":"Equal","isMain":false,"metricName":"# contigs (>= 25000 bp)"},{"values":[0,0,0,0,0,0,0,0],"quality":"Equal","isMain":false,"metricName":"# contigs (>= 50000 bp)"},{"values":[1391,1716,1168,1291,1391,1897,1391,1391],"quality":"More is better","isMain":true,"metricName":"Largest contig"},{"values":[15871,8585,18764,10451,20848,20386,21399,21399],"quality":"More is better","isMain":true,"metricName":"Total length"},{"values":[3949,4254,3254,1291,5844,4190,4808,4808],"quality":"More is better","isMain":true,"metricName":"Total length (>= 1000 bp)"},{"values":[0,0,0,0,0,0,0,0],"quality":"More is better","isMain":false,"metricName":"Total length (>= 5000 bp)"},{"values":[0,0,0,0,0,0,0,0],"quality":"More is better","isMain":true,"metricName":"Total length (>= 10000 bp)"},{"values":[0,0,0,0,0,0,0,0],"quality":"More is better","isMain":false,"metricName":"Total length (>= 25000 bp)"},{"values":[0,0,0,0,0,0,0,0],"quality":"More is better","isMain":true,"metricName":"Total length (>= 50000 bp)"},{"values":[803,852,743,682,805,751,803,803],"quality":"More is better","isMain":false,"metricName":"N50"},{"values":[682,575,634,560,694,667,682,682],"quality":"More is better","isMain":false,"metricName":"N75"},{"values":[8,4,11,7,11,11,11,11],"quality":"Less is better","isMain":false,"metricName":"L50"},{"values":[13,7,18,11,18,18,18,18],"quality":"Less is better","isMain":false,"metricName":"L75"},{"values":["48.14","48.39","48.29","49.32","48.47","48.36","48.42","48.43"],"quality":"Equal","isMain":false,"metricName":"GC (%)"}]],["Predicted genes",[]],["Similarity statistics",[{"values":[0,0,0,0,0,0,0,0],"quality":"Equal","isMain":false,"metricName":"# similar correct contigs"},{"values":[0,0,0,0,0,0,0,0],"quality":"Equal","isMain":false,"metricName":"# similar misassembled blocks"}]],["Reference statistics",[{"values":[156922,156922,156922,156922,156922,156922,156922,156922],"quality":"Equal","isMain":false,"metricName":"Reference length"},{"values":[1,1,1,1,1,1,1,1],"quality":"Equal","isMain":false,"metricName":"Reference fragments"},{"values":["49.26","49.26","49.26","49.26","49.26","49.26","49.26","49.26"],"quality":"Equal","isMain":false,"metricName":"Reference GC (%)"}]]],"referenceName":"gi\_115298502\_ref\_NC","date":"18 July 2018, Wednesday, 16:08:47","order":[0,1,2,3,4,5,6,7],"assembliesNames":["CLC","IDBA\_UD","MEGAHIT","SOAPdenovo2","SPAdes","SPAdes\_meta","SPAdes\_sc","SPAdes\_sc\_careful"]},{"assembliesWithNs":null,"minContig":500,"report":[["Genome statistics",[{"values":["86.229","32.283","96.184","64.489","82.063","99.622","74.364","93.679","61.490","63.351","58.189","1.966","99.296"],"quality":"More is better","isMain":true,"metricName":"Genome fraction (%)"},{"values":["1.123","1.011","1.015","1.011","1.064","2.046","1.118","1.041","1.000","1.013","1.013","1.000","1.792"],"quality":"Less is better","isMain":true,"metricName":"Duplication ratio"},{"values":[6646,2810,12708,3885,4835,12301,5442,8814,11188,3705,5132,561,15257],"quality":"More is better","isMain":true,"metricName":"Largest alignment"},{"values":[27654,9314,27854,18603,24905,57991,23373,27750,17549,18299,16826,561,50779],"quality":"More is better","isMain":true,"metricName":"Total aligned length"},{"values":[8910,null,8334,955,1315,88168,2594,5656,3324,879,847,null,106226],"quality":"More is better","isMain":false,"metricName":"NG50"},{"values":[6646,null,2755,null,597,88168,2093,2966,null,null,null,null,106226],"quality":"More is better","isMain":false,"metricName":"NG75"},{"values":[997,2192,8334,1206,1436,902,2679,5656,11188,1454,3705,561,null],"quality":"More is better","isMain":false,"metricName":"NA50"},{"values":[null,780,8334,965,805,null,2412,2966,3324,879,893,561,null],"quality":"More is better","isMain":false,"metricName":"NA75"},{"values":[2862,null,8334,955,1315,10024,2594,5656,3324,879,847,null,15257],"quality":"More is better","isMain":true,"metricName":"NGA50"},{"values":[877,null,2755,null,597,10024,2093,2966,null,null,null,null,12301],"quality":"More is better","isMain":false,"metricName":"NGA75"},{"values":[2,null,2,10,7,1,4,2,2,8,7,null,1],"quality":"Less is better","isMain":false,"metricName":"LG50"},{"values":[3,null,3,null,16,1,7,4,null,null,null,null,1],"quality":"Less is better","isMain":false,"metricName":"LG75"},{"values":[6,2,2,5,6,11,3,2,1,4,2,1,null],"quality":"Less is better","isMain":false,"metricName":"LA50"},{"values":[null,5,2,9,12,null,5,4,2,8,5,1,null],"quality":"Less is better","isMain":false,"metricName":"LA75"},{"values":[4,null,2,10,7,2,4,2,2,8,7,null,1],"quality":"Less is better","isMain":true,"metricName":"LGA50"},{"values":[8,null,3,null,16,2,7,4,null,null,null,null,2],"quality":"Less is better","isMain":false,"metricName":"LGA75"}]],["Misassemblies",[{"values":[0,0,0,0,0,0,0,0,0,0,0,0,0],"quality":"Less is better","isMain":true,"metricName":"# misassemblies"},{"values":[0,0,0,0,0,0,0,0,0,0,0,0,0],"quality":"Less is better","isMain":false,"metricName":" # relocations"},{"values":[0,0,0,0,0,0,0,0,0,0,0,0,0],"quality":"Less is better","isMain":false,"metricName":" # translocations"},{"values":[0,0,0,0,0,0,0,0,0,0,0,0,0],"quality":"Less is better","isMain":false,"metricName":" # inversions"},{"values":[0,0,0,0,0,0,0,0,0,0,0,0,0],"quality":"Less is better","isMain":false,"metricName":"# misassembled contigs"},{"values":[0,0,0,0,0,0,0,0,0,0,0,0,0],"quality":"Less is better","isMain":true,"metricName":"Misassembled contigs length"},{"values":[0,0,0,0,0,1,0,0,0,0,0,0,0],"quality":"Less is better","isMain":false,"metricName":"# local misassemblies"},{"values":[2,0,0,0,0,1,0,0,0,0,0,0,1],"quality":"Less is better","isMain":false,"metricName":"# unaligned mis. contigs"}]],["Unaligned",[{"values":[0,0,0,0,0,0,0,0,0,0,0,0,0],"quality":"Less is better","isMain":false,"metricName":"# fully unaligned contigs"},{"values":[0,0,0,0,0,0,0,0,0,0,0,0,0],"quality":"Less is better","isMain":false,"metricName":"Fully unaligned length"},{"values":[2,0,0,0,0,1,0,0,0,0,0,0,1],"quality":"Less is better","isMain":false,"metricName":"# partially unaligned contigs"},{"values":[11477,0,0,0,0,46964,0,0,0,0,0,0,55454],"quality":"Less is better","isMain":false,"metricName":"Partially unaligned length"}]],["Mismatches",[{"values":[15,0,21,0,22,309,30,8,196,124,6,0,133],"quality":"Less is better","isMain":false,"metricName":"# mismatches"},{"values":[0,0,0,0,0,9,0,0,3,2,0,0,7],"quality":"Less is better","isMain":false,"metricName":"# indels"},{"values":[0,0,0,0,0,15,0,0,3,2,0,0,13],"quality":"Less is better","isMain":false,"metricName":"Indels length"},{"values":["60.96","0.00","76.51","0.00","93.94","1086.88","141.36","29.92","1116.94","685.88","36.13","0.00","469.35"],"quality":"Less is better","isMain":true,"metricName":"# mismatches per 100 kbp"},{"values":["0.00","0.00","0.00","0.00","0.00","31.66","0.00","0.00","17.10","11.06","0.00","0.00","24.70"],"quality":"Less is better","isMain":true,"metricName":"# indels per 100 kbp"},{"values":[0,0,0,0,0,9,0,0,3,2,0,0,7],"quality":"Less is better","isMain":false,"metricName":" # indels (<= 5 bp)"},{"values":[0,0,0,0,0,0,0,0,0,0,0,0,0],"quality":"Less is better","isMain":false,"metricName":" # indels (> 5 bp)"},{"values":[111,0,0,0,0,1,0,0,0,0,0,0,50],"quality":"Less is better","isMain":false,"metricName":"# N's"},{"values":["283.71","0.00","0.00","0.00","0.00","0.95","0.00","0.00","0.00","0.00","0.00","0.00","47.07"],"quality":"Less is better","isMain":true,"metricName":"# N's per 100 kbp"}]],["Statistics without reference",[{"values":[14,8,6,16,22,9,8,10,4,13,11,1,1],"quality":"Equal","isMain":true,"metricName":"# contigs"},{"values":[5,2,4,8,9,4,8,6,4,7,4,0,1],"quality":"Equal","isMain":false,"metricName":"# contigs (>= 1000 bp)"},{"values":[3,0,2,0,0,2,1,2,1,0,1,0,1],"quality":"Equal","isMain":false,"metricName":"# contigs (>= 5000 bp)"},{"values":[1,0,1,0,0,1,0,0,1,0,0,0,1],"quality":"Equal","isMain":false,"metricName":"# contigs (>= 10000 bp)"},{"values":[0,0,0,0,0,1,0,0,0,0,0,0,1],"quality":"Equal","isMain":false,"metricName":"# contigs (>= 25000 bp)"},{"values":[0,0,0,0,0,1,0,0,0,0,0,0,1],"quality":"Equal","isMain":false,"metricName":"# contigs (>= 50000 bp)"},{"values":[10316,2810,12708,3885,4835,88168,5442,8814,11188,3705,5132,561,106226],"quality":"More is better","isMain":true,"metricName":"Largest contig"},{"values":[39124,9314,27854,18603,24911,105131,23724,27830,17549,18307,16826,561,106226],"quality":"More is better","isMain":true,"metricName":"Total length"},{"values":[32703,5002,26541,13082,16585,101012,23724,25457,17549,13490,11837,0,106226],"quality":"More is better","isMain":true,"metricName":"Total length (>= 1000 bp)"},{"values":[25872,0,21042,0,0,97965,5442,14470,11188,0,5132,0,106226],"quality":"More is better","isMain":false,"metricName":"Total length (>= 5000 bp)"},{"values":[10316,0,12708,0,0,88168,0,0,11188,0,0,0,106226],"quality":"More is better","isMain":true,"metricName":"Total length (>= 10000 bp)"},{"values":[0,0,0,0,0,88168,0,0,0,0,0,0,106226],"quality":"More is better","isMain":false,"metricName":"Total length (>= 25000 bp)"},{"values":[0,0,0,0,0,88168,0,0,0,0,0,0,106226],"quality":"More is better","isMain":true,"metricName":"Total length (>= 50000 bp)"},{"values":[6646,2192,8334,1206,1436,88168,3030,5656,11188,1454,3705,561,106226],"quality":"More is better","isMain":false,"metricName":"N50"},{"values":[3969,780,8334,965,805,88168,2412,2966,3324,879,893,561,106226],"quality":"More is better","isMain":false,"metricName":"N75"},{"values":[3,2,2,5,6,1,3,2,1,4,2,1,1],"quality":"Less is better","isMain":false,"metricName":"L50"},{"values":[4,5,2,9,12,1,5,4,2,8,5,1,1],"quality":"Less is better","isMain":false,"metricName":"L75"},{"values":["34.64","33.55","34.70","35.03","34.56","34.80","34.41","34.56","34.30","34.31","34.43","27.99","34.35"],"quality":"Equal","isMain":false,"metricName":"GC (%)"}]],["Predicted genes",[]],["Similarity statistics",[{"values":[0,0,0,0,0,0,0,0,0,0,0,0,0],"quality":"Equal","isMain":false,"metricName":"# similar correct contigs"},{"values":[0,0,0,0,0,0,0,0,0,0,0,0,0],"quality":"Equal","isMain":false,"metricName":"# similar misassembled blocks"}]],["Reference statistics",[{"values":[28538,28538,28538,28538,28538,28538,28538,28538,28538,28538,28538,28538,28538],"quality":"Equal","isMain":false,"metricName":"Reference length"},{"values":[1,1,1,1,1,1,1,1,1,1,1,1,1],"quality":"Equal","isMain":false,"metricName":"Reference fragments"},{"values":["34.68","34.68","34.68","34.68","34.68","34.68","34.68","34.68","34.68","34.68","34.68","34.68","34.68"],"quality":"Equal","isMain":false,"metricName":"Reference GC (%)"}]]],"referenceName":"gi\_115304210\_ref\_NC","date":"18 July 2018, Wednesday, 16:08:57","order":[0,1,2,3,4,5,6,7,8,9,10,11,12],"assembliesNames":["ABySS\_127","ABySS\_63","CLC","IDBA\_UD","MEGAHIT","MIRA","Ray\_Meta","SPAdes","SPAdes\_meta","SPAdes\_sc","SPAdes\_sc\_careful","Velvet","Geneious"]},{"assembliesWithNs":null,"minContig":500,"report":[["Genome statistics",[{"values":["99.634","100.000","100.000","100.000","12.865","92.358","100.000","100.000","100.000","100.000","100.000"],"quality":"More is better","isMain":true,"metricName":"Genome fraction (%)"},{"values":["1.004","1.004","1.005","1.007","1.001","1.014","1.005","1.002","1.002","1.002","1.001"],"quality":"Less is better","isMain":true,"metricName":"Duplication ratio"},{"values":[26440,26636,26678,26736,862,24861,26664,26592,26592,26592,26567],"quality":"More is better","isMain":true,"metricName":"Largest alignment"},{"values":[26440,26636,26678,26736,3418,24861,26664,26592,26592,26592,26567],"quality":"More is better","isMain":true,"metricName":"Total aligned length"},{"values":[26537,26636,26678,26736,null,27370,26664,26592,26592,26592,26567],"quality":"More is better","isMain":false,"metricName":"NG50"},{"values":[26537,26636,26678,26736,null,27370,26664,26592,26592,26592,26567],"quality":"More is better","isMain":false,"metricName":"NG75"},{"values":[26440,26636,26678,26736,614,24861,26664,26592,26592,26592,26567],"quality":"More is better","isMain":false,"metricName":"NA50"},{"values":[26440,26636,26678,26736,575,24861,26664,26592,26592,26592,26567],"quality":"More is better","isMain":false,"metricName":"NA75"},{"values":[26440,26636,26678,26736,null,24861,26664,26592,26592,26592,26567],"quality":"More is better","isMain":true,"metricName":"NGA50"},{"values":[26440,26636,26678,26736,null,24861,26664,26592,26592,26592,26567],"quality":"More is better","isMain":false,"metricName":"NGA75"},{"values":[1,1,1,1,null,1,1,1,1,1,1],"quality":"Less is better","isMain":false,"metricName":"LG50"},{"values":[1,1,1,1,null,1,1,1,1,1,1],"quality":"Less is better","isMain":false,"metricName":"LG75"},{"values":[1,1,1,1,3,1,1,1,1,1,1],"quality":"Less is better","isMain":false,"metricName":"LA50"},{"values":[1,1,1,1,4,1,1,1,1,1,1],"quality":"Less is better","isMain":false,"metricName":"LA75"},{"values":[1,1,1,1,null,1,1,1,1,1,1],"quality":"Less is better","isMain":true,"metricName":"LGA50"},{"values":[1,1,1,1,null,1,1,1,1,1,1],"quality":"Less is better","isMain":false,"metricName":"LGA75"}]],["Misassemblies",[{"values":[0,0,0,0,0,0,0,0,0,0,0],"quality":"Less is better","isMain":true,"metricName":"# misassemblies"},{"values":[0,0,0,0,0,0,0,0,0,0,0],"quality":"Less is better","isMain":false,"metricName":" # relocations"},{"values":[0,0,0,0,0,0,0,0,0,0,0],"quality":"Less is better","isMain":false,"metricName":" # translocations"},{"values":[0,0,0,0,0,0,0,0,0,0,0],"quality":"Less is better","isMain":false,"metricName":" # inversions"},{"values":[0,0,0,0,0,0,0,0,0,0,0],"quality":"Less is better","isMain":false,"metricName":"# misassembled contigs"},{"values":[0,0,0,0,0,0,0,0,0,0,0],"quality":"Less is better","isMain":true,"metricName":"Misassembled contigs length"},{"values":[0,1,1,1,0,1,1,0,0,0,0],"quality":"Less is better","isMain":false,"metricName":"# local misassemblies"},{"values":[0,0,0,0,0,0,0,0,0,0,0],"quality":"Less is better","isMain":false,"metricName":"# unaligned mis. contigs"}]],["Unaligned",[{"values":[0,0,0,0,0,0,0,0,0,0,0],"quality":"Less is better","isMain":false,"metricName":"# fully unaligned contigs"},{"values":[0,0,0,0,0,0,0,0,0,0,0],"quality":"Less is better","isMain":false,"metricName":"Fully unaligned length"},{"values":[0,0,0,0,0,1,0,0,0,0,0],"quality":"Less is better","isMain":false,"metricName":"# partially unaligned contigs"},{"values":[0,0,0,0,0,2509,0,0,0,0,0],"quality":"Less is better","isMain":false,"metricName":"Partially unaligned length"}]],["Mismatches",[{"values":[0,0,0,0,0,0,0,0,0,0,0],"quality":"Less is better","isMain":false,"metricName":"# mismatches"},{"values":[0,0,0,0,0,13,0,1,1,1,1],"quality":"Less is better","isMain":false,"metricName":"# indels"},{"values":[0,0,0,0,0,655,0,55,55,55,30],"quality":"Less is better","isMain":false,"metricName":"Indels length"},{"values":["0.00","0.00","0.00","0.00","0.00","0.00","0.00","0.00","0.00","0.00","0.00"],"quality":"Less is better","isMain":true,"metricName":"# mismatches per 100 kbp"},{"values":["0.00","0.00","0.00","0.00","0.00","53.04","0.00","3.77","3.77","3.77","3.77"],"quality":"Less is better","isMain":true,"metricName":"# indels per 100 kbp"},{"values":[0,0,0,0,0,0,0,0,0,0,0],"quality":"Less is better","isMain":false,"metricName":" # indels (<= 5 bp)"},{"values":[0,0,0,0,0,13,0,1,1,1,1],"quality":"Less is better","isMain":false,"metricName":" # indels (> 5 bp)"},{"values":[0,0,0,0,0,1437,0,0,0,0,0],"quality":"Less is better","isMain":false,"metricName":"# N's"},{"values":["0.00","0.00","0.00","0.00","0.00","5250.27","0.00","0.00","0.00","0.00","0.00"],"quality":"Less is better","isMain":true,"metricName":"# N's per 100 kbp"}]],["Statistics without reference",[{"values":[1,1,1,1,5,1,1,1,1,1,1],"quality":"Equal","isMain":true,"metricName":"# contigs"},{"values":[1,1,1,1,0,1,1,1,1,1,1],"quality":"Equal","isMain":false,"metricName":"# contigs (>= 1000 bp)"},{"values":[1,1,1,1,0,1,1,1,1,1,1],"quality":"Equal","isMain":false,"metricName":"# contigs (>= 5000 bp)"},{"values":[1,1,1,1,0,1,1,1,1,1,1],"quality":"Equal","isMain":false,"metricName":"# contigs (>= 10000 bp)"},{"values":[1,1,1,1,0,1,1,1,1,1,1],"quality":"Equal","isMain":false,"metricName":"# contigs (>= 25000 bp)"},{"values":[0,0,0,0,0,0,0,0,0,0,0],"quality":"Equal","isMain":false,"metricName":"# contigs (>= 50000 bp)"},{"values":[26537,26636,26678,26736,862,27370,26664,26592,26592,26592,26567],"quality":"More is better","isMain":true,"metricName":"Largest contig"},{"values":[26537,26636,26678,26736,3418,27370,26664,26592,26592,26592,26567],"quality":"More is better","isMain":true,"metricName":"Total length"},{"values":[26537,26636,26678,26736,0,27370,26664,26592,26592,26592,26567],"quality":"More is better","isMain":true,"metricName":"Total length (>= 1000 bp)"},{"values":[26537,26636,26678,26736,0,27370,26664,26592,26592,26592,26567],"quality":"More is better","isMain":false,"metricName":"Total length (>= 5000 bp)"},{"values":[26537,26636,26678,26736,0,27370,26664,26592,26592,26592,26567],"quality":"More is better","isMain":true,"metricName":"Total length (>= 10000 bp)"},{"values":[26537,26636,26678,26736,0,27370,26664,26592,26592,26592,26567],"quality":"More is better","isMain":false,"metricName":"Total length (>= 25000 bp)"},{"values":[0,0,0,0,0,0,0,0,0,0,0],"quality":"More is better","isMain":true,"metricName":"Total length (>= 50000 bp)"},{"values":[26537,26636,26678,26736,614,27370,26664,26592,26592,26592,26567],"quality":"More is better","isMain":false,"metricName":"N50"},{"values":[26537,26636,26678,26736,575,27370,26664,26592,26592,26592,26567],"quality":"More is better","isMain":false,"metricName":"N75"},{"values":[1,1,1,1,3,1,1,1,1,1,1],"quality":"Less is better","isMain":false,"metricName":"L50"},{"values":[1,1,1,1,4,1,1,1,1,1,1],"quality":"Less is better","isMain":false,"metricName":"L75"},{"values":["37.10","37.13","37.11","37.10","41.37","37.10","37.14","37.12","37.12","37.12","37.10"],"quality":"Equal","isMain":false,"metricName":"GC (%)"}]],["Predicted genes",[]],["Similarity statistics",[{"values":[0,0,0,0,0,0,0,0,0,0,0],"quality":"Equal","isMain":false,"metricName":"# similar correct contigs"},{"values":[0,0,0,0,0,0,0,0,0,0,0],"quality":"Equal","isMain":false,"metricName":"# similar misassembled blocks"}]],["Reference statistics",[{"values":[26537,26537,26537,26537,26537,26537,26537,26537,26537,26537,26537],"quality":"Equal","isMain":false,"metricName":"Reference length"},{"values":[1,1,1,1,1,1,1,1,1,1,1],"quality":"Equal","isMain":false,"metricName":"Reference fragments"},{"values":["37.10","37.10","37.10","37.10","37.10","37.10","37.10","37.10","37.10","37.10","37.10"],"quality":"Equal","isMain":false,"metricName":"Reference GC (%)"}]]],"referenceName":"gi\_115304270\_ref\_NC","date":"18 July 2018, Wednesday, 16:09:07","order":[0,1,2,3,4,5,6,7,8,9,10],"assembliesNames":["CLC","IDBA\_UD","MEGAHIT","MIRA","Ray\_Meta","SOAPdenovo2","SPAdes","SPAdes\_meta","SPAdes\_sc","SPAdes\_sc\_careful","Velvet"]},{"assembliesWithNs":null,"minContig":500,"report":[["Genome statistics",[{"values":["94.182","69.862","83.183","81.704","85.529","79.695","79.797","92.199","47.584","71.872","78.614","19.613","100.000"],"quality":"More is better","isMain":true,"metricName":"Genome fraction (%)"},{"values":["1.095","1.000","1.005","1.017","1.016","1.248","1.043","1.018","0.999","1.006","1.000","1.000","1.494"],"quality":"Less is better","isMain":true,"metricName":"Duplication ratio"},{"values":[6178,4853,4680,4565,4264,8004,4605,7066,5499,4511,5573,1048,8352],"quality":"More is better","isMain":true,"metricName":"Largest alignment"},{"values":[31456,21315,25496,25344,26513,29783,25396,28638,14500,22039,23986,5984,45591],"quality":"More is better","isMain":true,"metricName":"Total aligned length"},{"values":[5634,2113,3943,1915,3032,8576,4039,4642,5054,2860,3236,null,106226],"quality":"More is better","isMain":false,"metricName":"NG50"},{"values":[4117,null,2794,953,1379,4739,2584,3828,null,null,1169,null,106226],"quality":"More is better","isMain":false,"metricName":"NG75"},{"values":[5634,3442,4064,2210,3109,4739,4429,4642,5054,3358,3358,661,null],"quality":"More is better","isMain":false,"metricName":"NA50"},{"values":[4117,2113,3368,1702,2532,981,3532,4274,3947,2860,2860,622,null],"quality":"More is better","isMain":false,"metricName":"NA75"},{"values":[5634,2113,3943,1915,3032,7526,4039,4642,null,2860,3236,null,8004],"quality":"More is better","isMain":true,"metricName":"NGA50"},{"values":[4117,null,2794,953,1379,2862,2584,3828,null,null,1169,null,7340],"quality":"More is better","isMain":false,"metricName":"NGA75"},{"values":[3,5,4,5,5,2,4,3,3,5,4,null,1],"quality":"Less is better","isMain":false,"metricName":"LG50"},{"values":[5,null,6,10,8,3,6,5,null,null,8,null,1],"quality":"Less is better","isMain":false,"metricName":"LG75"},{"values":[3,3,3,4,4,3,3,3,2,3,3,4,null],"quality":"Less is better","isMain":false,"metricName":"LA50"},{"values":[5,5,5,7,6,8,5,4,3,5,5,6,null],"quality":"Less is better","isMain":false,"metricName":"LA75"},{"values":[3,5,4,5,5,2,4,3,null,5,4,null,2],"quality":"Less is better","isMain":true,"metricName":"LGA50"},{"values":[5,null,6,10,8,4,6,5,null,null,8,null,3],"quality":"Less is better","isMain":false,"metricName":"LGA75"}]],["Misassemblies",[{"values":[0,0,0,0,0,0,0,0,0,0,0,0,0],"quality":"Less is better","isMain":true,"metricName":"# misassemblies"},{"values":[0,0,0,0,0,0,0,0,0,0,0,0,0],"quality":"Less is better","isMain":false,"metricName":" # relocations"},{"values":[0,0,0,0,0,0,0,0,0,0,0,0,0],"quality":"Less is better","isMain":false,"metricName":" # translocations"},{"values":[0,0,0,0,0,0,0,0,0,0,0,0,0],"quality":"Less is better","isMain":false,"metricName":" # inversions"},{"values":[0,0,0,0,0,0,0,0,0,0,0,0,0],"quality":"Less is better","isMain":false,"metricName":"# misassembled contigs"},{"values":[0,0,0,0,0,0,0,0,0,0,0,0,0],"quality":"Less is better","isMain":true,"metricName":"Misassembled contigs length"},{"values":[0,0,0,0,0,1,0,0,1,0,0,0,0],"quality":"Less is better","isMain":false,"metricName":"# local misassemblies"},{"values":[0,0,0,0,0,0,0,0,0,0,0,0,2],"quality":"Less is better","isMain":false,"metricName":"# unaligned mis. contigs"}]],["Unaligned",[{"values":[0,0,0,0,0,0,0,0,0,0,0,0,0],"quality":"Less is better","isMain":false,"metricName":"# fully unaligned contigs"},{"values":[0,0,0,0,0,0,0,0,0,0,0,0,0],"quality":"Less is better","isMain":false,"metricName":"Fully unaligned length"},{"values":[0,0,0,0,0,2,0,0,2,0,0,0,3],"quality":"Less is better","isMain":false,"metricName":"# partially unaligned contigs"},{"values":[0,0,0,0,0,6686,0,0,3923,0,0,0,114320],"quality":"Less is better","isMain":false,"metricName":"Partially unaligned length"}]],["Mismatches",[{"values":[22,21,38,0,7,211,0,29,204,52,126,0,202],"quality":"Less is better","isMain":false,"metricName":"# mismatches"},{"values":[0,0,0,0,0,10,0,0,14,1,1,0,8],"quality":"Less is better","isMain":false,"metricName":"# indels"},{"values":[0,0,0,0,0,16,0,0,24,1,1,0,19],"quality":"Less is better","isMain":false,"metricName":"Indels length"},{"values":["76.56","98.52","149.73","0.00","26.83","867.78","0.00","103.09","1405.15","237.14","525.33","0.00","662.08"],"quality":"Less is better","isMain":true,"metricName":"# mismatches per 100 kbp"},{"values":["0.00","0.00","0.00","0.00","0.00","41.13","0.00","0.00","96.43","4.56","4.17","0.00","26.22"],"quality":"Less is better","isMain":true,"metricName":"# indels per 100 kbp"},{"values":[0,0,0,0,0,10,0,0,14,1,1,0,7],"quality":"Less is better","isMain":false,"metricName":" # indels (<= 5 bp)"},{"values":[0,0,0,0,0,0,0,0,0,0,0,0,1],"quality":"Less is better","isMain":false,"metricName":" # indels (> 5 bp)"},{"values":[0,0,0,0,0,9,0,0,0,0,0,0,286],"quality":"Less is better","isMain":false,"metricName":"# N's"},{"values":["0.00","0.00","0.00","0.00","0.00","24.30","0.00","0.00","0.00","0.00","0.00","0.00","178.85"],"quality":"Less is better","isMain":true,"metricName":"# N's per 100 kbp"}]],["Statistics without reference",[{"values":[7,11,8,13,11,10,7,8,3,8,8,8,4],"quality":"Equal","isMain":true,"metricName":"# contigs"},{"values":[6,5,8,9,8,7,7,6,3,7,8,2,4],"quality":"Equal","isMain":false,"metricName":"# contigs (>= 1000 bp)"},{"values":[3,0,0,0,0,2,0,2,3,0,1,0,3],"quality":"Equal","isMain":false,"metricName":"# contigs (>= 5000 bp)"},{"values":[0,0,0,0,0,1,0,0,0,0,0,0,2],"quality":"Equal","isMain":false,"metricName":"# contigs (>= 10000 bp)"},{"values":[0,0,0,0,0,0,0,0,0,0,0,0,2],"quality":"Equal","isMain":false,"metricName":"# contigs (>= 25000 bp)"},{"values":[0,0,0,0,0,0,0,0,0,0,0,0,1],"quality":"Equal","isMain":false,"metricName":"# contigs (>= 50000 bp)"},{"values":[6178,4853,4680,4565,4264,10863,4605,7066,7121,4513,5573,1048,106226],"quality":"More is better","isMain":true,"metricName":"Largest contig"},{"values":[31456,21315,25501,25344,26513,37041,25396,28638,18423,22051,23986,5984,159911],"quality":"More is better","isMain":true,"metricName":"Total length"},{"values":[30913,17106,25501,22174,23857,34701,25396,27514,18423,21455,23986,2074,159911],"quality":"More is better","isMain":true,"metricName":"Total length (>= 1000 bp)"},{"values":[17924,0,0,0,0,19439,0,13761,18423,0,5573,0,158562],"quality":"More is better","isMain":false,"metricName":"Total length (>= 5000 bp)"},{"values":[0,0,0,0,0,10863,0,0,0,0,0,0,153194],"quality":"More is better","isMain":true,"metricName":"Total length (>= 10000 bp)"},{"values":[0,0,0,0,0,0,0,0,0,0,0,0,153194],"quality":"More is better","isMain":false,"metricName":"Total length (>= 25000 bp)"},{"values":[0,0,0,0,0,0,0,0,0,0,0,0,106226],"quality":"More is better","isMain":true,"metricName":"Total length (>= 50000 bp)"},{"values":[5634,3442,4064,2210,3109,8576,4429,4642,6248,3358,3358,661,106226],"quality":"More is better","isMain":false,"metricName":"N50"},{"values":[4117,2113,3368,1702,2532,4523,3532,4274,5054,2860,2860,622,46968],"quality":"More is better","isMain":false,"metricName":"N75"},{"values":[3,3,3,4,4,2,3,3,2,3,3,4,1],"quality":"Less is better","isMain":false,"metricName":"L50"},{"values":[5,5,5,7,6,4,5,4,3,5,5,6,2],"quality":"Less is better","isMain":false,"metricName":"L75"},{"values":["33.70","34.06","33.98","33.91","33.96","33.72","33.38","33.93","33.82","33.94","34.16","33.77","34.45"],"quality":"Equal","isMain":false,"metricName":"GC (%)"}]],["Predicted genes",[]],["Similarity statistics",[{"values":[0,0,0,0,0,0,0,0,0,0,0,0,0],"quality":"Equal","isMain":false,"metricName":"# similar correct contigs"},{"values":[0,0,0,0,0,0,0,0,0,0,0,0,0],"quality":"Equal","isMain":false,"metricName":"# similar misassembled blocks"}]],["Reference statistics",[{"values":[30510,30510,30510,30510,30510,30510,30510,30510,30510,30510,30510,30510,30510],"quality":"Equal","isMain":false,"metricName":"Reference length"},{"values":[1,1,1,1,1,1,1,1,1,1,1,1,1],"quality":"Equal","isMain":false,"metricName":"Reference fragments"},{"values":["33.88","33.88","33.88","33.88","33.88","33.88","33.88","33.88","33.88","33.88","33.88","33.88","33.88"],"quality":"Equal","isMain":false,"metricName":"Reference GC (%)"}]]],"referenceName":"gi\_115315572\_ref\_NC","date":"18 July 2018, Wednesday, 16:09:18","order":[0,1,2,3,4,5,6,7,8,9,10,11,12],"assembliesNames":["ABySS\_127","ABySS\_63","CLC","IDBA\_UD","MEGAHIT","MIRA","Ray\_Meta","SPAdes","SPAdes\_meta","SPAdes\_sc","SPAdes\_sc\_careful","Velvet","Geneious"]},{"assembliesWithNs":null,"minContig":500,"report":[["Genome statistics",[{"values":["79.878","32.867","73.500","55.225","74.068","100.000","62.561","89.287","57.943","52.519","56.861","5.526","17.437"],"quality":"More is better","isMain":true,"metricName":"Genome fraction (%)"},{"values":["1.044","1.006","1.065","1.003","1.063","1.667","1.039","1.078","1.027","1.016","1.015","1.000","1.006"],"quality":"Less is better","isMain":true,"metricName":"Duplication ratio"},{"values":[4015,2848,3417,2533,4362,28432,7607,3249,4573,2873,2660,944,4815],"quality":"More is better","isMain":true,"metricName":"Largest alignment"},{"values":[22892,9077,21449,15213,21602,45635,17839,26417,16343,14648,15841,1517,4815],"quality":"More is better","isMain":true,"metricName":"Total aligned length"},{"values":[4015,null,1002,593,1017,88168,2686,1908,3665,519,595,null,null],"quality":"More is better","isMain":false,"metricName":"NG50"},{"values":[915,null,517,null,536,88168,null,890,1446,null,null,null,null],"quality":"More is better","isMain":false,"metricName":"NG75"},{"values":[889,928,1284,1184,1632,null,4793,1908,1817,1072,2200,944,4815],"quality":"More is better","isMain":false,"metricName":"NA50"},{"values":[500,632,957,834,749,null,2686,972,null,810,1072,573,4815],"quality":"More is better","isMain":false,"metricName":"NA75"},{"values":[915,null,964,593,1017,28432,2686,1908,1446,519,595,null,null],"quality":"More is better","isMain":true,"metricName":"NGA50"},{"values":[604,null,517,null,536,28432,null,890,null,null,null,null,null],"quality":"More is better","isMain":false,"metricName":"NGA75"},{"values":[2,null,8,12,7,1,3,6,3,12,9,null,null],"quality":"Less is better","isMain":false,"metricName":"LG50"},{"values":[6,null,17,null,18,1,null,12,6,null,null,null,null],"quality":"Less is better","isMain":false,"metricName":"LG75"},{"values":[8,3,5,5,5,null,2,6,5,4,4,1,1],"quality":"Less is better","isMain":false,"metricName":"LA50"},{"values":[19,6,10,8,10,null,3,11,null,8,6,2,1],"quality":"Less is better","isMain":false,"metricName":"LA75"},{"values":[6,null,8,12,7,1,3,6,6,12,9,null,null],"quality":"Less is better","isMain":true,"metricName":"LGA50"},{"values":[15,null,17,null,18,1,null,12,null,null,null,null,null],"quality":"Less is better","isMain":false,"metricName":"LGA75"}]],["Misassemblies",[{"values":[0,0,0,0,0,0,0,0,0,0,0,0,0],"quality":"Less is better","isMain":true,"metricName":"# misassemblies"},{"values":[0,0,0,0,0,0,0,0,0,0,0,0,0],"quality":"Less is better","isMain":false,"metricName":" # relocations"},{"values":[0,0,0,0,0,0,0,0,0,0,0,0,0],"quality":"Less is better","isMain":false,"metricName":" # translocations"},{"values":[0,0,0,0,0,0,0,0,0,0,0,0,0],"quality":"Less is better","isMain":false,"metricName":" # inversions"},{"values":[0,0,0,0,0,0,0,0,0,0,0,0,0],"quality":"Less is better","isMain":false,"metricName":"# misassembled contigs"},{"values":[0,0,0,0,0,0,0,0,0,0,0,0,0],"quality":"Less is better","isMain":true,"metricName":"Misassembled contigs length"},{"values":[0,0,0,0,0,1,0,0,0,0,0,0,0],"quality":"Less is better","isMain":false,"metricName":"# local misassemblies"},{"values":[0,0,0,0,0,0,0,0,1,0,0,0,0],"quality":"Less is better","isMain":false,"metricName":"# unaligned mis. contigs"}]],["Unaligned",[{"values":[0,0,0,0,0,0,0,0,0,0,0,0,0],"quality":"Less is better","isMain":false,"metricName":"# fully unaligned contigs"},{"values":[0,0,0,0,0,0,0,0,0,0,0,0,0],"quality":"Less is better","isMain":false,"metricName":"Fully unaligned length"},{"values":[1,0,0,0,0,2,0,0,2,0,0,0,1],"quality":"Less is better","isMain":false,"metricName":"# partially unaligned contigs"},{"values":[7406,0,0,0,0,81435,0,0,5776,0,0,0,553],"quality":"Less is better","isMain":false,"metricName":"Partially unaligned length"}]],["Mismatches",[{"values":[0,0,42,0,49,273,5,3,134,48,19,0,70],"quality":"Less is better","isMain":false,"metricName":"# mismatches"},{"values":[0,0,2,0,3,13,2,0,3,1,0,0,18],"quality":"Less is better","isMain":false,"metricName":"# indels"},{"values":[0,0,21,0,3,26,2,0,5,1,0,0,32],"quality":"Less is better","isMain":false,"metricName":"Indels length"},{"values":["0.00","0.00","208.15","0.00","240.98","994.43","29.11","12.24","842.40","332.92","121.72","0.00","1462.29"],"quality":"Less is better","isMain":true,"metricName":"# mismatches per 100 kbp"},{"values":["0.00","0.00","9.91","0.00","14.75","47.35","11.64","0.00","18.86","6.94","0.00","0.00","376.02"],"quality":"Less is better","isMain":true,"metricName":"# indels per 100 kbp"},{"values":[0,0,1,0,3,12,2,0,3,1,0,0,18],"quality":"Less is better","isMain":false,"metricName":" # indels (<= 5 bp)"},{"values":[0,0,1,0,0,1,0,0,0,0,0,0,0],"quality":"Less is better","isMain":false,"metricName":" # indels (> 5 bp)"},{"values":[109,0,0,0,0,3,0,0,0,0,0,0,19],"quality":"Less is better","isMain":false,"metricName":"# N's"},{"values":["359.76","0.00","0.00","0.00","0.00","2.36","0.00","0.00","0.00","0.00","0.00","0.00","353.95"],"quality":"Less is better","isMain":true,"metricName":"# N's per 100 kbp"}]],["Statistics without reference",[{"values":[19,10,18,14,19,8,5,20,7,13,12,2,1],"quality":"Equal","isMain":true,"metricName":"# contigs"},{"values":[4,1,8,6,7,6,4,10,7,7,7,0,1],"quality":"Equal","isMain":false,"metricName":"# contigs (>= 1000 bp)"},{"values":[1,0,0,0,0,2,1,0,1,0,0,0,1],"quality":"Equal","isMain":false,"metricName":"# contigs (>= 5000 bp)"},{"values":[1,0,0,0,0,2,0,0,0,0,0,0,0],"quality":"Equal","isMain":false,"metricName":"# contigs (>= 10000 bp)"},{"values":[0,0,0,0,0,2,0,0,0,0,0,0,0],"quality":"Equal","isMain":false,"metricName":"# contigs (>= 25000 bp)"},{"values":[0,0,0,0,0,1,0,0,0,0,0,0,0],"quality":"Equal","isMain":false,"metricName":"# contigs (>= 50000 bp)"},{"values":[10316,2848,3417,2533,4362,88168,7607,3249,7121,2873,2660,944,5368],"quality":"More is better","isMain":true,"metricName":"Largest contig"},{"values":[30298,9077,21489,15213,21606,127207,17839,26417,22119,14648,15841,1517,5368],"quality":"More is better","isMain":true,"metricName":"Total length"},{"values":[19520,2848,14501,9764,13908,125899,17146,19519,22119,10935,12945,0,5368],"quality":"More is better","isMain":true,"metricName":"Total length (>= 1000 bp)"},{"values":[10316,0,0,0,0,116737,7607,0,7121,0,0,0,5368],"quality":"More is better","isMain":false,"metricName":"Total length (>= 5000 bp)"},{"values":[10316,0,0,0,0,116737,0,0,0,0,0,0,0],"quality":"More is better","isMain":true,"metricName":"Total length (>= 10000 bp)"},{"values":[0,0,0,0,0,116737,0,0,0,0,0,0,0],"quality":"More is better","isMain":false,"metricName":"Total length (>= 25000 bp)"},{"values":[0,0,0,0,0,88168,0,0,0,0,0,0,0],"quality":"More is better","isMain":true,"metricName":"Total length (>= 50000 bp)"},{"values":[2923,928,1284,1184,1632,88168,4793,1908,4573,1072,2200,944,5368],"quality":"More is better","isMain":false,"metricName":"N50"},{"values":[889,632,957,834,749,28569,2686,972,2336,810,1072,573,5368],"quality":"More is better","isMain":false,"metricName":"N75"},{"values":[3,3,5,5,5,1,2,6,2,4,4,1,1],"quality":"Less is better","isMain":false,"metricName":"L50"},{"values":[8,6,10,8,10,2,3,11,4,8,6,2,1],"quality":"Less is better","isMain":false,"metricName":"L75"},{"values":["35.52","34.46","34.83","35.12","34.94","34.66","34.70","35.14","33.49","34.70","34.99","36.85","32.98"],"quality":"Equal","isMain":false,"metricName":"GC (%)"}]],["Predicted genes",[]],["Similarity statistics",[{"values":[0,0,0,0,0,0,0,0,0,0,0,0,0],"quality":"Equal","isMain":false,"metricName":"# similar correct contigs"},{"values":[0,0,0,0,0,0,0,0,0,0,0,0,0],"quality":"Equal","isMain":false,"metricName":"# similar misassembled blocks"}]],["Reference statistics",[{"values":[27453,27453,27453,27453,27453,27453,27453,27453,27453,27453,27453,27453,27453],"quality":"Equal","isMain":false,"metricName":"Reference length"},{"values":[1,1,1,1,1,1,1,1,1,1,1,1,1],"quality":"Equal","isMain":false,"metricName":"Reference fragments"},{"values":["34.94","34.94","34.94","34.94","34.94","34.94","34.94","34.94","34.94","34.94","34.94","34.94","34.94"],"quality":"Equal","isMain":false,"metricName":"Reference GC (%)"}]]],"referenceName":"gi\_115315628\_ref\_NC","date":"18 July 2018, Wednesday, 16:09:30","order":[0,1,2,3,4,5,6,7,8,9,10,11,12],"assembliesNames":["ABySS\_127","ABySS\_63","CLC","IDBA\_UD","MEGAHIT","MIRA","Ray\_Meta","SPAdes","SPAdes\_meta","SPAdes\_sc","SPAdes\_sc\_careful","Velvet","Geneious"]},{"assembliesWithNs":null,"minContig":500,"report":[["Genome statistics",[{"values":["38.331","18.173","32.343","37.270","43.205","67.637","2.178","15.965","40.069","20.793","30.650","30.650","20.467","16.259"],"quality":"More is better","isMain":true,"metricName":"Genome fraction (%)"},{"values":["1.002","1.004","1.008","1.000","1.007","1.169","1.000","1.066","1.000","1.000","1.003","1.003","1.004","1.000"],"quality":"Less is better","isMain":true,"metricName":"Duplication ratio"},{"values":[3355,1901,2400,4131,7534,5304,682,1700,8006,3930,3874,3874,3851,4817],"quality":"More is better","isMain":true,"metricName":"Largest alignment"},{"values":[23137,10989,19483,22451,26212,47628,1312,9666,24139,12526,18510,18510,12382,9793],"quality":"More is better","isMain":true,"metricName":"Total aligned length"},{"values":[null,null,null,null,null,153388,null,null,null,null,null,null,null,4817],"quality":"More is better","isMain":false,"metricName":"NG50"},{"values":[null,null,null,null,null,153388,null,null,null,null,null,null,null,null],"quality":"More is better","isMain":false,"metricName":"NG75"},{"values":[1263,930,1477,1204,1309,null,682,799,1994,3419,1387,1387,1367,null],"quality":"More is better","isMain":false,"metricName":"NA50"},{"values":[692,613,995,750,909,null,630,203,1387,1387,931,931,792,null],"quality":"More is better","isMain":false,"metricName":"NA75"},{"values":[null,null,null,null,null,1520,null,null,null,null,null,null,null,null],"quality":"More is better","isMain":true,"metricName":"NGA50"},{"values":[null,null,null,null,null,725,null,null,null,null,null,null,null,null],"quality":"More is better","isMain":false,"metricName":"NGA75"},{"values":[null,null,null,null,null,1,null,null,null,null,null,null,null,2],"quality":"Less is better","isMain":false,"metricName":"LG50"},{"values":[null,null,null,null,null,1,null,null,null,null,null,null,null,null],"quality":"Less is better","isMain":false,"metricName":"LG75"},{"values":[6,5,6,6,5,null,1,6,4,2,5,5,3,null],"quality":"Less is better","isMain":false,"metricName":"LA50"},{"values":[13,8,10,12,11,null,2,12,7,4,9,9,6,null],"quality":"Less is better","isMain":false,"metricName":"LA75"},{"values":[null,null,null,null,null,15,null,null,null,null,null,null,null,null],"quality":"Less is better","isMain":true,"metricName":"LGA50"},{"values":[null,null,null,null,null,28,null,null,null,null,null,null,null,null],"quality":"Less is better","isMain":false,"metricName":"LGA75"}]],["Misassemblies",[{"values":[0,0,0,0,0,0,0,0,0,0,0,0,0,0],"quality":"Less is better","isMain":true,"metricName":"# misassemblies"},{"values":[0,0,0,0,0,0,0,0,0,0,0,0,0,0],"quality":"Less is better","isMain":false,"metricName":" # relocations"},{"values":[0,0,0,0,0,0,0,0,0,0,0,0,0,0],"quality":"Less is better","isMain":false,"metricName":" # translocations"},{"values":[0,0,0,0,0,0,0,0,0,0,0,0,0,0],"quality":"Less is better","isMain":false,"metricName":" # inversions"},{"values":[0,0,0,0,0,0,0,0,0,0,0,0,0,0],"quality":"Less is better","isMain":false,"metricName":"# misassembled contigs"},{"values":[0,0,0,0,0,0,0,0,0,0,0,0,0,0],"quality":"Less is better","isMain":true,"metricName":"Misassembled contigs length"},{"values":[0,0,0,0,0,2,0,0,0,1,0,0,0,0],"quality":"Less is better","isMain":false,"metricName":"# local misassemblies"},{"values":[0,0,0,0,0,0,0,0,0,0,0,0,0,0],"quality":"Less is better","isMain":false,"metricName":"# unaligned mis. contigs"}]],["Unaligned",[{"values":[0,0,0,0,0,0,0,0,0,0,0,0,0,0],"quality":"Less is better","isMain":false,"metricName":"# fully unaligned contigs"},{"values":[0,0,0,0,0,0,0,0,0,0,0,0,0,0],"quality":"Less is better","isMain":false,"metricName":"Fully unaligned length"},{"values":[0,0,0,0,0,2,0,1,1,1,0,0,0,1],"quality":"Less is better","isMain":false,"metricName":"# partially unaligned contigs"},{"values":[0,0,0,0,0,152415,0,2569,720,894,0,0,0,22770],"quality":"Less is better","isMain":false,"metricName":"Partially unaligned length"}]],["Mismatches",[{"values":[4,0,31,12,10,183,0,0,27,22,6,6,2,43],"quality":"Less is better","isMain":false,"metricName":"# mismatches"},{"values":[0,0,0,2,0,7,0,3,2,1,0,0,0,1],"quality":"Less is better","isMain":false,"metricName":"# indels"},{"values":[0,0,0,2,0,7,0,158,2,1,0,0,0,1],"quality":"Less is better","isMain":false,"metricName":"Indels length"},{"values":["17.32","0.00","159.11","53.45","38.42","449.16","0.00","0.00","111.86","175.65","32.50","32.50","16.22","439.04"],"quality":"Less is better","isMain":true,"metricName":"# mismatches per 100 kbp"},{"values":["0.00","0.00","0.00","8.91","0.00","17.18","0.00","31.19","8.29","7.98","0.00","0.00","0.00","10.21"],"quality":"Less is better","isMain":true,"metricName":"# indels per 100 kbp"},{"values":[0,0,0,2,0,7,0,0,2,1,0,0,0,1],"quality":"Less is better","isMain":false,"metricName":" # indels (<= 5 bp)"},{"values":[0,0,0,0,0,0,0,3,0,0,0,0,0,0],"quality":"Less is better","isMain":false,"metricName":" # indels (> 5 bp)"},{"values":[0,0,0,0,0,24,0,1035,0,0,0,0,0,48],"quality":"Less is better","isMain":false,"metricName":"# N's"},{"values":["0.00","0.00","0.00","0.00","0.00","12.00","0.00","8070.80","0.00","0.00","0.00","0.00","0.00","147.41"],"quality":"Less is better","isMain":true,"metricName":"# N's per 100 kbp"}]],["Statistics without reference",[{"values":[22,13,15,21,20,31,2,12,13,6,15,15,10,3],"quality":"Equal","isMain":true,"metricName":"# contigs"},{"values":[8,3,9,6,8,23,0,4,9,4,7,7,5,3],"quality":"Equal","isMain":false,"metricName":"# contigs (>= 1000 bp)"},{"values":[0,0,0,0,1,2,0,0,1,0,0,0,0,1],"quality":"Equal","isMain":false,"metricName":"# contigs (>= 5000 bp)"},{"values":[0,0,0,0,0,1,0,0,0,0,0,0,0,1],"quality":"Equal","isMain":false,"metricName":"# contigs (>= 10000 bp)"},{"values":[0,0,0,0,0,1,0,0,0,0,0,0,0,1],"quality":"Equal","isMain":false,"metricName":"# contigs (>= 25000 bp)"},{"values":[0,0,0,0,0,1,0,0,0,0,0,0,0,0],"quality":"Equal","isMain":false,"metricName":"# contigs (>= 50000 bp)"},{"values":[3355,1901,2406,4131,7534,153388,682,3422,8006,3930,3874,3874,3851,26427],"quality":"More is better","isMain":true,"metricName":"Largest contig"},{"values":[23137,10989,19635,22451,26212,200046,1312,12824,24859,13420,18510,18510,12382,32563],"quality":"More is better","isMain":true,"metricName":"Total length"},{"values":[13943,4454,14682,12257,17461,193768,0,7508,22070,11930,12853,12853,8894,32563],"quality":"More is better","isMain":true,"metricName":"Total length (>= 1000 bp)"},{"values":[0,0,0,0,7534,158692,0,0,8006,0,0,0,0,26427],"quality":"More is better","isMain":false,"metricName":"Total length (>= 5000 bp)"},{"values":[0,0,0,0,0,153388,0,0,0,0,0,0,0,26427],"quality":"More is better","isMain":true,"metricName":"Total length (>= 10000 bp)"},{"values":[0,0,0,0,0,153388,0,0,0,0,0,0,0,26427],"quality":"More is better","isMain":false,"metricName":"Total length (>= 25000 bp)"},{"values":[0,0,0,0,0,153388,0,0,0,0,0,0,0,0],"quality":"More is better","isMain":true,"metricName":"Total length (>= 50000 bp)"},{"values":[1263,930,1477,1204,1309,153388,682,1243,2306,3419,1387,1387,1367,26427],"quality":"More is better","isMain":false,"metricName":"N50"},{"values":[692,613,995,750,909,153388,630,705,1387,3194,931,931,792,26427],"quality":"More is better","isMain":false,"metricName":"N75"},{"values":[6,5,6,6,5,1,1,3,3,2,5,5,3,1],"quality":"Less is better","isMain":false,"metricName":"L50"},{"values":[13,8,10,12,11,1,2,7,7,3,9,9,6,1],"quality":"Less is better","isMain":false,"metricName":"L75"},{"values":["50.28","50.52","49.92","49.95","50.43","50.74","38.34","45.88","50.03","49.28","50.05","50.05","49.59","50.39"],"quality":"Equal","isMain":false,"metricName":"GC (%)"}]],["Predicted genes",[]],["Similarity statistics",[{"values":[0,0,0,0,0,0,0,0,0,0,0,0,0,0],"quality":"Equal","isMain":false,"metricName":"# similar correct contigs"},{"values":[0,0,0,0,0,0,0,0,0,0,0,0,0,0],"quality":"Equal","isMain":false,"metricName":"# similar misassembled blocks"}]],["Reference statistics",[{"values":[60238,60238,60238,60238,60238,60238,60238,60238,60238,60238,60238,60238,60238,60238],"quality":"Equal","isMain":false,"metricName":"Reference length"},{"values":[1,1,1,1,1,1,1,1,1,1,1,1,1,1],"quality":"Equal","isMain":false,"metricName":"Reference fragments"},{"values":["49.07","49.07","49.07","49.07","49.07","49.07","49.07","49.07","49.07","49.07","49.07","49.07","49.07","49.07"],"quality":"Equal","isMain":false,"metricName":"Reference GC (%)"}]]],"referenceName":"gi\_116221992\_ref\_NC","date":"18 July 2018, Wednesday, 16:09:42","order":[0,1,2,3,4,5,6,7,8,9,10,11,12,13],"assembliesNames":["ABySS\_127","ABySS\_63","CLC","IDBA\_UD","MEGAHIT","MIRA","Ray\_Meta","SOAPdenovo2","SPAdes","SPAdes\_meta","SPAdes\_sc","SPAdes\_sc\_careful","Velvet","Geneious"]},{"assembliesWithNs":null,"minContig":500,"report":[["Genome statistics",[{"values":["99.679","92.646","95.437","93.156","88.644","100.000","93.445","2.564","97.620","99.006","95.115","86.138","46.783","100.000"],"quality":"More is better","isMain":true,"metricName":"Genome fraction (%)"},{"values":["1.039","1.015","1.010","1.043","1.067","1.844","1.134","1.121","1.050","1.013","1.030","1.024","1.003","1.730"],"quality":"Less is better","isMain":true,"metricName":"Duplication ratio"},{"values":[18611,25403,16673,6770,8156,81430,7501,749,13139,24149,8302,7311,3244,81430],"quality":"More is better","isMain":true,"metricName":"Largest alignment"},{"values":[171874,155959,159961,161118,156869,305566,175643,4254,170069,165626,162150,146272,77833,287014],"quality":"More is better","isMain":true,"metricName":"Total aligned length"},{"values":[9253,4083,6721,1873,2421,167653,2816,null,5296,11824,2371,1530,null,171313],"quality":"More is better","isMain":false,"metricName":"NG50"},{"values":[5010,1872,3045,1179,1233,167653,1955,null,2084,6810,1047,785,null,171313],"quality":"More is better","isMain":false,"metricName":"NG75"},{"values":[9021,4163,6721,1896,2452,17862,2758,608,5085,11295,2371,1838,878,26703],"quality":"More is better","isMain":false,"metricName":"NA50"},{"values":[4709,2068,3047,1210,1338,4878,1829,502,1840,6637,1125,920,671,11496],"quality":"More is better","isMain":false,"metricName":"NA75"},{"values":[9253,4083,6721,1873,2321,33704,2811,null,5296,11295,2371,1390,null,48260],"quality":"More is better","isMain":true,"metricName":"NGA50"},{"values":[5010,1872,3045,1179,1204,26703,1955,null,1984,6637,1046,755,null,48260],"quality":"More is better","isMain":false,"metricName":"NGA75"},{"values":[6,11,8,28,20,1,20,null,10,6,19,27,null,1],"quality":"Less is better","isMain":false,"metricName":"LG50"},{"values":[12,28,18,56,45,1,38,null,22,11,47,66,null,1],"quality":"Less is better","isMain":false,"metricName":"LG75"},{"values":[7,10,8,27,20,5,22,4,11,6,19,23,30,4],"quality":"Less is better","isMain":false,"metricName":"LA50"},{"values":[13,24,17,53,43,12,42,6,25,11,46,52,56,8],"quality":"Less is better","isMain":false,"metricName":"LA75"},{"values":[6,11,8,28,22,2,20,null,10,6,19,28,null,2],"quality":"Less is better","isMain":true,"metricName":"LGA50"},{"values":[12,28,18,56,47,3,38,null,23,11,48,68,null,2],"quality":"Less is better","isMain":false,"metricName":"LGA75"}]],["Misassemblies",[{"values":[0,0,0,0,1,7,0,0,0,0,0,0,0,9],"quality":"Less is better","isMain":true,"metricName":"# misassemblies"},{"values":[0,0,0,0,1,7,0,0,0,0,0,0,0,9],"quality":"Less is better","isMain":false,"metricName":" # relocations"},{"values":[0,0,0,0,0,0,0,0,0,0,0,0,0,0],"quality":"Less is better","isMain":false,"metricName":" # translocations"},{"values":[0,0,0,0,0,0,0,0,0,0,0,0,0,0],"quality":"Less is better","isMain":false,"metricName":" # inversions"},{"values":[0,0,0,0,1,4,0,0,0,0,0,0,0,2],"quality":"Less is better","isMain":false,"metricName":"# misassembled contigs"},{"values":[0,0,0,0,11039,279796,0,0,0,0,0,0,0,340604],"quality":"Less is better","isMain":true,"metricName":"Misassembled contigs length"},{"values":[0,0,0,0,0,19,1,0,0,1,1,0,0,25],"quality":"Less is better","isMain":false,"metricName":"# local misassemblies"},{"values":[0,0,0,0,0,0,0,0,0,0,0,0,0,0],"quality":"Less is better","isMain":false,"metricName":"# unaligned mis. contigs"}]],["Unaligned",[{"values":[0,0,0,0,0,0,0,0,0,0,0,0,0,0],"quality":"Less is better","isMain":false,"metricName":"# fully unaligned contigs"},{"values":[0,0,0,0,0,0,0,0,0,0,0,0,0,0],"quality":"Less is better","isMain":false,"metricName":"Fully unaligned length"},{"values":[0,0,0,0,1,4,0,0,2,2,1,2,0,2],"quality":"Less is better","isMain":false,"metricName":"# partially unaligned contigs"},{"values":[0,0,0,0,1117,45531,0,0,1407,1384,847,1904,0,53590],"quality":"Less is better","isMain":false,"metricName":"Partially unaligned length"}]],["Mismatches",[{"values":[2,29,9,11,279,1086,199,0,4,131,126,41,53,1443],"quality":"Less is better","isMain":false,"metricName":"# mismatches"},{"values":[0,0,0,0,8,32,8,4,0,8,3,3,2,46],"quality":"Less is better","isMain":false,"metricName":"# indels"},{"values":[0,0,0,0,13,37,12,204,0,20,52,52,2,80],"quality":"Less is better","isMain":false,"metricName":"Indels length"},{"values":["1.21","18.87","5.68","7.12","189.73","654.65","128.37","0.00","2.47","79.76","79.85","28.69","68.29","869.85"],"quality":"Less is better","isMain":true,"metricName":"# mismatches per 100 kbp"},{"values":["0.00","0.00","0.00","0.00","5.44","19.29","5.16","94.03","0.00","4.87","1.90","2.10","2.58","27.73"],"quality":"Less is better","isMain":true,"metricName":"# indels per 100 kbp"},{"values":[0,0,0,0,7,32,8,0,0,7,2,2,2,43],"quality":"Less is better","isMain":false,"metricName":" # indels (<= 5 bp)"},{"values":[0,0,0,0,1,0,0,4,0,1,1,1,0,3],"quality":"Less is better","isMain":false,"metricName":" # indels (> 5 bp)"},{"values":[0,17,0,0,0,14,0,515,1407,2085,1238,1904,0,253],"quality":"Less is better","isMain":false,"metricName":"# N's"},{"values":["0.00","10.90","0.00","0.00","0.00","3.98","0.00","10798.91","820.52","1243.33","757.63","1284.68","0.00","74.28"],"quality":"Less is better","isMain":true,"metricName":"# N's per 100 kbp"}]],["Statistics without reference",[{"values":[28,65,44,102,86,28,81,7,69,28,97,103,89,2],"quality":"Equal","isMain":true,"metricName":"# contigs"},{"values":[27,32,35,65,53,18,65,0,43,21,49,47,19,2],"quality":"Equal","isMain":false,"metricName":"# contigs (>= 1000 bp)"},{"values":[12,7,11,3,4,6,6,0,11,13,6,3,0,2],"quality":"Equal","isMain":false,"metricName":"# contigs (>= 5000 bp)"},{"values":[5,1,4,0,1,6,0,0,3,6,0,0,0,2],"quality":"Equal","isMain":false,"metricName":"# contigs (>= 10000 bp)"},{"values":[0,1,0,0,0,4,0,0,0,0,0,0,0,2],"quality":"Equal","isMain":false,"metricName":"# contigs (>= 25000 bp)"},{"values":[0,0,0,0,0,2,0,0,0,0,0,0,0,2],"quality":"Equal","isMain":false,"metricName":"# contigs (>= 50000 bp)"},{"values":[18611,25403,16673,6770,11039,167653,7501,967,13139,24149,8302,7311,3244,171313],"quality":"More is better","isMain":true,"metricName":"Largest contig"},{"values":[171874,155959,159961,161118,157988,351354,175839,4769,171476,167695,163405,148208,77834,340604],"quality":"More is better","isMain":true,"metricName":"Total length"},{"values":[171354,132757,153957,134679,133310,344425,163168,0,153963,162330,127368,108393,28851,340604],"quality":"More is better","isMain":true,"metricName":"Total length (>= 1000 bp)"},{"values":[127404,66787,101424,18472,33535,311543,36425,0,89821,140930,39730,18854,0,340604],"quality":"More is better","isMain":false,"metricName":"Total length (>= 5000 bp)"},{"values":[75975,25403,54325,0,11039,311543,0,0,36498,91348,0,0,0,340604],"quality":"More is better","isMain":true,"metricName":"Total length (>= 10000 bp)"},{"values":[0,25403,0,0,0,279796,0,0,0,0,0,0,0,340604],"quality":"More is better","isMain":false,"metricName":"Total length (>= 25000 bp)"},{"values":[0,0,0,0,0,221525,0,0,0,0,0,0,0,340604],"quality":"More is better","isMain":true,"metricName":"Total length (>= 50000 bp)"},{"values":[9021,4163,6721,1896,2452,53872,2763,718,5085,11824,2423,1892,878,171313],"quality":"More is better","isMain":false,"metricName":"N50"},{"values":[4709,2068,3047,1210,1342,28058,1840,544,1840,6810,1131,941,671,169291],"quality":"More is better","isMain":false,"metricName":"N75"},{"values":[7,10,8,27,19,2,22,3,11,6,18,22,30,1],"quality":"Less is better","isMain":false,"metricName":"L50"},{"values":[13,24,17,53,41,4,42,5,25,11,45,50,56,2],"quality":"Less is better","isMain":false,"metricName":"L75"},{"values":["35.36","35.34","35.37","35.40","35.12","35.46","35.29","34.88","35.31","35.30","35.37","35.41","35.26","35.32"],"quality":"Equal","isMain":false,"metricName":"GC (%)"}]],["Predicted genes",[]],["Similarity statistics",[{"values":[0,0,0,0,0,0,0,0,0,0,0,0,0,0],"quality":"Equal","isMain":false,"metricName":"# similar correct contigs"},{"values":[0,0,0,0,0,0,0,0,0,0,0,0,0,0],"quality":"Equal","isMain":false,"metricName":"# similar misassembled blocks"}]],["Reference statistics",[{"values":[165890,165890,165890,165890,165890,165890,165890,165890,165890,165890,165890,165890,165890,165890],"quality":"Equal","isMain":false,"metricName":"Reference length"},{"values":[1,1,1,1,1,1,1,1,1,1,1,1,1,1],"quality":"Equal","isMain":false,"metricName":"Reference fragments"},{"values":["35.34","35.34","35.34","35.34","35.34","35.34","35.34","35.34","35.34","35.34","35.34","35.34","35.34","35.34"],"quality":"Equal","isMain":false,"metricName":"Reference GC (%)"}]]],"referenceName":"gi\_116326222\_ref\_NC","date":"18 July 2018, Wednesday, 16:09:57","order":[0,1,2,3,4,5,6,7,8,9,10,11,12,13],"assembliesNames":["ABySS\_127","ABySS\_63","CLC","IDBA\_UD","MEGAHIT","MIRA","Ray\_Meta","SOAPdenovo2","SPAdes","SPAdes\_meta","SPAdes\_sc","SPAdes\_sc\_careful","Velvet","Geneious"]},{"assembliesWithNs":null,"minContig":500,"report":[["Genome statistics",[{"values":["11.368","9.389","71.051","62.583","67.595","29.429","62.189","76.880","75.162","75.590","75.590","4.523","6.853"],"quality":"More is better","isMain":true,"metricName":"Genome fraction (%)"},{"values":["1.000","1.000","1.012","1.000","1.001","1.027","1.002","1.002","1.005","1.002","1.002","1.000","1.000"],"quality":"Less is better","isMain":true,"metricName":"Duplication ratio"},{"values":[2257,1547,5191,5789,5789,3341,2991,5937,5937,5387,5387,1749,4203],"quality":"More is better","isMain":true,"metricName":"Largest alignment"},{"values":[19787,16343,124587,108966,117794,52595,108478,133889,131452,131770,131770,7872,11928],"quality":"More is better","isMain":true,"metricName":"Total aligned length"},{"values":[null,null,973,879,834,null,762,1380,1266,1325,1325,null,null],"quality":"More is better","isMain":false,"metricName":"NG50"},{"values":[null,null,null,null,null,null,null,599,563,563,563,null,null],"quality":"More is better","isMain":false,"metricName":"NG75"},{"values":[796,725,1437,1505,1492,1145,1202,1630,1589,1644,1644,1188,null],"quality":"More is better","isMain":false,"metricName":"NA50"},{"values":[639,599,854,1037,821,953,841,1037,990,1037,1037,686,null],"quality":"More is better","isMain":false,"metricName":"NA75"},{"values":[null,null,973,879,834,null,762,1351,1266,1292,1292,null,null],"quality":"More is better","isMain":true,"metricName":"NGA50"},{"values":[null,null,null,null,null,null,null,592,563,563,563,null,null],"quality":"More is better","isMain":false,"metricName":"NGA75"},{"values":[null,null,47,52,50,null,66,40,41,40,40,null,null],"quality":"Less is better","isMain":false,"metricName":"LG50"},{"values":[null,null,null,null,null,null,null,88,94,92,92,null,null],"quality":"Less is better","isMain":false,"metricName":"LG75"},{"values":[9,8,27,24,26,17,32,27,26,26,26,3,null],"quality":"Less is better","isMain":false,"metricName":"LA50"},{"values":[17,14,55,46,51,29,59,53,52,51,51,5,null],"quality":"Less is better","isMain":false,"metricName":"LA75"},{"values":[null,null,47,52,50,null,66,40,41,41,41,null,null],"quality":"Less is better","isMain":true,"metricName":"LGA50"},{"values":[null,null,null,null,null,null,null,90,95,92,92,null,null],"quality":"Less is better","isMain":false,"metricName":"LGA75"}]],["Misassemblies",[{"values":[0,0,1,0,0,0,0,0,1,0,0,0,0],"quality":"Less is better","isMain":true,"metricName":"# misassemblies"},{"values":[0,0,1,0,0,0,0,0,1,0,0,0,0],"quality":"Less is better","isMain":false,"metricName":" # relocations"},{"values":[0,0,0,0,0,0,0,0,0,0,0,0,0],"quality":"Less is better","isMain":false,"metricName":" # translocations"},{"values":[0,0,0,0,0,0,0,0,0,0,0,0,0],"quality":"Less is better","isMain":false,"metricName":" # inversions"},{"values":[0,0,1,0,0,0,0,0,1,0,0,0,0],"quality":"Less is better","isMain":false,"metricName":"# misassembled contigs"},{"values":[0,0,2159,0,0,0,0,0,4928,0,0,0,0],"quality":"Less is better","isMain":true,"metricName":"Misassembled contigs length"},{"values":[0,0,0,0,0,0,0,0,0,0,0,0,0],"quality":"Less is better","isMain":false,"metricName":"# local misassemblies"},{"values":[0,0,0,0,0,0,0,0,0,0,0,0,0],"quality":"Less is better","isMain":false,"metricName":"# unaligned mis. contigs"}]],["Unaligned",[{"values":[0,0,0,0,0,0,0,0,0,0,0,0,0],"quality":"Less is better","isMain":false,"metricName":"# fully unaligned contigs"},{"values":[0,0,0,0,0,0,0,0,0,0,0,0,0],"quality":"Less is better","isMain":false,"metricName":"Fully unaligned length"},{"values":[0,0,0,0,0,0,0,1,0,0,0,0,2],"quality":"Less is better","isMain":false,"metricName":"# partially unaligned contigs"},{"values":[0,0,0,0,0,0,0,1580,0,0,0,0,25262],"quality":"Less is better","isMain":false,"metricName":"Partially unaligned length"}]],["Mismatches",[{"values":[2,0,136,97,91,22,82,192,174,143,116,2,0],"quality":"Less is better","isMain":false,"metricName":"# mismatches"},{"values":[0,0,0,0,0,0,1,1,0,0,0,0,0],"quality":"Less is better","isMain":false,"metricName":"# indels"},{"values":[0,0,0,0,0,0,46,48,0,0,0,0,0],"quality":"Less is better","isMain":false,"metricName":"Indels length"},{"values":["10.11","0.00","109.97","89.05","77.34","42.95","75.75","143.48","133.00","108.69","88.16","25.41","0.00"],"quality":"Less is better","isMain":true,"metricName":"# mismatches per 100 kbp"},{"values":["0.00","0.00","0.00","0.00","0.00","0.00","0.92","0.75","0.00","0.00","0.00","0.00","0.00"],"quality":"Less is better","isMain":true,"metricName":"# indels per 100 kbp"},{"values":[0,0,0,0,0,0,0,0,0,0,0,0,0],"quality":"Less is better","isMain":false,"metricName":" # indels (<= 5 bp)"},{"values":[0,0,0,0,0,0,1,1,0,0,0,0,0],"quality":"Less is better","isMain":false,"metricName":" # indels (> 5 bp)"},{"values":[0,0,0,0,0,78,60,0,0,0,0,0,39],"quality":"Less is better","isMain":false,"metricName":"# N's"},{"values":["0.00","0.00","0.00","0.00","0.00","148.30","55.31","0.00","0.00","0.00","0.00","0.00","104.87"],"quality":"Less is better","isMain":true,"metricName":"# N's per 100 kbp"}]],["Statistics without reference",[{"values":[25,21,99,83,95,47,99,96,95,94,94,8,5],"quality":"Equal","isMain":true,"metricName":"# contigs"},{"values":[3,4,45,47,42,24,44,55,50,52,52,3,5],"quality":"Equal","isMain":false,"metricName":"# contigs (>= 1000 bp)"},{"values":[0,0,1,1,1,0,0,1,1,1,1,0,1],"quality":"Equal","isMain":false,"metricName":"# contigs (>= 5000 bp)"},{"values":[0,0,0,0,0,0,0,0,0,0,0,0,1],"quality":"Equal","isMain":false,"metricName":"# contigs (>= 10000 bp)"},{"values":[0,0,0,0,0,0,0,0,0,0,0,0,1],"quality":"Equal","isMain":false,"metricName":"# contigs (>= 25000 bp)"},{"values":[0,0,0,0,0,0,0,0,0,0,0,0,0],"quality":"Equal","isMain":false,"metricName":"# contigs (>= 50000 bp)"},{"values":[2257,1547,5191,5789,5789,3341,2991,5937,5937,5387,5387,1749,28523],"quality":"More is better","isMain":true,"metricName":"Largest contig"},{"values":[19787,16343,125157,108982,117796,52595,108478,135631,131452,131843,131843,7872,37190],"quality":"More is better","isMain":true,"metricName":"Total length"},{"values":[4822,5101,85885,82816,80102,34585,68546,105596,98068,100854,100854,4563,37190],"quality":"More is better","isMain":true,"metricName":"Total length (>= 1000 bp)"},{"values":[0,0,5191,5789,5789,0,0,5937,5937,5387,5387,0,28523],"quality":"More is better","isMain":false,"metricName":"Total length (>= 5000 bp)"},{"values":[0,0,0,0,0,0,0,0,0,0,0,0,28523],"quality":"More is better","isMain":true,"metricName":"Total length (>= 10000 bp)"},{"values":[0,0,0,0,0,0,0,0,0,0,0,0,28523],"quality":"More is better","isMain":false,"metricName":"Total length (>= 25000 bp)"},{"values":[0,0,0,0,0,0,0,0,0,0,0,0,0],"quality":"More is better","isMain":true,"metricName":"Total length (>= 50000 bp)"},{"values":[796,725,1437,1505,1492,1145,1202,1644,1589,1644,1644,1188,28523],"quality":"More is better","isMain":false,"metricName":"N50"},{"values":[639,599,870,1037,821,953,841,1082,990,1037,1037,686,28523],"quality":"More is better","isMain":false,"metricName":"N75"},{"values":[9,8,27,24,26,17,32,27,26,26,26,3,1],"quality":"Less is better","isMain":false,"metricName":"L50"},{"values":[17,14,54,46,51,29,59,52,51,51,51,5,1],"quality":"Less is better","isMain":false,"metricName":"L75"},{"values":["36.38","35.61","35.22","35.32","35.28","35.58","34.97","35.31","35.22","35.21","35.21","34.20","39.29"],"quality":"Equal","isMain":false,"metricName":"GC (%)"}]],["Predicted genes",[]],["Similarity statistics",[{"values":[0,0,0,0,0,0,0,0,0,0,0,0,0],"quality":"Equal","isMain":false,"metricName":"# similar correct contigs"},{"values":[0,0,0,0,0,0,0,0,0,0,0,0,0],"quality":"Equal","isMain":false,"metricName":"# similar misassembled blocks"}]],["Reference statistics",[{"values":[174059,174059,174059,174059,174059,174059,174059,174059,174059,174059,174059,174059,174059],"quality":"Equal","isMain":false,"metricName":"Reference length"},{"values":[1,1,1,1,1,1,1,1,1,1,1,1,1],"quality":"Equal","isMain":false,"metricName":"Reference fragments"},{"values":["35.23","35.23","35.23","35.23","35.23","35.23","35.23","35.23","35.23","35.23","35.23","35.23","35.23"],"quality":"Equal","isMain":false,"metricName":"Reference GC (%)"}]]],"referenceName":"gi\_116326687\_ref\_NC","date":"18 July 2018, Wednesday, 16:10:09","order":[0,1,2,3,4,5,6,7,8,9,10,11,12],"assembliesNames":["ABySS\_127","ABySS\_63","CLC","IDBA\_UD","MEGAHIT","MIRA","SOAPdenovo2","SPAdes","SPAdes\_meta","SPAdes\_sc","SPAdes\_sc\_careful","Velvet","Geneious"]},{"assembliesWithNs":null,"minContig":500,"report":[["Genome statistics",[{"values":["95.449","92.019","99.468","99.688","99.697","97.950","97.499","99.722","99.924","99.827","99.803","99.474","89.002"],"quality":"More is better","isMain":true,"metricName":"Genome fraction (%)"},{"values":["1.002","1.001","1.009","1.002","1.002","1.011","1.018","1.002","1.000","0.999","1.000","1.001","1.001"],"quality":"Less is better","isMain":true,"metricName":"Duplication ratio"},{"values":[7915,8093,39627,104015,54462,31005,38886,104015,104249,104249,104249,47498,41621],"quality":"More is better","isMain":true,"metricName":"Largest alignment"},{"values":[155009,149324,162370,161646,161653,160408,158503,161668,161944,161729,161745,161351,144280],"quality":"More is better","isMain":true,"metricName":"Total aligned length"},{"values":[2982,2855,31670,104300,49581,18565,21819,104300,104249,104249,104249,22465,22466],"quality":"More is better","isMain":false,"metricName":"NG50"},{"values":[1752,1502,22779,18568,24324,16984,7466,33275,33317,33102,33118,22146,18232],"quality":"More is better","isMain":false,"metricName":"NG75"},{"values":[3149,3085,31385,104015,49581,18565,21524,104015,104249,104249,104249,22465,31005],"quality":"More is better","isMain":false,"metricName":"NA50"},{"values":[1826,1715,22779,18568,24324,16984,7315,33275,33317,33102,33118,22146,18565],"quality":"More is better","isMain":false,"metricName":"NA75"},{"values":[2982,2855,31385,104015,49581,18565,21524,104015,104249,104249,104249,22465,22466],"quality":"More is better","isMain":true,"metricName":"NGA50"},{"values":[1752,1502,22779,18568,24324,16984,7026,33275,33317,33102,33118,22146,18232],"quality":"More is better","isMain":false,"metricName":"NGA75"},{"values":[17,18,3,1,2,4,3,1,1,1,1,3,3],"quality":"Less is better","isMain":false,"metricName":"LG50"},{"values":[35,38,4,2,3,6,7,2,2,2,2,4,5],"quality":"Less is better","isMain":false,"metricName":"LG75"},{"values":[16,16,3,1,2,4,3,1,1,1,1,3,2],"quality":"Less is better","isMain":false,"metricName":"LA50"},{"values":[32,32,4,2,3,6,7,2,2,2,2,4,4],"quality":"Less is better","isMain":false,"metricName":"LA75"},{"values":[17,18,3,1,2,4,3,1,1,1,1,3,3],"quality":"Less is better","isMain":true,"metricName":"LGA50"},{"values":[35,38,4,2,3,6,8,2,2,2,2,4,5],"quality":"Less is better","isMain":false,"metricName":"LGA75"}]],["Misassemblies",[{"values":[0,0,0,0,0,0,0,0,0,0,0,0,0],"quality":"Less is better","isMain":true,"metricName":"# misassemblies"},{"values":[0,0,0,0,0,0,0,0,0,0,0,0,0],"quality":"Less is better","isMain":false,"metricName":" # relocations"},{"values":[0,0,0,0,0,0,0,0,0,0,0,0,0],"quality":"Less is better","isMain":false,"metricName":" # translocations"},{"values":[0,0,0,0,0,0,0,0,0,0,0,0,0],"quality":"Less is better","isMain":false,"metricName":" # inversions"},{"values":[0,0,0,0,0,0,0,0,0,0,0,0,0],"quality":"Less is better","isMain":false,"metricName":"# misassembled contigs"},{"values":[0,0,0,0,0,0,0,0,0,0,0,0,0],"quality":"Less is better","isMain":true,"metricName":"Misassembled contigs length"},{"values":[0,0,0,0,0,0,0,0,0,0,1,0,0],"quality":"Less is better","isMain":false,"metricName":"# local misassemblies"},{"values":[0,0,0,0,0,0,0,0,0,0,0,0,0],"quality":"Less is better","isMain":false,"metricName":"# unaligned mis. contigs"}]],["Unaligned",[{"values":[0,0,0,0,0,0,0,0,0,0,0,0,0],"quality":"Less is better","isMain":false,"metricName":"# fully unaligned contigs"},{"values":[0,0,0,0,0,0,0,0,0,0,0,0,0],"quality":"Less is better","isMain":false,"metricName":"Fully unaligned length"},{"values":[0,0,0,0,0,0,0,0,0,0,0,0,0],"quality":"Less is better","isMain":false,"metricName":"# partially unaligned contigs"},{"values":[0,0,0,0,0,0,0,0,0,0,0,0,0],"quality":"Less is better","isMain":false,"metricName":"Partially unaligned length"}]],["Mismatches",[{"values":[17,5,6,5,6,0,6,21,13,9,16,15,0],"quality":"Less is better","isMain":false,"metricName":"# mismatches"},{"values":[0,1,2,0,0,1,31,1,1,4,9,8,1],"quality":"Less is better","isMain":false,"metricName":"# indels"},{"values":[0,51,2,0,0,42,1479,42,51,269,130,82,42],"quality":"Less is better","isMain":false,"metricName":"Indels length"},{"values":["10.99","3.35","3.72","3.09","3.71","0.00","3.80","12.99","8.03","5.56","9.89","9.30","0.00"],"quality":"Less is better","isMain":true,"metricName":"# mismatches per 100 kbp"},{"values":["0.00","0.67","1.24","0.00","0.00","0.63","19.61","0.62","0.62","2.47","5.56","4.96","0.69"],"quality":"Less is better","isMain":true,"metricName":"# indels per 100 kbp"},{"values":[0,0,2,0,0,0,0,0,0,0,7,7,0],"quality":"Less is better","isMain":false,"metricName":" # indels (<= 5 bp)"},{"values":[0,1,0,0,0,1,31,1,1,4,2,1,1],"quality":"Less is better","isMain":false,"metricName":" # indels (> 5 bp)"},{"values":[0,0,0,0,0,41,2817,0,0,10,0,0,33],"quality":"Less is better","isMain":false,"metricName":"# N's"},{"values":["0.00","0.00","0.00","0.00","0.00","25.54","1751.59","0.00","0.00","6.18","0.00","0.00","22.85"],"quality":"Less is better","isMain":true,"metricName":"# N's per 100 kbp"}]],["Statistics without reference",[{"values":[67,67,13,5,5,17,17,3,3,3,3,11,7],"quality":"Equal","isMain":true,"metricName":"# contigs"},{"values":[51,49,11,5,5,15,16,3,3,3,3,10,7],"quality":"Equal","isMain":false,"metricName":"# contigs (>= 1000 bp)"},{"values":[6,6,6,5,5,7,11,3,3,3,3,7,7],"quality":"Equal","isMain":false,"metricName":"# contigs (>= 5000 bp)"},{"values":[0,0,5,4,5,6,3,3,3,3,3,5,5],"quality":"Equal","isMain":false,"metricName":"# contigs (>= 10000 bp)"},{"values":[0,0,3,1,2,1,2,2,2,2,2,2,2],"quality":"Equal","isMain":false,"metricName":"# contigs (>= 25000 bp)"},{"values":[0,0,0,1,1,0,0,1,1,1,1,0,0],"quality":"Equal","isMain":false,"metricName":"# contigs (>= 50000 bp)"},{"values":[7915,8093,39627,104300,54718,31143,39085,104300,104249,104249,104249,47498,41621],"quality":"More is better","isMain":true,"metricName":"Largest contig"},{"values":[155009,149324,162655,161931,161909,160546,160825,161953,161944,161729,161745,161351,144418],"quality":"More is better","isMain":true,"metricName":"Total length"},{"values":[143613,136342,161038,161931,161909,159327,160301,161953,161944,161729,161745,160766,144418],"quality":"More is better","isMain":true,"metricName":"Total length (>= 1000 bp)"},{"values":[40139,39457,146782,161931,161909,137783,147848,161953,161944,161729,161745,155279,144418],"quality":"More is better","isMain":false,"metricName":"Total length (>= 5000 bp)"},{"values":[0,0,140917,152083,161909,132194,91911,161953,161944,161729,161745,138963,132027],"quality":"More is better","isMain":true,"metricName":"Total length (>= 10000 bp)"},{"values":[0,0,103564,104300,104299,31143,70092,137575,137566,137351,137367,79770,72764],"quality":"More is better","isMain":false,"metricName":"Total length (>= 25000 bp)"},{"values":[0,0,0,104300,54718,0,0,104300,104249,104249,104249,0,0],"quality":"More is better","isMain":true,"metricName":"Total length (>= 50000 bp)"},{"values":[3149,3085,31670,104300,49581,18565,21819,104300,104249,104249,104249,22465,31143],"quality":"More is better","isMain":false,"metricName":"N50"},{"values":[1826,1715,22779,18568,24324,16984,7466,33275,33317,33102,33118,22146,18565],"quality":"More is better","isMain":false,"metricName":"N75"},{"values":[16,16,3,1,2,4,3,1,1,1,1,3,2],"quality":"Less is better","isMain":false,"metricName":"L50"},{"values":[32,32,4,2,3,6,7,2,2,2,2,4,4],"quality":"Less is better","isMain":false,"metricName":"L75"},{"values":["45.94","45.95","45.91","45.95","45.96","46.00","46.05","45.95","45.96","45.94","45.95","45.96","46.24"],"quality":"Equal","isMain":false,"metricName":"GC (%)"}]],["Predicted genes",[]],["Similarity statistics",[{"values":[0,0,1,1,1,1,0,1,0,1,0,1,1],"quality":"Equal","isMain":false,"metricName":"# similar correct contigs"},{"values":[0,0,0,0,0,0,0,0,0,0,0,0,0],"quality":"Equal","isMain":false,"metricName":"# similar misassembled blocks"}]],["Reference statistics",[{"values":[162109,162109,162109,162109,162109,162109,162109,162109,162109,162109,162109,162109,162109],"quality":"Equal","isMain":false,"metricName":"Reference length"},{"values":[1,1,1,1,1,1,1,1,1,1,1,1,1],"quality":"Equal","isMain":false,"metricName":"Reference fragments"},{"values":["45.95","45.95","45.95","45.95","45.95","45.95","45.95","45.95","45.95","45.95","45.95","45.95","45.95"],"quality":"Equal","isMain":false,"metricName":"Reference GC (%)"}]]],"referenceName":"gi\_117530171\_ref\_NC","date":"18 July 2018, Wednesday, 16:10:21","order":[0,1,2,3,4,5,6,7,8,9,10,11,12],"assembliesNames":["ABySS\_127","ABySS\_63","CLC","IDBA\_UD","MEGAHIT","MIRA","SOAPdenovo2","SPAdes","SPAdes\_meta","SPAdes\_sc","SPAdes\_sc\_careful","Velvet","Geneious"]},{"assembliesWithNs":null,"minContig":500,"report":[["Genome statistics",[{"values":["29.339","11.733","28.614","20.946","21.430","94.540","18.606","31.988","13.592","20.978","17.652","19.320"],"quality":"More is better","isMain":true,"metricName":"Genome fraction (%)"},{"values":["1.000","1.000","1.000","1.000","1.000","1.504","1.051","1.011","1.000","1.000","1.000","1.000"],"quality":"Less is better","isMain":true,"metricName":"Duplication ratio"},{"values":[3365,3099,3667,3744,3784,19280,3620,3772,2769,2769,2769,3639],"quality":"More is better","isMain":true,"metricName":"Largest alignment"},{"values":[11043,4416,10770,7884,8066,53507,7088,12167,5116,7896,6644,7272],"quality":"More is better","isMain":true,"metricName":"Total aligned length"},{"values":[null,null,null,null,null,40228,null,null,null,null,null,null],"quality":"More is better","isMain":false,"metricName":"NG50"},{"values":[null,null,null,null,null,40228,null,null,null,null,null,null],"quality":"More is better","isMain":false,"metricName":"NG75"},{"values":[1128,3099,2829,2291,2184,15622,2225,2419,2769,2347,2347,3639],"quality":"More is better","isMain":false,"metricName":"NA50"},{"values":[876,746,2274,2291,822,1501,2225,785,2347,931,931,2282],"quality":"More is better","isMain":false,"metricName":"NA75"},{"values":[null,null,null,null,null,19280,null,null,null,null,null,null],"quality":"More is better","isMain":true,"metricName":"NGA50"},{"values":[null,null,null,null,null,15622,null,null,null,null,null,null],"quality":"More is better","isMain":false,"metricName":"NGA75"},{"values":[null,null,null,null,null,1,null,null,null,null,null,null],"quality":"Less is better","isMain":false,"metricName":"LG50"},{"values":[null,null,null,null,null,1,null,null,null,null,null,null],"quality":"Less is better","isMain":false,"metricName":"LG75"},{"values":[3,1,2,2,2,2,2,2,1,2,2,1],"quality":"Less is better","isMain":false,"metricName":"LA50"},{"values":[5,2,3,2,3,6,2,5,2,4,3,2],"quality":"Less is better","isMain":false,"metricName":"LA75"},{"values":[null,null,null,null,null,1,null,null,null,null,null,null],"quality":"Less is better","isMain":true,"metricName":"LGA50"},{"values":[null,null,null,null,null,2,null,null,null,null,null,null],"quality":"Less is better","isMain":false,"metricName":"LGA75"}]],["Misassemblies",[{"values":[0,0,0,0,0,1,0,0,0,0,0,0],"quality":"Less is better","isMain":true,"metricName":"# misassemblies"},{"values":[0,0,0,0,0,1,0,0,0,0,0,0],"quality":"Less is better","isMain":false,"metricName":" # relocations"},{"values":[0,0,0,0,0,0,0,0,0,0,0,0],"quality":"Less is better","isMain":false,"metricName":" # translocations"},{"values":[0,0,0,0,0,0,0,0,0,0,0,0],"quality":"Less is better","isMain":false,"metricName":" # inversions"},{"values":[0,0,0,0,0,1,0,0,0,0,0,0],"quality":"Less is better","isMain":false,"metricName":"# misassembled contigs"},{"values":[0,0,0,0,0,40228,0,0,0,0,0,0],"quality":"Less is better","isMain":true,"metricName":"Misassembled contigs length"},{"values":[0,0,0,0,0,2,0,0,0,0,0,0],"quality":"Less is better","isMain":false,"metricName":"# local misassemblies"},{"values":[0,0,0,0,0,0,0,0,0,0,0,0],"quality":"Less is better","isMain":false,"metricName":"# unaligned mis. contigs"}]],["Unaligned",[{"values":[0,0,0,0,0,0,0,0,0,0,0,0],"quality":"Less is better","isMain":false,"metricName":"# fully unaligned contigs"},{"values":[0,0,0,0,0,0,0,0,0,0,0,0],"quality":"Less is better","isMain":false,"metricName":"Fully unaligned length"},{"values":[0,0,0,0,0,1,0,0,0,1,1,0],"quality":"Less is better","isMain":false,"metricName":"# partially unaligned contigs"},{"values":[0,0,0,0,0,5326,0,0,0,1151,1151,0],"quality":"Less is better","isMain":false,"metricName":"Partially unaligned length"}]],["Mismatches",[{"values":[4,0,1,0,2,484,0,2,0,0,0,3],"quality":"Less is better","isMain":false,"metricName":"# mismatches"},{"values":[0,0,0,0,0,15,4,0,0,0,0,0],"quality":"Less is better","isMain":false,"metricName":"# indels"},{"values":[0,0,0,0,0,24,174,0,0,0,0,0],"quality":"Less is better","isMain":false,"metricName":"Indels length"},{"values":["36.22","0.00","9.29","0.00","24.80","1360.16","0.00","16.61","0.00","0.00","0.00","41.25"],"quality":"Less is better","isMain":true,"metricName":"# mismatches per 100 kbp"},{"values":["0.00","0.00","0.00","0.00","0.00","42.15","57.12","0.00","0.00","0.00","0.00","0.00"],"quality":"Less is better","isMain":true,"metricName":"# indels per 100 kbp"},{"values":[0,0,0,0,0,14,0,0,0,0,0,0],"quality":"Less is better","isMain":false,"metricName":" # indels (<= 5 bp)"},{"values":[0,0,0,0,0,1,4,0,0,0,0,0],"quality":"Less is better","isMain":false,"metricName":" # indels (> 5 bp)"},{"values":[0,0,0,0,0,11,364,0,0,0,0,0],"quality":"Less is better","isMain":false,"metricName":"# N's"},{"values":["0.00","0.00","0.00","0.00","0.00","18.70","4945.65","0.00","0.00","0.00","0.00","0.00"],"quality":"Less is better","isMain":true,"metricName":"# N's per 100 kbp"}]],["Statistics without reference",[{"values":[9,3,6,5,5,13,4,9,2,5,4,3],"quality":"Equal","isMain":true,"metricName":"# contigs"},{"values":[3,1,3,2,2,9,2,3,2,4,3,3],"quality":"Equal","isMain":false,"metricName":"# contigs (>= 1000 bp)"},{"values":[0,0,0,0,0,1,0,0,0,0,0,0],"quality":"Equal","isMain":false,"metricName":"# contigs (>= 5000 bp)"},{"values":[0,0,0,0,0,1,0,0,0,0,0,0],"quality":"Equal","isMain":false,"metricName":"# contigs (>= 10000 bp)"},{"values":[0,0,0,0,0,1,0,0,0,0,0,0],"quality":"Equal","isMain":false,"metricName":"# contigs (>= 25000 bp)"},{"values":[0,0,0,0,0,0,0,0,0,0,0,0],"quality":"Equal","isMain":false,"metricName":"# contigs (>= 50000 bp)"},{"values":[3365,3099,3667,3744,3784,40228,3755,3772,2769,2769,2769,3639],"quality":"More is better","isMain":true,"metricName":"Largest contig"},{"values":[11043,4416,10770,7884,8066,58833,7360,12167,5116,9047,7795,7272],"quality":"More is better","isMain":true,"metricName":"Total length"},{"values":[6587,3099,8770,6035,5968,55658,5980,8033,5116,8506,7198,7272],"quality":"More is better","isMain":true,"metricName":"Total length (>= 1000 bp)"},{"values":[0,0,0,0,0,40228,0,0,0,0,0,0],"quality":"More is better","isMain":false,"metricName":"Total length (>= 5000 bp)"},{"values":[0,0,0,0,0,40228,0,0,0,0,0,0],"quality":"More is better","isMain":true,"metricName":"Total length (>= 10000 bp)"},{"values":[0,0,0,0,0,40228,0,0,0,0,0,0],"quality":"More is better","isMain":false,"metricName":"Total length (>= 25000 bp)"},{"values":[0,0,0,0,0,0,0,0,0,0,0,0],"quality":"More is better","isMain":true,"metricName":"Total length (>= 50000 bp)"},{"values":[1128,3099,2829,2291,2184,40228,3755,2419,2769,2347,2347,3639],"quality":"More is better","isMain":false,"metricName":"N50"},{"values":[876,746,2274,2291,822,2632,2225,785,2347,2082,2082,2282],"quality":"More is better","isMain":false,"metricName":"N75"},{"values":[3,1,2,2,2,1,1,2,1,2,2,1],"quality":"Less is better","isMain":false,"metricName":"L50"},{"values":[5,2,3,2,3,3,2,5,2,3,3,2],"quality":"Less is better","isMain":false,"metricName":"L75"},{"values":["61.18","56.88","61.16","59.16","59.00","64.92","57.00","63.41","55.55","57.06","56.36","59.19"],"quality":"Equal","isMain":false,"metricName":"GC (%)"}]],["Predicted genes",[]],["Similarity statistics",[{"values":[0,0,0,0,0,0,0,0,0,0,0,0],"quality":"Equal","isMain":false,"metricName":"# similar correct contigs"},{"values":[0,0,0,0,0,0,0,0,0,0,0,0],"quality":"Equal","isMain":false,"metricName":"# similar misassembled blocks"}]],["Reference statistics",[{"values":[37639,37639,37639,37639,37639,37639,37639,37639,37639,37639,37639,37639],"quality":"Equal","isMain":false,"metricName":"Reference length"},{"values":[1,1,1,1,1,1,1,1,1,1,1,1],"quality":"Equal","isMain":false,"metricName":"Reference fragments"},{"values":["64.82","64.82","64.82","64.82","64.82","64.82","64.82","64.82","64.82","64.82","64.82","64.82"],"quality":"Equal","isMain":false,"metricName":"Reference GC (%)"}]]],"referenceName":"gi\_117676329\_ref\_NC","date":"18 July 2018, Wednesday, 16:10:31","order":[0,1,2,3,4,5,6,7,8,9,10,11],"assembliesNames":["ABySS\_127","ABySS\_63","CLC","IDBA\_UD","MEGAHIT","MIRA","SOAPdenovo2","SPAdes","SPAdes\_meta","SPAdes\_sc","SPAdes\_sc\_careful","Velvet"]},{"assembliesWithNs":null,"minContig":500,"report":[["Genome statistics",[{"values":["97.698","97.068","99.758","99.758","99.474","98.850","2.111","98.538","99.758","99.758","99.758","99.758","99.758","71.774"],"quality":"More is better","isMain":true,"metricName":"Genome fraction (%)"},{"values":["1.002","1.001","1.002","1.000","1.000","1.008","1.000","1.022","1.000","1.001","1.000","1.000","1.000","1.000"],"quality":"Less is better","isMain":true,"metricName":"Duplication ratio"},{"values":[17852,13705,58223,58223,58174,49102,796,33621,58223,58223,58223,58223,58223,50003],"quality":"More is better","isMain":true,"metricName":"Largest alignment"},{"values":[149200,148106,152304,152004,151571,151763,3216,150792,152004,152156,152004,152004,152004,109364],"quality":"More is better","isMain":true,"metricName":"Total aligned length"},{"values":[6106,7071,44938,44938,44832,44661,null,11445,44938,44938,44938,44938,44938,44661],"quality":"More is better","isMain":false,"metricName":"NG50"},{"values":[3844,4223,17667,33843,33683,26163,null,8540,33843,26502,33843,33843,33843,null],"quality":"More is better","isMain":false,"metricName":"NG75"},{"values":[6846,7646,44938,44938,44832,44661,596,11262,44938,44938,44938,44938,44938,44661],"quality":"More is better","isMain":false,"metricName":"NA50"},{"values":[4074,4724,17667,33843,33683,26163,586,8442,33843,26502,33843,33843,33843,44661],"quality":"More is better","isMain":false,"metricName":"NA75"},{"values":[6106,7071,44938,44938,44832,44661,null,11262,44938,44938,44938,44938,44938,44661],"quality":"More is better","isMain":true,"metricName":"NGA50"},{"values":[3844,4223,17667,33843,33683,26163,null,8442,33843,26502,33843,33843,33843,null],"quality":"More is better","isMain":false,"metricName":"NGA75"},{"values":[8,9,2,2,2,2,null,4,2,2,2,2,2,2],"quality":"Less is better","isMain":false,"metricName":"LG50"},{"values":[16,15,3,3,3,3,null,8,3,3,3,3,3,null],"quality":"Less is better","isMain":false,"metricName":"LG75"},{"values":[7,8,2,2,2,2,3,4,2,2,2,2,2,2],"quality":"Less is better","isMain":false,"metricName":"LA50"},{"values":[15,14,3,3,3,3,4,8,3,3,3,3,3,2],"quality":"Less is better","isMain":false,"metricName":"LA75"},{"values":[8,9,2,2,2,2,null,4,2,2,2,2,2,2],"quality":"Less is better","isMain":true,"metricName":"LGA50"},{"values":[16,15,3,3,3,3,null,8,3,3,3,3,3,null],"quality":"Less is better","isMain":false,"metricName":"LGA75"}]],["Misassemblies",[{"values":[0,0,0,0,0,0,0,0,0,0,0,0,0,0],"quality":"Less is better","isMain":true,"metricName":"# misassemblies"},{"values":[0,0,0,0,0,0,0,0,0,0,0,0,0,0],"quality":"Less is better","isMain":false,"metricName":" # relocations"},{"values":[0,0,0,0,0,0,0,0,0,0,0,0,0,0],"quality":"Less is better","isMain":false,"metricName":" # translocations"},{"values":[0,0,0,0,0,0,0,0,0,0,0,0,0,0],"quality":"Less is better","isMain":false,"metricName":" # inversions"},{"values":[0,0,0,0,0,0,0,0,0,0,0,0,0,0],"quality":"Less is better","isMain":false,"metricName":"# misassembled contigs"},{"values":[0,0,0,0,0,0,0,0,0,0,0,0,0,0],"quality":"Less is better","isMain":true,"metricName":"Misassembled contigs length"},{"values":[0,0,0,0,0,0,0,0,0,0,0,0,0,0],"quality":"Less is better","isMain":false,"metricName":"# local misassemblies"},{"values":[0,0,0,0,0,0,0,0,0,0,0,0,0,0],"quality":"Less is better","isMain":false,"metricName":"# unaligned mis. contigs"}]],["Unaligned",[{"values":[0,0,0,0,0,0,0,0,0,0,0,0,0,0],"quality":"Less is better","isMain":false,"metricName":"# fully unaligned contigs"},{"values":[0,0,0,0,0,0,0,0,0,0,0,0,0,0],"quality":"Less is better","isMain":false,"metricName":"Fully unaligned length"},{"values":[0,0,0,0,0,0,0,0,0,0,0,0,0,0],"quality":"Less is better","isMain":false,"metricName":"# partially unaligned contigs"},{"values":[0,0,0,0,0,0,0,0,0,0,0,0,0,0],"quality":"Less is better","isMain":false,"metricName":"Partially unaligned length"}]],["Mismatches",[{"values":[9,2,3,2,5,0,0,1,3,5,2,2,1,0],"quality":"Less is better","isMain":false,"metricName":"# mismatches"},{"values":[0,0,0,0,0,0,0,31,0,0,0,0,0,0],"quality":"Less is better","isMain":false,"metricName":"# indels"},{"values":[0,0,0,0,0,0,0,1517,0,0,0,0,0,0],"quality":"Less is better","isMain":false,"metricName":"Indels length"},{"values":["6.05","1.35","1.97","1.32","3.30","0.00","0.00","0.67","1.97","3.29","1.32","1.32","0.66","0.00"],"quality":"Less is better","isMain":true,"metricName":"# mismatches per 100 kbp"},{"values":["0.00","0.00","0.00","0.00","0.00","0.00","0.00","20.65","0.00","0.00","0.00","0.00","0.00","0.00"],"quality":"Less is better","isMain":true,"metricName":"# indels per 100 kbp"},{"values":[0,0,0,0,0,0,0,0,0,0,0,0,0,0],"quality":"Less is better","isMain":false,"metricName":" # indels (<= 5 bp)"},{"values":[0,0,0,0,0,0,0,31,0,0,0,0,0,0],"quality":"Less is better","isMain":false,"metricName":" # indels (> 5 bp)"},{"values":[0,0,0,0,0,15,0,3033,0,0,0,0,0,14],"quality":"Less is better","isMain":false,"metricName":"# N's"},{"values":["0.00","0.00","0.00","0.00","0.00","9.88","0.00","1976.55","0.00","0.00","0.00","0.00","0.00","12.80"],"quality":"Less is better","isMain":true,"metricName":"# N's per 100 kbp"}]],["Statistics without reference",[{"values":[31,29,5,4,4,8,5,16,4,5,4,4,4,3],"quality":"Equal","isMain":true,"metricName":"# contigs"},{"values":[28,27,5,4,4,7,0,16,4,5,4,4,4,3],"quality":"Equal","isMain":false,"metricName":"# contigs (>= 1000 bp)"},{"values":[9,13,5,4,4,6,0,10,4,5,4,4,4,3],"quality":"Equal","isMain":false,"metricName":"# contigs (>= 5000 bp)"},{"values":[3,2,5,4,4,4,0,5,4,4,4,4,4,3],"quality":"Equal","isMain":false,"metricName":"# contigs (>= 10000 bp)"},{"values":[0,0,2,3,3,3,0,1,3,3,3,3,3,2],"quality":"Equal","isMain":false,"metricName":"# contigs (>= 25000 bp)"},{"values":[0,0,1,1,1,0,0,0,1,1,1,1,1,1],"quality":"Equal","isMain":false,"metricName":"# contigs (>= 50000 bp)"},{"values":[17852,13705,58223,58223,58174,49102,796,34220,58223,58223,58223,58223,58223,50003],"quality":"More is better","isMain":true,"metricName":"Largest contig"},{"values":[149200,148106,152304,152004,151571,151763,3216,153449,152004,152156,152004,152004,152004,109364],"quality":"More is better","isMain":true,"metricName":"Total length"},{"values":[146949,146465,152304,152004,151571,150770,0,153449,152004,152156,152004,152004,152004,109364],"quality":"More is better","isMain":true,"metricName":"Total length (>= 1000 bp)"},{"values":[86994,106729,152304,152004,151571,149718,0,135326,152004,152156,152004,152004,152004,109364],"quality":"More is better","isMain":false,"metricName":"Total length (>= 5000 bp)"},{"values":[45067,25950,152304,152004,151571,134626,0,94694,152004,144663,152004,152004,152004,109364],"quality":"More is better","isMain":true,"metricName":"Total length (>= 10000 bp)"},{"values":[0,0,103161,137004,136689,119926,0,34220,137004,129663,137004,137004,137004,94664],"quality":"More is better","isMain":false,"metricName":"Total length (>= 25000 bp)"},{"values":[0,0,58223,58223,58174,0,0,0,58223,58223,58223,58223,58223,50003],"quality":"More is better","isMain":true,"metricName":"Total length (>= 50000 bp)"},{"values":[6846,7646,44938,44938,44832,44661,596,11445,44938,44938,44938,44938,44938,44661],"quality":"More is better","isMain":false,"metricName":"N50"},{"values":[4074,4724,17667,33843,33683,26163,586,8540,33843,26502,33843,33843,33843,44661],"quality":"More is better","isMain":false,"metricName":"N75"},{"values":[7,8,2,2,2,2,3,4,2,2,2,2,2,2],"quality":"Less is better","isMain":false,"metricName":"L50"},{"values":[15,14,3,3,3,3,4,8,3,3,3,3,3,2],"quality":"Less is better","isMain":false,"metricName":"L75"},{"values":["32.65","32.65","32.58","32.59","32.60","32.61","32.18","32.57","32.59","32.58","32.59","32.59","32.59","32.52"],"quality":"Equal","isMain":false,"metricName":"GC (%)"}]],["Predicted genes",[]],["Similarity statistics",[{"values":[0,0,3,3,3,2,0,1,3,3,3,3,3,2],"quality":"Equal","isMain":false,"metricName":"# similar correct contigs"},{"values":[0,0,0,0,0,0,0,0,0,0,0,0,0,0],"quality":"Equal","isMain":false,"metricName":"# similar misassembled blocks"}]],["Reference statistics",[{"values":[152372,152372,152372,152372,152372,152372,152372,152372,152372,152372,152372,152372,152372,152372],"quality":"Equal","isMain":false,"metricName":"Reference length"},{"values":[1,1,1,1,1,1,1,1,1,1,1,1,1,1],"quality":"Equal","isMain":false,"metricName":"Reference fragments"},{"values":["32.59","32.59","32.59","32.59","32.59","32.59","32.59","32.59","32.59","32.59","32.59","32.59","32.59","32.59"],"quality":"Equal","isMain":false,"metricName":"Reference GC (%)"}]]],"referenceName":"gi\_118197620\_ref\_NC","date":"18 July 2018, Wednesday, 16:10:43","order":[0,1,2,3,4,5,6,7,8,9,10,11,12,13],"assembliesNames":["ABySS\_127","ABySS\_63","CLC","IDBA\_UD","MEGAHIT","MIRA","Ray\_Meta","SOAPdenovo2","SPAdes","SPAdes\_meta","SPAdes\_sc","SPAdes\_sc\_careful","Velvet","Geneious"]},{"assembliesWithNs":null,"minContig":500,"report":[["Genome statistics",[{"values":["100.000","100.000","100.000","100.000","100.000","8.346","89.853","100.000","100.000","100.000","100.000","100.000"],"quality":"More is better","isMain":true,"metricName":"Genome fraction (%)"},{"values":["1.017","1.000","1.015","1.021","1.020","1.000","1.048","1.019","1.008","1.008","1.008","1.005"],"quality":"Less is better","isMain":true,"metricName":"Duplication ratio"},{"values":[6776,6662,6761,6803,6798,556,5986,6789,6717,6717,6717,6692],"quality":"More is better","isMain":true,"metricName":"Largest alignment"},{"values":[6776,6662,6761,6803,6798,556,5986,6789,6717,6717,6717,6692],"quality":"More is better","isMain":true,"metricName":"Total aligned length"},{"values":[6776,6662,6761,6803,6798,null,6271,6789,6717,6717,6717,6692],"quality":"More is better","isMain":false,"metricName":"NG50"},{"values":[6776,6662,6761,6803,6798,null,6271,6789,6717,6717,6717,6692],"quality":"More is better","isMain":false,"metricName":"NG75"},{"values":[6776,6662,6761,6803,6798,556,5986,6789,6717,6717,6717,6692],"quality":"More is better","isMain":false,"metricName":"NA50"},{"values":[6776,6662,6761,6803,6798,556,5986,6789,6717,6717,6717,6692],"quality":"More is better","isMain":false,"metricName":"NA75"},{"values":[6776,6662,6761,6803,6798,null,5986,6789,6717,6717,6717,6692],"quality":"More is better","isMain":true,"metricName":"NGA50"},{"values":[6776,6662,6761,6803,6798,null,5986,6789,6717,6717,6717,6692],"quality":"More is better","isMain":false,"metricName":"NGA75"},{"values":[1,1,1,1,1,null,1,1,1,1,1,1],"quality":"Less is better","isMain":false,"metricName":"LG50"},{"values":[1,1,1,1,1,null,1,1,1,1,1,1],"quality":"Less is better","isMain":false,"metricName":"LG75"},{"values":[1,1,1,1,1,1,1,1,1,1,1,1],"quality":"Less is better","isMain":false,"metricName":"LA50"},{"values":[1,1,1,1,1,1,1,1,1,1,1,1],"quality":"Less is better","isMain":false,"metricName":"LA75"},{"values":[1,1,1,1,1,null,1,1,1,1,1,1],"quality":"Less is better","isMain":true,"metricName":"LGA50"},{"values":[1,1,1,1,1,null,1,1,1,1,1,1],"quality":"Less is better","isMain":false,"metricName":"LGA75"}]],["Misassemblies",[{"values":[0,0,0,0,0,0,0,0,0,0,0,0],"quality":"Less is better","isMain":true,"metricName":"# misassemblies"},{"values":[0,0,0,0,0,0,0,0,0,0,0,0],"quality":"Less is better","isMain":false,"metricName":" # relocations"},{"values":[0,0,0,0,0,0,0,0,0,0,0,0],"quality":"Less is better","isMain":false,"metricName":" # translocations"},{"values":[0,0,0,0,0,0,0,0,0,0,0,0],"quality":"Less is better","isMain":false,"metricName":" # inversions"},{"values":[0,0,0,0,0,0,0,0,0,0,0,0],"quality":"Less is better","isMain":false,"metricName":"# misassembled contigs"},{"values":[0,0,0,0,0,0,0,0,0,0,0,0],"quality":"Less is better","isMain":true,"metricName":"Misassembled contigs length"},{"values":[1,0,1,1,1,0,0,1,0,0,0,0],"quality":"Less is better","isMain":false,"metricName":"# local misassemblies"},{"values":[0,0,0,0,0,0,0,0,0,0,0,0],"quality":"Less is better","isMain":false,"metricName":"# unaligned mis. contigs"}]],["Unaligned",[{"values":[0,0,0,0,0,0,0,0,0,0,0,0],"quality":"Less is better","isMain":false,"metricName":"# fully unaligned contigs"},{"values":[0,0,0,0,0,0,0,0,0,0,0,0],"quality":"Less is better","isMain":false,"metricName":"Fully unaligned length"},{"values":[0,0,0,0,0,0,0,0,0,0,0,0],"quality":"Less is better","isMain":false,"metricName":"# partially unaligned contigs"},{"values":[0,0,0,0,0,0,0,0,0,0,0,0],"quality":"Less is better","isMain":false,"metricName":"Partially unaligned length"}]],["Mismatches",[{"values":[0,0,0,0,0,0,0,0,0,0,0,0],"quality":"Less is better","isMain":false,"metricName":"# mismatches"},{"values":[0,0,0,0,0,0,2,0,1,1,1,1],"quality":"Less is better","isMain":false,"metricName":"# indels"},{"values":[0,0,0,0,0,0,101,0,55,55,55,30],"quality":"Less is better","isMain":false,"metricName":"Indels length"},{"values":["0.00","0.00","0.00","0.00","0.00","0.00","0.00","0.00","0.00","0.00","0.00","0.00"],"quality":"Less is better","isMain":true,"metricName":"# mismatches per 100 kbp"},{"values":["0.00","0.00","0.00","0.00","0.00","0.00","33.41","0.00","15.01","15.01","15.01","15.01"],"quality":"Less is better","isMain":true,"metricName":"# indels per 100 kbp"},{"values":[0,0,0,0,0,0,0,0,0,0,0,0],"quality":"Less is better","isMain":false,"metricName":" # indels (<= 5 bp)"},{"values":[0,0,0,0,0,0,2,0,1,1,1,1],"quality":"Less is better","isMain":false,"metricName":" # indels (> 5 bp)"},{"values":[0,0,0,0,0,0,285,0,0,0,0,0],"quality":"Less is better","isMain":false,"metricName":"# N's"},{"values":["0.00","0.00","0.00","0.00","0.00","0.00","4544.73","0.00","0.00","0.00","0.00","0.00"],"quality":"Less is better","isMain":true,"metricName":"# N's per 100 kbp"}]],["Statistics without reference",[{"values":[1,1,1,1,1,1,1,1,1,1,1,1],"quality":"Equal","isMain":true,"metricName":"# contigs"},{"values":[1,1,1,1,1,0,1,1,1,1,1,1],"quality":"Equal","isMain":false,"metricName":"# contigs (>= 1000 bp)"},{"values":[1,1,1,1,1,0,1,1,1,1,1,1],"quality":"Equal","isMain":false,"metricName":"# contigs (>= 5000 bp)"},{"values":[0,0,0,0,0,0,0,0,0,0,0,0],"quality":"Equal","isMain":false,"metricName":"# contigs (>= 10000 bp)"},{"values":[0,0,0,0,0,0,0,0,0,0,0,0],"quality":"Equal","isMain":false,"metricName":"# contigs (>= 25000 bp)"},{"values":[0,0,0,0,0,0,0,0,0,0,0,0],"quality":"Equal","isMain":false,"metricName":"# contigs (>= 50000 bp)"},{"values":[6776,6662,6761,6803,6798,556,6271,6789,6717,6717,6717,6692],"quality":"More is better","isMain":true,"metricName":"Largest contig"},{"values":[6776,6662,6761,6803,6798,556,6271,6789,6717,6717,6717,6692],"quality":"More is better","isMain":true,"metricName":"Total length"},{"values":[6776,6662,6761,6803,6798,0,6271,6789,6717,6717,6717,6692],"quality":"More is better","isMain":true,"metricName":"Total length (>= 1000 bp)"},{"values":[6776,6662,6761,6803,6798,0,6271,6789,6717,6717,6717,6692],"quality":"More is better","isMain":false,"metricName":"Total length (>= 5000 bp)"},{"values":[0,0,0,0,0,0,0,0,0,0,0,0],"quality":"More is better","isMain":true,"metricName":"Total length (>= 10000 bp)"},{"values":[0,0,0,0,0,0,0,0,0,0,0,0],"quality":"More is better","isMain":false,"metricName":"Total length (>= 25000 bp)"},{"values":[0,0,0,0,0,0,0,0,0,0,0,0],"quality":"More is better","isMain":true,"metricName":"Total length (>= 50000 bp)"},{"values":[6776,6662,6761,6803,6798,556,6271,6789,6717,6717,6717,6692],"quality":"More is better","isMain":false,"metricName":"N50"},{"values":[6776,6662,6761,6803,6798,556,6271,6789,6717,6717,6717,6692],"quality":"More is better","isMain":false,"metricName":"N75"},{"values":[1,1,1,1,1,1,1,1,1,1,1,1],"quality":"Less is better","isMain":false,"metricName":"L50"},{"values":[1,1,1,1,1,1,1,1,1,1,1,1],"quality":"Less is better","isMain":false,"metricName":"L75"},{"values":["62.60","62.61","62.56","62.52","62.59","62.77","62.68","62.69","62.66","62.66","62.66","62.61"],"quality":"Equal","isMain":false,"metricName":"GC (%)"}]],["Predicted genes",[]],["Similarity statistics",[{"values":[0,0,0,0,0,0,0,0,0,0,0,0],"quality":"Equal","isMain":false,"metricName":"# similar correct contigs"},{"values":[0,0,0,0,0,0,0,0,0,0,0,0],"quality":"Equal","isMain":false,"metricName":"# similar misassembled blocks"}]],["Reference statistics",[{"values":[6662,6662,6662,6662,6662,6662,6662,6662,6662,6662,6662,6662],"quality":"Equal","isMain":false,"metricName":"Reference length"},{"values":[1,1,1,1,1,1,1,1,1,1,1,1],"quality":"Equal","isMain":false,"metricName":"Reference fragments"},{"values":["62.61","62.61","62.61","62.61","62.61","62.61","62.61","62.61","62.61","62.61","62.61","62.61"],"quality":"Equal","isMain":false,"metricName":"Reference GC (%)"}]]],"referenceName":"gi\_118430711\_ref\_NC","date":"18 July 2018, Wednesday, 16:10:52","order":[0,1,2,3,4,5,6,7,8,9,10,11],"assembliesNames":["ABySS\_127","CLC","IDBA\_UD","MEGAHIT","MIRA","Ray\_Meta","SOAPdenovo2","SPAdes","SPAdes\_meta","SPAdes\_sc","SPAdes\_sc\_careful","Velvet"]},{"assembliesWithNs":null,"minContig":500,"report":[["Genome statistics",[{"values":["13.103","10.865","9.335","10.838","10.351","23.180","4.368","12.150","20.098","14.459","10.511","6.845","30.616"],"quality":"More is better","isMain":true,"metricName":"Genome fraction (%)"},{"values":["1.056","1.013","1.005","1.021","1.032","1.000","1.000","1.024","1.019","1.000","1.000","1.010","1.540"],"quality":"Less is better","isMain":true,"metricName":"Duplication ratio"},{"values":[1060,990,1583,1060,1072,3011,1079,1076,2687,2158,1488,958,3851],"quality":"More is better","isMain":true,"metricName":"Largest alignment"},{"values":[5969,4748,4048,4773,4605,9997,1884,5367,8830,6236,4533,2982,20337],"quality":"More is better","isMain":true,"metricName":"Total aligned length"},{"values":[null,null,null,null,null,null,null,null,null,null,null,null,36887],"quality":"More is better","isMain":false,"metricName":"NG50"},{"values":[null,null,null,null,null,null,null,null,null,null,null,null,36887],"quality":"More is better","isMain":false,"metricName":"NG75"},{"values":[764,635,1017,612,764,2812,1079,764,1256,1921,1060,849,null],"quality":"More is better","isMain":false,"metricName":"NA50"},{"values":[612,612,833,537,612,1447,805,612,833,1097,889,604,null],"quality":"More is better","isMain":false,"metricName":"NA75"},{"values":[null,null,null,null,null,null,null,null,null,null,null,null,null],"quality":"More is better","isMain":true,"metricName":"NGA50"},{"values":[null,null,null,null,null,null,null,null,null,null,null,null,1],"quality":"Less is better","isMain":false,"metricName":"LG50"},{"values":[null,null,null,null,null,null,null,null,null,null,null,null,1],"quality":"Less is better","isMain":false,"metricName":"LG75"},{"values":[4,3,2,3,3,2,1,3,3,2,2,2,null],"quality":"Less is better","isMain":false,"metricName":"LA50"},{"values":[6,5,3,5,4,4,2,5,6,3,3,3,null],"quality":"Less is better","isMain":false,"metricName":"LA75"}]],["Misassemblies",[{"values":[0,0,0,0,0,0,0,0,0,0,0,0,0],"quality":"Less is better","isMain":true,"metricName":"# misassemblies"},{"values":[0,0,0,0,0,0,0,0,0,0,0,0,0],"quality":"Less is better","isMain":false,"metricName":" # relocations"},{"values":[0,0,0,0,0,0,0,0,0,0,0,0,0],"quality":"Less is better","isMain":false,"metricName":" # translocations"},{"values":[0,0,0,0,0,0,0,0,0,0,0,0,0],"quality":"Less is better","isMain":false,"metricName":" # inversions"},{"values":[0,0,0,0,0,0,0,0,0,0,0,0,0],"quality":"Less is better","isMain":false,"metricName":"# misassembled contigs"},{"values":[0,0,0,0,0,0,0,0,0,0,0,0,0],"quality":"Less is better","isMain":true,"metricName":"Misassembled contigs length"},{"values":[0,0,0,0,0,1,0,0,0,0,0,0,1],"quality":"Less is better","isMain":false,"metricName":"# local misassemblies"},{"values":[0,0,0,0,0,0,0,0,0,0,0,0,2],"quality":"Less is better","isMain":false,"metricName":"# unaligned mis. contigs"}]],["Unaligned",[{"values":[0,0,0,0,0,0,0,0,0,0,0,0,0],"quality":"Less is better","isMain":false,"metricName":"# fully unaligned contigs"},{"values":[0,0,0,0,0,0,0,0,0,0,0,0,0],"quality":"Less is better","isMain":false,"metricName":"Fully unaligned length"},{"values":[0,0,0,0,0,1,0,0,2,0,0,0,4],"quality":"Less is better","isMain":false,"metricName":"# partially unaligned contigs"},{"values":[0,0,0,0,0,1555,0,0,1455,0,0,0,58581],"quality":"Less is better","isMain":false,"metricName":"Partially unaligned length"}]],["Mismatches",[{"values":[0,4,2,0,0,130,0,0,107,44,4,0,178],"quality":"Less is better","isMain":false,"metricName":"# mismatches"},{"values":[0,0,0,0,0,4,0,0,0,0,0,0,9],"quality":"Less is better","isMain":false,"metricName":"# indels"},{"values":[0,0,0,0,0,4,0,0,0,0,0,0,9],"quality":"Less is better","isMain":false,"metricName":"Indels length"},{"values":["0.00","85.36","49.68","0.00","0.00","1300.39","0.00","0.00","1234.43","705.58","88.24","0.00","1348.08"],"quality":"Less is better","isMain":true,"metricName":"# mismatches per 100 kbp"},{"values":["0.00","0.00","0.00","0.00","0.00","40.01","0.00","0.00","0.00","0.00","0.00","0.00","68.16"],"quality":"Less is better","isMain":true,"metricName":"# indels per 100 kbp"},{"values":[0,0,0,0,0,4,0,0,0,0,0,0,9],"quality":"Less is better","isMain":false,"metricName":" # indels (<= 5 bp)"},{"values":[0,0,0,0,0,0,0,0,0,0,0,0,0],"quality":"Less is better","isMain":false,"metricName":" # indels (> 5 bp)"},{"values":[0,0,0,0,0,2,0,0,0,0,0,0,543],"quality":"Less is better","isMain":false,"metricName":"# N's"},{"values":["0.00","0.00","0.00","0.00","0.00","17.31","0.00","0.00","0.00","0.00","0.00","0.00","688.06"],"quality":"Less is better","isMain":true,"metricName":"# N's per 100 kbp"}]],["Statistics without reference",[{"values":[8,7,4,7,6,5,2,7,7,4,5,4,4],"quality":"Equal","isMain":true,"metricName":"# contigs"},{"values":[1,0,2,1,2,4,1,2,5,4,2,0,4],"quality":"Equal","isMain":false,"metricName":"# contigs (>= 1000 bp)"},{"values":[0,0,0,0,0,0,0,0,0,0,0,0,3],"quality":"Equal","isMain":false,"metricName":"# contigs (>= 5000 bp)"},{"values":[0,0,0,0,0,0,0,0,0,0,0,0,2],"quality":"Equal","isMain":false,"metricName":"# contigs (>= 10000 bp)"},{"values":[0,0,0,0,0,0,0,0,0,0,0,0,2],"quality":"Equal","isMain":false,"metricName":"# contigs (>= 25000 bp)"},{"values":[0,0,0,0,0,0,0,0,0,0,0,0,0],"quality":"Equal","isMain":false,"metricName":"# contigs (>= 50000 bp)"},{"values":[1060,990,1583,1060,1072,4566,1079,1076,2687,2158,1488,958,36887],"quality":"More is better","isMain":true,"metricName":"Largest contig"},{"values":[5969,4748,4048,4773,4605,11552,1884,5367,10285,6236,4533,2982,78918],"quality":"More is better","isMain":true,"metricName":"Total length"},{"values":[1060,0,2600,1060,2132,10560,1079,2136,8882,6236,2548,0,78918],"quality":"More is better","isMain":true,"metricName":"Total length (>= 1000 bp)"},{"values":[0,0,0,0,0,0,0,0,0,0,0,0,74352],"quality":"More is better","isMain":false,"metricName":"Total length (>= 5000 bp)"},{"values":[0,0,0,0,0,0,0,0,0,0,0,0,65450],"quality":"More is better","isMain":true,"metricName":"Total length (>= 10000 bp)"},{"values":[0,0,0,0,0,0,0,0,0,0,0,0,65450],"quality":"More is better","isMain":false,"metricName":"Total length (>= 25000 bp)"},{"values":[0,0,0,0,0,0,0,0,0,0,0,0,0],"quality":"More is better","isMain":true,"metricName":"Total length (>= 50000 bp)"},{"values":[764,635,1017,612,764,2812,1079,764,1926,1921,1060,849,28563],"quality":"More is better","isMain":false,"metricName":"N50"},{"values":[612,612,833,537,612,1735,805,612,1097,1097,889,604,28563],"quality":"More is better","isMain":false,"metricName":"N75"},{"values":[4,3,2,3,3,2,1,3,3,2,2,2,2],"quality":"Less is better","isMain":false,"metricName":"L50"},{"values":[6,5,3,5,4,3,2,5,4,3,3,3,2],"quality":"Less is better","isMain":false,"metricName":"L75"},{"values":["33.42","32.18","32.56","32.96","32.62","33.17","32.75","33.67","33.00","32.63","32.19","32.06","34.90"],"quality":"Equal","isMain":false,"metricName":"GC (%)"}]],["Predicted genes",[]],["Similarity statistics",[{"values":[0,0,0,0,0,0,0,0,0,0,0,0,0],"quality":"Equal","isMain":false,"metricName":"# similar correct contigs"},{"values":[0,0,0,0,0,0,0,0,0,0,0,0,0],"quality":"Equal","isMain":false,"metricName":"# similar misassembled blocks"}]],["Reference statistics",[{"values":[43128,43128,43128,43128,43128,43128,43128,43128,43128,43128,43128,43128,43128],"quality":"Equal","isMain":false,"metricName":"Reference length"},{"values":[1,1,1,1,1,1,1,1,1,1,1,1,1],"quality":"Equal","isMain":false,"metricName":"Reference fragments"},{"values":["34.16","34.16","34.16","34.16","34.16","34.16","34.16","34.16","34.16","34.16","34.16","34.16","34.16"],"quality":"Equal","isMain":false,"metricName":"Reference GC (%)"}]]],"referenceName":"gi\_118430724\_ref\_NC","date":"18 July 2018, Wednesday, 16:11:03","order":[0,1,2,3,4,5,6,7,8,9,10,11,12],"assembliesNames":["ABySS\_127","ABySS\_63","CLC","IDBA\_UD","MEGAHIT","MIRA","Ray\_Meta","SPAdes","SPAdes\_meta","SPAdes\_sc","SPAdes\_sc\_careful","Velvet","Geneious"]},{"assembliesWithNs":null,"minContig":500,"report":[["Genome statistics",[{"values":["67.704","55.269","93.614","93.596","94.572","84.278","84.921","97.020","95.773","95.291","94.835","78.507","34.885"],"quality":"More is better","isMain":true,"metricName":"Genome fraction (%)"},{"values":["1.002","1.002","1.012","1.003","1.002","1.029","1.012","1.002","1.003","1.002","1.002","1.001","0.999"],"quality":"Less is better","isMain":true,"metricName":"Duplication ratio"},{"values":[5757,3008,21238,21238,21219,9286,9978,27427,27049,26741,26742,19469,13833],"quality":"More is better","isMain":true,"metricName":"Largest alignment"},{"values":[217977,177932,303941,301643,304261,278574,274094,312278,308447,306340,304835,252377,111981],"quality":"More is better","isMain":true,"metricName":"Total aligned length"},{"values":[930,596,4632,5135,5542,2247,2573,7417,6355,7311,7301,3821,null],"quality":"More is better","isMain":false,"metricName":"NG50"},{"values":[null,null,2267,2688,3096,1206,1165,3708,3671,3671,3671,991,null],"quality":"More is better","isMain":false,"metricName":"NG75"},{"values":[1402,1098,5024,5652,5677,2352,2914,7417,6355,7301,7301,4913,5512],"quality":"More is better","isMain":false,"metricName":"NA50"},{"values":[910,797,2607,3340,3484,1679,1655,3821,3788,3788,3788,2396,4598],"quality":"More is better","isMain":false,"metricName":"NA75"},{"values":[929,596,4632,5135,5542,2220,2573,7417,6337,6355,6355,3821,null],"quality":"More is better","isMain":true,"metricName":"NGA50"},{"values":[null,null,2267,2688,3096,1206,1052,3708,3627,3542,3540,959,null],"quality":"More is better","isMain":false,"metricName":"NGA75"},{"values":[99,143,20,18,18,48,38,12,14,13,13,25,null],"quality":"Less is better","isMain":false,"metricName":"LG50"},{"values":[null,null,44,38,36,94,83,28,31,29,29,66,null],"quality":"Less is better","isMain":false,"metricName":"LG75"},{"values":[54,57,18,16,16,39,30,12,14,13,13,17,8],"quality":"Less is better","isMain":false,"metricName":"LA50"},{"values":[102,104,39,34,33,73,61,26,29,28,28,34,13],"quality":"Less is better","isMain":false,"metricName":"LA75"},{"values":[99,143,20,18,18,48,38,12,15,14,14,25,null],"quality":"Less is better","isMain":true,"metricName":"LGA50"},{"values":[null,null,44,38,36,95,85,28,32,31,31,67,null],"quality":"Less is better","isMain":false,"metricName":"LGA75"}]],["Misassemblies",[{"values":[1,0,1,1,0,1,0,0,1,3,3,1,0],"quality":"Less is better","isMain":true,"metricName":"# misassemblies"},{"values":[1,0,1,1,0,1,0,0,1,3,3,1,0],"quality":"Less is better","isMain":false,"metricName":" # relocations"},{"values":[0,0,0,0,0,0,0,0,0,0,0,0,0],"quality":"Less is better","isMain":false,"metricName":" # translocations"},{"values":[0,0,0,0,0,0,0,0,0,0,0,0,0],"quality":"Less is better","isMain":false,"metricName":" # inversions"},{"values":[1,0,1,1,0,1,0,0,1,2,2,1,0],"quality":"Less is better","isMain":false,"metricName":"# misassembled contigs"},{"values":[1114,0,1095,1851,0,2579,0,0,10601,30895,30895,2203,0],"quality":"Less is better","isMain":true,"metricName":"Misassembled contigs length"},{"values":[0,0,0,2,0,2,0,3,1,7,7,2,2],"quality":"Less is better","isMain":false,"metricName":"# local misassemblies"},{"values":[0,0,0,0,0,0,0,0,0,0,0,0,0],"quality":"Less is better","isMain":false,"metricName":"# unaligned mis. contigs"}]],["Unaligned",[{"values":[0,0,0,0,0,0,0,0,0,0,0,0,0],"quality":"Less is better","isMain":false,"metricName":"# fully unaligned contigs"},{"values":[0,0,0,0,0,0,0,0,0,0,0,0,0],"quality":"Less is better","isMain":false,"metricName":"Fully unaligned length"},{"values":[0,0,0,0,0,0,1,0,0,0,0,0,1],"quality":"Less is better","isMain":false,"metricName":"# partially unaligned contigs"},{"values":[0,0,0,0,0,0,728,0,0,0,0,0,5532],"quality":"Less is better","isMain":false,"metricName":"Partially unaligned length"}]],["Mismatches",[{"values":[77,29,160,191,90,72,75,225,225,178,145,92,64],"quality":"Less is better","isMain":false,"metricName":"# mismatches"},{"values":[1,0,5,4,1,1,31,4,13,6,6,4,1],"quality":"Less is better","isMain":false,"metricName":"# indels"},{"values":[12,0,66,133,21,84,1538,174,330,146,150,74,84],"quality":"Less is better","isMain":false,"metricName":"Indels length"},{"values":["35.40","16.33","53.20","63.53","29.62","26.59","27.49","72.19","73.13","58.15","47.60","36.48","57.11"],"quality":"Less is better","isMain":true,"metricName":"# mismatches per 100 kbp"},{"values":["0.46","0.00","1.66","1.33","0.33","0.37","11.36","1.28","4.23","1.96","1.97","1.59","0.89"],"quality":"Less is better","isMain":true,"metricName":"# indels per 100 kbp"},{"values":[0,0,3,0,0,0,0,0,5,0,0,2,0],"quality":"Less is better","isMain":false,"metricName":" # indels (<= 5 bp)"},{"values":[1,0,2,4,1,1,31,4,8,6,6,2,1],"quality":"Less is better","isMain":false,"metricName":" # indels (> 5 bp)"},{"values":[0,0,0,0,0,214,3164,0,0,348,316,10,96],"quality":"Less is better","isMain":false,"metricName":"# N's"},{"values":["0.00","0.00","0.00","0.00","0.00","76.82","1143.45","0.00","0.00","113.46","103.54","3.96","81.69"],"quality":"Less is better","isMain":true,"metricName":"# N's per 100 kbp"}]],["Statistics without reference",[{"values":[177,174,105,81,79,140,129,65,68,69,68,82,22],"quality":"Equal","isMain":true,"metricName":"# contigs"},{"values":[94,72,67,68,65,99,89,56,60,56,56,64,22],"quality":"Equal","isMain":false,"metricName":"# contigs (>= 1000 bp)"},{"values":[1,0,18,19,22,5,11,20,21,19,19,16,12],"quality":"Equal","isMain":false,"metricName":"# contigs (>= 5000 bp)"},{"values":[0,0,4,4,5,0,0,9,6,7,7,3,1],"quality":"Equal","isMain":false,"metricName":"# contigs (>= 10000 bp)"},{"values":[0,0,0,0,0,0,0,1,1,1,1,0,0],"quality":"Equal","isMain":false,"metricName":"# contigs (>= 25000 bp)"},{"values":[0,0,0,0,0,0,0,0,0,0,0,0,0],"quality":"Equal","isMain":false,"metricName":"# contigs (>= 50000 bp)"},{"values":[5757,3008,21238,21238,21219,9286,9978,27427,27049,26751,26751,19469,13833],"quality":"More is better","isMain":true,"metricName":"Largest contig"},{"values":[217977,177932,304268,301684,304271,278575,276706,312280,308568,306726,305209,252393,117513],"quality":"More is better","isMain":true,"metricName":"Total length"},{"values":[156489,105377,276747,292935,295132,247074,247960,305885,303101,297481,296684,239023,117513],"quality":"More is better","isMain":true,"metricName":"Total length (>= 1000 bp)"},{"values":[5757,0,154947,168947,187740,33019,73327,210685,201596,200130,200139,123935,87484],"quality":"More is better","isMain":false,"metricName":"Total length (>= 5000 bp)"},{"values":[0,0,61195,62198,75789,0,0,140021,97362,116743,116738,41073,13833],"quality":"More is better","isMain":true,"metricName":"Total length (>= 10000 bp)"},{"values":[0,0,0,0,0,0,0,27427,27049,26751,26751,0,0],"quality":"More is better","isMain":false,"metricName":"Total length (>= 25000 bp)"},{"values":[0,0,0,0,0,0,0,0,0,0,0,0,0],"quality":"More is better","isMain":true,"metricName":"Total length (>= 50000 bp)"},{"values":[1402,1098,5024,5652,5677,2358,2914,7417,7417,7417,7417,4913,6348],"quality":"More is better","isMain":false,"metricName":"N50"},{"values":[914,797,2624,3340,3484,1734,1678,3821,3821,3821,3821,2396,4812],"quality":"More is better","isMain":false,"metricName":"N75"},{"values":[54,57,18,16,16,39,30,12,13,12,12,17,7],"quality":"Less is better","isMain":false,"metricName":"L50"},{"values":[102,104,39,34,33,72,60,26,28,26,26,34,13],"quality":"Less is better","isMain":false,"metricName":"L75"},{"values":["44.68","44.50","44.44","44.49","44.54","44.48","44.27","44.55","44.51","44.45","44.42","44.30","44.47"],"quality":"Equal","isMain":false,"metricName":"GC (%)"}]],["Predicted genes",[]],["Similarity statistics",[{"values":[0,0,2,2,2,0,0,2,2,2,2,0,0],"quality":"Equal","isMain":false,"metricName":"# similar correct contigs"},{"values":[0,0,0,0,0,0,0,0,0,0,0,0,0],"quality":"Equal","isMain":false,"metricName":"# similar misassembled blocks"}]],["Reference statistics",[{"values":[321240,321240,321240,321240,321240,321240,321240,321240,321240,321240,321240,321240,321240],"quality":"Equal","isMain":false,"metricName":"Reference length"},{"values":[1,1,1,1,1,1,1,1,1,1,1,1,1],"quality":"Equal","isMain":false,"metricName":"Reference fragments"},{"values":["44.61","44.61","44.61","44.61","44.61","44.61","44.61","44.61","44.61","44.61","44.61","44.61","44.61"],"quality":"Equal","isMain":false,"metricName":"Reference GC (%)"}]]],"referenceName":"gi\_118496614\_ref\_NC","date":"18 July 2018, Wednesday, 16:11:15","order":[0,1,2,3,4,5,6,7,8,9,10,11,12],"assembliesNames":["ABySS\_127","ABySS\_63","CLC","IDBA\_UD","MEGAHIT","MIRA","SOAPdenovo2","SPAdes","SPAdes\_meta","SPAdes\_sc","SPAdes\_sc\_careful","Velvet","Geneious"]},{"assembliesWithNs":null,"minContig":500,"report":[["Genome statistics",[{"values":["46.540","25.975","56.676","51.592","58.296","80.597","3.116","9.103","67.080","34.647","38.994","42.616","23.767","74.592"],"quality":"More is better","isMain":true,"metricName":"Genome fraction (%)"},{"values":["1.018","1.007","1.034","1.026","1.055","1.393","1.000","1.181","1.034","1.082","1.012","1.018","1.002","1.076"],"quality":"Less is better","isMain":true,"metricName":"Duplication ratio"},{"values":[6995,2031,2827,3938,3501,9299,1373,2055,10579,5071,4638,5195,1769,14321],"quality":"More is better","isMain":true,"metricName":"Largest alignment"},{"values":[20876,11459,25616,23315,26872,48603,1373,4011,30382,16038,17325,19107,10498,34860],"quality":"More is better","isMain":true,"metricName":"Total aligned length"},{"values":[null,null,579,544,1103,6565,null,null,1439,null,null,null,null,28563],"quality":"More is better","isMain":false,"metricName":"NG50"},{"values":[null,null,null,null,null,4478,null,null,null,null,null,null,null,28444],"quality":"More is better","isMain":false,"metricName":"NG75"},{"values":[1467,1819,1284,1425,1467,3274,1373,777,2549,3800,1889,1357,1059,null],"quality":"More is better","isMain":false,"metricName":"NA50"},{"values":[1117,989,900,1012,1215,594,1373,null,1439,1351,1258,859,811,null],"quality":"More is better","isMain":false,"metricName":"NA75"},{"values":[null,null,579,544,1103,5855,null,null,1439,null,null,null,null,5863],"quality":"More is better","isMain":true,"metricName":"NGA50"},{"values":[null,null,null,null,null,2708,null,null,null,null,null,null,null,747],"quality":"More is better","isMain":false,"metricName":"NGA75"},{"values":[null,null,18,15,12,3,null,null,7,null,null,null,null,1],"quality":"Less is better","isMain":false,"metricName":"LG50"},{"values":[null,null,null,null,null,5,null,null,null,null,null,null,null,2],"quality":"Less is better","isMain":false,"metricName":"LG75"},{"values":[4,3,8,5,6,5,1,3,3,2,4,4,4,null],"quality":"Less is better","isMain":false,"metricName":"LA50"},{"values":[7,6,14,9,11,18,1,null,7,5,7,9,7,null],"quality":"Less is better","isMain":false,"metricName":"LA75"},{"values":[null,null,18,15,12,3,null,null,7,null,null,null,null,3],"quality":"Less is better","isMain":true,"metricName":"LGA50"},{"values":[null,null,null,null,null,6,null,null,null,null,null,null,null,7],"quality":"Less is better","isMain":false,"metricName":"LGA75"}]],["Misassemblies",[{"values":[0,0,0,0,0,0,0,0,0,1,0,0,0,0],"quality":"Less is better","isMain":true,"metricName":"# misassemblies"},{"values":[0,0,0,0,0,0,0,0,0,1,0,0,0,0],"quality":"Less is better","isMain":false,"metricName":" # relocations"},{"values":[0,0,0,0,0,0,0,0,0,0,0,0,0,0],"quality":"Less is better","isMain":false,"metricName":" # translocations"},{"values":[0,0,0,0,0,0,0,0,0,0,0,0,0,0],"quality":"Less is better","isMain":false,"metricName":" # inversions"},{"values":[0,0,0,0,0,0,0,0,0,1,0,0,0,0],"quality":"Less is better","isMain":false,"metricName":"# misassembled contigs"},{"values":[0,0,0,0,0,0,0,0,0,6917,0,0,0,0],"quality":"Less is better","isMain":true,"metricName":"Misassembled contigs length"},{"values":[0,0,0,0,0,4,0,1,0,0,0,0,0,2],"quality":"Less is better","isMain":false,"metricName":"# local misassemblies"},{"values":[0,0,0,0,0,1,0,0,0,0,0,0,0,2],"quality":"Less is better","isMain":false,"metricName":"# unaligned mis. contigs"}]],["Unaligned",[{"values":[0,0,0,0,0,0,0,0,0,0,0,0,0,0],"quality":"Less is better","isMain":false,"metricName":"# fully unaligned contigs"},{"values":[0,0,0,0,0,0,0,0,0,0,0,0,0,0],"quality":"Less is better","isMain":false,"metricName":"Fully unaligned length"},{"values":[0,0,0,0,0,7,0,2,0,0,1,1,0,5],"quality":"Less is better","isMain":false,"metricName":"# partially unaligned contigs"},{"values":[0,0,0,0,0,12948,0,2771,0,0,1690,1690,0,49785],"quality":"Less is better","isMain":false,"metricName":"Partially unaligned length"}]],["Mismatches",[{"values":[0,0,17,19,33,165,0,0,14,173,59,61,3,341],"quality":"Less is better","isMain":false,"metricName":"# mismatches"},{"values":[0,0,1,0,0,0,0,8,0,8,2,1,0,6],"quality":"Less is better","isMain":false,"metricName":"# indels"},{"values":[0,0,1,0,0,0,0,386,0,8,4,3,0,11],"quality":"Less is better","isMain":false,"metricName":"Indels length"},{"values":["0.00","0.00","68.08","83.58","128.47","464.63","0.00","0.00","47.37","1133.24","343.40","324.87","28.65","1037.55"],"quality":"Less is better","isMain":true,"metricName":"# mismatches per 100 kbp"},{"values":["0.00","0.00","4.00","0.00","0.00","0.00","0.00","199.45","0.00","52.40","11.64","5.33","0.00","18.26"],"quality":"Less is better","isMain":true,"metricName":"# indels per 100 kbp"},{"values":[0,0,1,0,0,0,0,0,0,8,2,1,0,5],"quality":"Less is better","isMain":false,"metricName":" # indels (<= 5 bp)"},{"values":[0,0,0,0,0,0,0,8,0,0,0,0,0,1],"quality":"Less is better","isMain":false,"metricName":" # indels (> 5 bp)"},{"values":[0,0,0,0,0,20,0,2164,0,0,0,0,0,925],"quality":"Less is better","isMain":false,"metricName":"# N's"},{"values":["0.00","0.00","0.00","0.00","0.00","32.03","0.00","28826.43","0.00","0.00","0.00","0.00","0.00","1086.41"],"quality":"Less is better","isMain":true,"metricName":"# N's per 100 kbp"}]],["Statistics without reference",[{"values":[14,9,24,17,19,21,1,4,16,7,11,14,11,7],"quality":"Equal","isMain":true,"metricName":"# contigs"},{"values":[7,5,12,9,12,15,1,4,9,4,7,7,4,7],"quality":"Equal","isMain":false,"metricName":"# contigs (>= 1000 bp)"},{"values":[1,0,0,0,0,4,0,0,1,1,0,1,0,4],"quality":"Equal","isMain":false,"metricName":"# contigs (>= 5000 bp)"},{"values":[0,0,0,0,0,0,0,0,1,0,0,0,0,3],"quality":"Equal","isMain":false,"metricName":"# contigs (>= 10000 bp)"},{"values":[0,0,0,0,0,0,0,0,0,0,0,0,0,2],"quality":"Equal","isMain":false,"metricName":"# contigs (>= 25000 bp)"},{"values":[0,0,0,0,0,0,0,0,0,0,0,0,0,0],"quality":"Equal","isMain":false,"metricName":"# contigs (>= 50000 bp)"},{"values":[6995,2031,2939,3938,3501,9755,1373,3304,10744,6917,4639,5195,1769,28563],"quality":"More is better","isMain":true,"metricName":"Largest contig"},{"values":[20876,11520,25817,23315,27086,62433,1373,7507,30547,16512,19070,20797,10498,85143],"quality":"More is better","isMain":true,"metricName":"Total length"},{"values":[15731,8556,17714,17939,22740,58260,1373,7507,25850,13971,16335,16148,5637,85143],"quality":"More is better","isMain":true,"metricName":"Total length (>= 1000 bp)"},{"values":[6995,0,0,0,0,31835,0,0,10744,6917,0,5195,0,77901],"quality":"More is better","isMain":false,"metricName":"Total length (>= 5000 bp)"},{"values":[0,0,0,0,0,0,0,0,10744,0,0,0,0,71328],"quality":"More is better","isMain":true,"metricName":"Total length (>= 10000 bp)"},{"values":[0,0,0,0,0,0,0,0,0,0,0,0,0,57007],"quality":"More is better","isMain":false,"metricName":"Total length (>= 25000 bp)"},{"values":[0,0,0,0,0,0,0,0,0,0,0,0,0,0],"quality":"More is better","isMain":true,"metricName":"Total length (>= 50000 bp)"},{"values":[1467,1880,1284,1425,1508,5832,1373,1698,2549,3800,2032,2032,1059,28444],"quality":"More is better","isMain":false,"metricName":"N50"},{"values":[1117,989,900,1012,1219,2708,1373,1336,1439,1898,1300,1218,811,14321],"quality":"More is better","isMain":false,"metricName":"N75"},{"values":[4,3,8,5,5,4,1,2,3,2,3,3,4,2],"quality":"Less is better","isMain":false,"metricName":"L50"},{"values":[7,6,14,9,10,9,1,3,7,3,6,7,7,3],"quality":"Less is better","isMain":false,"metricName":"L75"},{"values":["32.22","29.74","32.65","32.12","32.37","33.44","26.80","29.12","32.56","32.34","32.20","32.42","29.80","34.64"],"quality":"Equal","isMain":false,"metricName":"GC (%)"}]],["Predicted genes",[]],["Similarity statistics",[{"values":[0,0,0,0,0,0,0,0,0,0,0,0,0,0],"quality":"Equal","isMain":false,"metricName":"# similar correct contigs"},{"values":[0,0,0,0,0,0,0,0,0,0,0,0,0,0],"quality":"Equal","isMain":false,"metricName":"# similar misassembled blocks"}]],["Reference statistics",[{"values":[44061,44061,44061,44061,44061,44061,44061,44061,44061,44061,44061,44061,44061,44061],"quality":"Equal","isMain":false,"metricName":"Reference length"},{"values":[1,1,1,1,1,1,1,1,1,1,1,1,1,1],"quality":"Equal","isMain":false,"metricName":"Reference fragments"},{"values":["32.99","32.99","32.99","32.99","32.99","32.99","32.99","32.99","32.99","32.99","32.99","32.99","32.99","32.99"],"quality":"Equal","isMain":false,"metricName":"Reference GC (%)"}]]],"referenceName":"gi\_118725053\_ref\_NC","date":"18 July 2018, Wednesday, 16:11:26","order":[0,1,2,3,4,5,6,7,8,9,10,11,12,13],"assembliesNames":["ABySS\_127","ABySS\_63","CLC","IDBA\_UD","MEGAHIT","MIRA","Ray\_Meta","SOAPdenovo2","SPAdes","SPAdes\_meta","SPAdes\_sc","SPAdes\_sc\_careful","Velvet","Geneious"]},{"assembliesWithNs":null,"minContig":500,"report":[["Genome statistics",[{"values":["19.326","4.385","27.956","16.631","29.804","57.871","1.155","39.795","7.796","24.219","28.299","28.705","16.709"],"quality":"More is better","isMain":true,"metricName":"Genome fraction (%)"},{"values":["1.000","1.000","1.005","1.000","1.042","1.294","1.201","1.000","1.000","1.000","1.000","1.002","1.191"],"quality":"Less is better","isMain":true,"metricName":"Duplication ratio"},{"values":[3020,783,2484,3547,3304,3689,518,7111,2751,5052,7111,6282,3698],"quality":"More is better","isMain":true,"metricName":"Largest alignment"},{"values":[8669,1967,12607,7460,13933,33176,518,17852,3498,10863,12697,12905,8927],"quality":"More is better","isMain":true,"metricName":"Total aligned length"},{"values":[null,null,null,null,null,9674,null,null,null,null,null,null,42152],"quality":"More is better","isMain":false,"metricName":"NG50"},{"values":[null,null,null,null,null,8776,null,null,null,null,null,null,42152],"quality":"More is better","isMain":false,"metricName":"NG75"},{"values":[734,677,1344,807,885,565,518,2504,2751,1104,7111,2746,null],"quality":"More is better","isMain":false,"metricName":"NA50"},{"values":[592,507,602,680,577,null,518,1295,747,675,693,962,null],"quality":"More is better","isMain":false,"metricName":"NA75"},{"values":[null,null,null,null,null,995,null,null,null,null,null,null,null],"quality":"More is better","isMain":true,"metricName":"NGA50"},{"values":[null,null,null,null,null,2,null,null,null,null,null,null,1],"quality":"Less is better","isMain":false,"metricName":"LG50"},{"values":[null,null,null,null,null,3,null,null,null,null,null,null,1],"quality":"Less is better","isMain":false,"metricName":"LG75"},{"values":[3,2,4,2,4,23,1,2,1,2,1,2,null],"quality":"Less is better","isMain":false,"metricName":"LA50"},{"values":[6,3,8,4,9,null,1,5,2,5,5,3,null],"quality":"Less is better","isMain":false,"metricName":"LA75"},{"values":[null,null,null,null,null,12,null,null,null,null,null,null,null],"quality":"Less is better","isMain":true,"metricName":"LGA50"}]],["Misassemblies",[{"values":[0,0,0,0,0,0,0,0,0,0,0,0,0],"quality":"Less is better","isMain":true,"metricName":"# misassemblies"},{"values":[0,0,0,0,0,0,0,0,0,0,0,0,0],"quality":"Less is better","isMain":false,"metricName":" # relocations"},{"values":[0,0,0,0,0,0,0,0,0,0,0,0,0],"quality":"Less is better","isMain":false,"metricName":" # translocations"},{"values":[0,0,0,0,0,0,0,0,0,0,0,0,0],"quality":"Less is better","isMain":false,"metricName":" # inversions"},{"values":[0,0,0,0,0,0,0,0,0,0,0,0,0],"quality":"Less is better","isMain":false,"metricName":"# misassembled contigs"},{"values":[0,0,0,0,0,0,0,0,0,0,0,0,0],"quality":"Less is better","isMain":true,"metricName":"Misassembled contigs length"},{"values":[0,0,0,0,0,0,0,0,1,0,0,0,0],"quality":"Less is better","isMain":false,"metricName":"# local misassemblies"},{"values":[0,0,0,0,0,2,0,0,0,0,0,0,2],"quality":"Less is better","isMain":false,"metricName":"# unaligned mis. contigs"}]],["Unaligned",[{"values":[0,0,0,0,0,0,0,0,0,0,0,0,0],"quality":"Less is better","isMain":false,"metricName":"# fully unaligned contigs"},{"values":[0,0,0,0,0,0,0,0,0,0,0,0,0],"quality":"Less is better","isMain":false,"metricName":"Fully unaligned length"},{"values":[0,0,0,0,0,4,0,0,1,0,0,0,2],"quality":"Less is better","isMain":false,"metricName":"# partially unaligned contigs"},{"values":[0,0,0,0,0,26859,0,0,555,0,0,0,69953],"quality":"Less is better","isMain":false,"metricName":"Partially unaligned length"}]],["Mismatches",[{"values":[0,0,16,0,3,144,0,3,10,11,7,4,30],"quality":"Less is better","isMain":false,"metricName":"# mismatches"},{"values":[0,0,1,0,0,5,1,1,1,1,1,0,0],"quality":"Less is better","isMain":false,"metricName":"# indels"},{"values":[0,0,1,0,0,5,48,1,1,1,1,0,0],"quality":"Less is better","isMain":false,"metricName":"Indels length"},{"values":["0.00","0.00","127.59","0.00","22.44","554.72","0.00","16.81","285.96","101.25","55.14","31.07","400.27"],"quality":"Less is better","isMain":true,"metricName":"# mismatches per 100 kbp"},{"values":["0.00","0.00","7.97","0.00","0.00","19.26","193.05","5.60","28.60","9.20","7.88","0.00","0.00"],"quality":"Less is better","isMain":true,"metricName":"# indels per 100 kbp"},{"values":[0,0,1,0,0,5,0,1,1,1,1,0,0],"quality":"Less is better","isMain":false,"metricName":" # indels (<= 5 bp)"},{"values":[0,0,0,0,0,0,1,0,0,0,0,0,0],"quality":"Less is better","isMain":false,"metricName":" # indels (> 5 bp)"},{"values":[0,0,0,0,0,9,104,0,0,0,0,0,468],"quality":"Less is better","isMain":false,"metricName":"# N's"},{"values":["0.00","0.00","0.00","0.00","0.00","14.89","16720.26","0.00","0.00","0.00","0.00","0.00","593.31"],"quality":"Less is better","isMain":true,"metricName":"# N's per 100 kbp"}]],["Statistics without reference",[{"values":[10,3,13,7,15,26,1,9,2,9,9,8,3],"quality":"Equal","isMain":true,"metricName":"# contigs"},{"values":[1,0,5,1,3,11,0,7,1,2,1,2,3],"quality":"Equal","isMain":false,"metricName":"# contigs (>= 1000 bp)"},{"values":[0,0,0,0,0,3,0,1,0,1,1,1,2],"quality":"Equal","isMain":false,"metricName":"# contigs (>= 5000 bp)"},{"values":[0,0,0,0,0,1,0,0,0,0,0,0,2],"quality":"Equal","isMain":false,"metricName":"# contigs (>= 10000 bp)"},{"values":[0,0,0,0,0,0,0,0,0,0,0,0,2],"quality":"Equal","isMain":false,"metricName":"# contigs (>= 25000 bp)"},{"values":[0,0,0,0,0,0,0,0,0,0,0,0,0],"quality":"Equal","isMain":false,"metricName":"# contigs (>= 50000 bp)"},{"values":[3020,783,2484,3547,3304,16098,622,7111,3306,5052,7111,6282,42152],"quality":"More is better","isMain":true,"metricName":"Largest contig"},{"values":[8669,1967,12607,7460,13933,60449,622,17852,4053,10863,12697,12905,78880],"quality":"More is better","isMain":true,"metricName":"Total length"},{"values":[3020,0,7828,3547,6327,50131,0,16410,3306,6156,7111,9028,78880],"quality":"More is better","isMain":true,"metricName":"Total length (>= 1000 bp)"},{"values":[0,0,0,0,0,34548,0,7111,0,5052,7111,6282,75182],"quality":"More is better","isMain":false,"metricName":"Total length (>= 5000 bp)"},{"values":[0,0,0,0,0,16098,0,0,0,0,0,0,75182],"quality":"More is better","isMain":true,"metricName":"Total length (>= 10000 bp)"},{"values":[0,0,0,0,0,0,0,0,0,0,0,0,75182],"quality":"More is better","isMain":false,"metricName":"Total length (>= 25000 bp)"},{"values":[0,0,0,0,0,0,0,0,0,0,0,0,0],"quality":"More is better","isMain":true,"metricName":"Total length (>= 50000 bp)"},{"values":[734,677,1344,807,885,8776,622,2504,3306,1104,7111,2746,42152],"quality":"More is better","isMain":false,"metricName":"N50"},{"values":[592,507,602,680,577,1596,622,1295,3306,675,693,962,33030],"quality":"More is better","isMain":false,"metricName":"N75"},{"values":[3,2,4,2,4,3,1,2,1,2,1,2,1],"quality":"Less is better","isMain":false,"metricName":"L50"},{"values":[6,3,8,4,9,8,1,5,1,5,5,3,2],"quality":"Less is better","isMain":false,"metricName":"L75"},{"values":["33.78","31.77","33.10","33.91","34.06","33.57","31.85","33.89","32.35","33.97","33.24","33.37","34.51"],"quality":"Equal","isMain":false,"metricName":"GC (%)"}]],["Predicted genes",[]],["Similarity statistics",[{"values":[0,0,0,0,0,0,0,0,0,0,0,0,0],"quality":"Equal","isMain":false,"metricName":"# similar correct contigs"},{"values":[0,0,0,0,0,0,0,0,0,0,0,0,0],"quality":"Equal","isMain":false,"metricName":"# similar misassembled blocks"}]],["Reference statistics",[{"values":[44857,44857,44857,44857,44857,44857,44857,44857,44857,44857,44857,44857,44857],"quality":"Equal","isMain":false,"metricName":"Reference length"},{"values":[1,1,1,1,1,1,1,1,1,1,1,1,1],"quality":"Equal","isMain":false,"metricName":"Reference fragments"},{"values":["33.47","33.47","33.47","33.47","33.47","33.47","33.47","33.47","33.47","33.47","33.47","33.47","33.47"],"quality":"Equal","isMain":false,"metricName":"Reference GC (%)"}]]],"referenceName":"gi\_119443652\_ref\_NC","date":"18 July 2018, Wednesday, 16:11:37","order":[0,1,2,3,4,5,6,7,8,9,10,11,12],"assembliesNames":["ABySS\_127","ABySS\_63","CLC","IDBA\_UD","MEGAHIT","MIRA","SOAPdenovo2","SPAdes","SPAdes\_meta","SPAdes\_sc","SPAdes\_sc\_careful","Velvet","Geneious"]},{"assembliesWithNs":null,"minContig":500,"report":[["Genome statistics",[{"values":["100.000","100.000","100.000","99.766","83.011","62.060","100.000","100.000","100.000","100.000","100.000"],"quality":"More is better","isMain":true,"metricName":"Genome fraction (%)"},{"values":["1.000","1.001","1.002","1.005","1.019","1.020","1.002","1.001","1.001","1.001","1.000"],"quality":"Less is better","isMain":true,"metricName":"Duplication ratio"},{"values":[70153,70252,70294,69989,3577,21931,70280,70208,70208,70208,70183],"quality":"More is better","isMain":true,"metricName":"Largest alignment"},{"values":[70153,70252,70294,69989,59353,44093,70280,70208,70208,70208,70183],"quality":"More is better","isMain":true,"metricName":"Total aligned length"},{"values":[70153,70252,70294,70325,1387,29197,70280,70208,70208,70208,70183],"quality":"More is better","isMain":false,"metricName":"NG50"},{"values":[70153,70252,70294,70325,785,29197,70280,70208,70208,70208,70183],"quality":"More is better","isMain":false,"metricName":"NG75"},{"values":[70153,70252,70294,69989,1729,1808,70280,70208,70208,70208,70183],"quality":"More is better","isMain":false,"metricName":"NA50"},{"values":[70153,70252,70294,69989,903,null,70280,70208,70208,70208,70183],"quality":"More is better","isMain":false,"metricName":"NA75"},{"values":[70153,70252,70294,69989,1387,1808,70280,70208,70208,70208,70183],"quality":"More is better","isMain":true,"metricName":"NGA50"},{"values":[70153,70252,70294,69989,785,null,70280,70208,70208,70208,70183],"quality":"More is better","isMain":false,"metricName":"NGA75"},{"values":[1,1,1,1,16,2,1,1,1,1,1],"quality":"Less is better","isMain":false,"metricName":"LG50"},{"values":[1,1,1,1,34,2,1,1,1,1,1],"quality":"Less is better","isMain":false,"metricName":"LG75"},{"values":[1,1,1,1,12,5,1,1,1,1,1],"quality":"Less is better","isMain":false,"metricName":"LA50"},{"values":[1,1,1,1,24,null,1,1,1,1,1],"quality":"Less is better","isMain":false,"metricName":"LA75"},{"values":[1,1,1,1,16,5,1,1,1,1,1],"quality":"Less is better","isMain":true,"metricName":"LGA50"},{"values":[1,1,1,1,34,null,1,1,1,1,1],"quality":"Less is better","isMain":false,"metricName":"LGA75"}]],["Misassemblies",[{"values":[0,0,0,0,0,0,0,0,0,0,0],"quality":"Less is better","isMain":true,"metricName":"# misassemblies"},{"values":[0,0,0,0,0,0,0,0,0,0,0],"quality":"Less is better","isMain":false,"metricName":" # relocations"},{"values":[0,0,0,0,0,0,0,0,0,0,0],"quality":"Less is better","isMain":false,"metricName":" # translocations"},{"values":[0,0,0,0,0,0,0,0,0,0,0],"quality":"Less is better","isMain":false,"metricName":" # inversions"},{"values":[0,0,0,0,0,0,0,0,0,0,0],"quality":"Less is better","isMain":false,"metricName":"# misassembled contigs"},{"values":[0,0,0,0,0,0,0,0,0,0,0],"quality":"Less is better","isMain":true,"metricName":"Misassembled contigs length"},{"values":[0,1,1,0,0,2,1,0,0,0,0],"quality":"Less is better","isMain":false,"metricName":"# local misassemblies"},{"values":[0,0,0,0,0,1,0,0,0,0,0],"quality":"Less is better","isMain":false,"metricName":"# unaligned mis. contigs"}]],["Unaligned",[{"values":[0,0,0,0,0,0,0,0,0,0,0],"quality":"Less is better","isMain":false,"metricName":"# fully unaligned contigs"},{"values":[0,0,0,0,0,0,0,0,0,0,0],"quality":"Less is better","isMain":false,"metricName":"Fully unaligned length"},{"values":[0,0,0,0,0,2,0,0,0,0,0],"quality":"Less is better","isMain":false,"metricName":"# partially unaligned contigs"},{"values":[0,0,0,0,0,25703,0,0,0,0,0],"quality":"Less is better","isMain":false,"metricName":"Partially unaligned length"}]],["Mismatches",[{"values":[0,0,0,0,0,0,0,0,0,0,0],"quality":"Less is better","isMain":false,"metricName":"# mismatches"},{"values":[0,0,0,0,0,24,0,1,1,1,1],"quality":"Less is better","isMain":false,"metricName":"# indels"},{"values":[0,0,0,0,0,1122,0,55,55,55,30],"quality":"Less is better","isMain":false,"metricName":"Indels length"},{"values":["0.00","0.00","0.00","0.00","0.00","0.00","0.00","0.00","0.00","0.00","0.00"],"quality":"Less is better","isMain":true,"metricName":"# mismatches per 100 kbp"},{"values":["0.00","0.00","0.00","0.00","0.00","55.13","0.00","1.43","1.43","1.43","1.43"],"quality":"Less is better","isMain":true,"metricName":"# indels per 100 kbp"},{"values":[0,0,0,0,0,0,0,0,0,0,0],"quality":"Less is better","isMain":false,"metricName":" # indels (<= 5 bp)"},{"values":[0,0,0,0,0,24,0,1,1,1,1],"quality":"Less is better","isMain":false,"metricName":" # indels (> 5 bp)"},{"values":[0,0,0,0,0,5409,0,0,0,0,0],"quality":"Less is better","isMain":false,"metricName":"# N's"},{"values":["0.00","0.00","0.00","0.00","0.00","7716.23","0.00","0.00","0.00","0.00","0.00"],"quality":"Less is better","isMain":true,"metricName":"# N's per 100 kbp"}]],["Statistics without reference",[{"values":[1,1,1,1,44,8,1,1,1,1,1],"quality":"Equal","isMain":true,"metricName":"# contigs"},{"values":[1,1,1,1,23,4,1,1,1,1,1],"quality":"Equal","isMain":false,"metricName":"# contigs (>= 1000 bp)"},{"values":[1,1,1,1,0,3,1,1,1,1,1],"quality":"Equal","isMain":false,"metricName":"# contigs (>= 5000 bp)"},{"values":[1,1,1,1,0,2,1,1,1,1,1],"quality":"Equal","isMain":false,"metricName":"# contigs (>= 10000 bp)"},{"values":[1,1,1,1,0,2,1,1,1,1,1],"quality":"Equal","isMain":false,"metricName":"# contigs (>= 25000 bp)"},{"values":[1,1,1,1,0,0,1,1,1,1,1],"quality":"Equal","isMain":false,"metricName":"# contigs (>= 50000 bp)"},{"values":[70153,70252,70294,70325,3577,29941,70280,70208,70208,70208,70183],"quality":"More is better","isMain":true,"metricName":"Largest contig"},{"values":[70153,70252,70294,70325,59353,70099,70280,70208,70208,70208,70183],"quality":"More is better","isMain":true,"metricName":"Total length"},{"values":[70153,70252,70294,70325,43875,67413,70280,70208,70208,70208,70183],"quality":"More is better","isMain":true,"metricName":"Total length (>= 1000 bp)"},{"values":[70153,70252,70294,70325,0,65512,70280,70208,70208,70208,70183],"quality":"More is better","isMain":false,"metricName":"Total length (>= 5000 bp)"},{"values":[70153,70252,70294,70325,0,59138,70280,70208,70208,70208,70183],"quality":"More is better","isMain":true,"metricName":"Total length (>= 10000 bp)"},{"values":[70153,70252,70294,70325,0,59138,70280,70208,70208,70208,70183],"quality":"More is better","isMain":false,"metricName":"Total length (>= 25000 bp)"},{"values":[70153,70252,70294,70325,0,0,70280,70208,70208,70208,70183],"quality":"More is better","isMain":true,"metricName":"Total length (>= 50000 bp)"},{"values":[70153,70252,70294,70325,1729,29197,70280,70208,70208,70208,70183],"quality":"More is better","isMain":false,"metricName":"N50"},{"values":[70153,70252,70294,70325,903,29197,70280,70208,70208,70208,70183],"quality":"More is better","isMain":false,"metricName":"N75"},{"values":[1,1,1,1,12,2,1,1,1,1,1],"quality":"Less is better","isMain":false,"metricName":"L50"},{"values":[1,1,1,1,24,2,1,1,1,1,1],"quality":"Less is better","isMain":false,"metricName":"L75"},{"values":["41.30","41.31","41.31","41.29","41.31","41.22","41.30","41.30","41.30","41.30","41.30"],"quality":"Equal","isMain":false,"metricName":"GC (%)"}]],["Predicted genes",[]],["Similarity statistics",[{"values":[0,0,0,0,0,0,0,0,0,0,0],"quality":"Equal","isMain":false,"metricName":"# similar correct contigs"},{"values":[0,0,0,0,0,0,0,0,0,0,0],"quality":"Equal","isMain":false,"metricName":"# similar misassembled blocks"}]],["Reference statistics",[{"values":[70153,70153,70153,70153,70153,70153,70153,70153,70153,70153,70153],"quality":"Equal","isMain":false,"metricName":"Reference length"},{"values":[1,1,1,1,1,1,1,1,1,1,1],"quality":"Equal","isMain":false,"metricName":"Reference fragments"},{"values":["41.30","41.30","41.30","41.30","41.30","41.30","41.30","41.30","41.30","41.30","41.30"],"quality":"Equal","isMain":false,"metricName":"Reference GC (%)"}]]],"referenceName":"gi\_119952179\_ref\_NC","date":"18 July 2018, Wednesday, 16:11:47","order":[0,1,2,3,4,5,6,7,8,9,10],"assembliesNames":["CLC","IDBA\_UD","MEGAHIT","MIRA","Ray\_Meta","SOAPdenovo2","SPAdes","SPAdes\_meta","SPAdes\_sc","SPAdes\_sc\_careful","Velvet"]},{"assembliesWithNs":null,"minContig":500,"report":[["Genome statistics",[{"values":["69.500","49.452","86.142","69.493","81.962","100.000","1.398","26.285","91.578","24.348","53.086","55.046","32.213","99.988"],"quality":"More is better","isMain":true,"metricName":"Genome fraction (%)"},{"values":["1.010","1.005","1.000","1.010","1.018","1.372","1.000","1.273","1.000","1.000","1.006","1.008","1.003","1.507"],"quality":"Less is better","isMain":true,"metricName":"Duplication ratio"},{"values":[13127,3051,16636,5102,7697,40776,607,2499,31176,7763,3407,3463,2182,50789],"quality":"More is better","isMain":true,"metricName":"Largest alignment"},{"values":[30473,21586,37403,30473,36231,59576,607,11488,39763,10570,23184,24090,14022,65421],"quality":"More is better","isMain":true,"metricName":"Total aligned length"},{"values":[2366,null,13566,1515,5208,44069,null,null,31176,null,607,675,null,56062],"quality":"More is better","isMain":false,"metricName":"NG50"},{"values":[null,null,1927,null,1088,44069,null,null,2430,null,null,null,null,56062],"quality":"More is better","isMain":false,"metricName":"NG75"},{"values":[5165,2006,13566,2579,5198,5524,607,211,31176,7763,1842,1842,1512,50789],"quality":"More is better","isMain":false,"metricName":"NA50"},{"values":[2366,1392,13566,1015,1745,null,607,null,31176,2807,1434,935,921,null],"quality":"More is better","isMain":false,"metricName":"NA75"},{"values":[2366,null,13566,1515,5198,40776,null,null,31176,null,607,675,null,50789],"quality":"More is better","isMain":true,"metricName":"NGA50"},{"values":[null,null,1927,null,864,40776,null,null,2430,null,null,null,null,50789],"quality":"More is better","isMain":false,"metricName":"NGA75"},{"values":[4,null,2,8,4,1,null,null,1,null,14,13,null,1],"quality":"Less is better","isMain":false,"metricName":"LG50"},{"values":[null,null,4,null,8,1,null,null,2,null,null,null,null,1],"quality":"Less is better","isMain":false,"metricName":"LG75"},{"values":[2,5,2,4,4,2,1,9,1,1,5,5,4,1],"quality":"Less is better","isMain":false,"metricName":"LA50"},{"values":[4,8,2,9,7,null,1,null,1,2,8,9,7,null],"quality":"Less is better","isMain":false,"metricName":"LA75"},{"values":[4,null,2,8,4,1,null,null,1,null,14,13,null,1],"quality":"Less is better","isMain":true,"metricName":"LGA50"},{"values":[null,null,4,null,10,1,null,null,2,null,null,null,null,1],"quality":"Less is better","isMain":false,"metricName":"LGA75"}]],["Misassemblies",[{"values":[0,0,0,0,0,0,0,0,0,0,0,0,0,0],"quality":"Less is better","isMain":true,"metricName":"# misassemblies"},{"values":[0,0,0,0,0,0,0,0,0,0,0,0,0,0],"quality":"Less is better","isMain":false,"metricName":" # relocations"},{"values":[0,0,0,0,0,0,0,0,0,0,0,0,0,0],"quality":"Less is better","isMain":false,"metricName":" # translocations"},{"values":[0,0,0,0,0,0,0,0,0,0,0,0,0,0],"quality":"Less is better","isMain":false,"metricName":" # inversions"},{"values":[0,0,0,0,0,0,0,0,0,0,0,0,0,0],"quality":"Less is better","isMain":false,"metricName":"# misassembled contigs"},{"values":[0,0,0,0,0,0,0,0,0,0,0,0,0,0],"quality":"Less is better","isMain":true,"metricName":"Misassembled contigs length"},{"values":[0,0,0,0,0,2,0,0,0,0,0,0,0,2],"quality":"Less is better","isMain":false,"metricName":"# local misassemblies"},{"values":[0,0,0,0,0,1,0,3,0,0,0,0,0,1],"quality":"Less is better","isMain":false,"metricName":"# unaligned mis. contigs"}]],["Unaligned",[{"values":[0,0,0,0,0,0,0,0,0,0,0,0,0,0],"quality":"Less is better","isMain":false,"metricName":"# fully unaligned contigs"},{"values":[0,0,0,0,0,0,0,0,0,0,0,0,0,0],"quality":"Less is better","isMain":false,"metricName":"Fully unaligned length"},{"values":[0,0,0,0,1,2,0,4,0,0,0,0,0,2],"quality":"Less is better","isMain":false,"metricName":"# partially unaligned contigs"},{"values":[0,0,0,0,2509,23641,0,6325,0,0,0,0,0,25621],"quality":"Less is better","isMain":false,"metricName":"Partially unaligned length"}]],["Mismatches",[{"values":[2,0,48,2,23,397,0,0,1,95,2,1,0,498],"quality":"Less is better","isMain":false,"metricName":"# mismatches"},{"values":[0,0,0,0,0,8,0,12,0,10,0,0,0,11],"quality":"Less is better","isMain":false,"metricName":"# indels"},{"values":[0,0,0,0,0,8,0,601,0,10,0,0,0,11],"quality":"Less is better","isMain":false,"metricName":"Indels length"},{"values":["6.63","0.00","128.33","6.63","64.63","914.33","0.00","0.00","2.51","898.60","8.68","4.18","0.00","1147.07"],"quality":"Less is better","isMain":true,"metricName":"# mismatches per 100 kbp"},{"values":["0.00","0.00","0.00","0.00","0.00","18.42","0.00","105.14","0.00","94.59","0.00","0.00","0.00","25.34"],"quality":"Less is better","isMain":true,"metricName":"# indels per 100 kbp"},{"values":[0,0,0,0,0,8,0,0,0,10,0,0,0,11],"quality":"Less is better","isMain":false,"metricName":" # indels (<= 5 bp)"},{"values":[0,0,0,0,0,0,0,12,0,0,0,0,0,0],"quality":"Less is better","isMain":false,"metricName":" # indels (> 5 bp)"},{"values":[0,0,0,0,0,0,0,4935,0,0,0,0,0,182],"quality":"Less is better","isMain":false,"metricName":"# N's"},{"values":["0.00","0.00","0.00","0.00","0.00","0.00","0.00","23669.06","0.00","0.00","0.00","0.00","0.00","199.91"],"quality":"Less is better","isMain":true,"metricName":"# N's per 100 kbp"}]],["Statistics without reference",[{"values":[8,15,7,18,15,6,1,13,7,2,16,17,12,2],"quality":"Equal","isMain":true,"metricName":"# contigs"},{"values":[8,8,6,9,8,5,0,9,6,2,8,8,6,2],"quality":"Equal","isMain":false,"metricName":"# contigs (>= 1000 bp)"},{"values":[2,0,2,2,4,2,0,0,1,1,0,0,0,2],"quality":"Equal","isMain":false,"metricName":"# contigs (>= 5000 bp)"},{"values":[1,0,2,0,0,2,0,0,1,0,0,0,0,2],"quality":"Equal","isMain":false,"metricName":"# contigs (>= 10000 bp)"},{"values":[0,0,0,0,0,2,0,0,1,0,0,0,0,2],"quality":"Equal","isMain":false,"metricName":"# contigs (>= 25000 bp)"},{"values":[0,0,0,0,0,0,0,0,0,0,0,0,0,1],"quality":"Equal","isMain":false,"metricName":"# contigs (>= 50000 bp)"},{"values":[13127,3051,16636,5102,7707,44069,607,4335,31176,7763,3407,3463,2182,56062],"quality":"More is better","isMain":true,"metricName":"Largest contig"},{"values":[30473,21586,37403,30473,38740,83217,607,20850,39763,10570,23184,24090,14022,91042],"quality":"More is better","isMain":true,"metricName":"Total length"},{"values":[30473,16743,36827,23764,33585,82611,0,17826,39195,10570,17618,17894,9775,91042],"quality":"More is better","isMain":true,"metricName":"Total length (>= 1000 bp)"},{"values":[18292,0,30202,10118,26345,73818,0,0,31176,7763,0,0,0,91042],"quality":"More is better","isMain":false,"metricName":"Total length (>= 5000 bp)"},{"values":[13127,0,30202,0,0,73818,0,0,31176,0,0,0,0,91042],"quality":"More is better","isMain":true,"metricName":"Total length (>= 10000 bp)"},{"values":[0,0,0,0,0,73818,0,0,31176,0,0,0,0,91042],"quality":"More is better","isMain":false,"metricName":"Total length (>= 25000 bp)"},{"values":[0,0,0,0,0,0,0,0,0,0,0,0,0,56062],"quality":"More is better","isMain":true,"metricName":"Total length (>= 50000 bp)"},{"values":[5165,2006,13566,2579,5733,44069,607,1746,31176,7763,1842,1842,1512,56062],"quality":"More is better","isMain":false,"metricName":"N50"},{"values":[2366,1392,13566,1015,2011,29749,607,1226,31176,2807,1434,935,921,34980],"quality":"More is better","isMain":false,"metricName":"N75"},{"values":[2,5,2,4,3,1,1,4,1,1,5,5,4,1],"quality":"Less is better","isMain":false,"metricName":"L50"},{"values":[4,8,2,9,6,2,1,7,1,2,8,9,7,2],"quality":"Less is better","isMain":false,"metricName":"L75"},{"values":["34.18","33.64","34.75","34.16","34.78","34.82","33.94","34.11","34.61","33.84","33.70","33.81","32.98","34.75"],"quality":"Equal","isMain":false,"metricName":"GC (%)"}]],["Predicted genes",[]],["Similarity statistics",[{"values":[0,0,0,0,0,0,0,0,0,0,0,0,0,0],"quality":"Equal","isMain":false,"metricName":"# similar correct contigs"},{"values":[0,0,0,0,0,0,0,0,0,0,0,0,0,0],"quality":"Equal","isMain":false,"metricName":"# similar misassembled blocks"}]],["Reference statistics",[{"values":[43420,43420,43420,43420,43420,43420,43420,43420,43420,43420,43420,43420,43420,43420],"quality":"Equal","isMain":false,"metricName":"Reference length"},{"values":[1,1,1,1,1,1,1,1,1,1,1,1,1,1],"quality":"Equal","isMain":false,"metricName":"Reference fragments"},{"values":["34.68","34.68","34.68","34.68","34.68","34.68","34.68","34.68","34.68","34.68","34.68","34.68","34.68","34.68"],"quality":"Equal","isMain":false,"metricName":"Reference GC (%)"}]]],"referenceName":"gi\_119953678\_ref\_NC","date":"18 July 2018, Wednesday, 16:11:58","order":[0,1,2,3,4,5,6,7,8,9,10,11,12,13],"assembliesNames":["ABySS\_127","ABySS\_63","CLC","IDBA\_UD","MEGAHIT","MIRA","Ray\_Meta","SOAPdenovo2","SPAdes","SPAdes\_meta","SPAdes\_sc","SPAdes\_sc\_careful","Velvet","Geneious"]},{"assembliesWithNs":null,"minContig":500,"report":[["Genome statistics",[{"values":["97.144","93.464","99.019","99.611","99.694","98.658","93.835","99.209","99.578","98.811","98.811","96.999","55.799"],"quality":"More is better","isMain":true,"metricName":"Genome fraction (%)"},{"values":["1.001","1.002","1.006","1.001","1.000","1.023","1.032","1.000","1.001","1.002","1.002","1.001","1.001"],"quality":"Less is better","isMain":true,"metricName":"Duplication ratio"},{"values":[17271,10518,40294,47952,46206,42314,15485,64369,64436,56631,56568,17366,42314],"quality":"More is better","isMain":true,"metricName":"Largest alignment"},{"values":[280075,269854,286934,287122,287301,290731,272258,285787,287038,285179,285114,279694,160679],"quality":"More is better","isMain":true,"metricName":"Total aligned length"},{"values":[4547,3618,17425,32128,34853,28866,6641,42592,38869,33433,33431,9319,14561],"quality":"More is better","isMain":false,"metricName":"NG50"},{"values":[2659,1806,10065,20464,29113,13465,3661,39059,29161,29161,29161,4452,null],"quality":"More is better","isMain":false,"metricName":"NG75"},{"values":[4983,3822,17425,32128,34853,28866,6504,42592,38846,33433,33431,9968,32545],"quality":"More is better","isMain":false,"metricName":"NA50"},{"values":[3039,2454,10065,20464,29113,13465,3600,39036,29161,29161,29161,4737,30087],"quality":"More is better","isMain":false,"metricName":"NA75"},{"values":[4547,3618,17425,32128,34853,28866,6417,42592,38846,33433,33431,9319,14561],"quality":"More is better","isMain":true,"metricName":"NGA50"},{"values":[2659,1806,10065,20464,29113,13465,3481,39036,29161,29161,29161,4452,null],"quality":"More is better","isMain":false,"metricName":"NGA75"},{"values":[18,27,6,4,4,4,15,3,3,4,4,12,5],"quality":"Less is better","isMain":false,"metricName":"LG50"},{"values":[37,53,11,7,6,9,29,5,6,6,6,22,null],"quality":"Less is better","isMain":false,"metricName":"LG75"},{"values":[17,25,6,4,4,4,15,3,3,4,4,11,3],"quality":"Less is better","isMain":false,"metricName":"LA50"},{"values":[35,47,11,7,6,9,29,5,6,6,6,21,4],"quality":"Less is better","isMain":false,"metricName":"LA75"},{"values":[18,27,6,4,4,4,16,3,3,4,4,12,5],"quality":"Less is better","isMain":true,"metricName":"LGA50"},{"values":[37,53,11,7,6,9,30,5,6,6,6,22,null],"quality":"Less is better","isMain":false,"metricName":"LGA75"}]],["Misassemblies",[{"values":[0,0,0,0,0,0,0,0,0,0,0,0,0],"quality":"Less is better","isMain":true,"metricName":"# misassemblies"},{"values":[0,0,0,0,0,0,0,0,0,0,0,0,0],"quality":"Less is better","isMain":false,"metricName":" # relocations"},{"values":[0,0,0,0,0,0,0,0,0,0,0,0,0],"quality":"Less is better","isMain":false,"metricName":" # translocations"},{"values":[0,0,0,0,0,0,0,0,0,0,0,0,0],"quality":"Less is better","isMain":false,"metricName":" # inversions"},{"values":[0,0,0,0,0,0,0,0,0,0,0,0,0],"quality":"Less is better","isMain":false,"metricName":"# misassembled contigs"},{"values":[0,0,0,0,0,0,0,0,0,0,0,0,0],"quality":"Less is better","isMain":true,"metricName":"Misassembled contigs length"},{"values":[0,0,0,0,1,2,0,1,1,5,5,0,1],"quality":"Less is better","isMain":false,"metricName":"# local misassemblies"},{"values":[0,0,0,0,0,0,1,0,0,0,0,0,0],"quality":"Less is better","isMain":false,"metricName":"# unaligned mis. contigs"}]],["Unaligned",[{"values":[0,0,0,0,0,0,0,0,0,0,0,0,0],"quality":"Less is better","isMain":false,"metricName":"# fully unaligned contigs"},{"values":[0,0,0,0,0,0,0,0,0,0,0,0,0],"quality":"Less is better","isMain":false,"metricName":"Fully unaligned length"},{"values":[0,0,0,0,0,0,5,0,0,2,2,0,0],"quality":"Less is better","isMain":false,"metricName":"# partially unaligned contigs"},{"values":[0,0,0,0,0,0,4132,0,0,1506,1506,0,0],"quality":"Less is better","isMain":false,"metricName":"Partially unaligned length"}]],["Mismatches",[{"values":[21,5,17,8,9,7,9,26,54,35,25,9,1],"quality":"Less is better","isMain":false,"metricName":"# mismatches"},{"values":[0,0,0,1,1,3,89,0,4,5,4,4,2],"quality":"Less is better","isMain":false,"metricName":"# indels"},{"values":[0,0,0,27,1,121,4315,0,27,91,52,90,120],"quality":"Less is better","isMain":false,"metricName":"Indels length"},{"values":["7.50","1.86","5.96","2.79","3.13","2.46","3.33","9.10","18.83","12.30","8.78","3.22","0.62"],"quality":"Less is better","isMain":true,"metricName":"# mismatches per 100 kbp"},{"values":["0.00","0.00","0.00","0.35","0.35","1.06","32.93","0.00","1.39","1.76","1.41","1.43","1.24"],"quality":"Less is better","isMain":true,"metricName":"# indels per 100 kbp"},{"values":[0,0,0,0,1,1,0,0,2,2,2,1,0],"quality":"Less is better","isMain":false,"metricName":" # indels (<= 5 bp)"},{"values":[0,0,0,1,0,2,89,0,2,3,2,3,2],"quality":"Less is better","isMain":false,"metricName":" # indels (> 5 bp)"},{"values":[0,0,0,0,0,59,9754,0,100,1529,1499,10,136],"quality":"Less is better","isMain":false,"metricName":"# N's"},{"values":["0.00","0.00","0.00","0.00","0.00","20.29","3445.69","0.00","34.82","533.30","522.95","3.57","84.50"],"quality":"Less is better","isMain":true,"metricName":"# N's per 100 kbp"}]],["Statistics without reference",[{"values":[83,108,28,17,15,28,64,8,10,16,16,51,6],"quality":"Equal","isMain":true,"metricName":"# contigs"},{"values":[67,73,24,14,15,21,54,8,10,11,11,44,6],"quality":"Equal","isMain":false,"metricName":"# contigs (>= 1000 bp)"},{"values":[16,11,19,11,11,13,25,8,10,11,11,20,5],"quality":"Equal","isMain":false,"metricName":"# contigs (>= 5000 bp)"},{"values":[4,1,11,9,8,10,4,7,8,7,7,10,5],"quality":"Equal","isMain":false,"metricName":"# contigs (>= 10000 bp)"},{"values":[0,0,3,6,6,4,0,6,6,6,6,0,4],"quality":"Equal","isMain":false,"metricName":"# contigs (>= 25000 bp)"},{"values":[0,0,0,0,0,0,0,1,1,1,1,0,0],"quality":"Equal","isMain":false,"metricName":"# contigs (>= 50000 bp)"},{"values":[17271,10518,40294,47952,46206,42314,16065,64369,64536,56631,56568,17366,42314],"quality":"More is better","isMain":true,"metricName":"Largest contig"},{"values":[280098,269854,286966,287122,287301,290731,283078,285810,287161,286708,286643,279733,160942],"quality":"More is better","isMain":true,"metricName":"Total length"},{"values":[268340,244224,283798,285151,287301,285519,275665,285810,287161,283326,283261,275015,160942],"quality":"More is better","isMain":true,"metricName":"Total length (>= 1000 bp)"},{"values":[137140,79001,274432,278158,277869,264903,202590,285810,287161,283326,283261,207625,156169],"quality":"More is better","isMain":false,"metricName":"Total length (>= 5000 bp)"},{"values":[51140,10518,216218,259153,250482,240355,57118,276497,268026,248468,248403,133751,156169],"quality":"More is better","isMain":true,"metricName":"Total length (>= 10000 bp)"},{"values":[0,0,104979,214251,223698,149270,0,262740,243085,234711,234646,0,141608],"quality":"More is better","isMain":false,"metricName":"Total length (>= 25000 bp)"},{"values":[0,0,0,0,0,0,0,64369,64536,56631,56568,0,0],"quality":"More is better","isMain":true,"metricName":"Total length (>= 50000 bp)"},{"values":[4983,3822,17425,32128,34853,28866,6641,42592,38869,33433,33431,9968,32545],"quality":"More is better","isMain":false,"metricName":"N50"},{"values":[3039,2454,10065,20464,29113,13465,4044,39059,29161,29161,29161,4737,30087],"quality":"More is better","isMain":false,"metricName":"N75"},{"values":[17,25,6,4,4,4,15,3,3,4,4,11,3],"quality":"Less is better","isMain":false,"metricName":"L50"},{"values":[35,47,11,7,6,9,28,5,6,6,6,21,4],"quality":"Less is better","isMain":false,"metricName":"L75"},{"values":["49.39","49.49","49.40","49.40","49.40","49.42","49.25","49.38","49.40","49.37","49.37","49.24","49.79"],"quality":"Equal","isMain":false,"metricName":"GC (%)"}]],["Predicted genes",[]],["Similarity statistics",[{"values":[0,0,1,2,2,2,1,2,2,2,2,1,0],"quality":"Equal","isMain":false,"metricName":"# similar correct contigs"},{"values":[0,0,0,0,0,0,0,0,0,0,0,0,0],"quality":"Equal","isMain":false,"metricName":"# similar misassembled blocks"}]],["Reference statistics",[{"values":[288047,288047,288047,288047,288047,288047,288047,288047,288047,288047,288047,288047,288047],"quality":"Equal","isMain":false,"metricName":"Reference length"},{"values":[1,1,1,1,1,1,1,1,1,1,1,1,1],"quality":"Equal","isMain":false,"metricName":"Reference fragments"},{"values":["49.40","49.40","49.40","49.40","49.40","49.40","49.40","49.40","49.40","49.40","49.40","49.40","49.40"],"quality":"Equal","isMain":false,"metricName":"Reference GC (%)"}]]],"referenceName":"gi\_119953744\_ref\_NC","date":"18 July 2018, Wednesday, 16:12:10","order":[0,1,2,3,4,5,6,7,8,9,10,11,12],"assembliesNames":["ABySS\_127","ABySS\_63","CLC","IDBA\_UD","MEGAHIT","MIRA","SOAPdenovo2","SPAdes","SPAdes\_meta","SPAdes\_sc","SPAdes\_sc\_careful","Velvet","Geneious"]},{"assembliesWithNs":null,"minContig":500,"report":[["Genome statistics",[{"values":["99.816","99.535","100.000","95.422","99.850","100.000","98.281","4.444","99.180","100.000","97.255","92.814","40.603","100.000"],"quality":"More is better","isMain":true,"metricName":"Genome fraction (%)"},{"values":["1.003","1.037","1.042","1.034","1.090","1.810","1.032","1.000","1.023","1.004","1.028","1.019","1.002","1.530"],"quality":"Less is better","isMain":true,"metricName":"Duplication ratio"},{"values":[43960,18760,32356,5864,15070,27981,15661,714,34309,16086,14052,14349,3047,33206],"quality":"More is better","isMain":true,"metricName":"Largest alignment"},{"values":[43960,45438,45906,43438,47915,79703,44664,1957,44696,44206,43492,41426,17915,67385],"quality":"More is better","isMain":true,"metricName":"Total aligned length"},{"values":[44091,11654,32356,2856,7707,44069,12142,null,34309,14480,9843,4062,null,56062],"quality":"More is better","isMain":false,"metricName":"NG50"},{"values":[44091,9558,6371,1593,5599,44069,11700,null,34309,13640,3673,1375,null,56062],"quality":"More is better","isMain":false,"metricName":"NG75"},{"values":[43960,11654,32356,3642,5764,8042,12142,704,34309,14480,9283,4062,1053,17964],"quality":"More is better","isMain":false,"metricName":"NA50"},{"values":[43960,9558,6371,1786,2458,758,11700,539,34309,13640,3673,1627,820,null],"quality":"More is better","isMain":false,"metricName":"NA75"},{"values":[43960,11654,32356,2856,5764,27981,12142,null,34309,14480,9283,4062,null,33206],"quality":"More is better","isMain":true,"metricName":"NGA50"},{"values":[43960,9558,6371,1593,3922,15660,11700,null,34309,13640,3673,1375,null,33206],"quality":"More is better","isMain":false,"metricName":"NGA75"},{"values":[1,2,1,6,2,1,2,null,1,2,2,3,null,1],"quality":"Less is better","isMain":false,"metricName":"LG50"},{"values":[1,3,2,11,4,1,3,null,1,3,4,8,null,1],"quality":"Less is better","isMain":false,"metricName":"LG75"},{"values":[1,2,1,5,3,3,2,2,1,2,2,3,5,2],"quality":"Less is better","isMain":false,"metricName":"LA50"},{"values":[1,3,2,10,6,13,3,3,1,3,4,7,10,null],"quality":"Less is better","isMain":false,"metricName":"LA75"},{"values":[1,2,1,6,3,1,2,null,1,2,2,3,null,1],"quality":"Less is better","isMain":true,"metricName":"LGA50"},{"values":[1,3,2,11,5,2,3,null,1,3,4,8,null,1],"quality":"Less is better","isMain":false,"metricName":"LGA75"}]],["Misassemblies",[{"values":[0,0,0,0,0,1,0,0,0,0,0,0,0,1],"quality":"Less is better","isMain":true,"metricName":"# misassemblies"},{"values":[0,0,0,0,0,1,0,0,0,0,0,0,0,1],"quality":"Less is better","isMain":false,"metricName":" # relocations"},{"values":[0,0,0,0,0,0,0,0,0,0,0,0,0,0],"quality":"Less is better","isMain":false,"metricName":" # translocations"},{"values":[0,0,0,0,0,0,0,0,0,0,0,0,0,0],"quality":"Less is better","isMain":false,"metricName":" # inversions"},{"values":[0,0,0,0,0,1,0,0,0,0,0,0,0,1],"quality":"Less is better","isMain":false,"metricName":"# misassembled contigs"},{"values":[0,0,0,0,0,44069,0,0,0,0,0,0,0,56062],"quality":"Less is better","isMain":true,"metricName":"Misassembled contigs length"},{"values":[0,0,0,0,1,2,0,0,0,0,0,0,0,3],"quality":"Less is better","isMain":false,"metricName":"# local misassemblies"},{"values":[0,0,0,0,0,0,0,0,0,0,0,0,0,0],"quality":"Less is better","isMain":false,"metricName":"# unaligned mis. contigs"}]],["Unaligned",[{"values":[0,0,0,0,0,0,0,0,0,0,0,0,0,0],"quality":"Less is better","isMain":false,"metricName":"# fully unaligned contigs"},{"values":[0,0,0,0,0,0,0,0,0,0,0,0,0,0],"quality":"Less is better","isMain":false,"metricName":"Fully unaligned length"},{"values":[0,0,0,0,1,2,0,0,0,0,0,0,0,2],"quality":"Less is better","isMain":false,"metricName":"# partially unaligned contigs"},{"values":[0,0,0,0,1943,22135,0,0,0,0,0,0,0,23657],"quality":"Less is better","isMain":false,"metricName":"Partially unaligned length"}]],["Mismatches",[{"values":[0,0,0,0,167,398,1,2,0,1,0,2,2,194],"quality":"Less is better","isMain":false,"metricName":"# mismatches"},{"values":[0,0,0,0,0,7,0,0,0,0,0,1,0,5],"quality":"Less is better","isMain":false,"metricName":"# indels"},{"values":[0,0,0,0,0,7,0,0,0,0,0,1,0,5],"quality":"Less is better","isMain":false,"metricName":"Indels length"},{"values":["0.00","0.00","0.00","0.00","379.76","903.70","2.31","102.20","0.00","2.27","0.00","4.89","11.18","440.50"],"quality":"Less is better","isMain":true,"metricName":"# mismatches per 100 kbp"},{"values":["0.00","0.00","0.00","0.00","0.00","15.89","0.00","0.00","0.00","0.00","0.00","2.45","0.00","11.35"],"quality":"Less is better","isMain":true,"metricName":"# indels per 100 kbp"},{"values":[0,0,0,0,0,7,0,0,0,0,0,1,0,5],"quality":"Less is better","isMain":false,"metricName":" # indels (<= 5 bp)"},{"values":[0,0,0,0,0,0,0,0,0,0,0,0,0,0],"quality":"Less is better","isMain":false,"metricName":" # indels (> 5 bp)"},{"values":[0,8,0,0,0,1,0,0,0,0,560,204,0,182],"quality":"Less is better","isMain":false,"metricName":"# N's"},{"values":["0.00","17.61","0.00","0.00","0.00","0.98","0.00","0.00","0.00","0.00","1271.22","490.00","0.00","199.91"],"quality":"Less is better","isMain":true,"metricName":"# N's per 100 kbp"}]],["Statistics without reference",[{"values":[1,5,8,20,13,17,4,3,10,3,14,19,17,2],"quality":"Equal","isMain":true,"metricName":"# contigs"},{"values":[1,4,6,16,11,8,4,0,7,3,8,9,5,2],"quality":"Equal","isMain":false,"metricName":"# contigs (>= 1000 bp)"},{"values":[1,3,2,1,4,3,4,0,1,3,3,2,0,2],"quality":"Equal","isMain":false,"metricName":"# contigs (>= 5000 bp)"},{"values":[1,2,1,0,1,2,3,0,1,3,1,1,0,2],"quality":"Equal","isMain":false,"metricName":"# contigs (>= 10000 bp)"},{"values":[1,0,1,0,0,2,0,0,1,0,0,0,0,2],"quality":"Equal","isMain":false,"metricName":"# contigs (>= 25000 bp)"},{"values":[0,0,0,0,0,0,0,0,0,0,0,0,0,1],"quality":"Equal","isMain":false,"metricName":"# contigs (>= 50000 bp)"},{"values":[44091,18760,32356,5864,15070,44069,15663,714,34309,16086,14052,14349,3047,56062],"quality":"More is better","isMain":true,"metricName":"Largest contig"},{"values":[44091,45438,45906,43438,49858,101838,44666,1957,44696,44206,44052,41633,17915,91042],"quality":"More is better","isMain":true,"metricName":"Total length"},{"values":[44091,44910,44742,40455,48466,95382,44666,0,42924,44206,39661,35163,9104,91042],"quality":"More is better","isMain":true,"metricName":"Total length (>= 1000 bp)"},{"values":[44091,39972,38727,5864,34176,80282,44666,0,34309,44206,30298,20752,0,91042],"quality":"More is better","isMain":false,"metricName":"Total length (>= 5000 bp)"},{"values":[44091,30414,32356,0,15070,73818,39505,0,34309,44206,14052,14349,0,91042],"quality":"More is better","isMain":true,"metricName":"Total length (>= 10000 bp)"},{"values":[44091,0,32356,0,0,73818,0,0,34309,0,0,0,0,91042],"quality":"More is better","isMain":false,"metricName":"Total length (>= 25000 bp)"},{"values":[0,0,0,0,0,0,0,0,0,0,0,0,0,56062],"quality":"More is better","isMain":true,"metricName":"Total length (>= 50000 bp)"},{"values":[44091,11654,32356,3642,5800,29749,12142,704,34309,14480,9843,4062,1053,56062],"quality":"More is better","isMain":false,"metricName":"N50"},{"values":[44091,9558,6371,1786,3922,6464,11700,539,34309,13640,3673,1627,820,34980],"quality":"More is better","isMain":false,"metricName":"N75"},{"values":[1,2,1,5,3,2,2,2,1,2,2,3,5,1],"quality":"Less is better","isMain":false,"metricName":"L50"},{"values":[1,3,2,10,5,3,3,3,1,3,4,7,10,2],"quality":"Less is better","isMain":false,"metricName":"L75"},{"values":["34.91","34.93","34.74","34.82","34.92","34.73","34.99","34.90","34.92","34.93","34.85","34.82","34.07","34.75"],"quality":"Equal","isMain":false,"metricName":"GC (%)"}]],["Predicted genes",[]],["Similarity statistics",[{"values":[0,0,0,0,0,0,0,0,0,0,0,0,0,0],"quality":"Equal","isMain":false,"metricName":"# similar correct contigs"},{"values":[0,0,0,0,0,0,0,0,0,0,0,0,0,0],"quality":"Equal","isMain":false,"metricName":"# similar misassembled blocks"}]],["Reference statistics",[{"values":[44041,44041,44041,44041,44041,44041,44041,44041,44041,44041,44041,44041,44041,44041],"quality":"Equal","isMain":false,"metricName":"Reference length"},{"values":[1,1,1,1,1,1,1,1,1,1,1,1,1,1],"quality":"Equal","isMain":false,"metricName":"Reference fragments"},{"values":["34.91","34.91","34.91","34.91","34.91","34.91","34.91","34.91","34.91","34.91","34.91","34.91","34.91","34.91"],"quality":"Equal","isMain":false,"metricName":"Reference GC (%)"}]]],"referenceName":"gi\_119967833\_ref\_NC","date":"18 July 2018, Wednesday, 16:12:22","order":[0,1,2,3,4,5,6,7,8,9,10,11,12,13],"assembliesNames":["ABySS\_127","ABySS\_63","CLC","IDBA\_UD","MEGAHIT","MIRA","Ray\_Meta","SOAPdenovo2","SPAdes","SPAdes\_meta","SPAdes\_sc","SPAdes\_sc\_careful","Velvet","Geneious"]},{"assembliesWithNs":null,"minContig":500,"report":[["Genome statistics",[{"values":["100.000","100.000","100.000","100.000","100.000","100.000","100.000"],"quality":"More is better","isMain":true,"metricName":"Genome fraction (%)"},{"values":["1.000","243.432","1.028","1.012","1.012","1.012","1.670"],"quality":"Less is better","isMain":true,"metricName":"Duplication ratio"},{"values":[4594,1847,4721,4649,4649,4649,4127],"quality":"More is better","isMain":true,"metricName":"Largest alignment"},{"values":[4594,1116731,4721,4649,4649,4649,7670],"quality":"More is better","isMain":true,"metricName":"Total aligned length"},{"values":[4594,1502,4721,4649,4649,4649,7670],"quality":"More is better","isMain":false,"metricName":"NG50"},{"values":[4594,1458,4721,4649,4649,4649,7670],"quality":"More is better","isMain":false,"metricName":"NG75"},{"values":[4594,719,4721,4649,4649,4649,4127],"quality":"More is better","isMain":false,"metricName":"NA50"},{"values":[4594,576,4721,4649,4649,4649,3543],"quality":"More is better","isMain":false,"metricName":"NA75"},{"values":[4594,1502,4721,4649,4649,4649,4127],"quality":"More is better","isMain":true,"metricName":"NGA50"},{"values":[4594,1458,4721,4649,4649,4649,4127],"quality":"More is better","isMain":false,"metricName":"NGA75"},{"values":[1,2,1,1,1,1,1],"quality":"Less is better","isMain":false,"metricName":"LG50"},{"values":[1,3,1,1,1,1,1],"quality":"Less is better","isMain":false,"metricName":"LG75"},{"values":[1,634,1,1,1,1,1],"quality":"Less is better","isMain":false,"metricName":"LA50"},{"values":[1,1085,1,1,1,1,2],"quality":"Less is better","isMain":false,"metricName":"LA75"},{"values":[1,2,1,1,1,1,1],"quality":"Less is better","isMain":true,"metricName":"LGA50"},{"values":[1,3,1,1,1,1,1],"quality":"Less is better","isMain":false,"metricName":"LGA75"}]],["Misassemblies",[{"values":[0,0,0,0,0,0,1],"quality":"Less is better","isMain":true,"metricName":"# misassemblies"},{"values":[0,0,0,0,0,0,1],"quality":"Less is better","isMain":false,"metricName":" # relocations"},{"values":[0,0,0,0,0,0,0],"quality":"Less is better","isMain":false,"metricName":" # translocations"},{"values":[0,0,0,0,0,0,0],"quality":"Less is better","isMain":false,"metricName":" # inversions"},{"values":[0,0,0,0,0,0,1],"quality":"Less is better","isMain":false,"metricName":"# misassembled contigs"},{"values":[0,0,0,0,0,0,7670],"quality":"Less is better","isMain":true,"metricName":"Misassembled contigs length"},{"values":[0,0,1,0,0,0,0],"quality":"Less is better","isMain":false,"metricName":"# local misassemblies"},{"values":[0,0,0,0,0,0,0],"quality":"Less is better","isMain":false,"metricName":"# unaligned mis. contigs"}]],["Unaligned",[{"values":[0,0,0,0,0,0,0],"quality":"Less is better","isMain":false,"metricName":"# fully unaligned contigs"},{"values":[0,0,0,0,0,0,0],"quality":"Less is better","isMain":false,"metricName":"Fully unaligned length"},{"values":[0,0,0,0,0,0,0],"quality":"Less is better","isMain":false,"metricName":"# partially unaligned contigs"},{"values":[0,0,0,0,0,0,0],"quality":"Less is better","isMain":false,"metricName":"Partially unaligned length"}]],["Mismatches",[{"values":[0,12,0,0,0,0,2],"quality":"Less is better","isMain":false,"metricName":"# mismatches"},{"values":[0,0,0,1,1,1,0],"quality":"Less is better","isMain":false,"metricName":"# indels"},{"values":[0,0,0,55,55,55,0],"quality":"Less is better","isMain":false,"metricName":"Indels length"},{"values":["0.00","261.21","0.00","0.00","0.00","0.00","43.54"],"quality":"Less is better","isMain":true,"metricName":"# mismatches per 100 kbp"},{"values":["0.00","0.00","0.00","21.77","21.77","21.77","0.00"],"quality":"Less is better","isMain":true,"metricName":"# indels per 100 kbp"},{"values":[0,0,0,0,0,0,0],"quality":"Less is better","isMain":false,"metricName":" # indels (<= 5 bp)"},{"values":[0,0,0,1,1,1,0],"quality":"Less is better","isMain":false,"metricName":" # indels (> 5 bp)"},{"values":[0,37,0,0,0,0,0],"quality":"Less is better","isMain":false,"metricName":"# N's"},{"values":["0.00","3.31","0.00","0.00","0.00","0.00","0.00"],"quality":"Less is better","isMain":true,"metricName":"# N's per 100 kbp"}]],["Statistics without reference",[{"values":[1,1589,1,1,1,1,1],"quality":"Equal","isMain":true,"metricName":"# contigs"},{"values":[1,127,1,1,1,1,1],"quality":"Equal","isMain":false,"metricName":"# contigs (>= 1000 bp)"},{"values":[0,0,0,0,0,0,1],"quality":"Equal","isMain":false,"metricName":"# contigs (>= 5000 bp)"},{"values":[0,0,0,0,0,0,0],"quality":"Equal","isMain":false,"metricName":"# contigs (>= 10000 bp)"},{"values":[0,0,0,0,0,0,0],"quality":"Equal","isMain":false,"metricName":"# contigs (>= 25000 bp)"},{"values":[0,0,0,0,0,0,0],"quality":"Equal","isMain":false,"metricName":"# contigs (>= 50000 bp)"},{"values":[4594,1847,4721,4649,4649,4649,7670],"quality":"More is better","isMain":true,"metricName":"Largest contig"},{"values":[4594,1118327,4721,4649,4649,4649,7670],"quality":"More is better","isMain":true,"metricName":"Total length"},{"values":[4594,143797,4721,4649,4649,4649,7670],"quality":"More is better","isMain":true,"metricName":"Total length (>= 1000 bp)"},{"values":[0,0,0,0,0,0,7670],"quality":"More is better","isMain":false,"metricName":"Total length (>= 5000 bp)"},{"values":[0,0,0,0,0,0,0],"quality":"More is better","isMain":true,"metricName":"Total length (>= 10000 bp)"},{"values":[0,0,0,0,0,0,0],"quality":"More is better","isMain":false,"metricName":"Total length (>= 25000 bp)"},{"values":[0,0,0,0,0,0,0],"quality":"More is better","isMain":true,"metricName":"Total length (>= 50000 bp)"},{"values":[4594,721,4721,4649,4649,4649,7670],"quality":"More is better","isMain":false,"metricName":"N50"},{"values":[4594,577,4721,4649,4649,4649,7670],"quality":"More is better","isMain":false,"metricName":"N75"},{"values":[1,634,1,1,1,1,1],"quality":"Less is better","isMain":false,"metricName":"L50"},{"values":[1,1083,1,1,1,1,1],"quality":"Less is better","isMain":false,"metricName":"L75"},{"values":["46.10","46.07","45.88","46.14","46.14","46.14","46.48"],"quality":"Equal","isMain":false,"metricName":"GC (%)"}]],["Predicted genes",[]],["Similarity statistics",[{"values":[0,0,0,0,0,0,0],"quality":"Equal","isMain":false,"metricName":"# similar correct contigs"},{"values":[0,0,0,0,0,0,0],"quality":"Equal","isMain":false,"metricName":"# similar misassembled blocks"}]],["Reference statistics",[{"values":[4594,4594,4594,4594,4594,4594,4594],"quality":"Equal","isMain":false,"metricName":"Reference length"},{"values":[1,1,1,1,1,1,1],"quality":"Equal","isMain":false,"metricName":"Reference fragments"},{"values":["46.10","46.10","46.10","46.10","46.10","46.10","46.10"],"quality":"Equal","isMain":false,"metricName":"Reference GC (%)"}]]],"referenceName":"gi\_12085135\_ref\_NC","date":"18 July 2018, Wednesday, 16:12:49","order":[0,1,2,3,4,5,6],"assembliesNames":["CLC","MIRA","SPAdes","SPAdes\_meta","SPAdes\_sc","SPAdes\_sc\_careful","Geneious"]},{"assembliesWithNs":null,"minContig":500,"report":[["Genome statistics",[{"values":["100.000","63.240","2.513","100.000","100.000","100.000","100.000","100.000","100.000","2.499","100.000"],"quality":"More is better","isMain":true,"metricName":"Genome fraction (%)"},{"values":["1.000","1.007","1.000","325.750","1.002","1.006","1.003","1.003","1.003","1.000","2.981"],"quality":"Less is better","isMain":true,"metricName":"Duplication ratio"},{"values":[21129,1310,531,21602,21178,21256,21184,21184,21184,528,21129],"quality":"More is better","isMain":true,"metricName":"Largest alignment"},{"values":[21129,13461,531,6881197,21178,21256,21184,21184,21184,528,62994],"quality":"More is better","isMain":true,"metricName":"Total aligned length"},{"values":[21129,607,null,26186,21178,21256,21184,21184,21184,null,63664],"quality":"More is better","isMain":false,"metricName":"NG50"},{"values":[21129,null,null,26186,21178,21256,21184,21184,21184,null,63664],"quality":"More is better","isMain":false,"metricName":"NG75"},{"values":[21129,828,531,3759,21178,21256,21184,21184,21184,528,21129],"quality":"More is better","isMain":false,"metricName":"NA50"},{"values":[21129,690,531,2339,21178,21256,21184,21184,21184,528,15647],"quality":"More is better","isMain":false,"metricName":"NA75"},{"values":[21129,607,null,21602,21178,21256,21184,21184,21184,null,21129],"quality":"More is better","isMain":true,"metricName":"NGA50"},{"values":[21129,null,null,21602,21178,21256,21184,21184,21184,null,21129],"quality":"More is better","isMain":false,"metricName":"NGA75"},{"values":[1,12,null,1,1,1,1,1,1,null,1],"quality":"Less is better","isMain":false,"metricName":"LG50"},{"values":[1,null,null,1,1,1,1,1,1,null,1],"quality":"Less is better","isMain":false,"metricName":"LG75"},{"values":[1,7,1,508,1,1,1,1,1,1,2],"quality":"Less is better","isMain":false,"metricName":"LA50"},{"values":[1,11,1,1098,1,1,1,1,1,1,3],"quality":"Less is better","isMain":false,"metricName":"LA75"},{"values":[1,12,null,1,1,1,1,1,1,null,1],"quality":"Less is better","isMain":true,"metricName":"LGA50"},{"values":[1,null,null,1,1,1,1,1,1,null,1],"quality":"Less is better","isMain":false,"metricName":"LGA75"}]],["Misassemblies",[{"values":[0,0,0,8,0,0,0,0,0,0,3],"quality":"Less is better","isMain":true,"metricName":"# misassemblies"},{"values":[0,0,0,8,0,0,0,0,0,0,3],"quality":"Less is better","isMain":false,"metricName":" # relocations"},{"values":[0,0,0,0,0,0,0,0,0,0,0],"quality":"Less is better","isMain":false,"metricName":" # translocations"},{"values":[0,0,0,0,0,0,0,0,0,0,0],"quality":"Less is better","isMain":false,"metricName":" # inversions"},{"values":[0,0,0,8,0,0,0,0,0,0,1],"quality":"Less is better","isMain":false,"metricName":"# misassembled contigs"},{"values":[0,0,0,189753,0,0,0,0,0,0,63664],"quality":"Less is better","isMain":true,"metricName":"Misassembled contigs length"},{"values":[0,0,0,1,0,1,0,0,0,0,0],"quality":"Less is better","isMain":false,"metricName":"# local misassemblies"},{"values":[0,0,0,0,0,0,0,0,0,0,0],"quality":"Less is better","isMain":false,"metricName":"# unaligned mis. contigs"}]],["Unaligned",[{"values":[0,0,0,0,0,0,0,0,0,0,0],"quality":"Less is better","isMain":false,"metricName":"# fully unaligned contigs"},{"values":[0,0,0,0,0,0,0,0,0,0,0],"quality":"Less is better","isMain":false,"metricName":"Fully unaligned length"},{"values":[0,0,0,0,0,0,0,0,0,0,1],"quality":"Less is better","isMain":false,"metricName":"# partially unaligned contigs"},{"values":[0,0,0,0,0,0,0,0,0,0,670],"quality":"Less is better","isMain":false,"metricName":"Partially unaligned length"}]],["Mismatches",[{"values":[0,0,3,68,0,1,2,1,0,0,96],"quality":"Less is better","isMain":false,"metricName":"# mismatches"},{"values":[0,0,0,6,1,0,1,1,1,0,0],"quality":"Less is better","isMain":false,"metricName":"# indels"},{"values":[0,0,0,6,49,0,55,55,55,0,0],"quality":"Less is better","isMain":false,"metricName":"Indels length"},{"values":["0.00","0.00","564.97","321.83","0.00","4.73","9.47","4.73","0.00","0.00","454.35"],"quality":"Less is better","isMain":true,"metricName":"# mismatches per 100 kbp"},{"values":["0.00","0.00","0.00","28.40","4.73","0.00","4.73","4.73","4.73","0.00","0.00"],"quality":"Less is better","isMain":true,"metricName":"# indels per 100 kbp"},{"values":[0,0,0,6,0,0,0,0,0,0,0],"quality":"Less is better","isMain":false,"metricName":" # indels (<= 5 bp)"},{"values":[0,0,0,0,1,0,1,1,1,0,0],"quality":"Less is better","isMain":false,"metricName":" # indels (> 5 bp)"},{"values":[0,0,0,2570,0,0,0,0,1,0,20],"quality":"Less is better","isMain":false,"metricName":"# N's"},{"values":["0.00","0.00","0.00","37.34","0.00","0.00","0.00","0.00","4.72","0.00","31.41"],"quality":"Less is better","isMain":true,"metricName":"# N's per 100 kbp"}]],["Statistics without reference",[{"values":[1,17,1,2042,1,1,1,1,1,1,1],"quality":"Equal","isMain":true,"metricName":"# contigs"},{"values":[1,2,0,2020,1,1,1,1,1,0,1],"quality":"Equal","isMain":false,"metricName":"# contigs (>= 1000 bp)"},{"values":[1,0,0,298,1,1,1,1,1,0,1],"quality":"Equal","isMain":false,"metricName":"# contigs (>= 5000 bp)"},{"values":[1,0,0,72,1,1,1,1,1,0,1],"quality":"Equal","isMain":false,"metricName":"# contigs (>= 10000 bp)"},{"values":[0,0,0,2,0,0,0,0,0,0,1],"quality":"Equal","isMain":false,"metricName":"# contigs (>= 25000 bp)"},{"values":[0,0,0,0,0,0,0,0,0,0,1],"quality":"Equal","isMain":false,"metricName":"# contigs (>= 50000 bp)"},{"values":[21129,1310,531,26186,21178,21256,21184,21184,21184,528,63664],"quality":"More is better","isMain":true,"metricName":"Largest contig"},{"values":[21129,13461,531,6882766,21178,21256,21184,21184,21184,528,63664],"quality":"More is better","isMain":true,"metricName":"Total length"},{"values":[21129,2574,0,6864388,21178,21256,21184,21184,21184,0,63664],"quality":"More is better","isMain":true,"metricName":"Total length (>= 1000 bp)"},{"values":[21129,0,0,2580192,21178,21256,21184,21184,21184,0,63664],"quality":"More is better","isMain":false,"metricName":"Total length (>= 5000 bp)"},{"values":[21129,0,0,1040959,21178,21256,21184,21184,21184,0,63664],"quality":"More is better","isMain":true,"metricName":"Total length (>= 10000 bp)"},{"values":[0,0,0,51538,0,0,0,0,0,0,63664],"quality":"More is better","isMain":false,"metricName":"Total length (>= 25000 bp)"},{"values":[0,0,0,0,0,0,0,0,0,0,63664],"quality":"More is better","isMain":true,"metricName":"Total length (>= 50000 bp)"},{"values":[21129,828,531,3768,21178,21256,21184,21184,21184,528,63664],"quality":"More is better","isMain":false,"metricName":"N50"},{"values":[21129,690,531,2340,21178,21256,21184,21184,21184,528,63664],"quality":"More is better","isMain":false,"metricName":"N75"},{"values":[1,7,1,500,1,1,1,1,1,1,1],"quality":"Less is better","isMain":false,"metricName":"L50"},{"values":[1,11,1,1089,1,1,1,1,1,1,1],"quality":"Less is better","isMain":false,"metricName":"L75"},{"values":["34.66","34.13","34.84","34.78","34.68","34.69","34.63","34.63","34.63","35.04","34.75"],"quality":"Equal","isMain":false,"metricName":"GC (%)"}]],["Predicted genes",[]],["Similarity statistics",[{"values":[0,0,0,0,0,0,0,0,0,0,0],"quality":"Equal","isMain":false,"metricName":"# similar correct contigs"},{"values":[0,0,0,0,0,0,0,0,0,0,0],"quality":"Equal","isMain":false,"metricName":"# similar misassembled blocks"}]],["Reference statistics",[{"values":[21129,21129,21129,21129,21129,21129,21129,21129,21129,21129,21129],"quality":"Equal","isMain":false,"metricName":"Reference length"},{"values":[1,1,1,1,1,1,1,1,1,1,1],"quality":"Equal","isMain":false,"metricName":"Reference fragments"},{"values":["34.65","34.65","34.65","34.65","34.65","34.65","34.65","34.65","34.65","34.65","34.65"],"quality":"Equal","isMain":false,"metricName":"Reference GC (%)"}]]],"referenceName":"gi\_12248100\_ref\_NC","date":"18 July 2018, Wednesday, 16:13:40","order":[0,1,2,3,4,5,6,7,8,9,10],"assembliesNames":["CLC","IDBA\_UD","MEGAHIT","MIRA","Ray\_Meta","SPAdes","SPAdes\_meta","SPAdes\_sc","SPAdes\_sc\_careful","Velvet","Geneious"]},{"assembliesWithNs":null,"minContig":500,"report":[["Genome statistics",[{"values":["3.962","5.390","5.732","4.560","22.062","20.351","8.168","1.692","1.692","3.220","5.173","35.842"],"quality":"More is better","isMain":true,"metricName":"Genome fraction (%)"},{"values":["1.000","1.217","1.000","1.001","1.000","1.000","1.000","1.000","1.000","1.000","1.000","1.360"],"quality":"Less is better","isMain":true,"metricName":"Duplication ratio"},{"values":[606,749,787,749,2243,4874,777,732,732,732,1197,3720],"quality":"More is better","isMain":true,"metricName":"Largest alignment"},{"values":[1714,2837,2481,1974,9546,8805,3534,732,732,1393,2238,21089],"quality":"More is better","isMain":true,"metricName":"Total aligned length"},{"values":[null,null,null,null,null,null,null,null,null,null,null,36887],"quality":"More is better","isMain":false,"metricName":"NG50"},{"values":[null,null,null,null,null,null,null,null,null,null,null,36887],"quality":"More is better","isMain":false,"metricName":"NG75"},{"values":[571,507,648,632,1113,4874,749,732,732,732,1197,null],"quality":"More is better","isMain":false,"metricName":"NA50"},{"values":[537,506,531,593,850,2936,692,732,732,661,536,null],"quality":"More is better","isMain":false,"metricName":"NA75"},{"values":[null,null,null,null,null,null,null,null,null,null,null,null],"quality":"More is better","isMain":true,"metricName":"NGA50"},{"values":[null,null,null,null,null,null,null,null,null,null,null,1],"quality":"Less is better","isMain":false,"metricName":"LG50"},{"values":[null,null,null,null,null,null,null,null,null,null,null,1],"quality":"Less is better","isMain":false,"metricName":"LG75"},{"values":[2,3,2,2,3,1,3,1,1,1,1,null],"quality":"Less is better","isMain":false,"metricName":"LA50"},{"values":[3,4,3,3,6,2,4,1,1,2,2,null],"quality":"Less is better","isMain":false,"metricName":"LA75"}]],["Misassemblies",[{"values":[0,0,0,0,0,0,0,0,0,0,0,0],"quality":"Less is better","isMain":true,"metricName":"# misassemblies"},{"values":[0,0,0,0,0,0,0,0,0,0,0,0],"quality":"Less is better","isMain":false,"metricName":" # relocations"},{"values":[0,0,0,0,0,0,0,0,0,0,0,0],"quality":"Less is better","isMain":false,"metricName":" # translocations"},{"values":[0,0,0,0,0,0,0,0,0,0,0,0],"quality":"Less is better","isMain":false,"metricName":" # inversions"},{"values":[0,0,0,0,0,0,0,0,0,0,0,0],"quality":"Less is better","isMain":false,"metricName":"# misassembled contigs"},{"values":[0,0,0,0,0,0,0,0,0,0,0,0],"quality":"Less is better","isMain":true,"metricName":"Misassembled contigs length"},{"values":[0,0,0,0,0,0,0,0,0,0,0,0],"quality":"Less is better","isMain":false,"metricName":"# local misassemblies"},{"values":[0,0,0,0,0,0,0,0,0,0,0,3],"quality":"Less is better","isMain":false,"metricName":"# unaligned mis. contigs"}]],["Unaligned",[{"values":[0,0,0,0,0,0,0,0,0,0,0,0],"quality":"Less is better","isMain":false,"metricName":"# fully unaligned contigs"},{"values":[0,0,0,0,0,0,0,0,0,0,0,0],"quality":"Less is better","isMain":false,"metricName":"Fully unaligned length"},{"values":[0,0,0,0,1,0,0,0,0,0,0,3],"quality":"Less is better","isMain":false,"metricName":"# partially unaligned contigs"},{"values":[0,0,0,0,850,0,0,0,0,0,0,40441],"quality":"Less is better","isMain":false,"metricName":"Partially unaligned length"}]],["Mismatches",[{"values":[0,7,6,9,121,121,0,0,0,0,0,259],"quality":"Less is better","isMain":false,"metricName":"# mismatches"},{"values":[0,1,1,1,1,0,0,0,0,0,0,3],"quality":"Less is better","isMain":false,"metricName":"# indels"},{"values":[0,1,1,1,1,0,0,0,0,0,0,3],"quality":"Less is better","isMain":false,"metricName":"Indels length"},{"values":["0.00","300.17","241.94","456.16","1267.68","1374.22","0.00","0.00","0.00","0.00","0.00","1670.21"],"quality":"Less is better","isMain":true,"metricName":"# mismatches per 100 kbp"},{"values":["0.00","42.88","40.32","50.68","10.48","0.00","0.00","0.00","0.00","0.00","0.00","19.35"],"quality":"Less is better","isMain":true,"metricName":"# indels per 100 kbp"},{"values":[0,1,1,1,1,0,0,0,0,0,0,3],"quality":"Less is better","isMain":false,"metricName":" # indels (<= 5 bp)"},{"values":[0,0,0,0,0,0,0,0,0,0,0,0],"quality":"Less is better","isMain":false,"metricName":" # indels (> 5 bp)"},{"values":[0,0,0,0,3,0,0,0,0,0,0,280],"quality":"Less is better","isMain":false,"metricName":"# N's"},{"values":["0.00","0.00","0.00","0.00","28.86","0.00","0.00","0.00","0.00","0.00","0.00","455.06"],"quality":"Less is better","isMain":true,"metricName":"# N's per 100 kbp"}]],["Statistics without reference",[{"values":[3,5,4,3,8,3,5,1,1,2,3,3],"quality":"Equal","isMain":true,"metricName":"# contigs"},{"values":[0,0,0,0,6,2,0,0,0,0,1,3],"quality":"Equal","isMain":false,"metricName":"# contigs (>= 1000 bp)"},{"values":[0,0,0,0,0,0,0,0,0,0,0,2],"quality":"Equal","isMain":false,"metricName":"# contigs (>= 5000 bp)"},{"values":[0,0,0,0,0,0,0,0,0,0,0,2],"quality":"Equal","isMain":false,"metricName":"# contigs (>= 10000 bp)"},{"values":[0,0,0,0,0,0,0,0,0,0,0,1],"quality":"Equal","isMain":false,"metricName":"# contigs (>= 25000 bp)"},{"values":[0,0,0,0,0,0,0,0,0,0,0,0],"quality":"Equal","isMain":false,"metricName":"# contigs (>= 50000 bp)"},{"values":[606,749,787,749,2243,4874,777,732,732,732,1197,36887],"quality":"More is better","isMain":true,"metricName":"Largest contig"},{"values":[1714,2837,2481,1974,10396,8805,3534,732,732,1393,2238,61530],"quality":"More is better","isMain":true,"metricName":"Total length"},{"values":[0,0,0,0,8766,7810,0,0,0,0,1197,61530],"quality":"More is better","isMain":true,"metricName":"Total length (>= 1000 bp)"},{"values":[0,0,0,0,0,0,0,0,0,0,0,58257],"quality":"More is better","isMain":false,"metricName":"Total length (>= 5000 bp)"},{"values":[0,0,0,0,0,0,0,0,0,0,0,58257],"quality":"More is better","isMain":true,"metricName":"Total length (>= 10000 bp)"},{"values":[0,0,0,0,0,0,0,0,0,0,0,36887],"quality":"More is better","isMain":false,"metricName":"Total length (>= 25000 bp)"},{"values":[0,0,0,0,0,0,0,0,0,0,0,0],"quality":"More is better","isMain":true,"metricName":"Total length (>= 50000 bp)"},{"values":[571,507,648,632,1386,4874,749,732,732,732,1197,36887],"quality":"More is better","isMain":false,"metricName":"N50"},{"values":[537,506,531,593,1037,2936,692,732,732,661,536,21370],"quality":"More is better","isMain":false,"metricName":"N75"},{"values":[2,3,2,2,3,1,3,1,1,1,1,1],"quality":"Less is better","isMain":false,"metricName":"L50"},{"values":[3,4,3,3,6,2,4,1,1,2,2,2],"quality":"Less is better","isMain":false,"metricName":"L75"},{"values":["36.87","29.82","31.92","31.16","35.36","33.02","33.90","28.42","28.42","30.44","30.03","34.26"],"quality":"Equal","isMain":false,"metricName":"GC (%)"}]],["Predicted genes",[]],["Similarity statistics",[{"values":[0,0,0,0,0,0,0,0,0,0,0,0],"quality":"Equal","isMain":false,"metricName":"# similar correct contigs"},{"values":[0,0,0,0,0,0,0,0,0,0,0,0],"quality":"Equal","isMain":false,"metricName":"# similar misassembled blocks"}]],["Reference statistics",[{"values":[43265,43265,43265,43265,43265,43265,43265,43265,43265,43265,43265,43265],"quality":"Equal","isMain":false,"metricName":"Reference length"},{"values":[1,1,1,1,1,1,1,1,1,1,1,1],"quality":"Equal","isMain":false,"metricName":"Reference fragments"},{"values":["34.28","34.28","34.28","34.28","34.28","34.28","34.28","34.28","34.28","34.28","34.28","34.28"],"quality":"Equal","isMain":false,"metricName":"Reference GC (%)"}]]],"referenceName":"gi\_122891714\_ref\_NC","date":"18 July 2018, Wednesday, 16:13:50","order":[0,1,2,3,4,5,6,7,8,9,10,11],"assembliesNames":["ABySS\_127","CLC","IDBA\_UD","MEGAHIT","MIRA","Ray\_Meta","SPAdes","SPAdes\_meta","SPAdes\_sc","SPAdes\_sc\_careful","Velvet","Geneious"]},{"assembliesWithNs":null,"minContig":500,"report":[["Genome statistics",[{"values":["67.943","47.486","53.184","51.093","60.263","83.473","62.737","10.372","69.703","45.040","59.688","61.028","25.579","58.560"],"quality":"More is better","isMain":true,"metricName":"Genome fraction (%)"},{"values":["1.005","1.003","1.013","1.014","1.016","1.250","1.040","1.005","1.013","1.000","1.004","1.004","1.000","1.121"],"quality":"Less is better","isMain":true,"metricName":"Duplication ratio"},{"values":[11172,6492,6958,6740,6820,11765,4672,1275,10511,6569,9250,9250,2877,15800],"quality":"More is better","isMain":true,"metricName":"Largest alignment"},{"values":[29556,20615,23305,22414,26506,45146,28236,4510,30550,19494,25945,26524,11071,28403],"quality":"More is better","isMain":true,"metricName":"Total aligned length"},{"values":[3466,null,636,682,2058,47368,1582,null,2219,null,1921,1904,null,60961],"quality":"More is better","isMain":false,"metricName":"NG50"},{"values":[null,null,null,null,null,47368,null,null,null,null,null,null,null,60961],"quality":"More is better","isMain":false,"metricName":"NG75"},{"values":[4839,4066,2794,2481,2565,null,2653,674,3777,4885,4258,4203,2014,null],"quality":"More is better","isMain":false,"metricName":"NA50"},{"values":[3466,2234,1245,1218,1250,null,1582,609,2219,1921,1921,1904,1836,null],"quality":"More is better","isMain":false,"metricName":"NA75"},{"values":[3466,null,545,682,1250,4107,1582,null,2219,null,1921,1904,null,2885],"quality":"More is better","isMain":true,"metricName":"NGA50"},{"values":[null,null,null,null,null,2910,null,null,null,null,null,null,null,null],"quality":"More is better","isMain":false,"metricName":"NGA75"},{"values":[4,null,11,11,6,1,7,null,5,null,5,5,null,1],"quality":"Less is better","isMain":false,"metricName":"LG50"},{"values":[null,null,null,null,null,1,null,null,null,null,null,null,null,1],"quality":"Less is better","isMain":false,"metricName":"LG75"},{"values":[2,2,3,3,4,null,4,3,3,2,2,3,3,null],"quality":"Less is better","isMain":false,"metricName":"LA50"},{"values":[4,4,7,6,8,null,7,5,5,4,5,5,4,null],"quality":"Less is better","isMain":false,"metricName":"LA75"},{"values":[4,null,11,11,8,4,7,null,5,null,5,5,null,3],"quality":"Less is better","isMain":true,"metricName":"LGA50"},{"values":[null,null,null,null,null,7,null,null,null,null,null,null,null,null],"quality":"Less is better","isMain":false,"metricName":"LGA75"}]],["Misassemblies",[{"values":[0,0,0,0,0,0,0,0,0,0,0,0,0,0],"quality":"Less is better","isMain":true,"metricName":"# misassemblies"},{"values":[0,0,0,0,0,0,0,0,0,0,0,0,0,0],"quality":"Less is better","isMain":false,"metricName":" # relocations"},{"values":[0,0,0,0,0,0,0,0,0,0,0,0,0,0],"quality":"Less is better","isMain":false,"metricName":" # translocations"},{"values":[0,0,0,0,0,0,0,0,0,0,0,0,0,0],"quality":"Less is better","isMain":false,"metricName":" # inversions"},{"values":[0,0,0,0,0,0,0,0,0,0,0,0,0,0],"quality":"Less is better","isMain":false,"metricName":"# misassembled contigs"},{"values":[0,0,0,0,0,0,0,0,0,0,0,0,0,0],"quality":"Less is better","isMain":true,"metricName":"Misassembled contigs length"},{"values":[0,0,0,0,0,0,0,0,0,0,1,1,0,1],"quality":"Less is better","isMain":false,"metricName":"# local misassemblies"},{"values":[0,0,0,0,0,2,0,0,0,0,0,0,0,3],"quality":"Less is better","isMain":false,"metricName":"# unaligned mis. contigs"}]],["Unaligned",[{"values":[0,0,0,0,0,0,0,0,0,0,0,0,0,0],"quality":"Less is better","isMain":false,"metricName":"# fully unaligned contigs"},{"values":[0,0,0,0,0,0,0,0,0,0,0,0,0,0],"quality":"Less is better","isMain":false,"metricName":"Fully unaligned length"},{"values":[0,0,1,0,1,7,0,0,0,1,1,1,0,6],"quality":"Less is better","isMain":false,"metricName":"# partially unaligned contigs"},{"values":[0,0,606,0,3068,60522,0,0,0,1072,969,969,0,136217],"quality":"Less is better","isMain":false,"metricName":"Partially unaligned length"}]],["Mismatches",[{"values":[0,1,15,2,30,129,6,0,11,129,28,8,1,95],"quality":"Less is better","isMain":false,"metricName":"# mismatches"},{"values":[0,0,0,0,0,1,0,0,0,2,0,1,0,5],"quality":"Less is better","isMain":false,"metricName":"# indels"},{"values":[0,0,0,0,0,1,0,0,0,2,0,1,0,6],"quality":"Less is better","isMain":false,"metricName":"Indels length"},{"values":["0.00","4.87","65.16","9.04","115.02","357.05","22.10","0.00","36.46","661.74","108.38","30.29","9.03","374.81"],"quality":"Less is better","isMain":true,"metricName":"# mismatches per 100 kbp"},{"values":["0.00","0.00","0.00","0.00","0.00","2.77","0.00","0.00","0.00","10.26","0.00","3.79","0.00","19.73"],"quality":"Less is better","isMain":true,"metricName":"# indels per 100 kbp"},{"values":[0,0,0,0,0,1,0,0,0,2,0,1,0,5],"quality":"Less is better","isMain":false,"metricName":" # indels (<= 5 bp)"},{"values":[0,0,0,0,0,0,0,0,0,0,0,0,0,0],"quality":"Less is better","isMain":false,"metricName":" # indels (> 5 bp)"},{"values":[0,3,0,0,0,12,0,0,0,0,0,0,0,931],"quality":"Less is better","isMain":false,"metricName":"# N's"},{"values":["0.00","14.55","0.00","0.00","0.00","11.36","0.00","0.00","0.00","0.00","0.00","0.00","0.00","565.54"],"quality":"Less is better","isMain":true,"metricName":"# N's per 100 kbp"}]],["Statistics without reference",[{"values":[11,9,15,12,15,18,12,6,13,6,9,11,6,6],"quality":"Equal","isMain":true,"metricName":"# contigs"},{"values":[5,5,7,7,8,13,10,1,7,6,6,5,5,6],"quality":"Equal","isMain":false,"metricName":"# contigs (>= 1000 bp)"},{"values":[1,1,1,1,1,5,0,0,1,1,1,1,0,5],"quality":"Equal","isMain":false,"metricName":"# contigs (>= 5000 bp)"},{"values":[1,0,0,0,0,2,0,0,1,0,1,1,0,4],"quality":"Equal","isMain":false,"metricName":"# contigs (>= 10000 bp)"},{"values":[0,0,0,0,0,1,0,0,0,0,0,0,0,3],"quality":"Equal","isMain":false,"metricName":"# contigs (>= 25000 bp)"},{"values":[0,0,0,0,0,0,0,0,0,0,0,0,0,1],"quality":"Equal","isMain":false,"metricName":"# contigs (>= 50000 bp)"},{"values":[11172,6492,6958,6740,6820,47368,4672,1275,10511,6569,10219,10219,2877,60961],"quality":"More is better","isMain":true,"metricName":"Largest contig"},{"values":[29556,20615,23933,22414,29576,105668,28236,4510,30550,20569,26914,27493,11071,164620],"quality":"More is better","isMain":true,"metricName":"Total length"},{"values":[25751,17876,19145,18782,25310,102171,26469,1275,26817,20569,24854,23078,10431,164620],"quality":"More is better","isMain":true,"metricName":"Total length (>= 1000 bp)"},{"values":[11172,6492,6958,6740,6820,79319,0,0,10511,6569,10219,10219,0,161816],"quality":"More is better","isMain":false,"metricName":"Total length (>= 5000 bp)"},{"values":[11172,0,0,0,0,60062,0,0,10511,0,10219,10219,0,156102],"quality":"More is better","isMain":true,"metricName":"Total length (>= 10000 bp)"},{"values":[0,0,0,0,0,47368,0,0,0,0,0,0,0,131708],"quality":"More is better","isMain":false,"metricName":"Total length (>= 25000 bp)"},{"values":[0,0,0,0,0,0,0,0,0,0,0,0,0,60961],"quality":"More is better","isMain":true,"metricName":"Total length (>= 50000 bp)"},{"values":[4839,4066,2794,2481,3037,12694,2653,674,3777,4885,4258,4403,2014,42152],"quality":"More is better","isMain":false,"metricName":"N50"},{"values":[3466,2234,1245,1218,2058,5222,1582,609,2219,1926,2349,2349,1836,28595],"quality":"More is better","isMain":false,"metricName":"N75"},{"values":[2,2,3,3,4,2,4,3,3,2,2,2,3,2],"quality":"Less is better","isMain":false,"metricName":"L50"},{"values":[4,4,7,6,6,5,7,5,5,4,4,4,4,3],"quality":"Less is better","isMain":false,"metricName":"L75"},{"values":["34.59","34.23","35.06","34.86","34.88","35.40","35.32","34.61","34.79","34.98","34.95","35.42","33.68","34.97"],"quality":"Equal","isMain":false,"metricName":"GC (%)"}]],["Predicted genes",[]],["Similarity statistics",[{"values":[0,0,0,0,0,0,0,0,0,0,0,0,0,0],"quality":"Equal","isMain":false,"metricName":"# similar correct contigs"},{"values":[0,0,0,0,0,0,0,0,0,0,0,0,0,0],"quality":"Equal","isMain":false,"metricName":"# similar misassembled blocks"}]],["Reference statistics",[{"values":[43282,43282,43282,43282,43282,43282,43282,43282,43282,43282,43282,43282,43282,43282],"quality":"Equal","isMain":false,"metricName":"Reference length"},{"values":[1,1,1,1,1,1,1,1,1,1,1,1,1,1],"quality":"Equal","isMain":false,"metricName":"Reference fragments"},{"values":["34.90","34.90","34.90","34.90","34.90","34.90","34.90","34.90","34.90","34.90","34.90","34.90","34.90","34.90"],"quality":"Equal","isMain":false,"metricName":"Reference GC (%)"}]]],"referenceName":"gi\_122891784\_ref\_NC","date":"18 July 2018, Wednesday, 16:14:01","order":[0,1,2,3,4,5,6,7,8,9,10,11,12,13],"assembliesNames":["ABySS\_127","ABySS\_63","CLC","IDBA\_UD","MEGAHIT","MIRA","Ray\_Meta","SOAPdenovo2","SPAdes","SPAdes\_meta","SPAdes\_sc","SPAdes\_sc\_careful","Velvet","Geneious"]},{"assembliesWithNs":null,"minContig":500,"report":[["Genome statistics",[{"values":["87.022","72.734","87.851","84.724","85.467","100.000","8.490","22.943","90.120","86.676","75.653","80.698","29.731","100.000"],"quality":"More is better","isMain":true,"metricName":"Genome fraction (%)"},{"values":["1.010","1.015","1.001","1.028","1.023","0.993","1.039","1.454","1.007","1.000","1.008","1.010","1.003","1.006"],"quality":"Less is better","isMain":true,"metricName":"Duplication ratio"},{"values":[26287,7528,21461,15582,15398,70134,995,1222,29553,45080,7661,11729,3275,75680],"quality":"More is better","isMain":true,"metricName":"Largest alignment"},{"values":[121139,101753,121151,119992,120498,149016,12116,31681,125056,119426,105150,112357,41113,150898],"quality":"More is better","isMain":true,"metricName":"Total aligned length"},{"values":[8265,2117,10979,5746,6692,124838,null,749,9987,24870,2444,2392,null,125392],"quality":"More is better","isMain":false,"metricName":"NG50"},{"values":[3480,null,5765,2207,3537,124838,null,null,7865,9194,730,861,null,125392],"quality":"More is better","isMain":false,"metricName":"NG75"},{"values":[9354,3147,11660,7034,7118,35474,745,null,10520,24870,3863,3060,921,21924],"quality":"More is better","isMain":false,"metricName":"NA50"},{"values":[5984,2104,7760,3256,5501,null,594,null,9356,20431,2103,1907,676,null],"quality":"More is better","isMain":false,"metricName":"NA75"},{"values":[8265,2117,10979,5746,6692,70134,null,null,9987,24870,2444,2392,null,75680],"quality":"More is better","isMain":true,"metricName":"NGA50"},{"values":[3480,null,5765,2207,3537,35474,null,null,7865,9194,730,861,null,35475],"quality":"More is better","isMain":false,"metricName":"NGA75"},{"values":[6,18,5,8,8,1,null,49,5,2,16,15,null,1],"quality":"Less is better","isMain":false,"metricName":"LG50"},{"values":[11,null,9,17,14,1,null,null,9,5,39,37,null,1],"quality":"Less is better","isMain":false,"metricName":"LG75"},{"values":[5,11,4,6,7,2,7,null,4,2,10,10,15,3],"quality":"Less is better","isMain":false,"metricName":"LA50"},{"values":[9,21,7,12,11,null,12,null,8,3,20,22,27,null],"quality":"Less is better","isMain":false,"metricName":"LA75"},{"values":[6,18,5,8,8,1,null,null,5,2,16,15,null,1],"quality":"Less is better","isMain":true,"metricName":"LGA50"},{"values":[11,null,9,17,14,2,null,null,9,5,39,37,null,2],"quality":"Less is better","isMain":false,"metricName":"LGA75"}]],["Misassemblies",[{"values":[0,0,0,0,0,0,0,0,0,0,0,0,0,1],"quality":"Less is better","isMain":true,"metricName":"# misassemblies"},{"values":[0,0,0,0,0,0,0,0,0,0,0,0,0,1],"quality":"Less is better","isMain":false,"metricName":" # relocations"},{"values":[0,0,0,0,0,0,0,0,0,0,0,0,0,0],"quality":"Less is better","isMain":false,"metricName":" # translocations"},{"values":[0,0,0,0,0,0,0,0,0,0,0,0,0,0],"quality":"Less is better","isMain":false,"metricName":" # inversions"},{"values":[0,0,0,0,0,0,0,0,0,0,0,0,0,1],"quality":"Less is better","isMain":false,"metricName":"# misassembled contigs"},{"values":[0,0,0,0,0,0,0,0,0,0,0,0,0,99767],"quality":"Less is better","isMain":true,"metricName":"Misassembled contigs length"},{"values":[0,0,0,0,0,0,0,1,0,0,0,0,0,1],"quality":"Less is better","isMain":false,"metricName":"# local misassemblies"},{"values":[0,0,0,0,0,1,0,30,0,0,0,0,0,1],"quality":"Less is better","isMain":false,"metricName":"# unaligned mis. contigs"}]],["Unaligned",[{"values":[0,0,0,0,0,0,0,0,0,0,0,0,0,0],"quality":"Less is better","isMain":false,"metricName":"# fully unaligned contigs"},{"values":[0,0,0,0,0,0,0,0,0,0,0,0,0,0],"quality":"Less is better","isMain":false,"metricName":"Fully unaligned length"},{"values":[0,0,0,0,0,1,0,31,0,0,0,0,0,2],"quality":"Less is better","isMain":false,"metricName":"# partially unaligned contigs"},{"values":[0,0,0,0,0,64044,0,35331,0,0,0,0,0,86569],"quality":"Less is better","isMain":false,"metricName":"Partially unaligned length"}]],["Mismatches",[{"values":[134,27,15,31,49,128,11,12,22,231,74,41,15,150],"quality":"Less is better","isMain":false,"metricName":"# mismatches"},{"values":[13,8,6,5,3,23,6,34,6,12,3,9,0,22],"quality":"Less is better","isMain":false,"metricName":"# indels"},{"values":[28,9,18,5,16,43,17,1607,23,40,33,41,0,24],"quality":"Less is better","isMain":false,"metricName":"Indels length"},{"values":["111.73","26.93","12.39","26.55","41.60","92.87","94.01","37.95","17.71","193.37","70.97","36.86","36.61","108.84"],"quality":"Less is better","isMain":true,"metricName":"# mismatches per 100 kbp"},{"values":["10.84","7.98","4.96","4.28","2.55","16.69","51.28","107.53","4.83","10.05","2.88","8.09","0.00","15.96"],"quality":"Less is better","isMain":true,"metricName":"# indels per 100 kbp"},{"values":[12,8,5,5,2,21,5,1,5,11,1,7,0,22],"quality":"Less is better","isMain":false,"metricName":" # indels (<= 5 bp)"},{"values":[1,0,1,0,1,2,1,33,1,1,2,2,0,0],"quality":"Less is better","isMain":false,"metricName":" # indels (> 5 bp)"},{"values":[0,0,0,0,0,0,0,28270,0,0,0,0,0,5],"quality":"Less is better","isMain":false,"metricName":"# N's"},{"values":["0.00","0.00","0.00","0.00","0.00","0.00","0.00","34765.18","0.00","0.00","0.00","0.00","0.00","2.22"],"quality":"Less is better","isMain":true,"metricName":"# N's per 100 kbp"}]],["Statistics without reference",[{"values":[21,46,15,33,27,8,17,68,16,8,42,51,45,2],"quality":"Equal","isMain":true,"metricName":"# contigs"},{"values":[18,32,12,24,21,3,0,35,14,6,33,35,12,2],"quality":"Equal","isMain":false,"metricName":"# contigs (>= 1000 bp)"},{"values":[10,3,10,9,11,2,0,0,10,6,5,5,0,2],"quality":"Equal","isMain":false,"metricName":"# contigs (>= 5000 bp)"},{"values":[2,0,5,2,1,2,0,0,4,4,0,1,0,2],"quality":"Equal","isMain":false,"metricName":"# contigs (>= 10000 bp)"},{"values":[1,0,0,0,0,2,0,0,1,1,0,0,0,2],"quality":"Equal","isMain":false,"metricName":"# contigs (>= 25000 bp)"},{"values":[0,0,0,0,0,2,0,0,0,0,0,0,0,2],"quality":"Equal","isMain":false,"metricName":"# contigs (>= 50000 bp)"},{"values":[26287,7528,21461,15582,15398,124838,995,4004,29553,45080,7661,11729,3275,125392],"quality":"More is better","isMain":true,"metricName":"Largest contig"},{"values":[121139,101753,121151,119992,120498,200952,12161,81317,125056,119426,105150,112363,41113,225159],"quality":"More is better","isMain":true,"metricName":"Total length"},{"values":[119042,92090,119134,112977,115764,197443,0,56653,124027,118115,98599,101687,18621,225159],"quality":"More is better","isMain":true,"metricName":"Total length (>= 1000 bp)"},{"values":[100787,19586,112734,80198,91307,194972,0,0,115213,118115,31523,37532,0,225159],"quality":"More is better","isMain":false,"metricName":"Total length (>= 5000 bp)"},{"values":[37289,0,76959,27513,15398,194972,0,0,62793,100746,0,11729,0,225159],"quality":"More is better","isMain":true,"metricName":"Total length (>= 10000 bp)"},{"values":[26287,0,0,0,0,194972,0,0,29553,45080,0,0,0,225159],"quality":"More is better","isMain":false,"metricName":"Total length (>= 25000 bp)"},{"values":[0,0,0,0,0,194972,0,0,0,0,0,0,0,225159],"quality":"More is better","isMain":true,"metricName":"Total length (>= 50000 bp)"},{"values":[9354,3147,11660,7034,7118,124838,761,1371,10520,24870,3863,3064,921,125392],"quality":"More is better","isMain":false,"metricName":"N50"},{"values":[5984,2104,7760,3256,5501,70134,594,960,9356,20431,2103,1907,676,99767],"quality":"More is better","isMain":false,"metricName":"N75"},{"values":[5,11,4,6,7,1,7,21,4,2,10,10,15,1],"quality":"Less is better","isMain":false,"metricName":"L50"},{"values":[9,21,7,12,11,2,12,40,8,3,20,22,27,2],"quality":"Less is better","isMain":false,"metricName":"L75"},{"values":["74.75","74.95","74.86","74.72","74.82","74.08","72.86","75.22","74.77","74.63","75.06","74.93","74.12","73.83"],"quality":"Equal","isMain":false,"metricName":"GC (%)"}]],["Predicted genes",[]],["Similarity statistics",[{"values":[0,0,0,0,0,0,0,0,0,0,0,0,0,0],"quality":"Equal","isMain":false,"metricName":"# similar correct contigs"},{"values":[0,0,0,0,0,0,0,0,0,0,0,0,0,0],"quality":"Equal","isMain":false,"metricName":"# similar misassembled blocks"}]],["Reference statistics",[{"values":[137821,137821,137821,137821,137821,137821,137821,137821,137821,137821,137821,137821,137821,137821],"quality":"Equal","isMain":false,"metricName":"Reference length"},{"values":[1,1,1,1,1,1,1,1,1,1,1,1,1,1],"quality":"Equal","isMain":false,"metricName":"Reference fragments"},{"values":["74.80","74.80","74.80","74.80","74.80","74.80","74.80","74.80","74.80","74.80","74.80","74.80","74.80","74.80"],"quality":"Equal","isMain":false,"metricName":"Reference GC (%)"}]]],"referenceName":"gi\_123318702\_ref\_NC","date":"18 July 2018, Wednesday, 16:14:14","order":[0,1,2,3,4,5,6,7,8,9,10,11,12,13],"assembliesNames":["ABySS\_127","ABySS\_63","CLC","IDBA\_UD","MEGAHIT","MIRA","Ray\_Meta","SOAPdenovo2","SPAdes","SPAdes\_meta","SPAdes\_sc","SPAdes\_sc\_careful","Velvet","Geneious"]},{"assembliesWithNs":null,"minContig":500,"report":[["Genome statistics",[{"values":["42.600","39.751","91.682","67.503","75.746","100.000","47.582","68.172","35.833","60.240","47.709","75.337","94.799"],"quality":"More is better","isMain":true,"metricName":"Genome fraction (%)"},{"values":["1.100","1.040","1.012","1.025","1.038","7.243","1.000","1.021","1.000","1.005","1.003","1.002","1.500"],"quality":"Less is better","isMain":true,"metricName":"Duplication ratio"},{"values":[3589,5157,7740,6595,5936,11700,8500,7770,4987,7626,7626,5364,27026],"quality":"More is better","isMain":true,"metricName":"Largest alignment"},{"values":[16256,14219,32190,23992,27258,251207,16503,24152,12428,21003,16602,26174,49312],"quality":"More is better","isMain":true,"metricName":"Total aligned length"},{"values":[null,null,5936,1364,2929,34722,null,5071,null,4658,null,1421,49412],"quality":"More is better","isMain":false,"metricName":"NG50"},{"values":[null,null,3194,null,1086,34722,null,null,null,null,null,702,49412],"quality":"More is better","isMain":false,"metricName":"NG75"},{"values":[3295,5117,5936,5936,4973,1959,8500,5936,4658,5936,4658,3490,13230],"quality":"More is better","isMain":false,"metricName":"NA50"},{"values":[2395,3185,4861,1364,2769,1561,5638,5071,4658,4658,1535,1239,null],"quality":"More is better","isMain":false,"metricName":"NA75"},{"values":[null,null,5936,1364,2929,8265,null,5071,null,4658,null,1421,27026],"quality":"More is better","isMain":true,"metricName":"NGA50"},{"values":[null,null,3194,null,1086,7037,null,null,null,null,null,702,27026],"quality":"More is better","isMain":false,"metricName":"NGA75"},{"values":[null,null,3,4,4,1,null,3,null,3,null,5,1],"quality":"Less is better","isMain":false,"metricName":"LG50"},{"values":[null,null,5,null,8,1,null,null,null,null,null,14,1],"quality":"Less is better","isMain":false,"metricName":"LG75"},{"values":[3,2,3,2,3,42,1,2,2,2,2,3,2],"quality":"Less is better","isMain":false,"metricName":"LA50"},{"values":[4,3,4,4,5,81,2,3,2,3,3,7,null],"quality":"Less is better","isMain":false,"metricName":"LA75"},{"values":[null,null,3,4,4,2,null,3,null,3,null,5,1],"quality":"Less is better","isMain":true,"metricName":"LGA50"},{"values":[null,null,5,null,8,3,null,null,null,null,null,14,1],"quality":"Less is better","isMain":false,"metricName":"LGA75"}]],["Misassemblies",[{"values":[0,0,0,0,0,0,0,0,0,0,0,0,0],"quality":"Less is better","isMain":true,"metricName":"# misassemblies"},{"values":[0,0,0,0,0,0,0,0,0,0,0,0,0],"quality":"Less is better","isMain":false,"metricName":" # relocations"},{"values":[0,0,0,0,0,0,0,0,0,0,0,0,0],"quality":"Less is better","isMain":false,"metricName":" # translocations"},{"values":[0,0,0,0,0,0,0,0,0,0,0,0,0],"quality":"Less is better","isMain":false,"metricName":" # inversions"},{"values":[0,0,0,0,0,0,0,0,0,0,0,0,0],"quality":"Less is better","isMain":false,"metricName":"# misassembled contigs"},{"values":[0,0,0,0,0,0,0,0,0,0,0,0,0],"quality":"Less is better","isMain":true,"metricName":"Misassembled contigs length"},{"values":[0,0,0,0,1,0,0,0,0,0,0,0,5],"quality":"Less is better","isMain":false,"metricName":"# local misassemblies"},{"values":[0,0,0,0,0,1,0,0,0,0,0,0,0],"quality":"Less is better","isMain":false,"metricName":"# unaligned mis. contigs"}]],["Unaligned",[{"values":[0,0,0,0,0,0,0,0,0,0,0,0,0],"quality":"Less is better","isMain":false,"metricName":"# fully unaligned contigs"},{"values":[0,0,0,0,0,0,0,0,0,0,0,0,0],"quality":"Less is better","isMain":false,"metricName":"Fully unaligned length"},{"values":[0,1,0,0,0,1,0,0,0,0,0,0,2],"quality":"Less is better","isMain":false,"metricName":"# partially unaligned contigs"},{"values":[0,1537,0,0,0,17385,0,0,0,0,0,0,23277],"quality":"Less is better","isMain":false,"metricName":"Partially unaligned length"}]],["Mismatches",[{"values":[0,0,3,0,5,112,0,0,1,3,0,14,59],"quality":"Less is better","isMain":false,"metricName":"# mismatches"},{"values":[0,0,0,0,0,1,0,0,0,0,0,1,1],"quality":"Less is better","isMain":false,"metricName":"# indels"},{"values":[0,0,0,0,0,1,0,0,0,0,0,1,1],"quality":"Less is better","isMain":false,"metricName":"Indels length"},{"values":["0.00","0.00","9.43","0.00","19.03","322.92","0.00","0.00","8.05","14.36","0.00","53.58","179.45"],"quality":"Less is better","isMain":true,"metricName":"# mismatches per 100 kbp"},{"values":["0.00","0.00","0.00","0.00","0.00","2.88","0.00","0.00","0.00","0.00","0.00","3.83","3.04"],"quality":"Less is better","isMain":true,"metricName":"# indels per 100 kbp"},{"values":[0,0,0,0,0,1,0,0,0,0,0,1,1],"quality":"Less is better","isMain":false,"metricName":" # indels (<= 5 bp)"},{"values":[0,0,0,0,0,0,0,0,0,0,0,0,0],"quality":"Less is better","isMain":false,"metricName":" # indels (> 5 bp)"},{"values":[0,437,0,0,0,223,0,0,0,0,0,0,145],"quality":"Less is better","isMain":false,"metricName":"# N's"},{"values":["0.00","2751.89","0.00","0.00","0.00","83.02","0.00","0.00","0.00","0.00","0.00","0.00","199.75"],"quality":"Less is better","isMain":true,"metricName":"# N's per 100 kbp"}]],["Statistics without reference",[{"values":[8,5,9,11,9,121,4,7,5,6,6,14,3],"quality":"Equal","isMain":true,"metricName":"# contigs"},{"values":[5,4,6,6,8,114,3,6,4,5,5,9,3],"quality":"Equal","isMain":false,"metricName":"# contigs (>= 1000 bp)"},{"values":[0,2,3,2,2,3,2,3,0,2,1,1,3],"quality":"Equal","isMain":false,"metricName":"# contigs (>= 5000 bp)"},{"values":[0,0,0,0,0,2,0,0,0,0,0,0,2],"quality":"Equal","isMain":false,"metricName":"# contigs (>= 10000 bp)"},{"values":[0,0,0,0,0,1,0,0,0,0,0,0,1],"quality":"Equal","isMain":false,"metricName":"# contigs (>= 25000 bp)"},{"values":[0,0,0,0,0,0,0,0,0,0,0,0,0],"quality":"Equal","isMain":false,"metricName":"# contigs (>= 50000 bp)"},{"values":[3589,5157,7740,6595,5936,34722,8500,7770,4987,7626,7626,5364,49412],"quality":"More is better","isMain":true,"metricName":"Largest contig"},{"values":[16256,15880,32190,23992,27258,268604,16503,24152,12428,21003,16602,26174,72589],"quality":"More is better","isMain":true,"metricName":"Total length"},{"values":[14237,15283,30589,21052,26503,263622,15562,23425,11845,20420,16019,22084,72589],"quality":"More is better","isMain":true,"metricName":"Total length (>= 1000 bp)"},{"values":[0,10274,20374,12531,11106,54687,14138,18777,0,13562,7626,5364,72589],"quality":"More is better","isMain":false,"metricName":"Total length (>= 5000 bp)"},{"values":[0,0,0,0,0,46422,0,0,0,0,0,0,62642],"quality":"More is better","isMain":true,"metricName":"Total length (>= 10000 bp)"},{"values":[0,0,0,0,0,34722,0,0,0,0,0,0,49412],"quality":"More is better","isMain":false,"metricName":"Total length (>= 25000 bp)"},{"values":[0,0,0,0,0,0,0,0,0,0,0,0,0],"quality":"More is better","isMain":true,"metricName":"Total length (>= 50000 bp)"},{"values":[3295,5117,5936,5936,4973,2085,8500,5936,4658,5936,4658,3490,49412],"quality":"More is better","isMain":false,"metricName":"N50"},{"values":[2395,3309,4861,1364,2769,1659,5638,5071,4658,4658,1535,1239,13230],"quality":"More is better","isMain":false,"metricName":"N75"},{"values":[3,2,3,2,3,32,1,2,2,2,2,3,1],"quality":"Less is better","isMain":false,"metricName":"L50"},{"values":[4,3,4,4,5,68,2,3,2,3,3,7,2],"quality":"Less is better","isMain":false,"metricName":"L75"},{"values":["44.97","45.35","44.11","45.23","45.03","42.79","45.76","44.68","44.86","44.87","45.39","43.70","44.64"],"quality":"Equal","isMain":false,"metricName":"GC (%)"}]],["Predicted genes",[]],["Similarity statistics",[{"values":[0,0,0,0,0,0,0,0,0,0,0,0,0],"quality":"Equal","isMain":false,"metricName":"# similar correct contigs"},{"values":[0,0,0,0,0,0,0,0,0,0,0,0,0],"quality":"Equal","isMain":false,"metricName":"# similar misassembled blocks"}]],["Reference statistics",[{"values":[34683,34683,34683,34683,34683,34683,34683,34683,34683,34683,34683,34683,34683],"quality":"Equal","isMain":false,"metricName":"Reference length"},{"values":[1,1,1,1,1,1,1,1,1,1,1,1,1],"quality":"Equal","isMain":false,"metricName":"Reference fragments"},{"values":["44.38","44.38","44.38","44.38","44.38","44.38","44.38","44.38","44.38","44.38","44.38","44.38","44.38"],"quality":"Equal","isMain":false,"metricName":"Reference GC (%)"}]]],"referenceName":"gi\_124300942\_ref\_NC","date":"18 July 2018, Wednesday, 16:14:26","order":[0,1,2,3,4,5,6,7,8,9,10,11,12],"assembliesNames":["ABySS\_127","ABySS\_63","CLC","IDBA\_UD","MEGAHIT","MIRA","Ray\_Meta","SPAdes","SPAdes\_meta","SPAdes\_sc","SPAdes\_sc\_careful","Velvet","Geneious"]},{"assembliesWithNs":null,"minContig":500,"report":[["Genome statistics",[{"values":["95.129","99.936","100.000","95.925","100.000","31.614","85.196","100.000","99.809","92.869","92.869","87.806","100.000"],"quality":"More is better","isMain":true,"metricName":"Genome fraction (%)"},{"values":["1.000","1.001","1.027","1.000","1.139","1.000","1.016","1.040","1.001","1.000","1.000","1.011","1.124"],"quality":"Less is better","isMain":true,"metricName":"Duplication ratio"},{"values":[2988,3141,3227,3013,3141,993,2719,3268,3137,2917,2917,2758,3141],"quality":"More is better","isMain":true,"metricName":"Largest alignment"},{"values":[2988,3141,3227,3013,3531,993,2719,3268,3137,2917,2917,2758,3530],"quality":"More is better","isMain":true,"metricName":"Total aligned length"},{"values":[2988,3141,3227,3013,3577,null,3267,3268,3137,2917,2917,2789,5297],"quality":"More is better","isMain":false,"metricName":"NG50"},{"values":[2988,3141,3227,3013,3577,null,3267,3268,3137,2917,2917,2789,5297],"quality":"More is better","isMain":false,"metricName":"NG75"},{"values":[2988,3141,3227,3013,3141,993,2719,3268,3137,2917,2917,2758,3141],"quality":"More is better","isMain":false,"metricName":"NA50"},{"values":[2988,3141,3227,3013,3141,993,2719,3268,3137,2917,2917,2758,null],"quality":"More is better","isMain":false,"metricName":"NA75"},{"values":[2988,3141,3227,3013,3141,null,2719,3268,3137,2917,2917,2758,3141],"quality":"More is better","isMain":true,"metricName":"NGA50"},{"values":[2988,3141,3227,3013,3141,null,2719,3268,3137,2917,2917,2758,3141],"quality":"More is better","isMain":false,"metricName":"NGA75"},{"values":[1,1,1,1,1,null,1,1,1,1,1,1,1],"quality":"Less is better","isMain":false,"metricName":"LG50"},{"values":[1,1,1,1,1,null,1,1,1,1,1,1,1],"quality":"Less is better","isMain":false,"metricName":"LG75"},{"values":[1,1,1,1,1,1,1,1,1,1,1,1,1],"quality":"Less is better","isMain":false,"metricName":"LA50"},{"values":[1,1,1,1,1,1,1,1,1,1,1,1,null],"quality":"Less is better","isMain":false,"metricName":"LA75"},{"values":[1,1,1,1,1,null,1,1,1,1,1,1,1],"quality":"Less is better","isMain":true,"metricName":"LGA50"},{"values":[1,1,1,1,1,null,1,1,1,1,1,1,1],"quality":"Less is better","isMain":false,"metricName":"LGA75"}]],["Misassemblies",[{"values":[0,0,0,0,1,0,0,0,0,0,0,0,1],"quality":"Less is better","isMain":true,"metricName":"# misassemblies"},{"values":[0,0,0,0,1,0,0,0,0,0,0,0,1],"quality":"Less is better","isMain":false,"metricName":" # relocations"},{"values":[0,0,0,0,0,0,0,0,0,0,0,0,0],"quality":"Less is better","isMain":false,"metricName":" # translocations"},{"values":[0,0,0,0,0,0,0,0,0,0,0,0,0],"quality":"Less is better","isMain":false,"metricName":" # inversions"},{"values":[0,0,0,0,1,0,0,0,0,0,0,0,1],"quality":"Less is better","isMain":false,"metricName":"# misassembled contigs"},{"values":[0,0,0,0,3577,0,0,0,0,0,0,0,5297],"quality":"Less is better","isMain":true,"metricName":"Misassembled contigs length"},{"values":[0,0,1,0,0,0,1,1,0,0,0,0,0],"quality":"Less is better","isMain":false,"metricName":"# local misassemblies"},{"values":[0,0,0,0,0,0,0,0,0,0,0,0,0],"quality":"Less is better","isMain":false,"metricName":"# unaligned mis. contigs"}]],["Unaligned",[{"values":[0,0,0,0,0,0,0,0,0,0,0,0,0],"quality":"Less is better","isMain":false,"metricName":"# fully unaligned contigs"},{"values":[0,0,0,0,0,0,0,0,0,0,0,0,0],"quality":"Less is better","isMain":false,"metricName":"Fully unaligned length"},{"values":[0,0,0,0,0,0,1,0,0,0,0,0,1],"quality":"Less is better","isMain":false,"metricName":"# partially unaligned contigs"},{"values":[0,0,0,0,0,0,548,0,0,0,0,0,1767],"quality":"Less is better","isMain":false,"metricName":"Partially unaligned length"}]],["Mismatches",[{"values":[0,0,0,1,0,0,0,0,0,0,0,0,0],"quality":"Less is better","isMain":false,"metricName":"# mismatches"},{"values":[0,1,1,0,0,0,1,0,1,0,0,0,0],"quality":"Less is better","isMain":false,"metricName":"# indels"},{"values":[0,2,12,0,0,0,43,0,2,0,0,0,0],"quality":"Less is better","isMain":false,"metricName":"Indels length"},{"values":["0.00","0.00","0.00","33.19","0.00","0.00","0.00","0.00","0.00","0.00","0.00","0.00","0.00"],"quality":"Less is better","isMain":true,"metricName":"# mismatches per 100 kbp"},{"values":["0.00","31.86","31.84","0.00","0.00","0.00","37.37","0.00","31.90","0.00","0.00","0.00","0.00"],"quality":"Less is better","isMain":true,"metricName":"# indels per 100 kbp"},{"values":[0,1,0,0,0,0,0,0,1,0,0,0,0],"quality":"Less is better","isMain":false,"metricName":" # indels (<= 5 bp)"},{"values":[0,0,1,0,0,0,1,0,0,0,0,0,0],"quality":"Less is better","isMain":false,"metricName":" # indels (> 5 bp)"},{"values":[0,0,0,0,0,0,553,0,0,0,0,0,2],"quality":"Less is better","isMain":false,"metricName":"# N's"},{"values":["0.00","0.00","0.00","0.00","0.00","0.00","16926.84","0.00","0.00","0.00","0.00","0.00","37.76"],"quality":"Less is better","isMain":true,"metricName":"# N's per 100 kbp"}]],["Statistics without reference",[{"values":[1,1,1,1,1,1,1,1,1,1,1,1,1],"quality":"Equal","isMain":true,"metricName":"# contigs"},{"values":[1,1,1,1,1,0,1,1,1,1,1,1,1],"quality":"Equal","isMain":false,"metricName":"# contigs (>= 1000 bp)"},{"values":[0,0,0,0,0,0,0,0,0,0,0,0,1],"quality":"Equal","isMain":false,"metricName":"# contigs (>= 5000 bp)"},{"values":[0,0,0,0,0,0,0,0,0,0,0,0,0],"quality":"Equal","isMain":false,"metricName":"# contigs (>= 10000 bp)"},{"values":[0,0,0,0,0,0,0,0,0,0,0,0,0],"quality":"Equal","isMain":false,"metricName":"# contigs (>= 25000 bp)"},{"values":[0,0,0,0,0,0,0,0,0,0,0,0,0],"quality":"Equal","isMain":false,"metricName":"# contigs (>= 50000 bp)"},{"values":[2988,3141,3227,3013,3577,993,3267,3268,3137,2917,2917,2789,5297],"quality":"More is better","isMain":true,"metricName":"Largest contig"},{"values":[2988,3141,3227,3013,3577,993,3267,3268,3137,2917,2917,2789,5297],"quality":"More is better","isMain":true,"metricName":"Total length"},{"values":[2988,3141,3227,3013,3577,0,3267,3268,3137,2917,2917,2789,5297],"quality":"More is better","isMain":true,"metricName":"Total length (>= 1000 bp)"},{"values":[0,0,0,0,0,0,0,0,0,0,0,0,5297],"quality":"More is better","isMain":false,"metricName":"Total length (>= 5000 bp)"},{"values":[0,0,0,0,0,0,0,0,0,0,0,0,0],"quality":"More is better","isMain":true,"metricName":"Total length (>= 10000 bp)"},{"values":[0,0,0,0,0,0,0,0,0,0,0,0,0],"quality":"More is better","isMain":false,"metricName":"Total length (>= 25000 bp)"},{"values":[0,0,0,0,0,0,0,0,0,0,0,0,0],"quality":"More is better","isMain":true,"metricName":"Total length (>= 50000 bp)"},{"values":[2988,3141,3227,3013,3577,993,3267,3268,3137,2917,2917,2789,5297],"quality":"More is better","isMain":false,"metricName":"N50"},{"values":[2988,3141,3227,3013,3577,993,3267,3268,3137,2917,2917,2789,5297],"quality":"More is better","isMain":false,"metricName":"N75"},{"values":[1,1,1,1,1,1,1,1,1,1,1,1,1],"quality":"Less is better","isMain":false,"metricName":"L50"},{"values":[1,1,1,1,1,1,1,1,1,1,1,1,1],"quality":"Less is better","isMain":false,"metricName":"L75"},{"values":["34.94","35.53","35.92","34.95","37.18","33.13","33.46","35.40","35.51","34.52","34.52","33.99","42.66"],"quality":"Equal","isMain":false,"metricName":"GC (%)"}]],["Predicted genes",[]],["Similarity statistics",[{"values":[0,0,0,0,0,0,0,0,0,0,0,0,0],"quality":"Equal","isMain":false,"metricName":"# similar correct contigs"},{"values":[0,0,0,0,0,0,0,0,0,0,0,0,0],"quality":"Equal","isMain":false,"metricName":"# similar misassembled blocks"}]],["Reference statistics",[{"values":[3141,3141,3141,3141,3141,3141,3141,3141,3141,3141,3141,3141,3141],"quality":"Equal","isMain":false,"metricName":"Reference length"},{"values":[1,1,1,1,1,1,1,1,1,1,1,1,1],"quality":"Equal","isMain":false,"metricName":"Reference fragments"},{"values":["35.53","35.53","35.53","35.53","35.53","35.53","35.53","35.53","35.53","35.53","35.53","35.53","35.53"],"quality":"Equal","isMain":false,"metricName":"Reference GC (%)"}]]],"referenceName":"gi\_124378202\_ref\_NC","date":"18 July 2018, Wednesday, 16:14:36","order":[0,1,2,3,4,5,6,7,8,9,10,11,12],"assembliesNames":["ABySS\_63","CLC","IDBA\_UD","MEGAHIT","MIRA","Ray\_Meta","SOAPdenovo2","SPAdes","SPAdes\_meta","SPAdes\_sc","SPAdes\_sc\_careful","Velvet","Geneious"]},{"assembliesWithNs":null,"minContig":500,"report":[["Genome statistics",[{"values":["8.232","4.874","9.304","0.419","7.092","10.230","10.218","9.919","9.919"],"quality":"More is better","isMain":true,"metricName":"Genome fraction (%)"},{"values":["1.000","1.000","1.000","1.000","1.000","1.000","1.000","1.000","1.000"],"quality":"Less is better","isMain":true,"metricName":"Duplication ratio"},{"values":[1557,1479,1479,745,1054,1637,1637,1637,1637],"quality":"More is better","isMain":true,"metricName":"Largest alignment"},{"values":[14642,8673,16550,745,12615,18197,18175,17644,17644],"quality":"More is better","isMain":true,"metricName":"Total aligned length"},{"values":[769,810,771,745,747,884,884,979,979],"quality":"More is better","isMain":false,"metricName":"NA50"},{"values":[621,602,621,745,613,632,632,676,676],"quality":"More is better","isMain":false,"metricName":"NA75"},{"values":[null,null,null,null,null,null,null,null,null],"quality":"More is better","isMain":true,"metricName":"NGA50"},{"values":[8,4,8,1,8,8,8,7,7],"quality":"Less is better","isMain":false,"metricName":"LA50"},{"values":[13,8,14,1,12,14,14,13,13],"quality":"Less is better","isMain":false,"metricName":"LA75"}]],["Misassemblies",[{"values":[0,0,0,0,0,0,0,0,0],"quality":"Less is better","isMain":true,"metricName":"# misassemblies"},{"values":[0,0,0,0,0,0,0,0,0],"quality":"Less is better","isMain":false,"metricName":" # relocations"},{"values":[0,0,0,0,0,0,0,0,0],"quality":"Less is better","isMain":false,"metricName":" # translocations"},{"values":[0,0,0,0,0,0,0,0,0],"quality":"Less is better","isMain":false,"metricName":" # inversions"},{"values":[0,0,0,0,0,0,0,0,0],"quality":"Less is better","isMain":false,"metricName":"# misassembled contigs"},{"values":[0,0,0,0,0,0,0,0,0],"quality":"Less is better","isMain":true,"metricName":"Misassembled contigs length"},{"values":[0,1,2,0,1,1,1,1,1],"quality":"Less is better","isMain":false,"metricName":"# local misassemblies"},{"values":[0,0,0,0,0,0,0,0,0],"quality":"Less is better","isMain":false,"metricName":"# unaligned mis. contigs"}]],["Unaligned",[{"values":[0,0,0,0,0,0,0,0,0],"quality":"Less is better","isMain":false,"metricName":"# fully unaligned contigs"},{"values":[0,0,0,0,0,0,0,0,0],"quality":"Less is better","isMain":false,"metricName":"Fully unaligned length"},{"values":[0,0,0,0,0,0,0,0,0],"quality":"Less is better","isMain":false,"metricName":"# partially unaligned contigs"},{"values":[0,0,0,0,0,0,0,0,0],"quality":"Less is better","isMain":false,"metricName":"Partially unaligned length"}]],["Mismatches",[{"values":[24,10,34,0,18,36,34,32,32],"quality":"Less is better","isMain":false,"metricName":"# mismatches"},{"values":[0,0,1,0,0,0,0,0,0],"quality":"Less is better","isMain":false,"metricName":"# indels"},{"values":[0,0,69,0,0,0,0,0,0],"quality":"Less is better","isMain":false,"metricName":"Indels length"},{"values":["163.91","115.34","205.44","0.00","142.69","197.83","187.07","181.36","181.36"],"quality":"Less is better","isMain":true,"metricName":"# mismatches per 100 kbp"},{"values":["0.00","0.00","6.04","0.00","0.00","0.00","0.00","0.00","0.00"],"quality":"Less is better","isMain":true,"metricName":"# indels per 100 kbp"},{"values":[0,0,0,0,0,0,0,0,0],"quality":"Less is better","isMain":false,"metricName":" # indels (<= 5 bp)"},{"values":[0,0,1,0,0,0,0,0,0],"quality":"Less is better","isMain":false,"metricName":" # indels (> 5 bp)"},{"values":[0,0,0,3,0,0,0,0,0],"quality":"Less is better","isMain":false,"metricName":"# N's"},{"values":["0.00","0.00","0.00","402.68","0.00","0.00","0.00","0.00","0.00"],"quality":"Less is better","isMain":true,"metricName":"# N's per 100 kbp"}]],["Statistics without reference",[{"values":[19,11,21,1,17,21,21,20,20],"quality":"Equal","isMain":true,"metricName":"# contigs"},{"values":[3,3,3,0,2,6,6,6,6],"quality":"Equal","isMain":false,"metricName":"# contigs (>= 1000 bp)"},{"values":[0,0,0,0,0,0,0,0,0],"quality":"Equal","isMain":false,"metricName":"# contigs (>= 5000 bp)"},{"values":[0,0,0,0,0,0,0,0,0],"quality":"Equal","isMain":false,"metricName":"# contigs (>= 10000 bp)"},{"values":[0,0,0,0,0,0,0,0,0],"quality":"Equal","isMain":false,"metricName":"# contigs (>= 25000 bp)"},{"values":[0,0,0,0,0,0,0,0,0],"quality":"Equal","isMain":false,"metricName":"# contigs (>= 50000 bp)"},{"values":[1557,1479,1479,745,1054,1637,1637,1637,1637],"quality":"More is better","isMain":true,"metricName":"Largest contig"},{"values":[14642,8673,16550,745,12615,18197,18175,17644,17644],"quality":"More is better","isMain":true,"metricName":"Total length"},{"values":[3670,3627,4084,0,2080,7908,7908,7908,7908],"quality":"More is better","isMain":true,"metricName":"Total length (>= 1000 bp)"},{"values":[0,0,0,0,0,0,0,0,0],"quality":"More is better","isMain":false,"metricName":"Total length (>= 5000 bp)"},{"values":[0,0,0,0,0,0,0,0,0],"quality":"More is better","isMain":true,"metricName":"Total length (>= 10000 bp)"},{"values":[0,0,0,0,0,0,0,0,0],"quality":"More is better","isMain":false,"metricName":"Total length (>= 25000 bp)"},{"values":[0,0,0,0,0,0,0,0,0],"quality":"More is better","isMain":true,"metricName":"Total length (>= 50000 bp)"},{"values":[769,810,771,745,747,884,884,979,979],"quality":"More is better","isMain":false,"metricName":"N50"},{"values":[621,602,621,745,613,632,632,676,676],"quality":"More is better","isMain":false,"metricName":"N75"},{"values":[8,4,8,1,8,8,8,7,7],"quality":"Less is better","isMain":false,"metricName":"L50"},{"values":[13,8,14,1,12,14,14,13,13],"quality":"Less is better","isMain":false,"metricName":"L75"},{"values":["44.14","46.72","44.92","53.10","45.10","45.09","45.09","45.28","45.28"],"quality":"Equal","isMain":false,"metricName":"GC (%)"}]],["Predicted genes",[]],["Similarity statistics",[{"values":[0,0,0,0,0,0,0,0,0],"quality":"Equal","isMain":false,"metricName":"# similar correct contigs"},{"values":[0,0,0,0,0,0,0,0,0],"quality":"Equal","isMain":false,"metricName":"# similar misassembled blocks"}]],["Reference statistics",[{"values":[177874,177874,177874,177874,177874,177874,177874,177874,177874],"quality":"Equal","isMain":false,"metricName":"Reference length"},{"values":[1,1,1,1,1,1,1,1,1],"quality":"Equal","isMain":false,"metricName":"Reference fragments"},{"values":["44.06","44.06","44.06","44.06","44.06","44.06","44.06","44.06","44.06"],"quality":"Equal","isMain":false,"metricName":"Reference GC (%)"}]]],"referenceName":"gi\_125745044\_ref\_NC","date":"18 July 2018, Wednesday, 16:14:45","order":[0,1,2,3,4,5,6,7,8],"assembliesNames":["CLC","IDBA\_UD","MEGAHIT","MIRA","SOAPdenovo2","SPAdes","SPAdes\_meta","SPAdes\_sc","SPAdes\_sc\_careful"]},{"assembliesWithNs":null,"minContig":500,"report":[["Genome statistics",[{"values":["100.000","100.000","100.000","100.000","100.000","27.204","100.000","100.000","100.000","100.000","100.000"],"quality":"More is better","isMain":true,"metricName":"Genome fraction (%)"},{"values":["1.000","1.003","1.004","1.008","1.003","1.000","1.003","1.001","1.001","1.001","1.001"],"quality":"Less is better","isMain":true,"metricName":"Duplication ratio"},{"values":[38197,38296,38338,38488,38300,1285,38324,38252,38252,38252,38227],"quality":"More is better","isMain":true,"metricName":"Largest alignment"},{"values":[38197,38296,38338,38488,38300,10391,38324,38252,38252,38252,38227],"quality":"More is better","isMain":true,"metricName":"Total aligned length"},{"values":[38197,38296,38338,38488,38300,null,38324,38252,38252,38252,38227],"quality":"More is better","isMain":false,"metricName":"NG50"},{"values":[38197,38296,38338,38488,38300,null,38324,38252,38252,38252,38227],"quality":"More is better","isMain":false,"metricName":"NG75"},{"values":[38197,38296,38338,38488,38300,758,38324,38252,38252,38252,38227],"quality":"More is better","isMain":false,"metricName":"NA50"},{"values":[38197,38296,38338,38488,38300,687,38324,38252,38252,38252,38227],"quality":"More is better","isMain":false,"metricName":"NA75"},{"values":[38197,38296,38338,38488,38300,null,38324,38252,38252,38252,38227],"quality":"More is better","isMain":true,"metricName":"NGA50"},{"values":[38197,38296,38338,38488,38300,null,38324,38252,38252,38252,38227],"quality":"More is better","isMain":false,"metricName":"NGA75"},{"values":[1,1,1,1,1,null,1,1,1,1,1],"quality":"Less is better","isMain":false,"metricName":"LG50"},{"values":[1,1,1,1,1,null,1,1,1,1,1],"quality":"Less is better","isMain":false,"metricName":"LG75"},{"values":[1,1,1,1,1,5,1,1,1,1,1],"quality":"Less is better","isMain":false,"metricName":"LA50"},{"values":[1,1,1,1,1,9,1,1,1,1,1],"quality":"Less is better","isMain":false,"metricName":"LA75"},{"values":[1,1,1,1,1,null,1,1,1,1,1],"quality":"Less is better","isMain":true,"metricName":"LGA50"},{"values":[1,1,1,1,1,null,1,1,1,1,1],"quality":"Less is better","isMain":false,"metricName":"LGA75"}]],["Misassemblies",[{"values":[0,0,0,0,0,0,0,0,0,0,0],"quality":"Less is better","isMain":true,"metricName":"# misassemblies"},{"values":[0,0,0,0,0,0,0,0,0,0,0],"quality":"Less is better","isMain":false,"metricName":" # relocations"},{"values":[0,0,0,0,0,0,0,0,0,0,0],"quality":"Less is better","isMain":false,"metricName":" # translocations"},{"values":[0,0,0,0,0,0,0,0,0,0,0],"quality":"Less is better","isMain":false,"metricName":" # inversions"},{"values":[0,0,0,0,0,0,0,0,0,0,0],"quality":"Less is better","isMain":false,"metricName":"# misassembled contigs"},{"values":[0,0,0,0,0,0,0,0,0,0,0],"quality":"Less is better","isMain":true,"metricName":"Misassembled contigs length"},{"values":[0,1,1,1,1,0,1,0,0,0,0],"quality":"Less is better","isMain":false,"metricName":"# local misassemblies"},{"values":[0,0,0,0,0,0,0,0,0,0,0],"quality":"Less is better","isMain":false,"metricName":"# unaligned mis. contigs"}]],["Unaligned",[{"values":[0,0,0,0,0,0,0,0,0,0,0],"quality":"Less is better","isMain":false,"metricName":"# fully unaligned contigs"},{"values":[0,0,0,0,0,0,0,0,0,0,0],"quality":"Less is better","isMain":false,"metricName":"Fully unaligned length"},{"values":[0,0,0,0,0,0,0,0,0,0,0],"quality":"Less is better","isMain":false,"metricName":"# partially unaligned contigs"},{"values":[0,0,0,0,0,0,0,0,0,0,0],"quality":"Less is better","isMain":false,"metricName":"Partially unaligned length"}]],["Mismatches",[{"values":[0,0,0,0,0,2,0,0,0,0,0],"quality":"Less is better","isMain":false,"metricName":"# mismatches"},{"values":[0,0,0,0,0,0,0,1,1,1,1],"quality":"Less is better","isMain":false,"metricName":"# indels"},{"values":[0,0,0,0,0,0,0,55,55,55,30],"quality":"Less is better","isMain":false,"metricName":"Indels length"},{"values":["0.00","0.00","0.00","0.00","0.00","19.25","0.00","0.00","0.00","0.00","0.00"],"quality":"Less is better","isMain":true,"metricName":"# mismatches per 100 kbp"},{"values":["0.00","0.00","0.00","0.00","0.00","0.00","0.00","2.62","2.62","2.62","2.62"],"quality":"Less is better","isMain":true,"metricName":"# indels per 100 kbp"},{"values":[0,0,0,0,0,0,0,0,0,0,0],"quality":"Less is better","isMain":false,"metricName":" # indels (<= 5 bp)"},{"values":[0,0,0,0,0,0,0,1,1,1,1],"quality":"Less is better","isMain":false,"metricName":" # indels (> 5 bp)"},{"values":[0,0,0,0,0,0,0,0,0,0,0],"quality":"Less is better","isMain":false,"metricName":"# N's"},{"values":["0.00","0.00","0.00","0.00","0.00","0.00","0.00","0.00","0.00","0.00","0.00"],"quality":"Less is better","isMain":true,"metricName":"# N's per 100 kbp"}]],["Statistics without reference",[{"values":[1,1,1,1,1,13,1,1,1,1,1],"quality":"Equal","isMain":true,"metricName":"# contigs"},{"values":[1,1,1,1,1,3,1,1,1,1,1],"quality":"Equal","isMain":false,"metricName":"# contigs (>= 1000 bp)"},{"values":[1,1,1,1,1,0,1,1,1,1,1],"quality":"Equal","isMain":false,"metricName":"# contigs (>= 5000 bp)"},{"values":[1,1,1,1,1,0,1,1,1,1,1],"quality":"Equal","isMain":false,"metricName":"# contigs (>= 10000 bp)"},{"values":[1,1,1,1,1,0,1,1,1,1,1],"quality":"Equal","isMain":false,"metricName":"# contigs (>= 25000 bp)"},{"values":[0,0,0,0,0,0,0,0,0,0,0],"quality":"Equal","isMain":false,"metricName":"# contigs (>= 50000 bp)"},{"values":[38197,38296,38338,38488,38300,1285,38324,38252,38252,38252,38227],"quality":"More is better","isMain":true,"metricName":"Largest contig"},{"values":[38197,38296,38338,38488,38300,10391,38324,38252,38252,38252,38227],"quality":"More is better","isMain":true,"metricName":"Total length"},{"values":[38197,38296,38338,38488,38300,3748,38324,38252,38252,38252,38227],"quality":"More is better","isMain":true,"metricName":"Total length (>= 1000 bp)"},{"values":[38197,38296,38338,38488,38300,0,38324,38252,38252,38252,38227],"quality":"More is better","isMain":false,"metricName":"Total length (>= 5000 bp)"},{"values":[38197,38296,38338,38488,38300,0,38324,38252,38252,38252,38227],"quality":"More is better","isMain":true,"metricName":"Total length (>= 10000 bp)"},{"values":[38197,38296,38338,38488,38300,0,38324,38252,38252,38252,38227],"quality":"More is better","isMain":false,"metricName":"Total length (>= 25000 bp)"},{"values":[0,0,0,0,0,0,0,0,0,0,0],"quality":"More is better","isMain":true,"metricName":"Total length (>= 50000 bp)"},{"values":[38197,38296,38338,38488,38300,758,38324,38252,38252,38252,38227],"quality":"More is better","isMain":false,"metricName":"N50"},{"values":[38197,38296,38338,38488,38300,687,38324,38252,38252,38252,38227],"quality":"More is better","isMain":false,"metricName":"N75"},{"values":[1,1,1,1,1,5,1,1,1,1,1],"quality":"Less is better","isMain":false,"metricName":"L50"},{"values":[1,1,1,1,1,9,1,1,1,1,1],"quality":"Less is better","isMain":false,"metricName":"L75"},{"values":["56.95","56.96","56.98","56.93","56.93","56.73","56.96","56.96","56.96","56.96","56.96"],"quality":"Equal","isMain":false,"metricName":"GC (%)"}]],["Predicted genes",[]],["Similarity statistics",[{"values":[0,0,0,0,0,0,0,0,0,0,0],"quality":"Equal","isMain":false,"metricName":"# similar correct contigs"},{"values":[0,0,0,0,0,0,0,0,0,0,0],"quality":"Equal","isMain":false,"metricName":"# similar misassembled blocks"}]],["Reference statistics",[{"values":[38197,38197,38197,38197,38197,38197,38197,38197,38197,38197,38197],"quality":"Equal","isMain":false,"metricName":"Reference length"},{"values":[1,1,1,1,1,1,1,1,1,1,1],"quality":"Equal","isMain":false,"metricName":"Reference fragments"},{"values":["56.95","56.95","56.95","56.95","56.95","56.95","56.95","56.95","56.95","56.95","56.95"],"quality":"Equal","isMain":false,"metricName":"Reference GC (%)"}]]],"referenceName":"gi\_126010846\_ref\_NC","date":"18 July 2018, Wednesday, 16:14:54","order":[0,1,2,3,4,5,6,7,8,9,10],"assembliesNames":["CLC","IDBA\_UD","MEGAHIT","MIRA","Ray\_Meta","SOAPdenovo2","SPAdes","SPAdes\_meta","SPAdes\_sc","SPAdes\_sc\_careful","Velvet"]},{"assembliesWithNs":null,"minContig":500,"report":[["Genome statistics",[{"values":["99.558","98.988","100.000","100.000","100.000","99.874","3.273","98.678","100.000","100.000","100.000","100.000","100.000","99.874"],"quality":"More is better","isMain":true,"metricName":"Genome fraction (%)"},{"values":["1.001","1.000","1.004","1.001","1.002","1.009","1.000","1.036","1.002","1.001","1.001","1.001","1.000","1.000"],"quality":"Less is better","isMain":true,"metricName":"Duplication ratio"},{"values":[48417,20659,72713,72514,72556,72324,658,34234,72542,72470,72470,72470,72441,72324],"quality":"More is better","isMain":true,"metricName":"Largest alignment"},{"values":[72144,71688,72713,72514,72556,72962,2370,72447,72542,72470,72470,72470,72441,72324],"quality":"More is better","isMain":true,"metricName":"Total aligned length"},{"values":[48417,12580,72715,72514,72556,72324,null,28326,72542,72470,72470,72470,72441,72324],"quality":"More is better","isMain":false,"metricName":"NG50"},{"values":[13496,9790,72715,72514,72556,72324,null,28326,72542,72470,72470,72470,72441,72324],"quality":"More is better","isMain":false,"metricName":"NG75"},{"values":[48417,12580,72713,72514,72556,72324,594,27428,72542,72470,72470,72470,72441,72324],"quality":"More is better","isMain":false,"metricName":"NA50"},{"values":[13496,9790,72713,72514,72556,72324,589,27428,72542,72470,72470,72470,72441,72324],"quality":"More is better","isMain":false,"metricName":"NA75"},{"values":[48417,12580,72713,72514,72556,72324,null,27428,72542,72470,72470,72470,72441,72324],"quality":"More is better","isMain":true,"metricName":"NGA50"},{"values":[13496,9790,72713,72514,72556,72324,null,27428,72542,72470,72470,72470,72441,72324],"quality":"More is better","isMain":false,"metricName":"NGA75"},{"values":[1,3,1,1,1,1,null,2,1,1,1,1,1,1],"quality":"Less is better","isMain":false,"metricName":"LG50"},{"values":[2,4,1,1,1,1,null,2,1,1,1,1,1,1],"quality":"Less is better","isMain":false,"metricName":"LG75"},{"values":[1,3,1,1,1,1,2,2,1,1,1,1,1,1],"quality":"Less is better","isMain":false,"metricName":"LA50"},{"values":[2,4,1,1,1,1,3,2,1,1,1,1,1,1],"quality":"Less is better","isMain":false,"metricName":"LA75"},{"values":[1,3,1,1,1,1,null,2,1,1,1,1,1,1],"quality":"Less is better","isMain":true,"metricName":"LGA50"},{"values":[2,4,1,1,1,1,null,2,1,1,1,1,1,1],"quality":"Less is better","isMain":false,"metricName":"LGA75"}]],["Misassemblies",[{"values":[0,0,0,0,0,0,0,0,0,0,0,0,0,0],"quality":"Less is better","isMain":true,"metricName":"# misassemblies"},{"values":[0,0,0,0,0,0,0,0,0,0,0,0,0,0],"quality":"Less is better","isMain":false,"metricName":" # relocations"},{"values":[0,0,0,0,0,0,0,0,0,0,0,0,0,0],"quality":"Less is better","isMain":false,"metricName":" # translocations"},{"values":[0,0,0,0,0,0,0,0,0,0,0,0,0,0],"quality":"Less is better","isMain":false,"metricName":" # inversions"},{"values":[0,0,0,0,0,0,0,0,0,0,0,0,0,0],"quality":"Less is better","isMain":false,"metricName":"# misassembled contigs"},{"values":[0,0,0,0,0,0,0,0,0,0,0,0,0,0],"quality":"Less is better","isMain":true,"metricName":"Misassembled contigs length"},{"values":[0,0,1,1,1,0,0,0,1,0,0,0,0,0],"quality":"Less is better","isMain":false,"metricName":"# local misassemblies"},{"values":[0,0,0,0,0,0,0,0,0,0,0,0,0,0],"quality":"Less is better","isMain":false,"metricName":"# unaligned mis. contigs"}]],["Unaligned",[{"values":[0,0,0,0,0,0,0,0,0,0,0,0,0,0],"quality":"Less is better","isMain":false,"metricName":"# fully unaligned contigs"},{"values":[0,0,0,0,0,0,0,0,0,0,0,0,0,0],"quality":"Less is better","isMain":false,"metricName":"Fully unaligned length"},{"values":[0,0,0,0,0,0,0,0,0,0,0,0,0,0],"quality":"Less is better","isMain":false,"metricName":"# partially unaligned contigs"},{"values":[0,0,0,0,0,0,0,0,0,0,0,0,0,0],"quality":"Less is better","isMain":false,"metricName":"Partially unaligned length"}]],["Mismatches",[{"values":[4,0,0,0,0,0,0,0,0,0,0,0,0,0],"quality":"Less is better","isMain":false,"metricName":"# mismatches"},{"values":[0,0,0,0,0,0,0,35,0,1,1,1,1,0],"quality":"Less is better","isMain":false,"metricName":"# indels"},{"values":[0,0,0,0,0,0,0,1720,0,55,55,55,26,0],"quality":"Less is better","isMain":false,"metricName":"Indels length"},{"values":["5.55","0.00","0.00","0.00","0.00","0.00","0.00","0.00","0.00","0.00","0.00","0.00","0.00","0.00"],"quality":"Less is better","isMain":true,"metricName":"# mismatches per 100 kbp"},{"values":["0.00","0.00","0.00","0.00","0.00","0.00","0.00","48.98","0.00","1.38","1.38","1.38","1.38","0.00"],"quality":"Less is better","isMain":true,"metricName":"# indels per 100 kbp"},{"values":[0,0,0,0,0,0,0,0,0,0,0,0,0,0],"quality":"Less is better","isMain":false,"metricName":" # indels (<= 5 bp)"},{"values":[0,0,0,0,0,0,0,35,0,1,1,1,1,0],"quality":"Less is better","isMain":false,"metricName":" # indels (> 5 bp)"},{"values":[0,0,0,0,0,2,0,2806,0,0,0,0,0,2],"quality":"Less is better","isMain":false,"metricName":"# N's"},{"values":["0.00","0.00","0.00","0.00","0.00","2.74","0.00","3789.84","0.00","0.00","0.00","0.00","0.00","2.77"],"quality":"Less is better","isMain":true,"metricName":"# N's per 100 kbp"}]],["Statistics without reference",[{"values":[4,7,1,1,1,2,4,5,1,1,1,1,1,1],"quality":"Equal","isMain":true,"metricName":"# contigs"},{"values":[4,7,1,1,1,1,0,5,1,1,1,1,1,1],"quality":"Equal","isMain":false,"metricName":"# contigs (>= 1000 bp)"},{"values":[3,6,1,1,1,1,0,3,1,1,1,1,1,1],"quality":"Equal","isMain":false,"metricName":"# contigs (>= 5000 bp)"},{"values":[2,3,1,1,1,1,0,2,1,1,1,1,1,1],"quality":"Equal","isMain":false,"metricName":"# contigs (>= 10000 bp)"},{"values":[1,0,1,1,1,1,0,2,1,1,1,1,1,1],"quality":"Equal","isMain":false,"metricName":"# contigs (>= 25000 bp)"},{"values":[0,0,1,1,1,1,0,0,1,1,1,1,1,1],"quality":"Equal","isMain":false,"metricName":"# contigs (>= 50000 bp)"},{"values":[48417,20659,72715,72514,72556,72324,658,34709,72542,72470,72470,72470,72441,72324],"quality":"More is better","isMain":true,"metricName":"Largest contig"},{"values":[72144,71688,72715,72514,72556,72962,2370,74040,72542,72470,72470,72470,72441,72324],"quality":"More is better","isMain":true,"metricName":"Total length"},{"values":[72144,71688,72715,72514,72556,72324,0,74040,72542,72470,72470,72470,72441,72324],"quality":"More is better","isMain":true,"metricName":"Total length (>= 1000 bp)"},{"values":[70937,70547,72715,72514,72556,72324,0,69013,72542,72470,72470,72470,72441,72324],"quality":"More is better","isMain":false,"metricName":"Total length (>= 5000 bp)"},{"values":[61913,48360,72715,72514,72556,72324,0,63035,72542,72470,72470,72470,72441,72324],"quality":"More is better","isMain":true,"metricName":"Total length (>= 10000 bp)"},{"values":[48417,0,72715,72514,72556,72324,0,63035,72542,72470,72470,72470,72441,72324],"quality":"More is better","isMain":false,"metricName":"Total length (>= 25000 bp)"},{"values":[0,0,72715,72514,72556,72324,0,0,72542,72470,72470,72470,72441,72324],"quality":"More is better","isMain":true,"metricName":"Total length (>= 50000 bp)"},{"values":[48417,12580,72715,72514,72556,72324,594,28326,72542,72470,72470,72470,72441,72324],"quality":"More is better","isMain":false,"metricName":"N50"},{"values":[13496,9790,72715,72514,72556,72324,589,28326,72542,72470,72470,72470,72441,72324],"quality":"More is better","isMain":false,"metricName":"N75"},{"values":[1,3,1,1,1,1,2,2,1,1,1,1,1,1],"quality":"Less is better","isMain":false,"metricName":"L50"},{"values":[2,4,1,1,1,1,3,2,1,1,1,1,1,1],"quality":"Less is better","isMain":false,"metricName":"L75"},{"values":["55.89","55.87","55.91","55.89","55.89","55.92","58.06","55.89","55.90","55.89","55.89","55.89","55.90","55.89"],"quality":"Equal","isMain":false,"metricName":"GC (%)"}]],["Predicted genes",[]],["Similarity statistics",[{"values":[0,0,0,0,0,0,0,0,0,0,0,0,0,0],"quality":"Equal","isMain":false,"metricName":"# similar correct contigs"},{"values":[0,0,0,0,0,0,0,0,0,0,0,0,0,0],"quality":"Equal","isMain":false,"metricName":"# similar misassembled blocks"}]],["Reference statistics",[{"values":[72415,72415,72415,72415,72415,72415,72415,72415,72415,72415,72415,72415,72415,72415],"quality":"Equal","isMain":false,"metricName":"Reference length"},{"values":[1,1,1,1,1,1,1,1,1,1,1,1,1,1],"quality":"Equal","isMain":false,"metricName":"Reference fragments"},{"values":["55.89","55.89","55.89","55.89","55.89","55.89","55.89","55.89","55.89","55.89","55.89","55.89","55.89","55.89"],"quality":"Equal","isMain":false,"metricName":"Reference GC (%)"}]]],"referenceName":"gi\_126010934\_ref\_NC","date":"18 July 2018, Wednesday, 16:15:06","order":[0,1,2,3,4,5,6,7,8,9,10,11,12,13],"assembliesNames":["ABySS\_127","ABySS\_63","CLC","IDBA\_UD","MEGAHIT","MIRA","Ray\_Meta","SOAPdenovo2","SPAdes","SPAdes\_meta","SPAdes\_sc","SPAdes\_sc\_careful","Velvet","Geneious"]},{"assembliesWithNs":null,"minContig":500,"report":[["Genome statistics",[{"values":["100.000","75.156","91.087","78.766","86.854","99.073","77.310","13.979","95.323","60.639","72.420","79.524","48.902","100.000"],"quality":"More is better","isMain":true,"metricName":"Genome fraction (%)"},{"values":["1.094","1.099","1.009","1.013","1.011","1.117","1.074","1.000","1.007","1.000","1.000","1.000","1.004","1.174"],"quality":"Less is better","isMain":true,"metricName":"Duplication ratio"},{"values":[18858,7859,11859,13659,15048,29462,4525,1047,21533,12666,14442,14442,2356,29462],"quality":"More is better","isMain":true,"metricName":"Largest alignment"},{"values":[42136,31833,35411,30747,33844,42631,31992,5386,36980,23361,27902,30639,18925,45224],"quality":"More is better","isMain":true,"metricName":"Total aligned length"},{"values":[11730,5364,8301,5690,8539,31842,1895,null,21533,3165,7838,7838,null,40600],"quality":"More is better","isMain":false,"metricName":"NG50"},{"values":[11730,2136,5010,638,5613,31842,1228,null,8511,null,null,1408,null,40600],"quality":"More is better","isMain":false,"metricName":"NG75"},{"values":[11730,5364,8301,5690,8539,29462,2066,882,21533,12666,14442,7838,1720,12819],"quality":"More is better","isMain":false,"metricName":"NA50"},{"values":[11548,4840,5010,2434,5613,10580,1457,661,8511,5300,7838,3679,1374,null],"quality":"More is better","isMain":false,"metricName":"NA75"},{"values":[11730,5364,8301,5690,8539,29462,1895,null,21533,3165,7838,7838,null,29462],"quality":"More is better","isMain":true,"metricName":"NGA50"},{"values":[11730,2136,5010,638,5613,29462,1228,null,8511,null,null,1408,null,29462],"quality":"More is better","isMain":false,"metricName":"NGA75"},{"values":[2,3,2,2,2,1,8,null,1,3,2,2,null,1],"quality":"Less is better","isMain":false,"metricName":"LG50"},{"values":[2,6,4,8,3,1,14,null,2,null,null,5,null,1],"quality":"Less is better","isMain":false,"metricName":"LG75"},{"values":[2,3,2,2,2,1,6,3,1,1,1,2,5,2],"quality":"Less is better","isMain":false,"metricName":"LA50"},{"values":[3,4,4,4,3,2,11,5,2,2,2,3,8,null],"quality":"Less is better","isMain":false,"metricName":"LA75"},{"values":[2,3,2,2,2,1,8,null,1,3,2,2,null,1],"quality":"Less is better","isMain":true,"metricName":"LGA50"},{"values":[2,6,4,8,3,1,14,null,2,null,null,5,null,1],"quality":"Less is better","isMain":false,"metricName":"LGA75"}]],["Misassemblies",[{"values":[0,0,0,0,0,0,0,0,0,0,0,0,0,1],"quality":"Less is better","isMain":true,"metricName":"# misassemblies"},{"values":[0,0,0,0,0,0,0,0,0,0,0,0,0,1],"quality":"Less is better","isMain":false,"metricName":" # relocations"},{"values":[0,0,0,0,0,0,0,0,0,0,0,0,0,0],"quality":"Less is better","isMain":false,"metricName":" # translocations"},{"values":[0,0,0,0,0,0,0,0,0,0,0,0,0,0],"quality":"Less is better","isMain":false,"metricName":" # inversions"},{"values":[0,0,0,0,0,0,0,0,0,0,0,0,0,1],"quality":"Less is better","isMain":false,"metricName":"# misassembled contigs"},{"values":[0,0,0,0,0,0,0,0,0,0,0,0,0,40600],"quality":"Less is better","isMain":true,"metricName":"Misassembled contigs length"},{"values":[0,0,0,0,0,0,0,0,0,0,0,0,0,0],"quality":"Less is better","isMain":false,"metricName":"# local misassemblies"},{"values":[0,0,0,0,0,0,0,0,0,0,0,0,0,1],"quality":"Less is better","isMain":false,"metricName":"# unaligned mis. contigs"}]],["Unaligned",[{"values":[0,0,0,0,0,0,0,0,0,0,0,0,0,0],"quality":"Less is better","isMain":false,"metricName":"# fully unaligned contigs"},{"values":[0,0,0,0,0,0,0,0,0,0,0,0,0,0],"quality":"Less is better","isMain":false,"metricName":"Fully unaligned length"},{"values":[0,0,0,0,0,1,0,0,0,0,0,0,0,2],"quality":"Less is better","isMain":false,"metricName":"# partially unaligned contigs"},{"values":[0,0,0,0,0,2380,0,0,0,0,0,0,0,33212],"quality":"Less is better","isMain":false,"metricName":"Partially unaligned length"}]],["Mismatches",[{"values":[217,23,2,0,5,5,0,0,1,15,33,6,0,42],"quality":"Less is better","isMain":false,"metricName":"# mismatches"},{"values":[6,0,0,0,0,0,0,0,0,3,2,0,0,2],"quality":"Less is better","isMain":false,"metricName":"# indels"},{"values":[13,0,0,0,0,0,0,0,0,4,2,0,0,3],"quality":"Less is better","isMain":false,"metricName":"Indels length"},{"values":["563.23","79.43","5.70","0.00","14.94","13.10","0.00","0.00","2.72","64.20","118.27","19.58","0.00","109.01"],"quality":"Less is better","isMain":true,"metricName":"# mismatches per 100 kbp"},{"values":["15.57","0.00","0.00","0.00","0.00","0.00","0.00","0.00","0.00","12.84","7.17","0.00","0.00","5.19"],"quality":"Less is better","isMain":true,"metricName":"# indels per 100 kbp"},{"values":[6,0,0,0,0,0,0,0,0,3,2,0,0,2],"quality":"Less is better","isMain":false,"metricName":" # indels (<= 5 bp)"},{"values":[0,0,0,0,0,0,0,0,0,0,0,0,0,0],"quality":"Less is better","isMain":false,"metricName":" # indels (> 5 bp)"},{"values":[0,2,0,0,0,1,0,0,0,46,0,0,0,111],"quality":"Less is better","isMain":false,"metricName":"# N's"},{"values":["0.00","6.28","0.00","0.00","0.00","2.22","0.00","0.00","0.00","196.91","0.00","0.00","0.00","141.52"],"quality":"Less is better","isMain":true,"metricName":"# N's per 100 kbp"}]],["Statistics without reference",[{"values":[3,8,6,11,5,5,18,7,4,4,4,6,13,2],"quality":"Equal","isMain":true,"metricName":"# contigs"},{"values":[3,6,6,6,5,3,15,1,4,4,4,6,10,2],"quality":"Equal","isMain":false,"metricName":"# contigs (>= 1000 bp)"},{"values":[3,3,4,2,3,2,0,0,3,2,2,2,0,2],"quality":"Equal","isMain":false,"metricName":"# contigs (>= 5000 bp)"},{"values":[3,0,1,1,1,2,0,0,1,1,1,1,0,2],"quality":"Equal","isMain":false,"metricName":"# contigs (>= 10000 bp)"},{"values":[0,0,0,0,0,1,0,0,0,0,0,0,0,2],"quality":"Equal","isMain":false,"metricName":"# contigs (>= 25000 bp)"},{"values":[0,0,0,0,0,0,0,0,0,0,0,0,0,0],"quality":"Equal","isMain":false,"metricName":"# contigs (>= 50000 bp)"},{"values":[18858,7859,11859,13659,15048,31842,4525,1047,21533,12666,14442,14442,2356,40600],"quality":"More is better","isMain":true,"metricName":"Largest contig"},{"values":[42136,31833,35411,30747,33844,45011,31992,5386,36980,23361,27902,30639,18925,78436],"quality":"More is better","isMain":true,"metricName":"Total length"},{"values":[42136,30739,35411,27728,33844,43847,30028,1047,36980,23361,27902,30639,17037,78436],"quality":"More is better","isMain":true,"metricName":"Total length (>= 1000 bp)"},{"values":[42136,19317,30574,19349,29200,42422,0,0,35143,17966,22280,22280,0,78436],"quality":"More is better","isMain":false,"metricName":"Total length (>= 5000 bp)"},{"values":[42136,0,11859,13659,15048,42422,0,0,21533,12666,14442,14442,0,78436],"quality":"More is better","isMain":true,"metricName":"Total length (>= 10000 bp)"},{"values":[0,0,0,0,0,31842,0,0,0,0,0,0,0,78436],"quality":"More is better","isMain":false,"metricName":"Total length (>= 25000 bp)"},{"values":[0,0,0,0,0,0,0,0,0,0,0,0,0,0],"quality":"More is better","isMain":true,"metricName":"Total length (>= 50000 bp)"},{"values":[11730,5364,8301,5690,8539,31842,2066,882,21533,12666,14442,7838,1720,40600],"quality":"More is better","isMain":false,"metricName":"N50"},{"values":[11548,4840,5010,2434,5613,10580,1457,661,8511,5300,7838,3679,1374,37836],"quality":"More is better","isMain":false,"metricName":"N75"},{"values":[2,3,2,2,2,1,6,3,1,1,1,2,5,1],"quality":"Less is better","isMain":false,"metricName":"L50"},{"values":[3,4,4,4,3,2,11,5,2,2,2,3,8,2],"quality":"Less is better","isMain":false,"metricName":"L75"},{"values":["37.30","37.46","37.67","37.69","37.46","36.97","37.71","37.73","37.49","38.32","37.97","38.07","37.82","37.62"],"quality":"Equal","isMain":false,"metricName":"GC (%)"}]],["Predicted genes",[]],["Similarity statistics",[{"values":[0,0,0,0,0,0,0,0,0,0,0,0,0,0],"quality":"Equal","isMain":false,"metricName":"# similar correct contigs"},{"values":[0,0,0,0,0,0,0,0,0,0,0,0,0,0],"quality":"Equal","isMain":false,"metricName":"# similar misassembled blocks"}]],["Reference statistics",[{"values":[38528,38528,38528,38528,38528,38528,38528,38528,38528,38528,38528,38528,38528,38528],"quality":"Equal","isMain":false,"metricName":"Reference length"},{"values":[1,1,1,1,1,1,1,1,1,1,1,1,1,1],"quality":"Equal","isMain":false,"metricName":"Reference fragments"},{"values":["37.48","37.48","37.48","37.48","37.48","37.48","37.48","37.48","37.48","37.48","37.48","37.48","37.48","37.48"],"quality":"Equal","isMain":false,"metricName":"Reference GC (%)"}]]],"referenceName":"gi\_126011062\_ref\_NC","date":"18 July 2018, Wednesday, 16:15:17","order":[0,1,2,3,4,5,6,7,8,9,10,11,12,13],"assembliesNames":["ABySS\_127","ABySS\_63","CLC","IDBA\_UD","MEGAHIT","MIRA","Ray\_Meta","SOAPdenovo2","SPAdes","SPAdes\_meta","SPAdes\_sc","SPAdes\_sc\_careful","Velvet","Geneious"]},{"assembliesWithNs":null,"minContig":500,"report":[["Genome statistics",[{"values":["95.059","100.000","92.943","93.618","94.583","98.973","97.932","100.000","100.000","99.008","99.008","82.336","98.973"],"quality":"More is better","isMain":true,"metricName":"Genome fraction (%)"},{"values":["1.062","1.076","1.012","1.018","1.028","1.055","1.000","1.019","1.018","1.011","1.011","1.002","1.000"],"quality":"Less is better","isMain":true,"metricName":"Duplication ratio"},{"values":[9079,10008,7961,8115,8199,14361,8834,14681,12558,14421,14422,6829,14361],"quality":"More is better","isMain":true,"metricName":"Largest alignment"},{"values":[14602,15612,13644,13824,14106,15144,14210,14681,14675,14421,14422,11975,14361],"quality":"More is better","isMain":true,"metricName":"Total aligned length"},{"values":[13756,10008,7961,8115,8199,14361,8834,14781,12658,14521,14521,1482,14361],"quality":"More is better","isMain":false,"metricName":"NG50"},{"values":[13756,4661,4501,1719,1803,14361,5376,14781,12658,14521,14521,1133,14361],"quality":"More is better","isMain":false,"metricName":"NG75"},{"values":[9079,10008,7961,8115,8199,14361,8834,14681,12558,14421,14422,6829,14361],"quality":"More is better","isMain":false,"metricName":"NA50"},{"values":[4627,4661,4501,1719,1803,14361,5376,14681,12558,14421,14422,1463,14361],"quality":"More is better","isMain":false,"metricName":"NA75"},{"values":[9079,10008,7961,8115,8199,14361,8834,14681,12558,14421,14422,1482,14361],"quality":"More is better","isMain":true,"metricName":"NGA50"},{"values":[4627,4661,4501,1719,1803,14361,5376,14681,12558,14421,14422,1133,14361],"quality":"More is better","isMain":false,"metricName":"NGA75"},{"values":[1,1,1,1,1,1,1,1,1,1,1,2,1],"quality":"Less is better","isMain":false,"metricName":"LG50"},{"values":[1,2,2,3,3,1,2,1,1,1,1,4,1],"quality":"Less is better","isMain":false,"metricName":"LG75"},{"values":[1,1,1,1,1,1,1,1,1,1,1,1,1],"quality":"Less is better","isMain":false,"metricName":"LA50"},{"values":[2,2,2,3,3,1,2,1,1,1,1,3,1],"quality":"Less is better","isMain":false,"metricName":"LA75"},{"values":[1,1,1,1,1,1,1,1,1,1,1,2,1],"quality":"Less is better","isMain":true,"metricName":"LGA50"},{"values":[2,2,2,3,3,1,2,1,1,1,1,4,1],"quality":"Less is better","isMain":false,"metricName":"LGA75"}]],["Misassemblies",[{"values":[1,0,0,0,0,0,0,0,0,0,0,0,0],"quality":"Less is better","isMain":true,"metricName":"# misassemblies"},{"values":[1,0,0,0,0,0,0,0,0,0,0,0,0],"quality":"Less is better","isMain":false,"metricName":" # relocations"},{"values":[0,0,0,0,0,0,0,0,0,0,0,0,0],"quality":"Less is better","isMain":false,"metricName":" # translocations"},{"values":[0,0,0,0,0,0,0,0,0,0,0,0,0],"quality":"Less is better","isMain":false,"metricName":" # inversions"},{"values":[1,0,0,0,0,0,0,0,0,0,0,0,0],"quality":"Less is better","isMain":false,"metricName":"# misassembled contigs"},{"values":[13756,0,0,0,0,0,0,0,0,0,0,0,0],"quality":"Less is better","isMain":true,"metricName":"Misassembled contigs length"},{"values":[0,0,0,0,0,0,0,1,1,1,1,0,0],"quality":"Less is better","isMain":false,"metricName":"# local misassemblies"},{"values":[0,0,0,0,0,0,0,0,0,0,0,0,0],"quality":"Less is better","isMain":false,"metricName":"# unaligned mis. contigs"}]],["Unaligned",[{"values":[0,0,0,0,0,0,0,0,0,0,0,0,0],"quality":"Less is better","isMain":false,"metricName":"# fully unaligned contigs"},{"values":[0,0,0,0,0,0,0,0,0,0,0,0,0],"quality":"Less is better","isMain":false,"metricName":"Fully unaligned length"},{"values":[0,0,0,0,0,0,0,0,0,0,0,0,0],"quality":"Less is better","isMain":false,"metricName":"# partially unaligned contigs"},{"values":[0,0,0,0,0,0,0,0,0,0,0,0,0],"quality":"Less is better","isMain":false,"metricName":"Partially unaligned length"}]],["Mismatches",[{"values":[0,0,1,0,0,1,0,0,0,0,0,0,1],"quality":"Less is better","isMain":false,"metricName":"# mismatches"},{"values":[0,0,0,0,0,0,0,0,0,0,0,0,0],"quality":"Less is better","isMain":false,"metricName":"# indels"},{"values":[0,0,0,0,0,0,0,0,0,0,0,0,0],"quality":"Less is better","isMain":false,"metricName":"Indels length"},{"values":["0.00","0.00","7.42","0.00","0.00","6.96","0.00","0.00","0.00","0.00","0.00","0.00","6.96"],"quality":"Less is better","isMain":true,"metricName":"# mismatches per 100 kbp"},{"values":["0.00","0.00","0.00","0.00","0.00","0.00","0.00","0.00","0.00","0.00","0.00","0.00","0.00"],"quality":"Less is better","isMain":true,"metricName":"# indels per 100 kbp"},{"values":[0,0,0,0,0,0,0,0,0,0,0,0,0],"quality":"Less is better","isMain":false,"metricName":" # indels (<= 5 bp)"},{"values":[0,0,0,0,0,0,0,0,0,0,0,0,0],"quality":"Less is better","isMain":false,"metricName":" # indels (> 5 bp)"},{"values":[50,1,0,0,0,2,0,100,100,100,99,0,2],"quality":"Less is better","isMain":false,"metricName":"# N's"},{"values":["341.25","6.41","0.00","0.00","0.00","13.21","0.00","676.54","676.82","688.66","681.77","0.00","13.93"],"quality":"Less is better","isMain":true,"metricName":"# N's per 100 kbp"}]],["Statistics without reference",[{"values":[2,3,3,5,5,2,2,1,2,1,1,5,1],"quality":"Equal","isMain":true,"metricName":"# contigs"},{"values":[1,2,3,5,5,1,2,1,2,1,1,5,1],"quality":"Equal","isMain":false,"metricName":"# contigs (>= 1000 bp)"},{"values":[1,1,1,1,1,1,2,1,1,1,1,1,1],"quality":"Equal","isMain":false,"metricName":"# contigs (>= 5000 bp)"},{"values":[1,1,0,0,0,1,0,1,1,1,1,0,1],"quality":"Equal","isMain":false,"metricName":"# contigs (>= 10000 bp)"},{"values":[0,0,0,0,0,0,0,0,0,0,0,0,0],"quality":"Equal","isMain":false,"metricName":"# contigs (>= 25000 bp)"},{"values":[0,0,0,0,0,0,0,0,0,0,0,0,0],"quality":"Equal","isMain":false,"metricName":"# contigs (>= 50000 bp)"},{"values":[13756,10008,7961,8115,8199,14361,8834,14781,12658,14521,14521,6829,14361],"quality":"More is better","isMain":true,"metricName":"Largest contig"},{"values":[14652,15612,13644,13824,14106,15144,14210,14781,14775,14521,14521,11975,14361],"quality":"More is better","isMain":true,"metricName":"Total length"},{"values":[13756,14669,13644,13824,14106,14361,14210,14781,14775,14521,14521,11975,14361],"quality":"More is better","isMain":true,"metricName":"Total length (>= 1000 bp)"},{"values":[13756,10008,7961,8115,8199,14361,14210,14781,12658,14521,14521,6829,14361],"quality":"More is better","isMain":false,"metricName":"Total length (>= 5000 bp)"},{"values":[13756,10008,0,0,0,14361,0,14781,12658,14521,14521,0,14361],"quality":"More is better","isMain":true,"metricName":"Total length (>= 10000 bp)"},{"values":[0,0,0,0,0,0,0,0,0,0,0,0,0],"quality":"More is better","isMain":false,"metricName":"Total length (>= 25000 bp)"},{"values":[0,0,0,0,0,0,0,0,0,0,0,0,0],"quality":"More is better","isMain":true,"metricName":"Total length (>= 50000 bp)"},{"values":[13756,10008,7961,8115,8199,14361,8834,14781,12658,14521,14521,6829,14361],"quality":"More is better","isMain":false,"metricName":"N50"},{"values":[13756,4661,4501,1719,1803,14361,5376,14781,12658,14521,14521,1463,14361],"quality":"More is better","isMain":false,"metricName":"N75"},{"values":[1,1,1,1,1,1,1,1,1,1,1,1,1],"quality":"Less is better","isMain":false,"metricName":"L50"},{"values":[1,2,2,3,3,1,2,1,1,1,1,3,1],"quality":"Less is better","isMain":false,"metricName":"L75"},{"values":["34.34","34.80","33.47","33.55","33.69","34.73","33.98","34.26","34.15","34.28","34.28","33.78","34.15"],"quality":"Equal","isMain":false,"metricName":"GC (%)"}]],["Predicted genes",[]],["Similarity statistics",[{"values":[0,0,0,0,0,0,0,0,0,0,0,0,0],"quality":"Equal","isMain":false,"metricName":"# similar correct contigs"},{"values":[0,0,0,0,0,0,0,0,0,0,0,0,0],"quality":"Equal","isMain":false,"metricName":"# similar misassembled blocks"}]],["Reference statistics",[{"values":[14510,14510,14510,14510,14510,14510,14510,14510,14510,14510,14510,14510,14510],"quality":"Equal","isMain":false,"metricName":"Reference length"},{"values":[1,1,1,1,1,1,1,1,1,1,1,1,1],"quality":"Equal","isMain":false,"metricName":"Reference fragments"},{"values":["34.24","34.24","34.24","34.24","34.24","34.24","34.24","34.24","34.24","34.24","34.24","34.24","34.24"],"quality":"Equal","isMain":false,"metricName":"Reference GC (%)"}]]],"referenceName":"gi\_13095658\_ref\_NC","date":"18 July 2018, Wednesday, 16:15:28","order":[0,1,2,3,4,5,6,7,8,9,10,11,12],"assembliesNames":["ABySS\_127","ABySS\_63","CLC","IDBA\_UD","MEGAHIT","MIRA","Ray\_Meta","SPAdes","SPAdes\_meta","SPAdes\_sc","SPAdes\_sc\_careful","Velvet","Geneious"]},{"assembliesWithNs":null,"minContig":500,"report":[["Genome statistics",[{"values":["86.037","61.202","90.548","74.357","84.518","100.000","99.024","88.491","77.824","81.485","79.627","53.731","100.000"],"quality":"More is better","isMain":true,"metricName":"Genome fraction (%)"},{"values":["1.000","1.001","1.017","1.009","1.011","1.414","1.066","1.028","1.000","1.008","1.004","1.003","1.583"],"quality":"Less is better","isMain":true,"metricName":"Duplication ratio"},{"values":[18785,13907,17150,13652,13940,20202,21268,17147,16829,14131,13939,5594,20202],"quality":"More is better","isMain":true,"metricName":"Largest alignment"},{"values":[30576,21773,32685,26657,30330,49215,37516,32337,27657,29178,28408,19155,56265],"quality":"More is better","isMain":true,"metricName":"Total aligned length"},{"values":[18785,3826,5797,2600,4658,79060,21268,5464,3541,5524,5816,635,136335],"quality":"More is better","isMain":false,"metricName":"NG50"},{"values":[2497,null,1793,510,1063,79060,6027,876,676,891,642,null,136335],"quality":"More is better","isMain":false,"metricName":"NG75"},{"values":[18785,13907,17150,13652,4658,null,21268,17147,16829,5524,5816,5063,null],"quality":"More is better","isMain":false,"metricName":"NA50"},{"values":[5878,956,1915,2600,1250,null,3718,2161,2248,2248,2030,2061,null],"quality":"More is better","isMain":false,"metricName":"NA75"},{"values":[18785,956,5745,2600,4658,20202,21268,5464,3541,5524,5816,635,20202],"quality":"More is better","isMain":true,"metricName":"NGA50"},{"values":[2497,null,1456,510,1063,8301,6027,876,665,891,642,null,6592],"quality":"More is better","isMain":false,"metricName":"NGA75"},{"values":[1,2,2,3,2,1,1,2,2,2,2,7,1],"quality":"Less is better","isMain":false,"metricName":"LG50"},{"values":[3,null,4,11,8,1,2,5,6,7,9,null,1],"quality":"Less is better","isMain":false,"metricName":"LG75"},{"values":[1,1,1,1,2,null,1,1,1,2,2,2,null],"quality":"Less is better","isMain":false,"metricName":"LA50"},{"values":[2,3,3,3,5,null,3,3,3,3,3,4,null],"quality":"Less is better","isMain":false,"metricName":"LA75"},{"values":[1,3,2,3,2,1,1,2,2,2,2,7,1],"quality":"Less is better","isMain":true,"metricName":"LGA50"},{"values":[3,null,5,11,8,2,2,5,6,7,9,null,2],"quality":"Less is better","isMain":false,"metricName":"LGA75"}]],["Misassemblies",[{"values":[0,0,0,0,0,0,0,0,0,0,0,0,0],"quality":"Less is better","isMain":true,"metricName":"# misassemblies"},{"values":[0,0,0,0,0,0,0,0,0,0,0,0,0],"quality":"Less is better","isMain":false,"metricName":" # relocations"},{"values":[0,0,0,0,0,0,0,0,0,0,0,0,0],"quality":"Less is better","isMain":false,"metricName":" # translocations"},{"values":[0,0,0,0,0,0,0,0,0,0,0,0,0],"quality":"Less is better","isMain":false,"metricName":" # inversions"},{"values":[0,0,0,0,0,0,0,0,0,0,0,0,0],"quality":"Less is better","isMain":false,"metricName":"# misassembled contigs"},{"values":[0,0,0,0,0,0,0,0,0,0,0,0,0],"quality":"Less is better","isMain":true,"metricName":"Misassembled contigs length"},{"values":[0,0,0,0,0,1,0,0,0,0,0,0,0],"quality":"Less is better","isMain":false,"metricName":"# local misassemblies"},{"values":[0,0,0,0,0,1,0,0,0,0,0,0,2],"quality":"Less is better","isMain":false,"metricName":"# unaligned mis. contigs"}]],["Unaligned",[{"values":[0,0,0,0,0,0,0,0,0,0,0,0,0],"quality":"Less is better","isMain":false,"metricName":"# fully unaligned contigs"},{"values":[0,0,0,0,0,0,0,0,0,0,0,0,0],"quality":"Less is better","isMain":false,"metricName":"Fully unaligned length"},{"values":[0,1,0,0,0,2,0,0,0,0,0,0,2],"quality":"Less is better","isMain":false,"metricName":"# partially unaligned contigs"},{"values":[0,1996,0,0,0,64642,0,0,0,0,0,0,171495],"quality":"Less is better","isMain":false,"metricName":"Partially unaligned length"}]],["Mismatches",[{"values":[0,0,7,0,54,79,6,1,6,32,20,3,118],"quality":"Less is better","isMain":false,"metricName":"# mismatches"},{"values":[0,0,0,0,2,16,0,0,0,0,1,0,4],"quality":"Less is better","isMain":false,"metricName":"# indels"},{"values":[0,0,0,0,3,18,0,0,0,0,9,0,4],"quality":"Less is better","isMain":false,"metricName":"Indels length"},{"values":["0.00","0.00","21.75","0.00","179.78","222.30","17.05","3.18","21.69","110.50","70.68","15.71","332.04"],"quality":"Less is better","isMain":true,"metricName":"# mismatches per 100 kbp"},{"values":["0.00","0.00","0.00","0.00","6.66","45.02","0.00","0.00","0.00","0.00","3.53","0.00","11.26"],"quality":"Less is better","isMain":true,"metricName":"# indels per 100 kbp"},{"values":[0,0,0,0,2,16,0,0,0,0,0,0,4],"quality":"Less is better","isMain":false,"metricName":" # indels (<= 5 bp)"},{"values":[0,0,0,0,0,0,0,0,0,0,1,0,0],"quality":"Less is better","isMain":false,"metricName":" # indels (> 5 bp)"},{"values":[0,50,0,0,0,4,0,0,0,0,0,0,265],"quality":"Less is better","isMain":false,"metricName":"# N's"},{"values":["0.00","210.36","0.00","0.00","0.00","3.48","0.00","0.00","0.00","0.00","0.00","0.00","116.36"],"quality":"Less is better","isMain":true,"metricName":"# N's per 100 kbp"}]],["Statistics without reference",[{"values":[7,8,11,11,12,5,7,13,7,11,11,9,2],"quality":"Equal","isMain":true,"metricName":"# contigs"},{"values":[4,2,6,5,8,4,6,4,5,6,5,4,2],"quality":"Equal","isMain":false,"metricName":"# contigs (>= 1000 bp)"},{"values":[2,1,2,1,1,3,2,2,1,2,2,2,2],"quality":"Equal","isMain":false,"metricName":"# contigs (>= 5000 bp)"},{"values":[1,1,1,1,1,3,1,1,1,1,1,0,2],"quality":"Equal","isMain":false,"metricName":"# contigs (>= 10000 bp)"},{"values":[0,0,0,0,0,1,0,0,0,0,0,0,2],"quality":"Equal","isMain":false,"metricName":"# contigs (>= 25000 bp)"},{"values":[0,0,0,0,0,1,0,0,0,0,0,0,2],"quality":"Equal","isMain":false,"metricName":"# contigs (>= 50000 bp)"},{"values":[18785,15903,17150,13652,13940,79060,21268,17147,16829,14131,13939,5594,136335],"quality":"More is better","isMain":true,"metricName":"Largest contig"},{"values":[30576,23769,32737,26657,30353,114901,37516,32337,27668,29178,28408,19155,227745],"quality":"More is better","isMain":true,"metricName":"Total length"},{"values":[28779,19729,29566,22426,27014,113970,36609,26071,26358,26009,24045,15500,227745],"quality":"More is better","isMain":true,"metricName":"Total length (>= 1000 bp)"},{"values":[24663,15903,22947,13652,13940,110478,27295,22611,16829,19655,19755,10657,227745],"quality":"More is better","isMain":false,"metricName":"Total length (>= 5000 bp)"},{"values":[18785,15903,17150,13652,13940,110478,21268,17147,16829,14131,13939,0,227745],"quality":"More is better","isMain":true,"metricName":"Total length (>= 10000 bp)"},{"values":[0,0,0,0,0,79060,0,0,0,0,0,0,227745],"quality":"More is better","isMain":false,"metricName":"Total length (>= 25000 bp)"},{"values":[0,0,0,0,0,79060,0,0,0,0,0,0,227745],"quality":"More is better","isMain":true,"metricName":"Total length (>= 50000 bp)"},{"values":[18785,15903,17150,13652,4658,79060,21268,17147,16829,5524,5816,5063,136335],"quality":"More is better","isMain":false,"metricName":"N50"},{"values":[5878,3826,1915,2600,1250,20875,3718,2161,2248,2248,2030,2061,91410],"quality":"More is better","isMain":false,"metricName":"N75"},{"values":[1,1,1,1,2,1,1,1,1,2,2,2,1],"quality":"Less is better","isMain":false,"metricName":"L50"},{"values":[2,2,3,3,5,2,3,3,3,3,3,4,2],"quality":"Less is better","isMain":false,"metricName":"L75"},{"values":["35.34","35.82","35.43","35.41","35.34","35.79","35.10","35.24","35.00","35.09","35.19","34.93","35.67"],"quality":"Equal","isMain":false,"metricName":"GC (%)"}]],["Predicted genes",[]],["Similarity statistics",[{"values":[0,0,0,0,0,0,0,0,0,0,0,0,0],"quality":"Equal","isMain":false,"metricName":"# similar correct contigs"},{"values":[0,0,0,0,0,0,0,0,0,0,0,0,0],"quality":"Equal","isMain":false,"metricName":"# similar misassembled blocks"}]],["Reference statistics",[{"values":[35538,35538,35538,35538,35538,35538,35538,35538,35538,35538,35538,35538,35538],"quality":"Equal","isMain":false,"metricName":"Reference length"},{"values":[1,1,1,1,1,1,1,1,1,1,1,1,1],"quality":"Equal","isMain":false,"metricName":"Reference fragments"},{"values":["35.19","35.19","35.19","35.19","35.19","35.19","35.19","35.19","35.19","35.19","35.19","35.19","35.19"],"quality":"Equal","isMain":false,"metricName":"Reference GC (%)"}]]],"referenceName":"gi\_13095680\_ref\_NC","date":"18 July 2018, Wednesday, 16:15:40","order":[0,1,2,3,4,5,6,7,8,9,10,11,12],"assembliesNames":["ABySS\_127","ABySS\_63","CLC","IDBA\_UD","MEGAHIT","MIRA","Ray\_Meta","SPAdes","SPAdes\_meta","SPAdes\_sc","SPAdes\_sc\_careful","Velvet","Geneious"]},{"assembliesWithNs":null,"minContig":500,"report":[["Genome statistics",[{"values":["87.154","46.532","82.435","63.989","80.408","98.661","37.928","7.152","80.939","37.871","53.688","55.450","22.169","97.483"],"quality":"More is better","isMain":true,"metricName":"Genome fraction (%)"},{"values":["1.081","1.003","1.003","1.024","1.041","1.400","1.032","1.000","1.015","1.000","1.000","0.999","1.000","1.641"],"quality":"Less is better","isMain":true,"metricName":"Duplication ratio"},{"values":[10574,4404,4453,2614,4083,12847,2032,938,8486,5738,4389,8471,1994,27481],"quality":"More is better","isMain":true,"metricName":"Largest alignment"},{"values":[39396,19519,34585,27413,35027,57750,16371,2992,34368,15844,22460,23166,9274,66845],"quality":"More is better","isMain":true,"metricName":"Total aligned length"},{"values":[42775,null,2048,821,1587,79060,null,null,3971,null,954,1146,null,136335],"quality":"More is better","isMain":false,"metricName":"NG50"},{"values":[42775,null,923,null,644,79060,null,null,645,null,null,null,null,136335],"quality":"More is better","isMain":false,"metricName":"NG75"},{"values":[643,1667,2218,1617,1707,null,1166,581,5580,3858,3604,4695,904,null],"quality":"More is better","isMain":false,"metricName":"NA50"},{"values":[null,934,1132,821,1162,null,827,546,2206,2437,1594,1594,626,null],"quality":"More is better","isMain":false,"metricName":"NA75"},{"values":[5047,null,2048,821,1587,9485,null,null,3971,null,954,1146,null,27481],"quality":"More is better","isMain":true,"metricName":"NGA50"},{"values":[1004,null,923,null,644,6455,null,null,645,null,null,null,null,13587],"quality":"More is better","isMain":false,"metricName":"NGA75"},{"values":[1,null,7,13,10,1,null,null,4,null,10,7,null,1],"quality":"Less is better","isMain":false,"metricName":"LG50"},{"values":[1,null,15,null,20,1,null,null,10,null,null,null,null,1],"quality":"Less is better","isMain":false,"metricName":"LG75"},{"values":[12,4,6,7,8,null,6,3,3,2,3,2,4,null],"quality":"Less is better","isMain":false,"metricName":"LA50"},{"values":[null,8,10,13,14,null,10,4,5,3,6,5,7,null],"quality":"Less is better","isMain":false,"metricName":"LA75"},{"values":[3,null,7,13,10,2,null,null,4,null,10,7,null,1],"quality":"Less is better","isMain":true,"metricName":"LGA50"},{"values":[7,null,15,null,20,4,null,null,10,null,null,null,null,2],"quality":"Less is better","isMain":false,"metricName":"LGA75"}]],["Misassemblies",[{"values":[0,0,0,0,0,0,0,0,0,0,0,0,0,0],"quality":"Less is better","isMain":true,"metricName":"# misassemblies"},{"values":[0,0,0,0,0,0,0,0,0,0,0,0,0,0],"quality":"Less is better","isMain":false,"metricName":" # relocations"},{"values":[0,0,0,0,0,0,0,0,0,0,0,0,0,0],"quality":"Less is better","isMain":false,"metricName":" # translocations"},{"values":[0,0,0,0,0,0,0,0,0,0,0,0,0,0],"quality":"Less is better","isMain":false,"metricName":" # inversions"},{"values":[0,0,0,0,0,0,0,0,0,0,0,0,0,0],"quality":"Less is better","isMain":false,"metricName":"# misassembled contigs"},{"values":[0,0,0,0,0,0,0,0,0,0,0,0,0,0],"quality":"Less is better","isMain":true,"metricName":"Misassembled contigs length"},{"values":[1,0,0,0,0,0,0,0,0,0,0,0,0,0],"quality":"Less is better","isMain":false,"metricName":"# local misassemblies"},{"values":[1,0,0,0,0,1,0,0,0,0,0,0,0,2],"quality":"Less is better","isMain":false,"metricName":"# unaligned mis. contigs"}]],["Unaligned",[{"values":[0,0,0,0,0,0,0,0,0,0,0,0,0,0],"quality":"Less is better","isMain":false,"metricName":"# fully unaligned contigs"},{"values":[0,0,0,0,0,0,0,0,0,0,0,0,0,0],"quality":"Less is better","isMain":false,"metricName":"Fully unaligned length"},{"values":[2,0,0,0,0,1,0,1,0,0,0,0,0,2],"quality":"Less is better","isMain":false,"metricName":"# partially unaligned contigs"},{"values":[30764,0,0,0,0,69838,0,635,0,0,0,0,0,188458],"quality":"Less is better","isMain":false,"metricName":"Partially unaligned length"}]],["Mismatches",[{"values":[170,60,26,0,36,100,6,0,7,56,69,92,2,92],"quality":"Less is better","isMain":false,"metricName":"# mismatches"},{"values":[7,0,0,0,0,0,0,0,0,4,2,8,0,8],"quality":"Less is better","isMain":false,"metricName":"# indels"},{"values":[8,0,0,0,0,0,0,0,0,5,2,31,0,8],"quality":"Less is better","isMain":false,"metricName":"Indels length"},{"values":["466.26","308.23","75.39","0.00","107.02","242.28","37.81","0.00","20.67","353.47","307.21","396.60","21.57","225.60"],"quality":"Less is better","isMain":true,"metricName":"# mismatches per 100 kbp"},{"values":["19.20","0.00","0.00","0.00","0.00","0.00","0.00","0.00","0.00","25.25","8.90","34.49","0.00","19.62"],"quality":"Less is better","isMain":true,"metricName":"# indels per 100 kbp"},{"values":[7,0,0,0,0,0,0,0,0,4,2,5,0,8],"quality":"Less is better","isMain":false,"metricName":" # indels (<= 5 bp)"},{"values":[0,0,0,0,0,0,0,0,0,0,0,3,0,0],"quality":"Less is better","isMain":false,"metricName":" # indels (> 5 bp)"},{"values":[50,2,0,0,0,6,0,179,0,0,0,0,0,421],"quality":"Less is better","isMain":false,"metricName":"# N's"},{"values":["71.27","10.25","0.00","0.00","0.00","4.70","0.00","4935.21","0.00","0.00","0.00","0.00","0.00","164.86"],"quality":"Less is better","isMain":true,"metricName":"# N's per 100 kbp"}]],["Statistics without reference",[{"values":[16,15,20,23,26,15,15,5,15,7,11,9,11,3],"quality":"Equal","isMain":true,"metricName":"# contigs"},{"values":[7,6,12,12,15,9,9,0,6,4,8,9,2,3],"quality":"Equal","isMain":false,"metricName":"# contigs (>= 1000 bp)"},{"values":[3,0,0,0,0,5,0,0,3,1,0,1,0,3],"quality":"Equal","isMain":false,"metricName":"# contigs (>= 5000 bp)"},{"values":[1,0,0,0,0,2,0,0,0,0,0,0,0,3],"quality":"Equal","isMain":false,"metricName":"# contigs (>= 10000 bp)"},{"values":[1,0,0,0,0,1,0,0,0,0,0,0,0,3],"quality":"Equal","isMain":false,"metricName":"# contigs (>= 25000 bp)"},{"values":[0,0,0,0,0,1,0,0,0,0,0,0,0,2],"quality":"Equal","isMain":false,"metricName":"# contigs (>= 50000 bp)"},{"values":[42775,4404,4453,2614,4083,79060,2032,938,8486,5738,4389,8474,1994,136335],"quality":"More is better","isMain":true,"metricName":"Largest contig"},{"values":[70160,19519,34585,27413,35027,127602,16371,3627,34368,15844,22463,23169,9274,255362],"quality":"More is better","isMain":true,"metricName":"Total length"},{"values":[64150,12794,28570,20243,27860,123386,12130,0,28512,13309,19908,23169,3007,255362],"quality":"More is better","isMain":true,"metricName":"Total length (>= 1000 bp)"},{"values":[54068,0,0,0,0,114438,0,0,20535,5738,0,8474,0,255362],"quality":"More is better","isMain":false,"metricName":"Total length (>= 5000 bp)"},{"values":[42775,0,0,0,0,91907,0,0,0,0,0,0,0,255362],"quality":"More is better","isMain":true,"metricName":"Total length (>= 10000 bp)"},{"values":[42775,0,0,0,0,79060,0,0,0,0,0,0,0,255362],"quality":"More is better","isMain":false,"metricName":"Total length (>= 25000 bp)"},{"values":[0,0,0,0,0,79060,0,0,0,0,0,0,0,227745],"quality":"More is better","isMain":true,"metricName":"Total length (>= 50000 bp)"},{"values":[42775,1667,2218,1617,1707,79060,1166,757,5580,3858,3604,4695,904,136335],"quality":"More is better","isMain":false,"metricName":"N50"},{"values":[5047,934,1132,821,1162,9485,827,581,2206,2437,1594,1594,626,91410],"quality":"More is better","isMain":false,"metricName":"N75"},{"values":[1,4,6,7,8,1,6,3,3,2,3,2,4,1],"quality":"Less is better","isMain":false,"metricName":"L50"},{"values":[3,8,10,13,14,3,10,4,5,3,6,5,7,2],"quality":"Less is better","isMain":false,"metricName":"L75"},{"values":["35.60","35.97","35.15","35.49","35.38","35.58","35.75","35.06","35.39","35.67","35.25","35.22","34.70","35.66"],"quality":"Equal","isMain":false,"metricName":"GC (%)"}]],["Predicted genes",[]],["Similarity statistics",[{"values":[0,0,0,0,0,0,0,0,0,0,0,0,0,0],"quality":"Equal","isMain":false,"metricName":"# similar correct contigs"},{"values":[0,0,0,0,0,0,0,0,0,0,0,0,0,0],"quality":"Equal","isMain":false,"metricName":"# similar misassembled blocks"}]],["Reference statistics",[{"values":[41834,41834,41834,41834,41834,41834,41834,41834,41834,41834,41834,41834,41834,41834],"quality":"Equal","isMain":false,"metricName":"Reference length"},{"values":[1,1,1,1,1,1,1,1,1,1,1,1,1,1],"quality":"Equal","isMain":false,"metricName":"Reference fragments"},{"values":["35.32","35.32","35.32","35.32","35.32","35.32","35.32","35.32","35.32","35.32","35.32","35.32","35.32","35.32"],"quality":"Equal","isMain":false,"metricName":"Reference GC (%)"}]]],"referenceName":"gi\_13095743\_ref\_NC","date":"18 July 2018, Wednesday, 16:15:52","order":[0,1,2,3,4,5,6,7,8,9,10,11,12,13],"assembliesNames":["ABySS\_127","ABySS\_63","CLC","IDBA\_UD","MEGAHIT","MIRA","Ray\_Meta","SOAPdenovo2","SPAdes","SPAdes\_meta","SPAdes\_sc","SPAdes\_sc\_careful","Velvet","Geneious"]},{"assembliesWithNs":null,"minContig":500,"report":[["Genome statistics",[{"values":["100.000","84.040","94.909","86.925","90.492","100.000","95.943","94.771","94.449","86.606","88.097","52.995","100.000"],"quality":"More is better","isMain":true,"metricName":"Genome fraction (%)"},{"values":["1.076","1.002","1.056","1.021","1.024","1.244","1.115","1.007","1.003","1.003","1.005","1.000","1.308"],"quality":"Less is better","isMain":true,"metricName":"Duplication ratio"},{"values":[21097,12206,11505,11623,12711,8403,18145,16844,21630,12003,16177,7468,26532],"quality":"More is better","isMain":true,"metricName":"Largest alignment"},{"values":[39739,31114,37015,32792,34179,45738,39370,35271,35005,32108,32715,19581,48315],"quality":"More is better","isMain":true,"metricName":"Total aligned length"},{"values":[42775,6625,8853,2660,3870,79060,14443,14292,21630,8596,9777,685,136335],"quality":"More is better","isMain":false,"metricName":"NG50"},{"values":[42775,1901,4166,1114,1985,79060,14443,14292,12781,4380,2161,null,136335],"quality":"More is better","isMain":false,"metricName":"NG75"},{"values":[14279,2961,8853,2660,3805,null,14443,14292,21630,4380,9777,3372,null],"quality":"More is better","isMain":false,"metricName":"NA50"},{"values":[14279,1601,4166,1321,1985,null,1415,14292,12781,2333,9777,1112,null],"quality":"More is better","isMain":false,"metricName":"NA75"},{"values":[21097,2844,8853,2660,3805,7076,14443,14292,21630,4380,9777,685,26532],"quality":"More is better","isMain":true,"metricName":"NGA50"},{"values":[14279,954,4166,1114,1413,4350,14443,14292,12781,2333,2161,null,9505],"quality":"More is better","isMain":false,"metricName":"NGA75"},{"values":[1,2,2,3,3,1,2,2,1,2,2,10,1],"quality":"Less is better","isMain":false,"metricName":"LG50"},{"values":[1,6,4,10,6,1,2,2,2,4,3,null,1],"quality":"Less is better","isMain":false,"metricName":"LG75"},{"values":[2,3,2,3,3,null,2,2,1,3,2,2,null],"quality":"Less is better","isMain":false,"metricName":"LA50"},{"values":[2,7,4,7,6,null,5,2,2,6,2,5,null],"quality":"Less is better","isMain":false,"metricName":"LA75"},{"values":[1,4,2,3,3,3,2,2,1,3,2,10,1],"quality":"Less is better","isMain":true,"metricName":"LGA50"},{"values":[2,8,4,10,7,5,2,2,2,6,3,null,2],"quality":"Less is better","isMain":false,"metricName":"LGA75"}]],["Misassemblies",[{"values":[1,0,0,0,0,0,0,0,0,0,0,0,0],"quality":"Less is better","isMain":true,"metricName":"# misassemblies"},{"values":[1,0,0,0,0,0,0,0,0,0,0,0,0],"quality":"Less is better","isMain":false,"metricName":" # relocations"},{"values":[0,0,0,0,0,0,0,0,0,0,0,0,0],"quality":"Less is better","isMain":false,"metricName":" # translocations"},{"values":[0,0,0,0,0,0,0,0,0,0,0,0,0],"quality":"Less is better","isMain":false,"metricName":" # inversions"},{"values":[1,0,0,0,0,0,0,0,0,0,0,0,0],"quality":"Less is better","isMain":false,"metricName":"# misassembled contigs"},{"values":[42775,0,0,0,0,0,0,0,0,0,0,0,0],"quality":"Less is better","isMain":true,"metricName":"Misassembled contigs length"},{"values":[0,0,0,0,0,0,0,0,0,0,0,0,0],"quality":"Less is better","isMain":false,"metricName":"# local misassemblies"},{"values":[0,0,0,0,0,2,1,0,0,0,0,0,2],"quality":"Less is better","isMain":false,"metricName":"# unaligned mis. contigs"}]],["Unaligned",[{"values":[0,0,0,0,0,0,0,0,0,0,0,0,0],"quality":"Less is better","isMain":false,"metricName":"# fully unaligned contigs"},{"values":[0,0,0,0,0,0,0,0,0,0,0,0,0],"quality":"Less is better","isMain":false,"metricName":"Fully unaligned length"},{"values":[1,1,0,0,0,2,1,0,0,1,0,0,2],"quality":"Less is better","isMain":false,"metricName":"# partially unaligned contigs"},{"values":[7399,3541,0,0,0,95542,9819,0,0,6075,0,0,179430],"quality":"Less is better","isMain":false,"metricName":"Partially unaligned length"}]],["Mismatches",[{"values":[54,33,1,0,5,41,5,0,91,57,20,0,45],"quality":"Less is better","isMain":false,"metricName":"# mismatches"},{"values":[5,0,0,0,0,0,0,0,3,2,1,0,0],"quality":"Less is better","isMain":false,"metricName":"# indels"},{"values":[38,0,0,0,0,0,0,0,3,2,1,0,0],"quality":"Less is better","isMain":false,"metricName":"Indels length"},{"values":["146.15","106.27","2.85","0.00","14.95","110.96","14.10","0.00","260.76","178.12","61.44","0.00","121.79"],"quality":"Less is better","isMain":true,"metricName":"# mismatches per 100 kbp"},{"values":["13.53","0.00","0.00","0.00","0.00","0.00","0.00","0.00","8.60","6.25","3.07","0.00","0.00"],"quality":"Less is better","isMain":true,"metricName":"# indels per 100 kbp"},{"values":[2,0,0,0,0,0,0,0,3,2,1,0,0],"quality":"Less is better","isMain":false,"metricName":" # indels (<= 5 bp)"},{"values":[3,0,0,0,0,0,0,0,0,0,0,0,0],"quality":"Less is better","isMain":false,"metricName":" # indels (> 5 bp)"},{"values":[51,0,0,0,0,7,0,0,0,0,0,0,265],"quality":"Less is better","isMain":false,"metricName":"# N's"},{"values":["108.19","0.00","0.00","0.00","0.00","4.95","0.00","0.00","0.00","0.00","0.00","0.00","116.36"],"quality":"Less is better","isMain":true,"metricName":"# N's per 100 kbp"}]],["Statistics without reference",[{"values":[2,12,9,15,12,16,6,5,3,8,7,11,2],"quality":"Equal","isMain":true,"metricName":"# contigs"},{"values":[2,7,7,11,10,8,5,4,2,7,5,5,2],"quality":"Equal","isMain":false,"metricName":"# contigs (>= 1000 bp)"},{"values":[1,2,3,1,1,4,3,2,2,3,2,1,2],"quality":"Equal","isMain":false,"metricName":"# contigs (>= 5000 bp)"},{"values":[1,1,1,1,1,2,3,2,2,1,1,0,2],"quality":"Equal","isMain":false,"metricName":"# contigs (>= 10000 bp)"},{"values":[1,0,0,0,0,2,0,0,0,0,0,0,2],"quality":"Equal","isMain":false,"metricName":"# contigs (>= 25000 bp)"},{"values":[0,0,0,0,0,1,0,0,0,0,0,0,2],"quality":"Equal","isMain":false,"metricName":"# contigs (>= 50000 bp)"},{"values":[42775,12206,11505,11623,12711,79060,18301,16844,21630,12003,16177,7468,136335],"quality":"More is better","isMain":true,"metricName":"Largest contig"},{"values":[47138,34655,37015,32792,34244,141512,49345,35271,35005,38183,32715,19581,227745],"quality":"More is better","isMain":true,"metricName":"Total length"},{"values":[47138,30789,35499,29890,32798,135523,48480,34744,34411,37565,31338,15053,227745],"quality":"More is better","isMain":true,"metricName":"Total length (>= 1000 bp)"},{"values":[42775,18831,25485,11623,12711,125394,44097,31136,34411,25612,25954,7468,227745],"quality":"More is better","isMain":false,"metricName":"Total length (>= 5000 bp)"},{"values":[42775,12206,11505,11623,12711,109838,44097,31136,34411,12003,16177,0,227745],"quality":"More is better","isMain":true,"metricName":"Total length (>= 10000 bp)"},{"values":[42775,0,0,0,0,109838,0,0,0,0,0,0,227745],"quality":"More is better","isMain":false,"metricName":"Total length (>= 25000 bp)"},{"values":[0,0,0,0,0,79060,0,0,0,0,0,0,227745],"quality":"More is better","isMain":true,"metricName":"Total length (>= 50000 bp)"},{"values":[42775,6625,8853,2660,3870,79060,14443,14292,21630,8596,9777,3372,136335],"quality":"More is better","isMain":false,"metricName":"N50"},{"values":[42775,2651,4166,1321,2154,30778,11353,14292,12781,4380,9777,1112,91410],"quality":"More is better","isMain":false,"metricName":"N75"},{"values":[1,2,2,3,3,1,2,2,1,2,2,2,1],"quality":"Less is better","isMain":false,"metricName":"L50"},{"values":[1,5,4,7,5,2,3,2,2,4,2,5,2],"quality":"Less is better","isMain":false,"metricName":"L75"},{"values":["36.00","35.67","35.55","35.48","35.57","35.74","35.69","35.64","35.55","35.65","35.53","35.00","35.67"],"quality":"Equal","isMain":false,"metricName":"GC (%)"}]],["Predicted genes",[]],["Similarity statistics",[{"values":[0,0,0,0,0,0,0,0,0,0,0,0,0],"quality":"Equal","isMain":false,"metricName":"# similar correct contigs"},{"values":[0,0,0,0,0,0,0,0,0,0,0,0,0],"quality":"Equal","isMain":false,"metricName":"# similar misassembled blocks"}]],["Reference statistics",[{"values":[36949,36949,36949,36949,36949,36949,36949,36949,36949,36949,36949,36949,36949],"quality":"Equal","isMain":false,"metricName":"Reference length"},{"values":[1,1,1,1,1,1,1,1,1,1,1,1,1],"quality":"Equal","isMain":false,"metricName":"Reference fragments"},{"values":["35.68","35.68","35.68","35.68","35.68","35.68","35.68","35.68","35.68","35.68","35.68","35.68","35.68"],"quality":"Equal","isMain":false,"metricName":"Reference GC (%)"}]]],"referenceName":"gi\_13095805\_ref\_NC","date":"18 July 2018, Wednesday, 16:16:04","order":[0,1,2,3,4,5,6,7,8,9,10,11,12],"assembliesNames":["ABySS\_127","ABySS\_63","CLC","IDBA\_UD","MEGAHIT","MIRA","Ray\_Meta","SPAdes","SPAdes\_meta","SPAdes\_sc","SPAdes\_sc\_careful","Velvet","Geneious"]},{"assembliesWithNs":null,"minContig":500,"report":[["Genome statistics",[{"values":["42.736","49.723","92.532","96.156","91.562","72.755","84.455","98.542","89.858","97.286","97.286","61.496"],"quality":"More is better","isMain":true,"metricName":"Genome fraction (%)"},{"values":["1.000","1.000","1.002","1.001","1.000","1.000","1.017","1.001","1.001","1.001","1.001","1.001"],"quality":"Less is better","isMain":true,"metricName":"Duplication ratio"},{"values":[1663,1829,3730,3730,3661,3412,3473,3730,3730,3730,3730,3521],"quality":"More is better","isMain":true,"metricName":"Largest alignment"},{"values":[6392,7437,13861,14392,13695,10882,12632,14749,13450,14561,14561,9208],"quality":"More is better","isMain":true,"metricName":"Total aligned length"},{"values":[null,null,3495,3363,3132,2160,1852,3495,3495,3495,3495,975],"quality":"More is better","isMain":false,"metricName":"NG50"},{"values":[null,null,1083,1647,1468,null,1045,1647,1647,1647,1647,null],"quality":"More is better","isMain":false,"metricName":"NG75"},{"values":[1053,1332,3521,3521,3473,3186,3033,3495,3521,3495,3495,1790],"quality":"More is better","isMain":false,"metricName":"NA50"},{"values":[921,1185,3495,1647,1468,2160,1751,1647,3495,1647,1647,975],"quality":"More is better","isMain":false,"metricName":"NA75"},{"values":[null,null,3495,3363,3132,2160,1825,3495,3495,3495,3495,975],"quality":"More is better","isMain":true,"metricName":"NGA50"},{"values":[null,null,1083,1647,1468,null,904,1647,1647,1647,1647,null],"quality":"More is better","isMain":false,"metricName":"NGA75"},{"values":[null,null,3,3,3,3,3,3,3,3,3,4],"quality":"Less is better","isMain":false,"metricName":"LG50"},{"values":[null,null,4,4,4,null,5,4,4,4,4,null],"quality":"Less is better","isMain":false,"metricName":"LG75"},{"values":[3,3,2,2,2,2,2,3,2,3,3,2],"quality":"Less is better","isMain":false,"metricName":"LA50"},{"values":[4,4,3,4,4,3,4,4,3,4,4,4],"quality":"Less is better","isMain":false,"metricName":"LA75"},{"values":[null,null,3,3,3,3,3,3,3,3,3,4],"quality":"Less is better","isMain":true,"metricName":"LGA50"},{"values":[null,null,4,4,4,null,6,4,4,4,4,null],"quality":"Less is better","isMain":false,"metricName":"LGA75"}]],["Misassemblies",[{"values":[0,0,0,0,0,0,0,0,0,0,0,0],"quality":"Less is better","isMain":true,"metricName":"# misassemblies"},{"values":[0,0,0,0,0,0,0,0,0,0,0,0],"quality":"Less is better","isMain":false,"metricName":" # relocations"},{"values":[0,0,0,0,0,0,0,0,0,0,0,0],"quality":"Less is better","isMain":false,"metricName":" # translocations"},{"values":[0,0,0,0,0,0,0,0,0,0,0,0],"quality":"Less is better","isMain":false,"metricName":" # inversions"},{"values":[0,0,0,0,0,0,0,0,0,0,0,0],"quality":"Less is better","isMain":false,"metricName":"# misassembled contigs"},{"values":[0,0,0,0,0,0,0,0,0,0,0,0],"quality":"Less is better","isMain":true,"metricName":"Misassembled contigs length"},{"values":[0,0,0,0,0,0,0,0,0,0,0,0],"quality":"Less is better","isMain":false,"metricName":"# local misassemblies"},{"values":[0,0,0,0,0,0,0,0,0,0,0,0],"quality":"Less is better","isMain":false,"metricName":"# unaligned mis. contigs"}]],["Unaligned",[{"values":[0,0,0,0,0,0,0,0,0,0,0,0],"quality":"Less is better","isMain":false,"metricName":"# fully unaligned contigs"},{"values":[0,0,0,0,0,0,0,0,0,0,0,0],"quality":"Less is better","isMain":false,"metricName":"Fully unaligned length"},{"values":[0,0,0,0,0,0,0,0,0,0,0,0],"quality":"Less is better","isMain":false,"metricName":"# partially unaligned contigs"},{"values":[0,0,0,0,0,0,0,0,0,0,0,0],"quality":"Less is better","isMain":false,"metricName":"Partially unaligned length"}]],["Mismatches",[{"values":[2,0,8,3,4,0,6,10,5,6,5,3],"quality":"Less is better","isMain":false,"metricName":"# mismatches"},{"values":[0,0,0,0,0,0,2,0,0,0,0,0],"quality":"Less is better","isMain":false,"metricName":"# indels"},{"values":[0,0,0,0,0,0,105,0,0,0,0,0],"quality":"Less is better","isMain":false,"metricName":"Indels length"},{"values":["31.29","0.00","57.80","20.86","29.21","0.00","47.50","67.85","37.20","41.23","34.36","32.62"],"quality":"Less is better","isMain":true,"metricName":"# mismatches per 100 kbp"},{"values":["0.00","0.00","0.00","0.00","0.00","0.00","15.83","0.00","0.00","0.00","0.00","0.00"],"quality":"Less is better","isMain":true,"metricName":"# indels per 100 kbp"},{"values":[0,0,0,0,0,0,0,0,0,0,0,0],"quality":"Less is better","isMain":false,"metricName":" # indels (<= 5 bp)"},{"values":[0,0,0,0,0,0,2,0,0,0,0,0],"quality":"Less is better","isMain":false,"metricName":" # indels (> 5 bp)"},{"values":[0,0,0,0,0,9,217,0,0,0,0,0],"quality":"Less is better","isMain":false,"metricName":"# N's"},{"values":["0.00","0.00","0.00","0.00","0.00","82.71","1688.85","0.00","0.00","0.00","0.00","0.00"],"quality":"Less is better","isMain":true,"metricName":"# N's per 100 kbp"}]],["Statistics without reference",[{"values":[6,6,7,6,6,5,7,6,5,6,6,6],"quality":"Equal","isMain":true,"metricName":"# contigs"},{"values":[3,4,4,5,5,4,5,6,5,6,6,3],"quality":"Equal","isMain":false,"metricName":"# contigs (>= 1000 bp)"},{"values":[0,0,0,0,0,0,0,0,0,0,0,0],"quality":"Equal","isMain":false,"metricName":"# contigs (>= 5000 bp)"},{"values":[0,0,0,0,0,0,0,0,0,0,0,0],"quality":"Equal","isMain":false,"metricName":"# contigs (>= 10000 bp)"},{"values":[0,0,0,0,0,0,0,0,0,0,0,0],"quality":"Equal","isMain":false,"metricName":"# contigs (>= 25000 bp)"},{"values":[0,0,0,0,0,0,0,0,0,0,0,0],"quality":"Equal","isMain":false,"metricName":"# contigs (>= 50000 bp)"},{"values":[1663,1829,3730,3730,3661,3412,3473,3730,3730,3730,3730,3521],"quality":"More is better","isMain":true,"metricName":"Largest contig"},{"values":[6392,7437,13861,14392,13695,10882,12849,14749,13450,14561,14561,9208],"quality":"More is better","isMain":true,"metricName":"Total length"},{"values":[4142,6009,11829,13436,12776,10129,11228,14749,13450,14561,14561,6732],"quality":"More is better","isMain":true,"metricName":"Total length (>= 1000 bp)"},{"values":[0,0,0,0,0,0,0,0,0,0,0,0],"quality":"More is better","isMain":false,"metricName":"Total length (>= 5000 bp)"},{"values":[0,0,0,0,0,0,0,0,0,0,0,0],"quality":"More is better","isMain":true,"metricName":"Total length (>= 10000 bp)"},{"values":[0,0,0,0,0,0,0,0,0,0,0,0],"quality":"More is better","isMain":false,"metricName":"Total length (>= 25000 bp)"},{"values":[0,0,0,0,0,0,0,0,0,0,0,0],"quality":"More is better","isMain":true,"metricName":"Total length (>= 50000 bp)"},{"values":[1053,1332,3521,3521,3473,3186,3033,3495,3521,3495,3495,1790],"quality":"More is better","isMain":false,"metricName":"N50"},{"values":[921,1185,3495,1647,1468,2160,1825,1647,3495,1647,1647,975],"quality":"More is better","isMain":false,"metricName":"N75"},{"values":[3,3,2,2,2,2,2,3,2,3,3,2],"quality":"Less is better","isMain":false,"metricName":"L50"},{"values":[4,4,3,4,4,3,4,4,3,4,4,4],"quality":"Less is better","isMain":false,"metricName":"L75"},{"values":["36.66","37.07","35.82","35.81","35.79","36.25","35.79","35.84","35.88","35.84","35.85","36.33"],"quality":"Equal","isMain":false,"metricName":"GC (%)"}]],["Predicted genes",[]],["Similarity statistics",[{"values":[0,0,0,0,0,0,0,0,0,0,0,0],"quality":"Equal","isMain":false,"metricName":"# similar correct contigs"},{"values":[0,0,0,0,0,0,0,0,0,0,0,0],"quality":"Equal","isMain":false,"metricName":"# similar misassembled blocks"}]],["Reference statistics",[{"values":[14957,14957,14957,14957,14957,14957,14957,14957,14957,14957,14957,14957],"quality":"Equal","isMain":false,"metricName":"Reference length"},{"values":[1,1,1,1,1,1,1,1,1,1,1,1],"quality":"Equal","isMain":false,"metricName":"Reference fragments"},{"values":["35.88","35.88","35.88","35.88","35.88","35.88","35.88","35.88","35.88","35.88","35.88","35.88"],"quality":"Equal","isMain":false,"metricName":"Reference GC (%)"}]]],"referenceName":"gi\_13095862\_ref\_NC","date":"18 July 2018, Wednesday, 16:16:14","order":[0,1,2,3,4,5,6,7,8,9,10,11],"assembliesNames":["ABySS\_127","ABySS\_63","CLC","IDBA\_UD","MEGAHIT","MIRA","SOAPdenovo2","SPAdes","SPAdes\_meta","SPAdes\_sc","SPAdes\_sc\_careful","Velvet"]},{"assembliesWithNs":null,"minContig":500,"report":[["Genome statistics",[{"values":["99.987","98.933","100.000","99.631","100.000","100.000","72.699","65.334","100.000","89.565","100.000","100.000","90.171"],"quality":"More is better","isMain":true,"metricName":"Genome fraction (%)"},{"values":["1.006","1.003","1.000","1.003","1.007","1.043","1.080","1.191","1.003","1.000","1.004","1.004","1.004"],"quality":"Less is better","isMain":true,"metricName":"Duplication ratio"},{"values":[13737,9433,15179,13683,13767,15499,2091,2301,15223,13595,15234,15234,3692],"quality":"More is better","isMain":true,"metricName":"Largest alignment"},{"values":[15273,15065,15179,15165,15279,15499,11921,10075,15223,13595,15234,15234,13745],"quality":"More is better","isMain":true,"metricName":"Total aligned length"},{"values":[13737,9433,15179,13683,13767,15827,829,1610,15223,13595,15234,15234,2492],"quality":"More is better","isMain":false,"metricName":"NG50"},{"values":[13737,4224,15179,13683,13767,15827,526,844,15223,13595,15234,15234,1247],"quality":"More is better","isMain":false,"metricName":"NG75"},{"values":[13737,9433,15179,13683,13767,15499,1194,1474,15223,13595,15234,15234,3621],"quality":"More is better","isMain":false,"metricName":"NA50"},{"values":[13737,4224,15179,13683,13767,15499,771,709,15223,13595,15234,15234,1551],"quality":"More is better","isMain":false,"metricName":"NA75"},{"values":[13737,9433,15179,13683,13767,15499,829,1047,15223,13595,15234,15234,2492],"quality":"More is better","isMain":true,"metricName":"NGA50"},{"values":[13737,4224,15179,13683,13767,15499,526,null,15223,13595,15234,15234,1247],"quality":"More is better","isMain":false,"metricName":"NGA75"},{"values":[1,1,1,1,1,1,6,4,1,1,1,1,3],"quality":"Less is better","isMain":false,"metricName":"LG50"},{"values":[1,2,1,1,1,1,11,7,1,1,1,1,5],"quality":"Less is better","isMain":false,"metricName":"LG75"},{"values":[1,1,1,1,1,1,4,4,1,1,1,1,2],"quality":"Less is better","isMain":false,"metricName":"LA50"},{"values":[1,2,1,1,1,1,8,7,1,1,1,1,4],"quality":"Less is better","isMain":false,"metricName":"LA75"},{"values":[1,1,1,1,1,1,6,5,1,1,1,1,3],"quality":"Less is better","isMain":true,"metricName":"LGA50"},{"values":[1,2,1,1,1,1,11,null,1,1,1,1,5],"quality":"Less is better","isMain":false,"metricName":"LGA75"}]],["Misassemblies",[{"values":[0,0,0,0,0,0,0,0,0,0,0,0,0],"quality":"Less is better","isMain":true,"metricName":"# misassemblies"},{"values":[0,0,0,0,0,0,0,0,0,0,0,0,0],"quality":"Less is better","isMain":false,"metricName":" # relocations"},{"values":[0,0,0,0,0,0,0,0,0,0,0,0,0],"quality":"Less is better","isMain":false,"metricName":" # translocations"},{"values":[0,0,0,0,0,0,0,0,0,0,0,0,0],"quality":"Less is better","isMain":false,"metricName":" # inversions"},{"values":[0,0,0,0,0,0,0,0,0,0,0,0,0],"quality":"Less is better","isMain":false,"metricName":"# misassembled contigs"},{"values":[0,0,0,0,0,0,0,0,0,0,0,0,0],"quality":"Less is better","isMain":true,"metricName":"Misassembled contigs length"},{"values":[0,0,0,0,0,1,0,0,0,0,0,0,0],"quality":"Less is better","isMain":false,"metricName":"# local misassemblies"},{"values":[0,0,0,0,0,0,0,0,0,0,0,0,0],"quality":"Less is better","isMain":false,"metricName":"# unaligned mis. contigs"}]],["Unaligned",[{"values":[0,0,0,0,0,0,0,0,0,0,0,0,0],"quality":"Less is better","isMain":false,"metricName":"# fully unaligned contigs"},{"values":[0,0,0,0,0,0,0,0,0,0,0,0,0],"quality":"Less is better","isMain":false,"metricName":"Fully unaligned length"},{"values":[0,0,0,0,0,0,0,1,0,0,0,0,0],"quality":"Less is better","isMain":false,"metricName":"# partially unaligned contigs"},{"values":[0,0,0,0,0,0,0,683,0,0,0,0,0],"quality":"Less is better","isMain":false,"metricName":"Partially unaligned length"}]],["Mismatches",[{"values":[0,0,5,0,1,3,0,0,1,2,3,16,0],"quality":"Less is better","isMain":false,"metricName":"# mismatches"},{"values":[0,0,0,0,0,2,1,12,1,0,1,3,0],"quality":"Less is better","isMain":false,"metricName":"# indels"},{"values":[0,0,0,0,0,2,29,581,44,0,55,57,0],"quality":"Less is better","isMain":false,"metricName":"Indels length"},{"values":["0.00","0.00","32.94","0.00","6.59","19.76","0.00","0.00","6.59","14.71","19.76","105.41","0.00"],"quality":"Less is better","isMain":true,"metricName":"# mismatches per 100 kbp"},{"values":["0.00","0.00","0.00","0.00","0.00","13.18","9.06","121.00","6.59","0.00","6.59","19.76","0.00"],"quality":"Less is better","isMain":true,"metricName":"# indels per 100 kbp"},{"values":[0,0,0,0,0,2,0,0,0,0,0,2,0],"quality":"Less is better","isMain":false,"metricName":" # indels (<= 5 bp)"},{"values":[0,0,0,0,0,0,1,12,1,0,1,1,0],"quality":"Less is better","isMain":false,"metricName":" # indels (> 5 bp)"},{"values":[0,0,0,0,0,0,0,1852,0,0,0,0,0],"quality":"Less is better","isMain":false,"metricName":"# N's"},{"values":["0.00","0.00","0.00","0.00","0.00","0.00","0.00","14821.93","0.00","0.00","0.00","0.00","0.00"],"quality":"Less is better","isMain":true,"metricName":"# N's per 100 kbp"}]],["Statistics without reference",[{"values":[2,3,1,2,2,1,12,8,1,1,1,1,6],"quality":"Equal","isMain":true,"metricName":"# contigs"},{"values":[2,3,1,2,2,1,4,6,1,1,1,1,6],"quality":"Equal","isMain":false,"metricName":"# contigs (>= 1000 bp)"},{"values":[1,1,1,1,1,1,0,0,1,1,1,1,0],"quality":"Equal","isMain":false,"metricName":"# contigs (>= 5000 bp)"},{"values":[1,0,1,1,1,1,0,0,1,1,1,1,0],"quality":"Equal","isMain":false,"metricName":"# contigs (>= 10000 bp)"},{"values":[0,0,0,0,0,0,0,0,0,0,0,0,0],"quality":"Equal","isMain":false,"metricName":"# contigs (>= 25000 bp)"},{"values":[0,0,0,0,0,0,0,0,0,0,0,0,0],"quality":"Equal","isMain":false,"metricName":"# contigs (>= 50000 bp)"},{"values":[13737,9433,15179,13683,13767,15827,2091,2371,15223,13595,15234,15234,3692],"quality":"More is better","isMain":true,"metricName":"Largest contig"},{"values":[15273,15065,15179,15165,15279,15827,11921,12495,15223,13595,15234,15234,13745],"quality":"More is better","isMain":true,"metricName":"Total length"},{"values":[15273,15065,15179,15165,15279,15827,6285,10942,15223,13595,15234,15234,13745],"quality":"More is better","isMain":true,"metricName":"Total length (>= 1000 bp)"},{"values":[13737,9433,15179,13683,13767,15827,0,0,15223,13595,15234,15234,0],"quality":"More is better","isMain":false,"metricName":"Total length (>= 5000 bp)"},{"values":[13737,0,15179,13683,13767,15827,0,0,15223,13595,15234,15234,0],"quality":"More is better","isMain":true,"metricName":"Total length (>= 10000 bp)"},{"values":[0,0,0,0,0,0,0,0,0,0,0,0,0],"quality":"More is better","isMain":false,"metricName":"Total length (>= 25000 bp)"},{"values":[0,0,0,0,0,0,0,0,0,0,0,0,0],"quality":"More is better","isMain":true,"metricName":"Total length (>= 50000 bp)"},{"values":[13737,9433,15179,13683,13767,15827,1194,2007,15223,13595,15234,15234,3621],"quality":"More is better","isMain":false,"metricName":"N50"},{"values":[13737,4224,15179,13683,13767,15827,771,1606,15223,13595,15234,15234,1551],"quality":"More is better","isMain":false,"metricName":"N75"},{"values":[1,1,1,1,1,1,4,3,1,1,1,1,2],"quality":"Less is better","isMain":false,"metricName":"L50"},{"values":[1,2,1,1,1,1,8,5,1,1,1,1,4],"quality":"Less is better","isMain":false,"metricName":"L75"},{"values":["33.04","33.06","33.01","33.06","33.02","32.95","32.56","33.40","33.06","33.59","33.01","33.01","33.07"],"quality":"Equal","isMain":false,"metricName":"GC (%)"}]],["Predicted genes",[]],["Similarity statistics",[{"values":[0,0,0,0,0,0,0,0,0,0,0,0,0],"quality":"Equal","isMain":false,"metricName":"# similar correct contigs"},{"values":[0,0,0,0,0,0,0,0,0,0,0,0,0],"quality":"Equal","isMain":false,"metricName":"# similar misassembled blocks"}]],["Reference statistics",[{"values":[15179,15179,15179,15179,15179,15179,15179,15179,15179,15179,15179,15179,15179],"quality":"Equal","isMain":false,"metricName":"Reference length"},{"values":[1,1,1,1,1,1,1,1,1,1,1,1,1],"quality":"Equal","isMain":false,"metricName":"Reference fragments"},{"values":["33.04","33.04","33.04","33.04","33.04","33.04","33.04","33.04","33.04","33.04","33.04","33.04","33.04"],"quality":"Equal","isMain":false,"metricName":"Reference GC (%)"}]]],"referenceName":"gi\_13095891\_ref\_NC","date":"18 July 2018, Wednesday, 16:16:25","order":[0,1,2,3,4,5,6,7,8,9,10,11,12],"assembliesNames":["ABySS\_127","ABySS\_63","CLC","IDBA\_UD","MEGAHIT","MIRA","Ray\_Meta","SOAPdenovo2","SPAdes","SPAdes\_meta","SPAdes\_sc","SPAdes\_sc\_careful","Velvet"]},{"assembliesWithNs":null,"minContig":500,"report":[["Genome statistics",[{"values":["90.018","88.030","91.695","92.370","92.342","91.580","2.304","89.576","92.357","92.357","92.358","92.358","91.421","89.955"],"quality":"More is better","isMain":true,"metricName":"Genome fraction (%)"},{"values":["1.002","1.002","1.007","1.000","1.001","1.008","1.005","1.019","1.000","1.000","1.001","1.001","1.000","1.000"],"quality":"Less is better","isMain":true,"metricName":"Duplication ratio"},{"values":[32725,15465,69237,106333,83830,70325,795,27297,106333,68435,68356,68366,26365,71616],"quality":"More is better","isMain":true,"metricName":"Largest alignment"},{"values":[266241,260244,272516,272708,272784,272555,6814,266023,272589,272605,272566,272482,269931,265500],"quality":"More is better","isMain":true,"metricName":"Total aligned length"},{"values":[5148,4329,15166,59817,45217,39173,null,8118,85806,51337,49772,49725,14381,44965],"quality":"More is better","isMain":false,"metricName":"NG50"},{"values":[2602,1828,11033,37264,35331,25871,null,4177,85806,45261,25983,25983,8684,40220],"quality":"More is better","isMain":false,"metricName":"NG75"},{"values":[5204,5454,18854,59817,59838,37273,654,8819,85805,51337,49415,49368,14381,70325],"quality":"More is better","isMain":false,"metricName":"NA50"},{"values":[3128,2956,13733,35190,35331,27399,528,5029,45261,45261,45254,45245,10251,32489],"quality":"More is better","isMain":false,"metricName":"NA75"},{"values":[5148,4329,15166,59817,45217,32489,null,8008,85805,51337,49415,49368,14381,44965],"quality":"More is better","isMain":true,"metricName":"NGA50"},{"values":[2602,1828,11033,35190,35331,25871,null,3297,45261,45261,25982,25982,8684,27264],"quality":"More is better","isMain":false,"metricName":"NGA75"},{"values":[17,19,5,2,3,3,null,10,2,3,3,3,8,3],"quality":"Less is better","isMain":false,"metricName":"LG50"},{"values":[39,43,10,3,4,6,null,21,2,4,5,5,15,4],"quality":"Less is better","isMain":false,"metricName":"LG75"},{"values":[15,15,4,2,2,3,5,9,2,3,3,3,8,2],"quality":"Less is better","isMain":false,"metricName":"LA50"},{"values":[31,32,9,4,4,5,8,19,3,4,4,4,13,4],"quality":"Less is better","isMain":false,"metricName":"LA75"},{"values":[17,19,5,2,3,4,null,10,2,3,3,3,8,3],"quality":"Less is better","isMain":true,"metricName":"LGA50"},{"values":[39,43,10,4,4,6,null,23,3,4,5,5,15,5],"quality":"Less is better","isMain":false,"metricName":"LGA75"}]],["Misassemblies",[{"values":[1,0,0,1,0,1,0,0,1,0,0,0,0,1],"quality":"Less is better","isMain":true,"metricName":"# misassemblies"},{"values":[1,0,0,1,0,1,0,0,1,0,0,0,0,1],"quality":"Less is better","isMain":false,"metricName":" # relocations"},{"values":[0,0,0,0,0,0,0,0,0,0,0,0,0,0],"quality":"Less is better","isMain":false,"metricName":" # translocations"},{"values":[0,0,0,0,0,0,0,0,0,0,0,0,0,0],"quality":"Less is better","isMain":false,"metricName":" # inversions"},{"values":[1,0,0,1,0,1,0,0,1,0,0,0,0,1],"quality":"Less is better","isMain":false,"metricName":"# misassembled contigs"},{"values":[33856,0,0,141523,0,40220,0,0,141523,0,0,0,0,40220],"quality":"Less is better","isMain":true,"metricName":"Misassembled contigs length"},{"values":[0,0,0,0,0,0,0,2,0,0,0,0,0,0],"quality":"Less is better","isMain":false,"metricName":"# local misassemblies"},{"values":[0,0,0,0,0,0,0,0,0,0,0,0,0,0],"quality":"Less is better","isMain":false,"metricName":"# unaligned mis. contigs"}]],["Unaligned",[{"values":[0,0,0,0,0,0,0,0,0,0,0,0,0,0],"quality":"Less is better","isMain":false,"metricName":"# fully unaligned contigs"},{"values":[0,0,0,0,0,0,0,0,0,0,0,0,0,0],"quality":"Less is better","isMain":false,"metricName":"Fully unaligned length"},{"values":[0,0,0,0,0,0,0,3,0,0,0,0,0,0],"quality":"Less is better","isMain":false,"metricName":"# partially unaligned contigs"},{"values":[0,0,0,0,0,0,0,3986,0,0,0,0,0,0],"quality":"Less is better","isMain":false,"metricName":"Partially unaligned length"}]],["Mismatches",[{"values":[13,5,7,5,8,6,10,5,12,15,8,4,8,1],"quality":"Less is better","isMain":false,"metricName":"# mismatches"},{"values":[2,0,0,2,1,0,0,70,1,6,15,13,1,0],"quality":"Less is better","isMain":false,"metricName":"# indels"},{"values":[26,0,0,26,26,0,0,3291,36,29,229,172,66,0],"quality":"Less is better","isMain":false,"metricName":"Indels length"},{"values":["4.89","1.92","2.59","1.83","2.94","2.22","147.06","1.89","4.40","5.50","2.93","1.47","2.96","0.38"],"quality":"Less is better","isMain":true,"metricName":"# mismatches per 100 kbp"},{"values":["0.75","0.00","0.00","0.73","0.37","0.00","0.00","26.48","0.37","2.20","5.50","4.77","0.37","0.00"],"quality":"Less is better","isMain":true,"metricName":"# indels per 100 kbp"},{"values":[1,0,0,1,0,0,0,0,0,3,4,4,0,0],"quality":"Less is better","isMain":false,"metricName":" # indels (<= 5 bp)"},{"values":[1,0,0,1,1,0,0,70,1,3,11,9,1,0],"quality":"Less is better","isMain":false,"metricName":" # indels (> 5 bp)"},{"values":[0,0,0,0,0,41,0,6860,0,0,50,0,0,47],"quality":"Less is better","isMain":false,"metricName":"# N's"},{"values":["0.00","0.00","0.00","0.00","0.00","15.04","0.00","2508.42","0.00","0.00","18.32","0.00","0.00","17.70"],"quality":"Less is better","isMain":true,"metricName":"# N's per 100 kbp"}]],["Statistics without reference",[{"values":[67,75,20,5,6,17,11,44,3,6,7,7,29,7],"quality":"Equal","isMain":true,"metricName":"# contigs"},{"values":[59,63,19,5,6,14,0,40,3,6,7,7,29,7],"quality":"Equal","isMain":false,"metricName":"# contigs (>= 1000 bp)"},{"values":[17,15,14,5,6,9,0,19,3,6,7,7,17,6],"quality":"Equal","isMain":false,"metricName":"# contigs (>= 5000 bp)"},{"values":[4,4,11,4,6,6,0,8,3,6,7,7,13,5],"quality":"Equal","isMain":false,"metricName":"# contigs (>= 10000 bp)"},{"values":[1,0,2,4,5,6,0,2,3,5,5,5,2,5],"quality":"Equal","isMain":false,"metricName":"# contigs (>= 25000 bp)"},{"values":[0,0,1,2,2,1,0,0,2,3,2,2,0,2],"quality":"Equal","isMain":false,"metricName":"# contigs (>= 50000 bp)"},{"values":[33856,15465,69237,141523,83830,70325,795,27668,141523,68435,68356,68366,26365,71616],"quality":"More is better","isMain":true,"metricName":"Largest contig"},{"values":[266241,260244,272517,272709,272914,272555,6833,273479,272590,272606,272924,272840,269932,265500],"quality":"More is better","isMain":true,"metricName":"Total length"},{"values":[260638,251271,271603,272709,272914,270733,0,270582,272590,272606,272924,272840,269932,265500],"quality":"More is better","isMain":true,"metricName":"Total length (>= 1000 bp)"},{"values":[149494,130879,258786,272709,272914,258593,0,213057,272590,272606,272924,272840,238885,261810],"quality":"More is better","isMain":false,"metricName":"Total length (>= 5000 bp)"},{"values":[65682,52120,236523,264664,272914,240261,0,137289,272590,272606,272924,272840,206619,254390],"quality":"More is better","isMain":true,"metricName":"Total length (>= 10000 bp)"},{"values":[33856,0,102392,264664,250304,240261,0,54251,272590,250819,240730,240662,52290,254390],"quality":"More is better","isMain":false,"metricName":"Total length (>= 25000 bp)"},{"values":[0,0,69237,201340,143668,70325,0,0,227329,179542,119721,119709,0,141941],"quality":"More is better","isMain":true,"metricName":"Total length (>= 50000 bp)"},{"values":[5746,5454,18854,141523,59838,39173,654,10921,141523,51337,49772,49725,14381,70325],"quality":"More is better","isMain":false,"metricName":"N50"},{"values":[3128,2956,13733,37264,35331,27399,528,5247,85806,45261,45254,45245,10251,40220],"quality":"More is better","isMain":false,"metricName":"N75"},{"values":[14,15,4,1,2,3,5,8,1,3,3,3,8,2],"quality":"Less is better","isMain":false,"metricName":"L50"},{"values":[31,32,9,3,4,5,8,18,2,4,4,4,13,4],"quality":"Less is better","isMain":false,"metricName":"L75"},{"values":["60.01","60.06","59.98","59.95","59.94","59.96","55.88","60.07","59.95","59.95","59.91","59.92","60.05","59.94"],"quality":"Equal","isMain":false,"metricName":"GC (%)"}]],["Predicted genes",[]],["Similarity statistics",[{"values":[0,0,0,0,0,0,0,0,0,0,0,0,0,0],"quality":"Equal","isMain":false,"metricName":"# similar correct contigs"},{"values":[0,0,0,0,0,0,0,0,0,0,0,0,0,0],"quality":"Equal","isMain":false,"metricName":"# similar misassembled blocks"}]],["Reference statistics",[{"values":[295146,295146,295146,295146,295146,295146,295146,295146,295146,295146,295146,295146,295146,295146],"quality":"Equal","isMain":false,"metricName":"Reference length"},{"values":[1,1,1,1,1,1,1,1,1,1,1,1,1,1],"quality":"Equal","isMain":false,"metricName":"Reference fragments"},{"values":["59.23","59.23","59.23","59.23","59.23","59.23","59.23","59.23","59.23","59.23","59.23","59.23","59.23","59.23"],"quality":"Equal","isMain":false,"metricName":"Reference GC (%)"}]]],"referenceName":"gi\_131840030\_ref\_NC","date":"18 July 2018, Wednesday, 16:16:39","order":[0,1,2,3,4,5,6,7,8,9,10,11,12,13],"assembliesNames":["ABySS\_127","ABySS\_63","CLC","IDBA\_UD","MEGAHIT","MIRA","Ray\_Meta","SOAPdenovo2","SPAdes","SPAdes\_meta","SPAdes\_sc","SPAdes\_sc\_careful","Velvet","Geneious"]},{"assembliesWithNs":null,"minContig":500,"report":[["Genome statistics",[{"values":["86.896","80.419","89.270","89.378","90.072","95.275","0.539","82.134","93.575","89.799","88.836","88.346","85.508","68.262"],"quality":"More is better","isMain":true,"metricName":"Genome fraction (%)"},{"values":["1.004","1.003","1.007","1.002","1.004","1.067","1.000","1.027","1.003","1.001","1.002","1.002","1.001","1.009"],"quality":"Less is better","isMain":true,"metricName":"Duplication ratio"},{"values":[9579,7964,27271,32637,33291,25784,772,14845,44655,44691,32566,33150,15092,49498],"quality":"More is better","isMain":true,"metricName":"Largest alignment"},{"values":[292828,270720,301534,300512,303235,340621,1808,277102,314788,301524,298726,296973,287218,230660],"quality":"More is better","isMain":true,"metricName":"Total aligned length"},{"values":[2778,2050,12003,11676,12252,10343,null,4499,21466,15076,11987,11488,7161,10946],"quality":"More is better","isMain":false,"metricName":"NG50"},{"values":[1351,853,6324,6688,8699,5330,null,2418,12111,10759,9017,9017,3316,null],"quality":"More is better","isMain":false,"metricName":"NG75"},{"values":[3060,2691,12006,12211,12497,10343,522,4805,15092,16197,12136,12136,9571,18530],"quality":"More is better","isMain":false,"metricName":"NA50"},{"values":[1935,1645,9587,9640,9649,5330,514,3493,11310,11310,10629,9972,4746,9826],"quality":"More is better","isMain":false,"metricName":"NA75"},{"values":[2778,2050,12003,11676,12252,10343,null,4204,15092,15076,11987,11488,7161,9826],"quality":"More is better","isMain":true,"metricName":"NGA50"},{"values":[1351,853,6324,6688,8699,5330,null,2084,10692,10551,6688,6688,3316,null],"quality":"More is better","isMain":false,"metricName":"NGA75"},{"values":[39,49,10,10,9,12,null,23,6,8,10,10,15,7],"quality":"Less is better","isMain":false,"metricName":"LG50"},{"values":[80,108,19,18,17,24,null,47,11,14,17,17,32,null],"quality":"Less is better","isMain":false,"metricName":"LG75"},{"values":[32,35,9,8,8,12,2,18,7,6,8,8,13,4],"quality":"Less is better","isMain":false,"metricName":"LA50"},{"values":[61,69,15,15,15,24,3,36,13,12,15,15,23,8],"quality":"Less is better","isMain":false,"metricName":"LA75"},{"values":[39,49,10,10,9,12,null,23,7,8,10,10,15,8],"quality":"Less is better","isMain":true,"metricName":"LGA50"},{"values":[80,108,19,18,17,24,null,49,14,14,18,18,32,null],"quality":"Less is better","isMain":false,"metricName":"LGA75"}]],["Misassemblies",[{"values":[0,0,0,0,1,1,0,0,4,2,1,1,0,2],"quality":"Less is better","isMain":true,"metricName":"# misassemblies"},{"values":[0,0,0,0,1,1,0,0,4,2,1,1,0,2],"quality":"Less is better","isMain":false,"metricName":" # relocations"},{"values":[0,0,0,0,0,0,0,0,0,0,0,0,0,0],"quality":"Less is better","isMain":false,"metricName":" # translocations"},{"values":[0,0,0,0,0,0,0,0,0,0,0,0,0,0],"quality":"Less is better","isMain":false,"metricName":" # inversions"},{"values":[0,0,0,0,1,1,0,0,2,2,1,1,0,2],"quality":"Less is better","isMain":false,"metricName":"# misassembled contigs"},{"values":[0,0,0,0,36772,13459,0,0,61263,22395,11482,11482,0,54268],"quality":"Less is better","isMain":true,"metricName":"Misassembled contigs length"},{"values":[0,0,2,0,3,8,0,1,2,2,5,5,1,7],"quality":"Less is better","isMain":false,"metricName":"# local misassemblies"},{"values":[0,0,0,0,0,0,0,2,0,0,0,0,0,0],"quality":"Less is better","isMain":false,"metricName":"# unaligned mis. contigs"}]],["Unaligned",[{"values":[0,0,0,0,0,0,0,0,0,0,0,0,0,0],"quality":"Less is better","isMain":false,"metricName":"# fully unaligned contigs"},{"values":[0,0,0,0,0,0,0,0,0,0,0,0,0,0],"quality":"Less is better","isMain":false,"metricName":"Fully unaligned length"},{"values":[0,0,0,0,0,0,0,5,0,0,0,0,0,1],"quality":"Less is better","isMain":false,"metricName":"# partially unaligned contigs"},{"values":[0,0,0,0,0,0,0,4289,0,0,0,0,0,590],"quality":"Less is better","isMain":false,"metricName":"Partially unaligned length"}]],["Mismatches",[{"values":[22,35,65,32,57,297,1,20,136,124,78,46,27,296],"quality":"Less is better","isMain":false,"metricName":"# mismatches"},{"values":[0,0,5,0,0,40,0,77,13,8,8,4,2,68],"quality":"Less is better","isMain":false,"metricName":"# indels"},{"values":[0,0,7,0,0,186,0,3772,32,69,132,85,11,443],"quality":"Less is better","isMain":false,"metricName":"Indels length"},{"values":["7.54","12.97","21.70","10.67","18.86","92.89","55.31","7.26","43.31","41.15","26.16","15.52","9.41","129.21"],"quality":"Less is better","isMain":true,"metricName":"# mismatches per 100 kbp"},{"values":["0.00","0.00","1.67","0.00","0.00","12.51","0.00","27.94","4.14","2.65","2.68","1.35","0.70","29.68"],"quality":"Less is better","isMain":true,"metricName":"# indels per 100 kbp"},{"values":[0,0,5,0,0,34,0,0,12,5,2,1,1,51],"quality":"Less is better","isMain":false,"metricName":" # indels (<= 5 bp)"},{"values":[0,0,0,0,0,6,0,77,1,3,6,3,1,17],"quality":"Less is better","isMain":false,"metricName":" # indels (> 5 bp)"},{"values":[0,0,0,0,0,99,0,8970,100,110,30,0,10,264],"quality":"Less is better","isMain":false,"metricName":"# N's"},{"values":["0.00","0.00","0.00","0.00","0.00","29.03","0.00","3120.15","31.76","36.47","10.04","0.00","3.48","113.94"],"quality":"Less is better","isMain":true,"metricName":"# N's per 100 kbp"}]],["Statistics without reference",[{"values":[131,137,36,38,35,63,3,72,24,24,34,33,56,17],"quality":"Equal","isMain":true,"metricName":"# contigs"},{"values":[92,99,32,30,28,50,0,64,19,23,29,29,44,17],"quality":"Equal","isMain":false,"metricName":"# contigs (>= 1000 bp)"},{"values":[7,4,23,20,21,27,0,17,17,18,20,20,22,14],"quality":"Equal","isMain":false,"metricName":"# contigs (>= 5000 bp)"},{"values":[0,0,13,14,14,12,0,6,13,16,16,15,11,7],"quality":"Equal","isMain":false,"metricName":"# contigs (>= 10000 bp)"},{"values":[0,0,2,2,2,1,0,0,4,3,2,2,0,3],"quality":"Equal","isMain":false,"metricName":"# contigs (>= 25000 bp)"},{"values":[0,0,0,0,0,0,0,0,0,0,0,0,0,0],"quality":"Equal","isMain":false,"metricName":"# contigs (>= 50000 bp)"},{"values":[9579,7964,27271,32637,36772,25784,772,15042,44655,44711,32566,33150,15092,49498],"quality":"More is better","isMain":true,"metricName":"Largest contig"},{"values":[292828,270720,301538,300512,303606,341043,1808,287486,314888,301654,298795,297077,287324,231701],"quality":"More is better","isMain":true,"metricName":"Total length"},{"values":[265661,243725,298611,295414,298585,332047,0,281634,311588,300884,295550,294071,277750,231701],"quality":"More is better","isMain":true,"metricName":"Total length (>= 1000 bp)"},{"values":[49330,25979,278563,265765,280603,272527,0,143951,308301,287492,272681,271240,211813,220544],"quality":"More is better","isMain":false,"metricName":"Total length (>= 5000 bp)"},{"values":[0,0,207299,219730,229002,175211,0,71534,280221,274447,244183,232770,133854,176282],"quality":"More is better","isMain":true,"metricName":"Total length (>= 10000 bp)"},{"values":[0,0,52722,60438,64106,25784,0,0,138135,97665,59870,60454,0,121094],"quality":"More is better","isMain":false,"metricName":"Total length (>= 25000 bp)"},{"values":[0,0,0,0,0,0,0,0,0,0,0,0,0,0],"quality":"More is better","isMain":true,"metricName":"Total length (>= 50000 bp)"},{"values":[3060,2691,12006,12211,12497,10343,522,5074,23352,16197,12136,12136,9828,30787],"quality":"More is better","isMain":false,"metricName":"N50"},{"values":[1935,1645,9587,9640,10757,5330,514,3588,15076,12039,11091,10629,4746,10946],"quality":"More is better","isMain":false,"metricName":"N75"},{"values":[32,35,9,8,8,12,2,17,5,6,8,8,12,3],"quality":"Less is better","isMain":false,"metricName":"L50"},{"values":[61,69,15,15,14,24,3,35,10,12,15,15,23,7],"quality":"Less is better","isMain":false,"metricName":"L75"},{"values":["51.81","51.84","51.86","51.80","51.85","51.78","55.03","51.81","51.77","51.85","51.87","51.85","51.85","51.32"],"quality":"Equal","isMain":false,"metricName":"GC (%)"}]],["Predicted genes",[]],["Similarity statistics",[{"values":[0,0,2,2,2,0,0,0,2,2,2,2,2,0],"quality":"Equal","isMain":false,"metricName":"# similar correct contigs"},{"values":[0,0,0,0,0,0,0,0,0,0,0,0,0,0],"quality":"Equal","isMain":false,"metricName":"# similar misassembled blocks"}]],["Reference statistics",[{"values":[335593,335593,335593,335593,335593,335593,335593,335593,335593,335593,335593,335593,335593,335593],"quality":"Equal","isMain":false,"metricName":"Reference length"},{"values":[1,1,1,1,1,1,1,1,1,1,1,1,1,1],"quality":"Equal","isMain":false,"metricName":"Reference fragments"},{"values":["51.73","51.73","51.73","51.73","51.73","51.73","51.73","51.73","51.73","51.73","51.73","51.73","51.73","51.73"],"quality":"Equal","isMain":false,"metricName":"Reference GC (%)"}]]],"referenceName":"gi\_13242472\_ref\_NC","date":"18 July 2018, Wednesday, 16:16:53","order":[0,1,2,3,4,5,6,7,8,9,10,11,12,13],"assembliesNames":["ABySS\_127","ABySS\_63","CLC","IDBA\_UD","MEGAHIT","MIRA","Ray\_Meta","SOAPdenovo2","SPAdes","SPAdes\_meta","SPAdes\_sc","SPAdes\_sc\_careful","Velvet","Geneious"]},{"assembliesWithNs":null,"minContig":500,"report":[["Genome statistics",[{"values":["23.160","14.281","77.938","74.415","75.057","40.769","71.432","79.340","80.934","81.164","81.164","9.088","5.363"],"quality":"More is better","isMain":true,"metricName":"Genome fraction (%)"},{"values":["1.012","1.000","1.010","1.010","1.008","1.015","1.002","1.010","1.005","1.007","1.007","1.000","1.014"],"quality":"Less is better","isMain":true,"metricName":"Duplication ratio"},{"values":[2480,1999,8195,8195,7935,3208,4927,8195,8195,8195,8195,2159,3209],"quality":"More is better","isMain":true,"metricName":"Largest alignment"},{"values":[43390,26601,146122,138933,140275,76750,133228,148003,151094,151427,151427,16927,9991],"quality":"More is better","isMain":true,"metricName":"Total aligned length"},{"values":[null,null,1600,1619,1513,null,1093,1911,1901,1619,1619,null,null],"quality":"More is better","isMain":false,"metricName":"NG50"},{"values":[null,null,614,537,531,null,null,769,831,717,717,null,null],"quality":"More is better","isMain":false,"metricName":"NG75"},{"values":[810,812,2108,2108,2148,1325,1767,2290,2198,2183,2183,1319,2337],"quality":"More is better","isMain":false,"metricName":"NA50"},{"values":[618,629,1074,1146,933,882,1016,1239,1146,1086,1086,767,1685],"quality":"More is better","isMain":false,"metricName":"NA75"},{"values":[null,null,1597,1616,1513,null,1093,1874,1851,1619,1619,null,null],"quality":"More is better","isMain":true,"metricName":"NGA50"},{"values":[null,null,601,null,509,null,null,717,740,699,699,null,null],"quality":"More is better","isMain":false,"metricName":"NGA75"},{"values":[null,null,33,32,31,null,45,28,29,32,32,null,null],"quality":"Less is better","isMain":false,"metricName":"LG50"},{"values":[null,null,81,82,86,null,null,67,69,75,75,null,null],"quality":"Less is better","isMain":false,"metricName":"LG75"},{"values":[18,12,22,20,20,19,26,20,22,23,23,5,2],"quality":"Less is better","isMain":false,"metricName":"LA50"},{"values":[33,21,46,42,44,37,51,41,45,48,48,10,4],"quality":"Less is better","isMain":false,"metricName":"LA75"},{"values":[null,null,33,33,32,null,45,29,30,32,32,null,null],"quality":"Less is better","isMain":true,"metricName":"LGA50"},{"values":[null,null,82,null,92,null,null,70,71,76,76,null,null],"quality":"Less is better","isMain":false,"metricName":"LGA75"}]],["Misassemblies",[{"values":[0,0,0,3,3,1,0,3,1,0,0,0,1],"quality":"Less is better","isMain":true,"metricName":"# misassemblies"},{"values":[0,0,0,3,3,1,0,3,1,0,0,0,1],"quality":"Less is better","isMain":false,"metricName":" # relocations"},{"values":[0,0,0,0,0,0,0,0,0,0,0,0,0],"quality":"Less is better","isMain":false,"metricName":" # translocations"},{"values":[0,0,0,0,0,0,0,0,0,0,0,0,0],"quality":"Less is better","isMain":false,"metricName":" # inversions"},{"values":[0,0,0,3,3,1,0,2,1,0,0,0,1],"quality":"Less is better","isMain":false,"metricName":"# misassembled contigs"},{"values":[0,0,0,8409,8320,3753,0,7466,5934,0,0,0,3754],"quality":"Less is better","isMain":true,"metricName":"Misassembled contigs length"},{"values":[0,0,0,0,0,0,0,1,0,0,0,0,0],"quality":"Less is better","isMain":false,"metricName":"# local misassemblies"},{"values":[0,0,0,0,0,0,0,0,0,0,0,0,0],"quality":"Less is better","isMain":false,"metricName":"# unaligned mis. contigs"}]],["Unaligned",[{"values":[0,0,0,0,0,0,0,0,0,0,0,0,0],"quality":"Less is better","isMain":false,"metricName":"# fully unaligned contigs"},{"values":[0,0,0,0,0,0,0,0,0,0,0,0,0],"quality":"Less is better","isMain":false,"metricName":"Fully unaligned length"},{"values":[0,0,0,0,1,0,0,0,1,0,0,0,1],"quality":"Less is better","isMain":false,"metricName":"# partially unaligned contigs"},{"values":[0,0,0,0,1356,0,0,0,718,0,0,0,938],"quality":"Less is better","isMain":false,"metricName":"Partially unaligned length"}]],["Mismatches",[{"values":[30,30,161,119,139,148,75,154,145,163,149,8,67],"quality":"Less is better","isMain":false,"metricName":"# mismatches"},{"values":[0,0,0,2,4,5,2,0,1,2,2,0,3],"quality":"Less is better","isMain":false,"metricName":"# indels"},{"values":[0,0,0,2,60,20,103,0,1,58,58,0,3],"quality":"Less is better","isMain":false,"metricName":"Indels length"},{"values":["69.54","112.78","110.91","85.85","99.43","194.90","56.37","104.21","96.19","107.82","98.56","47.26","670.67"],"quality":"Less is better","isMain":true,"metricName":"# mismatches per 100 kbp"},{"values":["0.00","0.00","0.00","1.44","2.86","6.58","1.50","0.00","0.66","1.32","1.32","0.00","30.03"],"quality":"Less is better","isMain":true,"metricName":"# indels per 100 kbp"},{"values":[0,0,0,2,3,4,0,0,1,1,1,0,3],"quality":"Less is better","isMain":false,"metricName":" # indels (<= 5 bp)"},{"values":[0,0,0,0,1,1,2,0,0,1,1,0,0],"quality":"Less is better","isMain":false,"metricName":" # indels (> 5 bp)"},{"values":[0,0,0,0,0,94,103,0,0,0,0,0,130],"quality":"Less is better","isMain":false,"metricName":"# N's"},{"values":["0.00","0.00","0.00","0.00","0.00","121.94","77.25","0.00","0.00","0.00","0.00","0.00","1174.35"],"quality":"Less is better","isMain":true,"metricName":"# N's per 100 kbp"}]],["Statistics without reference",[{"values":[52,32,93,82,90,64,97,82,88,95,95,17,5],"quality":"Equal","isMain":true,"metricName":"# contigs"},{"values":[14,5,52,50,43,32,52,54,55,55,55,5,5],"quality":"Equal","isMain":false,"metricName":"# contigs (>= 1000 bp)"},{"values":[0,0,2,3,3,0,0,4,3,2,2,0,0],"quality":"Equal","isMain":false,"metricName":"# contigs (>= 5000 bp)"},{"values":[0,0,0,0,0,0,0,0,0,0,0,0,0],"quality":"Equal","isMain":false,"metricName":"# contigs (>= 10000 bp)"},{"values":[0,0,0,0,0,0,0,0,0,0,0,0,0],"quality":"Equal","isMain":false,"metricName":"# contigs (>= 25000 bp)"},{"values":[0,0,0,0,0,0,0,0,0,0,0,0,0],"quality":"Equal","isMain":false,"metricName":"# contigs (>= 50000 bp)"},{"values":[2643,1999,8195,8195,7935,3753,4927,8195,8195,8195,8195,2159,3754],"quality":"More is better","isMain":true,"metricName":"Largest contig"},{"values":[43675,26601,146593,139934,142307,77084,133333,149308,152221,152168,152168,16927,11070],"quality":"More is better","isMain":true,"metricName":"Total length"},{"values":[19192,7803,116847,116318,109420,54348,101184,128564,127533,122730,122730,9021,11070],"quality":"More is better","isMain":true,"metricName":"Total length (>= 1000 bp)"},{"values":[0,0,14153,19932,19460,0,0,26037,20087,14153,14153,0,0],"quality":"More is better","isMain":false,"metricName":"Total length (>= 5000 bp)"},{"values":[0,0,0,0,0,0,0,0,0,0,0,0,0],"quality":"More is better","isMain":true,"metricName":"Total length (>= 10000 bp)"},{"values":[0,0,0,0,0,0,0,0,0,0,0,0,0],"quality":"More is better","isMain":false,"metricName":"Total length (>= 25000 bp)"},{"values":[0,0,0,0,0,0,0,0,0,0,0,0,0],"quality":"More is better","isMain":true,"metricName":"Total length (>= 50000 bp)"},{"values":[895,812,2108,2108,2201,1325,1767,2290,2198,2183,2183,1319,2337],"quality":"More is better","isMain":false,"metricName":"N50"},{"values":[618,629,1086,1225,1069,934,1016,1366,1164,1124,1124,767,1685],"quality":"More is better","isMain":false,"metricName":"N75"},{"values":[17,12,22,19,19,19,26,19,21,23,23,5,2],"quality":"Less is better","isMain":false,"metricName":"L50"},{"values":[33,21,46,40,41,36,51,40,43,47,47,10,4],"quality":"Less is better","isMain":false,"metricName":"L75"},{"values":["46.10","45.04","45.56","45.82","45.68","46.36","45.48","45.70","45.67","45.64","45.63","45.86","48.83"],"quality":"Equal","isMain":false,"metricName":"GC (%)"}]],["Predicted genes",[]],["Similarity statistics",[{"values":[0,0,0,0,0,0,0,0,0,0,0,0,0],"quality":"Equal","isMain":false,"metricName":"# similar correct contigs"},{"values":[0,0,0,0,0,0,0,0,0,0,0,0,0],"quality":"Equal","isMain":false,"metricName":"# similar misassembled blocks"}]],["Reference statistics",[{"values":[186262,186262,186262,186262,186262,186262,186262,186262,186262,186262,186262,186262,186262],"quality":"Equal","isMain":false,"metricName":"Reference length"},{"values":[1,1,1,1,1,1,1,1,1,1,1,1,1],"quality":"Equal","isMain":false,"metricName":"Reference fragments"},{"values":["45.87","45.87","45.87","45.87","45.87","45.87","45.87","45.87","45.87","45.87","45.87","45.87","45.87"],"quality":"Equal","isMain":false,"metricName":"Reference GC (%)"}]]],"referenceName":"gi\_134287156\_ref\_NC","date":"18 July 2018, Wednesday, 16:17:05","order":[0,1,2,3,4,5,6,7,8,9,10,11,12],"assembliesNames":["ABySS\_127","ABySS\_63","CLC","IDBA\_UD","MEGAHIT","MIRA","SOAPdenovo2","SPAdes","SPAdes\_meta","SPAdes\_sc","SPAdes\_sc\_careful","Velvet","Geneious"]},{"assembliesWithNs":null,"minContig":500,"report":[["Genome statistics",[{"values":["45.424","69.134","94.338","92.143","89.071","100.000","93.392","98.691","98.997","90.613","91.323","76.310","100.000"],"quality":"More is better","isMain":true,"metricName":"Genome fraction (%)"},{"values":["1.017","1.042","1.042","1.013","1.022","3.336","1.032","1.031","1.002","1.018","1.019","1.001","2.465"],"quality":"Less is better","isMain":true,"metricName":"Duplication ratio"},{"values":[3312,13586,20819,20725,21057,47416,28391,23126,51099,20565,20565,10670,59054],"quality":"More is better","isMain":true,"metricName":"Largest alignment"},{"values":[26129,40205,55501,52749,51445,188541,54467,57449,56099,52135,52591,43174,139350],"quality":"More is better","isMain":true,"metricName":"Total aligned length"},{"values":[null,8819,10135,4214,5452,116371,28391,10661,51099,10517,10517,4022,213470],"quality":"More is better","isMain":false,"metricName":"NG50"},{"values":[null,2345,1740,1554,1554,116371,4868,10345,51099,2706,2706,521,213470],"quality":"More is better","isMain":false,"metricName":"NG75"},{"values":[884,2513,10135,4214,5974,2690,28391,10661,51099,10517,10517,4509,6611],"quality":"More is better","isMain":false,"metricName":"NA50"},{"values":[669,119,1740,1809,2165,null,4868,10345,null,4207,9596,1266,null],"quality":"More is better","isMain":false,"metricName":"NA75"},{"values":[null,2153,10135,4214,5452,47416,28391,10661,51099,10517,10517,4022,59054],"quality":"More is better","isMain":true,"metricName":"NGA50"},{"values":[null,null,1740,1554,1554,47416,4868,10345,51099,2706,2706,521,59054],"quality":"More is better","isMain":false,"metricName":"NGA75"},{"values":[null,3,2,3,3,1,1,2,1,2,2,4,1],"quality":"Less is better","isMain":false,"metricName":"LG50"},{"values":[null,5,5,8,7,1,3,3,1,5,4,19,1],"quality":"Less is better","isMain":false,"metricName":"LG75"},{"values":[7,4,2,3,2,10,1,2,1,2,2,3,3],"quality":"Less is better","isMain":false,"metricName":"LA50"},{"values":[16,17,5,7,5,null,3,3,null,4,3,7,null],"quality":"Less is better","isMain":false,"metricName":"LA75"},{"values":[null,5,2,3,3,1,1,2,1,2,2,4,1],"quality":"Less is better","isMain":true,"metricName":"LGA50"},{"values":[null,null,5,8,7,1,3,3,1,5,4,19,1],"quality":"Less is better","isMain":false,"metricName":"LGA75"}]],["Misassemblies",[{"values":[0,0,0,0,0,7,0,0,0,0,0,0,7],"quality":"Less is better","isMain":true,"metricName":"# misassemblies"},{"values":[0,0,0,0,0,7,0,0,0,0,0,0,7],"quality":"Less is better","isMain":false,"metricName":" # relocations"},{"values":[0,0,0,0,0,0,0,0,0,0,0,0,0],"quality":"Less is better","isMain":false,"metricName":" # translocations"},{"values":[0,0,0,0,0,0,0,0,0,0,0,0,0],"quality":"Less is better","isMain":false,"metricName":" # inversions"},{"values":[0,0,0,0,0,3,0,0,0,0,0,0,1],"quality":"Less is better","isMain":false,"metricName":"# misassembled contigs"},{"values":[0,0,0,0,0,211022,0,0,0,0,0,0,213470],"quality":"Less is better","isMain":true,"metricName":"Misassembled contigs length"},{"values":[0,1,0,0,0,7,0,1,0,0,0,0,7],"quality":"Less is better","isMain":false,"metricName":"# local misassemblies"},{"values":[0,2,0,0,0,0,0,0,0,0,0,0,0],"quality":"Less is better","isMain":false,"metricName":"# unaligned mis. contigs"}]],["Unaligned",[{"values":[0,0,0,0,0,0,0,0,0,0,0,0,0],"quality":"Less is better","isMain":false,"metricName":"# fully unaligned contigs"},{"values":[0,0,0,0,0,0,0,0,0,0,0,0,0],"quality":"Less is better","isMain":false,"metricName":"Fully unaligned length"},{"values":[0,2,0,0,0,3,0,0,1,0,0,0,2],"quality":"Less is better","isMain":false,"metricName":"# partially unaligned contigs"},{"values":[0,12802,0,0,0,79220,0,0,25270,0,0,0,79242],"quality":"Less is better","isMain":false,"metricName":"Partially unaligned length"}]],["Mismatches",[{"values":[0,0,17,0,11,68,0,0,36,57,80,8,54],"quality":"Less is better","isMain":false,"metricName":"# mismatches"},{"values":[0,2,0,0,3,4,0,0,4,5,5,0,4],"quality":"Less is better","isMain":false,"metricName":"# indels"},{"values":[0,2,0,0,7,16,0,0,4,6,6,0,12],"quality":"Less is better","isMain":false,"metricName":"Indels length"},{"values":["0.00","0.00","31.87","0.00","21.84","120.27","0.00","0.00","64.32","111.26","154.94","18.54","95.51"],"quality":"Less is better","isMain":true,"metricName":"# mismatches per 100 kbp"},{"values":["0.00","5.12","0.00","0.00","5.96","7.07","0.00","0.00","7.15","9.76","9.68","0.00","7.07"],"quality":"Less is better","isMain":true,"metricName":"# indels per 100 kbp"},{"values":[0,2,0,0,3,3,0,0,4,5,5,0,3],"quality":"Less is better","isMain":false,"metricName":" # indels (<= 5 bp)"},{"values":[0,0,0,0,0,1,0,0,0,0,0,0,1],"quality":"Less is better","isMain":false,"metricName":" # indels (> 5 bp)"},{"values":[0,1385,0,0,0,82,0,100,0,0,0,0,232],"quality":"Less is better","isMain":false,"metricName":"# N's"},{"values":["0.00","2588.20","0.00","0.00","0.00","30.62","0.00","173.76","0.00","0.00","0.00","0.00","106.13"],"quality":"Less is better","isMain":true,"metricName":"# N's per 100 kbp"}]],["Statistics without reference",[{"values":[26,15,20,21,16,47,7,16,3,14,14,20,2],"quality":"Equal","isMain":true,"metricName":"# contigs"},{"values":[5,7,8,10,11,35,7,9,3,9,7,9,2],"quality":"Equal","isMain":false,"metricName":"# contigs (>= 1000 bp)"},{"values":[0,4,3,2,3,3,2,3,2,2,3,2,2],"quality":"Equal","isMain":false,"metricName":"# contigs (>= 5000 bp)"},{"values":[0,2,2,1,1,3,2,3,2,2,2,1,1],"quality":"Equal","isMain":false,"metricName":"# contigs (>= 10000 bp)"},{"values":[0,0,0,0,0,3,1,0,2,0,0,0,1],"quality":"Equal","isMain":false,"metricName":"# contigs (>= 25000 bp)"},{"values":[0,0,0,0,0,2,0,0,1,0,0,0,1],"quality":"Equal","isMain":false,"metricName":"# contigs (>= 50000 bp)"},{"values":[3312,13773,20819,20725,21057,116371,28391,23226,51099,20565,20565,10670,213470],"quality":"More is better","isMain":true,"metricName":"Largest contig"},{"values":[26129,53512,55591,52749,51445,267819,54467,57549,81369,52135,52607,43177,218592],"quality":"More is better","isMain":true,"metricName":"Total length"},{"values":[11460,47407,47228,44783,47273,258855,54467,51912,81369,48926,48167,35718,218592],"quality":"More is better","isMain":true,"metricName":"Total length (>= 1000 bp)"},{"values":[0,41386,36690,25989,32483,211022,38785,44232,79378,31082,40694,20071,218592],"quality":"More is better","isMain":false,"metricName":"Total length (>= 5000 bp)"},{"values":[0,24727,30954,20725,21057,211022,38785,44232,79378,31082,31082,10670,213470],"quality":"More is better","isMain":true,"metricName":"Total length (>= 10000 bp)"},{"values":[0,0,0,0,0,211022,28391,0,79378,0,0,0,213470],"quality":"More is better","isMain":false,"metricName":"Total length (>= 25000 bp)"},{"values":[0,0,0,0,0,178762,0,0,51099,0,0,0,213470],"quality":"More is better","isMain":true,"metricName":"Total length (>= 50000 bp)"},{"values":[884,8819,10135,4214,5974,62391,28391,10661,51099,10517,10517,4509,213470],"quality":"More is better","isMain":false,"metricName":"N50"},{"values":[669,7840,1740,1809,2165,32260,4868,10345,28279,4207,9612,1266,213470],"quality":"More is better","isMain":false,"metricName":"N75"},{"values":[7,3,2,3,2,2,1,2,1,2,2,3,1],"quality":"Less is better","isMain":false,"metricName":"L50"},{"values":[16,4,5,7,5,3,3,3,2,4,3,7,1],"quality":"Less is better","isMain":false,"metricName":"L75"},{"values":["28.68","28.65","28.75","28.71","29.10","28.89","28.62","28.65","29.18","28.69","28.77","28.69","28.90"],"quality":"Equal","isMain":false,"metricName":"GC (%)"}]],["Predicted genes",[]],["Similarity statistics",[{"values":[0,0,0,0,0,0,0,0,0,0,0,0,0],"quality":"Equal","isMain":false,"metricName":"# similar correct contigs"},{"values":[0,0,0,0,0,0,0,0,0,0,0,0,0],"quality":"Equal","isMain":false,"metricName":"# similar misassembled blocks"}]],["Reference statistics",[{"values":[56538,56538,56538,56538,56538,56538,56538,56538,56538,56538,56538,56538,56538],"quality":"Equal","isMain":false,"metricName":"Reference length"},{"values":[1,1,1,1,1,1,1,1,1,1,1,1,1],"quality":"Equal","isMain":false,"metricName":"Reference fragments"},{"values":["28.73","28.73","28.73","28.73","28.73","28.73","28.73","28.73","28.73","28.73","28.73","28.73","28.73"],"quality":"Equal","isMain":false,"metricName":"Reference GC (%)"}]]],"referenceName":"gi\_134287335\_ref\_NC","date":"18 July 2018, Wednesday, 16:17:16","order":[0,1,2,3,4,5,6,7,8,9,10,11,12],"assembliesNames":["ABySS\_127","ABySS\_63","CLC","IDBA\_UD","MEGAHIT","MIRA","Ray\_Meta","SPAdes","SPAdes\_meta","SPAdes\_sc","SPAdes\_sc\_careful","Velvet","Geneious"]},{"assembliesWithNs":null,"minContig":500,"report":[["Genome statistics",[{"values":["43.486","40.472","68.763","45.078","58.962","81.376","17.582","31.842","56.462","40.033","36.421","42.832","39.095","100.000"],"quality":"More is better","isMain":true,"metricName":"Genome fraction (%)"},{"values":["1.008","1.000","1.016","1.005","1.013","1.092","1.001","1.053","1.000","1.000","1.000","1.000","1.000","1.369"],"quality":"Less is better","isMain":true,"metricName":"Duplication ratio"},{"values":[13676,8664,13967,8825,13853,15792,1267,7343,14209,13314,8575,8575,8467,32344],"quality":"More is better","isMain":true,"metricName":"Largest alignment"},{"values":[16076,14854,25645,16618,21915,32602,6458,11901,20716,14688,13363,15715,14344,50235],"quality":"More is better","isMain":true,"metricName":"Total aligned length"},{"values":[null,null,2504,null,2026,8830,null,null,785,null,null,null,null,63637],"quality":"More is better","isMain":false,"metricName":"NG50"},{"values":[null,null,null,null,null,6851,null,null,null,null,null,null,null,63637],"quality":"More is better","isMain":false,"metricName":"NG75"},{"values":[13676,8664,13967,8825,13853,8830,670,7343,14209,13314,8575,8575,8467,32344],"quality":"More is better","isMain":false,"metricName":"NA50"},{"values":[13676,4802,1576,4876,3936,8830,579,4558,1809,13314,4788,4788,4656,17891],"quality":"More is better","isMain":false,"metricName":"NA75"},{"values":[null,null,2504,null,2026,8830,null,null,785,null,null,null,null,32344],"quality":"More is better","isMain":true,"metricName":"NGA50"},{"values":[null,null,null,null,null,6850,null,null,null,null,null,null,null,32344],"quality":"More is better","isMain":false,"metricName":"NGA75"},{"values":[null,null,3,null,3,2,null,null,5,null,null,null,null,1],"quality":"Less is better","isMain":false,"metricName":"LG50"},{"values":[null,null,null,null,null,3,null,null,null,null,null,null,null,1],"quality":"Less is better","isMain":false,"metricName":"LG75"},{"values":[1,1,1,1,1,2,4,1,1,1,1,1,1,1],"quality":"Less is better","isMain":false,"metricName":"LA50"},{"values":[1,2,4,2,2,2,7,2,2,1,2,2,2,2],"quality":"Less is better","isMain":false,"metricName":"LA75"},{"values":[null,null,3,null,3,2,null,null,5,null,null,null,null,1],"quality":"Less is better","isMain":true,"metricName":"LGA50"},{"values":[null,null,null,null,null,3,null,null,null,null,null,null,null,1],"quality":"Less is better","isMain":false,"metricName":"LGA75"}]],["Misassemblies",[{"values":[0,0,0,0,0,0,0,0,0,0,0,0,0,1],"quality":"Less is better","isMain":true,"metricName":"# misassemblies"},{"values":[0,0,0,0,0,0,0,0,0,0,0,0,0,1],"quality":"Less is better","isMain":false,"metricName":" # relocations"},{"values":[0,0,0,0,0,0,0,0,0,0,0,0,0,0],"quality":"Less is better","isMain":false,"metricName":" # translocations"},{"values":[0,0,0,0,0,0,0,0,0,0,0,0,0,0],"quality":"Less is better","isMain":false,"metricName":" # inversions"},{"values":[0,0,0,0,0,0,0,0,0,0,0,0,0,1],"quality":"Less is better","isMain":false,"metricName":"# misassembled contigs"},{"values":[0,0,0,0,0,0,0,0,0,0,0,0,0,63637],"quality":"Less is better","isMain":true,"metricName":"Misassembled contigs length"},{"values":[0,0,0,0,0,0,0,0,0,0,0,0,0,2],"quality":"Less is better","isMain":false,"metricName":"# local misassemblies"},{"values":[0,0,0,0,0,0,0,0,0,0,0,0,0,0],"quality":"Less is better","isMain":false,"metricName":"# unaligned mis. contigs"}]],["Unaligned",[{"values":[0,0,0,0,0,0,0,0,0,0,0,0,0,0],"quality":"Less is better","isMain":false,"metricName":"# fully unaligned contigs"},{"values":[0,0,0,0,0,0,0,0,0,0,0,0,0,0],"quality":"Less is better","isMain":false,"metricName":"Fully unaligned length"},{"values":[0,0,0,0,0,0,0,1,0,0,0,0,0,1],"quality":"Less is better","isMain":false,"metricName":"# partially unaligned contigs"},{"values":[0,0,0,0,0,0,0,1705,0,0,0,0,0,13402],"quality":"Less is better","isMain":false,"metricName":"Partially unaligned length"}]],["Mismatches",[{"values":[0,0,5,3,31,51,0,3,4,0,1,0,0,79],"quality":"Less is better","isMain":false,"metricName":"# mismatches"},{"values":[0,0,0,0,0,0,0,13,0,0,0,0,0,0],"quality":"Less is better","isMain":false,"metricName":"# indels"},{"values":[0,0,0,0,0,0,0,605,0,0,0,0,0,0],"quality":"Less is better","isMain":false,"metricName":"Indels length"},{"values":["0.00","0.00","19.82","18.14","143.30","170.81","0.00","25.68","19.31","0.00","7.48","0.00","0.00","215.32"],"quality":"Less is better","isMain":true,"metricName":"# mismatches per 100 kbp"},{"values":["0.00","0.00","0.00","0.00","0.00","0.00","0.00","111.27","0.00","0.00","0.00","0.00","0.00","0.00"],"quality":"Less is better","isMain":true,"metricName":"# indels per 100 kbp"},{"values":[0,0,0,0,0,0,0,0,0,0,0,0,0,0],"quality":"Less is better","isMain":false,"metricName":" # indels (<= 5 bp)"},{"values":[0,0,0,0,0,0,0,13,0,0,0,0,0,0],"quality":"Less is better","isMain":false,"metricName":" # indels (> 5 bp)"},{"values":[0,0,0,0,0,1,0,1548,0,0,0,0,0,363],"quality":"Less is better","isMain":false,"metricName":"# N's"},{"values":["0.00","0.00","0.00","0.00","0.00","3.07","0.00","11049.25","0.00","0.00","0.00","0.00","0.00","570.42"],"quality":"Less is better","isMain":true,"metricName":"# N's per 100 kbp"}]],["Statistics without reference",[{"values":[4,3,10,5,6,4,9,2,8,2,2,4,3,1],"quality":"Equal","isMain":true,"metricName":"# contigs"},{"values":[1,3,6,3,3,4,1,2,3,2,2,3,3,1],"quality":"Equal","isMain":false,"metricName":"# contigs (>= 1000 bp)"},{"values":[1,1,1,1,1,3,0,2,1,1,1,1,1,1],"quality":"Equal","isMain":false,"metricName":"# contigs (>= 5000 bp)"},{"values":[1,0,1,0,1,1,0,0,1,1,0,0,0,1],"quality":"Equal","isMain":false,"metricName":"# contigs (>= 10000 bp)"},{"values":[0,0,0,0,0,0,0,0,0,0,0,0,0,1],"quality":"Equal","isMain":false,"metricName":"# contigs (>= 25000 bp)"},{"values":[0,0,0,0,0,0,0,0,0,0,0,0,0,1],"quality":"Equal","isMain":false,"metricName":"# contigs (>= 50000 bp)"},{"values":[13676,8664,13967,8825,13853,15792,1267,7747,14209,13314,8575,8575,8467,63637],"quality":"More is better","isMain":true,"metricName":"Largest contig"},{"values":[16076,14854,25645,16618,21916,32603,6458,14010,20716,14688,13363,15715,14344,63637],"quality":"More is better","isMain":true,"metricName":"Total length"},{"values":[13676,14854,23116,15502,19815,32603,1267,14010,17056,14688,13363,14962,14344,63637],"quality":"More is better","isMain":true,"metricName":"Total length (>= 1000 bp)"},{"values":[13676,8664,13967,8825,13853,31473,0,14010,14209,13314,8575,8575,8467,63637],"quality":"More is better","isMain":false,"metricName":"Total length (>= 5000 bp)"},{"values":[13676,0,13967,0,13853,15792,0,0,14209,13314,0,0,0,63637],"quality":"More is better","isMain":true,"metricName":"Total length (>= 10000 bp)"},{"values":[0,0,0,0,0,0,0,0,0,0,0,0,0,63637],"quality":"More is better","isMain":false,"metricName":"Total length (>= 25000 bp)"},{"values":[0,0,0,0,0,0,0,0,0,0,0,0,0,63637],"quality":"More is better","isMain":true,"metricName":"Total length (>= 50000 bp)"},{"values":[13676,8664,13967,8825,13853,8830,670,7747,14209,13314,8575,8575,8467,63637],"quality":"More is better","isMain":false,"metricName":"N50"},{"values":[13676,4802,1576,4876,3936,8830,579,6263,1809,13314,4788,4788,4656,63637],"quality":"More is better","isMain":false,"metricName":"N75"},{"values":[1,1,1,1,1,2,4,1,1,1,1,1,1,1],"quality":"Less is better","isMain":false,"metricName":"L50"},{"values":[1,2,4,2,2,2,7,2,2,1,2,2,2,1],"quality":"Less is better","isMain":false,"metricName":"L75"},{"values":["61.59","60.69","63.76","61.29","62.54","64.49","61.06","60.63","62.57","60.63","60.52","61.02","60.53","64.92"],"quality":"Equal","isMain":false,"metricName":"GC (%)"}]],["Predicted genes",[]],["Similarity statistics",[{"values":[0,0,0,0,0,0,0,0,0,0,0,0,0,0],"quality":"Equal","isMain":false,"metricName":"# similar correct contigs"},{"values":[0,0,0,0,0,0,0,0,0,0,0,0,0,0],"quality":"Equal","isMain":false,"metricName":"# similar misassembled blocks"}]],["Reference statistics",[{"values":[36690,36690,36690,36690,36690,36690,36690,36690,36690,36690,36690,36690,36690,36690],"quality":"Equal","isMain":false,"metricName":"Reference length"},{"values":[1,1,1,1,1,1,1,1,1,1,1,1,1,1],"quality":"Equal","isMain":false,"metricName":"Reference fragments"},{"values":["64.62","64.62","64.62","64.62","64.62","64.62","64.62","64.62","64.62","64.62","64.62","64.62","64.62","64.62"],"quality":"Equal","isMain":false,"metricName":"Reference GC (%)"}]]],"referenceName":"gi\_134288688\_ref\_NC","date":"18 July 2018, Wednesday, 16:17:28","order":[0,1,2,3,4,5,6,7,8,9,10,11,12,13],"assembliesNames":["ABySS\_127","ABySS\_63","CLC","IDBA\_UD","MEGAHIT","MIRA","Ray\_Meta","SOAPdenovo2","SPAdes","SPAdes\_meta","SPAdes\_sc","SPAdes\_sc\_careful","Velvet","Geneious"]},{"assembliesWithNs":null,"minContig":500,"report":[["Genome statistics",[{"values":["86.990","94.709","89.779","97.619","87.323","100.000","99.161","10.761","96.640","99.085","100.000","94.813","82.927","100.000"],"quality":"More is better","isMain":true,"metricName":"Genome fraction (%)"},{"values":["1.005","1.005","1.023","1.035","1.045","1.302","1.059","1.000","1.088","1.006","1.008","1.029","1.004","1.379"],"quality":"Less is better","isMain":true,"metricName":"Duplication ratio"},{"values":[12816,9805,4529,4475,6885,31235,13832,1207,7588,17281,20750,16334,3958,31238],"quality":"More is better","isMain":true,"metricName":"Largest alignment"},{"values":[31252,34036,32817,36106,32620,46539,37536,3846,37577,35634,36016,34217,29751,49289],"quality":"More is better","isMain":true,"metricName":"Total aligned length"},{"values":[6349,7183,3363,3417,3651,40228,11052,null,4025,11418,20750,6716,1387,63637],"quality":"More is better","isMain":false,"metricName":"NG50"},{"values":[2102,5775,1265,2456,1592,40228,5533,null,1809,11418,6716,4473,786,63637],"quality":"More is better","isMain":false,"metricName":"NG75"},{"values":[6349,7351,3794,3417,4297,31235,11052,926,4004,11418,20750,6716,1625,18051],"quality":"More is better","isMain":false,"metricName":"NA50"},{"values":[2946,5775,2326,2456,3648,8993,5533,883,821,11418,6716,4473,1221,18051],"quality":"More is better","isMain":false,"metricName":"NA75"},{"values":[6349,7183,3363,3417,3651,31235,11052,null,4004,11418,20750,6716,1387,31238],"quality":"More is better","isMain":true,"metricName":"NGA50"},{"values":[2102,5775,1265,2456,1591,31235,5533,null,1809,11418,6716,4473,786,31238],"quality":"More is better","isMain":false,"metricName":"NGA75"},{"values":[2,3,5,5,4,1,2,null,3,2,1,2,8,1],"quality":"Less is better","isMain":false,"metricName":"LG50"},{"values":[5,4,9,8,7,1,3,null,7,2,2,3,16,1],"quality":"Less is better","isMain":false,"metricName":"LG75"},{"values":[2,2,4,5,3,1,2,2,4,2,1,2,6,2],"quality":"Less is better","isMain":false,"metricName":"LA50"},{"values":[4,4,7,8,5,2,3,3,10,2,2,3,11,2],"quality":"Less is better","isMain":false,"metricName":"LA75"},{"values":[2,3,5,5,4,1,2,null,4,2,1,2,8,1],"quality":"Less is better","isMain":true,"metricName":"LGA50"},{"values":[5,4,9,8,7,1,3,null,7,2,2,3,16,1],"quality":"Less is better","isMain":false,"metricName":"LGA75"}]],["Misassemblies",[{"values":[0,0,0,0,0,1,0,0,0,0,0,0,0,1],"quality":"Less is better","isMain":true,"metricName":"# misassemblies"},{"values":[0,0,0,0,0,1,0,0,0,0,0,0,0,1],"quality":"Less is better","isMain":false,"metricName":" # relocations"},{"values":[0,0,0,0,0,0,0,0,0,0,0,0,0,0],"quality":"Less is better","isMain":false,"metricName":" # translocations"},{"values":[0,0,0,0,0,0,0,0,0,0,0,0,0,0],"quality":"Less is better","isMain":false,"metricName":" # inversions"},{"values":[0,0,0,0,0,1,0,0,0,0,0,0,0,1],"quality":"Less is better","isMain":false,"metricName":"# misassembled contigs"},{"values":[0,0,0,0,0,40228,0,0,0,0,0,0,0,63637],"quality":"Less is better","isMain":true,"metricName":"Misassembled contigs length"},{"values":[0,0,0,0,0,0,0,0,0,0,0,0,0,2],"quality":"Less is better","isMain":false,"metricName":"# local misassemblies"},{"values":[0,0,0,0,0,0,0,0,0,0,0,0,0,0],"quality":"Less is better","isMain":false,"metricName":"# unaligned mis. contigs"}]],["Unaligned",[{"values":[0,0,0,0,0,0,0,0,0,0,0,0,0,0],"quality":"Less is better","isMain":false,"metricName":"# fully unaligned contigs"},{"values":[0,0,0,0,0,0,0,0,0,0,0,0,0,0],"quality":"Less is better","isMain":false,"metricName":"Fully unaligned length"},{"values":[0,0,0,0,0,0,0,0,1,0,0,0,0,1],"quality":"Less is better","isMain":false,"metricName":"# partially unaligned contigs"},{"values":[0,0,0,0,0,0,0,0,1373,0,0,0,0,14348],"quality":"Less is better","isMain":false,"metricName":"Partially unaligned length"}]],["Mismatches",[{"values":[5,5,14,5,34,114,24,0,5,16,5,3,8,89],"quality":"Less is better","isMain":false,"metricName":"# mismatches"},{"values":[0,0,0,0,0,0,0,0,0,0,0,0,0,2],"quality":"Less is better","isMain":false,"metricName":"# indels"},{"values":[0,0,0,0,0,0,0,0,0,0,0,0,0,3],"quality":"Less is better","isMain":false,"metricName":"Indels length"},{"values":["16.08","14.77","43.63","14.33","108.94","318.96","67.72","0.00","14.48","45.18","13.99","8.85","26.99","249.01"],"quality":"Less is better","isMain":true,"metricName":"# mismatches per 100 kbp"},{"values":["0.00","0.00","0.00","0.00","0.00","0.00","0.00","0.00","0.00","0.00","0.00","0.00","0.00","5.60"],"quality":"Less is better","isMain":true,"metricName":"# indels per 100 kbp"},{"values":[0,0,0,0,0,0,0,0,0,0,0,0,0,2],"quality":"Less is better","isMain":false,"metricName":" # indels (<= 5 bp)"},{"values":[0,0,0,0,0,0,0,0,0,0,0,0,0,0],"quality":"Less is better","isMain":false,"metricName":" # indels (> 5 bp)"},{"values":[0,74,0,0,0,3,0,0,1373,0,0,653,0,363],"quality":"Less is better","isMain":false,"metricName":"# N's"},{"values":["0.00","217.42","0.00","0.00","0.00","6.45","0.00","0.00","3525.03","0.00","0.00","1872.51","0.00","570.42"],"quality":"Less is better","isMain":true,"metricName":"# N's per 100 kbp"}]],["Statistics without reference",[{"values":[8,8,16,17,12,5,6,4,22,5,5,7,20,1],"quality":"Equal","isMain":true,"metricName":"# contigs"},{"values":[6,5,9,12,9,3,6,1,7,5,5,7,13,1],"quality":"Equal","isMain":false,"metricName":"# contigs (>= 1000 bp)"},{"values":[2,4,0,0,2,1,3,0,2,2,2,2,0,1],"quality":"Equal","isMain":false,"metricName":"# contigs (>= 5000 bp)"},{"values":[1,0,0,0,0,1,2,0,0,2,1,1,0,1],"quality":"Equal","isMain":false,"metricName":"# contigs (>= 10000 bp)"},{"values":[0,0,0,0,0,1,0,0,0,0,0,0,0,1],"quality":"Equal","isMain":false,"metricName":"# contigs (>= 25000 bp)"},{"values":[0,0,0,0,0,0,0,0,0,0,0,0,0,1],"quality":"Equal","isMain":false,"metricName":"# contigs (>= 50000 bp)"},{"values":[12816,9805,4529,4475,6885,40228,13832,1207,7588,17281,20750,16990,3958,63637],"quality":"More is better","isMain":true,"metricName":"Largest contig"},{"values":[31252,34036,32823,36106,32621,46539,37536,3846,38950,35634,36016,34873,29751,63637],"quality":"More is better","isMain":true,"metricName":"Total length"},{"values":[29861,31890,28062,32966,30542,45298,37536,1207,28350,35634,36016,34873,24887,63637],"quality":"More is better","isMain":true,"metricName":"Total length (>= 1000 bp)"},{"values":[19165,30114,0,0,13034,40228,30417,0,14003,28699,27466,23706,0,63637],"quality":"More is better","isMain":false,"metricName":"Total length (>= 5000 bp)"},{"values":[12816,0,0,0,0,40228,24884,0,0,28699,20750,16990,0,63637],"quality":"More is better","isMain":true,"metricName":"Total length (>= 10000 bp)"},{"values":[0,0,0,0,0,40228,0,0,0,0,0,0,0,63637],"quality":"More is better","isMain":false,"metricName":"Total length (>= 25000 bp)"},{"values":[0,0,0,0,0,0,0,0,0,0,0,0,0,63637],"quality":"More is better","isMain":true,"metricName":"Total length (>= 50000 bp)"},{"values":[6349,7351,3794,3417,4297,40228,11052,926,4004,11418,20750,6716,1625,63637],"quality":"More is better","isMain":false,"metricName":"N50"},{"values":[2946,5775,2326,2456,3648,40228,5533,883,906,11418,6716,4473,1221,63637],"quality":"More is better","isMain":false,"metricName":"N75"},{"values":[2,2,4,5,3,1,2,2,4,2,1,2,6,1],"quality":"Less is better","isMain":false,"metricName":"L50"},{"values":[4,4,7,8,5,1,3,3,8,2,2,3,11,1],"quality":"Less is better","isMain":false,"metricName":"L75"},{"values":["65.05","65.37","65.38","65.45","65.46","65.90","65.56","58.74","65.43","65.52","65.44","65.36","64.94","64.92"],"quality":"Equal","isMain":false,"metricName":"GC (%)"}]],["Predicted genes",[]],["Similarity statistics",[{"values":[0,0,0,0,0,0,0,0,0,0,0,0,0,0],"quality":"Equal","isMain":false,"metricName":"# similar correct contigs"},{"values":[0,0,0,0,0,0,0,0,0,0,0,0,0,0],"quality":"Equal","isMain":false,"metricName":"# similar misassembled blocks"}]],["Reference statistics",[{"values":[35741,35741,35741,35741,35741,35741,35741,35741,35741,35741,35741,35741,35741,35741],"quality":"Equal","isMain":false,"metricName":"Reference length"},{"values":[1,1,1,1,1,1,1,1,1,1,1,1,1,1],"quality":"Equal","isMain":false,"metricName":"Reference fragments"},{"values":["65.43","65.43","65.43","65.43","65.43","65.43","65.43","65.43","65.43","65.43","65.43","65.43","65.43","65.43"],"quality":"Equal","isMain":false,"metricName":"Reference GC (%)"}]]],"referenceName":"gi\_134288739\_ref\_NC","date":"18 July 2018, Wednesday, 16:17:39","order":[0,1,2,3,4,5,6,7,8,9,10,11,12,13],"assembliesNames":["ABySS\_127","ABySS\_63","CLC","IDBA\_UD","MEGAHIT","MIRA","Ray\_Meta","SOAPdenovo2","SPAdes","SPAdes\_meta","SPAdes\_sc","SPAdes\_sc\_careful","Velvet","Geneious"]},{"assembliesWithNs":null,"minContig":500,"report":[["Genome statistics",[{"values":["100.000","99.469","98.451","99.239","100.000","100.000","100.000","100.000","100.000","97.020","97.094","37.019","79.645"],"quality":"More is better","isMain":true,"metricName":"Genome fraction (%)"},{"values":["1.528","1.011","1.000","1.024","1.008","1.020","1.001","1.003","1.003","1.009","1.007","1.001","1.000"],"quality":"Less is better","isMain":true,"metricName":"Duplication ratio"},{"values":[37178,27706,36020,9594,23079,38205,37495,37573,30741,21805,22136,4846,29824],"quality":"More is better","isMain":true,"metricName":"Largest alignment"},{"values":[57201,37653,36866,38039,37728,38205,37495,37573,37556,36660,36578,13878,29824],"quality":"More is better","isMain":true,"metricName":"Total aligned length"},{"values":[57201,27706,36020,4293,23079,38205,37495,37573,30741,21805,22155,null,66553],"quality":"More is better","isMain":false,"metricName":"NG50"},{"values":[57201,9947,36020,2671,14649,38205,37495,37573,30741,4150,6318,null,66553],"quality":"More is better","isMain":false,"metricName":"NG75"},{"values":[37178,27706,36020,4293,23079,38205,37495,37573,30741,21805,22136,1296,null],"quality":"More is better","isMain":false,"metricName":"NA50"},{"values":[20023,9947,36020,2025,14649,38205,37495,37573,30741,3579,6318,680,null],"quality":"More is better","isMain":false,"metricName":"NA75"},{"values":[37178,27706,36020,4293,23079,38205,37495,37573,30741,21805,22136,null,29824],"quality":"More is better","isMain":true,"metricName":"NGA50"},{"values":[37178,9947,36020,2671,14649,38205,37495,37573,30741,3579,6318,null,29824],"quality":"More is better","isMain":false,"metricName":"NGA75"},{"values":[1,1,1,3,1,1,1,1,1,1,1,null,1],"quality":"Less is better","isMain":false,"metricName":"LG50"},{"values":[1,2,1,5,2,1,1,1,1,3,2,null,1],"quality":"Less is better","isMain":false,"metricName":"LG75"},{"values":[1,1,1,3,1,1,1,1,1,1,1,3,null],"quality":"Less is better","isMain":false,"metricName":"LA50"},{"values":[2,2,1,6,2,1,1,1,1,3,2,7,null],"quality":"Less is better","isMain":false,"metricName":"LA75"},{"values":[1,1,1,3,1,1,1,1,1,1,1,null,1],"quality":"Less is better","isMain":true,"metricName":"LGA50"},{"values":[1,2,1,5,2,1,1,1,1,3,2,null,1],"quality":"Less is better","isMain":false,"metricName":"LGA75"}]],["Misassemblies",[{"values":[1,0,0,0,0,0,0,0,0,0,0,0,0],"quality":"Less is better","isMain":true,"metricName":"# misassemblies"},{"values":[1,0,0,0,0,0,0,0,0,0,0,0,0],"quality":"Less is better","isMain":false,"metricName":" # relocations"},{"values":[0,0,0,0,0,0,0,0,0,0,0,0,0],"quality":"Less is better","isMain":false,"metricName":" # translocations"},{"values":[0,0,0,0,0,0,0,0,0,0,0,0,0],"quality":"Less is better","isMain":false,"metricName":" # inversions"},{"values":[1,0,0,0,0,0,0,0,0,0,0,0,0],"quality":"Less is better","isMain":false,"metricName":"# misassembled contigs"},{"values":[57201,0,0,0,0,0,0,0,0,0,0,0,0],"quality":"Less is better","isMain":true,"metricName":"Misassembled contigs length"},{"values":[0,0,0,0,0,1,0,1,0,0,0,0,0],"quality":"Less is better","isMain":false,"metricName":"# local misassemblies"},{"values":[0,0,0,0,0,0,0,0,0,0,0,0,0],"quality":"Less is better","isMain":false,"metricName":"# unaligned mis. contigs"}]],["Unaligned",[{"values":[0,0,0,0,0,0,0,0,0,0,0,0,0],"quality":"Less is better","isMain":false,"metricName":"# fully unaligned contigs"},{"values":[0,0,0,0,0,0,0,0,0,0,0,0,0],"quality":"Less is better","isMain":false,"metricName":"Fully unaligned length"},{"values":[0,0,0,0,0,0,0,0,0,1,0,0,1],"quality":"Less is better","isMain":false,"metricName":"# partially unaligned contigs"},{"values":[0,0,0,0,0,0,0,0,0,571,0,0,36729],"quality":"Less is better","isMain":false,"metricName":"Partially unaligned length"}]],["Mismatches",[{"values":[0,0,0,0,2,1,0,0,0,0,0,0,2],"quality":"Less is better","isMain":false,"metricName":"# mismatches"},{"values":[0,0,0,0,0,0,1,0,0,0,0,0,0],"quality":"Less is better","isMain":false,"metricName":"# indels"},{"values":[0,0,0,0,0,0,49,0,0,0,0,0,0],"quality":"Less is better","isMain":false,"metricName":"Indels length"},{"values":["0.00","0.00","0.00","0.00","5.34","2.67","0.00","0.00","0.00","0.00","0.00","0.00","6.71"],"quality":"Less is better","isMain":true,"metricName":"# mismatches per 100 kbp"},{"values":["0.00","0.00","0.00","0.00","0.00","0.00","2.67","0.00","0.00","0.00","0.00","0.00","0.00"],"quality":"Less is better","isMain":true,"metricName":"# indels per 100 kbp"},{"values":[0,0,0,0,0,0,0,0,0,0,0,0,0],"quality":"Less is better","isMain":false,"metricName":" # indels (<= 5 bp)"},{"values":[0,0,0,0,0,0,1,0,0,0,0,0,0],"quality":"Less is better","isMain":false,"metricName":" # indels (> 5 bp)"},{"values":[0,21,0,0,0,0,0,0,0,571,0,0,525],"quality":"Less is better","isMain":false,"metricName":"# N's"},{"values":["0.00","55.77","0.00","0.00","0.00","0.00","0.00","0.00","0.00","1533.67","0.00","0.00","788.84"],"quality":"Less is better","isMain":true,"metricName":"# N's per 100 kbp"}]],["Statistics without reference",[{"values":[1,2,2,12,2,1,1,1,2,7,8,12,1],"quality":"Equal","isMain":true,"metricName":"# contigs"},{"values":[1,2,1,10,2,1,1,1,2,6,4,3,1],"quality":"Equal","isMain":false,"metricName":"# contigs (>= 1000 bp)"},{"values":[1,2,1,2,2,1,1,1,2,1,2,0,1],"quality":"Equal","isMain":false,"metricName":"# contigs (>= 5000 bp)"},{"values":[1,1,1,0,2,1,1,1,1,1,1,0,1],"quality":"Equal","isMain":false,"metricName":"# contigs (>= 10000 bp)"},{"values":[1,1,1,0,0,1,1,1,1,0,0,0,1],"quality":"Equal","isMain":false,"metricName":"# contigs (>= 25000 bp)"},{"values":[1,0,0,0,0,0,0,0,0,0,0,0,1],"quality":"Equal","isMain":false,"metricName":"# contigs (>= 50000 bp)"},{"values":[57201,27706,36020,9594,23079,38205,37495,37573,30741,21805,22155,4846,66553],"quality":"More is better","isMain":true,"metricName":"Largest contig"},{"values":[57201,37653,36866,38039,37728,38205,37495,37573,37556,37231,36597,13878,66553],"quality":"More is better","isMain":true,"metricName":"Total length"},{"values":[57201,37653,36020,36840,37728,38205,37495,37573,37556,36550,33916,8011,66553],"quality":"More is better","isMain":true,"metricName":"Total length (>= 1000 bp)"},{"values":[57201,37653,36020,18178,37728,38205,37495,37573,37556,21805,28473,0,66553],"quality":"More is better","isMain":false,"metricName":"Total length (>= 5000 bp)"},{"values":[57201,27706,36020,0,37728,38205,37495,37573,30741,21805,22155,0,66553],"quality":"More is better","isMain":true,"metricName":"Total length (>= 10000 bp)"},{"values":[57201,27706,36020,0,0,38205,37495,37573,30741,0,0,0,66553],"quality":"More is better","isMain":false,"metricName":"Total length (>= 25000 bp)"},{"values":[57201,0,0,0,0,0,0,0,0,0,0,0,66553],"quality":"More is better","isMain":true,"metricName":"Total length (>= 50000 bp)"},{"values":[57201,27706,36020,4293,23079,38205,37495,37573,30741,21805,22155,1296,66553],"quality":"More is better","isMain":false,"metricName":"N50"},{"values":[57201,9947,36020,2025,14649,38205,37495,37573,30741,4150,6318,680,66553],"quality":"More is better","isMain":false,"metricName":"N75"},{"values":[1,1,1,3,1,1,1,1,1,1,1,3,1],"quality":"Less is better","isMain":false,"metricName":"L50"},{"values":[1,2,1,6,2,1,1,1,1,3,2,7,1],"quality":"Less is better","isMain":false,"metricName":"L75"},{"values":["62.63","63.01","63.04","63.02","63.07","63.11","63.06","63.05","63.04","62.95","62.96","61.99","62.80"],"quality":"Equal","isMain":false,"metricName":"GC (%)"}]],["Predicted genes",[]],["Similarity statistics",[{"values":[0,0,0,0,0,0,0,0,0,0,0,0,0],"quality":"Equal","isMain":false,"metricName":"# similar correct contigs"},{"values":[0,0,0,0,0,0,0,0,0,0,0,0,0],"quality":"Equal","isMain":false,"metricName":"# similar misassembled blocks"}]],["Reference statistics",[{"values":[37446,37446,37446,37446,37446,37446,37446,37446,37446,37446,37446,37446,37446],"quality":"Equal","isMain":false,"metricName":"Reference length"},{"values":[1,1,1,1,1,1,1,1,1,1,1,1,1],"quality":"Equal","isMain":false,"metricName":"Reference fragments"},{"values":["63.05","63.05","63.05","63.05","63.05","63.05","63.05","63.05","63.05","63.05","63.05","63.05","63.05"],"quality":"Equal","isMain":false,"metricName":"Reference GC (%)"}]]],"referenceName":"gi\_134288788\_ref\_NC","date":"18 July 2018, Wednesday, 16:17:50","order":[0,1,2,3,4,5,6,7,8,9,10,11,12],"assembliesNames":["ABySS\_127","ABySS\_63","CLC","IDBA\_UD","MEGAHIT","MIRA","Ray\_Meta","SPAdes","SPAdes\_meta","SPAdes\_sc","SPAdes\_sc\_careful","Velvet","Geneious"]},{"assembliesWithNs":null,"minContig":500,"report":[["Genome statistics",[{"values":["99.756","98.676","100.000","99.985","100.000","100.000","2.906","98.536","100.000","99.889","99.889","99.889","99.502","100.000"],"quality":"More is better","isMain":true,"metricName":"Genome fraction (%)"},{"values":["1.003","1.001","1.000","1.000","1.002","1.019","1.003","1.032","1.001","1.001","1.001","1.001","1.000","1.003"],"quality":"Less is better","isMain":true,"metricName":"Duplication ratio"},{"values":[26302,15106,96944,96929,97101,97200,618,49140,97031,96887,96887,96887,56605,97200],"quality":"More is better","isMain":true,"metricName":"Largest alignment"},{"values":[97018,95779,96944,96929,97101,98798,2825,96219,97031,96887,96887,96887,96493,97200],"quality":"More is better","isMain":true,"metricName":"Total aligned length"},{"values":[9676,6956,96944,96929,97101,97200,null,50190,97031,96887,96887,96887,56605,97200],"quality":"More is better","isMain":false,"metricName":"NG50"},{"values":[7301,4193,96944,96929,97101,97200,null,27847,97031,96887,96887,96887,36383,97200],"quality":"More is better","isMain":false,"metricName":"NG75"},{"values":[9676,6956,96944,96929,97101,97200,578,26880,97031,96887,96887,96887,56605,97200],"quality":"More is better","isMain":false,"metricName":"NA50"},{"values":[7301,4193,96944,96929,97101,97200,549,26880,97031,96887,96887,96887,36383,97200],"quality":"More is better","isMain":false,"metricName":"NA75"},{"values":[9676,6956,96944,96929,97101,97200,null,49140,97031,96887,96887,96887,56605,97200],"quality":"More is better","isMain":true,"metricName":"NGA50"},{"values":[7301,4193,96944,96929,97101,97200,null,26880,97031,96887,96887,96887,36383,97200],"quality":"More is better","isMain":false,"metricName":"NGA75"},{"values":[3,5,1,1,1,1,null,1,1,1,1,1,1,1],"quality":"Less is better","isMain":false,"metricName":"LG50"},{"values":[6,10,1,1,1,1,null,2,1,1,1,1,2,1],"quality":"Less is better","isMain":false,"metricName":"LG75"},{"values":[3,5,1,1,1,1,3,2,1,1,1,1,1,1],"quality":"Less is better","isMain":false,"metricName":"LA50"},{"values":[6,10,1,1,1,1,4,2,1,1,1,1,2,1],"quality":"Less is better","isMain":false,"metricName":"LA75"},{"values":[3,5,1,1,1,1,null,1,1,1,1,1,1,1],"quality":"Less is better","isMain":true,"metricName":"LGA50"},{"values":[6,10,1,1,1,1,null,2,1,1,1,1,2,1],"quality":"Less is better","isMain":false,"metricName":"LGA75"}]],["Misassemblies",[{"values":[0,0,0,0,0,0,0,0,0,0,0,0,0,0],"quality":"Less is better","isMain":true,"metricName":"# misassemblies"},{"values":[0,0,0,0,0,0,0,0,0,0,0,0,0,0],"quality":"Less is better","isMain":false,"metricName":" # relocations"},{"values":[0,0,0,0,0,0,0,0,0,0,0,0,0,0],"quality":"Less is better","isMain":false,"metricName":" # translocations"},{"values":[0,0,0,0,0,0,0,0,0,0,0,0,0,0],"quality":"Less is better","isMain":false,"metricName":" # inversions"},{"values":[0,0,0,0,0,0,0,0,0,0,0,0,0,0],"quality":"Less is better","isMain":false,"metricName":"# misassembled contigs"},{"values":[0,0,0,0,0,0,0,0,0,0,0,0,0,0],"quality":"Less is better","isMain":true,"metricName":"Misassembled contigs length"},{"values":[0,0,0,0,0,1,0,0,1,1,1,1,0,1],"quality":"Less is better","isMain":false,"metricName":"# local misassemblies"},{"values":[0,0,0,0,0,0,0,0,0,0,0,0,0,0],"quality":"Less is better","isMain":false,"metricName":"# unaligned mis. contigs"}]],["Unaligned",[{"values":[0,0,0,0,0,0,0,0,0,0,0,0,0,0],"quality":"Less is better","isMain":false,"metricName":"# fully unaligned contigs"},{"values":[0,0,0,0,0,0,0,0,0,0,0,0,0,0],"quality":"Less is better","isMain":false,"metricName":"Fully unaligned length"},{"values":[0,0,0,0,0,0,0,0,0,0,0,0,0,0],"quality":"Less is better","isMain":false,"metricName":"# partially unaligned contigs"},{"values":[0,0,0,0,0,0,0,0,0,0,0,0,0,0],"quality":"Less is better","isMain":false,"metricName":"Partially unaligned length"}]],["Mismatches",[{"values":[9,0,0,0,0,0,0,0,1,1,1,1,0,0],"quality":"Less is better","isMain":false,"metricName":"# mismatches"},{"values":[1,0,0,0,3,0,0,35,1,3,3,3,0,0],"quality":"Less is better","isMain":false,"metricName":"# indels"},{"values":[40,0,0,0,36,0,0,1692,40,3,3,3,0,0],"quality":"Less is better","isMain":false,"metricName":"Indels length"},{"values":["9.31","0.00","0.00","0.00","0.00","0.00","0.00","0.00","1.03","1.03","1.03","1.03","0.00","0.00"],"quality":"Less is better","isMain":true,"metricName":"# mismatches per 100 kbp"},{"values":["1.03","0.00","0.00","0.00","3.09","0.00","0.00","36.64","1.03","3.10","3.10","3.10","0.00","0.00"],"quality":"Less is better","isMain":true,"metricName":"# indels per 100 kbp"},{"values":[0,0,0,0,1,0,0,0,0,3,3,3,0,0],"quality":"Less is better","isMain":false,"metricName":" # indels (<= 5 bp)"},{"values":[1,0,0,0,2,0,0,35,1,0,0,0,0,0],"quality":"Less is better","isMain":false,"metricName":" # indels (> 5 bp)"},{"values":[0,0,0,0,0,3,0,3211,0,0,0,0,0,3],"quality":"Less is better","isMain":false,"metricName":"# N's"},{"values":["0.00","0.00","0.00","0.00","0.00","3.04","0.00","3256.86","0.00","0.00","0.00","0.00","0.00","3.09"],"quality":"Less is better","isMain":true,"metricName":"# N's per 100 kbp"}]],["Statistics without reference",[{"values":[13,19,1,1,1,3,5,6,1,1,1,1,3,1],"quality":"Equal","isMain":true,"metricName":"# contigs"},{"values":[11,17,1,1,1,2,0,6,1,1,1,1,3,1],"quality":"Equal","isMain":false,"metricName":"# contigs (>= 1000 bp)"},{"values":[7,8,1,1,1,1,0,3,1,1,1,1,2,1],"quality":"Equal","isMain":false,"metricName":"# contigs (>= 5000 bp)"},{"values":[2,2,1,1,1,1,0,2,1,1,1,1,2,1],"quality":"Equal","isMain":false,"metricName":"# contigs (>= 10000 bp)"},{"values":[1,0,1,1,1,1,0,2,1,1,1,1,2,1],"quality":"Equal","isMain":false,"metricName":"# contigs (>= 25000 bp)"},{"values":[0,0,1,1,1,1,0,1,1,1,1,1,1,1],"quality":"Equal","isMain":false,"metricName":"# contigs (>= 50000 bp)"},{"values":[26302,15106,96944,96929,97101,97200,618,50190,97031,96887,96887,96887,56605,97200],"quality":"More is better","isMain":true,"metricName":"Largest contig"},{"values":[97018,95779,96944,96929,97101,98798,2825,98592,97031,96887,96887,96887,96493,97200],"quality":"More is better","isMain":true,"metricName":"Total length"},{"values":[95604,94604,96944,96929,97101,98249,0,98592,97031,96887,96887,96887,96493,97200],"quality":"More is better","isMain":true,"metricName":"Total length (>= 1000 bp)"},{"values":[83305,66361,96944,96929,97101,97200,0,85136,97031,96887,96887,96887,92988,97200],"quality":"More is better","isMain":false,"metricName":"Total length (>= 5000 bp)"},{"values":[44207,26434,96944,96929,97101,97200,0,78037,97031,96887,96887,96887,92988,97200],"quality":"More is better","isMain":true,"metricName":"Total length (>= 10000 bp)"},{"values":[26302,0,96944,96929,97101,97200,0,78037,97031,96887,96887,96887,92988,97200],"quality":"More is better","isMain":false,"metricName":"Total length (>= 25000 bp)"},{"values":[0,0,96944,96929,97101,97200,0,50190,97031,96887,96887,96887,56605,97200],"quality":"More is better","isMain":true,"metricName":"Total length (>= 50000 bp)"},{"values":[9676,6956,96944,96929,97101,97200,578,50190,97031,96887,96887,96887,56605,97200],"quality":"More is better","isMain":false,"metricName":"N50"},{"values":[7301,4193,96944,96929,97101,97200,549,27847,97031,96887,96887,96887,36383,97200],"quality":"More is better","isMain":false,"metricName":"N75"},{"values":[3,5,1,1,1,1,3,1,1,1,1,1,1,1],"quality":"Less is better","isMain":false,"metricName":"L50"},{"values":[6,10,1,1,1,1,4,2,1,1,1,1,2,1],"quality":"Less is better","isMain":false,"metricName":"L75"},{"values":["27.97","27.98","27.99","27.99","27.97","27.93","27.58","28.01","27.97","28.01","28.01","28.01","27.99","27.93"],"quality":"Equal","isMain":false,"metricName":"GC (%)"}]],["Predicted genes",[]],["Similarity statistics",[{"values":[0,0,0,0,0,0,0,0,0,0,0,0,0,0],"quality":"Equal","isMain":false,"metricName":"# similar correct contigs"},{"values":[0,0,0,0,0,0,0,0,0,0,0,0,0,0],"quality":"Equal","isMain":false,"metricName":"# similar misassembled blocks"}]],["Reference statistics",[{"values":[96944,96944,96944,96944,96944,96944,96944,96944,96944,96944,96944,96944,96944,96944],"quality":"Equal","isMain":false,"metricName":"Reference length"},{"values":[1,1,1,1,1,1,1,1,1,1,1,1,1,1],"quality":"Equal","isMain":false,"metricName":"Reference fragments"},{"values":["27.99","27.99","27.99","27.99","27.99","27.99","27.99","27.99","27.99","27.99","27.99","27.99","27.99","27.99"],"quality":"Equal","isMain":false,"metricName":"Reference GC (%)"}]]],"referenceName":"gi\_134303398\_ref\_NC","date":"18 July 2018, Wednesday, 16:18:01","order":[0,1,2,3,4,5,6,7,8,9,10,11,12,13],"assembliesNames":["ABySS\_127","ABySS\_63","CLC","IDBA\_UD","MEGAHIT","MIRA","Ray\_Meta","SOAPdenovo2","SPAdes","SPAdes\_meta","SPAdes\_sc","SPAdes\_sc\_careful","Velvet","Geneious"]},{"assembliesWithNs":null,"minContig":500,"report":[["Genome statistics",[{"values":["74.994","43.354","76.514","59.919","71.747","100.000","74.838","84.121","43.091","57.699","51.553","24.109","94.724"],"quality":"More is better","isMain":true,"metricName":"Genome fraction (%)"},{"values":["1.086","1.000","1.008","1.011","1.035","1.460","1.070","1.079","1.070","1.021","1.020","1.005","1.157"],"quality":"Less is better","isMain":true,"metricName":"Duplication ratio"},{"values":[5031,2776,10841,2528,4239,34603,9431,7047,10517,6192,6192,1387,27147],"quality":"More is better","isMain":true,"metricName":"Largest alignment"},{"values":[31242,16625,29581,23234,28419,55996,30717,34796,17670,22568,20159,9290,42030],"quality":"More is better","isMain":true,"metricName":"Total aligned length"},{"values":[4343,null,4785,617,999,79060,3545,6101,9651,1527,791,null,136335],"quality":"More is better","isMain":false,"metricName":"NG50"},{"values":[2081,null,626,null,null,79060,2272,5587,null,null,null,null,136335],"quality":"More is better","isMain":false,"metricName":"NG75"},{"values":[3035,2048,7671,1077,1654,null,4240,5286,3509,3562,1785,836,null],"quality":"More is better","isMain":false,"metricName":"NA50"},{"values":[962,892,4785,776,795,null,2410,677,null,988,791,712,null],"quality":"More is better","isMain":false,"metricName":"NA75"},{"values":[3035,null,4785,617,999,34603,3545,5286,null,844,525,null,27147],"quality":"More is better","isMain":true,"metricName":"NGA50"},{"values":[745,null,626,null,null,34603,2272,1788,null,null,null,null,3642],"quality":"More is better","isMain":false,"metricName":"NGA75"},{"values":[4,null,3,16,9,1,4,3,2,5,7,null,1],"quality":"Less is better","isMain":false,"metricName":"LG50"},{"values":[7,null,8,null,null,1,7,4,null,null,null,null,1],"quality":"Less is better","isMain":false,"metricName":"LG75"},{"values":[5,4,2,7,6,null,3,4,2,3,3,5,null],"quality":"Less is better","isMain":false,"metricName":"LA50"},{"values":[9,7,3,13,12,null,5,10,null,7,7,7,null],"quality":"Less is better","isMain":false,"metricName":"LA75"},{"values":[5,null,3,16,9,1,4,4,null,8,11,null,1],"quality":"Less is better","isMain":true,"metricName":"LGA50"},{"values":[11,null,8,null,null,1,7,7,null,null,null,null,2],"quality":"Less is better","isMain":false,"metricName":"LGA75"}]],["Misassemblies",[{"values":[0,0,0,0,0,2,0,0,0,0,0,0,0],"quality":"Less is better","isMain":true,"metricName":"# misassemblies"},{"values":[0,0,0,0,0,2,0,0,0,0,0,0,0],"quality":"Less is better","isMain":false,"metricName":" # relocations"},{"values":[0,0,0,0,0,0,0,0,0,0,0,0,0],"quality":"Less is better","isMain":false,"metricName":" # translocations"},{"values":[0,0,0,0,0,0,0,0,0,0,0,0,0],"quality":"Less is better","isMain":false,"metricName":" # inversions"},{"values":[0,0,0,0,0,1,0,0,0,0,0,0,0],"quality":"Less is better","isMain":false,"metricName":"# misassembled contigs"},{"values":[0,0,0,0,0,79060,0,0,0,0,0,0,0],"quality":"Less is better","isMain":true,"metricName":"Misassembled contigs length"},{"values":[0,0,0,0,0,10,0,1,0,0,0,0,0],"quality":"Less is better","isMain":false,"metricName":"# local misassemblies"},{"values":[0,0,0,0,0,0,0,0,1,1,1,0,1],"quality":"Less is better","isMain":false,"metricName":"# unaligned mis. contigs"}]],["Unaligned",[{"values":[0,0,0,0,0,0,0,0,0,0,0,0,0],"quality":"Less is better","isMain":false,"metricName":"# fully unaligned contigs"},{"values":[0,0,0,0,0,0,0,0,0,0,0,0,0],"quality":"Less is better","isMain":false,"metricName":"Fully unaligned length"},{"values":[1,0,0,0,0,3,0,2,1,1,1,0,1],"quality":"Less is better","isMain":false,"metricName":"# partially unaligned contigs"},{"values":[5246,0,0,0,0,56865,0,8557,6550,2439,2439,0,94316],"quality":"Less is better","isMain":false,"metricName":"Partially unaligned length"}]],["Mismatches",[{"values":[0,5,4,0,17,71,16,10,99,48,25,0,94],"quality":"Less is better","isMain":false,"metricName":"# mismatches"},{"values":[0,0,0,0,1,3,0,0,1,0,0,0,15],"quality":"Less is better","isMain":false,"metricName":"# indels"},{"values":[0,0,0,0,1,160,0,0,1,0,0,0,18],"quality":"Less is better","isMain":false,"metricName":"Indels length"},{"values":["0.00","30.08","13.63","0.00","61.79","185.15","55.75","31.00","599.13","216.94","126.46","0.00","258.78"],"quality":"Less is better","isMain":true,"metricName":"# mismatches per 100 kbp"},{"values":["0.00","0.00","0.00","0.00","3.63","7.82","0.00","0.00","6.05","0.00","0.00","0.00","41.30"],"quality":"Less is better","isMain":true,"metricName":"# indels per 100 kbp"},{"values":[0,0,0,0,1,0,0,0,1,0,0,0,15],"quality":"Less is better","isMain":false,"metricName":" # indels (<= 5 bp)"},{"values":[0,0,0,0,0,3,0,0,0,0,0,0,0],"quality":"Less is better","isMain":false,"metricName":" # indels (> 5 bp)"},{"values":[61,68,0,0,0,6,0,0,0,0,0,0,199],"quality":"Less is better","isMain":false,"metricName":"# N's"},{"values":["167.18","409.02","0.00","0.00","0.00","5.32","0.00","0.00","0.00","0.00","0.00","0.00","145.96"],"quality":"Less is better","isMain":true,"metricName":"# N's per 100 kbp"}]],["Statistics without reference",[{"values":[14,12,9,22,22,8,8,13,4,12,12,11,1],"quality":"Equal","isMain":true,"metricName":"# contigs"},{"values":[8,6,6,9,8,5,8,8,3,6,6,3,1],"quality":"Equal","isMain":false,"metricName":"# contigs (>= 1000 bp)"},{"values":[2,0,2,0,0,3,1,5,2,1,1,0,1],"quality":"Equal","isMain":false,"metricName":"# contigs (>= 5000 bp)"},{"values":[0,0,1,0,0,3,0,1,1,0,0,0,1],"quality":"Equal","isMain":false,"metricName":"# contigs (>= 10000 bp)"},{"values":[0,0,0,0,0,1,0,0,0,0,0,0,1],"quality":"Equal","isMain":false,"metricName":"# contigs (>= 25000 bp)"},{"values":[0,0,0,0,0,1,0,0,0,0,0,0,1],"quality":"Equal","isMain":false,"metricName":"# contigs (>= 50000 bp)"},{"values":[7725,2776,10841,2528,4239,79060,9431,11066,10520,6192,6192,1387,136335],"quality":"More is better","isMain":true,"metricName":"Largest contig"},{"values":[36488,16625,29581,23234,28470,112861,30717,43353,24223,25019,22607,9290,136335],"quality":"More is better","isMain":true,"metricName":"Total length"},{"values":[32005,12269,27818,14388,18229,110535,30717,40145,23680,20707,18936,3732,136335],"quality":"More is better","isMain":true,"metricName":"Total length (>= 1000 bp)"},{"values":[12756,0,18512,0,0,107256,9431,35061,20171,6192,6192,0,136335],"quality":"More is better","isMain":false,"metricName":"Total length (>= 5000 bp)"},{"values":[0,0,10841,0,0,107256,0,11066,10520,0,0,0,136335],"quality":"More is better","isMain":true,"metricName":"Total length (>= 10000 bp)"},{"values":[0,0,0,0,0,79060,0,0,0,0,0,0,136335],"quality":"More is better","isMain":false,"metricName":"Total length (>= 25000 bp)"},{"values":[0,0,0,0,0,79060,0,0,0,0,0,0,136335],"quality":"More is better","isMain":true,"metricName":"Total length (>= 50000 bp)"},{"values":[4343,2048,7671,1077,1654,79060,4240,6101,9651,3699,3708,836,136335],"quality":"More is better","isMain":false,"metricName":"N50"},{"values":[3035,892,4785,776,795,17653,2410,5286,9651,1527,1527,712,136335],"quality":"More is better","isMain":false,"metricName":"N75"},{"values":[4,4,2,7,6,1,3,3,2,3,3,5,1],"quality":"Less is better","isMain":false,"metricName":"L50"},{"values":[6,7,3,13,12,2,5,5,2,5,5,7,1],"quality":"Less is better","isMain":false,"metricName":"L75"},{"values":["36.67","36.30","36.73","36.77","36.78","35.49","36.80","36.00","36.16","36.18","36.19","35.55","35.76"],"quality":"Equal","isMain":false,"metricName":"GC (%)"}]],["Predicted genes",[]],["Similarity statistics",[{"values":[0,0,0,0,0,0,0,0,0,0,0,0,0],"quality":"Equal","isMain":false,"metricName":"# similar correct contigs"},{"values":[0,0,0,0,0,0,0,0,0,0,0,0,0],"quality":"Equal","isMain":false,"metricName":"# similar misassembled blocks"}]],["Reference statistics",[{"values":[38347,38347,38347,38347,38347,38347,38347,38347,38347,38347,38347,38347,38347],"quality":"Equal","isMain":false,"metricName":"Reference length"},{"values":[1,1,1,1,1,1,1,1,1,1,1,1,1],"quality":"Equal","isMain":false,"metricName":"Reference fragments"},{"values":["36.22","36.22","36.22","36.22","36.22","36.22","36.22","36.22","36.22","36.22","36.22","36.22","36.22"],"quality":"Equal","isMain":false,"metricName":"Reference GC (%)"}]]],"referenceName":"gi\_13487801\_ref\_NC","date":"18 July 2018, Wednesday, 16:18:13","order":[0,1,2,3,4,5,6,7,8,9,10,11,12],"assembliesNames":["ABySS\_127","ABySS\_63","CLC","IDBA\_UD","MEGAHIT","MIRA","Ray\_Meta","SPAdes","SPAdes\_meta","SPAdes\_sc","SPAdes\_sc\_careful","Velvet","Geneious"]},{"assembliesWithNs":null,"minContig":500,"report":[["Genome statistics",[{"values":["92.086","52.863","92.929","74.695","83.155","100.000","84.218","87.709","92.793","85.620","71.590","35.862","100.000"],"quality":"More is better","isMain":true,"metricName":"Genome fraction (%)"},{"values":["1.014","1.046","1.145","1.021","1.025","1.422","1.000","1.008","1.002","1.008","1.008","1.002","1.409"],"quality":"Less is better","isMain":true,"metricName":"Duplication ratio"},{"values":[15867,6034,16664,4455,6952,28151,22877,21993,26900,7803,7803,3825,28151],"quality":"More is better","isMain":true,"metricName":"Largest alignment"},{"values":[35776,21184,40766,29207,32646,54473,32253,33844,35561,33061,27637,13759,53960],"quality":"More is better","isMain":true,"metricName":"Total aligned length"},{"values":[9992,829,5935,1547,2159,77112,22877,21993,26900,4182,1971,null,77772],"quality":"More is better","isMain":false,"metricName":"NG50"},{"values":[8746,null,4013,547,1074,77112,6913,9318,7355,2532,null,null,77772],"quality":"More is better","isMain":false,"metricName":"NG75"},{"values":[9992,5236,5935,2800,4496,14344,22877,21993,26900,4182,4181,1569,14344],"quality":"More is better","isMain":false,"metricName":"NA50"},{"values":[8746,2730,4013,920,1426,null,6913,9318,26900,2747,1971,860,null],"quality":"More is better","isMain":false,"metricName":"NA75"},{"values":[9992,829,5935,1547,2159,28151,22877,21993,26900,4182,1971,null,28151],"quality":"More is better","isMain":true,"metricName":"NGA50"},{"values":[8746,null,4013,547,1074,14344,6913,9318,7355,2532,null,null,14344],"quality":"More is better","isMain":false,"metricName":"NGA75"},{"values":[2,7,2,7,4,1,1,1,1,3,5,null,1],"quality":"Less is better","isMain":false,"metricName":"LG50"},{"values":[3,null,4,20,11,1,2,2,2,6,null,null,1],"quality":"Less is better","isMain":false,"metricName":"LG75"},{"values":[2,2,2,4,3,2,1,1,1,3,3,3,2],"quality":"Less is better","isMain":false,"metricName":"LA50"},{"values":[3,4,4,9,7,null,2,2,1,5,5,7,null],"quality":"Less is better","isMain":false,"metricName":"LA75"},{"values":[2,7,2,7,4,1,1,1,1,3,5,null,1],"quality":"Less is better","isMain":true,"metricName":"LGA50"},{"values":[3,null,4,20,11,2,2,2,2,6,null,null,2],"quality":"Less is better","isMain":false,"metricName":"LGA75"}]],["Misassemblies",[{"values":[0,0,0,0,0,4,0,0,0,0,0,0,4],"quality":"Less is better","isMain":true,"metricName":"# misassemblies"},{"values":[0,0,0,0,0,4,0,0,0,0,0,0,4],"quality":"Less is better","isMain":false,"metricName":" # relocations"},{"values":[0,0,0,0,0,0,0,0,0,0,0,0,0],"quality":"Less is better","isMain":false,"metricName":" # translocations"},{"values":[0,0,0,0,0,0,0,0,0,0,0,0,0],"quality":"Less is better","isMain":false,"metricName":" # inversions"},{"values":[0,0,0,0,0,1,0,0,0,0,0,0,1],"quality":"Less is better","isMain":false,"metricName":"# misassembled contigs"},{"values":[0,0,0,0,0,77112,0,0,0,0,0,0,77772],"quality":"Less is better","isMain":true,"metricName":"Misassembled contigs length"},{"values":[0,0,0,0,0,8,0,0,0,0,0,0,8],"quality":"Less is better","isMain":false,"metricName":"# local misassemblies"},{"values":[0,0,0,0,0,0,0,0,0,0,0,0,0],"quality":"Less is better","isMain":false,"metricName":"# unaligned mis. contigs"}]],["Unaligned",[{"values":[0,0,0,0,0,0,0,0,0,0,0,0,0],"quality":"Less is better","isMain":false,"metricName":"# fully unaligned contigs"},{"values":[0,0,0,0,0,0,0,0,0,0,0,0,0],"quality":"Less is better","isMain":false,"metricName":"Fully unaligned length"},{"values":[0,0,0,0,0,1,0,0,0,0,0,0,1],"quality":"Less is better","isMain":false,"metricName":"# partially unaligned contigs"},{"values":[0,0,0,0,0,23812,0,0,0,0,0,0,23812],"quality":"Less is better","isMain":false,"metricName":"Partially unaligned length"}]],["Mismatches",[{"values":[70,28,14,0,3,125,1,5,248,187,27,1,125],"quality":"Less is better","isMain":false,"metricName":"# mismatches"},{"values":[0,0,0,0,0,4,0,0,4,2,0,0,4],"quality":"Less is better","isMain":false,"metricName":"# indels"},{"values":[0,0,0,0,0,4,0,0,6,4,0,0,4],"quality":"Less is better","isMain":false,"metricName":"Indels length"},{"values":["198.49","138.31","39.34","0.00","9.42","326.40","3.10","14.89","697.86","570.30","98.48","7.28","326.40"],"quality":"Less is better","isMain":true,"metricName":"# mismatches per 100 kbp"},{"values":["0.00","0.00","0.00","0.00","0.00","10.44","0.00","0.00","11.26","6.10","0.00","0.00","10.44"],"quality":"Less is better","isMain":true,"metricName":"# indels per 100 kbp"},{"values":[0,0,0,0,0,4,0,0,4,2,0,0,4],"quality":"Less is better","isMain":false,"metricName":" # indels (<= 5 bp)"},{"values":[0,0,0,0,0,0,0,0,0,0,0,0,0],"quality":"Less is better","isMain":false,"metricName":" # indels (> 5 bp)"},{"values":[0,20,0,0,0,0,0,0,0,0,0,0,16],"quality":"Less is better","isMain":false,"metricName":"# N's"},{"values":["0.00","94.41","0.00","0.00","0.00","0.00","0.00","0.00","0.00","0.00","0.00","0.00","20.57"],"quality":"Less is better","isMain":true,"metricName":"# N's per 100 kbp"}]],["Statistics without reference",[{"values":[5,9,13,20,16,2,3,6,4,10,11,11,1],"quality":"Equal","isMain":true,"metricName":"# contigs"},{"values":[3,5,8,7,11,2,3,2,2,8,9,4,1],"quality":"Equal","isMain":false,"metricName":"# contigs (>= 1000 bp)"},{"values":[3,2,2,0,2,1,2,2,2,2,1,0,1],"quality":"Equal","isMain":false,"metricName":"# contigs (>= 5000 bp)"},{"values":[1,0,1,0,0,1,1,1,1,0,0,0,1],"quality":"Equal","isMain":false,"metricName":"# contigs (>= 10000 bp)"},{"values":[0,0,0,0,0,1,0,0,1,0,0,0,1],"quality":"Equal","isMain":false,"metricName":"# contigs (>= 25000 bp)"},{"values":[0,0,0,0,0,1,0,0,0,0,0,0,1],"quality":"Equal","isMain":false,"metricName":"# contigs (>= 50000 bp)"},{"values":[15867,6034,16664,4455,6952,77112,22877,21993,26900,7803,7803,3825,77772],"quality":"More is better","isMain":true,"metricName":"Largest contig"},{"values":[35776,21184,40766,29207,32648,78285,32253,33844,35608,33061,27637,13759,77772],"quality":"More is better","isMain":true,"metricName":"Total length"},{"values":[34605,18176,37921,20379,28953,78285,32253,31311,34255,31865,26154,8159,77772],"quality":"More is better","isMain":true,"metricName":"Total length (>= 1000 bp)"},{"values":[34605,11270,22599,0,12574,77112,29790,31311,34255,15095,7803,0,77772],"quality":"More is better","isMain":false,"metricName":"Total length (>= 5000 bp)"},{"values":[15867,0,16664,0,0,77112,22877,21993,26900,0,0,0,77772],"quality":"More is better","isMain":true,"metricName":"Total length (>= 10000 bp)"},{"values":[0,0,0,0,0,77112,0,0,26900,0,0,0,77772],"quality":"More is better","isMain":false,"metricName":"Total length (>= 25000 bp)"},{"values":[0,0,0,0,0,77112,0,0,0,0,0,0,77772],"quality":"More is better","isMain":true,"metricName":"Total length (>= 50000 bp)"},{"values":[9992,5236,5935,2800,4496,77112,22877,21993,26900,4182,4181,1569,77772],"quality":"More is better","isMain":false,"metricName":"N50"},{"values":[8746,2730,4013,920,1426,77112,6913,9318,26900,2747,1971,860,77772],"quality":"More is better","isMain":false,"metricName":"N75"},{"values":[2,2,2,4,3,1,1,1,1,3,3,3,1],"quality":"Less is better","isMain":false,"metricName":"L50"},{"values":[3,4,4,9,7,1,2,2,1,5,5,7,1],"quality":"Less is better","isMain":false,"metricName":"L75"},{"values":["46.56","45.32","46.84","46.25","46.39","47.09","46.33","46.40","46.38","46.45","46.22","43.72","47.09"],"quality":"Equal","isMain":false,"metricName":"GC (%)"}]],["Predicted genes",[]],["Similarity statistics",[{"values":[0,0,0,0,0,0,0,0,0,0,0,0,0],"quality":"Equal","isMain":false,"metricName":"# similar correct contigs"},{"values":[0,0,0,0,0,0,0,0,0,0,0,0,0],"quality":"Equal","isMain":false,"metricName":"# similar misassembled blocks"}]],["Reference statistics",[{"values":[38297,38297,38297,38297,38297,38297,38297,38297,38297,38297,38297,38297,38297],"quality":"Equal","isMain":false,"metricName":"Reference length"},{"values":[1,1,1,1,1,1,1,1,1,1,1,1,1],"quality":"Equal","isMain":false,"metricName":"Reference fragments"},{"values":["46.69","46.69","46.69","46.69","46.69","46.69","46.69","46.69","46.69","46.69","46.69","46.69","46.69"],"quality":"Equal","isMain":false,"metricName":"Reference GC (%)"}]]],"referenceName":"gi\_13559823\_ref\_NC","date":"18 July 2018, Wednesday, 16:18:24","order":[0,1,2,3,4,5,6,7,8,9,10,11,12],"assembliesNames":["ABySS\_127","ABySS\_63","CLC","IDBA\_UD","MEGAHIT","MIRA","Ray\_Meta","SPAdes","SPAdes\_meta","SPAdes\_sc","SPAdes\_sc\_careful","Velvet","Geneious"]},{"assembliesWithNs":null,"minContig":500,"report":[["Genome statistics",[{"values":["89.075","42.270","94.627","72.212","88.568","99.788","64.675","7.688","91.839","69.838","53.097","47.747","22.975","99.788"],"quality":"More is better","isMain":true,"metricName":"Genome fraction (%)"},{"values":["1.017","1.003","1.002","1.023","1.084","1.296","1.040","1.000","1.000","0.999","1.003","1.000","1.000","1.095"],"quality":"Less is better","isMain":true,"metricName":"Duplication ratio"},{"values":[6399,5219,10783,3387,5409,10325,5274,1233,11022,16933,7523,7523,2969,39826],"quality":"More is better","isMain":true,"metricName":"Largest alignment"},{"values":[34111,15971,35658,27818,36171,48585,25342,2896,34593,26270,20058,17985,8654,41155],"quality":"More is better","isMain":true,"metricName":"Total aligned length"},{"values":[4519,null,8554,818,1798,10231,1254,null,9881,9651,762,null,null,69446],"quality":"More is better","isMain":false,"metricName":"NG50"},{"values":[4193,null,4995,null,897,6976,null,null,4773,null,null,null,null,69446],"quality":"More is better","isMain":false,"metricName":"NG75"},{"values":[4400,1703,8514,1491,1798,6974,1619,592,9881,16933,4655,4655,1504,39826],"quality":"More is better","isMain":false,"metricName":"NA50"},{"values":[1421,886,8343,792,901,2823,1254,544,5058,8148,3518,3515,808,null],"quality":"More is better","isMain":false,"metricName":"NA75"},{"values":[4400,null,8514,818,1798,10231,1254,null,9881,8148,762,null,null,39826],"quality":"More is better","isMain":true,"metricName":"NGA50"},{"values":[1421,null,4995,null,897,6490,null,null,4773,null,null,null,null,39826],"quality":"More is better","isMain":false,"metricName":"NGA75"},{"values":[3,null,2,12,6,2,9,null,2,2,6,null,null,1],"quality":"Less is better","isMain":false,"metricName":"LG50"},{"values":[5,null,4,null,14,3,null,null,4,null,null,null,null,1],"quality":"Less is better","isMain":false,"metricName":"LG75"},{"values":[4,3,2,7,6,3,4,2,2,1,2,2,2,1],"quality":"Less is better","isMain":false,"metricName":"LA50"},{"values":[7,6,3,15,13,6,9,3,3,2,3,3,5,null],"quality":"Less is better","isMain":false,"metricName":"LA75"},{"values":[4,null,2,12,6,2,9,null,2,2,6,null,null,1],"quality":"Less is better","isMain":true,"metricName":"LGA50"},{"values":[7,null,4,null,14,4,null,null,4,null,null,null,null,1],"quality":"Less is better","isMain":false,"metricName":"LGA75"}]],["Misassemblies",[{"values":[0,0,0,0,0,0,0,0,0,0,0,0,0,1],"quality":"Less is better","isMain":true,"metricName":"# misassemblies"},{"values":[0,0,0,0,0,0,0,0,0,0,0,0,0,1],"quality":"Less is better","isMain":false,"metricName":" # relocations"},{"values":[0,0,0,0,0,0,0,0,0,0,0,0,0,0],"quality":"Less is better","isMain":false,"metricName":" # translocations"},{"values":[0,0,0,0,0,0,0,0,0,0,0,0,0,0],"quality":"Less is better","isMain":false,"metricName":" # inversions"},{"values":[0,0,0,0,0,0,0,0,0,0,0,0,0,1],"quality":"Less is better","isMain":false,"metricName":"# misassembled contigs"},{"values":[0,0,0,0,0,0,0,0,0,0,0,0,0,69446],"quality":"Less is better","isMain":true,"metricName":"Misassembled contigs length"},{"values":[0,0,0,0,0,0,0,0,0,1,0,0,0,3],"quality":"Less is better","isMain":false,"metricName":"# local misassemblies"},{"values":[0,0,0,0,0,0,0,0,0,0,0,0,0,0],"quality":"Less is better","isMain":false,"metricName":"# unaligned mis. contigs"}]],["Unaligned",[{"values":[0,0,0,0,0,0,0,0,0,0,0,0,0,0],"quality":"Less is better","isMain":false,"metricName":"# fully unaligned contigs"},{"values":[0,0,0,0,0,0,0,0,0,0,0,0,0,0],"quality":"Less is better","isMain":false,"metricName":"Fully unaligned length"},{"values":[1,0,0,0,0,1,0,0,0,1,0,0,0,1],"quality":"Less is better","isMain":false,"metricName":"# partially unaligned contigs"},{"values":[3357,0,0,0,0,3775,0,0,0,1503,0,0,0,28291],"quality":"Less is better","isMain":false,"metricName":"Partially unaligned length"}]],["Mismatches",[{"values":[73,36,17,2,80,25,0,0,13,365,83,2,0,0],"quality":"Less is better","isMain":false,"metricName":"# mismatches"},{"values":[0,2,0,0,0,1,1,0,0,7,1,0,0,3],"quality":"Less is better","isMain":false,"metricName":"# indels"},{"values":[0,3,0,0,0,80,80,0,0,38,3,0,0,82],"quality":"Less is better","isMain":false,"metricName":"Indels length"},{"values":["217.57","226.10","47.70","7.35","239.80","66.51","0.00","0.00","37.58","1387.52","415.00","11.12","0.00","0.00"],"quality":"Less is better","isMain":true,"metricName":"# mismatches per 100 kbp"},{"values":["0.00","12.56","0.00","0.00","0.00","2.66","4.10","0.00","0.00","26.61","5.00","0.00","0.00","7.98"],"quality":"Less is better","isMain":true,"metricName":"# indels per 100 kbp"},{"values":[0,2,0,0,0,0,0,0,0,6,1,0,0,2],"quality":"Less is better","isMain":false,"metricName":" # indels (<= 5 bp)"},{"values":[0,0,0,0,0,1,1,0,0,1,0,0,0,1],"quality":"Less is better","isMain":false,"metricName":" # indels (> 5 bp)"},{"values":[50,1,0,0,0,3,0,0,0,0,0,0,0,133],"quality":"Less is better","isMain":false,"metricName":"# N's"},{"values":["133.45","6.26","0.00","0.00","0.00","5.72","0.00","0.00","0.00","0.00","0.00","0.00","0.00","191.52"],"quality":"Less is better","isMain":true,"metricName":"# N's per 100 kbp"}]],["Statistics without reference",[{"values":[13,12,7,25,25,15,15,4,7,4,8,6,7,1],"quality":"Equal","isMain":true,"metricName":"# contigs"},{"values":[8,4,5,10,10,8,13,1,5,2,5,4,3,1],"quality":"Equal","isMain":false,"metricName":"# contigs (>= 1000 bp)"},{"values":[2,1,3,0,1,4,1,0,3,2,1,1,0,1],"quality":"Equal","isMain":false,"metricName":"# contigs (>= 5000 bp)"},{"values":[0,0,1,0,0,2,0,0,1,1,0,0,0,1],"quality":"Equal","isMain":false,"metricName":"# contigs (>= 10000 bp)"},{"values":[0,0,0,0,0,0,0,0,0,0,0,0,0,1],"quality":"Equal","isMain":false,"metricName":"# contigs (>= 25000 bp)"},{"values":[0,0,0,0,0,0,0,0,0,0,0,0,0,1],"quality":"Equal","isMain":false,"metricName":"# contigs (>= 50000 bp)"},{"values":[9756,5219,10783,3387,5409,14100,5274,1233,11022,16933,7523,7523,2969,69446],"quality":"More is better","isMain":true,"metricName":"Largest contig"},{"values":[37468,15971,35698,27818,36171,52487,25342,2896,34593,27773,20058,17985,8654,69446],"quality":"More is better","isMain":true,"metricName":"Total length"},{"values":[33847,10204,34051,17370,25061,47459,23981,1233,32989,26584,18159,16796,5578,69446],"quality":"More is better","isMain":true,"metricName":"Total length (>= 1000 bp)"},{"values":[15365,5219,27680,0,5409,37869,5274,0,25961,26584,7523,7523,0,69446],"quality":"More is better","isMain":false,"metricName":"Total length (>= 5000 bp)"},{"values":[0,0,10783,0,0,24331,0,0,11022,16933,0,0,0,69446],"quality":"More is better","isMain":true,"metricName":"Total length (>= 10000 bp)"},{"values":[0,0,0,0,0,0,0,0,0,0,0,0,0,69446],"quality":"More is better","isMain":false,"metricName":"Total length (>= 25000 bp)"},{"values":[0,0,0,0,0,0,0,0,0,0,0,0,0,69446],"quality":"More is better","isMain":true,"metricName":"Total length (>= 50000 bp)"},{"values":[4519,1703,8554,1491,1798,6976,1619,592,9881,16933,4655,4655,1504,69446],"quality":"More is better","isMain":false,"metricName":"N50"},{"values":[4193,886,8343,792,901,4335,1254,544,5058,9651,3518,3515,808,69446],"quality":"More is better","isMain":false,"metricName":"N75"},{"values":[3,3,2,7,6,3,4,2,2,1,2,2,2,1],"quality":"Less is better","isMain":false,"metricName":"L50"},{"values":[5,6,3,15,13,5,9,3,3,2,3,3,5,1],"quality":"Less is better","isMain":false,"metricName":"L75"},{"values":["35.74","32.74","35.39","35.01","35.36","35.51","34.92","29.80","35.41","35.52","34.30","33.85","31.66","35.31"],"quality":"Equal","isMain":false,"metricName":"GC (%)"}]],["Predicted genes",[]],["Similarity statistics",[{"values":[0,0,0,0,0,0,0,0,0,0,0,0,0,0],"quality":"Equal","isMain":false,"metricName":"# similar correct contigs"},{"values":[0,0,0,0,0,0,0,0,0,0,0,0,0,0],"quality":"Equal","isMain":false,"metricName":"# similar misassembled blocks"}]],["Reference statistics",[{"values":[37667,37667,37667,37667,37667,37667,37667,37667,37667,37667,37667,37667,37667,37667],"quality":"Equal","isMain":false,"metricName":"Reference length"},{"values":[1,1,1,1,1,1,1,1,1,1,1,1,1,1],"quality":"Equal","isMain":false,"metricName":"Reference fragments"},{"values":["35.38","35.38","35.38","35.38","35.38","35.38","35.38","35.38","35.38","35.38","35.38","35.38","35.38","35.38"],"quality":"Equal","isMain":false,"metricName":"Reference GC (%)"}]]],"referenceName":"gi\_13786531\_ref\_NC","date":"18 July 2018, Wednesday, 16:18:36","order":[0,1,2,3,4,5,6,7,8,9,10,11,12,13],"assembliesNames":["ABySS\_127","ABySS\_63","CLC","IDBA\_UD","MEGAHIT","MIRA","Ray\_Meta","SOAPdenovo2","SPAdes","SPAdes\_meta","SPAdes\_sc","SPAdes\_sc\_careful","Velvet","Geneious"]},{"assembliesWithNs":null,"minContig":500,"report":[["Genome statistics",[{"values":["61.707","48.500","56.607","54.144","70.598","74.244","0.333","30.861","50.994","75.248","59.296","68.846","53.609","56.506"],"quality":"More is better","isMain":true,"metricName":"Genome fraction (%)"},{"values":["1.009","1.001","1.019","1.003","1.010","1.056","1.000","1.037","1.007","1.001","1.011","1.013","1.003","1.000"],"quality":"Less is better","isMain":true,"metricName":"Duplication ratio"},{"values":[6176,7103,12031,43859,18417,44589,576,4455,44716,44356,44235,43204,20195,45880],"quality":"More is better","isMain":true,"metricName":"Largest alignment"},{"values":[107524,83909,99313,93821,123189,135409,576,53745,88725,130143,102773,119209,92871,97657],"quality":"More is better","isMain":true,"metricName":"Total aligned length"},{"values":[1386,null,1592,3168,7293,7209,null,null,1655,23930,7379,7496,1054,8032],"quality":"More is better","isMain":false,"metricName":"NG50"},{"values":[null,null,null,null,null,834,null,null,null,1585,null,null,null,null],"quality":"More is better","isMain":false,"metricName":"NG75"},{"values":[2371,2229,4153,16858,8419,10895,576,1129,44716,33233,11854,8897,11372,13587],"quality":"More is better","isMain":false,"metricName":"NA50"},{"values":[1484,1490,2292,7707,4707,4729,576,null,9314,23930,7059,7029,7766,9294],"quality":"More is better","isMain":false,"metricName":"NA75"},{"values":[1386,null,1157,1634,5936,7209,null,null,1655,23930,6934,7029,1054,8032],"quality":"More is better","isMain":true,"metricName":"NGA50"},{"values":[null,null,null,null,null,834,null,null,null,1009,null,null,null,null],"quality":"More is better","isMain":false,"metricName":"NGA75"},{"values":[32,null,19,6,8,5,null,null,6,3,5,5,12,5],"quality":"Less is better","isMain":false,"metricName":"LG50"},{"values":[null,null,null,null,null,22,null,null,null,9,null,null,null,null],"quality":"Less is better","isMain":false,"metricName":"LG75"},{"values":[14,12,7,2,5,3,1,17,1,2,2,3,3,2],"quality":"Less is better","isMain":false,"metricName":"LA50"},{"values":[28,23,16,4,10,7,1,null,3,3,5,7,6,4],"quality":"Less is better","isMain":false,"metricName":"LA75"},{"values":[32,null,21,8,8,5,null,null,6,3,6,7,12,5],"quality":"Less is better","isMain":true,"metricName":"LGA50"},{"values":[null,null,null,null,null,22,null,null,null,11,null,null,null,null],"quality":"Less is better","isMain":false,"metricName":"LGA75"}]],["Misassemblies",[{"values":[0,0,0,0,0,0,0,0,0,0,1,1,0,1],"quality":"Less is better","isMain":true,"metricName":"# misassemblies"},{"values":[0,0,0,0,0,0,0,0,0,0,1,1,0,1],"quality":"Less is better","isMain":false,"metricName":" # relocations"},{"values":[0,0,0,0,0,0,0,0,0,0,0,0,0,0],"quality":"Less is better","isMain":false,"metricName":" # translocations"},{"values":[0,0,0,0,0,0,0,0,0,0,0,0,0,0],"quality":"Less is better","isMain":false,"metricName":" # inversions"},{"values":[0,0,0,0,0,0,0,0,0,0,1,1,0,1],"quality":"Less is better","isMain":false,"metricName":"# misassembled contigs"},{"values":[0,0,0,0,0,0,0,0,0,0,19234,19235,0,46538],"quality":"Less is better","isMain":true,"metricName":"Misassembled contigs length"},{"values":[0,0,0,0,0,0,0,2,0,2,2,2,1,1],"quality":"Less is better","isMain":false,"metricName":"# local misassemblies"},{"values":[0,0,1,0,0,0,0,1,0,0,0,0,0,0],"quality":"Less is better","isMain":false,"metricName":"# unaligned mis. contigs"}]],["Unaligned",[{"values":[0,0,0,0,0,0,0,0,0,0,0,0,0,0],"quality":"Less is better","isMain":false,"metricName":"# fully unaligned contigs"},{"values":[0,0,0,0,0,0,0,0,0,0,0,0,0,0],"quality":"Less is better","isMain":false,"metricName":"Fully unaligned length"},{"values":[0,0,1,1,1,0,0,10,0,1,1,0,0,0],"quality":"Less is better","isMain":false,"metricName":"# partially unaligned contigs"},{"values":[0,0,2065,3275,4269,0,0,16602,0,2610,1139,0,0,0],"quality":"Less is better","isMain":false,"metricName":"Partially unaligned length"}]],["Mismatches",[{"values":[104,79,264,214,264,205,0,42,267,385,263,240,125,105],"quality":"Less is better","isMain":false,"metricName":"# mismatches"},{"values":[4,2,9,5,25,11,0,23,25,28,16,15,4,36],"quality":"Less is better","isMain":false,"metricName":"# indels"},{"values":[35,2,12,20,80,90,0,1036,188,247,178,174,9,114],"quality":"Less is better","isMain":false,"metricName":"Indels length"},{"values":["97.55","94.28","269.95","228.77","216.45","159.82","0.00","78.77","303.07","296.15","256.73","201.78","134.96","107.56"],"quality":"Less is better","isMain":true,"metricName":"# mismatches per 100 kbp"},{"values":["3.75","2.39","9.20","5.35","20.50","8.58","0.00","43.14","28.38","21.54","15.62","12.61","4.32","36.88"],"quality":"Less is better","isMain":true,"metricName":"# indels per 100 kbp"},{"values":[3,2,9,4,23,9,0,2,19,20,13,12,4,32],"quality":"Less is better","isMain":false,"metricName":" # indels (<= 5 bp)"},{"values":[1,0,0,1,2,2,0,21,6,8,3,3,0,4],"quality":"Less is better","isMain":false,"metricName":" # indels (> 5 bp)"},{"values":[0,1,0,0,0,124,0,5057,0,120,1915,1297,10,435],"quality":"Less is better","isMain":false,"metricName":"# N's"},{"values":["0.00","1.19","0.00","0.00","0.00","91.52","0.00","7032.11","0.00","90.39","1829.24","1076.27","10.77","445.44"],"quality":"Less is better","isMain":true,"metricName":"# N's per 100 kbp"}]],["Statistics without reference",[{"values":[57,48,39,14,23,29,1,45,8,11,15,20,20,6],"quality":"Equal","isMain":true,"metricName":"# contigs"},{"values":[40,29,22,11,22,17,0,24,7,11,12,15,12,6],"quality":"Equal","isMain":false,"metricName":"# contigs (>= 1000 bp)"},{"values":[4,1,5,5,9,6,0,2,5,5,5,7,6,6],"quality":"Equal","isMain":false,"metricName":"# contigs (>= 5000 bp)"},{"values":[0,0,2,2,4,3,0,0,2,3,2,2,3,3],"quality":"Equal","isMain":false,"metricName":"# contigs (>= 10000 bp)"},{"values":[0,0,0,1,0,1,0,0,1,2,1,1,0,1],"quality":"Equal","isMain":false,"metricName":"# contigs (>= 25000 bp)"},{"values":[0,0,0,0,0,0,0,0,0,0,0,0,0,0],"quality":"Equal","isMain":false,"metricName":"# contigs (>= 50000 bp)"},{"values":[6176,7103,12031,43859,18417,44590,576,5555,44723,44356,44235,43376,20195,46538],"quality":"More is better","isMain":true,"metricName":"Largest contig"},{"values":[107524,83909,101749,97096,127461,135491,576,71913,88732,132753,104688,120509,92885,97657],"quality":"More is better","isMain":true,"metricName":"Total length"},{"values":[96130,70809,89994,95147,126492,125899,0,56904,88067,132753,102479,116899,86898,97657],"quality":"More is better","isMain":true,"metricName":"Total length (>= 1000 bp)"},{"values":[23524,7103,44121,83432,93769,97482,0,10999,84819,122930,88296,101560,74863,97657],"quality":"More is better","isMain":false,"metricName":"Total length (>= 5000 bp)"},{"values":[0,0,23590,60717,57071,76738,0,0,60514,104129,63469,62611,50502,73122],"quality":"More is better","isMain":true,"metricName":"Total length (>= 10000 bp)"},{"values":[0,0,0,43859,0,44590,0,0,44723,80199,44235,43376,0,46538],"quality":"More is better","isMain":false,"metricName":"Total length (>= 25000 bp)"},{"values":[0,0,0,0,0,0,0,0,0,0,0,0,0,0],"quality":"More is better","isMain":true,"metricName":"Total length (>= 50000 bp)"},{"values":[2371,2229,4786,16858,8419,10895,576,2258,44723,35843,19234,19235,11383,13587],"quality":"More is better","isMain":false,"metricName":"N50"},{"values":[1484,1490,2308,7707,4798,4751,576,1120,9314,23930,7496,7379,7769,9294],"quality":"More is better","isMain":false,"metricName":"N75"},{"values":[14,12,7,2,5,3,1,10,1,2,2,2,3,2],"quality":"Less is better","isMain":false,"metricName":"L50"},{"values":[28,23,15,4,10,7,1,22,3,3,4,6,6,4],"quality":"Less is better","isMain":false,"metricName":"L75"},{"values":["58.03","57.69","57.47","56.98","57.56","58.14","84.55","57.13","56.17","57.81","56.56","56.97","56.96","57.37"],"quality":"Equal","isMain":false,"metricName":"GC (%)"}]],["Predicted genes",[]],["Similarity statistics",[{"values":[0,0,0,0,0,0,0,0,0,0,0,0,0,0],"quality":"Equal","isMain":false,"metricName":"# similar correct contigs"},{"values":[0,0,0,0,0,0,0,0,0,0,0,0,0,0],"quality":"Equal","isMain":false,"metricName":"# similar misassembled blocks"}]],["Reference statistics",[{"values":[172764,172764,172764,172764,172764,172764,172764,172764,172764,172764,172764,172764,172764,172764],"quality":"Equal","isMain":false,"metricName":"Reference length"},{"values":[1,1,1,1,1,1,1,1,1,1,1,1,1,1],"quality":"Equal","isMain":false,"metricName":"Reference fragments"},{"values":["59.54","59.54","59.54","59.54","59.54","59.54","59.54","59.54","59.54","59.54","59.54","59.54","59.54","59.54"],"quality":"Equal","isMain":false,"metricName":"Reference GC (%)"}]]],"referenceName":"gi\_139424470\_ref\_NC","date":"18 July 2018, Wednesday, 16:18:49","order":[0,1,2,3,4,5,6,7,8,9,10,11,12,13],"assembliesNames":["ABySS\_127","ABySS\_63","CLC","IDBA\_UD","MEGAHIT","MIRA","Ray\_Meta","SOAPdenovo2","SPAdes","SPAdes\_meta","SPAdes\_sc","SPAdes\_sc\_careful","Velvet","Geneious"]},{"assembliesWithNs":null,"minContig":500,"report":[["Genome statistics",[{"values":["99.724","98.658","98.657","98.532","98.334","100.000","96.504","47.751","99.099","98.980","99.123","99.123","96.225","99.827"],"quality":"More is better","isMain":true,"metricName":"Genome fraction (%)"},{"values":["1.006","1.000","1.000","1.002","1.002","1.004","1.037","1.000","1.006","1.000","1.001","1.001","1.000","1.002"],"quality":"Less is better","isMain":true,"metricName":"Duplication ratio"},{"values":[95546,93824,93604,88197,93182,94284,53979,1566,95171,93228,93517,93517,62784,138015],"quality":"More is better","isMain":true,"metricName":"Largest alignment"},{"values":[138339,136147,136045,136211,135952,138574,138007,65904,136768,136513,136856,136856,132791,138015],"quality":"More is better","isMain":true,"metricName":"Total aligned length"},{"values":[95596,93824,93702,88197,93182,94284,32225,null,95940,93313,93517,93517,34670,138015],"quality":"More is better","isMain":false,"metricName":"NG50"},{"values":[36062,35623,35935,35561,35393,36610,25560,null,41081,42509,35862,35862,25160,138015],"quality":"More is better","isMain":false,"metricName":"NG75"},{"values":[95546,93824,93604,88197,93182,94284,32225,790,95171,93228,93517,93517,34670,138015],"quality":"More is better","isMain":false,"metricName":"NA50"},{"values":[36062,35623,35935,35561,35393,36610,25537,663,41081,42509,35862,35862,25160,138015],"quality":"More is better","isMain":false,"metricName":"NA75"},{"values":[95546,93824,93604,88197,93182,94284,32225,null,95171,93228,93517,93517,34670,138015],"quality":"More is better","isMain":true,"metricName":"NGA50"},{"values":[36062,35623,35935,35561,35393,36610,25537,null,41081,42509,35862,35862,25160,138015],"quality":"More is better","isMain":false,"metricName":"NGA75"},{"values":[1,1,1,1,1,1,2,null,1,1,1,1,2,1],"quality":"Less is better","isMain":false,"metricName":"LG50"},{"values":[2,2,2,2,2,2,3,null,2,2,2,2,3,1],"quality":"Less is better","isMain":false,"metricName":"LG75"},{"values":[1,1,1,1,1,1,2,31,1,1,1,1,2,1],"quality":"Less is better","isMain":false,"metricName":"LA50"},{"values":[2,2,2,2,2,2,3,55,2,2,2,2,3,1],"quality":"Less is better","isMain":false,"metricName":"LA75"},{"values":[1,1,1,1,1,1,2,null,1,1,1,1,2,1],"quality":"Less is better","isMain":true,"metricName":"LGA50"},{"values":[2,2,2,2,2,2,3,null,2,2,2,2,3,1],"quality":"Less is better","isMain":false,"metricName":"LGA75"}]],["Misassemblies",[{"values":[0,0,0,0,0,0,1,0,0,0,0,0,0,0],"quality":"Less is better","isMain":true,"metricName":"# misassemblies"},{"values":[0,0,0,0,0,0,1,0,0,0,0,0,0,0],"quality":"Less is better","isMain":false,"metricName":" # relocations"},{"values":[0,0,0,0,0,0,0,0,0,0,0,0,0,0],"quality":"Less is better","isMain":false,"metricName":" # translocations"},{"values":[0,0,0,0,0,0,0,0,0,0,0,0,0,0],"quality":"Less is better","isMain":false,"metricName":" # inversions"},{"values":[0,0,0,0,0,0,1,0,0,0,0,0,0,0],"quality":"Less is better","isMain":false,"metricName":"# misassembled contigs"},{"values":[0,0,0,0,0,0,6481,0,0,0,0,0,0,0],"quality":"Less is better","isMain":true,"metricName":"Misassembled contigs length"},{"values":[1,0,0,1,1,0,2,0,0,1,0,0,0,1],"quality":"Less is better","isMain":false,"metricName":"# local misassemblies"},{"values":[0,0,0,0,0,0,0,0,0,0,0,0,0,0],"quality":"Less is better","isMain":false,"metricName":"# unaligned mis. contigs"}]],["Unaligned",[{"values":[0,0,0,0,0,0,0,0,0,0,0,0,0,0],"quality":"Less is better","isMain":false,"metricName":"# fully unaligned contigs"},{"values":[0,0,0,0,0,0,0,0,0,0,0,0,0,0],"quality":"Less is better","isMain":false,"metricName":"Fully unaligned length"},{"values":[0,0,0,0,0,0,0,0,0,0,0,0,0,0],"quality":"Less is better","isMain":false,"metricName":"# partially unaligned contigs"},{"values":[0,0,0,0,0,0,0,0,0,0,0,0,0,0],"quality":"Less is better","isMain":false,"metricName":"Partially unaligned length"}]],["Mismatches",[{"values":[1,1,15,1,1,2,19,2,4,40,0,0,0,1],"quality":"Less is better","isMain":false,"metricName":"# mismatches"},{"values":[2,4,15,5,2,3,14,0,8,17,3,3,0,1],"quality":"Less is better","isMain":false,"metricName":"# indels"},{"values":[146,30,105,93,23,68,171,0,223,62,37,37,0,1],"quality":"Less is better","isMain":false,"metricName":"Indels length"},{"values":["0.73","0.73","11.02","0.74","0.74","1.45","14.27","3.04","2.93","29.29","0.00","0.00","0.00","0.73"],"quality":"Less is better","isMain":true,"metricName":"# mismatches per 100 kbp"},{"values":["1.45","2.94","11.02","3.68","1.47","2.17","10.51","0.00","5.85","12.45","2.19","2.19","0.00","0.73"],"quality":"Less is better","isMain":true,"metricName":"# indels per 100 kbp"},{"values":[0,2,13,1,1,0,7,0,2,15,1,1,0,1],"quality":"Less is better","isMain":false,"metricName":" # indels (<= 5 bp)"},{"values":[2,2,2,4,1,3,7,0,6,2,2,2,0,0],"quality":"Less is better","isMain":false,"metricName":" # indels (> 5 bp)"},{"values":[50,50,0,0,0,0,0,0,0,0,0,0,0,0],"quality":"Less is better","isMain":false,"metricName":"# N's"},{"values":["36.13","36.73","0.00","0.00","0.00","0.00","0.00","0.00","0.00","0.00","0.00","0.00","0.00","0.00"],"quality":"Less is better","isMain":true,"metricName":"# N's per 100 kbp"}]],["Statistics without reference",[{"values":[4,5,3,7,6,4,8,83,3,3,6,6,6,1],"quality":"Equal","isMain":true,"metricName":"# contigs"},{"values":[3,3,3,4,4,3,8,16,2,2,4,4,5,1],"quality":"Equal","isMain":false,"metricName":"# contigs (>= 1000 bp)"},{"values":[3,3,3,4,2,3,6,0,2,2,2,2,3,1],"quality":"Equal","isMain":false,"metricName":"# contigs (>= 5000 bp)"},{"values":[2,2,2,2,2,2,3,0,2,2,2,2,3,1],"quality":"Equal","isMain":false,"metricName":"# contigs (>= 10000 bp)"},{"values":[2,2,2,2,2,2,3,0,2,2,2,2,3,1],"quality":"Equal","isMain":false,"metricName":"# contigs (>= 25000 bp)"},{"values":[1,1,1,1,1,1,1,0,1,1,1,1,1,1],"quality":"Equal","isMain":false,"metricName":"# contigs (>= 50000 bp)"},{"values":[95596,93824,93702,88197,93182,94284,53979,1566,95940,93313,93517,93517,62784,138015],"quality":"More is better","isMain":true,"metricName":"Largest contig"},{"values":[138389,136147,136169,136211,135952,138574,138110,65904,137537,136598,136856,136856,132791,138015],"quality":"More is better","isMain":true,"metricName":"Total length"},{"values":[137491,134877,136169,133889,134513,137651,138110,20168,137021,135822,135584,135584,132275,138015],"quality":"More is better","isMain":true,"metricName":"Total length (>= 1000 bp)"},{"values":[137491,134877,136169,133889,128575,137651,132039,0,137021,135822,129379,129379,122614,138015],"quality":"More is better","isMain":false,"metricName":"Total length (>= 5000 bp)"},{"values":[131658,129447,129637,123758,128575,130894,111764,0,137021,135822,129379,129379,122614,138015],"quality":"More is better","isMain":true,"metricName":"Total length (>= 10000 bp)"},{"values":[131658,129447,129637,123758,128575,130894,111764,0,137021,135822,129379,129379,122614,138015],"quality":"More is better","isMain":false,"metricName":"Total length (>= 25000 bp)"},{"values":[95596,93824,93702,88197,93182,94284,53979,0,95940,93313,93517,93517,62784,138015],"quality":"More is better","isMain":true,"metricName":"Total length (>= 50000 bp)"},{"values":[95596,93824,93702,88197,93182,94284,32225,790,95940,93313,93517,93517,34670,138015],"quality":"More is better","isMain":false,"metricName":"N50"},{"values":[36062,35623,35935,35561,35393,36610,25560,663,41081,42509,35862,35862,25160,138015],"quality":"More is better","isMain":false,"metricName":"N75"},{"values":[1,1,1,1,1,1,2,31,1,1,1,1,2,1],"quality":"Less is better","isMain":false,"metricName":"L50"},{"values":[2,2,2,2,2,2,3,55,2,2,2,2,3,1],"quality":"Less is better","isMain":false,"metricName":"L75"},{"values":["53.68","53.57","53.55","53.51","53.50","53.84","53.57","53.19","53.68","53.60","53.64","53.64","53.33","53.74"],"quality":"Equal","isMain":false,"metricName":"GC (%)"}]],["Predicted genes",[]],["Similarity statistics",[{"values":[2,2,1,2,2,2,1,0,1,1,2,2,1,1],"quality":"Equal","isMain":false,"metricName":"# similar correct contigs"},{"values":[0,0,0,0,0,0,0,0,0,0,0,0,0,0],"quality":"Equal","isMain":false,"metricName":"# similar misassembled blocks"}]],["Reference statistics",[{"values":[137969,137969,137969,137969,137969,137969,137969,137969,137969,137969,137969,137969,137969,137969],"quality":"Equal","isMain":false,"metricName":"Reference length"},{"values":[1,1,1,1,1,1,1,1,1,1,1,1,1,1],"quality":"Equal","isMain":false,"metricName":"Reference fragments"},{"values":["53.76","53.76","53.76","53.76","53.76","53.76","53.76","53.76","53.76","53.76","53.76","53.76","53.76","53.76"],"quality":"Equal","isMain":false,"metricName":"Reference GC (%)"}]]],"referenceName":"gi\_139472801\_ref\_NC","date":"18 July 2018, Wednesday, 16:19:02","order":[0,1,2,3,4,5,6,7,8,9,10,11,12,13],"assembliesNames":["ABySS\_127","ABySS\_63","CLC","IDBA\_UD","MEGAHIT","MIRA","Ray\_Meta","SOAPdenovo2","SPAdes","SPAdes\_meta","SPAdes\_sc","SPAdes\_sc\_careful","Velvet","Geneious"]},{"assembliesWithNs":null,"minContig":500,"report":[["Genome statistics",[{"values":["97.759","94.961","99.362","99.628","99.709","98.936","91.537","99.572","99.867","99.587","99.548","99.027","56.376"],"quality":"More is better","isMain":true,"metricName":"Genome fraction (%)"},{"values":["1.001","1.002","1.009","1.002","1.000","1.012","1.027","1.000","1.000","0.999","1.000","1.001","1.000"],"quality":"Less is better","isMain":true,"metricName":"Duplication ratio"},{"values":[16248,13761,100872,58345,133675,40440,31270,128295,128253,127980,126501,43761,48463],"quality":"More is better","isMain":true,"metricName":"Largest alignment"},{"values":[191750,186280,196277,195162,195359,196197,181001,195021,195558,194891,194936,194177,110418],"quality":"More is better","isMain":true,"metricName":"Total aligned length"},{"values":[6872,5562,100872,49042,133675,18584,11558,128295,128253,127980,126501,14727,21515],"quality":"More is better","isMain":false,"metricName":"NG50"},{"values":[2890,2454,14670,26337,26651,13640,3878,26805,26805,26769,26769,7511,null],"quality":"More is better","isMain":false,"metricName":"NG75"},{"values":[6872,5562,100872,49042,133675,18584,8432,128295,128253,127980,126501,14727,40440],"quality":"More is better","isMain":false,"metricName":"NA50"},{"values":[3368,2695,14670,26337,26651,13640,3343,26805,26805,26769,26769,9890,40440],"quality":"More is better","isMain":false,"metricName":"NA75"},{"values":[6872,5562,100872,49042,133675,18584,8264,128295,128253,127980,126501,14727,21515],"quality":"More is better","isMain":true,"metricName":"NGA50"},{"values":[2890,2454,14670,26337,26651,13640,3331,26805,26805,26769,26769,7511,null],"quality":"More is better","isMain":false,"metricName":"NGA75"},{"values":[9,10,1,2,1,4,6,1,1,1,1,5,3],"quality":"Less is better","isMain":false,"metricName":"LG50"},{"values":[20,24,4,4,2,7,13,2,2,2,2,9,null],"quality":"Less is better","isMain":false,"metricName":"LG75"},{"values":[9,10,1,2,1,4,6,1,1,1,1,5,2],"quality":"Less is better","isMain":false,"metricName":"LA50"},{"values":[19,22,4,4,2,7,15,2,2,2,2,8,2],"quality":"Less is better","isMain":false,"metricName":"LA75"},{"values":[9,10,1,2,1,4,7,1,1,1,1,5,3],"quality":"Less is better","isMain":true,"metricName":"LGA50"},{"values":[20,24,4,4,2,7,16,2,2,2,2,9,null],"quality":"Less is better","isMain":false,"metricName":"LGA75"}]],["Misassemblies",[{"values":[0,0,0,0,0,0,0,0,0,0,0,0,0],"quality":"Less is better","isMain":true,"metricName":"# misassemblies"},{"values":[0,0,0,0,0,0,0,0,0,0,0,0,0],"quality":"Less is better","isMain":false,"metricName":" # relocations"},{"values":[0,0,0,0,0,0,0,0,0,0,0,0,0],"quality":"Less is better","isMain":false,"metricName":" # translocations"},{"values":[0,0,0,0,0,0,0,0,0,0,0,0,0],"quality":"Less is better","isMain":false,"metricName":" # inversions"},{"values":[0,0,0,0,0,0,0,0,0,0,0,0,0],"quality":"Less is better","isMain":false,"metricName":"# misassembled contigs"},{"values":[0,0,0,0,0,0,0,0,0,0,0,0,0],"quality":"Less is better","isMain":true,"metricName":"Misassembled contigs length"},{"values":[0,0,0,0,0,0,1,0,0,2,2,0,0],"quality":"Less is better","isMain":false,"metricName":"# local misassemblies"},{"values":[0,0,0,0,0,0,0,0,0,0,0,0,0],"quality":"Less is better","isMain":false,"metricName":"# unaligned mis. contigs"}]],["Unaligned",[{"values":[0,0,0,0,0,0,0,0,0,0,0,0,0],"quality":"Less is better","isMain":false,"metricName":"# fully unaligned contigs"},{"values":[0,0,0,0,0,0,0,0,0,0,0,0,0],"quality":"Less is better","isMain":false,"metricName":"Fully unaligned length"},{"values":[0,0,1,0,0,0,2,0,0,0,0,0,0],"quality":"Less is better","isMain":false,"metricName":"# partially unaligned contigs"},{"values":[0,0,1260,0,0,0,8494,0,0,0,0,0,0],"quality":"Less is better","isMain":false,"metricName":"Partially unaligned length"}]],["Mismatches",[{"values":[14,3,25,7,17,4,8,16,7,6,7,11,0],"quality":"Less is better","isMain":false,"metricName":"# mismatches"},{"values":[1,0,2,2,1,0,58,1,6,7,8,1,0],"quality":"Less is better","isMain":false,"metricName":"# indels"},{"values":[21,0,4,84,21,0,2797,21,141,117,180,12,0],"quality":"Less is better","isMain":false,"metricName":"Indels length"},{"values":["7.31","1.61","12.85","3.59","8.71","2.06","4.46","8.20","3.58","3.08","3.59","5.67","0.00"],"quality":"Less is better","isMain":true,"metricName":"# mismatches per 100 kbp"},{"values":["0.52","0.00","1.03","1.02","0.51","0.00","32.35","0.51","3.07","3.59","4.10","0.52","0.00"],"quality":"Less is better","isMain":true,"metricName":"# indels per 100 kbp"},{"values":[0,0,2,0,0,0,0,0,2,2,2,0,0],"quality":"Less is better","isMain":false,"metricName":" # indels (<= 5 bp)"},{"values":[1,0,0,2,1,0,58,1,4,5,6,1,0],"quality":"Less is better","isMain":false,"metricName":" # indels (> 5 bp)"},{"values":[0,0,0,0,0,24,5830,0,0,20,0,0,18],"quality":"Less is better","isMain":false,"metricName":"# N's"},{"values":["0.00","0.00","0.00","0.00","0.00","12.23","3026.89","0.00","0.00","10.26","0.00","0.00","16.30"],"quality":"Less is better","isMain":true,"metricName":"# N's per 100 kbp"}]],["Statistics without reference",[{"values":[47,51,15,7,5,15,34,5,4,4,5,20,3],"quality":"Equal","isMain":true,"metricName":"# contigs"},{"values":[40,45,9,6,4,14,32,5,4,4,5,18,3],"quality":"Equal","isMain":false,"metricName":"# contigs (>= 1000 bp)"},{"values":[12,11,6,6,4,10,9,5,4,4,4,13,3],"quality":"Equal","isMain":false,"metricName":"# contigs (>= 5000 bp)"},{"values":[5,5,6,6,4,10,6,4,4,4,4,7,3],"quality":"Equal","isMain":false,"metricName":"# contigs (>= 10000 bp)"},{"values":[0,0,2,4,2,2,1,2,2,2,2,1,2],"quality":"Equal","isMain":false,"metricName":"# contigs (>= 25000 bp)"},{"values":[0,0,1,1,1,0,0,1,1,1,1,0,0],"quality":"Equal","isMain":false,"metricName":"# contigs (>= 50000 bp)"},{"values":[16248,13761,100872,58345,133675,40440,31790,128295,128253,127980,126501,43761,48463],"quality":"More is better","isMain":true,"metricName":"Largest contig"},{"values":[191750,186280,197537,195451,195366,196197,192607,195021,195558,194891,194936,194177,110418],"quality":"More is better","isMain":true,"metricName":"Total length"},{"values":[186711,182034,193538,194858,194598,195520,191358,195021,195558,194891,194936,192830,110418],"quality":"More is better","isMain":true,"metricName":"Total length (>= 1000 bp)"},{"values":[117204,103313,188421,194858,194598,187087,129872,195021,195558,194891,193404,179602,110418],"quality":"More is better","isMain":false,"metricName":"Total length (>= 5000 bp)"},{"values":[66377,62429,188421,194858,194598,187087,105239,189384,195558,194891,193404,136635,110418],"quality":"More is better","isMain":true,"metricName":"Total length (>= 10000 bp)"},{"values":[0,0,127942,160529,160326,65677,31790,155100,155058,154749,153270,43761,88903],"quality":"More is better","isMain":false,"metricName":"Total length (>= 25000 bp)"},{"values":[0,0,100872,58345,133675,0,0,128295,128253,127980,126501,0,0],"quality":"More is better","isMain":true,"metricName":"Total length (>= 50000 bp)"},{"values":[6872,5562,100872,49042,133675,18584,11558,128295,128253,127980,126501,14727,40440],"quality":"More is better","isMain":false,"metricName":"N50"},{"values":[3368,2695,14670,26337,26651,13640,3878,26805,26805,26769,26769,9890,40440],"quality":"More is better","isMain":false,"metricName":"N75"},{"values":[9,10,1,2,1,4,6,1,1,1,1,5,2],"quality":"Less is better","isMain":false,"metricName":"L50"},{"values":[19,22,4,4,2,7,13,2,2,2,2,8,2],"quality":"Less is better","isMain":false,"metricName":"L75"},{"values":["66.64","66.60","66.55","66.55","66.60","66.63","66.48","66.56","66.60","66.58","66.58","66.54","65.62"],"quality":"Equal","isMain":false,"metricName":"GC (%)"}]],["Predicted genes",[]],["Similarity statistics",[{"values":[0,0,3,3,3,1,0,3,3,3,3,2,0],"quality":"Equal","isMain":false,"metricName":"# similar correct contigs"},{"values":[0,0,0,0,0,0,0,0,0,0,0,0,0],"quality":"Equal","isMain":false,"metricName":"# similar misassembled blocks"}]],["Reference statistics",[{"values":[195859,195859,195859,195859,195859,195859,195859,195859,195859,195859,195859,195859,195859],"quality":"Equal","isMain":false,"metricName":"Reference length"},{"values":[1,1,1,1,1,1,1,1,1,1,1,1,1],"quality":"Equal","isMain":false,"metricName":"Reference fragments"},{"values":["66.61","66.61","66.61","66.61","66.61","66.61","66.61","66.61","66.61","66.61","66.61","66.61","66.61"],"quality":"Equal","isMain":false,"metricName":"Reference GC (%)"}]]],"referenceName":"gi\_14250967\_ref\_NC","date":"18 July 2018, Wednesday, 16:19:14","order":[0,1,2,3,4,5,6,7,8,9,10,11,12],"assembliesNames":["ABySS\_127","ABySS\_63","CLC","IDBA\_UD","MEGAHIT","MIRA","SOAPdenovo2","SPAdes","SPAdes\_meta","SPAdes\_sc","SPAdes\_sc\_careful","Velvet","Geneious"]},{"assembliesWithNs":null,"minContig":500,"report":[["Genome statistics",[{"values":["74.537","51.109","78.434","64.413","68.030","97.835","51.699","15.119","88.783","35.187","57.458","56.741","29.008","1.282","98.770"],"quality":"More is better","isMain":true,"metricName":"Genome fraction (%)"},{"values":["1.011","1.002","1.009","1.020","1.014","1.098","1.028","1.016","1.034","1.029","1.000","1.000","1.000","1.000","1.576"],"quality":"Less is better","isMain":true,"metricName":"Duplication ratio"},{"values":[8372,5376,7719,4198,7708,12356,4814,1022,14680,4011,7319,7867,3656,513,14422],"quality":"More is better","isMain":true,"metricName":"Largest alignment"},{"values":[30148,20479,31626,26279,27589,42955,21261,6048,36632,14131,22982,22695,11604,513,62256],"quality":"More is better","isMain":true,"metricName":"Total aligned length"},{"values":[4927,588,3579,1156,2342,8918,809,null,11066,null,4197,4197,null,null,136335],"quality":"More is better","isMain":false,"metricName":"NG50"},{"values":[731,null,995,null,null,5238,null,null,7021,null,null,null,null,null,136335],"quality":"More is better","isMain":false,"metricName":"NG75"},{"values":[4927,3888,4543,2568,4368,8661,1554,700,5273,2361,4669,4669,1925,513,null],"quality":"More is better","isMain":false,"metricName":"NA50"},{"values":[733,1623,2435,1156,2342,5238,901,574,2387,1399,3398,3398,1179,513,null],"quality":"More is better","isMain":false,"metricName":"NA75"},{"values":[3275,588,3579,1156,2342,8918,809,null,5273,null,2209,3398,null,null,11645],"quality":"More is better","isMain":true,"metricName":"NGA50"},{"values":[529,null,995,null,null,5238,null,null,2652,null,null,null,null,null,10487],"quality":"More is better","isMain":false,"metricName":"NGA75"},{"values":[3,10,4,8,5,2,13,null,2,null,4,4,null,null,1],"quality":"Less is better","isMain":false,"metricName":"LG50"},{"values":[9,null,10,null,null,4,null,null,3,null,null,null,null,null,1],"quality":"Less is better","isMain":false,"metricName":"LG75"},{"values":[3,3,3,4,3,3,4,4,3,3,2,2,3,1,null],"quality":"Less is better","isMain":false,"metricName":"LA50"},{"values":[7,4,6,8,5,4,9,7,6,5,4,4,4,1,null],"quality":"Less is better","isMain":false,"metricName":"LA75"},{"values":[4,10,4,8,5,2,13,null,3,null,5,4,null,null,2],"quality":"Less is better","isMain":true,"metricName":"LGA50"},{"values":[14,null,10,null,null,4,null,null,5,null,null,null,null,null,3],"quality":"Less is better","isMain":false,"metricName":"LGA75"}]],["Misassemblies",[{"values":[0,0,0,0,0,0,0,0,0,0,0,0,0,0,1],"quality":"Less is better","isMain":true,"metricName":"# misassemblies"},{"values":[0,0,0,0,0,0,0,0,0,0,0,0,0,0,1],"quality":"Less is better","isMain":false,"metricName":" # relocations"},{"values":[0,0,0,0,0,0,0,0,0,0,0,0,0,0,0],"quality":"Less is better","isMain":false,"metricName":" # translocations"},{"values":[0,0,0,0,0,0,0,0,0,0,0,0,0,0,0],"quality":"Less is better","isMain":false,"metricName":" # inversions"},{"values":[0,0,0,0,0,0,0,0,0,0,0,0,0,0,1],"quality":"Less is better","isMain":false,"metricName":"# misassembled contigs"},{"values":[0,0,0,0,0,0,0,0,0,0,0,0,0,0,26308],"quality":"Less is better","isMain":true,"metricName":"Misassembled contigs length"},{"values":[0,0,0,0,0,0,0,0,1,1,0,0,0,0,3],"quality":"Less is better","isMain":false,"metricName":"# local misassemblies"},{"values":[0,0,0,0,0,0,0,0,0,0,0,0,0,0,2],"quality":"Less is better","isMain":false,"metricName":"# unaligned mis. contigs"}]],["Unaligned",[{"values":[0,0,0,0,0,0,0,0,0,0,0,0,0,0,0],"quality":"Less is better","isMain":false,"metricName":"# fully unaligned contigs"},{"values":[0,0,0,0,0,0,0,0,0,0,0,0,0,0,0],"quality":"Less is better","isMain":false,"metricName":"Fully unaligned length"},{"values":[1,0,0,0,0,0,0,0,2,0,1,1,0,0,4],"quality":"Less is better","isMain":false,"metricName":"# partially unaligned contigs"},{"values":[3650,0,0,0,0,0,0,0,7524,0,799,799,0,0,189461],"quality":"Less is better","isMain":false,"metricName":"Partially unaligned length"}]],["Mismatches",[{"values":[0,1,52,8,24,33,10,0,38,49,61,61,0,6,331],"quality":"Less is better","isMain":false,"metricName":"# mismatches"},{"values":[0,0,2,0,0,0,2,1,0,0,2,1,0,0,23],"quality":"Less is better","isMain":false,"metricName":"# indels"},{"values":[0,0,2,0,0,0,3,45,0,0,3,3,0,0,29],"quality":"Less is better","isMain":false,"metricName":"Indels length"},{"values":["0.00","4.89","165.73","31.05","88.19","84.32","48.35","0.00","106.99","348.11","265.39","268.75","0.00","1169.59","837.74"],"quality":"Less is better","isMain":true,"metricName":"# mismatches per 100 kbp"},{"values":["0.00","0.00","6.37","0.00","0.00","0.00","9.67","16.53","0.00","0.00","8.70","4.41","0.00","0.00","58.21"],"quality":"Less is better","isMain":true,"metricName":"# indels per 100 kbp"},{"values":[0,0,2,0,0,0,2,0,0,0,2,1,0,0,23],"quality":"Less is better","isMain":false,"metricName":" # indels (<= 5 bp)"},{"values":[0,0,0,0,0,0,0,1,0,0,0,0,0,0,0],"quality":"Less is better","isMain":false,"metricName":" # indels (> 5 bp)"},{"values":[50,0,0,0,0,0,0,98,100,0,0,0,0,0,366],"quality":"Less is better","isMain":false,"metricName":"# N's"},{"values":["147.94","0.00","0.00","0.00","0.00","0.00","0.00","1594.53","225.96","0.00","0.00","0.00","0.00","0.00","145.40"],"quality":"Less is better","isMain":true,"metricName":"# N's per 100 kbp"}]],["Statistics without reference",[{"values":[14,10,12,16,12,12,15,9,10,7,6,6,8,1,4],"quality":"Equal","isMain":true,"metricName":"# contigs"},{"values":[5,5,9,9,7,7,7,1,7,6,6,6,4,0,4],"quality":"Equal","isMain":false,"metricName":"# contigs (>= 1000 bp)"},{"values":[1,1,2,0,1,4,0,0,3,0,1,1,0,0,4],"quality":"Equal","isMain":false,"metricName":"# contigs (>= 5000 bp)"},{"values":[1,0,0,0,0,1,0,0,2,0,0,0,0,0,4],"quality":"Equal","isMain":false,"metricName":"# contigs (>= 10000 bp)"},{"values":[0,0,0,0,0,0,0,0,0,0,0,0,0,0,3],"quality":"Equal","isMain":false,"metricName":"# contigs (>= 25000 bp)"},{"values":[0,0,0,0,0,0,0,0,0,0,0,0,0,0,2],"quality":"Equal","isMain":false,"metricName":"# contigs (>= 50000 bp)"},{"values":[12022,5376,7719,4198,7708,12356,4814,1022,14780,4011,7319,7867,3656,513,136335],"quality":"More is better","isMain":true,"metricName":"Largest contig"},{"values":[33798,20479,31670,26279,27592,42955,21261,6146,44256,14488,23781,23494,11604,513,251717],"quality":"More is better","isMain":true,"metricName":"Total length"},{"values":[27674,17185,29517,21192,24343,39231,14844,1022,42162,13615,23781,23494,8878,0,251717],"quality":"More is better","isMain":true,"metricName":"Total length (>= 1000 bp)"},{"values":[12022,5376,12770,0,7708,35173,0,0,32867,0,7319,7867,0,0,251717],"quality":"More is better","isMain":false,"metricName":"Total length (>= 5000 bp)"},{"values":[12022,0,0,0,0,12356,0,0,25846,0,0,0,0,0,251717],"quality":"More is better","isMain":true,"metricName":"Total length (>= 10000 bp)"},{"values":[0,0,0,0,0,0,0,0,0,0,0,0,0,0,232089],"quality":"More is better","isMain":false,"metricName":"Total length (>= 25000 bp)"},{"values":[0,0,0,0,0,0,0,0,0,0,0,0,0,0,205781],"quality":"More is better","isMain":true,"metricName":"Total length (>= 50000 bp)"},{"values":[4939,3888,4582,2568,4368,8661,1554,700,11066,2361,4669,4669,1925,513,136335],"quality":"More is better","isMain":false,"metricName":"N50"},{"values":[2511,1623,2435,1156,2342,5238,901,574,3045,1936,4197,4197,1179,513,69446],"quality":"More is better","isMain":false,"metricName":"N75"},{"values":[2,3,3,4,3,3,4,4,2,3,2,2,3,1,1],"quality":"Less is better","isMain":false,"metricName":"L50"},{"values":[5,4,6,8,5,4,9,7,4,4,4,4,4,1,2],"quality":"Less is better","isMain":false,"metricName":"L75"},{"values":["34.86","34.35","34.73","34.50","34.43","34.88","34.02","33.50","34.70","33.68","34.38","34.29","32.63","45.81","34.94"],"quality":"Equal","isMain":false,"metricName":"GC (%)"}]],["Predicted genes",[]],["Similarity statistics",[{"values":[0,0,0,0,0,0,0,0,0,0,0,0,0,0,0],"quality":"Equal","isMain":false,"metricName":"# similar correct contigs"},{"values":[0,0,0,0,0,0,0,0,0,0,0,0,0,0,0],"quality":"Equal","isMain":false,"metricName":"# similar misassembled blocks"}]],["Reference statistics",[{"values":[40003,40003,40003,40003,40003,40003,40003,40003,40003,40003,40003,40003,40003,40003,40003],"quality":"Equal","isMain":false,"metricName":"Reference length"},{"values":[1,1,1,1,1,1,1,1,1,1,1,1,1,1,1],"quality":"Equal","isMain":false,"metricName":"Reference fragments"},{"values":["34.97","34.97","34.97","34.97","34.97","34.97","34.97","34.97","34.97","34.97","34.97","34.97","34.97","34.97","34.97"],"quality":"Equal","isMain":false,"metricName":"Reference GC (%)"}]]],"referenceName":"gi\_14251125\_ref\_NC","date":"18 July 2018, Wednesday, 16:19:28","order":[0,1,2,3,4,5,6,7,8,9,10,11,12,13,14],"assembliesNames":["ABySS\_127","ABySS\_63","CLC","IDBA\_UD","MEGAHIT","MIRA","Ray\_Meta","SOAPdenovo2","SPAdes","SPAdes\_meta","SPAdes\_sc","SPAdes\_sc\_careful","Velvet","VICUNA","Geneious"]},{"assembliesWithNs":null,"minContig":500,"report":[["Genome statistics",[{"values":["100.000","100.000","100.000","100.000","35.134","73.021","100.000","100.000","100.000","100.000","100.000"],"quality":"More is better","isMain":true,"metricName":"Genome fraction (%)"},{"values":["1.000","1.003","1.004","1.004","1.005","1.040","1.003","1.001","1.001","1.001","1.003"],"quality":"Less is better","isMain":true,"metricName":"Duplication ratio"},{"values":[38760,38859,38901,38931,1812,16189,38887,38815,38815,38815,14114],"quality":"More is better","isMain":true,"metricName":"Largest alignment"},{"values":[38760,38859,38901,38931,13643,28684,38887,38815,38815,38815,38874],"quality":"More is better","isMain":true,"metricName":"Total aligned length"},{"values":[38760,38859,38901,38931,null,10249,38887,38815,38815,38815,12135],"quality":"More is better","isMain":false,"metricName":"NG50"},{"values":[38760,38859,38901,38931,null,9699,38887,38815,38815,38815,9441],"quality":"More is better","isMain":false,"metricName":"NG75"},{"values":[38760,38859,38901,38931,917,6468,38887,38815,38815,38815,12135],"quality":"More is better","isMain":false,"metricName":"NA50"},{"values":[38760,38859,38901,38931,673,null,38887,38815,38815,38815,9441],"quality":"More is better","isMain":false,"metricName":"NA75"},{"values":[38760,38859,38901,38931,null,6468,38887,38815,38815,38815,12135],"quality":"More is better","isMain":true,"metricName":"NGA50"},{"values":[38760,38859,38901,38931,null,null,38887,38815,38815,38815,9441],"quality":"More is better","isMain":false,"metricName":"NGA75"},{"values":[1,1,1,1,null,2,1,1,1,1,2],"quality":"Less is better","isMain":false,"metricName":"LG50"},{"values":[1,1,1,1,null,3,1,1,1,1,3],"quality":"Less is better","isMain":false,"metricName":"LG75"},{"values":[1,1,1,1,6,2,1,1,1,1,2],"quality":"Less is better","isMain":false,"metricName":"LA50"},{"values":[1,1,1,1,11,null,1,1,1,1,3],"quality":"Less is better","isMain":false,"metricName":"LA75"},{"values":[1,1,1,1,null,2,1,1,1,1,2],"quality":"Less is better","isMain":true,"metricName":"LGA50"},{"values":[1,1,1,1,null,null,1,1,1,1,3],"quality":"Less is better","isMain":false,"metricName":"LGA75"}]],["Misassemblies",[{"values":[0,0,0,0,0,0,0,0,0,0,0],"quality":"Less is better","isMain":true,"metricName":"# misassemblies"},{"values":[0,0,0,0,0,0,0,0,0,0,0],"quality":"Less is better","isMain":false,"metricName":" # relocations"},{"values":[0,0,0,0,0,0,0,0,0,0,0],"quality":"Less is better","isMain":false,"metricName":" # translocations"},{"values":[0,0,0,0,0,0,0,0,0,0,0],"quality":"Less is better","isMain":false,"metricName":" # inversions"},{"values":[0,0,0,0,0,0,0,0,0,0,0],"quality":"Less is better","isMain":false,"metricName":"# misassembled contigs"},{"values":[0,0,0,0,0,0,0,0,0,0,0],"quality":"Less is better","isMain":true,"metricName":"Misassembled contigs length"},{"values":[0,1,1,1,0,0,1,0,0,0,0],"quality":"Less is better","isMain":false,"metricName":"# local misassemblies"},{"values":[0,0,0,0,0,2,0,0,0,0,0],"quality":"Less is better","isMain":false,"metricName":"# unaligned mis. contigs"}]],["Unaligned",[{"values":[0,0,0,0,0,0,0,0,0,0,0],"quality":"Less is better","isMain":false,"metricName":"# fully unaligned contigs"},{"values":[0,0,0,0,0,0,0,0,0,0,0],"quality":"Less is better","isMain":false,"metricName":"Fully unaligned length"},{"values":[0,0,0,0,0,3,0,0,0,0,0],"quality":"Less is better","isMain":false,"metricName":"# partially unaligned contigs"},{"values":[0,0,0,0,0,10799,0,0,0,0,0],"quality":"Less is better","isMain":false,"metricName":"Partially unaligned length"}]],["Mismatches",[{"values":[0,0,0,0,0,0,0,0,0,0,0],"quality":"Less is better","isMain":false,"metricName":"# mismatches"},{"values":[0,0,0,0,0,15,0,1,1,1,0],"quality":"Less is better","isMain":false,"metricName":"# indels"},{"values":[0,0,0,0,0,665,0,55,55,55,0],"quality":"Less is better","isMain":false,"metricName":"Indels length"},{"values":["0.00","0.00","0.00","0.00","0.00","0.00","0.00","0.00","0.00","0.00","0.00"],"quality":"Less is better","isMain":true,"metricName":"# mismatches per 100 kbp"},{"values":["0.00","0.00","0.00","0.00","0.00","53.00","0.00","2.58","2.58","2.58","0.00"],"quality":"Less is better","isMain":true,"metricName":"# indels per 100 kbp"},{"values":[0,0,0,0,0,0,0,0,0,0,0],"quality":"Less is better","isMain":false,"metricName":" # indels (<= 5 bp)"},{"values":[0,0,0,0,0,15,0,1,1,1,0],"quality":"Less is better","isMain":false,"metricName":" # indels (> 5 bp)"},{"values":[0,0,0,0,0,3243,0,0,0,0,0],"quality":"Less is better","isMain":false,"metricName":"# N's"},{"values":["0.00","0.00","0.00","0.00","0.00","8063.15","0.00","0.00","0.00","0.00","0.00"],"quality":"Less is better","isMain":true,"metricName":"# N's per 100 kbp"}]],["Statistics without reference",[{"values":[1,1,1,1,16,4,1,1,1,1,5],"quality":"Equal","isMain":true,"metricName":"# contigs"},{"values":[1,1,1,1,4,4,1,1,1,1,4],"quality":"Equal","isMain":false,"metricName":"# contigs (>= 1000 bp)"},{"values":[1,1,1,1,0,3,1,1,1,1,3],"quality":"Equal","isMain":false,"metricName":"# contigs (>= 5000 bp)"},{"values":[1,1,1,1,0,2,1,1,1,1,2],"quality":"Equal","isMain":false,"metricName":"# contigs (>= 10000 bp)"},{"values":[1,1,1,1,0,0,1,1,1,1,0],"quality":"Equal","isMain":false,"metricName":"# contigs (>= 25000 bp)"},{"values":[0,0,0,0,0,0,0,0,0,0,0],"quality":"Equal","isMain":false,"metricName":"# contigs (>= 50000 bp)"},{"values":[38760,38859,38901,38931,1812,16926,38887,38815,38815,38815,14114],"quality":"More is better","isMain":true,"metricName":"Largest contig"},{"values":[38760,38859,38901,38931,13688,40220,38887,38815,38815,38815,38874],"quality":"More is better","isMain":true,"metricName":"Total length"},{"values":[38760,38859,38901,38931,4990,40220,38887,38815,38815,38815,38141],"quality":"More is better","isMain":true,"metricName":"Total length (>= 1000 bp)"},{"values":[38760,38859,38901,38931,0,36874,38887,38815,38815,38815,35690],"quality":"More is better","isMain":false,"metricName":"Total length (>= 5000 bp)"},{"values":[38760,38859,38901,38931,0,27175,38887,38815,38815,38815,26249],"quality":"More is better","isMain":true,"metricName":"Total length (>= 10000 bp)"},{"values":[38760,38859,38901,38931,0,0,38887,38815,38815,38815,0],"quality":"More is better","isMain":false,"metricName":"Total length (>= 25000 bp)"},{"values":[0,0,0,0,0,0,0,0,0,0,0],"quality":"More is better","isMain":true,"metricName":"Total length (>= 50000 bp)"},{"values":[38760,38859,38901,38931,917,10249,38887,38815,38815,38815,12135],"quality":"More is better","isMain":false,"metricName":"N50"},{"values":[38760,38859,38901,38931,673,9699,38887,38815,38815,38815,9441],"quality":"More is better","isMain":false,"metricName":"N75"},{"values":[1,1,1,1,6,2,1,1,1,1,2],"quality":"Less is better","isMain":false,"metricName":"L50"},{"values":[1,1,1,1,11,3,1,1,1,1,3],"quality":"Less is better","isMain":false,"metricName":"L75"},{"values":["65.35","65.34","65.37","65.34","64.90","65.29","65.36","65.36","65.36","65.36","65.34"],"quality":"Equal","isMain":false,"metricName":"GC (%)"}]],["Predicted genes",[]],["Similarity statistics",[{"values":[0,0,0,0,0,0,0,0,0,0,0],"quality":"Equal","isMain":false,"metricName":"# similar correct contigs"},{"values":[0,0,0,0,0,0,0,0,0,0,0],"quality":"Equal","isMain":false,"metricName":"# similar misassembled blocks"}]],["Reference statistics",[{"values":[38760,38760,38760,38760,38760,38760,38760,38760,38760,38760,38760],"quality":"Equal","isMain":false,"metricName":"Reference length"},{"values":[1,1,1,1,1,1,1,1,1,1,1],"quality":"Equal","isMain":false,"metricName":"Reference fragments"},{"values":["65.35","65.35","65.35","65.35","65.35","65.35","65.35","65.35","65.35","65.35","65.35"],"quality":"Equal","isMain":false,"metricName":"Reference GC (%)"}]]],"referenceName":"gi\_145708074\_ref\_NC","date":"18 July 2018, Wednesday, 16:19:38","order":[0,1,2,3,4,5,6,7,8,9,10],"assembliesNames":["CLC","IDBA\_UD","MEGAHIT","MIRA","Ray\_Meta","SOAPdenovo2","SPAdes","SPAdes\_meta","SPAdes\_sc","SPAdes\_sc\_careful","Velvet"]},{"assembliesWithNs":null,"minContig":500,"report":[["Genome statistics",[{"values":["72.193","96.696","100.000","100.000","100.000","100.000","100.000","100.000","100.000","100.000","99.998"],"quality":"More is better","isMain":true,"metricName":"Genome fraction (%)"},{"values":["1.026","1.047","1.000","1.002","1.003","1.019","1.002","1.001","1.001","1.001","1.000"],"quality":"Less is better","isMain":true,"metricName":"Duplication ratio"},{"values":[7987,10725,52414,52513,52555,53394,52541,52469,52469,52469,52413],"quality":"More is better","isMain":true,"metricName":"Largest alignment"},{"values":[38840,52823,52414,52513,52555,53394,52541,52469,52469,52469,52413],"quality":"More is better","isMain":true,"metricName":"Total aligned length"},{"values":[926,10406,52414,52513,52555,53394,52541,52469,52469,52469,52413],"quality":"More is better","isMain":false,"metricName":"NG50"},{"values":[null,8814,52414,52513,52555,53394,52541,52469,52469,52469,52413],"quality":"More is better","isMain":false,"metricName":"NG75"},{"values":[2346,10248,52414,52513,52555,53394,52541,52469,52469,52469,52413],"quality":"More is better","isMain":false,"metricName":"NA50"},{"values":[853,8814,52414,52513,52555,53394,52541,52469,52469,52469,52413],"quality":"More is better","isMain":false,"metricName":"NA75"},{"values":[926,10248,52414,52513,52555,53394,52541,52469,52469,52469,52413],"quality":"More is better","isMain":true,"metricName":"NGA50"},{"values":[null,8814,52414,52513,52555,53394,52541,52469,52469,52469,52413],"quality":"More is better","isMain":false,"metricName":"NGA75"},{"values":[8,3,1,1,1,1,1,1,1,1,1],"quality":"Less is better","isMain":false,"metricName":"LG50"},{"values":[null,4,1,1,1,1,1,1,1,1,1],"quality":"Less is better","isMain":false,"metricName":"LG75"},{"values":[4,3,1,1,1,1,1,1,1,1,1],"quality":"Less is better","isMain":false,"metricName":"LA50"},{"values":[11,4,1,1,1,1,1,1,1,1,1],"quality":"Less is better","isMain":false,"metricName":"LA75"},{"values":[8,3,1,1,1,1,1,1,1,1,1],"quality":"Less is better","isMain":true,"metricName":"LGA50"},{"values":[null,4,1,1,1,1,1,1,1,1,1],"quality":"Less is better","isMain":false,"metricName":"LGA75"}]],["Misassemblies",[{"values":[0,0,0,0,0,0,0,0,0,0,0],"quality":"Less is better","isMain":true,"metricName":"# misassemblies"},{"values":[0,0,0,0,0,0,0,0,0,0,0],"quality":"Less is better","isMain":false,"metricName":" # relocations"},{"values":[0,0,0,0,0,0,0,0,0,0,0],"quality":"Less is better","isMain":false,"metricName":" # translocations"},{"values":[0,0,0,0,0,0,0,0,0,0,0],"quality":"Less is better","isMain":false,"metricName":" # inversions"},{"values":[0,0,0,0,0,0,0,0,0,0,0],"quality":"Less is better","isMain":false,"metricName":"# misassembled contigs"},{"values":[0,0,0,0,0,0,0,0,0,0,0],"quality":"Less is better","isMain":true,"metricName":"Misassembled contigs length"},{"values":[0,0,0,1,1,1,1,0,0,0,0],"quality":"Less is better","isMain":false,"metricName":"# local misassemblies"},{"values":[0,0,0,0,0,0,0,0,0,0,0],"quality":"Less is better","isMain":false,"metricName":"# unaligned mis. contigs"}]],["Unaligned",[{"values":[0,0,0,0,0,0,0,0,0,0,0],"quality":"Less is better","isMain":false,"metricName":"# fully unaligned contigs"},{"values":[0,0,0,0,0,0,0,0,0,0,0],"quality":"Less is better","isMain":false,"metricName":"Fully unaligned length"},{"values":[0,0,0,0,0,0,0,0,0,0,0],"quality":"Less is better","isMain":false,"metricName":"# partially unaligned contigs"},{"values":[0,0,0,0,0,0,0,0,0,0,0],"quality":"Less is better","isMain":false,"metricName":"Partially unaligned length"}]],["Mismatches",[{"values":[0,0,0,0,0,2,0,0,0,0,0],"quality":"Less is better","isMain":false,"metricName":"# mismatches"},{"values":[0,0,0,0,0,0,0,1,1,1,0],"quality":"Less is better","isMain":false,"metricName":"# indels"},{"values":[0,0,0,0,0,0,0,55,55,55,0],"quality":"Less is better","isMain":false,"metricName":"Indels length"},{"values":["0.00","0.00","0.00","0.00","0.00","3.82","0.00","0.00","0.00","0.00","0.00"],"quality":"Less is better","isMain":true,"metricName":"# mismatches per 100 kbp"},{"values":["0.00","0.00","0.00","0.00","0.00","0.00","0.00","1.91","1.91","1.91","0.00"],"quality":"Less is better","isMain":true,"metricName":"# indels per 100 kbp"},{"values":[0,0,0,0,0,0,0,0,0,0,0],"quality":"Less is better","isMain":false,"metricName":" # indels (<= 5 bp)"},{"values":[0,0,0,0,0,0,0,1,1,1,0],"quality":"Less is better","isMain":false,"metricName":" # indels (> 5 bp)"},{"values":[1,835,0,0,0,1,0,0,0,0,0],"quality":"Less is better","isMain":false,"metricName":"# N's"},{"values":["2.57","1573.01","0.00","0.00","0.00","1.87","0.00","0.00","0.00","0.00","0.00"],"quality":"Less is better","isMain":true,"metricName":"# N's per 100 kbp"}]],["Statistics without reference",[{"values":[26,11,1,1,1,1,1,1,1,1,1],"quality":"Equal","isMain":true,"metricName":"# contigs"},{"values":[7,7,1,1,1,1,1,1,1,1,1],"quality":"Equal","isMain":false,"metricName":"# contigs (>= 1000 bp)"},{"values":[2,4,1,1,1,1,1,1,1,1,1],"quality":"Equal","isMain":false,"metricName":"# contigs (>= 5000 bp)"},{"values":[0,3,1,1,1,1,1,1,1,1,1],"quality":"Equal","isMain":false,"metricName":"# contigs (>= 10000 bp)"},{"values":[0,0,1,1,1,1,1,1,1,1,1],"quality":"Equal","isMain":false,"metricName":"# contigs (>= 25000 bp)"},{"values":[0,0,1,1,1,1,1,1,1,1,1],"quality":"Equal","isMain":false,"metricName":"# contigs (>= 50000 bp)"},{"values":[7987,10725,52414,52513,52555,53394,52541,52469,52469,52469,52413],"quality":"More is better","isMain":true,"metricName":"Largest contig"},{"values":[38840,53083,52414,52513,52555,53394,52541,52469,52469,52469,52413],"quality":"More is better","isMain":true,"metricName":"Total length"},{"values":[25622,50657,52414,52513,52555,53394,52541,52469,52469,52469,52413],"quality":"More is better","isMain":true,"metricName":"Total length (>= 1000 bp)"},{"values":[14967,40374,52414,52513,52555,53394,52541,52469,52469,52469,52413],"quality":"More is better","isMain":false,"metricName":"Total length (>= 5000 bp)"},{"values":[0,31560,52414,52513,52555,53394,52541,52469,52469,52469,52413],"quality":"More is better","isMain":true,"metricName":"Total length (>= 10000 bp)"},{"values":[0,0,52414,52513,52555,53394,52541,52469,52469,52469,52413],"quality":"More is better","isMain":false,"metricName":"Total length (>= 25000 bp)"},{"values":[0,0,52414,52513,52555,53394,52541,52469,52469,52469,52413],"quality":"More is better","isMain":true,"metricName":"Total length (>= 50000 bp)"},{"values":[2346,10406,52414,52513,52555,53394,52541,52469,52469,52469,52413],"quality":"More is better","isMain":false,"metricName":"N50"},{"values":[853,8814,52414,52513,52555,53394,52541,52469,52469,52469,52413],"quality":"More is better","isMain":false,"metricName":"N75"},{"values":[4,3,1,1,1,1,1,1,1,1,1],"quality":"Less is better","isMain":false,"metricName":"L50"},{"values":[11,4,1,1,1,1,1,1,1,1,1],"quality":"Less is better","isMain":false,"metricName":"L75"},{"values":["56.17","56.31","56.29","56.28","56.28","56.25","56.30","56.29","56.29","56.29","56.29"],"quality":"Equal","isMain":false,"metricName":"GC (%)"}]],["Predicted genes",[]],["Similarity statistics",[{"values":[0,0,0,0,0,0,0,0,0,0,0],"quality":"Equal","isMain":false,"metricName":"# similar correct contigs"},{"values":[0,0,0,0,0,0,0,0,0,0,0],"quality":"Equal","isMain":false,"metricName":"# similar misassembled blocks"}]],["Reference statistics",[{"values":[52414,52414,52414,52414,52414,52414,52414,52414,52414,52414,52414],"quality":"Equal","isMain":false,"metricName":"Reference length"},{"values":[1,1,1,1,1,1,1,1,1,1,1],"quality":"Equal","isMain":false,"metricName":"Reference fragments"},{"values":["56.29","56.29","56.29","56.29","56.29","56.29","56.29","56.29","56.29","56.29","56.29"],"quality":"Equal","isMain":false,"metricName":"Reference GC (%)"}]]],"referenceName":"gi\_146329912\_ref\_NC","date":"18 July 2018, Wednesday, 16:19:51","order":[0,1,2,3,4,5,6,7,8,9,10],"assembliesNames":["ABySS\_127","ABySS\_63","CLC","IDBA\_UD","MEGAHIT","MIRA","SPAdes","SPAdes\_meta","SPAdes\_sc","SPAdes\_sc\_careful","Velvet"]},{"assembliesWithNs":null,"minContig":500,"report":[["Genome statistics",[{"values":["74.679","65.842","95.066","95.096","95.575","89.047","90.564","96.304","97.874","96.540","95.891","88.201","30.447"],"quality":"More is better","isMain":true,"metricName":"Genome fraction (%)"},{"values":["1.002","1.001","1.009","1.002","1.006","1.015","1.016","1.002","1.003","1.002","1.001","1.001","1.010"],"quality":"Less is better","isMain":true,"metricName":"Duplication ratio"},{"values":[3960,3270,18695,18870,18666,12890,15035,20298,19185,18647,18647,16211,8132],"quality":"More is better","isMain":true,"metricName":"Largest alignment"},{"values":[92788,81755,119049,118311,119293,112191,113014,119717,121773,120011,119184,109558,37795],"quality":"More is better","isMain":true,"metricName":"Total aligned length"},{"values":[1080,872,10010,7327,9862,4070,3905,13476,9760,10010,10010,8156,null],"quality":"More is better","isMain":false,"metricName":"NG50"},{"values":[null,null,4615,2950,3962,1451,2017,4895,4615,4024,4024,2644,null],"quality":"More is better","isMain":false,"metricName":"NG75"},{"values":[1354,1393,10943,7327,9862,4111,4020,13476,9760,10010,10010,8862,4746],"quality":"More is better","isMain":false,"metricName":"NA50"},{"values":[980,868,4895,4024,4430,1866,2361,7060,4615,4615,4615,4895,4111],"quality":"More is better","isMain":false,"metricName":"NA75"},{"values":[1080,868,10010,7327,9862,4070,3905,13476,9760,10010,10010,8156,null],"quality":"More is better","isMain":true,"metricName":"NGA50"},{"values":[null,null,4615,2950,3962,1444,2017,4895,4615,4024,4024,2644,null],"quality":"More is better","isMain":false,"metricName":"NGA75"},{"values":[36,39,5,5,5,10,10,4,5,5,5,6,null],"quality":"Less is better","isMain":false,"metricName":"LG50"},{"values":[null,null,9,11,10,24,21,8,9,10,10,12,null],"quality":"Less is better","isMain":false,"metricName":"LG75"},{"values":[23,20,4,5,5,9,9,4,5,5,5,5,4],"quality":"Less is better","isMain":false,"metricName":"LA50"},{"values":[43,39,8,10,9,19,18,7,9,9,9,9,6],"quality":"Less is better","isMain":false,"metricName":"LA75"},{"values":[36,39,5,5,5,10,10,4,5,5,5,6,null],"quality":"Less is better","isMain":true,"metricName":"LGA50"},{"values":[null,null,9,11,10,25,21,8,9,10,10,12,null],"quality":"Less is better","isMain":false,"metricName":"LGA75"}]],["Misassemblies",[{"values":[0,0,0,0,0,2,0,0,0,1,1,0,1],"quality":"Less is better","isMain":true,"metricName":"# misassemblies"},{"values":[0,0,0,0,0,2,0,0,0,1,1,0,1],"quality":"Less is better","isMain":false,"metricName":" # relocations"},{"values":[0,0,0,0,0,0,0,0,0,0,0,0,0],"quality":"Less is better","isMain":false,"metricName":" # translocations"},{"values":[0,0,0,0,0,0,0,0,0,0,0,0,0],"quality":"Less is better","isMain":false,"metricName":" # inversions"},{"values":[0,0,0,0,0,2,0,0,0,1,1,0,1],"quality":"Less is better","isMain":false,"metricName":"# misassembled contigs"},{"values":[0,0,0,0,0,10540,0,0,0,3145,3145,0,7381],"quality":"Less is better","isMain":true,"metricName":"Misassembled contigs length"},{"values":[0,0,0,0,1,0,0,0,1,0,0,0,1],"quality":"Less is better","isMain":false,"metricName":"# local misassemblies"},{"values":[0,0,0,0,0,0,0,0,0,0,0,0,0],"quality":"Less is better","isMain":false,"metricName":"# unaligned mis. contigs"}]],["Unaligned",[{"values":[0,0,0,0,0,0,0,0,0,0,0,0,0],"quality":"Less is better","isMain":false,"metricName":"# fully unaligned contigs"},{"values":[0,0,0,0,0,0,0,0,0,0,0,0,0],"quality":"Less is better","isMain":false,"metricName":"Fully unaligned length"},{"values":[0,0,0,0,0,0,0,0,0,0,0,0,0],"quality":"Less is better","isMain":false,"metricName":"# partially unaligned contigs"},{"values":[0,0,0,0,0,0,0,0,0,0,0,0,0],"quality":"Less is better","isMain":false,"metricName":"Partially unaligned length"}]],["Mismatches",[{"values":[11,7,28,21,51,8,24,56,45,39,30,28,8],"quality":"Less is better","isMain":false,"metricName":"# mismatches"},{"values":[0,0,0,0,6,0,16,0,0,1,0,0,4],"quality":"Less is better","isMain":false,"metricName":"# indels"},{"values":[0,0,0,0,34,0,791,0,0,21,0,0,4],"quality":"Less is better","isMain":false,"metricName":"Indels length"},{"values":["11.87","8.57","23.73","17.79","42.99","7.24","21.35","46.85","37.04","32.55","25.21","25.58","21.17"],"quality":"Less is better","isMain":true,"metricName":"# mismatches per 100 kbp"},{"values":["0.00","0.00","0.00","0.00","5.06","0.00","14.23","0.00","0.00","0.83","0.00","0.00","10.58"],"quality":"Less is better","isMain":true,"metricName":"# indels per 100 kbp"},{"values":[0,0,0,0,4,0,0,0,0,0,0,0,4],"quality":"Less is better","isMain":false,"metricName":" # indels (<= 5 bp)"},{"values":[0,0,0,0,2,0,16,0,0,1,0,0,0],"quality":"Less is better","isMain":false,"metricName":" # indels (> 5 bp)"},{"values":[0,0,0,0,0,78,1651,0,100,0,0,0,213],"quality":"Less is better","isMain":false,"metricName":"# N's"},{"values":["0.00","0.00","0.00","0.00","0.00","69.52","1445.03","0.00","82.05","0.00","0.00","0.00","557.87"],"quality":"Less is better","isMain":true,"metricName":"# N's per 100 kbp"}]],["Statistics without reference",[{"values":[76,68,25,24,23,43,38,17,19,22,21,21,7],"quality":"Equal","isMain":true,"metricName":"# contigs"},{"values":[42,32,20,21,21,31,30,16,19,20,20,19,7],"quality":"Equal","isMain":false,"metricName":"# contigs (>= 1000 bp)"},{"values":[0,0,7,7,7,6,5,7,7,6,6,8,3],"quality":"Equal","isMain":false,"metricName":"# contigs (>= 5000 bp)"},{"values":[0,0,5,4,4,1,1,5,4,5,5,4,0],"quality":"Equal","isMain":false,"metricName":"# contigs (>= 10000 bp)"},{"values":[0,0,0,0,0,0,0,0,0,0,0,0,0],"quality":"Equal","isMain":false,"metricName":"# contigs (>= 25000 bp)"},{"values":[0,0,0,0,0,0,0,0,0,0,0,0,0],"quality":"Equal","isMain":false,"metricName":"# contigs (>= 50000 bp)"},{"values":[3960,3270,18695,18870,18666,12890,15035,20298,19185,18647,18647,16211,8132],"quality":"More is better","isMain":true,"metricName":"Largest contig"},{"values":[92846,81782,119049,118316,119293,112191,114254,119717,121873,120013,119186,109558,38181],"quality":"More is better","isMain":true,"metricName":"Total length"},{"values":[68918,55855,115718,116307,118175,102617,107539,119177,121873,118253,118232,107773,38181],"quality":"More is better","isMain":true,"metricName":"Total length (>= 1000 bp)"},{"values":[0,0,85240,78186,81546,46511,43639,90738,85944,76526,76526,78594,20886],"quality":"More is better","isMain":false,"metricName":"Total length (>= 5000 bp)"},{"values":[0,0,69798,57015,57563,12890,15035,73918,60482,69466,69466,48737,0],"quality":"More is better","isMain":true,"metricName":"Total length (>= 10000 bp)"},{"values":[0,0,0,0,0,0,0,0,0,0,0,0,0],"quality":"More is better","isMain":false,"metricName":"Total length (>= 25000 bp)"},{"values":[0,0,0,0,0,0,0,0,0,0,0,0,0],"quality":"More is better","isMain":true,"metricName":"Total length (>= 50000 bp)"},{"values":[1354,1393,10943,7327,9862,4111,4020,13476,9760,10010,10010,8862,5373],"quality":"More is better","isMain":false,"metricName":"N50"},{"values":[980,872,4895,4024,4430,1866,2436,7060,4615,4615,4615,4895,4498],"quality":"More is better","isMain":false,"metricName":"N75"},{"values":[23,20,4,5,5,9,9,4,5,5,5,5,3],"quality":"Less is better","isMain":false,"metricName":"L50"},{"values":[43,39,8,10,9,19,17,7,9,9,9,9,5],"quality":"Less is better","isMain":false,"metricName":"L75"},{"values":["38.37","38.36","38.90","38.84","38.82","38.65","38.87","38.87","38.84","38.90","38.93","38.89","38.53"],"quality":"Equal","isMain":false,"metricName":"GC (%)"}]],["Predicted genes",[]],["Similarity statistics",[{"values":[0,0,2,2,2,0,0,1,2,2,2,2,0],"quality":"Equal","isMain":false,"metricName":"# similar correct contigs"},{"values":[0,0,0,0,0,0,0,0,0,0,0,0,0],"quality":"Equal","isMain":false,"metricName":"# similar misassembled blocks"}]],["Reference statistics",[{"values":[124121,124121,124121,124121,124121,124121,124121,124121,124121,124121,124121,124121,124121],"quality":"Equal","isMain":false,"metricName":"Reference length"},{"values":[1,1,1,1,1,1,1,1,1,1,1,1,1],"quality":"Equal","isMain":false,"metricName":"Reference fragments"},{"values":["38.83","38.83","38.83","38.83","38.83","38.83","38.83","38.83","38.83","38.83","38.83","38.83","38.83"],"quality":"Equal","isMain":false,"metricName":"Reference GC (%)"}]]],"referenceName":"gi\_148368821\_ref\_NC","date":"18 July 2018, Wednesday, 16:20:02","order":[0,1,2,3,4,5,6,7,8,9,10,11,12],"assembliesNames":["ABySS\_127","ABySS\_63","CLC","IDBA\_UD","MEGAHIT","MIRA","SOAPdenovo2","SPAdes","SPAdes\_meta","SPAdes\_sc","SPAdes\_sc\_careful","Velvet","Geneious"]},{"assembliesWithNs":null,"minContig":500,"report":[["Genome statistics",[{"values":["99.124","89.388","98.101","94.273","98.528","100.000","49.036","50.665","100.000","94.656","89.715","92.429","73.050","100.000"],"quality":"More is better","isMain":true,"metricName":"Genome fraction (%)"},{"values":["1.013","1.010","1.003","1.006","1.011","1.078","1.025","1.079","1.047","1.008","1.009","1.006","1.001","1.052"],"quality":"Less is better","isMain":true,"metricName":"Duplication ratio"},{"values":[21880,14423,43611,16713,19564,24701,1830,7171,27332,19912,16474,18518,6910,24701],"quality":"More is better","isMain":true,"metricName":"Largest alignment"},{"values":[47215,42446,46128,44614,46817,50676,23627,24010,49241,44618,42549,43735,34395,49444],"quality":"More is better","isMain":true,"metricName":"Total aligned length"},{"values":[11072,5632,43744,7355,11123,153388,505,2046,57853,16600,24723,24723,4597,179797],"quality":"More is better","isMain":false,"metricName":"NG50"},{"values":[5929,3822,43744,3430,5959,153388,null,null,57853,16600,16474,18518,null,179797],"quality":"More is better","isMain":false,"metricName":"NG75"},{"values":[11072,7503,43611,7355,11123,null,983,1642,21053,16600,13870,13870,5464,null],"quality":"More is better","isMain":false,"metricName":"NA50"},{"values":[5929,3822,43611,3760,5959,null,765,null,21053,16600,1062,1174,3046,null],"quality":"More is better","isMain":false,"metricName":"NA75"},{"values":[11072,5632,43611,7355,11123,24701,505,314,27332,16600,13870,13870,4597,24701],"quality":"More is better","isMain":true,"metricName":"NGA50"},{"values":[5929,3822,43611,3430,5959,14568,null,null,21053,16600,1441,2079,null,14635],"quality":"More is better","isMain":false,"metricName":"NGA75"},{"values":[2,3,1,2,2,1,25,5,1,2,1,1,4,1],"quality":"Less is better","isMain":false,"metricName":"LG50"},{"values":[3,5,1,5,3,1,null,null,1,2,2,2,null,1],"quality":"Less is better","isMain":false,"metricName":"LG75"},{"values":[2,2,1,2,2,null,9,5,2,2,2,2,3,null],"quality":"Less is better","isMain":false,"metricName":"LA50"},{"values":[3,5,1,4,3,null,16,null,2,2,8,7,5,null],"quality":"Less is better","isMain":false,"metricName":"LA75"},{"values":[2,3,1,2,2,1,25,12,1,2,2,2,4,1],"quality":"Less is better","isMain":true,"metricName":"LGA50"},{"values":[3,5,1,5,3,2,null,null,2,2,5,4,null,2],"quality":"Less is better","isMain":false,"metricName":"LGA75"}]],["Misassemblies",[{"values":[0,0,0,0,0,0,0,0,1,0,0,0,0,0],"quality":"Less is better","isMain":true,"metricName":"# misassemblies"},{"values":[0,0,0,0,0,0,0,0,1,0,0,0,0,0],"quality":"Less is better","isMain":false,"metricName":" # relocations"},{"values":[0,0,0,0,0,0,0,0,0,0,0,0,0,0],"quality":"Less is better","isMain":false,"metricName":" # translocations"},{"values":[0,0,0,0,0,0,0,0,0,0,0,0,0,0],"quality":"Less is better","isMain":false,"metricName":" # inversions"},{"values":[0,0,0,0,0,0,0,0,1,0,0,0,0,0],"quality":"Less is better","isMain":false,"metricName":"# misassembled contigs"},{"values":[0,0,0,0,0,0,0,0,57853,0,0,0,0,0],"quality":"Less is better","isMain":true,"metricName":"Misassembled contigs length"},{"values":[0,0,0,0,0,0,0,3,0,1,0,0,0,0],"quality":"Less is better","isMain":false,"metricName":"# local misassemblies"},{"values":[0,0,0,0,0,1,0,1,0,0,0,0,0,1],"quality":"Less is better","isMain":false,"metricName":"# unaligned mis. contigs"}]],["Unaligned",[{"values":[0,0,0,0,0,0,0,0,0,0,0,0,0,0],"quality":"Less is better","isMain":false,"metricName":"# fully unaligned contigs"},{"values":[0,0,0,0,0,0,0,0,0,0,0,0,0,0],"quality":"Less is better","isMain":false,"metricName":"Fully unaligned length"},{"values":[0,0,0,0,0,1,0,4,1,0,1,1,0,1],"quality":"Less is better","isMain":false,"metricName":"# partially unaligned contigs"},{"values":[0,0,0,0,0,104011,0,7890,9468,0,10853,10853,0,130353],"quality":"Less is better","isMain":false,"metricName":"Partially unaligned length"}]],["Mismatches",[{"values":[0,8,9,1,25,10,13,0,120,146,58,22,0,12],"quality":"Less is better","isMain":false,"metricName":"# mismatches"},{"values":[0,2,0,0,0,0,0,18,3,6,3,1,0,0],"quality":"Less is better","isMain":false,"metricName":"# indels"},{"values":[0,2,0,0,0,0,0,886,3,6,3,1,0,0],"quality":"Less is better","isMain":false,"metricName":"Indels length"},{"values":["0.00","19.03","19.51","2.26","53.96","21.27","56.38","0.00","255.21","328.03","137.49","50.62","0.00","25.52"],"quality":"Less is better","isMain":true,"metricName":"# mismatches per 100 kbp"},{"values":["0.00","4.76","0.00","0.00","0.00","0.00","0.00","75.56","6.38","13.48","7.11","2.30","0.00","0.00"],"quality":"Less is better","isMain":true,"metricName":"# indels per 100 kbp"},{"values":[0,2,0,0,0,0,0,0,3,6,3,1,0,0],"quality":"Less is better","isMain":false,"metricName":" # indels (<= 5 bp)"},{"values":[0,0,0,0,0,0,0,18,0,0,0,0,0,0],"quality":"Less is better","isMain":false,"metricName":" # indels (> 5 bp)"},{"values":[0,0,0,0,0,0,0,4941,0,0,0,0,0,192],"quality":"Less is better","isMain":false,"metricName":"# N's"},{"values":["0.00","0.00","0.00","0.00","0.00","0.00","0.00","14706.67","0.00","0.00","0.00","0.00","0.00","106.79"],"quality":"Less is better","isMain":true,"metricName":"# N's per 100 kbp"}]],["Statistics without reference",[{"values":[8,11,2,11,7,3,25,14,2,4,11,10,11,1],"quality":"Equal","isMain":true,"metricName":"# contigs"},{"values":[6,9,2,8,7,1,8,8,1,3,9,9,8,1],"quality":"Equal","isMain":false,"metricName":"# contigs (>= 1000 bp)"},{"values":[3,3,1,3,3,1,0,2,1,3,2,2,3,1],"quality":"Equal","isMain":false,"metricName":"# contigs (>= 5000 bp)"},{"values":[2,1,1,1,2,1,0,1,1,2,2,2,0,1],"quality":"Equal","isMain":false,"metricName":"# contigs (>= 10000 bp)"},{"values":[0,0,1,0,0,1,0,0,1,0,0,0,0,1],"quality":"Equal","isMain":false,"metricName":"# contigs (>= 25000 bp)"},{"values":[0,0,0,0,0,1,0,0,1,0,0,0,0,1],"quality":"Equal","isMain":false,"metricName":"# contigs (>= 50000 bp)"},{"values":[21880,14423,43744,16713,19564,153388,1830,10070,57853,20141,24723,24723,6910,179797],"quality":"More is better","isMain":true,"metricName":"Largest contig"},{"values":[47215,42446,46261,44614,46817,154687,23627,33597,58709,44847,53402,54588,34395,179797],"quality":"More is better","isMain":true,"metricName":"Total length"},{"values":[45746,41184,46261,42601,46817,153388,11117,29542,57853,44224,52147,54054,32630,179797],"quality":"More is better","isMain":true,"metricName":"Total length (>= 1000 bp)"},{"values":[38881,27558,43744,29774,36646,153388,0,16072,57853,44224,41197,43241,18982,179797],"quality":"More is better","isMain":false,"metricName":"Total length (>= 5000 bp)"},{"values":[32952,14423,43744,16713,30687,153388,0,10070,57853,36741,41197,43241,0,179797],"quality":"More is better","isMain":true,"metricName":"Total length (>= 10000 bp)"},{"values":[0,0,43744,0,0,153388,0,0,57853,0,0,0,0,179797],"quality":"More is better","isMain":false,"metricName":"Total length (>= 25000 bp)"},{"values":[0,0,0,0,0,153388,0,0,57853,0,0,0,0,179797],"quality":"More is better","isMain":true,"metricName":"Total length (>= 50000 bp)"},{"values":[11072,7503,43744,7355,11123,153388,983,3570,57853,16600,16474,18518,5464,179797],"quality":"More is better","isMain":false,"metricName":"N50"},{"values":[5929,3822,43744,3760,5959,153388,765,1770,57853,16600,16474,18518,3046,179797],"quality":"More is better","isMain":false,"metricName":"N75"},{"values":[2,2,1,2,2,1,9,3,1,2,2,2,3,1],"quality":"Less is better","isMain":false,"metricName":"L50"},{"values":[3,5,1,4,3,1,16,6,1,2,2,2,5,1],"quality":"Less is better","isMain":false,"metricName":"L75"},{"values":["49.09","48.37","49.12","48.63","48.99","50.97","47.40","47.33","48.69","48.80","49.27","49.66","47.11","50.51"],"quality":"Equal","isMain":false,"metricName":"GC (%)"}]],["Predicted genes",[]],["Similarity statistics",[{"values":[0,0,0,0,0,0,0,0,0,0,0,0,0,0],"quality":"Equal","isMain":false,"metricName":"# similar correct contigs"},{"values":[0,0,0,0,0,0,0,0,0,0,0,0,0,0],"quality":"Equal","isMain":false,"metricName":"# similar misassembled blocks"}]],["Reference statistics",[{"values":[47021,47021,47021,47021,47021,47021,47021,47021,47021,47021,47021,47021,47021,47021],"quality":"Equal","isMain":false,"metricName":"Reference length"},{"values":[1,1,1,1,1,1,1,1,1,1,1,1,1,1],"quality":"Equal","isMain":false,"metricName":"Reference fragments"},{"values":["49.12","49.12","49.12","49.12","49.12","49.12","49.12","49.12","49.12","49.12","49.12","49.12","49.12","49.12"],"quality":"Equal","isMain":false,"metricName":"Reference GC (%)"}]]],"referenceName":"gi\_148609382\_ref\_NC","date":"18 July 2018, Wednesday, 16:20:14","order":[0,1,2,3,4,5,6,7,8,9,10,11,12,13],"assembliesNames":["ABySS\_127","ABySS\_63","CLC","IDBA\_UD","MEGAHIT","MIRA","Ray\_Meta","SOAPdenovo2","SPAdes","SPAdes\_meta","SPAdes\_sc","SPAdes\_sc\_careful","Velvet","Geneious"]},{"assembliesWithNs":null,"minContig":500,"report":[["Genome statistics",[{"values":["31.677","11.894","35.421","18.131","33.157","76.892","22.000","55.558","25.474","33.082","18.749","6.397","38.175"],"quality":"More is better","isMain":true,"metricName":"Genome fraction (%)"},{"values":["1.010","1.012","1.011","1.013","1.050","1.642","1.031","1.110","1.008","1.016","1.011","1.000","1.398"],"quality":"Less is better","isMain":true,"metricName":"Duplication ratio"},{"values":[2481,1363,1788,1217,2026,3970,4026,2127,4146,2666,2064,1122,3507],"quality":"More is better","isMain":true,"metricName":"Largest alignment"},{"values":[14031,5279,15710,8057,15272,54953,9946,27052,11231,14736,8279,2806,23412],"quality":"More is better","isMain":true,"metricName":"Total aligned length"},{"values":[null,null,null,null,null,46439,null,618,null,null,null,null,92482],"quality":"More is better","isMain":false,"metricName":"NG50"},{"values":[null,null,null,null,null,46439,null,null,null,null,null,null,92482],"quality":"More is better","isMain":false,"metricName":"NG75"},{"values":[676,1175,1092,829,703,null,1709,1166,2512,1169,1169,1056,null],"quality":"More is better","isMain":false,"metricName":"NA50"},{"values":[590,641,758,642,599,null,1123,679,1166,876,786,1056,null],"quality":"More is better","isMain":false,"metricName":"NA75"},{"values":[null,null,null,null,null,2218,null,618,null,null,null,null,464],"quality":"More is better","isMain":true,"metricName":"NGA50"},{"values":[null,null,null,null,null,1461,null,null,null,null,null,null,null],"quality":"More is better","isMain":false,"metricName":"NGA75"},{"values":[null,null,null,null,null,1,null,19,null,null,null,null,1],"quality":"Less is better","isMain":false,"metricName":"LG50"},{"values":[null,null,null,null,null,1,null,null,null,null,null,null,1],"quality":"Less is better","isMain":false,"metricName":"LG75"},{"values":[6,3,6,4,8,null,2,9,2,4,3,2,null],"quality":"Less is better","isMain":false,"metricName":"LA50"},{"values":[11,5,11,7,13,null,4,16,4,8,5,2,null],"quality":"Less is better","isMain":false,"metricName":"LA75"},{"values":[null,null,null,null,null,7,null,19,null,null,null,null,13],"quality":"Less is better","isMain":true,"metricName":"LGA50"},{"values":[null,null,null,null,null,14,null,null,null,null,null,null,null],"quality":"Less is better","isMain":false,"metricName":"LGA75"}]],["Misassemblies",[{"values":[0,1,0,0,0,0,0,0,0,0,0,0,0],"quality":"Less is better","isMain":true,"metricName":"# misassemblies"},{"values":[0,1,0,0,0,0,0,0,0,0,0,0,0],"quality":"Less is better","isMain":false,"metricName":" # relocations"},{"values":[0,0,0,0,0,0,0,0,0,0,0,0,0],"quality":"Less is better","isMain":false,"metricName":" # translocations"},{"values":[0,0,0,0,0,0,0,0,0,0,0,0,0],"quality":"Less is better","isMain":false,"metricName":" # inversions"},{"values":[0,1,0,0,0,0,0,0,0,0,0,0,0],"quality":"Less is better","isMain":false,"metricName":"# misassembled contigs"},{"values":[0,3680,0,0,0,0,0,0,0,0,0,0,0],"quality":"Less is better","isMain":true,"metricName":"Misassembled contigs length"},{"values":[0,0,0,0,0,1,0,0,0,0,0,0,0],"quality":"Less is better","isMain":false,"metricName":"# local misassemblies"},{"values":[0,0,0,0,0,4,0,0,0,0,0,0,4],"quality":"Less is better","isMain":false,"metricName":"# unaligned mis. contigs"}]],["Unaligned",[{"values":[0,0,0,0,0,0,0,0,0,0,0,0,0],"quality":"Less is better","isMain":false,"metricName":"# fully unaligned contigs"},{"values":[0,0,0,0,0,0,0,0,0,0,0,0,0],"quality":"Less is better","isMain":false,"metricName":"Fully unaligned length"},{"values":[0,1,0,0,0,7,0,0,0,0,0,0,4],"quality":"Less is better","isMain":false,"metricName":"# partially unaligned contigs"},{"values":[0,1226,0,0,0,120389,0,0,0,0,0,0,126686],"quality":"Less is better","isMain":false,"metricName":"Partially unaligned length"}]],["Mismatches",[{"values":[0,0,27,11,15,446,23,135,139,104,37,0,187],"quality":"Less is better","isMain":false,"metricName":"# mismatches"},{"values":[0,0,0,0,0,14,1,0,2,5,0,0,8],"quality":"Less is better","isMain":false,"metricName":"# indels"},{"values":[0,0,0,0,0,14,1,0,2,5,0,0,9],"quality":"Less is better","isMain":false,"metricName":"Indels length"},{"values":["0.00","0.00","173.78","138.31","103.14","1322.34","238.34","553.96","1243.96","716.70","449.90","0.00","1116.75"],"quality":"Less is better","isMain":true,"metricName":"# mismatches per 100 kbp"},{"values":["0.00","0.00","0.00","0.00","0.00","41.51","10.36","0.00","17.90","34.46","0.00","0.00","47.78"],"quality":"Less is better","isMain":true,"metricName":"# indels per 100 kbp"},{"values":[0,0,0,0,0,14,1,0,2,5,0,0,8],"quality":"Less is better","isMain":false,"metricName":" # indels (<= 5 bp)"},{"values":[0,0,0,0,0,0,0,0,0,0,0,0,0],"quality":"Less is better","isMain":false,"metricName":" # indels (> 5 bp)"},{"values":[0,108,0,0,0,28,0,0,0,0,0,0,877],"quality":"Less is better","isMain":false,"metricName":"# N's"},{"values":["0.00","1660.26","0.00","0.00","0.00","15.93","0.00","0.00","0.00","0.00","0.00","0.00","584.28"],"quality":"Less is better","isMain":true,"metricName":"# N's per 100 kbp"}]],["Statistics without reference",[{"values":[17,4,17,10,20,32,7,27,7,13,8,3,5],"quality":"Equal","isMain":true,"metricName":"# contigs"},{"values":[4,2,7,2,3,19,4,11,4,6,4,2,5],"quality":"Equal","isMain":false,"metricName":"# contigs (>= 1000 bp)"},{"values":[0,0,0,0,0,4,0,0,0,0,0,0,3],"quality":"Equal","isMain":false,"metricName":"# contigs (>= 5000 bp)"},{"values":[0,0,0,0,0,4,0,0,0,0,0,0,3],"quality":"Equal","isMain":false,"metricName":"# contigs (>= 10000 bp)"},{"values":[0,0,0,0,0,3,0,0,0,0,0,0,2],"quality":"Equal","isMain":false,"metricName":"# contigs (>= 25000 bp)"},{"values":[0,0,0,0,0,0,0,0,0,0,0,0,1],"quality":"Equal","isMain":false,"metricName":"# contigs (>= 50000 bp)"},{"values":[2481,3680,1788,1217,2026,46439,4026,2127,4146,2666,2064,1122,92482],"quality":"More is better","isMain":true,"metricName":"Largest contig"},{"values":[14031,6505,15710,8057,15272,175780,9946,27052,11266,14738,8313,2806,150098],"quality":"More is better","isMain":true,"metricName":"Total length"},{"values":[6211,5043,8963,2383,4504,167401,8096,16671,9218,9758,5589,2178,150098],"quality":"More is better","isMain":true,"metricName":"Total length (>= 1000 bp)"},{"values":[0,0,0,0,0,134634,0,0,0,0,0,0,143488],"quality":"More is better","isMain":false,"metricName":"Total length (>= 5000 bp)"},{"values":[0,0,0,0,0,134634,0,0,0,0,0,0,143488],"quality":"More is better","isMain":true,"metricName":"Total length (>= 10000 bp)"},{"values":[0,0,0,0,0,118536,0,0,0,0,0,0,132329],"quality":"More is better","isMain":false,"metricName":"Total length (>= 25000 bp)"},{"values":[0,0,0,0,0,0,0,0,0,0,0,0,92482],"quality":"More is better","isMain":true,"metricName":"Total length (>= 50000 bp)"},{"values":[676,3680,1092,829,703,43885,1709,1166,2512,1169,1169,1056,92482],"quality":"More is better","isMain":false,"metricName":"N50"},{"values":[590,1363,758,642,599,16098,1123,679,1166,876,786,1056,39847],"quality":"More is better","isMain":false,"metricName":"N75"},{"values":[6,1,6,4,8,2,2,9,2,4,3,2,1],"quality":"Less is better","isMain":false,"metricName":"L50"},{"values":[11,2,11,7,13,4,4,16,4,8,5,2,2],"quality":"Less is better","isMain":false,"metricName":"L75"},{"values":["33.98","31.01","33.77","32.49","33.20","34.38","32.30","33.89","33.31","33.13","32.80","28.69","34.08"],"quality":"Equal","isMain":false,"metricName":"GC (%)"}]],["Predicted genes",[]],["Similarity statistics",[{"values":[0,0,0,0,0,0,0,0,0,0,0,0,0],"quality":"Equal","isMain":false,"metricName":"# similar correct contigs"},{"values":[0,0,0,0,0,0,0,0,0,0,0,0,0],"quality":"Equal","isMain":false,"metricName":"# similar misassembled blocks"}]],["Reference statistics",[{"values":[43864,43864,43864,43864,43864,43864,43864,43864,43864,43864,43864,43864,43864],"quality":"Equal","isMain":false,"metricName":"Reference length"},{"values":[1,1,1,1,1,1,1,1,1,1,1,1,1],"quality":"Equal","isMain":false,"metricName":"Reference fragments"},{"values":["34.10","34.10","34.10","34.10","34.10","34.10","34.10","34.10","34.10","34.10","34.10","34.10","34.10"],"quality":"Equal","isMain":false,"metricName":"Reference GC (%)"}]]],"referenceName":"gi\_148717842\_ref\_NC","date":"18 July 2018, Wednesday, 16:20:25","order":[0,1,2,3,4,5,6,7,8,9,10,11,12],"assembliesNames":["ABySS\_127","ABySS\_63","CLC","IDBA\_UD","MEGAHIT","MIRA","Ray\_Meta","SPAdes","SPAdes\_meta","SPAdes\_sc","SPAdes\_sc\_careful","Velvet","Geneious"]},{"assembliesWithNs":null,"minContig":500,"report":[["Genome statistics",[{"values":["99.520","100.000","100.000","99.403","99.585","100.000","86.636","51.930","99.524","100.000","100.000","100.000","98.327"],"quality":"More is better","isMain":true,"metricName":"Genome fraction (%)"},{"values":["1.000","1.000","1.000","1.000","1.000","1.007","1.002","1.002","1.000","1.001","1.001","1.001","1.000"],"quality":"Less is better","isMain":true,"metricName":"Duplication ratio"},{"values":[45992,46214,46214,45938,46022,46520,15232,1561,45994,46269,46269,46269,23406],"quality":"More is better","isMain":true,"metricName":"Largest alignment"},{"values":[45992,46214,46214,45938,46022,46520,40107,24039,45994,46269,46269,46269,45461],"quality":"More is better","isMain":true,"metricName":"Total aligned length"},{"values":[45992,46214,46214,45938,46022,46520,10049,520,45994,46269,46269,46269,23406],"quality":"More is better","isMain":false,"metricName":"NG50"},{"values":[45992,46214,46214,45938,46022,46520,6181,null,45994,46269,46269,46269,20466],"quality":"More is better","isMain":false,"metricName":"NG75"},{"values":[45992,46214,46214,45938,46022,46520,10049,794,45994,46269,46269,46269,23406],"quality":"More is better","isMain":false,"metricName":"NA50"},{"values":[45992,46214,46214,45938,46022,46520,8645,632,45994,46269,46269,46269,20466],"quality":"More is better","isMain":false,"metricName":"NA75"},{"values":[45992,46214,46214,45938,46022,46520,10049,520,45994,46269,46269,46269,23406],"quality":"More is better","isMain":true,"metricName":"NGA50"},{"values":[45992,46214,46214,45938,46022,46520,6181,null,45994,46269,46269,46269,20466],"quality":"More is better","isMain":false,"metricName":"NGA75"},{"values":[1,1,1,1,1,1,2,29,1,1,1,1,1],"quality":"Less is better","isMain":false,"metricName":"LG50"},{"values":[1,1,1,1,1,1,4,null,1,1,1,1,2],"quality":"Less is better","isMain":false,"metricName":"LG75"},{"values":[1,1,1,1,1,1,2,12,1,1,1,1,1],"quality":"Less is better","isMain":false,"metricName":"LA50"},{"values":[1,1,1,1,1,1,3,20,1,1,1,1,2],"quality":"Less is better","isMain":false,"metricName":"LA75"},{"values":[1,1,1,1,1,1,2,29,1,1,1,1,1],"quality":"Less is better","isMain":true,"metricName":"LGA50"},{"values":[1,1,1,1,1,1,4,null,1,1,1,1,2],"quality":"Less is better","isMain":false,"metricName":"LGA75"}]],["Misassemblies",[{"values":[0,0,0,0,0,0,0,0,0,0,0,0,0],"quality":"Less is better","isMain":true,"metricName":"# misassemblies"},{"values":[0,0,0,0,0,0,0,0,0,0,0,0,0],"quality":"Less is better","isMain":false,"metricName":" # relocations"},{"values":[0,0,0,0,0,0,0,0,0,0,0,0,0],"quality":"Less is better","isMain":false,"metricName":" # translocations"},{"values":[0,0,0,0,0,0,0,0,0,0,0,0,0],"quality":"Less is better","isMain":false,"metricName":" # inversions"},{"values":[0,0,0,0,0,0,0,0,0,0,0,0,0],"quality":"Less is better","isMain":false,"metricName":"# misassembled contigs"},{"values":[0,0,0,0,0,0,0,0,0,0,0,0,0],"quality":"Less is better","isMain":true,"metricName":"Misassembled contigs length"},{"values":[0,0,0,0,0,1,0,0,0,0,0,0,0],"quality":"Less is better","isMain":false,"metricName":"# local misassemblies"},{"values":[0,0,0,0,0,0,0,0,0,0,0,0,0],"quality":"Less is better","isMain":false,"metricName":"# unaligned mis. contigs"}]],["Unaligned",[{"values":[0,0,0,0,0,0,0,0,0,0,0,0,0],"quality":"Less is better","isMain":false,"metricName":"# fully unaligned contigs"},{"values":[0,0,0,0,0,0,0,0,0,0,0,0,0],"quality":"Less is better","isMain":false,"metricName":"Fully unaligned length"},{"values":[0,0,0,0,0,0,0,0,0,0,0,0,0],"quality":"Less is better","isMain":false,"metricName":"# partially unaligned contigs"},{"values":[0,0,0,0,0,0,0,0,0,0,0,0,0],"quality":"Less is better","isMain":false,"metricName":"Partially unaligned length"}]],["Mismatches",[{"values":[0,0,0,0,0,0,0,3,0,0,0,0,0],"quality":"Less is better","isMain":false,"metricName":"# mismatches"},{"values":[0,0,0,0,0,0,0,0,0,1,1,1,0],"quality":"Less is better","isMain":false,"metricName":"# indels"},{"values":[0,0,0,0,0,0,0,0,0,55,55,55,0],"quality":"Less is better","isMain":false,"metricName":"Indels length"},{"values":["0.00","0.00","0.00","0.00","0.00","0.00","0.00","12.50","0.00","0.00","0.00","0.00","0.00"],"quality":"Less is better","isMain":true,"metricName":"# mismatches per 100 kbp"},{"values":["0.00","0.00","0.00","0.00","0.00","0.00","0.00","0.00","0.00","2.16","2.16","2.16","0.00"],"quality":"Less is better","isMain":true,"metricName":"# indels per 100 kbp"},{"values":[0,0,0,0,0,0,0,0,0,0,0,0,0],"quality":"Less is better","isMain":false,"metricName":" # indels (<= 5 bp)"},{"values":[0,0,0,0,0,0,0,0,0,1,1,1,0],"quality":"Less is better","isMain":false,"metricName":" # indels (> 5 bp)"},{"values":[0,0,0,0,0,1,0,0,0,0,0,0,0],"quality":"Less is better","isMain":false,"metricName":"# N's"},{"values":["0.00","0.00","0.00","0.00","0.00","2.15","0.00","0.00","0.00","0.00","0.00","0.00","0.00"],"quality":"Less is better","isMain":true,"metricName":"# N's per 100 kbp"}]],["Statistics without reference",[{"values":[1,1,1,1,1,1,4,30,1,1,1,1,3],"quality":"Equal","isMain":true,"metricName":"# contigs"},{"values":[1,1,1,1,1,1,4,7,1,1,1,1,3],"quality":"Equal","isMain":false,"metricName":"# contigs (>= 1000 bp)"},{"values":[1,1,1,1,1,1,4,0,1,1,1,1,2],"quality":"Equal","isMain":false,"metricName":"# contigs (>= 5000 bp)"},{"values":[1,1,1,1,1,1,2,0,1,1,1,1,2],"quality":"Equal","isMain":false,"metricName":"# contigs (>= 10000 bp)"},{"values":[1,1,1,1,1,1,0,0,1,1,1,1,0],"quality":"Equal","isMain":false,"metricName":"# contigs (>= 25000 bp)"},{"values":[0,0,0,0,0,0,0,0,0,0,0,0,0],"quality":"Equal","isMain":false,"metricName":"# contigs (>= 50000 bp)"},{"values":[45992,46214,46214,45938,46022,46520,15232,1561,45994,46269,46269,46269,23406],"quality":"More is better","isMain":true,"metricName":"Largest contig"},{"values":[45992,46214,46214,45938,46022,46520,40107,24039,45994,46269,46269,46269,45461],"quality":"More is better","isMain":true,"metricName":"Total length"},{"values":[45992,46214,46214,45938,46022,46520,40107,8410,45994,46269,46269,46269,45461],"quality":"More is better","isMain":true,"metricName":"Total length (>= 1000 bp)"},{"values":[45992,46214,46214,45938,46022,46520,40107,0,45994,46269,46269,46269,43872],"quality":"More is better","isMain":false,"metricName":"Total length (>= 5000 bp)"},{"values":[45992,46214,46214,45938,46022,46520,25281,0,45994,46269,46269,46269,43872],"quality":"More is better","isMain":true,"metricName":"Total length (>= 10000 bp)"},{"values":[45992,46214,46214,45938,46022,46520,0,0,45994,46269,46269,46269,0],"quality":"More is better","isMain":false,"metricName":"Total length (>= 25000 bp)"},{"values":[0,0,0,0,0,0,0,0,0,0,0,0,0],"quality":"More is better","isMain":true,"metricName":"Total length (>= 50000 bp)"},{"values":[45992,46214,46214,45938,46022,46520,10049,794,45994,46269,46269,46269,23406],"quality":"More is better","isMain":false,"metricName":"N50"},{"values":[45992,46214,46214,45938,46022,46520,8645,632,45994,46269,46269,46269,20466],"quality":"More is better","isMain":false,"metricName":"N75"},{"values":[1,1,1,1,1,1,2,12,1,1,1,1,1],"quality":"Less is better","isMain":false,"metricName":"L50"},{"values":[1,1,1,1,1,1,3,20,1,1,1,1,2],"quality":"Less is better","isMain":false,"metricName":"L75"},{"values":["54.93","54.97","54.97","54.93","54.93","54.94","54.83","55.06","54.93","54.98","54.98","54.98","54.91"],"quality":"Equal","isMain":false,"metricName":"GC (%)"}]],["Predicted genes",[]],["Similarity statistics",[{"values":[1,1,0,1,1,0,0,0,1,1,1,1,0],"quality":"Equal","isMain":false,"metricName":"# similar correct contigs"},{"values":[0,0,0,0,0,0,0,0,0,0,0,0,0],"quality":"Equal","isMain":false,"metricName":"# similar misassembled blocks"}]],["Reference statistics",[{"values":[46214,46214,46214,46214,46214,46214,46214,46214,46214,46214,46214,46214,46214],"quality":"Equal","isMain":false,"metricName":"Reference length"},{"values":[1,1,1,1,1,1,1,1,1,1,1,1,1],"quality":"Equal","isMain":false,"metricName":"Reference fragments"},{"values":["54.97","54.97","54.97","54.97","54.97","54.97","54.97","54.97","54.97","54.97","54.97","54.97","54.97"],"quality":"Equal","isMain":false,"metricName":"Reference GC (%)"}]]],"referenceName":"gi\_148724443\_ref\_NC","date":"18 July 2018, Wednesday, 16:20:37","order":[0,1,2,3,4,5,6,7,8,9,10,11,12],"assembliesNames":["ABySS\_127","ABySS\_63","CLC","IDBA\_UD","MEGAHIT","MIRA","Ray\_Meta","SOAPdenovo2","SPAdes","SPAdes\_meta","SPAdes\_sc","SPAdes\_sc\_careful","Velvet"]},{"assembliesWithNs":null,"minContig":500,"report":[["Genome statistics",[{"values":["99.929","100.000","100.000","100.000","100.000","100.000","100.000","100.000","100.000","100.000"],"quality":"More is better","isMain":true,"metricName":"Genome fraction (%)"},{"values":["1.001","1.003","1.005","1.020","1.016","1.004","1.002","1.002","1.002","1.002"],"quality":"Less is better","isMain":true,"metricName":"Duplication ratio"},{"values":[29718,29838,29880,30319,30219,29866,29794,29794,29794,29253],"quality":"More is better","isMain":true,"metricName":"Largest alignment"},{"values":[29718,29838,29880,30319,30219,29866,29794,29794,29794,29791],"quality":"More is better","isMain":true,"metricName":"Total aligned length"},{"values":[29739,29838,29880,30319,30219,29866,29794,29794,29794,29253],"quality":"More is better","isMain":false,"metricName":"NG50"},{"values":[29739,29838,29880,30319,30219,29866,29794,29794,29794,29253],"quality":"More is better","isMain":false,"metricName":"NG75"},{"values":[29718,29838,29880,30319,30219,29866,29794,29794,29794,29253],"quality":"More is better","isMain":false,"metricName":"NA50"},{"values":[29718,29838,29880,30319,30219,29866,29794,29794,29794,29253],"quality":"More is better","isMain":false,"metricName":"NA75"},{"values":[29718,29838,29880,30319,30219,29866,29794,29794,29794,29253],"quality":"More is better","isMain":true,"metricName":"NGA50"},{"values":[29718,29838,29880,30319,30219,29866,29794,29794,29794,29253],"quality":"More is better","isMain":false,"metricName":"NGA75"},{"values":[1,1,1,1,1,1,1,1,1,1],"quality":"Less is better","isMain":false,"metricName":"LG50"},{"values":[1,1,1,1,1,1,1,1,1,1],"quality":"Less is better","isMain":false,"metricName":"LG75"},{"values":[1,1,1,1,1,1,1,1,1,1],"quality":"Less is better","isMain":false,"metricName":"LA50"},{"values":[1,1,1,1,1,1,1,1,1,1],"quality":"Less is better","isMain":false,"metricName":"LA75"},{"values":[1,1,1,1,1,1,1,1,1,1],"quality":"Less is better","isMain":true,"metricName":"LGA50"},{"values":[1,1,1,1,1,1,1,1,1,1],"quality":"Less is better","isMain":false,"metricName":"LGA75"}]],["Misassemblies",[{"values":[0,0,0,0,0,0,0,0,0,0],"quality":"Less is better","isMain":true,"metricName":"# misassemblies"},{"values":[0,0,0,0,0,0,0,0,0,0],"quality":"Less is better","isMain":false,"metricName":" # relocations"},{"values":[0,0,0,0,0,0,0,0,0,0],"quality":"Less is better","isMain":false,"metricName":" # translocations"},{"values":[0,0,0,0,0,0,0,0,0,0],"quality":"Less is better","isMain":false,"metricName":" # inversions"},{"values":[0,0,0,0,0,0,0,0,0,0],"quality":"Less is better","isMain":false,"metricName":"# misassembled contigs"},{"values":[0,0,0,0,0,0,0,0,0,0],"quality":"Less is better","isMain":true,"metricName":"Misassembled contigs length"},{"values":[0,1,1,1,1,1,0,0,0,0],"quality":"Less is better","isMain":false,"metricName":"# local misassemblies"},{"values":[0,0,0,0,0,0,0,0,0,0],"quality":"Less is better","isMain":false,"metricName":"# unaligned mis. contigs"}]],["Unaligned",[{"values":[0,0,0,0,0,0,0,0,0,0],"quality":"Less is better","isMain":false,"metricName":"# fully unaligned contigs"},{"values":[0,0,0,0,0,0,0,0,0,0],"quality":"Less is better","isMain":false,"metricName":"Fully unaligned length"},{"values":[0,0,0,0,0,0,0,0,0,0],"quality":"Less is better","isMain":false,"metricName":"# partially unaligned contigs"},{"values":[0,0,0,0,0,0,0,0,0,0],"quality":"Less is better","isMain":false,"metricName":"Partially unaligned length"}]],["Mismatches",[{"values":[0,0,0,0,0,0,0,0,0,1],"quality":"Less is better","isMain":false,"metricName":"# mismatches"},{"values":[0,0,0,0,1,0,1,1,1,0],"quality":"Less is better","isMain":false,"metricName":"# indels"},{"values":[0,0,0,0,58,0,55,55,55,0],"quality":"Less is better","isMain":false,"metricName":"Indels length"},{"values":["0.00","0.00","0.00","0.00","0.00","0.00","0.00","0.00","0.00","3.36"],"quality":"Less is better","isMain":true,"metricName":"# mismatches per 100 kbp"},{"values":["0.00","0.00","0.00","0.00","3.36","0.00","3.36","3.36","3.36","0.00"],"quality":"Less is better","isMain":true,"metricName":"# indels per 100 kbp"},{"values":[0,0,0,0,0,0,0,0,0,0],"quality":"Less is better","isMain":false,"metricName":" # indels (<= 5 bp)"},{"values":[0,0,0,0,1,0,1,1,1,0],"quality":"Less is better","isMain":false,"metricName":" # indels (> 5 bp)"},{"values":[0,0,0,2,0,0,0,0,0,0],"quality":"Less is better","isMain":false,"metricName":"# N's"},{"values":["0.00","0.00","0.00","6.60","0.00","0.00","0.00","0.00","0.00","0.00"],"quality":"Less is better","isMain":true,"metricName":"# N's per 100 kbp"}]],["Statistics without reference",[{"values":[1,1,1,1,1,1,1,1,1,2],"quality":"Equal","isMain":true,"metricName":"# contigs"},{"values":[1,1,1,1,1,1,1,1,1,1],"quality":"Equal","isMain":false,"metricName":"# contigs (>= 1000 bp)"},{"values":[1,1,1,1,1,1,1,1,1,1],"quality":"Equal","isMain":false,"metricName":"# contigs (>= 5000 bp)"},{"values":[1,1,1,1,1,1,1,1,1,1],"quality":"Equal","isMain":false,"metricName":"# contigs (>= 10000 bp)"},{"values":[1,1,1,1,1,1,1,1,1,1],"quality":"Equal","isMain":false,"metricName":"# contigs (>= 25000 bp)"},{"values":[0,0,0,0,0,0,0,0,0,0],"quality":"Equal","isMain":false,"metricName":"# contigs (>= 50000 bp)"},{"values":[29739,29838,29880,30319,30219,29866,29794,29794,29794,29253],"quality":"More is better","isMain":true,"metricName":"Largest contig"},{"values":[29739,29838,29880,30319,30219,29866,29794,29794,29794,29791],"quality":"More is better","isMain":true,"metricName":"Total length"},{"values":[29739,29838,29880,30319,30219,29866,29794,29794,29794,29253],"quality":"More is better","isMain":true,"metricName":"Total length (>= 1000 bp)"},{"values":[29739,29838,29880,30319,30219,29866,29794,29794,29794,29253],"quality":"More is better","isMain":false,"metricName":"Total length (>= 5000 bp)"},{"values":[29739,29838,29880,30319,30219,29866,29794,29794,29794,29253],"quality":"More is better","isMain":true,"metricName":"Total length (>= 10000 bp)"},{"values":[29739,29838,29880,30319,30219,29866,29794,29794,29794,29253],"quality":"More is better","isMain":false,"metricName":"Total length (>= 25000 bp)"},{"values":[0,0,0,0,0,0,0,0,0,0],"quality":"More is better","isMain":true,"metricName":"Total length (>= 50000 bp)"},{"values":[29739,29838,29880,30319,30219,29866,29794,29794,29794,29253],"quality":"More is better","isMain":false,"metricName":"N50"},{"values":[29739,29838,29880,30319,30219,29866,29794,29794,29794,29253],"quality":"More is better","isMain":false,"metricName":"N75"},{"values":[1,1,1,1,1,1,1,1,1,1],"quality":"Less is better","isMain":false,"metricName":"L50"},{"values":[1,1,1,1,1,1,1,1,1,1],"quality":"Less is better","isMain":false,"metricName":"L75"},{"values":["54.03","54.00","54.00","53.88","53.98","54.03","54.01","54.03","54.01","54.05"],"quality":"Equal","isMain":false,"metricName":"GC (%)"}]],["Predicted genes",[]],["Similarity statistics",[{"values":[0,0,0,0,0,0,0,0,0,0],"quality":"Equal","isMain":false,"metricName":"# similar correct contigs"},{"values":[0,0,0,0,0,0,0,0,0,0],"quality":"Equal","isMain":false,"metricName":"# similar misassembled blocks"}]],["Reference statistics",[{"values":[29739,29739,29739,29739,29739,29739,29739,29739,29739,29739],"quality":"Equal","isMain":false,"metricName":"Reference length"},{"values":[1,1,1,1,1,1,1,1,1,1],"quality":"Equal","isMain":false,"metricName":"Reference fragments"},{"values":["54.02","54.02","54.02","54.02","54.02","54.02","54.02","54.02","54.02","54.02"],"quality":"Equal","isMain":false,"metricName":"Reference GC (%)"}]]],"referenceName":"gi\_148727082\_ref\_NC","date":"18 July 2018, Wednesday, 16:20:47","order":[0,1,2,3,4,5,6,7,8,9],"assembliesNames":["CLC","IDBA\_UD","MEGAHIT","MIRA","Ray\_Meta","SPAdes","SPAdes\_meta","SPAdes\_sc","SPAdes\_sc\_careful","Velvet"]},{"assembliesWithNs":null,"minContig":500,"report":[["Genome statistics",[{"values":["30.813","23.178","35.959","46.723","41.015","54.000","10.469","43.969","38.396","39.737","38.953","14.686","24.632"],"quality":"More is better","isMain":true,"metricName":"Genome fraction (%)"},{"values":["1.018","1.026","1.002","1.010","1.008","1.127","1.204","0.999","1.001","1.005","1.005","1.050","1.000"],"quality":"Less is better","isMain":true,"metricName":"Duplication ratio"},{"values":[1892,1493,3133,2496,2728,2607,2092,2728,3169,2327,2327,1427,6255],"quality":"More is better","isMain":true,"metricName":"Largest alignment"},{"values":[13970,10320,16010,21017,18399,27101,4662,19541,17094,17748,17398,6538,10971],"quality":"More is better","isMain":true,"metricName":"Total aligned length"},{"values":[null,null,null,null,null,1207,null,567,null,null,null,null,49757],"quality":"More is better","isMain":false,"metricName":"NG50"},{"values":[null,null,null,null,null,null,null,null,null,null,null,null,49757],"quality":"More is better","isMain":false,"metricName":"NG75"},{"values":[843,838,1483,1053,1262,1732,636,1334,2728,1219,1082,1089,null],"quality":"More is better","isMain":false,"metricName":"NA50"},{"values":[682,713,875,816,783,1132,null,564,1471,912,720,618,null],"quality":"More is better","isMain":false,"metricName":"NA75"},{"values":[null,null,null,null,null,1122,null,null,null,null,null,null,null],"quality":"More is better","isMain":true,"metricName":"NGA50"},{"values":[null,null,null,null,null,12,null,12,null,null,null,null,1],"quality":"Less is better","isMain":false,"metricName":"LG50"},{"values":[null,null,null,null,null,null,null,null,null,null,null,null,1],"quality":"Less is better","isMain":false,"metricName":"LG75"},{"values":[6,6,4,7,5,7,3,6,3,5,6,4,null],"quality":"Less is better","isMain":false,"metricName":"LA50"},{"values":[11,9,7,12,10,12,null,13,6,9,10,6,null],"quality":"Less is better","isMain":false,"metricName":"LA75"},{"values":[null,null,null,null,null,13,null,null,null,null,null,null,null],"quality":"Less is better","isMain":true,"metricName":"LGA50"}]],["Misassemblies",[{"values":[0,0,0,0,0,0,0,0,0,0,0,0,0],"quality":"Less is better","isMain":true,"metricName":"# misassemblies"},{"values":[0,0,0,0,0,0,0,0,0,0,0,0,0],"quality":"Less is better","isMain":false,"metricName":" # relocations"},{"values":[0,0,0,0,0,0,0,0,0,0,0,0,0],"quality":"Less is better","isMain":false,"metricName":" # translocations"},{"values":[0,0,0,0,0,0,0,0,0,0,0,0,0],"quality":"Less is better","isMain":false,"metricName":" # inversions"},{"values":[0,0,0,0,0,0,0,0,0,0,0,0,0],"quality":"Less is better","isMain":false,"metricName":"# misassembled contigs"},{"values":[0,0,0,0,0,0,0,0,0,0,0,0,0],"quality":"Less is better","isMain":true,"metricName":"Misassembled contigs length"},{"values":[0,0,0,0,0,0,0,0,0,0,0,0,0],"quality":"Less is better","isMain":false,"metricName":"# local misassemblies"},{"values":[0,0,0,0,0,0,0,0,0,0,0,0,1],"quality":"Less is better","isMain":false,"metricName":"# unaligned mis. contigs"}]],["Unaligned",[{"values":[0,0,0,0,0,0,0,0,0,0,0,0,0],"quality":"Less is better","isMain":false,"metricName":"# fully unaligned contigs"},{"values":[0,0,0,0,0,0,0,0,0,0,0,0,0],"quality":"Less is better","isMain":false,"metricName":"Fully unaligned length"},{"values":[0,0,0,0,0,1,2,2,0,0,0,1,2],"quality":"Less is better","isMain":false,"metricName":"# partially unaligned contigs"},{"values":[0,0,0,0,0,1008,1145,4573,0,0,0,1108,43924],"quality":"Less is better","isMain":false,"metricName":"Partially unaligned length"}]],["Mismatches",[{"values":[4,8,152,24,60,166,0,116,137,53,16,13,40],"quality":"Less is better","isMain":false,"metricName":"# mismatches"},{"values":[0,1,7,0,2,1,5,4,0,2,1,0,3],"quality":"Less is better","isMain":false,"metricName":"# indels"},{"values":[0,1,7,0,2,3,211,34,0,2,1,0,5],"quality":"Less is better","isMain":false,"metricName":"Indels length"},{"values":["29.16","77.53","949.47","115.38","328.59","690.49","0.00","592.59","801.45","299.59","92.26","198.84","364.76"],"quality":"Less is better","isMain":true,"metricName":"# mismatches per 100 kbp"},{"values":["0.00","9.69","43.73","0.00","10.95","4.16","107.27","20.43","0.00","11.31","5.77","0.00","27.36"],"quality":"Less is better","isMain":true,"metricName":"# indels per 100 kbp"},{"values":[0,1,7,0,2,1,1,2,0,2,1,0,3],"quality":"Less is better","isMain":false,"metricName":" # indels (<= 5 bp)"},{"values":[0,0,0,0,0,0,4,2,0,0,0,0,0],"quality":"Less is better","isMain":false,"metricName":" # indels (> 5 bp)"},{"values":[0,0,0,0,0,58,1574,0,0,0,0,10,798],"quality":"Less is better","isMain":false,"metricName":"# N's"},{"values":["0.00","0.00","0.00","0.00","0.00","206.34","23301.26","0.00","0.00","0.00","0.00","125.44","1453.68"],"quality":"Less is better","isMain":true,"metricName":"# N's per 100 kbp"}]],["Statistics without reference",[{"values":[16,13,13,20,17,19,8,15,9,15,17,7,3],"quality":"Equal","isMain":true,"metricName":"# contigs"},{"values":[4,1,5,8,5,14,1,7,8,8,7,5,3],"quality":"Equal","isMain":false,"metricName":"# contigs (>= 1000 bp)"},{"values":[0,0,0,0,0,0,0,1,0,0,0,0,1],"quality":"Equal","isMain":false,"metricName":"# contigs (>= 5000 bp)"},{"values":[0,0,0,0,0,0,0,0,0,0,0,0,1],"quality":"Equal","isMain":false,"metricName":"# contigs (>= 10000 bp)"},{"values":[0,0,0,0,0,0,0,0,0,0,0,0,1],"quality":"Equal","isMain":false,"metricName":"# contigs (>= 25000 bp)"},{"values":[0,0,0,0,0,0,0,0,0,0,0,0,0],"quality":"Equal","isMain":false,"metricName":"# contigs (>= 50000 bp)"},{"values":[1892,1493,3133,2496,2728,2607,2212,5587,3169,2345,2345,1454,49757],"quality":"More is better","isMain":true,"metricName":"Largest contig"},{"values":[13970,10590,16039,21017,18401,28109,6755,24128,17112,17777,17427,7972,54895],"quality":"More is better","isMain":true,"metricName":"Total length"},{"values":[5607,1493,10301,12537,9920,24527,2212,18945,16578,12795,10824,6661,54895],"quality":"More is better","isMain":true,"metricName":"Total length (>= 1000 bp)"},{"values":[0,0,0,0,0,0,0,5587,0,0,0,0,49757],"quality":"More is better","isMain":false,"metricName":"Total length (>= 5000 bp)"},{"values":[0,0,0,0,0,0,0,0,0,0,0,0,49757],"quality":"More is better","isMain":true,"metricName":"Total length (>= 10000 bp)"},{"values":[0,0,0,0,0,0,0,0,0,0,0,0,49757],"quality":"More is better","isMain":false,"metricName":"Total length (>= 25000 bp)"},{"values":[0,0,0,0,0,0,0,0,0,0,0,0,0],"quality":"More is better","isMain":true,"metricName":"Total length (>= 50000 bp)"},{"values":[843,838,1483,1053,1262,1884,725,2728,2728,1230,1082,1369,49757],"quality":"More is better","isMain":false,"metricName":"N50"},{"values":[682,713,875,816,783,1207,636,1097,1471,912,720,1089,49757],"quality":"More is better","isMain":false,"metricName":"N75"},{"values":[6,6,4,7,5,7,3,3,3,5,6,3,1],"quality":"Less is better","isMain":false,"metricName":"L50"},{"values":[11,9,7,12,10,12,5,7,6,9,10,5,1],"quality":"Less is better","isMain":false,"metricName":"L75"},{"values":["51.78","52.28","51.35","52.11","51.85","52.08","51.59","52.03","52.09","51.85","51.74","51.04","51.01"],"quality":"Equal","isMain":false,"metricName":"GC (%)"}]],["Predicted genes",[]],["Similarity statistics",[{"values":[0,0,0,0,0,0,0,0,0,0,0,0,0],"quality":"Equal","isMain":false,"metricName":"# similar correct contigs"},{"values":[0,0,0,0,0,0,0,0,0,0,0,0,0],"quality":"Equal","isMain":false,"metricName":"# similar misassembled blocks"}]],["Reference statistics",[{"values":[44520,44520,44520,44520,44520,44520,44520,44520,44520,44520,44520,44520,44520],"quality":"Equal","isMain":false,"metricName":"Reference length"},{"values":[1,1,1,1,1,1,1,1,1,1,1,1,1],"quality":"Equal","isMain":false,"metricName":"Reference fragments"},{"values":["51.90","51.90","51.90","51.90","51.90","51.90","51.90","51.90","51.90","51.90","51.90","51.90","51.90"],"quality":"Equal","isMain":false,"metricName":"Reference GC (%)"}]]],"referenceName":"gi\_148727181\_ref\_NC","date":"18 July 2018, Wednesday, 16:20:58","order":[0,1,2,3,4,5,6,7,8,9,10,11,12],"assembliesNames":["ABySS\_127","ABySS\_63","CLC","IDBA\_UD","MEGAHIT","MIRA","SOAPdenovo2","SPAdes","SPAdes\_meta","SPAdes\_sc","SPAdes\_sc\_careful","Velvet","Geneious"]},{"assembliesWithNs":null,"minContig":500,"report":[["Genome statistics",[{"values":["20.801","13.314","90.030","83.393","85.253","43.203","80.488","91.644","91.109","91.644","91.644","5.783","7.895"],"quality":"More is better","isMain":true,"metricName":"Genome fraction (%)"},{"values":["1.000","1.000","1.011","1.000","1.000","1.023","1.002","1.000","1.000","1.000","1.000","1.000","1.000"],"quality":"Less is better","isMain":true,"metricName":"Duplication ratio"},{"values":[1078,1359,5673,5505,5385,3566,4033,5673,5567,5673,5673,1265,2337],"quality":"More is better","isMain":true,"metricName":"Largest alignment"},{"values":[10380,6644,45432,41618,42558,22049,40226,45752,45482,45752,45752,2886,3940],"quality":"More is better","isMain":true,"metricName":"Total aligned length"},{"values":[null,null,1757,2337,1734,null,1629,2528,2528,2528,2528,null,null],"quality":"More is better","isMain":false,"metricName":"NG50"},{"values":[null,null,900,949,767,null,687,1140,976,1140,1140,null,null],"quality":"More is better","isMain":false,"metricName":"NG75"},{"values":[726,904,1786,2389,2301,1307,1787,2561,2561,2561,2561,1047,2337],"quality":"More is better","isMain":false,"metricName":"NA50"},{"values":[577,624,1140,1364,1285,1011,1094,1364,1364,1364,1364,1047,1603],"quality":"More is better","isMain":false,"metricName":"NA75"},{"values":[null,null,1757,2337,1734,null,1629,2528,2528,2528,2528,null,null],"quality":"More is better","isMain":true,"metricName":"NGA50"},{"values":[null,null,900,949,767,null,687,1140,976,1140,1140,null,null],"quality":"More is better","isMain":false,"metricName":"NGA75"},{"values":[null,null,10,7,8,null,10,7,7,7,7,null,null],"quality":"Less is better","isMain":false,"metricName":"LG50"},{"values":[null,null,19,16,19,null,23,15,15,15,15,null,null],"quality":"Less is better","isMain":false,"metricName":"LG75"},{"values":[6,3,9,6,6,6,8,6,6,6,6,2,1],"quality":"Less is better","isMain":false,"metricName":"LA50"},{"values":[11,6,16,11,13,11,15,12,12,12,12,2,2],"quality":"Less is better","isMain":false,"metricName":"LA75"},{"values":[null,null,10,7,8,null,10,7,7,7,7,null,null],"quality":"Less is better","isMain":true,"metricName":"LGA50"},{"values":[null,null,19,16,19,null,23,15,15,15,15,null,null],"quality":"Less is better","isMain":false,"metricName":"LGA75"}]],["Misassemblies",[{"values":[0,0,0,0,0,0,0,0,0,0,0,0,0],"quality":"Less is better","isMain":true,"metricName":"# misassemblies"},{"values":[0,0,0,0,0,0,0,0,0,0,0,0,0],"quality":"Less is better","isMain":false,"metricName":" # relocations"},{"values":[0,0,0,0,0,0,0,0,0,0,0,0,0],"quality":"Less is better","isMain":false,"metricName":" # translocations"},{"values":[0,0,0,0,0,0,0,0,0,0,0,0,0],"quality":"Less is better","isMain":false,"metricName":" # inversions"},{"values":[0,0,0,0,0,0,0,0,0,0,0,0,0],"quality":"Less is better","isMain":false,"metricName":"# misassembled contigs"},{"values":[0,0,0,0,0,0,0,0,0,0,0,0,0],"quality":"Less is better","isMain":true,"metricName":"Misassembled contigs length"},{"values":[0,0,0,0,0,0,0,0,0,0,0,0,0],"quality":"Less is better","isMain":false,"metricName":"# local misassemblies"},{"values":[0,0,0,0,0,0,0,0,0,0,0,0,0],"quality":"Less is better","isMain":false,"metricName":"# unaligned mis. contigs"}]],["Unaligned",[{"values":[0,0,0,0,0,0,0,0,0,0,0,0,0],"quality":"Less is better","isMain":false,"metricName":"# fully unaligned contigs"},{"values":[0,0,0,0,0,0,0,0,0,0,0,0,0],"quality":"Less is better","isMain":false,"metricName":"Fully unaligned length"},{"values":[0,0,0,0,0,0,0,0,0,0,0,0,0],"quality":"Less is better","isMain":false,"metricName":"# partially unaligned contigs"},{"values":[0,0,0,0,0,0,0,0,0,0,0,0,0],"quality":"Less is better","isMain":false,"metricName":"Partially unaligned length"}]],["Mismatches",[{"values":[0,3,45,20,23,9,22,44,33,31,23,0,1],"quality":"Less is better","isMain":false,"metricName":"# mismatches"},{"values":[0,0,0,0,0,0,0,0,0,0,0,0,0],"quality":"Less is better","isMain":false,"metricName":"# indels"},{"values":[0,0,0,0,0,0,0,0,0,0,0,0,0],"quality":"Less is better","isMain":false,"metricName":"Indels length"},{"values":["0.00","45.15","100.16","48.06","54.06","41.75","54.77","96.21","72.58","67.79","50.29","0.00","25.38"],"quality":"Less is better","isMain":true,"metricName":"# mismatches per 100 kbp"},{"values":["0.00","0.00","0.00","0.00","0.00","0.00","0.00","0.00","0.00","0.00","0.00","0.00","0.00"],"quality":"Less is better","isMain":true,"metricName":"# indels per 100 kbp"},{"values":[0,0,0,0,0,0,0,0,0,0,0,0,0],"quality":"Less is better","isMain":false,"metricName":" # indels (<= 5 bp)"},{"values":[0,0,0,0,0,0,0,0,0,0,0,0,0],"quality":"Less is better","isMain":false,"metricName":" # indels (> 5 bp)"},{"values":[0,0,0,0,0,29,0,0,0,0,0,0,8],"quality":"Less is better","isMain":false,"metricName":"# N's"},{"values":["0.00","0.00","0.00","0.00","0.00","131.53","0.00","0.00","0.00","0.00","0.00","0.00","203.05"],"quality":"Less is better","isMain":true,"metricName":"# N's per 100 kbp"}]],["Statistics without reference",[{"values":[15,8,30,22,26,18,27,25,25,25,25,3,2],"quality":"Equal","isMain":true,"metricName":"# contigs"},{"values":[2,2,17,14,14,12,16,15,14,15,15,2,2],"quality":"Equal","isMain":false,"metricName":"# contigs (>= 1000 bp)"},{"values":[0,0,1,2,2,0,0,2,2,2,2,0,0],"quality":"Equal","isMain":false,"metricName":"# contigs (>= 5000 bp)"},{"values":[0,0,0,0,0,0,0,0,0,0,0,0,0],"quality":"Equal","isMain":false,"metricName":"# contigs (>= 10000 bp)"},{"values":[0,0,0,0,0,0,0,0,0,0,0,0,0],"quality":"Equal","isMain":false,"metricName":"# contigs (>= 25000 bp)"},{"values":[0,0,0,0,0,0,0,0,0,0,0,0,0],"quality":"Equal","isMain":false,"metricName":"# contigs (>= 50000 bp)"},{"values":[1078,1359,5673,5505,5385,3566,4033,5673,5567,5673,5673,1265,2337],"quality":"More is better","isMain":true,"metricName":"Largest contig"},{"values":[10380,6644,45435,41618,42558,22049,40226,45752,45482,45752,45752,2886,3940],"quality":"More is better","isMain":true,"metricName":"Total length"},{"values":[2118,2550,35881,35650,33737,17874,32144,38189,36943,38189,38189,2312,3940],"quality":"More is better","isMain":true,"metricName":"Total length (>= 1000 bp)"},{"values":[0,0,5673,11001,10658,0,0,11169,11063,11169,11169,0,0],"quality":"More is better","isMain":false,"metricName":"Total length (>= 5000 bp)"},{"values":[0,0,0,0,0,0,0,0,0,0,0,0,0],"quality":"More is better","isMain":true,"metricName":"Total length (>= 10000 bp)"},{"values":[0,0,0,0,0,0,0,0,0,0,0,0,0],"quality":"More is better","isMain":false,"metricName":"Total length (>= 25000 bp)"},{"values":[0,0,0,0,0,0,0,0,0,0,0,0,0],"quality":"More is better","isMain":true,"metricName":"Total length (>= 50000 bp)"},{"values":[726,904,1786,2389,2301,1307,1787,2561,2561,2561,2561,1047,2337],"quality":"More is better","isMain":false,"metricName":"N50"},{"values":[577,624,1140,1364,1285,1011,1094,1364,1364,1364,1364,1047,1603],"quality":"More is better","isMain":false,"metricName":"N75"},{"values":[6,3,9,6,6,6,8,6,6,6,6,2,1],"quality":"Less is better","isMain":false,"metricName":"L50"},{"values":[11,6,16,11,13,11,15,12,12,12,12,2,2],"quality":"Less is better","isMain":false,"metricName":"L75"},{"values":["42.32","42.52","42.65","42.77","42.65","42.30","42.54","42.69","42.72","42.69","42.69","43.24","39.09"],"quality":"Equal","isMain":false,"metricName":"GC (%)"}]],["Predicted genes",[]],["Similarity statistics",[{"values":[0,0,0,0,0,0,0,0,0,0,0,0,0],"quality":"Equal","isMain":false,"metricName":"# similar correct contigs"},{"values":[0,0,0,0,0,0,0,0,0,0,0,0,0],"quality":"Equal","isMain":false,"metricName":"# similar misassembled blocks"}]],["Reference statistics",[{"values":[49902,49902,49902,49902,49902,49902,49902,49902,49902,49902,49902,49902,49902],"quality":"Equal","isMain":false,"metricName":"Reference length"},{"values":[1,1,1,1,1,1,1,1,1,1,1,1,1],"quality":"Equal","isMain":false,"metricName":"Reference fragments"},{"values":["42.68","42.68","42.68","42.68","42.68","42.68","42.68","42.68","42.68","42.68","42.68","42.68","42.68"],"quality":"Equal","isMain":false,"metricName":"Reference GC (%)"}]]],"referenceName":"gi\_148734472\_ref\_NC","date":"18 July 2018, Wednesday, 16:21:09","order":[0,1,2,3,4,5,6,7,8,9,10,11,12],"assembliesNames":["ABySS\_127","ABySS\_63","CLC","IDBA\_UD","MEGAHIT","MIRA","SOAPdenovo2","SPAdes","SPAdes\_meta","SPAdes\_sc","SPAdes\_sc\_careful","Velvet","Geneious"]},{"assembliesWithNs":null,"minContig":500,"report":[["Genome statistics",[{"values":["71.997","66.921","71.174","62.742","67.545","99.082","12.403","28.833","72.992","48.860","59.128","51.475","39.304","100.000"],"quality":"More is better","isMain":true,"metricName":"Genome fraction (%)"},{"values":["1.016","1.059","1.001","1.012","1.016","1.450","1.006","1.060","1.002","1.002","1.007","1.006","1.003","1.503"],"quality":"Less is better","isMain":true,"metricName":"Duplication ratio"},{"values":[9216,11965,12933,9058,9658,21454,1260,3694,9630,6775,5410,5410,3497,21454],"quality":"More is better","isMain":true,"metricName":"Largest alignment"},{"values":[35593,34508,34687,30903,33418,69758,6074,14198,35611,23837,28989,25209,19177,73011],"quality":"More is better","isMain":true,"metricName":"Total aligned length"},{"values":[27368,4071,3951,1442,2610,13449,null,null,3526,null,1256,559,null,54122],"quality":"More is better","isMain":false,"metricName":"NG50"},{"values":[9216,710,null,null,null,12094,null,null,null,null,null,null,null,54122],"quality":"More is better","isMain":false,"metricName":"NG75"},{"values":[2666,4071,4066,4105,4325,5044,734,907,4276,3547,3547,3728,1856,3918],"quality":"More is better","isMain":false,"metricName":"NA50"},{"values":[null,1281,3118,2385,2502,2465,596,null,2724,1933,1488,2236,1177,null],"quality":"More is better","isMain":false,"metricName":"NA75"},{"values":[3870,2272,3951,1442,2610,13449,null,null,3526,null,1256,559,null,14398],"quality":"More is better","isMain":true,"metricName":"NGA50"},{"values":[null,null,null,null,null,5044,null,null,null,null,null,null,null,9939],"quality":"More is better","isMain":false,"metricName":"NGA75"},{"values":[1,3,4,6,5,2,null,null,5,null,9,10,null,1],"quality":"Less is better","isMain":false,"metricName":"LG50"},{"values":[2,11,null,null,null,3,null,null,null,null,null,null,null,1],"quality":"Less is better","isMain":false,"metricName":"LG75"},{"values":[5,3,3,3,3,3,4,5,3,3,4,3,4,7],"quality":"Less is better","isMain":false,"metricName":"LA50"},{"values":[null,7,5,5,6,8,6,null,6,5,7,5,7,null],"quality":"Less is better","isMain":false,"metricName":"LA75"},{"values":[4,5,4,6,5,2,null,null,5,null,9,10,null,2],"quality":"Less is better","isMain":true,"metricName":"LGA50"},{"values":[null,null,null,null,null,3,null,null,null,null,null,null,null,3],"quality":"Less is better","isMain":false,"metricName":"LGA75"}]],["Misassemblies",[{"values":[0,0,0,0,0,0,0,0,0,0,0,0,0,0],"quality":"Less is better","isMain":true,"metricName":"# misassemblies"},{"values":[0,0,0,0,0,0,0,0,0,0,0,0,0,0],"quality":"Less is better","isMain":false,"metricName":" # relocations"},{"values":[0,0,0,0,0,0,0,0,0,0,0,0,0,0],"quality":"Less is better","isMain":false,"metricName":" # translocations"},{"values":[0,0,0,0,0,0,0,0,0,0,0,0,0,0],"quality":"Less is better","isMain":false,"metricName":" # inversions"},{"values":[0,0,0,0,0,0,0,0,0,0,0,0,0,0],"quality":"Less is better","isMain":false,"metricName":"# misassembled contigs"},{"values":[0,0,0,0,0,0,0,0,0,0,0,0,0,0],"quality":"Less is better","isMain":true,"metricName":"Misassembled contigs length"},{"values":[0,1,0,0,0,0,0,0,0,0,0,0,0,0],"quality":"Less is better","isMain":false,"metricName":"# local misassemblies"},{"values":[0,0,0,0,0,1,0,4,0,0,0,0,0,2],"quality":"Less is better","isMain":false,"metricName":"# unaligned mis. contigs"}]],["Unaligned",[{"values":[0,0,0,0,0,0,0,0,0,0,0,0,0,0],"quality":"Less is better","isMain":false,"metricName":"# fully unaligned contigs"},{"values":[0,0,0,0,0,0,0,0,0,0,0,0,0,0],"quality":"Less is better","isMain":false,"metricName":"Fully unaligned length"},{"values":[1,1,0,0,0,1,0,4,0,0,0,0,0,2],"quality":"Less is better","isMain":false,"metricName":"# partially unaligned contigs"},{"values":[19335,4148,0,0,0,8156,0,7220,0,0,0,0,0,52733],"quality":"Less is better","isMain":false,"metricName":"Partially unaligned length"}]],["Mismatches",[{"values":[27,169,6,6,16,654,0,0,15,93,31,13,0,603],"quality":"Less is better","isMain":false,"metricName":"# mismatches"},{"values":[0,0,0,0,0,15,0,7,0,0,0,0,0,15],"quality":"Less is better","isMain":false,"metricName":"# indels"},{"values":[0,0,0,0,0,16,0,338,0,0,0,0,0,19],"quality":"Less is better","isMain":false,"metricName":"Indels length"},{"values":["77.05","518.83","17.32","19.65","48.67","1356.09","0.00","0.00","42.22","391.05","107.71","51.89","0.00","1238.85"],"quality":"Less is better","isMain":true,"metricName":"# mismatches per 100 kbp"},{"values":["0.00","0.00","0.00","0.00","0.00","31.10","0.00","49.88","0.00","0.00","0.00","0.00","0.00","30.82"],"quality":"Less is better","isMain":true,"metricName":"# indels per 100 kbp"},{"values":[0,0,0,0,0,15,0,0,0,0,0,0,0,15],"quality":"Less is better","isMain":false,"metricName":" # indels (<= 5 bp)"},{"values":[0,0,0,0,0,0,0,7,0,0,0,0,0,0],"quality":"Less is better","isMain":false,"metricName":" # indels (> 5 bp)"},{"values":[50,1,0,0,0,2,0,4044,0,0,0,0,0,226],"quality":"Less is better","isMain":false,"metricName":"# N's"},{"values":["91.03","2.59","0.00","0.00","0.00","2.56","0.00","18296.99","0.00","0.00","0.00","0.00","0.00","179.49"],"quality":"Less is better","isMain":true,"metricName":"# N's per 100 kbp"}]],["Statistics without reference",[{"values":[13,14,11,12,12,15,8,10,13,11,14,11,12,5],"quality":"Equal","isMain":true,"metricName":"# contigs"},{"values":[7,9,8,9,9,13,1,9,8,7,11,9,8,5],"quality":"Equal","isMain":false,"metricName":"# contigs (>= 1000 bp)"},{"values":[2,2,1,1,1,5,0,0,1,1,1,1,0,5],"quality":"Equal","isMain":false,"metricName":"# contigs (>= 5000 bp)"},{"values":[1,1,1,0,0,3,0,0,0,0,0,0,0,4],"quality":"Equal","isMain":false,"metricName":"# contigs (>= 10000 bp)"},{"values":[1,0,0,0,0,0,0,0,0,0,0,0,0,2],"quality":"Equal","isMain":false,"metricName":"# contigs (>= 25000 bp)"},{"values":[0,0,0,0,0,0,0,0,0,0,0,0,0,1],"quality":"Equal","isMain":false,"metricName":"# contigs (>= 50000 bp)"},{"values":[27368,16113,12933,9058,9658,21619,1260,4444,9630,6775,5410,5410,3497,54122],"quality":"More is better","isMain":true,"metricName":"Largest contig"},{"values":[54928,38656,34687,30903,33418,78079,6074,22102,35611,23838,28989,25209,19184,125911],"quality":"More is better","isMain":true,"metricName":"Total length"},{"values":[51614,35551,32514,28864,31823,76829,1260,21422,32821,20980,27243,24131,16003,125911],"quality":"More is better","isMain":true,"metricName":"Total length (>= 1000 bp)"},{"values":[36584,21537,12933,9058,9658,57219,0,0,9630,6775,5410,5410,0,125911],"quality":"More is better","isMain":false,"metricName":"Total length (>= 5000 bp)"},{"values":[27368,16113,12933,0,0,47162,0,0,0,0,0,0,0,120543],"quality":"More is better","isMain":true,"metricName":"Total length (>= 10000 bp)"},{"values":[27368,0,0,0,0,0,0,0,0,0,0,0,0,84526],"quality":"More is better","isMain":false,"metricName":"Total length (>= 25000 bp)"},{"values":[0,0,0,0,0,0,0,0,0,0,0,0,0,54122],"quality":"More is better","isMain":true,"metricName":"Total length (>= 50000 bp)"},{"values":[9216,5424,4066,4105,4325,12094,734,3348,4276,3547,3547,3728,1856,30404],"quality":"More is better","isMain":false,"metricName":"N50"},{"values":[3870,2272,3118,2385,2502,4621,596,2292,2724,1933,1488,2236,1177,21619],"quality":"More is better","isMain":false,"metricName":"N75"},{"values":[2,2,3,3,3,3,4,3,3,3,4,3,4,2],"quality":"Less is better","isMain":false,"metricName":"L50"},{"values":[4,5,5,5,6,6,6,5,6,5,7,5,7,3],"quality":"Less is better","isMain":false,"metricName":"L75"},{"values":["60.12","59.68","59.77","59.29","59.68","61.04","55.24","58.32","59.84","59.37","59.39","58.58","57.57","60.95"],"quality":"Equal","isMain":false,"metricName":"GC (%)"}]],["Predicted genes",[]],["Similarity statistics",[{"values":[0,0,0,0,0,0,0,0,0,0,0,0,0,0],"quality":"Equal","isMain":false,"metricName":"# similar correct contigs"},{"values":[0,0,0,0,0,0,0,0,0,0,0,0,0,0],"quality":"Equal","isMain":false,"metricName":"# similar misassembled blocks"}]],["Reference statistics",[{"values":[48674,48674,48674,48674,48674,48674,48674,48674,48674,48674,48674,48674,48674,48674],"quality":"Equal","isMain":false,"metricName":"Reference length"},{"values":[1,1,1,1,1,1,1,1,1,1,1,1,1,1],"quality":"Equal","isMain":false,"metricName":"Reference fragments"},{"values":["60.45","60.45","60.45","60.45","60.45","60.45","60.45","60.45","60.45","60.45","60.45","60.45","60.45","60.45"],"quality":"Equal","isMain":false,"metricName":"Reference GC (%)"}]]],"referenceName":"gi\_148747798\_ref\_NC","date":"18 July 2018, Wednesday, 16:21:21","order":[0,1,2,3,4,5,6,7,8,9,10,11,12,13],"assembliesNames":["ABySS\_127","ABySS\_63","CLC","IDBA\_UD","MEGAHIT","MIRA","Ray\_Meta","SOAPdenovo2","SPAdes","SPAdes\_meta","SPAdes\_sc","SPAdes\_sc\_careful","Velvet","Geneious"]},{"assembliesWithNs":null,"minContig":500,"report":[["Genome statistics",[{"values":["100.000","100.000","100.000","100.000","100.000","4.648","100.000","100.000","100.000","100.000","100.000"],"quality":"More is better","isMain":true,"metricName":"Genome fraction (%)"},{"values":["1.000","1.003","1.004","1.010","1.001","1.000","1.004","1.002","1.002","1.002","1.001"],"quality":"Less is better","isMain":true,"metricName":"Duplication ratio"},{"values":[34659,34758,34800,34994,34706,555,34786,34714,34714,34714,34689],"quality":"More is better","isMain":true,"metricName":"Largest alignment"},{"values":[34659,34758,34800,34994,34706,1611,34786,34714,34714,34714,34689],"quality":"More is better","isMain":true,"metricName":"Total aligned length"},{"values":[34659,34758,34800,34994,34706,null,34786,34714,34714,34714,34689],"quality":"More is better","isMain":false,"metricName":"NG50"},{"values":[34659,34758,34800,34994,34706,null,34786,34714,34714,34714,34689],"quality":"More is better","isMain":false,"metricName":"NG75"},{"values":[34659,34758,34800,34994,34706,553,34786,34714,34714,34714,34689],"quality":"More is better","isMain":false,"metricName":"NA50"},{"values":[34659,34758,34800,34994,34706,503,34786,34714,34714,34714,34689],"quality":"More is better","isMain":false,"metricName":"NA75"},{"values":[34659,34758,34800,34994,34706,null,34786,34714,34714,34714,34689],"quality":"More is better","isMain":true,"metricName":"NGA50"},{"values":[34659,34758,34800,34994,34706,null,34786,34714,34714,34714,34689],"quality":"More is better","isMain":false,"metricName":"NGA75"},{"values":[1,1,1,1,1,null,1,1,1,1,1],"quality":"Less is better","isMain":false,"metricName":"LG50"},{"values":[1,1,1,1,1,null,1,1,1,1,1],"quality":"Less is better","isMain":false,"metricName":"LG75"},{"values":[1,1,1,1,1,2,1,1,1,1,1],"quality":"Less is better","isMain":false,"metricName":"LA50"},{"values":[1,1,1,1,1,3,1,1,1,1,1],"quality":"Less is better","isMain":false,"metricName":"LA75"},{"values":[1,1,1,1,1,null,1,1,1,1,1],"quality":"Less is better","isMain":true,"metricName":"LGA50"},{"values":[1,1,1,1,1,null,1,1,1,1,1],"quality":"Less is better","isMain":false,"metricName":"LGA75"}]],["Misassemblies",[{"values":[0,0,0,0,0,0,0,0,0,0,0],"quality":"Less is better","isMain":true,"metricName":"# misassemblies"},{"values":[0,0,0,0,0,0,0,0,0,0,0],"quality":"Less is better","isMain":false,"metricName":" # relocations"},{"values":[0,0,0,0,0,0,0,0,0,0,0],"quality":"Less is better","isMain":false,"metricName":" # translocations"},{"values":[0,0,0,0,0,0,0,0,0,0,0],"quality":"Less is better","isMain":false,"metricName":" # inversions"},{"values":[0,0,0,0,0,0,0,0,0,0,0],"quality":"Less is better","isMain":false,"metricName":"# misassembled contigs"},{"values":[0,0,0,0,0,0,0,0,0,0,0],"quality":"Less is better","isMain":true,"metricName":"Misassembled contigs length"},{"values":[0,1,1,1,0,0,1,0,0,0,0],"quality":"Less is better","isMain":false,"metricName":"# local misassemblies"},{"values":[0,0,0,0,0,0,0,0,0,0,0],"quality":"Less is better","isMain":false,"metricName":"# unaligned mis. contigs"}]],["Unaligned",[{"values":[0,0,0,0,0,0,0,0,0,0,0],"quality":"Less is better","isMain":false,"metricName":"# fully unaligned contigs"},{"values":[0,0,0,0,0,0,0,0,0,0,0],"quality":"Less is better","isMain":false,"metricName":"Fully unaligned length"},{"values":[0,0,0,0,0,0,0,0,0,0,0],"quality":"Less is better","isMain":false,"metricName":"# partially unaligned contigs"},{"values":[0,0,0,0,0,0,0,0,0,0,0],"quality":"Less is better","isMain":false,"metricName":"Partially unaligned length"}]],["Mismatches",[{"values":[0,0,0,0,0,0,0,0,0,0,0],"quality":"Less is better","isMain":false,"metricName":"# mismatches"},{"values":[0,0,0,0,1,0,0,1,1,1,1],"quality":"Less is better","isMain":false,"metricName":"# indels"},{"values":[0,0,0,0,47,0,0,55,55,55,30],"quality":"Less is better","isMain":false,"metricName":"Indels length"},{"values":["0.00","0.00","0.00","0.00","0.00","0.00","0.00","0.00","0.00","0.00","0.00"],"quality":"Less is better","isMain":true,"metricName":"# mismatches per 100 kbp"},{"values":["0.00","0.00","0.00","0.00","2.89","0.00","0.00","2.89","2.89","2.89","2.89"],"quality":"Less is better","isMain":true,"metricName":"# indels per 100 kbp"},{"values":[0,0,0,0,0,0,0,0,0,0,0],"quality":"Less is better","isMain":false,"metricName":" # indels (<= 5 bp)"},{"values":[0,0,0,0,1,0,0,1,1,1,1],"quality":"Less is better","isMain":false,"metricName":" # indels (> 5 bp)"},{"values":[0,0,0,1,0,0,0,0,0,0,0],"quality":"Less is better","isMain":false,"metricName":"# N's"},{"values":["0.00","0.00","0.00","2.86","0.00","0.00","0.00","0.00","0.00","0.00","0.00"],"quality":"Less is better","isMain":true,"metricName":"# N's per 100 kbp"}]],["Statistics without reference",[{"values":[1,1,1,1,1,3,1,1,1,1,1],"quality":"Equal","isMain":true,"metricName":"# contigs"},{"values":[1,1,1,1,1,0,1,1,1,1,1],"quality":"Equal","isMain":false,"metricName":"# contigs (>= 1000 bp)"},{"values":[1,1,1,1,1,0,1,1,1,1,1],"quality":"Equal","isMain":false,"metricName":"# contigs (>= 5000 bp)"},{"values":[1,1,1,1,1,0,1,1,1,1,1],"quality":"Equal","isMain":false,"metricName":"# contigs (>= 10000 bp)"},{"values":[1,1,1,1,1,0,1,1,1,1,1],"quality":"Equal","isMain":false,"metricName":"# contigs (>= 25000 bp)"},{"values":[0,0,0,0,0,0,0,0,0,0,0],"quality":"Equal","isMain":false,"metricName":"# contigs (>= 50000 bp)"},{"values":[34659,34758,34800,34994,34706,555,34786,34714,34714,34714,34689],"quality":"More is better","isMain":true,"metricName":"Largest contig"},{"values":[34659,34758,34800,34994,34706,1611,34786,34714,34714,34714,34689],"quality":"More is better","isMain":true,"metricName":"Total length"},{"values":[34659,34758,34800,34994,34706,0,34786,34714,34714,34714,34689],"quality":"More is better","isMain":true,"metricName":"Total length (>= 1000 bp)"},{"values":[34659,34758,34800,34994,34706,0,34786,34714,34714,34714,34689],"quality":"More is better","isMain":false,"metricName":"Total length (>= 5000 bp)"},{"values":[34659,34758,34800,34994,34706,0,34786,34714,34714,34714,34689],"quality":"More is better","isMain":true,"metricName":"Total length (>= 10000 bp)"},{"values":[34659,34758,34800,34994,34706,0,34786,34714,34714,34714,34689],"quality":"More is better","isMain":false,"metricName":"Total length (>= 25000 bp)"},{"values":[0,0,0,0,0,0,0,0,0,0,0],"quality":"More is better","isMain":true,"metricName":"Total length (>= 50000 bp)"},{"values":[34659,34758,34800,34994,34706,553,34786,34714,34714,34714,34689],"quality":"More is better","isMain":false,"metricName":"N50"},{"values":[34659,34758,34800,34994,34706,503,34786,34714,34714,34714,34689],"quality":"More is better","isMain":false,"metricName":"N75"},{"values":[1,1,1,1,1,2,1,1,1,1,1],"quality":"Less is better","isMain":false,"metricName":"L50"},{"values":[1,1,1,1,1,3,1,1,1,1,1],"quality":"Less is better","isMain":false,"metricName":"L75"},{"values":["47.82","47.79","47.78","47.77","47.79","48.98","47.83","47.80","47.81","47.81","47.82"],"quality":"Equal","isMain":false,"metricName":"GC (%)"}]],["Predicted genes",[]],["Similarity statistics",[{"values":[0,0,0,0,0,0,0,0,0,0,0],"quality":"Equal","isMain":false,"metricName":"# similar correct contigs"},{"values":[0,0,0,0,0,0,0,0,0,0,0],"quality":"Equal","isMain":false,"metricName":"# similar misassembled blocks"}]],["Reference statistics",[{"values":[34659,34659,34659,34659,34659,34659,34659,34659,34659,34659,34659],"quality":"Equal","isMain":false,"metricName":"Reference length"},{"values":[1,1,1,1,1,1,1,1,1,1,1],"quality":"Equal","isMain":false,"metricName":"Reference fragments"},{"values":["47.82","47.82","47.82","47.82","47.82","47.82","47.82","47.82","47.82","47.82","47.82"],"quality":"Equal","isMain":false,"metricName":"Reference GC (%)"}]]],"referenceName":"gi\_148750834\_ref\_NC","date":"18 July 2018, Wednesday, 16:21:31","order":[0,1,2,3,4,5,6,7,8,9,10],"assembliesNames":["CLC","IDBA\_UD","MEGAHIT","MIRA","Ray\_Meta","SOAPdenovo2","SPAdes","SPAdes\_meta","SPAdes\_sc","SPAdes\_sc\_careful","Velvet"]},{"assembliesWithNs":null,"minContig":500,"report":[["Genome statistics",[{"values":["100.000","100.000","100.000","100.000","100.000","28.821","100.000","100.000","100.000","100.000","100.000"],"quality":"More is better","isMain":true,"metricName":"Genome fraction (%)"},{"values":["1.000","1.003","1.003","1.006","1.007","1.000","1.003","1.001","1.001","1.001","1.001"],"quality":"Less is better","isMain":true,"metricName":"Duplication ratio"},{"values":[46365,46514,46506,46653,46704,1196,46492,46420,46420,46420,46395],"quality":"More is better","isMain":true,"metricName":"Largest alignment"},{"values":[46365,46514,46506,46653,46704,13363,46492,46420,46420,46420,46395],"quality":"More is better","isMain":true,"metricName":"Total aligned length"},{"values":[46365,46514,46506,46653,46704,null,46492,46420,46420,46420,46395],"quality":"More is better","isMain":false,"metricName":"NG50"},{"values":[46365,46514,46506,46653,46704,null,46492,46420,46420,46420,46395],"quality":"More is better","isMain":false,"metricName":"NG75"},{"values":[46365,46514,46506,46653,46704,611,46492,46420,46420,46420,46395],"quality":"More is better","isMain":false,"metricName":"NA50"},{"values":[46365,46514,46506,46653,46704,559,46492,46420,46420,46420,46395],"quality":"More is better","isMain":false,"metricName":"NA75"},{"values":[46365,46514,46506,46653,46704,null,46492,46420,46420,46420,46395],"quality":"More is better","isMain":true,"metricName":"NGA50"},{"values":[46365,46514,46506,46653,46704,null,46492,46420,46420,46420,46395],"quality":"More is better","isMain":false,"metricName":"NGA75"},{"values":[1,1,1,1,1,null,1,1,1,1,1],"quality":"Less is better","isMain":false,"metricName":"LG50"},{"values":[1,1,1,1,1,null,1,1,1,1,1],"quality":"Less is better","isMain":false,"metricName":"LG75"},{"values":[1,1,1,1,1,9,1,1,1,1,1],"quality":"Less is better","isMain":false,"metricName":"LA50"},{"values":[1,1,1,1,1,15,1,1,1,1,1],"quality":"Less is better","isMain":false,"metricName":"LA75"},{"values":[1,1,1,1,1,null,1,1,1,1,1],"quality":"Less is better","isMain":true,"metricName":"LGA50"},{"values":[1,1,1,1,1,null,1,1,1,1,1],"quality":"Less is better","isMain":false,"metricName":"LGA75"}]],["Misassemblies",[{"values":[0,0,0,0,0,0,0,0,0,0,0],"quality":"Less is better","isMain":true,"metricName":"# misassemblies"},{"values":[0,0,0,0,0,0,0,0,0,0,0],"quality":"Less is better","isMain":false,"metricName":" # relocations"},{"values":[0,0,0,0,0,0,0,0,0,0,0],"quality":"Less is better","isMain":false,"metricName":" # translocations"},{"values":[0,0,0,0,0,0,0,0,0,0,0],"quality":"Less is better","isMain":false,"metricName":" # inversions"},{"values":[0,0,0,0,0,0,0,0,0,0,0],"quality":"Less is better","isMain":false,"metricName":"# misassembled contigs"},{"values":[0,0,0,0,0,0,0,0,0,0,0],"quality":"Less is better","isMain":true,"metricName":"Misassembled contigs length"},{"values":[0,1,1,1,1,0,1,0,0,0,0],"quality":"Less is better","isMain":false,"metricName":"# local misassemblies"},{"values":[0,0,0,0,0,0,0,0,0,0,0],"quality":"Less is better","isMain":false,"metricName":"# unaligned mis. contigs"}]],["Unaligned",[{"values":[0,0,0,0,0,0,0,0,0,0,0],"quality":"Less is better","isMain":false,"metricName":"# fully unaligned contigs"},{"values":[0,0,0,0,0,0,0,0,0,0,0],"quality":"Less is better","isMain":false,"metricName":"Fully unaligned length"},{"values":[0,0,0,0,0,0,0,0,0,0,0],"quality":"Less is better","isMain":false,"metricName":"# partially unaligned contigs"},{"values":[0,0,0,0,0,0,0,0,0,0,0],"quality":"Less is better","isMain":false,"metricName":"Partially unaligned length"}]],["Mismatches",[{"values":[0,1,0,0,0,2,0,0,0,0,0],"quality":"Less is better","isMain":false,"metricName":"# mismatches"},{"values":[0,0,0,0,0,0,0,1,1,1,1],"quality":"Less is better","isMain":false,"metricName":"# indels"},{"values":[0,0,0,0,0,0,0,55,55,55,30],"quality":"Less is better","isMain":false,"metricName":"Indels length"},{"values":["0.00","2.16","0.00","0.00","0.00","14.97","0.00","0.00","0.00","0.00","0.00"],"quality":"Less is better","isMain":true,"metricName":"# mismatches per 100 kbp"},{"values":["0.00","0.00","0.00","0.00","0.00","0.00","0.00","2.16","2.16","2.16","2.16"],"quality":"Less is better","isMain":true,"metricName":"# indels per 100 kbp"},{"values":[0,0,0,0,0,0,0,0,0,0,0],"quality":"Less is better","isMain":false,"metricName":" # indels (<= 5 bp)"},{"values":[0,0,0,0,0,0,0,1,1,1,1],"quality":"Less is better","isMain":false,"metricName":" # indels (> 5 bp)"},{"values":[0,0,0,0,0,0,0,0,0,0,0],"quality":"Less is better","isMain":false,"metricName":"# N's"},{"values":["0.00","0.00","0.00","0.00","0.00","0.00","0.00","0.00","0.00","0.00","0.00"],"quality":"Less is better","isMain":true,"metricName":"# N's per 100 kbp"}]],["Statistics without reference",[{"values":[1,1,1,1,1,21,1,1,1,1,1],"quality":"Equal","isMain":true,"metricName":"# contigs"},{"values":[1,1,1,1,1,1,1,1,1,1,1],"quality":"Equal","isMain":false,"metricName":"# contigs (>= 1000 bp)"},{"values":[1,1,1,1,1,0,1,1,1,1,1],"quality":"Equal","isMain":false,"metricName":"# contigs (>= 5000 bp)"},{"values":[1,1,1,1,1,0,1,1,1,1,1],"quality":"Equal","isMain":false,"metricName":"# contigs (>= 10000 bp)"},{"values":[1,1,1,1,1,0,1,1,1,1,1],"quality":"Equal","isMain":false,"metricName":"# contigs (>= 25000 bp)"},{"values":[0,0,0,0,0,0,0,0,0,0,0],"quality":"Equal","isMain":false,"metricName":"# contigs (>= 50000 bp)"},{"values":[46365,46514,46506,46653,46704,1196,46492,46420,46420,46420,46395],"quality":"More is better","isMain":true,"metricName":"Largest contig"},{"values":[46365,46514,46506,46653,46704,13363,46492,46420,46420,46420,46395],"quality":"More is better","isMain":true,"metricName":"Total length"},{"values":[46365,46514,46506,46653,46704,1196,46492,46420,46420,46420,46395],"quality":"More is better","isMain":true,"metricName":"Total length (>= 1000 bp)"},{"values":[46365,46514,46506,46653,46704,0,46492,46420,46420,46420,46395],"quality":"More is better","isMain":false,"metricName":"Total length (>= 5000 bp)"},{"values":[46365,46514,46506,46653,46704,0,46492,46420,46420,46420,46395],"quality":"More is better","isMain":true,"metricName":"Total length (>= 10000 bp)"},{"values":[46365,46514,46506,46653,46704,0,46492,46420,46420,46420,46395],"quality":"More is better","isMain":false,"metricName":"Total length (>= 25000 bp)"},{"values":[0,0,0,0,0,0,0,0,0,0,0],"quality":"More is better","isMain":true,"metricName":"Total length (>= 50000 bp)"},{"values":[46365,46514,46506,46653,46704,611,46492,46420,46420,46420,46395],"quality":"More is better","isMain":false,"metricName":"N50"},{"values":[46365,46514,46506,46653,46704,559,46492,46420,46420,46420,46395],"quality":"More is better","isMain":false,"metricName":"N75"},{"values":[1,1,1,1,1,9,1,1,1,1,1],"quality":"Less is better","isMain":false,"metricName":"L50"},{"values":[1,1,1,1,1,15,1,1,1,1,1],"quality":"Less is better","isMain":false,"metricName":"L75"},{"values":["68.31","68.29","68.30","68.33","68.34","66.78","68.33","68.32","68.32","68.32","68.32"],"quality":"Equal","isMain":false,"metricName":"GC (%)"}]],["Predicted genes",[]],["Similarity statistics",[{"values":[0,0,0,0,0,0,0,0,0,0,0],"quality":"Equal","isMain":false,"metricName":"# similar correct contigs"},{"values":[0,0,0,0,0,0,0,0,0,0,0],"quality":"Equal","isMain":false,"metricName":"# similar misassembled blocks"}]],["Reference statistics",[{"values":[46365,46365,46365,46365,46365,46365,46365,46365,46365,46365,46365],"quality":"Equal","isMain":false,"metricName":"Reference length"},{"values":[1,1,1,1,1,1,1,1,1,1,1],"quality":"Equal","isMain":false,"metricName":"Reference fragments"},{"values":["68.31","68.31","68.31","68.31","68.31","68.31","68.31","68.31","68.31","68.31","68.31"],"quality":"Equal","isMain":false,"metricName":"Reference GC (%)"}]]],"referenceName":"gi\_149882781\_ref\_NC","date":"18 July 2018, Wednesday, 16:21:41","order":[0,1,2,3,4,5,6,7,8,9,10],"assembliesNames":["CLC","IDBA\_UD","MEGAHIT","MIRA","Ray\_Meta","SOAPdenovo2","SPAdes","SPAdes\_meta","SPAdes\_sc","SPAdes\_sc\_careful","Velvet"]},{"assembliesWithNs":null,"minContig":500,"report":[["Genome statistics",[{"values":["82.734","66.392","85.153","83.120","79.444","99.968","99.567","86.115","89.842","63.446","70.324","24.997","89.547"],"quality":"More is better","isMain":true,"metricName":"Genome fraction (%)"},{"values":["1.092","1.047","1.027","1.021","1.094","1.172","1.076","1.047","1.014","1.011","1.013","1.003","1.000"],"quality":"Less is better","isMain":true,"metricName":"Duplication ratio"},{"values":[10233,4558,10338,3115,4609,42429,14676,12065,19963,4676,4913,2104,42427],"quality":"More is better","isMain":true,"metricName":"Largest alignment"},{"values":[42749,32939,41423,40216,41181,55526,50761,42708,42734,30390,33759,11874,42427],"quality":"More is better","isMain":true,"metricName":"Total aligned length"},{"values":[5252,1628,1970,1380,1572,45108,13044,3110,12873,1301,1399,null,45106],"quality":"More is better","isMain":false,"metricName":"NG50"},{"values":[873,null,817,672,886,45108,9283,831,2814,null,null,null,45106],"quality":"More is better","isMain":false,"metricName":"NG75"},{"values":[5252,2077,2859,1837,1572,42429,13044,3110,12873,3403,2840,1151,42427],"quality":"More is better","isMain":false,"metricName":"NA50"},{"values":[1389,1464,1348,992,1291,5028,6656,1509,12873,1348,1348,668,42427],"quality":"More is better","isMain":false,"metricName":"NA75"},{"values":[4743,1628,1970,1380,1523,42429,13044,3110,12873,1301,1399,null,42427],"quality":"More is better","isMain":true,"metricName":"NGA50"},{"values":[856,null,817,672,886,42429,9283,831,2814,null,null,null,42427],"quality":"More is better","isMain":false,"metricName":"NGA75"},{"values":[3,9,6,12,9,1,2,4,2,9,8,null,1],"quality":"Less is better","isMain":false,"metricName":"LG50"},{"values":[7,null,14,24,19,1,3,11,3,null,null,null,1],"quality":"Less is better","isMain":false,"metricName":"LG75"},{"values":[3,6,4,9,8,1,2,4,2,4,5,4,1],"quality":"Less is better","isMain":false,"metricName":"LA50"},{"values":[6,10,10,17,15,2,4,8,2,8,9,8,1],"quality":"Less is better","isMain":false,"metricName":"LA75"},{"values":[4,9,6,12,10,1,2,4,2,9,8,null,1],"quality":"Less is better","isMain":true,"metricName":"LGA50"},{"values":[9,null,14,24,20,1,3,11,3,null,null,null,1],"quality":"Less is better","isMain":false,"metricName":"LGA75"}]],["Misassemblies",[{"values":[1,0,0,0,1,0,0,0,0,0,0,0,0],"quality":"Less is better","isMain":true,"metricName":"# misassemblies"},{"values":[1,0,0,0,1,0,0,0,0,0,0,0,0],"quality":"Less is better","isMain":false,"metricName":" # relocations"},{"values":[0,0,0,0,0,0,0,0,0,0,0,0,0],"quality":"Less is better","isMain":false,"metricName":" # translocations"},{"values":[0,0,0,0,0,0,0,0,0,0,0,0,0],"quality":"Less is better","isMain":false,"metricName":" # inversions"},{"values":[1,0,0,0,1,0,0,0,0,0,0,0,0],"quality":"Less is better","isMain":false,"metricName":"# misassembled contigs"},{"values":[15026,0,0,0,2638,0,0,0,0,0,0,0,0],"quality":"Less is better","isMain":true,"metricName":"Misassembled contigs length"},{"values":[1,0,0,0,0,0,0,0,0,0,0,0,0],"quality":"Less is better","isMain":false,"metricName":"# local misassemblies"},{"values":[0,0,0,0,0,0,0,0,0,0,0,0,0],"quality":"Less is better","isMain":false,"metricName":"# unaligned mis. contigs"}]],["Unaligned",[{"values":[0,0,0,0,0,0,0,0,0,0,0,0,0],"quality":"Less is better","isMain":false,"metricName":"# fully unaligned contigs"},{"values":[0,0,0,0,0,0,0,0,0,0,0,0,0],"quality":"Less is better","isMain":false,"metricName":"Fully unaligned length"},{"values":[1,0,0,0,0,2,0,0,0,0,0,0,1],"quality":"Less is better","isMain":false,"metricName":"# partially unaligned contigs"},{"values":[598,0,0,0,0,3184,0,0,0,0,0,0,2679],"quality":"Less is better","isMain":false,"metricName":"Partially unaligned length"}]],["Mismatches",[{"values":[70,1,15,0,26,45,56,3,58,39,0,5,27],"quality":"Less is better","isMain":false,"metricName":"# mismatches"},{"values":[2,1,0,0,0,2,0,0,0,0,0,0,1],"quality":"Less is better","isMain":false,"metricName":"# indels"},{"values":[52,1,0,0,0,2,0,0,0,0,0,0,2],"quality":"Less is better","isMain":false,"metricName":"Indels length"},{"values":["178.57","3.18","37.18","0.00","69.07","95.00","118.70","7.35","136.25","129.73","0.00","42.22","63.64"],"quality":"Less is better","isMain":true,"metricName":"# mismatches per 100 kbp"},{"values":["5.10","3.18","0.00","0.00","0.00","4.22","0.00","0.00","0.00","0.00","0.00","0.00","2.36"],"quality":"Less is better","isMain":true,"metricName":"# indels per 100 kbp"},{"values":[1,1,0,0,0,2,0,0,0,0,0,0,1],"quality":"Less is better","isMain":false,"metricName":" # indels (<= 5 bp)"},{"values":[1,0,0,0,0,0,0,0,0,0,0,0,0],"quality":"Less is better","isMain":false,"metricName":" # indels (> 5 bp)"},{"values":[100,81,0,0,0,2,0,0,428,0,0,0,36],"quality":"Less is better","isMain":false,"metricName":"# N's"},{"values":["230.43","245.91","0.00","0.00","0.00","3.41","0.00","0.00","991.61","0.00","0.00","0.00","79.81"],"quality":"Less is better","isMain":true,"metricName":"# N's per 100 kbp"}]],["Statistics without reference",[{"values":[18,22,22,32,27,12,6,21,7,17,20,12,1],"quality":"Equal","isMain":true,"metricName":"# contigs"},{"values":[6,10,13,16,16,4,6,10,7,10,9,5,1],"quality":"Equal","isMain":false,"metricName":"# contigs (>= 1000 bp)"},{"values":[3,0,2,0,0,2,5,1,2,0,0,0,1],"quality":"Equal","isMain":false,"metricName":"# contigs (>= 5000 bp)"},{"values":[1,0,1,0,0,1,2,1,2,0,0,0,1],"quality":"Equal","isMain":false,"metricName":"# contigs (>= 10000 bp)"},{"values":[0,0,0,0,0,1,0,0,0,0,0,0,1],"quality":"Equal","isMain":false,"metricName":"# contigs (>= 25000 bp)"},{"values":[0,0,0,0,0,0,0,0,0,0,0,0,0],"quality":"Equal","isMain":false,"metricName":"# contigs (>= 50000 bp)"},{"values":[15026,4558,10338,3115,4609,45108,14676,12065,20391,4676,4913,2104,45106],"quality":"More is better","isMain":true,"metricName":"Largest contig"},{"values":[43397,32939,41423,40216,41182,58710,50761,42708,43162,30392,33761,11874,45106],"quality":"More is better","isMain":true,"metricName":"Total length"},{"values":[35182,25224,35323,29305,33164,53306,50761,35533,43162,25317,25598,7250,45106],"quality":"More is better","isMain":true,"metricName":"Total length (>= 1000 bp)"},{"values":[28450,0,15457,0,0,50641,49700,12065,33264,0,0,0,45106],"quality":"More is better","isMain":false,"metricName":"Total length (>= 5000 bp)"},{"values":[15026,0,10338,0,0,45108,27720,12065,33264,0,0,0,45106],"quality":"More is better","isMain":true,"metricName":"Total length (>= 10000 bp)"},{"values":[0,0,0,0,0,45108,0,0,0,0,0,0,45106],"quality":"More is better","isMain":false,"metricName":"Total length (>= 25000 bp)"},{"values":[0,0,0,0,0,0,0,0,0,0,0,0,0],"quality":"More is better","isMain":true,"metricName":"Total length (>= 50000 bp)"},{"values":[8172,2077,2859,1837,1999,45108,13044,3110,12873,3403,2840,1151,45106],"quality":"More is better","isMain":false,"metricName":"N50"},{"values":[4287,1464,1348,992,1291,45108,6656,1509,12873,1348,1348,668,45106],"quality":"More is better","isMain":false,"metricName":"N75"},{"values":[2,6,4,9,8,1,2,4,2,4,5,4,1],"quality":"Less is better","isMain":false,"metricName":"L50"},{"values":[4,10,10,17,14,1,4,8,2,8,9,8,1],"quality":"Less is better","isMain":false,"metricName":"L75"},{"values":["63.32","63.40","63.70","63.50","63.58","63.75","63.71","63.44","63.64","63.19","63.40","62.78","63.72"],"quality":"Equal","isMain":false,"metricName":"GC (%)"}]],["Predicted genes",[]],["Similarity statistics",[{"values":[0,0,0,0,0,0,0,0,0,0,0,0,0],"quality":"Equal","isMain":false,"metricName":"# similar correct contigs"},{"values":[0,0,0,0,0,0,0,0,0,0,0,0,0],"quality":"Equal","isMain":false,"metricName":"# similar misassembled blocks"}]],["Reference statistics",[{"values":[47382,47382,47382,47382,47382,47382,47382,47382,47382,47382,47382,47382,47382],"quality":"Equal","isMain":false,"metricName":"Reference length"},{"values":[1,1,1,1,1,1,1,1,1,1,1,1,1],"quality":"Equal","isMain":false,"metricName":"Reference fragments"},{"values":["63.64","63.64","63.64","63.64","63.64","63.64","63.64","63.64","63.64","63.64","63.64","63.64","63.64"],"quality":"Equal","isMain":false,"metricName":"Reference GC (%)"}]]],"referenceName":"gi\_149882900\_ref\_NC","date":"18 July 2018, Wednesday, 16:21:52","order":[0,1,2,3,4,5,6,7,8,9,10,11,12],"assembliesNames":["ABySS\_127","ABySS\_63","CLC","IDBA\_UD","MEGAHIT","MIRA","Ray\_Meta","SPAdes","SPAdes\_meta","SPAdes\_sc","SPAdes\_sc\_careful","Velvet","Geneious"]},{"assembliesWithNs":null,"minContig":500,"report":[["Genome statistics",[{"values":["93.873","89.089","99.132","99.419","99.465","98.703","1.055","90.056","99.266","99.562","98.135","97.890","97.076","74.310"],"quality":"More is better","isMain":true,"metricName":"Genome fraction (%)"},{"values":["1.003","1.005","1.005","1.001","1.000","1.026","1.001","1.026","1.000","1.000","1.003","1.003","1.002","1.000"],"quality":"Less is better","isMain":true,"metricName":"Duplication ratio"},{"values":[10481,9840,89991,84739,109942,42813,601,9390,110154,84944,58075,58075,19395,42813],"quality":"More is better","isMain":true,"metricName":"Largest alignment"},{"values":[200043,190189,211725,211478,211365,215169,2244,192786,210949,211613,208911,208414,206704,157865],"quality":"More is better","isMain":true,"metricName":"Total aligned length"},{"values":[2852,1997,37877,27087,109942,17819,null,5045,110154,55510,53358,53358,7137,17819],"quality":"More is better","isMain":false,"metricName":"NG50"},{"values":[1471,1150,17940,17254,23897,10670,null,2729,58779,25244,16568,16592,3822,null],"quality":"More is better","isMain":false,"metricName":"NG75"},{"values":[2898,2305,37877,27087,109942,17819,589,4550,110154,55510,53358,53358,7139,38517],"quality":"More is better","isMain":false,"metricName":"NA50"},{"values":[1814,1398,17940,17254,48748,10670,533,2604,58779,25244,16568,16592,4631,17819],"quality":"More is better","isMain":false,"metricName":"NA75"},{"values":[2852,1997,37877,27087,109942,17819,null,4286,110154,55510,53358,53358,7137,17819],"quality":"More is better","isMain":true,"metricName":"NGA50"},{"values":[1471,1150,17940,17254,23897,10670,null,2258,58779,25244,16568,16592,3822,null],"quality":"More is better","isMain":false,"metricName":"NGA75"},{"values":[24,30,2,2,1,4,null,15,1,2,2,2,10,4],"quality":"Less is better","isMain":false,"metricName":"LG50"},{"values":[49,64,4,5,3,8,null,30,2,3,4,4,19,null],"quality":"Less is better","isMain":false,"metricName":"LG75"},{"values":[22,25,2,2,1,4,2,16,1,2,2,2,9,2],"quality":"Less is better","isMain":false,"metricName":"LA50"},{"values":[43,52,4,5,2,8,3,31,2,3,4,4,18,4],"quality":"Less is better","isMain":false,"metricName":"LA75"},{"values":[24,30,2,2,1,4,null,17,1,2,2,2,10,4],"quality":"Less is better","isMain":true,"metricName":"LGA50"},{"values":[49,64,4,5,3,8,null,33,2,3,4,4,19,null],"quality":"Less is better","isMain":false,"metricName":"LGA75"}]],["Misassemblies",[{"values":[0,0,0,0,0,1,0,0,1,0,0,0,0,0],"quality":"Less is better","isMain":true,"metricName":"# misassemblies"},{"values":[0,0,0,0,0,1,0,0,1,0,0,0,0,0],"quality":"Less is better","isMain":false,"metricName":" # relocations"},{"values":[0,0,0,0,0,0,0,0,0,0,0,0,0,0],"quality":"Less is better","isMain":false,"metricName":" # translocations"},{"values":[0,0,0,0,0,0,0,0,0,0,0,0,0,0],"quality":"Less is better","isMain":false,"metricName":" # inversions"},{"values":[0,0,0,0,0,1,0,0,1,0,0,0,0,0],"quality":"Less is better","isMain":false,"metricName":"# misassembled contigs"},{"values":[0,0,0,0,0,3952,0,0,42016,0,0,0,0,0],"quality":"Less is better","isMain":true,"metricName":"Misassembled contigs length"},{"values":[0,0,0,0,0,0,0,0,1,0,3,3,0,1],"quality":"Less is better","isMain":false,"metricName":"# local misassemblies"},{"values":[0,0,0,0,0,0,0,0,0,0,0,0,0,0],"quality":"Less is better","isMain":false,"metricName":"# unaligned mis. contigs"}]],["Unaligned",[{"values":[0,0,0,0,0,0,0,0,0,0,0,0,0,0],"quality":"Less is better","isMain":false,"metricName":"# fully unaligned contigs"},{"values":[0,0,0,0,0,0,0,0,0,0,0,0,0,0],"quality":"Less is better","isMain":false,"metricName":"Fully unaligned length"},{"values":[0,0,0,0,0,0,0,5,0,0,0,0,0,0],"quality":"Less is better","isMain":false,"metricName":"# partially unaligned contigs"},{"values":[0,0,0,0,0,0,0,7406,0,0,0,0,0,0],"quality":"Less is better","isMain":false,"metricName":"Partially unaligned length"}]],["Mismatches",[{"values":[25,3,29,5,10,8,0,12,34,27,8,11,10,33],"quality":"Less is better","isMain":false,"metricName":"# mismatches"},{"values":[0,0,0,0,0,2,0,49,2,3,4,2,0,3],"quality":"Less is better","isMain":false,"metricName":"# indels"},{"values":[0,0,0,0,0,2,0,2403,2,26,102,75,0,9],"quality":"Less is better","isMain":false,"metricName":"Indels length"},{"values":["12.53","1.58","13.77","2.37","4.73","3.81","0.00","6.27","16.12","12.76","3.84","5.29","4.85","20.90"],"quality":"Less is better","isMain":true,"metricName":"# mismatches per 100 kbp"},{"values":["0.00","0.00","0.00","0.00","0.00","0.95","0.00","25.61","0.95","1.42","1.92","0.96","0.00","1.90"],"quality":"Less is better","isMain":true,"metricName":"# indels per 100 kbp"},{"values":[0,0,0,0,0,2,0,0,2,1,1,0,0,3],"quality":"Less is better","isMain":false,"metricName":" # indels (<= 5 bp)"},{"values":[0,0,0,0,0,0,0,49,0,2,3,2,0,0],"quality":"Less is better","isMain":false,"metricName":" # indels (> 5 bp)"},{"values":[0,0,0,0,0,52,0,5742,0,0,219,218,0,71],"quality":"Less is better","isMain":false,"metricName":"# N's"},{"values":["0.00","0.00","0.00","0.00","0.00","24.17","0.00","2818.20","0.00","0.00","104.69","104.46","0.00","44.98"],"quality":"Less is better","isMain":true,"metricName":"# N's per 100 kbp"}]],["Statistics without reference",[{"values":[93,102,20,13,7,27,4,57,3,8,11,10,40,7],"quality":"Equal","isMain":true,"metricName":"# contigs"},{"values":[64,72,17,12,6,19,0,50,3,7,9,9,35,7],"quality":"Equal","isMain":false,"metricName":"# contigs (>= 1000 bp)"},{"values":[7,5,6,9,5,12,0,15,3,5,7,7,17,6],"quality":"Equal","isMain":false,"metricName":"# contigs (>= 5000 bp)"},{"values":[1,0,5,6,4,8,0,1,3,5,6,6,5,6],"quality":"Equal","isMain":false,"metricName":"# contigs (>= 10000 bp)"},{"values":[0,0,2,2,2,2,0,0,3,3,3,3,0,2],"quality":"Equal","isMain":false,"metricName":"# contigs (>= 25000 bp)"},{"values":[0,0,1,1,1,0,0,0,2,2,2,2,0,0],"quality":"Equal","isMain":false,"metricName":"# contigs (>= 50000 bp)"},{"values":[10481,9840,89991,84768,109942,42813,601,10425,110154,84944,58128,58128,19395,42813],"quality":"More is better","isMain":true,"metricName":"Largest contig"},{"values":[200043,190189,211725,211507,211365,215169,2244,203747,210949,211613,209183,208685,206704,157865],"quality":"More is better","isMain":true,"metricName":"Total length"},{"values":[178905,168007,209938,210750,210528,210000,0,198790,210949,211024,207835,207859,203414,157865],"quality":"More is better","isMain":true,"metricName":"Total length (>= 1000 bp)"},{"values":[47756,31342,186955,202073,207141,187588,0,106934,210949,206374,203849,203873,152314,154087],"quality":"More is better","isMain":false,"metricName":"Total length (>= 5000 bp)"},{"values":[10481,0,179030,183641,200393,161654,0,10425,210949,206374,195002,195026,71545,154087],"quality":"More is better","isMain":true,"metricName":"Total length (>= 10000 bp)"},{"values":[0,0,127868,111855,158690,70812,0,0,210949,165698,149120,149120,0,81330],"quality":"More is better","isMain":false,"metricName":"Total length (>= 25000 bp)"},{"values":[0,0,89991,84768,109942,0,0,0,168933,140454,111486,111486,0,0],"quality":"More is better","isMain":true,"metricName":"Total length (>= 50000 bp)"},{"values":[2898,2305,37877,27087,109942,17819,589,5404,110154,55510,53358,53358,7139,38517],"quality":"More is better","isMain":false,"metricName":"N50"},{"values":[1814,1398,17940,17254,48748,10670,533,2759,58779,25244,16568,16592,4631,17819],"quality":"More is better","isMain":false,"metricName":"N75"},{"values":[22,25,2,2,1,4,2,14,1,2,2,2,9,2],"quality":"Less is better","isMain":false,"metricName":"L50"},{"values":[43,52,4,5,2,8,3,28,2,3,4,4,18,4],"quality":"Less is better","isMain":false,"metricName":"L75"},{"values":["28.57","28.37","28.56","28.54","28.56","28.68","29.77","28.34","28.54","28.56","28.39","28.37","28.34","28.44"],"quality":"Equal","isMain":false,"metricName":"GC (%)"}]],["Predicted genes",[]],["Similarity statistics",[{"values":[0,0,0,0,0,0,0,0,0,0,0,0,0,0],"quality":"Equal","isMain":false,"metricName":"# similar correct contigs"},{"values":[0,0,0,0,0,0,0,0,0,0,0,0,0,0],"quality":"Equal","isMain":false,"metricName":"# similar misassembled blocks"}]],["Reference statistics",[{"values":[212482,212482,212482,212482,212482,212482,212482,212482,212482,212482,212482,212482,212482,212482],"quality":"Equal","isMain":false,"metricName":"Reference length"},{"values":[1,1,1,1,1,1,1,1,1,1,1,1,1,1],"quality":"Equal","isMain":false,"metricName":"Reference fragments"},{"values":["28.63","28.63","28.63","28.63","28.63","28.63","28.63","28.63","28.63","28.63","28.63","28.63","28.63","28.63"],"quality":"Equal","isMain":false,"metricName":"Reference GC (%)"}]]],"referenceName":"gi\_15078713\_ref\_NC","date":"18 July 2018, Wednesday, 16:22:06","order":[0,1,2,3,4,5,6,7,8,9,10,11,12,13],"assembliesNames":["ABySS\_127","ABySS\_63","CLC","IDBA\_UD","MEGAHIT","MIRA","Ray\_Meta","SOAPdenovo2","SPAdes","SPAdes\_meta","SPAdes\_sc","SPAdes\_sc\_careful","Velvet","Geneious"]},{"assembliesWithNs":null,"minContig":500,"report":[["Genome statistics",[{"values":["100.000","86.390","2.924","100.000","100.000","100.000","100.000","100.000","99.977","100.000"],"quality":"More is better","isMain":true,"metricName":"Genome fraction (%)"},{"values":["1.017","1.000","1.000","248.955","1.003","1.007","1.003","1.003","1.000","4.272"],"quality":"Less is better","isMain":true,"metricName":"Duplication ratio"},{"values":[17171,3134,502,17171,17220,17298,17171,17171,17167,17171],"quality":"More is better","isMain":true,"metricName":"Largest alignment"},{"values":[17171,14834,502,4273227,17220,17298,17171,17171,17167,73354],"quality":"More is better","isMain":true,"metricName":"Total aligned length"},{"values":[17463,2194,null,38321,17220,17298,17225,17225,17169,73354],"quality":"More is better","isMain":false,"metricName":"NG50"},{"values":[17463,894,null,38321,17220,17298,17225,17225,17169,73354],"quality":"More is better","isMain":false,"metricName":"NG75"},{"values":[17171,2361,502,3480,17220,17298,17171,17171,17167,17171],"quality":"More is better","isMain":false,"metricName":"NA50"},{"values":[17171,2057,502,2347,17220,17298,17171,17171,17167,17169],"quality":"More is better","isMain":false,"metricName":"NA75"},{"values":[17171,2194,null,17171,17220,17298,17171,17171,17167,17171],"quality":"More is better","isMain":true,"metricName":"NGA50"},{"values":[17171,894,null,17171,17220,17298,17171,17171,17167,17171],"quality":"More is better","isMain":false,"metricName":"NGA75"},{"values":[1,4,null,1,1,1,1,1,1,1],"quality":"Less is better","isMain":false,"metricName":"LG50"},{"values":[1,6,null,1,1,1,1,1,1,1],"quality":"Less is better","isMain":false,"metricName":"LG75"},{"values":[1,3,1,347,1,1,1,1,1,3],"quality":"Less is better","isMain":false,"metricName":"LA50"},{"values":[1,5,1,723,1,1,1,1,1,4],"quality":"Less is better","isMain":false,"metricName":"LA75"},{"values":[1,4,null,1,1,1,1,1,1,1],"quality":"Less is better","isMain":true,"metricName":"LGA50"},{"values":[1,6,null,1,1,1,1,1,1,1],"quality":"Less is better","isMain":false,"metricName":"LGA75"}]],["Misassemblies",[{"values":[0,0,0,18,0,0,0,0,0,5],"quality":"Less is better","isMain":true,"metricName":"# misassemblies"},{"values":[0,0,0,14,0,0,0,0,0,4],"quality":"Less is better","isMain":false,"metricName":" # relocations"},{"values":[0,0,0,0,0,0,0,0,0,0],"quality":"Less is better","isMain":false,"metricName":" # translocations"},{"values":[0,0,0,4,0,0,0,0,0,1],"quality":"Less is better","isMain":false,"metricName":" # inversions"},{"values":[0,0,0,16,0,0,0,0,0,1],"quality":"Less is better","isMain":false,"metricName":"# misassembled contigs"},{"values":[0,0,0,246455,0,0,0,0,0,73354],"quality":"Less is better","isMain":true,"metricName":"Misassembled contigs length"},{"values":[0,0,0,0,0,1,0,0,0,0],"quality":"Less is better","isMain":false,"metricName":"# local misassemblies"},{"values":[0,0,0,0,0,0,0,0,0,0],"quality":"Less is better","isMain":false,"metricName":"# unaligned mis. contigs"}]],["Unaligned",[{"values":[0,0,0,0,0,0,0,0,0,0],"quality":"Less is better","isMain":false,"metricName":"# fully unaligned contigs"},{"values":[0,0,0,0,0,0,0,0,0,0],"quality":"Less is better","isMain":false,"metricName":"Fully unaligned length"},{"values":[0,0,0,0,0,0,0,0,0,0],"quality":"Less is better","isMain":false,"metricName":"# partially unaligned contigs"},{"values":[0,0,0,0,0,0,0,0,0,0],"quality":"Less is better","isMain":false,"metricName":"Partially unaligned length"}]],["Mismatches",[{"values":[4,0,4,38,0,0,1,0,0,58],"quality":"Less is better","isMain":false,"metricName":"# mismatches"},{"values":[0,0,0,8,1,0,0,0,0,2],"quality":"Less is better","isMain":false,"metricName":"# indels"},{"values":[0,0,0,8,49,0,0,0,0,2],"quality":"Less is better","isMain":false,"metricName":"Indels length"},{"values":["23.30","0.00","796.81","221.30","0.00","0.00","5.82","0.00","0.00","337.78"],"quality":"Less is better","isMain":true,"metricName":"# mismatches per 100 kbp"},{"values":["0.00","0.00","0.00","46.59","5.82","0.00","0.00","0.00","0.00","11.65"],"quality":"Less is better","isMain":true,"metricName":"# indels per 100 kbp"},{"values":[0,0,0,8,0,0,0,0,0,2],"quality":"Less is better","isMain":false,"metricName":" # indels (<= 5 bp)"},{"values":[0,0,0,0,1,0,0,0,0,0],"quality":"Less is better","isMain":false,"metricName":" # indels (> 5 bp)"},{"values":[0,0,0,2135,0,0,0,0,0,59],"quality":"Less is better","isMain":false,"metricName":"# N's"},{"values":["0.00","0.00","0.00","49.94","0.00","0.00","0.00","0.00","0.00","80.43"],"quality":"Less is better","isMain":true,"metricName":"# N's per 100 kbp"}]],["Statistics without reference",[{"values":[1,8,1,1303,1,1,1,1,1,1],"quality":"Equal","isMain":true,"metricName":"# contigs"},{"values":[1,5,0,1289,1,1,1,1,1,1],"quality":"Equal","isMain":false,"metricName":"# contigs (>= 1000 bp)"},{"values":[1,0,0,168,1,1,1,1,1,1],"quality":"Equal","isMain":false,"metricName":"# contigs (>= 5000 bp)"},{"values":[1,0,0,42,1,1,1,1,1,1],"quality":"Equal","isMain":false,"metricName":"# contigs (>= 10000 bp)"},{"values":[0,0,0,4,0,0,0,0,0,1],"quality":"Equal","isMain":false,"metricName":"# contigs (>= 25000 bp)"},{"values":[0,0,0,0,0,0,0,0,0,1],"quality":"Equal","isMain":false,"metricName":"# contigs (>= 50000 bp)"},{"values":[17463,3134,502,38321,17220,17298,17225,17225,17169,73354],"quality":"More is better","isMain":true,"metricName":"Largest contig"},{"values":[17463,14834,502,4274813,17220,17298,17225,17225,17169,73354],"quality":"More is better","isMain":true,"metricName":"Total length"},{"values":[17463,12206,0,4263441,17220,17298,17225,17225,17169,73354],"quality":"More is better","isMain":true,"metricName":"Total length (>= 1000 bp)"},{"values":[17463,0,0,1457274,17220,17298,17225,17225,17169,73354],"quality":"More is better","isMain":false,"metricName":"Total length (>= 5000 bp)"},{"values":[17463,0,0,626632,17220,17298,17225,17225,17169,73354],"quality":"More is better","isMain":true,"metricName":"Total length (>= 10000 bp)"},{"values":[0,0,0,122235,0,0,0,0,0,73354],"quality":"More is better","isMain":false,"metricName":"Total length (>= 25000 bp)"},{"values":[0,0,0,0,0,0,0,0,0,73354],"quality":"More is better","isMain":true,"metricName":"Total length (>= 50000 bp)"},{"values":[17463,2361,502,3501,17220,17298,17225,17225,17169,73354],"quality":"More is better","isMain":false,"metricName":"N50"},{"values":[17463,2057,502,2357,17220,17298,17225,17225,17169,73354],"quality":"More is better","isMain":false,"metricName":"N75"},{"values":[1,3,1,335,1,1,1,1,1,1],"quality":"Less is better","isMain":false,"metricName":"L50"},{"values":[1,5,1,710,1,1,1,1,1,1],"quality":"Less is better","isMain":false,"metricName":"L75"},{"values":["49.43","50.11","49.80","49.54","49.45","49.50","49.53","49.52","49.50","49.66"],"quality":"Equal","isMain":false,"metricName":"GC (%)"}]],["Predicted genes",[]],["Similarity statistics",[{"values":[0,0,0,0,0,0,0,0,0,0],"quality":"Equal","isMain":false,"metricName":"# similar correct contigs"},{"values":[0,0,0,0,0,0,0,0,0,0],"quality":"Equal","isMain":false,"metricName":"# similar misassembled blocks"}]],["Reference statistics",[{"values":[17171,17171,17171,17171,17171,17171,17171,17171,17171,17171],"quality":"Equal","isMain":false,"metricName":"Reference length"},{"values":[1,1,1,1,1,1,1,1,1,1],"quality":"Equal","isMain":false,"metricName":"Reference fragments"},{"values":["49.50","49.50","49.50","49.50","49.50","49.50","49.50","49.50","49.50","49.50"],"quality":"Equal","isMain":false,"metricName":"Reference GC (%)"}]]],"referenceName":"gi\_151266287\_ref\_NC","date":"18 July 2018, Wednesday, 16:22:21","order":[0,1,2,3,4,5,6,7,8,9],"assembliesNames":["CLC","IDBA\_UD","MEGAHIT","MIRA","Ray\_Meta","SPAdes","SPAdes\_meta","SPAdes\_sc","SPAdes\_sc\_careful","Geneious"]},{"assembliesWithNs":null,"minContig":500,"report":[["Genome statistics",[{"values":["100.000","100.000","100.000","100.000","59.809","92.716","100.000","100.000","100.000","100.000","100.000"],"quality":"More is better","isMain":true,"metricName":"Genome fraction (%)"},{"values":["1.000","1.002","1.003","1.007","1.005","1.016","1.003","1.001","1.001","1.001","1.001"],"quality":"Less is better","isMain":true,"metricName":"Duplication ratio"},{"values":[49534,49633,49675,49884,1850,46639,49661,49589,49589,49589,49564],"quality":"More is better","isMain":true,"metricName":"Largest alignment"},{"values":[49534,49633,49675,49884,29783,46639,49661,49589,49589,49589,49564],"quality":"More is better","isMain":true,"metricName":"Total aligned length"},{"values":[49534,49633,49675,49884,639,50969,49661,49589,49589,49589,49564],"quality":"More is better","isMain":false,"metricName":"NG50"},{"values":[49534,49633,49675,49884,null,50969,49661,49589,49589,49589,49564],"quality":"More is better","isMain":false,"metricName":"NG75"},{"values":[49534,49633,49675,49884,931,46639,49661,49589,49589,49589,49564],"quality":"More is better","isMain":false,"metricName":"NA50"},{"values":[49534,49633,49675,49884,686,46639,49661,49589,49589,49589,49564],"quality":"More is better","isMain":false,"metricName":"NA75"},{"values":[49534,49633,49675,49884,639,46639,49661,49589,49589,49589,49564],"quality":"More is better","isMain":true,"metricName":"NGA50"},{"values":[49534,49633,49675,49884,null,46639,49661,49589,49589,49589,49564],"quality":"More is better","isMain":false,"metricName":"NGA75"},{"values":[1,1,1,1,25,1,1,1,1,1,1],"quality":"Less is better","isMain":false,"metricName":"LG50"},{"values":[1,1,1,1,null,1,1,1,1,1,1],"quality":"Less is better","isMain":false,"metricName":"LG75"},{"values":[1,1,1,1,12,1,1,1,1,1,1],"quality":"Less is better","isMain":false,"metricName":"LA50"},{"values":[1,1,1,1,21,1,1,1,1,1,1],"quality":"Less is better","isMain":false,"metricName":"LA75"},{"values":[1,1,1,1,25,1,1,1,1,1,1],"quality":"Less is better","isMain":true,"metricName":"LGA50"},{"values":[1,1,1,1,null,1,1,1,1,1,1],"quality":"Less is better","isMain":false,"metricName":"LGA75"}]],["Misassemblies",[{"values":[0,0,0,0,0,0,0,0,0,0,0],"quality":"Less is better","isMain":true,"metricName":"# misassemblies"},{"values":[0,0,0,0,0,0,0,0,0,0,0],"quality":"Less is better","isMain":false,"metricName":" # relocations"},{"values":[0,0,0,0,0,0,0,0,0,0,0],"quality":"Less is better","isMain":false,"metricName":" # translocations"},{"values":[0,0,0,0,0,0,0,0,0,0,0],"quality":"Less is better","isMain":false,"metricName":" # inversions"},{"values":[0,0,0,0,0,0,0,0,0,0,0],"quality":"Less is better","isMain":false,"metricName":"# misassembled contigs"},{"values":[0,0,0,0,0,0,0,0,0,0,0],"quality":"Less is better","isMain":true,"metricName":"Misassembled contigs length"},{"values":[0,1,1,1,0,1,1,0,0,0,0],"quality":"Less is better","isMain":false,"metricName":"# local misassemblies"},{"values":[0,0,0,0,0,0,0,0,0,0,0],"quality":"Less is better","isMain":false,"metricName":"# unaligned mis. contigs"}]],["Unaligned",[{"values":[0,0,0,0,0,0,0,0,0,0,0],"quality":"Less is better","isMain":false,"metricName":"# fully unaligned contigs"},{"values":[0,0,0,0,0,0,0,0,0,0,0],"quality":"Less is better","isMain":false,"metricName":"Fully unaligned length"},{"values":[0,0,0,0,0,1,0,0,0,0,0],"quality":"Less is better","isMain":false,"metricName":"# partially unaligned contigs"},{"values":[0,0,0,0,0,4330,0,0,0,0,0],"quality":"Less is better","isMain":false,"metricName":"Partially unaligned length"}]],["Mismatches",[{"values":[0,0,0,1,0,0,0,0,0,0,0],"quality":"Less is better","isMain":false,"metricName":"# mismatches"},{"values":[0,0,0,0,0,29,0,1,1,1,1],"quality":"Less is better","isMain":false,"metricName":"# indels"},{"values":[0,0,0,0,0,1379,0,55,55,55,30],"quality":"Less is better","isMain":false,"metricName":"Indels length"},{"values":["0.00","0.00","0.00","2.02","0.00","0.00","0.00","0.00","0.00","0.00","0.00"],"quality":"Less is better","isMain":true,"metricName":"# mismatches per 100 kbp"},{"values":["0.00","0.00","0.00","0.00","0.00","63.15","0.00","2.02","2.02","2.02","2.02"],"quality":"Less is better","isMain":true,"metricName":"# indels per 100 kbp"},{"values":[0,0,0,0,0,0,0,0,0,0,0],"quality":"Less is better","isMain":false,"metricName":" # indels (<= 5 bp)"},{"values":[0,0,0,0,0,29,0,1,1,1,1],"quality":"Less is better","isMain":false,"metricName":" # indels (> 5 bp)"},{"values":[0,0,0,2,0,2882,0,0,0,0,0],"quality":"Less is better","isMain":false,"metricName":"# N's"},{"values":["0.00","0.00","0.00","4.01","0.00","5654.42","0.00","0.00","0.00","0.00","0.00"],"quality":"Less is better","isMain":true,"metricName":"# N's per 100 kbp"}]],["Statistics without reference",[{"values":[1,1,1,1,33,1,1,1,1,1,1],"quality":"Equal","isMain":true,"metricName":"# contigs"},{"values":[1,1,1,1,9,1,1,1,1,1,1],"quality":"Equal","isMain":false,"metricName":"# contigs (>= 1000 bp)"},{"values":[1,1,1,1,0,1,1,1,1,1,1],"quality":"Equal","isMain":false,"metricName":"# contigs (>= 5000 bp)"},{"values":[1,1,1,1,0,1,1,1,1,1,1],"quality":"Equal","isMain":false,"metricName":"# contigs (>= 10000 bp)"},{"values":[1,1,1,1,0,1,1,1,1,1,1],"quality":"Equal","isMain":false,"metricName":"# contigs (>= 25000 bp)"},{"values":[0,0,0,0,0,1,0,0,0,0,0],"quality":"Equal","isMain":false,"metricName":"# contigs (>= 50000 bp)"},{"values":[49534,49633,49675,49884,1850,50969,49661,49589,49589,49589,49564],"quality":"More is better","isMain":true,"metricName":"Largest contig"},{"values":[49534,49633,49675,49884,29783,50969,49661,49589,49589,49589,49564],"quality":"More is better","isMain":true,"metricName":"Total length"},{"values":[49534,49633,49675,49884,12776,50969,49661,49589,49589,49589,49564],"quality":"More is better","isMain":true,"metricName":"Total length (>= 1000 bp)"},{"values":[49534,49633,49675,49884,0,50969,49661,49589,49589,49589,49564],"quality":"More is better","isMain":false,"metricName":"Total length (>= 5000 bp)"},{"values":[49534,49633,49675,49884,0,50969,49661,49589,49589,49589,49564],"quality":"More is better","isMain":true,"metricName":"Total length (>= 10000 bp)"},{"values":[49534,49633,49675,49884,0,50969,49661,49589,49589,49589,49564],"quality":"More is better","isMain":false,"metricName":"Total length (>= 25000 bp)"},{"values":[0,0,0,0,0,50969,0,0,0,0,0],"quality":"More is better","isMain":true,"metricName":"Total length (>= 50000 bp)"},{"values":[49534,49633,49675,49884,931,50969,49661,49589,49589,49589,49564],"quality":"More is better","isMain":false,"metricName":"N50"},{"values":[49534,49633,49675,49884,686,50969,49661,49589,49589,49589,49564],"quality":"More is better","isMain":false,"metricName":"N75"},{"values":[1,1,1,1,12,1,1,1,1,1,1],"quality":"Less is better","isMain":false,"metricName":"L50"},{"values":[1,1,1,1,21,1,1,1,1,1,1],"quality":"Less is better","isMain":false,"metricName":"L75"},{"values":["67.75","67.75","67.73","67.74","67.90","67.78","67.75","67.75","67.75","67.75","67.75"],"quality":"Equal","isMain":false,"metricName":"GC (%)"}]],["Predicted genes",[]],["Similarity statistics",[{"values":[0,0,0,0,0,0,0,0,0,0,0],"quality":"Equal","isMain":false,"metricName":"# similar correct contigs"},{"values":[0,0,0,0,0,0,0,0,0,0,0],"quality":"Equal","isMain":false,"metricName":"# similar misassembled blocks"}]],["Reference statistics",[{"values":[49534,49534,49534,49534,49534,49534,49534,49534,49534,49534,49534],"quality":"Equal","isMain":false,"metricName":"Reference length"},{"values":[1,1,1,1,1,1,1,1,1,1,1],"quality":"Equal","isMain":false,"metricName":"Reference fragments"},{"values":["67.75","67.75","67.75","67.75","67.75","67.75","67.75","67.75","67.75","67.75","67.75"],"quality":"Equal","isMain":false,"metricName":"Reference GC (%)"}]]],"referenceName":"gi\_15320570\_ref\_NC","date":"18 July 2018, Wednesday, 16:22:31","order":[0,1,2,3,4,5,6,7,8,9,10],"assembliesNames":["CLC","IDBA\_UD","MEGAHIT","MIRA","Ray\_Meta","SOAPdenovo2","SPAdes","SPAdes\_meta","SPAdes\_sc","SPAdes\_sc\_careful","Velvet"]},{"assembliesWithNs":null,"minContig":500,"report":[["Genome statistics",[{"values":["12.865","4.738","14.056","1.146","9.880","17.564","17.663","18.195","18.195"],"quality":"More is better","isMain":true,"metricName":"Genome fraction (%)"},{"values":["1.016","1.000","1.012","1.000","1.030","1.044","1.017","1.027","1.027"],"quality":"Less is better","isMain":true,"metricName":"Duplication ratio"},{"values":[1077,916,1111,750,947,1111,1111,1111,1111],"quality":"More is better","isMain":true,"metricName":"Largest alignment"},{"values":[15256,5618,16668,1359,11716,20828,20945,21576,21576],"quality":"More is better","isMain":true,"metricName":"Total aligned length"},{"values":[685,690,677,750,660,717,749,717,717],"quality":"More is better","isMain":false,"metricName":"NA50"},{"values":[591,550,583,609,552,660,677,591,591],"quality":"More is better","isMain":false,"metricName":"NA75"},{"values":[null,null,null,null,null,null,null,null,null],"quality":"More is better","isMain":true,"metricName":"NGA50"},{"values":[10,5,11,1,8,13,12,14,14],"quality":"Less is better","isMain":false,"metricName":"LA50"},{"values":[16,7,18,2,13,20,20,23,23],"quality":"Less is better","isMain":false,"metricName":"LA75"}]],["Misassemblies",[{"values":[0,0,0,0,0,0,0,0,0],"quality":"Less is better","isMain":true,"metricName":"# misassemblies"},{"values":[0,0,0,0,0,0,0,0,0],"quality":"Less is better","isMain":false,"metricName":" # relocations"},{"values":[0,0,0,0,0,0,0,0,0],"quality":"Less is better","isMain":false,"metricName":" # translocations"},{"values":[0,0,0,0,0,0,0,0,0],"quality":"Less is better","isMain":false,"metricName":" # inversions"},{"values":[0,0,0,0,0,0,0,0,0],"quality":"Less is better","isMain":false,"metricName":"# misassembled contigs"},{"values":[0,0,0,0,0,0,0,0,0],"quality":"Less is better","isMain":true,"metricName":"Misassembled contigs length"},{"values":[0,0,0,0,0,1,0,0,0],"quality":"Less is better","isMain":false,"metricName":"# local misassemblies"},{"values":[0,0,0,0,0,0,0,0,0],"quality":"Less is better","isMain":false,"metricName":"# unaligned mis. contigs"}]],["Unaligned",[{"values":[0,0,0,0,0,0,0,0,0],"quality":"Less is better","isMain":false,"metricName":"# fully unaligned contigs"},{"values":[0,0,0,0,0,0,0,0,0],"quality":"Less is better","isMain":false,"metricName":"Fully unaligned length"},{"values":[0,1,0,0,0,0,0,2,2],"quality":"Less is better","isMain":false,"metricName":"# partially unaligned contigs"},{"values":[0,1095,0,0,0,0,0,1667,1667],"quality":"Less is better","isMain":false,"metricName":"Partially unaligned length"}]],["Mismatches",[{"values":[23,3,30,0,25,61,40,62,64],"quality":"Less is better","isMain":false,"metricName":"# mismatches"},{"values":[0,0,0,0,0,0,0,2,2],"quality":"Less is better","isMain":false,"metricName":"# indels"},{"values":[0,0,0,0,0,0,0,2,2],"quality":"Less is better","isMain":false,"metricName":"Indels length"},{"values":["150.76","53.40","179.99","0.00","213.38","292.87","190.98","287.36","296.63"],"quality":"Less is better","isMain":true,"metricName":"# mismatches per 100 kbp"},{"values":["0.00","0.00","0.00","0.00","0.00","0.00","0.00","9.27","9.27"],"quality":"Less is better","isMain":true,"metricName":"# indels per 100 kbp"},{"values":[0,0,0,0,0,0,0,2,2],"quality":"Less is better","isMain":false,"metricName":" # indels (<= 5 bp)"},{"values":[0,0,0,0,0,0,0,0,0],"quality":"Less is better","isMain":false,"metricName":" # indels (> 5 bp)"},{"values":[0,0,0,2,0,0,0,0,0],"quality":"Less is better","isMain":false,"metricName":"# N's"},{"values":["0.00","0.00","0.00","147.17","0.00","0.00","0.00","0.00","0.00"],"quality":"Less is better","isMain":true,"metricName":"# N's per 100 kbp"}]],["Statistics without reference",[{"values":[22,8,25,2,18,28,28,30,30],"quality":"Equal","isMain":true,"metricName":"# contigs"},{"values":[1,1,1,0,1,4,4,6,6],"quality":"Equal","isMain":false,"metricName":"# contigs (>= 1000 bp)"},{"values":[0,0,0,0,0,0,0,0,0],"quality":"Equal","isMain":false,"metricName":"# contigs (>= 5000 bp)"},{"values":[0,0,0,0,0,0,0,0,0],"quality":"Equal","isMain":false,"metricName":"# contigs (>= 10000 bp)"},{"values":[0,0,0,0,0,0,0,0,0],"quality":"Equal","isMain":false,"metricName":"# contigs (>= 25000 bp)"},{"values":[0,0,0,0,0,0,0,0,0],"quality":"Equal","isMain":false,"metricName":"# contigs (>= 50000 bp)"},{"values":[1077,1672,1111,750,1159,1624,1159,1436,1436],"quality":"More is better","isMain":true,"metricName":"Largest contig"},{"values":[15498,6713,16875,1359,12066,21748,21295,23835,23835],"quality":"More is better","isMain":true,"metricName":"Total length"},{"values":[1077,1672,1111,0,1159,4817,4352,6869,6869],"quality":"More is better","isMain":true,"metricName":"Total length (>= 1000 bp)"},{"values":[0,0,0,0,0,0,0,0,0],"quality":"More is better","isMain":false,"metricName":"Total length (>= 5000 bp)"},{"values":[0,0,0,0,0,0,0,0,0],"quality":"More is better","isMain":true,"metricName":"Total length (>= 10000 bp)"},{"values":[0,0,0,0,0,0,0,0,0],"quality":"More is better","isMain":false,"metricName":"Total length (>= 25000 bp)"},{"values":[0,0,0,0,0,0,0,0,0],"quality":"More is better","isMain":true,"metricName":"Total length (>= 50000 bp)"},{"values":[685,829,684,750,660,749,749,803,803],"quality":"More is better","isMain":false,"metricName":"N50"},{"values":[619,690,591,609,552,679,677,679,679],"quality":"More is better","isMain":false,"metricName":"N75"},{"values":[10,3,11,1,8,12,12,12,12],"quality":"Less is better","isMain":false,"metricName":"L50"},{"values":[16,6,18,2,13,19,20,21,21],"quality":"Less is better","isMain":false,"metricName":"L75"},{"values":["40.84","42.71","41.06","39.87","41.92","40.82","40.90","41.04","41.04"],"quality":"Equal","isMain":false,"metricName":"GC (%)"}]],["Predicted genes",[]],["Similarity statistics",[{"values":[0,0,0,0,0,0,0,0,0],"quality":"Equal","isMain":false,"metricName":"# similar correct contigs"},{"values":[0,0,0,0,0,0,0,0,0],"quality":"Equal","isMain":false,"metricName":"# similar misassembled blocks"}]],["Reference statistics",[{"values":[118584,118584,118584,118584,118584,118584,118584,118584,118584],"quality":"Equal","isMain":false,"metricName":"Reference length"},{"values":[1,1,1,1,1,1,1,1,1],"quality":"Equal","isMain":false,"metricName":"Reference fragments"},{"values":["40.68","40.68","40.68","40.68","40.68","40.68","40.68","40.68","40.68"],"quality":"Equal","isMain":false,"metricName":"Reference GC (%)"}]]],"referenceName":"gi\_15320657\_ref\_NC","date":"18 July 2018, Wednesday, 16:22:40","order":[0,1,2,3,4,5,6,7,8],"assembliesNames":["CLC","IDBA\_UD","MEGAHIT","MIRA","SOAPdenovo2","SPAdes","SPAdes\_meta","SPAdes\_sc","SPAdes\_sc\_careful"]},{"assembliesWithNs":null,"minContig":500,"report":[["Genome statistics",[{"values":["0.472","7.502","5.962","13.598","0.545","7.312","16.242","15.633","16.388","16.388","0.423"],"quality":"More is better","isMain":true,"metricName":"Genome fraction (%)"},{"values":["1.000","1.054","1.114","1.000","1.000","1.000","1.020","1.002","1.038","1.038","1.000"],"quality":"Less is better","isMain":true,"metricName":"Duplication ratio"},{"values":[598,1022,873,975,690,1017,1273,1098,1022,1022,536],"quality":"More is better","isMain":true,"metricName":"Largest alignment"},{"values":[598,9500,7551,17219,690,9259,20658,19827,20752,20752,536],"quality":"More is better","isMain":true,"metricName":"Total aligned length"},{"values":[598,723,601,678,690,661,780,780,723,723,536],"quality":"More is better","isMain":false,"metricName":"NA50"},{"values":[598,570,464,569,690,604,617,636,617,617,536],"quality":"More is better","isMain":false,"metricName":"NA75"},{"values":[null,null,null,null,null,null,null,null,null,null,null],"quality":"More is better","isMain":true,"metricName":"NGA50"},{"values":[1,7,7,11,1,6,13,12,14,14,1],"quality":"Less is better","isMain":false,"metricName":"LA50"},{"values":[1,11,11,19,1,10,20,19,22,22,1],"quality":"Less is better","isMain":false,"metricName":"LA75"}]],["Misassemblies",[{"values":[0,0,0,0,0,0,0,0,0,0,0],"quality":"Less is better","isMain":true,"metricName":"# misassemblies"},{"values":[0,0,0,0,0,0,0,0,0,0,0],"quality":"Less is better","isMain":false,"metricName":" # relocations"},{"values":[0,0,0,0,0,0,0,0,0,0,0],"quality":"Less is better","isMain":false,"metricName":" # translocations"},{"values":[0,0,0,0,0,0,0,0,0,0,0],"quality":"Less is better","isMain":false,"metricName":" # inversions"},{"values":[0,0,0,0,0,0,0,0,0,0,0],"quality":"Less is better","isMain":false,"metricName":"# misassembled contigs"},{"values":[0,0,0,0,0,0,0,0,0,0,0],"quality":"Less is better","isMain":true,"metricName":"Misassembled contigs length"},{"values":[0,0,0,0,0,0,1,0,0,0,0],"quality":"Less is better","isMain":false,"metricName":"# local misassemblies"},{"values":[0,0,0,0,0,0,0,0,0,0,0],"quality":"Less is better","isMain":false,"metricName":"# unaligned mis. contigs"}]],["Unaligned",[{"values":[0,0,0,0,0,0,0,0,0,0,0],"quality":"Less is better","isMain":false,"metricName":"# fully unaligned contigs"},{"values":[0,0,0,0,0,0,0,0,0,0,0],"quality":"Less is better","isMain":false,"metricName":"Fully unaligned length"},{"values":[0,1,1,0,0,0,1,0,1,1,0],"quality":"Less is better","isMain":false,"metricName":"# partially unaligned contigs"},{"values":[0,634,877,0,0,0,790,0,790,790,0],"quality":"Less is better","isMain":false,"metricName":"Partially unaligned length"}]],["Mismatches",[{"values":[0,42,38,52,0,42,58,36,86,72,0],"quality":"Less is better","isMain":false,"metricName":"# mismatches"},{"values":[0,2,1,2,0,0,0,0,0,0,0],"quality":"Less is better","isMain":false,"metricName":"# indels"},{"values":[0,2,1,2,0,0,0,0,0,0,0],"quality":"Less is better","isMain":false,"metricName":"Indels length"},{"values":["0.00","442.11","503.31","301.99","0.00","453.61","282.01","181.85","414.42","346.95","0.00"],"quality":"Less is better","isMain":true,"metricName":"# mismatches per 100 kbp"},{"values":["0.00","21.05","13.25","11.62","0.00","0.00","0.00","0.00","0.00","0.00","0.00"],"quality":"Less is better","isMain":true,"metricName":"# indels per 100 kbp"},{"values":[0,2,1,2,0,0,0,0,0,0,0],"quality":"Less is better","isMain":false,"metricName":" # indels (<= 5 bp)"},{"values":[0,0,0,0,0,0,0,0,0,0,0],"quality":"Less is better","isMain":false,"metricName":" # indels (> 5 bp)"},{"values":[0,0,0,0,0,0,0,0,0,0,0],"quality":"Less is better","isMain":false,"metricName":"# N's"},{"values":["0.00","0.00","0.00","0.00","0.00","0.00","0.00","0.00","0.00","0.00","0.00"],"quality":"Less is better","isMain":true,"metricName":"# N's per 100 kbp"}]],["Statistics without reference",[{"values":[1,14,13,26,1,14,28,27,29,29,1],"quality":"Equal","isMain":true,"metricName":"# contigs"},{"values":[0,2,1,0,0,1,3,2,3,3,0],"quality":"Equal","isMain":false,"metricName":"# contigs (>= 1000 bp)"},{"values":[0,0,0,0,0,0,0,0,0,0,0],"quality":"Equal","isMain":false,"metricName":"# contigs (>= 5000 bp)"},{"values":[0,0,0,0,0,0,0,0,0,0,0],"quality":"Equal","isMain":false,"metricName":"# contigs (>= 10000 bp)"},{"values":[0,0,0,0,0,0,0,0,0,0,0],"quality":"Equal","isMain":false,"metricName":"# contigs (>= 25000 bp)"},{"values":[0,0,0,0,0,0,0,0,0,0,0],"quality":"Equal","isMain":false,"metricName":"# contigs (>= 50000 bp)"},{"values":[598,1133,1478,975,690,1017,1289,1098,1289,1289,536],"quality":"More is better","isMain":true,"metricName":"Largest contig"},{"values":[598,10649,9290,17219,690,9259,21764,19827,22329,22329,536],"quality":"More is better","isMain":true,"metricName":"Total length"},{"values":[0,2155,1478,0,0,1017,3584,2120,3374,3374,0],"quality":"More is better","isMain":true,"metricName":"Total length (>= 1000 bp)"},{"values":[0,0,0,0,0,0,0,0,0,0,0],"quality":"More is better","isMain":false,"metricName":"Total length (>= 5000 bp)"},{"values":[0,0,0,0,0,0,0,0,0,0,0],"quality":"More is better","isMain":true,"metricName":"Total length (>= 10000 bp)"},{"values":[0,0,0,0,0,0,0,0,0,0,0],"quality":"More is better","isMain":false,"metricName":"Total length (>= 25000 bp)"},{"values":[0,0,0,0,0,0,0,0,0,0,0],"quality":"More is better","isMain":true,"metricName":"Total length (>= 50000 bp)"},{"values":[598,785,684,678,690,661,806,780,785,785,536],"quality":"More is better","isMain":false,"metricName":"N50"},{"values":[598,621,581,569,690,604,678,636,675,675,536],"quality":"More is better","isMain":false,"metricName":"N75"},{"values":[1,6,6,11,1,6,12,12,13,13,1],"quality":"Less is better","isMain":false,"metricName":"L50"},{"values":[1,10,9,19,1,10,19,19,20,20,1],"quality":"Less is better","isMain":false,"metricName":"L75"},{"values":["52.01","53.89","53.61","53.99","55.07","55.00","53.17","53.64","53.19","53.21","57.09"],"quality":"Equal","isMain":false,"metricName":"GC (%)"}]],["Predicted genes",[]],["Similarity statistics",[{"values":[0,0,0,0,0,0,0,0,0,0,0],"quality":"Equal","isMain":false,"metricName":"# similar correct contigs"},{"values":[0,0,0,0,0,0,0,0,0,0,0],"quality":"Equal","isMain":false,"metricName":"# similar misassembled blocks"}]],["Reference statistics",[{"values":[126629,126629,126629,126629,126629,126629,126629,126629,126629,126629,126629],"quality":"Equal","isMain":false,"metricName":"Reference length"},{"values":[1,1,1,1,1,1,1,1,1,1,1],"quality":"Equal","isMain":false,"metricName":"Reference fragments"},{"values":["53.47","53.47","53.47","53.47","53.47","53.47","53.47","53.47","53.47","53.47","53.47"],"quality":"Equal","isMain":false,"metricName":"Reference GC (%)"}]]],"referenceName":"gi\_154551150\_ref\_NC","date":"18 July 2018, Wednesday, 16:22:50","order":[0,1,2,3,4,5,6,7,8,9,10],"assembliesNames":["ABySS\_127","CLC","IDBA\_UD","MEGAHIT","MIRA","SOAPdenovo2","SPAdes","SPAdes\_meta","SPAdes\_sc","SPAdes\_sc\_careful","Velvet"]},{"assembliesWithNs":null,"minContig":500,"report":[["Genome statistics",[{"values":["100.000","93.428","42.819","100.000","100.000","100.000","100.000","99.726","99.726","23.054","100.000"],"quality":"More is better","isMain":true,"metricName":"Genome fraction (%)"},{"values":["1.073","1.027","1.000","141.005","1.996","1.003","1.001","1.000","1.000","1.001","2.116"],"quality":"Less is better","isMain":true,"metricName":"Duplication ratio"},{"values":[39104,6869,1334,31167,39104,39231,39159,38997,38997,1025,39104],"quality":"More is better","isMain":true,"metricName":"Largest alignment"},{"values":[41950,37516,16744,5512339,78060,39231,39159,38997,38997,9027,82760],"quality":"More is better","isMain":true,"metricName":"Total aligned length"},{"values":[39104,3928,null,41658,78060,39231,39159,38997,38997,null,179797],"quality":"More is better","isMain":false,"metricName":"NG50"},{"values":[39104,1847,null,41658,78060,39231,39159,38997,38997,null,179797],"quality":"More is better","isMain":false,"metricName":"NG75"},{"values":[39104,3928,808,3167,39104,39231,39159,38997,38997,639,null],"quality":"More is better","isMain":false,"metricName":"NA50"},{"values":[39104,1847,718,2212,33632,39231,39159,38997,38997,541,null],"quality":"More is better","isMain":false,"metricName":"NA75"},{"values":[39104,3928,null,31167,39104,39231,39159,38997,38997,null,39104],"quality":"More is better","isMain":true,"metricName":"NGA50"},{"values":[39104,1847,null,31167,39104,39231,39159,38997,38997,null,39104],"quality":"More is better","isMain":false,"metricName":"NGA75"},{"values":[1,4,null,1,1,1,1,1,1,null,1],"quality":"Less is better","isMain":false,"metricName":"LG50"},{"values":[1,8,null,1,1,1,1,1,1,null,1],"quality":"Less is better","isMain":false,"metricName":"LG75"},{"values":[1,4,9,531,1,1,1,1,1,6,null],"quality":"Less is better","isMain":false,"metricName":"LA50"},{"values":[1,8,15,1057,2,1,1,1,1,10,null],"quality":"Less is better","isMain":false,"metricName":"LA75"},{"values":[1,4,null,1,1,1,1,1,1,null,1],"quality":"Less is better","isMain":true,"metricName":"LGA50"},{"values":[1,8,null,1,1,1,1,1,1,null,1],"quality":"Less is better","isMain":false,"metricName":"LGA75"}]],["Misassemblies",[{"values":[0,0,0,1,2,0,0,0,0,0,2],"quality":"Less is better","isMain":true,"metricName":"# misassemblies"},{"values":[0,0,0,1,2,0,0,0,0,0,2],"quality":"Less is better","isMain":false,"metricName":" # relocations"},{"values":[0,0,0,0,0,0,0,0,0,0,0],"quality":"Less is better","isMain":false,"metricName":" # translocations"},{"values":[0,0,0,0,0,0,0,0,0,0,0],"quality":"Less is better","isMain":false,"metricName":" # inversions"},{"values":[0,0,0,1,1,0,0,0,0,0,1],"quality":"Less is better","isMain":false,"metricName":"# misassembled contigs"},{"values":[0,0,0,41658,78060,0,0,0,0,0,67995],"quality":"Less is better","isMain":true,"metricName":"Misassembled contigs length"},{"values":[0,0,0,0,0,1,0,0,0,0,0],"quality":"Less is better","isMain":false,"metricName":"# local misassemblies"},{"values":[0,0,0,1,0,0,0,0,0,0,2],"quality":"Less is better","isMain":false,"metricName":"# unaligned mis. contigs"}]],["Unaligned",[{"values":[0,0,0,0,0,0,0,0,0,0,0],"quality":"Less is better","isMain":false,"metricName":"# fully unaligned contigs"},{"values":[0,0,0,0,0,0,0,0,0,0,0],"quality":"Less is better","isMain":false,"metricName":"Fully unaligned length"},{"values":[0,1,0,4,0,0,0,0,0,0,3],"quality":"Less is better","isMain":false,"metricName":"# partially unaligned contigs"},{"values":[0,1382,0,4639,0,0,0,0,0,0,192209],"quality":"Less is better","isMain":false,"metricName":"Partially unaligned length"}]],["Mismatches",[{"values":[0,0,2,78,0,0,6,0,0,1,20],"quality":"Less is better","isMain":false,"metricName":"# mismatches"},{"values":[0,0,0,5,0,0,1,0,0,0,0],"quality":"Less is better","isMain":false,"metricName":"# indels"},{"values":[0,0,0,5,0,0,55,0,0,0,0],"quality":"Less is better","isMain":false,"metricName":"Indels length"},{"values":["0.00","0.00","11.94","199.47","0.00","0.00","15.34","0.00","0.00","11.09","51.15"],"quality":"Less is better","isMain":true,"metricName":"# mismatches per 100 kbp"},{"values":["0.00","0.00","0.00","12.79","0.00","0.00","2.56","0.00","0.00","0.00","0.00"],"quality":"Less is better","isMain":true,"metricName":"# indels per 100 kbp"},{"values":[0,0,0,5,0,0,0,0,0,0,0],"quality":"Less is better","isMain":false,"metricName":" # indels (<= 5 bp)"},{"values":[0,0,0,0,0,0,1,0,0,0,0],"quality":"Less is better","isMain":false,"metricName":" # indels (> 5 bp)"},{"values":[0,0,0,2948,0,0,0,0,0,0,455],"quality":"Less is better","isMain":false,"metricName":"# N's"},{"values":["0.00","0.00","0.00","53.42","0.00","0.00","0.00","0.00","0.00","0.00","165.47"],"quality":"Less is better","isMain":true,"metricName":"# N's per 100 kbp"}]],["Statistics without reference",[{"values":[6,18,21,1857,1,1,1,1,1,14,4],"quality":"Equal","isMain":true,"metricName":"# contigs"},{"values":[1,11,3,1831,1,1,1,1,1,1,4],"quality":"Equal","isMain":false,"metricName":"# contigs (>= 1000 bp)"},{"values":[1,2,0,193,1,1,1,1,1,0,4],"quality":"Equal","isMain":false,"metricName":"# contigs (>= 5000 bp)"},{"values":[1,0,0,26,1,1,1,1,1,0,3],"quality":"Equal","isMain":false,"metricName":"# contigs (>= 10000 bp)"},{"values":[1,0,0,1,1,1,1,1,1,0,2],"quality":"Equal","isMain":false,"metricName":"# contigs (>= 25000 bp)"},{"values":[0,0,0,0,1,0,0,0,0,0,2],"quality":"Equal","isMain":false,"metricName":"# contigs (>= 50000 bp)"},{"values":[39104,6869,1334,41658,78060,39231,39159,38997,38997,1025,179797],"quality":"More is better","isMain":true,"metricName":"Largest contig"},{"values":[41950,38898,16744,5518493,78060,39231,39159,38997,38997,9027,274972],"quality":"More is better","isMain":true,"metricName":"Total length"},{"values":[39104,33868,3444,5495051,78060,39231,39159,38997,38997,1025,274972],"quality":"More is better","isMain":true,"metricName":"Total length (>= 1000 bp)"},{"values":[39104,13592,0,1457352,78060,39231,39159,38997,38997,0,274972],"quality":"More is better","isMain":false,"metricName":"Total length (>= 5000 bp)"},{"values":[39104,0,0,345897,78060,39231,39159,38997,38997,0,268491],"quality":"More is better","isMain":true,"metricName":"Total length (>= 10000 bp)"},{"values":[39104,0,0,41658,78060,39231,39159,38997,38997,0,247792],"quality":"More is better","isMain":false,"metricName":"Total length (>= 25000 bp)"},{"values":[0,0,0,0,78060,0,0,0,0,0,247792],"quality":"More is better","isMain":true,"metricName":"Total length (>= 50000 bp)"},{"values":[39104,3928,808,3168,78060,39231,39159,38997,38997,639,179797],"quality":"More is better","isMain":false,"metricName":"N50"},{"values":[39104,1847,718,2215,78060,39231,39159,38997,38997,541,67995],"quality":"More is better","isMain":false,"metricName":"N75"},{"values":[1,4,9,529,1,1,1,1,1,6,1],"quality":"Less is better","isMain":false,"metricName":"L50"},{"values":[1,8,15,1054,1,1,1,1,1,10,2],"quality":"Less is better","isMain":false,"metricName":"L75"},{"values":["49.18","48.79","48.57","49.05","48.98","48.96","48.99","48.98","48.98","48.83","49.98"],"quality":"Equal","isMain":false,"metricName":"GC (%)"}]],["Predicted genes",[]],["Similarity statistics",[{"values":[0,0,0,0,0,0,0,0,0,0,0],"quality":"Equal","isMain":false,"metricName":"# similar correct contigs"},{"values":[0,0,0,0,0,0,0,0,0,0,0],"quality":"Equal","isMain":false,"metricName":"# similar misassembled blocks"}]],["Reference statistics",[{"values":[39104,39104,39104,39104,39104,39104,39104,39104,39104,39104,39104],"quality":"Equal","isMain":false,"metricName":"Reference length"},{"values":[1,1,1,1,1,1,1,1,1,1,1],"quality":"Equal","isMain":false,"metricName":"Reference fragments"},{"values":["48.97","48.97","48.97","48.97","48.97","48.97","48.97","48.97","48.97","48.97","48.97"],"quality":"Equal","isMain":false,"metricName":"Reference GC (%)"}]]],"referenceName":"gi\_155370093\_ref\_NC","date":"18 July 2018, Wednesday, 16:23:07","order":[0,1,2,3,4,5,6,7,8,9,10],"assembliesNames":["CLC","IDBA\_UD","MEGAHIT","MIRA","Ray\_Meta","SPAdes","SPAdes\_meta","SPAdes\_sc","SPAdes\_sc\_careful","Velvet","Geneious"]},{"assembliesWithNs":null,"minContig":500,"report":[["Genome statistics",[{"values":["8.066","2.813","60.438","54.675","62.524","21.498","47.739","70.676","72.124","71.026","71.567","2.695"],"quality":"More is better","isMain":true,"metricName":"Genome fraction (%)"},{"values":["1.000","1.000","1.027","1.006","1.000","1.007","1.002","1.007","1.004","1.006","1.006","1.000"],"quality":"Less is better","isMain":true,"metricName":"Duplication ratio"},{"values":[1925,948,3553,3542,3331,2598,2563,3542,3466,3542,3542,2006],"quality":"More is better","isMain":true,"metricName":"Largest alignment"},{"values":[19008,6628,144649,128948,147376,50660,112672,166767,170653,167466,168795,6350],"quality":"More is better","isMain":true,"metricName":"Total aligned length"},{"values":[null,null,926,675,782,null,null,1282,1040,1167,1197,null],"quality":"More is better","isMain":false,"metricName":"NG50"},{"values":[null,null,null,null,null,null,null,688,null,560,589,null],"quality":"More is better","isMain":false,"metricName":"NG75"},{"values":[651,804,1079,1048,1156,1116,912,1254,1282,1266,1266,1308],"quality":"More is better","isMain":false,"metricName":"NA50"},{"values":[579,661,705,709,774,924,654,747,870,836,833,613],"quality":"More is better","isMain":false,"metricName":"NA75"},{"values":[null,null,746,597,743,null,null,1018,1018,1002,1004,null],"quality":"More is better","isMain":true,"metricName":"NGA50"},{"values":[null,null,72,91,84,null,null,51,70,61,59,null],"quality":"Less is better","isMain":false,"metricName":"LG50"},{"values":[null,null,null,null,null,null,null,112,null,134,130,null],"quality":"Less is better","isMain":false,"metricName":"LG75"},{"values":[11,4,50,42,43,18,43,52,46,47,48,2],"quality":"Less is better","isMain":false,"metricName":"LA50"},{"values":[18,7,96,81,85,31,81,102,86,91,93,5],"quality":"Less is better","isMain":false,"metricName":"LA75"},{"values":[null,null,91,103,90,null,null,72,73,72,72,null],"quality":"Less is better","isMain":true,"metricName":"LGA50"}]],["Misassemblies",[{"values":[0,0,0,0,0,0,0,0,0,0,0,0],"quality":"Less is better","isMain":true,"metricName":"# misassemblies"},{"values":[0,0,0,0,0,0,0,0,0,0,0,0],"quality":"Less is better","isMain":false,"metricName":" # relocations"},{"values":[0,0,0,0,0,0,0,0,0,0,0,0],"quality":"Less is better","isMain":false,"metricName":" # translocations"},{"values":[0,0,0,0,0,0,0,0,0,0,0,0],"quality":"Less is better","isMain":false,"metricName":" # inversions"},{"values":[0,0,0,0,0,0,0,0,0,0,0,0],"quality":"Less is better","isMain":false,"metricName":"# misassembled contigs"},{"values":[0,0,0,0,0,0,0,0,0,0,0,0],"quality":"Less is better","isMain":true,"metricName":"Misassembled contigs length"},{"values":[0,0,0,1,0,0,0,0,0,0,0,0],"quality":"Less is better","isMain":false,"metricName":"# local misassemblies"},{"values":[0,0,1,0,0,0,0,0,0,0,0,0],"quality":"Less is better","isMain":false,"metricName":"# unaligned mis. contigs"}]],["Unaligned",[{"values":[0,0,0,0,0,0,0,0,0,0,0,0],"quality":"Less is better","isMain":false,"metricName":"# fully unaligned contigs"},{"values":[0,0,0,0,0,0,0,0,0,0,0,0],"quality":"Less is better","isMain":false,"metricName":"Fully unaligned length"},{"values":[0,0,9,6,3,0,0,11,2,6,7,0],"quality":"Less is better","isMain":false,"metricName":"# partially unaligned contigs"},{"values":[0,0,15971,7900,4858,0,0,24637,2920,11505,12787,0],"quality":"Less is better","isMain":false,"metricName":"Partially unaligned length"}]],["Mismatches",[{"values":[7,0,532,281,198,47,90,614,246,359,410,2],"quality":"Less is better","isMain":false,"metricName":"# mismatches"},{"values":[0,0,25,9,0,6,1,7,0,3,3,0],"quality":"Less is better","isMain":false,"metricName":"# indels"},{"values":[0,0,35,16,0,6,44,13,0,47,47,0],"quality":"Less is better","isMain":false,"metricName":"Indels length"},{"values":["36.83","0.00","373.55","218.10","134.39","92.78","80.00","368.67","144.74","214.49","243.12","31.50"],"quality":"Less is better","isMain":true,"metricName":"# mismatches per 100 kbp"},{"values":["0.00","0.00","17.55","6.99","0.00","11.84","0.89","4.20","0.00","1.79","1.78","0.00"],"quality":"Less is better","isMain":true,"metricName":"# indels per 100 kbp"},{"values":[0,0,25,9,0,6,0,7,0,2,2,0],"quality":"Less is better","isMain":false,"metricName":" # indels (<= 5 bp)"},{"values":[0,0,0,0,0,0,1,0,0,1,1,0],"quality":"Less is better","isMain":false,"metricName":" # indels (> 5 bp)"},{"values":[0,0,0,0,0,92,56,0,0,0,0,0],"quality":"Less is better","isMain":false,"metricName":"# N's"},{"values":["0.00","0.00","0.00","0.00","0.00","180.33","49.70","0.00","0.00","0.00","0.00","0.00"],"quality":"Less is better","isMain":true,"metricName":"# N's per 100 kbp"}]],["Statistics without reference",[{"values":[27,9,135,124,137,48,129,137,144,139,140,7],"quality":"Equal","isMain":true,"metricName":"# contigs"},{"values":[2,0,62,50,53,25,37,77,74,73,74,2],"quality":"Equal","isMain":false,"metricName":"# contigs (>= 1000 bp)"},{"values":[0,0,0,0,0,0,0,2,0,0,0,0],"quality":"Equal","isMain":false,"metricName":"# contigs (>= 5000 bp)"},{"values":[0,0,0,0,0,0,0,0,0,0,0,0],"quality":"Equal","isMain":false,"metricName":"# contigs (>= 10000 bp)"},{"values":[0,0,0,0,0,0,0,0,0,0,0,0],"quality":"Equal","isMain":false,"metricName":"# contigs (>= 25000 bp)"},{"values":[0,0,0,0,0,0,0,0,0,0,0,0],"quality":"Equal","isMain":false,"metricName":"# contigs (>= 50000 bp)"},{"values":[1925,948,4174,4208,4158,2598,2563,7056,4348,4348,4348,2006],"quality":"More is better","isMain":true,"metricName":"Largest contig"},{"values":[19008,6628,162288,137571,152234,51018,112672,192289,173574,179888,182503,6350],"quality":"More is better","isMain":true,"metricName":"Total length"},{"values":[3383,0,108916,84611,91747,33267,50794,148359,122251,131611,134226,3314],"quality":"More is better","isMain":true,"metricName":"Total length (>= 1000 bp)"},{"values":[0,0,0,0,0,0,0,13984,0,0,0,0],"quality":"More is better","isMain":false,"metricName":"Total length (>= 5000 bp)"},{"values":[0,0,0,0,0,0,0,0,0,0,0,0],"quality":"More is better","isMain":true,"metricName":"Total length (>= 10000 bp)"},{"values":[0,0,0,0,0,0,0,0,0,0,0,0],"quality":"More is better","isMain":false,"metricName":"Total length (>= 25000 bp)"},{"values":[0,0,0,0,0,0,0,0,0,0,0,0],"quality":"More is better","isMain":true,"metricName":"Total length (>= 50000 bp)"},{"values":[651,804,1308,1264,1199,1116,912,1512,1312,1405,1425,1308],"quality":"More is better","isMain":false,"metricName":"N50"},{"values":[579,661,874,828,800,924,654,1029,882,946,962,613],"quality":"More is better","isMain":false,"metricName":"N75"},{"values":[11,4,38,36,39,18,43,35,44,39,39,2],"quality":"Less is better","isMain":false,"metricName":"L50"},{"values":[18,7,76,71,79,31,81,73,83,77,77,5],"quality":"Less is better","isMain":false,"metricName":"L75"},{"values":["58.33","60.37","57.80","57.43","57.37","57.89","56.55","58.15","57.26","57.67","57.79","63.81"],"quality":"Equal","isMain":false,"metricName":"GC (%)"}]],["Predicted genes",[]],["Similarity statistics",[{"values":[0,0,0,0,0,0,0,0,0,0,0,0],"quality":"Equal","isMain":false,"metricName":"# similar correct contigs"},{"values":[0,0,0,0,0,0,0,0,0,0,0,0],"quality":"Equal","isMain":false,"metricName":"# similar misassembled blocks"}]],["Reference statistics",[{"values":[235646,235646,235646,235646,235646,235646,235646,235646,235646,235646,235646,235646],"quality":"Equal","isMain":false,"metricName":"Reference length"},{"values":[1,1,1,1,1,1,1,1,1,1,1,1],"quality":"Equal","isMain":false,"metricName":"Reference fragments"},{"values":["57.48","57.48","57.48","57.48","57.48","57.48","57.48","57.48","57.48","57.48","57.48","57.48"],"quality":"Equal","isMain":false,"metricName":"Reference GC (%)"}]]],"referenceName":"gi\_155573622\_ref\_NC","date":"18 July 2018, Wednesday, 16:23:20","order":[0,1,2,3,4,5,6,7,8,9,10,11],"assembliesNames":["ABySS\_127","ABySS\_63","CLC","IDBA\_UD","MEGAHIT","MIRA","SOAPdenovo2","SPAdes","SPAdes\_meta","SPAdes\_sc","SPAdes\_sc\_careful","Velvet"]},{"assembliesWithNs":null,"minContig":500,"report":[["Genome statistics",[{"values":["5.575","3.103","53.420","46.867","57.782","16.261","47.845","67.517","64.826","66.416","66.416","0.685"],"quality":"More is better","isMain":true,"metricName":"Genome fraction (%)"},{"values":["0.999","1.000","1.012","1.002","1.000","1.000","1.009","1.000","1.005","1.005","1.005","1.000"],"quality":"Less is better","isMain":true,"metricName":"Duplication ratio"},{"values":[1584,858,4127,3289,3983,1583,2206,4127,2889,4127,4127,954],"quality":"More is better","isMain":true,"metricName":"Largest alignment"},{"values":[7763,4324,75180,65327,80543,22658,66797,94114,90788,92868,92868,954],"quality":"More is better","isMain":true,"metricName":"Total aligned length"},{"values":[null,null,631,866,653,null,null,968,834,968,968,null],"quality":"More is better","isMain":false,"metricName":"NG50"},{"values":[601,585,1034,906,984,1110,1014,1206,1180,1206,1206,954],"quality":"More is better","isMain":false,"metricName":"NA50"},{"values":[592,555,631,null,735,907,664,823,840,795,795,954],"quality":"More is better","isMain":false,"metricName":"NA75"},{"values":[null,null,578,null,640,null,null,884,834,834,834,null],"quality":"More is better","isMain":true,"metricName":"NGA50"},{"values":[null,null,51,39,61,null,null,44,50,44,44,null],"quality":"Less is better","isMain":false,"metricName":"LG50"},{"values":[5,4,24,34,27,9,24,27,27,27,27,1],"quality":"Less is better","isMain":false,"metricName":"LA50"},{"values":[8,5,50,null,51,15,44,51,49,51,51,1],"quality":"Less is better","isMain":false,"metricName":"LA75"},{"values":[null,null,63,null,63,null,null,47,51,47,47,null],"quality":"Less is better","isMain":true,"metricName":"LGA50"}]],["Misassemblies",[{"values":[0,0,0,1,0,0,0,0,1,0,0,0],"quality":"Less is better","isMain":true,"metricName":"# misassemblies"},{"values":[0,0,0,1,0,0,0,0,1,0,0,0],"quality":"Less is better","isMain":false,"metricName":" # relocations"},{"values":[0,0,0,0,0,0,0,0,0,0,0,0],"quality":"Less is better","isMain":false,"metricName":" # translocations"},{"values":[0,0,0,0,0,0,0,0,0,0,0,0],"quality":"Less is better","isMain":false,"metricName":" # inversions"},{"values":[0,0,0,1,0,0,0,0,1,0,0,0],"quality":"Less is better","isMain":false,"metricName":"# misassembled contigs"},{"values":[0,0,0,920,0,0,0,0,1566,0,0,0],"quality":"Less is better","isMain":true,"metricName":"Misassembled contigs length"},{"values":[0,0,0,0,0,0,0,0,0,1,1,0],"quality":"Less is better","isMain":false,"metricName":"# local misassemblies"},{"values":[0,0,0,0,0,0,0,0,0,0,0,0],"quality":"Less is better","isMain":false,"metricName":"# unaligned mis. contigs"}]],["Unaligned",[{"values":[0,0,0,0,0,0,0,0,0,0,0,0],"quality":"Less is better","isMain":false,"metricName":"# fully unaligned contigs"},{"values":[0,0,0,0,0,0,0,0,0,0,0,0],"quality":"Less is better","isMain":false,"metricName":"Fully unaligned length"},{"values":[0,0,2,4,2,0,0,2,0,3,3,0],"quality":"Less is better","isMain":false,"metricName":"# partially unaligned contigs"},{"values":[0,0,7291,22513,1544,0,0,3854,0,4397,4397,0],"quality":"Less is better","isMain":false,"metricName":"Partially unaligned length"}]],["Mismatches",[{"values":[26,2,125,49,76,11,53,114,109,102,89,1],"quality":"Less is better","isMain":false,"metricName":"# mismatches"},{"values":[1,0,9,1,1,0,3,0,0,3,3,0],"quality":"Less is better","isMain":false,"metricName":"# indels"},{"values":[6,0,14,13,13,0,104,0,0,89,89,0],"quality":"Less is better","isMain":false,"metricName":"Indels length"},{"values":["334.66","46.25","167.93","75.03","94.39","48.55","79.50","121.17","120.67","110.22","96.17","104.82"],"quality":"Less is better","isMain":true,"metricName":"# mismatches per 100 kbp"},{"values":["12.87","0.00","12.09","1.53","1.24","0.00","4.50","0.00","0.00","3.24","3.24","0.00"],"quality":"Less is better","isMain":true,"metricName":"# indels per 100 kbp"},{"values":[0,0,9,0,0,0,0,0,0,1,1,0],"quality":"Less is better","isMain":false,"metricName":" # indels (<= 5 bp)"},{"values":[1,0,0,1,1,0,3,0,0,2,2,0],"quality":"Less is better","isMain":false,"metricName":" # indels (> 5 bp)"},{"values":[0,0,0,0,0,39,163,0,0,0,0,0],"quality":"Less is better","isMain":false,"metricName":"# N's"},{"values":["0.00","0.00","0.00","0.00","0.00","172.12","242.30","0.00","0.00","0.00","0.00","0.00"],"quality":"Less is better","isMain":true,"metricName":"# N's per 100 kbp"}]],["Statistics without reference",[{"values":[11,7,73,66,82,22,72,83,81,83,83,1],"quality":"Equal","isMain":true,"metricName":"# contigs"},{"values":[1,0,26,30,28,11,26,41,39,41,41,0],"quality":"Equal","isMain":false,"metricName":"# contigs (>= 1000 bp)"},{"values":[0,0,1,1,0,0,0,0,0,0,0,0],"quality":"Equal","isMain":false,"metricName":"# contigs (>= 5000 bp)"},{"values":[0,0,0,1,0,0,0,0,0,0,0,0],"quality":"Equal","isMain":false,"metricName":"# contigs (>= 10000 bp)"},{"values":[0,0,0,0,0,0,0,0,0,0,0,0],"quality":"Equal","isMain":false,"metricName":"# contigs (>= 25000 bp)"},{"values":[0,0,0,0,0,0,0,0,0,0,0,0],"quality":"Equal","isMain":false,"metricName":"# contigs (>= 50000 bp)"},{"values":[1584,858,8066,20296,3983,1583,2206,4127,2889,4127,4127,954],"quality":"More is better","isMain":true,"metricName":"Largest contig"},{"values":[7763,4324,82621,87980,82087,22658,67272,97968,90788,97372,97372,954],"quality":"More is better","isMain":true,"metricName":"Total length"},{"values":[1584,0,50274,61826,43084,13430,36490,67584,60047,67192,67192,0],"quality":"More is better","isMain":true,"metricName":"Total length (>= 1000 bp)"},{"values":[0,0,8066,20296,0,0,0,0,0,0,0,0],"quality":"More is better","isMain":false,"metricName":"Total length (>= 5000 bp)"},{"values":[0,0,0,20296,0,0,0,0,0,0,0,0],"quality":"More is better","isMain":true,"metricName":"Total length (>= 10000 bp)"},{"values":[0,0,0,0,0,0,0,0,0,0,0,0],"quality":"More is better","isMain":false,"metricName":"Total length (>= 25000 bp)"},{"values":[0,0,0,0,0,0,0,0,0,0,0,0],"quality":"More is better","isMain":true,"metricName":"Total length (>= 50000 bp)"},{"values":[601,585,1238,1274,1060,1110,1018,1265,1214,1265,1265,954],"quality":"More is better","isMain":false,"metricName":"N50"},{"values":[592,555,789,927,770,907,664,890,846,884,884,954],"quality":"More is better","isMain":false,"metricName":"N75"},{"values":[5,4,19,15,27,9,24,25,26,25,25,1],"quality":"Less is better","isMain":false,"metricName":"L50"},{"values":[8,5,40,35,50,15,44,48,48,48,48,1],"quality":"Less is better","isMain":false,"metricName":"L75"},{"values":["43.58","43.04","42.92","42.82","43.48","44.71","43.73","43.27","43.45","43.30","43.30","50.84"],"quality":"Equal","isMain":false,"metricName":"GC (%)"}]],["Predicted genes",[]],["Similarity statistics",[{"values":[0,0,0,0,0,0,0,0,0,0,0,0],"quality":"Equal","isMain":false,"metricName":"# similar correct contigs"},{"values":[0,0,0,0,0,0,0,0,0,0,0,0],"quality":"Equal","isMain":false,"metricName":"# similar misassembled blocks"}]],["Reference statistics",[{"values":[139342,139342,139342,139342,139342,139342,139342,139342,139342,139342,139342,139342],"quality":"Equal","isMain":false,"metricName":"Reference length"},{"values":[1,1,1,1,1,1,1,1,1,1,1,1],"quality":"Equal","isMain":false,"metricName":"Reference fragments"},{"values":["42.77","42.77","42.77","42.77","42.77","42.77","42.77","42.77","42.77","42.77","42.77","42.77"],"quality":"Equal","isMain":false,"metricName":"Reference GC (%)"}]]],"referenceName":"gi\_15617474\_ref\_NC","date":"18 July 2018, Wednesday, 16:23:31","order":[0,1,2,3,4,5,6,7,8,9,10,11],"assembliesNames":["ABySS\_127","ABySS\_63","CLC","IDBA\_UD","MEGAHIT","MIRA","SOAPdenovo2","SPAdes","SPAdes\_meta","SPAdes\_sc","SPAdes\_sc\_careful","Velvet"]},{"assembliesWithNs":null,"minContig":500,"report":[["Genome statistics",[{"values":["97.098","97.069","97.051","97.086","97.105","99.633","76.448","71.964","97.099","97.066","97.041","97.041","97.055","97.159"],"quality":"More is better","isMain":true,"metricName":"Genome fraction (%)"},{"values":["1.000","1.000","1.000","1.000","1.001","1.002","1.022","1.024","1.001","1.000","1.000","1.000","1.000","1.001"],"quality":"Less is better","isMain":true,"metricName":"Duplication ratio"},{"values":[212595,212531,206034,206188,206272,212728,6523,18061,206244,206100,196339,196339,206050,212728],"quality":"More is better","isMain":true,"metricName":"Largest alignment"},{"values":[212595,212531,212513,212667,212751,218236,171130,159586,212723,212579,212524,212524,212529,212728],"quality":"More is better","isMain":true,"metricName":"Total aligned length"},{"values":[212595,212531,206034,206188,206272,212966,1101,8155,206244,206100,212524,212524,206050,212966],"quality":"More is better","isMain":false,"metricName":"NG50"},{"values":[212595,212531,206034,206188,206272,212966,561,4945,206244,206100,212524,212524,206050,212966],"quality":"More is better","isMain":false,"metricName":"NG75"},{"values":[212595,212531,206034,206188,206272,212728,1375,4848,206244,206100,196339,196339,206050,212728],"quality":"More is better","isMain":false,"metricName":"NA50"},{"values":[212595,212531,206034,206188,206272,212728,863,689,206244,206100,196339,196339,206050,212728],"quality":"More is better","isMain":false,"metricName":"NA75"},{"values":[212595,212531,206034,206188,206272,212728,1101,4696,206244,206100,196339,196339,206050,212728],"quality":"More is better","isMain":true,"metricName":"NGA50"},{"values":[212595,212531,206034,206188,206272,212728,561,null,206244,206100,196339,196339,206050,212728],"quality":"More is better","isMain":false,"metricName":"NGA75"},{"values":[1,1,1,1,1,1,61,7,1,1,1,1,1,1],"quality":"Less is better","isMain":false,"metricName":"LG50"},{"values":[1,1,1,1,1,1,132,15,1,1,1,1,1,1],"quality":"Less is better","isMain":false,"metricName":"LG75"},{"values":[1,1,1,1,1,1,41,10,1,1,1,1,1,1],"quality":"Less is better","isMain":false,"metricName":"LA50"},{"values":[1,1,1,1,1,1,80,34,1,1,1,1,1,1],"quality":"Less is better","isMain":false,"metricName":"LA75"},{"values":[1,1,1,1,1,1,61,11,1,1,1,1,1,1],"quality":"Less is better","isMain":true,"metricName":"LGA50"},{"values":[1,1,1,1,1,1,132,null,1,1,1,1,1,1],"quality":"Less is better","isMain":false,"metricName":"LGA75"}]],["Misassemblies",[{"values":[0,0,0,0,0,0,0,0,0,0,1,1,0,0],"quality":"Less is better","isMain":true,"metricName":"# misassemblies"},{"values":[0,0,0,0,0,0,0,0,0,0,1,1,0,0],"quality":"Less is better","isMain":false,"metricName":" # relocations"},{"values":[0,0,0,0,0,0,0,0,0,0,0,0,0,0],"quality":"Less is better","isMain":false,"metricName":" # translocations"},{"values":[0,0,0,0,0,0,0,0,0,0,0,0,0,0],"quality":"Less is better","isMain":false,"metricName":" # inversions"},{"values":[0,0,0,0,0,0,0,0,0,0,1,1,0,0],"quality":"Less is better","isMain":false,"metricName":"# misassembled contigs"},{"values":[0,0,0,0,0,0,0,0,0,0,212524,212524,0,0],"quality":"Less is better","isMain":true,"metricName":"Misassembled contigs length"},{"values":[0,0,0,0,0,0,0,10,0,0,0,0,0,0],"quality":"Less is better","isMain":false,"metricName":"# local misassemblies"},{"values":[0,0,0,0,0,0,0,2,0,0,0,0,0,0],"quality":"Less is better","isMain":false,"metricName":"# unaligned mis. contigs"}]],["Unaligned",[{"values":[0,0,0,0,0,0,0,0,0,0,0,0,0,0],"quality":"Less is better","isMain":false,"metricName":"# fully unaligned contigs"},{"values":[0,0,0,0,0,0,0,0,0,0,0,0,0,0],"quality":"Less is better","isMain":false,"metricName":"Fully unaligned length"},{"values":[0,0,0,0,0,0,0,12,0,0,0,0,0,0],"quality":"Less is better","isMain":false,"metricName":"# partially unaligned contigs"},{"values":[0,0,0,0,0,0,0,44585,0,0,0,0,0,0],"quality":"Less is better","isMain":false,"metricName":"Partially unaligned length"}]],["Mismatches",[{"values":[0,0,0,0,0,0,0,0,0,0,0,0,0,0],"quality":"Less is better","isMain":false,"metricName":"# mismatches"},{"values":[0,0,0,0,0,0,0,80,0,0,0,0,0,0],"quality":"Less is better","isMain":false,"metricName":"# indels"},{"values":[0,0,0,0,0,0,0,3739,0,0,0,0,0,0],"quality":"Less is better","isMain":false,"metricName":"Indels length"},{"values":["0.00","0.00","0.00","0.00","0.00","0.00","0.00","0.00","0.00","0.00","0.00","0.00","0.00","0.00"],"quality":"Less is better","isMain":true,"metricName":"# mismatches per 100 kbp"},{"values":["0.00","0.00","0.00","0.00","0.00","0.00","0.00","50.77","0.00","0.00","0.00","0.00","0.00","0.00"],"quality":"Less is better","isMain":true,"metricName":"# indels per 100 kbp"},{"values":[0,0,0,0,0,0,0,0,0,0,0,0,0,0],"quality":"Less is better","isMain":false,"metricName":" # indels (<= 5 bp)"},{"values":[0,0,0,0,0,0,0,80,0,0,0,0,0,0],"quality":"Less is better","isMain":false,"metricName":" # indels (> 5 bp)"},{"values":[0,0,0,0,0,0,0,13897,0,0,0,0,0,0],"quality":"Less is better","isMain":false,"metricName":"# N's"},{"values":["0.00","0.00","0.00","0.00","0.00","0.00","0.00","6747.46","0.00","0.00","0.00","0.00","0.00","0.00"],"quality":"Less is better","isMain":true,"metricName":"# N's per 100 kbp"}]],["Statistics without reference",[{"values":[1,1,2,2,2,3,144,38,2,2,1,1,2,1],"quality":"Equal","isMain":true,"metricName":"# contigs"},{"values":[1,1,2,2,2,2,73,27,2,2,1,1,2,1],"quality":"Equal","isMain":false,"metricName":"# contigs (>= 1000 bp)"},{"values":[1,1,2,2,2,1,1,14,2,2,1,1,2,1],"quality":"Equal","isMain":false,"metricName":"# contigs (>= 5000 bp)"},{"values":[1,1,1,1,1,1,0,5,1,1,1,1,1,1],"quality":"Equal","isMain":false,"metricName":"# contigs (>= 10000 bp)"},{"values":[1,1,1,1,1,1,0,2,1,1,1,1,1,1],"quality":"Equal","isMain":false,"metricName":"# contigs (>= 25000 bp)"},{"values":[1,1,1,1,1,1,0,0,1,1,1,1,1,1],"quality":"Equal","isMain":false,"metricName":"# contigs (>= 50000 bp)"},{"values":[212595,212531,206034,206188,206272,212966,6523,27762,206244,206100,212524,212524,206050,212966],"quality":"More is better","isMain":true,"metricName":"Largest contig"},{"values":[212595,212531,212513,212667,212751,218474,171131,205959,212723,212579,212524,212524,212529,212966],"quality":"More is better","isMain":true,"metricName":"Total length"},{"values":[212595,212531,212513,212667,212751,217910,122382,197788,212723,212579,212524,212524,212529,212966],"quality":"More is better","isMain":true,"metricName":"Total length (>= 1000 bp)"},{"values":[212595,212531,212513,212667,212751,212966,6523,159429,212723,212579,212524,212524,212529,212966],"quality":"More is better","isMain":false,"metricName":"Total length (>= 5000 bp)"},{"values":[212595,212531,206034,206188,206272,212966,0,95971,206244,206100,212524,212524,206050,212966],"quality":"More is better","isMain":true,"metricName":"Total length (>= 10000 bp)"},{"values":[212595,212531,206034,206188,206272,212966,0,54971,206244,206100,212524,212524,206050,212966],"quality":"More is better","isMain":false,"metricName":"Total length (>= 25000 bp)"},{"values":[212595,212531,206034,206188,206272,212966,0,0,206244,206100,212524,212524,206050,212966],"quality":"More is better","isMain":true,"metricName":"Total length (>= 50000 bp)"},{"values":[212595,212531,206034,206188,206272,212966,1375,8674,206244,206100,212524,212524,206050,212966],"quality":"More is better","isMain":false,"metricName":"N50"},{"values":[212595,212531,206034,206188,206272,212966,863,5664,206244,206100,212524,212524,206050,212966],"quality":"More is better","isMain":false,"metricName":"N75"},{"values":[1,1,1,1,1,1,41,6,1,1,1,1,1,1],"quality":"Less is better","isMain":false,"metricName":"L50"},{"values":[1,1,1,1,1,1,80,14,1,1,1,1,1,1],"quality":"Less is better","isMain":false,"metricName":"L75"},{"values":["41.84","41.84","41.84","41.84","41.84","41.78","41.75","41.86","41.84","41.84","41.84","41.84","41.84","41.85"],"quality":"Equal","isMain":false,"metricName":"GC (%)"}]],["Predicted genes",[]],["Similarity statistics",[{"values":[1,1,1,1,1,1,0,0,1,1,0,0,1,1],"quality":"Equal","isMain":false,"metricName":"# similar correct contigs"},{"values":[0,0,0,0,0,0,0,0,0,0,0,0,0,0],"quality":"Equal","isMain":false,"metricName":"# similar misassembled blocks"}]],["Reference statistics",[{"values":[218948,218948,218948,218948,218948,218948,218948,218948,218948,218948,218948,218948,218948,218948],"quality":"Equal","isMain":false,"metricName":"Reference length"},{"values":[1,1,1,1,1,1,1,1,1,1,1,1,1,1],"quality":"Equal","isMain":false,"metricName":"Reference fragments"},{"values":["41.80","41.80","41.80","41.80","41.80","41.80","41.80","41.80","41.80","41.80","41.80","41.80","41.80","41.80"],"quality":"Equal","isMain":false,"metricName":"Reference GC (%)"}]]],"referenceName":"gi\_156563990\_ref\_NC","date":"18 July 2018, Wednesday, 16:23:44","order":[0,1,2,3,4,5,6,7,8,9,10,11,12,13],"assembliesNames":["ABySS\_127","ABySS\_63","CLC","IDBA\_UD","MEGAHIT","MIRA","Ray\_Meta","SOAPdenovo2","SPAdes","SPAdes\_meta","SPAdes\_sc","SPAdes\_sc\_careful","Velvet","Geneious"]},{"assembliesWithNs":null,"minContig":500,"report":[["Genome statistics",[{"values":["27.894","22.832","30.529","30.290","27.609","56.053","29.314","28.758","22.883","32.639","38.109","21.600","43.459"],"quality":"More is better","isMain":true,"metricName":"Genome fraction (%)"},{"values":["1.011","1.007","1.002","1.043","1.020","1.546","1.004","1.064","1.000","1.021","1.028","1.007","1.270"],"quality":"Less is better","isMain":true,"metricName":"Duplication ratio"},{"values":[4865,4388,1818,1953,2207,9311,5040,2195,5959,3027,3027,1695,9555],"quality":"More is better","isMain":true,"metricName":"Largest alignment"},{"values":[11676,9516,12663,13080,11665,35430,12188,12670,9475,13790,16220,9004,22848],"quality":"More is better","isMain":true,"metricName":"Total aligned length"},{"values":[null,null,null,null,null,1774,null,null,null,null,null,null,60961],"quality":"More is better","isMain":false,"metricName":"NG50"},{"values":[null,null,null,null,null,927,null,null,null,null,null,null,60961],"quality":"More is better","isMain":false,"metricName":"NG75"},{"values":[1621,2068,1182,1039,1319,1766,3816,1448,5959,1217,1258,902,null],"quality":"More is better","isMain":false,"metricName":"NA50"},{"values":[737,1553,993,694,802,927,2042,721,1792,807,777,757,null],"quality":"More is better","isMain":false,"metricName":"NA75"},{"values":[null,null,null,null,null,1203,null,null,null,null,null,null,1128],"quality":"More is better","isMain":true,"metricName":"NGA50"},{"values":[null,null,null,null,null,854,null,null,null,null,null,null,null],"quality":"More is better","isMain":false,"metricName":"NGA75"},{"values":[null,null,null,null,null,6,null,null,null,null,null,null,1],"quality":"Less is better","isMain":false,"metricName":"LG50"},{"values":[null,null,null,null,null,15,null,null,null,null,null,null,1],"quality":"Less is better","isMain":false,"metricName":"LG75"},{"values":[3,2,5,5,4,6,2,4,1,4,4,4,null],"quality":"Less is better","isMain":false,"metricName":"LA50"},{"values":[6,3,7,9,7,14,3,7,2,8,9,7,null],"quality":"Less is better","isMain":false,"metricName":"LA75"},{"values":[null,null,null,null,null,7,null,null,null,null,null,null,6],"quality":"Less is better","isMain":true,"metricName":"LGA50"},{"values":[null,null,null,null,null,17,null,null,null,null,null,null,null],"quality":"Less is better","isMain":false,"metricName":"LGA75"}]],["Misassemblies",[{"values":[0,0,0,0,0,0,0,0,0,0,0,0,0],"quality":"Less is better","isMain":true,"metricName":"# misassemblies"},{"values":[0,0,0,0,0,0,0,0,0,0,0,0,0],"quality":"Less is better","isMain":false,"metricName":" # relocations"},{"values":[0,0,0,0,0,0,0,0,0,0,0,0,0],"quality":"Less is better","isMain":false,"metricName":" # translocations"},{"values":[0,0,0,0,0,0,0,0,0,0,0,0,0],"quality":"Less is better","isMain":false,"metricName":" # inversions"},{"values":[0,0,0,0,0,0,0,0,0,0,0,0,0],"quality":"Less is better","isMain":false,"metricName":"# misassembled contigs"},{"values":[0,0,0,0,0,0,0,0,0,0,0,0,0],"quality":"Less is better","isMain":true,"metricName":"Misassembled contigs length"},{"values":[0,0,0,0,0,1,0,0,0,0,0,0,1],"quality":"Less is better","isMain":false,"metricName":"# local misassemblies"},{"values":[0,0,0,0,0,0,0,0,0,0,0,0,2],"quality":"Less is better","isMain":false,"metricName":"# unaligned mis. contigs"}]],["Unaligned",[{"values":[0,0,0,0,0,0,0,0,0,0,0,0,0],"quality":"Less is better","isMain":false,"metricName":"# fully unaligned contigs"},{"values":[0,0,0,0,0,0,0,0,0,0,0,0,0],"quality":"Less is better","isMain":false,"metricName":"Fully unaligned length"},{"values":[1,0,0,0,0,2,0,0,0,1,1,0,4],"quality":"Less is better","isMain":false,"metricName":"# partially unaligned contigs"},{"values":[2770,0,0,0,0,2002,0,0,0,761,761,0,76489],"quality":"Less is better","isMain":false,"metricName":"Partially unaligned length"}]],["Mismatches",[{"values":[0,1,22,0,11,178,23,5,22,46,34,5,148],"quality":"Less is better","isMain":false,"metricName":"# mismatches"},{"values":[0,0,0,0,0,16,0,0,0,2,4,0,9],"quality":"Less is better","isMain":false,"metricName":"# indels"},{"values":[0,0,0,0,0,25,0,0,0,2,4,0,17],"quality":"Less is better","isMain":false,"metricName":"Indels length"},{"values":["0.00","10.58","174.04","0.00","96.22","766.91","189.49","41.99","232.19","340.36","215.46","55.90","822.45"],"quality":"Less is better","isMain":true,"metricName":"# mismatches per 100 kbp"},{"values":["0.00","0.00","0.00","0.00","0.00","68.94","0.00","0.00","0.00","14.80","25.35","0.00","50.01"],"quality":"Less is better","isMain":true,"metricName":"# indels per 100 kbp"},{"values":[0,0,0,0,0,15,0,0,0,2,4,0,8],"quality":"Less is better","isMain":false,"metricName":" # indels (<= 5 bp)"},{"values":[0,0,0,0,0,1,0,0,0,0,0,0,1],"quality":"Less is better","isMain":false,"metricName":" # indels (> 5 bp)"},{"values":[50,12,0,0,0,8,0,0,0,0,0,0,148],"quality":"Less is better","isMain":false,"metricName":"# N's"},{"values":["346.12","126.10","0.00","0.00","0.00","21.12","0.00","0.00","0.00","0.00","0.00","0.00","148.99"],"quality":"Less is better","isMain":true,"metricName":"# N's per 100 kbp"}]],["Statistics without reference",[{"values":[7,5,12,14,11,23,4,11,3,11,13,10,5],"quality":"Equal","isMain":true,"metricName":"# contigs"},{"values":[3,3,6,5,5,13,4,5,3,6,6,3,5],"quality":"Equal","isMain":false,"metricName":"# contigs (>= 1000 bp)"},{"values":[0,0,0,0,0,1,1,0,1,0,0,0,4],"quality":"Equal","isMain":false,"metricName":"# contigs (>= 5000 bp)"},{"values":[0,0,0,0,0,0,0,0,0,0,0,0,4],"quality":"Equal","isMain":false,"metricName":"# contigs (>= 10000 bp)"},{"values":[0,0,0,0,0,0,0,0,0,0,0,0,1],"quality":"Equal","isMain":false,"metricName":"# contigs (>= 25000 bp)"},{"values":[0,0,0,0,0,0,0,0,0,0,0,0,1],"quality":"Equal","isMain":false,"metricName":"# contigs (>= 50000 bp)"},{"values":[4865,4388,1818,1953,2207,9755,5040,2195,5959,3788,3788,1695,60961],"quality":"More is better","isMain":true,"metricName":"Largest contig"},{"values":[14446,9516,12664,13080,11665,37876,12188,12670,9475,14554,16981,9004,99337],"quality":"More is better","isMain":true,"metricName":"Total length"},{"values":[11250,8009,8511,6913,7451,29905,12188,8695,9475,10693,11621,3975,99337],"quality":"More is better","isMain":true,"metricName":"Total length (>= 1000 bp)"},{"values":[0,0,0,0,0,9755,5040,0,5959,0,0,0,97770],"quality":"More is better","isMain":false,"metricName":"Total length (>= 5000 bp)"},{"values":[0,0,0,0,0,0,0,0,0,0,0,0,97770],"quality":"More is better","isMain":true,"metricName":"Total length (>= 10000 bp)"},{"values":[0,0,0,0,0,0,0,0,0,0,0,0,60961],"quality":"More is better","isMain":false,"metricName":"Total length (>= 25000 bp)"},{"values":[0,0,0,0,0,0,0,0,0,0,0,0,60961],"quality":"More is better","isMain":true,"metricName":"Total length (>= 50000 bp)"},{"values":[4764,2068,1182,1039,1319,1877,3816,1448,5959,1565,1258,902,60961],"quality":"More is better","isMain":false,"metricName":"N50"},{"values":[1621,1553,993,694,802,1042,2042,721,1792,951,868,757,13847],"quality":"More is better","isMain":false,"metricName":"N75"},{"values":[2,2,5,5,4,5,2,4,1,3,4,4,1],"quality":"Less is better","isMain":false,"metricName":"L50"},{"values":[3,3,7,9,7,12,3,7,2,7,8,7,2],"quality":"Less is better","isMain":false,"metricName":"L75"},{"values":["33.80","33.40","33.44","33.33","32.82","32.96","33.47","33.12","32.16","33.97","33.58","33.94","34.20"],"quality":"Equal","isMain":false,"metricName":"GC (%)"}]],["Predicted genes",[]],["Similarity statistics",[{"values":[0,0,0,0,0,0,0,0,0,0,0,0,0],"quality":"Equal","isMain":false,"metricName":"# similar correct contigs"},{"values":[0,0,0,0,0,0,0,0,0,0,0,0,0],"quality":"Equal","isMain":false,"metricName":"# similar misassembled blocks"}]],["Reference statistics",[{"values":[41407,41407,41407,41407,41407,41407,41407,41407,41407,41407,41407,41407,41407],"quality":"Equal","isMain":false,"metricName":"Reference length"},{"values":[1,1,1,1,1,1,1,1,1,1,1,1,1],"quality":"Equal","isMain":false,"metricName":"Reference fragments"},{"values":["33.62","33.62","33.62","33.62","33.62","33.62","33.62","33.62","33.62","33.62","33.62","33.62","33.62"],"quality":"Equal","isMain":false,"metricName":"Reference GC (%)"}]]],"referenceName":"gi\_157102936\_ref\_NC","date":"18 July 2018, Wednesday, 16:23:57","order":[0,1,2,3,4,5,6,7,8,9,10,11,12],"assembliesNames":["ABySS\_127","ABySS\_63","CLC","IDBA\_UD","MEGAHIT","MIRA","Ray\_Meta","SPAdes","SPAdes\_meta","SPAdes\_sc","SPAdes\_sc\_careful","Velvet","Geneious"]},{"assembliesWithNs":null,"minContig":500,"report":[["Genome statistics",[{"values":["31.634","1.678","55.944","37.854","46.473","95.222","44.119","41.326","14.970","41.927","22.496","9.026","100.000"],"quality":"More is better","isMain":true,"metricName":"Genome fraction (%)"},{"values":["1.008","1.000","1.095","1.019","1.043","3.425","1.000","1.032","1.000","1.009","1.005","1.001","2.579"],"quality":"Less is better","isMain":true,"metricName":"Duplication ratio"},{"values":[6548,767,5806,1368,4719,39941,4153,6382,3730,9005,4340,1108,39939],"quality":"More is better","isMain":true,"metricName":"Largest alignment"},{"values":[14574,767,27934,17627,22159,148692,20167,19486,6844,19330,10337,4129,117864],"quality":"More is better","isMain":true,"metricName":"Total aligned length"},{"values":[null,null,732,null,null,73503,null,null,null,null,null,null,92482],"quality":"More is better","isMain":false,"metricName":"NG50"},{"values":[null,null,null,null,null,73503,null,null,null,null,null,null,92482],"quality":"More is better","isMain":false,"metricName":"NG75"},{"values":[792,767,1357,816,1452,7476,3118,1318,3730,4415,2317,669,2972],"quality":"More is better","isMain":false,"metricName":"NA50"},{"values":[null,767,774,713,894,null,2574,695,2294,2294,2058,547,null],"quality":"More is better","isMain":false,"metricName":"NA75"},{"values":[null,null,732,null,null,39941,null,null,null,null,null,null,39939],"quality":"More is better","isMain":true,"metricName":"NGA50"},{"values":[null,null,null,null,null,39941,null,null,null,null,null,null,39939],"quality":"More is better","isMain":false,"metricName":"NGA75"},{"values":[null,null,16,null,null,1,null,null,null,null,null,null,1],"quality":"Less is better","isMain":false,"metricName":"LG50"},{"values":[null,null,null,null,null,1,null,null,null,null,null,null,1],"quality":"Less is better","isMain":false,"metricName":"LG75"},{"values":[4,1,6,9,5,5,3,3,1,2,2,3,8],"quality":"Less is better","isMain":false,"metricName":"LA50"},{"values":[null,1,14,15,10,null,5,7,2,3,3,5,null],"quality":"Less is better","isMain":false,"metricName":"LA75"},{"values":[null,null,16,null,null,1,null,null,null,null,null,null,1],"quality":"Less is better","isMain":true,"metricName":"LGA50"},{"values":[null,null,null,null,null,1,null,null,null,null,null,null,1],"quality":"Less is better","isMain":false,"metricName":"LGA75"}]],["Misassemblies",[{"values":[0,0,0,0,0,2,0,0,0,0,0,0,1],"quality":"Less is better","isMain":true,"metricName":"# misassemblies"},{"values":[0,0,0,0,0,2,0,0,0,0,0,0,1],"quality":"Less is better","isMain":false,"metricName":" # relocations"},{"values":[0,0,0,0,0,0,0,0,0,0,0,0,0],"quality":"Less is better","isMain":false,"metricName":" # translocations"},{"values":[0,0,0,0,0,0,0,0,0,0,0,0,0],"quality":"Less is better","isMain":false,"metricName":" # inversions"},{"values":[0,0,0,0,0,2,0,0,0,0,0,0,1],"quality":"Less is better","isMain":false,"metricName":"# misassembled contigs"},{"values":[0,0,0,0,0,119942,0,0,0,0,0,0,73504],"quality":"Less is better","isMain":true,"metricName":"Misassembled contigs length"},{"values":[0,0,0,0,0,1,0,0,0,0,0,0,1],"quality":"Less is better","isMain":false,"metricName":"# local misassemblies"},{"values":[0,0,0,0,0,1,0,0,0,0,0,0,3],"quality":"Less is better","isMain":false,"metricName":"# unaligned mis. contigs"}]],["Unaligned",[{"values":[0,0,0,0,0,0,0,0,0,0,0,0,0],"quality":"Less is better","isMain":false,"metricName":"# fully unaligned contigs"},{"values":[0,0,0,0,0,0,0,0,0,0,0,0,0],"quality":"Less is better","isMain":false,"metricName":"Fully unaligned length"},{"values":[1,0,0,0,0,4,0,0,0,1,0,0,5],"quality":"Less is better","isMain":false,"metricName":"# partially unaligned contigs"},{"values":[5766,0,0,0,0,64255,0,0,0,676,0,0,110203],"quality":"Less is better","isMain":false,"metricName":"Partially unaligned length"}]],["Mismatches",[{"values":[0,0,57,0,62,490,24,14,39,86,64,5,119],"quality":"Less is better","isMain":false,"metricName":"# mismatches"},{"values":[0,0,0,0,0,13,0,0,1,2,1,0,9],"quality":"Less is better","isMain":false,"metricName":"# indels"},{"values":[0,0,0,0,0,46,0,0,1,2,1,0,41],"quality":"Less is better","isMain":false,"metricName":"Indels length"},{"values":["0.00","0.00","222.90","0.00","291.86","1125.76","119.01","74.11","569.93","448.73","622.39","121.18","260.34"],"quality":"Less is better","isMain":true,"metricName":"# mismatches per 100 kbp"},{"values":["0.00","0.00","0.00","0.00","0.00","29.87","0.00","0.00","14.61","10.44","9.72","0.00","19.69"],"quality":"Less is better","isMain":true,"metricName":"# indels per 100 kbp"},{"values":[0,0,0,0,0,12,0,0,1,2,1,0,8],"quality":"Less is better","isMain":false,"metricName":" # indels (<= 5 bp)"},{"values":[0,0,0,0,0,1,0,0,0,0,0,0,1],"quality":"Less is better","isMain":false,"metricName":" # indels (> 5 bp)"},{"values":[50,0,0,0,0,33,0,0,0,676,0,0,1138],"quality":"Less is better","isMain":false,"metricName":"# N's"},{"values":["245.82","0.00","0.00","0.00","0.00","15.47","0.00","0.00","0.00","3378.65","0.00","0.00","498.98"],"quality":"Less is better","isMain":true,"metricName":"# N's per 100 kbp"}]],["Statistics without reference",[{"values":[11,1,24,22,18,23,8,15,3,7,6,6,5],"quality":"Equal","isMain":true,"metricName":"# contigs"},{"values":[2,0,9,4,7,18,6,4,2,4,3,1,5],"quality":"Equal","isMain":false,"metricName":"# contigs (>= 1000 bp)"},{"values":[2,0,1,0,0,5,0,1,0,1,0,0,5],"quality":"Equal","isMain":false,"metricName":"# contigs (>= 5000 bp)"},{"values":[0,0,0,0,0,4,0,0,0,0,0,0,4],"quality":"Equal","isMain":false,"metricName":"# contigs (>= 10000 bp)"},{"values":[0,0,0,0,0,3,0,0,0,0,0,0,4],"quality":"Equal","isMain":false,"metricName":"# contigs (>= 25000 bp)"},{"values":[0,0,0,0,0,1,0,0,0,0,0,0,2],"quality":"Equal","isMain":false,"metricName":"# contigs (>= 50000 bp)"},{"values":[8110,767,5806,1368,4719,73503,4153,6382,3730,9681,4340,1108,92482],"quality":"More is better","isMain":true,"metricName":"Largest contig"},{"values":[20340,767,28008,17627,22162,213323,20167,19486,6845,20008,10337,4129,228067],"quality":"More is better","isMain":true,"metricName":"Total length"},{"values":[14658,0,17464,4828,13855,208994,18475,12240,6024,18039,8715,1108,228067],"quality":"More is better","isMain":true,"metricName":"Total length (>= 1000 bp)"},{"values":[14658,0,5806,0,0,184706,0,6382,0,9681,0,0,228067],"quality":"More is better","isMain":false,"metricName":"Total length (>= 5000 bp)"},{"values":[0,0,0,0,0,177230,0,0,0,0,0,0,223025],"quality":"More is better","isMain":true,"metricName":"Total length (>= 10000 bp)"},{"values":[0,0,0,0,0,163827,0,0,0,0,0,0,223025],"quality":"More is better","isMain":false,"metricName":"Total length (>= 25000 bp)"},{"values":[0,0,0,0,0,73503,0,0,0,0,0,0,165986],"quality":"More is better","isMain":true,"metricName":"Total length (>= 50000 bp)"},{"values":[6548,767,1357,816,1452,46439,3118,1318,3730,4416,2317,669,73504],"quality":"More is better","isMain":false,"metricName":"N50"},{"values":[940,767,843,713,894,43885,2574,695,2294,2294,2058,547,28595],"quality":"More is better","isMain":false,"metricName":"N75"},{"values":[2,1,6,9,5,2,3,3,1,2,2,3,2],"quality":"Less is better","isMain":false,"metricName":"L50"},{"values":[3,1,13,15,10,3,5,7,2,3,3,5,3],"quality":"Less is better","isMain":false,"metricName":"L75"},{"values":["34.24","30.25","34.11","33.70","33.53","33.55","33.56","34.30","33.03","34.03","34.27","31.80","33.64"],"quality":"Equal","isMain":false,"metricName":"GC (%)"}]],["Predicted genes",[]],["Similarity statistics",[{"values":[0,0,0,0,0,0,0,0,0,0,0,0,0],"quality":"Equal","isMain":false,"metricName":"# similar correct contigs"},{"values":[0,0,0,0,0,0,0,0,0,0,0,0,0],"quality":"Equal","isMain":false,"metricName":"# similar misassembled blocks"}]],["Reference statistics",[{"values":[45710,45710,45710,45710,45710,45710,45710,45710,45710,45710,45710,45710,45710],"quality":"Equal","isMain":false,"metricName":"Reference length"},{"values":[1,1,1,1,1,1,1,1,1,1,1,1,1],"quality":"Equal","isMain":false,"metricName":"Reference fragments"},{"values":["33.59","33.59","33.59","33.59","33.59","33.59","33.59","33.59","33.59","33.59","33.59","33.59","33.59"],"quality":"Equal","isMain":false,"metricName":"Reference GC (%)"}]]],"referenceName":"gi\_157102937\_ref\_NC","date":"18 July 2018, Wednesday, 16:24:09","order":[0,1,2,3,4,5,6,7,8,9,10,11,12],"assembliesNames":["ABySS\_127","ABySS\_63","CLC","IDBA\_UD","MEGAHIT","MIRA","Ray\_Meta","SPAdes","SPAdes\_meta","SPAdes\_sc","SPAdes\_sc\_careful","Velvet","Geneious"]},{"assembliesWithNs":null,"minContig":500,"report":[["Genome statistics",[{"values":["3.310","10.706","4.623","6.753","73.753","5.252","3.338","5.843","8.693","7.356","5.409","2.257","67.154"],"quality":"More is better","isMain":true,"metricName":"Genome fraction (%)"},{"values":["1.000","1.006","1.000","1.000","1.074","1.000","1.000","1.052","1.000","1.000","1.000","1.000","1.043"],"quality":"Less is better","isMain":true,"metricName":"Duplication ratio"},{"values":[844,1740,1396,1743,24548,1548,893,1455,1524,1607,1575,947,23725],"quality":"More is better","isMain":true,"metricName":"Largest alignment"},{"values":[1389,4519,1940,2834,33006,2204,1401,2579,3649,3087,2270,947,29380],"quality":"More is better","isMain":true,"metricName":"Total aligned length"},{"values":[null,null,null,null,46439,null,null,null,null,null,null,null,92482],"quality":"More is better","isMain":false,"metricName":"NG50"},{"values":[null,null,null,null,46439,null,null,null,null,null,null,null,92482],"quality":"More is better","isMain":false,"metricName":"NG75"},{"values":[844,1696,1396,1743,null,1548,893,1455,998,1607,1575,947,null],"quality":"More is better","isMain":false,"metricName":"NA50"},{"values":[545,null,544,586,null,656,508,610,null,785,695,947,null],"quality":"More is better","isMain":false,"metricName":"NA75"},{"values":[null,null,null,null,24548,null,null,null,null,null,null,null,23725],"quality":"More is better","isMain":true,"metricName":"NGA50"},{"values":[null,null,null,null,677,null,null,null,null,null,null,null,null],"quality":"More is better","isMain":false,"metricName":"NGA75"},{"values":[null,null,null,null,1,null,null,null,null,null,null,null,1],"quality":"Less is better","isMain":false,"metricName":"LG50"},{"values":[null,null,null,null,1,null,null,null,null,null,null,null,1],"quality":"Less is better","isMain":false,"metricName":"LG75"},{"values":[1,2,1,1,null,1,1,1,2,1,1,1,null],"quality":"Less is better","isMain":false,"metricName":"LA50"},{"values":[2,null,2,2,null,2,2,2,null,2,2,1,null],"quality":"Less is better","isMain":false,"metricName":"LA75"},{"values":[null,null,null,null,1,null,null,null,null,null,null,null,1],"quality":"Less is better","isMain":true,"metricName":"LGA50"},{"values":[null,null,null,null,7,null,null,null,null,null,null,null,null],"quality":"Less is better","isMain":false,"metricName":"LGA75"}]],["Misassemblies",[{"values":[0,1,0,0,0,0,0,0,1,0,0,0,0],"quality":"Less is better","isMain":true,"metricName":"# misassemblies"},{"values":[0,1,0,0,0,0,0,0,1,0,0,0,0],"quality":"Less is better","isMain":false,"metricName":" # relocations"},{"values":[0,0,0,0,0,0,0,0,0,0,0,0,0],"quality":"Less is better","isMain":false,"metricName":" # translocations"},{"values":[0,0,0,0,0,0,0,0,0,0,0,0,0],"quality":"Less is better","isMain":false,"metricName":" # inversions"},{"values":[0,1,0,0,0,0,0,0,1,0,0,0,0],"quality":"Less is better","isMain":false,"metricName":"# misassembled contigs"},{"values":[0,3606,0,0,0,0,0,0,3306,0,0,0,0],"quality":"Less is better","isMain":true,"metricName":"Misassembled contigs length"},{"values":[0,0,0,0,2,0,0,0,1,0,0,0,0],"quality":"Less is better","isMain":false,"metricName":"# local misassemblies"},{"values":[0,0,0,0,1,0,0,0,0,0,0,0,1],"quality":"Less is better","isMain":false,"metricName":"# unaligned mis. contigs"}]],["Unaligned",[{"values":[0,0,0,0,0,0,0,0,0,0,0,0,0],"quality":"Less is better","isMain":false,"metricName":"# fully unaligned contigs"},{"values":[0,0,0,0,0,0,0,0,0,0,0,0,0],"quality":"Less is better","isMain":false,"metricName":"Fully unaligned length"},{"values":[0,1,0,0,2,0,0,0,1,0,0,0,2],"quality":"Less is better","isMain":false,"metricName":"# partially unaligned contigs"},{"values":[0,1671,0,0,62148,0,0,0,1287,0,0,0,65313],"quality":"Less is better","isMain":false,"metricName":"Partially unaligned length"}]],["Mismatches",[{"values":[0,3,0,7,129,0,0,1,23,16,18,0,152],"quality":"Less is better","isMain":false,"metricName":"# mismatches"},{"values":[0,0,0,0,10,0,0,0,1,0,0,0,12],"quality":"Less is better","isMain":false,"metricName":"# indels"},{"values":[0,0,0,0,10,0,0,0,1,0,0,0,12],"quality":"Less is better","isMain":false,"metricName":"Indels length"},{"values":["0.00","66.77","0.00","247.00","416.79","0.00","0.00","40.78","630.48","518.30","792.95","0.00","539.35"],"quality":"Less is better","isMain":true,"metricName":"# mismatches per 100 kbp"},{"values":["0.00","0.00","0.00","0.00","32.31","0.00","0.00","0.00","27.41","0.00","0.00","0.00","42.58"],"quality":"Less is better","isMain":true,"metricName":"# indels per 100 kbp"},{"values":[0,0,0,0,10,0,0,0,1,0,0,0,12],"quality":"Less is better","isMain":false,"metricName":" # indels (<= 5 bp)"},{"values":[0,0,0,0,0,0,0,0,0,0,0,0,0],"quality":"Less is better","isMain":false,"metricName":" # indels (> 5 bp)"},{"values":[0,50,0,0,10,0,0,0,0,0,0,0,573],"quality":"Less is better","isMain":false,"metricName":"# N's"},{"values":["0.00","807.75","0.00","0.00","10.48","0.00","0.00","0.00","0.00","0.00","0.00","0.00","605.11"],"quality":"Less is better","isMain":true,"metricName":"# N's per 100 kbp"}]],["Statistics without reference",[{"values":[2,3,2,3,7,2,2,3,3,3,2,1,2],"quality":"Equal","isMain":true,"metricName":"# contigs"},{"values":[0,2,1,1,5,1,0,1,1,1,1,0,2],"quality":"Equal","isMain":false,"metricName":"# contigs (>= 1000 bp)"},{"values":[0,0,0,0,2,0,0,0,0,0,0,0,1],"quality":"Equal","isMain":false,"metricName":"# contigs (>= 5000 bp)"},{"values":[0,0,0,0,2,0,0,0,0,0,0,0,1],"quality":"Equal","isMain":false,"metricName":"# contigs (>= 10000 bp)"},{"values":[0,0,0,0,2,0,0,0,0,0,0,0,1],"quality":"Equal","isMain":false,"metricName":"# contigs (>= 25000 bp)"},{"values":[0,0,0,0,0,0,0,0,0,0,0,0,1],"quality":"Equal","isMain":false,"metricName":"# contigs (>= 50000 bp)"},{"values":[844,3606,1396,1743,46439,1548,893,1455,3306,1607,1575,947,92482],"quality":"More is better","isMain":true,"metricName":"Largest contig"},{"values":[1389,6190,1940,2834,95400,2204,1401,2579,4936,3087,2270,947,94693],"quality":"More is better","isMain":true,"metricName":"Total length"},{"values":[0,5346,1396,1743,94091,1548,0,1455,3306,1607,1575,0,94693],"quality":"More is better","isMain":true,"metricName":"Total length (>= 1000 bp)"},{"values":[0,0,0,0,90324,0,0,0,0,0,0,0,92482],"quality":"More is better","isMain":false,"metricName":"Total length (>= 5000 bp)"},{"values":[0,0,0,0,90324,0,0,0,0,0,0,0,92482],"quality":"More is better","isMain":true,"metricName":"Total length (>= 10000 bp)"},{"values":[0,0,0,0,90324,0,0,0,0,0,0,0,92482],"quality":"More is better","isMain":false,"metricName":"Total length (>= 25000 bp)"},{"values":[0,0,0,0,0,0,0,0,0,0,0,0,92482],"quality":"More is better","isMain":true,"metricName":"Total length (>= 50000 bp)"},{"values":[844,3606,1396,1743,43885,1548,893,1455,3306,1607,1575,947,92482],"quality":"More is better","isMain":false,"metricName":"N50"},{"values":[545,1740,544,586,43885,656,508,610,998,785,695,947,92482],"quality":"More is better","isMain":false,"metricName":"N75"},{"values":[1,1,1,1,2,1,1,1,1,1,1,1,1],"quality":"Less is better","isMain":false,"metricName":"L50"},{"values":[2,2,2,2,2,2,2,2,2,2,2,1,1],"quality":"Less is better","isMain":false,"metricName":"L75"},{"values":["29.81","33.13","32.16","32.46","33.76","31.35","31.83","31.33","32.74","32.49","32.47","32.21","33.87"],"quality":"Equal","isMain":false,"metricName":"GC (%)"}]],["Predicted genes",[]],["Similarity statistics",[{"values":[0,0,0,0,0,0,0,0,0,0,0,0,0],"quality":"Equal","isMain":false,"metricName":"# similar correct contigs"},{"values":[0,0,0,0,0,0,0,0,0,0,0,0,0],"quality":"Equal","isMain":false,"metricName":"# similar misassembled blocks"}]],["Reference statistics",[{"values":[41966,41966,41966,41966,41966,41966,41966,41966,41966,41966,41966,41966,41966],"quality":"Equal","isMain":false,"metricName":"Reference length"},{"values":[1,1,1,1,1,1,1,1,1,1,1,1,1],"quality":"Equal","isMain":false,"metricName":"Reference fragments"},{"values":["33.46","33.46","33.46","33.46","33.46","33.46","33.46","33.46","33.46","33.46","33.46","33.46","33.46"],"quality":"Equal","isMain":false,"metricName":"Reference GC (%)"}]]],"referenceName":"gi\_157102938\_ref\_NC","date":"18 July 2018, Wednesday, 16:24:20","order":[0,1,2,3,4,5,6,7,8,9,10,11,12],"assembliesNames":["ABySS\_127","ABySS\_63","IDBA\_UD","MEGAHIT","MIRA","Ray\_Meta","SOAPdenovo2","SPAdes","SPAdes\_meta","SPAdes\_sc","SPAdes\_sc\_careful","Velvet","Geneious"]},{"assembliesWithNs":null,"minContig":500,"report":[["Genome statistics",[{"values":["5.422","8.226","2.227","10.098","14.733","11.489","24.382","8.404","8.469"],"quality":"More is better","isMain":true,"metricName":"Genome fraction (%)"},{"values":["1.000","1.000","1.000","1.007","1.000","1.106","1.008","1.006","1.006"],"quality":"Less is better","isMain":true,"metricName":"Duplication ratio"},{"values":[981,1446,688,930,1554,946,1604,1029,1029],"quality":"More is better","isMain":true,"metricName":"Largest alignment"},{"values":[4565,6926,1875,8503,12405,9679,20700,7118,7173],"quality":"More is better","isMain":true,"metricName":"Total aligned length"},{"values":[638,1027,615,646,1133,null,710,688,688],"quality":"More is better","isMain":false,"metricName":"NA50"},{"values":[582,614,572,570,809,null,581,659,659],"quality":"More is better","isMain":false,"metricName":"NA75"},{"values":[null,null,null,null,null,null,null,null,null],"quality":"More is better","isMain":true,"metricName":"NGA50"},{"values":[4,3,2,6,5,null,10,5,5],"quality":"Less is better","isMain":false,"metricName":"LA50"},{"values":[5,6,3,10,8,null,18,7,7],"quality":"Less is better","isMain":false,"metricName":"LA75"}]],["Misassemblies",[{"values":[0,0,0,0,0,0,0,0,0],"quality":"Less is better","isMain":true,"metricName":"# misassemblies"},{"values":[0,0,0,0,0,0,0,0,0],"quality":"Less is better","isMain":false,"metricName":" # relocations"},{"values":[0,0,0,0,0,0,0,0,0],"quality":"Less is better","isMain":false,"metricName":" # translocations"},{"values":[0,0,0,0,0,0,0,0,0],"quality":"Less is better","isMain":false,"metricName":" # inversions"},{"values":[0,0,0,0,0,0,0,0,0],"quality":"Less is better","isMain":false,"metricName":"# misassembled contigs"},{"values":[0,0,0,0,0,0,0,0,0],"quality":"Less is better","isMain":true,"metricName":"Misassembled contigs length"},{"values":[0,0,0,0,0,1,0,0,0],"quality":"Less is better","isMain":false,"metricName":"# local misassemblies"},{"values":[0,0,0,0,0,4,0,0,0],"quality":"Less is better","isMain":false,"metricName":"# unaligned mis. contigs"}]],["Unaligned",[{"values":[0,0,0,0,0,0,0,0,0],"quality":"Less is better","isMain":false,"metricName":"# fully unaligned contigs"},{"values":[0,0,0,0,0,0,0,0,0],"quality":"Less is better","isMain":false,"metricName":"Fully unaligned length"},{"values":[0,0,0,0,0,4,0,0,0],"quality":"Less is better","isMain":false,"metricName":"# partially unaligned contigs"},{"values":[0,0,0,0,0,16989,0,0,0],"quality":"Less is better","isMain":false,"metricName":"Partially unaligned length"}]],["Mismatches",[{"values":[1,42,1,10,4,101,50,12,12],"quality":"Less is better","isMain":false,"metricName":"# mismatches"},{"values":[0,0,0,0,0,2,0,0,0],"quality":"Less is better","isMain":false,"metricName":"# indels"},{"values":[0,0,0,0,0,5,0,0,0],"quality":"Less is better","isMain":false,"metricName":"Indels length"},{"values":["21.91","606.41","53.33","117.61","32.25","1044.04","243.55","169.59","168.28"],"quality":"Less is better","isMain":true,"metricName":"# mismatches per 100 kbp"},{"values":["0.00","0.00","0.00","0.00","0.00","20.67","0.00","0.00","0.00"],"quality":"Less is better","isMain":true,"metricName":"# indels per 100 kbp"},{"values":[0,0,0,0,0,2,0,0,0],"quality":"Less is better","isMain":false,"metricName":" # indels (<= 5 bp)"},{"values":[0,0,0,0,0,0,0,0,0],"quality":"Less is better","isMain":false,"metricName":" # indels (> 5 bp)"},{"values":[0,0,0,0,14,7537,0,0,0],"quality":"Less is better","isMain":false,"metricName":"# N's"},{"values":["0.00","0.00","0.00","0.00","112.86","27218.23","0.00","0.00","0.00"],"quality":"Less is better","isMain":true,"metricName":"# N's per 100 kbp"}]],["Statistics without reference",[{"values":[7,8,3,13,12,9,27,10,10],"quality":"Equal","isMain":true,"metricName":"# contigs"},{"values":[0,3,0,0,6,6,4,1,1],"quality":"Equal","isMain":false,"metricName":"# contigs (>= 1000 bp)"},{"values":[0,0,0,0,0,2,0,0,0],"quality":"Equal","isMain":false,"metricName":"# contigs (>= 5000 bp)"},{"values":[0,0,0,0,0,1,0,0,0],"quality":"Equal","isMain":false,"metricName":"# contigs (>= 10000 bp)"},{"values":[0,0,0,0,0,0,0,0,0],"quality":"Equal","isMain":false,"metricName":"# contigs (>= 25000 bp)"},{"values":[0,0,0,0,0,0,0,0,0],"quality":"Equal","isMain":false,"metricName":"# contigs (>= 50000 bp)"},{"values":[981,1446,688,930,1554,11292,1604,1029,1029],"quality":"More is better","isMain":true,"metricName":"Largest contig"},{"values":[4565,6926,1875,8565,12405,27691,20703,7118,7174],"quality":"More is better","isMain":true,"metricName":"Total length"},{"values":[0,3787,0,0,7677,25786,5672,1029,1029],"quality":"More is better","isMain":true,"metricName":"Total length (>= 1000 bp)"},{"values":[0,0,0,0,0,18427,0,0,0],"quality":"More is better","isMain":false,"metricName":"Total length (>= 5000 bp)"},{"values":[0,0,0,0,0,11292,0,0,0],"quality":"More is better","isMain":true,"metricName":"Total length (>= 10000 bp)"},{"values":[0,0,0,0,0,0,0,0,0],"quality":"More is better","isMain":false,"metricName":"Total length (>= 25000 bp)"},{"values":[0,0,0,0,0,0,0,0,0],"quality":"More is better","isMain":true,"metricName":"Total length (>= 50000 bp)"},{"values":[638,1027,615,646,1133,7135,710,688,688],"quality":"More is better","isMain":false,"metricName":"N50"},{"values":[582,614,572,570,809,2961,581,659,659],"quality":"More is better","isMain":false,"metricName":"N75"},{"values":[4,3,2,6,5,2,10,5,5],"quality":"Less is better","isMain":false,"metricName":"L50"},{"values":[5,6,3,10,8,3,18,7,7],"quality":"Less is better","isMain":false,"metricName":"L75"},{"values":["60.42","56.35","56.16","57.47","60.32","54.83","57.74","52.50","52.56"],"quality":"Equal","isMain":false,"metricName":"GC (%)"}]],["Predicted genes",[]],["Similarity statistics",[{"values":[0,0,0,0,0,0,0,0,0],"quality":"Equal","isMain":false,"metricName":"# similar correct contigs"},{"values":[0,0,0,0,0,0,0,0,0],"quality":"Equal","isMain":false,"metricName":"# similar misassembled blocks"}]],["Reference statistics",[{"values":[84201,84201,84201,84201,84201,84201,84201,84201,84201],"quality":"Equal","isMain":false,"metricName":"Reference length"},{"values":[1,1,1,1,1,1,1,1,1],"quality":"Equal","isMain":false,"metricName":"Reference fragments"},{"values":["57.76","57.76","57.76","57.76","57.76","57.76","57.76","57.76","57.76"],"quality":"Equal","isMain":false,"metricName":"Reference GC (%)"}]]],"referenceName":"gi\_157265294\_ref\_NC","date":"18 July 2018, Wednesday, 16:24:29","order":[0,1,2,3,4,5,6,7,8],"assembliesNames":["ABySS\_127","CLC","IDBA\_UD","MEGAHIT","MIRA","SOAPdenovo2","SPAdes","SPAdes\_sc","SPAdes\_sc\_careful"]},{"assembliesWithNs":null,"minContig":500,"report":[["Genome statistics",[{"values":["100.000","99.222","99.349","99.782","100.000","100.000","22.968","42.745","100.000","100.000","99.813","95.280","97.452","100.000"],"quality":"More is better","isMain":true,"metricName":"Genome fraction (%)"},{"values":["1.008","1.007","1.002","1.002","1.018","1.016","1.001","1.084","1.002","1.001","1.007","1.013","1.000","1.016"],"quality":"Less is better","isMain":true,"metricName":"Duplication ratio"},{"values":[47760,16337,52516,31652,26811,58314,1091,7780,83446,83374,32460,12855,18419,58314],"quality":"More is better","isMain":true,"metricName":"Largest alignment"},{"values":[83949,83291,82926,83335,84838,84661,19150,35824,83446,83374,83708,79991,81226,84661],"quality":"More is better","isMain":true,"metricName":"Total aligned length"},{"values":[47760,10069,52516,26111,16274,84661,null,5963,83446,83374,21337,6910,11091,84661],"quality":"More is better","isMain":false,"metricName":"NG50"},{"values":[23814,7640,16452,18152,9609,84661,null,2431,83446,83374,9940,2719,3794,84661],"quality":"More is better","isMain":false,"metricName":"NG75"},{"values":[47760,10069,52516,26111,16274,58314,629,null,83446,83374,21337,6456,11091,58314],"quality":"More is better","isMain":false,"metricName":"NA50"},{"values":[23814,7640,16452,18152,9609,26347,546,null,83446,83374,9940,2984,3794,26347],"quality":"More is better","isMain":false,"metricName":"NA75"},{"values":[47760,10069,52516,26111,16274,58314,null,null,83446,83374,21337,6456,11091,58314],"quality":"More is better","isMain":true,"metricName":"NGA50"},{"values":[23814,7640,16452,18152,9609,26347,null,null,83446,83374,9940,2719,3794,26347],"quality":"More is better","isMain":false,"metricName":"NGA75"},{"values":[1,3,1,2,2,1,null,5,1,1,2,5,3,1],"quality":"Less is better","isMain":false,"metricName":"LG50"},{"values":[2,6,2,3,4,1,null,11,1,1,3,10,7,1],"quality":"Less is better","isMain":false,"metricName":"LG75"},{"values":[1,3,1,2,2,1,12,null,1,1,2,5,3,1],"quality":"Less is better","isMain":false,"metricName":"LA50"},{"values":[2,6,2,3,4,2,20,null,1,1,3,9,7,2],"quality":"Less is better","isMain":false,"metricName":"LA75"},{"values":[1,3,1,2,2,1,null,null,1,1,2,5,3,1],"quality":"Less is better","isMain":true,"metricName":"LGA50"},{"values":[2,6,2,3,4,2,null,null,1,1,3,10,7,2],"quality":"Less is better","isMain":false,"metricName":"LGA75"}]],["Misassemblies",[{"values":[0,0,0,0,0,1,0,0,0,0,0,0,0,1],"quality":"Less is better","isMain":true,"metricName":"# misassemblies"},{"values":[0,0,0,0,0,1,0,0,0,0,0,0,0,1],"quality":"Less is better","isMain":false,"metricName":" # relocations"},{"values":[0,0,0,0,0,0,0,0,0,0,0,0,0,0],"quality":"Less is better","isMain":false,"metricName":" # translocations"},{"values":[0,0,0,0,0,0,0,0,0,0,0,0,0,0],"quality":"Less is better","isMain":false,"metricName":" # inversions"},{"values":[0,0,0,0,0,1,0,0,0,0,0,0,0,1],"quality":"Less is better","isMain":false,"metricName":"# misassembled contigs"},{"values":[0,0,0,0,0,84661,0,0,0,0,0,0,0,84661],"quality":"Less is better","isMain":true,"metricName":"Misassembled contigs length"},{"values":[0,0,0,0,0,0,0,4,1,0,0,0,0,0],"quality":"Less is better","isMain":false,"metricName":"# local misassemblies"},{"values":[0,0,0,0,0,0,0,9,0,0,0,0,0,0],"quality":"Less is better","isMain":false,"metricName":"# unaligned mis. contigs"}]],["Unaligned",[{"values":[0,0,0,0,0,0,0,0,0,0,0,0,0,0],"quality":"Less is better","isMain":false,"metricName":"# fully unaligned contigs"},{"values":[0,0,0,0,0,0,0,0,0,0,0,0,0,0],"quality":"Less is better","isMain":false,"metricName":"Fully unaligned length"},{"values":[0,0,0,0,0,0,0,14,0,0,0,0,0,0],"quality":"Less is better","isMain":false,"metricName":"# partially unaligned contigs"},{"values":[0,0,0,0,0,0,0,36179,0,0,0,0,0,0],"quality":"Less is better","isMain":false,"metricName":"Partially unaligned length"}]],["Mismatches",[{"values":[0,0,0,0,57,0,0,1,0,2,0,12,3,0],"quality":"Less is better","isMain":false,"metricName":"# mismatches"},{"values":[0,0,0,0,0,0,0,41,0,1,0,0,0,0],"quality":"Less is better","isMain":false,"metricName":"# indels"},{"values":[0,0,0,0,0,0,0,1970,0,55,0,0,0,0],"quality":"Less is better","isMain":false,"metricName":"Indels length"},{"values":["0.00","0.00","0.00","0.00","68.41","0.00","0.00","2.81","0.00","2.40","0.00","15.12","3.69","0.00"],"quality":"Less is better","isMain":true,"metricName":"# mismatches per 100 kbp"},{"values":["0.00","0.00","0.00","0.00","0.00","0.00","0.00","115.12","0.00","1.20","0.00","0.00","0.00","0.00"],"quality":"Less is better","isMain":true,"metricName":"# indels per 100 kbp"},{"values":[0,0,0,0,0,0,0,0,0,0,0,0,0,0],"quality":"Less is better","isMain":false,"metricName":" # indels (<= 5 bp)"},{"values":[0,0,0,0,0,0,0,41,0,1,0,0,0,0],"quality":"Less is better","isMain":false,"metricName":" # indels (> 5 bp)"},{"values":[0,0,0,0,0,0,0,20524,0,0,0,453,0,404],"quality":"Less is better","isMain":false,"metricName":"# N's"},{"values":["0.00","0.00","0.00","0.00","0.00","0.00","0.00","27442.54","0.00","0.00","0.00","563.12","0.00","477.20"],"quality":"Less is better","isMain":true,"metricName":"# N's per 100 kbp"}]],["Statistics without reference",[{"values":[5,12,4,4,8,1,29,18,1,1,11,26,14,1],"quality":"Equal","isMain":true,"metricName":"# contigs"},{"values":[4,12,4,4,8,1,2,16,1,1,10,15,12,1],"quality":"Equal","isMain":false,"metricName":"# contigs (>= 1000 bp)"},{"values":[3,6,3,4,7,1,0,6,1,1,4,5,5,1],"quality":"Equal","isMain":false,"metricName":"# contigs (>= 5000 bp)"},{"values":[2,3,2,3,3,1,0,2,1,1,2,1,3,1],"quality":"Equal","isMain":false,"metricName":"# contigs (>= 10000 bp)"},{"values":[1,0,1,2,1,1,0,0,1,1,1,0,0,1],"quality":"Equal","isMain":false,"metricName":"# contigs (>= 25000 bp)"},{"values":[0,0,1,0,0,1,0,0,1,1,0,0,0,1],"quality":"Equal","isMain":false,"metricName":"# contigs (>= 50000 bp)"},{"values":[47760,16337,52516,31652,26811,84661,1091,11329,83446,83374,32460,12855,18419,84661],"quality":"More is better","isMain":true,"metricName":"Largest contig"},{"values":[83949,83291,82926,83335,84838,84661,19150,74789,83446,83374,83708,80445,81226,84661],"quality":"More is better","isMain":true,"metricName":"Total length"},{"values":[83274,83291,82926,83335,84838,84661,2100,73640,83446,83374,82798,72489,79803,84661],"quality":"More is better","isMain":true,"metricName":"Total length (>= 1000 bp)"},{"values":[80639,66772,78144,83335,82173,84661,0,47555,83446,83374,69617,45515,55880,84661],"quality":"More is better","isMain":false,"metricName":"Total length (>= 5000 bp)"},{"values":[71574,41779,68968,75915,54790,84661,0,22621,83446,83374,53797,12855,43709,84661],"quality":"More is better","isMain":true,"metricName":"Total length (>= 10000 bp)"},{"values":[47760,0,52516,57763,26811,84661,0,0,83446,83374,32460,0,0,84661],"quality":"More is better","isMain":false,"metricName":"Total length (>= 25000 bp)"},{"values":[0,0,52516,0,0,84661,0,0,83446,83374,0,0,0,84661],"quality":"More is better","isMain":true,"metricName":"Total length (>= 50000 bp)"},{"values":[47760,10069,52516,26111,16274,84661,629,5963,83446,83374,21337,6910,11091,84661],"quality":"More is better","isMain":false,"metricName":"N50"},{"values":[23814,7640,16452,18152,9609,84661,546,4486,83446,83374,9940,2984,3794,84661],"quality":"More is better","isMain":false,"metricName":"N75"},{"values":[1,3,1,2,2,1,12,5,1,1,2,5,3,1],"quality":"Less is better","isMain":false,"metricName":"L50"},{"values":[2,6,2,3,4,1,20,8,1,1,3,9,7,1],"quality":"Less is better","isMain":false,"metricName":"L75"},{"values":["57.78","57.72","57.80","57.74","57.80","57.81","57.51","57.89","57.76","57.76","57.77","57.78","57.76","57.83"],"quality":"Equal","isMain":false,"metricName":"GC (%)"}]],["Predicted genes",[]],["Similarity statistics",[{"values":[0,0,0,0,0,0,0,0,0,0,0,0,0,0],"quality":"Equal","isMain":false,"metricName":"# similar correct contigs"},{"values":[0,0,0,0,0,0,0,0,0,0,0,0,0,0],"quality":"Equal","isMain":false,"metricName":"# similar misassembled blocks"}]],["Reference statistics",[{"values":[83319,83319,83319,83319,83319,83319,83319,83319,83319,83319,83319,83319,83319,83319],"quality":"Equal","isMain":false,"metricName":"Reference length"},{"values":[1,1,1,1,1,1,1,1,1,1,1,1,1,1],"quality":"Equal","isMain":false,"metricName":"Reference fragments"},{"values":["57.76","57.76","57.76","57.76","57.76","57.76","57.76","57.76","57.76","57.76","57.76","57.76","57.76","57.76"],"quality":"Equal","isMain":false,"metricName":"Reference GC (%)"}]]],"referenceName":"gi\_157265412\_ref\_NC","date":"18 July 2018, Wednesday, 16:24:41","order":[0,1,2,3,4,5,6,7,8,9,10,11,12,13],"assembliesNames":["ABySS\_127","ABySS\_63","CLC","IDBA\_UD","MEGAHIT","MIRA","Ray\_Meta","SOAPdenovo2","SPAdes","SPAdes\_meta","SPAdes\_sc","SPAdes\_sc\_careful","Velvet","Geneious"]},{"assembliesWithNs":null,"minContig":500,"report":[["Genome statistics",[{"values":["75.868","60.400","96.874","97.377","96.302","89.341","94.915","98.335","96.357","97.950","97.964","85.637","49.091"],"quality":"More is better","isMain":true,"metricName":"Genome fraction (%)"},{"values":["1.001","1.000","1.010","1.004","1.000","1.025","1.009","1.001","1.001","1.001","1.001","1.001","1.000"],"quality":"Less is better","isMain":true,"metricName":"Duplication ratio"},{"values":[2846,3087,11894,12180,12181,11425,12326,18697,17438,17834,17851,10569,11795],"quality":"More is better","isMain":true,"metricName":"Largest alignment"},{"values":[53593,42640,69026,69008,67980,64634,67156,69446,68051,69204,69221,60505,34648],"quality":"More is better","isMain":true,"metricName":"Total aligned length"},{"values":[1159,716,7952,5489,8177,2700,4576,10573,10573,10573,10573,4429,null],"quality":"More is better","isMain":false,"metricName":"NG50"},{"values":[533,null,2433,3987,3897,1540,2433,4429,4429,4429,4429,1300,null],"quality":"More is better","isMain":false,"metricName":"NG75"},{"values":[1381,1173,7952,5489,8177,3366,4576,10573,10573,10573,10573,4763,9269],"quality":"More is better","isMain":false,"metricName":"NA50"},{"values":[1032,892,2433,4429,3897,1703,3987,10569,4429,4429,4429,2262,3682],"quality":"More is better","isMain":false,"metricName":"NA75"},{"values":[1159,716,7952,5489,8177,2700,4576,10573,10573,10573,10573,4429,null],"quality":"More is better","isMain":true,"metricName":"NGA50"},{"values":[533,null,2433,3987,3897,1540,2433,4429,4429,4429,4429,1300,null],"quality":"More is better","isMain":false,"metricName":"NGA75"},{"values":[20,27,4,4,4,7,5,3,3,3,3,5,null],"quality":"Less is better","isMain":false,"metricName":"LG50"},{"values":[41,null,8,8,7,15,10,5,5,5,5,12,null],"quality":"Less is better","isMain":false,"metricName":"LG75"},{"values":[13,12,4,4,4,6,5,3,3,3,3,4,2],"quality":"Less is better","isMain":false,"metricName":"LA50"},{"values":[24,23,8,7,7,13,9,4,5,5,5,8,4],"quality":"Less is better","isMain":false,"metricName":"LA75"},{"values":[20,27,4,4,4,7,5,3,3,3,3,5,null],"quality":"Less is better","isMain":true,"metricName":"LGA50"},{"values":[41,null,8,8,7,15,10,5,5,5,5,12,null],"quality":"Less is better","isMain":false,"metricName":"LGA75"}]],["Misassemblies",[{"values":[0,0,0,0,0,0,0,0,0,0,0,0,0],"quality":"Less is better","isMain":true,"metricName":"# misassemblies"},{"values":[0,0,0,0,0,0,0,0,0,0,0,0,0],"quality":"Less is better","isMain":false,"metricName":" # relocations"},{"values":[0,0,0,0,0,0,0,0,0,0,0,0,0],"quality":"Less is better","isMain":false,"metricName":" # translocations"},{"values":[0,0,0,0,0,0,0,0,0,0,0,0,0],"quality":"Less is better","isMain":false,"metricName":" # inversions"},{"values":[0,0,0,0,0,0,0,0,0,0,0,0,0],"quality":"Less is better","isMain":false,"metricName":"# misassembled contigs"},{"values":[0,0,0,0,0,0,0,0,0,0,0,0,0],"quality":"Less is better","isMain":true,"metricName":"Misassembled contigs length"},{"values":[0,0,0,1,0,0,0,2,0,4,4,0,0],"quality":"Less is better","isMain":false,"metricName":"# local misassemblies"},{"values":[0,0,0,0,0,0,0,0,0,0,0,0,0],"quality":"Less is better","isMain":false,"metricName":"# unaligned mis. contigs"}]],["Unaligned",[{"values":[0,0,0,0,0,0,0,0,0,0,0,0,0],"quality":"Less is better","isMain":false,"metricName":"# fully unaligned contigs"},{"values":[0,0,0,0,0,0,0,0,0,0,0,0,0],"quality":"Less is better","isMain":false,"metricName":"Fully unaligned length"},{"values":[0,0,0,0,0,0,0,0,0,0,0,0,0],"quality":"Less is better","isMain":false,"metricName":"# partially unaligned contigs"},{"values":[0,0,0,0,0,0,0,0,0,0,0,0,0],"quality":"Less is better","isMain":false,"metricName":"Partially unaligned length"}]],["Mismatches",[{"values":[9,1,19,14,10,11,13,30,16,25,9,9,3],"quality":"Less is better","isMain":false,"metricName":"# mismatches"},{"values":[0,0,0,2,0,0,6,1,1,4,1,0,0],"quality":"Less is better","isMain":false,"metricName":"# indels"},{"values":[0,0,0,18,0,0,254,1,40,65,10,0,0],"quality":"Less is better","isMain":false,"metricName":"Indels length"},{"values":["16.81","2.35","27.79","20.37","14.71","17.44","19.41","43.23","23.53","36.16","13.02","14.89","8.66"],"quality":"Less is better","isMain":true,"metricName":"# mismatches per 100 kbp"},{"values":["0.00","0.00","0.00","2.91","0.00","0.00","8.96","1.44","1.47","5.79","1.45","0.00","0.00"],"quality":"Less is better","isMain":true,"metricName":"# indels per 100 kbp"},{"values":[0,0,0,0,0,0,0,1,0,2,0,0,0],"quality":"Less is better","isMain":false,"metricName":" # indels (<= 5 bp)"},{"values":[0,0,0,2,0,0,6,0,1,2,1,0,0],"quality":"Less is better","isMain":false,"metricName":" # indels (> 5 bp)"},{"values":[0,0,0,0,0,37,564,0,0,20,0,0,30],"quality":"Less is better","isMain":false,"metricName":"# N's"},{"values":["0.00","0.00","0.00","0.00","0.00","57.25","834.76","0.00","0.00","28.90","0.00","0.00","86.59"],"quality":"Less is better","isMain":true,"metricName":"# N's per 100 kbp"}]],["Statistics without reference",[{"values":[42,38,18,16,15,27,20,11,11,12,12,20,6],"quality":"Equal","isMain":true,"metricName":"# contigs"},{"values":[25,16,15,14,13,19,16,10,10,10,10,13,6],"quality":"Equal","isMain":false,"metricName":"# contigs (>= 1000 bp)"},{"values":[0,0,4,4,4,4,4,4,4,4,4,3,2],"quality":"Equal","isMain":false,"metricName":"# contigs (>= 5000 bp)"},{"values":[0,0,2,3,3,1,1,4,4,4,4,2,1],"quality":"Equal","isMain":false,"metricName":"# contigs (>= 10000 bp)"},{"values":[0,0,0,0,0,0,0,0,0,0,0,0,0],"quality":"Equal","isMain":false,"metricName":"# contigs (>= 25000 bp)"},{"values":[0,0,0,0,0,0,0,0,0,0,0,0,0],"quality":"Equal","isMain":false,"metricName":"# contigs (>= 50000 bp)"},{"values":[2846,3087,11894,12180,12181,11425,12326,18697,17438,17834,17851,10569,11795],"quality":"More is better","isMain":true,"metricName":"Largest contig"},{"values":[53593,42640,69027,69013,67980,64634,67564,69446,68051,69204,69221,60505,34648],"quality":"More is better","isMain":true,"metricName":"Total length"},{"values":[41348,25871,66652,67401,66193,58657,64243,68543,67148,67680,67697,55284,34648],"quality":"More is better","isMain":true,"metricName":"Total length (>= 1000 bp)"},{"values":[0,0,40278,38811,41324,27833,30973,52184,50789,51321,51338,26638,21064],"quality":"More is better","isMain":false,"metricName":"Total length (>= 5000 bp)"},{"values":[0,0,22467,33322,33147,11425,12326,52184,50789,51321,51338,21101,11795],"quality":"More is better","isMain":true,"metricName":"Total length (>= 10000 bp)"},{"values":[0,0,0,0,0,0,0,0,0,0,0,0,0],"quality":"More is better","isMain":false,"metricName":"Total length (>= 25000 bp)"},{"values":[0,0,0,0,0,0,0,0,0,0,0,0,0],"quality":"More is better","isMain":true,"metricName":"Total length (>= 50000 bp)"},{"values":[1381,1173,7952,5489,8177,3366,4576,10573,10573,10573,10573,4763,9269],"quality":"More is better","isMain":false,"metricName":"N50"},{"values":[1032,892,2433,4429,3897,1703,3987,10569,4429,4429,4429,2262,3682],"quality":"More is better","isMain":false,"metricName":"N75"},{"values":[13,12,4,4,4,6,5,3,3,3,3,4,2],"quality":"Less is better","isMain":false,"metricName":"L50"},{"values":[24,23,8,7,7,13,9,4,5,5,5,8,4],"quality":"Less is better","isMain":false,"metricName":"L75"},{"values":["50.96","50.69","50.97","50.95","50.94","50.93","51.00","50.95","50.91","50.94","50.94","50.97","50.14"],"quality":"Equal","isMain":false,"metricName":"GC (%)"}]],["Predicted genes",[]],["Similarity statistics",[{"values":[0,0,2,3,3,0,1,3,3,3,3,2,0],"quality":"Equal","isMain":false,"metricName":"# similar correct contigs"},{"values":[0,0,0,0,0,0,0,0,0,0,0,0,0],"quality":"Equal","isMain":false,"metricName":"# similar misassembled blocks"}]],["Reference statistics",[{"values":[70579,70579,70579,70579,70579,70579,70579,70579,70579,70579,70579,70579,70579],"quality":"Equal","isMain":false,"metricName":"Reference length"},{"values":[1,1,1,1,1,1,1,1,1,1,1,1,1],"quality":"Equal","isMain":false,"metricName":"Reference fragments"},{"values":["50.92","50.92","50.92","50.92","50.92","50.92","50.92","50.92","50.92","50.92","50.92","50.92","50.92"],"quality":"Equal","isMain":false,"metricName":"Reference GC (%)"}]]],"referenceName":"gi\_157310906\_ref\_NC","date":"18 July 2018, Wednesday, 16:24:53","order":[0,1,2,3,4,5,6,7,8,9,10,11,12],"assembliesNames":["ABySS\_127","ABySS\_63","CLC","IDBA\_UD","MEGAHIT","MIRA","SOAPdenovo2","SPAdes","SPAdes\_meta","SPAdes\_sc","SPAdes\_sc\_careful","Velvet","Geneious"]},{"assembliesWithNs":null,"minContig":500,"report":[["Genome statistics",[{"values":["100.000","98.172","100.000","98.845","100.000","100.000","100.000","25.525","100.000","99.360","99.291","99.292","95.743","100.000"],"quality":"More is better","isMain":true,"metricName":"Genome fraction (%)"},{"values":["1.002","1.000","1.000","1.001","1.006","1.009","1.004","1.000","1.002","1.000","1.001","1.001","1.001","1.004"],"quality":"Less is better","isMain":true,"metricName":"Duplication ratio"},{"values":[77806,77785,79232,75372,76862,79333,79570,1237,77808,78726,77778,77161,33429,79526],"quality":"More is better","isMain":true,"metricName":"Largest alignment"},{"values":[79398,77785,79232,78416,79740,79929,79570,20224,79402,78726,78726,78781,75897,79526],"quality":"More is better","isMain":true,"metricName":"Total aligned length"},{"values":[77806,77785,79232,75372,76862,79333,79570,null,77808,78726,77778,77161,21633,79526],"quality":"More is better","isMain":false,"metricName":"NG50"},{"values":[77806,77785,79232,75372,76862,79333,79570,null,77808,78726,77778,77161,11275,79526],"quality":"More is better","isMain":false,"metricName":"NG75"},{"values":[77806,77785,79232,75372,76862,79333,79570,729,77808,78726,77778,77161,21633,79526],"quality":"More is better","isMain":false,"metricName":"NA50"},{"values":[77806,77785,79232,75372,76862,79333,79570,621,77808,78726,77778,77161,11275,79526],"quality":"More is better","isMain":false,"metricName":"NA75"},{"values":[77806,77785,79232,75372,76862,79333,79570,null,77808,78726,77778,77161,21633,79526],"quality":"More is better","isMain":true,"metricName":"NGA50"},{"values":[77806,77785,79232,75372,76862,79333,79570,null,77808,78726,77778,77161,11275,79526],"quality":"More is better","isMain":false,"metricName":"NGA75"},{"values":[1,1,1,1,1,1,1,null,1,1,1,1,2,1],"quality":"Less is better","isMain":false,"metricName":"LG50"},{"values":[1,1,1,1,1,1,1,null,1,1,1,1,3,1],"quality":"Less is better","isMain":false,"metricName":"LG75"},{"values":[1,1,1,1,1,1,1,12,1,1,1,1,2,1],"quality":"Less is better","isMain":false,"metricName":"LA50"},{"values":[1,1,1,1,1,1,1,19,1,1,1,1,3,1],"quality":"Less is better","isMain":false,"metricName":"LA75"},{"values":[1,1,1,1,1,1,1,null,1,1,1,1,2,1],"quality":"Less is better","isMain":true,"metricName":"LGA50"},{"values":[1,1,1,1,1,1,1,null,1,1,1,1,3,1],"quality":"Less is better","isMain":false,"metricName":"LGA75"}]],["Misassemblies",[{"values":[0,0,0,0,0,0,0,0,0,0,0,0,0,0],"quality":"Less is better","isMain":true,"metricName":"# misassemblies"},{"values":[0,0,0,0,0,0,0,0,0,0,0,0,0,0],"quality":"Less is better","isMain":false,"metricName":" # relocations"},{"values":[0,0,0,0,0,0,0,0,0,0,0,0,0,0],"quality":"Less is better","isMain":false,"metricName":" # translocations"},{"values":[0,0,0,0,0,0,0,0,0,0,0,0,0,0],"quality":"Less is better","isMain":false,"metricName":" # inversions"},{"values":[0,0,0,0,0,0,0,0,0,0,0,0,0,0],"quality":"Less is better","isMain":false,"metricName":"# misassembled contigs"},{"values":[0,0,0,0,0,0,0,0,0,0,0,0,0,0],"quality":"Less is better","isMain":true,"metricName":"Misassembled contigs length"},{"values":[0,0,0,0,0,1,1,0,0,0,0,0,0,1],"quality":"Less is better","isMain":false,"metricName":"# local misassemblies"},{"values":[0,0,0,0,0,0,0,0,0,0,0,0,0,0],"quality":"Less is better","isMain":false,"metricName":"# unaligned mis. contigs"}]],["Unaligned",[{"values":[0,0,0,0,0,0,0,0,0,0,0,0,0,0],"quality":"Less is better","isMain":false,"metricName":"# fully unaligned contigs"},{"values":[0,0,0,0,0,0,0,0,0,0,0,0,0,0],"quality":"Less is better","isMain":false,"metricName":"Fully unaligned length"},{"values":[0,0,0,0,0,0,0,0,0,0,0,0,0,0],"quality":"Less is better","isMain":false,"metricName":"# partially unaligned contigs"},{"values":[0,0,0,0,0,0,0,0,0,0,0,0,0,0],"quality":"Less is better","isMain":false,"metricName":"Partially unaligned length"}]],["Mismatches",[{"values":[0,22,0,1,1,1,0,0,0,22,23,0,0,0],"quality":"Less is better","isMain":false,"metricName":"# mismatches"},{"values":[0,1,0,0,0,0,0,0,0,1,1,0,0,0],"quality":"Less is better","isMain":false,"metricName":"# indels"},{"values":[0,1,0,0,0,0,0,0,0,1,1,0,0,0],"quality":"Less is better","isMain":false,"metricName":"Indels length"},{"values":["0.00","28.28","0.00","1.28","1.26","1.26","0.00","0.00","0.00","27.95","29.24","0.00","0.00","0.00"],"quality":"Less is better","isMain":true,"metricName":"# mismatches per 100 kbp"},{"values":["0.00","1.29","0.00","0.00","0.00","0.00","0.00","0.00","0.00","1.27","1.27","0.00","0.00","0.00"],"quality":"Less is better","isMain":true,"metricName":"# indels per 100 kbp"},{"values":[0,1,0,0,0,0,0,0,0,1,1,0,0,0],"quality":"Less is better","isMain":false,"metricName":" # indels (<= 5 bp)"},{"values":[0,0,0,0,0,0,0,0,0,0,0,0,0,0],"quality":"Less is better","isMain":false,"metricName":" # indels (> 5 bp)"},{"values":[0,8,0,0,0,0,0,0,0,0,0,0,0,1],"quality":"Less is better","isMain":false,"metricName":"# N's"},{"values":["0.00","10.28","0.00","0.00","0.00","0.00","0.00","0.00","0.00","0.00","0.00","0.00","0.00","1.26"],"quality":"Less is better","isMain":true,"metricName":"# N's per 100 kbp"}]],["Statistics without reference",[{"values":[2,1,1,4,4,2,1,28,2,1,2,3,5,1],"quality":"Equal","isMain":true,"metricName":"# contigs"},{"values":[2,1,1,2,3,1,1,2,2,1,1,1,5,1],"quality":"Equal","isMain":false,"metricName":"# contigs (>= 1000 bp)"},{"values":[1,1,1,1,1,1,1,0,1,1,1,1,4,1],"quality":"Equal","isMain":false,"metricName":"# contigs (>= 5000 bp)"},{"values":[1,1,1,1,1,1,1,0,1,1,1,1,3,1],"quality":"Equal","isMain":false,"metricName":"# contigs (>= 10000 bp)"},{"values":[1,1,1,1,1,1,1,0,1,1,1,1,1,1],"quality":"Equal","isMain":false,"metricName":"# contigs (>= 25000 bp)"},{"values":[1,1,1,1,1,1,1,0,1,1,1,1,0,1],"quality":"Equal","isMain":false,"metricName":"# contigs (>= 50000 bp)"},{"values":[77806,77785,79232,75372,76862,79333,79570,1237,77808,78726,77778,77161,33429,79526],"quality":"More is better","isMain":true,"metricName":"Largest contig"},{"values":[79398,77785,79232,78416,79740,79929,79570,20224,79402,78726,78726,78781,75897,79526],"quality":"More is better","isMain":true,"metricName":"Total length"},{"values":[79398,77785,79232,76752,79092,79333,79570,2375,79402,78726,77778,77161,75897,79526],"quality":"More is better","isMain":true,"metricName":"Total length (>= 1000 bp)"},{"values":[77806,77785,79232,75372,76862,79333,79570,0,77808,78726,77778,77161,74863,79526],"quality":"More is better","isMain":false,"metricName":"Total length (>= 5000 bp)"},{"values":[77806,77785,79232,75372,76862,79333,79570,0,77808,78726,77778,77161,66337,79526],"quality":"More is better","isMain":true,"metricName":"Total length (>= 10000 bp)"},{"values":[77806,77785,79232,75372,76862,79333,79570,0,77808,78726,77778,77161,33429,79526],"quality":"More is better","isMain":false,"metricName":"Total length (>= 25000 bp)"},{"values":[77806,77785,79232,75372,76862,79333,79570,0,77808,78726,77778,77161,0,79526],"quality":"More is better","isMain":true,"metricName":"Total length (>= 50000 bp)"},{"values":[77806,77785,79232,75372,76862,79333,79570,729,77808,78726,77778,77161,21633,79526],"quality":"More is better","isMain":false,"metricName":"N50"},{"values":[77806,77785,79232,75372,76862,79333,79570,621,77808,78726,77778,77161,11275,79526],"quality":"More is better","isMain":false,"metricName":"N75"},{"values":[1,1,1,1,1,1,1,12,1,1,1,1,2,1],"quality":"Less is better","isMain":false,"metricName":"L50"},{"values":[1,1,1,1,1,1,1,19,1,1,1,1,3,1],"quality":"Less is better","isMain":false,"metricName":"L75"},{"values":["35.13","35.09","35.11","35.11","35.14","35.12","35.10","35.61","35.13","35.11","35.12","35.12","35.06","35.13"],"quality":"Equal","isMain":false,"metricName":"GC (%)"}]],["Predicted genes",[]],["Similarity statistics",[{"values":[2,2,2,2,2,0,0,0,2,2,2,2,0,0],"quality":"Equal","isMain":false,"metricName":"# similar correct contigs"},{"values":[0,0,0,0,0,0,0,0,0,0,0,0,0,0],"quality":"Equal","isMain":false,"metricName":"# similar misassembled blocks"}]],["Reference statistics",[{"values":[79232,79232,79232,79232,79232,79232,79232,79232,79232,79232,79232,79232,79232,79232],"quality":"Equal","isMain":false,"metricName":"Reference length"},{"values":[1,1,1,1,1,1,1,1,1,1,1,1,1,1],"quality":"Equal","isMain":false,"metricName":"Reference fragments"},{"values":["35.11","35.11","35.11","35.11","35.11","35.11","35.11","35.11","35.11","35.11","35.11","35.11","35.11","35.11"],"quality":"Equal","isMain":false,"metricName":"Reference GC (%)"}]]],"referenceName":"gi\_157311004\_ref\_NC","date":"18 July 2018, Wednesday, 16:25:04","order":[0,1,2,3,4,5,6,7,8,9,10,11,12,13],"assembliesNames":["ABySS\_127","ABySS\_63","CLC","IDBA\_UD","MEGAHIT","MIRA","Ray\_Meta","SOAPdenovo2","SPAdes","SPAdes\_meta","SPAdes\_sc","SPAdes\_sc\_careful","Velvet","Geneious"]},{"assembliesWithNs":null,"minContig":500,"report":[["Genome statistics",[{"values":["2.526","8.330","100.000","93.414","100.000","100.000","88.493","99.425","100.000","89.788","88.443","61.311","100.000"],"quality":"More is better","isMain":true,"metricName":"Genome fraction (%)"},{"values":["1.000","1.000","1.043","1.038","1.019","3.081","1.060","1.038","1.094","1.023","1.014","1.000","1.671"],"quality":"Less is better","isMain":true,"metricName":"Duplication ratio"},{"values":[522,643,14324,5465,23546,41293,17419,9154,27938,10167,9507,4300,36936],"quality":"More is better","isMain":true,"metricName":"Largest alignment"},{"values":[1024,3377,42275,39297,41308,124710,38014,41829,44356,36894,36349,24855,67747],"quality":"More is better","isMain":true,"metricName":"Total aligned length"},{"values":[null,null,7544,1712,30728,41293,15985,7626,34643,3370,3986,825,41554],"quality":"More is better","isMain":false,"metricName":"NG50"},{"values":[null,null,5115,1025,30728,41293,15985,5913,34643,1010,1087,null,41554],"quality":"More is better","isMain":false,"metricName":"NG75"},{"values":[522,567,7544,1771,6448,1880,15985,7626,3597,3370,3986,1127,36936],"quality":"More is better","isMain":false,"metricName":"NA50"},{"values":[502,545,5115,1032,3183,1428,15985,5580,null,1616,1327,846,9794],"quality":"More is better","isMain":false,"metricName":"NA75"},{"values":[null,null,7544,1712,23546,41293,15985,7626,27938,3370,3986,825,36936],"quality":"More is better","isMain":true,"metricName":"NGA50"},{"values":[null,null,5115,1025,4452,41293,15985,5913,9367,1010,931,null,36936],"quality":"More is better","isMain":false,"metricName":"NGA75"},{"values":[null,null,2,7,1,1,2,3,1,3,3,14,1],"quality":"Less is better","isMain":false,"metricName":"LG50"},{"values":[null,null,4,15,1,1,2,4,1,8,8,null,1],"quality":"Less is better","isMain":false,"metricName":"LG75"},{"values":[1,3,2,6,2,10,2,3,3,3,3,6,1],"quality":"Less is better","isMain":false,"metricName":"LA50"},{"values":[2,5,4,14,4,29,2,5,null,6,6,12,3],"quality":"Less is better","isMain":false,"metricName":"LA75"},{"values":[null,null,2,7,1,1,2,3,1,3,3,14,1],"quality":"Less is better","isMain":true,"metricName":"LGA50"},{"values":[null,null,4,15,3,1,2,4,2,8,9,null,1],"quality":"Less is better","isMain":false,"metricName":"LGA75"}]],["Misassemblies",[{"values":[0,0,0,0,0,0,0,0,0,0,0,0,2],"quality":"Less is better","isMain":true,"metricName":"# misassemblies"},{"values":[0,0,0,0,0,0,0,0,0,0,0,0,2],"quality":"Less is better","isMain":false,"metricName":" # relocations"},{"values":[0,0,0,0,0,0,0,0,0,0,0,0,0],"quality":"Less is better","isMain":false,"metricName":" # translocations"},{"values":[0,0,0,0,0,0,0,0,0,0,0,0,0],"quality":"Less is better","isMain":false,"metricName":" # inversions"},{"values":[0,0,0,0,0,0,0,0,0,0,0,0,2],"quality":"Less is better","isMain":false,"metricName":"# misassembled contigs"},{"values":[0,0,0,0,0,0,0,0,0,0,0,0,61859],"quality":"Less is better","isMain":true,"metricName":"Misassembled contigs length"},{"values":[0,0,0,0,0,1,0,0,0,0,0,0,2],"quality":"Less is better","isMain":false,"metricName":"# local misassemblies"},{"values":[0,0,0,0,0,0,0,0,0,0,0,0,0],"quality":"Less is better","isMain":false,"metricName":"# unaligned mis. contigs"}]],["Unaligned",[{"values":[0,0,0,0,0,0,0,0,0,0,0,0,0],"quality":"Less is better","isMain":false,"metricName":"# fully unaligned contigs"},{"values":[0,0,0,0,0,0,0,0,0,0,0,0,0],"quality":"Less is better","isMain":false,"metricName":"Fully unaligned length"},{"values":[0,0,0,0,1,0,0,0,1,0,1,0,1],"quality":"Less is better","isMain":false,"metricName":"# partially unaligned contigs"},{"values":[0,0,0,0,7182,0,0,0,31046,0,660,0,3906],"quality":"Less is better","isMain":false,"metricName":"Partially unaligned length"}]],["Mismatches",[{"values":[0,0,6,0,4,14,2,0,13,7,9,36,5],"quality":"Less is better","isMain":false,"metricName":"# mismatches"},{"values":[0,0,0,0,0,1,0,0,0,0,0,0,1],"quality":"Less is better","isMain":false,"metricName":"# indels"},{"values":[0,0,0,0,0,1,0,0,0,0,0,0,1],"quality":"Less is better","isMain":false,"metricName":"Indels length"},{"values":["0.00","0.00","14.80","0.00","9.87","34.53","5.58","0.00","32.07","19.23","25.10","144.84","12.33"],"quality":"Less is better","isMain":true,"metricName":"# mismatches per 100 kbp"},{"values":["0.00","0.00","0.00","0.00","0.00","2.47","0.00","0.00","0.00","0.00","0.00","0.00","2.47"],"quality":"Less is better","isMain":true,"metricName":"# indels per 100 kbp"},{"values":[0,0,0,0,0,1,0,0,0,0,0,0,1],"quality":"Less is better","isMain":false,"metricName":" # indels (<= 5 bp)"},{"values":[0,0,0,0,0,0,0,0,0,0,0,0,0],"quality":"Less is better","isMain":false,"metricName":" # indels (> 5 bp)"},{"values":[0,0,0,0,0,66,0,0,0,334,660,0,168],"quality":"Less is better","isMain":false,"metricName":"# N's"},{"values":["0.00","0.00","0.00","0.00","0.00","52.84","0.00","0.00","0.00","897.17","1783.35","0.00","234.46"],"quality":"Less is better","isMain":true,"metricName":"# N's per 100 kbp"}]],["Statistics without reference",[{"values":[2,6,9,27,6,55,3,10,5,17,16,21,3],"quality":"Equal","isMain":true,"metricName":"# contigs"},{"values":[0,0,7,15,6,50,3,7,5,8,8,9,3],"quality":"Equal","isMain":false,"metricName":"# contigs (>= 1000 bp)"},{"values":[0,0,4,1,2,2,2,5,3,2,2,0,3],"quality":"Equal","isMain":false,"metricName":"# contigs (>= 5000 bp)"},{"values":[0,0,1,0,1,1,2,0,2,1,1,0,2],"quality":"Equal","isMain":false,"metricName":"# contigs (>= 10000 bp)"},{"values":[0,0,0,0,1,1,0,0,2,0,0,0,1],"quality":"Equal","isMain":false,"metricName":"# contigs (>= 25000 bp)"},{"values":[0,0,0,0,0,0,0,0,0,0,0,0,0],"quality":"Equal","isMain":false,"metricName":"# contigs (>= 50000 bp)"},{"values":[522,643,14324,5465,30728,41293,17419,9154,34643,10167,10167,4300,41554],"quality":"More is better","isMain":true,"metricName":"Largest contig"},{"values":[1024,3377,42275,39297,48490,124906,38014,41829,75402,37228,37009,24855,71653],"quality":"More is better","isMain":true,"metricName":"Total length"},{"values":[0,0,41212,31015,48490,120348,38014,39780,75402,30745,31043,16366,71653],"quality":"More is better","isMain":true,"metricName":"Total length (>= 1000 bp)"},{"values":[0,0,32166,5465,37176,46685,33404,36644,71948,18502,18502,0,71653],"quality":"More is better","isMain":false,"metricName":"Total length (>= 5000 bp)"},{"values":[0,0,14324,0,30728,41293,33404,0,62581,10167,10167,0,61859],"quality":"More is better","isMain":true,"metricName":"Total length (>= 10000 bp)"},{"values":[0,0,0,0,30728,41293,0,0,62581,0,0,0,41554],"quality":"More is better","isMain":false,"metricName":"Total length (>= 25000 bp)"},{"values":[0,0,0,0,0,0,0,0,0,0,0,0,0],"quality":"More is better","isMain":true,"metricName":"Total length (>= 50000 bp)"},{"values":[522,567,7544,1771,30728,1880,15985,7626,27938,3370,3986,1127,41554],"quality":"More is better","isMain":false,"metricName":"N50"},{"values":[502,545,5115,1032,6448,1428,15985,5580,27938,1616,1327,846,20305],"quality":"More is better","isMain":false,"metricName":"N75"},{"values":[1,3,2,6,1,10,2,3,2,3,3,6,1],"quality":"Less is better","isMain":false,"metricName":"L50"},{"values":[2,5,4,14,2,29,2,5,2,6,6,12,2],"quality":"Less is better","isMain":false,"metricName":"L75"},{"values":["46.39","40.04","39.62","39.72","39.33","39.47","39.63","39.64","39.05","39.29","39.56","39.27","39.74"],"quality":"Equal","isMain":false,"metricName":"GC (%)"}]],["Predicted genes",[]],["Similarity statistics",[{"values":[0,0,0,0,0,0,0,0,0,0,0,0,0],"quality":"Equal","isMain":false,"metricName":"# similar correct contigs"},{"values":[0,0,0,0,0,0,0,0,0,0,0,0,0],"quality":"Equal","isMain":false,"metricName":"# similar misassembled blocks"}]],["Reference statistics",[{"values":[40539,40539,40539,40539,40539,40539,40539,40539,40539,40539,40539,40539,40539],"quality":"Equal","isMain":false,"metricName":"Reference length"},{"values":[1,1,1,1,1,1,1,1,1,1,1,1,1],"quality":"Equal","isMain":false,"metricName":"Reference fragments"},{"values":["39.65","39.65","39.65","39.65","39.65","39.65","39.65","39.65","39.65","39.65","39.65","39.65","39.65"],"quality":"Equal","isMain":false,"metricName":"Reference GC (%)"}]]],"referenceName":"gi\_157311135\_ref\_NC","date":"18 July 2018, Wednesday, 16:25:16","order":[0,1,2,3,4,5,6,7,8,9,10,11,12],"assembliesNames":["ABySS\_127","ABySS\_63","CLC","IDBA\_UD","MEGAHIT","MIRA","Ray\_Meta","SPAdes","SPAdes\_meta","SPAdes\_sc","SPAdes\_sc\_careful","Velvet","Geneious"]},{"assembliesWithNs":null,"minContig":500,"report":[["Genome statistics",[{"values":["26.995","7.195","10.351","23.903","29.542","54.471","8.223","23.756","12.635","18.846","17.997","6.541","34.834"],"quality":"More is better","isMain":true,"metricName":"Genome fraction (%)"},{"values":["1.011","1.015","1.000","1.014","1.025","1.201","1.280","1.013","1.000","1.021","1.022","1.000","1.221"],"quality":"Less is better","isMain":true,"metricName":"Duplication ratio"},{"values":[1482,941,968,2564,2592,5565,862,2521,1908,2629,2131,1914,5088],"quality":"More is better","isMain":true,"metricName":"Largest alignment"},{"values":[16018,4285,6074,14212,17481,38395,4826,14124,7416,11061,10563,3839,24971],"quality":"More is better","isMain":true,"metricName":"Total aligned length"},{"values":[null,null,null,null,null,56138,null,null,null,null,null,null,112476],"quality":"More is better","isMain":false,"metricName":"NG50"},{"values":[null,null,null,null,null,56138,null,null,null,null,null,null,112476],"quality":"More is better","isMain":false,"metricName":"NG75"},{"values":[826,702,706,986,1166,null,730,1846,901,842,829,817,null],"quality":"More is better","isMain":false,"metricName":"NA50"},{"values":[648,622,698,671,655,null,116,768,768,768,768,591,null],"quality":"More is better","isMain":false,"metricName":"NA75"},{"values":[null,null,null,null,null,1303,null,null,null,null,null,null,null],"quality":"More is better","isMain":true,"metricName":"NGA50"},{"values":[null,null,null,null,null,1,null,null,null,null,null,null,1],"quality":"Less is better","isMain":false,"metricName":"LG50"},{"values":[null,null,null,null,null,1,null,null,null,null,null,null,1],"quality":"Less is better","isMain":false,"metricName":"LG75"},{"values":[8,3,4,5,6,null,4,4,3,4,5,2,null],"quality":"Less is better","isMain":false,"metricName":"LA50"},{"values":[13,5,6,9,11,null,8,8,6,8,8,3,null],"quality":"Less is better","isMain":false,"metricName":"LA75"},{"values":[null,null,null,null,null,11,null,null,null,null,null,null,null],"quality":"Less is better","isMain":true,"metricName":"LGA50"}]],["Misassemblies",[{"values":[0,0,1,0,0,0,0,0,0,0,0,0,0],"quality":"Less is better","isMain":true,"metricName":"# misassemblies"},{"values":[0,0,1,0,0,0,0,0,0,0,0,0,0],"quality":"Less is better","isMain":false,"metricName":" # relocations"},{"values":[0,0,0,0,0,0,0,0,0,0,0,0,0],"quality":"Less is better","isMain":false,"metricName":" # translocations"},{"values":[0,0,0,0,0,0,0,0,0,0,0,0,0],"quality":"Less is better","isMain":false,"metricName":" # inversions"},{"values":[0,0,1,0,0,0,0,0,0,0,0,0,0],"quality":"Less is better","isMain":false,"metricName":"# misassembled contigs"},{"values":[0,0,1340,0,0,0,0,0,0,0,0,0,0],"quality":"Less is better","isMain":true,"metricName":"Misassembled contigs length"},{"values":[0,0,0,0,0,0,0,0,0,0,0,0,1],"quality":"Less is better","isMain":false,"metricName":"# local misassemblies"},{"values":[0,0,0,0,0,1,1,0,0,0,0,0,3],"quality":"Less is better","isMain":false,"metricName":"# unaligned mis. contigs"}]],["Unaligned",[{"values":[0,0,0,0,0,0,0,0,0,0,0,0,0],"quality":"Less is better","isMain":false,"metricName":"# fully unaligned contigs"},{"values":[0,0,0,0,0,0,0,0,0,0,0,0,0],"quality":"Less is better","isMain":false,"metricName":"Fully unaligned length"},{"values":[0,0,0,0,1,2,0,1,0,0,0,0,6],"quality":"Less is better","isMain":false,"metricName":"# partially unaligned contigs"},{"values":[0,0,0,0,913,67349,0,1647,0,0,0,0,197302],"quality":"Less is better","isMain":false,"metricName":"Partially unaligned length"}]],["Mismatches",[{"values":[18,10,10,20,43,389,11,145,15,38,28,10,256],"quality":"Less is better","isMain":false,"metricName":"# mismatches"},{"values":[0,0,1,1,1,6,2,18,0,0,0,0,7],"quality":"Less is better","isMain":false,"metricName":"# indels"},{"values":[0,0,1,1,1,57,93,26,0,0,0,0,82],"quality":"Less is better","isMain":false,"metricName":"Indels length"},{"values":["113.61","236.80","164.61","142.56","248.00","1216.77","227.93","1039.95","202.27","343.55","265.08","260.48","1252.14"],"quality":"Less is better","isMain":true,"metricName":"# mismatches per 100 kbp"},{"values":["0.00","0.00","16.46","7.13","5.77","18.77","41.44","129.10","0.00","0.00","0.00","0.00","34.24"],"quality":"Less is better","isMain":true,"metricName":"# indels per 100 kbp"},{"values":[0,0,1,1,1,5,0,18,0,0,0,0,6],"quality":"Less is better","isMain":false,"metricName":" # indels (<= 5 bp)"},{"values":[0,0,0,0,0,1,2,0,0,0,0,0,1],"quality":"Less is better","isMain":false,"metricName":" # indels (> 5 bp)"},{"values":[0,0,0,0,0,50,1149,0,0,0,0,0,628],"quality":"Less is better","isMain":false,"metricName":"# N's"},{"values":["0.00","0.00","0.00","0.00","0.00","47.28","18598.25","0.00","0.00","0.00","0.00","0.00","282.54"],"quality":"Less is better","isMain":true,"metricName":"# N's per 100 kbp"}]],["Statistics without reference",[{"values":[20,6,8,15,16,20,8,11,8,11,11,4,6],"quality":"Equal","isMain":true,"metricName":"# contigs"},{"values":[4,0,1,4,8,15,1,5,1,2,2,1,6],"quality":"Equal","isMain":false,"metricName":"# contigs (>= 1000 bp)"},{"values":[0,0,0,0,0,2,0,0,0,0,0,0,3],"quality":"Equal","isMain":false,"metricName":"# contigs (>= 5000 bp)"},{"values":[0,0,0,0,0,2,0,0,0,0,0,0,2],"quality":"Equal","isMain":false,"metricName":"# contigs (>= 10000 bp)"},{"values":[0,0,0,0,0,1,0,0,0,0,0,0,2],"quality":"Equal","isMain":false,"metricName":"# contigs (>= 25000 bp)"},{"values":[0,0,0,0,0,1,0,0,0,0,0,0,2],"quality":"Equal","isMain":false,"metricName":"# contigs (>= 50000 bp)"},{"values":[1482,941,1340,2564,2592,56138,1064,4168,1908,2629,2131,1914,112476],"quality":"More is better","isMain":true,"metricName":"Largest contig"},{"values":[16018,4285,6074,14221,18677,105751,6178,15773,7416,11297,10799,3839,222273],"quality":"More is better","isMain":true,"metricName":"Total length"},{"values":[4906,0,1340,6509,13420,102198,1064,11616,1908,3964,3466,1914,222273],"quality":"More is better","isMain":true,"metricName":"Total length (>= 1000 bp)"},{"values":[0,0,0,0,0,74674,0,0,0,0,0,0,210428],"quality":"More is better","isMain":false,"metricName":"Total length (>= 5000 bp)"},{"values":[0,0,0,0,0,74674,0,0,0,0,0,0,205011],"quality":"More is better","isMain":true,"metricName":"Total length (>= 10000 bp)"},{"values":[0,0,0,0,0,56138,0,0,0,0,0,0,205011],"quality":"More is better","isMain":false,"metricName":"Total length (>= 25000 bp)"},{"values":[0,0,0,0,0,56138,0,0,0,0,0,0,205011],"quality":"More is better","isMain":true,"metricName":"Total length (>= 50000 bp)"},{"values":[826,702,706,986,1311,56138,849,2154,901,933,842,817,112476],"quality":"More is better","isMain":false,"metricName":"N50"},{"values":[648,622,698,671,889,3999,663,841,768,775,775,591,92535],"quality":"More is better","isMain":false,"metricName":"N75"},{"values":[8,3,4,5,5,1,4,3,3,4,5,2,1],"quality":"Less is better","isMain":false,"metricName":"L50"},{"values":[13,5,6,9,9,4,6,6,6,8,8,3,2],"quality":"Less is better","isMain":false,"metricName":"L75"},{"values":["62.13","61.54","62.45","62.66","62.02","61.20","62.10","62.19","63.16","62.06","62.09","61.24","61.34"],"quality":"Equal","isMain":false,"metricName":"GC (%)"}]],["Predicted genes",[]],["Similarity statistics",[{"values":[0,0,0,0,0,0,0,0,0,0,0,0,0],"quality":"Equal","isMain":false,"metricName":"# similar correct contigs"},{"values":[0,0,0,0,0,0,0,0,0,0,0,0,0],"quality":"Equal","isMain":false,"metricName":"# similar misassembled blocks"}]],["Reference statistics",[{"values":[58692,58692,58692,58692,58692,58692,58692,58692,58692,58692,58692,58692,58692],"quality":"Equal","isMain":false,"metricName":"Reference length"},{"values":[1,1,1,1,1,1,1,1,1,1,1,1,1],"quality":"Equal","isMain":false,"metricName":"Reference fragments"},{"values":["61.74","61.74","61.74","61.74","61.74","61.74","61.74","61.74","61.74","61.74","61.74","61.74","61.74"],"quality":"Equal","isMain":false,"metricName":"Reference GC (%)"}]]],"referenceName":"gi\_157311189\_ref\_NC","date":"18 July 2018, Wednesday, 16:25:28","order":[0,1,2,3,4,5,6,7,8,9,10,11,12],"assembliesNames":["ABySS\_127","ABySS\_63","CLC","IDBA\_UD","MEGAHIT","MIRA","SOAPdenovo2","SPAdes","SPAdes\_meta","SPAdes\_sc","SPAdes\_sc\_careful","Velvet","Geneious"]},{"assembliesWithNs":null,"minContig":500,"report":[["Genome statistics",[{"values":["15.885","7.872","32.331","26.220","43.499","42.008","8.845","41.380","12.329","13.783","12.093","22.156","23.330"],"quality":"More is better","isMain":true,"metricName":"Genome fraction (%)"},{"values":["1.005","1.000","1.006","1.032","1.033","1.076","1.115","1.088","1.001","1.003","1.004","1.035","1.001"],"quality":"Less is better","isMain":true,"metricName":"Duplication ratio"},{"values":[1944,1488,6059,7464,6067,15860,2394,37384,2978,3228,2126,5528,14268],"quality":"More is better","isMain":true,"metricName":"Largest alignment"},{"values":[26221,12932,53428,44433,73802,74172,14578,73982,20269,22711,19936,37180,38352],"quality":"More is better","isMain":true,"metricName":"Total aligned length"},{"values":[null,null,null,null,null,null,null,530,null,null,null,null,null],"quality":"More is better","isMain":false,"metricName":"NG50"},{"values":[799,774,1699,2477,1693,2301,712,5016,1565,1202,952,2186,9374],"quality":"More is better","isMain":false,"metricName":"NA50"},{"values":[591,684,911,790,967,1101,null,672,846,883,731,1118,9262],"quality":"More is better","isMain":false,"metricName":"NA75"},{"values":[null,null,null,null,null,null,null,null,null,null,null,null,null],"quality":"More is better","isMain":true,"metricName":"NGA50"},{"values":[null,null,null,null,null,null,null,30,null,null,null,null,null],"quality":"Less is better","isMain":false,"metricName":"LG50"},{"values":[12,6,10,5,14,8,10,2,5,6,8,6,2],"quality":"Less is better","isMain":false,"metricName":"LA50"},{"values":[22,11,21,15,30,21,null,16,10,11,14,12,3],"quality":"Less is better","isMain":false,"metricName":"LA75"}]],["Misassemblies",[{"values":[0,0,0,0,0,0,0,0,0,0,0,2,0],"quality":"Less is better","isMain":true,"metricName":"# misassemblies"},{"values":[0,0,0,0,0,0,0,0,0,0,0,2,0],"quality":"Less is better","isMain":false,"metricName":" # relocations"},{"values":[0,0,0,0,0,0,0,0,0,0,0,0,0],"quality":"Less is better","isMain":false,"metricName":" # translocations"},{"values":[0,0,0,0,0,0,0,0,0,0,0,0,0],"quality":"Less is better","isMain":false,"metricName":" # inversions"},{"values":[0,0,0,0,0,0,0,0,0,0,0,2,0],"quality":"Less is better","isMain":false,"metricName":"# misassembled contigs"},{"values":[0,0,0,0,0,0,0,0,0,0,0,8430,0],"quality":"Less is better","isMain":true,"metricName":"Misassembled contigs length"},{"values":[0,0,0,0,0,2,0,7,0,0,0,5,5],"quality":"Less is better","isMain":false,"metricName":"# local misassemblies"},{"values":[0,0,0,0,0,0,1,0,0,0,0,0,0],"quality":"Less is better","isMain":false,"metricName":"# unaligned mis. contigs"}]],["Unaligned",[{"values":[0,0,0,0,0,0,0,0,0,0,0,0,0],"quality":"Less is better","isMain":false,"metricName":"# fully unaligned contigs"},{"values":[0,0,0,0,0,0,0,0,0,0,0,0,0],"quality":"Less is better","isMain":false,"metricName":"Fully unaligned length"},{"values":[0,0,0,1,0,1,4,2,0,0,0,2,2],"quality":"Less is better","isMain":false,"metricName":"# partially unaligned contigs"},{"values":[0,0,0,898,0,728,3796,10697,0,0,0,1539,5297],"quality":"Less is better","isMain":false,"metricName":"Partially unaligned length"}]],["Mismatches",[{"values":[57,61,383,443,454,562,59,1116,122,89,42,436,372],"quality":"Less is better","isMain":false,"metricName":"# mismatches"},{"values":[0,0,28,30,8,46,8,71,0,2,4,28,39],"quality":"Less is better","isMain":false,"metricName":"# indels"},{"values":[0,0,33,37,10,72,397,86,0,2,4,84,66],"quality":"Less is better","isMain":false,"metricName":"Indels length"},{"values":["218.43","471.70","721.14","1028.53","635.36","814.41","406.06","1641.78","602.38","393.07","211.43","1197.93","970.67"],"quality":"Less is better","isMain":true,"metricName":"# mismatches per 100 kbp"},{"values":["0.00","0.00","52.72","69.65","11.20","66.66","55.06","104.45","0.00","8.83","20.14","76.93","101.76"],"quality":"Less is better","isMain":true,"metricName":"# indels per 100 kbp"},{"values":[0,0,28,30,8,44,0,71,0,2,4,27,37],"quality":"Less is better","isMain":false,"metricName":" # indels (<= 5 bp)"},{"values":[0,0,0,0,0,2,8,0,0,0,0,1,2],"quality":"Less is better","isMain":false,"metricName":" # indels (> 5 bp)"},{"values":[0,0,0,0,0,178,2111,0,0,0,0,50,475],"quality":"Less is better","isMain":false,"metricName":"# N's"},{"values":["0.00","0.00","0.00","0.00","0.00","237.46","10553.94","0.00","0.00","0.00","0.00","127.47","1088.23"],"quality":"Less is better","isMain":true,"metricName":"# N's per 100 kbp"}]],["Statistics without reference",[{"values":[33,16,40,30,55,40,21,34,17,19,21,22,4],"quality":"Equal","isMain":true,"metricName":"# contigs"},{"values":[6,4,18,10,29,28,7,9,6,7,5,13,4],"quality":"Equal","isMain":false,"metricName":"# contigs (>= 1000 bp)"},{"values":[0,0,1,3,1,2,0,3,0,0,0,2,4],"quality":"Equal","isMain":false,"metricName":"# contigs (>= 5000 bp)"},{"values":[0,0,0,0,0,1,0,1,0,0,0,0,1],"quality":"Equal","isMain":false,"metricName":"# contigs (>= 10000 bp)"},{"values":[0,0,0,0,0,0,0,1,0,0,0,0,0],"quality":"Equal","isMain":false,"metricName":"# contigs (>= 25000 bp)"},{"values":[0,0,0,0,0,0,0,0,0,0,0,0,0],"quality":"Equal","isMain":false,"metricName":"# contigs (>= 50000 bp)"},{"values":[1944,1488,6059,7464,6067,16588,2394,43840,2978,3228,2126,6734,18963],"quality":"More is better","isMain":true,"metricName":"Largest contig"},{"values":[26221,12932,53431,45334,73803,74961,20002,84685,20274,22721,19939,39226,43649],"quality":"More is better","isMain":true,"metricName":"Total length"},{"values":[8340,4878,37515,31200,55143,65286,9990,68959,11947,13388,7727,33522,43649],"quality":"More is better","isMain":true,"metricName":"Total length (>= 1000 bp)"},{"values":[0,0,6059,18117,6067,22036,0,56445,0,0,0,12102,43649],"quality":"More is better","isMain":false,"metricName":"Total length (>= 5000 bp)"},{"values":[0,0,0,0,0,16588,0,43840,0,0,0,0,18963],"quality":"More is better","isMain":true,"metricName":"Total length (>= 10000 bp)"},{"values":[0,0,0,0,0,0,0,43840,0,0,0,0,0],"quality":"More is better","isMain":false,"metricName":"Total length (>= 25000 bp)"},{"values":[0,0,0,0,0,0,0,0,0,0,0,0,0],"quality":"More is better","isMain":true,"metricName":"Total length (>= 50000 bp)"},{"values":[799,774,1699,2477,1693,2383,981,43840,1567,1202,952,2341,9976],"quality":"More is better","isMain":false,"metricName":"N50"},{"values":[591,684,911,892,967,1133,719,2483,846,883,731,1424,9262],"quality":"More is better","isMain":false,"metricName":"N75"},{"values":[12,6,10,5,14,7,8,1,5,6,8,5,2],"quality":"Less is better","isMain":false,"metricName":"L50"},{"values":[22,11,21,13,30,20,13,5,10,11,14,10,3],"quality":"Less is better","isMain":false,"metricName":"L75"},{"values":["40.46","40.14","40.26","39.97","40.13","40.21","40.01","40.63","39.47","39.05","39.32","40.59","39.97"],"quality":"Equal","isMain":false,"metricName":"GC (%)"}]],["Predicted genes",[]],["Similarity statistics",[{"values":[0,0,0,0,0,0,0,0,0,0,0,0,0],"quality":"Equal","isMain":false,"metricName":"# similar correct contigs"},{"values":[0,0,0,0,0,0,0,0,0,0,0,0,0],"quality":"Equal","isMain":false,"metricName":"# similar misassembled blocks"}]],["Reference statistics",[{"values":[164270,164270,164270,164270,164270,164270,164270,164270,164270,164270,164270,164270,164270],"quality":"Equal","isMain":false,"metricName":"Reference length"},{"values":[1,1,1,1,1,1,1,1,1,1,1,1,1],"quality":"Equal","isMain":false,"metricName":"Reference fragments"},{"values":["40.50","40.50","40.50","40.50","40.50","40.50","40.50","40.50","40.50","40.50","40.50","40.50","40.50"],"quality":"Equal","isMain":false,"metricName":"Reference GC (%)"}]]],"referenceName":"gi\_157311299\_ref\_NC","date":"18 July 2018, Wednesday, 16:25:40","order":[0,1,2,3,4,5,6,7,8,9,10,11,12],"assembliesNames":["ABySS\_127","ABySS\_63","CLC","IDBA\_UD","MEGAHIT","MIRA","SOAPdenovo2","SPAdes","SPAdes\_meta","SPAdes\_sc","SPAdes\_sc\_careful","Velvet","Geneious"]},{"assembliesWithNs":null,"minContig":500,"report":[["Genome statistics",[{"values":["100.000","100.000","98.858","98.304","99.995","100.000","100.000","99.174","98.690","97.039","100.000","65.791","100.000"],"quality":"More is better","isMain":true,"metricName":"Genome fraction (%)"},{"values":["1.015","1.024","1.000","1.009","1.019","1.042","1.046","1.000","1.000","1.003","1.006","1.002","1.044"],"quality":"Less is better","isMain":true,"metricName":"Duplication ratio"},{"values":[39442,18038,36283,14698,15551,27913,22473,38546,38357,16069,21514,4222,26305],"quality":"More is better","isMain":true,"metricName":"Largest alignment"},{"values":[39442,39810,38423,38557,39621,40213,40657,38546,38357,37825,39087,25594,40581],"quality":"More is better","isMain":true,"metricName":"Total aligned length"},{"values":[39442,11941,36283,9609,9421,28184,22473,38546,38357,10944,21514,1228,192212],"quality":"More is better","isMain":false,"metricName":"NG50"},{"values":[39442,11941,36283,7584,7668,7887,18184,38546,38357,9427,15696,null,192212],"quality":"More is better","isMain":false,"metricName":"NG75"},{"values":[39442,11941,36283,9609,7668,27913,22473,38546,38357,10944,21514,2209,null],"quality":"More is better","isMain":false,"metricName":"NA50"},{"values":[39442,11941,36283,7584,4169,7887,18184,38546,38357,9427,15696,1228,null],"quality":"More is better","isMain":false,"metricName":"NA75"},{"values":[39442,11941,36283,9609,7668,27913,22473,38546,38357,10944,21514,1228,26305],"quality":"More is better","isMain":true,"metricName":"NGA50"},{"values":[39442,11941,36283,7584,4327,7887,18184,38546,38357,9427,15696,null,13376],"quality":"More is better","isMain":false,"metricName":"NGA75"},{"values":[1,2,1,2,2,1,1,1,1,2,1,9,1],"quality":"Less is better","isMain":false,"metricName":"LG50"},{"values":[1,2,1,3,3,2,2,1,1,3,2,null,1],"quality":"Less is better","isMain":false,"metricName":"LG75"},{"values":[1,2,1,2,2,1,1,1,1,2,1,4,null],"quality":"Less is better","isMain":false,"metricName":"LA50"},{"values":[1,2,1,3,5,2,2,1,1,3,2,9,null],"quality":"Less is better","isMain":false,"metricName":"LA75"},{"values":[1,2,1,2,2,1,1,1,1,2,1,9,1],"quality":"Less is better","isMain":true,"metricName":"LGA50"},{"values":[1,2,1,3,4,2,2,1,1,3,2,null,2],"quality":"Less is better","isMain":false,"metricName":"LGA75"}]],["Misassemblies",[{"values":[0,0,0,0,0,0,0,0,0,0,0,0,0],"quality":"Less is better","isMain":true,"metricName":"# misassemblies"},{"values":[0,0,0,0,0,0,0,0,0,0,0,0,0],"quality":"Less is better","isMain":false,"metricName":" # relocations"},{"values":[0,0,0,0,0,0,0,0,0,0,0,0,0],"quality":"Less is better","isMain":false,"metricName":" # translocations"},{"values":[0,0,0,0,0,0,0,0,0,0,0,0,0],"quality":"Less is better","isMain":false,"metricName":" # inversions"},{"values":[0,0,0,0,0,0,0,0,0,0,0,0,0],"quality":"Less is better","isMain":false,"metricName":"# misassembled contigs"},{"values":[0,0,0,0,0,0,0,0,0,0,0,0,0],"quality":"Less is better","isMain":true,"metricName":"Misassembled contigs length"},{"values":[1,0,0,0,0,0,0,0,0,0,0,0,0],"quality":"Less is better","isMain":false,"metricName":"# local misassemblies"},{"values":[0,0,0,0,0,0,0,0,0,0,0,0,1],"quality":"Less is better","isMain":false,"metricName":"# unaligned mis. contigs"}]],["Unaligned",[{"values":[0,0,0,0,0,0,0,0,0,0,0,0,0],"quality":"Less is better","isMain":false,"metricName":"# fully unaligned contigs"},{"values":[0,0,0,0,0,0,0,0,0,0,0,0,0],"quality":"Less is better","isMain":false,"metricName":"Fully unaligned length"},{"values":[0,0,0,0,2,0,0,0,0,0,0,0,1],"quality":"Less is better","isMain":false,"metricName":"# partially unaligned contigs"},{"values":[0,0,0,0,5410,0,0,0,0,0,0,0,151631],"quality":"Less is better","isMain":false,"metricName":"Partially unaligned length"}]],["Mismatches",[{"values":[0,28,0,1,1,0,0,0,13,9,25,0,4],"quality":"Less is better","isMain":false,"metricName":"# mismatches"},{"values":[0,2,0,0,1,0,0,0,1,1,0,0,0],"quality":"Less is better","isMain":false,"metricName":"# indels"},{"values":[0,2,0,0,1,0,0,0,1,1,0,0,0],"quality":"Less is better","isMain":false,"metricName":"Indels length"},{"values":["0.00","72.04","0.00","2.62","2.57","0.00","0.00","0.00","33.89","23.86","64.32","0.00","10.29"],"quality":"Less is better","isMain":true,"metricName":"# mismatches per 100 kbp"},{"values":["0.00","5.15","0.00","0.00","2.57","0.00","0.00","0.00","2.61","2.65","0.00","0.00","0.00"],"quality":"Less is better","isMain":true,"metricName":"# indels per 100 kbp"},{"values":[0,2,0,0,1,0,0,0,1,1,0,0,0],"quality":"Less is better","isMain":false,"metricName":" # indels (<= 5 bp)"},{"values":[0,0,0,0,0,0,0,0,0,0,0,0,0],"quality":"Less is better","isMain":false,"metricName":" # indels (> 5 bp)"},{"values":[0,0,0,0,0,0,0,0,0,0,0,0,17],"quality":"Less is better","isMain":false,"metricName":"# N's"},{"values":["0.00","0.00","0.00","0.00","0.00","0.00","0.00","0.00","0.00","0.00","0.00","0.00","8.84"],"quality":"Less is better","isMain":true,"metricName":"# N's per 100 kbp"}]],["Statistics without reference",[{"values":[1,5,2,7,7,3,2,1,1,4,4,16,1],"quality":"Equal","isMain":true,"metricName":"# contigs"},{"values":[1,3,2,6,6,3,2,1,1,4,3,12,1],"quality":"Equal","isMain":false,"metricName":"# contigs (>= 1000 bp)"},{"values":[1,3,1,3,4,2,2,1,1,3,2,0,1],"quality":"Equal","isMain":false,"metricName":"# contigs (>= 5000 bp)"},{"values":[1,2,1,1,1,1,2,1,1,2,2,0,1],"quality":"Equal","isMain":false,"metricName":"# contigs (>= 10000 bp)"},{"values":[1,0,1,0,0,1,0,1,1,0,0,0,1],"quality":"Equal","isMain":false,"metricName":"# contigs (>= 25000 bp)"},{"values":[0,0,0,0,0,0,0,0,0,0,0,0,1],"quality":"Equal","isMain":false,"metricName":"# contigs (>= 50000 bp)"},{"values":[39442,18038,36283,14698,15551,28184,22473,38546,38357,16069,21514,4222,192212],"quality":"More is better","isMain":true,"metricName":"Largest contig"},{"values":[39442,39810,38423,38557,45031,40484,40657,38546,38357,37825,39087,25634,192212],"quality":"More is better","isMain":true,"metricName":"Total length"},{"values":[39442,38399,38423,37653,44456,40484,40657,38546,38357,37825,38335,23449,192212],"quality":"More is better","isMain":true,"metricName":"Total length (>= 1000 bp)"},{"values":[39442,38399,36283,31891,37987,36071,40657,38546,38357,36440,37210,0,192212],"quality":"More is better","isMain":false,"metricName":"Total length (>= 5000 bp)"},{"values":[39442,29979,36283,14698,15551,28184,40657,38546,38357,27013,37210,0,192212],"quality":"More is better","isMain":true,"metricName":"Total length (>= 10000 bp)"},{"values":[39442,0,36283,0,0,28184,0,38546,38357,0,0,0,192212],"quality":"More is better","isMain":false,"metricName":"Total length (>= 25000 bp)"},{"values":[0,0,0,0,0,0,0,0,0,0,0,0,192212],"quality":"More is better","isMain":true,"metricName":"Total length (>= 50000 bp)"},{"values":[39442,11941,36283,9609,9421,28184,22473,38546,38357,10944,21514,2209,192212],"quality":"More is better","isMain":false,"metricName":"N50"},{"values":[39442,11941,36283,7584,5347,7887,18184,38546,38357,9427,15696,1228,192212],"quality":"More is better","isMain":false,"metricName":"N75"},{"values":[1,2,1,2,2,1,1,1,1,2,1,4,1],"quality":"Less is better","isMain":false,"metricName":"L50"},{"values":[1,2,1,3,4,2,2,1,1,3,2,9,1],"quality":"Less is better","isMain":false,"metricName":"L75"},{"values":["36.63","36.66","36.73","36.74","36.69","36.73","36.70","36.67","36.74","36.75","36.63","36.40","35.60"],"quality":"Equal","isMain":false,"metricName":"GC (%)"}]],["Predicted genes",[]],["Similarity statistics",[{"values":[0,0,0,0,0,0,0,0,0,0,0,0,0],"quality":"Equal","isMain":false,"metricName":"# similar correct contigs"},{"values":[0,0,0,0,0,0,0,0,0,0,0,0,0],"quality":"Equal","isMain":false,"metricName":"# similar misassembled blocks"}]],["Reference statistics",[{"values":[38867,38867,38867,38867,38867,38867,38867,38867,38867,38867,38867,38867,38867],"quality":"Equal","isMain":false,"metricName":"Reference length"},{"values":[1,1,1,1,1,1,1,1,1,1,1,1,1],"quality":"Equal","isMain":false,"metricName":"Reference fragments"},{"values":["36.68","36.68","36.68","36.68","36.68","36.68","36.68","36.68","36.68","36.68","36.68","36.68","36.68"],"quality":"Equal","isMain":false,"metricName":"Reference GC (%)"}]]],"referenceName":"gi\_157324960\_ref\_NC","date":"18 July 2018, Wednesday, 16:25:52","order":[0,1,2,3,4,5,6,7,8,9,10,11,12],"assembliesNames":["ABySS\_127","ABySS\_63","CLC","IDBA\_UD","MEGAHIT","MIRA","Ray\_Meta","SPAdes","SPAdes\_meta","SPAdes\_sc","SPAdes\_sc\_careful","Velvet","Geneious"]},{"assembliesWithNs":null,"minContig":500,"report":[["Genome statistics",[{"values":["100.000","93.039","99.259","99.353","97.836","100.000","55.801","37.128","99.665","81.270","95.201","95.184","74.332","95.055"],"quality":"More is better","isMain":true,"metricName":"Genome fraction (%)"},{"values":["1.036","1.004","1.001","1.007","1.000","1.034","1.010","1.135","1.006","1.009","1.002","1.004","1.001","1.039"],"quality":"Less is better","isMain":true,"metricName":"Duplication ratio"},{"values":[35217,26509,22939,33833,33916,35050,2635,11924,34948,33241,28016,32756,12755,32063],"quality":"More is better","isMain":true,"metricName":"Largest alignment"},{"values":[44199,39847,42381,42660,41737,44091,24028,16078,42764,34663,40701,40764,31734,42120],"quality":"More is better","isMain":true,"metricName":"Total aligned length"},{"values":[35217,26509,22939,33833,33916,36517,661,4765,34948,33561,28016,32756,4600,192212],"quality":"More is better","isMain":false,"metricName":"NG50"},{"values":[35217,2628,18353,33833,33916,36517,null,566,34948,33561,5736,32756,null,192212],"quality":"More is better","isMain":false,"metricName":"NG75"},{"values":[35217,26509,22939,33833,33916,35050,897,null,34948,33241,28016,32756,6304,null],"quality":"More is better","isMain":false,"metricName":"NA50"},{"values":[35217,4761,18353,33833,33916,9041,789,null,34948,33241,5736,32756,2362,null],"quality":"More is better","isMain":false,"metricName":"NA75"},{"values":[35217,26509,22939,33833,33916,35050,661,null,34948,33241,28016,32756,4600,32063],"quality":"More is better","isMain":true,"metricName":"NGA50"},{"values":[35217,2628,18353,33833,33916,35050,null,null,34948,33241,5736,32756,null,32063],"quality":"More is better","isMain":false,"metricName":"NGA75"},{"values":[1,1,1,1,1,1,21,2,1,1,1,1,3,1],"quality":"Less is better","isMain":false,"metricName":"LG50"},{"values":[1,3,2,1,1,1,null,9,1,1,2,1,null,1],"quality":"Less is better","isMain":false,"metricName":"LG75"},{"values":[1,1,1,1,1,1,9,null,1,1,1,1,2,null],"quality":"Less is better","isMain":false,"metricName":"LA50"},{"values":[1,2,2,1,1,2,16,null,1,1,2,1,4,null],"quality":"Less is better","isMain":false,"metricName":"LA75"},{"values":[1,1,1,1,1,1,21,null,1,1,1,1,3,1],"quality":"Less is better","isMain":true,"metricName":"LGA50"},{"values":[1,3,2,1,1,1,null,null,1,1,2,1,null,1],"quality":"Less is better","isMain":false,"metricName":"LGA75"}]],["Misassemblies",[{"values":[0,0,0,0,0,0,0,0,0,0,0,0,0,0],"quality":"Less is better","isMain":true,"metricName":"# misassemblies"},{"values":[0,0,0,0,0,0,0,0,0,0,0,0,0,0],"quality":"Less is better","isMain":false,"metricName":" # relocations"},{"values":[0,0,0,0,0,0,0,0,0,0,0,0,0,0],"quality":"Less is better","isMain":false,"metricName":" # translocations"},{"values":[0,0,0,0,0,0,0,0,0,0,0,0,0,0],"quality":"Less is better","isMain":false,"metricName":" # inversions"},{"values":[0,0,0,0,0,0,0,0,0,0,0,0,0,0],"quality":"Less is better","isMain":false,"metricName":"# misassembled contigs"},{"values":[0,0,0,0,0,0,0,0,0,0,0,0,0,0],"quality":"Less is better","isMain":true,"metricName":"Misassembled contigs length"},{"values":[0,0,0,0,0,0,0,2,0,1,0,0,0,0],"quality":"Less is better","isMain":false,"metricName":"# local misassemblies"},{"values":[0,0,0,0,0,0,0,4,0,0,0,0,0,1],"quality":"Less is better","isMain":false,"metricName":"# unaligned mis. contigs"}]],["Unaligned",[{"values":[0,0,0,0,0,0,0,0,0,0,0,0,0,0],"quality":"Less is better","isMain":false,"metricName":"# fully unaligned contigs"},{"values":[0,0,0,0,0,0,0,0,0,0,0,0,0,0],"quality":"Less is better","isMain":false,"metricName":"Fully unaligned length"},{"values":[0,0,0,0,0,2,0,4,0,0,0,0,0,1],"quality":"Less is better","isMain":false,"metricName":"# partially unaligned contigs"},{"values":[0,0,0,0,0,3952,0,14269,0,0,0,0,0,150092],"quality":"Less is better","isMain":false,"metricName":"Partially unaligned length"}]],["Mismatches",[{"values":[81,47,14,0,0,0,0,1,0,247,15,0,0,71],"quality":"Less is better","isMain":false,"metricName":"# mismatches"},{"values":[0,0,0,0,0,0,0,7,0,7,1,0,0,5],"quality":"Less is better","isMain":false,"metricName":"# indels"},{"values":[0,0,0,0,0,0,0,325,0,7,15,0,0,5],"quality":"Less is better","isMain":false,"metricName":"Indels length"},{"values":["189.90","118.44","33.07","0.00","0.00","0.00","0.00","6.31","0.00","712.55","36.94","0.00","0.00","175.12"],"quality":"Less is better","isMain":true,"metricName":"# mismatches per 100 kbp"},{"values":["0.00","0.00","0.00","0.00","0.00","0.00","0.00","44.20","0.00","20.19","2.46","0.00","0.00","12.33"],"quality":"Less is better","isMain":true,"metricName":"# indels per 100 kbp"},{"values":[0,0,0,0,0,0,0,0,0,7,0,0,0,5],"quality":"Less is better","isMain":false,"metricName":" # indels (<= 5 bp)"},{"values":[0,0,0,0,0,0,0,7,0,0,1,0,0,0],"quality":"Less is better","isMain":false,"metricName":" # indels (> 5 bp)"},{"values":[0,0,0,0,0,0,0,4035,0,0,0,0,0,17],"quality":"Less is better","isMain":false,"metricName":"# N's"},{"values":["0.00","0.00","0.00","0.00","0.00","0.00","0.00","12514.73","0.00","0.00","0.00","0.00","0.00","8.84"],"quality":"Less is better","isMain":true,"metricName":"# N's per 100 kbp"}]],["Statistics without reference",[{"values":[2,8,3,5,3,2,25,9,3,2,5,5,9,1],"quality":"Equal","isMain":true,"metricName":"# contigs"},{"values":[2,5,3,4,3,2,5,5,3,2,5,5,6,1],"quality":"Equal","isMain":false,"metricName":"# contigs (>= 1000 bp)"},{"values":[2,1,2,1,2,2,0,1,2,1,2,1,2,1],"quality":"Equal","isMain":false,"metricName":"# contigs (>= 5000 bp)"},{"values":[1,1,2,1,1,2,0,1,1,1,1,1,1,1],"quality":"Equal","isMain":false,"metricName":"# contigs (>= 10000 bp)"},{"values":[1,1,0,1,1,1,0,0,1,1,1,1,0,1],"quality":"Equal","isMain":false,"metricName":"# contigs (>= 25000 bp)"},{"values":[0,0,0,0,0,0,0,0,0,0,0,0,0,1],"quality":"Equal","isMain":false,"metricName":"# contigs (>= 50000 bp)"},{"values":[35217,26509,22939,33833,33916,36517,2635,20434,34948,33561,28016,32756,12755,192212],"quality":"More is better","isMain":true,"metricName":"Largest contig"},{"values":[44199,39847,42381,42660,41737,48043,24028,32242,42764,34983,40701,40764,31734,192212],"quality":"More is better","isMain":true,"metricName":"Total length"},{"values":[44199,37166,42381,41678,41737,48043,8659,29151,42764,34983,40701,40764,29416,192212],"quality":"More is better","isMain":true,"metricName":"Total length (>= 1000 bp)"},{"values":[44199,26509,41292,33833,40671,48043,0,20434,41675,33561,33752,32756,19059,192212],"quality":"More is better","isMain":false,"metricName":"Total length (>= 5000 bp)"},{"values":[35217,26509,41292,33833,33916,48043,0,20434,34948,33561,28016,32756,12755,192212],"quality":"More is better","isMain":true,"metricName":"Total length (>= 10000 bp)"},{"values":[35217,26509,0,33833,33916,36517,0,0,34948,33561,28016,32756,0,192212],"quality":"More is better","isMain":false,"metricName":"Total length (>= 25000 bp)"},{"values":[0,0,0,0,0,0,0,0,0,0,0,0,0,192212],"quality":"More is better","isMain":true,"metricName":"Total length (>= 50000 bp)"},{"values":[35217,26509,22939,33833,33916,36517,897,20434,34948,33561,28016,32756,6304,192212],"quality":"More is better","isMain":false,"metricName":"N50"},{"values":[35217,4761,18353,33833,33916,36517,789,4765,34948,33561,5736,32756,2362,192212],"quality":"More is better","isMain":false,"metricName":"N75"},{"values":[1,1,1,1,1,1,9,1,1,1,1,1,2,1],"quality":"Less is better","isMain":false,"metricName":"L50"},{"values":[1,2,2,1,1,1,16,2,1,1,2,1,4,1],"quality":"Less is better","isMain":false,"metricName":"L75"},{"values":["35.16","34.99","35.10","35.15","35.06","35.27","35.12","35.42","35.13","34.91","35.18","35.15","35.36","35.60"],"quality":"Equal","isMain":false,"metricName":"GC (%)"}]],["Predicted genes",[]],["Similarity statistics",[{"values":[0,0,0,0,0,0,0,0,0,0,0,0,0,0],"quality":"Equal","isMain":false,"metricName":"# similar correct contigs"},{"values":[0,0,0,0,0,0,0,0,0,0,0,0,0,0],"quality":"Equal","isMain":false,"metricName":"# similar misassembled blocks"}]],["Reference statistics",[{"values":[42653,42653,42653,42653,42653,42653,42653,42653,42653,42653,42653,42653,42653,42653],"quality":"Equal","isMain":false,"metricName":"Reference length"},{"values":[1,1,1,1,1,1,1,1,1,1,1,1,1,1],"quality":"Equal","isMain":false,"metricName":"Reference fragments"},{"values":["35.10","35.10","35.10","35.10","35.10","35.10","35.10","35.10","35.10","35.10","35.10","35.10","35.10","35.10"],"quality":"Equal","isMain":false,"metricName":"Reference GC (%)"}]]],"referenceName":"gi\_157325217\_ref\_NC","date":"18 July 2018, Wednesday, 16:26:05","order":[0,1,2,3,4,5,6,7,8,9,10,11,12,13],"assembliesNames":["ABySS\_127","ABySS\_63","CLC","IDBA\_UD","MEGAHIT","MIRA","Ray\_Meta","SOAPdenovo2","SPAdes","SPAdes\_meta","SPAdes\_sc","SPAdes\_sc\_careful","Velvet","Geneious"]},{"assembliesWithNs":null,"minContig":500,"report":[["Genome statistics",[{"values":["100.000","100.000","100.000","100.000","100.000","100.000","100.000","100.000","99.547","99.547","100.000","98.825"],"quality":"More is better","isMain":true,"metricName":"Genome fraction (%)"},{"values":["1.237","1.005","1.020","1.002","1.003","1.018","1.009","1.003","1.000","1.000","1.001","1.001"],"quality":"Less is better","isMain":true,"metricName":"Duplication ratio"},{"values":[32004,48399,48172,48271,48313,49058,48624,48299,47954,47954,48227,30328],"quality":"More is better","isMain":true,"metricName":"Largest alignment"},{"values":[59580,48399,49127,48271,48313,49058,48624,48299,47954,47954,48227,47641],"quality":"More is better","isMain":true,"metricName":"Total aligned length"},{"values":[59580,48399,48172,48271,48313,49058,48624,48299,47954,47954,48227,30328],"quality":"More is better","isMain":false,"metricName":"NG50"},{"values":[59580,48399,48172,48271,48313,49058,48624,48299,47954,47954,48227,9670],"quality":"More is better","isMain":false,"metricName":"NG75"},{"values":[32004,48399,48172,48271,48313,49058,48624,48299,47954,47954,48227,30328],"quality":"More is better","isMain":false,"metricName":"NA50"},{"values":[27576,48399,48172,48271,48313,49058,48624,48299,47954,47954,48227,9670],"quality":"More is better","isMain":false,"metricName":"NA75"},{"values":[32004,48399,48172,48271,48313,49058,48624,48299,47954,47954,48227,30328],"quality":"More is better","isMain":true,"metricName":"NGA50"},{"values":[27576,48399,48172,48271,48313,49058,48624,48299,47954,47954,48227,9670],"quality":"More is better","isMain":false,"metricName":"NGA75"},{"values":[1,1,1,1,1,1,1,1,1,1,1,1],"quality":"Less is better","isMain":false,"metricName":"LG50"},{"values":[1,1,1,1,1,1,1,1,1,1,1,2],"quality":"Less is better","isMain":false,"metricName":"LG75"},{"values":[1,1,1,1,1,1,1,1,1,1,1,1],"quality":"Less is better","isMain":false,"metricName":"LA50"},{"values":[2,1,1,1,1,1,1,1,1,1,1,2],"quality":"Less is better","isMain":false,"metricName":"LA75"},{"values":[1,1,1,1,1,1,1,1,1,1,1,1],"quality":"Less is better","isMain":true,"metricName":"LGA50"},{"values":[2,1,1,1,1,1,1,1,1,1,1,2],"quality":"Less is better","isMain":false,"metricName":"LGA75"}]],["Misassemblies",[{"values":[1,0,0,0,0,0,0,0,0,0,0,0],"quality":"Less is better","isMain":true,"metricName":"# misassemblies"},{"values":[1,0,0,0,0,0,0,0,0,0,0,0],"quality":"Less is better","isMain":false,"metricName":" # relocations"},{"values":[0,0,0,0,0,0,0,0,0,0,0,0],"quality":"Less is better","isMain":false,"metricName":" # translocations"},{"values":[0,0,0,0,0,0,0,0,0,0,0,0],"quality":"Less is better","isMain":false,"metricName":" # inversions"},{"values":[1,0,0,0,0,0,0,0,0,0,0,0],"quality":"Less is better","isMain":false,"metricName":"# misassembled contigs"},{"values":[59580,0,0,0,0,0,0,0,0,0,0,0],"quality":"Less is better","isMain":true,"metricName":"Misassembled contigs length"},{"values":[0,1,0,1,1,1,1,1,0,0,0,0],"quality":"Less is better","isMain":false,"metricName":"# local misassemblies"},{"values":[0,0,0,0,0,0,0,0,0,0,0,0],"quality":"Less is better","isMain":false,"metricName":"# unaligned mis. contigs"}]],["Unaligned",[{"values":[0,0,0,0,0,0,0,0,0,0,0,0],"quality":"Less is better","isMain":false,"metricName":"# fully unaligned contigs"},{"values":[0,0,0,0,0,0,0,0,0,0,0,0],"quality":"Less is better","isMain":false,"metricName":"Fully unaligned length"},{"values":[0,0,0,0,0,0,0,0,0,0,0,0],"quality":"Less is better","isMain":false,"metricName":"# partially unaligned contigs"},{"values":[0,0,0,0,0,0,0,0,0,0,0,0],"quality":"Less is better","isMain":false,"metricName":"Partially unaligned length"}]],["Mismatches",[{"values":[0,0,0,0,0,3,0,0,0,0,0,3],"quality":"Less is better","isMain":false,"metricName":"# mismatches"},{"values":[0,0,0,0,0,0,0,0,0,0,1,0],"quality":"Less is better","isMain":false,"metricName":"# indels"},{"values":[0,0,0,0,0,0,0,0,0,0,55,0],"quality":"Less is better","isMain":false,"metricName":"Indels length"},{"values":["0.00","0.00","0.00","0.00","0.00","6.23","0.00","0.00","0.00","0.00","0.00","6.30"],"quality":"Less is better","isMain":true,"metricName":"# mismatches per 100 kbp"},{"values":["0.00","0.00","0.00","0.00","0.00","0.00","0.00","0.00","0.00","0.00","2.08","0.00"],"quality":"Less is better","isMain":true,"metricName":"# indels per 100 kbp"},{"values":[0,0,0,0,0,0,0,0,0,0,0,0],"quality":"Less is better","isMain":false,"metricName":" # indels (<= 5 bp)"},{"values":[0,0,0,0,0,0,0,0,0,0,1,0],"quality":"Less is better","isMain":false,"metricName":" # indels (> 5 bp)"},{"values":[0,1,0,0,0,1,0,0,0,0,0,0],"quality":"Less is better","isMain":false,"metricName":"# N's"},{"values":["0.00","2.07","0.00","0.00","0.00","2.04","0.00","0.00","0.00","0.00","0.00","0.00"],"quality":"Less is better","isMain":true,"metricName":"# N's per 100 kbp"}]],["Statistics without reference",[{"values":[1,1,2,1,1,1,1,1,1,1,1,4],"quality":"Equal","isMain":true,"metricName":"# contigs"},{"values":[1,1,1,1,1,1,1,1,1,1,1,4],"quality":"Equal","isMain":false,"metricName":"# contigs (>= 1000 bp)"},{"values":[1,1,1,1,1,1,1,1,1,1,1,3],"quality":"Equal","isMain":false,"metricName":"# contigs (>= 5000 bp)"},{"values":[1,1,1,1,1,1,1,1,1,1,1,1],"quality":"Equal","isMain":false,"metricName":"# contigs (>= 10000 bp)"},{"values":[1,1,1,1,1,1,1,1,1,1,1,1],"quality":"Equal","isMain":false,"metricName":"# contigs (>= 25000 bp)"},{"values":[1,0,0,0,0,0,0,0,0,0,0,0],"quality":"Equal","isMain":false,"metricName":"# contigs (>= 50000 bp)"},{"values":[59580,48399,48172,48271,48313,49058,48624,48299,47954,47954,48227,30328],"quality":"More is better","isMain":true,"metricName":"Largest contig"},{"values":[59580,48399,49127,48271,48313,49058,48624,48299,47954,47954,48227,47641],"quality":"More is better","isMain":true,"metricName":"Total length"},{"values":[59580,48399,48172,48271,48313,49058,48624,48299,47954,47954,48227,47641],"quality":"More is better","isMain":true,"metricName":"Total length (>= 1000 bp)"},{"values":[59580,48399,48172,48271,48313,49058,48624,48299,47954,47954,48227,46225],"quality":"More is better","isMain":false,"metricName":"Total length (>= 5000 bp)"},{"values":[59580,48399,48172,48271,48313,49058,48624,48299,47954,47954,48227,30328],"quality":"More is better","isMain":true,"metricName":"Total length (>= 10000 bp)"},{"values":[59580,48399,48172,48271,48313,49058,48624,48299,47954,47954,48227,30328],"quality":"More is better","isMain":false,"metricName":"Total length (>= 25000 bp)"},{"values":[59580,0,0,0,0,0,0,0,0,0,0,0],"quality":"More is better","isMain":true,"metricName":"Total length (>= 50000 bp)"},{"values":[59580,48399,48172,48271,48313,49058,48624,48299,47954,47954,48227,30328],"quality":"More is better","isMain":false,"metricName":"N50"},{"values":[59580,48399,48172,48271,48313,49058,48624,48299,47954,47954,48227,9670],"quality":"More is better","isMain":false,"metricName":"N75"},{"values":[1,1,1,1,1,1,1,1,1,1,1,1],"quality":"Less is better","isMain":false,"metricName":"L50"},{"values":[1,1,1,1,1,1,1,1,1,1,1,2],"quality":"Less is better","isMain":false,"metricName":"L75"},{"values":["36.02","36.19","36.22","36.21","36.18","36.15","36.23","36.20","36.23","36.23","36.21","36.22"],"quality":"Equal","isMain":false,"metricName":"GC (%)"}]],["Predicted genes",[]],["Similarity statistics",[{"values":[0,0,0,0,0,0,0,0,0,0,0,0],"quality":"Equal","isMain":false,"metricName":"# similar correct contigs"},{"values":[0,0,0,0,0,0,0,0,0,0,0,0],"quality":"Equal","isMain":false,"metricName":"# similar misassembled blocks"}]],["Reference statistics",[{"values":[48172,48172,48172,48172,48172,48172,48172,48172,48172,48172,48172,48172],"quality":"Equal","isMain":false,"metricName":"Reference length"},{"values":[1,1,1,1,1,1,1,1,1,1,1,1],"quality":"Equal","isMain":false,"metricName":"Reference fragments"},{"values":["36.21","36.21","36.21","36.21","36.21","36.21","36.21","36.21","36.21","36.21","36.21","36.21"],"quality":"Equal","isMain":false,"metricName":"Reference GC (%)"}]]],"referenceName":"gi\_157325284\_ref\_NC","date":"18 July 2018, Wednesday, 16:26:17","order":[0,1,2,3,4,5,6,7,8,9,10,11],"assembliesNames":["ABySS\_127","ABySS\_63","CLC","IDBA\_UD","MEGAHIT","MIRA","Ray\_Meta","SPAdes","SPAdes\_meta","SPAdes\_sc","SPAdes\_sc\_careful","Velvet"]},{"assembliesWithNs":null,"minContig":500,"report":[["Genome statistics",[{"values":["100.000","100.000","99.872","100.000","100.000","100.000","19.753","100.000","100.000","100.000","100.000","99.833"],"quality":"More is better","isMain":true,"metricName":"Genome fraction (%)"},{"values":["1.002","1.000","1.004","1.004","1.010","1.002","1.000","1.004","1.000","1.000","1.000","1.000"],"quality":"Less is better","isMain":true,"metricName":"Duplication ratio"},{"values":[19409,35822,35776,35963,36173,35900,1217,35949,19285,19285,19285,19345],"quality":"More is better","isMain":true,"metricName":"Largest alignment"},{"values":[35890,35822,35776,35963,36173,35900,7076,35949,35822,35822,35822,35762],"quality":"More is better","isMain":true,"metricName":"Total aligned length"},{"values":[19409,35822,35921,35963,36173,35900,null,35949,19285,19285,19285,19345],"quality":"More is better","isMain":false,"metricName":"NG50"},{"values":[16481,35822,35921,35963,36173,35900,null,35949,16537,16537,16537,16417],"quality":"More is better","isMain":false,"metricName":"NG75"},{"values":[19409,35822,35776,35963,36173,35900,857,35949,19285,19285,19285,19345],"quality":"More is better","isMain":false,"metricName":"NA50"},{"values":[16481,35822,35776,35963,36173,35900,532,35949,16537,16537,16537,16417],"quality":"More is better","isMain":false,"metricName":"NA75"},{"values":[19409,35822,35776,35963,36173,35900,null,35949,19285,19285,19285,19345],"quality":"More is better","isMain":true,"metricName":"NGA50"},{"values":[16481,35822,35776,35963,36173,35900,null,35949,16537,16537,16537,16417],"quality":"More is better","isMain":false,"metricName":"NGA75"},{"values":[1,1,1,1,1,1,null,1,1,1,1,1],"quality":"Less is better","isMain":false,"metricName":"LG50"},{"values":[2,1,1,1,1,1,null,1,2,2,2,2],"quality":"Less is better","isMain":false,"metricName":"LG75"},{"values":[1,1,1,1,1,1,4,1,1,1,1,1],"quality":"Less is better","isMain":false,"metricName":"LA50"},{"values":[2,1,1,1,1,1,7,1,2,2,2,2],"quality":"Less is better","isMain":false,"metricName":"LA75"},{"values":[1,1,1,1,1,1,null,1,1,1,1,1],"quality":"Less is better","isMain":true,"metricName":"LGA50"},{"values":[2,1,1,1,1,1,null,1,2,2,2,2],"quality":"Less is better","isMain":false,"metricName":"LGA75"}]],["Misassemblies",[{"values":[0,0,0,0,0,0,0,0,0,0,0,0],"quality":"Less is better","isMain":true,"metricName":"# misassemblies"},{"values":[0,0,0,0,0,0,0,0,0,0,0,0],"quality":"Less is better","isMain":false,"metricName":" # relocations"},{"values":[0,0,0,0,0,0,0,0,0,0,0,0],"quality":"Less is better","isMain":false,"metricName":" # translocations"},{"values":[0,0,0,0,0,0,0,0,0,0,0,0],"quality":"Less is better","isMain":false,"metricName":" # inversions"},{"values":[0,0,0,0,0,0,0,0,0,0,0,0],"quality":"Less is better","isMain":false,"metricName":"# misassembled contigs"},{"values":[0,0,0,0,0,0,0,0,0,0,0,0],"quality":"Less is better","isMain":true,"metricName":"Misassembled contigs length"},{"values":[0,0,0,1,1,0,0,1,0,0,0,0],"quality":"Less is better","isMain":false,"metricName":"# local misassemblies"},{"values":[0,0,0,0,0,0,0,0,0,0,0,0],"quality":"Less is better","isMain":false,"metricName":"# unaligned mis. contigs"}]],["Unaligned",[{"values":[0,0,0,0,0,0,0,0,0,0,0,0],"quality":"Less is better","isMain":false,"metricName":"# fully unaligned contigs"},{"values":[0,0,0,0,0,0,0,0,0,0,0,0],"quality":"Less is better","isMain":false,"metricName":"Fully unaligned length"},{"values":[0,0,0,0,0,0,0,0,0,0,0,0],"quality":"Less is better","isMain":false,"metricName":"# partially unaligned contigs"},{"values":[0,0,0,0,0,0,0,0,0,0,0,0],"quality":"Less is better","isMain":false,"metricName":"Partially unaligned length"}]],["Mismatches",[{"values":[0,0,0,0,1,0,2,0,0,0,0,0],"quality":"Less is better","isMain":false,"metricName":"# mismatches"},{"values":[0,0,0,0,0,1,0,0,0,0,0,0],"quality":"Less is better","isMain":false,"metricName":"# indels"},{"values":[0,0,0,0,0,78,0,0,0,0,0,0],"quality":"Less is better","isMain":false,"metricName":"Indels length"},{"values":["0.00","0.00","0.00","0.00","2.79","0.00","28.26","0.00","0.00","0.00","0.00","0.00"],"quality":"Less is better","isMain":true,"metricName":"# mismatches per 100 kbp"},{"values":["0.00","0.00","0.00","0.00","0.00","2.79","0.00","0.00","0.00","0.00","0.00","0.00"],"quality":"Less is better","isMain":true,"metricName":"# indels per 100 kbp"},{"values":[0,0,0,0,0,0,0,0,0,0,0,0],"quality":"Less is better","isMain":false,"metricName":" # indels (<= 5 bp)"},{"values":[0,0,0,0,0,1,0,0,0,0,0,0],"quality":"Less is better","isMain":false,"metricName":" # indels (> 5 bp)"},{"values":[0,0,0,0,0,0,0,0,0,0,0,0],"quality":"Less is better","isMain":false,"metricName":"# N's"},{"values":["0.00","0.00","0.00","0.00","0.00","0.00","0.00","0.00","0.00","0.00","0.00","0.00"],"quality":"Less is better","isMain":true,"metricName":"# N's per 100 kbp"}]],["Statistics without reference",[{"values":[2,1,1,1,1,1,10,1,2,2,2,2],"quality":"Equal","isMain":true,"metricName":"# contigs"},{"values":[2,1,1,1,1,1,1,1,2,2,2,2],"quality":"Equal","isMain":false,"metricName":"# contigs (>= 1000 bp)"},{"values":[2,1,1,1,1,1,0,1,2,2,2,2],"quality":"Equal","isMain":false,"metricName":"# contigs (>= 5000 bp)"},{"values":[2,1,1,1,1,1,0,1,2,2,2,2],"quality":"Equal","isMain":false,"metricName":"# contigs (>= 10000 bp)"},{"values":[0,1,1,1,1,1,0,1,0,0,0,0],"quality":"Equal","isMain":false,"metricName":"# contigs (>= 25000 bp)"},{"values":[0,0,0,0,0,0,0,0,0,0,0,0],"quality":"Equal","isMain":false,"metricName":"# contigs (>= 50000 bp)"},{"values":[19409,35822,35921,35963,36173,35900,1217,35949,19285,19285,19285,19345],"quality":"More is better","isMain":true,"metricName":"Largest contig"},{"values":[35890,35822,35921,35963,36173,35900,7076,35949,35822,35822,35822,35762],"quality":"More is better","isMain":true,"metricName":"Total length"},{"values":[35890,35822,35921,35963,36173,35900,1217,35949,35822,35822,35822,35762],"quality":"More is better","isMain":true,"metricName":"Total length (>= 1000 bp)"},{"values":[35890,35822,35921,35963,36173,35900,0,35949,35822,35822,35822,35762],"quality":"More is better","isMain":false,"metricName":"Total length (>= 5000 bp)"},{"values":[35890,35822,35921,35963,36173,35900,0,35949,35822,35822,35822,35762],"quality":"More is better","isMain":true,"metricName":"Total length (>= 10000 bp)"},{"values":[0,35822,35921,35963,36173,35900,0,35949,0,0,0,0],"quality":"More is better","isMain":false,"metricName":"Total length (>= 25000 bp)"},{"values":[0,0,0,0,0,0,0,0,0,0,0,0],"quality":"More is better","isMain":true,"metricName":"Total length (>= 50000 bp)"},{"values":[19409,35822,35921,35963,36173,35900,857,35949,19285,19285,19285,19345],"quality":"More is better","isMain":false,"metricName":"N50"},{"values":[16481,35822,35921,35963,36173,35900,532,35949,16537,16537,16537,16417],"quality":"More is better","isMain":false,"metricName":"N75"},{"values":[1,1,1,1,1,1,4,1,1,1,1,1],"quality":"Less is better","isMain":false,"metricName":"L50"},{"values":[2,1,1,1,1,1,7,1,2,2,2,2],"quality":"Less is better","isMain":false,"metricName":"L75"},{"values":["40.77","40.82","40.83","40.83","40.83","40.77","40.52","40.80","40.82","40.82","40.82","40.86"],"quality":"Equal","isMain":false,"metricName":"GC (%)"}]],["Predicted genes",[]],["Similarity statistics",[{"values":[2,2,0,0,0,2,0,0,2,2,2,2],"quality":"Equal","isMain":false,"metricName":"# similar correct contigs"},{"values":[0,0,0,0,0,0,0,0,0,0,0,0],"quality":"Equal","isMain":false,"metricName":"# similar misassembled blocks"}]],["Reference statistics",[{"values":[35822,35822,35822,35822,35822,35822,35822,35822,35822,35822,35822,35822],"quality":"Equal","isMain":false,"metricName":"Reference length"},{"values":[1,1,1,1,1,1,1,1,1,1,1,1],"quality":"Equal","isMain":false,"metricName":"Reference fragments"},{"values":["40.82","40.82","40.82","40.82","40.82","40.82","40.82","40.82","40.82","40.82","40.82","40.82"],"quality":"Equal","isMain":false,"metricName":"Reference GC (%)"}]]],"referenceName":"gi\_157325365\_ref\_NC","date":"18 July 2018, Wednesday, 16:26:29","order":[0,1,2,3,4,5,6,7,8,9,10,11],"assembliesNames":["ABySS\_63","CLC","IDBA\_UD","MEGAHIT","MIRA","Ray\_Meta","SOAPdenovo2","SPAdes","SPAdes\_meta","SPAdes\_sc","SPAdes\_sc\_careful","Velvet"]},{"assembliesWithNs":null,"minContig":500,"report":[["Genome statistics",[{"values":["100.000","100.000","96.860","97.477","99.449","95.680","100.000","97.810","100.000","97.432","97.432","89.191","100.000"],"quality":"More is better","isMain":true,"metricName":"Genome fraction (%)"},{"values":["1.204","1.029","1.003","1.009","1.014","1.048","1.085","1.000","1.002","1.003","1.000","1.001","1.105"],"quality":"Less is better","isMain":true,"metricName":"Duplication ratio"},{"values":[37090,33869,23267,22104,22188,35652,34471,37289,38199,33733,37145,20045,30908],"quality":"More is better","isMain":true,"metricName":"Largest alignment"},{"values":[45898,39178,37032,37485,38456,38243,41348,37289,38199,37255,37145,34053,42145],"quality":"More is better","isMain":true,"metricName":"Total aligned length"},{"values":[37090,33919,23267,22104,22188,36517,34471,37289,65947,33733,37145,20045,192212],"quality":"More is better","isMain":false,"metricName":"NG50"},{"values":[37090,33919,7933,3599,7187,36517,34471,37289,65947,33733,37145,1062,192212],"quality":"More is better","isMain":false,"metricName":"NG75"},{"values":[37090,33869,23267,22104,22188,2591,34471,37289,38199,33733,37145,20045,null],"quality":"More is better","isMain":false,"metricName":"NA50"},{"values":[37090,33869,7933,3599,7187,null,34471,37289,null,33733,37145,1267,null],"quality":"More is better","isMain":false,"metricName":"NA75"},{"values":[37090,33869,23267,22104,22188,35652,34471,37289,38199,33733,37145,20045,30908],"quality":"More is better","isMain":true,"metricName":"NGA50"},{"values":[37090,33869,7933,3599,7187,35652,34471,37289,38199,33733,37145,1062,30908],"quality":"More is better","isMain":false,"metricName":"NGA75"},{"values":[1,1,1,1,1,1,1,1,1,1,1,1,1],"quality":"Less is better","isMain":false,"metricName":"LG50"},{"values":[1,1,2,3,2,1,1,1,1,1,1,7,1],"quality":"Less is better","isMain":false,"metricName":"LG75"},{"values":[1,1,1,1,1,2,1,1,1,1,1,1,null],"quality":"Less is better","isMain":false,"metricName":"LA50"},{"values":[1,1,2,3,2,null,1,1,null,1,1,5,null],"quality":"Less is better","isMain":false,"metricName":"LA75"},{"values":[1,1,1,1,1,1,1,1,1,1,1,1,1],"quality":"Less is better","isMain":true,"metricName":"LGA50"},{"values":[1,1,2,3,2,1,1,1,1,1,1,7,1],"quality":"Less is better","isMain":false,"metricName":"LGA75"}]],["Misassemblies",[{"values":[0,0,0,0,0,0,0,0,0,0,0,0,0],"quality":"Less is better","isMain":true,"metricName":"# misassemblies"},{"values":[0,0,0,0,0,0,0,0,0,0,0,0,0],"quality":"Less is better","isMain":false,"metricName":" # relocations"},{"values":[0,0,0,0,0,0,0,0,0,0,0,0,0],"quality":"Less is better","isMain":false,"metricName":" # translocations"},{"values":[0,0,0,0,0,0,0,0,0,0,0,0,0],"quality":"Less is better","isMain":false,"metricName":" # inversions"},{"values":[0,0,0,0,0,0,0,0,0,0,0,0,0],"quality":"Less is better","isMain":false,"metricName":"# misassembled contigs"},{"values":[0,0,0,0,0,0,0,0,0,0,0,0,0],"quality":"Less is better","isMain":true,"metricName":"Misassembled contigs length"},{"values":[0,1,0,0,0,0,0,0,0,0,0,0,0],"quality":"Less is better","isMain":false,"metricName":"# local misassemblies"},{"values":[0,0,0,0,0,0,0,0,0,0,0,0,1],"quality":"Less is better","isMain":false,"metricName":"# unaligned mis. contigs"}]],["Unaligned",[{"values":[0,0,0,0,0,0,0,0,0,0,0,0,0],"quality":"Less is better","isMain":false,"metricName":"# fully unaligned contigs"},{"values":[0,0,0,0,0,0,0,0,0,0,0,0,0],"quality":"Less is better","isMain":false,"metricName":"Fully unaligned length"},{"values":[0,0,0,0,0,1,0,0,1,0,0,0,1],"quality":"Less is better","isMain":false,"metricName":"# partially unaligned contigs"},{"values":[0,0,0,0,0,33926,0,0,27748,0,0,0,150067],"quality":"Less is better","isMain":false,"metricName":"Partially unaligned length"}]],["Mismatches",[{"values":[0,0,0,0,1,6,0,0,13,9,0,2,2],"quality":"Less is better","isMain":false,"metricName":"# mismatches"},{"values":[0,0,0,0,0,0,0,0,1,0,0,0,0],"quality":"Less is better","isMain":false,"metricName":"# indels"},{"values":[0,0,0,0,0,0,0,0,75,0,0,0,0],"quality":"Less is better","isMain":false,"metricName":"Indels length"},{"values":["0.00","0.00","0.00","0.00","2.64","16.45","0.00","0.00","34.10","24.23","0.00","5.88","5.25"],"quality":"Less is better","isMain":true,"metricName":"# mismatches per 100 kbp"},{"values":["0.00","0.00","0.00","0.00","0.00","0.00","0.00","0.00","2.62","0.00","0.00","0.00","0.00"],"quality":"Less is better","isMain":true,"metricName":"# indels per 100 kbp"},{"values":[0,0,0,0,0,0,0,0,0,0,0,0,0],"quality":"Less is better","isMain":false,"metricName":" # indels (<= 5 bp)"},{"values":[0,0,0,0,0,0,0,0,1,0,0,0,0],"quality":"Less is better","isMain":false,"metricName":" # indels (> 5 bp)"},{"values":[0,165,0,0,0,0,0,0,0,0,0,0,17],"quality":"Less is better","isMain":false,"metricName":"# N's"},{"values":["0.00","420.62","0.00","0.00","0.00","0.00","0.00","0.00","0.00","0.00","0.00","0.00","8.84"],"quality":"Less is better","isMain":true,"metricName":"# N's per 100 kbp"}]],["Statistics without reference",[{"values":[3,2,4,7,6,2,2,1,1,3,1,13,1],"quality":"Equal","isMain":true,"metricName":"# contigs"},{"values":[3,2,4,6,5,2,2,1,1,2,1,8,1],"quality":"Equal","isMain":false,"metricName":"# contigs (>= 1000 bp)"},{"values":[1,2,2,1,2,2,2,1,1,1,1,1,1],"quality":"Equal","isMain":false,"metricName":"# contigs (>= 5000 bp)"},{"values":[1,1,1,1,1,2,1,1,1,1,1,1,1],"quality":"Equal","isMain":false,"metricName":"# contigs (>= 10000 bp)"},{"values":[1,1,0,0,0,2,1,1,1,1,1,0,1],"quality":"Equal","isMain":false,"metricName":"# contigs (>= 25000 bp)"},{"values":[0,0,0,0,0,0,0,0,1,0,0,0,1],"quality":"Equal","isMain":false,"metricName":"# contigs (>= 50000 bp)"},{"values":[37090,33919,23267,22104,22188,36517,34471,37289,65947,33733,37145,20045,192212],"quality":"More is better","isMain":true,"metricName":"Largest contig"},{"values":[45898,39228,37032,37485,38456,72169,41348,37289,65947,37255,37145,34053,192212],"quality":"More is better","isMain":true,"metricName":"Total length"},{"values":[45898,39228,37032,36597,37484,72169,41348,37289,65947,36635,37145,29928,192212],"quality":"More is better","isMain":true,"metricName":"Total length (>= 1000 bp)"},{"values":[37090,39228,31200,22104,29375,72169,41348,37289,65947,33733,37145,20045,192212],"quality":"More is better","isMain":false,"metricName":"Total length (>= 5000 bp)"},{"values":[37090,33919,23267,22104,22188,72169,34471,37289,65947,33733,37145,20045,192212],"quality":"More is better","isMain":true,"metricName":"Total length (>= 10000 bp)"},{"values":[37090,33919,0,0,0,72169,34471,37289,65947,33733,37145,0,192212],"quality":"More is better","isMain":false,"metricName":"Total length (>= 25000 bp)"},{"values":[0,0,0,0,0,0,0,0,65947,0,0,0,192212],"quality":"More is better","isMain":true,"metricName":"Total length (>= 50000 bp)"},{"values":[37090,33919,23267,22104,22188,36517,34471,37289,65947,33733,37145,20045,192212],"quality":"More is better","isMain":false,"metricName":"N50"},{"values":[37090,33919,7933,3599,7187,35652,34471,37289,65947,33733,37145,1267,192212],"quality":"More is better","isMain":false,"metricName":"N75"},{"values":[1,1,1,1,1,1,1,1,1,1,1,1,1],"quality":"Less is better","isMain":false,"metricName":"L50"},{"values":[1,1,2,3,2,2,1,1,1,1,1,5,1],"quality":"Less is better","isMain":false,"metricName":"L75"},{"values":["35.61","35.53","35.43","35.44","35.52","35.23","35.58","35.45","35.87","35.44","35.46","35.50","35.60"],"quality":"Equal","isMain":false,"metricName":"GC (%)"}]],["Predicted genes",[]],["Similarity statistics",[{"values":[0,0,0,0,0,0,0,0,0,0,0,0,0],"quality":"Equal","isMain":false,"metricName":"# similar correct contigs"},{"values":[0,0,0,0,0,0,0,0,0,0,0,0,0],"quality":"Equal","isMain":false,"metricName":"# similar misassembled blocks"}]],["Reference statistics",[{"values":[38124,38124,38124,38124,38124,38124,38124,38124,38124,38124,38124,38124,38124],"quality":"Equal","isMain":false,"metricName":"Reference length"},{"values":[1,1,1,1,1,1,1,1,1,1,1,1,1],"quality":"Equal","isMain":false,"metricName":"Reference fragments"},{"values":["35.50","35.50","35.50","35.50","35.50","35.50","35.50","35.50","35.50","35.50","35.50","35.50","35.50"],"quality":"Equal","isMain":false,"metricName":"Reference GC (%)"}]]],"referenceName":"gi\_157325422\_ref\_NC","date":"18 July 2018, Wednesday, 16:26:41","order":[0,1,2,3,4,5,6,7,8,9,10,11,12],"assembliesNames":["ABySS\_127","ABySS\_63","CLC","IDBA\_UD","MEGAHIT","MIRA","Ray\_Meta","SPAdes","SPAdes\_meta","SPAdes\_sc","SPAdes\_sc\_careful","Velvet","Geneious"]},{"assembliesWithNs":null,"minContig":500,"report":[["Genome statistics",[{"values":["55.175","16.879","40.402","43.228","48.856","94.015","10.116","50.438","6.658","20.637","21.415","14.069","79.191"],"quality":"More is better","isMain":true,"metricName":"Genome fraction (%)"},{"values":["1.026","1.007","1.034","1.012","1.053","1.130","1.224","1.040","1.016","1.002","1.002","1.004","1.069"],"quality":"Less is better","isMain":true,"metricName":"Duplication ratio"},{"values":[2576,1260,3705,2516,2591,13929,1085,5596,1410,2006,2006,1767,18867],"quality":"More is better","isMain":true,"metricName":"Largest alignment"},{"values":[29034,8717,21009,22440,26048,54332,5187,26745,3469,10600,10999,7214,43391],"quality":"More is better","isMain":true,"metricName":"Total aligned length"},{"values":[586,null,null,null,556,4965,null,669,null,null,null,null,8012],"quality":"More is better","isMain":false,"metricName":"NG50"},{"values":[null,null,null,null,null,4408,null,null,null,null,null,null,6323],"quality":"More is better","isMain":false,"metricName":"NG75"},{"values":[1236,916,1424,1213,1110,4657,445,2572,782,1139,1139,730,7016],"quality":"More is better","isMain":false,"metricName":"NA50"},{"values":[750,604,698,772,616,3364,413,1752,658,780,594,609,5702],"quality":"More is better","isMain":false,"metricName":"NA75"},{"values":[586,null,null,null,548,4657,null,604,null,null,null,null,7016],"quality":"More is better","isMain":true,"metricName":"NGA50"},{"values":[null,null,null,null,null,3582,null,null,null,null,null,null,5523],"quality":"More is better","isMain":false,"metricName":"NGA75"},{"values":[23,null,null,null,24,4,null,11,null,null,null,null,2],"quality":"Less is better","isMain":false,"metricName":"LG50"},{"values":[null,null,null,null,null,6,null,null,null,null,null,null,4],"quality":"Less is better","isMain":false,"metricName":"LG75"},{"values":[9,5,5,7,9,4,5,4,2,4,4,3,2],"quality":"Less is better","isMain":false,"metricName":"LA50"},{"values":[17,8,10,12,17,8,8,8,3,7,8,6,4],"quality":"Less is better","isMain":false,"metricName":"LA75"},{"values":[23,null,null,null,26,4,null,13,null,null,null,null,2],"quality":"Less is better","isMain":true,"metricName":"LGA50"},{"values":[null,null,null,null,null,7,null,null,null,null,null,null,5],"quality":"Less is better","isMain":false,"metricName":"LGA75"}]],["Misassemblies",[{"values":[0,0,0,0,0,0,0,0,0,0,0,0,0],"quality":"Less is better","isMain":true,"metricName":"# misassemblies"},{"values":[0,0,0,0,0,0,0,0,0,0,0,0,0],"quality":"Less is better","isMain":false,"metricName":" # relocations"},{"values":[0,0,0,0,0,0,0,0,0,0,0,0,0],"quality":"Less is better","isMain":false,"metricName":" # translocations"},{"values":[0,0,0,0,0,0,0,0,0,0,0,0,0],"quality":"Less is better","isMain":false,"metricName":" # inversions"},{"values":[0,0,0,0,0,0,0,0,0,0,0,0,0],"quality":"Less is better","isMain":false,"metricName":"# misassembled contigs"},{"values":[0,0,0,0,0,0,0,0,0,0,0,0,0],"quality":"Less is better","isMain":true,"metricName":"Misassembled contigs length"},{"values":[0,0,0,0,2,5,0,1,0,0,0,0,7],"quality":"Less is better","isMain":false,"metricName":"# local misassemblies"},{"values":[0,0,0,0,0,0,0,1,0,0,0,0,0],"quality":"Less is better","isMain":false,"metricName":"# unaligned mis. contigs"}]],["Unaligned",[{"values":[0,0,0,0,0,0,0,0,0,0,0,0,0],"quality":"Less is better","isMain":false,"metricName":"# fully unaligned contigs"},{"values":[0,0,0,0,0,0,0,0,0,0,0,0,0],"quality":"Less is better","isMain":false,"metricName":"Fully unaligned length"},{"values":[0,0,0,0,1,4,0,1,0,0,0,0,3],"quality":"Less is better","isMain":false,"metricName":"# partially unaligned contigs"},{"values":[0,0,0,0,621,3219,0,1308,0,0,0,0,3295],"quality":"Less is better","isMain":false,"metricName":"Partially unaligned length"}]],["Mismatches",[{"values":[58,1,163,85,96,122,0,111,41,46,44,3,136],"quality":"Less is better","isMain":false,"metricName":"# mismatches"},{"values":[1,0,3,1,4,7,4,3,0,0,0,0,14],"quality":"Less is better","isMain":false,"metricName":"# indels"},{"values":[1,0,3,1,6,8,186,3,0,0,0,0,17],"quality":"Less is better","isMain":false,"metricName":"Indels length"},{"values":["205.00","11.55","786.79","383.47","383.20","253.07","0.00","429.18","1200.94","434.70","400.69","41.59","334.92"],"quality":"Less is better","isMain":true,"metricName":"# mismatches per 100 kbp"},{"values":["3.53","0.00","14.48","4.51","15.97","14.52","77.12","11.60","0.00","0.00","0.00","0.00","34.48"],"quality":"Less is better","isMain":true,"metricName":"# indels per 100 kbp"},{"values":[1,0,3,1,4,7,0,3,0,0,0,0,14],"quality":"Less is better","isMain":false,"metricName":" # indels (<= 5 bp)"},{"values":[0,0,0,0,0,0,4,0,0,0,0,0,0],"quality":"Less is better","isMain":false,"metricName":" # indels (> 5 bp)"},{"values":[0,0,0,0,0,47,1065,0,0,0,0,0,304],"quality":"Less is better","isMain":false,"metricName":"# N's"},{"values":["0.00","0.00","0.00","0.00","0.00","81.43","16776.94","0.00","0.00","0.00","0.00","0.00","651.16"],"quality":"Less is better","isMain":true,"metricName":"# N's per 100 kbp"}]],["Statistics without reference",[{"values":[29,11,18,21,26,13,9,15,4,11,12,9,5],"quality":"Equal","isMain":true,"metricName":"# contigs"},{"values":[10,1,7,8,10,11,1,9,1,5,5,2,5],"quality":"Equal","isMain":false,"metricName":"# contigs (>= 1000 bp)"},{"values":[0,0,0,0,0,3,0,1,0,0,0,0,5],"quality":"Equal","isMain":false,"metricName":"# contigs (>= 5000 bp)"},{"values":[0,0,0,0,0,1,0,0,0,0,0,0,1],"quality":"Equal","isMain":false,"metricName":"# contigs (>= 10000 bp)"},{"values":[0,0,0,0,0,0,0,0,0,0,0,0,0],"quality":"Equal","isMain":false,"metricName":"# contigs (>= 25000 bp)"},{"values":[0,0,0,0,0,0,0,0,0,0,0,0,0],"quality":"Equal","isMain":false,"metricName":"# contigs (>= 50000 bp)"},{"values":[2576,1260,3705,2516,3212,13929,1085,5737,1410,2006,2006,1797,18867],"quality":"More is better","isMain":true,"metricName":"Largest contig"},{"values":[29034,8717,21418,22441,26990,57716,6348,28199,3469,10600,10999,7244,46686],"quality":"More is better","isMain":true,"metricName":"Total length"},{"values":[15973,1260,14514,13411,16467,55838,1085,24216,1410,6684,6735,2989,46686],"quality":"More is better","isMain":true,"metricName":"Total length (>= 1000 bp)"},{"values":[0,0,0,0,0,25485,0,5737,0,0,0,0,46686],"quality":"More is better","isMain":false,"metricName":"Total length (>= 5000 bp)"},{"values":[0,0,0,0,0,13929,0,0,0,0,0,0,18867],"quality":"More is better","isMain":true,"metricName":"Total length (>= 10000 bp)"},{"values":[0,0,0,0,0,0,0,0,0,0,0,0,0],"quality":"More is better","isMain":false,"metricName":"Total length (>= 25000 bp)"},{"values":[0,0,0,0,0,0,0,0,0,0,0,0,0],"quality":"More is better","isMain":true,"metricName":"Total length (>= 50000 bp)"},{"values":[1236,916,1424,1213,1177,4965,667,2572,782,1139,1139,730,8012],"quality":"More is better","isMain":false,"metricName":"N50"},{"values":[750,604,738,772,749,4276,595,1752,658,780,594,609,6323],"quality":"More is better","isMain":false,"metricName":"N75"},{"values":[9,5,5,7,8,4,4,4,2,4,4,3,2],"quality":"Less is better","isMain":false,"metricName":"L50"},{"values":[17,8,9,12,15,7,7,8,3,7,8,6,4],"quality":"Less is better","isMain":false,"metricName":"L75"},{"values":["64.20","63.20","63.57","63.79","63.67","63.41","64.57","63.32","65.72","64.26","64.18","64.90","63.58"],"quality":"Equal","isMain":false,"metricName":"GC (%)"}]],["Predicted genes",[]],["Similarity statistics",[{"values":[0,0,0,0,0,0,0,0,0,0,0,0,0],"quality":"Equal","isMain":false,"metricName":"# similar correct contigs"},{"values":[0,0,0,0,0,0,0,0,0,0,0,0,0],"quality":"Equal","isMain":false,"metricName":"# similar misassembled blocks"}]],["Reference statistics",[{"values":[51277,51277,51277,51277,51277,51277,51277,51277,51277,51277,51277,51277,51277],"quality":"Equal","isMain":false,"metricName":"Reference length"},{"values":[1,1,1,1,1,1,1,1,1,1,1,1,1],"quality":"Equal","isMain":false,"metricName":"Reference fragments"},{"values":["63.70","63.70","63.70","63.70","63.70","63.70","63.70","63.70","63.70","63.70","63.70","63.70","63.70"],"quality":"Equal","isMain":false,"metricName":"Reference GC (%)"}]]],"referenceName":"gi\_157786393\_ref\_NC","date":"18 July 2018, Wednesday, 16:26:53","order":[0,1,2,3,4,5,6,7,8,9,10,11,12],"assembliesNames":["ABySS\_127","ABySS\_63","CLC","IDBA\_UD","MEGAHIT","MIRA","SOAPdenovo2","SPAdes","SPAdes\_meta","SPAdes\_sc","SPAdes\_sc\_careful","Velvet","Geneious"]},{"assembliesWithNs":null,"minContig":500,"report":[["Genome statistics",[{"values":["71.556","44.396","80.411","70.809","74.468","92.098","18.119","64.017","71.993","45.384","57.760","39.842","81.444"],"quality":"More is better","isMain":true,"metricName":"Genome fraction (%)"},{"values":["1.043","1.006","1.020","1.037","1.116","1.233","1.399","1.072","1.041","1.047","1.014","1.006","1.230"],"quality":"Less is better","isMain":true,"metricName":"Duplication ratio"},{"values":[2535,2525,4109,3455,3992,13880,1161,7715,6145,3068,2564,1881,17658],"quality":"More is better","isMain":true,"metricName":"Largest alignment"},{"values":[27129,16226,29846,26716,30269,40903,6614,24956,26707,16871,21322,14590,36460],"quality":"More is better","isMain":true,"metricName":"Total aligned length"},{"values":[907,null,1435,1329,1445,5962,null,7254,1906,null,863,null,19751],"quality":"More is better","isMain":false,"metricName":"NG50"},{"values":[null,null,775,null,731,4243,null,2903,null,null,null,null,10568],"quality":"More is better","isMain":false,"metricName":"NG75"},{"values":[1104,865,1777,1950,1749,4243,454,2712,2991,1289,1292,1061,9810],"quality":"More is better","isMain":false,"metricName":"NA50"},{"values":[775,652,1084,1119,1061,2134,null,null,1040,899,899,852,3611],"quality":"More is better","isMain":false,"metricName":"NA75"},{"values":[907,null,1435,1329,1445,5962,null,2712,1906,null,863,null,9810],"quality":"More is better","isMain":true,"metricName":"NGA50"},{"values":[null,null,775,null,731,3782,null,null,null,null,null,null,9810],"quality":"More is better","isMain":false,"metricName":"NGA75"},{"values":[14,null,8,8,8,2,null,2,5,null,13,null,1],"quality":"Less is better","isMain":false,"metricName":"LG50"},{"values":[null,null,17,null,17,4,null,4,null,null,null,null,2],"quality":"Less is better","isMain":false,"metricName":"LG75"},{"values":[9,7,6,5,6,3,8,4,3,4,6,6,2],"quality":"Less is better","isMain":false,"metricName":"LA50"},{"values":[17,12,11,10,12,7,null,null,7,8,11,9,4],"quality":"Less is better","isMain":false,"metricName":"LA75"},{"values":[14,null,8,8,8,2,null,4,5,null,13,null,2],"quality":"Less is better","isMain":true,"metricName":"LGA50"},{"values":[null,null,17,null,17,4,null,null,null,null,null,null,2],"quality":"Less is better","isMain":false,"metricName":"LGA75"}]],["Misassemblies",[{"values":[0,0,0,0,0,0,0,1,0,0,0,0,0],"quality":"Less is better","isMain":true,"metricName":"# misassemblies"},{"values":[0,0,0,0,0,0,0,1,0,0,0,0,0],"quality":"Less is better","isMain":false,"metricName":" # relocations"},{"values":[0,0,0,0,0,0,0,0,0,0,0,0,0],"quality":"Less is better","isMain":false,"metricName":" # translocations"},{"values":[0,0,0,0,0,0,0,0,0,0,0,0,0],"quality":"Less is better","isMain":false,"metricName":" # inversions"},{"values":[0,0,0,0,0,0,0,1,0,0,0,0,0],"quality":"Less is better","isMain":false,"metricName":"# misassembled contigs"},{"values":[0,0,0,0,0,0,0,7254,0,0,0,0,0],"quality":"Less is better","isMain":true,"metricName":"Misassembled contigs length"},{"values":[0,0,0,0,0,2,0,0,0,0,0,0,2],"quality":"Less is better","isMain":false,"metricName":"# local misassemblies"},{"values":[0,0,0,0,0,0,2,0,0,0,0,0,0],"quality":"Less is better","isMain":false,"metricName":"# unaligned mis. contigs"}]],["Unaligned",[{"values":[0,0,0,0,0,0,0,0,0,0,0,0,0],"quality":"Less is better","isMain":false,"metricName":"# fully unaligned contigs"},{"values":[0,0,0,0,0,0,0,0,0,0,0,0,0],"quality":"Less is better","isMain":false,"metricName":"Fully unaligned length"},{"values":[0,0,0,0,0,2,2,2,0,0,0,0,5],"quality":"Less is better","isMain":false,"metricName":"# partially unaligned contigs"},{"values":[0,0,0,0,0,4190,1543,10294,0,0,0,0,7379],"quality":"Less is better","isMain":false,"metricName":"Partially unaligned length"}]],["Mismatches",[{"values":[8,33,76,106,127,303,6,303,195,76,99,33,413],"quality":"Less is better","isMain":false,"metricName":"# mismatches"},{"values":[0,0,0,0,0,2,7,3,0,0,0,0,6],"quality":"Less is better","isMain":false,"metricName":"# indels"},{"values":[0,0,0,0,0,2,352,5,0,0,0,0,8],"quality":"Less is better","isMain":false,"metricName":"Indels length"},{"values":["30.71","204.16","259.59","411.16","468.41","903.61","90.95","1299.98","743.93","459.94","470.76","227.49","1392.78"],"quality":"Less is better","isMain":true,"metricName":"# mismatches per 100 kbp"},{"values":["0.00","0.00","0.00","0.00","0.00","5.96","106.11","12.87","0.00","0.00","0.00","0.00","20.23"],"quality":"Less is better","isMain":true,"metricName":"# indels per 100 kbp"},{"values":[0,0,0,0,0,2,0,3,0,0,0,0,6],"quality":"Less is better","isMain":false,"metricName":" # indels (<= 5 bp)"},{"values":[0,0,0,0,0,0,7,0,0,0,0,0,0],"quality":"Less is better","isMain":false,"metricName":" # indels (> 5 bp)"},{"values":[0,0,0,0,0,34,2929,0,428,428,0,0,295],"quality":"Less is better","isMain":false,"metricName":"# N's"},{"values":["0.00","0.00","0.00","0.00","0.00","74.65","27195.91","0.00","1569.26","2474.13","0.00","0.00","672.92"],"quality":"Less is better","isMain":true,"metricName":"# N's per 100 kbp"}]],["Statistics without reference",[{"values":[27,19,20,19,21,13,13,7,14,13,18,14,5],"quality":"Equal","isMain":true,"metricName":"# contigs"},{"values":[12,4,13,10,13,8,2,6,8,7,9,6,5],"quality":"Equal","isMain":false,"metricName":"# contigs (>= 1000 bp)"},{"values":[0,0,0,0,0,2,0,2,2,0,0,0,4],"quality":"Equal","isMain":false,"metricName":"# contigs (>= 5000 bp)"},{"values":[0,0,0,0,0,1,0,1,0,0,0,0,2],"quality":"Equal","isMain":false,"metricName":"# contigs (>= 10000 bp)"},{"values":[0,0,0,0,0,0,0,0,0,0,0,0,0],"quality":"Equal","isMain":false,"metricName":"# contigs (>= 25000 bp)"},{"values":[0,0,0,0,0,0,0,0,0,0,0,0,0],"quality":"Equal","isMain":false,"metricName":"# contigs (>= 50000 bp)"},{"values":[2535,2525,4109,3455,3992,15973,1161,16634,6573,3496,2564,1881,19751],"quality":"More is better","isMain":true,"metricName":"Largest contig"},{"values":[27181,16262,29848,26729,30270,45547,10770,35279,27274,17299,21322,14590,43839],"quality":"More is better","isMain":true,"metricName":"Total length"},{"values":[16755,6088,24557,20619,24369,41684,2301,34748,22869,12921,14768,8327,43839],"quality":"More is better","isMain":true,"metricName":"Total length (>= 1000 bp)"},{"values":[0,0,0,0,0,21935,0,23888,12275,0,0,0,41348],"quality":"More is better","isMain":false,"metricName":"Total length (>= 5000 bp)"},{"values":[0,0,0,0,0,15973,0,16634,0,0,0,0,30319],"quality":"More is better","isMain":true,"metricName":"Total length (>= 10000 bp)"},{"values":[0,0,0,0,0,0,0,0,0,0,0,0,0],"quality":"More is better","isMain":false,"metricName":"Total length (>= 25000 bp)"},{"values":[0,0,0,0,0,0,0,0,0,0,0,0,0],"quality":"More is better","isMain":true,"metricName":"Total length (>= 50000 bp)"},{"values":[1104,865,1777,1950,1749,4322,874,7254,2991,1289,1292,1061,10568],"quality":"More is better","isMain":false,"metricName":"N50"},{"values":[775,652,1084,1119,1061,3782,799,2913,1600,899,899,852,5708],"quality":"More is better","isMain":false,"metricName":"N75"},{"values":[9,7,6,5,6,3,6,2,3,4,6,6,2],"quality":"Less is better","isMain":false,"metricName":"L50"},{"values":[17,12,11,10,12,5,9,3,6,8,11,9,3],"quality":"Less is better","isMain":false,"metricName":"L75"},{"values":["64.07","62.69","64.12","64.18","63.84","63.90","62.06","63.93","64.27","63.71","63.79","63.47","63.65"],"quality":"Equal","isMain":false,"metricName":"GC (%)"}]],["Predicted genes",[]],["Similarity statistics",[{"values":[0,0,0,0,0,0,0,0,0,0,0,0,0],"quality":"Equal","isMain":false,"metricName":"# similar correct contigs"},{"values":[0,0,0,0,0,0,0,0,0,0,0,0,0],"quality":"Equal","isMain":false,"metricName":"# similar misassembled blocks"}]],["Reference statistics",[{"values":[36409,36409,36409,36409,36409,36409,36409,36409,36409,36409,36409,36409,36409],"quality":"Equal","isMain":false,"metricName":"Reference length"},{"values":[1,1,1,1,1,1,1,1,1,1,1,1,1],"quality":"Equal","isMain":false,"metricName":"Reference fragments"},{"values":["64.19","64.19","64.19","64.19","64.19","64.19","64.19","64.19","64.19","64.19","64.19","64.19","64.19"],"quality":"Equal","isMain":false,"metricName":"Reference GC (%)"}]]],"referenceName":"gi\_157834972\_ref\_NC","date":"18 July 2018, Wednesday, 16:27:05","order":[0,1,2,3,4,5,6,7,8,9,10,11,12],"assembliesNames":["ABySS\_127","ABySS\_63","CLC","IDBA\_UD","MEGAHIT","MIRA","SOAPdenovo2","SPAdes","SPAdes\_meta","SPAdes\_sc","SPAdes\_sc\_careful","Velvet","Geneious"]},{"assembliesWithNs":null,"minContig":500,"report":[["Genome statistics",[{"values":["7.647","3.881","50.367","43.622","55.045","15.356","39.596","61.594","61.929","60.312","60.270","0.861"],"quality":"More is better","isMain":true,"metricName":"Genome fraction (%)"},{"values":["1.000","1.000","1.022","1.016","1.004","1.001","1.005","1.009","1.007","1.015","1.015","1.010"],"quality":"Less is better","isMain":true,"metricName":"Duplication ratio"},{"values":[1056,790,3674,2289,3511,1651,2266,3675,3674,3674,3674,648],"quality":"More is better","isMain":true,"metricName":"Largest alignment"},{"values":[11055,5610,73654,63396,79669,22200,57395,89192,90084,87332,87273,1245],"quality":"More is better","isMain":true,"metricName":"Total aligned length"},{"values":[null,null,603,null,628,null,null,906,831,876,876,null],"quality":"More is better","isMain":false,"metricName":"NG50"},{"values":[728,607,1018,1011,1058,996,947,1487,1318,1409,1409,648],"quality":"More is better","isMain":false,"metricName":"NA50"},{"values":[613,547,729,741,759,797,705,1004,912,969,969,597],"quality":"More is better","isMain":false,"metricName":"NA75"},{"values":[null,null,533,null,617,null,null,869,831,851,851,null],"quality":"More is better","isMain":true,"metricName":"NGA50"},{"values":[null,null,59,null,63,null,null,43,50,45,45,null],"quality":"Less is better","isMain":false,"metricName":"LG50"},{"values":[7,4,24,24,25,9,21,21,25,21,21,1],"quality":"Less is better","isMain":false,"metricName":"LA50"},{"values":[11,7,47,43,47,15,39,39,45,40,40,2],"quality":"Less is better","isMain":false,"metricName":"LA75"},{"values":[null,null,68,null,64,null,null,44,50,46,46,null],"quality":"Less is better","isMain":true,"metricName":"LGA50"}]],["Misassemblies",[{"values":[0,0,0,0,0,0,0,0,0,0,0,0],"quality":"Less is better","isMain":true,"metricName":"# misassemblies"},{"values":[0,0,0,0,0,0,0,0,0,0,0,0],"quality":"Less is better","isMain":false,"metricName":" # relocations"},{"values":[0,0,0,0,0,0,0,0,0,0,0,0],"quality":"Less is better","isMain":false,"metricName":" # translocations"},{"values":[0,0,0,0,0,0,0,0,0,0,0,0],"quality":"Less is better","isMain":false,"metricName":" # inversions"},{"values":[0,0,0,0,0,0,0,0,0,0,0,0],"quality":"Less is better","isMain":false,"metricName":"# misassembled contigs"},{"values":[0,0,0,0,0,0,0,0,0,0,0,0],"quality":"Less is better","isMain":true,"metricName":"Misassembled contigs length"},{"values":[0,0,0,0,0,0,0,0,0,0,0,0],"quality":"Less is better","isMain":false,"metricName":"# local misassemblies"},{"values":[0,0,0,0,0,0,0,0,0,0,0,0],"quality":"Less is better","isMain":false,"metricName":"# unaligned mis. contigs"}]],["Unaligned",[{"values":[0,0,0,0,0,0,0,0,0,0,0,0],"quality":"Less is better","isMain":false,"metricName":"# fully unaligned contigs"},{"values":[0,0,0,0,0,0,0,0,0,0,0,0],"quality":"Less is better","isMain":false,"metricName":"Fully unaligned length"},{"values":[0,0,2,2,1,0,0,1,0,1,1,0],"quality":"Less is better","isMain":false,"metricName":"# partially unaligned contigs"},{"values":[0,0,4900,2670,595,0,0,559,0,559,559,0],"quality":"Less is better","isMain":false,"metricName":"Partially unaligned length"}]],["Mismatches",[{"values":[1,0,443,315,183,12,57,691,191,705,659,2],"quality":"Less is better","isMain":false,"metricName":"# mismatches"},{"values":[0,0,7,5,3,0,3,29,0,31,33,0],"quality":"Less is better","isMain":false,"metricName":"# indels"},{"values":[0,0,10,10,8,0,144,62,0,69,71,0],"quality":"Less is better","isMain":false,"metricName":"Indels length"},{"values":["9.05","0.00","608.41","499.51","229.97","54.05","99.58","776.03","213.34","808.58","756.35","160.64"],"quality":"Less is better","isMain":true,"metricName":"# mismatches per 100 kbp"},{"values":["0.00","0.00","9.61","7.93","3.77","0.00","5.24","32.57","0.00","35.55","37.87","0.00"],"quality":"Less is better","isMain":true,"metricName":"# indels per 100 kbp"},{"values":[0,0,7,5,2,0,0,27,0,29,31,0],"quality":"Less is better","isMain":false,"metricName":" # indels (<= 5 bp)"},{"values":[0,0,0,0,1,0,3,2,0,2,2,0],"quality":"Less is better","isMain":false,"metricName":" # indels (> 5 bp)"},{"values":[0,0,0,0,0,36,202,0,0,0,0,0],"quality":"Less is better","isMain":false,"metricName":"# N's"},{"values":["0.00","0.00","0.00","0.00","0.00","162.00","351.29","0.00","0.00","0.00","0.00","0.00"],"quality":"Less is better","isMain":true,"metricName":"# N's per 100 kbp"}]],["Statistics without reference",[{"values":[16,9,71,65,77,23,63,69,77,68,68,2],"quality":"Equal","isMain":true,"metricName":"# contigs"},{"values":[1,0,27,28,29,8,19,41,41,41,41,0],"quality":"Equal","isMain":false,"metricName":"# contigs (>= 1000 bp)"},{"values":[0,0,0,0,0,0,0,0,0,0,0,0],"quality":"Equal","isMain":false,"metricName":"# contigs (>= 5000 bp)"},{"values":[0,0,0,0,0,0,0,0,0,0,0,0],"quality":"Equal","isMain":false,"metricName":"# contigs (>= 10000 bp)"},{"values":[0,0,0,0,0,0,0,0,0,0,0,0],"quality":"Equal","isMain":false,"metricName":"# contigs (>= 25000 bp)"},{"values":[0,0,0,0,0,0,0,0,0,0,0,0],"quality":"Equal","isMain":false,"metricName":"# contigs (>= 50000 bp)"},{"values":[1056,790,4464,2289,3528,1651,2266,3697,3696,3696,3696,660],"quality":"More is better","isMain":true,"metricName":"Largest contig"},{"values":[11055,5610,79311,66747,80527,22222,57502,90425,90111,89038,88969,1257],"quality":"More is better","isMain":true,"metricName":"Total length"},{"values":[1056,0,47894,40025,45662,10559,26980,70736,64323,69418,69418,0],"quality":"More is better","isMain":true,"metricName":"Total length (>= 1000 bp)"},{"values":[0,0,0,0,0,0,0,0,0,0,0,0],"quality":"More is better","isMain":false,"metricName":"Total length (>= 5000 bp)"},{"values":[0,0,0,0,0,0,0,0,0,0,0,0],"quality":"More is better","isMain":true,"metricName":"Total length (>= 10000 bp)"},{"values":[0,0,0,0,0,0,0,0,0,0,0,0],"quality":"More is better","isMain":false,"metricName":"Total length (>= 25000 bp)"},{"values":[0,0,0,0,0,0,0,0,0,0,0,0],"quality":"More is better","isMain":true,"metricName":"Total length (>= 50000 bp)"},{"values":[728,607,1216,1116,1062,996,947,1540,1318,1516,1516,660],"quality":"More is better","isMain":false,"metricName":"N50"},{"values":[613,547,778,866,761,797,705,1012,912,1011,1011,597],"quality":"More is better","isMain":false,"metricName":"N75"},{"values":[7,4,20,22,24,9,21,21,25,21,21,1],"quality":"Less is better","isMain":false,"metricName":"L50"},{"values":[11,7,41,39,46,15,39,39,45,39,39,2],"quality":"Less is better","isMain":false,"metricName":"L75"},{"values":["28.39","29.25","27.63","27.88","27.11","27.84","26.83","27.37","27.13","27.40","27.40","27.37"],"quality":"Equal","isMain":false,"metricName":"GC (%)"}]],["Predicted genes",[]],["Similarity statistics",[{"values":[0,0,0,0,0,0,0,0,0,0,0,0],"quality":"Equal","isMain":false,"metricName":"# similar correct contigs"},{"values":[0,0,0,0,0,0,0,0,0,0,0,0],"quality":"Equal","isMain":false,"metricName":"# similar misassembled blocks"}]],["Reference statistics",[{"values":[144565,144565,144565,144565,144565,144565,144565,144565,144565,144565,144565,144565],"quality":"Equal","isMain":false,"metricName":"Reference length"},{"values":[1,1,1,1,1,1,1,1,1,1,1,1],"quality":"Equal","isMain":false,"metricName":"Reference fragments"},{"values":["26.99","26.99","26.99","26.99","26.99","26.99","26.99","26.99","26.99","26.99","26.99","26.99"],"quality":"Equal","isMain":false,"metricName":"Reference GC (%)"}]]],"referenceName":"gi\_157939619\_ref\_NC","date":"18 July 2018, Wednesday, 16:27:17","order":[0,1,2,3,4,5,6,7,8,9,10,11],"assembliesNames":["ABySS\_127","ABySS\_63","CLC","IDBA\_UD","MEGAHIT","MIRA","SOAPdenovo2","SPAdes","SPAdes\_meta","SPAdes\_sc","SPAdes\_sc\_careful","Velvet"]},{"assembliesWithNs":null,"minContig":500,"report":[["Genome statistics",[{"values":["57.645","37.485","62.239","56.095","68.256","77.540","0.284","22.830","66.454","57.718","58.685","54.529","38.169","64.920"],"quality":"More is better","isMain":true,"metricName":"Genome fraction (%)"},{"values":["1.033","1.008","1.019","1.017","1.037","1.194","1.000","1.135","1.044","1.012","1.016","1.013","1.003","1.092"],"quality":"Less is better","isMain":true,"metricName":"Duplication ratio"},{"values":[4806,6504,7359,6826,5712,26284,528,3635,16812,11282,7944,6977,5036,27055],"quality":"More is better","isMain":true,"metricName":"Largest alignment"},{"values":[218903,139315,233199,210352,260718,340505,1046,84549,254447,215240,219359,203148,141172,259825],"quality":"More is better","isMain":true,"metricName":"Total aligned length"},{"values":[745,null,1028,764,1316,26899,null,null,6546,1082,971,736,null,40395],"quality":"More is better","isMain":false,"metricName":"NG50"},{"values":[null,null,null,null,null,8228,null,null,574,null,null,null,null,28463],"quality":"More is better","isMain":false,"metricName":"NG75"},{"values":[1412,1263,1843,1756,2016,1735,528,613,5723,2164,2136,1828,1239,1513],"quality":"More is better","isMain":false,"metricName":"NA50"},{"values":[865,815,1111,1091,1194,null,518,null,2789,1265,1133,995,888,null],"quality":"More is better","isMain":false,"metricName":"NA75"},{"values":[731,null,1027,764,1290,3255,null,null,3767,1021,911,677,null,4853],"quality":"More is better","isMain":true,"metricName":"NGA50"},{"values":[null,null,null,null,null,1355,null,null,null,null,null,null,null,null],"quality":"More is better","isMain":false,"metricName":"NGA75"},{"values":[121,null,89,103,74,4,null,null,18,73,77,99,null,2],"quality":"Less is better","isMain":false,"metricName":"LG50"},{"values":[null,null,null,null,null,11,null,null,57,null,null,null,null,5],"quality":"Less is better","isMain":false,"metricName":"LG75"},{"values":[48,36,41,36,41,55,1,55,17,29,30,32,36,36],"quality":"Less is better","isMain":false,"metricName":"LA50"},{"values":[97,70,81,74,84,null,2,null,34,63,66,70,70,null],"quality":"Less is better","isMain":false,"metricName":"LA75"},{"values":[122,null,89,103,75,26,null,null,26,79,83,105,null,16],"quality":"Less is better","isMain":true,"metricName":"LGA50"},{"values":[null,null,null,null,null,71,null,null,null,null,null,null,null,null],"quality":"Less is better","isMain":false,"metricName":"LGA75"}]],["Misassemblies",[{"values":[1,0,0,0,1,8,0,0,6,0,1,0,0,9],"quality":"Less is better","isMain":true,"metricName":"# misassemblies"},{"values":[1,0,0,0,1,8,0,0,6,0,1,0,0,9],"quality":"Less is better","isMain":false,"metricName":" # relocations"},{"values":[0,0,0,0,0,0,0,0,0,0,0,0,0,0],"quality":"Less is better","isMain":false,"metricName":" # translocations"},{"values":[0,0,0,0,0,0,0,0,0,0,0,0,0,0],"quality":"Less is better","isMain":false,"metricName":" # inversions"},{"values":[1,0,0,0,1,6,0,0,4,0,1,0,0,6],"quality":"Less is better","isMain":false,"metricName":"# misassembled contigs"},{"values":[2067,0,0,0,1207,53596,0,0,56189,0,5247,0,0,116175],"quality":"Less is better","isMain":true,"metricName":"Misassembled contigs length"},{"values":[0,0,0,2,1,8,0,2,6,1,1,1,0,12],"quality":"Less is better","isMain":false,"metricName":"# local misassemblies"},{"values":[0,0,0,0,0,2,0,12,0,0,0,0,0,3],"quality":"Less is better","isMain":false,"metricName":"# unaligned mis. contigs"}]],["Unaligned",[{"values":[0,0,0,0,0,0,0,0,0,0,0,0,0,0],"quality":"Less is better","isMain":false,"metricName":"# fully unaligned contigs"},{"values":[0,0,0,0,0,0,0,0,0,0,0,0,0,0],"quality":"Less is better","isMain":false,"metricName":"Fully unaligned length"},{"values":[0,0,0,0,0,11,0,29,8,1,3,3,0,14],"quality":"Less is better","isMain":false,"metricName":"# partially unaligned contigs"},{"values":[0,0,0,0,0,162093,0,29539,22185,6062,3870,3782,0,242913],"quality":"Less is better","isMain":false,"metricName":"Partially unaligned length"}]],["Mismatches",[{"values":[473,246,274,352,463,1573,3,42,2445,880,1015,575,246,2034],"quality":"Less is better","isMain":false,"metricName":"# mismatches"},{"values":[25,6,10,19,31,116,0,57,177,37,47,31,5,213],"quality":"Less is better","isMain":false,"metricName":"# indels"},{"values":[60,18,30,86,185,275,0,2806,501,124,225,160,27,566],"quality":"Less is better","isMain":false,"metricName":"Indels length"},{"values":["222.56","178.00","119.41","170.20","183.99","550.24","286.81","49.90","997.93","413.54","469.12","286.01","174.81","849.81"],"quality":"Less is better","isMain":true,"metricName":"# mismatches per 100 kbp"},{"values":["11.76","4.34","4.36","9.19","12.32","40.58","0.00","67.72","72.24","17.39","21.72","15.42","3.55","88.99"],"quality":"Less is better","isMain":true,"metricName":"# indels per 100 kbp"},{"values":[21,5,9,15,21,105,0,0,155,35,38,22,4,194],"quality":"Less is better","isMain":false,"metricName":" # indels (<= 5 bp)"},{"values":[4,1,1,4,10,11,0,57,22,2,9,9,1,19],"quality":"Less is better","isMain":false,"metricName":" # indels (> 5 bp)"},{"values":[0,0,0,0,0,215,0,19905,0,0,2357,2755,0,2685],"quality":"Less is better","isMain":false,"metricName":"# N's"},{"values":["0.00","0.00","0.00","0.00","0.00","42.70","0.00","15911.65","0.00","0.00","1053.43","1328.14","0.00","532.54"],"quality":"Less is better","isMain":true,"metricName":"# N's per 100 kbp"}]],["Statistics without reference",[{"values":[178,125,155,145,169,129,2,113,59,120,133,137,122,29],"quality":"Equal","isMain":true,"metricName":"# contigs"},{"values":[81,55,91,81,93,96,0,51,51,80,73,68,56,29],"quality":"Equal","isMain":false,"metricName":"# contigs (>= 1000 bp)"},{"values":[0,2,1,3,3,16,0,1,21,7,6,3,1,19],"quality":"Equal","isMain":false,"metricName":"# contigs (>= 5000 bp)"},{"values":[0,0,0,0,0,7,0,0,7,1,0,0,0,12],"quality":"Equal","isMain":false,"metricName":"# contigs (>= 10000 bp)"},{"values":[0,0,0,0,0,4,0,0,0,0,0,0,0,5],"quality":"Equal","isMain":false,"metricName":"# contigs (>= 25000 bp)"},{"values":[0,0,0,0,0,1,0,0,0,0,0,0,0,1],"quality":"Equal","isMain":false,"metricName":"# contigs (>= 50000 bp)"},{"values":[4806,6504,7359,6826,5712,115158,528,5074,20214,11282,7944,7445,5036,171838],"quality":"More is better","isMain":true,"metricName":"Largest contig"},{"values":[219440,139315,233779,210366,260875,503518,1046,125097,277942,221481,223746,207433,141175,504185],"quality":"More is better","isMain":true,"metricName":"Total length"},{"values":[150983,91815,187255,165690,206413,477774,0,80746,272664,191792,180514,158015,93021,504185],"quality":"More is better","isMain":true,"metricName":"Total length (>= 1000 bp)"},{"values":[0,11558,7359,17733,16416,314534,0,5074,204136,47542,38990,19697,5036,470987],"quality":"More is better","isMain":false,"metricName":"Total length (>= 5000 bp)"},{"values":[0,0,0,0,0,246361,0,0,100841,11282,0,0,0,419804],"quality":"More is better","isMain":true,"metricName":"Total length (>= 10000 bp)"},{"values":[0,0,0,0,0,200272,0,0,0,0,0,0,0,301990],"quality":"More is better","isMain":false,"metricName":"Total length (>= 25000 bp)"},{"values":[0,0,0,0,0,115158,0,0,0,0,0,0,0,171838],"quality":"More is better","isMain":true,"metricName":"Total length (>= 50000 bp)"},{"values":[1427,1263,1843,1756,2016,9858,528,1179,7714,2263,2298,1931,1239,29758],"quality":"More is better","isMain":false,"metricName":"N50"},{"values":[872,815,1137,1091,1198,2756,518,841,3807,1382,1213,1041,888,17218],"quality":"More is better","isMain":false,"metricName":"N75"},{"values":[48,36,41,36,41,8,1,35,12,27,27,30,36,4],"quality":"Less is better","isMain":false,"metricName":"L50"},{"values":[96,70,80,74,84,33,2,66,23,59,62,66,70,9],"quality":"Less is better","isMain":false,"metricName":"L75"},{"values":["40.48","40.18","40.60","40.15","40.48","40.59","44.65","39.54","40.79","40.40","40.16","40.04","39.81","40.53"],"quality":"Equal","isMain":false,"metricName":"GC (%)"}]],["Predicted genes",[]],["Similarity statistics",[{"values":[0,0,0,0,0,0,0,0,0,0,0,0,0,0],"quality":"Equal","isMain":false,"metricName":"# similar correct contigs"},{"values":[0,0,0,0,0,0,0,0,0,0,0,0,0,0],"quality":"Equal","isMain":false,"metricName":"# similar misassembled blocks"}]],["Reference statistics",[{"values":[368683,368683,368683,368683,368683,368683,368683,368683,368683,368683,368683,368683,368683,368683],"quality":"Equal","isMain":false,"metricName":"Reference length"},{"values":[1,1,1,1,1,1,1,1,1,1,1,1,1,1],"quality":"Equal","isMain":false,"metricName":"Reference fragments"},{"values":["40.69","40.69","40.69","40.69","40.69","40.69","40.69","40.69","40.69","40.69","40.69","40.69","40.69","40.69"],"quality":"Equal","isMain":false,"metricName":"Reference GC (%)"}]]],"referenceName":"gi\_157952304\_ref\_NC","date":"18 July 2018, Wednesday, 16:27:33","order":[0,1,2,3,4,5,6,7,8,9,10,11,12,13],"assembliesNames":["ABySS\_127","ABySS\_63","CLC","IDBA\_UD","MEGAHIT","MIRA","Ray\_Meta","SOAPdenovo2","SPAdes","SPAdes\_meta","SPAdes\_sc","SPAdes\_sc\_careful","Velvet","Geneious"]},{"assembliesWithNs":null,"minContig":500,"report":[["Genome statistics",[{"values":["68.766","55.789","72.911","66.591","71.533","89.267","0.341","19.055","70.016","58.381","53.253","52.665","37.828","82.949"],"quality":"More is better","isMain":true,"metricName":"Genome fraction (%)"},{"values":["1.031","1.010","1.020","1.021","1.034","1.186","1.000","1.283","1.021","1.015","1.016","1.017","1.005","1.126"],"quality":"Less is better","isMain":true,"metricName":"Duplication ratio"},{"values":[6296,5492,8053,7550,6852,29862,639,2840,22416,8958,7026,7540,4693,31472],"quality":"More is better","isMain":true,"metricName":"Largest alignment"},{"values":[243808,194227,255803,234254,254796,363837,1175,65984,246315,204200,186272,183633,130899,321044],"quality":"More is better","isMain":true,"metricName":"Total aligned length"},{"values":[1156,719,1657,1224,2565,73723,null,null,7619,1223,647,633,null,73724],"quality":"More is better","isMain":false,"metricName":"NG50"},{"values":[null,null,null,null,1237,26899,null,null,1247,null,null,null,null,40395],"quality":"More is better","isMain":false,"metricName":"NG75"},{"values":[1815,1593,2138,2044,1853,3309,639,174,6199,2014,1876,1858,1165,2732],"quality":"More is better","isMain":false,"metricName":"NA50"},{"values":[1060,926,1336,1192,712,null,536,null,2828,1348,1030,1011,796,null],"quality":"More is better","isMain":false,"metricName":"NA75"},{"values":[1138,719,1657,1213,1699,7226,null,null,4255,1144,606,600,null,12109],"quality":"More is better","isMain":true,"metricName":"NGA50"},{"values":[null,null,null,null,null,4577,null,null,null,null,null,null,null,5861],"quality":"More is better","isMain":false,"metricName":"NGA75"},{"values":[82,112,62,71,30,2,null,null,13,69,94,99,null,2],"quality":"Less is better","isMain":false,"metricName":"LG50"},{"values":[null,null,null,null,77,5,null,null,43,null,null,null,null,3],"quality":"Less is better","isMain":false,"metricName":"LG75"},{"values":[47,38,39,37,54,35,1,93,13,28,30,30,32,28],"quality":"Less is better","isMain":false,"metricName":"LA50"},{"values":[92,79,75,75,125,null,2,null,29,59,63,64,66,null],"quality":"Less is better","isMain":false,"metricName":"LA75"},{"values":[82,112,62,72,61,14,null,null,21,72,101,105,null,9],"quality":"Less is better","isMain":true,"metricName":"LGA50"},{"values":[null,null,null,null,null,28,null,null,null,null,null,null,null,20],"quality":"Less is better","isMain":false,"metricName":"LGA75"}]],["Misassemblies",[{"values":[0,0,1,0,0,11,0,0,6,0,0,0,1,13],"quality":"Less is better","isMain":true,"metricName":"# misassemblies"},{"values":[0,0,1,0,0,11,0,0,6,0,0,0,1,13],"quality":"Less is better","isMain":false,"metricName":" # relocations"},{"values":[0,0,0,0,0,0,0,0,0,0,0,0,0,0],"quality":"Less is better","isMain":false,"metricName":" # translocations"},{"values":[0,0,0,0,0,0,0,0,0,0,0,0,0,0],"quality":"Less is better","isMain":false,"metricName":" # inversions"},{"values":[0,0,1,0,0,9,0,0,5,0,0,0,1,10],"quality":"Less is better","isMain":false,"metricName":"# misassembled contigs"},{"values":[0,0,905,0,0,106759,0,0,74779,0,0,0,1153,118007],"quality":"Less is better","isMain":true,"metricName":"Misassembled contigs length"},{"values":[0,0,2,1,1,15,0,3,4,4,0,0,0,16],"quality":"Less is better","isMain":false,"metricName":"# local misassemblies"},{"values":[0,0,0,0,1,2,0,20,1,0,0,0,0,2],"quality":"Less is better","isMain":false,"metricName":"# unaligned mis. contigs"}]],["Unaligned",[{"values":[0,0,0,0,0,0,0,0,0,0,0,0,0,0],"quality":"Less is better","isMain":false,"metricName":"# fully unaligned contigs"},{"values":[0,0,0,0,0,0,0,0,0,0,0,0,0,0],"quality":"Less is better","isMain":false,"metricName":"Fully unaligned length"},{"values":[0,0,0,1,1,10,0,24,7,2,3,1,0,15],"quality":"Less is better","isMain":false,"metricName":"# partially unaligned contigs"},{"values":[0,0,0,935,63365,206388,0,38360,19784,3313,4670,2425,0,262226],"quality":"Less is better","isMain":false,"metricName":"Partially unaligned length"}]],["Mismatches",[{"values":[233,446,295,471,422,2259,1,40,1469,653,438,386,293,1972],"quality":"Less is better","isMain":false,"metricName":"# mismatches"},{"values":[10,13,7,15,27,91,0,72,69,29,9,14,13,135],"quality":"Less is better","isMain":false,"metricName":"# indels"},{"values":[73,42,32,60,60,225,0,3529,348,129,109,115,170,383],"quality":"Less is better","isMain":false,"metricName":"Indels length"},{"values":["98.30","231.93","117.38","205.20","171.15","734.17","85.11","60.90","608.68","324.50","238.62","212.63","224.71","689.71"],"quality":"Less is better","isMain":true,"metricName":"# mismatches per 100 kbp"},{"values":["4.22","6.76","2.79","6.54","10.95","29.57","0.00","109.62","28.59","14.41","4.90","7.71","9.97","47.22"],"quality":"Less is better","isMain":true,"metricName":"# indels per 100 kbp"},{"values":[7,11,4,12,24,83,0,0,51,22,6,11,10,123],"quality":"Less is better","isMain":false,"metricName":" # indels (<= 5 bp)"},{"values":[3,2,3,3,3,8,0,72,18,7,3,3,3,12],"quality":"Less is better","isMain":false,"metricName":" # indels (> 5 bp)"},{"values":[0,0,0,0,0,287,0,26848,0,507,1185,1499,30,2998],"quality":"Less is better","isMain":false,"metricName":"# N's"},{"values":["0.00","0.00","0.00","0.00","0.00","50.23","0.00","21896.18","0.00","244.32","619.92","801.77","22.90","513.15"],"quality":"Less is better","isMain":true,"metricName":"# N's per 100 kbp"}]],["Statistics without reference",[{"values":[170,148,149,152,151,80,2,97,52,112,127,125,118,27],"quality":"Equal","isMain":true,"metricName":"# contigs"},{"values":[98,72,97,85,89,61,0,45,44,78,63,66,46,27],"quality":"Equal","isMain":false,"metricName":"# contigs (>= 1000 bp)"},{"values":[1,1,5,4,4,27,0,1,18,6,5,5,0,20],"quality":"Equal","isMain":false,"metricName":"# contigs (>= 5000 bp)"},{"values":[0,0,0,0,1,13,0,0,9,0,0,0,0,15],"quality":"Equal","isMain":false,"metricName":"# contigs (>= 10000 bp)"},{"values":[0,0,0,0,1,5,0,0,0,0,0,0,0,6],"quality":"Equal","isMain":false,"metricName":"# contigs (>= 25000 bp)"},{"values":[0,0,0,0,1,2,0,0,0,0,0,0,0,2],"quality":"Equal","isMain":false,"metricName":"# contigs (>= 50000 bp)"},{"values":[6296,5492,8053,7550,65564,115158,639,6886,22416,8958,7026,7944,4693,171838],"quality":"More is better","isMain":true,"metricName":"Largest contig"},{"values":[244278,194237,256317,235225,318346,571410,1175,122615,266129,207513,191153,186961,130994,584240],"quality":"More is better","isMain":true,"metricName":"Total length"},{"values":[190687,139195,219038,188693,272345,556983,0,85923,260848,182934,147612,146002,81624,584240],"quality":"More is better","isMain":true,"metricName":"Total length (>= 1000 bp)"},{"values":[6296,5492,33011,25752,84757,482286,0,6886,201539,44028,30284,31688,0,563141],"quality":"More is better","isMain":false,"metricName":"Total length (>= 5000 bp)"},{"values":[0,0,0,0,65564,386873,0,0,139435,0,0,0,0,529743],"quality":"More is better","isMain":true,"metricName":"Total length (>= 10000 bp)"},{"values":[0,0,0,0,65564,274104,0,0,0,0,0,0,0,374311],"quality":"More is better","isMain":false,"metricName":"Total length (>= 25000 bp)"},{"values":[0,0,0,0,65564,188881,0,0,0,0,0,0,0,245562],"quality":"More is better","isMain":true,"metricName":"Total length (>= 50000 bp)"},{"values":[1815,1593,2138,2059,2806,21227,639,1339,10930,2149,2014,1876,1165,31536],"quality":"More is better","isMain":false,"metricName":"N50"},{"values":[1079,926,1336,1199,1637,6865,536,880,5507,1433,1106,1030,814,20059],"quality":"More is better","isMain":false,"metricName":"N75"},{"values":[47,38,39,36,25,6,1,25,9,26,28,28,32,4],"quality":"Less is better","isMain":false,"metricName":"L50"},{"values":[91,79,75,74,63,19,2,52,18,56,60,61,65,9],"quality":"Less is better","isMain":false,"metricName":"L75"},{"values":["40.67","40.23","40.37","40.58","40.29","40.66","44.17","39.67","40.53","40.48","40.11","40.14","39.86","40.56"],"quality":"Equal","isMain":false,"metricName":"GC (%)"}]],["Predicted genes",[]],["Similarity statistics",[{"values":[0,0,0,0,0,0,0,0,0,0,0,0,0,0],"quality":"Equal","isMain":false,"metricName":"# similar correct contigs"},{"values":[0,0,0,0,0,0,0,0,0,0,0,0,0,0],"quality":"Equal","isMain":false,"metricName":"# similar misassembled blocks"}]],["Reference statistics",[{"values":[344691,344691,344691,344691,344691,344691,344691,344691,344691,344691,344691,344691,344691,344691],"quality":"Equal","isMain":false,"metricName":"Reference length"},{"values":[1,1,1,1,1,1,1,1,1,1,1,1,1,1],"quality":"Equal","isMain":false,"metricName":"Reference fragments"},{"values":["40.76","40.76","40.76","40.76","40.76","40.76","40.76","40.76","40.76","40.76","40.76","40.76","40.76","40.76"],"quality":"Equal","isMain":false,"metricName":"Reference GC (%)"}]]],"referenceName":"gi\_157953191\_ref\_NC","date":"18 July 2018, Wednesday, 16:27:49","order":[0,1,2,3,4,5,6,7,8,9,10,11,12,13],"assembliesNames":["ABySS\_127","ABySS\_63","CLC","IDBA\_UD","MEGAHIT","MIRA","Ray\_Meta","SOAPdenovo2","SPAdes","SPAdes\_meta","SPAdes\_sc","SPAdes\_sc\_careful","Velvet","Geneious"]},{"assembliesWithNs":null,"minContig":500,"report":[["Genome statistics",[{"values":["100.000","99.985","100.000","100.000","100.000","83.282","76.362","100.000","100.000","100.000","100.000","98.266"],"quality":"More is better","isMain":true,"metricName":"Genome fraction (%)"},{"values":["1.000","1.000","1.001","1.001","1.002","1.022","1.034","1.001","1.000","1.000","1.000","1.001"],"quality":"Less is better","isMain":true,"metricName":"Duplication ratio"},{"values":[142122,142050,142171,142213,142365,4059,8842,142199,142108,142108,142108,83236],"quality":"More is better","isMain":true,"metricName":"Largest alignment"},{"values":[142122,142050,142171,142213,142365,120965,109672,142199,142108,142108,142108,139753],"quality":"More is better","isMain":true,"metricName":"Total aligned length"},{"values":[142122,142072,142171,142213,142365,1183,5659,142199,142108,142108,142108,83236],"quality":"More is better","isMain":false,"metricName":"NG50"},{"values":[142122,142072,142171,142213,142365,694,1906,142199,142108,142108,142108,36403],"quality":"More is better","isMain":false,"metricName":"NG75"},{"values":[142122,142050,142171,142213,142365,1433,3596,142199,142108,142108,142108,83236],"quality":"More is better","isMain":false,"metricName":"NA50"},{"values":[142122,142050,142171,142213,142365,944,1019,142199,142108,142108,142108,36403],"quality":"More is better","isMain":false,"metricName":"NA75"},{"values":[142122,142050,142171,142213,142365,1183,3373,142199,142108,142108,142108,83236],"quality":"More is better","isMain":true,"metricName":"NGA50"},{"values":[142122,142050,142171,142213,142365,694,656,142199,142108,142108,142108,36403],"quality":"More is better","isMain":false,"metricName":"NGA75"},{"values":[1,1,1,1,1,34,9,1,1,1,1,1],"quality":"Less is better","isMain":false,"metricName":"LG50"},{"values":[1,1,1,1,1,73,19,1,1,1,1,2],"quality":"Less is better","isMain":false,"metricName":"LG75"},{"values":[1,1,1,1,1,26,11,1,1,1,1,1],"quality":"Less is better","isMain":false,"metricName":"LA50"},{"values":[1,1,1,1,1,53,25,1,1,1,1,2],"quality":"Less is better","isMain":false,"metricName":"LA75"},{"values":[1,1,1,1,1,34,13,1,1,1,1,1],"quality":"Less is better","isMain":true,"metricName":"LGA50"},{"values":[1,1,1,1,1,73,37,1,1,1,1,2],"quality":"Less is better","isMain":false,"metricName":"LGA75"}]],["Misassemblies",[{"values":[0,0,0,0,0,0,1,0,0,0,0,0],"quality":"Less is better","isMain":true,"metricName":"# misassemblies"},{"values":[0,0,0,0,0,0,1,0,0,0,0,0],"quality":"Less is better","isMain":false,"metricName":" # relocations"},{"values":[0,0,0,0,0,0,0,0,0,0,0,0],"quality":"Less is better","isMain":false,"metricName":" # translocations"},{"values":[0,0,0,0,0,0,0,0,0,0,0,0],"quality":"Less is better","isMain":false,"metricName":" # inversions"},{"values":[0,0,0,0,0,0,1,0,0,0,0,0],"quality":"Less is better","isMain":false,"metricName":"# misassembled contigs"},{"values":[0,0,0,0,0,0,9609,0,0,0,0,0],"quality":"Less is better","isMain":true,"metricName":"Misassembled contigs length"},{"values":[0,0,1,1,1,0,4,1,0,0,0,0],"quality":"Less is better","isMain":false,"metricName":"# local misassemblies"},{"values":[0,0,0,0,0,0,0,0,0,0,0,0],"quality":"Less is better","isMain":false,"metricName":"# unaligned mis. contigs"}]],["Unaligned",[{"values":[0,0,0,0,0,0,0,0,0,0,0,0],"quality":"Less is better","isMain":false,"metricName":"# fully unaligned contigs"},{"values":[0,0,0,0,0,0,0,0,0,0,0,0],"quality":"Less is better","isMain":false,"metricName":"Fully unaligned length"},{"values":[0,0,0,0,0,0,7,0,0,0,0,0],"quality":"Less is better","isMain":false,"metricName":"# partially unaligned contigs"},{"values":[0,0,0,0,0,0,16683,0,0,0,0,0],"quality":"Less is better","isMain":false,"metricName":"Partially unaligned length"}]],["Mismatches",[{"values":[0,1,0,0,0,0,13,0,0,0,0,0],"quality":"Less is better","isMain":false,"metricName":"# mismatches"},{"values":[1,0,0,0,0,0,61,0,1,1,1,0],"quality":"Less is better","isMain":false,"metricName":"# indels"},{"values":[50,0,0,0,0,0,2855,0,36,36,36,0],"quality":"Less is better","isMain":false,"metricName":"Indels length"},{"values":["0.00","0.70","0.00","0.00","0.00","0.00","11.98","0.00","0.00","0.00","0.00","0.00"],"quality":"Less is better","isMain":true,"metricName":"# mismatches per 100 kbp"},{"values":["0.70","0.00","0.00","0.00","0.00","0.00","56.23","0.00","0.70","0.70","0.70","0.00"],"quality":"Less is better","isMain":true,"metricName":"# indels per 100 kbp"},{"values":[0,0,0,0,0,0,0,0,0,0,0,0],"quality":"Less is better","isMain":false,"metricName":" # indels (<= 5 bp)"},{"values":[1,0,0,0,0,0,61,0,1,1,1,0],"quality":"Less is better","isMain":false,"metricName":" # indels (> 5 bp)"},{"values":[0,0,0,0,0,0,8640,0,0,0,0,0],"quality":"Less is better","isMain":false,"metricName":"# N's"},{"values":["0.00","0.00","0.00","0.00","0.00","0.00","6707.50","0.00","0.00","0.00","0.00","0.00"],"quality":"Less is better","isMain":true,"metricName":"# N's per 100 kbp"}]],["Statistics without reference",[{"values":[1,1,1,1,1,96,42,1,1,1,1,13],"quality":"Equal","isMain":true,"metricName":"# contigs"},{"values":[1,1,1,1,1,45,25,1,1,1,1,7],"quality":"Equal","isMain":false,"metricName":"# contigs (>= 1000 bp)"},{"values":[1,1,1,1,1,0,10,1,1,1,1,3],"quality":"Equal","isMain":false,"metricName":"# contigs (>= 5000 bp)"},{"values":[1,1,1,1,1,0,2,1,1,1,1,2],"quality":"Equal","isMain":false,"metricName":"# contigs (>= 10000 bp)"},{"values":[1,1,1,1,1,0,0,1,1,1,1,2],"quality":"Equal","isMain":false,"metricName":"# contigs (>= 25000 bp)"},{"values":[1,1,1,1,1,0,0,1,1,1,1,1],"quality":"Equal","isMain":false,"metricName":"# contigs (>= 50000 bp)"},{"values":[142122,142072,142171,142213,142365,4059,12203,142199,142108,142108,142108,83236],"quality":"More is better","isMain":true,"metricName":"Largest contig"},{"values":[142122,142072,142171,142213,142365,120965,128811,142199,142108,142108,142108,139753],"quality":"More is better","isMain":true,"metricName":"Total length"},{"values":[142122,142072,142171,142213,142365,83541,116492,142199,142108,142108,142108,135309],"quality":"More is better","isMain":true,"metricName":"Total length (>= 1000 bp)"},{"values":[142122,142072,142171,142213,142365,0,80436,142199,142108,142108,142108,128948],"quality":"More is better","isMain":false,"metricName":"Total length (>= 5000 bp)"},{"values":[142122,142072,142171,142213,142365,0,23862,142199,142108,142108,142108,119639],"quality":"More is better","isMain":true,"metricName":"Total length (>= 10000 bp)"},{"values":[142122,142072,142171,142213,142365,0,0,142199,142108,142108,142108,119639],"quality":"More is better","isMain":false,"metricName":"Total length (>= 25000 bp)"},{"values":[142122,142072,142171,142213,142365,0,0,142199,142108,142108,142108,83236],"quality":"More is better","isMain":true,"metricName":"Total length (>= 50000 bp)"},{"values":[142122,142072,142171,142213,142365,1433,5889,142199,142108,142108,142108,83236],"quality":"More is better","isMain":false,"metricName":"N50"},{"values":[142122,142072,142171,142213,142365,944,3098,142199,142108,142108,142108,36403],"quality":"More is better","isMain":false,"metricName":"N75"},{"values":[1,1,1,1,1,26,8,1,1,1,1,1],"quality":"Less is better","isMain":false,"metricName":"L50"},{"values":[1,1,1,1,1,53,15,1,1,1,1,2],"quality":"Less is better","isMain":false,"metricName":"L75"},{"values":["35.74","35.74","35.74","35.73","35.73","35.73","35.91","35.74","35.74","35.74","35.74","35.81"],"quality":"Equal","isMain":false,"metricName":"GC (%)"}]],["Predicted genes",[]],["Similarity statistics",[{"values":[0,0,0,0,0,0,0,0,0,0,0,0],"quality":"Equal","isMain":false,"metricName":"# similar correct contigs"},{"values":[0,0,0,0,0,0,0,0,0,0,0,0],"quality":"Equal","isMain":false,"metricName":"# similar misassembled blocks"}]],["Reference statistics",[{"values":[142072,142072,142072,142072,142072,142072,142072,142072,142072,142072,142072,142072],"quality":"Equal","isMain":false,"metricName":"Reference length"},{"values":[1,1,1,1,1,1,1,1,1,1,1,1],"quality":"Equal","isMain":false,"metricName":"Reference fragments"},{"values":["35.74","35.74","35.74","35.74","35.74","35.74","35.74","35.74","35.74","35.74","35.74","35.74"],"quality":"Equal","isMain":false,"metricName":"Reference GC (%)"}]]],"referenceName":"gi\_158079296\_ref\_NC","date":"18 July 2018, Wednesday, 16:28:01","order":[0,1,2,3,4,5,6,7,8,9,10,11],"assembliesNames":["ABySS\_63","CLC","IDBA\_UD","MEGAHIT","MIRA","Ray\_Meta","SOAPdenovo2","SPAdes","SPAdes\_meta","SPAdes\_sc","SPAdes\_sc\_careful","Velvet"]},{"assembliesWithNs":null,"minContig":500,"report":[["Genome statistics",[{"values":["99.601","99.289","98.844","98.061","98.267","100.000","99.465","11.993","99.998","99.998","99.998","99.998","97.487","100.000"],"quality":"More is better","isMain":true,"metricName":"Genome fraction (%)"},{"values":["1.000","1.000","1.001","1.000","1.000","1.016","1.000","1.000","1.000","1.000","1.000","1.000","1.001","1.009"],"quality":"Less is better","isMain":true,"metricName":"Duplication ratio"},{"values":[40737,40609,39953,40107,40191,40369,40681,840,40899,40899,40899,40899,17376,41251],"quality":"More is better","isMain":true,"metricName":"Largest alignment"},{"values":[40737,40609,40449,40107,40191,41542,40681,4905,40899,40899,40899,40899,39921,41251],"quality":"More is better","isMain":true,"metricName":"Total aligned length"},{"values":[40737,40609,39953,40107,40191,40369,40681,null,40899,40899,40899,40899,16704,41251],"quality":"More is better","isMain":false,"metricName":"NG50"},{"values":[40737,40609,39953,40107,40191,40369,40681,null,40899,40899,40899,40899,16704,41251],"quality":"More is better","isMain":false,"metricName":"NG75"},{"values":[40737,40609,39953,40107,40191,40369,40681,583,40899,40899,40899,40899,16704,41251],"quality":"More is better","isMain":false,"metricName":"NA50"},{"values":[40737,40609,39953,40107,40191,40369,40681,550,40899,40899,40899,40899,16704,41251],"quality":"More is better","isMain":false,"metricName":"NA75"},{"values":[40737,40609,39953,40107,40191,40369,40681,null,40899,40899,40899,40899,16704,41251],"quality":"More is better","isMain":true,"metricName":"NGA50"},{"values":[40737,40609,39953,40107,40191,40369,40681,null,40899,40899,40899,40899,16704,41251],"quality":"More is better","isMain":false,"metricName":"NGA75"},{"values":[1,1,1,1,1,1,1,null,1,1,1,1,2,1],"quality":"Less is better","isMain":false,"metricName":"LG50"},{"values":[1,1,1,1,1,1,1,null,1,1,1,1,2,1],"quality":"Less is better","isMain":false,"metricName":"LG75"},{"values":[1,1,1,1,1,1,1,4,1,1,1,1,2,1],"quality":"Less is better","isMain":false,"metricName":"LA50"},{"values":[1,1,1,1,1,1,1,6,1,1,1,1,2,1],"quality":"Less is better","isMain":false,"metricName":"LA75"},{"values":[1,1,1,1,1,1,1,null,1,1,1,1,2,1],"quality":"Less is better","isMain":true,"metricName":"LGA50"},{"values":[1,1,1,1,1,1,1,null,1,1,1,1,2,1],"quality":"Less is better","isMain":false,"metricName":"LGA75"}]],["Misassemblies",[{"values":[0,0,0,0,0,0,0,0,0,0,0,0,0,0],"quality":"Less is better","isMain":true,"metricName":"# misassemblies"},{"values":[0,0,0,0,0,0,0,0,0,0,0,0,0,0],"quality":"Less is better","isMain":false,"metricName":" # relocations"},{"values":[0,0,0,0,0,0,0,0,0,0,0,0,0,0],"quality":"Less is better","isMain":false,"metricName":" # translocations"},{"values":[0,0,0,0,0,0,0,0,0,0,0,0,0,0],"quality":"Less is better","isMain":false,"metricName":" # inversions"},{"values":[0,0,0,0,0,0,0,0,0,0,0,0,0,0],"quality":"Less is better","isMain":false,"metricName":"# misassembled contigs"},{"values":[0,0,0,0,0,0,0,0,0,0,0,0,0,0],"quality":"Less is better","isMain":true,"metricName":"Misassembled contigs length"},{"values":[0,0,0,0,0,0,0,0,0,0,0,0,0,1],"quality":"Less is better","isMain":false,"metricName":"# local misassemblies"},{"values":[0,0,0,0,0,0,0,0,0,0,0,0,0,0],"quality":"Less is better","isMain":false,"metricName":"# unaligned mis. contigs"}]],["Unaligned",[{"values":[0,0,0,0,0,0,0,0,0,0,0,0,0,0],"quality":"Less is better","isMain":false,"metricName":"# fully unaligned contigs"},{"values":[0,0,0,0,0,0,0,0,0,0,0,0,0,0],"quality":"Less is better","isMain":false,"metricName":"Fully unaligned length"},{"values":[0,0,0,0,0,0,0,0,0,0,0,0,0,0],"quality":"Less is better","isMain":false,"metricName":"# partially unaligned contigs"},{"values":[0,0,0,0,0,0,0,0,0,0,0,0,0,0],"quality":"Less is better","isMain":false,"metricName":"Partially unaligned length"}]],["Mismatches",[{"values":[0,0,0,0,0,3,0,2,0,0,0,0,0,3],"quality":"Less is better","isMain":false,"metricName":"# mismatches"},{"values":[1,0,0,0,0,1,1,0,0,0,0,0,0,1],"quality":"Less is better","isMain":false,"metricName":"# indels"},{"values":[27,0,0,0,0,1,27,0,0,0,0,0,0,1],"quality":"Less is better","isMain":false,"metricName":"Indels length"},{"values":["0.00","0.00","0.00","0.00","0.00","7.33","0.00","40.77","0.00","0.00","0.00","0.00","0.00","7.33"],"quality":"Less is better","isMain":true,"metricName":"# mismatches per 100 kbp"},{"values":["2.45","0.00","0.00","0.00","0.00","2.44","2.46","0.00","0.00","0.00","0.00","0.00","0.00","2.44"],"quality":"Less is better","isMain":true,"metricName":"# indels per 100 kbp"},{"values":[0,0,0,0,0,1,0,0,0,0,0,0,0,1],"quality":"Less is better","isMain":false,"metricName":" # indels (<= 5 bp)"},{"values":[1,0,0,0,0,0,1,0,0,0,0,0,0,0],"quality":"Less is better","isMain":false,"metricName":" # indels (> 5 bp)"},{"values":[0,0,0,0,0,0,0,0,0,0,0,0,0,0],"quality":"Less is better","isMain":false,"metricName":"# N's"},{"values":["0.00","0.00","0.00","0.00","0.00","0.00","0.00","0.00","0.00","0.00","0.00","0.00","0.00","0.00"],"quality":"Less is better","isMain":true,"metricName":"# N's per 100 kbp"}]],["Statistics without reference",[{"values":[1,1,2,1,1,2,1,8,1,1,1,1,3,1],"quality":"Equal","isMain":true,"metricName":"# contigs"},{"values":[1,1,1,1,1,2,1,0,1,1,1,1,3,1],"quality":"Equal","isMain":false,"metricName":"# contigs (>= 1000 bp)"},{"values":[1,1,1,1,1,1,1,0,1,1,1,1,3,1],"quality":"Equal","isMain":false,"metricName":"# contigs (>= 5000 bp)"},{"values":[1,1,1,1,1,1,1,0,1,1,1,1,2,1],"quality":"Equal","isMain":false,"metricName":"# contigs (>= 10000 bp)"},{"values":[1,1,1,1,1,1,1,0,1,1,1,1,0,1],"quality":"Equal","isMain":false,"metricName":"# contigs (>= 25000 bp)"},{"values":[0,0,0,0,0,0,0,0,0,0,0,0,0,0],"quality":"Equal","isMain":false,"metricName":"# contigs (>= 50000 bp)"},{"values":[40737,40609,39953,40107,40191,40369,40681,840,40899,40899,40899,40899,17376,41251],"quality":"More is better","isMain":true,"metricName":"Largest contig"},{"values":[40737,40609,40471,40107,40191,41542,40681,4905,40899,40899,40899,40899,39921,41251],"quality":"More is better","isMain":true,"metricName":"Total length"},{"values":[40737,40609,39953,40107,40191,41542,40681,0,40899,40899,40899,40899,39921,41251],"quality":"More is better","isMain":true,"metricName":"Total length (>= 1000 bp)"},{"values":[40737,40609,39953,40107,40191,40369,40681,0,40899,40899,40899,40899,39921,41251],"quality":"More is better","isMain":false,"metricName":"Total length (>= 5000 bp)"},{"values":[40737,40609,39953,40107,40191,40369,40681,0,40899,40899,40899,40899,34080,41251],"quality":"More is better","isMain":true,"metricName":"Total length (>= 10000 bp)"},{"values":[40737,40609,39953,40107,40191,40369,40681,0,40899,40899,40899,40899,0,41251],"quality":"More is better","isMain":false,"metricName":"Total length (>= 25000 bp)"},{"values":[0,0,0,0,0,0,0,0,0,0,0,0,0,0],"quality":"More is better","isMain":true,"metricName":"Total length (>= 50000 bp)"},{"values":[40737,40609,39953,40107,40191,40369,40681,583,40899,40899,40899,40899,16704,41251],"quality":"More is better","isMain":false,"metricName":"N50"},{"values":[40737,40609,39953,40107,40191,40369,40681,550,40899,40899,40899,40899,16704,41251],"quality":"More is better","isMain":false,"metricName":"N75"},{"values":[1,1,1,1,1,1,1,4,1,1,1,1,2,1],"quality":"Less is better","isMain":false,"metricName":"L50"},{"values":[1,1,1,1,1,1,1,6,1,1,1,1,2,1],"quality":"Less is better","isMain":false,"metricName":"L75"},{"values":["33.34","33.31","33.30","33.28","33.29","33.37","33.33","32.09","33.35","33.35","33.35","33.35","33.26","33.37"],"quality":"Equal","isMain":false,"metricName":"GC (%)"}]],["Predicted genes",[]],["Similarity statistics",[{"values":[1,1,1,1,1,1,1,0,1,1,1,1,0,0],"quality":"Equal","isMain":false,"metricName":"# similar correct contigs"},{"values":[0,0,0,0,0,0,0,0,0,0,0,0,0,0],"quality":"Equal","isMain":false,"metricName":"# similar misassembled blocks"}]],["Reference statistics",[{"values":[40900,40900,40900,40900,40900,40900,40900,40900,40900,40900,40900,40900,40900,40900],"quality":"Equal","isMain":false,"metricName":"Reference length"},{"values":[1,1,1,1,1,1,1,1,1,1,1,1,1,1],"quality":"Equal","isMain":false,"metricName":"Reference fragments"},{"values":["33.35","33.35","33.35","33.35","33.35","33.35","33.35","33.35","33.35","33.35","33.35","33.35","33.35","33.35"],"quality":"Equal","isMain":false,"metricName":"Reference GC (%)"}]]],"referenceName":"gi\_158267608\_ref\_NC","date":"18 July 2018, Wednesday, 16:28:13","order":[0,1,2,3,4,5,6,7,8,9,10,11,12,13],"assembliesNames":["ABySS\_127","ABySS\_63","CLC","IDBA\_UD","MEGAHIT","MIRA","Ray\_Meta","SOAPdenovo2","SPAdes","SPAdes\_meta","SPAdes\_sc","SPAdes\_sc\_careful","Velvet","Geneious"]},{"assembliesWithNs":null,"minContig":500,"report":[["Genome statistics",[{"values":["29.317","21.068","82.182","79.523","76.763","49.605","75.522","80.009","83.740","84.570","84.570","22.679"],"quality":"More is better","isMain":true,"metricName":"Genome fraction (%)"},{"values":["1.000","1.000","1.010","1.000","1.000","1.004","1.002","1.000","1.006","1.000","1.000","1.000"],"quality":"Less is better","isMain":true,"metricName":"Duplication ratio"},{"values":[1393,1883,4926,5713,5561,2808,4275,5713,5713,5713,5713,2170],"quality":"More is better","isMain":true,"metricName":"Largest alignment"},{"values":[12194,8763,34516,33080,31928,20632,31473,33282,34969,35179,35179,9433],"quality":"More is better","isMain":true,"metricName":"Total aligned length"},{"values":[null,null,1876,2212,1648,null,1316,2363,2363,2363,2363,null],"quality":"More is better","isMain":false,"metricName":"NG50"},{"values":[null,null,955,878,764,null,592,955,1020,1478,1478,null],"quality":"More is better","isMain":false,"metricName":"NG75"},{"values":[1051,950,2254,2227,2079,1321,2156,2416,2416,2416,2416,1389],"quality":"More is better","isMain":false,"metricName":"NA50"},{"values":[666,831,1498,1376,1342,1163,1222,1876,1534,1876,1876,946],"quality":"More is better","isMain":false,"metricName":"NA75"},{"values":[null,null,1876,1701,1648,null,1316,2363,2363,2363,2363,null],"quality":"More is better","isMain":true,"metricName":"NGA50"},{"values":[null,null,955,875,764,null,592,955,1020,1478,1478,null],"quality":"More is better","isMain":false,"metricName":"NGA75"},{"values":[null,null,7,6,7,null,9,6,6,6,6,null],"quality":"Less is better","isMain":false,"metricName":"LG50"},{"values":[null,null,15,14,16,null,19,13,13,12,12,null],"quality":"Less is better","isMain":false,"metricName":"LG75"},{"values":[5,4,6,5,5,6,6,5,5,5,5,3],"quality":"Less is better","isMain":false,"metricName":"LA50"},{"values":[9,6,11,10,9,10,11,8,10,9,9,5],"quality":"Less is better","isMain":false,"metricName":"LA75"},{"values":[null,null,7,7,7,null,9,6,6,6,6,null],"quality":"Less is better","isMain":true,"metricName":"LGA50"},{"values":[null,null,15,15,16,null,19,13,13,12,12,null],"quality":"Less is better","isMain":false,"metricName":"LGA75"}]],["Misassemblies",[{"values":[0,0,0,0,0,0,0,0,0,0,0,0],"quality":"Less is better","isMain":true,"metricName":"# misassemblies"},{"values":[0,0,0,0,0,0,0,0,0,0,0,0],"quality":"Less is better","isMain":false,"metricName":" # relocations"},{"values":[0,0,0,0,0,0,0,0,0,0,0,0],"quality":"Less is better","isMain":false,"metricName":" # translocations"},{"values":[0,0,0,0,0,0,0,0,0,0,0,0],"quality":"Less is better","isMain":false,"metricName":" # inversions"},{"values":[0,0,0,0,0,0,0,0,0,0,0,0],"quality":"Less is better","isMain":false,"metricName":"# misassembled contigs"},{"values":[0,0,0,0,0,0,0,0,0,0,0,0],"quality":"Less is better","isMain":true,"metricName":"Misassembled contigs length"},{"values":[0,0,0,0,0,0,0,0,0,1,1,0],"quality":"Less is better","isMain":false,"metricName":"# local misassemblies"},{"values":[0,0,0,0,0,0,0,0,0,0,0,0],"quality":"Less is better","isMain":false,"metricName":"# unaligned mis. contigs"}]],["Unaligned",[{"values":[0,0,0,0,0,0,0,0,0,0,0,0],"quality":"Less is better","isMain":false,"metricName":"# fully unaligned contigs"},{"values":[0,0,0,0,0,0,0,0,0,0,0,0],"quality":"Less is better","isMain":false,"metricName":"Fully unaligned length"},{"values":[0,0,1,1,0,0,0,0,0,0,0,0],"quality":"Less is better","isMain":false,"metricName":"# partially unaligned contigs"},{"values":[0,0,569,956,0,0,0,0,0,0,0,0],"quality":"Less is better","isMain":false,"metricName":"Partially unaligned length"}]],["Mismatches",[{"values":[4,1,30,44,15,4,16,34,37,31,28,4],"quality":"Less is better","isMain":false,"metricName":"# mismatches"},{"values":[0,0,0,2,0,0,1,0,0,0,0,0],"quality":"Less is better","isMain":false,"metricName":"# indels"},{"values":[0,0,0,2,0,0,40,0,0,0,0,0],"quality":"Less is better","isMain":false,"metricName":"Indels length"},{"values":["32.80","11.41","87.77","133.03","46.98","19.39","50.94","102.17","106.23","88.13","79.60","42.40"],"quality":"Less is better","isMain":true,"metricName":"# mismatches per 100 kbp"},{"values":["0.00","0.00","0.00","6.05","0.00","0.00","3.18","0.00","0.00","0.00","0.00","0.00"],"quality":"Less is better","isMain":true,"metricName":"# indels per 100 kbp"},{"values":[0,0,0,2,0,0,0,0,0,0,0,0],"quality":"Less is better","isMain":false,"metricName":" # indels (<= 5 bp)"},{"values":[0,0,0,0,0,0,1,0,0,0,0,0],"quality":"Less is better","isMain":false,"metricName":" # indels (> 5 bp)"},{"values":[0,0,0,0,0,21,42,0,0,0,0,0],"quality":"Less is better","isMain":false,"metricName":"# N's"},{"values":["0.00","0.00","0.00","0.00","0.00","101.41","133.45","0.00","0.00","0.00","0.00","0.00"],"quality":"Less is better","isMain":true,"metricName":"# N's per 100 kbp"}]],["Statistics without reference",[{"values":[14,9,19,17,17,16,19,15,17,16,16,7],"quality":"Equal","isMain":true,"metricName":"# contigs"},{"values":[6,2,14,13,13,12,14,12,13,13,13,4],"quality":"Equal","isMain":false,"metricName":"# contigs (>= 1000 bp)"},{"values":[0,0,0,1,1,0,0,1,1,1,1,0],"quality":"Equal","isMain":false,"metricName":"# contigs (>= 5000 bp)"},{"values":[0,0,0,0,0,0,0,0,0,0,0,0],"quality":"Equal","isMain":false,"metricName":"# contigs (>= 10000 bp)"},{"values":[0,0,0,0,0,0,0,0,0,0,0,0],"quality":"Equal","isMain":false,"metricName":"# contigs (>= 25000 bp)"},{"values":[0,0,0,0,0,0,0,0,0,0,0,0],"quality":"Equal","isMain":false,"metricName":"# contigs (>= 50000 bp)"},{"values":[1393,1883,4926,5713,5561,2808,4275,5713,5713,5713,5713,2170],"quality":"More is better","isMain":true,"metricName":"Largest contig"},{"values":[12194,8763,35085,34036,31928,20709,31473,33282,35023,35179,35179,9433],"quality":"More is better","isMain":true,"metricName":"Total length"},{"values":[7217,3083,30898,30859,28663,17757,27652,30833,31675,32730,32730,6960],"quality":"More is better","isMain":true,"metricName":"Total length (>= 1000 bp)"},{"values":[0,0,0,5713,5561,0,0,5713,5713,5713,5713,0],"quality":"More is better","isMain":false,"metricName":"Total length (>= 5000 bp)"},{"values":[0,0,0,0,0,0,0,0,0,0,0,0],"quality":"More is better","isMain":true,"metricName":"Total length (>= 10000 bp)"},{"values":[0,0,0,0,0,0,0,0,0,0,0,0],"quality":"More is better","isMain":false,"metricName":"Total length (>= 25000 bp)"},{"values":[0,0,0,0,0,0,0,0,0,0,0,0],"quality":"More is better","isMain":true,"metricName":"Total length (>= 50000 bp)"},{"values":[1051,950,2254,2227,2079,1321,2156,2416,2416,2416,2416,1389],"quality":"More is better","isMain":false,"metricName":"N50"},{"values":[666,831,1498,1444,1342,1163,1222,1876,1534,1876,1876,946],"quality":"More is better","isMain":false,"metricName":"N75"},{"values":[5,4,6,5,5,6,6,5,5,5,5,3],"quality":"Less is better","isMain":false,"metricName":"L50"},{"values":[9,6,11,9,9,10,11,8,10,9,9,5],"quality":"Less is better","isMain":false,"metricName":"L75"},{"values":["60.75","60.15","60.84","60.77","60.81","60.80","60.62","60.89","60.82","60.85","60.85","61.09"],"quality":"Equal","isMain":false,"metricName":"GC (%)"}]],["Predicted genes",[]],["Similarity statistics",[{"values":[0,0,0,0,0,0,0,0,0,0,0,0],"quality":"Equal","isMain":false,"metricName":"# similar correct contigs"},{"values":[0,0,0,0,0,0,0,0,0,0,0,0],"quality":"Equal","isMain":false,"metricName":"# similar misassembled blocks"}]],["Reference statistics",[{"values":[41593,41593,41593,41593,41593,41593,41593,41593,41593,41593,41593,41593],"quality":"Equal","isMain":false,"metricName":"Reference length"},{"values":[1,1,1,1,1,1,1,1,1,1,1,1],"quality":"Equal","isMain":false,"metricName":"Reference fragments"},{"values":["60.90","60.90","60.90","60.90","60.90","60.90","60.90","60.90","60.90","60.90","60.90","60.90"],"quality":"Equal","isMain":false,"metricName":"Reference GC (%)"}]]],"referenceName":"gi\_158345134\_ref\_NC","date":"18 July 2018, Wednesday, 16:28:25","order":[0,1,2,3,4,5,6,7,8,9,10,11],"assembliesNames":["ABySS\_127","ABySS\_63","CLC","IDBA\_UD","MEGAHIT","MIRA","SOAPdenovo2","SPAdes","SPAdes\_meta","SPAdes\_sc","SPAdes\_sc\_careful","Velvet"]},{"assembliesWithNs":null,"minContig":500,"report":[["Genome statistics",[{"values":["100.000","99.951","100.000","100.000","100.000","100.000","100.000","100.000","100.000","100.000","99.701"],"quality":"More is better","isMain":true,"metricName":"Genome fraction (%)"},{"values":["1.001","1.000","1.002","1.003","1.009","1.011","1.003","1.001","1.001","1.001","1.001"],"quality":"Less is better","isMain":true,"metricName":"Duplication ratio"},{"values":[40904,40843,40962,41004,41230,41316,40990,40890,40890,40918,37486],"quality":"More is better","isMain":true,"metricName":"Largest alignment"},{"values":[40904,40843,40962,41004,41230,41316,40990,40890,40890,40918,40766],"quality":"More is better","isMain":true,"metricName":"Total aligned length"},{"values":[40904,40863,40962,41004,41230,41316,40990,40890,40890,40918,37486],"quality":"More is better","isMain":false,"metricName":"NG50"},{"values":[40904,40863,40962,41004,41230,41316,40990,40890,40890,40918,37486],"quality":"More is better","isMain":false,"metricName":"NG75"},{"values":[40904,40843,40962,41004,41230,41316,40990,40890,40890,40918,37486],"quality":"More is better","isMain":false,"metricName":"NA50"},{"values":[40904,40843,40962,41004,41230,41316,40990,40890,40890,40918,37486],"quality":"More is better","isMain":false,"metricName":"NA75"},{"values":[40904,40843,40962,41004,41230,41316,40990,40890,40890,40918,37486],"quality":"More is better","isMain":true,"metricName":"NGA50"},{"values":[40904,40843,40962,41004,41230,41316,40990,40890,40890,40918,37486],"quality":"More is better","isMain":false,"metricName":"NGA75"},{"values":[1,1,1,1,1,1,1,1,1,1,1],"quality":"Less is better","isMain":false,"metricName":"LG50"},{"values":[1,1,1,1,1,1,1,1,1,1,1],"quality":"Less is better","isMain":false,"metricName":"LG75"},{"values":[1,1,1,1,1,1,1,1,1,1,1],"quality":"Less is better","isMain":false,"metricName":"LA50"},{"values":[1,1,1,1,1,1,1,1,1,1,1],"quality":"Less is better","isMain":false,"metricName":"LA75"},{"values":[1,1,1,1,1,1,1,1,1,1,1],"quality":"Less is better","isMain":true,"metricName":"LGA50"},{"values":[1,1,1,1,1,1,1,1,1,1,1],"quality":"Less is better","isMain":false,"metricName":"LGA75"}]],["Misassemblies",[{"values":[0,0,0,0,0,0,0,0,0,0,0],"quality":"Less is better","isMain":true,"metricName":"# misassemblies"},{"values":[0,0,0,0,0,0,0,0,0,0,0],"quality":"Less is better","isMain":false,"metricName":" # relocations"},{"values":[0,0,0,0,0,0,0,0,0,0,0],"quality":"Less is better","isMain":false,"metricName":" # translocations"},{"values":[0,0,0,0,0,0,0,0,0,0,0],"quality":"Less is better","isMain":false,"metricName":" # inversions"},{"values":[0,0,0,0,0,0,0,0,0,0,0],"quality":"Less is better","isMain":false,"metricName":"# misassembled contigs"},{"values":[0,0,0,0,0,0,0,0,0,0,0],"quality":"Less is better","isMain":true,"metricName":"Misassembled contigs length"},{"values":[0,0,1,1,1,1,1,0,0,0,0],"quality":"Less is better","isMain":false,"metricName":"# local misassemblies"},{"values":[0,0,0,0,0,0,0,0,0,0,0],"quality":"Less is better","isMain":false,"metricName":"# unaligned mis. contigs"}]],["Unaligned",[{"values":[0,0,0,0,0,0,0,0,0,0,0],"quality":"Less is better","isMain":false,"metricName":"# fully unaligned contigs"},{"values":[0,0,0,0,0,0,0,0,0,0,0],"quality":"Less is better","isMain":false,"metricName":"Fully unaligned length"},{"values":[0,0,0,0,0,0,0,0,0,0,0],"quality":"Less is better","isMain":false,"metricName":"# partially unaligned contigs"},{"values":[0,0,0,0,0,0,0,0,0,0,0],"quality":"Less is better","isMain":false,"metricName":"Partially unaligned length"}]],["Mismatches",[{"values":[0,0,0,0,0,0,0,0,0,0,1],"quality":"Less is better","isMain":false,"metricName":"# mismatches"},{"values":[1,0,0,0,0,0,0,1,1,1,0],"quality":"Less is better","isMain":false,"metricName":"# indels"},{"values":[41,0,0,0,0,0,0,27,27,55,0],"quality":"Less is better","isMain":false,"metricName":"Indels length"},{"values":["0.00","0.00","0.00","0.00","0.00","0.00","0.00","0.00","0.00","0.00","2.45"],"quality":"Less is better","isMain":true,"metricName":"# mismatches per 100 kbp"},{"values":["2.45","0.00","0.00","0.00","0.00","0.00","0.00","2.45","2.45","2.45","0.00"],"quality":"Less is better","isMain":true,"metricName":"# indels per 100 kbp"},{"values":[0,0,0,0,0,0,0,0,0,0,0],"quality":"Less is better","isMain":false,"metricName":" # indels (<= 5 bp)"},{"values":[1,0,0,0,0,0,0,1,1,1,0],"quality":"Less is better","isMain":false,"metricName":" # indels (> 5 bp)"},{"values":[0,0,0,0,0,0,0,0,0,0,0],"quality":"Less is better","isMain":false,"metricName":"# N's"},{"values":["0.00","0.00","0.00","0.00","0.00","0.00","0.00","0.00","0.00","0.00","0.00"],"quality":"Less is better","isMain":true,"metricName":"# N's per 100 kbp"}]],["Statistics without reference",[{"values":[1,1,1,1,1,1,1,1,1,1,2],"quality":"Equal","isMain":true,"metricName":"# contigs"},{"values":[1,1,1,1,1,1,1,1,1,1,2],"quality":"Equal","isMain":false,"metricName":"# contigs (>= 1000 bp)"},{"values":[1,1,1,1,1,1,1,1,1,1,1],"quality":"Equal","isMain":false,"metricName":"# contigs (>= 5000 bp)"},{"values":[1,1,1,1,1,1,1,1,1,1,1],"quality":"Equal","isMain":false,"metricName":"# contigs (>= 10000 bp)"},{"values":[1,1,1,1,1,1,1,1,1,1,1],"quality":"Equal","isMain":false,"metricName":"# contigs (>= 25000 bp)"},{"values":[0,0,0,0,0,0,0,0,0,0,0],"quality":"Equal","isMain":false,"metricName":"# contigs (>= 50000 bp)"},{"values":[40904,40863,40962,41004,41230,41316,40990,40890,40890,40918,37486],"quality":"More is better","isMain":true,"metricName":"Largest contig"},{"values":[40904,40863,40962,41004,41230,41316,40990,40890,40890,40918,40766],"quality":"More is better","isMain":true,"metricName":"Total length"},{"values":[40904,40863,40962,41004,41230,41316,40990,40890,40890,40918,40766],"quality":"More is better","isMain":true,"metricName":"Total length (>= 1000 bp)"},{"values":[40904,40863,40962,41004,41230,41316,40990,40890,40890,40918,37486],"quality":"More is better","isMain":false,"metricName":"Total length (>= 5000 bp)"},{"values":[40904,40863,40962,41004,41230,41316,40990,40890,40890,40918,37486],"quality":"More is better","isMain":true,"metricName":"Total length (>= 10000 bp)"},{"values":[40904,40863,40962,41004,41230,41316,40990,40890,40890,40918,37486],"quality":"More is better","isMain":false,"metricName":"Total length (>= 25000 bp)"},{"values":[0,0,0,0,0,0,0,0,0,0,0],"quality":"More is better","isMain":true,"metricName":"Total length (>= 50000 bp)"},{"values":[40904,40863,40962,41004,41230,41316,40990,40890,40890,40918,37486],"quality":"More is better","isMain":false,"metricName":"N50"},{"values":[40904,40863,40962,41004,41230,41316,40990,40890,40890,40918,37486],"quality":"More is better","isMain":false,"metricName":"N75"},{"values":[1,1,1,1,1,1,1,1,1,1,1],"quality":"Less is better","isMain":false,"metricName":"L50"},{"values":[1,1,1,1,1,1,1,1,1,1,1],"quality":"Less is better","isMain":false,"metricName":"L75"},{"values":["44.80","44.79","44.81","44.79","44.68","44.82","44.79","44.80","44.80","44.80","44.75"],"quality":"Equal","isMain":false,"metricName":"GC (%)"}]],["Predicted genes",[]],["Similarity statistics",[{"values":[0,0,0,0,0,0,0,0,0,0,0],"quality":"Equal","isMain":false,"metricName":"# similar correct contigs"},{"values":[0,0,0,0,0,0,0,0,0,0,0],"quality":"Equal","isMain":false,"metricName":"# similar misassembled blocks"}]],["Reference statistics",[{"values":[40863,40863,40863,40863,40863,40863,40863,40863,40863,40863,40863],"quality":"Equal","isMain":false,"metricName":"Reference length"},{"values":[1,1,1,1,1,1,1,1,1,1,1],"quality":"Equal","isMain":false,"metricName":"Reference fragments"},{"values":["44.79","44.79","44.79","44.79","44.79","44.79","44.79","44.79","44.79","44.79","44.79"],"quality":"Equal","isMain":false,"metricName":"Reference GC (%)"}]]],"referenceName":"gi\_158380386\_ref\_NC","date":"18 July 2018, Wednesday, 16:28:35","order":[0,1,2,3,4,5,6,7,8,9,10],"assembliesNames":["ABySS\_63","CLC","IDBA\_UD","MEGAHIT","MIRA","Ray\_Meta","SPAdes","SPAdes\_meta","SPAdes\_sc","SPAdes\_sc\_careful","Velvet"]},{"assembliesWithNs":null,"minContig":500,"report":[["Genome statistics",[{"values":["99.997","100.000","100.000","100.000","100.000","12.041","97.955","100.000","100.000","100.000","100.000","100.000","100.000"],"quality":"More is better","isMain":true,"metricName":"Genome fraction (%)"},{"values":["1.000","1.000","1.003","1.004","1.025","1.000","1.045","1.004","1.002","1.002","1.002","1.001","1.008"],"quality":"Less is better","isMain":true,"metricName":"Duplication ratio"},{"values":[33984,33985,34084,34126,34240,923,33522,34112,34040,34040,34040,34015,34240],"quality":"More is better","isMain":true,"metricName":"Largest alignment"},{"values":[33984,33985,34084,34126,34851,4092,33522,34112,34040,34040,34040,34015,34240],"quality":"More is better","isMain":true,"metricName":"Total aligned length"},{"values":[33984,33985,34084,34126,34240,null,34781,34112,34040,34040,34040,34015,34240],"quality":"More is better","isMain":false,"metricName":"NG50"},{"values":[33984,33985,34084,34126,34240,null,34781,34112,34040,34040,34040,34015,34240],"quality":"More is better","isMain":false,"metricName":"NG75"},{"values":[33984,33985,34084,34126,34240,750,33522,34112,34040,34040,34040,34015,34240],"quality":"More is better","isMain":false,"metricName":"NA50"},{"values":[33984,33985,34084,34126,34240,528,33522,34112,34040,34040,34040,34015,34240],"quality":"More is better","isMain":false,"metricName":"NA75"},{"values":[33984,33985,34084,34126,34240,null,33522,34112,34040,34040,34040,34015,34240],"quality":"More is better","isMain":true,"metricName":"NGA50"},{"values":[33984,33985,34084,34126,34240,null,33522,34112,34040,34040,34040,34015,34240],"quality":"More is better","isMain":false,"metricName":"NGA75"},{"values":[1,1,1,1,1,null,1,1,1,1,1,1,1],"quality":"Less is better","isMain":false,"metricName":"LG50"},{"values":[1,1,1,1,1,null,1,1,1,1,1,1,1],"quality":"Less is better","isMain":false,"metricName":"LG75"},{"values":[1,1,1,1,1,3,1,1,1,1,1,1,1],"quality":"Less is better","isMain":false,"metricName":"LA50"},{"values":[1,1,1,1,1,5,1,1,1,1,1,1,1],"quality":"Less is better","isMain":false,"metricName":"LA75"},{"values":[1,1,1,1,1,null,1,1,1,1,1,1,1],"quality":"Less is better","isMain":true,"metricName":"LGA50"},{"values":[1,1,1,1,1,null,1,1,1,1,1,1,1],"quality":"Less is better","isMain":false,"metricName":"LGA75"}]],["Misassemblies",[{"values":[0,0,0,0,0,0,0,0,0,0,0,0,0],"quality":"Less is better","isMain":true,"metricName":"# misassemblies"},{"values":[0,0,0,0,0,0,0,0,0,0,0,0,0],"quality":"Less is better","isMain":false,"metricName":" # relocations"},{"values":[0,0,0,0,0,0,0,0,0,0,0,0,0],"quality":"Less is better","isMain":false,"metricName":" # translocations"},{"values":[0,0,0,0,0,0,0,0,0,0,0,0,0],"quality":"Less is better","isMain":false,"metricName":" # inversions"},{"values":[0,0,0,0,0,0,0,0,0,0,0,0,0],"quality":"Less is better","isMain":false,"metricName":"# misassembled contigs"},{"values":[0,0,0,0,0,0,0,0,0,0,0,0,0],"quality":"Less is better","isMain":true,"metricName":"Misassembled contigs length"},{"values":[0,0,1,1,1,0,0,1,0,0,0,0,1],"quality":"Less is better","isMain":false,"metricName":"# local misassemblies"},{"values":[0,0,0,0,0,0,0,0,0,0,0,0,0],"quality":"Less is better","isMain":false,"metricName":"# unaligned mis. contigs"}]],["Unaligned",[{"values":[0,0,0,0,0,0,0,0,0,0,0,0,0],"quality":"Less is better","isMain":false,"metricName":"# fully unaligned contigs"},{"values":[0,0,0,0,0,0,0,0,0,0,0,0,0],"quality":"Less is better","isMain":false,"metricName":"Fully unaligned length"},{"values":[0,0,0,0,0,0,0,0,0,0,0,0,0],"quality":"Less is better","isMain":false,"metricName":"# partially unaligned contigs"},{"values":[0,0,0,0,0,0,0,0,0,0,0,0,0],"quality":"Less is better","isMain":false,"metricName":"Partially unaligned length"}]],["Mismatches",[{"values":[0,0,0,0,0,0,0,0,0,0,0,0,0],"quality":"Less is better","isMain":false,"metricName":"# mismatches"},{"values":[0,0,0,0,0,0,17,0,1,1,1,1,0],"quality":"Less is better","isMain":false,"metricName":"# indels"},{"values":[0,0,0,0,0,0,820,0,55,55,55,30,0],"quality":"Less is better","isMain":false,"metricName":"Indels length"},{"values":["0.00","0.00","0.00","0.00","0.00","0.00","0.00","0.00","0.00","0.00","0.00","0.00","0.00"],"quality":"Less is better","isMain":true,"metricName":"# mismatches per 100 kbp"},{"values":["0.00","0.00","0.00","0.00","0.00","0.00","51.07","0.00","2.94","2.94","2.94","2.94","0.00"],"quality":"Less is better","isMain":true,"metricName":"# indels per 100 kbp"},{"values":[0,0,0,0,0,0,0,0,0,0,0,0,0],"quality":"Less is better","isMain":false,"metricName":" # indels (<= 5 bp)"},{"values":[0,0,0,0,0,0,17,0,1,1,1,1,0],"quality":"Less is better","isMain":false,"metricName":" # indels (> 5 bp)"},{"values":[0,0,0,0,2,0,1564,0,0,0,0,0,2],"quality":"Less is better","isMain":false,"metricName":"# N's"},{"values":["0.00","0.00","0.00","0.00","5.74","0.00","4496.71","0.00","0.00","0.00","0.00","0.00","5.84"],"quality":"Less is better","isMain":true,"metricName":"# N's per 100 kbp"}]],["Statistics without reference",[{"values":[1,1,1,1,2,6,1,1,1,1,1,1,1],"quality":"Equal","isMain":true,"metricName":"# contigs"},{"values":[1,1,1,1,1,0,1,1,1,1,1,1,1],"quality":"Equal","isMain":false,"metricName":"# contigs (>= 1000 bp)"},{"values":[1,1,1,1,1,0,1,1,1,1,1,1,1],"quality":"Equal","isMain":false,"metricName":"# contigs (>= 5000 bp)"},{"values":[1,1,1,1,1,0,1,1,1,1,1,1,1],"quality":"Equal","isMain":false,"metricName":"# contigs (>= 10000 bp)"},{"values":[1,1,1,1,1,0,1,1,1,1,1,1,1],"quality":"Equal","isMain":false,"metricName":"# contigs (>= 25000 bp)"},{"values":[0,0,0,0,0,0,0,0,0,0,0,0,0],"quality":"Equal","isMain":false,"metricName":"# contigs (>= 50000 bp)"},{"values":[33984,33985,34084,34126,34240,923,34781,34112,34040,34040,34040,34015,34240],"quality":"More is better","isMain":true,"metricName":"Largest contig"},{"values":[33984,33985,34084,34126,34851,4092,34781,34112,34040,34040,34040,34015,34240],"quality":"More is better","isMain":true,"metricName":"Total length"},{"values":[33984,33985,34084,34126,34240,0,34781,34112,34040,34040,34040,34015,34240],"quality":"More is better","isMain":true,"metricName":"Total length (>= 1000 bp)"},{"values":[33984,33985,34084,34126,34240,0,34781,34112,34040,34040,34040,34015,34240],"quality":"More is better","isMain":false,"metricName":"Total length (>= 5000 bp)"},{"values":[33984,33985,34084,34126,34240,0,34781,34112,34040,34040,34040,34015,34240],"quality":"More is better","isMain":true,"metricName":"Total length (>= 10000 bp)"},{"values":[33984,33985,34084,34126,34240,0,34781,34112,34040,34040,34040,34015,34240],"quality":"More is better","isMain":false,"metricName":"Total length (>= 25000 bp)"},{"values":[0,0,0,0,0,0,0,0,0,0,0,0,0],"quality":"More is better","isMain":true,"metricName":"Total length (>= 50000 bp)"},{"values":[33984,33985,34084,34126,34240,750,34781,34112,34040,34040,34040,34015,34240],"quality":"More is better","isMain":false,"metricName":"N50"},{"values":[33984,33985,34084,34126,34240,528,34781,34112,34040,34040,34040,34015,34240],"quality":"More is better","isMain":false,"metricName":"N75"},{"values":[1,1,1,1,1,3,1,1,1,1,1,1,1],"quality":"Less is better","isMain":false,"metricName":"L50"},{"values":[1,1,1,1,1,5,1,1,1,1,1,1,1],"quality":"Less is better","isMain":false,"metricName":"L75"},{"values":["61.74","61.74","61.75","61.78","61.76","62.27","61.75","61.78","61.74","61.74","61.74","61.74","61.67"],"quality":"Equal","isMain":false,"metricName":"GC (%)"}]],["Predicted genes",[]],["Similarity statistics",[{"values":[0,0,0,0,0,0,0,0,0,0,0,0,0],"quality":"Equal","isMain":false,"metricName":"# similar correct contigs"},{"values":[0,0,0,0,0,0,0,0,0,0,0,0,0],"quality":"Equal","isMain":false,"metricName":"# similar misassembled blocks"}]],["Reference statistics",[{"values":[33985,33985,33985,33985,33985,33985,33985,33985,33985,33985,33985,33985,33985],"quality":"Equal","isMain":false,"metricName":"Reference length"},{"values":[1,1,1,1,1,1,1,1,1,1,1,1,1],"quality":"Equal","isMain":false,"metricName":"Reference fragments"},{"values":["61.74","61.74","61.74","61.74","61.74","61.74","61.74","61.74","61.74","61.74","61.74","61.74","61.74"],"quality":"Equal","isMain":false,"metricName":"Reference GC (%)"}]]],"referenceName":"gi\_158518661\_ref\_NC","date":"18 July 2018, Wednesday, 16:28:48","order":[0,1,2,3,4,5,6,7,8,9,10,11,12],"assembliesNames":["ABySS\_127","CLC","IDBA\_UD","MEGAHIT","MIRA","Ray\_Meta","SOAPdenovo2","SPAdes","SPAdes\_meta","SPAdes\_sc","SPAdes\_sc\_careful","Velvet","Geneious"]},{"assembliesWithNs":null,"minContig":500,"report":[["Genome statistics",[{"values":["93.887","76.366","100.000","92.303","99.056","100.000","11.941","30.295","99.869","82.171","83.602","89.886","51.519","100.000"],"quality":"More is better","isMain":true,"metricName":"Genome fraction (%)"},{"values":["1.057","1.006","1.000","1.009","1.060","1.005","1.000","1.108","1.000","1.000","1.006","1.009","1.002","1.014"],"quality":"Less is better","isMain":true,"metricName":"Duplication ratio"},{"values":[51186,9251,63879,13208,52196,64169,1264,5611,63795,47471,12049,12049,5892,64794],"quality":"More is better","isMain":true,"metricName":"Largest alignment"},{"values":[63384,49058,63879,59479,67095,64169,7628,19546,63795,52502,53724,57956,32964,64794],"quality":"More is better","isMain":true,"metricName":"Total aligned length"},{"values":[116165,2851,63879,6662,52196,64169,null,6255,63795,51774,4582,4582,570,126913],"quality":"More is better","isMain":false,"metricName":"NG50"},{"values":[116165,626,63879,2748,52196,64169,null,1476,63795,51774,1254,1703,null,126913],"quality":"More is better","isMain":false,"metricName":"NG75"},{"values":[2146,6267,63879,7194,52196,64169,634,null,63795,47471,6816,6816,2014,64794],"quality":"More is better","isMain":false,"metricName":"NA50"},{"values":[null,2380,63879,2855,52196,64169,613,null,63795,47471,2309,1836,979,null],"quality":"More is better","isMain":false,"metricName":"NA75"},{"values":[51186,2851,63879,6662,52196,64169,null,null,63795,47471,4582,4582,570,64794],"quality":"More is better","isMain":true,"metricName":"NGA50"},{"values":[51186,626,63879,2748,52196,64169,null,null,63795,5031,1254,1703,null,64794],"quality":"More is better","isMain":false,"metricName":"NGA75"},{"values":[1,5,1,4,1,1,null,4,1,1,4,4,20,1],"quality":"Less is better","isMain":false,"metricName":"LG50"},{"values":[1,17,1,7,1,1,null,12,1,1,11,10,null,1],"quality":"Less is better","isMain":false,"metricName":"LG75"},{"values":[4,4,1,3,1,1,5,null,1,1,3,3,4,1],"quality":"Less is better","isMain":false,"metricName":"LA50"},{"values":[null,7,1,6,1,1,8,null,1,1,6,8,10,null],"quality":"Less is better","isMain":false,"metricName":"LA75"},{"values":[1,5,1,4,1,1,null,null,1,1,4,4,20,1],"quality":"Less is better","isMain":true,"metricName":"LGA50"},{"values":[1,17,1,7,1,1,null,null,1,2,11,10,null,1],"quality":"Less is better","isMain":false,"metricName":"LGA75"}]],["Misassemblies",[{"values":[0,0,0,0,0,0,0,0,0,0,0,0,0,0],"quality":"Less is better","isMain":true,"metricName":"# misassemblies"},{"values":[0,0,0,0,0,0,0,0,0,0,0,0,0,0],"quality":"Less is better","isMain":false,"metricName":" # relocations"},{"values":[0,0,0,0,0,0,0,0,0,0,0,0,0,0],"quality":"Less is better","isMain":false,"metricName":" # translocations"},{"values":[0,0,0,0,0,0,0,0,0,0,0,0,0,0],"quality":"Less is better","isMain":false,"metricName":" # inversions"},{"values":[0,0,0,0,0,0,0,0,0,0,0,0,0,0],"quality":"Less is better","isMain":false,"metricName":"# misassembled contigs"},{"values":[0,0,0,0,0,0,0,0,0,0,0,0,0,0],"quality":"Less is better","isMain":true,"metricName":"Misassembled contigs length"},{"values":[0,0,0,0,0,1,0,1,0,2,0,0,0,1],"quality":"Less is better","isMain":false,"metricName":"# local misassemblies"},{"values":[1,0,0,0,0,0,0,11,0,0,0,0,0,0],"quality":"Less is better","isMain":false,"metricName":"# unaligned mis. contigs"}]],["Unaligned",[{"values":[0,0,0,0,0,0,0,0,0,0,0,0,0,0],"quality":"Less is better","isMain":false,"metricName":"# fully unaligned contigs"},{"values":[0,0,0,0,0,0,0,0,0,0,0,0,0,0],"quality":"Less is better","isMain":false,"metricName":"Fully unaligned length"},{"values":[1,0,0,0,0,0,0,16,0,1,0,0,0,1],"quality":"Less is better","isMain":false,"metricName":"# partially unaligned contigs"},{"values":[59530,0,0,0,0,0,0,33407,0,4303,0,0,0,62119],"quality":"Less is better","isMain":false,"metricName":"Partially unaligned length"}]],["Mismatches",[{"values":[1,0,6,1,2,0,0,0,0,655,31,1,0,0],"quality":"Less is better","isMain":false,"metricName":"# mismatches"},{"values":[3,0,0,0,0,0,0,10,0,36,0,2,0,0],"quality":"Less is better","isMain":false,"metricName":"# indels"},{"values":[4,0,0,0,0,0,0,475,0,46,0,6,0,0],"quality":"Less is better","isMain":false,"metricName":"Indels length"},{"values":["1.67","0.00","9.39","1.70","3.16","0.00","0.00","0.00","0.00","1247.86","58.05","1.74","0.00","0.00"],"quality":"Less is better","isMain":true,"metricName":"# mismatches per 100 kbp"},{"values":["5.00","0.00","0.00","0.00","0.00","0.00","0.00","51.67","0.00","68.58","0.00","3.48","0.00","0.00"],"quality":"Less is better","isMain":true,"metricName":"# indels per 100 kbp"},{"values":[3,0,0,0,0,0,0,0,0,36,0,2,0,0],"quality":"Less is better","isMain":false,"metricName":" # indels (<= 5 bp)"},{"values":[0,0,0,0,0,0,0,10,0,0,0,0,0,0],"quality":"Less is better","isMain":false,"metricName":" # indels (> 5 bp)"},{"values":[0,0,0,0,0,0,0,14200,0,0,0,0,0,18],"quality":"Less is better","isMain":false,"metricName":"# N's"},{"values":["0.00","0.00","0.00","0.00","0.00","0.00","0.00","25892.09","0.00","0.00","0.00","0.00","0.00","14.18"],"quality":"Less is better","isMain":true,"metricName":"# N's per 100 kbp"}]],["Statistics without reference",[{"values":[3,18,1,15,5,1,11,18,1,2,19,23,21,1],"quality":"Equal","isMain":true,"metricName":"# contigs"},{"values":[3,11,1,11,5,1,1,16,1,2,11,12,9,1],"quality":"Equal","isMain":false,"metricName":"# contigs (>= 1000 bp)"},{"values":[1,4,1,5,2,1,0,4,1,2,3,3,2,1],"quality":"Equal","isMain":false,"metricName":"# contigs (>= 5000 bp)"},{"values":[1,0,1,2,1,1,0,0,1,1,2,2,0,1],"quality":"Equal","isMain":false,"metricName":"# contigs (>= 10000 bp)"},{"values":[1,0,1,0,1,1,0,0,1,1,0,0,0,1],"quality":"Equal","isMain":false,"metricName":"# contigs (>= 25000 bp)"},{"values":[1,0,1,0,1,1,0,0,1,1,0,0,0,1],"quality":"Equal","isMain":false,"metricName":"# contigs (>= 50000 bp)"},{"values":[116165,9251,63879,13208,52196,64169,1264,9658,63795,51774,12049,12049,5892,126913],"quality":"More is better","isMain":true,"metricName":"Largest contig"},{"values":[122914,49058,63879,59479,67095,64169,7628,54843,63795,56808,53727,57959,32964,126913],"quality":"More is better","isMain":true,"metricName":"Total length"},{"values":[122914,44009,63879,56878,67095,64169,1264,53027,63795,56808,48953,50794,24623,126913],"quality":"More is better","isMain":true,"metricName":"Total length (>= 1000 bp)"},{"values":[116165,29258,63879,42920,60593,64169,0,32069,63795,56808,30123,30123,11187,126913],"quality":"More is better","isMain":false,"metricName":"Total length (>= 5000 bp)"},{"values":[116165,0,63879,24033,52196,64169,0,0,63795,51774,23307,23307,0,126913],"quality":"More is better","isMain":true,"metricName":"Total length (>= 10000 bp)"},{"values":[116165,0,63879,0,52196,64169,0,0,63795,51774,0,0,0,126913],"quality":"More is better","isMain":false,"metricName":"Total length (>= 25000 bp)"},{"values":[116165,0,63879,0,52196,64169,0,0,63795,51774,0,0,0,126913],"quality":"More is better","isMain":true,"metricName":"Total length (>= 50000 bp)"},{"values":[116165,6267,63879,7194,52196,64169,634,6255,63795,51774,6816,6816,2014,126913],"quality":"More is better","isMain":false,"metricName":"N50"},{"values":[116165,2380,63879,2855,52196,64169,613,2051,63795,51774,2309,1836,979,126913],"quality":"More is better","isMain":false,"metricName":"N75"},{"values":[1,4,1,3,1,1,5,4,1,1,3,3,4,1],"quality":"Less is better","isMain":false,"metricName":"L50"},{"values":[1,7,1,6,1,1,8,8,1,1,6,8,10,1],"quality":"Less is better","isMain":false,"metricName":"L75"},{"values":["65.75","65.12","65.30","65.25","65.32","65.30","63.82","64.66","65.31","65.05","65.22","65.30","64.11","65.75"],"quality":"Equal","isMain":false,"metricName":"GC (%)"}]],["Predicted genes",[]],["Similarity statistics",[{"values":[0,0,0,0,0,0,0,0,0,0,0,0,0,0],"quality":"Equal","isMain":false,"metricName":"# similar correct contigs"},{"values":[0,0,0,0,0,0,0,0,0,0,0,0,0,0],"quality":"Equal","isMain":false,"metricName":"# similar misassembled blocks"}]],["Reference statistics",[{"values":[63879,63879,63879,63879,63879,63879,63879,63879,63879,63879,63879,63879,63879,63879],"quality":"Equal","isMain":false,"metricName":"Reference length"},{"values":[1,1,1,1,1,1,1,1,1,1,1,1,1,1],"quality":"Equal","isMain":false,"metricName":"Reference fragments"},{"values":["65.31","65.31","65.31","65.31","65.31","65.31","65.31","65.31","65.31","65.31","65.31","65.31","65.31","65.31"],"quality":"Equal","isMain":false,"metricName":"Reference GC (%)"}]]],"referenceName":"gi\_158997720\_ref\_NC","date":"18 July 2018, Wednesday, 16:29:02","order":[0,1,2,3,4,5,6,7,8,9,10,11,12,13],"assembliesNames":["ABySS\_127","ABySS\_63","CLC","IDBA\_UD","MEGAHIT","MIRA","Ray\_Meta","SOAPdenovo2","SPAdes","SPAdes\_meta","SPAdes\_sc","SPAdes\_sc\_careful","Velvet","Geneious"]},{"assembliesWithNs":null,"minContig":500,"report":[["Genome statistics",[{"values":["69.833","57.299","95.900","90.641","90.809","81.389","87.439","92.792","92.785","92.423","92.423","75.869","17.251"],"quality":"More is better","isMain":true,"metricName":"Genome fraction (%)"},{"values":["1.002","1.000","1.042","1.000","1.002","1.038","1.000","1.005","1.000","1.000","1.000","1.003","1.000"],"quality":"Less is better","isMain":true,"metricName":"Duplication ratio"},{"values":[2492,2492,7865,7708,7799,3458,7533,7865,7865,7865,7865,4120,2575],"quality":"More is better","isMain":true,"metricName":"Largest alignment"},{"values":[10424,8553,14915,13530,13555,12615,13052,13851,13850,13796,13796,11354,2575],"quality":"More is better","isMain":true,"metricName":"Total aligned length"},{"values":[737,748,7865,7708,7799,1383,7533,7865,7865,7865,7865,3774,null],"quality":"More is better","isMain":false,"metricName":"NG50"},{"values":[null,null,1980,5082,5079,766,946,5156,5092,5038,5038,730,null],"quality":"More is better","isMain":false,"metricName":"NG75"},{"values":[1533,2087,7865,7708,7799,3378,7533,7865,7865,7865,7865,3774,2575],"quality":"More is better","isMain":false,"metricName":"NA50"},{"values":[737,784,1980,5082,5048,1269,2917,5093,5092,5038,5038,1487,2575],"quality":"More is better","isMain":false,"metricName":"NA75"},{"values":[737,748,7865,7708,7799,1383,7533,7865,7865,7865,7865,3774,null],"quality":"More is better","isMain":true,"metricName":"NGA50"},{"values":[null,null,1980,5082,5048,766,946,5093,5092,5038,5038,730,null],"quality":"More is better","isMain":false,"metricName":"NGA75"},{"values":[5,6,1,1,1,3,1,1,1,1,1,2,null],"quality":"Less is better","isMain":false,"metricName":"LG50"},{"values":[null,null,3,2,2,6,3,2,2,2,2,5,null],"quality":"Less is better","isMain":false,"metricName":"LG75"},{"values":[3,2,1,1,1,2,1,1,1,1,1,2,1],"quality":"Less is better","isMain":false,"metricName":"LA50"},{"values":[5,4,3,2,2,4,2,2,2,2,2,3,1],"quality":"Less is better","isMain":false,"metricName":"LA75"},{"values":[5,6,1,1,1,3,1,1,1,1,1,2,null],"quality":"Less is better","isMain":true,"metricName":"LGA50"},{"values":[null,null,3,2,2,6,3,2,2,2,2,5,null],"quality":"Less is better","isMain":false,"metricName":"LGA75"}]],["Misassemblies",[{"values":[0,0,0,0,0,0,0,0,0,0,0,0,0],"quality":"Less is better","isMain":true,"metricName":"# misassemblies"},{"values":[0,0,0,0,0,0,0,0,0,0,0,0,0],"quality":"Less is better","isMain":false,"metricName":" # relocations"},{"values":[0,0,0,0,0,0,0,0,0,0,0,0,0],"quality":"Less is better","isMain":false,"metricName":" # translocations"},{"values":[0,0,0,0,0,0,0,0,0,0,0,0,0],"quality":"Less is better","isMain":false,"metricName":" # inversions"},{"values":[0,0,0,0,0,0,0,0,0,0,0,0,0],"quality":"Less is better","isMain":false,"metricName":"# misassembled contigs"},{"values":[0,0,0,0,0,0,0,0,0,0,0,0,0],"quality":"Less is better","isMain":true,"metricName":"Misassembled contigs length"},{"values":[0,0,0,0,0,0,0,0,0,0,0,0,0],"quality":"Less is better","isMain":false,"metricName":"# local misassemblies"},{"values":[0,0,0,0,0,0,0,0,0,0,0,0,0],"quality":"Less is better","isMain":false,"metricName":"# unaligned mis. contigs"}]],["Unaligned",[{"values":[0,0,0,0,0,0,0,0,0,0,0,0,0],"quality":"Less is better","isMain":false,"metricName":"# fully unaligned contigs"},{"values":[0,0,0,0,0,0,0,0,0,0,0,0,0],"quality":"Less is better","isMain":false,"metricName":"Fully unaligned length"},{"values":[0,0,0,0,0,0,0,0,0,0,0,0,0],"quality":"Less is better","isMain":false,"metricName":"# partially unaligned contigs"},{"values":[0,0,0,0,0,0,0,0,0,0,0,0,0],"quality":"Less is better","isMain":false,"metricName":"Partially unaligned length"}]],["Mismatches",[{"values":[4,1,4,1,1,2,2,7,5,4,3,1,0],"quality":"Less is better","isMain":false,"metricName":"# mismatches"},{"values":[0,0,0,0,0,0,0,0,0,0,0,0,0],"quality":"Less is better","isMain":false,"metricName":"# indels"},{"values":[0,0,0,0,0,0,0,0,0,0,0,0,0],"quality":"Less is better","isMain":false,"metricName":"Indels length"},{"values":["38.37","11.69","27.94","7.39","7.38","16.46","15.32","50.54","36.10","28.99","21.75","8.83","0.00"],"quality":"Less is better","isMain":true,"metricName":"# mismatches per 100 kbp"},{"values":["0.00","0.00","0.00","0.00","0.00","0.00","0.00","0.00","0.00","0.00","0.00","0.00","0.00"],"quality":"Less is better","isMain":true,"metricName":"# indels per 100 kbp"},{"values":[0,0,0,0,0,0,0,0,0,0,0,0,0],"quality":"Less is better","isMain":false,"metricName":" # indels (<= 5 bp)"},{"values":[0,0,0,0,0,0,0,0,0,0,0,0,0],"quality":"Less is better","isMain":false,"metricName":" # indels (> 5 bp)"},{"values":[0,0,0,0,0,9,0,0,0,0,0,0,9],"quality":"Less is better","isMain":false,"metricName":"# N's"},{"values":["0.00","0.00","0.00","0.00","0.00","71.34","0.00","0.00","0.00","0.00","0.00","0.00","349.51"],"quality":"Less is better","isMain":true,"metricName":"# N's per 100 kbp"}]],["Statistics without reference",[{"values":[9,7,5,3,3,8,5,3,3,3,3,5,1],"quality":"Equal","isMain":true,"metricName":"# contigs"},{"values":[4,3,4,2,2,5,2,2,2,2,2,4,1],"quality":"Equal","isMain":false,"metricName":"# contigs (>= 1000 bp)"},{"values":[0,0,1,2,2,0,1,2,2,2,2,0,0],"quality":"Equal","isMain":false,"metricName":"# contigs (>= 5000 bp)"},{"values":[0,0,0,0,0,0,0,0,0,0,0,0,0],"quality":"Equal","isMain":false,"metricName":"# contigs (>= 10000 bp)"},{"values":[0,0,0,0,0,0,0,0,0,0,0,0,0],"quality":"Equal","isMain":false,"metricName":"# contigs (>= 25000 bp)"},{"values":[0,0,0,0,0,0,0,0,0,0,0,0,0],"quality":"Equal","isMain":false,"metricName":"# contigs (>= 50000 bp)"},{"values":[2492,2492,7865,7708,7799,3458,7533,7865,7865,7865,7865,4120,2575],"quality":"More is better","isMain":true,"metricName":"Largest contig"},{"values":[10440,8553,14915,13530,13586,12615,13052,13914,13850,13796,13796,11354,2575],"quality":"More is better","isMain":true,"metricName":"Total length"},{"values":[7150,5681,14022,12790,12878,10575,10450,13021,12957,12903,12903,10624,2575],"quality":"More is better","isMain":true,"metricName":"Total length (>= 1000 bp)"},{"values":[0,0,7865,12790,12878,0,7533,13021,12957,12903,12903,0,0],"quality":"More is better","isMain":false,"metricName":"Total length (>= 5000 bp)"},{"values":[0,0,0,0,0,0,0,0,0,0,0,0,0],"quality":"More is better","isMain":true,"metricName":"Total length (>= 10000 bp)"},{"values":[0,0,0,0,0,0,0,0,0,0,0,0,0],"quality":"More is better","isMain":false,"metricName":"Total length (>= 25000 bp)"},{"values":[0,0,0,0,0,0,0,0,0,0,0,0,0],"quality":"More is better","isMain":true,"metricName":"Total length (>= 50000 bp)"},{"values":[1533,2087,7865,7708,7799,3378,7533,7865,7865,7865,7865,3774,2575],"quality":"More is better","isMain":false,"metricName":"N50"},{"values":[737,784,1980,5082,5079,1269,2917,5156,5092,5038,5038,1487,2575],"quality":"More is better","isMain":false,"metricName":"N75"},{"values":[3,2,1,1,1,2,1,1,1,1,1,2,1],"quality":"Less is better","isMain":false,"metricName":"L50"},{"values":[5,4,3,2,2,4,2,2,2,2,2,3,1],"quality":"Less is better","isMain":false,"metricName":"L75"},{"values":["48.06","47.74","48.07","48.18","48.26","47.57","48.15","48.22","48.20","48.16","48.17","48.29","46.41"],"quality":"Equal","isMain":false,"metricName":"GC (%)"}]],["Predicted genes",[]],["Similarity statistics",[{"values":[0,0,0,0,0,0,0,0,0,0,0,0,0],"quality":"Equal","isMain":false,"metricName":"# similar correct contigs"},{"values":[0,0,0,0,0,0,0,0,0,0,0,0,0],"quality":"Equal","isMain":false,"metricName":"# similar misassembled blocks"}]],["Reference statistics",[{"values":[14927,14927,14927,14927,14927,14927,14927,14927,14927,14927,14927,14927,14927],"quality":"Equal","isMain":false,"metricName":"Reference length"},{"values":[1,1,1,1,1,1,1,1,1,1,1,1,1],"quality":"Equal","isMain":false,"metricName":"Reference fragments"},{"values":["48.10","48.10","48.10","48.10","48.10","48.10","48.10","48.10","48.10","48.10","48.10","48.10","48.10"],"quality":"Equal","isMain":false,"metricName":"Reference GC (%)"}]]],"referenceName":"gi\_159192286\_ref\_NC","date":"18 July 2018, Wednesday, 16:29:14","order":[0,1,2,3,4,5,6,7,8,9,10,11,12],"assembliesNames":["ABySS\_127","ABySS\_63","CLC","IDBA\_UD","MEGAHIT","MIRA","SOAPdenovo2","SPAdes","SPAdes\_meta","SPAdes\_sc","SPAdes\_sc\_careful","Velvet","Geneious"]},{"assembliesWithNs":null,"minContig":500,"report":[["Genome statistics",[{"values":["95.230","100.000","94.549","94.865","100.000","96.735","91.754","100.000","100.000","100.000","100.000","94.585","1.991","96.735"],"quality":"More is better","isMain":true,"metricName":"Genome fraction (%)"},{"values":["1.003","1.003","1.001","1.006","1.011","1.025","1.000","1.005","1.002","1.002","1.002","1.002","1.116","1.000"],"quality":"Less is better","isMain":true,"metricName":"Duplication ratio"},{"values":[23479,24655,21968,22123,22207,21972,22622,24655,24651,24655,24655,21985,491,23850],"quality":"More is better","isMain":true,"metricName":"Largest alignment"},{"values":[23479,24655,23333,23488,24937,24436,22622,24655,24651,24655,24655,23350,491,23850],"quality":"More is better","isMain":true,"metricName":"Total aligned length"},{"values":[23542,24717,21968,22123,22207,21972,22622,24781,24705,24709,24709,21985,null,23850],"quality":"More is better","isMain":false,"metricName":"NG50"},{"values":[23542,24717,21968,22123,22207,21972,22622,24781,24705,24709,24709,21985,null,23850],"quality":"More is better","isMain":false,"metricName":"NG75"},{"values":[23479,24655,21968,22123,22207,21972,22622,24655,24651,24655,24655,21985,491,23850],"quality":"More is better","isMain":false,"metricName":"NA50"},{"values":[23479,24655,21968,22123,22207,21972,22622,24655,24651,24655,24655,21985,491,23850],"quality":"More is better","isMain":false,"metricName":"NA75"},{"values":[23479,24655,21968,22123,22207,21972,22622,24655,24651,24655,24655,21985,null,23850],"quality":"More is better","isMain":true,"metricName":"NGA50"},{"values":[23479,24655,21968,22123,22207,21972,22622,24655,24651,24655,24655,21985,null,23850],"quality":"More is better","isMain":false,"metricName":"NGA75"},{"values":[1,1,1,1,1,1,1,1,1,1,1,1,null,1],"quality":"Less is better","isMain":false,"metricName":"LG50"},{"values":[1,1,1,1,1,1,1,1,1,1,1,1,null,1],"quality":"Less is better","isMain":false,"metricName":"LG75"},{"values":[1,1,1,1,1,1,1,1,1,1,1,1,1,1],"quality":"Less is better","isMain":false,"metricName":"LA50"},{"values":[1,1,1,1,1,1,1,1,1,1,1,1,1,1],"quality":"Less is better","isMain":false,"metricName":"LA75"},{"values":[1,1,1,1,1,1,1,1,1,1,1,1,null,1],"quality":"Less is better","isMain":true,"metricName":"LGA50"},{"values":[1,1,1,1,1,1,1,1,1,1,1,1,null,1],"quality":"Less is better","isMain":false,"metricName":"LGA75"}]],["Misassemblies",[{"values":[0,0,0,0,0,0,0,0,0,0,0,0,0,0],"quality":"Less is better","isMain":true,"metricName":"# misassemblies"},{"values":[0,0,0,0,0,0,0,0,0,0,0,0,0,0],"quality":"Less is better","isMain":false,"metricName":" # relocations"},{"values":[0,0,0,0,0,0,0,0,0,0,0,0,0,0],"quality":"Less is better","isMain":false,"metricName":" # translocations"},{"values":[0,0,0,0,0,0,0,0,0,0,0,0,0,0],"quality":"Less is better","isMain":false,"metricName":" # inversions"},{"values":[0,0,0,0,0,0,0,0,0,0,0,0,0,0],"quality":"Less is better","isMain":false,"metricName":"# misassembled contigs"},{"values":[0,0,0,0,0,0,0,0,0,0,0,0,0,0],"quality":"Less is better","isMain":true,"metricName":"Misassembled contigs length"},{"values":[0,0,0,0,0,0,0,0,0,0,0,0,0,0],"quality":"Less is better","isMain":false,"metricName":"# local misassemblies"},{"values":[0,0,0,0,0,0,0,0,0,0,0,0,0,0],"quality":"Less is better","isMain":false,"metricName":"# unaligned mis. contigs"}]],["Unaligned",[{"values":[0,0,0,0,0,0,0,0,0,0,0,0,0,0],"quality":"Less is better","isMain":false,"metricName":"# fully unaligned contigs"},{"values":[0,0,0,0,0,0,0,0,0,0,0,0,0,0],"quality":"Less is better","isMain":false,"metricName":"Fully unaligned length"},{"values":[0,0,0,0,0,0,0,0,0,0,0,0,0,0],"quality":"Less is better","isMain":false,"metricName":"# partially unaligned contigs"},{"values":[0,0,0,0,0,0,0,0,0,0,0,0,0,0],"quality":"Less is better","isMain":false,"metricName":"Partially unaligned length"}]],["Mismatches",[{"values":[0,0,0,0,0,0,0,0,0,0,0,0,0,0],"quality":"Less is better","isMain":false,"metricName":"# mismatches"},{"values":[0,0,0,0,0,0,0,0,2,0,0,0,0,0],"quality":"Less is better","isMain":false,"metricName":"# indels"},{"values":[0,0,0,0,0,0,0,0,4,0,0,0,0,0],"quality":"Less is better","isMain":false,"metricName":"Indels length"},{"values":["0.00","0.00","0.00","0.00","0.00","0.00","0.00","0.00","0.00","0.00","0.00","0.00","0.00","0.00"],"quality":"Less is better","isMain":true,"metricName":"# mismatches per 100 kbp"},{"values":["0.00","0.00","0.00","0.00","0.00","0.00","0.00","0.00","8.11","0.00","0.00","0.00","0.00","0.00"],"quality":"Less is better","isMain":true,"metricName":"# indels per 100 kbp"},{"values":[0,0,0,0,0,0,0,0,2,0,0,0,0,0],"quality":"Less is better","isMain":false,"metricName":" # indels (<= 5 bp)"},{"values":[0,0,0,0,0,0,0,0,0,0,0,0,0,0],"quality":"Less is better","isMain":false,"metricName":" # indels (> 5 bp)"},{"values":[0,2,0,0,0,0,0,0,0,0,0,0,0,0],"quality":"Less is better","isMain":false,"metricName":"# N's"},{"values":["0.00","8.09","0.00","0.00","0.00","0.00","0.00","0.00","0.00","0.00","0.00","0.00","0.00","0.00"],"quality":"Less is better","isMain":true,"metricName":"# N's per 100 kbp"}]],["Statistics without reference",[{"values":[1,1,2,2,2,3,1,1,1,1,1,2,1,1],"quality":"Equal","isMain":true,"metricName":"# contigs"},{"values":[1,1,2,2,2,2,1,1,1,1,1,2,0,1],"quality":"Equal","isMain":false,"metricName":"# contigs (>= 1000 bp)"},{"values":[1,1,1,1,1,1,1,1,1,1,1,1,0,1],"quality":"Equal","isMain":false,"metricName":"# contigs (>= 5000 bp)"},{"values":[1,1,1,1,1,1,1,1,1,1,1,1,0,1],"quality":"Equal","isMain":false,"metricName":"# contigs (>= 10000 bp)"},{"values":[0,0,0,0,0,0,0,0,0,0,0,0,0,0],"quality":"Equal","isMain":false,"metricName":"# contigs (>= 25000 bp)"},{"values":[0,0,0,0,0,0,0,0,0,0,0,0,0,0],"quality":"Equal","isMain":false,"metricName":"# contigs (>= 50000 bp)"},{"values":[23542,24717,21968,22123,22207,21972,22622,24781,24705,24709,24709,21985,548,23850],"quality":"More is better","isMain":true,"metricName":"Largest contig"},{"values":[23542,24717,23344,23527,24937,24436,22622,24781,24705,24709,24709,23365,548,23850],"quality":"More is better","isMain":true,"metricName":"Total length"},{"values":[23542,24717,23344,23527,24937,23880,22622,24781,24705,24709,24709,23365,0,23850],"quality":"More is better","isMain":true,"metricName":"Total length (>= 1000 bp)"},{"values":[23542,24717,21968,22123,22207,21972,22622,24781,24705,24709,24709,21985,0,23850],"quality":"More is better","isMain":false,"metricName":"Total length (>= 5000 bp)"},{"values":[23542,24717,21968,22123,22207,21972,22622,24781,24705,24709,24709,21985,0,23850],"quality":"More is better","isMain":true,"metricName":"Total length (>= 10000 bp)"},{"values":[0,0,0,0,0,0,0,0,0,0,0,0,0,0],"quality":"More is better","isMain":false,"metricName":"Total length (>= 25000 bp)"},{"values":[0,0,0,0,0,0,0,0,0,0,0,0,0,0],"quality":"More is better","isMain":true,"metricName":"Total length (>= 50000 bp)"},{"values":[23542,24717,21968,22123,22207,21972,22622,24781,24705,24709,24709,21985,548,23850],"quality":"More is better","isMain":false,"metricName":"N50"},{"values":[23542,24717,21968,22123,22207,21972,22622,24781,24705,24709,24709,21985,548,23850],"quality":"More is better","isMain":false,"metricName":"N75"},{"values":[1,1,1,1,1,1,1,1,1,1,1,1,1,1],"quality":"Less is better","isMain":false,"metricName":"L50"},{"values":[1,1,1,1,1,1,1,1,1,1,1,1,1,1],"quality":"Less is better","isMain":false,"metricName":"L75"},{"values":["39.56","39.11","39.64","39.61","39.08","39.34","40.08","39.03","39.11","39.11","39.11","39.66","22.81","39.39"],"quality":"Equal","isMain":false,"metricName":"GC (%)"}]],["Predicted genes",[]],["Similarity statistics",[{"values":[1,1,0,0,1,0,1,1,1,1,1,0,0,1],"quality":"Equal","isMain":false,"metricName":"# similar correct contigs"},{"values":[0,0,0,0,0,0,0,0,0,0,0,0,0,0],"quality":"Equal","isMain":false,"metricName":"# similar misassembled blocks"}]],["Reference statistics",[{"values":[24655,24655,24655,24655,24655,24655,24655,24655,24655,24655,24655,24655,24655,24655],"quality":"Equal","isMain":false,"metricName":"Reference length"},{"values":[1,1,1,1,1,1,1,1,1,1,1,1,1,1],"quality":"Equal","isMain":false,"metricName":"Reference fragments"},{"values":["39.13","39.13","39.13","39.13","39.13","39.13","39.13","39.13","39.13","39.13","39.13","39.13","39.13","39.13"],"quality":"Equal","isMain":false,"metricName":"Reference GC (%)"}]]],"referenceName":"gi\_159341058\_ref\_NC","date":"18 July 2018, Wednesday, 16:29:27","order":[0,1,2,3,4,5,6,7,8,9,10,11,12,13],"assembliesNames":["ABySS\_127","ABySS\_63","CLC","IDBA\_UD","MEGAHIT","MIRA","Ray\_Meta","SPAdes","SPAdes\_meta","SPAdes\_sc","SPAdes\_sc\_careful","Velvet","VICUNA","Geneious"]},{"assembliesWithNs":null,"minContig":500,"report":[["Genome statistics",[{"values":["100.000","88.691","100.000","91.385","97.738","100.000","13.774","100.000","100.000","86.366","80.407","15.680","100.000"],"quality":"More is better","isMain":true,"metricName":"Genome fraction (%)"},{"values":["1.002","1.038","1.000","1.000","1.029","1.093","1.693","1.000","1.079","1.016","1.016","1.000","1.029"],"quality":"Less is better","isMain":true,"metricName":"Duplication ratio"},{"values":[6159,1851,7870,3024,3681,8097,150,5907,5834,2388,2278,618,8097],"quality":"More is better","isMain":true,"metricName":"Largest alignment"},{"values":[7888,7242,7870,7192,7911,8600,1084,7870,8039,6907,6431,1234,8097],"quality":"More is better","isMain":true,"metricName":"Total aligned length"},{"values":[6159,900,7870,1675,1759,8097,603,5907,5834,1755,1146,null,8097],"quality":"More is better","isMain":false,"metricName":"NG50"},{"values":[6159,682,7870,1627,1682,8097,null,5907,2660,1153,552,null,8097],"quality":"More is better","isMain":false,"metricName":"NG75"},{"values":[6159,989,7870,1675,1759,8097,null,5907,5834,1755,1611,618,8097],"quality":"More is better","isMain":false,"metricName":"NA50"},{"values":[6159,737,7870,1627,1681,8097,null,5907,2205,1611,1146,616,8097],"quality":"More is better","isMain":false,"metricName":"NA75"},{"values":[6159,900,7870,1675,1759,8097,null,5907,5834,1755,1146,null,8097],"quality":"More is better","isMain":true,"metricName":"NGA50"},{"values":[6159,682,7870,1627,1681,8097,null,5907,2205,1153,552,null,8097],"quality":"More is better","isMain":false,"metricName":"NGA75"},{"values":[1,4,1,2,2,1,4,1,1,2,3,null,1],"quality":"Less is better","isMain":false,"metricName":"LG50"},{"values":[1,6,1,3,3,1,null,1,2,4,5,null,1],"quality":"Less is better","isMain":false,"metricName":"LG75"},{"values":[1,3,1,2,2,1,null,1,1,2,2,1,1],"quality":"Less is better","isMain":false,"metricName":"LA50"},{"values":[1,5,1,3,3,1,null,1,2,3,3,2,1],"quality":"Less is better","isMain":false,"metricName":"LA75"},{"values":[1,4,1,2,2,1,null,1,1,2,3,null,1],"quality":"Less is better","isMain":true,"metricName":"LGA50"},{"values":[1,6,1,3,3,1,null,1,2,4,5,null,1],"quality":"Less is better","isMain":false,"metricName":"LGA75"}]],["Misassemblies",[{"values":[0,0,0,0,0,0,0,0,0,0,0,0,0],"quality":"Less is better","isMain":true,"metricName":"# misassemblies"},{"values":[0,0,0,0,0,0,0,0,0,0,0,0,0],"quality":"Less is better","isMain":false,"metricName":" # relocations"},{"values":[0,0,0,0,0,0,0,0,0,0,0,0,0],"quality":"Less is better","isMain":false,"metricName":" # translocations"},{"values":[0,0,0,0,0,0,0,0,0,0,0,0,0],"quality":"Less is better","isMain":false,"metricName":" # inversions"},{"values":[0,0,0,0,0,0,0,0,0,0,0,0,0],"quality":"Less is better","isMain":false,"metricName":"# misassembled contigs"},{"values":[0,0,0,0,0,0,0,0,0,0,0,0,0],"quality":"Less is better","isMain":true,"metricName":"Misassembled contigs length"},{"values":[0,0,0,0,0,1,0,0,1,0,0,0,1],"quality":"Less is better","isMain":false,"metricName":"# local misassemblies"},{"values":[0,0,0,0,0,0,3,0,0,0,0,0,0],"quality":"Less is better","isMain":false,"metricName":"# unaligned mis. contigs"}]],["Unaligned",[{"values":[0,0,0,0,0,0,0,0,0,0,0,0,0],"quality":"Less is better","isMain":false,"metricName":"# fully unaligned contigs"},{"values":[0,0,0,0,0,0,0,0,0,0,0,0,0],"quality":"Less is better","isMain":false,"metricName":"Fully unaligned length"},{"values":[0,0,0,0,0,0,2,0,0,0,0,0,0],"quality":"Less is better","isMain":false,"metricName":"# partially unaligned contigs"},{"values":[0,0,0,0,0,0,2112,0,0,0,0,0,0],"quality":"Less is better","isMain":false,"metricName":"Partially unaligned length"}]],["Mismatches",[{"values":[0,0,0,2,2,0,0,0,44,11,0,0,0],"quality":"Less is better","isMain":false,"metricName":"# mismatches"},{"values":[0,0,0,0,0,0,0,0,2,0,0,0,0],"quality":"Less is better","isMain":false,"metricName":"# indels"},{"values":[0,0,0,0,0,0,0,0,7,0,0,0,0],"quality":"Less is better","isMain":false,"metricName":"Indels length"},{"values":["0.00","0.00","0.00","27.81","26.00","0.00","0.00","0.00","559.09","161.84","0.00","0.00","0.00"],"quality":"Less is better","isMain":true,"metricName":"# mismatches per 100 kbp"},{"values":["0.00","0.00","0.00","0.00","0.00","0.00","0.00","0.00","25.41","0.00","0.00","0.00","0.00"],"quality":"Less is better","isMain":true,"metricName":"# indels per 100 kbp"},{"values":[0,0,0,0,0,0,0,0,1,0,0,0,0],"quality":"Less is better","isMain":false,"metricName":" # indels (<= 5 bp)"},{"values":[0,0,0,0,0,0,0,0,1,0,0,0,0],"quality":"Less is better","isMain":false,"metricName":" # indels (> 5 bp)"},{"values":[0,0,0,0,0,0,1916,0,0,0,0,0,0],"quality":"Less is better","isMain":false,"metricName":"# N's"},{"values":["0.00","0.00","0.00","0.00","0.00","0.00","48543.20","0.00","0.00","0.00","0.00","0.00","0.00"],"quality":"Less is better","isMain":true,"metricName":"# N's per 100 kbp"}]],["Statistics without reference",[{"values":[2,8,1,4,4,2,4,2,2,4,5,2,1],"quality":"Equal","isMain":true,"metricName":"# contigs"},{"values":[2,2,1,3,3,1,2,2,2,4,3,0,1],"quality":"Equal","isMain":false,"metricName":"# contigs (>= 1000 bp)"},{"values":[1,0,1,0,0,1,0,1,1,0,0,0,1],"quality":"Equal","isMain":false,"metricName":"# contigs (>= 5000 bp)"},{"values":[0,0,0,0,0,0,0,0,0,0,0,0,0],"quality":"Equal","isMain":false,"metricName":"# contigs (>= 10000 bp)"},{"values":[0,0,0,0,0,0,0,0,0,0,0,0,0],"quality":"Equal","isMain":false,"metricName":"# contigs (>= 25000 bp)"},{"values":[0,0,0,0,0,0,0,0,0,0,0,0,0],"quality":"Equal","isMain":false,"metricName":"# contigs (>= 50000 bp)"},{"values":[6159,1851,7870,3024,3681,8097,1663,5907,5834,2388,2278,618,8097],"quality":"More is better","isMain":true,"metricName":"Largest contig"},{"values":[7888,7242,7870,7192,7912,8600,3947,7870,8494,6907,6431,1234,8097],"quality":"More is better","isMain":true,"metricName":"Total length"},{"values":[7888,2870,7870,6326,7122,8097,2685,7870,8494,6907,5035,0,8097],"quality":"More is better","isMain":true,"metricName":"Total length (>= 1000 bp)"},{"values":[6159,0,7870,0,0,8097,0,5907,5834,0,0,0,8097],"quality":"More is better","isMain":false,"metricName":"Total length (>= 5000 bp)"},{"values":[0,0,0,0,0,0,0,0,0,0,0,0,0],"quality":"More is better","isMain":true,"metricName":"Total length (>= 10000 bp)"},{"values":[0,0,0,0,0,0,0,0,0,0,0,0,0],"quality":"More is better","isMain":false,"metricName":"Total length (>= 25000 bp)"},{"values":[0,0,0,0,0,0,0,0,0,0,0,0,0],"quality":"More is better","isMain":true,"metricName":"Total length (>= 50000 bp)"},{"values":[6159,989,7870,1675,1759,8097,1022,5907,5834,1755,1611,618,8097],"quality":"More is better","isMain":false,"metricName":"N50"},{"values":[6159,737,7870,1627,1682,8097,659,5907,2660,1611,1146,616,8097],"quality":"More is better","isMain":false,"metricName":"N75"},{"values":[1,3,1,2,2,1,2,1,1,2,2,1,1],"quality":"Less is better","isMain":false,"metricName":"L50"},{"values":[1,5,1,3,3,1,3,1,2,3,3,2,1],"quality":"Less is better","isMain":false,"metricName":"L75"},{"values":["22.19","22.25","22.21","22.15","21.84","22.14","23.58","22.21","21.91","22.66","22.61","25.12","22.05"],"quality":"Equal","isMain":false,"metricName":"GC (%)"}]],["Predicted genes",[]],["Similarity statistics",[{"values":[0,0,0,0,0,0,0,0,0,0,0,0,0],"quality":"Equal","isMain":false,"metricName":"# similar correct contigs"},{"values":[0,0,0,0,0,0,0,0,0,0,0,0,0],"quality":"Equal","isMain":false,"metricName":"# similar misassembled blocks"}]],["Reference statistics",[{"values":[7870,7870,7870,7870,7870,7870,7870,7870,7870,7870,7870,7870,7870],"quality":"Equal","isMain":false,"metricName":"Reference length"},{"values":[1,1,1,1,1,1,1,1,1,1,1,1,1],"quality":"Equal","isMain":false,"metricName":"Reference fragments"},{"values":["22.21","22.21","22.21","22.21","22.21","22.21","22.21","22.21","22.21","22.21","22.21","22.21","22.21"],"quality":"Equal","isMain":false,"metricName":"Reference GC (%)"}]]],"referenceName":"gi\_160688402\_ref\_NC","date":"18 July 2018, Wednesday, 16:29:39","order":[0,1,2,3,4,5,6,7,8,9,10,11,12],"assembliesNames":["ABySS\_127","ABySS\_63","CLC","IDBA\_UD","MEGAHIT","MIRA","SOAPdenovo2","SPAdes","SPAdes\_meta","SPAdes\_sc","SPAdes\_sc\_careful","Velvet","Geneious"]},{"assembliesWithNs":null,"minContig":500,"report":[["Genome statistics",[{"values":["99.535","99.348","100.000","100.000","99.853","74.856","59.897","100.000","100.000","99.960","99.960","98.892"],"quality":"More is better","isMain":true,"metricName":"Genome fraction (%)"},{"values":["1.000","1.000","1.002","1.003","1.006","1.012","1.036","1.003","1.000","1.000","1.000","1.001"],"quality":"Less is better","isMain":true,"metricName":"Duplication ratio"},{"values":[42769,42689,43068,43110,43156,3597,4926,43096,42987,42952,42952,32889],"quality":"More is better","isMain":true,"metricName":"Largest alignment"},{"values":[42769,42689,43068,43110,43156,32540,25981,43096,42987,42952,42952,42549],"quality":"More is better","isMain":true,"metricName":"Total aligned length"},{"values":[42769,42689,43068,43110,43156,1233,1452,43096,42987,42952,42952,32889],"quality":"More is better","isMain":false,"metricName":"NG50"},{"values":[42769,42689,43068,43110,43156,538,null,43096,42987,42952,42952,32889],"quality":"More is better","isMain":false,"metricName":"NG75"},{"values":[42769,42689,43068,43110,43156,1509,1563,43096,42987,42952,42952,32889],"quality":"More is better","isMain":false,"metricName":"NA50"},{"values":[42769,42689,43068,43110,43156,818,551,43096,42987,42952,42952,32889],"quality":"More is better","isMain":false,"metricName":"NA75"},{"values":[42769,42689,43068,43110,43156,1233,633,43096,42987,42952,42952,32889],"quality":"More is better","isMain":true,"metricName":"NGA50"},{"values":[42769,42689,43068,43110,43156,538,null,43096,42987,42952,42952,32889],"quality":"More is better","isMain":false,"metricName":"NGA75"},{"values":[1,1,1,1,1,12,6,1,1,1,1,1],"quality":"Less is better","isMain":false,"metricName":"LG50"},{"values":[1,1,1,1,1,27,null,1,1,1,1,1],"quality":"Less is better","isMain":false,"metricName":"LG75"},{"values":[1,1,1,1,1,9,5,1,1,1,1,1],"quality":"Less is better","isMain":false,"metricName":"LA50"},{"values":[1,1,1,1,1,15,14,1,1,1,1,1],"quality":"Less is better","isMain":false,"metricName":"LA75"},{"values":[1,1,1,1,1,12,11,1,1,1,1,1],"quality":"Less is better","isMain":true,"metricName":"LGA50"},{"values":[1,1,1,1,1,27,null,1,1,1,1,1],"quality":"Less is better","isMain":false,"metricName":"LGA75"}]],["Misassemblies",[{"values":[0,0,0,0,0,0,1,0,0,0,0,0],"quality":"Less is better","isMain":true,"metricName":"# misassemblies"},{"values":[0,0,0,0,0,0,1,0,0,0,0,0],"quality":"Less is better","isMain":false,"metricName":" # relocations"},{"values":[0,0,0,0,0,0,0,0,0,0,0,0],"quality":"Less is better","isMain":false,"metricName":" # translocations"},{"values":[0,0,0,0,0,0,0,0,0,0,0,0],"quality":"Less is better","isMain":false,"metricName":" # inversions"},{"values":[0,0,0,0,0,0,1,0,0,0,0,0],"quality":"Less is better","isMain":false,"metricName":"# misassembled contigs"},{"values":[0,0,0,0,0,0,10424,0,0,0,0,0],"quality":"Less is better","isMain":true,"metricName":"Misassembled contigs length"},{"values":[0,0,1,1,0,0,0,1,0,0,0,0],"quality":"Less is better","isMain":false,"metricName":"# local misassemblies"},{"values":[0,0,0,0,0,0,0,0,0,0,0,0],"quality":"Less is better","isMain":false,"metricName":"# unaligned mis. contigs"}]],["Unaligned",[{"values":[0,0,0,0,0,0,0,0,0,0,0,0],"quality":"Less is better","isMain":false,"metricName":"# fully unaligned contigs"},{"values":[0,0,0,0,0,0,0,0,0,0,0,0],"quality":"Less is better","isMain":false,"metricName":"Fully unaligned length"},{"values":[0,0,0,0,0,0,2,0,0,0,0,0],"quality":"Less is better","isMain":false,"metricName":"# partially unaligned contigs"},{"values":[0,0,0,0,0,0,4225,0,0,0,0,0],"quality":"Less is better","isMain":false,"metricName":"Partially unaligned length"}]],["Mismatches",[{"values":[0,0,0,0,2,0,0,0,0,0,0,0],"quality":"Less is better","isMain":false,"metricName":"# mismatches"},{"values":[0,0,0,0,1,0,8,0,1,0,0,0],"quality":"Less is better","isMain":false,"metricName":"# indels"},{"values":[0,0,0,0,63,0,385,0,18,0,0,0],"quality":"Less is better","isMain":false,"metricName":"Indels length"},{"values":["0.00","0.00","0.00","0.00","4.66","0.00","0.00","0.00","0.00","0.00","0.00","0.00"],"quality":"Less is better","isMain":true,"metricName":"# mismatches per 100 kbp"},{"values":["0.00","0.00","0.00","0.00","2.33","0.00","31.08","0.00","2.33","0.00","0.00","0.00"],"quality":"Less is better","isMain":true,"metricName":"# indels per 100 kbp"},{"values":[0,0,0,0,0,0,0,0,0,0,0,0],"quality":"Less is better","isMain":false,"metricName":" # indels (<= 5 bp)"},{"values":[0,0,0,0,1,0,8,0,1,0,0,0],"quality":"Less is better","isMain":false,"metricName":" # indels (> 5 bp)"},{"values":[0,0,0,0,0,0,1527,0,0,0,0,0],"quality":"Less is better","isMain":false,"metricName":"# N's"},{"values":["0.00","0.00","0.00","0.00","0.00","0.00","4944.79","0.00","0.00","0.00","0.00","0.00"],"quality":"Less is better","isMain":true,"metricName":"# N's per 100 kbp"}]],["Statistics without reference",[{"values":[1,1,1,1,1,27,18,1,1,1,1,4],"quality":"Equal","isMain":true,"metricName":"# contigs"},{"values":[1,1,1,1,1,14,7,1,1,1,1,4],"quality":"Equal","isMain":false,"metricName":"# contigs (>= 1000 bp)"},{"values":[1,1,1,1,1,0,2,1,1,1,1,2],"quality":"Equal","isMain":false,"metricName":"# contigs (>= 5000 bp)"},{"values":[1,1,1,1,1,0,1,1,1,1,1,1],"quality":"Equal","isMain":false,"metricName":"# contigs (>= 10000 bp)"},{"values":[1,1,1,1,1,0,0,1,1,1,1,1],"quality":"Equal","isMain":false,"metricName":"# contigs (>= 25000 bp)"},{"values":[0,0,0,0,0,0,0,0,0,0,0,0],"quality":"Equal","isMain":false,"metricName":"# contigs (>= 50000 bp)"},{"values":[42769,42689,43068,43110,43156,3597,10424,43096,42987,42952,42952,32889],"quality":"More is better","isMain":true,"metricName":"Largest contig"},{"values":[42769,42689,43068,43110,43156,32540,30881,43096,42987,42952,42952,42549],"quality":"More is better","isMain":true,"metricName":"Total length"},{"values":[42769,42689,43068,43110,43156,23647,23886,43096,42987,42952,42952,42549],"quality":"More is better","isMain":true,"metricName":"Total length (>= 1000 bp)"},{"values":[42769,42689,43068,43110,43156,0,15522,43096,42987,42952,42952,38660],"quality":"More is better","isMain":false,"metricName":"Total length (>= 5000 bp)"},{"values":[42769,42689,43068,43110,43156,0,10424,43096,42987,42952,42952,32889],"quality":"More is better","isMain":true,"metricName":"Total length (>= 10000 bp)"},{"values":[42769,42689,43068,43110,43156,0,0,43096,42987,42952,42952,32889],"quality":"More is better","isMain":false,"metricName":"Total length (>= 25000 bp)"},{"values":[0,0,0,0,0,0,0,0,0,0,0,0],"quality":"More is better","isMain":true,"metricName":"Total length (>= 50000 bp)"},{"values":[42769,42689,43068,43110,43156,1509,5098,43096,42987,42952,42952,32889],"quality":"More is better","isMain":false,"metricName":"N50"},{"values":[42769,42689,43068,43110,43156,818,1288,43096,42987,42952,42952,32889],"quality":"More is better","isMain":false,"metricName":"N75"},{"values":[1,1,1,1,1,9,2,1,1,1,1,1],"quality":"Less is better","isMain":false,"metricName":"L50"},{"values":[1,1,1,1,1,15,7,1,1,1,1,1],"quality":"Less is better","isMain":false,"metricName":"L75"},{"values":["56.16","56.16","56.17","56.15","56.15","56.35","56.13","56.16","56.16","56.16","56.16","56.11"],"quality":"Equal","isMain":false,"metricName":"GC (%)"}]],["Predicted genes",[]],["Similarity statistics",[{"values":[1,1,0,0,1,0,0,0,1,1,1,1],"quality":"Equal","isMain":false,"metricName":"# similar correct contigs"},{"values":[0,0,0,0,0,0,0,0,0,0,0,0],"quality":"Equal","isMain":false,"metricName":"# similar misassembled blocks"}]],["Reference statistics",[{"values":[42969,42969,42969,42969,42969,42969,42969,42969,42969,42969,42969,42969],"quality":"Equal","isMain":false,"metricName":"Reference length"},{"values":[1,1,1,1,1,1,1,1,1,1,1,1],"quality":"Equal","isMain":false,"metricName":"Reference fragments"},{"values":["56.16","56.16","56.16","56.16","56.16","56.16","56.16","56.16","56.16","56.16","56.16","56.16"],"quality":"Equal","isMain":false,"metricName":"Reference GC (%)"}]]],"referenceName":"gi\_160693214\_ref\_NC","date":"18 July 2018, Wednesday, 16:29:50","order":[0,1,2,3,4,5,6,7,8,9,10,11],"assembliesNames":["ABySS\_63","CLC","IDBA\_UD","MEGAHIT","MIRA","Ray\_Meta","SOAPdenovo2","SPAdes","SPAdes\_meta","SPAdes\_sc","SPAdes\_sc\_careful","Velvet"]},{"assembliesWithNs":null,"minContig":500,"report":[["Genome statistics",[{"values":["99.997","99.397","100.000","100.000","100.000","100.000","6.748","98.349","100.000","100.000","100.000","100.000","99.914"],"quality":"More is better","isMain":true,"metricName":"Genome fraction (%)"},{"values":["1.000","1.007","1.000","1.003","1.004","1.006","1.000","1.028","1.003","1.001","1.001","1.001","1.000"],"quality":"Less is better","isMain":true,"metricName":"Duplication ratio"},{"values":[37312,37088,37313,37412,37454,37540,709,36837,37440,37368,37368,37368,37281],"quality":"More is better","isMain":true,"metricName":"Largest alignment"},{"values":[37312,37088,37313,37412,37454,37540,2518,36837,37440,37368,37368,37368,37281],"quality":"More is better","isMain":true,"metricName":"Total aligned length"},{"values":[37312,37363,37313,37412,37454,37540,null,37711,37440,37368,37368,37368,37281],"quality":"More is better","isMain":false,"metricName":"NG50"},{"values":[37312,37363,37313,37412,37454,37540,null,37711,37440,37368,37368,37368,37281],"quality":"More is better","isMain":false,"metricName":"NG75"},{"values":[37312,37088,37313,37412,37454,37540,684,36837,37440,37368,37368,37368,37281],"quality":"More is better","isMain":false,"metricName":"NA50"},{"values":[37312,37088,37313,37412,37454,37540,580,36837,37440,37368,37368,37368,37281],"quality":"More is better","isMain":false,"metricName":"NA75"},{"values":[37312,37088,37313,37412,37454,37540,null,36837,37440,37368,37368,37368,37281],"quality":"More is better","isMain":true,"metricName":"NGA50"},{"values":[37312,37088,37313,37412,37454,37540,null,36837,37440,37368,37368,37368,37281],"quality":"More is better","isMain":false,"metricName":"NGA75"},{"values":[1,1,1,1,1,1,null,1,1,1,1,1,1],"quality":"Less is better","isMain":false,"metricName":"LG50"},{"values":[1,1,1,1,1,1,null,1,1,1,1,1,1],"quality":"Less is better","isMain":false,"metricName":"LG75"},{"values":[1,1,1,1,1,1,2,1,1,1,1,1,1],"quality":"Less is better","isMain":false,"metricName":"LA50"},{"values":[1,1,1,1,1,1,3,1,1,1,1,1,1],"quality":"Less is better","isMain":false,"metricName":"LA75"},{"values":[1,1,1,1,1,1,null,1,1,1,1,1,1],"quality":"Less is better","isMain":true,"metricName":"LGA50"},{"values":[1,1,1,1,1,1,null,1,1,1,1,1,1],"quality":"Less is better","isMain":false,"metricName":"LGA75"}]],["Misassemblies",[{"values":[0,0,0,0,0,0,0,0,0,0,0,0,0],"quality":"Less is better","isMain":true,"metricName":"# misassemblies"},{"values":[0,0,0,0,0,0,0,0,0,0,0,0,0],"quality":"Less is better","isMain":false,"metricName":" # relocations"},{"values":[0,0,0,0,0,0,0,0,0,0,0,0,0],"quality":"Less is better","isMain":false,"metricName":" # translocations"},{"values":[0,0,0,0,0,0,0,0,0,0,0,0,0],"quality":"Less is better","isMain":false,"metricName":" # inversions"},{"values":[0,0,0,0,0,0,0,0,0,0,0,0,0],"quality":"Less is better","isMain":false,"metricName":"# misassembled contigs"},{"values":[0,0,0,0,0,0,0,0,0,0,0,0,0],"quality":"Less is better","isMain":true,"metricName":"Misassembled contigs length"},{"values":[0,0,0,1,1,1,0,0,1,0,0,0,0],"quality":"Less is better","isMain":false,"metricName":"# local misassemblies"},{"values":[0,0,0,0,0,0,0,0,0,0,0,0,0],"quality":"Less is better","isMain":false,"metricName":"# unaligned mis. contigs"}]],["Unaligned",[{"values":[0,0,0,0,0,0,0,0,0,0,0,0,0],"quality":"Less is better","isMain":false,"metricName":"# fully unaligned contigs"},{"values":[0,0,0,0,0,0,0,0,0,0,0,0,0],"quality":"Less is better","isMain":false,"metricName":"Fully unaligned length"},{"values":[0,0,0,0,0,0,0,0,0,0,0,0,0],"quality":"Less is better","isMain":false,"metricName":"# partially unaligned contigs"},{"values":[0,0,0,0,0,0,0,0,0,0,0,0,0],"quality":"Less is better","isMain":false,"metricName":"Partially unaligned length"}]],["Mismatches",[{"values":[1,0,0,0,0,0,0,0,0,0,0,0,0],"quality":"Less is better","isMain":false,"metricName":"# mismatches"},{"values":[0,0,0,0,0,0,0,10,0,1,1,1,0],"quality":"Less is better","isMain":false,"metricName":"# indels"},{"values":[0,0,0,0,0,0,0,491,0,55,55,55,0],"quality":"Less is better","isMain":false,"metricName":"Indels length"},{"values":["2.68","0.00","0.00","0.00","0.00","0.00","0.00","0.00","0.00","0.00","0.00","0.00","0.00"],"quality":"Less is better","isMain":true,"metricName":"# mismatches per 100 kbp"},{"values":["0.00","0.00","0.00","0.00","0.00","0.00","0.00","27.25","0.00","2.68","2.68","2.68","0.00"],"quality":"Less is better","isMain":true,"metricName":"# indels per 100 kbp"},{"values":[0,0,0,0,0,0,0,0,0,0,0,0,0],"quality":"Less is better","isMain":false,"metricName":" # indels (<= 5 bp)"},{"values":[0,0,0,0,0,0,0,10,0,1,1,1,0],"quality":"Less is better","isMain":false,"metricName":" # indels (> 5 bp)"},{"values":[0,0,0,0,0,3,0,1041,0,0,0,0,0],"quality":"Less is better","isMain":false,"metricName":"# N's"},{"values":["0.00","0.00","0.00","0.00","0.00","7.99","0.00","2760.47","0.00","0.00","0.00","0.00","0.00"],"quality":"Less is better","isMain":true,"metricName":"# N's per 100 kbp"}]],["Statistics without reference",[{"values":[1,1,1,1,1,1,4,1,1,1,1,1,1],"quality":"Equal","isMain":true,"metricName":"# contigs"},{"values":[1,1,1,1,1,1,0,1,1,1,1,1,1],"quality":"Equal","isMain":false,"metricName":"# contigs (>= 1000 bp)"},{"values":[1,1,1,1,1,1,0,1,1,1,1,1,1],"quality":"Equal","isMain":false,"metricName":"# contigs (>= 5000 bp)"},{"values":[1,1,1,1,1,1,0,1,1,1,1,1,1],"quality":"Equal","isMain":false,"metricName":"# contigs (>= 10000 bp)"},{"values":[1,1,1,1,1,1,0,1,1,1,1,1,1],"quality":"Equal","isMain":false,"metricName":"# contigs (>= 25000 bp)"},{"values":[0,0,0,0,0,0,0,0,0,0,0,0,0],"quality":"Equal","isMain":false,"metricName":"# contigs (>= 50000 bp)"},{"values":[37312,37363,37313,37412,37454,37540,709,37711,37440,37368,37368,37368,37281],"quality":"More is better","isMain":true,"metricName":"Largest contig"},{"values":[37312,37363,37313,37412,37454,37540,2518,37711,37440,37368,37368,37368,37281],"quality":"More is better","isMain":true,"metricName":"Total length"},{"values":[37312,37363,37313,37412,37454,37540,0,37711,37440,37368,37368,37368,37281],"quality":"More is better","isMain":true,"metricName":"Total length (>= 1000 bp)"},{"values":[37312,37363,37313,37412,37454,37540,0,37711,37440,37368,37368,37368,37281],"quality":"More is better","isMain":false,"metricName":"Total length (>= 5000 bp)"},{"values":[37312,37363,37313,37412,37454,37540,0,37711,37440,37368,37368,37368,37281],"quality":"More is better","isMain":true,"metricName":"Total length (>= 10000 bp)"},{"values":[37312,37363,37313,37412,37454,37540,0,37711,37440,37368,37368,37368,37281],"quality":"More is better","isMain":false,"metricName":"Total length (>= 25000 bp)"},{"values":[0,0,0,0,0,0,0,0,0,0,0,0,0],"quality":"More is better","isMain":true,"metricName":"Total length (>= 50000 bp)"},{"values":[37312,37363,37313,37412,37454,37540,684,37711,37440,37368,37368,37368,37281],"quality":"More is better","isMain":false,"metricName":"N50"},{"values":[37312,37363,37313,37412,37454,37540,580,37711,37440,37368,37368,37368,37281],"quality":"More is better","isMain":false,"metricName":"N75"},{"values":[1,1,1,1,1,1,2,1,1,1,1,1,1],"quality":"Less is better","isMain":false,"metricName":"L50"},{"values":[1,1,1,1,1,1,3,1,1,1,1,1,1],"quality":"Less is better","isMain":false,"metricName":"L75"},{"values":["40.88","40.89","40.89","40.89","40.89","40.90","37.97","40.88","40.91","40.91","40.89","40.89","40.89"],"quality":"Equal","isMain":false,"metricName":"GC (%)"}]],["Predicted genes",[]],["Similarity statistics",[{"values":[0,0,0,0,0,0,0,0,0,0,0,0,0],"quality":"Equal","isMain":false,"metricName":"# similar correct contigs"},{"values":[0,0,0,0,0,0,0,0,0,0,0,0,0],"quality":"Equal","isMain":false,"metricName":"# similar misassembled blocks"}]],["Reference statistics",[{"values":[37313,37313,37313,37313,37313,37313,37313,37313,37313,37313,37313,37313,37313],"quality":"Equal","isMain":false,"metricName":"Reference length"},{"values":[1,1,1,1,1,1,1,1,1,1,1,1,1],"quality":"Equal","isMain":false,"metricName":"Reference fragments"},{"values":["40.89","40.89","40.89","40.89","40.89","40.89","40.89","40.89","40.89","40.89","40.89","40.89","40.89"],"quality":"Equal","isMain":false,"metricName":"Reference GC (%)"}]]],"referenceName":"gi\_160700594\_ref\_NC","date":"18 July 2018, Wednesday, 16:30:02","order":[0,1,2,3,4,5,6,7,8,9,10,11,12],"assembliesNames":["ABySS\_127","ABySS\_63","CLC","IDBA\_UD","MEGAHIT","MIRA","Ray\_Meta","SOAPdenovo2","SPAdes","SPAdes\_meta","SPAdes\_sc","SPAdes\_sc\_careful","Velvet"]},{"assembliesWithNs":null,"minContig":500,"report":[["Genome statistics",[{"values":["99.958","100.000","100.000","100.000","94.205","75.896","100.000","100.000","100.000","100.000","100.000"],"quality":"More is better","isMain":true,"metricName":"Genome fraction (%)"},{"values":["1.000","1.002","1.003","1.007","1.011","1.001","1.002","1.001","1.001","1.001","1.001"],"quality":"Less is better","isMain":true,"metricName":"Duplication ratio"},{"values":[52953,53074,53116,53369,19356,3411,53102,53030,53030,53030,53005],"quality":"More is better","isMain":true,"metricName":"Largest alignment"},{"values":[52953,53074,53116,53369,50451,40252,53102,53030,53030,53030,53005],"quality":"More is better","isMain":true,"metricName":"Total aligned length"},{"values":[52975,53074,53116,53369,10292,895,53102,53030,53030,53030,53005],"quality":"More is better","isMain":false,"metricName":"NG50"},{"values":[52975,53074,53116,53369,3124,524,53102,53030,53030,53030,53005],"quality":"More is better","isMain":false,"metricName":"NG75"},{"values":[52953,53074,53116,53369,10292,1237,53102,53030,53030,53030,53005],"quality":"More is better","isMain":false,"metricName":"NA50"},{"values":[52953,53074,53116,53369,8242,823,53102,53030,53030,53030,53005],"quality":"More is better","isMain":false,"metricName":"NA75"},{"values":[52953,53074,53116,53369,10292,895,53102,53030,53030,53030,53005],"quality":"More is better","isMain":true,"metricName":"NGA50"},{"values":[52953,53074,53116,53369,3124,524,53102,53030,53030,53030,53005],"quality":"More is better","isMain":false,"metricName":"NGA75"},{"values":[1,1,1,1,2,18,1,1,1,1,1],"quality":"Less is better","isMain":false,"metricName":"LG50"},{"values":[1,1,1,1,4,37,1,1,1,1,1],"quality":"Less is better","isMain":false,"metricName":"LG75"},{"values":[1,1,1,1,2,12,1,1,1,1,1],"quality":"Less is better","isMain":false,"metricName":"LA50"},{"values":[1,1,1,1,3,22,1,1,1,1,1],"quality":"Less is better","isMain":false,"metricName":"LA75"},{"values":[1,1,1,1,2,18,1,1,1,1,1],"quality":"Less is better","isMain":true,"metricName":"LGA50"},{"values":[1,1,1,1,4,37,1,1,1,1,1],"quality":"Less is better","isMain":false,"metricName":"LGA75"}]],["Misassemblies",[{"values":[0,0,0,0,0,0,0,0,0,0,0],"quality":"Less is better","isMain":true,"metricName":"# misassemblies"},{"values":[0,0,0,0,0,0,0,0,0,0,0],"quality":"Less is better","isMain":false,"metricName":" # relocations"},{"values":[0,0,0,0,0,0,0,0,0,0,0],"quality":"Less is better","isMain":false,"metricName":" # translocations"},{"values":[0,0,0,0,0,0,0,0,0,0,0],"quality":"Less is better","isMain":false,"metricName":" # inversions"},{"values":[0,0,0,0,0,0,0,0,0,0,0],"quality":"Less is better","isMain":false,"metricName":"# misassembled contigs"},{"values":[0,0,0,0,0,0,0,0,0,0,0],"quality":"Less is better","isMain":true,"metricName":"Misassembled contigs length"},{"values":[0,1,1,1,0,0,1,0,0,0,0],"quality":"Less is better","isMain":false,"metricName":"# local misassemblies"},{"values":[0,0,0,0,0,0,0,0,0,0,0],"quality":"Less is better","isMain":false,"metricName":"# unaligned mis. contigs"}]],["Unaligned",[{"values":[0,0,0,0,0,0,0,0,0,0,0],"quality":"Less is better","isMain":false,"metricName":"# fully unaligned contigs"},{"values":[0,0,0,0,0,0,0,0,0,0,0],"quality":"Less is better","isMain":false,"metricName":"Fully unaligned length"},{"values":[0,0,0,0,0,0,0,0,0,0,0],"quality":"Less is better","isMain":false,"metricName":"# partially unaligned contigs"},{"values":[0,0,0,0,0,0,0,0,0,0,0],"quality":"Less is better","isMain":false,"metricName":"Partially unaligned length"}]],["Mismatches",[{"values":[0,0,0,0,0,0,0,0,0,0,0],"quality":"Less is better","isMain":false,"metricName":"# mismatches"},{"values":[0,0,0,0,0,0,0,1,1,1,1],"quality":"Less is better","isMain":false,"metricName":"# indels"},{"values":[0,0,0,0,0,0,0,55,55,55,30],"quality":"Less is better","isMain":false,"metricName":"Indels length"},{"values":["0.00","0.00","0.00","0.00","0.00","0.00","0.00","0.00","0.00","0.00","0.00"],"quality":"Less is better","isMain":true,"metricName":"# mismatches per 100 kbp"},{"values":["0.00","0.00","0.00","0.00","0.00","0.00","0.00","1.89","1.89","1.89","1.89"],"quality":"Less is better","isMain":true,"metricName":"# indels per 100 kbp"},{"values":[0,0,0,0,0,0,0,0,0,0,0],"quality":"Less is better","isMain":false,"metricName":" # indels (<= 5 bp)"},{"values":[0,0,0,0,0,0,0,1,1,1,1],"quality":"Less is better","isMain":false,"metricName":" # indels (> 5 bp)"},{"values":[0,0,0,0,0,0,0,0,0,0,0],"quality":"Less is better","isMain":false,"metricName":"# N's"},{"values":["0.00","0.00","0.00","0.00","0.00","0.00","0.00","0.00","0.00","0.00","0.00"],"quality":"Less is better","isMain":true,"metricName":"# N's per 100 kbp"}]],["Statistics without reference",[{"values":[1,1,1,1,11,37,1,1,1,1,1],"quality":"Equal","isMain":true,"metricName":"# contigs"},{"values":[1,1,1,1,9,17,1,1,1,1,1],"quality":"Equal","isMain":false,"metricName":"# contigs (>= 1000 bp)"},{"values":[1,1,1,1,3,0,1,1,1,1,1],"quality":"Equal","isMain":false,"metricName":"# contigs (>= 5000 bp)"},{"values":[1,1,1,1,2,0,1,1,1,1,1],"quality":"Equal","isMain":false,"metricName":"# contigs (>= 10000 bp)"},{"values":[1,1,1,1,0,0,1,1,1,1,1],"quality":"Equal","isMain":false,"metricName":"# contigs (>= 25000 bp)"},{"values":[1,1,1,1,0,0,1,1,1,1,1],"quality":"Equal","isMain":false,"metricName":"# contigs (>= 50000 bp)"},{"values":[52975,53074,53116,53369,19356,3411,53102,53030,53030,53030,53005],"quality":"More is better","isMain":true,"metricName":"Largest contig"},{"values":[52975,53074,53116,53369,50451,40252,53102,53030,53030,53030,53005],"quality":"More is better","isMain":true,"metricName":"Total length"},{"values":[52975,53074,53116,53369,48873,26323,53102,53030,53030,53030,53005],"quality":"More is better","isMain":true,"metricName":"Total length (>= 1000 bp)"},{"values":[52975,53074,53116,53369,37890,0,53102,53030,53030,53030,53005],"quality":"More is better","isMain":false,"metricName":"Total length (>= 5000 bp)"},{"values":[52975,53074,53116,53369,29648,0,53102,53030,53030,53030,53005],"quality":"More is better","isMain":true,"metricName":"Total length (>= 10000 bp)"},{"values":[52975,53074,53116,53369,0,0,53102,53030,53030,53030,53005],"quality":"More is better","isMain":false,"metricName":"Total length (>= 25000 bp)"},{"values":[52975,53074,53116,53369,0,0,53102,53030,53030,53030,53005],"quality":"More is better","isMain":true,"metricName":"Total length (>= 50000 bp)"},{"values":[52975,53074,53116,53369,10292,1237,53102,53030,53030,53030,53005],"quality":"More is better","isMain":false,"metricName":"N50"},{"values":[52975,53074,53116,53369,8242,823,53102,53030,53030,53030,53005],"quality":"More is better","isMain":false,"metricName":"N75"},{"values":[1,1,1,1,2,12,1,1,1,1,1],"quality":"Less is better","isMain":false,"metricName":"L50"},{"values":[1,1,1,1,3,22,1,1,1,1,1],"quality":"Less is better","isMain":false,"metricName":"L75"},{"values":["44.02","44.00","44.01","44.04","43.98","43.95","44.02","44.03","44.03","44.03","44.02"],"quality":"Equal","isMain":false,"metricName":"GC (%)"}]],["Predicted genes",[]],["Similarity statistics",[{"values":[0,0,0,0,0,0,0,0,0,0,0],"quality":"Equal","isMain":false,"metricName":"# similar correct contigs"},{"values":[0,0,0,0,0,0,0,0,0,0,0],"quality":"Equal","isMain":false,"metricName":"# similar misassembled blocks"}]],["Reference statistics",[{"values":[52975,52975,52975,52975,52975,52975,52975,52975,52975,52975,52975],"quality":"Equal","isMain":false,"metricName":"Reference length"},{"values":[1,1,1,1,1,1,1,1,1,1,1],"quality":"Equal","isMain":false,"metricName":"Reference fragments"},{"values":["44.02","44.02","44.02","44.02","44.02","44.02","44.02","44.02","44.02","44.02","44.02"],"quality":"Equal","isMain":false,"metricName":"Reference GC (%)"}]]],"referenceName":"gi\_161617915\_ref\_NC","date":"18 July 2018, Wednesday, 16:30:13","order":[0,1,2,3,4,5,6,7,8,9,10],"assembliesNames":["CLC","IDBA\_UD","MEGAHIT","MIRA","Ray\_Meta","SOAPdenovo2","SPAdes","SPAdes\_meta","SPAdes\_sc","SPAdes\_sc\_careful","Velvet"]},{"assembliesWithNs":null,"minContig":500,"report":[["Genome statistics",[{"values":["96.577","94.627","100.000","95.636","92.919","100.000","49.755","19.772","100.000","100.000","99.412","91.368","78.566","100.000"],"quality":"More is better","isMain":true,"metricName":"Genome fraction (%)"},{"values":["1.041","1.022","1.003","1.023","1.032","1.026","1.031","1.226","1.009","1.003","1.015","1.021","1.004","1.026"],"quality":"Less is better","isMain":true,"metricName":"Duplication ratio"},{"values":[9673,7890,22282,12538,12470,39711,1933,1745,45873,29720,16542,5116,3360,39714],"quality":"More is better","isMain":true,"metricName":"Largest alignment"},{"values":[58983,56751,58839,57390,56227,60204,30098,11645,59171,58828,59143,54698,46269,60211],"quality":"More is better","isMain":true,"metricName":"Total aligned length"},{"values":[5562,3053,14140,6618,5968,60204,510,null,45873,29720,8149,3107,1253,60211],"quality":"More is better","isMain":false,"metricName":"NG50"},{"values":[2092,1469,11700,3973,3293,60204,null,null,45873,26170,4268,1094,555,60211],"quality":"More is better","isMain":false,"metricName":"NG75"},{"values":[5562,3053,14140,6618,7237,39711,813,null,45873,29720,8102,3107,1906,39714],"quality":"More is better","isMain":false,"metricName":"NA50"},{"values":[2092,2023,11700,4338,3293,20493,671,null,45873,26170,3749,1217,1041,20497],"quality":"More is better","isMain":false,"metricName":"NA75"},{"values":[5562,3053,14140,6618,5968,39711,510,null,45873,29720,8102,3107,1253,39714],"quality":"More is better","isMain":true,"metricName":"NGA50"},{"values":[2092,1469,11700,3973,3293,20493,null,null,45873,26170,4129,1094,555,20497],"quality":"More is better","isMain":false,"metricName":"NGA75"},{"values":[4,7,2,4,4,1,36,null,1,1,3,7,14,1],"quality":"Less is better","isMain":false,"metricName":"LG50"},{"values":[9,14,3,7,7,1,null,null,1,2,5,17,31,1],"quality":"Less is better","isMain":false,"metricName":"LG75"},{"values":[4,7,2,4,3,1,14,null,1,1,3,7,9,1],"quality":"Less is better","isMain":false,"metricName":"LA50"},{"values":[9,13,3,6,7,2,24,null,1,2,6,15,18,2],"quality":"Less is better","isMain":false,"metricName":"LA75"},{"values":[4,7,2,4,4,1,36,null,1,1,3,7,14,1],"quality":"Less is better","isMain":true,"metricName":"LGA50"},{"values":[9,14,3,7,7,2,null,null,1,2,5,17,31,2],"quality":"Less is better","isMain":false,"metricName":"LGA75"}]],["Misassemblies",[{"values":[0,0,0,0,0,1,0,0,0,0,0,0,0,1],"quality":"Less is better","isMain":true,"metricName":"# misassemblies"},{"values":[0,0,0,0,0,1,0,0,0,0,0,0,0,1],"quality":"Less is better","isMain":false,"metricName":" # relocations"},{"values":[0,0,0,0,0,0,0,0,0,0,0,0,0,0],"quality":"Less is better","isMain":false,"metricName":" # translocations"},{"values":[0,0,0,0,0,0,0,0,0,0,0,0,0,0],"quality":"Less is better","isMain":false,"metricName":" # inversions"},{"values":[0,0,0,0,0,1,0,0,0,0,0,0,0,1],"quality":"Less is better","isMain":false,"metricName":"# misassembled contigs"},{"values":[0,0,0,0,0,60204,0,0,0,0,0,0,0,60211],"quality":"Less is better","isMain":true,"metricName":"Misassembled contigs length"},{"values":[0,0,0,0,0,0,0,0,0,0,0,0,0,0],"quality":"Less is better","isMain":false,"metricName":"# local misassemblies"},{"values":[0,0,0,0,0,0,0,12,0,0,0,0,0,0],"quality":"Less is better","isMain":false,"metricName":"# unaligned mis. contigs"}]],["Unaligned",[{"values":[0,0,0,0,0,0,0,0,0,0,0,0,0,0],"quality":"Less is better","isMain":false,"metricName":"# fully unaligned contigs"},{"values":[0,0,0,0,0,0,0,0,0,0,0,0,0,0],"quality":"Less is better","isMain":false,"metricName":"Fully unaligned length"},{"values":[0,0,0,0,0,0,0,9,0,0,2,0,0,0],"quality":"Less is better","isMain":false,"metricName":"# partially unaligned contigs"},{"values":[0,0,0,0,0,0,0,12240,0,0,2455,0,0,0],"quality":"Less is better","isMain":false,"metricName":"Partially unaligned length"}]],["Mismatches",[{"values":[0,0,0,0,44,5,5,0,0,0,0,4,3,12],"quality":"Less is better","isMain":false,"metricName":"# mismatches"},{"values":[0,0,0,0,3,1,0,9,0,0,0,0,0,6],"quality":"Less is better","isMain":false,"metricName":"# indels"},{"values":[0,0,0,0,3,1,0,430,0,0,0,0,0,8],"quality":"Less is better","isMain":false,"metricName":"Indels length"},{"values":["0.00","0.00","0.00","0.00","80.72","8.52","17.13","0.00","0.00","0.00","0.00","7.46","6.51","20.46"],"quality":"Less is better","isMain":true,"metricName":"# mismatches per 100 kbp"},{"values":["0.00","0.00","0.00","0.00","5.50","1.70","0.00","77.59","0.00","0.00","0.00","0.00","0.00","10.23"],"quality":"Less is better","isMain":true,"metricName":"# indels per 100 kbp"},{"values":[0,0,0,0,3,1,0,0,0,0,0,0,0,6],"quality":"Less is better","isMain":false,"metricName":" # indels (<= 5 bp)"},{"values":[0,0,0,0,0,0,0,9,0,0,0,0,0,0],"quality":"Less is better","isMain":false,"metricName":" # indels (> 5 bp)"},{"values":[0,0,0,0,0,3,0,8800,0,0,2455,0,0,431],"quality":"Less is better","isMain":false,"metricName":"# N's"},{"values":["0.00","0.00","0.00","0.00","0.00","4.98","0.00","33250.21","0.00","0.00","3981.96","0.00","0.00","715.82"],"quality":"Less is better","isMain":true,"metricName":"# N's per 100 kbp"}]],["Statistics without reference",[{"values":[19,27,8,14,14,1,37,19,4,3,14,31,35,1],"quality":"Equal","isMain":true,"metricName":"# contigs"},{"values":[16,18,7,9,14,1,8,10,4,3,9,18,18,1],"quality":"Equal","isMain":false,"metricName":"# contigs (>= 1000 bp)"},{"values":[4,2,3,4,4,1,0,0,3,2,4,1,0,1],"quality":"Equal","isMain":false,"metricName":"# contigs (>= 5000 bp)"},{"values":[0,0,3,1,1,1,0,0,1,2,1,0,0,1],"quality":"Equal","isMain":false,"metricName":"# contigs (>= 10000 bp)"},{"values":[0,0,0,0,0,1,0,0,1,2,0,0,0,1],"quality":"Equal","isMain":false,"metricName":"# contigs (>= 25000 bp)"},{"values":[0,0,0,0,0,1,0,0,0,0,0,0,0,1],"quality":"Equal","isMain":false,"metricName":"# contigs (>= 50000 bp)"},{"values":[9673,7890,22282,12538,12470,60204,1933,4140,45873,29720,18478,5116,3360,60211],"quality":"More is better","isMain":true,"metricName":"Largest contig"},{"values":[58983,56751,58839,57390,56227,60204,30100,26466,59171,58828,61653,54699,46277,60211],"quality":"More is better","isMain":true,"metricName":"Total length"},{"values":[56322,49799,58240,53958,56227,60204,10281,21021,59171,58828,58225,45168,34770,60211],"quality":"More is better","isMain":true,"metricName":"Total length (>= 1000 bp)"},{"values":[30525,12922,48122,34314,34503,60204,0,0,57766,55890,43522,5116,0,60211],"quality":"More is better","isMain":false,"metricName":"Total length (>= 5000 bp)"},{"values":[0,0,48122,12538,12470,60204,0,0,45873,55890,18478,0,0,60211],"quality":"More is better","isMain":true,"metricName":"Total length (>= 10000 bp)"},{"values":[0,0,0,0,0,60204,0,0,45873,55890,0,0,0,60211],"quality":"More is better","isMain":false,"metricName":"Total length (>= 25000 bp)"},{"values":[0,0,0,0,0,60204,0,0,0,0,0,0,0,60211],"quality":"More is better","isMain":true,"metricName":"Total length (>= 50000 bp)"},{"values":[5562,3053,14140,6618,7237,60204,813,1936,45873,29720,8149,3107,1906,60211],"quality":"More is better","isMain":false,"metricName":"N50"},{"values":[2092,2023,11700,4338,3293,60204,671,1313,45873,26170,4268,1217,1041,60211],"quality":"More is better","isMain":false,"metricName":"N75"},{"values":[4,7,2,4,3,1,14,5,1,1,3,7,9,1],"quality":"Less is better","isMain":false,"metricName":"L50"},{"values":[9,13,3,6,7,1,24,9,1,2,5,15,18,1],"quality":"Less is better","isMain":false,"metricName":"L75"},{"values":["64.32","64.30","64.25","64.42","64.30","64.20","64.03","63.67","64.27","64.26","64.24","64.14","64.36","64.28"],"quality":"Equal","isMain":false,"metricName":"GC (%)"}]],["Predicted genes",[]],["Similarity statistics",[{"values":[0,0,0,0,0,0,0,0,0,0,0,0,0,0],"quality":"Equal","isMain":false,"metricName":"# similar correct contigs"},{"values":[0,0,0,0,0,0,0,0,0,0,0,0,0,0],"quality":"Equal","isMain":false,"metricName":"# similar misassembled blocks"}]],["Reference statistics",[{"values":[58663,58663,58663,58663,58663,58663,58663,58663,58663,58663,58663,58663,58663,58663],"quality":"Equal","isMain":false,"metricName":"Reference length"},{"values":[1,1,1,1,1,1,1,1,1,1,1,1,1,1],"quality":"Equal","isMain":false,"metricName":"Reference fragments"},{"values":["64.26","64.26","64.26","64.26","64.26","64.26","64.26","64.26","64.26","64.26","64.26","64.26","64.26","64.26"],"quality":"Equal","isMain":false,"metricName":"Reference GC (%)"}]]],"referenceName":"gi\_162135082\_ref\_NC","date":"18 July 2018, Wednesday, 16:30:27","order":[0,1,2,3,4,5,6,7,8,9,10,11,12,13],"assembliesNames":["ABySS\_127","ABySS\_63","CLC","IDBA\_UD","MEGAHIT","MIRA","Ray\_Meta","SOAPdenovo2","SPAdes","SPAdes\_meta","SPAdes\_sc","SPAdes\_sc\_careful","Velvet","Geneious"]},{"assembliesWithNs":null,"minContig":500,"report":[["Genome statistics",[{"values":["50.229","22.329","61.370","42.110","51.066","92.665","57.429","78.136","32.076","48.048","51.047","11.372","96.094"],"quality":"More is better","isMain":true,"metricName":"Genome fraction (%)"},{"values":["1.010","1.010","1.070","1.017","1.038","2.055","1.038","1.087","1.000","1.005","1.007","1.006","1.724"],"quality":"Less is better","isMain":true,"metricName":"Duplication ratio"},{"values":[6746,2757,3415,3208,3292,22021,5876,6519,3739,5058,4915,1405,29428],"quality":"More is better","isMain":true,"metricName":"Largest alignment"},{"values":[21768,9698,27750,18413,22478,81520,25637,36515,13795,20776,22120,4920,71256],"quality":"More is better","isMain":true,"metricName":"Total aligned length"},{"values":[502,null,1071,null,548,28212,1366,1765,null,null,825,null,39847],"quality":"More is better","isMain":false,"metricName":"NG50"},{"values":[null,null,null,null,null,9683,null,948,null,null,null,null,39847],"quality":"More is better","isMain":false,"metricName":"NG75"},{"values":[1826,2305,1693,923,1208,3823,2785,3210,2756,2529,1201,855,452],"quality":"More is better","isMain":false,"metricName":"NA50"},{"values":[919,685,987,823,814,856,1401,1412,2682,1180,939,735,null],"quality":"More is better","isMain":false,"metricName":"NA75"},{"values":[502,null,987,null,520,22021,1366,1765,null,null,538,null,29428],"quality":"More is better","isMain":true,"metricName":"NGA50"},{"values":[null,null,null,null,null,6501,null,948,null,null,null,null,11819],"quality":"More is better","isMain":false,"metricName":"NGA75"},{"values":[14,null,12,null,19,1,8,6,null,null,10,null,1],"quality":"Less is better","isMain":false,"metricName":"LG50"},{"values":[null,null,null,null,null,2,null,13,null,null,null,null,1],"quality":"Less is better","isMain":false,"metricName":"LG75"},{"values":[3,2,6,6,7,6,4,4,3,3,5,3,13],"quality":"Less is better","isMain":false,"metricName":"LA50"},{"values":[7,5,12,11,12,23,7,9,4,6,10,4,null],"quality":"Less is better","isMain":false,"metricName":"LA75"},{"values":[14,null,12,null,20,1,8,6,null,null,15,null,1],"quality":"Less is better","isMain":true,"metricName":"LGA50"},{"values":[null,null,null,null,null,3,null,13,null,null,null,null,2],"quality":"Less is better","isMain":false,"metricName":"LGA75"}]],["Misassemblies",[{"values":[0,0,0,0,0,1,0,0,0,0,0,0,1],"quality":"Less is better","isMain":true,"metricName":"# misassemblies"},{"values":[0,0,0,0,0,1,0,0,0,0,0,0,1],"quality":"Less is better","isMain":false,"metricName":" # relocations"},{"values":[0,0,0,0,0,0,0,0,0,0,0,0,0],"quality":"Less is better","isMain":false,"metricName":" # translocations"},{"values":[0,0,0,0,0,0,0,0,0,0,0,0,0],"quality":"Less is better","isMain":false,"metricName":" # inversions"},{"values":[0,0,0,0,0,1,0,0,0,0,0,0,1],"quality":"Less is better","isMain":false,"metricName":"# misassembled contigs"},{"values":[0,0,0,0,0,28212,0,0,0,0,0,0,39847],"quality":"Less is better","isMain":true,"metricName":"Misassembled contigs length"},{"values":[1,0,0,0,0,3,1,0,0,0,0,0,2],"quality":"Less is better","isMain":false,"metricName":"# local misassemblies"},{"values":[0,0,0,0,0,1,0,0,0,0,1,0,3],"quality":"Less is better","isMain":false,"metricName":"# unaligned mis. contigs"}]],["Unaligned",[{"values":[0,0,0,0,0,0,0,0,0,0,0,0,0],"quality":"Less is better","isMain":false,"metricName":"# fully unaligned contigs"},{"values":[0,0,0,0,0,0,0,0,0,0,0,0,0],"quality":"Less is better","isMain":false,"metricName":"Fully unaligned length"},{"values":[0,0,0,0,0,5,1,0,1,0,1,0,6],"quality":"Less is better","isMain":false,"metricName":"# partially unaligned contigs"},{"values":[0,0,0,0,0,19528,936,0,2328,0,2435,0,69086],"quality":"Less is better","isMain":false,"metricName":"Partially unaligned length"}]],["Mismatches",[{"values":[48,1,44,6,17,688,121,43,50,115,185,0,722],"quality":"Less is better","isMain":false,"metricName":"# mismatches"},{"values":[2,0,0,0,0,20,0,0,1,2,2,0,14],"quality":"Less is better","isMain":false,"metricName":"# indels"},{"values":[2,0,0,0,0,24,0,0,1,2,2,0,17],"quality":"Less is better","isMain":false,"metricName":"Indels length"},{"values":["222.18","10.41","166.69","33.13","77.40","1726.21","489.86","127.95","362.42","556.47","842.59","0.00","1746.87"],"quality":"Less is better","isMain":true,"metricName":"# mismatches per 100 kbp"},{"values":["9.26","0.00","0.00","0.00","0.00","50.18","0.00","0.00","7.25","9.68","9.11","0.00","33.87"],"quality":"Less is better","isMain":true,"metricName":"# indels per 100 kbp"},{"values":[2,0,0,0,0,20,0,0,1,2,2,0,14],"quality":"Less is better","isMain":false,"metricName":" # indels (<= 5 bp)"},{"values":[0,0,0,0,0,0,0,0,0,0,0,0,0],"quality":"Less is better","isMain":false,"metricName":" # indels (> 5 bp)"},{"values":[50,1,0,0,0,34,0,0,0,0,0,0,980],"quality":"Less is better","isMain":false,"metricName":"# N's"},{"values":["229.17","10.31","0.00","0.00","0.00","33.52","0.00","0.00","0.00","0.00","0.00","0.00","698.29"],"quality":"Less is better","isMain":true,"metricName":"# N's per 100 kbp"}]],["Statistics without reference",[{"values":[14,9,22,18,21,29,11,19,6,12,14,6,6],"quality":"Equal","isMain":true,"metricName":"# contigs"},{"values":[4,2,13,4,9,20,11,12,4,8,8,1,6],"quality":"Equal","isMain":false,"metricName":"# contigs (>= 1000 bp)"},{"values":[1,0,0,0,0,6,1,1,1,1,0,0,5],"quality":"Equal","isMain":false,"metricName":"# contigs (>= 5000 bp)"},{"values":[0,0,0,0,0,1,0,0,0,0,0,0,4],"quality":"Equal","isMain":false,"metricName":"# contigs (>= 10000 bp)"},{"values":[0,0,0,0,0,1,0,0,0,0,0,0,3],"quality":"Equal","isMain":false,"metricName":"# contigs (>= 25000 bp)"},{"values":[0,0,0,0,0,0,0,0,0,0,0,0,0],"quality":"Equal","isMain":false,"metricName":"# contigs (>= 50000 bp)"},{"values":[6796,2757,3415,3208,3292,28212,6812,6519,5010,5058,4915,1405,39847],"quality":"More is better","isMain":true,"metricName":"Largest contig"},{"values":[21818,9698,28245,18413,22798,101418,26577,36515,16123,20776,24555,4920,140342],"quality":"More is better","isMain":true,"metricName":"Total length"},{"values":[14156,5062,22759,7854,14750,94359,26577,31530,14672,17895,20197,1405,140342],"quality":"More is better","isMain":true,"metricName":"Total length (>= 1000 bp)"},{"values":[6796,0,0,0,0,68328,6812,6519,5010,5058,0,0,139097],"quality":"More is better","isMain":false,"metricName":"Total length (>= 5000 bp)"},{"values":[0,0,0,0,0,28212,0,0,0,0,0,0,130195],"quality":"More is better","isMain":true,"metricName":"Total length (>= 10000 bp)"},{"values":[0,0,0,0,0,28212,0,0,0,0,0,0,112973],"quality":"More is better","isMain":false,"metricName":"Total length (>= 25000 bp)"},{"values":[0,0,0,0,0,0,0,0,0,0,0,0,0],"quality":"More is better","isMain":true,"metricName":"Total length (>= 50000 bp)"},{"values":[1826,2305,1693,923,1208,8212,2785,3210,3739,2529,3723,855,36887],"quality":"More is better","isMain":false,"metricName":"N50"},{"values":[919,685,1071,823,814,3823,1880,1412,2756,1180,1180,735,36239],"quality":"More is better","isMain":false,"metricName":"N75"},{"values":[3,2,6,6,7,4,4,4,2,3,3,3,2],"quality":"Less is better","isMain":false,"metricName":"L50"},{"values":[7,5,12,11,12,8,6,9,4,6,7,4,3],"quality":"Less is better","isMain":false,"metricName":"L75"},{"values":["35.37","33.61","35.84","35.24","35.41","35.43","35.73","35.60","34.71","35.31","35.28","32.52","35.20"],"quality":"Equal","isMain":false,"metricName":"GC (%)"}]],["Predicted genes",[]],["Similarity statistics",[{"values":[0,0,0,0,0,0,0,0,0,0,0,0,0],"quality":"Equal","isMain":false,"metricName":"# similar correct contigs"},{"values":[0,0,0,0,0,0,0,0,0,0,0,0,0],"quality":"Equal","isMain":false,"metricName":"# similar misassembled blocks"}]],["Reference statistics",[{"values":[43011,43011,43011,43011,43011,43011,43011,43011,43011,43011,43011,43011,43011],"quality":"Equal","isMain":false,"metricName":"Reference length"},{"values":[1,1,1,1,1,1,1,1,1,1,1,1,1],"quality":"Equal","isMain":false,"metricName":"Reference fragments"},{"values":["35.63","35.63","35.63","35.63","35.63","35.63","35.63","35.63","35.63","35.63","35.63","35.63","35.63"],"quality":"Equal","isMain":false,"metricName":"Reference GC (%)"}]]],"referenceName":"gi\_162290107\_ref\_NC","date":"18 July 2018, Wednesday, 16:30:40","order":[0,1,2,3,4,5,6,7,8,9,10,11,12],"assembliesNames":["ABySS\_127","ABySS\_63","CLC","IDBA\_UD","MEGAHIT","MIRA","Ray\_Meta","SPAdes","SPAdes\_meta","SPAdes\_sc","SPAdes\_sc\_careful","Velvet","Geneious"]},{"assembliesWithNs":null,"minContig":500,"report":[["Genome statistics",[{"values":["99.914","99.953","99.535","99.732","99.707","100.000","94.701","61.351","99.691","99.762","99.840","100.000","98.381","100.000"],"quality":"More is better","isMain":true,"metricName":"Genome fraction (%)"},{"values":["1.000","1.000","1.000","1.001","1.000","1.004","1.039","1.004","1.000","1.000","1.000","1.001","1.002","1.002"],"quality":"Less is better","isMain":true,"metricName":"Duplication ratio"},{"values":[177148,135081,176475,176922,176780,177332,29558,2704,176752,176878,177017,177410,66599,177643],"quality":"More is better","isMain":true,"metricName":"Largest alignment"},{"values":[177148,177274,176475,176922,176780,178034,174494,108895,176752,176878,177017,177410,174726,177643],"quality":"More is better","isMain":true,"metricName":"Total aligned length"},{"values":[177148,135081,176475,176922,176780,177340,8435,618,176752,176878,177017,177510,23595,177643],"quality":"More is better","isMain":false,"metricName":"NG50"},{"values":[177148,135081,176475,176922,176780,177340,3904,null,176752,176878,177017,177510,8265,177643],"quality":"More is better","isMain":false,"metricName":"NG75"},{"values":[177148,135081,176475,176922,176780,177332,8435,862,176752,176878,177017,177410,23595,177643],"quality":"More is better","isMain":false,"metricName":"NA50"},{"values":[177148,135081,176475,176922,176780,177332,3904,656,176752,176878,177017,177410,10165,177643],"quality":"More is better","isMain":false,"metricName":"NA75"},{"values":[177148,135081,176475,176922,176780,177332,8435,612,176752,176878,177017,177410,23595,177643],"quality":"More is better","isMain":true,"metricName":"NGA50"},{"values":[177148,135081,176475,176922,176780,177332,3904,null,176752,176878,177017,177410,8265,177643],"quality":"More is better","isMain":false,"metricName":"NGA75"},{"values":[1,1,1,1,1,1,5,91,1,1,1,1,2,1],"quality":"Less is better","isMain":false,"metricName":"LG50"},{"values":[1,1,1,1,1,1,13,null,1,1,1,1,6,1],"quality":"Less is better","isMain":false,"metricName":"LG75"},{"values":[1,1,1,1,1,1,5,43,1,1,1,1,2,1],"quality":"Less is better","isMain":false,"metricName":"LA50"},{"values":[1,1,1,1,1,1,13,81,1,1,1,1,5,1],"quality":"Less is better","isMain":false,"metricName":"LA75"},{"values":[1,1,1,1,1,1,5,91,1,1,1,1,2,1],"quality":"Less is better","isMain":true,"metricName":"LGA50"},{"values":[1,1,1,1,1,1,13,null,1,1,1,1,6,1],"quality":"Less is better","isMain":false,"metricName":"LGA75"}]],["Misassemblies",[{"values":[0,0,0,0,0,0,0,0,0,0,0,0,0,0],"quality":"Less is better","isMain":true,"metricName":"# misassemblies"},{"values":[0,0,0,0,0,0,0,0,0,0,0,0,0,0],"quality":"Less is better","isMain":false,"metricName":" # relocations"},{"values":[0,0,0,0,0,0,0,0,0,0,0,0,0,0],"quality":"Less is better","isMain":false,"metricName":" # translocations"},{"values":[0,0,0,0,0,0,0,0,0,0,0,0,0,0],"quality":"Less is better","isMain":false,"metricName":" # inversions"},{"values":[0,0,0,0,0,0,0,0,0,0,0,0,0,0],"quality":"Less is better","isMain":false,"metricName":"# misassembled contigs"},{"values":[0,0,0,0,0,0,0,0,0,0,0,0,0,0],"quality":"Less is better","isMain":true,"metricName":"Misassembled contigs length"},{"values":[0,0,0,1,0,1,0,0,0,0,0,1,0,2],"quality":"Less is better","isMain":false,"metricName":"# local misassemblies"},{"values":[0,0,0,0,0,0,0,0,0,0,0,0,0,0],"quality":"Less is better","isMain":false,"metricName":"# unaligned mis. contigs"}]],["Unaligned",[{"values":[0,0,0,0,0,0,0,0,0,0,0,0,0,0],"quality":"Less is better","isMain":false,"metricName":"# fully unaligned contigs"},{"values":[0,0,0,0,0,0,0,0,0,0,0,0,0,0],"quality":"Less is better","isMain":false,"metricName":"Fully unaligned length"},{"values":[0,0,0,0,0,0,0,0,0,0,0,0,0,0],"quality":"Less is better","isMain":false,"metricName":"# partially unaligned contigs"},{"values":[0,0,0,0,0,0,0,0,0,0,0,0,0,0],"quality":"Less is better","isMain":false,"metricName":"Partially unaligned length"}]],["Mismatches",[{"values":[0,0,0,4,0,0,0,8,0,0,0,0,0,0],"quality":"Less is better","isMain":false,"metricName":"# mismatches"},{"values":[0,1,0,1,0,0,0,2,0,0,0,0,0,0],"quality":"Less is better","isMain":false,"metricName":"# indels"},{"values":[0,1,0,1,0,0,0,90,0,0,0,0,0,0],"quality":"Less is better","isMain":false,"metricName":"Indels length"},{"values":["0.00","0.00","0.00","2.26","0.00","0.00","0.00","7.35","0.00","0.00","0.00","0.00","0.00","0.00"],"quality":"Less is better","isMain":true,"metricName":"# mismatches per 100 kbp"},{"values":["0.00","0.56","0.00","0.57","0.00","0.00","0.00","1.84","0.00","0.00","0.00","0.00","0.00","0.00"],"quality":"Less is better","isMain":true,"metricName":"# indels per 100 kbp"},{"values":[0,1,0,1,0,0,0,0,0,0,0,0,0,0],"quality":"Less is better","isMain":false,"metricName":" # indels (<= 5 bp)"},{"values":[0,0,0,0,0,0,0,2,0,0,0,0,0,0],"quality":"Less is better","isMain":false,"metricName":" # indels (> 5 bp)"},{"values":[0,0,0,0,0,0,0,324,0,0,0,100,0,10],"quality":"Less is better","isMain":false,"metricName":"# N's"},{"values":["0.00","0.00","0.00","0.00","0.00","0.00","0.00","296.65","0.00","0.00","0.00","56.33","0.00","5.63"],"quality":"Less is better","isMain":true,"metricName":"# N's per 100 kbp"}]],["Statistics without reference",[{"values":[1,2,1,1,1,2,32,127,1,1,1,1,21,1],"quality":"Equal","isMain":true,"metricName":"# contigs"},{"values":[1,2,1,1,1,1,29,29,1,1,1,1,16,1],"quality":"Equal","isMain":false,"metricName":"# contigs (>= 1000 bp)"},{"values":[1,2,1,1,1,1,12,0,1,1,1,1,7,1],"quality":"Equal","isMain":false,"metricName":"# contigs (>= 5000 bp)"},{"values":[1,2,1,1,1,1,3,0,1,1,1,1,5,1],"quality":"Equal","isMain":false,"metricName":"# contigs (>= 10000 bp)"},{"values":[1,2,1,1,1,1,1,0,1,1,1,1,1,1],"quality":"Equal","isMain":false,"metricName":"# contigs (>= 25000 bp)"},{"values":[1,1,1,1,1,1,0,0,1,1,1,1,1,1],"quality":"Equal","isMain":false,"metricName":"# contigs (>= 50000 bp)"},{"values":[177148,135081,176475,176922,176780,177340,29558,2704,176752,176878,177017,177510,66599,177643],"quality":"More is better","isMain":true,"metricName":"Largest contig"},{"values":[177148,177274,176475,176922,176780,178042,174494,109221,176752,176878,177017,177510,174780,177643],"quality":"More is better","isMain":true,"metricName":"Total length"},{"values":[177148,177274,176475,176922,176780,177340,172172,42098,176752,176878,177017,177510,171205,177643],"quality":"More is better","isMain":true,"metricName":"Total length (>= 1000 bp)"},{"values":[177148,177274,176475,176922,176780,177340,129952,0,176752,176878,177017,177510,145325,177643],"quality":"More is better","isMain":false,"metricName":"Total length (>= 5000 bp)"},{"values":[177148,177274,176475,176922,176780,177340,72986,0,176752,176878,177017,177510,131825,177643],"quality":"More is better","isMain":true,"metricName":"Total length (>= 10000 bp)"},{"values":[177148,177274,176475,176922,176780,177340,29558,0,176752,176878,177017,177510,66599,177643],"quality":"More is better","isMain":false,"metricName":"Total length (>= 25000 bp)"},{"values":[177148,135081,176475,176922,176780,177340,0,0,176752,176878,177017,177510,66599,177643],"quality":"More is better","isMain":true,"metricName":"Total length (>= 50000 bp)"},{"values":[177148,135081,176475,176922,176780,177340,8435,862,176752,176878,177017,177510,23595,177643],"quality":"More is better","isMain":false,"metricName":"N50"},{"values":[177148,135081,176475,176922,176780,177340,3904,660,176752,176878,177017,177510,10165,177643],"quality":"More is better","isMain":false,"metricName":"N75"},{"values":[1,1,1,1,1,1,5,43,1,1,1,1,2,1],"quality":"Less is better","isMain":false,"metricName":"L50"},{"values":[1,1,1,1,1,1,13,80,1,1,1,1,5,1],"quality":"Less is better","isMain":false,"metricName":"L75"},{"values":["40.62","40.63","40.60","40.61","40.61","40.66","40.56","40.43","40.61","40.62","40.62","40.64","40.54","40.65"],"quality":"Equal","isMain":false,"metricName":"GC (%)"}]],["Predicted genes",[]],["Similarity statistics",[{"values":[0,0,0,0,0,0,0,0,0,0,0,0,0,0],"quality":"Equal","isMain":false,"metricName":"# similar correct contigs"},{"values":[0,0,0,0,0,0,0,0,0,0,0,0,0,0],"quality":"Equal","isMain":false,"metricName":"# similar misassembled blocks"}]],["Reference statistics",[{"values":[177300,177300,177300,177300,177300,177300,177300,177300,177300,177300,177300,177300,177300,177300],"quality":"Equal","isMain":false,"metricName":"Reference length"},{"values":[1,1,1,1,1,1,1,1,1,1,1,1,1,1],"quality":"Equal","isMain":false,"metricName":"Reference fragments"},{"values":["40.63","40.63","40.63","40.63","40.63","40.63","40.63","40.63","40.63","40.63","40.63","40.63","40.63","40.63"],"quality":"Equal","isMain":false,"metricName":"Reference GC (%)"}]]],"referenceName":"gi\_162290187\_ref\_NC","date":"18 July 2018, Wednesday, 16:30:55","order":[0,1,2,3,4,5,6,7,8,9,10,11,12,13],"assembliesNames":["ABySS\_127","ABySS\_63","CLC","IDBA\_UD","MEGAHIT","MIRA","Ray\_Meta","SOAPdenovo2","SPAdes","SPAdes\_meta","SPAdes\_sc","SPAdes\_sc\_careful","Velvet","Geneious"]},{"assembliesWithNs":null,"minContig":500,"report":[["Genome statistics",[{"values":["99.683","99.680","99.832","100.000","99.680","100.000","100.000","8.527","99.680","99.683","99.683","99.683","99.615","100.000"],"quality":"More is better","isMain":true,"metricName":"Genome fraction (%)"},{"values":["1.000","1.000","1.000","1.003","1.000","1.015","1.001","1.000","1.003","1.004","1.004","1.004","1.001","1.008"],"quality":"Less is better","isMain":true,"metricName":"Duplication ratio"},{"values":[36780,34839,36833,36994,34839,29812,36944,804,36777,36828,36828,36830,31859,37175],"quality":"More is better","isMain":true,"metricName":"Largest alignment"},{"values":[36780,36777,36833,36994,36777,37445,36944,3146,36777,36828,36828,36830,36781,37175],"quality":"More is better","isMain":true,"metricName":"Total aligned length"},{"values":[36780,34839,36833,36994,34839,29812,36944,null,36893,36937,36937,36939,31859,37175],"quality":"More is better","isMain":false,"metricName":"NG50"},{"values":[36780,34839,36833,36994,34839,29812,36944,null,36893,36937,36937,36939,31859,37175],"quality":"More is better","isMain":false,"metricName":"NG75"},{"values":[36780,34839,36833,36994,34839,29812,36944,611,36777,36828,36828,36830,31859,37175],"quality":"More is better","isMain":false,"metricName":"NA50"},{"values":[36780,34839,36833,36994,34839,29812,36944,578,36777,36828,36828,36830,31859,37175],"quality":"More is better","isMain":false,"metricName":"NA75"},{"values":[36780,34839,36833,36994,34839,29812,36944,null,36777,36828,36828,36830,31859,37175],"quality":"More is better","isMain":true,"metricName":"NGA50"},{"values":[36780,34839,36833,36994,34839,29812,36944,null,36777,36828,36828,36830,31859,37175],"quality":"More is better","isMain":false,"metricName":"NGA75"},{"values":[1,1,1,1,1,1,1,null,1,1,1,1,1,1],"quality":"Less is better","isMain":false,"metricName":"LG50"},{"values":[1,1,1,1,1,1,1,null,1,1,1,1,1,1],"quality":"Less is better","isMain":false,"metricName":"LG75"},{"values":[1,1,1,1,1,1,1,3,1,1,1,1,1,1],"quality":"Less is better","isMain":false,"metricName":"LA50"},{"values":[1,1,1,1,1,1,1,4,1,1,1,1,1,1],"quality":"Less is better","isMain":false,"metricName":"LA75"},{"values":[1,1,1,1,1,1,1,null,1,1,1,1,1,1],"quality":"Less is better","isMain":true,"metricName":"LGA50"},{"values":[1,1,1,1,1,1,1,null,1,1,1,1,1,1],"quality":"Less is better","isMain":false,"metricName":"LGA75"}]],["Misassemblies",[{"values":[0,0,0,0,0,0,0,0,0,0,0,0,0,0],"quality":"Less is better","isMain":true,"metricName":"# misassemblies"},{"values":[0,0,0,0,0,0,0,0,0,0,0,0,0,0],"quality":"Less is better","isMain":false,"metricName":" # relocations"},{"values":[0,0,0,0,0,0,0,0,0,0,0,0,0,0],"quality":"Less is better","isMain":false,"metricName":" # translocations"},{"values":[0,0,0,0,0,0,0,0,0,0,0,0,0,0],"quality":"Less is better","isMain":false,"metricName":" # inversions"},{"values":[0,0,0,0,0,0,0,0,0,0,0,0,0,0],"quality":"Less is better","isMain":false,"metricName":"# misassembled contigs"},{"values":[0,0,0,0,0,0,0,0,0,0,0,0,0,0],"quality":"Less is better","isMain":true,"metricName":"Misassembled contigs length"},{"values":[0,0,0,1,0,0,0,0,0,0,0,0,0,1],"quality":"Less is better","isMain":false,"metricName":"# local misassemblies"},{"values":[0,0,0,0,0,0,0,0,0,0,0,0,0,0],"quality":"Less is better","isMain":false,"metricName":"# unaligned mis. contigs"}]],["Unaligned",[{"values":[0,0,0,0,0,0,0,0,0,0,0,0,0,0],"quality":"Less is better","isMain":false,"metricName":"# fully unaligned contigs"},{"values":[0,0,0,0,0,0,0,0,0,0,0,0,0,0],"quality":"Less is better","isMain":false,"metricName":"Fully unaligned length"},{"values":[0,0,0,0,0,0,0,0,0,0,0,0,0,0],"quality":"Less is better","isMain":false,"metricName":"# partially unaligned contigs"},{"values":[0,0,0,0,0,0,0,0,0,0,0,0,0,0],"quality":"Less is better","isMain":false,"metricName":"Partially unaligned length"}]],["Mismatches",[{"values":[0,0,2,2,0,0,0,2,0,0,0,0,0,0],"quality":"Less is better","isMain":false,"metricName":"# mismatches"},{"values":[1,0,0,0,0,0,1,0,1,2,2,1,0,0],"quality":"Less is better","isMain":false,"metricName":"# indels"},{"values":[2,0,0,0,0,0,49,0,1,10,10,8,0,0],"quality":"Less is better","isMain":false,"metricName":"Indels length"},{"values":["0.00","0.00","5.43","5.42","0.00","0.00","0.00","63.57","0.00","0.00","0.00","0.00","0.00","0.00"],"quality":"Less is better","isMain":true,"metricName":"# mismatches per 100 kbp"},{"values":["2.72","0.00","0.00","0.00","0.00","0.00","2.71","0.00","2.72","5.44","5.44","2.72","0.00","0.00"],"quality":"Less is better","isMain":true,"metricName":"# indels per 100 kbp"},{"values":[1,0,0,0,0,0,0,0,1,1,1,0,0,0],"quality":"Less is better","isMain":false,"metricName":" # indels (<= 5 bp)"},{"values":[0,0,0,0,0,0,1,0,0,1,1,1,0,0],"quality":"Less is better","isMain":false,"metricName":" # indels (> 5 bp)"},{"values":[3,0,3,0,0,3,0,0,116,109,109,110,0,3],"quality":"Less is better","isMain":false,"metricName":"# N's"},{"values":["8.16","0.00","8.14","0.00","0.00","8.01","0.00","0.00","314.42","295.10","295.10","297.79","0.00","8.07"],"quality":"Less is better","isMain":true,"metricName":"# N's per 100 kbp"}]],["Statistics without reference",[{"values":[1,2,1,1,2,3,1,5,1,1,1,1,2,1],"quality":"Equal","isMain":true,"metricName":"# contigs"},{"values":[1,2,1,1,2,3,1,0,1,1,1,1,2,1],"quality":"Equal","isMain":false,"metricName":"# contigs (>= 1000 bp)"},{"values":[1,1,1,1,1,2,1,0,1,1,1,1,1,1],"quality":"Equal","isMain":false,"metricName":"# contigs (>= 5000 bp)"},{"values":[1,1,1,1,1,1,1,0,1,1,1,1,1,1],"quality":"Equal","isMain":false,"metricName":"# contigs (>= 10000 bp)"},{"values":[1,1,1,1,1,1,1,0,1,1,1,1,1,1],"quality":"Equal","isMain":false,"metricName":"# contigs (>= 25000 bp)"},{"values":[0,0,0,0,0,0,0,0,0,0,0,0,0,0],"quality":"Equal","isMain":false,"metricName":"# contigs (>= 50000 bp)"},{"values":[36780,34839,36833,36994,34839,29812,36944,804,36893,36937,36937,36939,31859,37175],"quality":"More is better","isMain":true,"metricName":"Largest contig"},{"values":[36780,36777,36833,36994,36777,37445,36944,3146,36893,36937,36937,36939,36781,37175],"quality":"More is better","isMain":true,"metricName":"Total length"},{"values":[36780,36777,36833,36994,36777,37445,36944,0,36893,36937,36937,36939,36781,37175],"quality":"More is better","isMain":true,"metricName":"Total length (>= 1000 bp)"},{"values":[36780,34839,36833,36994,34839,34890,36944,0,36893,36937,36937,36939,31859,37175],"quality":"More is better","isMain":false,"metricName":"Total length (>= 5000 bp)"},{"values":[36780,34839,36833,36994,34839,29812,36944,0,36893,36937,36937,36939,31859,37175],"quality":"More is better","isMain":true,"metricName":"Total length (>= 10000 bp)"},{"values":[36780,34839,36833,36994,34839,29812,36944,0,36893,36937,36937,36939,31859,37175],"quality":"More is better","isMain":false,"metricName":"Total length (>= 25000 bp)"},{"values":[0,0,0,0,0,0,0,0,0,0,0,0,0,0],"quality":"More is better","isMain":true,"metricName":"Total length (>= 50000 bp)"},{"values":[36780,34839,36833,36994,34839,29812,36944,611,36893,36937,36937,36939,31859,37175],"quality":"More is better","isMain":false,"metricName":"N50"},{"values":[36780,34839,36833,36994,34839,29812,36944,578,36893,36937,36937,36939,31859,37175],"quality":"More is better","isMain":false,"metricName":"N75"},{"values":[1,1,1,1,1,1,1,3,1,1,1,1,1,1],"quality":"Less is better","isMain":false,"metricName":"L50"},{"values":[1,1,1,1,1,1,1,4,1,1,1,1,1,1],"quality":"Less is better","isMain":false,"metricName":"L75"},{"values":["37.98","37.98","38.00","37.96","37.98","37.93","37.97","38.24","37.98","37.98","37.98","37.98","37.96","37.88"],"quality":"Equal","isMain":false,"metricName":"GC (%)"}]],["Predicted genes",[]],["Similarity statistics",[{"values":[0,0,0,0,0,0,0,0,0,0,0,0,0,0],"quality":"Equal","isMain":false,"metricName":"# similar correct contigs"},{"values":[0,0,0,0,0,0,0,0,0,0,0,0,0,0],"quality":"Equal","isMain":false,"metricName":"# similar misassembled blocks"}]],["Reference statistics",[{"values":[36895,36895,36895,36895,36895,36895,36895,36895,36895,36895,36895,36895,36895,36895],"quality":"Equal","isMain":false,"metricName":"Reference length"},{"values":[1,1,1,1,1,1,1,1,1,1,1,1,1,1],"quality":"Equal","isMain":false,"metricName":"Reference fragments"},{"values":["37.98","37.98","37.98","37.98","37.98","37.98","37.98","37.98","37.98","37.98","37.98","37.98","37.98","37.98"],"quality":"Equal","isMain":false,"metricName":"Reference GC (%)"}]]],"referenceName":"gi\_162329424\_ref\_NC","date":"18 July 2018, Wednesday, 16:31:08","order":[0,1,2,3,4,5,6,7,8,9,10,11,12,13],"assembliesNames":["ABySS\_127","ABySS\_63","CLC","IDBA\_UD","MEGAHIT","MIRA","Ray\_Meta","SOAPdenovo2","SPAdes","SPAdes\_meta","SPAdes\_sc","SPAdes\_sc\_careful","Velvet","Geneious"]},{"assembliesWithNs":null,"minContig":500,"report":[["Genome statistics",[{"values":["0.806","0.341","40.623","29.698","41.227","3.337","34.806","47.827","46.731","46.894","46.894"],"quality":"More is better","isMain":true,"metricName":"Genome fraction (%)"},{"values":["1.000","1.000","1.012","1.010","1.007","1.000","1.002","1.006","1.002","1.023","1.023"],"quality":"Less is better","isMain":true,"metricName":"Duplication ratio"},{"values":[751,533,1896,1822,1609,1095,1620,1989,1896,1896,1896],"quality":"More is better","isMain":true,"metricName":"Largest alignment"},{"values":[1259,533,63788,46432,64450,5211,54365,74760,73096,73303,73303],"quality":"More is better","isMain":true,"metricName":"Total aligned length"},{"values":[751,533,794,760,757,890,716,891,868,856,856],"quality":"More is better","isMain":false,"metricName":"NA50"},{"values":[508,533,646,632,624,825,607,677,681,665,665],"quality":"More is better","isMain":false,"metricName":"NA75"},{"values":[null,null,null,null,null,null,null,null,null,null,null],"quality":"More is better","isMain":true,"metricName":"NGA50"},{"values":[1,1,31,24,32,3,28,31,32,33,33],"quality":"Less is better","isMain":false,"metricName":"LA50"},{"values":[2,1,54,41,56,5,49,56,55,59,59],"quality":"Less is better","isMain":false,"metricName":"LA75"}]],["Misassemblies",[{"values":[0,0,0,0,0,0,0,1,0,0,0],"quality":"Less is better","isMain":true,"metricName":"# misassemblies"},{"values":[0,0,0,0,0,0,0,1,0,0,0],"quality":"Less is better","isMain":false,"metricName":" # relocations"},{"values":[0,0,0,0,0,0,0,0,0,0,0],"quality":"Less is better","isMain":false,"metricName":" # translocations"},{"values":[0,0,0,0,0,0,0,0,0,0,0],"quality":"Less is better","isMain":false,"metricName":" # inversions"},{"values":[0,0,0,0,0,0,0,1,0,0,0],"quality":"Less is better","isMain":false,"metricName":"# misassembled contigs"},{"values":[0,0,0,0,0,0,0,771,0,0,0],"quality":"Less is better","isMain":true,"metricName":"Misassembled contigs length"},{"values":[0,0,0,0,0,0,0,0,0,0,0],"quality":"Less is better","isMain":false,"metricName":"# local misassemblies"},{"values":[0,0,0,0,0,0,0,0,0,0,0],"quality":"Less is better","isMain":false,"metricName":"# unaligned mis. contigs"}]],["Unaligned",[{"values":[0,0,0,0,0,0,0,0,0,0,0],"quality":"Less is better","isMain":false,"metricName":"# fully unaligned contigs"},{"values":[0,0,0,0,0,0,0,0,0,0,0],"quality":"Less is better","isMain":false,"metricName":"Fully unaligned length"},{"values":[0,0,0,2,0,0,0,2,0,3,3],"quality":"Less is better","isMain":false,"metricName":"# partially unaligned contigs"},{"values":[0,0,0,1411,0,0,0,1408,0,2450,2450],"quality":"Less is better","isMain":false,"metricName":"Partially unaligned length"}]],["Mismatches",[{"values":[0,0,77,38,54,0,52,117,90,116,99],"quality":"Less is better","isMain":false,"metricName":"# mismatches"},{"values":[0,0,0,0,1,0,1,0,1,1,1],"quality":"Less is better","isMain":false,"metricName":"# indels"},{"values":[0,0,0,0,79,0,57,0,79,79,79],"quality":"Less is better","isMain":false,"metricName":"Indels length"},{"values":["0.00","0.00","121.37","81.93","83.87","0.00","95.66","156.64","123.31","158.39","135.17"],"quality":"Less is better","isMain":true,"metricName":"# mismatches per 100 kbp"},{"values":["0.00","0.00","0.00","0.00","1.55","0.00","1.84","0.00","1.37","1.37","1.37"],"quality":"Less is better","isMain":true,"metricName":"# indels per 100 kbp"},{"values":[0,0,0,0,0,0,0,0,0,0,0],"quality":"Less is better","isMain":false,"metricName":" # indels (<= 5 bp)"},{"values":[0,0,0,0,1,0,1,0,1,1,1],"quality":"Less is better","isMain":false,"metricName":" # indels (> 5 bp)"},{"values":[0,0,0,0,0,13,106,0,0,0,0],"quality":"Less is better","isMain":false,"metricName":"# N's"},{"values":["0.00","0.00","0.00","0.00","0.00","249.47","194.60","0.00","0.00","0.00","0.00"],"quality":"Less is better","isMain":true,"metricName":"# N's per 100 kbp"}]],["Statistics without reference",[{"values":[2,1,81,60,84,6,73,86,86,87,87],"quality":"Equal","isMain":true,"metricName":"# contigs"},{"values":[0,0,14,8,11,1,10,24,22,24,24],"quality":"Equal","isMain":false,"metricName":"# contigs (>= 1000 bp)"},{"values":[0,0,0,0,0,0,0,0,0,0,0],"quality":"Equal","isMain":false,"metricName":"# contigs (>= 5000 bp)"},{"values":[0,0,0,0,0,0,0,0,0,0,0],"quality":"Equal","isMain":false,"metricName":"# contigs (>= 10000 bp)"},{"values":[0,0,0,0,0,0,0,0,0,0,0],"quality":"Equal","isMain":false,"metricName":"# contigs (>= 25000 bp)"},{"values":[0,0,0,0,0,0,0,0,0,0,0],"quality":"Equal","isMain":false,"metricName":"# contigs (>= 50000 bp)"},{"values":[751,533,1896,1838,1609,1095,1620,1989,1896,2033,2033],"quality":"More is better","isMain":true,"metricName":"Largest contig"},{"values":[1259,533,64203,48277,64860,5211,54471,76573,73096,77365,77365],"quality":"More is better","isMain":true,"metricName":"Total length"},{"values":[0,0,17787,11675,14082,1095,12812,32458,28007,32634,32634],"quality":"More is better","isMain":true,"metricName":"Total length (>= 1000 bp)"},{"values":[0,0,0,0,0,0,0,0,0,0,0],"quality":"More is better","isMain":false,"metricName":"Total length (>= 5000 bp)"},{"values":[0,0,0,0,0,0,0,0,0,0,0],"quality":"More is better","isMain":true,"metricName":"Total length (>= 10000 bp)"},{"values":[0,0,0,0,0,0,0,0,0,0,0],"quality":"More is better","isMain":false,"metricName":"Total length (>= 25000 bp)"},{"values":[0,0,0,0,0,0,0,0,0,0,0],"quality":"More is better","isMain":true,"metricName":"Total length (>= 50000 bp)"},{"values":[751,533,794,769,783,890,716,917,868,917,917],"quality":"More is better","isMain":false,"metricName":"N50"},{"values":[508,533,655,646,629,825,607,707,681,690,690],"quality":"More is better","isMain":false,"metricName":"N75"},{"values":[1,1,31,23,32,3,28,31,32,31,31],"quality":"Less is better","isMain":false,"metricName":"L50"},{"values":[2,1,54,40,56,5,49,55,55,55,55],"quality":"Less is better","isMain":false,"metricName":"L75"},{"values":["42.10","44.09","39.71","39.74","39.75","38.05","39.55","39.66","39.89","39.68","39.67"],"quality":"Equal","isMain":false,"metricName":"GC (%)"}]],["Predicted genes",[]],["Similarity statistics",[{"values":[0,0,0,0,0,0,0,0,0,0,0],"quality":"Equal","isMain":false,"metricName":"# similar correct contigs"},{"values":[0,0,0,0,0,0,0,0,0,0,0],"quality":"Equal","isMain":false,"metricName":"# similar misassembled blocks"}]],["Reference statistics",[{"values":[156179,156179,156179,156179,156179,156179,156179,156179,156179,156179,156179],"quality":"Equal","isMain":false,"metricName":"Reference length"},{"values":[1,1,1,1,1,1,1,1,1,1,1],"quality":"Equal","isMain":false,"metricName":"Reference fragments"},{"values":["39.88","39.88","39.88","39.88","39.88","39.88","39.88","39.88","39.88","39.88","39.88"],"quality":"Equal","isMain":false,"metricName":"Reference GC (%)"}]]],"referenceName":"gi\_165969010\_ref\_NC","date":"18 July 2018, Wednesday, 16:31:20","order":[0,1,2,3,4,5,6,7,8,9,10],"assembliesNames":["ABySS\_127","ABySS\_63","CLC","IDBA\_UD","MEGAHIT","MIRA","SOAPdenovo2","SPAdes","SPAdes\_meta","SPAdes\_sc","SPAdes\_sc\_careful"]},{"assembliesWithNs":null,"minContig":500,"report":[["Genome statistics",[{"values":["62.172","42.446","95.871","94.465","93.418","82.577","93.303","95.871","95.404","95.337","95.337","67.574","19.973"],"quality":"More is better","isMain":true,"metricName":"Genome fraction (%)"},{"values":["1.001","1.002","1.001","1.001","1.001","1.022","1.008","1.000","1.005","1.000","1.000","1.000","1.000"],"quality":"Less is better","isMain":true,"metricName":"Duplication ratio"},{"values":[3597,2720,11074,11074,11004,6237,7250,11074,11074,11074,11074,11074,3367],"quality":"More is better","isMain":true,"metricName":"Largest alignment"},{"values":[20614,14084,31788,31300,30953,27966,30922,31766,31757,31589,31589,22390,6618],"quality":"More is better","isMain":true,"metricName":"Total aligned length"},{"values":[766,null,10214,8163,8163,2390,3659,10214,6446,10214,10214,3661,null],"quality":"More is better","isMain":false,"metricName":"NG50"},{"values":[null,null,3661,3026,2981,1182,1791,3661,3661,3661,3661,null,null],"quality":"More is better","isMain":false,"metricName":"NG75"},{"values":[1226,1009,10214,8163,8163,2492,3659,10214,6446,10214,10214,4871,3367],"quality":"More is better","isMain":false,"metricName":"NA50"},{"values":[801,818,3661,3026,2981,1465,1791,3661,3661,3661,3661,3661,3251],"quality":"More is better","isMain":false,"metricName":"NA75"},{"values":[766,null,10214,8163,8163,2390,3659,10214,6446,10214,10214,3661,null],"quality":"More is better","isMain":true,"metricName":"NGA50"},{"values":[null,null,3661,3026,2981,1182,1711,3661,3661,3661,3661,null,null],"quality":"More is better","isMain":false,"metricName":"NGA75"},{"values":[13,null,2,2,2,5,3,2,2,2,2,3,null],"quality":"Less is better","isMain":false,"metricName":"LG50"},{"values":[null,null,3,4,4,11,7,3,4,3,3,null,null],"quality":"Less is better","isMain":false,"metricName":"LG75"},{"values":[6,5,2,2,2,4,3,2,2,2,2,2,1],"quality":"Less is better","isMain":false,"metricName":"LA50"},{"values":[11,9,3,4,4,8,6,3,4,3,3,3,2],"quality":"Less is better","isMain":false,"metricName":"LA75"},{"values":[13,null,2,2,2,5,3,2,2,2,2,3,null],"quality":"Less is better","isMain":true,"metricName":"LGA50"},{"values":[null,null,3,4,4,11,7,3,4,3,3,null,null],"quality":"Less is better","isMain":false,"metricName":"LGA75"}]],["Misassemblies",[{"values":[0,0,0,0,0,0,0,0,0,0,0,0,0],"quality":"Less is better","isMain":true,"metricName":"# misassemblies"},{"values":[0,0,0,0,0,0,0,0,0,0,0,0,0],"quality":"Less is better","isMain":false,"metricName":" # relocations"},{"values":[0,0,0,0,0,0,0,0,0,0,0,0,0],"quality":"Less is better","isMain":false,"metricName":" # translocations"},{"values":[0,0,0,0,0,0,0,0,0,0,0,0,0],"quality":"Less is better","isMain":false,"metricName":" # inversions"},{"values":[0,0,0,0,0,0,0,0,0,0,0,0,0],"quality":"Less is better","isMain":false,"metricName":"# misassembled contigs"},{"values":[0,0,0,0,0,0,0,0,0,0,0,0,0],"quality":"Less is better","isMain":true,"metricName":"Misassembled contigs length"},{"values":[0,0,0,0,0,0,0,0,0,0,0,0,0],"quality":"Less is better","isMain":false,"metricName":"# local misassemblies"},{"values":[0,0,0,0,0,0,0,0,0,0,0,0,0],"quality":"Less is better","isMain":false,"metricName":"# unaligned mis. contigs"}]],["Unaligned",[{"values":[0,0,0,0,0,0,0,0,0,0,0,0,0],"quality":"Less is better","isMain":false,"metricName":"# fully unaligned contigs"},{"values":[0,0,0,0,0,0,0,0,0,0,0,0,0],"quality":"Less is better","isMain":false,"metricName":"Fully unaligned length"},{"values":[0,0,0,0,0,0,0,0,0,1,1,0,0],"quality":"Less is better","isMain":false,"metricName":"# partially unaligned contigs"},{"values":[0,0,0,0,0,0,0,0,0,910,910,0,0],"quality":"Less is better","isMain":false,"metricName":"Partially unaligned length"}]],["Mismatches",[{"values":[3,2,17,9,10,5,7,17,17,9,6,4,0],"quality":"Less is better","isMain":false,"metricName":"# mismatches"},{"values":[0,0,0,0,0,0,2,0,0,0,0,0,0],"quality":"Less is better","isMain":false,"metricName":"# indels"},{"values":[0,0,0,0,0,0,106,0,0,0,0,0,0],"quality":"Less is better","isMain":false,"metricName":"Indels length"},{"values":["14.56","14.22","53.52","28.75","32.31","18.27","22.64","53.52","53.78","28.49","18.99","17.87","0.00"],"quality":"Less is better","isMain":true,"metricName":"# mismatches per 100 kbp"},{"values":["0.00","0.00","0.00","0.00","0.00","0.00","6.47","0.00","0.00","0.00","0.00","0.00","0.00"],"quality":"Less is better","isMain":true,"metricName":"# indels per 100 kbp"},{"values":[0,0,0,0,0,0,0,0,0,0,0,0,0],"quality":"Less is better","isMain":false,"metricName":" # indels (<= 5 bp)"},{"values":[0,0,0,0,0,0,2,0,0,0,0,0,0],"quality":"Less is better","isMain":false,"metricName":" # indels (> 5 bp)"},{"values":[0,0,0,0,0,35,228,0,0,0,0,0,7],"quality":"Less is better","isMain":false,"metricName":"# N's"},{"values":["0.00","0.00","0.00","0.00","0.00","125.15","731.73","0.00","0.00","0.00","0.00","0.00","105.77"],"quality":"Less is better","isMain":true,"metricName":"# N's per 100 kbp"}]],["Statistics without reference",[{"values":[19,14,8,8,8,14,12,7,8,7,7,5,2],"quality":"Equal","isMain":true,"metricName":"# contigs"},{"values":[8,6,6,7,7,12,11,6,7,6,6,5,2],"quality":"Equal","isMain":false,"metricName":"# contigs (>= 1000 bp)"},{"values":[0,0,2,2,2,1,2,2,2,2,2,1,0],"quality":"Equal","isMain":false,"metricName":"# contigs (>= 5000 bp)"},{"values":[0,0,2,1,1,0,0,2,1,2,2,1,0],"quality":"Equal","isMain":false,"metricName":"# contigs (>= 10000 bp)"},{"values":[0,0,0,0,0,0,0,0,0,0,0,0,0],"quality":"Equal","isMain":false,"metricName":"# contigs (>= 25000 bp)"},{"values":[0,0,0,0,0,0,0,0,0,0,0,0,0],"quality":"Equal","isMain":false,"metricName":"# contigs (>= 50000 bp)"},{"values":[3597,2720,11074,11074,11004,6237,7250,11074,11074,11074,11074,11074,3367],"quality":"More is better","isMain":true,"metricName":"Largest contig"},{"values":[20623,14093,31788,31332,30985,27966,31159,31766,31757,32499,32499,22390,6618],"quality":"More is better","isMain":true,"metricName":"Total length"},{"values":[13123,8266,30535,30672,30429,26516,30540,31106,31097,31839,31839,22390,6618],"quality":"More is better","isMain":true,"metricName":"Total length (>= 1000 bp)"},{"values":[0,0,21288,19237,19167,6237,13814,21288,17520,21288,21288,11074,0],"quality":"More is better","isMain":false,"metricName":"Total length (>= 5000 bp)"},{"values":[0,0,21288,11074,11004,0,0,21288,11074,21288,21288,11074,0],"quality":"More is better","isMain":true,"metricName":"Total length (>= 10000 bp)"},{"values":[0,0,0,0,0,0,0,0,0,0,0,0,0],"quality":"More is better","isMain":false,"metricName":"Total length (>= 25000 bp)"},{"values":[0,0,0,0,0,0,0,0,0,0,0,0,0],"quality":"More is better","isMain":true,"metricName":"Total length (>= 50000 bp)"},{"values":[1226,1009,10214,8163,8163,2492,3659,10214,6446,10214,10214,4871,3367],"quality":"More is better","isMain":false,"metricName":"N50"},{"values":[801,818,3661,3026,2981,1465,1837,3661,3661,3661,3661,3661,3251],"quality":"More is better","isMain":false,"metricName":"N75"},{"values":[6,5,2,2,2,4,3,2,2,2,2,2,1],"quality":"Less is better","isMain":false,"metricName":"L50"},{"values":[11,9,3,4,4,8,6,3,4,3,3,3,2],"quality":"Less is better","isMain":false,"metricName":"L75"},{"values":["48.16","47.45","48.94","48.92","48.96","48.68","48.96","48.94","48.98","49.35","49.35","49.02","50.36"],"quality":"Equal","isMain":false,"metricName":"GC (%)"}]],["Predicted genes",[]],["Similarity statistics",[{"values":[0,0,1,1,1,0,0,1,1,1,1,1,0],"quality":"Equal","isMain":false,"metricName":"# similar correct contigs"},{"values":[0,0,0,0,0,0,0,0,0,0,0,0,0],"quality":"Equal","isMain":false,"metricName":"# similar misassembled blocks"}]],["Reference statistics",[{"values":[33134,33134,33134,33134,33134,33134,33134,33134,33134,33134,33134,33134,33134],"quality":"Equal","isMain":false,"metricName":"Reference length"},{"values":[1,1,1,1,1,1,1,1,1,1,1,1,1],"quality":"Equal","isMain":false,"metricName":"Reference fragments"},{"values":["48.89","48.89","48.89","48.89","48.89","48.89","48.89","48.89","48.89","48.89","48.89","48.89","48.89"],"quality":"Equal","isMain":false,"metricName":"Reference GC (%)"}]]],"referenceName":"gi\_165970234\_ref\_NC","date":"18 July 2018, Wednesday, 16:31:34","order":[0,1,2,3,4,5,6,7,8,9,10,11,12],"assembliesNames":["ABySS\_127","ABySS\_63","CLC","IDBA\_UD","MEGAHIT","MIRA","SOAPdenovo2","SPAdes","SPAdes\_meta","SPAdes\_sc","SPAdes\_sc\_careful","Velvet","Geneious"]},{"assembliesWithNs":null,"minContig":500,"report":[["Genome statistics",[{"values":["99.560","98.840","100.000","99.751","99.859","99.682","12.269","98.337","99.823","99.638","99.638","99.638","99.573","99.682"],"quality":"More is better","isMain":true,"metricName":"Genome fraction (%)"},{"values":["1.001","1.000","1.004","1.000","1.000","1.000","1.000","1.033","1.000","1.000","1.000","1.000","1.000","1.000"],"quality":"Less is better","isMain":true,"metricName":"Duplication ratio"},{"values":[45903,22588,77854,77361,77445,77307,1459,44645,77417,77273,77273,77273,46897,77307],"quality":"More is better","isMain":true,"metricName":"Largest alignment"},{"values":[77294,76689,77854,77361,77445,77307,9515,77123,77417,77273,77273,77273,77250,77307],"quality":"More is better","isMain":true,"metricName":"Total aligned length"},{"values":[45903,20045,77854,77361,77445,77307,null,45686,77417,77273,77273,77273,46897,77307],"quality":"More is better","isMain":false,"metricName":"NG50"},{"values":[16130,16080,77854,77361,77445,77307,null,31781,77417,77273,77273,77273,30353,77307],"quality":"More is better","isMain":false,"metricName":"NG75"},{"values":[45903,20045,77854,77361,77445,77307,636,44645,77417,77273,77273,77273,46897,77307],"quality":"More is better","isMain":false,"metricName":"NA50"},{"values":[16130,16080,77854,77361,77445,77307,521,31152,77417,77273,77273,77273,30353,77307],"quality":"More is better","isMain":false,"metricName":"NA75"},{"values":[45903,20045,77854,77361,77445,77307,null,44645,77417,77273,77273,77273,46897,77307],"quality":"More is better","isMain":true,"metricName":"NGA50"},{"values":[16130,16080,77854,77361,77445,77307,null,31152,77417,77273,77273,77273,30353,77307],"quality":"More is better","isMain":false,"metricName":"NGA75"},{"values":[1,2,1,1,1,1,null,1,1,1,1,1,1,1],"quality":"Less is better","isMain":false,"metricName":"LG50"},{"values":[2,3,1,1,1,1,null,2,1,1,1,1,2,1],"quality":"Less is better","isMain":false,"metricName":"LG75"},{"values":[1,2,1,1,1,1,6,1,1,1,1,1,1,1],"quality":"Less is better","isMain":false,"metricName":"LA50"},{"values":[2,3,1,1,1,1,10,2,1,1,1,1,2,1],"quality":"Less is better","isMain":false,"metricName":"LA75"},{"values":[1,2,1,1,1,1,null,1,1,1,1,1,1,1],"quality":"Less is better","isMain":true,"metricName":"LGA50"},{"values":[2,3,1,1,1,1,null,2,1,1,1,1,2,1],"quality":"Less is better","isMain":false,"metricName":"LGA75"}]],["Misassemblies",[{"values":[0,0,0,0,0,0,0,0,0,0,0,0,0,0],"quality":"Less is better","isMain":true,"metricName":"# misassemblies"},{"values":[0,0,0,0,0,0,0,0,0,0,0,0,0,0],"quality":"Less is better","isMain":false,"metricName":" # relocations"},{"values":[0,0,0,0,0,0,0,0,0,0,0,0,0,0],"quality":"Less is better","isMain":false,"metricName":" # translocations"},{"values":[0,0,0,0,0,0,0,0,0,0,0,0,0,0],"quality":"Less is better","isMain":false,"metricName":" # inversions"},{"values":[0,0,0,0,0,0,0,0,0,0,0,0,0,0],"quality":"Less is better","isMain":false,"metricName":"# misassembled contigs"},{"values":[0,0,0,0,0,0,0,0,0,0,0,0,0,0],"quality":"Less is better","isMain":true,"metricName":"Misassembled contigs length"},{"values":[0,0,1,0,0,0,0,0,0,0,0,0,0,0],"quality":"Less is better","isMain":false,"metricName":"# local misassemblies"},{"values":[0,0,0,0,0,0,0,0,0,0,0,0,0,0],"quality":"Less is better","isMain":false,"metricName":"# unaligned mis. contigs"}]],["Unaligned",[{"values":[0,0,0,0,0,0,0,0,0,0,0,0,0,0],"quality":"Less is better","isMain":false,"metricName":"# fully unaligned contigs"},{"values":[0,0,0,0,0,0,0,0,0,0,0,0,0,0],"quality":"Less is better","isMain":false,"metricName":"Fully unaligned length"},{"values":[0,0,0,0,0,0,0,0,0,0,0,0,0,0],"quality":"Less is better","isMain":false,"metricName":"# partially unaligned contigs"},{"values":[0,0,0,0,0,0,0,0,0,0,0,0,0,0],"quality":"Less is better","isMain":false,"metricName":"Partially unaligned length"}]],["Mismatches",[{"values":[3,0,0,0,0,0,0,0,0,0,0,0,0,0],"quality":"Less is better","isMain":false,"metricName":"# mismatches"},{"values":[0,0,0,0,0,0,0,34,0,0,0,0,0,0],"quality":"Less is better","isMain":false,"metricName":"# indels"},{"values":[0,0,0,0,0,0,0,1635,0,0,0,0,0,0],"quality":"Less is better","isMain":false,"metricName":"Indels length"},{"values":["3.89","0.00","0.00","0.00","0.00","0.00","0.00","0.00","0.00","0.00","0.00","0.00","0.00","0.00"],"quality":"Less is better","isMain":true,"metricName":"# mismatches per 100 kbp"},{"values":["0.00","0.00","0.00","0.00","0.00","0.00","0.00","44.58","0.00","0.00","0.00","0.00","0.00","0.00"],"quality":"Less is better","isMain":true,"metricName":"# indels per 100 kbp"},{"values":[0,0,0,0,0,0,0,0,0,0,0,0,0,0],"quality":"Less is better","isMain":false,"metricName":" # indels (<= 5 bp)"},{"values":[0,0,0,0,0,0,0,34,0,0,0,0,0,0],"quality":"Less is better","isMain":false,"metricName":" # indels (> 5 bp)"},{"values":[0,0,0,0,0,0,0,2852,0,0,0,0,0,0],"quality":"Less is better","isMain":false,"metricName":"# N's"},{"values":["0.00","0.00","0.00","0.00","0.00","0.00","0.00","3619.61","0.00","0.00","0.00","0.00","0.00","0.00"],"quality":"Less is better","isMain":true,"metricName":"# N's per 100 kbp"}]],["Statistics without reference",[{"values":[5,8,1,1,1,1,14,3,1,1,1,1,2,1],"quality":"Equal","isMain":true,"metricName":"# contigs"},{"values":[5,8,1,1,1,1,2,3,1,1,1,1,2,1],"quality":"Equal","isMain":false,"metricName":"# contigs (>= 1000 bp)"},{"values":[3,4,1,1,1,1,0,2,1,1,1,1,2,1],"quality":"Equal","isMain":false,"metricName":"# contigs (>= 5000 bp)"},{"values":[2,3,1,1,1,1,0,2,1,1,1,1,2,1],"quality":"Equal","isMain":false,"metricName":"# contigs (>= 10000 bp)"},{"values":[1,0,1,1,1,1,0,2,1,1,1,1,2,1],"quality":"Equal","isMain":false,"metricName":"# contigs (>= 25000 bp)"},{"values":[0,0,1,1,1,1,0,0,1,1,1,1,0,1],"quality":"Equal","isMain":false,"metricName":"# contigs (>= 50000 bp)"},{"values":[45903,22588,77854,77361,77445,77307,1459,45686,77417,77273,77273,77273,46897,77307],"quality":"More is better","isMain":true,"metricName":"Largest contig"},{"values":[77294,76689,77854,77361,77445,77307,9515,78793,77417,77273,77273,77273,77250,77307],"quality":"More is better","isMain":true,"metricName":"Total length"},{"values":[77294,76689,77854,77361,77445,77307,2490,78793,77417,77273,77273,77273,77250,77307],"quality":"More is better","isMain":true,"metricName":"Total length (>= 1000 bp)"},{"values":[70369,66985,77854,77361,77445,77307,0,77467,77417,77273,77273,77273,77250,77307],"quality":"More is better","isMain":false,"metricName":"Total length (>= 5000 bp)"},{"values":[62033,58713,77854,77361,77445,77307,0,77467,77417,77273,77273,77273,77250,77307],"quality":"More is better","isMain":true,"metricName":"Total length (>= 10000 bp)"},{"values":[45903,0,77854,77361,77445,77307,0,77467,77417,77273,77273,77273,77250,77307],"quality":"More is better","isMain":false,"metricName":"Total length (>= 25000 bp)"},{"values":[0,0,77854,77361,77445,77307,0,0,77417,77273,77273,77273,0,77307],"quality":"More is better","isMain":true,"metricName":"Total length (>= 50000 bp)"},{"values":[45903,20045,77854,77361,77445,77307,636,45686,77417,77273,77273,77273,46897,77307],"quality":"More is better","isMain":false,"metricName":"N50"},{"values":[16130,16080,77854,77361,77445,77307,521,31781,77417,77273,77273,77273,30353,77307],"quality":"More is better","isMain":false,"metricName":"N75"},{"values":[1,2,1,1,1,1,6,1,1,1,1,1,1,1],"quality":"Less is better","isMain":false,"metricName":"L50"},{"values":[2,3,1,1,1,1,10,2,1,1,1,1,2,1],"quality":"Less is better","isMain":false,"metricName":"L75"},{"values":["42.27","42.25","42.28","42.26","42.27","42.28","42.21","42.30","42.27","42.26","42.26","42.26","42.26","42.28"],"quality":"Equal","isMain":false,"metricName":"GC (%)"}]],["Predicted genes",[]],["Similarity statistics",[{"values":[0,0,0,0,0,0,0,0,0,0,0,0,0,0],"quality":"Equal","isMain":false,"metricName":"# similar correct contigs"},{"values":[0,0,0,0,0,0,0,0,0,0,0,0,0,0],"quality":"Equal","isMain":false,"metricName":"# similar misassembled blocks"}]],["Reference statistics",[{"values":[77554,77554,77554,77554,77554,77554,77554,77554,77554,77554,77554,77554,77554,77554],"quality":"Equal","isMain":false,"metricName":"Reference length"},{"values":[1,1,1,1,1,1,1,1,1,1,1,1,1,1],"quality":"Equal","isMain":false,"metricName":"Reference fragments"},{"values":["42.27","42.27","42.27","42.27","42.27","42.27","42.27","42.27","42.27","42.27","42.27","42.27","42.27","42.27"],"quality":"Equal","isMain":false,"metricName":"Reference GC (%)"}]]],"referenceName":"gi\_167583555\_ref\_NC","date":"18 July 2018, Wednesday, 16:31:48","order":[0,1,2,3,4,5,6,7,8,9,10,11,12,13],"assembliesNames":["ABySS\_127","ABySS\_63","CLC","IDBA\_UD","MEGAHIT","MIRA","Ray\_Meta","SOAPdenovo2","SPAdes","SPAdes\_meta","SPAdes\_sc","SPAdes\_sc\_careful","Velvet","Geneious"]},{"assembliesWithNs":null,"minContig":500,"report":[["Genome statistics",[{"values":["100.000","99.733","99.952","99.627","99.812","100.000","100.000","1.238","99.750","100.000","100.000","100.000","96.756","100.000"],"quality":"More is better","isMain":true,"metricName":"Genome fraction (%)"},{"values":["1.006","1.000","1.000","1.000","1.000","1.016","1.008","1.000","1.000","1.001","1.001","1.001","1.001","1.010"],"quality":"Less is better","isMain":true,"metricName":"Duplication ratio"},{"values":[45625,45503,45603,45455,45539,45625,45625,565,45511,45665,45665,45665,12398,45635],"quality":"More is better","isMain":true,"metricName":"Largest alignment"},{"values":[45625,45503,45603,45455,45539,46227,45625,565,45511,45665,45665,45665,44183,45635],"quality":"More is better","isMain":true,"metricName":"Total aligned length"},{"values":[45877,45503,45625,45455,45539,45764,45993,null,45511,45665,45665,45665,11905,46079],"quality":"More is better","isMain":false,"metricName":"NG50"},{"values":[45877,45503,45625,45455,45539,45764,45993,null,45511,45665,45665,45665,4406,46079],"quality":"More is better","isMain":false,"metricName":"NG75"},{"values":[45625,45503,45603,45455,45539,45625,45625,565,45511,45665,45665,45665,11905,45635],"quality":"More is better","isMain":false,"metricName":"NA50"},{"values":[45625,45503,45603,45455,45539,45625,45625,565,45511,45665,45665,45665,4724,45635],"quality":"More is better","isMain":false,"metricName":"NA75"},{"values":[45625,45503,45603,45455,45539,45625,45625,null,45511,45665,45665,45665,11905,45635],"quality":"More is better","isMain":true,"metricName":"NGA50"},{"values":[45625,45503,45603,45455,45539,45625,45625,null,45511,45665,45665,45665,4406,45635],"quality":"More is better","isMain":false,"metricName":"NGA75"},{"values":[1,1,1,1,1,1,1,null,1,1,1,1,2,1],"quality":"Less is better","isMain":false,"metricName":"LG50"},{"values":[1,1,1,1,1,1,1,null,1,1,1,1,5,1],"quality":"Less is better","isMain":false,"metricName":"LG75"},{"values":[1,1,1,1,1,1,1,1,1,1,1,1,2,1],"quality":"Less is better","isMain":false,"metricName":"LA50"},{"values":[1,1,1,1,1,1,1,1,1,1,1,1,4,1],"quality":"Less is better","isMain":false,"metricName":"LA75"},{"values":[1,1,1,1,1,1,1,null,1,1,1,1,2,1],"quality":"Less is better","isMain":true,"metricName":"LGA50"},{"values":[1,1,1,1,1,1,1,null,1,1,1,1,5,1],"quality":"Less is better","isMain":false,"metricName":"LGA75"}]],["Misassemblies",[{"values":[0,0,0,0,0,0,0,0,0,0,0,0,0,0],"quality":"Less is better","isMain":true,"metricName":"# misassemblies"},{"values":[0,0,0,0,0,0,0,0,0,0,0,0,0,0],"quality":"Less is better","isMain":false,"metricName":" # relocations"},{"values":[0,0,0,0,0,0,0,0,0,0,0,0,0,0],"quality":"Less is better","isMain":false,"metricName":" # translocations"},{"values":[0,0,0,0,0,0,0,0,0,0,0,0,0,0],"quality":"Less is better","isMain":false,"metricName":" # inversions"},{"values":[0,0,0,0,0,0,0,0,0,0,0,0,0,0],"quality":"Less is better","isMain":false,"metricName":"# misassembled contigs"},{"values":[0,0,0,0,0,0,0,0,0,0,0,0,0,0],"quality":"Less is better","isMain":true,"metricName":"Misassembled contigs length"},{"values":[0,0,0,0,0,0,0,0,0,0,0,0,0,0],"quality":"Less is better","isMain":false,"metricName":"# local misassemblies"},{"values":[0,0,0,0,0,0,0,0,0,0,0,0,0,0],"quality":"Less is better","isMain":false,"metricName":"# unaligned mis. contigs"}]],["Unaligned",[{"values":[0,0,0,0,0,0,0,0,0,0,0,0,0,0],"quality":"Less is better","isMain":false,"metricName":"# fully unaligned contigs"},{"values":[0,0,0,0,0,0,0,0,0,0,0,0,0,0],"quality":"Less is better","isMain":false,"metricName":"Fully unaligned length"},{"values":[0,0,0,0,0,0,0,0,0,0,0,0,0,0],"quality":"Less is better","isMain":false,"metricName":"# partially unaligned contigs"},{"values":[0,0,0,0,0,0,0,0,0,0,0,0,0,0],"quality":"Less is better","isMain":false,"metricName":"Partially unaligned length"}]],["Mismatches",[{"values":[0,0,0,0,0,0,0,0,0,0,0,0,0,0],"quality":"Less is better","isMain":false,"metricName":"# mismatches"},{"values":[0,0,0,0,0,1,0,0,0,1,1,1,0,7],"quality":"Less is better","isMain":false,"metricName":"# indels"},{"values":[0,0,0,0,0,38,0,0,0,40,40,40,0,10],"quality":"Less is better","isMain":false,"metricName":"Indels length"},{"values":["0.00","0.00","0.00","0.00","0.00","0.00","0.00","0.00","0.00","0.00","0.00","0.00","0.00","0.00"],"quality":"Less is better","isMain":true,"metricName":"# mismatches per 100 kbp"},{"values":["0.00","0.00","0.00","0.00","0.00","2.19","0.00","0.00","0.00","2.19","2.19","2.19","0.00","15.34"],"quality":"Less is better","isMain":true,"metricName":"# indels per 100 kbp"},{"values":[0,0,0,0,0,0,0,0,0,0,0,0,0,7],"quality":"Less is better","isMain":false,"metricName":" # indels (<= 5 bp)"},{"values":[0,0,0,0,0,1,0,0,0,1,1,1,0,0],"quality":"Less is better","isMain":false,"metricName":" # indels (> 5 bp)"},{"values":[0,0,0,0,0,2,0,0,0,0,0,0,0,493],"quality":"Less is better","isMain":false,"metricName":"# N's"},{"values":["0.00","0.00","0.00","0.00","0.00","4.31","0.00","0.00","0.00","0.00","0.00","0.00","0.00","1069.90"],"quality":"Less is better","isMain":true,"metricName":"# N's per 100 kbp"}]],["Statistics without reference",[{"values":[1,1,1,1,1,2,1,1,1,1,1,1,8,1],"quality":"Equal","isMain":true,"metricName":"# contigs"},{"values":[1,1,1,1,1,1,1,0,1,1,1,1,7,1],"quality":"Equal","isMain":false,"metricName":"# contigs (>= 1000 bp)"},{"values":[1,1,1,1,1,1,1,0,1,1,1,1,2,1],"quality":"Equal","isMain":false,"metricName":"# contigs (>= 5000 bp)"},{"values":[1,1,1,1,1,1,1,0,1,1,1,1,2,1],"quality":"Equal","isMain":false,"metricName":"# contigs (>= 10000 bp)"},{"values":[1,1,1,1,1,1,1,0,1,1,1,1,0,1],"quality":"Equal","isMain":false,"metricName":"# contigs (>= 25000 bp)"},{"values":[0,0,0,0,0,0,0,0,0,0,0,0,0,0],"quality":"Equal","isMain":false,"metricName":"# contigs (>= 50000 bp)"},{"values":[45877,45503,45625,45455,45539,45764,45993,565,45511,45665,45665,45665,12398,46079],"quality":"More is better","isMain":true,"metricName":"Largest contig"},{"values":[45877,45503,45625,45455,45539,46366,45993,565,45511,45665,45665,45665,44183,46079],"quality":"More is better","isMain":true,"metricName":"Total length"},{"values":[45877,45503,45625,45455,45539,45764,45993,0,45511,45665,45665,45665,43355,46079],"quality":"More is better","isMain":true,"metricName":"Total length (>= 1000 bp)"},{"values":[45877,45503,45625,45455,45539,45764,45993,0,45511,45665,45665,45665,24303,46079],"quality":"More is better","isMain":false,"metricName":"Total length (>= 5000 bp)"},{"values":[45877,45503,45625,45455,45539,45764,45993,0,45511,45665,45665,45665,24303,46079],"quality":"More is better","isMain":true,"metricName":"Total length (>= 10000 bp)"},{"values":[45877,45503,45625,45455,45539,45764,45993,0,45511,45665,45665,45665,0,46079],"quality":"More is better","isMain":false,"metricName":"Total length (>= 25000 bp)"},{"values":[0,0,0,0,0,0,0,0,0,0,0,0,0,0],"quality":"More is better","isMain":true,"metricName":"Total length (>= 50000 bp)"},{"values":[45877,45503,45625,45455,45539,45764,45993,565,45511,45665,45665,45665,11905,46079],"quality":"More is better","isMain":false,"metricName":"N50"},{"values":[45877,45503,45625,45455,45539,45764,45993,565,45511,45665,45665,45665,4724,46079],"quality":"More is better","isMain":false,"metricName":"N75"},{"values":[1,1,1,1,1,1,1,1,1,1,1,1,2,1],"quality":"Less is better","isMain":false,"metricName":"L50"},{"values":[1,1,1,1,1,1,1,1,1,1,1,1,4,1],"quality":"Less is better","isMain":false,"metricName":"L75"},{"values":["52.27","52.25","52.25","52.23","52.24","52.29","52.28","48.67","52.23","52.23","52.23","52.23","52.30","52.34"],"quality":"Equal","isMain":false,"metricName":"GC (%)"}]],["Predicted genes",[]],["Similarity statistics",[{"values":[1,1,0,1,1,1,1,0,1,0,0,0,0,1],"quality":"Equal","isMain":false,"metricName":"# similar correct contigs"},{"values":[0,0,0,0,0,0,0,0,0,0,0,0,0,0],"quality":"Equal","isMain":false,"metricName":"# similar misassembled blocks"}]],["Reference statistics",[{"values":[45625,45625,45625,45625,45625,45625,45625,45625,45625,45625,45625,45625,45625,45625],"quality":"Equal","isMain":false,"metricName":"Reference length"},{"values":[1,1,1,1,1,1,1,1,1,1,1,1,1,1],"quality":"Equal","isMain":false,"metricName":"Reference fragments"},{"values":["52.25","52.25","52.25","52.25","52.25","52.25","52.25","52.25","52.25","52.25","52.25","52.25","52.25","52.25"],"quality":"Equal","isMain":false,"metricName":"Reference GC (%)"}]]],"referenceName":"gi\_167600373\_ref\_NC","date":"18 July 2018, Wednesday, 16:32:03","order":[0,1,2,3,4,5,6,7,8,9,10,11,12,13],"assembliesNames":["ABySS\_127","ABySS\_63","CLC","IDBA\_UD","MEGAHIT","MIRA","Ray\_Meta","SOAPdenovo2","SPAdes","SPAdes\_meta","SPAdes\_sc","SPAdes\_sc\_careful","Velvet","Geneious"]},{"assembliesWithNs":null,"minContig":500,"report":[["Genome statistics",[{"values":["98.530","98.204","98.002","98.395","98.609","99.549","76.639","74.050","99.997","100.000","100.000","100.000","97.962"],"quality":"More is better","isMain":true,"metricName":"Genome fraction (%)"},{"values":["1.000","1.001","1.000","1.000","1.000","1.013","1.006","1.027","1.006","1.006","1.005","1.005","1.000"],"quality":"Less is better","isMain":true,"metricName":"Duplication ratio"},{"values":[37867,30782,38461,38615,38699,39075,3348,4988,39371,39363,39345,39336,31456],"quality":"More is better","isMain":true,"metricName":"Largest alignment"},{"values":[38668,38572,38461,38615,38699,39581,30268,29347,39371,39363,39345,39336,38445],"quality":"More is better","isMain":true,"metricName":"Total aligned length"},{"values":[37867,30782,38461,38615,38699,39075,1197,4235,39471,39463,39445,39436,31456],"quality":"More is better","isMain":false,"metricName":"NG50"},{"values":[37867,30782,38461,38615,38699,39075,549,3375,39471,39463,39445,39436,31456],"quality":"More is better","isMain":false,"metricName":"NG75"},{"values":[37867,30782,38461,38615,38699,39075,1487,3465,39371,39363,39345,39336,31456],"quality":"More is better","isMain":false,"metricName":"NA50"},{"values":[37867,30782,38461,38615,38699,39075,914,751,39371,39363,39345,39336,31456],"quality":"More is better","isMain":false,"metricName":"NA75"},{"values":[37867,30782,38461,38615,38699,39075,1197,3465,39371,39363,39345,39336,31456],"quality":"More is better","isMain":true,"metricName":"NGA50"},{"values":[37867,30782,38461,38615,38699,39075,549,null,39371,39363,39345,39336,31456],"quality":"More is better","isMain":false,"metricName":"NGA75"},{"values":[1,1,1,1,1,1,11,4,1,1,1,1,1],"quality":"Less is better","isMain":false,"metricName":"LG50"},{"values":[1,1,1,1,1,1,24,6,1,1,1,1,1],"quality":"Less is better","isMain":false,"metricName":"LG75"},{"values":[1,1,1,1,1,1,8,5,1,1,1,1,1],"quality":"Less is better","isMain":false,"metricName":"LA50"},{"values":[1,1,1,1,1,1,14,9,1,1,1,1,1],"quality":"Less is better","isMain":false,"metricName":"LA75"},{"values":[1,1,1,1,1,1,11,5,1,1,1,1,1],"quality":"Less is better","isMain":true,"metricName":"LGA50"},{"values":[1,1,1,1,1,1,24,null,1,1,1,1,1],"quality":"Less is better","isMain":false,"metricName":"LGA75"}]],["Misassemblies",[{"values":[0,0,0,0,0,0,0,0,0,0,0,0,0],"quality":"Less is better","isMain":true,"metricName":"# misassemblies"},{"values":[0,0,0,0,0,0,0,0,0,0,0,0,0],"quality":"Less is better","isMain":false,"metricName":" # relocations"},{"values":[0,0,0,0,0,0,0,0,0,0,0,0,0],"quality":"Less is better","isMain":false,"metricName":" # translocations"},{"values":[0,0,0,0,0,0,0,0,0,0,0,0,0],"quality":"Less is better","isMain":false,"metricName":" # inversions"},{"values":[0,0,0,0,0,0,0,0,0,0,0,0,0],"quality":"Less is better","isMain":false,"metricName":"# misassembled contigs"},{"values":[0,0,0,0,0,0,0,0,0,0,0,0,0],"quality":"Less is better","isMain":true,"metricName":"Misassembled contigs length"},{"values":[0,0,0,0,0,1,0,0,1,1,1,1,0],"quality":"Less is better","isMain":false,"metricName":"# local misassemblies"},{"values":[0,0,0,0,0,0,0,1,0,0,0,0,0],"quality":"Less is better","isMain":false,"metricName":"# unaligned mis. contigs"}]],["Unaligned",[{"values":[0,0,0,0,0,0,0,0,0,0,0,0,0],"quality":"Less is better","isMain":false,"metricName":"# fully unaligned contigs"},{"values":[0,0,0,0,0,0,0,0,0,0,0,0,0],"quality":"Less is better","isMain":false,"metricName":"Fully unaligned length"},{"values":[0,0,0,0,0,0,0,3,0,0,0,0,0],"quality":"Less is better","isMain":false,"metricName":"# partially unaligned contigs"},{"values":[0,0,0,0,0,0,0,7249,0,0,0,0,0],"quality":"Less is better","isMain":false,"metricName":"Partially unaligned length"}]],["Mismatches",[{"values":[0,0,0,2,2,0,0,2,0,0,0,0,0],"quality":"Less is better","isMain":false,"metricName":"# mismatches"},{"values":[0,0,0,0,0,0,0,13,0,1,1,0,0],"quality":"Less is better","isMain":false,"metricName":"# indels"},{"values":[0,0,0,0,0,0,0,617,0,9,9,0,0],"quality":"Less is better","isMain":false,"metricName":"Indels length"},{"values":["0.00","0.00","0.00","5.18","5.17","0.00","0.00","6.88","0.00","0.00","0.00","0.00","0.00"],"quality":"Less is better","isMain":true,"metricName":"# mismatches per 100 kbp"},{"values":["0.00","0.00","0.00","0.00","0.00","0.00","0.00","44.73","0.00","2.55","2.55","0.00","0.00"],"quality":"Less is better","isMain":true,"metricName":"# indels per 100 kbp"},{"values":[0,0,0,0,0,0,0,0,0,0,0,0,0],"quality":"Less is better","isMain":false,"metricName":" # indels (<= 5 bp)"},{"values":[0,0,0,0,0,0,0,13,0,1,1,0,0],"quality":"Less is better","isMain":false,"metricName":" # indels (> 5 bp)"},{"values":[0,0,1,0,0,1,0,1926,100,110,110,101,0],"quality":"Less is better","isMain":false,"metricName":"# N's"},{"values":["0.00","0.00","2.60","0.00","0.00","2.53","0.00","5190.26","253.35","278.74","278.87","256.11","0.00"],"quality":"Less is better","isMain":true,"metricName":"# N's per 100 kbp"}]],["Statistics without reference",[{"values":[2,3,1,1,1,2,25,10,1,1,1,1,2],"quality":"Equal","isMain":true,"metricName":"# contigs"},{"values":[1,2,1,1,1,1,11,8,1,1,1,1,2],"quality":"Equal","isMain":false,"metricName":"# contigs (>= 1000 bp)"},{"values":[1,2,1,1,1,1,0,3,1,1,1,1,2],"quality":"Equal","isMain":false,"metricName":"# contigs (>= 5000 bp)"},{"values":[1,1,1,1,1,1,0,0,1,1,1,1,1],"quality":"Equal","isMain":false,"metricName":"# contigs (>= 10000 bp)"},{"values":[1,1,1,1,1,1,0,0,1,1,1,1,1],"quality":"Equal","isMain":false,"metricName":"# contigs (>= 25000 bp)"},{"values":[0,0,0,0,0,0,0,0,0,0,0,0,0],"quality":"Equal","isMain":false,"metricName":"# contigs (>= 50000 bp)"},{"values":[37867,30782,38461,38615,38699,39075,3348,8160,39471,39463,39445,39436,31456],"quality":"More is better","isMain":true,"metricName":"Largest contig"},{"values":[38668,38572,38461,38615,38699,39581,30268,37108,39471,39463,39445,39436,38445],"quality":"More is better","isMain":true,"metricName":"Total length"},{"values":[37867,37835,38461,38615,38699,39075,20047,35737,39471,39463,39445,39436,38445],"quality":"More is better","isMain":true,"metricName":"Total length (>= 1000 bp)"},{"values":[37867,37835,38461,38615,38699,39075,0,19495,39471,39463,39445,39436,38445],"quality":"More is better","isMain":false,"metricName":"Total length (>= 5000 bp)"},{"values":[37867,30782,38461,38615,38699,39075,0,0,39471,39463,39445,39436,31456],"quality":"More is better","isMain":true,"metricName":"Total length (>= 10000 bp)"},{"values":[37867,30782,38461,38615,38699,39075,0,0,39471,39463,39445,39436,31456],"quality":"More is better","isMain":false,"metricName":"Total length (>= 25000 bp)"},{"values":[0,0,0,0,0,0,0,0,0,0,0,0,0],"quality":"More is better","isMain":true,"metricName":"Total length (>= 50000 bp)"},{"values":[37867,30782,38461,38615,38699,39075,1487,5177,39471,39463,39445,39436,31456],"quality":"More is better","isMain":false,"metricName":"N50"},{"values":[37867,30782,38461,38615,38699,39075,914,3375,39471,39463,39445,39436,31456],"quality":"More is better","isMain":false,"metricName":"N75"},{"values":[1,1,1,1,1,1,8,3,1,1,1,1,1],"quality":"Less is better","isMain":false,"metricName":"L50"},{"values":[1,1,1,1,1,1,14,6,1,1,1,1,1],"quality":"Less is better","isMain":false,"metricName":"L75"},{"values":["58.90","58.87","58.88","58.90","58.90","59.00","59.12","58.89","59.00","58.96","58.96","58.96","58.90"],"quality":"Equal","isMain":false,"metricName":"GC (%)"}]],["Predicted genes",[]],["Similarity statistics",[{"values":[1,1,1,1,1,0,0,0,1,0,0,0,1],"quality":"Equal","isMain":false,"metricName":"# similar correct contigs"},{"values":[0,0,0,0,0,0,0,0,0,0,0,0,0],"quality":"Equal","isMain":false,"metricName":"# similar misassembled blocks"}]],["Reference statistics",[{"values":[39245,39245,39245,39245,39245,39245,39245,39245,39245,39245,39245,39245,39245],"quality":"Equal","isMain":false,"metricName":"Reference length"},{"values":[1,1,1,1,1,1,1,1,1,1,1,1,1],"quality":"Equal","isMain":false,"metricName":"Reference fragments"},{"values":["58.98","58.98","58.98","58.98","58.98","58.98","58.98","58.98","58.98","58.98","58.98","58.98","58.98"],"quality":"Equal","isMain":false,"metricName":"Reference GC (%)"}]]],"referenceName":"gi\_167832345\_ref\_NC","date":"18 July 2018, Wednesday, 16:32:16","order":[0,1,2,3,4,5,6,7,8,9,10,11,12],"assembliesNames":["ABySS\_127","ABySS\_63","CLC","IDBA\_UD","MEGAHIT","MIRA","Ray\_Meta","SOAPdenovo2","SPAdes","SPAdes\_meta","SPAdes\_sc","SPAdes\_sc\_careful","Velvet"]},{"assembliesWithNs":null,"minContig":500,"report":[["Genome statistics",[{"values":["97.468","80.634","82.542","82.784","83.132","91.299","84.586","1.543","83.781","88.637","79.471","77.805","55.792","100.000"],"quality":"More is better","isMain":true,"metricName":"Genome fraction (%)"},{"values":["1.014","1.023","1.000","1.012","1.012","1.093","1.012","1.000","1.007","1.000","1.003","1.000","1.008","1.055"],"quality":"Less is better","isMain":true,"metricName":"Duplication ratio"},{"values":[30256,17404,26422,12684,18099,8086,26523,630,26397,36189,15435,15061,4884,25696],"quality":"More is better","isMain":true,"metricName":"Largest alignment"},{"values":[40372,33678,33705,34214,34346,40763,34949,630,34465,36189,32561,31771,22936,43091],"quality":"More is better","isMain":true,"metricName":"Total aligned length"},{"values":[30256,8357,26422,3569,9421,6246,26523,null,26397,65947,9361,9361,802,192212],"quality":"More is better","isMain":false,"metricName":"NG50"},{"values":[10116,7034,7283,1382,6687,4089,7268,null,7493,65947,6210,7349,null,192212],"quality":"More is better","isMain":false,"metricName":"NG75"},{"values":[30256,17404,26422,7593,6687,3609,26523,630,26397,36189,9361,9361,1672,null],"quality":"More is better","isMain":false,"metricName":"NA50"},{"values":[10116,8357,26422,3455,1179,1689,26523,630,26397,null,9361,9361,1056,null],"quality":"More is better","isMain":false,"metricName":"NA75"},{"values":[30256,8357,26422,3569,6687,4089,26523,null,26397,36189,9361,9361,802,25696],"quality":"More is better","isMain":true,"metricName":"NGA50"},{"values":[10116,7034,7283,1382,3177,2986,7268,null,7493,36189,6210,7349,null,11502],"quality":"More is better","isMain":false,"metricName":"NGA75"},{"values":[1,2,1,3,2,3,1,null,1,1,2,2,12,1],"quality":"Less is better","isMain":false,"metricName":"LG50"},{"values":[2,3,2,6,3,5,2,null,2,1,3,3,null,1],"quality":"Less is better","isMain":false,"metricName":"LG75"},{"values":[1,1,1,2,2,5,1,1,1,1,2,2,4,null],"quality":"Less is better","isMain":false,"metricName":"LA50"},{"values":[2,2,1,4,5,10,1,1,1,null,2,2,9,null],"quality":"Less is better","isMain":false,"metricName":"LA75"},{"values":[1,2,1,3,2,4,1,null,1,1,2,2,12,1],"quality":"Less is better","isMain":true,"metricName":"LGA50"},{"values":[2,3,2,6,4,7,2,null,2,1,3,3,null,2],"quality":"Less is better","isMain":false,"metricName":"LGA75"}]],["Misassemblies",[{"values":[0,0,0,0,0,0,0,0,0,0,0,0,0,0],"quality":"Less is better","isMain":true,"metricName":"# misassemblies"},{"values":[0,0,0,0,0,0,0,0,0,0,0,0,0,0],"quality":"Less is better","isMain":false,"metricName":" # relocations"},{"values":[0,0,0,0,0,0,0,0,0,0,0,0,0,0],"quality":"Less is better","isMain":false,"metricName":" # translocations"},{"values":[0,0,0,0,0,0,0,0,0,0,0,0,0,0],"quality":"Less is better","isMain":false,"metricName":" # inversions"},{"values":[0,0,0,0,0,0,0,0,0,0,0,0,0,0],"quality":"Less is better","isMain":false,"metricName":"# misassembled contigs"},{"values":[0,0,0,0,0,0,0,0,0,0,0,0,0,0],"quality":"Less is better","isMain":true,"metricName":"Misassembled contigs length"},{"values":[0,0,0,0,0,0,0,0,0,0,0,0,0,0],"quality":"Less is better","isMain":false,"metricName":"# local misassemblies"},{"values":[0,0,0,0,0,0,0,0,0,0,0,0,0,1],"quality":"Less is better","isMain":false,"metricName":"# unaligned mis. contigs"}]],["Unaligned",[{"values":[0,0,0,0,0,0,0,0,0,0,0,0,0,0],"quality":"Less is better","isMain":false,"metricName":"# fully unaligned contigs"},{"values":[0,0,0,0,0,0,0,0,0,0,0,0,0,0],"quality":"Less is better","isMain":false,"metricName":"Fully unaligned length"},{"values":[0,0,0,0,2,1,0,0,0,1,0,0,0,1],"quality":"Less is better","isMain":false,"metricName":"# partially unaligned contigs"},{"values":[0,0,0,0,9360,7917,0,0,0,29758,0,0,0,149121],"quality":"Less is better","isMain":false,"metricName":"Partially unaligned length"}]],["Mismatches",[{"values":[50,110,0,1,2,2,0,0,0,201,24,0,2,5],"quality":"Less is better","isMain":false,"metricName":"# mismatches"},{"values":[1,3,0,0,1,1,0,0,0,9,0,0,0,0],"quality":"Less is better","isMain":false,"metricName":"# indels"},{"values":[3,5,0,0,1,1,0,0,0,15,0,0,0,0],"quality":"Less is better","isMain":false,"metricName":"Indels length"},{"values":["125.63","334.08","0.00","2.96","5.89","5.36","0.00","0.00","0.00","555.34","73.96","0.00","8.78","12.24"],"quality":"Less is better","isMain":true,"metricName":"# mismatches per 100 kbp"},{"values":["2.51","9.11","0.00","0.00","2.95","2.68","0.00","0.00","0.00","24.87","0.00","0.00","0.00","0.00"],"quality":"Less is better","isMain":true,"metricName":"# indels per 100 kbp"},{"values":[1,3,0,0,1,1,0,0,0,9,0,0,0,0],"quality":"Less is better","isMain":false,"metricName":" # indels (<= 5 bp)"},{"values":[0,0,0,0,0,0,0,0,0,0,0,0,0,0],"quality":"Less is better","isMain":false,"metricName":" # indels (> 5 bp)"},{"values":[0,5,0,0,0,0,0,0,0,0,0,0,0,17],"quality":"Less is better","isMain":false,"metricName":"# N's"},{"values":["0.00","14.85","0.00","0.00","0.00","0.00","0.00","0.00","0.00","0.00","0.00","0.00","0.00","8.84"],"quality":"Less is better","isMain":true,"metricName":"# N's per 100 kbp"}]],["Statistics without reference",[{"values":[2,4,2,9,6,13,3,1,3,1,5,3,16,1],"quality":"Equal","isMain":true,"metricName":"# contigs"},{"values":[2,3,2,6,5,11,3,0,2,1,3,3,9,1],"quality":"Equal","isMain":false,"metricName":"# contigs (>= 1000 bp)"},{"values":[2,3,2,2,4,3,2,0,2,1,3,3,0,1],"quality":"Equal","isMain":false,"metricName":"# contigs (>= 5000 bp)"},{"values":[2,1,1,1,1,1,1,0,1,1,1,1,0,1],"quality":"Equal","isMain":false,"metricName":"# contigs (>= 10000 bp)"},{"values":[1,0,1,0,0,0,1,0,1,1,0,0,0,1],"quality":"Equal","isMain":false,"metricName":"# contigs (>= 25000 bp)"},{"values":[0,0,0,0,0,0,0,0,0,1,0,0,0,1],"quality":"Equal","isMain":false,"metricName":"# contigs (>= 50000 bp)"},{"values":[30256,17404,26422,12684,18099,11526,26523,630,26397,65947,15435,15062,4884,192212],"quality":"More is better","isMain":true,"metricName":"Largest contig"},{"values":[40372,33678,33705,34214,43706,48680,34949,630,34465,65947,32561,31772,22974,192212],"quality":"More is better","isMain":true,"metricName":"Total length"},{"values":[40372,32795,33705,31788,42731,47546,34949,0,33890,65947,31006,31772,17939,192212],"quality":"More is better","isMain":true,"metricName":"Total length (>= 1000 bp)"},{"values":[40372,32795,33705,20277,39554,25858,33791,0,33890,65947,31006,31772,0,192212],"quality":"More is better","isMain":false,"metricName":"Total length (>= 5000 bp)"},{"values":[40372,17404,26422,12684,18099,11526,26523,0,26397,65947,15435,15062,0,192212],"quality":"More is better","isMain":true,"metricName":"Total length (>= 10000 bp)"},{"values":[30256,0,26422,0,0,0,26523,0,26397,65947,0,0,0,192212],"quality":"More is better","isMain":false,"metricName":"Total length (>= 25000 bp)"},{"values":[0,0,0,0,0,0,0,0,0,65947,0,0,0,192212],"quality":"More is better","isMain":true,"metricName":"Total length (>= 50000 bp)"},{"values":[30256,17404,26422,7593,9421,6246,26523,630,26397,65947,9361,9361,1672,192212],"quality":"More is better","isMain":false,"metricName":"N50"},{"values":[10116,8357,26422,3455,6687,2990,26523,630,26397,65947,9361,9361,1056,192212],"quality":"More is better","isMain":false,"metricName":"N75"},{"values":[1,1,1,2,2,3,1,1,1,1,2,2,4,1],"quality":"Less is better","isMain":false,"metricName":"L50"},{"values":[2,2,1,4,3,6,1,1,1,1,2,2,9,1],"quality":"Less is better","isMain":false,"metricName":"L75"},{"values":["36.02","36.06","36.08","36.05","36.12","36.00","36.07","36.51","35.96","35.87","35.98","36.13","35.50","35.60"],"quality":"Equal","isMain":false,"metricName":"GC (%)"}]],["Predicted genes",[]],["Similarity statistics",[{"values":[0,0,0,0,0,0,0,0,0,0,0,0,0,0],"quality":"Equal","isMain":false,"metricName":"# similar correct contigs"},{"values":[0,0,0,0,0,0,0,0,0,0,0,0,0,0],"quality":"Equal","isMain":false,"metricName":"# similar misassembled blocks"}]],["Reference statistics",[{"values":[40834,40834,40834,40834,40834,40834,40834,40834,40834,40834,40834,40834,40834,40834],"quality":"Equal","isMain":false,"metricName":"Reference length"},{"values":[1,1,1,1,1,1,1,1,1,1,1,1,1,1],"quality":"Equal","isMain":false,"metricName":"Reference fragments"},{"values":["36.11","36.11","36.11","36.11","36.11","36.11","36.11","36.11","36.11","36.11","36.11","36.11","36.11","36.11"],"quality":"Equal","isMain":false,"metricName":"Reference GC (%)"}]]],"referenceName":"gi\_16798783\_ref\_NC","date":"18 July 2018, Wednesday, 16:32:31","order":[0,1,2,3,4,5,6,7,8,9,10,11,12,13],"assembliesNames":["ABySS\_127","ABySS\_63","CLC","IDBA\_UD","MEGAHIT","MIRA","Ray\_Meta","SOAPdenovo2","SPAdes","SPAdes\_meta","SPAdes\_sc","SPAdes\_sc\_careful","Velvet","Geneious"]},{"assembliesWithNs":null,"minContig":500,"report":[["Genome statistics",[{"values":["93.340","53.710","95.968","76.167","93.484","100.000","46.248","10.010","96.115","54.675","74.830","78.291","22.052","95.335"],"quality":"More is better","isMain":true,"metricName":"Genome fraction (%)"},{"values":["1.025","1.005","1.001","1.011","1.047","1.225","1.015","1.102","1.015","1.002","1.000","1.000","1.004","1.000"],"quality":"Less is better","isMain":true,"metricName":"Duplication ratio"},{"values":[5406,5278,14335,5352,6853,33888,2310,2179,11654,10289,8572,7939,3637,33889],"quality":"More is better","isMain":true,"metricName":"Largest alignment"},{"values":[34015,19176,34155,27358,34798,42806,16692,3602,34670,19433,26597,27825,7866,33889],"quality":"More is better","isMain":true,"metricName":"Total aligned length"},{"values":[3139,526,8278,1117,2957,44926,null,null,9300,706,3339,5358,null,62660],"quality":"More is better","isMain":false,"metricName":"NG50"},{"values":[1946,null,5690,594,1582,44926,null,null,5895,null,666,2514,null,62660],"quality":"More is better","isMain":false,"metricName":"NG75"},{"values":[3541,1239,8278,2545,2957,33888,1248,2179,9300,10289,5417,3410,1605,33889],"quality":"More is better","isMain":false,"metricName":"NA50"},{"values":[2315,724,5690,937,1681,711,688,500,5895,5271,2514,3045,760,null],"quality":"More is better","isMain":false,"metricName":"NA75"},{"values":[3139,526,8278,1117,2957,33888,null,null,9300,706,3045,3339,null,33889],"quality":"More is better","isMain":true,"metricName":"NGA50"},{"values":[1946,null,5690,594,1582,33888,null,null,5895,null,null,655,null,33889],"quality":"More is better","isMain":false,"metricName":"NGA75"},{"values":[5,15,2,7,4,1,null,null,2,4,3,3,null,1],"quality":"Less is better","isMain":false,"metricName":"LG50"},{"values":[8,null,3,17,9,1,null,null,3,null,8,6,null,1],"quality":"Less is better","isMain":false,"metricName":"LG75"},{"values":[4,4,2,4,4,1,6,1,2,1,2,3,2,1],"quality":"Less is better","isMain":false,"metricName":"LA50"},{"values":[7,9,3,9,8,6,11,3,3,2,5,5,3,null],"quality":"Less is better","isMain":false,"metricName":"LA75"},{"values":[5,15,2,7,4,1,null,null,2,4,4,4,null,1],"quality":"Less is better","isMain":true,"metricName":"LGA50"},{"values":[8,null,3,17,9,1,null,null,3,null,null,8,null,1],"quality":"Less is better","isMain":false,"metricName":"LGA75"}]],["Misassemblies",[{"values":[0,0,0,0,0,0,0,0,0,0,0,0,0,0],"quality":"Less is better","isMain":true,"metricName":"# misassemblies"},{"values":[0,0,0,0,0,0,0,0,0,0,0,0,0,0],"quality":"Less is better","isMain":false,"metricName":" # relocations"},{"values":[0,0,0,0,0,0,0,0,0,0,0,0,0,0],"quality":"Less is better","isMain":false,"metricName":" # translocations"},{"values":[0,0,0,0,0,0,0,0,0,0,0,0,0,0],"quality":"Less is better","isMain":false,"metricName":" # inversions"},{"values":[0,0,0,0,0,0,0,0,0,0,0,0,0,0],"quality":"Less is better","isMain":false,"metricName":"# misassembled contigs"},{"values":[0,0,0,0,0,0,0,0,0,0,0,0,0,0],"quality":"Less is better","isMain":true,"metricName":"Misassembled contigs length"},{"values":[0,0,0,0,0,1,0,0,0,0,0,0,0,1],"quality":"Less is better","isMain":false,"metricName":"# local misassemblies"},{"values":[0,0,0,0,0,0,0,0,0,0,0,0,0,0],"quality":"Less is better","isMain":false,"metricName":"# unaligned mis. contigs"}]],["Unaligned",[{"values":[0,0,0,0,0,0,0,0,0,0,0,0,0,0],"quality":"Less is better","isMain":false,"metricName":"# fully unaligned contigs"},{"values":[0,0,0,0,0,0,0,0,0,0,0,0,0,0],"quality":"Less is better","isMain":false,"metricName":"Fully unaligned length"},{"values":[0,0,0,0,0,1,0,0,0,0,1,1,0,1],"quality":"Less is better","isMain":false,"metricName":"# partially unaligned contigs"},{"values":[0,0,0,0,0,11038,0,0,0,0,683,1948,0,28771],"quality":"Less is better","isMain":false,"metricName":"Partially unaligned length"}]],["Mismatches",[{"values":[0,0,1,0,53,178,2,0,0,180,63,49,0,137],"quality":"Less is better","isMain":false,"metricName":"# mismatches"},{"values":[0,0,1,0,2,11,0,2,0,5,3,2,0,4],"quality":"Less is better","isMain":false,"metricName":"# indels"},{"values":[0,0,1,0,2,22,0,93,0,8,6,2,0,4],"quality":"Less is better","isMain":false,"metricName":"Indels length"},{"values":["0.00","0.00","2.93","0.00","159.51","500.80","12.17","0.00","0.00","926.26","236.87","176.09","0.00","404.31"],"quality":"Less is better","isMain":true,"metricName":"# mismatches per 100 kbp"},{"values":["0.00","0.00","2.93","0.00","6.02","30.95","0.00","56.21","0.00","25.73","11.28","7.19","0.00","11.80"],"quality":"Less is better","isMain":true,"metricName":"# indels per 100 kbp"},{"values":[0,0,1,0,2,10,0,0,0,5,3,2,0,4],"quality":"Less is better","isMain":false,"metricName":" # indels (<= 5 bp)"},{"values":[0,0,0,0,0,1,0,2,0,0,0,0,0,0],"quality":"Less is better","isMain":false,"metricName":" # indels (> 5 bp)"},{"values":[0,0,0,0,0,3,0,314,0,0,0,0,0,276],"quality":"Less is better","isMain":false,"metricName":"# N's"},{"values":["0.00","0.00","0.00","0.00","0.00","5.50","0.00","8006.12","0.00","0.00","0.00","0.00","0.00","440.47"],"quality":"Less is better","isMain":true,"metricName":"# N's per 100 kbp"}]],["Statistics without reference",[{"values":[14,17,7,18,17,9,18,4,6,6,9,9,6,1],"quality":"Equal","isMain":true,"metricName":"# contigs"},{"values":[11,5,5,8,12,4,6,1,5,3,6,6,2,1],"quality":"Equal","isMain":false,"metricName":"# contigs (>= 1000 bp)"},{"values":[1,1,3,1,1,1,0,0,4,2,2,3,0,1],"quality":"Equal","isMain":false,"metricName":"# contigs (>= 5000 bp)"},{"values":[0,0,1,0,0,1,0,0,1,1,0,0,0,1],"quality":"Equal","isMain":false,"metricName":"# contigs (>= 10000 bp)"},{"values":[0,0,0,0,0,1,0,0,0,0,0,0,0,1],"quality":"Equal","isMain":false,"metricName":"# contigs (>= 25000 bp)"},{"values":[0,0,0,0,0,0,0,0,0,0,0,0,0,1],"quality":"Equal","isMain":false,"metricName":"# contigs (>= 50000 bp)"},{"values":[5406,5278,14335,5352,6853,44926,2310,2275,11654,10331,9255,7939,3637,62660],"quality":"More is better","isMain":true,"metricName":"Largest contig"},{"values":[34015,19176,34155,27358,34798,54572,16692,3922,34670,19475,27283,29773,7866,62660],"quality":"More is better","isMain":true,"metricName":"Total length"},{"values":[32079,11425,32536,19875,31685,51358,9102,2275,34129,17520,25364,27612,5242,62660],"quality":"More is better","isMain":true,"metricName":"Total length (>= 1000 bp)"},{"values":[5406,5278,28303,5352,6853,44926,0,0,32317,15602,14672,18714,0,62660],"quality":"More is better","isMain":false,"metricName":"Total length (>= 5000 bp)"},{"values":[0,0,14335,0,0,44926,0,0,11654,10331,0,0,0,62660],"quality":"More is better","isMain":true,"metricName":"Total length (>= 10000 bp)"},{"values":[0,0,0,0,0,44926,0,0,0,0,0,0,0,62660],"quality":"More is better","isMain":false,"metricName":"Total length (>= 25000 bp)"},{"values":[0,0,0,0,0,0,0,0,0,0,0,0,0,62660],"quality":"More is better","isMain":true,"metricName":"Total length (>= 50000 bp)"},{"values":[3541,1239,8278,2545,2957,44926,1248,2275,9300,10331,5417,5358,1605,62660],"quality":"More is better","isMain":false,"metricName":"N50"},{"values":[2315,724,5690,937,1681,44926,688,504,5895,5271,3045,3045,760,62660],"quality":"More is better","isMain":false,"metricName":"N75"},{"values":[4,4,2,4,4,1,6,1,2,1,2,3,2,1],"quality":"Less is better","isMain":false,"metricName":"L50"},{"values":[7,9,3,9,8,1,11,3,3,2,4,5,3,1],"quality":"Less is better","isMain":false,"metricName":"L75"},{"values":["39.97","39.93","39.92","40.10","40.06","38.64","40.14","37.25","39.77","39.60","40.26","40.01","38.53","38.97"],"quality":"Equal","isMain":false,"metricName":"GC (%)"}]],["Predicted genes",[]],["Similarity statistics",[{"values":[0,0,0,0,0,0,0,0,0,0,0,0,0,0],"quality":"Equal","isMain":false,"metricName":"# similar correct contigs"},{"values":[0,0,0,0,0,0,0,0,0,0,0,0,0,0],"quality":"Equal","isMain":false,"metricName":"# similar misassembled blocks"}]],["Reference statistics",[{"values":[35543,35543,35543,35543,35543,35543,35543,35543,35543,35543,35543,35543,35543,35543],"quality":"Equal","isMain":false,"metricName":"Reference length"},{"values":[1,1,1,1,1,1,1,1,1,1,1,1,1,1],"quality":"Equal","isMain":false,"metricName":"Reference fragments"},{"values":["39.79","39.79","39.79","39.79","39.79","39.79","39.79","39.79","39.79","39.79","39.79","39.79","39.79","39.79"],"quality":"Equal","isMain":false,"metricName":"Reference GC (%)"}]]],"referenceName":"gi\_168229273\_ref\_NC","date":"18 July 2018, Wednesday, 16:32:46","order":[0,1,2,3,4,5,6,7,8,9,10,11,12,13],"assembliesNames":["ABySS\_127","ABySS\_63","CLC","IDBA\_UD","MEGAHIT","MIRA","Ray\_Meta","SOAPdenovo2","SPAdes","SPAdes\_meta","SPAdes\_sc","SPAdes\_sc\_careful","Velvet","Geneious"]},{"assembliesWithNs":null,"minContig":500,"report":[["Genome statistics",[{"values":["98.686","99.047","99.945","99.899","99.806","99.366","2.774","98.170","99.945","99.945","99.945","99.945","99.856","99.366"],"quality":"More is better","isMain":true,"metricName":"Genome fraction (%)"},{"values":["1.000","1.001","1.005","1.000","1.000","1.047","1.000","1.022","1.000","1.000","1.000","1.000","1.000","1.000"],"quality":"Less is better","isMain":true,"metricName":"Duplication ratio"},{"values":[21495,13380,56982,62274,62216,56675,614,27127,62303,62303,62303,62303,56340,56675],"quality":"More is better","isMain":true,"metricName":"Largest alignment"},{"values":[61518,61784,62603,62274,62216,64829,1729,61374,62303,62303,62303,62303,62247,61942],"quality":"More is better","isMain":true,"metricName":"Total aligned length"},{"values":[16471,7979,56982,62274,62216,56675,null,17420,62303,62303,62303,62303,56340,56675],"quality":"More is better","isMain":false,"metricName":"NG50"},{"values":[13781,5653,56982,62274,62216,56675,null,7146,62303,62303,62303,62303,56340,56675],"quality":"More is better","isMain":false,"metricName":"NG75"},{"values":[16471,7979,56982,62274,62216,56675,589,17317,62303,62303,62303,62303,56340,56675],"quality":"More is better","isMain":false,"metricName":"NA50"},{"values":[13781,6170,56982,62274,62216,56675,526,7048,62303,62303,62303,62303,56340,56675],"quality":"More is better","isMain":false,"metricName":"NA75"},{"values":[16471,7979,56982,62274,62216,56675,null,17317,62303,62303,62303,62303,56340,56675],"quality":"More is better","isMain":true,"metricName":"NGA50"},{"values":[13781,5653,56982,62274,62216,56675,null,7048,62303,62303,62303,62303,56340,56675],"quality":"More is better","isMain":false,"metricName":"NGA75"},{"values":[2,3,1,1,1,1,null,2,1,1,1,1,1,1],"quality":"Less is better","isMain":false,"metricName":"LG50"},{"values":[3,6,1,1,1,1,null,3,1,1,1,1,1,1],"quality":"Less is better","isMain":false,"metricName":"LG75"},{"values":[2,3,1,1,1,1,2,2,1,1,1,1,1,1],"quality":"Less is better","isMain":false,"metricName":"LA50"},{"values":[3,5,1,1,1,1,3,3,1,1,1,1,1,1],"quality":"Less is better","isMain":false,"metricName":"LA75"},{"values":[2,3,1,1,1,1,null,2,1,1,1,1,1,1],"quality":"Less is better","isMain":true,"metricName":"LGA50"},{"values":[3,6,1,1,1,1,null,3,1,1,1,1,1,1],"quality":"Less is better","isMain":false,"metricName":"LGA75"}]],["Misassemblies",[{"values":[0,0,0,0,0,0,0,0,0,0,0,0,0,0],"quality":"Less is better","isMain":true,"metricName":"# misassemblies"},{"values":[0,0,0,0,0,0,0,0,0,0,0,0,0,0],"quality":"Less is better","isMain":false,"metricName":" # relocations"},{"values":[0,0,0,0,0,0,0,0,0,0,0,0,0,0],"quality":"Less is better","isMain":false,"metricName":" # translocations"},{"values":[0,0,0,0,0,0,0,0,0,0,0,0,0,0],"quality":"Less is better","isMain":false,"metricName":" # inversions"},{"values":[0,0,0,0,0,0,0,0,0,0,0,0,0,0],"quality":"Less is better","isMain":false,"metricName":"# misassembled contigs"},{"values":[0,0,0,0,0,0,0,0,0,0,0,0,0,0],"quality":"Less is better","isMain":true,"metricName":"Misassembled contigs length"},{"values":[0,0,0,0,0,0,0,0,0,0,0,0,0,0],"quality":"Less is better","isMain":false,"metricName":"# local misassemblies"},{"values":[0,0,0,0,0,0,0,0,0,0,0,0,0,0],"quality":"Less is better","isMain":false,"metricName":"# unaligned mis. contigs"}]],["Unaligned",[{"values":[0,0,0,0,0,0,0,0,0,0,0,0,0,0],"quality":"Less is better","isMain":false,"metricName":"# fully unaligned contigs"},{"values":[0,0,0,0,0,0,0,0,0,0,0,0,0,0],"quality":"Less is better","isMain":false,"metricName":"Fully unaligned length"},{"values":[0,0,0,0,0,0,0,0,0,0,0,0,0,0],"quality":"Less is better","isMain":false,"metricName":"# partially unaligned contigs"},{"values":[0,0,0,0,0,0,0,0,0,0,0,0,0,0],"quality":"Less is better","isMain":false,"metricName":"Partially unaligned length"}]],["Mismatches",[{"values":[1,0,0,0,0,0,0,0,1,0,0,0,0,0],"quality":"Less is better","isMain":false,"metricName":"# mismatches"},{"values":[0,0,0,0,0,0,0,14,0,0,0,0,0,0],"quality":"Less is better","isMain":false,"metricName":"# indels"},{"values":[0,0,0,0,0,0,0,666,0,0,0,0,0,0],"quality":"Less is better","isMain":false,"metricName":"Indels length"},{"values":["1.63","0.00","0.00","0.00","0.00","0.00","0.00","0.00","1.61","0.00","0.00","0.00","0.00","0.00"],"quality":"Less is better","isMain":true,"metricName":"# mismatches per 100 kbp"},{"values":["0.00","0.00","0.00","0.00","0.00","0.00","0.00","22.88","0.00","0.00","0.00","0.00","0.00","0.00"],"quality":"Less is better","isMain":true,"metricName":"# indels per 100 kbp"},{"values":[0,0,0,0,0,0,0,0,0,0,0,0,0,0],"quality":"Less is better","isMain":false,"metricName":" # indels (<= 5 bp)"},{"values":[0,0,0,0,0,0,0,14,0,0,0,0,0,0],"quality":"Less is better","isMain":false,"metricName":" # indels (> 5 bp)"},{"values":[0,0,0,0,0,6,0,1392,0,0,0,0,0,6],"quality":"Less is better","isMain":false,"metricName":"# N's"},{"values":["0.00","0.00","0.00","0.00","0.00","9.26","0.00","2225.78","0.00","0.00","0.00","0.00","0.00","9.69"],"quality":"Less is better","isMain":true,"metricName":"# N's per 100 kbp"}]],["Statistics without reference",[{"values":[6,10,2,1,1,5,3,5,1,1,1,1,2,2],"quality":"Equal","isMain":true,"metricName":"# contigs"},{"values":[5,9,2,1,1,3,0,5,1,1,1,1,2,2],"quality":"Equal","isMain":false,"metricName":"# contigs (>= 1000 bp)"},{"values":[4,6,2,1,1,2,0,4,1,1,1,1,2,2],"quality":"Equal","isMain":false,"metricName":"# contigs (>= 5000 bp)"},{"values":[3,2,1,1,1,1,0,2,1,1,1,1,1,1],"quality":"Equal","isMain":false,"metricName":"# contigs (>= 10000 bp)"},{"values":[0,0,1,1,1,1,0,1,1,1,1,1,1,1],"quality":"Equal","isMain":false,"metricName":"# contigs (>= 25000 bp)"},{"values":[0,0,1,1,1,1,0,0,1,1,1,1,1,1],"quality":"Equal","isMain":false,"metricName":"# contigs (>= 50000 bp)"},{"values":[21495,13380,56982,62274,62216,56675,614,27877,62303,62303,62303,62303,56340,56675],"quality":"More is better","isMain":true,"metricName":"Largest contig"},{"values":[61518,61784,62603,62274,62216,64829,1729,62540,62303,62303,62303,62303,62247,61942],"quality":"More is better","isMain":true,"metricName":"Total length"},{"values":[60832,61039,62603,62274,62216,63321,0,62540,62303,62303,62303,62303,62247,61942],"quality":"More is better","isMain":true,"metricName":"Total length (>= 1000 bp)"},{"values":[57128,52190,62603,62274,62216,61942,0,58425,62303,62303,62303,62303,62247,61942],"quality":"More is better","isMain":false,"metricName":"Total length (>= 5000 bp)"},{"values":[51747,24828,56982,62274,62216,56675,0,45297,62303,62303,62303,62303,56340,56675],"quality":"More is better","isMain":true,"metricName":"Total length (>= 10000 bp)"},{"values":[0,0,56982,62274,62216,56675,0,27877,62303,62303,62303,62303,56340,56675],"quality":"More is better","isMain":false,"metricName":"Total length (>= 25000 bp)"},{"values":[0,0,56982,62274,62216,56675,0,0,62303,62303,62303,62303,56340,56675],"quality":"More is better","isMain":true,"metricName":"Total length (>= 50000 bp)"},{"values":[16471,7979,56982,62274,62216,56675,589,17420,62303,62303,62303,62303,56340,56675],"quality":"More is better","isMain":false,"metricName":"N50"},{"values":[13781,6170,56982,62274,62216,56675,526,7146,62303,62303,62303,62303,56340,56675],"quality":"More is better","isMain":false,"metricName":"N75"},{"values":[2,3,1,1,1,1,2,2,1,1,1,1,1,1],"quality":"Less is better","isMain":false,"metricName":"L50"},{"values":[3,5,1,1,1,1,3,3,1,1,1,1,1,1],"quality":"Less is better","isMain":false,"metricName":"L75"},{"values":["63.93","63.88","63.94","63.93","63.92","63.95","65.70","63.97","63.94","63.93","63.93","63.93","63.95","63.90"],"quality":"Equal","isMain":false,"metricName":"GC (%)"}]],["Predicted genes",[]],["Similarity statistics",[{"values":[1,0,1,1,1,1,0,0,1,1,1,1,1,1],"quality":"Equal","isMain":false,"metricName":"# similar correct contigs"},{"values":[0,0,0,0,0,0,0,0,0,0,0,0,0,0],"quality":"Equal","isMain":false,"metricName":"# similar misassembled blocks"}]],["Reference statistics",[{"values":[62337,62337,62337,62337,62337,62337,62337,62337,62337,62337,62337,62337,62337,62337],"quality":"Equal","isMain":false,"metricName":"Reference length"},{"values":[1,1,1,1,1,1,1,1,1,1,1,1,1,1],"quality":"Equal","isMain":false,"metricName":"Reference fragments"},{"values":["63.93","63.93","63.93","63.93","63.93","63.93","63.93","63.93","63.93","63.93","63.93","63.93","63.93","63.93"],"quality":"Equal","isMain":false,"metricName":"Reference GC (%)"}]]],"referenceName":"gi\_168495103\_ref\_NC","date":"18 July 2018, Wednesday, 16:33:00","order":[0,1,2,3,4,5,6,7,8,9,10,11,12,13],"assembliesNames":["ABySS\_127","ABySS\_63","CLC","IDBA\_UD","MEGAHIT","MIRA","Ray\_Meta","SOAPdenovo2","SPAdes","SPAdes\_meta","SPAdes\_sc","SPAdes\_sc\_careful","Velvet","Geneious"]},{"assembliesWithNs":null,"minContig":500,"report":[["Genome statistics",[{"values":["7.506","2.672","6.883","5.919","7.813","7.312","7.872","7.872"],"quality":"More is better","isMain":true,"metricName":"Genome fraction (%)"},{"values":["1.015","1.000","1.000","1.000","1.023","1.000","1.000","1.000"],"quality":"Less is better","isMain":true,"metricName":"Duplication ratio"},{"values":[1268,1069,1181,1268,1290,1268,1290,1290],"quality":"More is better","isMain":true,"metricName":"Largest alignment"},{"values":[14263,5077,13079,11248,14847,13896,14959,14959],"quality":"More is better","isMain":true,"metricName":"Total aligned length"},{"values":[798,641,715,766,798,809,809,809],"quality":"More is better","isMain":false,"metricName":"NA50"},{"values":[653,504,644,631,702,702,702,702],"quality":"More is better","isMain":false,"metricName":"NA75"},{"values":[null,null,null,null,null,null,null,null],"quality":"More is better","isMain":true,"metricName":"NGA50"},{"values":[8,4,8,6,8,7,7,7],"quality":"Less is better","isMain":false,"metricName":"LA50"},{"values":[13,7,13,10,13,12,12,12],"quality":"Less is better","isMain":false,"metricName":"LA75"}]],["Misassemblies",[{"values":[0,0,0,0,0,0,0,0],"quality":"Less is better","isMain":true,"metricName":"# misassemblies"},{"values":[0,0,0,0,0,0,0,0],"quality":"Less is better","isMain":false,"metricName":" # relocations"},{"values":[0,0,0,0,0,0,0,0],"quality":"Less is better","isMain":false,"metricName":" # translocations"},{"values":[0,0,0,0,0,0,0,0],"quality":"Less is better","isMain":false,"metricName":" # inversions"},{"values":[0,0,0,0,0,0,0,0],"quality":"Less is better","isMain":false,"metricName":"# misassembled contigs"},{"values":[0,0,0,0,0,0,0,0],"quality":"Less is better","isMain":true,"metricName":"Misassembled contigs length"},{"values":[1,0,1,0,0,0,0,0],"quality":"Less is better","isMain":false,"metricName":"# local misassemblies"},{"values":[0,0,0,0,0,0,0,0],"quality":"Less is better","isMain":false,"metricName":"# unaligned mis. contigs"}]],["Unaligned",[{"values":[0,0,0,0,0,0,0,0],"quality":"Less is better","isMain":false,"metricName":"# fully unaligned contigs"},{"values":[0,0,0,0,0,0,0,0],"quality":"Less is better","isMain":false,"metricName":"Fully unaligned length"},{"values":[0,1,0,0,0,0,0,0],"quality":"Less is better","isMain":false,"metricName":"# partially unaligned contigs"},{"values":[0,745,0,0,0,0,0,0],"quality":"Less is better","isMain":false,"metricName":"Partially unaligned length"}]],["Mismatches",[{"values":[41,10,30,21,35,33,36,31],"quality":"Less is better","isMain":false,"metricName":"# mismatches"},{"values":[3,0,0,0,0,0,0,0],"quality":"Less is better","isMain":false,"metricName":"# indels"},{"values":[56,0,0,0,0,0,0,0],"quality":"Less is better","isMain":false,"metricName":"Indels length"},{"values":["287.46","196.97","229.38","186.70","235.74","237.48","240.66","207.23"],"quality":"Less is better","isMain":true,"metricName":"# mismatches per 100 kbp"},{"values":["21.03","0.00","0.00","0.00","0.00","0.00","0.00","0.00"],"quality":"Less is better","isMain":true,"metricName":"# indels per 100 kbp"},{"values":[2,0,0,0,0,0,0,0],"quality":"Less is better","isMain":false,"metricName":" # indels (<= 5 bp)"},{"values":[1,0,0,0,0,0,0,0],"quality":"Less is better","isMain":false,"metricName":" # indels (> 5 bp)"},{"values":[0,0,0,0,0,0,0,0],"quality":"Less is better","isMain":false,"metricName":"# N's"},{"values":["0.00","0.00","0.00","0.00","0.00","0.00","0.00","0.00"],"quality":"Less is better","isMain":true,"metricName":"# N's per 100 kbp"}]],["Statistics without reference",[{"values":[18,8,18,14,18,17,18,18],"quality":"Equal","isMain":true,"metricName":"# contigs"},{"values":[3,1,1,4,4,3,4,4],"quality":"Equal","isMain":false,"metricName":"# contigs (>= 1000 bp)"},{"values":[0,0,0,0,0,0,0,0],"quality":"Equal","isMain":false,"metricName":"# contigs (>= 5000 bp)"},{"values":[0,0,0,0,0,0,0,0],"quality":"Equal","isMain":false,"metricName":"# contigs (>= 10000 bp)"},{"values":[0,0,0,0,0,0,0,0],"quality":"Equal","isMain":false,"metricName":"# contigs (>= 25000 bp)"},{"values":[0,0,0,0,0,0,0,0],"quality":"Equal","isMain":false,"metricName":"# contigs (>= 50000 bp)"},{"values":[1268,1069,1181,1268,1290,1268,1290,1290],"quality":"More is better","isMain":true,"metricName":"Largest contig"},{"values":[14482,5822,13080,11248,15188,13896,14959,14959],"quality":"More is better","isMain":true,"metricName":"Total length"},{"values":[3669,1069,1181,4766,5037,3669,4959,4959],"quality":"More is better","isMain":true,"metricName":"Total length (>= 1000 bp)"},{"values":[0,0,0,0,0,0,0,0],"quality":"More is better","isMain":false,"metricName":"Total length (>= 5000 bp)"},{"values":[0,0,0,0,0,0,0,0],"quality":"More is better","isMain":true,"metricName":"Total length (>= 10000 bp)"},{"values":[0,0,0,0,0,0,0,0],"quality":"More is better","isMain":false,"metricName":"Total length (>= 25000 bp)"},{"values":[0,0,0,0,0,0,0,0],"quality":"More is better","isMain":true,"metricName":"Total length (>= 50000 bp)"},{"values":[798,956,715,766,798,809,809,809],"quality":"More is better","isMain":false,"metricName":"N50"},{"values":[653,538,644,631,715,702,702,702],"quality":"More is better","isMain":false,"metricName":"N75"},{"values":[8,3,8,6,8,7,7,7],"quality":"Less is better","isMain":false,"metricName":"L50"},{"values":[13,6,13,10,12,12,12,12],"quality":"Less is better","isMain":false,"metricName":"L75"},{"values":["27.50","26.54","27.52","26.75","27.75","26.78","27.24","27.23"],"quality":"Equal","isMain":false,"metricName":"GC (%)"}]],["Predicted genes",[]],["Similarity statistics",[{"values":[0,0,0,0,0,0,0,0],"quality":"Equal","isMain":false,"metricName":"# similar correct contigs"},{"values":[0,0,0,0,0,0,0,0],"quality":"Equal","isMain":false,"metricName":"# similar misassembled blocks"}]],["Reference statistics",[{"values":[190032,190032,190032,190032,190032,190032,190032,190032],"quality":"Equal","isMain":false,"metricName":"Reference length"},{"values":[1,1,1,1,1,1,1,1],"quality":"Equal","isMain":false,"metricName":"Reference fragments"},{"values":["27.99","27.99","27.99","27.99","27.99","27.99","27.99","27.99"],"quality":"Equal","isMain":false,"metricName":"Reference GC (%)"}]]],"referenceName":"gi\_168804017\_ref\_NC","date":"18 July 2018, Wednesday, 16:33:09","order":[0,1,2,3,4,5,6,7],"assembliesNames":["CLC","IDBA\_UD","MEGAHIT","SOAPdenovo2","SPAdes","SPAdes\_meta","SPAdes\_sc","SPAdes\_sc\_careful"]},{"assembliesWithNs":null,"minContig":500,"report":[["Genome statistics",[{"values":["100.000","95.370","100.000","97.367","99.869","100.000","43.110","62.760","99.951","100.000","96.964","98.165","88.458","100.000"],"quality":"More is better","isMain":true,"metricName":"Genome fraction (%)"},{"values":["1.008","1.002","1.000","1.002","1.003","1.035","1.030","1.032","1.000","1.003","1.001","1.004","1.004","1.035"],"quality":"Less is better","isMain":true,"metricName":"Duplication ratio"},{"values":[21757,19957,42723,19650,20082,28552,1428,12863,42702,41400,20646,20646,17664,28552],"quality":"More is better","isMain":true,"metricName":"Largest alignment"},{"values":[43046,40843,42723,41697,42793,44216,18970,27196,42702,42833,41481,42104,37940,44216],"quality":"More is better","isMain":true,"metricName":"Total aligned length"},{"values":[21757,18191,42723,18751,19723,153388,null,9244,42702,41400,18905,18905,5679,179797],"quality":"More is better","isMain":false,"metricName":"NG50"},{"values":[19856,18191,42723,18751,19723,153388,null,3681,42702,41400,18905,18905,2255,179797],"quality":"More is better","isMain":false,"metricName":"NG75"},{"values":[21757,18191,42723,18751,19723,null,779,3924,42702,41400,18905,18905,5679,null],"quality":"More is better","isMain":false,"metricName":"NA50"},{"values":[19856,18191,42723,18751,19723,null,610,null,42702,41400,18905,18905,2294,null],"quality":"More is better","isMain":false,"metricName":"NA75"},{"values":[21757,18191,42723,18751,19723,28552,null,3924,42702,41400,18905,18905,5679,28552],"quality":"More is better","isMain":true,"metricName":"NGA50"},{"values":[19856,18191,42723,18751,19723,15664,null,null,42702,41400,18905,18905,2255,15664],"quality":"More is better","isMain":false,"metricName":"NGA75"},{"values":[1,2,1,2,2,1,null,2,1,1,2,2,2,1],"quality":"Less is better","isMain":false,"metricName":"LG50"},{"values":[2,2,1,2,2,1,null,4,1,1,2,2,5,1],"quality":"Less is better","isMain":false,"metricName":"LG75"},{"values":[1,2,1,2,2,null,10,3,1,1,2,2,2,null],"quality":"Less is better","isMain":false,"metricName":"LA50"},{"values":[2,2,1,2,2,null,17,null,1,1,2,2,4,null],"quality":"Less is better","isMain":false,"metricName":"LA75"},{"values":[1,2,1,2,2,1,null,3,1,1,2,2,2,1],"quality":"Less is better","isMain":true,"metricName":"LGA50"},{"values":[2,2,1,2,2,2,null,null,1,1,2,2,5,2],"quality":"Less is better","isMain":false,"metricName":"LGA75"}]],["Misassemblies",[{"values":[0,0,0,0,0,0,0,0,0,0,0,0,0,0],"quality":"Less is better","isMain":true,"metricName":"# misassemblies"},{"values":[0,0,0,0,0,0,0,0,0,0,0,0,0,0],"quality":"Less is better","isMain":false,"metricName":" # relocations"},{"values":[0,0,0,0,0,0,0,0,0,0,0,0,0,0],"quality":"Less is better","isMain":false,"metricName":" # translocations"},{"values":[0,0,0,0,0,0,0,0,0,0,0,0,0,0],"quality":"Less is better","isMain":false,"metricName":" # inversions"},{"values":[0,0,0,0,0,0,0,0,0,0,0,0,0,0],"quality":"Less is better","isMain":false,"metricName":"# misassembled contigs"},{"values":[0,0,0,0,0,0,0,0,0,0,0,0,0,0],"quality":"Less is better","isMain":true,"metricName":"Misassembled contigs length"},{"values":[0,0,0,0,0,0,0,4,0,0,0,0,0,0],"quality":"Less is better","isMain":false,"metricName":"# local misassemblies"},{"values":[0,0,0,0,0,1,0,0,0,0,0,0,0,1],"quality":"Less is better","isMain":false,"metricName":"# unaligned mis. contigs"}]],["Unaligned",[{"values":[0,0,0,0,0,0,0,0,0,0,0,0,0,0],"quality":"Less is better","isMain":false,"metricName":"# fully unaligned contigs"},{"values":[0,0,0,0,0,0,0,0,0,0,0,0,0,0],"quality":"Less is better","isMain":false,"metricName":"Fully unaligned length"},{"values":[0,0,0,0,0,1,0,6,0,0,0,0,0,1],"quality":"Less is better","isMain":false,"metricName":"# partially unaligned contigs"},{"values":[0,0,0,0,0,109172,0,13817,0,0,0,0,0,135581],"quality":"Less is better","isMain":false,"metricName":"Partially unaligned length"}]],["Mismatches",[{"values":[0,3,0,1,2,84,0,0,0,58,0,3,0,84],"quality":"Less is better","isMain":false,"metricName":"# mismatches"},{"values":[0,0,0,0,0,1,1,23,0,0,0,0,0,1],"quality":"Less is better","isMain":false,"metricName":"# indels"},{"values":[0,0,0,0,0,1,1,1093,0,0,0,0,0,1],"quality":"Less is better","isMain":false,"metricName":"Indels length"},{"values":["0.00","7.36","0.00","2.40","4.69","196.62","0.00","0.00","0.00","135.76","0.00","7.15","0.00","196.62"],"quality":"Less is better","isMain":true,"metricName":"# mismatches per 100 kbp"},{"values":["0.00","0.00","0.00","0.00","0.00","2.34","5.43","85.78","0.00","0.00","0.00","0.00","0.00","2.34"],"quality":"Less is better","isMain":true,"metricName":"# indels per 100 kbp"},{"values":[0,0,0,0,0,1,1,0,0,0,0,0,0,1],"quality":"Less is better","isMain":false,"metricName":" # indels (<= 5 bp)"},{"values":[0,0,0,0,0,0,0,23,0,0,0,0,0,0],"quality":"Less is better","isMain":false,"metricName":" # indels (> 5 bp)"},{"values":[0,0,0,0,0,0,0,5100,0,0,0,0,0,192],"quality":"Less is better","isMain":false,"metricName":"# N's"},{"values":["0.00","0.00","0.00","0.00","0.00","0.00","0.00","12294.19","0.00","0.00","0.00","0.00","0.00","106.79"],"quality":"Less is better","isMain":true,"metricName":"# N's per 100 kbp"}]],["Statistics without reference",[{"values":[3,5,1,5,4,1,25,8,1,2,4,5,9,1],"quality":"Equal","isMain":true,"metricName":"# contigs"},{"values":[3,3,1,4,4,1,4,6,1,2,3,3,8,1],"quality":"Equal","isMain":false,"metricName":"# contigs (>= 1000 bp)"},{"values":[2,2,1,2,2,1,0,3,1,1,2,2,2,1],"quality":"Equal","isMain":false,"metricName":"# contigs (>= 5000 bp)"},{"values":[2,2,1,2,2,1,0,1,1,1,2,2,1,1],"quality":"Equal","isMain":false,"metricName":"# contigs (>= 10000 bp)"},{"values":[0,0,1,0,0,1,0,0,1,1,0,0,0,1],"quality":"Equal","isMain":false,"metricName":"# contigs (>= 25000 bp)"},{"values":[0,0,0,0,0,1,0,0,0,0,0,0,0,1],"quality":"Equal","isMain":false,"metricName":"# contigs (>= 50000 bp)"},{"values":[21757,19957,42723,19650,20082,153388,1428,16697,42702,41400,20646,20646,17664,179797],"quality":"More is better","isMain":true,"metricName":"Largest contig"},{"values":[43046,40843,42723,41697,42793,153388,18970,41483,42702,42833,41481,42104,37940,179797],"quality":"More is better","isMain":true,"metricName":"Total length"},{"values":[43046,39534,42723,41123,42793,153388,5001,39921,42702,42833,40923,40923,37016,179797],"quality":"More is better","isMain":true,"metricName":"Total length (>= 1000 bp)"},{"values":[41613,38148,42723,38401,39805,153388,0,31437,42702,41400,39551,39551,23343,179797],"quality":"More is better","isMain":false,"metricName":"Total length (>= 5000 bp)"},{"values":[41613,38148,42723,38401,39805,153388,0,16697,42702,41400,39551,39551,17664,179797],"quality":"More is better","isMain":true,"metricName":"Total length (>= 10000 bp)"},{"values":[0,0,42723,0,0,153388,0,0,42702,41400,0,0,0,179797],"quality":"More is better","isMain":false,"metricName":"Total length (>= 25000 bp)"},{"values":[0,0,0,0,0,153388,0,0,0,0,0,0,0,179797],"quality":"More is better","isMain":true,"metricName":"Total length (>= 50000 bp)"},{"values":[21757,18191,42723,18751,19723,153388,779,9244,42702,41400,18905,18905,5679,179797],"quality":"More is better","isMain":false,"metricName":"N50"},{"values":[19856,18191,42723,18751,19723,153388,610,5496,42702,41400,18905,18905,2294,179797],"quality":"More is better","isMain":false,"metricName":"N75"},{"values":[1,2,1,2,2,1,10,2,1,1,2,2,2,1],"quality":"Less is better","isMain":false,"metricName":"L50"},{"values":[2,2,1,2,2,1,17,3,1,1,2,2,4,1],"quality":"Less is better","isMain":false,"metricName":"L75"},{"values":["51.60","51.31","51.56","51.34","51.59","51.00","51.55","50.96","51.56","51.54","51.27","51.49","51.11","50.51"],"quality":"Equal","isMain":false,"metricName":"GC (%)"}]],["Predicted genes",[]],["Similarity statistics",[{"values":[1,1,0,1,1,0,0,0,1,1,1,1,1,0],"quality":"Equal","isMain":false,"metricName":"# similar correct contigs"},{"values":[0,0,0,0,0,0,0,0,0,0,0,0,0,0],"quality":"Equal","isMain":false,"metricName":"# similar misassembled blocks"}]],["Reference statistics",[{"values":[42723,42723,42723,42723,42723,42723,42723,42723,42723,42723,42723,42723,42723,42723],"quality":"Equal","isMain":false,"metricName":"Reference length"},{"values":[1,1,1,1,1,1,1,1,1,1,1,1,1,1],"quality":"Equal","isMain":false,"metricName":"Reference fragments"},{"values":["51.56","51.56","51.56","51.56","51.56","51.56","51.56","51.56","51.56","51.56","51.56","51.56","51.56","51.56"],"quality":"Equal","isMain":false,"metricName":"Reference GC (%)"}]]],"referenceName":"gi\_169257155\_ref\_NC","date":"18 July 2018, Wednesday, 16:33:24","order":[0,1,2,3,4,5,6,7,8,9,10,11,12,13],"assembliesNames":["ABySS\_127","ABySS\_63","CLC","IDBA\_UD","MEGAHIT","MIRA","Ray\_Meta","SOAPdenovo2","SPAdes","SPAdes\_meta","SPAdes\_sc","SPAdes\_sc\_careful","Velvet","Geneious"]},{"assembliesWithNs":null,"minContig":500,"report":[["Genome statistics",[{"values":["75.175","85.581","88.988","91.147","97.410","100.000","12.277","55.802","79.431","76.187","93.001","69.879","63.703","100.000"],"quality":"More is better","isMain":true,"metricName":"Genome fraction (%)"},{"values":["1.010","1.010","1.002","1.014","1.029","1.155","1.000","1.040","1.007","1.003","1.006","1.020","1.003","1.153"],"quality":"Less is better","isMain":true,"metricName":"Duplication ratio"},{"values":[15572,14799,23187,16223,19631,46358,1025,7269,23268,32659,15273,15131,5325,46358],"quality":"More is better","isMain":true,"metricName":"Largest alignment"},{"values":[36818,41913,43217,44816,48623,56031,5953,27359,38771,37055,45295,34184,30968,55929],"quality":"More is better","isMain":true,"metricName":"Total aligned length"},{"values":[9308,4593,14235,8749,8610,48674,null,2306,9310,32659,11657,9489,1864,48674],"quality":"More is better","isMain":false,"metricName":"NG50"},{"values":[609,2588,14235,8566,4593,48674,null,null,4409,1822,5033,null,null,48674],"quality":"More is better","isMain":false,"metricName":"NG75"},{"values":[9308,4593,23187,8566,8610,46358,606,1810,23268,32659,11571,9122,3447,46358],"quality":"More is better","isMain":false,"metricName":"NA50"},{"values":[5730,4149,14235,5106,4593,46358,527,731,9310,32659,8460,5033,1995,null],"quality":"More is better","isMain":false,"metricName":"NA75"},{"values":[9308,4593,14235,8749,8610,46358,null,1167,9310,32659,11571,9122,1864,46358],"quality":"More is better","isMain":true,"metricName":"NGA50"},{"values":[609,2588,14235,5106,4593,46358,null,null,4409,1822,5033,null,null,46358],"quality":"More is better","isMain":false,"metricName":"NGA75"},{"values":[2,3,2,2,2,1,null,6,2,1,2,2,8,1],"quality":"Less is better","isMain":false,"metricName":"LG50"},{"values":[6,7,2,3,4,1,null,null,3,3,4,null,null,1],"quality":"Less is better","isMain":false,"metricName":"LG75"},{"values":[2,3,1,3,2,1,4,5,1,1,2,2,4,1],"quality":"Less is better","isMain":false,"metricName":"LA50"},{"values":[3,5,2,4,4,1,7,12,2,1,3,3,7,null],"quality":"Less is better","isMain":false,"metricName":"LA75"},{"values":[2,3,2,2,2,1,null,10,2,1,2,2,8,1],"quality":"Less is better","isMain":true,"metricName":"LGA50"},{"values":[6,7,2,4,4,1,null,null,3,3,4,null,null,1],"quality":"Less is better","isMain":false,"metricName":"LGA75"}]],["Misassemblies",[{"values":[0,0,0,0,0,0,0,0,0,0,0,0,0,0],"quality":"Less is better","isMain":true,"metricName":"# misassemblies"},{"values":[0,0,0,0,0,0,0,0,0,0,0,0,0,0],"quality":"Less is better","isMain":false,"metricName":" # relocations"},{"values":[0,0,0,0,0,0,0,0,0,0,0,0,0,0],"quality":"Less is better","isMain":false,"metricName":" # translocations"},{"values":[0,0,0,0,0,0,0,0,0,0,0,0,0,0],"quality":"Less is better","isMain":false,"metricName":" # inversions"},{"values":[0,0,0,0,0,0,0,0,0,0,0,0,0,0],"quality":"Less is better","isMain":false,"metricName":"# misassembled contigs"},{"values":[0,0,0,0,0,0,0,0,0,0,0,0,0,0],"quality":"Less is better","isMain":true,"metricName":"Misassembled contigs length"},{"values":[0,0,0,0,0,1,0,0,0,0,0,1,0,1],"quality":"Less is better","isMain":false,"metricName":"# local misassemblies"},{"values":[0,0,0,0,0,0,0,0,0,0,0,0,0,1],"quality":"Less is better","isMain":false,"metricName":"# unaligned mis. contigs"}]],["Unaligned",[{"values":[0,0,0,0,0,0,0,0,0,0,0,0,0,0],"quality":"Less is better","isMain":false,"metricName":"# fully unaligned contigs"},{"values":[0,0,0,0,0,0,0,0,0,0,0,0,0,0],"quality":"Less is better","isMain":false,"metricName":"Fully unaligned length"},{"values":[0,0,0,1,0,1,0,7,0,0,0,0,0,2],"quality":"Less is better","isMain":false,"metricName":"# partially unaligned contigs"},{"values":[0,0,0,6689,0,2316,0,6538,0,0,0,0,0,29594],"quality":"Less is better","isMain":false,"metricName":"Partially unaligned length"}]],["Mismatches",[{"values":[0,32,0,38,8,31,15,1,0,46,27,29,6,31],"quality":"Less is better","isMain":false,"metricName":"# mismatches"},{"values":[0,0,0,0,0,2,0,14,0,1,1,1,0,2],"quality":"Less is better","isMain":false,"metricName":"# indels"},{"values":[0,0,0,0,0,67,0,676,0,1,18,12,0,67],"quality":"Less is better","isMain":false,"metricName":"Indels length"},{"values":["0.00","77.11","0.00","85.98","16.94","63.93","251.97","3.70","0.00","124.51","59.87","85.58","19.42","63.93"],"quality":"Less is better","isMain":true,"metricName":"# mismatches per 100 kbp"},{"values":["0.00","0.00","0.00","0.00","0.00","4.12","0.00","51.74","0.00","2.71","2.22","2.95","0.00","4.12"],"quality":"Less is better","isMain":true,"metricName":"# indels per 100 kbp"},{"values":[0,0,0,0,0,0,0,0,0,1,0,0,0,0],"quality":"Less is better","isMain":false,"metricName":" # indels (<= 5 bp)"},{"values":[0,0,0,0,0,2,0,14,0,0,1,1,0,2],"quality":"Less is better","isMain":false,"metricName":" # indels (> 5 bp)"},{"values":[0,0,0,0,0,1,0,2661,0,0,86,0,0,1],"quality":"Less is better","isMain":false,"metricName":"# N's"},{"values":["0.00","0.00","0.00","0.00","0.00","1.71","0.00","7673.67","0.00","0.00","189.51","0.00","0.00","1.17"],"quality":"Less is better","isMain":true,"metricName":"# N's per 100 kbp"}]],["Statistics without reference",[{"values":[6,10,5,7,12,6,9,15,5,3,7,6,13,2],"quality":"Equal","isMain":true,"metricName":"# contigs"},{"values":[5,9,3,6,7,5,1,13,4,3,6,5,10,2],"quality":"Equal","isMain":false,"metricName":"# contigs (>= 1000 bp)"},{"values":[3,2,2,4,3,2,0,1,2,1,4,3,1,2],"quality":"Equal","isMain":false,"metricName":"# contigs (>= 5000 bp)"},{"values":[1,1,2,1,1,1,0,0,1,1,2,1,0,2],"quality":"Equal","isMain":false,"metricName":"# contigs (>= 10000 bp)"},{"values":[0,0,0,0,0,1,0,0,0,1,0,0,0,2],"quality":"Equal","isMain":false,"metricName":"# contigs (>= 25000 bp)"},{"values":[0,0,0,0,0,0,0,0,0,0,0,0,0,0],"quality":"Equal","isMain":false,"metricName":"# contigs (>= 50000 bp)"},{"values":[15572,14799,23187,22912,19631,48674,1025,8178,23268,32659,15273,15131,5325,48674],"quality":"More is better","isMain":true,"metricName":"Largest contig"},{"values":[36818,41913,43217,51505,48625,58347,5953,34677,38771,37055,45381,34551,30968,85523],"quality":"More is better","isMain":true,"metricName":"Total length"},{"values":[36209,41393,42129,50911,45120,57831,1025,33477,38179,37055,44815,34045,28610,85523],"quality":"More is better","isMain":true,"metricName":"Total length (>= 1000 bp)"},{"values":[30610,19846,37422,45333,34001,54108,0,8178,32578,32659,40423,29653,5325,85523],"quality":"More is better","isMain":false,"metricName":"Total length (>= 5000 bp)"},{"values":[15572,14799,37422,22912,19631,48674,0,0,23268,32659,26930,15131,0,85523],"quality":"More is better","isMain":true,"metricName":"Total length (>= 10000 bp)"},{"values":[0,0,0,0,0,48674,0,0,0,32659,0,0,0,85523],"quality":"More is better","isMain":false,"metricName":"Total length (>= 25000 bp)"},{"values":[0,0,0,0,0,0,0,0,0,0,0,0,0,0],"quality":"More is better","isMain":true,"metricName":"Total length (>= 50000 bp)"},{"values":[9308,4593,23187,8749,8610,48674,606,2433,23268,32659,11657,9489,3447,48674],"quality":"More is better","isMain":false,"metricName":"N50"},{"values":[5730,4149,14235,8566,4593,48674,527,1810,9310,32659,8460,5033,1995,36849],"quality":"More is better","isMain":false,"metricName":"N75"},{"values":[2,3,1,2,2,1,4,4,1,1,2,2,4,1],"quality":"Less is better","isMain":false,"metricName":"L50"},{"values":[3,5,2,3,4,1,7,7,2,1,3,3,7,2],"quality":"Less is better","isMain":false,"metricName":"L75"},{"values":["51.20","50.96","51.49","51.24","51.07","50.67","49.35","50.44","51.26","51.42","50.68","51.03","49.84","51.30"],"quality":"Equal","isMain":false,"metricName":"GC (%)"}]],["Predicted genes",[]],["Similarity statistics",[{"values":[0,0,0,0,0,0,0,0,0,0,0,0,0,0],"quality":"Equal","isMain":false,"metricName":"# similar correct contigs"},{"values":[0,0,0,0,0,0,0,0,0,0,0,0,0,0],"quality":"Equal","isMain":false,"metricName":"# similar misassembled blocks"}]],["Reference statistics",[{"values":[48491,48491,48491,48491,48491,48491,48491,48491,48491,48491,48491,48491,48491,48491],"quality":"Equal","isMain":false,"metricName":"Reference length"},{"values":[1,1,1,1,1,1,1,1,1,1,1,1,1,1],"quality":"Equal","isMain":false,"metricName":"Reference fragments"},{"values":["51.10","51.10","51.10","51.10","51.10","51.10","51.10","51.10","51.10","51.10","51.10","51.10","51.10","51.10"],"quality":"Equal","isMain":false,"metricName":"Reference GC (%)"}]]],"referenceName":"gi\_169257208\_ref\_NC","date":"18 July 2018, Wednesday, 16:33:39","order":[0,1,2,3,4,5,6,7,8,9,10,11,12,13],"assembliesNames":["ABySS\_127","ABySS\_63","CLC","IDBA\_UD","MEGAHIT","MIRA","Ray\_Meta","SOAPdenovo2","SPAdes","SPAdes\_meta","SPAdes\_sc","SPAdes\_sc\_careful","Velvet","Geneious"]},{"assembliesWithNs":null,"minContig":500,"report":[["Genome statistics",[{"values":["96.994","73.501","88.309","72.821","75.065","100.000","40.212","42.736","97.705","96.189","68.331","89.193","78.209","100.000"],"quality":"More is better","isMain":true,"metricName":"Genome fraction (%)"},{"values":["1.013","1.002","1.001","1.018","1.005","1.170","1.054","1.092","1.011","1.004","1.007","1.011","1.005","1.158"],"quality":"Less is better","isMain":true,"metricName":"Duplication ratio"},{"values":[12667,11894,21673,12984,12454,31431,4378,8711,20289,30056,11880,12795,6050,34519],"quality":"More is better","isMain":true,"metricName":"Largest alignment"},{"values":[45030,33755,40503,33980,34565,53629,19427,20087,45296,44203,31543,40996,36044,53080],"quality":"More is better","isMain":true,"metricName":"Total aligned length"},{"values":[8221,6853,8907,12984,7025,48674,null,6731,10246,30056,6839,11657,3925,48674],"quality":"More is better","isMain":false,"metricName":"NG50"},{"values":[7733,null,5214,12984,721,48674,null,null,8223,11657,null,6839,864,48674],"quality":"More is better","isMain":false,"metricName":"NG75"},{"values":[8221,6975,21673,7140,7899,16080,1699,1522,10246,30056,6961,11319,4050,16080],"quality":"More is better","isMain":false,"metricName":"NA50"},{"values":[7733,6853,8907,null,7025,null,825,null,8223,11571,6839,7467,2520,null],"quality":"More is better","isMain":false,"metricName":"NA75"},{"values":[8221,6853,8907,7140,7025,31431,null,null,10246,30056,6839,11319,3566,34519],"quality":"More is better","isMain":true,"metricName":"NGA50"},{"values":[7733,null,5214,null,721,16080,null,null,8223,11571,null,6839,864,34519],"quality":"More is better","isMain":false,"metricName":"NGA75"},{"values":[3,3,2,2,3,1,null,3,2,1,3,2,5,1],"quality":"Less is better","isMain":false,"metricName":"LG50"},{"values":[4,null,3,2,7,1,null,null,3,2,null,4,10,1],"quality":"Less is better","isMain":false,"metricName":"LG75"},{"values":[3,2,1,3,2,2,4,4,2,1,2,2,4,2],"quality":"Less is better","isMain":false,"metricName":"LA50"},{"values":[4,3,2,null,3,null,9,null,3,2,3,3,7,null],"quality":"Less is better","isMain":false,"metricName":"LA75"},{"values":[3,3,2,3,3,1,null,null,2,1,3,2,6,1],"quality":"Less is better","isMain":true,"metricName":"LGA50"},{"values":[4,null,3,null,7,2,null,null,3,2,null,4,10,1],"quality":"Less is better","isMain":false,"metricName":"LGA75"}]],["Misassemblies",[{"values":[0,0,0,0,0,0,0,0,0,0,0,0,0,0],"quality":"Less is better","isMain":true,"metricName":"# misassemblies"},{"values":[0,0,0,0,0,0,0,0,0,0,0,0,0,0],"quality":"Less is better","isMain":false,"metricName":" # relocations"},{"values":[0,0,0,0,0,0,0,0,0,0,0,0,0,0],"quality":"Less is better","isMain":false,"metricName":" # translocations"},{"values":[0,0,0,0,0,0,0,0,0,0,0,0,0,0],"quality":"Less is better","isMain":false,"metricName":" # inversions"},{"values":[0,0,0,0,0,0,0,0,0,0,0,0,0,0],"quality":"Less is better","isMain":false,"metricName":"# misassembled contigs"},{"values":[0,0,0,0,0,0,0,0,0,0,0,0,0,0],"quality":"Less is better","isMain":true,"metricName":"Misassembled contigs length"},{"values":[0,0,0,0,0,0,0,2,0,0,0,0,0,1],"quality":"Less is better","isMain":false,"metricName":"# local misassemblies"},{"values":[0,0,0,0,0,1,0,2,0,0,0,0,0,1],"quality":"Less is better","isMain":false,"metricName":"# unaligned mis. contigs"}]],["Unaligned",[{"values":[0,0,0,0,0,0,0,0,0,0,0,0,0,0],"quality":"Less is better","isMain":false,"metricName":"# fully unaligned contigs"},{"values":[0,0,0,0,0,0,0,0,0,0,0,0,0,0],"quality":"Less is better","isMain":false,"metricName":"Fully unaligned length"},{"values":[0,0,0,1,0,1,0,3,0,0,0,0,0,2],"quality":"Less is better","isMain":false,"metricName":"# partially unaligned contigs"},{"values":[0,0,0,14031,0,30113,0,10109,0,0,0,0,0,32443],"quality":"Less is better","isMain":false,"metricName":"Partially unaligned length"}]],["Mismatches",[{"values":[2,2,3,74,12,26,21,0,1,32,8,12,9,26],"quality":"Less is better","isMain":false,"metricName":"# mismatches"},{"values":[0,0,0,2,0,0,0,14,0,1,0,1,0,0],"quality":"Less is better","isMain":false,"metricName":"# indels"},{"values":[0,0,0,2,0,0,0,671,0,51,0,51,0,0],"quality":"Less is better","isMain":false,"metricName":"Indels length"},{"values":["4.50","5.94","7.41","221.68","34.87","56.72","113.93","0.00","2.23","72.57","25.54","29.35","25.10","56.72"],"quality":"Less is better","isMain":true,"metricName":"# mismatches per 100 kbp"},{"values":["0.00","0.00","0.00","5.99","0.00","0.00","0.00","71.47","0.00","2.27","0.00","2.45","0.00","0.00"],"quality":"Less is better","isMain":true,"metricName":"# indels per 100 kbp"},{"values":[0,0,0,2,0,0,0,0,0,0,0,0,0,0],"quality":"Less is better","isMain":false,"metricName":" # indels (<= 5 bp)"},{"values":[0,0,0,0,0,0,0,14,0,1,0,1,0,0],"quality":"Less is better","isMain":false,"metricName":" # indels (> 5 bp)"},{"values":[0,0,0,0,0,0,0,4167,0,86,0,338,0,1],"quality":"Less is better","isMain":false,"metricName":"# N's"},{"values":["0.00","0.00","0.00","0.00","0.00","0.00","0.00","13231.93","0.00","194.18","0.00","817.73","0.00","1.17"],"quality":"Less is better","isMain":true,"metricName":"# N's per 100 kbp"}]],["Statistics without reference",[{"values":[7,6,4,6,7,5,15,9,6,3,8,5,12,2],"quality":"Equal","isMain":true,"metricName":"# contigs"},{"values":[7,6,4,5,5,4,6,5,5,3,5,5,9,2],"quality":"Equal","isMain":false,"metricName":"# contigs (>= 1000 bp)"},{"values":[4,3,3,3,3,2,0,3,3,2,3,4,1,2],"quality":"Equal","isMain":false,"metricName":"# contigs (>= 5000 bp)"},{"values":[1,1,1,2,1,2,0,1,2,2,1,2,0,2],"quality":"Equal","isMain":false,"metricName":"# contigs (>= 10000 bp)"},{"values":[0,0,0,0,0,2,0,0,0,1,0,0,0,2],"quality":"Equal","isMain":false,"metricName":"# contigs (>= 25000 bp)"},{"values":[0,0,0,0,0,0,0,0,0,0,0,0,0,0],"quality":"Equal","isMain":false,"metricName":"# contigs (>= 50000 bp)"},{"values":[12667,11894,21673,22912,12454,48674,4378,11657,20289,30056,11880,12795,6050,48674],"quality":"More is better","isMain":true,"metricName":"Largest contig"},{"values":[45030,33755,40503,48011,34565,83742,19427,31492,45296,44289,31543,41334,36046,85523],"quality":"More is better","isMain":true,"metricName":"Total length"},{"values":[45030,33755,40503,47237,32929,83027,12794,28198,44519,44289,29386,41334,33743,85523],"quality":"More is better","isMain":true,"metricName":"Total length (>= 1000 bp)"},{"values":[37270,25722,35794,43036,27378,80105,0,25658,38758,41713,25680,38758,6050,85523],"quality":"More is better","isMain":false,"metricName":"Total length (>= 5000 bp)"},{"values":[12667,11894,21673,35896,12454,80105,0,11657,30535,41713,11880,24452,0,85523],"quality":"More is better","isMain":true,"metricName":"Total length (>= 10000 bp)"},{"values":[0,0,0,0,0,80105,0,0,0,30056,0,0,0,85523],"quality":"More is better","isMain":false,"metricName":"Total length (>= 25000 bp)"},{"values":[0,0,0,0,0,0,0,0,0,0,0,0,0,0],"quality":"More is better","isMain":true,"metricName":"Total length (>= 50000 bp)"},{"values":[8221,6975,21673,12984,7899,48674,1699,7270,10246,30056,6961,11657,4050,48674],"quality":"More is better","isMain":false,"metricName":"N50"},{"values":[7733,6853,8907,7140,7025,31431,825,6731,8223,11657,6839,7467,2520,36849],"quality":"More is better","isMain":false,"metricName":"N75"},{"values":[3,2,1,2,2,1,4,2,2,1,2,2,4,1],"quality":"Less is better","isMain":false,"metricName":"L50"},{"values":[4,3,2,3,3,2,9,3,3,2,3,3,7,2],"quality":"Less is better","isMain":false,"metricName":"L75"},{"values":["51.14","50.86","50.99","51.96","51.37","51.39","50.88","51.07","51.30","51.29","51.68","50.95","50.93","51.30"],"quality":"Equal","isMain":false,"metricName":"GC (%)"}]],["Predicted genes",[]],["Similarity statistics",[{"values":[0,0,0,0,0,0,0,0,0,0,0,0,0,0],"quality":"Equal","isMain":false,"metricName":"# similar correct contigs"},{"values":[0,0,0,0,0,0,0,0,0,0,0,0,0,0],"quality":"Equal","isMain":false,"metricName":"# similar misassembled blocks"}]],["Reference statistics",[{"values":[45840,45840,45840,45840,45840,45840,45840,45840,45840,45840,45840,45840,45840,45840],"quality":"Equal","isMain":false,"metricName":"Reference length"},{"values":[1,1,1,1,1,1,1,1,1,1,1,1,1,1],"quality":"Equal","isMain":false,"metricName":"Reference fragments"},{"values":["51.10","51.10","51.10","51.10","51.10","51.10","51.10","51.10","51.10","51.10","51.10","51.10","51.10","51.10"],"quality":"Equal","isMain":false,"metricName":"Reference GC (%)"}]]],"referenceName":"gi\_169257267\_ref\_NC","date":"18 July 2018, Wednesday, 16:33:53","order":[0,1,2,3,4,5,6,7,8,9,10,11,12,13],"assembliesNames":["ABySS\_127","ABySS\_63","CLC","IDBA\_UD","MEGAHIT","MIRA","Ray\_Meta","SOAPdenovo2","SPAdes","SPAdes\_meta","SPAdes\_sc","SPAdes\_sc\_careful","Velvet","Geneious"]},{"assembliesWithNs":null,"minContig":500,"report":[["Genome statistics",[{"values":["100.000","100.000","100.000","98.755","98.960","100.000","100.000","100.000","100.000","100.000","100.000","62.501","100.000"],"quality":"More is better","isMain":true,"metricName":"Genome fraction (%)"},{"values":["1.017","1.092","1.000","1.013","1.004","1.030","1.013","1.006","1.003","1.005","1.003","1.002","1.027"],"quality":"Less is better","isMain":true,"metricName":"Duplication ratio"},{"values":[41588,37231,40881,12567,26524,27853,41404,40428,40284,36115,40284,4566,40154],"quality":"More is better","isMain":true,"metricName":"Largest alignment"},{"values":[41588,44657,40881,40897,40603,42100,41404,41135,40991,41101,40991,25601,41991],"quality":"More is better","isMain":true,"metricName":"Total aligned length"},{"values":[41588,37231,40881,10149,26524,27853,41404,40428,40284,36115,40284,787,78993],"quality":"More is better","isMain":false,"metricName":"NG50"},{"values":[41588,37231,40881,4263,10098,8636,41404,40428,40284,36115,40284,null,78993],"quality":"More is better","isMain":false,"metricName":"NG75"},{"values":[41588,37231,40881,10149,26524,27853,41404,40428,40284,36115,40284,1969,40154],"quality":"More is better","isMain":false,"metricName":"NA50"},{"values":[41588,37231,40881,4263,10098,8636,41404,40428,40284,36115,40284,826,null],"quality":"More is better","isMain":false,"metricName":"NA75"},{"values":[41588,37231,40881,10149,26524,27853,41404,40428,40284,36115,40284,787,40154],"quality":"More is better","isMain":true,"metricName":"NGA50"},{"values":[41588,37231,40881,4263,10098,8636,41404,40428,40284,36115,40284,null,40154],"quality":"More is better","isMain":false,"metricName":"NGA75"},{"values":[1,1,1,2,1,1,1,1,1,1,1,11,1],"quality":"Less is better","isMain":false,"metricName":"LG50"},{"values":[1,1,1,4,2,2,1,1,1,1,1,null,1],"quality":"Less is better","isMain":false,"metricName":"LG75"},{"values":[1,1,1,2,1,1,1,1,1,1,1,4,1],"quality":"Less is better","isMain":false,"metricName":"LA50"},{"values":[1,1,1,4,2,2,1,1,1,1,1,10,null],"quality":"Less is better","isMain":false,"metricName":"LA75"},{"values":[1,1,1,2,1,1,1,1,1,1,1,11,1],"quality":"Less is better","isMain":true,"metricName":"LGA50"},{"values":[1,1,1,4,2,2,1,1,1,1,1,null,1],"quality":"Less is better","isMain":false,"metricName":"LGA75"}]],["Misassemblies",[{"values":[0,0,0,0,0,0,0,0,0,0,0,0,1],"quality":"Less is better","isMain":true,"metricName":"# misassemblies"},{"values":[0,0,0,0,0,0,0,0,0,0,0,0,1],"quality":"Less is better","isMain":false,"metricName":" # relocations"},{"values":[0,0,0,0,0,0,0,0,0,0,0,0,0],"quality":"Less is better","isMain":false,"metricName":" # translocations"},{"values":[0,0,0,0,0,0,0,0,0,0,0,0,0],"quality":"Less is better","isMain":false,"metricName":" # inversions"},{"values":[0,0,0,0,0,0,0,0,0,0,0,0,1],"quality":"Less is better","isMain":false,"metricName":"# misassembled contigs"},{"values":[0,0,0,0,0,0,0,0,0,0,0,0,78993],"quality":"Less is better","isMain":true,"metricName":"Misassembled contigs length"},{"values":[1,0,0,0,0,0,1,0,0,0,0,0,0],"quality":"Less is better","isMain":false,"metricName":"# local misassemblies"},{"values":[0,0,0,0,0,0,0,0,0,0,0,0,0],"quality":"Less is better","isMain":false,"metricName":"# unaligned mis. contigs"}]],["Unaligned",[{"values":[0,0,0,0,0,0,0,0,0,0,0,0,0],"quality":"Less is better","isMain":false,"metricName":"# fully unaligned contigs"},{"values":[0,0,0,0,0,0,0,0,0,0,0,0,0],"quality":"Less is better","isMain":false,"metricName":"Fully unaligned length"},{"values":[0,0,0,0,0,0,0,0,0,0,0,0,1],"quality":"Less is better","isMain":false,"metricName":"# partially unaligned contigs"},{"values":[0,0,0,0,0,0,0,0,0,0,0,0,37002],"quality":"Less is better","isMain":false,"metricName":"Partially unaligned length"}]],["Mismatches",[{"values":[0,0,0,0,1,1,7,0,1,1,0,0,134],"quality":"Less is better","isMain":false,"metricName":"# mismatches"},{"values":[0,0,0,0,0,0,0,0,0,0,0,0,11],"quality":"Less is better","isMain":false,"metricName":"# indels"},{"values":[0,0,0,0,0,0,0,0,0,0,0,0,18],"quality":"Less is better","isMain":false,"metricName":"Indels length"},{"values":["0.00","0.00","0.00","0.00","2.47","2.45","17.12","0.00","2.45","2.45","0.00","0.00","327.78"],"quality":"Less is better","isMain":true,"metricName":"# mismatches per 100 kbp"},{"values":["0.00","0.00","0.00","0.00","0.00","0.00","0.00","0.00","0.00","0.00","0.00","0.00","26.91"],"quality":"Less is better","isMain":true,"metricName":"# indels per 100 kbp"},{"values":[0,0,0,0,0,0,0,0,0,0,0,0,11],"quality":"Less is better","isMain":false,"metricName":" # indels (<= 5 bp)"},{"values":[0,0,0,0,0,0,0,0,0,0,0,0,0],"quality":"Less is better","isMain":false,"metricName":" # indels (> 5 bp)"},{"values":[0,1,0,0,0,0,0,0,0,0,0,0,31],"quality":"Less is better","isMain":false,"metricName":"# N's"},{"values":["0.00","2.24","0.00","0.00","0.00","0.00","0.00","0.00","0.00","0.00","0.00","0.00","39.24"],"quality":"Less is better","isMain":true,"metricName":"# N's per 100 kbp"}]],["Statistics without reference",[{"values":[1,2,1,7,3,4,1,2,2,4,2,19,1],"quality":"Equal","isMain":true,"metricName":"# contigs"},{"values":[1,2,1,7,3,4,1,1,1,2,1,8,1],"quality":"Equal","isMain":false,"metricName":"# contigs (>= 1000 bp)"},{"values":[1,2,1,3,2,2,1,1,1,1,1,0,1],"quality":"Equal","isMain":false,"metricName":"# contigs (>= 5000 bp)"},{"values":[1,1,1,2,2,1,1,1,1,1,1,0,1],"quality":"Equal","isMain":false,"metricName":"# contigs (>= 10000 bp)"},{"values":[1,1,1,0,1,1,1,1,1,1,1,0,1],"quality":"Equal","isMain":false,"metricName":"# contigs (>= 25000 bp)"},{"values":[0,0,0,0,0,0,0,0,0,0,0,0,1],"quality":"Equal","isMain":false,"metricName":"# contigs (>= 50000 bp)"},{"values":[41588,37231,40881,12567,26524,27853,41404,40428,40284,36115,40284,4566,78993],"quality":"More is better","isMain":true,"metricName":"Largest contig"},{"values":[41588,44657,40881,40897,40603,42101,41404,41135,40991,41101,40991,25601,78993],"quality":"More is better","isMain":true,"metricName":"Total length"},{"values":[41588,44657,40881,40897,40603,42101,41404,40428,40284,39875,40284,18085,78993],"quality":"More is better","isMain":true,"metricName":"Total length (>= 1000 bp)"},{"values":[41588,44657,40881,28212,36622,36489,41404,40428,40284,36115,40284,0,78993],"quality":"More is better","isMain":false,"metricName":"Total length (>= 5000 bp)"},{"values":[41588,37231,40881,22716,36622,27853,41404,40428,40284,36115,40284,0,78993],"quality":"More is better","isMain":true,"metricName":"Total length (>= 10000 bp)"},{"values":[41588,37231,40881,0,26524,27853,41404,40428,40284,36115,40284,0,78993],"quality":"More is better","isMain":false,"metricName":"Total length (>= 25000 bp)"},{"values":[0,0,0,0,0,0,0,0,0,0,0,0,78993],"quality":"More is better","isMain":true,"metricName":"Total length (>= 50000 bp)"},{"values":[41588,37231,40881,10149,26524,27853,41404,40428,40284,36115,40284,1969,78993],"quality":"More is better","isMain":false,"metricName":"N50"},{"values":[41588,37231,40881,4263,10098,8636,41404,40428,40284,36115,40284,826,78993],"quality":"More is better","isMain":false,"metricName":"N75"},{"values":[1,1,1,2,1,1,1,1,1,1,1,4,1],"quality":"Less is better","isMain":false,"metricName":"L50"},{"values":[1,1,1,4,2,2,1,1,1,1,1,10,1],"quality":"Less is better","isMain":false,"metricName":"L75"},{"values":["36.40","36.46","36.40","36.40","36.39","36.48","36.40","36.41","36.41","36.41","36.41","35.58","36.63"],"quality":"Equal","isMain":false,"metricName":"GC (%)"}]],["Predicted genes",[]],["Similarity statistics",[{"values":[0,0,0,0,0,0,0,0,0,0,0,0,0],"quality":"Equal","isMain":false,"metricName":"# similar correct contigs"},{"values":[0,0,0,0,0,0,0,0,0,0,0,0,0],"quality":"Equal","isMain":false,"metricName":"# similar misassembled blocks"}]],["Reference statistics",[{"values":[40881,40881,40881,40881,40881,40881,40881,40881,40881,40881,40881,40881,40881],"quality":"Equal","isMain":false,"metricName":"Reference length"},{"values":[1,1,1,1,1,1,1,1,1,1,1,1,1],"quality":"Equal","isMain":false,"metricName":"Reference fragments"},{"values":["36.40","36.40","36.40","36.40","36.40","36.40","36.40","36.40","36.40","36.40","36.40","36.40","36.40"],"quality":"Equal","isMain":false,"metricName":"Reference GC (%)"}]]],"referenceName":"gi\_169794178\_ref\_NC","date":"18 July 2018, Wednesday, 16:34:06","order":[0,1,2,3,4,5,6,7,8,9,10,11,12],"assembliesNames":["ABySS\_127","ABySS\_63","CLC","IDBA\_UD","MEGAHIT","MIRA","Ray\_Meta","SPAdes","SPAdes\_meta","SPAdes\_sc","SPAdes\_sc\_careful","Velvet","Geneious"]},{"assembliesWithNs":null,"minContig":500,"report":[["Genome statistics",[{"values":["99.914","99.047","99.893","100.000","100.000","100.000","3.321","83.792","100.000","100.000","96.895","96.895","96.916"],"quality":"More is better","isMain":true,"metricName":"Genome fraction (%)"},{"values":["1.003","1.002","1.000","1.031","1.004","1.004","1.000","1.012","1.004","1.002","1.000","1.000","1.000"],"quality":"Less is better","isMain":true,"metricName":"Duplication ratio"},{"values":[20241,24875,33657,31149,33834,33837,581,26774,33820,33748,30951,30951,31011],"quality":"More is better","isMain":true,"metricName":"Largest alignment"},{"values":[33765,33440,33657,34325,33834,33837,1119,28562,33820,33748,32647,32647,32654],"quality":"More is better","isMain":true,"metricName":"Total aligned length"},{"values":[20241,24875,33657,31149,33834,33837,null,30232,33820,33748,30951,30951,31011],"quality":"More is better","isMain":false,"metricName":"NG50"},{"values":[7350,4425,33657,31149,33834,33837,null,30232,33820,33748,30951,30951,31011],"quality":"More is better","isMain":false,"metricName":"NG75"},{"values":[20241,24875,33657,31149,33834,33837,581,26774,33820,33748,30951,30951,31011],"quality":"More is better","isMain":false,"metricName":"NA50"},{"values":[7350,4425,33657,31149,33834,33837,538,26774,33820,33748,30951,30951,31011],"quality":"More is better","isMain":false,"metricName":"NA75"},{"values":[20241,24875,33657,31149,33834,33837,null,26774,33820,33748,30951,30951,31011],"quality":"More is better","isMain":true,"metricName":"NGA50"},{"values":[7350,4425,33657,31149,33834,33837,null,26774,33820,33748,30951,30951,31011],"quality":"More is better","isMain":false,"metricName":"NGA75"},{"values":[1,1,1,1,1,1,null,1,1,1,1,1,1],"quality":"Less is better","isMain":false,"metricName":"LG50"},{"values":[2,2,1,1,1,1,null,1,1,1,1,1,1],"quality":"Less is better","isMain":false,"metricName":"LG75"},{"values":[1,1,1,1,1,1,1,1,1,1,1,1,1],"quality":"Less is better","isMain":false,"metricName":"LA50"},{"values":[2,2,1,1,1,1,2,1,1,1,1,1,1],"quality":"Less is better","isMain":false,"metricName":"LA75"},{"values":[1,1,1,1,1,1,null,1,1,1,1,1,1],"quality":"Less is better","isMain":true,"metricName":"LGA50"},{"values":[2,2,1,1,1,1,null,1,1,1,1,1,1],"quality":"Less is better","isMain":false,"metricName":"LGA75"}]],["Misassemblies",[{"values":[0,0,0,0,0,0,0,0,0,0,0,0,0],"quality":"Less is better","isMain":true,"metricName":"# misassemblies"},{"values":[0,0,0,0,0,0,0,0,0,0,0,0,0],"quality":"Less is better","isMain":false,"metricName":" # relocations"},{"values":[0,0,0,0,0,0,0,0,0,0,0,0,0],"quality":"Less is better","isMain":false,"metricName":" # translocations"},{"values":[0,0,0,0,0,0,0,0,0,0,0,0,0],"quality":"Less is better","isMain":false,"metricName":" # inversions"},{"values":[0,0,0,0,0,0,0,0,0,0,0,0,0],"quality":"Less is better","isMain":false,"metricName":"# misassembled contigs"},{"values":[0,0,0,0,0,0,0,0,0,0,0,0,0],"quality":"Less is better","isMain":true,"metricName":"Misassembled contigs length"},{"values":[0,0,0,0,1,1,0,0,1,0,0,0,0],"quality":"Less is better","isMain":false,"metricName":"# local misassemblies"},{"values":[0,0,0,0,0,0,0,1,0,0,0,0,0],"quality":"Less is better","isMain":false,"metricName":"# unaligned mis. contigs"}]],["Unaligned",[{"values":[0,0,0,0,0,0,0,0,0,0,0,0,0],"quality":"Less is better","isMain":false,"metricName":"# fully unaligned contigs"},{"values":[0,0,0,0,0,0,0,0,0,0,0,0,0],"quality":"Less is better","isMain":false,"metricName":"Fully unaligned length"},{"values":[0,0,0,0,0,0,0,2,0,0,0,0,0],"quality":"Less is better","isMain":false,"metricName":"# partially unaligned contigs"},{"values":[0,0,0,0,0,0,0,5502,0,0,0,0,0],"quality":"Less is better","isMain":false,"metricName":"Partially unaligned length"}]],["Mismatches",[{"values":[2,0,1,1,0,1,0,0,0,0,0,0,0],"quality":"Less is better","isMain":false,"metricName":"# mismatches"},{"values":[0,0,0,0,0,0,0,17,0,1,0,0,0],"quality":"Less is better","isMain":false,"metricName":"# indels"},{"values":[0,0,0,0,0,0,0,815,0,55,0,0,0],"quality":"Less is better","isMain":false,"metricName":"Indels length"},{"values":["5.94","0.00","2.97","2.97","0.00","2.97","0.00","0.00","0.00","0.00","0.00","0.00","0.00"],"quality":"Less is better","isMain":true,"metricName":"# mismatches per 100 kbp"},{"values":["0.00","0.00","0.00","0.00","0.00","0.00","0.00","60.22","0.00","2.97","0.00","0.00","0.00"],"quality":"Less is better","isMain":true,"metricName":"# indels per 100 kbp"},{"values":[0,0,0,0,0,0,0,0,0,0,0,0,0],"quality":"Less is better","isMain":false,"metricName":" # indels (<= 5 bp)"},{"values":[0,0,0,0,0,0,0,17,0,1,0,0,0],"quality":"Less is better","isMain":false,"metricName":" # indels (> 5 bp)"},{"values":[0,0,0,0,0,0,0,2189,0,0,0,0,0],"quality":"Less is better","isMain":false,"metricName":"# N's"},{"values":["0.00","0.00","0.00","0.00","0.00","0.00","0.00","6426.14","0.00","0.00","0.00","0.00","0.00"],"quality":"Less is better","isMain":true,"metricName":"# N's per 100 kbp"}]],["Statistics without reference",[{"values":[5,5,1,3,1,1,2,2,1,1,2,2,2],"quality":"Equal","isMain":true,"metricName":"# contigs"},{"values":[5,4,1,3,1,1,0,2,1,1,2,2,2],"quality":"Equal","isMain":false,"metricName":"# contigs (>= 1000 bp)"},{"values":[2,1,1,1,1,1,0,1,1,1,1,1,1],"quality":"Equal","isMain":false,"metricName":"# contigs (>= 5000 bp)"},{"values":[1,1,1,1,1,1,0,1,1,1,1,1,1],"quality":"Equal","isMain":false,"metricName":"# contigs (>= 10000 bp)"},{"values":[0,0,1,1,1,1,0,1,1,1,1,1,1],"quality":"Equal","isMain":false,"metricName":"# contigs (>= 25000 bp)"},{"values":[0,0,0,0,0,0,0,0,0,0,0,0,0],"quality":"Equal","isMain":false,"metricName":"# contigs (>= 50000 bp)"},{"values":[20241,24875,33657,31149,33834,33837,581,30232,33820,33748,30951,30951,31011],"quality":"More is better","isMain":true,"metricName":"Largest contig"},{"values":[33765,33440,33657,34739,33834,33837,1119,34064,33820,33748,32647,32647,32654],"quality":"More is better","isMain":true,"metricName":"Total length"},{"values":[33765,32872,33657,34739,33834,33837,0,34064,33820,33748,32647,32647,32654],"quality":"More is better","isMain":true,"metricName":"Total length (>= 1000 bp)"},{"values":[27591,24875,33657,31149,33834,33837,0,30232,33820,33748,30951,30951,31011],"quality":"More is better","isMain":false,"metricName":"Total length (>= 5000 bp)"},{"values":[20241,24875,33657,31149,33834,33837,0,30232,33820,33748,30951,30951,31011],"quality":"More is better","isMain":true,"metricName":"Total length (>= 10000 bp)"},{"values":[0,0,33657,31149,33834,33837,0,30232,33820,33748,30951,30951,31011],"quality":"More is better","isMain":false,"metricName":"Total length (>= 25000 bp)"},{"values":[0,0,0,0,0,0,0,0,0,0,0,0,0],"quality":"More is better","isMain":true,"metricName":"Total length (>= 50000 bp)"},{"values":[20241,24875,33657,31149,33834,33837,581,30232,33820,33748,30951,30951,31011],"quality":"More is better","isMain":false,"metricName":"N50"},{"values":[7350,4425,33657,31149,33834,33837,538,30232,33820,33748,30951,30951,31011],"quality":"More is better","isMain":false,"metricName":"N75"},{"values":[1,1,1,1,1,1,1,1,1,1,1,1,1],"quality":"Less is better","isMain":false,"metricName":"L50"},{"values":[2,2,1,1,1,1,2,1,1,1,1,1,1],"quality":"Less is better","isMain":false,"metricName":"L75"},{"values":["52.50","52.54","52.51","52.33","52.51","52.46","52.19","52.68","52.52","52.48","52.66","52.66","52.66"],"quality":"Equal","isMain":false,"metricName":"GC (%)"}]],["Predicted genes",[]],["Similarity statistics",[{"values":[0,0,0,0,0,0,0,0,0,0,0,0,0],"quality":"Equal","isMain":false,"metricName":"# similar correct contigs"},{"values":[0,0,0,0,0,0,0,0,0,0,0,0,0],"quality":"Equal","isMain":false,"metricName":"# similar misassembled blocks"}]],["Reference statistics",[{"values":[33693,33693,33693,33693,33693,33693,33693,33693,33693,33693,33693,33693,33693],"quality":"Equal","isMain":false,"metricName":"Reference length"},{"values":[1,1,1,1,1,1,1,1,1,1,1,1,1],"quality":"Equal","isMain":false,"metricName":"Reference fragments"},{"values":["52.49","52.49","52.49","52.49","52.49","52.49","52.49","52.49","52.49","52.49","52.49","52.49","52.49"],"quality":"Equal","isMain":false,"metricName":"Reference GC (%)"}]]],"referenceName":"gi\_169936017\_ref\_NC","date":"18 July 2018, Wednesday, 16:34:20","order":[0,1,2,3,4,5,6,7,8,9,10,11,12],"assembliesNames":["ABySS\_127","ABySS\_63","CLC","IDBA\_UD","MEGAHIT","MIRA","Ray\_Meta","SOAPdenovo2","SPAdes","SPAdes\_meta","SPAdes\_sc","SPAdes\_sc\_careful","Velvet"]},{"assembliesWithNs":null,"minContig":500,"report":[["Genome statistics",[{"values":["100.000","100.000","100.000","100.000","100.000","69.290","65.928","100.000","100.000","100.000","100.000","100.000"],"quality":"More is better","isMain":true,"metricName":"Genome fraction (%)"},{"values":["1.002","1.000","1.002","1.003","1.006","1.018","1.016","1.003","1.001","1.001","1.001","1.001"],"quality":"Less is better","isMain":true,"metricName":"Duplication ratio"},{"values":[45139,45051,45150,45192,45340,3700,9642,45178,45106,45106,45106,40006],"quality":"More is better","isMain":true,"metricName":"Largest alignment"},{"values":[45139,45051,45150,45192,45340,31773,29896,45178,45106,45106,45106,45093],"quality":"More is better","isMain":true,"metricName":"Total aligned length"},{"values":[45139,45051,45150,45192,45340,1070,8767,45178,45106,45106,45106,40006],"quality":"More is better","isMain":false,"metricName":"NG50"},{"values":[45139,45051,45150,45192,45340,null,6688,45178,45106,45106,45106,40006],"quality":"More is better","isMain":false,"metricName":"NG75"},{"values":[45139,45051,45150,45192,45340,1423,4971,45178,45106,45106,45106,40006],"quality":"More is better","isMain":false,"metricName":"NA50"},{"values":[45139,45051,45150,45192,45340,1066,null,45178,45106,45106,45106,40006],"quality":"More is better","isMain":false,"metricName":"NA75"},{"values":[45139,45051,45150,45192,45340,1070,4971,45178,45106,45106,45106,40006],"quality":"More is better","isMain":true,"metricName":"NGA50"},{"values":[45139,45051,45150,45192,45340,null,null,45178,45106,45106,45106,40006],"quality":"More is better","isMain":false,"metricName":"NGA75"},{"values":[1,1,1,1,1,14,2,1,1,1,1,1],"quality":"Less is better","isMain":false,"metricName":"LG50"},{"values":[1,1,1,1,1,null,4,1,1,1,1,1],"quality":"Less is better","isMain":false,"metricName":"LG75"},{"values":[1,1,1,1,1,8,3,1,1,1,1,1],"quality":"Less is better","isMain":false,"metricName":"LA50"},{"values":[1,1,1,1,1,15,null,1,1,1,1,1],"quality":"Less is better","isMain":false,"metricName":"LA75"},{"values":[1,1,1,1,1,14,3,1,1,1,1,1],"quality":"Less is better","isMain":true,"metricName":"LGA50"},{"values":[1,1,1,1,1,null,null,1,1,1,1,1],"quality":"Less is better","isMain":false,"metricName":"LGA75"}]],["Misassemblies",[{"values":[0,0,0,0,0,0,0,0,0,0,0,0],"quality":"Less is better","isMain":true,"metricName":"# misassemblies"},{"values":[0,0,0,0,0,0,0,0,0,0,0,0],"quality":"Less is better","isMain":false,"metricName":" # relocations"},{"values":[0,0,0,0,0,0,0,0,0,0,0,0],"quality":"Less is better","isMain":false,"metricName":" # translocations"},{"values":[0,0,0,0,0,0,0,0,0,0,0,0],"quality":"Less is better","isMain":false,"metricName":" # inversions"},{"values":[0,0,0,0,0,0,0,0,0,0,0,0],"quality":"Less is better","isMain":false,"metricName":"# misassembled contigs"},{"values":[0,0,0,0,0,0,0,0,0,0,0,0],"quality":"Less is better","isMain":true,"metricName":"Misassembled contigs length"},{"values":[1,0,1,1,1,0,3,1,0,0,0,0],"quality":"Less is better","isMain":false,"metricName":"# local misassemblies"},{"values":[0,0,0,0,0,0,1,0,0,0,0,0],"quality":"Less is better","isMain":false,"metricName":"# unaligned mis. contigs"}]],["Unaligned",[{"values":[0,0,0,0,0,0,0,0,0,0,0,0],"quality":"Less is better","isMain":false,"metricName":"# fully unaligned contigs"},{"values":[0,0,0,0,0,0,0,0,0,0,0,0],"quality":"Less is better","isMain":false,"metricName":"Fully unaligned length"},{"values":[0,0,0,0,0,0,3,0,0,0,0,0],"quality":"Less is better","isMain":false,"metricName":"# partially unaligned contigs"},{"values":[0,0,0,0,0,0,12912,0,0,0,0,0],"quality":"Less is better","isMain":false,"metricName":"Partially unaligned length"}]],["Mismatches",[{"values":[0,0,0,0,0,0,3,0,0,0,0,0],"quality":"Less is better","isMain":false,"metricName":"# mismatches"},{"values":[0,0,0,0,0,0,17,0,1,1,1,0],"quality":"Less is better","isMain":false,"metricName":"# indels"},{"values":[0,0,0,0,0,0,843,0,55,55,55,0],"quality":"Less is better","isMain":false,"metricName":"Indels length"},{"values":["0.00","0.00","0.00","0.00","0.00","0.00","10.10","0.00","0.00","0.00","0.00","0.00"],"quality":"Less is better","isMain":true,"metricName":"# mismatches per 100 kbp"},{"values":["0.00","0.00","0.00","0.00","0.00","0.00","57.24","0.00","2.22","2.22","2.22","0.00"],"quality":"Less is better","isMain":true,"metricName":"# indels per 100 kbp"},{"values":[0,0,0,0,0,0,0,0,0,0,0,0],"quality":"Less is better","isMain":false,"metricName":" # indels (<= 5 bp)"},{"values":[0,0,0,0,0,0,17,0,1,1,1,0],"quality":"Less is better","isMain":false,"metricName":" # indels (> 5 bp)"},{"values":[0,0,0,0,1,0,3791,0,0,0,0,0],"quality":"Less is better","isMain":false,"metricName":"# N's"},{"values":["0.00","0.00","0.00","0.00","2.21","0.00","8796.23","0.00","0.00","0.00","0.00","0.00"],"quality":"Less is better","isMain":true,"metricName":"# N's per 100 kbp"}]],["Statistics without reference",[{"values":[1,1,1,1,1,25,8,1,1,1,1,2],"quality":"Equal","isMain":true,"metricName":"# contigs"},{"values":[1,1,1,1,1,16,8,1,1,1,1,2],"quality":"Equal","isMain":false,"metricName":"# contigs (>= 1000 bp)"},{"values":[1,1,1,1,1,0,4,1,1,1,1,2],"quality":"Equal","isMain":false,"metricName":"# contigs (>= 5000 bp)"},{"values":[1,1,1,1,1,0,1,1,1,1,1,1],"quality":"Equal","isMain":false,"metricName":"# contigs (>= 10000 bp)"},{"values":[1,1,1,1,1,0,0,1,1,1,1,1],"quality":"Equal","isMain":false,"metricName":"# contigs (>= 25000 bp)"},{"values":[0,0,0,0,0,0,0,0,0,0,0,0],"quality":"Equal","isMain":false,"metricName":"# contigs (>= 50000 bp)"},{"values":[45139,45051,45150,45192,45340,3700,14572,45178,45106,45106,45106,40006],"quality":"More is better","isMain":true,"metricName":"Largest contig"},{"values":[45139,45051,45150,45192,45340,31773,43098,45178,45106,45106,45106,45093],"quality":"More is better","isMain":true,"metricName":"Total length"},{"values":[45139,45051,45150,45192,45340,25287,43098,45178,45106,45106,45106,45093],"quality":"More is better","isMain":true,"metricName":"Total length (>= 1000 bp)"},{"values":[45139,45051,45150,45192,45340,0,38769,45178,45106,45106,45106,45093],"quality":"More is better","isMain":false,"metricName":"Total length (>= 5000 bp)"},{"values":[45139,45051,45150,45192,45340,0,14572,45178,45106,45106,45106,40006],"quality":"More is better","isMain":true,"metricName":"Total length (>= 10000 bp)"},{"values":[45139,45051,45150,45192,45340,0,0,45178,45106,45106,45106,40006],"quality":"More is better","isMain":false,"metricName":"Total length (>= 25000 bp)"},{"values":[0,0,0,0,0,0,0,0,0,0,0,0],"quality":"More is better","isMain":true,"metricName":"Total length (>= 50000 bp)"},{"values":[45139,45051,45150,45192,45340,1423,8767,45178,45106,45106,45106,40006],"quality":"More is better","isMain":false,"metricName":"N50"},{"values":[45139,45051,45150,45192,45340,1066,6688,45178,45106,45106,45106,40006],"quality":"More is better","isMain":false,"metricName":"N75"},{"values":[1,1,1,1,1,8,2,1,1,1,1,1],"quality":"Less is better","isMain":false,"metricName":"L50"},{"values":[1,1,1,1,1,15,4,1,1,1,1,1],"quality":"Less is better","isMain":false,"metricName":"L75"},{"values":["46.11","46.13","46.13","46.11","46.11","46.15","46.28","46.12","46.11","46.12","46.12","46.11"],"quality":"Equal","isMain":false,"metricName":"GC (%)"}]],["Predicted genes",[]],["Similarity statistics",[{"values":[0,0,0,0,0,0,0,0,0,0,0,0],"quality":"Equal","isMain":false,"metricName":"# similar correct contigs"},{"values":[0,0,0,0,0,0,0,0,0,0,0,0],"quality":"Equal","isMain":false,"metricName":"# similar misassembled blocks"}]],["Reference statistics",[{"values":[45051,45051,45051,45051,45051,45051,45051,45051,45051,45051,45051,45051],"quality":"Equal","isMain":false,"metricName":"Reference length"},{"values":[1,1,1,1,1,1,1,1,1,1,1,1],"quality":"Equal","isMain":false,"metricName":"Reference fragments"},{"values":["46.13","46.13","46.13","46.13","46.13","46.13","46.13","46.13","46.13","46.13","46.13","46.13"],"quality":"Equal","isMain":false,"metricName":"Reference GC (%)"}]]],"referenceName":"gi\_170676276\_ref\_NC","date":"18 July 2018, Wednesday, 16:34:33","order":[0,1,2,3,4,5,6,7,8,9,10,11],"assembliesNames":["ABySS\_127","CLC","IDBA\_UD","MEGAHIT","MIRA","Ray\_Meta","SOAPdenovo2","SPAdes","SPAdes\_meta","SPAdes\_sc","SPAdes\_sc\_careful","Velvet"]},{"assembliesWithNs":null,"minContig":500,"report":[["Genome statistics",[{"values":["75.345","62.383","64.552","68.717","67.452","37.214","30.771","12.074","89.423","54.772","66.789","67.649","47.758","98.266"],"quality":"More is better","isMain":true,"metricName":"Genome fraction (%)"},{"values":["1.011","1.005","1.027","1.029","1.027","1.312","1.085","1.229","1.002","1.002","1.006","1.009","1.003","1.160"],"quality":"Less is better","isMain":true,"metricName":"Duplication ratio"},{"values":[8318,11045,16402,6051,11619,12675,1272,1554,31577,18682,11718,11718,6003,36756],"quality":"More is better","isMain":true,"metricName":"Largest alignment"},{"values":[48283,39742,41607,44612,43755,30479,21162,7654,56796,34778,42606,43262,30375,72281],"quality":"More is better","isMain":true,"metricName":"Total aligned length"},{"values":[3811,1388,3699,1381,2477,667,null,null,8552,3069,9623,5311,null,179797],"quality":"More is better","isMain":false,"metricName":"NG50"},{"values":[593,null,null,null,null,null,null,null,3081,null,2952,3070,null,179797],"quality":"More is better","isMain":false,"metricName":"NG75"},{"values":[5763,2818,7735,2805,8165,4873,848,null,31577,18682,7485,4153,1477,null],"quality":"More is better","isMain":false,"metricName":"NA50"},{"values":[3132,1553,3699,1267,2475,908,726,null,6178,8769,567,723,1064,null],"quality":"More is better","isMain":false,"metricName":"NA75"},{"values":[3811,1388,3699,1381,2475,null,null,null,8552,3069,4845,3070,null,36756],"quality":"More is better","isMain":true,"metricName":"NGA50"},{"values":[593,null,null,null,null,null,null,null,3081,null,null,null,null,14037],"quality":"More is better","isMain":false,"metricName":"NGA75"},{"values":[5,10,4,10,5,11,null,null,2,4,2,3,null,1],"quality":"Less is better","isMain":false,"metricName":"LG50"},{"values":[15,null,null,null,null,null,null,null,4,null,5,6,null,1],"quality":"Less is better","isMain":false,"metricName":"LG75"},{"values":[4,4,2,6,3,2,11,null,1,1,3,5,5,null],"quality":"Less is better","isMain":false,"metricName":"LA50"},{"values":[6,8,4,12,5,7,18,null,3,2,12,13,11,null],"quality":"Less is better","isMain":false,"metricName":"LA75"},{"values":[5,10,4,10,5,null,null,null,2,4,4,6,null,1],"quality":"Less is better","isMain":true,"metricName":"LGA50"},{"values":[15,null,null,null,null,null,null,null,4,null,null,null,null,2],"quality":"Less is better","isMain":false,"metricName":"LGA75"}]],["Misassemblies",[{"values":[0,0,0,0,0,0,0,0,0,0,0,0,0,2],"quality":"Less is better","isMain":true,"metricName":"# misassemblies"},{"values":[0,0,0,0,0,0,0,0,0,0,0,0,0,2],"quality":"Less is better","isMain":false,"metricName":" # relocations"},{"values":[0,0,0,0,0,0,0,0,0,0,0,0,0,0],"quality":"Less is better","isMain":false,"metricName":" # translocations"},{"values":[0,0,0,0,0,0,0,0,0,0,0,0,0,0],"quality":"Less is better","isMain":false,"metricName":" # inversions"},{"values":[0,0,0,0,0,0,0,0,0,0,0,0,0,1],"quality":"Less is better","isMain":false,"metricName":"# misassembled contigs"},{"values":[0,0,0,0,0,0,0,0,0,0,0,0,0,58921],"quality":"Less is better","isMain":true,"metricName":"Misassembled contigs length"},{"values":[0,0,0,0,0,1,0,0,1,0,0,0,0,1],"quality":"Less is better","isMain":false,"metricName":"# local misassemblies"},{"values":[0,0,0,0,0,0,0,3,0,0,0,0,0,1],"quality":"Less is better","isMain":false,"metricName":"# unaligned mis. contigs"}]],["Unaligned",[{"values":[0,0,0,0,0,0,0,0,0,0,0,0,0,0],"quality":"Less is better","isMain":false,"metricName":"# fully unaligned contigs"},{"values":[0,0,0,0,0,0,0,0,0,0,0,0,0,0],"quality":"Less is better","isMain":false,"metricName":"Fully unaligned length"},{"values":[0,0,0,0,0,2,0,8,1,0,1,1,0,3],"quality":"Less is better","isMain":false,"metricName":"# partially unaligned contigs"},{"values":[0,0,0,0,0,2718,0,13012,1156,0,13005,13005,0,185196],"quality":"Less is better","isMain":false,"metricName":"Partially unaligned length"}]],["Mismatches",[{"values":[13,5,19,12,58,60,2,0,22,12,29,32,4,206],"quality":"Less is better","isMain":false,"metricName":"# mismatches"},{"values":[1,0,0,1,1,5,2,10,5,0,6,5,1,26],"quality":"Less is better","isMain":false,"metricName":"# indels"},{"values":[2,0,0,1,1,5,2,479,8,0,7,6,1,29],"quality":"Less is better","isMain":false,"metricName":"Indels length"},{"values":["27.22","12.64","46.43","27.55","135.64","254.32","10.25","0.00","38.81","34.56","68.49","74.62","13.21","330.68"],"quality":"Less is better","isMain":true,"metricName":"# mismatches per 100 kbp"},{"values":["2.09","0.00","0.00","2.30","2.34","21.19","10.25","130.65","8.82","0.00","14.17","11.66","3.30","41.74"],"quality":"Less is better","isMain":true,"metricName":"# indels per 100 kbp"},{"values":[1,0,0,1,1,5,2,0,5,0,6,5,1,26],"quality":"Less is better","isMain":false,"metricName":" # indels (<= 5 bp)"},{"values":[0,0,0,0,0,0,0,10,0,0,0,0,0,0],"quality":"Less is better","isMain":false,"metricName":" # indels (> 5 bp)"},{"values":[0,1,0,0,0,5,0,5631,34,0,0,0,0,809],"quality":"Less is better","isMain":false,"metricName":"# N's"},{"values":["0.00","2.52","0.00","0.00","0.00","14.85","0.00","25119.33","58.67","0.00","0.00","0.00","0.00","314.20"],"quality":"Less is better","isMain":true,"metricName":"# N's per 100 kbp"}]],["Statistics without reference",[{"values":[16,18,14,24,14,14,26,16,9,5,13,14,21,4],"quality":"Equal","isMain":true,"metricName":"# contigs"},{"values":[10,12,7,15,10,6,5,8,7,4,6,9,11,4],"quality":"Equal","isMain":false,"metricName":"# contigs (>= 1000 bp)"},{"values":[4,1,2,1,3,2,0,0,3,2,3,3,1,3],"quality":"Equal","isMain":false,"metricName":"# contigs (>= 5000 bp)"},{"values":[0,1,1,0,1,1,0,0,1,1,1,1,0,3],"quality":"Equal","isMain":false,"metricName":"# contigs (>= 10000 bp)"},{"values":[0,0,0,0,0,0,0,0,1,0,0,0,0,2],"quality":"Equal","isMain":false,"metricName":"# contigs (>= 25000 bp)"},{"values":[0,0,0,0,0,0,0,0,0,0,0,0,0,2],"quality":"Equal","isMain":false,"metricName":"# contigs (>= 50000 bp)"},{"values":[8318,11045,16402,6051,11619,14844,1272,3227,31577,18682,24723,24723,6003,179797],"quality":"More is better","isMain":true,"metricName":"Largest contig"},{"values":[48283,39742,42010,44826,43925,33668,21169,22417,57955,34778,55611,56267,30375,257477],"quality":"More is better","isMain":true,"metricName":"Total length"},{"values":[44097,35244,37071,38328,40823,28054,5707,16687,56749,34122,50828,52397,23722,257477],"quality":"More is better","isMain":true,"metricName":"Total length (>= 1000 bp)"},{"values":[29439,11045,24137,6051,28383,20188,0,0,46307,27451,41831,35587,6003,256420],"quality":"More is better","isMain":false,"metricName":"Total length (>= 5000 bp)"},{"values":[0,11045,16402,0,11619,14844,0,0,31577,18682,24723,24723,0,256420],"quality":"More is better","isMain":true,"metricName":"Total length (>= 10000 bp)"},{"values":[0,0,0,0,0,0,0,0,31577,0,0,0,0,238718],"quality":"More is better","isMain":false,"metricName":"Total length (>= 25000 bp)"},{"values":[0,0,0,0,0,0,0,0,0,0,0,0,0,238718],"quality":"More is better","isMain":true,"metricName":"Total length (>= 50000 bp)"},{"values":[5763,2818,7735,2805,8240,5344,848,1890,31577,18682,9623,5553,1477,179797],"quality":"More is better","isMain":false,"metricName":"N50"},{"values":[3132,1553,3699,1331,2477,1937,726,858,6178,8769,7485,4153,1064,58921],"quality":"More is better","isMain":false,"metricName":"N75"},{"values":[4,4,2,6,3,2,11,5,1,1,2,2,5,1],"quality":"Less is better","isMain":false,"metricName":"L50"},{"values":[6,8,4,11,5,5,18,9,3,2,3,5,11,2],"quality":"Less is better","isMain":false,"metricName":"L75"},{"values":["50.01","49.81","49.85","50.05","50.32","47.45","49.62","50.29","49.44","49.83","48.19","48.23","49.79","50.03"],"quality":"Equal","isMain":false,"metricName":"GC (%)"}]],["Predicted genes",[]],["Similarity statistics",[{"values":[0,0,0,0,0,0,0,0,0,0,0,0,0,0],"quality":"Equal","isMain":false,"metricName":"# similar correct contigs"},{"values":[0,0,0,0,0,0,0,0,0,0,0,0,0,0],"quality":"Equal","isMain":false,"metricName":"# similar misassembled blocks"}]],["Reference statistics",[{"values":[63395,63395,63395,63395,63395,63395,63395,63395,63395,63395,63395,63395,63395,63395],"quality":"Equal","isMain":false,"metricName":"Reference length"},{"values":[1,1,1,1,1,1,1,1,1,1,1,1,1,1],"quality":"Equal","isMain":false,"metricName":"Reference fragments"},{"values":["49.50","49.50","49.50","49.50","49.50","49.50","49.50","49.50","49.50","49.50","49.50","49.50","49.50","49.50"],"quality":"Equal","isMain":false,"metricName":"Reference GC (%)"}]]],"referenceName":"gi\_170783609\_ref\_NC","date":"18 July 2018, Wednesday, 16:34:49","order":[0,1,2,3,4,5,6,7,8,9,10,11,12,13],"assembliesNames":["ABySS\_127","ABySS\_63","CLC","IDBA\_UD","MEGAHIT","MIRA","Ray\_Meta","SOAPdenovo2","SPAdes","SPAdes\_meta","SPAdes\_sc","SPAdes\_sc\_careful","Velvet","Geneious"]},{"assembliesWithNs":null,"minContig":500,"report":[["Genome statistics",[{"values":["100.000","100.000","98.242","98.616","100.000","100.000","100.000","3.031","100.000","100.000","100.000","100.000","98.280","100.000"],"quality":"More is better","isMain":true,"metricName":"Genome fraction (%)"},{"values":["1.003","1.002","1.000","1.000","1.007","1.029","1.003","1.000","1.003","1.001","1.001","1.001","1.002","1.007"],"quality":"Less is better","isMain":true,"metricName":"Duplication ratio"},{"values":[41174,41174,40448,40602,40686,29507,41172,644,41170,41170,41170,41172,16933,41468],"quality":"More is better","isMain":true,"metricName":"Largest alignment"},{"values":[41174,41174,40448,40602,41454,42357,41172,1248,41170,41170,41170,41172,40541,41468],"quality":"More is better","isMain":true,"metricName":"Total aligned length"},{"values":[41300,41236,40448,40602,40686,29507,41282,null,41296,41224,41224,41226,12609,41468],"quality":"More is better","isMain":false,"metricName":"NG50"},{"values":[41300,41236,40448,40602,40686,7193,41282,null,41296,41224,41224,41226,7199,41468],"quality":"More is better","isMain":false,"metricName":"NG75"},{"values":[41174,41174,40448,40602,40686,29507,41172,644,41170,41170,41170,41172,12609,41468],"quality":"More is better","isMain":false,"metricName":"NA50"},{"values":[41174,41174,40448,40602,40686,7193,41172,604,41170,41170,41170,41172,7199,41468],"quality":"More is better","isMain":false,"metricName":"NA75"},{"values":[41174,41174,40448,40602,40686,29507,41172,null,41170,41170,41170,41172,12609,41468],"quality":"More is better","isMain":true,"metricName":"NGA50"},{"values":[41174,41174,40448,40602,40686,7193,41172,null,41170,41170,41170,41172,7199,41468],"quality":"More is better","isMain":false,"metricName":"NGA75"},{"values":[1,1,1,1,1,1,1,null,1,1,1,1,2,1],"quality":"Less is better","isMain":false,"metricName":"LG50"},{"values":[1,1,1,1,1,2,1,null,1,1,1,1,3,1],"quality":"Less is better","isMain":false,"metricName":"LG75"},{"values":[1,1,1,1,1,1,1,1,1,1,1,1,2,1],"quality":"Less is better","isMain":false,"metricName":"LA50"},{"values":[1,1,1,1,1,2,1,2,1,1,1,1,3,1],"quality":"Less is better","isMain":false,"metricName":"LA75"},{"values":[1,1,1,1,1,1,1,null,1,1,1,1,2,1],"quality":"Less is better","isMain":true,"metricName":"LGA50"},{"values":[1,1,1,1,1,2,1,null,1,1,1,1,3,1],"quality":"Less is better","isMain":false,"metricName":"LGA75"}]],["Misassemblies",[{"values":[0,0,0,0,0,0,0,0,0,0,0,0,0,0],"quality":"Less is better","isMain":true,"metricName":"# misassemblies"},{"values":[0,0,0,0,0,0,0,0,0,0,0,0,0,0],"quality":"Less is better","isMain":false,"metricName":" # relocations"},{"values":[0,0,0,0,0,0,0,0,0,0,0,0,0,0],"quality":"Less is better","isMain":false,"metricName":" # translocations"},{"values":[0,0,0,0,0,0,0,0,0,0,0,0,0,0],"quality":"Less is better","isMain":false,"metricName":" # inversions"},{"values":[0,0,0,0,0,0,0,0,0,0,0,0,0,0],"quality":"Less is better","isMain":false,"metricName":"# misassembled contigs"},{"values":[0,0,0,0,0,0,0,0,0,0,0,0,0,0],"quality":"Less is better","isMain":true,"metricName":"Misassembled contigs length"},{"values":[0,0,0,0,0,0,0,0,0,0,0,0,0,1],"quality":"Less is better","isMain":false,"metricName":"# local misassemblies"},{"values":[0,0,0,0,0,0,0,0,0,0,0,0,0,0],"quality":"Less is better","isMain":false,"metricName":"# unaligned mis. contigs"}]],["Unaligned",[{"values":[0,0,0,0,0,0,0,0,0,0,0,0,0,0],"quality":"Less is better","isMain":false,"metricName":"# fully unaligned contigs"},{"values":[0,0,0,0,0,0,0,0,0,0,0,0,0,0],"quality":"Less is better","isMain":false,"metricName":"Fully unaligned length"},{"values":[0,0,0,0,0,0,0,0,0,0,0,0,0,0],"quality":"Less is better","isMain":false,"metricName":"# partially unaligned contigs"},{"values":[0,0,0,0,0,0,0,0,0,0,0,0,0,0],"quality":"Less is better","isMain":false,"metricName":"Partially unaligned length"}]],["Mismatches",[{"values":[0,0,2,1,2,0,1,0,0,0,0,0,1,0],"quality":"Less is better","isMain":false,"metricName":"# mismatches"},{"values":[1,1,0,0,0,0,0,0,1,1,1,0,0,0],"quality":"Less is better","isMain":false,"metricName":"# indels"},{"values":[2,2,0,0,0,0,0,0,2,2,2,0,0,0],"quality":"Less is better","isMain":false,"metricName":"Indels length"},{"values":["0.00","0.00","4.94","2.46","4.86","0.00","2.43","0.00","0.00","0.00","0.00","0.00","2.47","0.00"],"quality":"Less is better","isMain":true,"metricName":"# mismatches per 100 kbp"},{"values":["2.43","2.43","0.00","0.00","0.00","0.00","0.00","0.00","2.43","2.43","2.43","0.00","0.00","0.00"],"quality":"Less is better","isMain":true,"metricName":"# indels per 100 kbp"},{"values":[1,1,0,0,0,0,0,0,1,1,1,0,0,0],"quality":"Less is better","isMain":false,"metricName":" # indels (<= 5 bp)"},{"values":[0,0,0,0,0,0,0,0,0,0,0,0,0,0],"quality":"Less is better","isMain":false,"metricName":" # indels (> 5 bp)"},{"values":[3,3,1,0,0,2,0,0,0,0,0,1,0,2],"quality":"Less is better","isMain":false,"metricName":"# N's"},{"values":["7.26","7.28","2.47","0.00","0.00","4.72","0.00","0.00","0.00","0.00","0.00","2.43","0.00","4.82"],"quality":"Less is better","isMain":true,"metricName":"# N's per 100 kbp"}]],["Statistics without reference",[{"values":[1,1,1,1,2,4,1,2,1,1,1,1,4,1],"quality":"Equal","isMain":true,"metricName":"# contigs"},{"values":[1,1,1,1,1,3,1,0,1,1,1,1,4,1],"quality":"Equal","isMain":false,"metricName":"# contigs (>= 1000 bp)"},{"values":[1,1,1,1,1,3,1,0,1,1,1,1,3,1],"quality":"Equal","isMain":false,"metricName":"# contigs (>= 5000 bp)"},{"values":[1,1,1,1,1,1,1,0,1,1,1,1,2,1],"quality":"Equal","isMain":false,"metricName":"# contigs (>= 10000 bp)"},{"values":[1,1,1,1,1,1,1,0,1,1,1,1,0,1],"quality":"Equal","isMain":false,"metricName":"# contigs (>= 25000 bp)"},{"values":[0,0,0,0,0,0,0,0,0,0,0,0,0,0],"quality":"Equal","isMain":false,"metricName":"# contigs (>= 50000 bp)"},{"values":[41300,41236,40448,40602,40686,29507,41282,644,41296,41224,41224,41226,16933,41468],"quality":"More is better","isMain":true,"metricName":"Largest contig"},{"values":[41300,41236,40448,40602,41454,42357,41282,1248,41296,41224,41224,41226,40541,41468],"quality":"More is better","isMain":true,"metricName":"Total length"},{"values":[41300,41236,40448,40602,40686,41781,41282,0,41296,41224,41224,41226,40541,41468],"quality":"More is better","isMain":true,"metricName":"Total length (>= 1000 bp)"},{"values":[41300,41236,40448,40602,40686,41781,41282,0,41296,41224,41224,41226,36741,41468],"quality":"More is better","isMain":false,"metricName":"Total length (>= 5000 bp)"},{"values":[41300,41236,40448,40602,40686,29507,41282,0,41296,41224,41224,41226,29542,41468],"quality":"More is better","isMain":true,"metricName":"Total length (>= 10000 bp)"},{"values":[41300,41236,40448,40602,40686,29507,41282,0,41296,41224,41224,41226,0,41468],"quality":"More is better","isMain":false,"metricName":"Total length (>= 25000 bp)"},{"values":[0,0,0,0,0,0,0,0,0,0,0,0,0,0],"quality":"More is better","isMain":true,"metricName":"Total length (>= 50000 bp)"},{"values":[41300,41236,40448,40602,40686,29507,41282,644,41296,41224,41224,41226,12609,41468],"quality":"More is better","isMain":false,"metricName":"N50"},{"values":[41300,41236,40448,40602,40686,7193,41282,604,41296,41224,41224,41226,7199,41468],"quality":"More is better","isMain":false,"metricName":"N75"},{"values":[1,1,1,1,1,1,1,1,1,1,1,1,2,1],"quality":"Less is better","isMain":false,"metricName":"L50"},{"values":[1,1,1,1,1,2,1,2,1,1,1,1,3,1],"quality":"Less is better","isMain":false,"metricName":"L75"},{"values":["34.67","34.66","34.57","34.58","34.69","34.64","34.67","35.02","34.67","34.65","34.65","34.65","34.56","34.69"],"quality":"Equal","isMain":false,"metricName":"GC (%)"}]],["Predicted genes",[]],["Similarity statistics",[{"values":[1,1,1,1,1,0,1,0,1,1,1,1,0,0],"quality":"Equal","isMain":false,"metricName":"# similar correct contigs"},{"values":[0,0,0,0,0,0,0,0,0,0,0,0,0,0],"quality":"Equal","isMain":false,"metricName":"# similar misassembled blocks"}]],["Reference statistics",[{"values":[41172,41172,41172,41172,41172,41172,41172,41172,41172,41172,41172,41172,41172,41172],"quality":"Equal","isMain":false,"metricName":"Reference length"},{"values":[1,1,1,1,1,1,1,1,1,1,1,1,1,1],"quality":"Equal","isMain":false,"metricName":"Reference fragments"},{"values":["34.64","34.64","34.64","34.64","34.64","34.64","34.64","34.64","34.64","34.64","34.64","34.64","34.64","34.64"],"quality":"Equal","isMain":false,"metricName":"Reference GC (%)"}]]],"referenceName":"gi\_171473648\_ref\_NC","date":"18 July 2018, Wednesday, 16:35:03","order":[0,1,2,3,4,5,6,7,8,9,10,11,12,13],"assembliesNames":["ABySS\_127","ABySS\_63","CLC","IDBA\_UD","MEGAHIT","MIRA","Ray\_Meta","SOAPdenovo2","SPAdes","SPAdes\_meta","SPAdes\_sc","SPAdes\_sc\_careful","Velvet","Geneious"]},{"assembliesWithNs":null,"minContig":500,"report":[["Genome statistics",[{"values":["100.000","100.000","100.000","100.000","100.000","3.243","100.000","100.000","100.000","100.000","100.000","100.000"],"quality":"More is better","isMain":true,"metricName":"Genome fraction (%)"},{"values":["1.000","1.003","1.004","1.016","1.001","1.000","1.004","1.002","1.002","1.002","1.001","1.008"],"quality":"Less is better","isMain":true,"metricName":"Duplication ratio"},{"values":[35580,35679,35721,35589,35628,595,35707,35635,35635,35635,20712,35866],"quality":"More is better","isMain":true,"metricName":"Largest alignment"},{"values":[35580,35679,35721,36158,35628,1154,35707,35635,35635,35635,35629,35866],"quality":"More is better","isMain":true,"metricName":"Total aligned length"},{"values":[35580,35679,35721,35589,35628,null,35707,35635,35635,35635,20712,35866],"quality":"More is better","isMain":false,"metricName":"NG50"},{"values":[35580,35679,35721,35589,35628,null,35707,35635,35635,35635,14917,35866],"quality":"More is better","isMain":false,"metricName":"NG75"},{"values":[35580,35679,35721,35589,35628,595,35707,35635,35635,35635,20712,35866],"quality":"More is better","isMain":false,"metricName":"NA50"},{"values":[35580,35679,35721,35589,35628,559,35707,35635,35635,35635,14917,35866],"quality":"More is better","isMain":false,"metricName":"NA75"},{"values":[35580,35679,35721,35589,35628,null,35707,35635,35635,35635,20712,35866],"quality":"More is better","isMain":true,"metricName":"NGA50"},{"values":[35580,35679,35721,35589,35628,null,35707,35635,35635,35635,14917,35866],"quality":"More is better","isMain":false,"metricName":"NGA75"},{"values":[1,1,1,1,1,null,1,1,1,1,1,1],"quality":"Less is better","isMain":false,"metricName":"LG50"},{"values":[1,1,1,1,1,null,1,1,1,1,2,1],"quality":"Less is better","isMain":false,"metricName":"LG75"},{"values":[1,1,1,1,1,1,1,1,1,1,1,1],"quality":"Less is better","isMain":false,"metricName":"LA50"},{"values":[1,1,1,1,1,2,1,1,1,1,2,1],"quality":"Less is better","isMain":false,"metricName":"LA75"},{"values":[1,1,1,1,1,null,1,1,1,1,1,1],"quality":"Less is better","isMain":true,"metricName":"LGA50"},{"values":[1,1,1,1,1,null,1,1,1,1,2,1],"quality":"Less is better","isMain":false,"metricName":"LGA75"}]],["Misassemblies",[{"values":[0,0,0,0,0,0,0,0,0,0,0,0],"quality":"Less is better","isMain":true,"metricName":"# misassemblies"},{"values":[0,0,0,0,0,0,0,0,0,0,0,0],"quality":"Less is better","isMain":false,"metricName":" # relocations"},{"values":[0,0,0,0,0,0,0,0,0,0,0,0],"quality":"Less is better","isMain":false,"metricName":" # translocations"},{"values":[0,0,0,0,0,0,0,0,0,0,0,0],"quality":"Less is better","isMain":false,"metricName":" # inversions"},{"values":[0,0,0,0,0,0,0,0,0,0,0,0],"quality":"Less is better","isMain":false,"metricName":"# misassembled contigs"},{"values":[0,0,0,0,0,0,0,0,0,0,0,0],"quality":"Less is better","isMain":true,"metricName":"Misassembled contigs length"},{"values":[0,1,1,0,0,0,1,0,0,0,0,1],"quality":"Less is better","isMain":false,"metricName":"# local misassemblies"},{"values":[0,0,0,0,0,0,0,0,0,0,0,0],"quality":"Less is better","isMain":false,"metricName":"# unaligned mis. contigs"}]],["Unaligned",[{"values":[0,0,0,0,0,0,0,0,0,0,0,0],"quality":"Less is better","isMain":false,"metricName":"# fully unaligned contigs"},{"values":[0,0,0,0,0,0,0,0,0,0,0,0],"quality":"Less is better","isMain":false,"metricName":"Fully unaligned length"},{"values":[0,0,0,0,0,0,0,0,0,0,0,0],"quality":"Less is better","isMain":false,"metricName":"# partially unaligned contigs"},{"values":[0,0,0,0,0,0,0,0,0,0,0,0],"quality":"Less is better","isMain":false,"metricName":"Partially unaligned length"}]],["Mismatches",[{"values":[0,0,0,0,0,0,0,0,0,0,0,0],"quality":"Less is better","isMain":false,"metricName":"# mismatches"},{"values":[0,0,0,1,1,0,0,1,1,1,0,0],"quality":"Less is better","isMain":false,"metricName":"# indels"},{"values":[0,0,0,9,48,0,0,55,55,55,0,0],"quality":"Less is better","isMain":false,"metricName":"Indels length"},{"values":["0.00","0.00","0.00","0.00","0.00","0.00","0.00","0.00","0.00","0.00","0.00","0.00"],"quality":"Less is better","isMain":true,"metricName":"# mismatches per 100 kbp"},{"values":["0.00","0.00","0.00","2.81","2.81","0.00","0.00","2.81","2.81","2.81","0.00","0.00"],"quality":"Less is better","isMain":true,"metricName":"# indels per 100 kbp"},{"values":[0,0,0,0,0,0,0,0,0,0,0,0],"quality":"Less is better","isMain":false,"metricName":" # indels (<= 5 bp)"},{"values":[0,0,0,1,1,0,0,1,1,1,0,0],"quality":"Less is better","isMain":false,"metricName":" # indels (> 5 bp)"},{"values":[0,0,0,0,0,0,0,0,0,0,0,0],"quality":"Less is better","isMain":false,"metricName":"# N's"},{"values":["0.00","0.00","0.00","0.00","0.00","0.00","0.00","0.00","0.00","0.00","0.00","0.00"],"quality":"Less is better","isMain":true,"metricName":"# N's per 100 kbp"}]],["Statistics without reference",[{"values":[1,1,1,2,1,2,1,1,1,1,2,1],"quality":"Equal","isMain":true,"metricName":"# contigs"},{"values":[1,1,1,1,1,0,1,1,1,1,2,1],"quality":"Equal","isMain":false,"metricName":"# contigs (>= 1000 bp)"},{"values":[1,1,1,1,1,0,1,1,1,1,2,1],"quality":"Equal","isMain":false,"metricName":"# contigs (>= 5000 bp)"},{"values":[1,1,1,1,1,0,1,1,1,1,2,1],"quality":"Equal","isMain":false,"metricName":"# contigs (>= 10000 bp)"},{"values":[1,1,1,1,1,0,1,1,1,1,0,1],"quality":"Equal","isMain":false,"metricName":"# contigs (>= 25000 bp)"},{"values":[0,0,0,0,0,0,0,0,0,0,0,0],"quality":"Equal","isMain":false,"metricName":"# contigs (>= 50000 bp)"},{"values":[35580,35679,35721,35589,35628,595,35707,35635,35635,35635,20712,35866],"quality":"More is better","isMain":true,"metricName":"Largest contig"},{"values":[35580,35679,35721,36158,35628,1154,35707,35635,35635,35635,35629,35866],"quality":"More is better","isMain":true,"metricName":"Total length"},{"values":[35580,35679,35721,35589,35628,0,35707,35635,35635,35635,35629,35866],"quality":"More is better","isMain":true,"metricName":"Total length (>= 1000 bp)"},{"values":[35580,35679,35721,35589,35628,0,35707,35635,35635,35635,35629,35866],"quality":"More is better","isMain":false,"metricName":"Total length (>= 5000 bp)"},{"values":[35580,35679,35721,35589,35628,0,35707,35635,35635,35635,35629,35866],"quality":"More is better","isMain":true,"metricName":"Total length (>= 10000 bp)"},{"values":[35580,35679,35721,35589,35628,0,35707,35635,35635,35635,0,35866],"quality":"More is better","isMain":false,"metricName":"Total length (>= 25000 bp)"},{"values":[0,0,0,0,0,0,0,0,0,0,0,0],"quality":"More is better","isMain":true,"metricName":"Total length (>= 50000 bp)"},{"values":[35580,35679,35721,35589,35628,595,35707,35635,35635,35635,20712,35866],"quality":"More is better","isMain":false,"metricName":"N50"},{"values":[35580,35679,35721,35589,35628,559,35707,35635,35635,35635,14917,35866],"quality":"More is better","isMain":false,"metricName":"N75"},{"values":[1,1,1,1,1,1,1,1,1,1,1,1],"quality":"Less is better","isMain":false,"metricName":"L50"},{"values":[1,1,1,1,1,2,1,1,1,1,2,1],"quality":"Less is better","isMain":false,"metricName":"L75"},{"values":["62.62","62.64","62.62","62.66","62.64","62.05","62.63","62.63","62.63","62.63","62.62","62.64"],"quality":"Equal","isMain":false,"metricName":"GC (%)"}]],["Predicted genes",[]],["Similarity statistics",[{"values":[0,0,0,0,0,0,0,0,0,0,0,0],"quality":"Equal","isMain":false,"metricName":"# similar correct contigs"},{"values":[0,0,0,0,0,0,0,0,0,0,0,0],"quality":"Equal","isMain":false,"metricName":"# similar misassembled blocks"}]],["Reference statistics",[{"values":[35580,35580,35580,35580,35580,35580,35580,35580,35580,35580,35580,35580],"quality":"Equal","isMain":false,"metricName":"Reference length"},{"values":[1,1,1,1,1,1,1,1,1,1,1,1],"quality":"Equal","isMain":false,"metricName":"Reference fragments"},{"values":["62.62","62.62","62.62","62.62","62.62","62.62","62.62","62.62","62.62","62.62","62.62","62.62"],"quality":"Equal","isMain":false,"metricName":"Reference GC (%)"}]]],"referenceName":"gi\_17313217\_ref\_NC","date":"18 July 2018, Wednesday, 16:35:16","order":[0,1,2,3,4,5,6,7,8,9,10,11],"assembliesNames":["CLC","IDBA\_UD","MEGAHIT","MIRA","Ray\_Meta","SOAPdenovo2","SPAdes","SPAdes\_meta","SPAdes\_sc","SPAdes\_sc\_careful","Velvet","Geneious"]},{"assembliesWithNs":null,"minContig":500,"report":[["Genome statistics",[{"values":["1.242","2.635","3.793","4.095","17.384","4.183","1.465","2.867","16.778"],"quality":"More is better","isMain":true,"metricName":"Genome fraction (%)"},{"values":["1.000","1.000","1.000","1.000","1.285","1.000","1.000","1.000","1.068"],"quality":"Less is better","isMain":true,"metricName":"Duplication ratio"},{"values":[535,587,570,689,1553,780,631,631,1553],"quality":"More is better","isMain":true,"metricName":"Largest alignment"},{"values":[535,1135,1634,1764,9139,1802,631,1235,7231],"quality":"More is better","isMain":true,"metricName":"Total aligned length"},{"values":[null,null,null,null,null,null,null,null,33030],"quality":"More is better","isMain":false,"metricName":"NG50"},{"values":[null,null,null,null,null,null,null,null,33030],"quality":"More is better","isMain":false,"metricName":"NG75"},{"values":[535,587,547,570,785,517,631,631,null],"quality":"More is better","isMain":false,"metricName":"NA50"},{"values":[535,548,517,505,null,505,631,604,null],"quality":"More is better","isMain":false,"metricName":"NA75"},{"values":[null,null,null,null,null,null,null,null,null],"quality":"More is better","isMain":true,"metricName":"NGA50"},{"values":[null,null,null,null,null,null,null,null,1],"quality":"Less is better","isMain":false,"metricName":"LG50"},{"values":[null,null,null,null,null,null,null,null,1],"quality":"Less is better","isMain":false,"metricName":"LG75"},{"values":[1,1,2,2,8,2,1,1,null],"quality":"Less is better","isMain":false,"metricName":"LA50"},{"values":[1,2,3,3,null,3,1,2,null],"quality":"Less is better","isMain":false,"metricName":"LA75"}]],["Misassemblies",[{"values":[0,0,0,0,0,0,0,0,0],"quality":"Less is better","isMain":true,"metricName":"# misassemblies"},{"values":[0,0,0,0,0,0,0,0,0],"quality":"Less is better","isMain":false,"metricName":" # relocations"},{"values":[0,0,0,0,0,0,0,0,0],"quality":"Less is better","isMain":false,"metricName":" # translocations"},{"values":[0,0,0,0,0,0,0,0,0],"quality":"Less is better","isMain":false,"metricName":" # inversions"},{"values":[0,0,0,0,0,0,0,0,0],"quality":"Less is better","isMain":false,"metricName":"# misassembled contigs"},{"values":[0,0,0,0,0,0,0,0,0],"quality":"Less is better","isMain":true,"metricName":"Misassembled contigs length"},{"values":[0,0,0,0,0,0,0,0,0],"quality":"Less is better","isMain":false,"metricName":"# local misassemblies"},{"values":[0,0,0,0,1,0,0,0,2],"quality":"Less is better","isMain":false,"metricName":"# unaligned mis. contigs"}]],["Unaligned",[{"values":[0,0,0,0,0,0,0,0,0],"quality":"Less is better","isMain":false,"metricName":"# fully unaligned contigs"},{"values":[0,0,0,0,0,0,0,0,0],"quality":"Less is better","isMain":false,"metricName":"Fully unaligned length"},{"values":[0,0,0,0,2,0,0,0,2],"quality":"Less is better","isMain":false,"metricName":"# partially unaligned contigs"},{"values":[0,0,0,0,7807,0,0,0,40498],"quality":"Less is better","isMain":false,"metricName":"Partially unaligned length"}]],["Mismatches",[{"values":[0,26,6,3,162,2,0,1,136],"quality":"Less is better","isMain":false,"metricName":"# mismatches"},{"values":[0,0,0,0,2,0,0,0,4],"quality":"Less is better","isMain":false,"metricName":"# indels"},{"values":[0,0,0,0,5,0,0,0,7],"quality":"Less is better","isMain":false,"metricName":"Indels length"},{"values":["0.00","2290.75","367.20","170.07","2163.17","110.99","0.00","80.97","1881.57"],"quality":"Less is better","isMain":true,"metricName":"# mismatches per 100 kbp"},{"values":["0.00","0.00","0.00","0.00","26.71","0.00","0.00","0.00","55.34"],"quality":"Less is better","isMain":true,"metricName":"# indels per 100 kbp"},{"values":[0,0,0,0,2,0,0,0,4],"quality":"Less is better","isMain":false,"metricName":" # indels (<= 5 bp)"},{"values":[0,0,0,0,0,0,0,0,0],"quality":"Less is better","isMain":false,"metricName":" # indels (> 5 bp)"},{"values":[0,0,0,0,11,0,0,0,39],"quality":"Less is better","isMain":false,"metricName":"# N's"},{"values":["0.00","0.00","0.00","0.00","63.10","0.00","0.00","0.00","80.89"],"quality":"Less is better","isMain":true,"metricName":"# N's per 100 kbp"}]],["Statistics without reference",[{"values":[1,2,3,3,6,3,1,2,4],"quality":"Equal","isMain":true,"metricName":"# contigs"},{"values":[0,0,0,0,5,0,0,0,4],"quality":"Equal","isMain":false,"metricName":"# contigs (>= 1000 bp)"},{"values":[0,0,0,0,1,0,0,0,2],"quality":"Equal","isMain":false,"metricName":"# contigs (>= 5000 bp)"},{"values":[0,0,0,0,0,0,0,0,2],"quality":"Equal","isMain":false,"metricName":"# contigs (>= 10000 bp)"},{"values":[0,0,0,0,0,0,0,0,1],"quality":"Equal","isMain":false,"metricName":"# contigs (>= 25000 bp)"},{"values":[0,0,0,0,0,0,0,0,0],"quality":"Equal","isMain":false,"metricName":"# contigs (>= 50000 bp)"},{"values":[535,587,570,689,9674,780,631,631,33030],"quality":"More is better","isMain":true,"metricName":"Largest contig"},{"values":[535,1135,1634,1764,17432,1802,631,1235,48215],"quality":"More is better","isMain":true,"metricName":"Total length"},{"values":[0,0,0,0,16647,0,0,0,48215],"quality":"More is better","isMain":true,"metricName":"Total length (>= 1000 bp)"},{"values":[0,0,0,0,9674,0,0,0,44833],"quality":"More is better","isMain":false,"metricName":"Total length (>= 5000 bp)"},{"values":[0,0,0,0,0,0,0,0,44833],"quality":"More is better","isMain":true,"metricName":"Total length (>= 10000 bp)"},{"values":[0,0,0,0,0,0,0,0,33030],"quality":"More is better","isMain":false,"metricName":"Total length (>= 25000 bp)"},{"values":[0,0,0,0,0,0,0,0,0],"quality":"More is better","isMain":true,"metricName":"Total length (>= 50000 bp)"},{"values":[535,587,547,570,9674,517,631,631,33030],"quality":"More is better","isMain":false,"metricName":"N50"},{"values":[535,548,517,505,1855,505,631,604,11803],"quality":"More is better","isMain":false,"metricName":"N75"},{"values":[1,1,2,2,1,2,1,1,1],"quality":"Less is better","isMain":false,"metricName":"L50"},{"values":[1,2,3,3,3,3,1,2,2],"quality":"Less is better","isMain":false,"metricName":"L75"},{"values":["36.82","37.09","37.64","36.73","34.65","34.02","35.82","34.74","33.97"],"quality":"Equal","isMain":false,"metricName":"GC (%)"}]],["Predicted genes",[]],["Similarity statistics",[{"values":[0,0,0,0,0,0,0,0,0],"quality":"Equal","isMain":false,"metricName":"# similar correct contigs"},{"values":[0,0,0,0,0,0,0,0,0],"quality":"Equal","isMain":false,"metricName":"# similar misassembled blocks"}]],["Reference statistics",[{"values":[43081,43081,43081,43081,43081,43081,43081,43081,43081],"quality":"Equal","isMain":false,"metricName":"Reference length"},{"values":[1,1,1,1,1,1,1,1,1],"quality":"Equal","isMain":false,"metricName":"Reference fragments"},{"values":["35.43","35.43","35.43","35.43","35.43","35.43","35.43","35.43","35.43"],"quality":"Equal","isMain":false,"metricName":"Reference GC (%)"}]]],"referenceName":"gi\_17426228\_ref\_NC","date":"18 July 2018, Wednesday, 16:35:26","order":[0,1,2,3,4,5,6,7,8],"assembliesNames":["ABySS\_127","CLC","IDBA\_UD","MEGAHIT","MIRA","SPAdes","SPAdes\_sc","SPAdes\_sc\_careful","Geneious"]},{"assembliesWithNs":null,"minContig":500,"report":[["Genome statistics",[{"values":["1.785","0.339","9.902","6.294","12.575","4.345","4.364","9.695","8.110","7.019","6.362","0.682"],"quality":"More is better","isMain":true,"metricName":"Genome fraction (%)"},{"values":["1.000","1.082","1.003","1.001","1.013","0.999","1.098","1.084","1.000","1.035","1.039","1.002"],"quality":"Less is better","isMain":true,"metricName":"Duplication ratio"},{"values":[1148,665,7172,2596,4113,1869,1363,1364,2239,1218,1218,784],"quality":"More is better","isMain":true,"metricName":"Largest alignment"},{"values":[3514,665,19488,12401,25016,8548,8589,19816,15947,13830,12525,1345],"quality":"More is better","isMain":true,"metricName":"Total aligned length"},{"values":[738,665,653,1215,979,1326,698,683,823,638,638,784],"quality":"More is better","isMain":false,"metricName":"NA50"},{"values":[553,665,null,649,634,1131,370,592,611,592,592,561],"quality":"More is better","isMain":false,"metricName":"NA75"},{"values":[null,null,null,null,null,null,null,null,null,null,null,null],"quality":"More is better","isMain":true,"metricName":"NGA50"},{"values":[2,1,11,4,7,3,6,12,7,9,8,1],"quality":"Less is better","isMain":false,"metricName":"LA50"},{"values":[4,1,null,8,15,5,11,20,13,15,14,2],"quality":"Less is better","isMain":false,"metricName":"LA75"}]],["Misassemblies",[{"values":[0,0,0,0,0,0,0,0,0,0,0,0],"quality":"Less is better","isMain":true,"metricName":"# misassemblies"},{"values":[0,0,0,0,0,0,0,0,0,0,0,0],"quality":"Less is better","isMain":false,"metricName":" # relocations"},{"values":[0,0,0,0,0,0,0,0,0,0,0,0],"quality":"Less is better","isMain":false,"metricName":" # translocations"},{"values":[0,0,0,0,0,0,0,0,0,0,0,0],"quality":"Less is better","isMain":false,"metricName":" # inversions"},{"values":[0,0,0,0,0,0,0,0,0,0,0,0],"quality":"Less is better","isMain":false,"metricName":"# misassembled contigs"},{"values":[0,0,0,0,0,0,0,0,0,0,0,0],"quality":"Less is better","isMain":true,"metricName":"Misassembled contigs length"},{"values":[0,0,0,0,0,0,0,0,0,0,0,0],"quality":"Less is better","isMain":false,"metricName":"# local misassemblies"},{"values":[0,0,0,0,0,0,0,0,0,0,0,0],"quality":"Less is better","isMain":false,"metricName":"# unaligned mis. contigs"}]],["Unaligned",[{"values":[0,0,0,0,0,0,0,0,0,0,0,0],"quality":"Less is better","isMain":false,"metricName":"# fully unaligned contigs"},{"values":[0,0,0,0,0,0,0,0,0,0,0,0],"quality":"Less is better","isMain":false,"metricName":"Fully unaligned length"},{"values":[0,0,1,1,1,0,2,0,0,0,0,0],"quality":"Less is better","isMain":false,"metricName":"# partially unaligned contigs"},{"values":[0,0,13637,1820,1748,0,1402,0,0,0,0,0],"quality":"Less is better","isMain":false,"metricName":"Partially unaligned length"}]],["Mismatches",[{"values":[2,10,272,204,261,28,58,96,161,71,64,24],"quality":"Less is better","isMain":false,"metricName":"# mismatches"},{"values":[0,2,12,16,39,1,7,4,25,17,11,2],"quality":"Less is better","isMain":false,"metricName":"# indels"},{"values":[0,2,32,26,104,6,253,16,54,29,18,3],"quality":"Less is better","isMain":false,"metricName":"Indels length"},{"values":["56.92","1499.25","1395.44","1646.36","1054.38","327.33","675.13","503.01","1008.39","513.82","510.98","1788.38"],"quality":"Less is better","isMain":true,"metricName":"# mismatches per 100 kbp"},{"values":["0.00","299.85","61.56","129.13","157.55","11.69","81.48","20.96","156.58","123.03","87.82","149.03"],"quality":"Less is better","isMain":true,"metricName":"# indels per 100 kbp"},{"values":[0,2,11,16,35,0,2,2,24,17,11,2],"quality":"Less is better","isMain":false,"metricName":" # indels (<= 5 bp)"},{"values":[0,0,1,0,4,1,5,2,1,0,0,0],"quality":"Less is better","isMain":false,"metricName":" # indels (> 5 bp)"},{"values":[0,0,0,0,0,9,1002,0,0,0,0,0],"quality":"Less is better","isMain":false,"metricName":"# N's"},{"values":["0.00","0.00","0.00","0.00","0.00","105.29","9249.52","0.00","0.00","0.00","0.00","0.00"],"quality":"Less is better","isMain":true,"metricName":"# N's per 100 kbp"}]],["Statistics without reference",[{"values":[5,1,16,10,23,7,12,27,19,20,18,2],"quality":"Equal","isMain":true,"metricName":"# contigs"},{"values":[1,0,6,6,7,5,4,3,3,2,2,0],"quality":"Equal","isMain":false,"metricName":"# contigs (>= 1000 bp)"},{"values":[0,0,1,0,0,0,0,0,0,0,0,0],"quality":"Equal","isMain":false,"metricName":"# contigs (>= 5000 bp)"},{"values":[0,0,1,0,0,0,0,0,0,0,0,0],"quality":"Equal","isMain":false,"metricName":"# contigs (>= 10000 bp)"},{"values":[0,0,0,0,0,0,0,0,0,0,0,0],"quality":"Equal","isMain":false,"metricName":"# contigs (>= 25000 bp)"},{"values":[0,0,0,0,0,0,0,0,0,0,0,0],"quality":"Equal","isMain":false,"metricName":"# contigs (>= 50000 bp)"},{"values":[1148,722,20809,2688,4113,1869,1489,1818,2239,1298,1298,784],"quality":"More is better","isMain":true,"metricName":"Largest contig"},{"values":[3514,722,33192,14224,26836,8548,10833,20690,15961,14299,13016,1345],"quality":"More is better","isMain":true,"metricName":"Total length"},{"values":[1148,0,26560,11538,15958,6946,5283,4352,4754,2468,2468,0],"quality":"More is better","isMain":true,"metricName":"Total length (>= 1000 bp)"},{"values":[0,0,20809,0,0,0,0,0,0,0,0,0],"quality":"More is better","isMain":false,"metricName":"Total length (>= 5000 bp)"},{"values":[0,0,20809,0,0,0,0,0,0,0,0,0],"quality":"More is better","isMain":true,"metricName":"Total length (>= 10000 bp)"},{"values":[0,0,0,0,0,0,0,0,0,0,0,0],"quality":"More is better","isMain":false,"metricName":"Total length (>= 25000 bp)"},{"values":[0,0,0,0,0,0,0,0,0,0,0,0],"quality":"More is better","isMain":true,"metricName":"Total length (>= 50000 bp)"},{"values":[738,722,20809,2006,1225,1326,906,763,823,683,683,784],"quality":"More is better","isMain":false,"metricName":"N50"},{"values":[553,722,1039,1119,770,1131,698,605,611,605,605,561],"quality":"More is better","isMain":false,"metricName":"N75"},{"values":[2,1,1,3,5,3,5,10,7,8,7,1],"quality":"Less is better","isMain":false,"metricName":"L50"},{"values":[4,1,5,6,12,5,8,18,13,14,13,2],"quality":"Less is better","isMain":false,"metricName":"L75"},{"values":["34.58","29.09","30.00","32.60","32.41","34.17","33.55","33.95","32.36","32.27","32.12","31.23"],"quality":"Equal","isMain":false,"metricName":"GC (%)"}]],["Predicted genes",[]],["Similarity statistics",[{"values":[0,0,0,0,0,0,0,0,0,0,0,0],"quality":"Equal","isMain":false,"metricName":"# similar correct contigs"},{"values":[0,0,0,0,0,0,0,0,0,0,0,0],"quality":"Equal","isMain":false,"metricName":"# similar misassembled blocks"}]],["Reference statistics",[{"values":[196858,196858,196858,196858,196858,196858,196858,196858,196858,196858,196858,196858],"quality":"Equal","isMain":false,"metricName":"Reference length"},{"values":[1,1,1,1,1,1,1,1,1,1,1,1],"quality":"Equal","isMain":false,"metricName":"Reference fragments"},{"values":["33.09","33.09","33.09","33.09","33.09","33.09","33.09","33.09","33.09","33.09","33.09","33.09"],"quality":"Equal","isMain":false,"metricName":"Reference GC (%)"}]]],"referenceName":"gi\_17974913\_ref\_NC","date":"18 July 2018, Wednesday, 16:35:40","order":[0,1,2,3,4,5,6,7,8,9,10,11],"assembliesNames":["ABySS\_127","ABySS\_63","CLC","IDBA\_UD","MEGAHIT","MIRA","SOAPdenovo2","SPAdes","SPAdes\_meta","SPAdes\_sc","SPAdes\_sc\_careful","Velvet"]},{"assembliesWithNs":null,"minContig":500,"report":[["Genome statistics",[{"values":["76.788","47.166","75.079","69.205","82.279","50.413","36.234","14.665","91.232","58.181","64.068","67.285","35.319","100.000"],"quality":"More is better","isMain":true,"metricName":"Genome fraction (%)"},{"values":["1.053","1.012","1.001","1.014","1.026","1.151","1.034","1.217","1.013","1.004","1.003","1.005","1.005","1.237"],"quality":"Less is better","isMain":true,"metricName":"Duplication ratio"},{"values":[8441,4027,8012,5666,8104,11483,3006,1517,12822,7823,10469,10789,2181,46009],"quality":"More is better","isMain":true,"metricName":"Largest alignment"},{"values":[43140,25468,40116,37451,45043,30959,19992,7827,49328,31163,34307,36079,18945,65998],"quality":"More is better","isMain":true,"metricName":"Total aligned length"},{"values":[4468,546,4671,1087,2568,1688,null,null,4898,1762,3363,3363,null,54122],"quality":"More is better","isMain":false,"metricName":"NG50"},{"values":[697,null,737,null,968,null,null,null,2089,null,null,null,null,54122],"quality":"More is better","isMain":false,"metricName":"NG75"},{"values":[6593,2106,7598,2484,4622,6539,901,226,5722,5802,7823,7823,1224,46009],"quality":"More is better","isMain":false,"metricName":"NA50"},{"values":[1922,931,2731,989,1389,5857,684,null,2136,4094,3363,3363,846,3748],"quality":"More is better","isMain":false,"metricName":"NA75"},{"values":[4468,null,4671,1087,2568,1688,null,null,4898,1762,3363,3363,null,46009],"quality":"More is better","isMain":true,"metricName":"NGA50"},{"values":[697,null,737,null,967,null,null,null,2089,null,null,null,null,46009],"quality":"More is better","isMain":false,"metricName":"NGA75"},{"values":[4,14,4,11,5,5,null,null,4,5,4,4,null,1],"quality":"Less is better","isMain":false,"metricName":"LG50"},{"values":[14,null,13,null,14,null,null,null,8,null,null,null,null,1],"quality":"Less is better","isMain":false,"metricName":"LG75"},{"values":[3,5,3,5,4,2,7,14,3,3,2,2,6,1],"quality":"Less is better","isMain":false,"metricName":"LA50"},{"values":[7,10,5,12,9,3,14,null,6,4,4,4,11,3],"quality":"Less is better","isMain":false,"metricName":"LA75"},{"values":[4,null,4,11,5,5,null,null,4,5,4,4,null,1],"quality":"Less is better","isMain":true,"metricName":"LGA50"},{"values":[14,null,13,null,14,null,null,null,8,null,null,null,null,1],"quality":"Less is better","isMain":false,"metricName":"LGA75"}]],["Misassemblies",[{"values":[0,0,0,0,0,0,0,0,0,0,0,0,0,1],"quality":"Less is better","isMain":true,"metricName":"# misassemblies"},{"values":[0,0,0,0,0,0,0,0,0,0,0,0,0,1],"quality":"Less is better","isMain":false,"metricName":" # relocations"},{"values":[0,0,0,0,0,0,0,0,0,0,0,0,0,0],"quality":"Less is better","isMain":false,"metricName":" # translocations"},{"values":[0,0,0,0,0,0,0,0,0,0,0,0,0,0],"quality":"Less is better","isMain":false,"metricName":" # inversions"},{"values":[0,0,0,0,0,0,0,0,0,0,0,0,0,1],"quality":"Less is better","isMain":false,"metricName":"# misassembled contigs"},{"values":[0,0,0,0,0,0,0,0,0,0,0,0,0,30404],"quality":"Less is better","isMain":true,"metricName":"Misassembled contigs length"},{"values":[0,0,0,0,0,1,0,0,0,0,0,0,0,5],"quality":"Less is better","isMain":false,"metricName":"# local misassemblies"},{"values":[0,0,0,0,0,0,0,3,0,0,0,0,0,0],"quality":"Less is better","isMain":false,"metricName":"# unaligned mis. contigs"}]],["Unaligned",[{"values":[0,0,0,0,0,0,0,0,0,0,0,0,0,0],"quality":"Less is better","isMain":false,"metricName":"# fully unaligned contigs"},{"values":[0,0,0,0,0,0,0,0,0,0,0,0,0,0],"quality":"Less is better","isMain":false,"metricName":"Fully unaligned length"},{"values":[0,1,0,0,0,1,0,3,0,0,0,0,0,2],"quality":"Less is better","isMain":false,"metricName":"# partially unaligned contigs"},{"values":[0,2656,0,0,0,611,0,5040,0,0,0,0,0,18528],"quality":"Less is better","isMain":false,"metricName":"Partially unaligned length"}]],["Mismatches",[{"values":[82,11,0,13,10,46,47,0,35,126,70,66,9,155],"quality":"Less is better","isMain":false,"metricName":"# mismatches"},{"values":[0,0,0,2,0,3,0,3,0,0,2,2,0,12],"quality":"Less is better","isMain":false,"metricName":"# indels"},{"values":[0,0,0,2,0,3,0,143,0,0,2,2,0,12],"quality":"Less is better","isMain":false,"metricName":"Indels length"},{"values":["200.08","43.70","0.00","35.20","22.77","170.96","243.03","0.00","71.88","405.76","204.71","183.78","47.74","290.41"],"quality":"Less is better","isMain":true,"metricName":"# mismatches per 100 kbp"},{"values":["0.00","0.00","0.00","5.41","0.00","11.15","0.00","38.33","0.00","0.00","5.85","5.57","0.00","22.48"],"quality":"Less is better","isMain":true,"metricName":"# indels per 100 kbp"},{"values":[0,0,0,2,0,3,0,0,0,0,2,2,0,12],"quality":"Less is better","isMain":false,"metricName":" # indels (<= 5 bp)"},{"values":[0,0,0,0,0,0,0,3,0,0,0,0,0,0],"quality":"Less is better","isMain":false,"metricName":" # indels (> 5 bp)"},{"values":[0,0,0,0,0,0,0,2710,0,0,0,0,0,139],"quality":"Less is better","isMain":false,"metricName":"# N's"},{"values":["0.00","0.00","0.00","0.00","0.00","0.00","0.00","18601.14","0.00","0.00","0.00","0.00","0.00","164.45"],"quality":"Less is better","isMain":true,"metricName":"# N's per 100 kbp"}]],["Statistics without reference",[{"values":[18,16,13,25,20,10,21,13,14,9,10,12,17,2],"quality":"Equal","isMain":true,"metricName":"# contigs"},{"values":[10,9,9,11,12,5,6,5,12,8,9,9,9,2],"quality":"Equal","isMain":false,"metricName":"# contigs (>= 1000 bp)"},{"values":[3,0,3,1,3,3,0,0,3,3,3,3,0,2],"quality":"Equal","isMain":false,"metricName":"# contigs (>= 5000 bp)"},{"values":[0,0,0,0,0,1,0,0,1,0,1,1,0,2],"quality":"Equal","isMain":false,"metricName":"# contigs (>= 10000 bp)"},{"values":[0,0,0,0,0,0,0,0,0,0,0,0,0,2],"quality":"Equal","isMain":false,"metricName":"# contigs (>= 25000 bp)"},{"values":[0,0,0,0,0,0,0,0,0,0,0,0,0,1],"quality":"Equal","isMain":false,"metricName":"# contigs (>= 50000 bp)"},{"values":[8441,4514,8012,5666,8104,12094,3006,2618,12822,7823,10469,10789,2181,54122],"quality":"More is better","isMain":true,"metricName":"Largest contig"},{"values":[43140,28124,40116,37451,45046,31570,19992,14569,49328,31163,34307,36079,18945,84526],"quality":"More is better","isMain":true,"metricName":"Total length"},{"values":[37497,23434,36778,27724,38104,28222,9136,9670,47840,30203,33754,34129,13294,84526],"quality":"More is better","isMain":true,"metricName":"Total length (>= 1000 bp)"},{"values":[22251,0,23587,5666,20427,24490,0,0,26630,20891,23665,24006,0,84526],"quality":"More is better","isMain":false,"metricName":"Total length (>= 5000 bp)"},{"values":[0,0,0,0,0,12094,0,0,12822,0,10469,10789,0,84526],"quality":"More is better","isMain":true,"metricName":"Total length (>= 10000 bp)"},{"values":[0,0,0,0,0,0,0,0,0,0,0,0,0,84526],"quality":"More is better","isMain":false,"metricName":"Total length (>= 25000 bp)"},{"values":[0,0,0,0,0,0,0,0,0,0,0,0,0,54122],"quality":"More is better","isMain":true,"metricName":"Total length (>= 50000 bp)"},{"values":[6593,3099,7598,2484,4622,6539,901,1610,5722,5802,7823,7823,1224,54122],"quality":"More is better","isMain":false,"metricName":"N50"},{"values":[1922,1380,2731,989,1389,5857,684,663,2136,4094,3363,3363,846,30404],"quality":"More is better","isMain":false,"metricName":"N75"},{"values":[3,4,3,5,4,2,7,4,3,3,2,2,6,1],"quality":"Less is better","isMain":false,"metricName":"L50"},{"values":[7,8,5,12,9,3,14,7,6,4,4,4,11,2],"quality":"Less is better","isMain":false,"metricName":"L75"},{"values":["61.30","61.06","61.52","61.17","61.47","59.92","61.62","60.85","61.20","61.26","61.41","61.36","59.64","60.23"],"quality":"Equal","isMain":false,"metricName":"GC (%)"}]],["Predicted genes",[]],["Similarity statistics",[{"values":[0,0,0,0,0,0,0,0,0,0,0,0,0,0],"quality":"Equal","isMain":false,"metricName":"# similar correct contigs"},{"values":[0,0,0,0,0,0,0,0,0,0,0,0,0,0],"quality":"Equal","isMain":false,"metricName":"# similar misassembled blocks"}]],["Reference statistics",[{"values":[53373,53373,53373,53373,53373,53373,53373,53373,53373,53373,53373,53373,53373,53373],"quality":"Equal","isMain":false,"metricName":"Reference length"},{"values":[1,1,1,1,1,1,1,1,1,1,1,1,1,1],"quality":"Equal","isMain":false,"metricName":"Reference fragments"},{"values":["61.19","61.19","61.19","61.19","61.19","61.19","61.19","61.19","61.19","61.19","61.19","61.19","61.19","61.19"],"quality":"Equal","isMain":false,"metricName":"Reference GC (%)"}]]],"referenceName":"gi\_17975161\_ref\_NC","date":"18 July 2018, Wednesday, 16:35:54","order":[0,1,2,3,4,5,6,7,8,9,10,11,12,13],"assembliesNames":["ABySS\_127","ABySS\_63","CLC","IDBA\_UD","MEGAHIT","MIRA","Ray\_Meta","SOAPdenovo2","SPAdes","SPAdes\_meta","SPAdes\_sc","SPAdes\_sc\_careful","Velvet","Geneious"]},{"assembliesWithNs":null,"minContig":500,"report":[["Genome statistics",[{"values":["9.959","5.342","68.170","61.321","59.347","22.940","60.017","70.319","70.319","70.319","70.319"],"quality":"More is better","isMain":true,"metricName":"Genome fraction (%)"},{"values":["1.000","1.000","1.001","1.000","1.000","1.000","1.001","1.001","1.001","1.001","1.001"],"quality":"Less is better","isMain":true,"metricName":"Duplication ratio"},{"values":[744,634,2972,2826,2711,2670,1795,2972,2972,2972,2972],"quality":"More is better","isMain":true,"metricName":"Largest alignment"},{"values":[3138,1683,21511,19321,18699,7228,18929,22188,22188,22188,22188],"quality":"More is better","isMain":true,"metricName":"Total aligned length"},{"values":[null,null,878,677,638,null,744,878,878,878,878],"quality":"More is better","isMain":false,"metricName":"NG50"},{"values":[622,537,1191,1191,1175,1522,965,1191,1191,1191,1191],"quality":"More is better","isMain":false,"metricName":"NA50"},{"values":[614,512,878,742,715,1270,777,876,876,876,876],"quality":"More is better","isMain":false,"metricName":"NA75"},{"values":[null,null,878,677,638,null,744,878,878,878,878],"quality":"More is better","isMain":true,"metricName":"NGA50"},{"values":[null,null,11,13,13,null,15,11,11,11,11],"quality":"Less is better","isMain":false,"metricName":"LG50"},{"values":[3,2,6,5,5,2,8,6,6,6,6],"quality":"Less is better","isMain":false,"metricName":"LA50"},{"values":[4,3,11,11,11,3,13,12,12,12,12],"quality":"Less is better","isMain":false,"metricName":"LA75"},{"values":[null,null,11,13,13,null,15,11,11,11,11],"quality":"Less is better","isMain":true,"metricName":"LGA50"}]],["Misassemblies",[{"values":[0,0,0,0,0,0,0,0,0,0,0],"quality":"Less is better","isMain":true,"metricName":"# misassemblies"},{"values":[0,0,0,0,0,0,0,0,0,0,0],"quality":"Less is better","isMain":false,"metricName":" # relocations"},{"values":[0,0,0,0,0,0,0,0,0,0,0],"quality":"Less is better","isMain":false,"metricName":" # translocations"},{"values":[0,0,0,0,0,0,0,0,0,0,0],"quality":"Less is better","isMain":false,"metricName":" # inversions"},{"values":[0,0,0,0,0,0,0,0,0,0,0],"quality":"Less is better","isMain":false,"metricName":"# misassembled contigs"},{"values":[0,0,0,0,0,0,0,0,0,0,0],"quality":"Less is better","isMain":true,"metricName":"Misassembled contigs length"},{"values":[0,0,0,0,0,0,0,0,0,0,0],"quality":"Less is better","isMain":false,"metricName":"# local misassemblies"},{"values":[0,0,0,0,0,0,0,0,0,0,0],"quality":"Less is better","isMain":false,"metricName":"# unaligned mis. contigs"}]],["Unaligned",[{"values":[0,0,0,0,0,0,0,0,0,0,0],"quality":"Less is better","isMain":false,"metricName":"# fully unaligned contigs"},{"values":[0,0,0,0,0,0,0,0,0,0,0],"quality":"Less is better","isMain":false,"metricName":"Fully unaligned length"},{"values":[0,0,0,0,0,0,0,0,0,0,0],"quality":"Less is better","isMain":false,"metricName":"# partially unaligned contigs"},{"values":[0,0,0,0,0,0,0,0,0,0,0],"quality":"Less is better","isMain":false,"metricName":"Partially unaligned length"}]],["Mismatches",[{"values":[2,0,25,20,15,1,17,26,23,23,20],"quality":"Less is better","isMain":false,"metricName":"# mismatches"},{"values":[0,0,0,0,0,0,0,0,0,0,0],"quality":"Less is better","isMain":false,"metricName":"# indels"},{"values":[0,0,0,0,0,0,0,0,0,0,0],"quality":"Less is better","isMain":false,"metricName":"Indels length"},{"values":["63.73","0.00","116.39","103.51","80.22","13.84","89.90","117.35","103.81","103.81","90.27"],"quality":"Less is better","isMain":true,"metricName":"# mismatches per 100 kbp"},{"values":["0.00","0.00","0.00","0.00","0.00","0.00","0.00","0.00","0.00","0.00","0.00"],"quality":"Less is better","isMain":true,"metricName":"# indels per 100 kbp"},{"values":[0,0,0,0,0,0,0,0,0,0,0],"quality":"Less is better","isMain":false,"metricName":" # indels (<= 5 bp)"},{"values":[0,0,0,0,0,0,0,0,0,0,0],"quality":"Less is better","isMain":false,"metricName":" # indels (> 5 bp)"},{"values":[0,0,0,0,0,13,0,0,0,0,0],"quality":"Less is better","isMain":false,"metricName":"# N's"},{"values":["0.00","0.00","0.00","0.00","0.00","179.86","0.00","0.00","0.00","0.00","0.00"],"quality":"Less is better","isMain":true,"metricName":"# N's per 100 kbp"}]],["Statistics without reference",[{"values":[5,3,18,18,18,5,20,19,19,19,19],"quality":"Equal","isMain":true,"metricName":"# contigs"},{"values":[0,0,9,6,7,4,7,9,9,9,9],"quality":"Equal","isMain":false,"metricName":"# contigs (>= 1000 bp)"},{"values":[0,0,0,0,0,0,0,0,0,0,0],"quality":"Equal","isMain":false,"metricName":"# contigs (>= 5000 bp)"},{"values":[0,0,0,0,0,0,0,0,0,0,0],"quality":"Equal","isMain":false,"metricName":"# contigs (>= 10000 bp)"},{"values":[0,0,0,0,0,0,0,0,0,0,0],"quality":"Equal","isMain":false,"metricName":"# contigs (>= 25000 bp)"},{"values":[0,0,0,0,0,0,0,0,0,0,0],"quality":"Equal","isMain":false,"metricName":"# contigs (>= 50000 bp)"},{"values":[744,634,2972,2826,2711,2670,1795,2972,2972,2972,2972],"quality":"More is better","isMain":true,"metricName":"Largest contig"},{"values":[3138,1683,21511,19321,18699,7228,18929,22188,22188,22188,22188],"quality":"More is better","isMain":true,"metricName":"Total length"},{"values":[0,0,14687,10867,11621,6511,9218,14687,14687,14687,14687],"quality":"More is better","isMain":true,"metricName":"Total length (>= 1000 bp)"},{"values":[0,0,0,0,0,0,0,0,0,0,0],"quality":"More is better","isMain":false,"metricName":"Total length (>= 5000 bp)"},{"values":[0,0,0,0,0,0,0,0,0,0,0],"quality":"More is better","isMain":true,"metricName":"Total length (>= 10000 bp)"},{"values":[0,0,0,0,0,0,0,0,0,0,0],"quality":"More is better","isMain":false,"metricName":"Total length (>= 25000 bp)"},{"values":[0,0,0,0,0,0,0,0,0,0,0],"quality":"More is better","isMain":true,"metricName":"Total length (>= 50000 bp)"},{"values":[622,537,1191,1191,1175,1522,965,1191,1191,1191,1191],"quality":"More is better","isMain":false,"metricName":"N50"},{"values":[614,512,878,742,715,1270,777,876,876,876,876],"quality":"More is better","isMain":false,"metricName":"N75"},{"values":[3,2,6,5,5,2,8,6,6,6,6],"quality":"Less is better","isMain":false,"metricName":"L50"},{"values":[4,3,11,11,11,3,13,12,12,12,12],"quality":"Less is better","isMain":false,"metricName":"L75"},{"values":["39.52","38.09","40.15","40.36","40.18","39.61","40.33","40.15","40.12","40.12","40.10"],"quality":"Equal","isMain":false,"metricName":"GC (%)"}]],["Predicted genes",[]],["Similarity statistics",[{"values":[0,0,0,0,0,0,0,0,0,0,0],"quality":"Equal","isMain":false,"metricName":"# similar correct contigs"},{"values":[0,0,0,0,0,0,0,0,0,0,0],"quality":"Equal","isMain":false,"metricName":"# similar misassembled blocks"}]],["Reference statistics",[{"values":[31508,31508,31508,31508,31508,31508,31508,31508,31508,31508,31508],"quality":"Equal","isMain":false,"metricName":"Reference length"},{"values":[1,1,1,1,1,1,1,1,1,1,1],"quality":"Equal","isMain":false,"metricName":"Reference fragments"},{"values":["39.94","39.94","39.94","39.94","39.94","39.94","39.94","39.94","39.94","39.94","39.94"],"quality":"Equal","isMain":false,"metricName":"Reference GC (%)"}]]],"referenceName":"gi\_17981815\_ref\_NC","date":"18 July 2018, Wednesday, 16:36:06","order":[0,1,2,3,4,5,6,7,8,9,10],"assembliesNames":["ABySS\_127","ABySS\_63","CLC","IDBA\_UD","MEGAHIT","MIRA","SOAPdenovo2","SPAdes","SPAdes\_meta","SPAdes\_sc","SPAdes\_sc\_careful"]},{"assembliesWithNs":null,"minContig":500,"report":[["Genome statistics",[{"values":["58.049","45.318","57.482","73.636","66.313","1.627","1.376","17.394","32.209","50.612","52.225","56.096","100.000"],"quality":"More is better","isMain":true,"metricName":"Genome fraction (%)"},{"values":["1.010","1.009","1.001","1.013","1.051","1.000","1.000","1.035","1.000","1.003","1.003","1.001","1.016"],"quality":"Less is better","isMain":true,"metricName":"Duplication ratio"},{"values":[16456,8823,16661,8143,8553,680,575,4438,6765,8119,8173,7262,42444],"quality":"More is better","isMain":true,"metricName":"Largest alignment"},{"values":[24493,19110,24047,31169,29116,680,575,7524,13462,21209,21883,23474,42444],"quality":"More is better","isMain":true,"metricName":"Total aligned length"},{"values":[4654,null,4355,3396,30728,null,null,null,null,864,1845,682,42444],"quality":"More is better","isMain":false,"metricName":"NG50"},{"values":[null,null,null,null,8553,null,null,null,null,null,null,null,42444],"quality":"More is better","isMain":false,"metricName":"NG75"},{"values":[16456,5761,16661,3843,1318,680,575,4438,6765,6627,6563,3152,42444],"quality":"More is better","isMain":false,"metricName":"NA50"},{"values":[4654,2384,4355,1447,null,680,575,2310,6697,4438,4438,1091,42444],"quality":"More is better","isMain":false,"metricName":"NA75"},{"values":[4654,null,4355,3396,7193,null,null,null,null,864,1845,682,42444],"quality":"More is better","isMain":true,"metricName":"NGA50"},{"values":[null,null,null,null,null,null,null,null,null,null,null,null,42444],"quality":"More is better","isMain":false,"metricName":"NGA75"},{"values":[2,null,2,4,1,null,null,null,null,5,4,9,1],"quality":"Less is better","isMain":false,"metricName":"LG50"},{"values":[null,null,null,null,2,null,null,null,null,null,null,null,1],"quality":"Less is better","isMain":false,"metricName":"LG75"},{"values":[1,2,1,3,5,1,1,1,1,2,2,3,1],"quality":"Less is better","isMain":false,"metricName":"LA50"},{"values":[2,3,2,6,null,1,1,2,2,3,3,6,1],"quality":"Less is better","isMain":false,"metricName":"LA75"},{"values":[2,null,2,4,3,null,null,null,null,5,4,9,1],"quality":"Less is better","isMain":true,"metricName":"LGA50"},{"values":[null,null,null,null,null,null,null,null,null,null,null,null,1],"quality":"Less is better","isMain":false,"metricName":"LGA75"}]],["Misassemblies",[{"values":[0,1,0,0,0,0,0,0,0,0,0,0,0],"quality":"Less is better","isMain":true,"metricName":"# misassemblies"},{"values":[0,1,0,0,0,0,0,0,0,0,0,0,0],"quality":"Less is better","isMain":false,"metricName":" # relocations"},{"values":[0,0,0,0,0,0,0,0,0,0,0,0,0],"quality":"Less is better","isMain":false,"metricName":" # translocations"},{"values":[0,0,0,0,0,0,0,0,0,0,0,0,0],"quality":"Less is better","isMain":false,"metricName":" # inversions"},{"values":[0,1,0,0,0,0,0,0,0,0,0,0,0],"quality":"Less is better","isMain":false,"metricName":"# misassembled contigs"},{"values":[0,9213,0,0,0,0,0,0,0,0,0,0,0],"quality":"Less is better","isMain":true,"metricName":"Misassembled contigs length"},{"values":[0,0,0,0,0,0,0,0,0,0,0,0,1],"quality":"Less is better","isMain":false,"metricName":"# local misassemblies"},{"values":[0,0,0,0,0,0,0,0,0,0,0,0,0],"quality":"Less is better","isMain":false,"metricName":"# unaligned mis. contigs"}]],["Unaligned",[{"values":[0,0,0,0,0,0,0,0,0,0,0,0,0],"quality":"Less is better","isMain":false,"metricName":"# fully unaligned contigs"},{"values":[0,0,0,0,0,0,0,0,0,0,0,0,0],"quality":"Less is better","isMain":false,"metricName":"Fully unaligned length"},{"values":[0,1,0,0,1,0,0,0,0,0,0,0,0],"quality":"Less is better","isMain":false,"metricName":"# partially unaligned contigs"},{"values":[0,1068,0,0,22470,0,0,0,0,0,0,0,0],"quality":"Less is better","isMain":false,"metricName":"Partially unaligned length"}]],["Mismatches",[{"values":[0,4,5,0,3,2,1,0,0,8,0,0,4],"quality":"Less is better","isMain":false,"metricName":"# mismatches"},{"values":[0,0,0,0,0,0,0,0,0,0,0,0,0],"quality":"Less is better","isMain":false,"metricName":"# indels"},{"values":[0,0,0,0,0,0,0,0,0,0,0,0,0],"quality":"Less is better","isMain":false,"metricName":"Indels length"},{"values":["0.00","21.12","20.81","0.00","10.82","294.12","173.91","0.00","0.00","37.82","0.00","0.00","9.57"],"quality":"Less is better","isMain":true,"metricName":"# mismatches per 100 kbp"},{"values":["0.00","0.00","0.00","0.00","0.00","0.00","0.00","0.00","0.00","0.00","0.00","0.00","0.00"],"quality":"Less is better","isMain":true,"metricName":"# indels per 100 kbp"},{"values":[0,0,0,0,0,0,0,0,0,0,0,0,0],"quality":"Less is better","isMain":false,"metricName":" # indels (<= 5 bp)"},{"values":[0,0,0,0,0,0,0,0,0,0,0,0,0],"quality":"Less is better","isMain":false,"metricName":" # indels (> 5 bp)"},{"values":[0,51,0,0,0,1,0,0,0,0,0,0,4],"quality":"Less is better","isMain":false,"metricName":"# N's"},{"values":["0.00","252.75","0.00","0.00","0.00","147.06","0.00","0.00","0.00","0.00","0.00","0.00","9.42"],"quality":"Less is better","isMain":true,"metricName":"# N's per 100 kbp"}]],["Statistics without reference",[{"values":[4,5,4,12,8,1,1,3,2,5,5,13,1],"quality":"Equal","isMain":true,"metricName":"# contigs"},{"values":[3,2,3,9,5,0,0,2,2,4,4,8,1],"quality":"Equal","isMain":false,"metricName":"# contigs (>= 1000 bp)"},{"values":[1,2,1,2,3,0,0,0,2,2,2,1,1],"quality":"Equal","isMain":false,"metricName":"# contigs (>= 5000 bp)"},{"values":[1,0,1,0,1,0,0,0,0,0,0,0,1],"quality":"Equal","isMain":false,"metricName":"# contigs (>= 10000 bp)"},{"values":[0,0,0,0,1,0,0,0,0,0,0,0,1],"quality":"Equal","isMain":false,"metricName":"# contigs (>= 25000 bp)"},{"values":[0,0,0,0,0,0,0,0,0,0,0,0,0],"quality":"Equal","isMain":false,"metricName":"# contigs (>= 50000 bp)"},{"values":[16456,9213,16661,8143,30728,680,575,4438,6765,8119,8178,7262,42444],"quality":"More is better","isMain":true,"metricName":"Largest contig"},{"values":[24493,20178,24047,31169,51586,680,575,7524,13462,21209,21888,23474,42444],"quality":"More is better","isMain":true,"metricName":"Total length"},{"values":[23767,18036,23488,28717,49499,0,0,6748,13462,20345,21024,20316,42444],"quality":"More is better","isMain":true,"metricName":"Total length (>= 1000 bp)"},{"values":[16456,18036,16661,13927,46474,0,0,0,13462,14746,14741,7262,42444],"quality":"More is better","isMain":false,"metricName":"Total length (>= 5000 bp)"},{"values":[16456,0,16661,0,30728,0,0,0,0,0,0,0,42444],"quality":"More is better","isMain":true,"metricName":"Total length (>= 10000 bp)"},{"values":[0,0,0,0,30728,0,0,0,0,0,0,0,42444],"quality":"More is better","isMain":false,"metricName":"Total length (>= 25000 bp)"},{"values":[0,0,0,0,0,0,0,0,0,0,0,0,0],"quality":"More is better","isMain":true,"metricName":"Total length (>= 50000 bp)"},{"values":[16456,8823,16661,3843,30728,680,575,4438,6765,6627,6563,3152,42444],"quality":"More is better","isMain":false,"metricName":"N50"},{"values":[4654,8823,4355,1447,8553,680,575,2310,6697,4438,4438,1091,42444],"quality":"More is better","isMain":false,"metricName":"N75"},{"values":[1,2,1,3,1,1,1,1,1,2,2,3,1],"quality":"Less is better","isMain":false,"metricName":"L50"},{"values":[2,2,2,6,2,1,1,2,2,3,3,6,1],"quality":"Less is better","isMain":false,"metricName":"L75"},{"values":["37.61","36.47","37.49","37.79","38.26","41.09","31.48","42.26","36.12","37.55","37.58","37.44","38.61"],"quality":"Equal","isMain":false,"metricName":"GC (%)"}]],["Predicted genes",[]],["Similarity statistics",[{"values":[0,0,0,0,0,0,0,0,0,0,0,0,0],"quality":"Equal","isMain":false,"metricName":"# similar correct contigs"},{"values":[0,0,0,0,0,0,0,0,0,0,0,0,0],"quality":"Equal","isMain":false,"metricName":"# similar misassembled blocks"}]],["Reference statistics",[{"values":[41796,41796,41796,41796,41796,41796,41796,41796,41796,41796,41796,41796,41796],"quality":"Equal","isMain":false,"metricName":"Reference length"},{"values":[1,1,1,1,1,1,1,1,1,1,1,1,1],"quality":"Equal","isMain":false,"metricName":"Reference fragments"},{"values":["38.57","38.57","38.57","38.57","38.57","38.57","38.57","38.57","38.57","38.57","38.57","38.57","38.57"],"quality":"Equal","isMain":false,"metricName":"Reference GC (%)"}]]],"referenceName":"gi\_18034800\_ref\_NC","date":"18 July 2018, Wednesday, 16:36:20","order":[0,1,2,3,4,5,6,7,8,9,10,11,12],"assembliesNames":["ABySS\_127","ABySS\_63","CLC","IDBA\_UD","MEGAHIT","MIRA","SOAPdenovo2","SPAdes","SPAdes\_meta","SPAdes\_sc","SPAdes\_sc\_careful","Velvet","Geneious"]},{"assembliesWithNs":null,"minContig":500,"report":[["Genome statistics",[{"values":["99.962","100.000","100.000","100.000","73.401","63.127","100.000","100.000","100.000","100.000","100.000"],"quality":"More is better","isMain":true,"metricName":"Genome fraction (%)"},{"values":["1.000","1.002","1.002","1.004","1.003","1.000","1.002","1.001","1.001","1.001","1.001"],"quality":"Less is better","isMain":true,"metricName":"Duplication ratio"},{"values":[57394,57515,57557,57647,7964,2649,57543,57471,57471,57471,57446],"quality":"More is better","isMain":true,"metricName":"Largest alignment"},{"values":[57394,57515,57557,57647,42280,36245,57543,57471,57471,57471,57446],"quality":"More is better","isMain":true,"metricName":"Total aligned length"},{"values":[57416,57515,57557,57647,4124,715,57543,57471,57471,57471,57446],"quality":"More is better","isMain":false,"metricName":"NG50"},{"values":[57416,57515,57557,57647,null,null,57543,57471,57471,57471,57446],"quality":"More is better","isMain":false,"metricName":"NG75"},{"values":[57394,57515,57557,57647,6842,1312,57543,57471,57471,57471,57446],"quality":"More is better","isMain":false,"metricName":"NA50"},{"values":[57394,57515,57557,57647,4124,670,57543,57471,57471,57471,57446],"quality":"More is better","isMain":false,"metricName":"NA75"},{"values":[57394,57515,57557,57647,4124,670,57543,57471,57471,57471,57446],"quality":"More is better","isMain":true,"metricName":"NGA50"},{"values":[57394,57515,57557,57647,null,null,57543,57471,57471,57471,57446],"quality":"More is better","isMain":false,"metricName":"NGA75"},{"values":[1,1,1,1,5,21,1,1,1,1,1],"quality":"Less is better","isMain":false,"metricName":"LG50"},{"values":[1,1,1,1,null,null,1,1,1,1,1],"quality":"Less is better","isMain":false,"metricName":"LG75"},{"values":[1,1,1,1,3,11,1,1,1,1,1],"quality":"Less is better","isMain":false,"metricName":"LA50"},{"values":[1,1,1,1,5,22,1,1,1,1,1],"quality":"Less is better","isMain":false,"metricName":"LA75"},{"values":[1,1,1,1,5,22,1,1,1,1,1],"quality":"Less is better","isMain":true,"metricName":"LGA50"},{"values":[1,1,1,1,null,null,1,1,1,1,1],"quality":"Less is better","isMain":false,"metricName":"LGA75"}]],["Misassemblies",[{"values":[0,0,0,0,0,0,0,0,0,0,0],"quality":"Less is better","isMain":true,"metricName":"# misassemblies"},{"values":[0,0,0,0,0,0,0,0,0,0,0],"quality":"Less is better","isMain":false,"metricName":" # relocations"},{"values":[0,0,0,0,0,0,0,0,0,0,0],"quality":"Less is better","isMain":false,"metricName":" # translocations"},{"values":[0,0,0,0,0,0,0,0,0,0,0],"quality":"Less is better","isMain":false,"metricName":" # inversions"},{"values":[0,0,0,0,0,0,0,0,0,0,0],"quality":"Less is better","isMain":false,"metricName":"# misassembled contigs"},{"values":[0,0,0,0,0,0,0,0,0,0,0],"quality":"Less is better","isMain":true,"metricName":"Misassembled contigs length"},{"values":[0,1,1,1,0,0,1,0,0,0,0],"quality":"Less is better","isMain":false,"metricName":"# local misassemblies"},{"values":[0,0,0,0,0,0,0,0,0,0,0],"quality":"Less is better","isMain":false,"metricName":"# unaligned mis. contigs"}]],["Unaligned",[{"values":[0,0,0,0,0,0,0,0,0,0,0],"quality":"Less is better","isMain":false,"metricName":"# fully unaligned contigs"},{"values":[0,0,0,0,0,0,0,0,0,0,0],"quality":"Less is better","isMain":false,"metricName":"Fully unaligned length"},{"values":[0,0,0,0,0,2,0,0,0,0,0],"quality":"Less is better","isMain":false,"metricName":"# partially unaligned contigs"},{"values":[0,0,0,0,0,1512,0,0,0,0,0],"quality":"Less is better","isMain":false,"metricName":"Partially unaligned length"}]],["Mismatches",[{"values":[0,0,0,0,0,0,0,0,0,0,0],"quality":"Less is better","isMain":false,"metricName":"# mismatches"},{"values":[0,0,0,0,0,0,0,1,1,1,1],"quality":"Less is better","isMain":false,"metricName":"# indels"},{"values":[0,0,0,0,0,0,0,55,55,55,30],"quality":"Less is better","isMain":false,"metricName":"Indels length"},{"values":["0.00","0.00","0.00","0.00","0.00","0.00","0.00","0.00","0.00","0.00","0.00"],"quality":"Less is better","isMain":true,"metricName":"# mismatches per 100 kbp"},{"values":["0.00","0.00","0.00","0.00","0.00","0.00","0.00","1.74","1.74","1.74","1.74"],"quality":"Less is better","isMain":true,"metricName":"# indels per 100 kbp"},{"values":[0,0,0,0,0,0,0,0,0,0,0],"quality":"Less is better","isMain":false,"metricName":" # indels (<= 5 bp)"},{"values":[0,0,0,0,0,0,0,1,1,1,1],"quality":"Less is better","isMain":false,"metricName":" # indels (> 5 bp)"},{"values":[0,0,0,0,0,388,0,0,0,0,0],"quality":"Less is better","isMain":false,"metricName":"# N's"},{"values":["0.00","0.00","0.00","0.00","0.00","1027.62","0.00","0.00","0.00","0.00","0.00"],"quality":"Less is better","isMain":true,"metricName":"# N's per 100 kbp"}]],["Statistics without reference",[{"values":[1,1,1,1,10,35,1,1,1,1,1],"quality":"Equal","isMain":true,"metricName":"# contigs"},{"values":[1,1,1,1,9,14,1,1,1,1,1],"quality":"Equal","isMain":false,"metricName":"# contigs (>= 1000 bp)"},{"values":[1,1,1,1,4,0,1,1,1,1,1],"quality":"Equal","isMain":false,"metricName":"# contigs (>= 5000 bp)"},{"values":[1,1,1,1,0,0,1,1,1,1,1],"quality":"Equal","isMain":false,"metricName":"# contigs (>= 10000 bp)"},{"values":[1,1,1,1,0,0,1,1,1,1,1],"quality":"Equal","isMain":false,"metricName":"# contigs (>= 25000 bp)"},{"values":[1,1,1,1,0,0,1,1,1,1,1],"quality":"Equal","isMain":false,"metricName":"# contigs (>= 50000 bp)"},{"values":[57416,57515,57557,57647,7964,2649,57543,57471,57471,57471,57446],"quality":"More is better","isMain":true,"metricName":"Largest contig"},{"values":[57416,57515,57557,57647,42280,37757,57543,57471,57471,57471,57446],"quality":"More is better","isMain":true,"metricName":"Total length"},{"values":[57416,57515,57557,57647,41733,23362,57543,57471,57471,57471,57446],"quality":"More is better","isMain":true,"metricName":"Total length (>= 1000 bp)"},{"values":[57416,57515,57557,57647,27824,0,57543,57471,57471,57471,57446],"quality":"More is better","isMain":false,"metricName":"Total length (>= 5000 bp)"},{"values":[57416,57515,57557,57647,0,0,57543,57471,57471,57471,57446],"quality":"More is better","isMain":true,"metricName":"Total length (>= 10000 bp)"},{"values":[57416,57515,57557,57647,0,0,57543,57471,57471,57471,57446],"quality":"More is better","isMain":false,"metricName":"Total length (>= 25000 bp)"},{"values":[57416,57515,57557,57647,0,0,57543,57471,57471,57471,57446],"quality":"More is better","isMain":true,"metricName":"Total length (>= 50000 bp)"},{"values":[57416,57515,57557,57647,6842,1410,57543,57471,57471,57471,57446],"quality":"More is better","isMain":false,"metricName":"N50"},{"values":[57416,57515,57557,57647,4124,815,57543,57471,57471,57471,57446],"quality":"More is better","isMain":false,"metricName":"N75"},{"values":[1,1,1,1,3,11,1,1,1,1,1],"quality":"Less is better","isMain":false,"metricName":"L50"},{"values":[1,1,1,1,5,20,1,1,1,1,1],"quality":"Less is better","isMain":false,"metricName":"L75"},{"values":["61.48","61.48","61.49","61.48","61.03","61.75","61.50","61.49","61.49","61.49","61.49"],"quality":"Equal","isMain":false,"metricName":"GC (%)"}]],["Predicted genes",[]],["Similarity statistics",[{"values":[0,0,0,0,0,0,0,0,0,0,0],"quality":"Equal","isMain":false,"metricName":"# similar correct contigs"},{"values":[0,0,0,0,0,0,0,0,0,0,0],"quality":"Equal","isMain":false,"metricName":"# similar misassembled blocks"}]],["Reference statistics",[{"values":[57416,57416,57416,57416,57416,57416,57416,57416,57416,57416,57416],"quality":"Equal","isMain":false,"metricName":"Reference length"},{"values":[1,1,1,1,1,1,1,1,1,1,1],"quality":"Equal","isMain":false,"metricName":"Reference fragments"},{"values":["61.49","61.49","61.49","61.49","61.49","61.49","61.49","61.49","61.49","61.49","61.49"],"quality":"Equal","isMain":false,"metricName":"Reference GC (%)"}]]],"referenceName":"gi\_18071213\_ref\_NC","date":"18 July 2018, Wednesday, 16:36:33","order":[0,1,2,3,4,5,6,7,8,9,10],"assembliesNames":["CLC","IDBA\_UD","MEGAHIT","MIRA","Ray\_Meta","SOAPdenovo2","SPAdes","SPAdes\_meta","SPAdes\_sc","SPAdes\_sc\_careful","Velvet"]},{"assembliesWithNs":null,"minContig":500,"report":[["Genome statistics",[{"values":["6.772","4.600","56.654","56.328","56.745","24.547","50.978","68.350","70.095","70.977","71.229","4.017","2.302"],"quality":"More is better","isMain":true,"metricName":"Genome fraction (%)"},{"values":["1.000","1.000","1.013","1.006","1.007","1.026","1.003","1.001","1.011","1.005","1.003","0.994","1.000"],"quality":"Less is better","isMain":true,"metricName":"Duplication ratio"},{"values":[811,855,2957,3532,3983,2118,2657,4176,4035,4176,4176,2171,2373],"quality":"More is better","isMain":true,"metricName":"Largest alignment"},{"values":[8863,6021,74647,73767,74791,32390,66814,89544,92255,92921,93259,5223,3013],"quality":"More is better","isMain":true,"metricName":"Total aligned length"},{"values":[null,null,871,695,704,null,536,1194,1133,1163,1163,null,null],"quality":"More is better","isMain":false,"metricName":"NG50"},{"values":[576,566,1407,1400,1290,1040,1025,1783,1508,1662,1662,755,2373],"quality":"More is better","isMain":false,"metricName":"NA50"},{"values":[546,520,909,822,876,829,765,1084,1025,1084,1084,599,640],"quality":"More is better","isMain":false,"metricName":"NA75"},{"values":[null,null,770,645,683,null,536,1137,1101,1163,1163,null,null],"quality":"More is better","isMain":true,"metricName":"NGA50"},{"values":[null,null,40,46,46,null,65,30,33,32,32,null,null],"quality":"Less is better","isMain":false,"metricName":"LG50"},{"values":[7,5,19,18,18,12,22,17,19,18,18,2,1],"quality":"Less is better","isMain":false,"metricName":"LA50"},{"values":[11,8,36,35,36,21,42,34,38,36,36,4,2],"quality":"Less is better","isMain":false,"metricName":"LA75"},{"values":[null,null,44,48,48,null,65,31,34,32,32,null,null],"quality":"Less is better","isMain":true,"metricName":"LGA50"}]],["Misassemblies",[{"values":[0,1,0,1,0,0,0,1,1,1,1,0,0],"quality":"Less is better","isMain":true,"metricName":"# misassemblies"},{"values":[0,1,0,1,0,0,0,1,1,1,1,0,0],"quality":"Less is better","isMain":false,"metricName":" # relocations"},{"values":[0,0,0,0,0,0,0,0,0,0,0,0,0],"quality":"Less is better","isMain":false,"metricName":" # translocations"},{"values":[0,0,0,0,0,0,0,0,0,0,0,0,0],"quality":"Less is better","isMain":false,"metricName":" # inversions"},{"values":[0,1,0,1,0,0,0,1,1,1,1,0,0],"quality":"Less is better","isMain":false,"metricName":"# misassembled contigs"},{"values":[0,551,0,1118,0,0,0,1308,996,1045,1018,0,0],"quality":"Less is better","isMain":true,"metricName":"Misassembled contigs length"},{"values":[0,0,0,1,1,1,0,2,0,0,0,0,0],"quality":"Less is better","isMain":false,"metricName":"# local misassemblies"},{"values":[0,0,0,0,0,0,0,0,0,0,0,0,0],"quality":"Less is better","isMain":false,"metricName":"# unaligned mis. contigs"}]],["Unaligned",[{"values":[0,0,0,0,0,0,0,0,0,0,0,0,0],"quality":"Less is better","isMain":false,"metricName":"# fully unaligned contigs"},{"values":[0,0,0,0,0,0,0,0,0,0,0,0,0],"quality":"Less is better","isMain":false,"metricName":"Fully unaligned length"},{"values":[0,0,4,1,1,0,0,1,0,0,0,0,1],"quality":"Less is better","isMain":false,"metricName":"# partially unaligned contigs"},{"values":[0,0,3508,506,888,0,0,1103,0,0,0,0,981],"quality":"Less is better","isMain":false,"metricName":"Partially unaligned length"}]],["Mismatches",[{"values":[1,3,263,243,226,51,127,338,286,241,226,8,3],"quality":"Less is better","isMain":false,"metricName":"# mismatches"},{"values":[0,1,19,6,6,0,5,10,12,11,12,3,0],"quality":"Less is better","isMain":false,"metricName":"# indels"},{"values":[0,1,45,40,21,0,69,56,92,76,84,48,0],"quality":"Less is better","isMain":false,"metricName":"Indels length"},{"values":["11.28","49.83","354.72","329.64","304.33","158.76","190.36","377.87","311.77","259.46","242.45","152.18","99.57"],"quality":"Less is better","isMain":true,"metricName":"# mismatches per 100 kbp"},{"values":["0.00","16.61","25.63","8.14","8.08","0.00","7.49","11.18","13.08","11.84","12.87","57.07","0.00"],"quality":"Less is better","isMain":true,"metricName":"# indels per 100 kbp"},{"values":[0,1,17,4,5,0,2,7,6,6,6,0,0],"quality":"Less is better","isMain":false,"metricName":" # indels (<= 5 bp)"},{"values":[0,0,2,2,1,0,3,3,6,5,6,3,0],"quality":"Less is better","isMain":false,"metricName":" # indels (> 5 bp)"},{"values":[0,0,0,0,0,85,106,0,0,0,0,0,13],"quality":"Less is better","isMain":false,"metricName":"# N's"},{"values":["0.00","0.00","0.00","0.00","0.00","257.80","158.40","0.00","0.00","0.00","0.00","0.00","325.49"],"quality":"Less is better","isMain":true,"metricName":"# N's per 100 kbp"}]],["Statistics without reference",[{"values":[15,10,59,61,63,33,67,60,68,66,66,6,2],"quality":"Equal","isMain":true,"metricName":"# contigs"},{"values":[0,0,33,29,27,13,22,38,38,40,40,1,2],"quality":"Equal","isMain":false,"metricName":"# contigs (>= 1000 bp)"},{"values":[0,0,0,0,0,0,0,0,0,0,0,0,0],"quality":"Equal","isMain":false,"metricName":"# contigs (>= 5000 bp)"},{"values":[0,0,0,0,0,0,0,0,0,0,0,0,0],"quality":"Equal","isMain":false,"metricName":"# contigs (>= 10000 bp)"},{"values":[0,0,0,0,0,0,0,0,0,0,0,0,0],"quality":"Equal","isMain":false,"metricName":"# contigs (>= 25000 bp)"},{"values":[0,0,0,0,0,0,0,0,0,0,0,0,0],"quality":"Equal","isMain":false,"metricName":"# contigs (>= 50000 bp)"},{"values":[811,855,4198,3532,3983,2471,2657,4742,4035,4176,4176,2171,2373],"quality":"More is better","isMain":true,"metricName":"Largest contig"},{"values":[8863,6021,78609,74638,75679,32971,66920,90685,92728,93328,93451,5223,3994],"quality":"More is better","isMain":true,"metricName":"Total length"},{"values":[0,0,59631,51943,49690,17978,33625,74331,70806,74302,74276,2171,3994],"quality":"More is better","isMain":true,"metricName":"Total length (>= 1000 bp)"},{"values":[0,0,0,0,0,0,0,0,0,0,0,0,0],"quality":"More is better","isMain":false,"metricName":"Total length (>= 5000 bp)"},{"values":[0,0,0,0,0,0,0,0,0,0,0,0,0],"quality":"More is better","isMain":true,"metricName":"Total length (>= 10000 bp)"},{"values":[0,0,0,0,0,0,0,0,0,0,0,0,0],"quality":"More is better","isMain":false,"metricName":"Total length (>= 25000 bp)"},{"values":[0,0,0,0,0,0,0,0,0,0,0,0,0],"quality":"More is better","isMain":true,"metricName":"Total length (>= 50000 bp)"},{"values":[576,566,1541,1491,1329,1040,1025,1796,1530,1662,1662,755,2373],"quality":"More is better","isMain":false,"metricName":"N50"},{"values":[546,538,1025,894,896,837,765,1133,1057,1084,1084,599,1621],"quality":"More is better","isMain":false,"metricName":"N75"},{"values":[7,5,17,17,17,12,22,16,19,18,18,2,1],"quality":"Less is better","isMain":false,"metricName":"L50"},{"values":[11,8,33,34,35,21,42,33,37,36,36,4,2],"quality":"Less is better","isMain":false,"metricName":"L75"},{"values":["38.51","37.34","39.79","39.49","39.72","40.25","39.43","39.40","39.42","39.20","39.23","38.54","39.39"],"quality":"Equal","isMain":false,"metricName":"GC (%)"}]],["Predicted genes",[]],["Similarity statistics",[{"values":[0,0,0,0,0,0,0,0,0,0,0,0,0],"quality":"Equal","isMain":false,"metricName":"# similar correct contigs"},{"values":[0,0,0,0,0,0,0,0,0,0,0,0,0],"quality":"Equal","isMain":false,"metricName":"# similar misassembled blocks"}]],["Reference statistics",[{"values":[130869,130869,130869,130869,130869,130869,130869,130869,130869,130869,130869,130869,130869],"quality":"Equal","isMain":false,"metricName":"Reference length"},{"values":[1,1,1,1,1,1,1,1,1,1,1,1,1],"quality":"Equal","isMain":false,"metricName":"Reference fragments"},{"values":["39.08","39.08","39.08","39.08","39.08","39.08","39.08","39.08","39.08","39.08","39.08","39.08","39.08"],"quality":"Equal","isMain":false,"metricName":"Reference GC (%)"}]]],"referenceName":"gi\_18138257\_ref\_NC","date":"18 July 2018, Wednesday, 16:36:48","order":[0,1,2,3,4,5,6,7,8,9,10,11,12],"assembliesNames":["ABySS\_127","ABySS\_63","CLC","IDBA\_UD","MEGAHIT","MIRA","SOAPdenovo2","SPAdes","SPAdes\_meta","SPAdes\_sc","SPAdes\_sc\_careful","Velvet","Geneious"]},{"assembliesWithNs":null,"minContig":500,"report":[["Genome statistics",[{"values":["100.000","99.154","100.000","99.248","100.000","100.000","100.000","7.812","100.000","99.638","100.000","100.000","94.988"],"quality":"More is better","isMain":true,"metricName":"Genome fraction (%)"},{"values":["1.002","1.001","1.000","1.000","1.003","1.022","1.010","1.000","1.002","1.000","1.001","1.001","1.001"],"quality":"Less is better","isMain":true,"metricName":"Duplication ratio"},{"values":[42679,41683,42575,42255,42709,42951,43011,804,42681,42421,42630,42630,15624],"quality":"More is better","isMain":true,"metricName":"Largest alignment"},{"values":[42679,42256,42575,42255,42709,43518,43011,3326,42681,42421,42630,42630,40494],"quality":"More is better","isMain":true,"metricName":"Total aligned length"},{"values":[42679,41683,42575,42255,42709,42951,43011,null,42681,42421,42630,42630,9791],"quality":"More is better","isMain":false,"metricName":"NG50"},{"values":[42679,41683,42575,42255,42709,42951,43011,null,42681,42421,42630,42630,2760],"quality":"More is better","isMain":false,"metricName":"NG75"},{"values":[42679,41683,42575,42255,42709,42951,43011,603,42681,42421,42630,42630,9791],"quality":"More is better","isMain":false,"metricName":"NA50"},{"values":[42679,41683,42575,42255,42709,42951,43011,586,42681,42421,42630,42630,5823],"quality":"More is better","isMain":false,"metricName":"NA75"},{"values":[42679,41683,42575,42255,42709,42951,43011,null,42681,42421,42630,42630,9791],"quality":"More is better","isMain":true,"metricName":"NGA50"},{"values":[42679,41683,42575,42255,42709,42951,43011,null,42681,42421,42630,42630,2760],"quality":"More is better","isMain":false,"metricName":"NGA75"},{"values":[1,1,1,1,1,1,1,null,1,1,1,1,2],"quality":"Less is better","isMain":false,"metricName":"LG50"},{"values":[1,1,1,1,1,1,1,null,1,1,1,1,4],"quality":"Less is better","isMain":false,"metricName":"LG75"},{"values":[1,1,1,1,1,1,1,3,1,1,1,1,2],"quality":"Less is better","isMain":false,"metricName":"LA50"},{"values":[1,1,1,1,1,1,1,4,1,1,1,1,3],"quality":"Less is better","isMain":false,"metricName":"LA75"},{"values":[1,1,1,1,1,1,1,null,1,1,1,1,2],"quality":"Less is better","isMain":true,"metricName":"LGA50"},{"values":[1,1,1,1,1,1,1,null,1,1,1,1,4],"quality":"Less is better","isMain":false,"metricName":"LGA75"}]],["Misassemblies",[{"values":[0,0,0,0,0,0,0,0,0,0,0,0,0],"quality":"Less is better","isMain":true,"metricName":"# misassemblies"},{"values":[0,0,0,0,0,0,0,0,0,0,0,0,0],"quality":"Less is better","isMain":false,"metricName":" # relocations"},{"values":[0,0,0,0,0,0,0,0,0,0,0,0,0],"quality":"Less is better","isMain":false,"metricName":" # translocations"},{"values":[0,0,0,0,0,0,0,0,0,0,0,0,0],"quality":"Less is better","isMain":false,"metricName":" # inversions"},{"values":[0,0,0,0,0,0,0,0,0,0,0,0,0],"quality":"Less is better","isMain":false,"metricName":"# misassembled contigs"},{"values":[0,0,0,0,0,0,0,0,0,0,0,0,0],"quality":"Less is better","isMain":true,"metricName":"Misassembled contigs length"},{"values":[1,0,0,0,1,1,1,0,1,0,0,0,0],"quality":"Less is better","isMain":false,"metricName":"# local misassemblies"},{"values":[0,0,0,0,0,0,0,0,0,0,0,0,0],"quality":"Less is better","isMain":false,"metricName":"# unaligned mis. contigs"}]],["Unaligned",[{"values":[0,0,0,0,0,0,0,0,0,0,0,0,0],"quality":"Less is better","isMain":false,"metricName":"# fully unaligned contigs"},{"values":[0,0,0,0,0,0,0,0,0,0,0,0,0],"quality":"Less is better","isMain":false,"metricName":"Fully unaligned length"},{"values":[0,0,0,0,0,0,0,0,0,0,0,0,0],"quality":"Less is better","isMain":false,"metricName":"# partially unaligned contigs"},{"values":[0,0,0,0,0,0,0,0,0,0,0,0,0],"quality":"Less is better","isMain":false,"metricName":"Partially unaligned length"}]],["Mismatches",[{"values":[0,0,0,0,0,1,0,0,0,0,0,0,0],"quality":"Less is better","isMain":false,"metricName":"# mismatches"},{"values":[0,0,0,0,0,0,0,0,0,0,1,1,0],"quality":"Less is better","isMain":false,"metricName":"# indels"},{"values":[0,0,0,0,0,0,0,0,0,0,55,55,0],"quality":"Less is better","isMain":false,"metricName":"Indels length"},{"values":["0.00","0.00","0.00","0.00","0.00","2.35","0.00","0.00","0.00","0.00","0.00","0.00","0.00"],"quality":"Less is better","isMain":true,"metricName":"# mismatches per 100 kbp"},{"values":["0.00","0.00","0.00","0.00","0.00","0.00","0.00","0.00","0.00","0.00","2.35","2.35","0.00"],"quality":"Less is better","isMain":true,"metricName":"# indels per 100 kbp"},{"values":[0,0,0,0,0,0,0,0,0,0,0,0,0],"quality":"Less is better","isMain":false,"metricName":" # indels (<= 5 bp)"},{"values":[0,0,0,0,0,0,0,0,0,0,1,1,0],"quality":"Less is better","isMain":false,"metricName":" # indels (> 5 bp)"},{"values":[0,0,0,0,0,0,0,0,0,0,0,0,0],"quality":"Less is better","isMain":false,"metricName":"# N's"},{"values":["0.00","0.00","0.00","0.00","0.00","0.00","0.00","0.00","0.00","0.00","0.00","0.00","0.00"],"quality":"Less is better","isMain":true,"metricName":"# N's per 100 kbp"}]],["Statistics without reference",[{"values":[1,2,1,1,1,2,1,5,1,1,1,1,9],"quality":"Equal","isMain":true,"metricName":"# contigs"},{"values":[1,1,1,1,1,1,1,0,1,1,1,1,6],"quality":"Equal","isMain":false,"metricName":"# contigs (>= 1000 bp)"},{"values":[1,1,1,1,1,1,1,0,1,1,1,1,3],"quality":"Equal","isMain":false,"metricName":"# contigs (>= 5000 bp)"},{"values":[1,1,1,1,1,1,1,0,1,1,1,1,1],"quality":"Equal","isMain":false,"metricName":"# contigs (>= 10000 bp)"},{"values":[1,1,1,1,1,1,1,0,1,1,1,1,0],"quality":"Equal","isMain":false,"metricName":"# contigs (>= 25000 bp)"},{"values":[0,0,0,0,0,0,0,0,0,0,0,0,0],"quality":"Equal","isMain":false,"metricName":"# contigs (>= 50000 bp)"},{"values":[42679,41683,42575,42255,42709,42951,43011,804,42681,42421,42630,42630,15624],"quality":"More is better","isMain":true,"metricName":"Largest contig"},{"values":[42679,42256,42575,42255,42709,43518,43011,3326,42681,42421,42630,42630,40494],"quality":"More is better","isMain":true,"metricName":"Total length"},{"values":[42679,41683,42575,42255,42709,42951,43011,0,42681,42421,42630,42630,38270],"quality":"More is better","isMain":true,"metricName":"Total length (>= 1000 bp)"},{"values":[42679,41683,42575,42255,42709,42951,43011,0,42681,42421,42630,42630,31238],"quality":"More is better","isMain":false,"metricName":"Total length (>= 5000 bp)"},{"values":[42679,41683,42575,42255,42709,42951,43011,0,42681,42421,42630,42630,15624],"quality":"More is better","isMain":true,"metricName":"Total length (>= 10000 bp)"},{"values":[42679,41683,42575,42255,42709,42951,43011,0,42681,42421,42630,42630,0],"quality":"More is better","isMain":false,"metricName":"Total length (>= 25000 bp)"},{"values":[0,0,0,0,0,0,0,0,0,0,0,0,0],"quality":"More is better","isMain":true,"metricName":"Total length (>= 50000 bp)"},{"values":[42679,41683,42575,42255,42709,42951,43011,603,42681,42421,42630,42630,9791],"quality":"More is better","isMain":false,"metricName":"N50"},{"values":[42679,41683,42575,42255,42709,42951,43011,586,42681,42421,42630,42630,5823],"quality":"More is better","isMain":false,"metricName":"N75"},{"values":[1,1,1,1,1,1,1,3,1,1,1,1,2],"quality":"Less is better","isMain":false,"metricName":"L50"},{"values":[1,1,1,1,1,1,1,4,1,1,1,1,3],"quality":"Less is better","isMain":false,"metricName":"L75"},{"values":["49.35","49.37","49.35","49.36","49.35","49.27","49.36","49.94","49.35","49.34","49.34","49.34","49.41"],"quality":"Equal","isMain":false,"metricName":"GC (%)"}]],["Predicted genes",[]],["Similarity statistics",[{"values":[0,0,0,0,0,0,0,0,0,0,0,0,0],"quality":"Equal","isMain":false,"metricName":"# similar correct contigs"},{"values":[0,0,0,0,0,0,0,0,0,0,0,0,0],"quality":"Equal","isMain":false,"metricName":"# similar misassembled blocks"}]],["Reference statistics",[{"values":[42575,42575,42575,42575,42575,42575,42575,42575,42575,42575,42575,42575,42575],"quality":"Equal","isMain":false,"metricName":"Reference length"},{"values":[1,1,1,1,1,1,1,1,1,1,1,1,1],"quality":"Equal","isMain":false,"metricName":"Reference fragments"},{"values":["49.35","49.35","49.35","49.35","49.35","49.35","49.35","49.35","49.35","49.35","49.35","49.35","49.35"],"quality":"Equal","isMain":false,"metricName":"Reference GC (%)"}]]],"referenceName":"gi\_18249864\_ref\_NC","date":"18 July 2018, Wednesday, 16:37:01","order":[0,1,2,3,4,5,6,7,8,9,10,11,12],"assembliesNames":["ABySS\_127","ABySS\_63","CLC","IDBA\_UD","MEGAHIT","MIRA","Ray\_Meta","SOAPdenovo2","SPAdes","SPAdes\_meta","SPAdes\_sc","SPAdes\_sc\_careful","Velvet"]},{"assembliesWithNs":null,"minContig":500,"report":[["Genome statistics",[{"values":["100.000","99.998","99.996","100.000","100.000","100.000","99.833","100.000","100.000","87.060","100.000"],"quality":"More is better","isMain":true,"metricName":"Genome fraction (%)"},{"values":["1.000","1.012","1.000","34.623","1.009","1.002","1.000","1.001","1.001","1.002","1.197"],"quality":"Less is better","isMain":true,"metricName":"Duplication ratio"},{"values":[55597,18919,55595,33288,29292,55724,55504,55641,55652,3730,34242],"quality":"More is better","isMain":true,"metricName":"Largest alignment"},{"values":[55597,56289,55595,1924818,56116,55724,55504,55641,55652,48455,66567],"quality":"More is better","isMain":true,"metricName":"Total aligned length"},{"values":[55597,13366,55595,60011,29292,55724,55504,55641,55652,1543,66567],"quality":"More is better","isMain":false,"metricName":"NG50"},{"values":[55597,10970,55595,60011,21923,55724,55504,55641,55652,944,66567],"quality":"More is better","isMain":false,"metricName":"NG75"},{"values":[55597,13366,55595,2424,29292,55724,55504,55641,55652,1852,34242],"quality":"More is better","isMain":false,"metricName":"NA50"},{"values":[55597,10970,55595,1770,21923,55724,55504,55641,55652,1169,32325],"quality":"More is better","isMain":false,"metricName":"NA75"},{"values":[55597,13366,55595,33288,29292,55724,55504,55641,55652,1543,34242],"quality":"More is better","isMain":true,"metricName":"NGA50"},{"values":[55597,10970,55595,26723,21923,55724,55504,55641,55652,944,32325],"quality":"More is better","isMain":false,"metricName":"NGA75"},{"values":[1,2,1,1,1,1,1,1,1,11,1],"quality":"Less is better","isMain":false,"metricName":"LG50"},{"values":[1,3,1,1,2,1,1,1,1,23,1],"quality":"Less is better","isMain":false,"metricName":"LG75"},{"values":[1,2,1,248,1,1,1,1,1,9,1],"quality":"Less is better","isMain":false,"metricName":"LA50"},{"values":[1,3,1,480,2,1,1,1,1,18,2],"quality":"Less is better","isMain":false,"metricName":"LA75"},{"values":[1,2,1,1,1,1,1,1,1,11,1],"quality":"Less is better","isMain":true,"metricName":"LGA50"},{"values":[1,3,1,2,2,1,1,1,1,23,2],"quality":"Less is better","isMain":false,"metricName":"LGA75"}]],["Misassemblies",[{"values":[0,0,0,1,0,0,0,0,0,0,1],"quality":"Less is better","isMain":true,"metricName":"# misassemblies"},{"values":[0,0,0,1,0,0,0,0,0,0,1],"quality":"Less is better","isMain":false,"metricName":" # relocations"},{"values":[0,0,0,0,0,0,0,0,0,0,0],"quality":"Less is better","isMain":false,"metricName":" # translocations"},{"values":[0,0,0,0,0,0,0,0,0,0,0],"quality":"Less is better","isMain":false,"metricName":" # inversions"},{"values":[0,0,0,1,0,0,0,0,0,0,1],"quality":"Less is better","isMain":false,"metricName":"# misassembled contigs"},{"values":[0,0,0,60011,0,0,0,0,0,0,66567],"quality":"Less is better","isMain":true,"metricName":"Misassembled contigs length"},{"values":[0,0,0,0,0,1,0,0,0,0,0],"quality":"Less is better","isMain":false,"metricName":"# local misassemblies"},{"values":[0,0,0,0,0,0,0,0,0,0,0],"quality":"Less is better","isMain":false,"metricName":"# unaligned mis. contigs"}]],["Unaligned",[{"values":[0,0,0,0,0,0,0,0,0,0,0],"quality":"Less is better","isMain":false,"metricName":"# fully unaligned contigs"},{"values":[0,0,0,0,0,0,0,0,0,0,0],"quality":"Less is better","isMain":false,"metricName":"Fully unaligned length"},{"values":[0,0,0,0,0,0,0,0,0,0,0],"quality":"Less is better","isMain":false,"metricName":"# partially unaligned contigs"},{"values":[0,0,0,0,0,0,0,0,0,0,0],"quality":"Less is better","isMain":false,"metricName":"Partially unaligned length"}]],["Mismatches",[{"values":[0,0,1,87,0,1,4,1,0,35,9],"quality":"Less is better","isMain":false,"metricName":"# mismatches"},{"values":[0,0,0,3,0,0,0,1,1,0,0],"quality":"Less is better","isMain":false,"metricName":"# indels"},{"values":[0,0,0,3,0,0,0,44,55,0,0],"quality":"Less is better","isMain":false,"metricName":"Indels length"},{"values":["0.00","0.00","1.80","156.48","0.00","1.80","7.21","1.80","0.00","72.31","16.19"],"quality":"Less is better","isMain":true,"metricName":"# mismatches per 100 kbp"},{"values":["0.00","0.00","0.00","5.40","0.00","0.00","0.00","1.80","1.80","0.00","0.00"],"quality":"Less is better","isMain":true,"metricName":"# indels per 100 kbp"},{"values":[0,0,0,3,0,0,0,0,0,0,0],"quality":"Less is better","isMain":false,"metricName":" # indels (<= 5 bp)"},{"values":[0,0,0,0,0,0,0,1,1,0,0],"quality":"Less is better","isMain":false,"metricName":" # indels (> 5 bp)"},{"values":[1,0,0,2042,0,0,0,0,1,0,39],"quality":"Less is better","isMain":false,"metricName":"# N's"},{"values":["1.80","0.00","0.00","106.08","0.00","0.00","0.00","0.00","1.80","0.00","58.59"],"quality":"Less is better","isMain":true,"metricName":"# N's per 100 kbp"}]],["Statistics without reference",[{"values":[1,8,1,831,3,1,1,1,1,32,1],"quality":"Equal","isMain":true,"metricName":"# contigs"},{"values":[1,8,1,792,3,1,1,1,1,22,1],"quality":"Equal","isMain":false,"metricName":"# contigs (>= 1000 bp)"},{"values":[1,3,1,32,2,1,1,1,1,0,1],"quality":"Equal","isMain":false,"metricName":"# contigs (>= 5000 bp)"},{"values":[1,3,1,5,2,1,1,1,1,0,1],"quality":"Equal","isMain":false,"metricName":"# contigs (>= 10000 bp)"},{"values":[1,0,1,1,1,1,1,1,1,0,1],"quality":"Equal","isMain":false,"metricName":"# contigs (>= 25000 bp)"},{"values":[1,0,1,1,0,1,1,1,1,0,1],"quality":"Equal","isMain":false,"metricName":"# contigs (>= 50000 bp)"},{"values":[55597,18919,55595,60011,29292,55724,55504,55641,55652,3730,66567],"quality":"More is better","isMain":true,"metricName":"Largest contig"},{"values":[55597,56289,55595,1924934,56116,55724,55504,55641,55652,48509,66567],"quality":"More is better","isMain":true,"metricName":"Total length"},{"values":[55597,56289,55595,1891469,56116,55724,55504,55641,55652,41371,66567],"quality":"More is better","isMain":true,"metricName":"Total length (>= 1000 bp)"},{"values":[55597,43255,55595,274326,51215,55724,55504,55641,55652,0,66567],"quality":"More is better","isMain":false,"metricName":"Total length (>= 5000 bp)"},{"values":[55597,43255,55595,115070,51215,55724,55504,55641,55652,0,66567],"quality":"More is better","isMain":true,"metricName":"Total length (>= 10000 bp)"},{"values":[55597,0,55595,60011,29292,55724,55504,55641,55652,0,66567],"quality":"More is better","isMain":false,"metricName":"Total length (>= 25000 bp)"},{"values":[55597,0,55595,60011,0,55724,55504,55641,55652,0,66567],"quality":"More is better","isMain":true,"metricName":"Total length (>= 50000 bp)"},{"values":[55597,13366,55595,2424,29292,55724,55504,55641,55652,1852,66567],"quality":"More is better","isMain":false,"metricName":"N50"},{"values":[55597,10970,55595,1770,21923,55724,55504,55641,55652,1169,66567],"quality":"More is better","isMain":false,"metricName":"N75"},{"values":[1,2,1,247,1,1,1,1,1,9,1],"quality":"Less is better","isMain":false,"metricName":"L50"},{"values":[1,3,1,479,2,1,1,1,1,18,1],"quality":"Less is better","isMain":false,"metricName":"L75"},{"values":["33.69","33.69","33.69","33.73","33.74","33.69","33.69","33.68","33.69","33.76","33.21"],"quality":"Equal","isMain":false,"metricName":"GC (%)"}]],["Predicted genes",[]],["Similarity statistics",[{"values":[0,0,0,0,0,0,0,0,0,0,0],"quality":"Equal","isMain":false,"metricName":"# similar correct contigs"},{"values":[0,0,0,0,0,0,0,0,0,0,0],"quality":"Equal","isMain":false,"metricName":"# similar misassembled blocks"}]],["Reference statistics",[{"values":[55597,55597,55597,55597,55597,55597,55597,55597,55597,55597,55597],"quality":"Equal","isMain":false,"metricName":"Reference length"},{"values":[1,1,1,1,1,1,1,1,1,1,1],"quality":"Equal","isMain":false,"metricName":"Reference fragments"},{"values":["33.69","33.69","33.69","33.69","33.69","33.69","33.69","33.69","33.69","33.69","33.69"],"quality":"Equal","isMain":false,"metricName":"Reference GC (%)"}]]],"referenceName":"gi\_182637477\_ref\_NC","date":"18 July 2018, Wednesday, 16:37:15","order":[0,1,2,3,4,5,6,7,8,9,10],"assembliesNames":["CLC","IDBA\_UD","MEGAHIT","MIRA","Ray\_Meta","SPAdes","SPAdes\_meta","SPAdes\_sc","SPAdes\_sc\_careful","Velvet","Geneious"]},{"assembliesWithNs":null,"minContig":500,"report":[["Genome statistics",[{"values":["93.717","87.616","96.103","95.224","99.088","99.208","50.607","99.793","70.776","93.176","93.176","69.439","89.524"],"quality":"More is better","isMain":true,"metricName":"Genome fraction (%)"},{"values":["1.004","1.007","1.013","1.003","1.004","1.011","1.092","1.002","1.000","1.007","1.007","1.003","1.000"],"quality":"Less is better","isMain":true,"metricName":"Duplication ratio"},{"values":[11699,8893,27154,29491,42241,34751,5722,63518,63649,15594,15594,9031,53488],"quality":"More is better","isMain":true,"metricName":"Largest alignment"},{"values":[104812,98255,108457,106426,110857,111682,57025,111370,78851,104529,104529,77593,99714],"quality":"More is better","isMain":true,"metricName":"Total aligned length"},{"values":[3267,1876,10059,12292,20976,30775,1005,63518,74507,3969,3969,1000,34751],"quality":"More is better","isMain":false,"metricName":"NG50"},{"values":[2007,1226,5599,7715,9580,18182,null,21541,14523,2042,2042,null,34751],"quality":"More is better","isMain":false,"metricName":"NG75"},{"values":[3578,2161,10059,12292,20976,30775,950,63518,63649,7906,7906,1801,53488],"quality":"More is better","isMain":false,"metricName":"NA50"},{"values":[2079,1379,5599,7715,9580,11475,455,21541,7022,2399,2399,979,34751],"quality":"More is better","isMain":false,"metricName":"NA75"},{"values":[3267,1876,10059,12292,20976,30775,304,63518,63649,3969,3969,1000,34751],"quality":"More is better","isMain":true,"metricName":"NGA50"},{"values":[2007,1226,5599,7715,9580,18182,null,21541,null,2042,2042,null,34751],"quality":"More is better","isMain":false,"metricName":"NGA75"},{"values":[9,16,4,3,2,2,25,1,1,5,5,22,2],"quality":"Less is better","isMain":false,"metricName":"LG50"},{"values":[20,35,7,6,4,3,null,2,2,15,15,null,2],"quality":"Less is better","isMain":false,"metricName":"LG75"},{"values":[8,13,4,3,2,2,17,1,1,4,4,11,1],"quality":"Less is better","isMain":false,"metricName":"LA50"},{"values":[18,27,7,6,4,4,41,2,3,13,13,25,2],"quality":"Less is better","isMain":false,"metricName":"LA75"},{"values":[9,16,4,3,2,2,46,1,1,5,5,22,2],"quality":"Less is better","isMain":true,"metricName":"LGA50"},{"values":[20,35,7,6,4,3,null,2,null,15,15,null,2],"quality":"Less is better","isMain":false,"metricName":"LGA75"}]],["Misassemblies",[{"values":[0,0,0,0,0,0,0,0,0,0,0,0,0],"quality":"Less is better","isMain":true,"metricName":"# misassemblies"},{"values":[0,0,0,0,0,0,0,0,0,0,0,0,0],"quality":"Less is better","isMain":false,"metricName":" # relocations"},{"values":[0,0,0,0,0,0,0,0,0,0,0,0,0],"quality":"Less is better","isMain":false,"metricName":" # translocations"},{"values":[0,0,0,0,0,0,0,0,0,0,0,0,0],"quality":"Less is better","isMain":false,"metricName":" # inversions"},{"values":[0,0,0,0,0,0,0,0,0,0,0,0,0],"quality":"Less is better","isMain":false,"metricName":"# misassembled contigs"},{"values":[0,0,0,0,0,0,0,0,0,0,0,0,0],"quality":"Less is better","isMain":true,"metricName":"Misassembled contigs length"},{"values":[0,0,0,0,0,0,0,0,4,0,0,0,0],"quality":"Less is better","isMain":false,"metricName":"# local misassemblies"},{"values":[0,0,0,0,0,0,4,0,0,0,0,0,0],"quality":"Less is better","isMain":false,"metricName":"# unaligned mis. contigs"}]],["Unaligned",[{"values":[0,0,0,0,0,0,0,0,0,0,0,0,0],"quality":"Less is better","isMain":false,"metricName":"# fully unaligned contigs"},{"values":[0,0,0,0,0,0,0,0,0,0,0,0,0],"quality":"Less is better","isMain":false,"metricName":"Fully unaligned length"},{"values":[0,0,0,0,0,0,7,0,2,0,0,0,0],"quality":"Less is better","isMain":false,"metricName":"# partially unaligned contigs"},{"values":[0,0,0,0,0,0,10000,0,17201,0,0,0,0],"quality":"Less is better","isMain":false,"metricName":"Partially unaligned length"}]],["Mismatches",[{"values":[7,2,49,5,10,6,3,17,673,6,4,5,3],"quality":"Less is better","isMain":false,"metricName":"# mismatches"},{"values":[0,0,2,0,0,0,39,0,21,0,0,0,0],"quality":"Less is better","isMain":false,"metricName":"# indels"},{"values":[0,0,2,0,0,0,1890,0,37,0,0,0,0],"quality":"Less is better","isMain":false,"metricName":"Indels length"},{"values":["6.71","2.05","45.78","4.71","9.06","5.43","5.32","15.29","853.71","5.78","3.85","6.46","3.01"],"quality":"Less is better","isMain":true,"metricName":"# mismatches per 100 kbp"},{"values":["0.00","0.00","1.87","0.00","0.00","0.00","69.19","0.00","26.64","0.00","0.00","0.00","0.00"],"quality":"Less is better","isMain":true,"metricName":"# indels per 100 kbp"},{"values":[0,0,2,0,0,0,0,0,19,0,0,0,0],"quality":"Less is better","isMain":false,"metricName":" # indels (<= 5 bp)"},{"values":[0,0,0,0,0,0,39,0,2,0,0,0,0],"quality":"Less is better","isMain":false,"metricName":" # indels (> 5 bp)"},{"values":[0,0,0,0,0,18,7071,0,0,0,0,0,20],"quality":"Less is better","isMain":false,"metricName":"# N's"},{"values":["0.00","0.00","0.00","0.00","0.00","16.12","9885.09","0.00","0.00","0.00","0.00","0.00","20.06"],"quality":"Less is better","isMain":true,"metricName":"# N's per 100 kbp"}]],["Statistics without reference",[{"values":[36,52,20,18,12,7,46,5,3,34,34,51,3],"quality":"Equal","isMain":true,"metricName":"# contigs"},{"values":[32,39,12,16,10,6,25,5,3,26,26,23,3],"quality":"Equal","isMain":false,"metricName":"# contigs (>= 1000 bp)"},{"values":[7,3,8,6,6,6,1,4,3,4,4,2,3],"quality":"Equal","isMain":false,"metricName":"# contigs (>= 5000 bp)"},{"values":[1,0,4,3,3,5,0,3,2,3,3,0,3],"quality":"Equal","isMain":false,"metricName":"# contigs (>= 10000 bp)"},{"values":[0,0,1,1,1,2,0,1,1,0,0,0,2],"quality":"Equal","isMain":false,"metricName":"# contigs (>= 25000 bp)"},{"values":[0,0,0,0,0,0,0,1,1,0,0,0,1],"quality":"Equal","isMain":false,"metricName":"# contigs (>= 50000 bp)"},{"values":[11699,8893,27154,29491,42241,34751,6218,63518,74507,15594,15594,9031,53488],"quality":"More is better","isMain":true,"metricName":"Largest contig"},{"values":[104812,98255,108457,106427,110860,111682,71532,111370,96052,104529,104529,77593,99714],"quality":"More is better","isMain":true,"metricName":"Total length"},{"values":[101539,88719,102720,105285,109276,110982,56336,111370,96052,98701,98701,56980,99714],"quality":"More is better","isMain":true,"metricName":"Total length (>= 1000 bp)"},{"values":[51023,21050,90559,86937,101367,110982,6218,109741,96052,52991,52991,15215,99714],"quality":"More is better","isMain":false,"metricName":"Total length (>= 5000 bp)"},{"values":[11699,0,62952,62059,77473,105969,0,100451,89030,45085,45085,0,99714],"quality":"More is better","isMain":true,"metricName":"Total length (>= 10000 bp)"},{"values":[0,0,27154,29491,42241,65526,0,63518,74507,0,0,0,88239],"quality":"More is better","isMain":false,"metricName":"Total length (>= 25000 bp)"},{"values":[0,0,0,0,0,0,0,63518,74507,0,0,0,53488],"quality":"More is better","isMain":true,"metricName":"Total length (>= 50000 bp)"},{"values":[3578,2161,10059,12292,20976,30775,1981,63518,74507,7906,7906,1801,53488],"quality":"More is better","isMain":false,"metricName":"N50"},{"values":[2079,1379,5599,7715,9580,11475,1105,21541,74507,2399,2399,979,34751],"quality":"More is better","isMain":false,"metricName":"N75"},{"values":[8,13,4,3,2,2,11,1,1,4,4,11,1],"quality":"Less is better","isMain":false,"metricName":"L50"},{"values":[18,27,7,6,4,4,23,2,1,13,13,25,2],"quality":"Less is better","isMain":false,"metricName":"L75"},{"values":["39.81","39.83","39.85","39.89","39.88","39.91","39.36","39.89","39.51","39.87","39.87","39.57","39.84"],"quality":"Equal","isMain":false,"metricName":"GC (%)"}]],["Predicted genes",[]],["Similarity statistics",[{"values":[0,0,0,0,0,0,0,0,0,0,0,0,0],"quality":"Equal","isMain":false,"metricName":"# similar correct contigs"},{"values":[0,0,0,0,0,0,0,0,0,0,0,0,0],"quality":"Equal","isMain":false,"metricName":"# similar misassembled blocks"}]],["Reference statistics",[{"values":[111382,111382,111382,111382,111382,111382,111382,111382,111382,111382,111382,111382,111382],"quality":"Equal","isMain":false,"metricName":"Reference length"},{"values":[1,1,1,1,1,1,1,1,1,1,1,1,1],"quality":"Equal","isMain":false,"metricName":"Reference fragments"},{"values":["39.90","39.90","39.90","39.90","39.90","39.90","39.90","39.90","39.90","39.90","39.90","39.90","39.90"],"quality":"Equal","isMain":false,"metricName":"Reference GC (%)"}]]],"referenceName":"gi\_182682799\_ref\_NC","date":"18 July 2018, Wednesday, 16:37:31","order":[0,1,2,3,4,5,6,7,8,9,10,11,12],"assembliesNames":["ABySS\_127","ABySS\_63","CLC","IDBA\_UD","MEGAHIT","MIRA","SOAPdenovo2","SPAdes","SPAdes\_meta","SPAdes\_sc","SPAdes\_sc\_careful","Velvet","Geneious"]},{"assembliesWithNs":null,"minContig":500,"report":[["Genome statistics",[{"values":["21.548","13.842","67.603","64.251","67.817","32.660","63.981","74.176","74.830","75.294","75.294","11.291","1.831"],"quality":"More is better","isMain":true,"metricName":"Genome fraction (%)"},{"values":["1.000","1.000","1.006","1.001","1.000","1.006","1.003","1.001","1.002","1.003","1.003","0.999","1.000"],"quality":"Less is better","isMain":true,"metricName":"Duplication ratio"},{"values":[4733,1734,6490,5882,6122,5853,4155,6490,6490,6490,6490,5854,2962],"quality":"More is better","isMain":true,"metricName":"Largest alignment"},{"values":[34859,22393,109998,103989,109712,53146,103633,120062,121279,121980,121980,18238,2962],"quality":"More is better","isMain":true,"metricName":"Total aligned length"},{"values":[null,null,1112,1052,1054,null,941,1275,1237,1272,1272,null,null],"quality":"More is better","isMain":false,"metricName":"NG50"},{"values":[null,null,null,null,null,null,null,null,null,601,601,null,null],"quality":"More is better","isMain":false,"metricName":"NG75"},{"values":[852,671,1706,1609,1613,1121,1422,1740,1706,1706,1706,3971,2962],"quality":"More is better","isMain":false,"metricName":"NA50"},{"values":[625,574,1043,1052,1006,893,966,1175,1149,1139,1139,875,2962],"quality":"More is better","isMain":false,"metricName":"NA75"},{"values":[null,null,1058,989,1050,null,941,1275,1237,1237,1237,null,null],"quality":"More is better","isMain":true,"metricName":"NGA50"},{"values":[null,null,null,null,null,null,null,null,null,458,458,null,null],"quality":"More is better","isMain":false,"metricName":"NGA75"},{"values":[null,null,41,44,42,null,51,38,39,39,39,null,null],"quality":"Less is better","isMain":false,"metricName":"LG50"},{"values":[null,null,null,null,null,null,null,null,null,82,82,null,null],"quality":"Less is better","isMain":false,"metricName":"LG75"},{"values":[13,12,23,22,22,15,26,24,25,25,25,2,1],"quality":"Less is better","isMain":false,"metricName":"LA50"},{"values":[25,21,45,42,45,28,48,46,47,48,48,6,1],"quality":"Less is better","isMain":false,"metricName":"LA75"},{"values":[null,null,43,45,43,null,51,38,39,39,39,null,null],"quality":"Less is better","isMain":true,"metricName":"LGA50"},{"values":[null,null,null,null,null,null,null,null,null,84,84,null,null],"quality":"Less is better","isMain":false,"metricName":"LGA75"}]],["Misassemblies",[{"values":[0,0,0,1,0,0,0,0,0,0,0,0,0],"quality":"Less is better","isMain":true,"metricName":"# misassemblies"},{"values":[0,0,0,1,0,0,0,0,0,0,0,0,0],"quality":"Less is better","isMain":false,"metricName":" # relocations"},{"values":[0,0,0,0,0,0,0,0,0,0,0,0,0],"quality":"Less is better","isMain":false,"metricName":" # translocations"},{"values":[0,0,0,0,0,0,0,0,0,0,0,0,0],"quality":"Less is better","isMain":false,"metricName":" # inversions"},{"values":[0,0,0,1,0,0,0,0,0,0,0,0,0],"quality":"Less is better","isMain":false,"metricName":"# misassembled contigs"},{"values":[0,0,0,1121,0,0,0,0,0,0,0,0,0],"quality":"Less is better","isMain":true,"metricName":"Misassembled contigs length"},{"values":[0,0,0,0,0,0,0,0,0,0,0,0,0],"quality":"Less is better","isMain":false,"metricName":"# local misassemblies"},{"values":[0,0,0,0,0,0,0,0,0,0,0,0,0],"quality":"Less is better","isMain":false,"metricName":"# unaligned mis. contigs"}]],["Unaligned",[{"values":[0,0,0,0,0,0,0,0,0,0,0,0,0],"quality":"Less is better","isMain":false,"metricName":"# fully unaligned contigs"},{"values":[0,0,0,0,0,0,0,0,0,0,0,0,0],"quality":"Less is better","isMain":false,"metricName":"Fully unaligned length"},{"values":[0,0,2,1,1,0,0,1,0,1,1,0,0],"quality":"Less is better","isMain":false,"metricName":"# partially unaligned contigs"},{"values":[0,0,1732,810,984,0,0,788,0,984,984,0,0],"quality":"Less is better","isMain":false,"metricName":"Partially unaligned length"}]],["Mismatches",[{"values":[5,4,102,76,76,7,74,138,111,106,90,3,0],"quality":"Less is better","isMain":false,"metricName":"# mismatches"},{"values":[0,0,0,0,0,0,3,0,0,0,0,1,0],"quality":"Less is better","isMain":false,"metricName":"# indels"},{"values":[0,0,0,0,0,0,137,0,0,0,0,28,0],"quality":"Less is better","isMain":false,"metricName":"Indels length"},{"values":["14.34","17.86","93.27","73.12","69.27","13.25","71.49","115.00","91.69","87.02","73.89","16.42","0.00"],"quality":"Less is better","isMain":true,"metricName":"# mismatches per 100 kbp"},{"values":["0.00","0.00","0.00","0.00","0.00","0.00","2.90","0.00","0.00","0.00","0.00","5.47","0.00"],"quality":"Less is better","isMain":true,"metricName":"# indels per 100 kbp"},{"values":[0,0,0,0,0,0,0,0,0,0,0,0,0],"quality":"Less is better","isMain":false,"metricName":" # indels (<= 5 bp)"},{"values":[0,0,0,0,0,0,3,0,0,0,0,1,0],"quality":"Less is better","isMain":false,"metricName":" # indels (> 5 bp)"},{"values":[0,0,2,0,0,59,254,0,0,0,0,0,4],"quality":"Less is better","isMain":false,"metricName":"# N's"},{"values":["0.00","0.00","1.79","0.00","0.00","111.01","244.63","0.00","0.00","0.00","0.00","0.00","135.04"],"quality":"Less is better","isMain":true,"metricName":"# N's per 100 kbp"}]],["Statistics without reference",[{"values":[40,31,81,75,81,46,83,81,84,85,85,13,1],"quality":"Equal","isMain":true,"metricName":"# contigs"},{"values":[7,3,45,46,46,18,46,53,53,53,53,3,1],"quality":"Equal","isMain":false,"metricName":"# contigs (>= 1000 bp)"},{"values":[0,0,1,1,2,1,0,2,2,2,2,1,0],"quality":"Equal","isMain":false,"metricName":"# contigs (>= 5000 bp)"},{"values":[0,0,0,0,0,0,0,0,0,0,0,0,0],"quality":"Equal","isMain":false,"metricName":"# contigs (>= 10000 bp)"},{"values":[0,0,0,0,0,0,0,0,0,0,0,0,0],"quality":"Equal","isMain":false,"metricName":"# contigs (>= 25000 bp)"},{"values":[0,0,0,0,0,0,0,0,0,0,0,0,0],"quality":"Equal","isMain":false,"metricName":"# contigs (>= 50000 bp)"},{"values":[4733,1734,6490,5901,6122,5853,4155,6490,6490,6490,6490,5869,2962],"quality":"More is better","isMain":true,"metricName":"Largest contig"},{"values":[34859,22393,111747,104818,110698,53146,103830,120850,121279,123107,123107,18253,2962],"quality":"More is better","isMain":true,"metricName":"Total length"},{"values":[12360,4193,85623,83012,85343,30733,76592,99369,97681,98370,98370,11040,2962],"quality":"More is better","isMain":true,"metricName":"Total length (>= 1000 bp)"},{"values":[0,0,6490,5901,12048,5853,0,12163,12163,12163,12163,5869,0],"quality":"More is better","isMain":false,"metricName":"Total length (>= 5000 bp)"},{"values":[0,0,0,0,0,0,0,0,0,0,0,0,0],"quality":"More is better","isMain":true,"metricName":"Total length (>= 10000 bp)"},{"values":[0,0,0,0,0,0,0,0,0,0,0,0,0],"quality":"More is better","isMain":false,"metricName":"Total length (>= 25000 bp)"},{"values":[0,0,0,0,0,0,0,0,0,0,0,0,0],"quality":"More is better","isMain":true,"metricName":"Total length (>= 50000 bp)"},{"values":[852,671,1740,1609,1613,1121,1424,1740,1706,1706,1706,3971,2962],"quality":"More is better","isMain":false,"metricName":"N50"},{"values":[625,574,1044,1105,1050,893,966,1182,1149,1149,1149,875,2962],"quality":"More is better","isMain":false,"metricName":"N75"},{"values":[13,12,23,22,22,15,25,24,25,25,25,2,1],"quality":"Less is better","isMain":false,"metricName":"L50"},{"values":[25,21,44,42,44,28,48,46,47,48,48,6,1],"quality":"Less is better","isMain":false,"metricName":"L75"},{"values":["44.34","45.66","43.24","43.17","43.05","43.77","43.44","43.12","43.18","43.02","43.02","47.12","42.09"],"quality":"Equal","isMain":false,"metricName":"GC (%)"}]],["Predicted genes",[]],["Similarity statistics",[{"values":[0,0,0,0,0,0,0,0,0,0,0,0,0],"quality":"Equal","isMain":false,"metricName":"# similar correct contigs"},{"values":[0,0,0,0,0,0,0,0,0,0,0,0,0],"quality":"Equal","isMain":false,"metricName":"# similar misassembled blocks"}]],["Reference statistics",[{"values":[161773,161773,161773,161773,161773,161773,161773,161773,161773,161773,161773,161773,161773],"quality":"Equal","isMain":false,"metricName":"Reference length"},{"values":[1,1,1,1,1,1,1,1,1,1,1,1,1],"quality":"Equal","isMain":false,"metricName":"Reference fragments"},{"values":["43.56","43.56","43.56","43.56","43.56","43.56","43.56","43.56","43.56","43.56","43.56","43.56","43.56"],"quality":"Equal","isMain":false,"metricName":"Reference GC (%)"}]]],"referenceName":"gi\_18450236\_ref\_NC","date":"18 July 2018, Wednesday, 16:37:46","order":[0,1,2,3,4,5,6,7,8,9,10,11,12],"assembliesNames":["ABySS\_127","ABySS\_63","CLC","IDBA\_UD","MEGAHIT","MIRA","SOAPdenovo2","SPAdes","SPAdes\_meta","SPAdes\_sc","SPAdes\_sc\_careful","Velvet","Geneious"]},{"assembliesWithNs":null,"minContig":500,"report":[["Genome statistics",[{"values":["99.826","99.985","98.924","100.000","100.000","100.000","3.684","75.167","100.000","100.000","100.000","100.000","97.197","100.000"],"quality":"More is better","isMain":true,"metricName":"Genome fraction (%)"},{"values":["1.001","1.003","1.001","1.002","1.003","1.014","1.000","1.063","1.002","1.011","1.007","1.002","1.004","1.004"],"quality":"Less is better","isMain":true,"metricName":"Duplication ratio"},{"values":[23543,30882,37508,52896,52938,53022,718,9028,52924,52960,46341,46199,9064,53022],"quality":"More is better","isMain":true,"metricName":"Largest alignment"},{"values":[52753,52938,52291,52896,52938,53546,1945,40633,52924,52960,53048,52906,51515,53022],"quality":"More is better","isMain":true,"metricName":"Total aligned length"},{"values":[17843,30882,37512,52896,52938,53022,null,5816,52924,53357,46473,46199,5624,53022],"quality":"More is better","isMain":false,"metricName":"NG50"},{"values":[17843,14302,12838,52896,52938,53022,null,2739,52924,53357,46473,46199,3583,53022],"quality":"More is better","isMain":false,"metricName":"NG75"},{"values":[17843,30882,37508,52896,52938,53022,688,5127,52924,52960,46341,46199,5624,53022],"quality":"More is better","isMain":false,"metricName":"NA50"},{"values":[17843,14302,12838,52896,52938,53022,539,2387,52924,52960,46341,46199,4893,53022],"quality":"More is better","isMain":false,"metricName":"NA75"},{"values":[17843,30882,37508,52896,52938,53022,null,5127,52924,52960,46341,46199,5624,53022],"quality":"More is better","isMain":true,"metricName":"NGA50"},{"values":[17843,14302,12838,52896,52938,53022,null,631,52924,52960,46341,46199,3583,53022],"quality":"More is better","isMain":false,"metricName":"NGA75"},{"values":[2,1,1,1,1,1,null,4,1,1,1,1,4,1],"quality":"Less is better","isMain":false,"metricName":"LG50"},{"values":[2,2,2,1,1,1,null,7,1,1,1,1,7,1],"quality":"Less is better","isMain":false,"metricName":"LG75"},{"values":[2,1,1,1,1,1,2,4,1,1,1,1,4,1],"quality":"Less is better","isMain":false,"metricName":"LA50"},{"values":[2,2,2,1,1,1,3,7,1,1,1,1,6,1],"quality":"Less is better","isMain":false,"metricName":"LA75"},{"values":[2,1,1,1,1,1,null,4,1,1,1,1,4,1],"quality":"Less is better","isMain":true,"metricName":"LGA50"},{"values":[2,2,2,1,1,1,null,9,1,1,1,1,7,1],"quality":"Less is better","isMain":false,"metricName":"LGA75"}]],["Misassemblies",[{"values":[0,0,0,0,0,0,0,0,0,0,0,0,0,0],"quality":"Less is better","isMain":true,"metricName":"# misassemblies"},{"values":[0,0,0,0,0,0,0,0,0,0,0,0,0,0],"quality":"Less is better","isMain":false,"metricName":" # relocations"},{"values":[0,0,0,0,0,0,0,0,0,0,0,0,0,0],"quality":"Less is better","isMain":false,"metricName":" # translocations"},{"values":[0,0,0,0,0,0,0,0,0,0,0,0,0,0],"quality":"Less is better","isMain":false,"metricName":" # inversions"},{"values":[0,0,0,0,0,0,0,0,0,0,0,0,0,0],"quality":"Less is better","isMain":false,"metricName":"# misassembled contigs"},{"values":[0,0,0,0,0,0,0,0,0,0,0,0,0,0],"quality":"Less is better","isMain":true,"metricName":"Misassembled contigs length"},{"values":[0,0,0,1,1,1,0,0,1,1,0,0,0,1],"quality":"Less is better","isMain":false,"metricName":"# local misassemblies"},{"values":[0,0,0,0,0,0,0,0,0,0,0,0,0,0],"quality":"Less is better","isMain":false,"metricName":"# unaligned mis. contigs"}]],["Unaligned",[{"values":[0,0,0,0,0,0,0,0,0,0,0,0,0,0],"quality":"Less is better","isMain":false,"metricName":"# fully unaligned contigs"},{"values":[0,0,0,0,0,0,0,0,0,0,0,0,0,0],"quality":"Less is better","isMain":false,"metricName":"Fully unaligned length"},{"values":[0,0,0,0,0,0,0,3,0,0,0,0,0,0],"quality":"Less is better","isMain":false,"metricName":"# partially unaligned contigs"},{"values":[0,0,0,0,0,0,0,6117,0,0,0,0,0,0],"quality":"Less is better","isMain":false,"metricName":"Partially unaligned length"}]],["Mismatches",[{"values":[5,1,9,0,0,0,0,0,2,1,0,0,0,0],"quality":"Less is better","isMain":false,"metricName":"# mismatches"},{"values":[0,0,0,0,0,0,0,31,0,0,0,0,0,0],"quality":"Less is better","isMain":false,"metricName":"# indels"},{"values":[0,0,0,0,0,0,0,1490,0,0,0,0,0,0],"quality":"Less is better","isMain":false,"metricName":"Indels length"},{"values":["9.49","1.89","17.23","0.00","0.00","0.00","0.00","0.00","3.79","1.89","0.00","0.00","0.00","0.00"],"quality":"Less is better","isMain":true,"metricName":"# mismatches per 100 kbp"},{"values":["0.00","0.00","0.00","0.00","0.00","0.00","0.00","78.11","0.00","0.00","0.00","0.00","0.00","0.00"],"quality":"Less is better","isMain":true,"metricName":"# indels per 100 kbp"},{"values":[0,0,0,0,0,0,0,0,0,0,0,0,0,0],"quality":"Less is better","isMain":false,"metricName":" # indels (<= 5 bp)"},{"values":[0,0,0,0,0,0,0,31,0,0,0,0,0,0],"quality":"Less is better","isMain":false,"metricName":" # indels (> 5 bp)"},{"values":[0,0,0,0,0,1,0,3371,0,0,0,0,0,1],"quality":"Less is better","isMain":false,"metricName":"# N's"},{"values":["0.00","0.00","0.00","0.00","0.00","1.87","0.00","6978.72","0.00","0.00","0.00","0.00","0.00","1.89"],"quality":"Less is better","isMain":true,"metricName":"# N's per 100 kbp"}]],["Statistics without reference",[{"values":[4,4,3,1,1,2,3,12,1,1,2,2,14,1],"quality":"Equal","isMain":true,"metricName":"# contigs"},{"values":[4,4,3,1,1,1,0,11,1,1,2,2,11,1],"quality":"Equal","isMain":false,"metricName":"# contigs (>= 1000 bp)"},{"values":[3,3,2,1,1,1,0,4,1,1,2,2,5,1],"quality":"Equal","isMain":false,"metricName":"# contigs (>= 5000 bp)"},{"values":[2,2,2,1,1,1,0,0,1,1,1,1,0,1],"quality":"Equal","isMain":false,"metricName":"# contigs (>= 10000 bp)"},{"values":[0,1,1,1,1,1,0,0,1,1,1,1,0,1],"quality":"Equal","isMain":false,"metricName":"# contigs (>= 25000 bp)"},{"values":[0,0,0,1,1,1,0,0,1,1,0,0,0,1],"quality":"Equal","isMain":false,"metricName":"# contigs (>= 50000 bp)"},{"values":[23543,30882,37512,52896,52938,53022,718,9115,52924,53357,46473,46199,9064,53022],"quality":"More is better","isMain":true,"metricName":"Largest contig"},{"values":[52753,52938,52295,52896,52938,53546,1945,48304,52924,53357,53180,52906,51515,53022],"quality":"More is better","isMain":true,"metricName":"Total length"},{"values":[52753,52938,52295,52896,52938,53022,0,47777,52924,53357,53180,52906,49550,53022],"quality":"More is better","isMain":true,"metricName":"Total length (>= 1000 bp)"},{"values":[51345,51763,50350,52896,52938,53022,0,30300,52924,53357,53180,52906,33948,53022],"quality":"More is better","isMain":false,"metricName":"Total length (>= 5000 bp)"},{"values":[41386,45184,50350,52896,52938,53022,0,0,52924,53357,46473,46199,0,53022],"quality":"More is better","isMain":true,"metricName":"Total length (>= 10000 bp)"},{"values":[0,30882,37512,52896,52938,53022,0,0,52924,53357,46473,46199,0,53022],"quality":"More is better","isMain":false,"metricName":"Total length (>= 25000 bp)"},{"values":[0,0,0,52896,52938,53022,0,0,52924,53357,0,0,0,53022],"quality":"More is better","isMain":true,"metricName":"Total length (>= 50000 bp)"},{"values":[17843,30882,37512,52896,52938,53022,688,6782,52924,53357,46473,46199,5624,53022],"quality":"More is better","isMain":false,"metricName":"N50"},{"values":[17843,14302,12838,52896,52938,53022,539,3111,52924,53357,46473,46199,4893,53022],"quality":"More is better","isMain":false,"metricName":"N75"},{"values":[2,1,1,1,1,1,2,3,1,1,1,1,4,1],"quality":"Less is better","isMain":false,"metricName":"L50"},{"values":[2,2,2,1,1,1,3,6,1,1,1,1,6,1],"quality":"Less is better","isMain":false,"metricName":"L75"},{"values":["68.12","68.11","68.12","68.11","68.12","68.13","67.30","68.13","68.12","68.08","68.11","68.12","68.15","68.10"],"quality":"Equal","isMain":false,"metricName":"GC (%)"}]],["Predicted genes",[]],["Similarity statistics",[{"values":[0,0,0,0,0,0,0,0,0,0,0,0,0,0],"quality":"Equal","isMain":false,"metricName":"# similar correct contigs"},{"values":[0,0,0,0,0,0,0,0,0,0,0,0,0,0],"quality":"Equal","isMain":false,"metricName":"# similar misassembled blocks"}]],["Reference statistics",[{"values":[52797,52797,52797,52797,52797,52797,52797,52797,52797,52797,52797,52797,52797,52797],"quality":"Equal","isMain":false,"metricName":"Reference length"},{"values":[1,1,1,1,1,1,1,1,1,1,1,1,1,1],"quality":"Equal","isMain":false,"metricName":"Reference fragments"},{"values":["68.11","68.11","68.11","68.11","68.11","68.11","68.11","68.11","68.11","68.11","68.11","68.11","68.11","68.11"],"quality":"Equal","isMain":false,"metricName":"Reference GC (%)"}]]],"referenceName":"gi\_18496886\_ref\_NC","date":"18 July 2018, Wednesday, 16:38:01","order":[0,1,2,3,4,5,6,7,8,9,10,11,12,13],"assembliesNames":["ABySS\_127","ABySS\_63","CLC","IDBA\_UD","MEGAHIT","MIRA","Ray\_Meta","SOAPdenovo2","SPAdes","SPAdes\_meta","SPAdes\_sc","SPAdes\_sc\_careful","Velvet","Geneious"]},{"assembliesWithNs":null,"minContig":500,"report":[["Genome statistics",[{"values":["4.888","1.935","49.519","39.857","54.049","15.731","45.232","61.150","61.190","61.697","61.458","0.418"],"quality":"More is better","isMain":true,"metricName":"Genome fraction (%)"},{"values":["1.000","1.000","1.005","1.013","1.004","1.000","1.011","1.005","1.001","1.013","1.013","1.000"],"quality":"Less is better","isMain":true,"metricName":"Duplication ratio"},{"values":[1097,679,3050,2885,3061,1753,2789,3123,2664,3123,3123,612],"quality":"More is better","isMain":true,"metricName":"Largest alignment"},{"values":[7159,2834,72877,58377,79157,23038,66395,89638,89697,90455,90106,612],"quality":"More is better","isMain":true,"metricName":"Total aligned length"},{"values":[null,null,526,null,590,null,null,826,711,799,795,null],"quality":"More is better","isMain":false,"metricName":"NG50"},{"values":[732,555,926,971,931,904,866,1063,1070,1070,1070,612],"quality":"More is better","isMain":false,"metricName":"NA50"},{"values":[603,515,691,733,682,700,705,826,799,826,826,612],"quality":"More is better","isMain":false,"metricName":"NA75"},{"values":[null,null,null,null,574,null,null,752,711,779,779,null],"quality":"More is better","isMain":true,"metricName":"NGA50"},{"values":[null,null,77,null,73,null,null,54,62,57,57,null],"quality":"Less is better","isMain":false,"metricName":"LG50"},{"values":[5,3,28,23,29,10,30,30,30,29,29,1],"quality":"Less is better","isMain":false,"metricName":"LA50"},{"values":[7,4,52,40,55,17,51,55,54,54,53,1],"quality":"Less is better","isMain":false,"metricName":"LA75"},{"values":[null,null,null,null,76,null,null,59,62,59,59,null],"quality":"Less is better","isMain":true,"metricName":"LGA50"}]],["Misassemblies",[{"values":[0,0,0,0,0,0,0,0,0,0,0,0],"quality":"Less is better","isMain":true,"metricName":"# misassemblies"},{"values":[0,0,0,0,0,0,0,0,0,0,0,0],"quality":"Less is better","isMain":false,"metricName":" # relocations"},{"values":[0,0,0,0,0,0,0,0,0,0,0,0],"quality":"Less is better","isMain":false,"metricName":" # translocations"},{"values":[0,0,0,0,0,0,0,0,0,0,0,0],"quality":"Less is better","isMain":false,"metricName":" # inversions"},{"values":[0,0,0,0,0,0,0,0,0,0,0,0],"quality":"Less is better","isMain":false,"metricName":"# misassembled contigs"},{"values":[0,0,0,0,0,0,0,0,0,0,0,0],"quality":"Less is better","isMain":true,"metricName":"Misassembled contigs length"},{"values":[0,0,0,0,0,0,0,0,0,0,0,0],"quality":"Less is better","isMain":false,"metricName":"# local misassemblies"},{"values":[0,0,0,0,0,0,0,0,0,0,0,0],"quality":"Less is better","isMain":false,"metricName":"# unaligned mis. contigs"}]],["Unaligned",[{"values":[0,0,0,0,0,0,0,0,0,0,0,0],"quality":"Less is better","isMain":false,"metricName":"# fully unaligned contigs"},{"values":[0,0,0,0,0,0,0,0,0,0,0,0],"quality":"Less is better","isMain":false,"metricName":"Fully unaligned length"},{"values":[0,0,1,0,1,0,0,1,0,1,0,0],"quality":"Less is better","isMain":false,"metricName":"# partially unaligned contigs"},{"values":[0,0,2138,0,1147,0,0,3513,0,658,0,0],"quality":"Less is better","isMain":false,"metricName":"Partially unaligned length"}]],["Mismatches",[{"values":[1,0,137,123,145,2,60,209,117,131,115,0],"quality":"Less is better","isMain":false,"metricName":"# mismatches"},{"values":[0,0,5,2,3,0,4,1,0,1,1,0],"quality":"Less is better","isMain":false,"metricName":"# indels"},{"values":[0,0,6,2,20,0,161,78,0,78,78,0],"quality":"Less is better","isMain":false,"metricName":"Indels length"},{"values":["13.97","0.00","188.91","210.72","183.18","8.68","90.57","233.37","130.56","144.98","127.77","0.00"],"quality":"Less is better","isMain":true,"metricName":"# mismatches per 100 kbp"},{"values":["0.00","0.00","6.89","3.43","3.79","0.00","6.04","1.12","0.00","1.11","1.11","0.00"],"quality":"Less is better","isMain":true,"metricName":"# indels per 100 kbp"},{"values":[0,0,5,2,2,0,0,0,0,0,0,0],"quality":"Less is better","isMain":false,"metricName":" # indels (<= 5 bp)"},{"values":[0,0,0,0,1,0,4,1,0,1,1,0],"quality":"Less is better","isMain":false,"metricName":" # indels (> 5 bp)"},{"values":[0,0,0,0,0,40,553,0,0,0,0,0],"quality":"Less is better","isMain":false,"metricName":"# N's"},{"values":["0.00","0.00","0.00","0.00","0.00","173.63","826.01","0.00","0.00","0.00","0.00","0.00"],"quality":"Less is better","isMain":true,"metricName":"# N's per 100 kbp"}]],["Statistics without reference",[{"values":[10,5,80,63,86,26,78,84,88,86,85,1],"quality":"Equal","isMain":true,"metricName":"# contigs"},{"values":[1,0,22,22,24,8,16,36,35,38,37,0],"quality":"Equal","isMain":false,"metricName":"# contigs (>= 1000 bp)"},{"values":[0,0,0,0,0,0,0,0,0,0,0,0],"quality":"Equal","isMain":false,"metricName":"# contigs (>= 5000 bp)"},{"values":[0,0,0,0,0,0,0,0,0,0,0,0],"quality":"Equal","isMain":false,"metricName":"# contigs (>= 10000 bp)"},{"values":[0,0,0,0,0,0,0,0,0,0,0,0],"quality":"Equal","isMain":false,"metricName":"# contigs (>= 25000 bp)"},{"values":[0,0,0,0,0,0,0,0,0,0,0,0],"quality":"Equal","isMain":false,"metricName":"# contigs (>= 50000 bp)"},{"values":[1097,679,3050,2885,3061,1753,2789,4402,2664,3123,3123,612],"quality":"More is better","isMain":true,"metricName":"Largest contig"},{"values":[7159,2834,75020,59159,80652,23038,66948,93528,89702,92211,91208,612],"quality":"More is better","isMain":true,"metricName":"Total length"},{"values":[1097,0,33908,29732,36888,10166,21386,57143,50697,56390,55387,0],"quality":"More is better","isMain":true,"metricName":"Total length (>= 1000 bp)"},{"values":[0,0,0,0,0,0,0,0,0,0,0,0],"quality":"More is better","isMain":false,"metricName":"Total length (>= 5000 bp)"},{"values":[0,0,0,0,0,0,0,0,0,0,0,0],"quality":"More is better","isMain":true,"metricName":"Total length (>= 10000 bp)"},{"values":[0,0,0,0,0,0,0,0,0,0,0,0],"quality":"More is better","isMain":false,"metricName":"Total length (>= 25000 bp)"},{"values":[0,0,0,0,0,0,0,0,0,0,0,0],"quality":"More is better","isMain":true,"metricName":"Total length (>= 50000 bp)"},{"values":[732,555,934,1015,956,904,877,1107,1070,1098,1107,612],"quality":"More is better","isMain":false,"metricName":"N50"},{"values":[603,515,710,733,697,700,710,894,799,845,841,612],"quality":"More is better","isMain":false,"metricName":"N75"},{"values":[5,3,26,22,28,10,29,27,30,29,28,1],"quality":"Less is better","isMain":false,"metricName":"L50"},{"values":[7,4,49,40,53,17,51,50,54,52,52,1],"quality":"Less is better","isMain":false,"metricName":"L75"},{"values":["28.19","27.88","27.14","27.46","27.44","27.85","27.47","27.05","27.33","27.35","27.35","26.96"],"quality":"Equal","isMain":false,"metricName":"GC (%)"}]],["Predicted genes",[]],["Similarity statistics",[{"values":[0,0,0,0,0,0,0,0,0,0,0,0],"quality":"Equal","isMain":false,"metricName":"# similar correct contigs"},{"values":[0,0,0,0,0,0,0,0,0,0,0,0],"quality":"Equal","isMain":false,"metricName":"# similar misassembled blocks"}]],["Reference statistics",[{"values":[146454,146454,146454,146454,146454,146454,146454,146454,146454,146454,146454,146454],"quality":"Equal","isMain":false,"metricName":"Reference length"},{"values":[1,1,1,1,1,1,1,1,1,1,1,1],"quality":"Equal","isMain":false,"metricName":"Reference fragments"},{"values":["27.40","27.40","27.40","27.40","27.40","27.40","27.40","27.40","27.40","27.40","27.40","27.40"],"quality":"Equal","isMain":false,"metricName":"Reference GC (%)"}]]],"referenceName":"gi\_18640086\_ref\_NC","date":"18 July 2018, Wednesday, 16:38:15","order":[0,1,2,3,4,5,6,7,8,9,10,11],"assembliesNames":["ABySS\_127","ABySS\_63","CLC","IDBA\_UD","MEGAHIT","MIRA","SOAPdenovo2","SPAdes","SPAdes\_meta","SPAdes\_sc","SPAdes\_sc\_careful","Velvet"]},{"assembliesWithNs":null,"minContig":500,"report":[["Genome statistics",[{"values":["0.634","2.069","1.593","2.017","3.677","1.251","10.011","4.344","5.141","6.203","2.001"],"quality":"More is better","isMain":true,"metricName":"Genome fraction (%)"},{"values":["1.000","1.001","1.000","1.006","1.012","1.080","1.009","1.009","1.005","1.036","1.015"],"quality":"Less is better","isMain":true,"metricName":"Duplication ratio"},{"values":[697,1009,3277,1352,4176,999,18703,4382,5179,7264,4176],"quality":"More is better","isMain":true,"metricName":"Largest alignment"},{"values":[1304,4260,3277,4170,7649,2573,20662,9010,10627,12808,4176],"quality":"More is better","isMain":true,"metricName":"Total aligned length"},{"values":[697,594,3277,1124,4176,591,18703,3442,3995,7264,4176],"quality":"More is better","isMain":false,"metricName":"NA50"},{"values":[607,513,3277,670,3473,530,18703,3442,670,4091,4176],"quality":"More is better","isMain":false,"metricName":"NA75"},{"values":[null,null,null,null,null,null,null,null,null,null,null],"quality":"More is better","isMain":true,"metricName":"NGA50"},{"values":[1,4,1,2,1,2,1,2,2,1,1],"quality":"Less is better","isMain":false,"metricName":"LA50"},{"values":[2,6,1,3,2,3,1,2,4,2,1],"quality":"Less is better","isMain":false,"metricName":"LA75"}]],["Misassemblies",[{"values":[0,0,0,0,0,0,0,0,0,0,0],"quality":"Less is better","isMain":true,"metricName":"# misassemblies"},{"values":[0,0,0,0,0,0,0,0,0,0,0],"quality":"Less is better","isMain":false,"metricName":" # relocations"},{"values":[0,0,0,0,0,0,0,0,0,0,0],"quality":"Less is better","isMain":false,"metricName":" # translocations"},{"values":[0,0,0,0,0,0,0,0,0,0,0],"quality":"Less is better","isMain":false,"metricName":" # inversions"},{"values":[0,0,0,0,0,0,0,0,0,0,0],"quality":"Less is better","isMain":false,"metricName":"# misassembled contigs"},{"values":[0,0,0,0,0,0,0,0,0,0,0],"quality":"Less is better","isMain":true,"metricName":"Misassembled contigs length"},{"values":[0,0,0,0,0,0,1,0,1,1,0],"quality":"Less is better","isMain":false,"metricName":"# local misassemblies"},{"values":[0,0,0,0,0,0,0,0,0,0,0],"quality":"Less is better","isMain":false,"metricName":"# unaligned mis. contigs"}]],["Unaligned",[{"values":[0,0,0,0,0,0,0,0,0,0,0],"quality":"Less is better","isMain":false,"metricName":"# fully unaligned contigs"},{"values":[0,0,0,0,0,0,0,0,0,0,0],"quality":"Less is better","isMain":false,"metricName":"Fully unaligned length"},{"values":[0,1,0,0,0,0,0,0,2,1,0],"quality":"Less is better","isMain":false,"metricName":"# partially unaligned contigs"},{"values":[0,839,0,0,0,0,0,0,3442,839,0],"quality":"Less is better","isMain":false,"metricName":"Partially unaligned length"}]],["Mismatches",[{"values":[5,45,16,60,56,26,337,134,158,197,56],"quality":"Less is better","isMain":false,"metricName":"# mismatches"},{"values":[0,5,2,14,14,2,61,24,41,53,13],"quality":"Less is better","isMain":false,"metricName":"# indels"},{"values":[0,6,2,31,112,2,312,101,166,241,70],"quality":"Less is better","isMain":false,"metricName":"Indels length"},{"values":["383.44","1057.33","488.25","1446.13","740.25","1010.49","1636.32","1499.38","1494.09","1543.89","1360.54"],"quality":"Less is better","isMain":true,"metricName":"# mismatches per 100 kbp"},{"values":["0.00","117.48","61.03","337.43","185.06","77.73","296.19","268.55","387.71","415.36","315.84"],"quality":"Less is better","isMain":true,"metricName":"# indels per 100 kbp"},{"values":[0,5,2,13,11,2,47,20,36,45,11],"quality":"Less is better","isMain":false,"metricName":" # indels (<= 5 bp)"},{"values":[0,0,0,1,3,0,14,4,5,8,2],"quality":"Less is better","isMain":false,"metricName":" # indels (> 5 bp)"},{"values":[0,0,0,0,20,70,0,0,1343,1246,11],"quality":"Less is better","isMain":false,"metricName":"# N's"},{"values":["0.00","0.00","0.00","0.00","261.23","2518.89","0.00","0.00","9544.45","8864.54","263.41"],"quality":"Less is better","isMain":true,"metricName":"# N's per 100 kbp"}]],["Statistics without reference",[{"values":[2,7,1,5,2,4,4,4,4,4,1],"quality":"Equal","isMain":true,"metricName":"# contigs"},{"values":[0,2,1,2,2,0,1,2,2,2,1],"quality":"Equal","isMain":false,"metricName":"# contigs (>= 1000 bp)"},{"values":[0,0,0,0,0,0,1,0,1,1,0],"quality":"Equal","isMain":false,"metricName":"# contigs (>= 5000 bp)"},{"values":[0,0,0,0,0,0,1,0,0,0,0],"quality":"Equal","isMain":false,"metricName":"# contigs (>= 10000 bp)"},{"values":[0,0,0,0,0,0,0,0,0,0,0],"quality":"Equal","isMain":false,"metricName":"# contigs (>= 25000 bp)"},{"values":[0,0,0,0,0,0,0,0,0,0,0],"quality":"Equal","isMain":false,"metricName":"# contigs (>= 50000 bp)"},{"values":[697,1196,3277,1352,4176,999,18822,4382,7686,7671,4176],"quality":"More is better","isMain":true,"metricName":"Largest contig"},{"values":[1304,5101,3277,4172,7656,2779,20783,9013,14071,14056,4176],"quality":"More is better","isMain":true,"metricName":"Total length"},{"values":[0,2205,3277,2476,7656,0,18822,7825,12616,12601,4176],"quality":"More is better","isMain":true,"metricName":"Total length (>= 1000 bp)"},{"values":[0,0,0,0,0,0,18822,0,7686,7671,0],"quality":"More is better","isMain":false,"metricName":"Total length (>= 5000 bp)"},{"values":[0,0,0,0,0,0,18822,0,0,0,0],"quality":"More is better","isMain":true,"metricName":"Total length (>= 10000 bp)"},{"values":[0,0,0,0,0,0,0,0,0,0,0],"quality":"More is better","isMain":false,"metricName":"Total length (>= 25000 bp)"},{"values":[0,0,0,0,0,0,0,0,0,0,0],"quality":"More is better","isMain":true,"metricName":"Total length (>= 50000 bp)"},{"values":[697,672,3277,1124,4176,659,18822,3443,7686,7671,4176],"quality":"More is better","isMain":false,"metricName":"N50"},{"values":[607,594,3277,672,3480,591,18822,3443,4930,4930,4176],"quality":"More is better","isMain":false,"metricName":"N75"},{"values":[1,3,1,2,1,2,1,2,1,1,1],"quality":"Less is better","isMain":false,"metricName":"L50"},{"values":[2,5,1,3,2,3,1,2,2,2,1],"quality":"Less is better","isMain":false,"metricName":"L75"},{"values":["33.59","31.43","34.42","31.90","32.99","31.64","32.78","31.58","31.73","31.73","32.53"],"quality":"Equal","isMain":false,"metricName":"GC (%)"}]],["Predicted genes",[]],["Similarity statistics",[{"values":[0,0,0,0,0,0,0,0,0,0,0],"quality":"Equal","isMain":false,"metricName":"# similar correct contigs"},{"values":[0,0,0,0,0,0,0,0,0,0,0],"quality":"Equal","isMain":false,"metricName":"# similar misassembled blocks"}]],["Reference statistics",[{"values":[205719,205719,205719,205719,205719,205719,205719,205719,205719,205719,205719],"quality":"Equal","isMain":false,"metricName":"Reference length"},{"values":[1,1,1,1,1,1,1,1,1,1,1],"quality":"Equal","isMain":false,"metricName":"Reference fragments"},{"values":["33.17","33.17","33.17","33.17","33.17","33.17","33.17","33.17","33.17","33.17","33.17"],"quality":"Equal","isMain":false,"metricName":"Reference GC (%)"}]]],"referenceName":"gi\_18640237\_ref\_NC","date":"18 July 2018, Wednesday, 16:38:29","order":[0,1,2,3,4,5,6,7,8,9,10],"assembliesNames":["ABySS\_127","CLC","IDBA\_UD","MEGAHIT","MIRA","SOAPdenovo2","SPAdes","SPAdes\_meta","SPAdes\_sc","SPAdes\_sc\_careful","Geneious"]},{"assembliesWithNs":null,"minContig":500,"report":[["Genome statistics",[{"values":["98.895","97.742","98.358","98.554","99.352","100.000","7.909","94.895","98.705","98.404","98.358","98.358","94.753"],"quality":"More is better","isMain":true,"metricName":"Genome fraction (%)"},{"values":["1.007","1.003","1.000","1.002","1.004","1.001","1.000","1.027","1.000","1.000","1.001","1.001","1.000"],"quality":"Less is better","isMain":true,"metricName":"Duplication ratio"},{"values":[15621,9381,37785,17430,19844,47899,863,16929,37995,37851,15624,15624,15574],"quality":"More is better","isMain":true,"metricName":"Largest alignment"},{"values":[47651,46917,47086,47254,47750,47899,3786,45687,47252,47108,47140,47140,45380],"quality":"More is better","isMain":true,"metricName":"Total aligned length"},{"values":[9509,8632,37785,15712,17738,47899,null,9499,37995,37851,11833,11833,11783],"quality":"More is better","isMain":false,"metricName":"NG50"},{"values":[8658,5743,37785,9455,17738,47899,null,4689,37995,37851,9257,9257,4712],"quality":"More is better","isMain":false,"metricName":"NG75"},{"values":[9509,8632,37785,15712,17738,47899,620,9386,37995,37851,11833,11833,11783],"quality":"More is better","isMain":false,"metricName":"NA50"},{"values":[8658,5743,37785,9455,17738,47899,570,4585,37995,37851,9257,9257,4712],"quality":"More is better","isMain":false,"metricName":"NA75"},{"values":[9509,8632,37785,15712,17738,47899,null,9386,37995,37851,11833,11833,11783],"quality":"More is better","isMain":true,"metricName":"NGA50"},{"values":[8658,5743,37785,9455,17738,47899,null,4585,37995,37851,9257,9257,4712],"quality":"More is better","isMain":false,"metricName":"NGA75"},{"values":[2,3,1,2,2,1,null,2,1,1,2,2,2],"quality":"Less is better","isMain":false,"metricName":"LG50"},{"values":[4,5,1,3,2,1,null,4,1,1,3,3,4],"quality":"Less is better","isMain":false,"metricName":"LG75"},{"values":[2,3,1,2,2,1,3,2,1,1,2,2,2],"quality":"Less is better","isMain":false,"metricName":"LA50"},{"values":[4,5,1,3,2,1,5,4,1,1,3,3,4],"quality":"Less is better","isMain":false,"metricName":"LA75"},{"values":[2,3,1,2,2,1,null,2,1,1,2,2,2],"quality":"Less is better","isMain":true,"metricName":"LGA50"},{"values":[4,5,1,3,2,1,null,4,1,1,3,3,4],"quality":"Less is better","isMain":false,"metricName":"LGA75"}]],["Misassemblies",[{"values":[0,0,0,0,0,0,0,0,0,0,0,0,0],"quality":"Less is better","isMain":true,"metricName":"# misassemblies"},{"values":[0,0,0,0,0,0,0,0,0,0,0,0,0],"quality":"Less is better","isMain":false,"metricName":" # relocations"},{"values":[0,0,0,0,0,0,0,0,0,0,0,0,0],"quality":"Less is better","isMain":false,"metricName":" # translocations"},{"values":[0,0,0,0,0,0,0,0,0,0,0,0,0],"quality":"Less is better","isMain":false,"metricName":" # inversions"},{"values":[0,0,0,0,0,0,0,0,0,0,0,0,0],"quality":"Less is better","isMain":false,"metricName":"# misassembled contigs"},{"values":[0,0,0,0,0,0,0,0,0,0,0,0,0],"quality":"Less is better","isMain":true,"metricName":"Misassembled contigs length"},{"values":[0,0,0,0,0,0,0,0,0,0,0,0,0],"quality":"Less is better","isMain":false,"metricName":"# local misassemblies"},{"values":[0,0,0,0,0,0,0,0,0,0,0,0,0],"quality":"Less is better","isMain":false,"metricName":"# unaligned mis. contigs"}]],["Unaligned",[{"values":[0,0,0,0,0,0,0,0,0,0,0,0,0],"quality":"Less is better","isMain":false,"metricName":"# fully unaligned contigs"},{"values":[0,0,0,0,0,0,0,0,0,0,0,0,0],"quality":"Less is better","isMain":false,"metricName":"Fully unaligned length"},{"values":[0,0,0,0,0,0,0,0,0,0,0,0,0],"quality":"Less is better","isMain":false,"metricName":"# partially unaligned contigs"},{"values":[0,0,0,0,0,0,0,0,0,0,0,0,0],"quality":"Less is better","isMain":false,"metricName":"Partially unaligned length"}]],["Mismatches",[{"values":[2,0,0,0,0,0,0,1,0,0,0,0,0],"quality":"Less is better","isMain":false,"metricName":"# mismatches"},{"values":[0,0,0,0,0,1,0,14,0,0,0,0,0],"quality":"Less is better","isMain":false,"metricName":"# indels"},{"values":[0,0,0,0,0,27,0,651,0,0,0,0,0],"quality":"Less is better","isMain":false,"metricName":"Indels length"},{"values":["4.22","0.00","0.00","0.00","0.00","0.00","0.00","2.20","0.00","0.00","0.00","0.00","0.00"],"quality":"Less is better","isMain":true,"metricName":"# mismatches per 100 kbp"},{"values":["0.00","0.00","0.00","0.00","0.00","2.09","0.00","30.82","0.00","0.00","0.00","0.00","0.00"],"quality":"Less is better","isMain":true,"metricName":"# indels per 100 kbp"},{"values":[0,0,0,0,0,0,0,0,0,0,0,0,0],"quality":"Less is better","isMain":false,"metricName":" # indels (<= 5 bp)"},{"values":[0,0,0,0,0,1,0,14,0,0,0,0,0],"quality":"Less is better","isMain":false,"metricName":" # indels (> 5 bp)"},{"values":[0,0,0,0,0,0,0,1292,0,0,0,0,0],"quality":"Less is better","isMain":false,"metricName":"# N's"},{"values":["0.00","0.00","0.00","0.00","0.00","0.00","0.00","2768.61","0.00","0.00","0.00","0.00","0.00"],"quality":"Less is better","isMain":true,"metricName":"# N's per 100 kbp"}]],["Statistics without reference",[{"values":[6,9,2,5,4,1,6,6,2,2,6,6,6],"quality":"Equal","isMain":true,"metricName":"# contigs"},{"values":[5,6,2,4,3,1,0,6,2,2,5,5,6],"quality":"Equal","isMain":false,"metricName":"# contigs (>= 1000 bp)"},{"values":[4,5,2,3,3,1,0,3,2,2,4,4,3],"quality":"Equal","isMain":false,"metricName":"# contigs (>= 5000 bp)"},{"values":[1,0,1,2,2,1,0,1,1,1,2,2,2],"quality":"Equal","isMain":false,"metricName":"# contigs (>= 10000 bp)"},{"values":[0,0,1,0,0,1,0,0,1,1,0,0,0],"quality":"Equal","isMain":false,"metricName":"# contigs (>= 25000 bp)"},{"values":[0,0,0,0,0,0,0,0,0,0,0,0,0],"quality":"Equal","isMain":false,"metricName":"# contigs (>= 50000 bp)"},{"values":[15621,9381,37785,17430,19844,47899,863,17581,37995,37851,15624,15624,15574],"quality":"More is better","isMain":true,"metricName":"Largest contig"},{"values":[47651,46917,47086,47254,47750,47899,3786,46666,47252,47108,47140,47140,45380],"quality":"More is better","isMain":true,"metricName":"Total length"},{"values":[46741,44528,47086,46709,47121,47899,0,46666,47252,47108,46301,46301,45380],"quality":"More is better","isMain":true,"metricName":"Total length (>= 1000 bp)"},{"values":[42575,40490,47086,42597,47121,47899,0,35575,47252,47108,42277,42277,32898],"quality":"More is better","isMain":false,"metricName":"Total length (>= 5000 bp)"},{"values":[15621,0,37785,33142,37582,47899,0,17581,37995,37851,27457,27457,27357],"quality":"More is better","isMain":true,"metricName":"Total length (>= 10000 bp)"},{"values":[0,0,37785,0,0,47899,0,0,37995,37851,0,0,0],"quality":"More is better","isMain":false,"metricName":"Total length (>= 25000 bp)"},{"values":[0,0,0,0,0,0,0,0,0,0,0,0,0],"quality":"More is better","isMain":true,"metricName":"Total length (>= 50000 bp)"},{"values":[9509,8632,37785,15712,17738,47899,620,9499,37995,37851,11833,11833,11783],"quality":"More is better","isMain":false,"metricName":"N50"},{"values":[8658,5743,37785,9455,17738,47899,570,8495,37995,37851,9257,9257,4712],"quality":"More is better","isMain":false,"metricName":"N75"},{"values":[2,3,1,2,2,1,3,2,1,1,2,2,2],"quality":"Less is better","isMain":false,"metricName":"L50"},{"values":[4,5,1,3,2,1,5,3,1,1,3,3,4],"quality":"Less is better","isMain":false,"metricName":"L75"},{"values":["53.24","53.28","53.24","53.25","53.25","53.21","53.20","53.33","53.20","53.22","53.23","53.23","53.32"],"quality":"Equal","isMain":false,"metricName":"GC (%)"}]],["Predicted genes",[]],["Similarity statistics",[{"values":[0,0,0,0,0,0,0,0,0,0,0,0,0],"quality":"Equal","isMain":false,"metricName":"# similar correct contigs"},{"values":[0,0,0,0,0,0,0,0,0,0,0,0,0],"quality":"Equal","isMain":false,"metricName":"# similar misassembled blocks"}]],["Reference statistics",[{"values":[47872,47872,47872,47872,47872,47872,47872,47872,47872,47872,47872,47872,47872],"quality":"Equal","isMain":false,"metricName":"Reference length"},{"values":[1,1,1,1,1,1,1,1,1,1,1,1,1],"quality":"Equal","isMain":false,"metricName":"Reference fragments"},{"values":["53.21","53.21","53.21","53.21","53.21","53.21","53.21","53.21","53.21","53.21","53.21","53.21","53.21"],"quality":"Equal","isMain":false,"metricName":"Reference GC (%)"}]]],"referenceName":"gi\_18640469\_ref\_NC","date":"18 July 2018, Wednesday, 16:38:43","order":[0,1,2,3,4,5,6,7,8,9,10,11,12],"assembliesNames":["ABySS\_127","ABySS\_63","CLC","IDBA\_UD","MEGAHIT","MIRA","Ray\_Meta","SOAPdenovo2","SPAdes","SPAdes\_meta","SPAdes\_sc","SPAdes\_sc\_careful","Velvet"]},{"assembliesWithNs":null,"minContig":500,"report":[["Genome statistics",[{"values":["13.706","7.400","12.878","12.253","13.633","13.083","14.859","14.874"],"quality":"More is better","isMain":true,"metricName":"Genome fraction (%)"},{"values":["1.001","1.002","1.000","1.000","1.000","1.000","1.000","1.000"],"quality":"Less is better","isMain":true,"metricName":"Duplication ratio"},{"values":[1517,1465,1491,1325,1813,1813,1813,1813],"quality":"More is better","isMain":true,"metricName":"Largest alignment"},{"values":[18354,9895,17220,16384,18230,17495,19869,19889],"quality":"More is better","isMain":true,"metricName":"Total aligned length"},{"values":[773,704,748,773,773,773,773,773],"quality":"More is better","isMain":false,"metricName":"NA50"},{"values":[651,618,625,615,625,651,625,625],"quality":"More is better","isMain":false,"metricName":"NA75"},{"values":[null,null,null,null,null,null,null,null],"quality":"More is better","isMain":true,"metricName":"NGA50"},{"values":[10,5,10,9,9,9,10,10],"quality":"Less is better","isMain":false,"metricName":"LA50"},{"values":[17,9,16,15,16,15,17,17],"quality":"Less is better","isMain":false,"metricName":"LA75"}]],["Misassemblies",[{"values":[0,0,0,0,0,0,0,0],"quality":"Less is better","isMain":true,"metricName":"# misassemblies"},{"values":[0,0,0,0,0,0,0,0],"quality":"Less is better","isMain":false,"metricName":" # relocations"},{"values":[0,0,0,0,0,0,0,0],"quality":"Less is better","isMain":false,"metricName":" # translocations"},{"values":[0,0,0,0,0,0,0,0],"quality":"Less is better","isMain":false,"metricName":" # inversions"},{"values":[0,0,0,0,0,0,0,0],"quality":"Less is better","isMain":false,"metricName":"# misassembled contigs"},{"values":[0,0,0,0,0,0,0,0],"quality":"Less is better","isMain":true,"metricName":"Misassembled contigs length"},{"values":[0,0,0,0,0,0,1,1],"quality":"Less is better","isMain":false,"metricName":"# local misassemblies"},{"values":[0,0,0,0,0,0,0,0],"quality":"Less is better","isMain":false,"metricName":"# unaligned mis. contigs"}]],["Unaligned",[{"values":[0,0,0,0,0,0,0,0],"quality":"Less is better","isMain":false,"metricName":"# fully unaligned contigs"},{"values":[0,0,0,0,0,0,0,0],"quality":"Less is better","isMain":false,"metricName":"Fully unaligned length"},{"values":[0,0,0,0,0,0,0,0],"quality":"Less is better","isMain":false,"metricName":"# partially unaligned contigs"},{"values":[0,0,0,0,0,0,0,0],"quality":"Less is better","isMain":false,"metricName":"Partially unaligned length"}]],["Mismatches",[{"values":[35,6,21,28,24,24,30,27],"quality":"Less is better","isMain":false,"metricName":"# mismatches"},{"values":[0,1,1,0,0,0,0,0],"quality":"Less is better","isMain":false,"metricName":"# indels"},{"values":[0,26,26,0,0,0,0,0],"quality":"Less is better","isMain":false,"metricName":"Indels length"},{"values":["190.96","60.64","121.95","170.90","131.65","137.18","150.99","135.75"],"quality":"Less is better","isMain":true,"metricName":"# mismatches per 100 kbp"},{"values":["0.00","10.11","5.81","0.00","0.00","0.00","0.00","0.00"],"quality":"Less is better","isMain":true,"metricName":"# indels per 100 kbp"},{"values":[0,0,0,0,0,0,0,0],"quality":"Less is better","isMain":false,"metricName":" # indels (<= 5 bp)"},{"values":[0,1,1,0,0,0,0,0],"quality":"Less is better","isMain":false,"metricName":" # indels (> 5 bp)"},{"values":[0,0,0,0,0,0,0,0],"quality":"Less is better","isMain":false,"metricName":"# N's"},{"values":["0.00","0.00","0.00","0.00","0.00","0.00","0.00","0.00"],"quality":"Less is better","isMain":true,"metricName":"# N's per 100 kbp"}]],["Statistics without reference",[{"values":[24,13,23,22,23,22,25,25],"quality":"Equal","isMain":true,"metricName":"# contigs"},{"values":[3,2,2,3,3,3,4,4],"quality":"Equal","isMain":false,"metricName":"# contigs (>= 1000 bp)"},{"values":[0,0,0,0,0,0,0,0],"quality":"Equal","isMain":false,"metricName":"# contigs (>= 5000 bp)"},{"values":[0,0,0,0,0,0,0,0],"quality":"Equal","isMain":false,"metricName":"# contigs (>= 10000 bp)"},{"values":[0,0,0,0,0,0,0,0],"quality":"Equal","isMain":false,"metricName":"# contigs (>= 25000 bp)"},{"values":[0,0,0,0,0,0,0,0],"quality":"Equal","isMain":false,"metricName":"# contigs (>= 50000 bp)"},{"values":[1517,1465,1491,1325,1813,1813,1813,1813],"quality":"More is better","isMain":true,"metricName":"Largest contig"},{"values":[18354,9914,17220,16384,18230,17495,19869,19889],"quality":"More is better","isMain":true,"metricName":"Total length"},{"values":[3557,2555,2644,3381,4347,4230,5455,5455],"quality":"More is better","isMain":true,"metricName":"Total length (>= 1000 bp)"},{"values":[0,0,0,0,0,0,0,0],"quality":"More is better","isMain":false,"metricName":"Total length (>= 5000 bp)"},{"values":[0,0,0,0,0,0,0,0],"quality":"More is better","isMain":true,"metricName":"Total length (>= 10000 bp)"},{"values":[0,0,0,0,0,0,0,0],"quality":"More is better","isMain":false,"metricName":"Total length (>= 25000 bp)"},{"values":[0,0,0,0,0,0,0,0],"quality":"More is better","isMain":true,"metricName":"Total length (>= 50000 bp)"},{"values":[773,704,748,773,773,773,773,773],"quality":"More is better","isMain":false,"metricName":"N50"},{"values":[651,618,625,615,625,651,625,625],"quality":"More is better","isMain":false,"metricName":"N75"},{"values":[10,5,10,9,9,9,10,10],"quality":"Less is better","isMain":false,"metricName":"L50"},{"values":[17,9,16,15,16,15,17,17],"quality":"Less is better","isMain":false,"metricName":"L75"},{"values":["48.94","47.75","49.76","48.98","49.33","49.31","49.52","49.53"],"quality":"Equal","isMain":false,"metricName":"GC (%)"}]],["Predicted genes",[]],["Similarity statistics",[{"values":[0,0,0,0,0,0,0,0],"quality":"Equal","isMain":false,"metricName":"# similar correct contigs"},{"values":[0,0,0,0,0,0,0,0],"quality":"Equal","isMain":false,"metricName":"# similar misassembled blocks"}]],["Reference statistics",[{"values":[133719,133719,133719,133719,133719,133719,133719,133719],"quality":"Equal","isMain":false,"metricName":"Reference length"},{"values":[1,1,1,1,1,1,1,1],"quality":"Equal","isMain":false,"metricName":"Reference fragments"},{"values":["52.47","52.47","52.47","52.47","52.47","52.47","52.47","52.47"],"quality":"Equal","isMain":false,"metricName":"Reference GC (%)"}]]],"referenceName":"gi\_18653809\_ref\_NC","date":"18 July 2018, Wednesday, 16:38:53","order":[0,1,2,3,4,5,6,7],"assembliesNames":["CLC","IDBA\_UD","MEGAHIT","SOAPdenovo2","SPAdes","SPAdes\_meta","SPAdes\_sc","SPAdes\_sc\_careful"]},{"assembliesWithNs":null,"minContig":500,"report":[["Genome statistics",[{"values":["48.354","16.542","49.206","27.635","44.608","96.166","56.127","1.319","62.749","22.750","34.211","48.604","7.431","40.984"],"quality":"More is better","isMain":true,"metricName":"Genome fraction (%)"},{"values":["1.007","1.000","1.030","1.014","1.036","1.742","1.059","1.000","1.037","1.037","1.011","1.019","1.000","1.204"],"quality":"Less is better","isMain":true,"metricName":"Duplication ratio"},{"values":[6002,2344,3247,2988,3891,12772,6469,585,5701,4142,2422,4391,1126,10008],"quality":"More is better","isMain":true,"metricName":"Largest alignment"},{"values":[21589,7336,22133,12420,20096,73615,26353,585,28840,10088,15334,21876,3295,21878],"quality":"More is better","isMain":true,"metricName":"Total aligned length"},{"values":[515,null,501,null,null,12207,1095,null,922,null,null,null,null,33030],"quality":"More is better","isMain":false,"metricName":"NG50"},{"values":[null,null,null,null,null,9674,null,null,null,null,null,null,null,28563],"quality":"More is better","isMain":false,"metricName":"NG75"},{"values":[880,1597,1155,999,974,4712,2755,585,1498,3611,1655,1097,858,null],"quality":"More is better","isMain":false,"metricName":"NA50"},{"values":[595,714,675,680,628,1135,1427,585,922,1048,1048,801,664,null],"quality":"More is better","isMain":false,"metricName":"NA75"},{"values":[null,null,null,null,null,7799,1095,null,922,null,null,null,null,null],"quality":"More is better","isMain":true,"metricName":"NGA50"},{"values":[null,null,null,null,null,6431,null,null,null,null,null,null,null,null],"quality":"More is better","isMain":false,"metricName":"NGA75"},{"values":[20,null,23,null,null,2,9,null,12,null,null,null,null,1],"quality":"Less is better","isMain":false,"metricName":"LG50"},{"values":[null,null,null,null,null,3,null,null,null,null,null,null,null,2],"quality":"Less is better","isMain":false,"metricName":"LG75"},{"values":[6,2,7,4,6,6,3,1,6,2,4,5,2,null],"quality":"Less is better","isMain":false,"metricName":"LA50"},{"values":[15,4,15,8,13,16,7,1,12,3,7,10,3,null],"quality":"Less is better","isMain":false,"metricName":"LA75"},{"values":[null,null,null,null,null,3,9,null,12,null,null,null,null,null],"quality":"Less is better","isMain":true,"metricName":"LGA50"},{"values":[null,null,null,null,null,4,null,null,null,null,null,null,null,null],"quality":"Less is better","isMain":false,"metricName":"LGA75"}]],["Misassemblies",[{"values":[0,0,0,0,0,0,0,0,0,0,0,0,0,0],"quality":"Less is better","isMain":true,"metricName":"# misassemblies"},{"values":[0,0,0,0,0,0,0,0,0,0,0,0,0,0],"quality":"Less is better","isMain":false,"metricName":" # relocations"},{"values":[0,0,0,0,0,0,0,0,0,0,0,0,0,0],"quality":"Less is better","isMain":false,"metricName":" # translocations"},{"values":[0,0,0,0,0,0,0,0,0,0,0,0,0,0],"quality":"Less is better","isMain":false,"metricName":" # inversions"},{"values":[0,0,0,0,0,0,0,0,0,0,0,0,0,0],"quality":"Less is better","isMain":false,"metricName":"# misassembled contigs"},{"values":[0,0,0,0,0,0,0,0,0,0,0,0,0,0],"quality":"Less is better","isMain":true,"metricName":"Misassembled contigs length"},{"values":[1,0,0,0,0,6,0,0,0,0,0,0,0,0],"quality":"Less is better","isMain":false,"metricName":"# local misassemblies"},{"values":[0,0,0,0,0,0,0,0,0,0,0,0,0,2],"quality":"Less is better","isMain":false,"metricName":"# unaligned mis. contigs"}]],["Unaligned",[{"values":[0,0,0,0,0,0,0,0,0,0,0,0,0,0],"quality":"Less is better","isMain":false,"metricName":"# fully unaligned contigs"},{"values":[0,0,0,0,0,0,0,0,0,0,0,0,0,0],"quality":"Less is better","isMain":false,"metricName":"Fully unaligned length"},{"values":[1,0,0,0,1,5,0,0,0,0,0,0,0,4],"quality":"Less is better","isMain":false,"metricName":"# partially unaligned contigs"},{"values":[2108,0,0,0,862,10463,0,0,0,0,0,0,0,62142],"quality":"Less is better","isMain":false,"metricName":"Partially unaligned length"}]],["Mismatches",[{"values":[10,0,45,0,19,361,166,0,14,107,56,100,3,204],"quality":"Less is better","isMain":false,"metricName":"# mismatches"},{"values":[0,1,0,0,0,5,3,0,0,0,1,0,0,13],"quality":"Less is better","isMain":false,"metricName":"# indels"},{"values":[0,1,0,0,0,5,6,0,0,0,1,0,0,18],"quality":"Less is better","isMain":false,"metricName":"Indels length"},{"values":["46.64","0.00","206.24","0.00","96.06","846.58","666.99","0.00","50.32","1060.67","369.15","463.99","91.05","1122.54"],"quality":"Less is better","isMain":true,"metricName":"# mismatches per 100 kbp"},{"values":["0.00","13.63","0.00","0.00","0.00","11.73","12.05","0.00","0.00","0.00","6.59","0.00","0.00","71.53"],"quality":"Less is better","isMain":true,"metricName":"# indels per 100 kbp"},{"values":[0,1,0,0,0,5,3,0,0,0,1,0,0,13],"quality":"Less is better","isMain":false,"metricName":" # indels (<= 5 bp)"},{"values":[0,0,0,0,0,0,0,0,0,0,0,0,0,0],"quality":"Less is better","isMain":false,"metricName":" # indels (> 5 bp)"},{"values":[50,0,0,0,0,14,0,0,0,0,0,0,0,466],"quality":"Less is better","isMain":false,"metricName":"# N's"},{"values":["211.00","0.00","0.00","0.00","0.00","16.52","0.00","0.00","0.00","0.00","0.00","0.00","0.00","554.63"],"quality":"Less is better","isMain":true,"metricName":"# N's per 100 kbp"}]],["Statistics without reference",[{"values":[22,7,23,13,20,29,13,1,21,5,12,19,4,4],"quality":"Equal","isMain":true,"metricName":"# contigs"},{"values":[2,2,7,3,6,18,10,0,11,4,7,7,1,4],"quality":"Equal","isMain":false,"metricName":"# contigs (>= 1000 bp)"},{"values":[1,0,0,0,0,6,1,0,1,0,0,0,0,4],"quality":"Equal","isMain":false,"metricName":"# contigs (>= 5000 bp)"},{"values":[0,0,0,0,0,2,0,0,0,0,0,0,0,4],"quality":"Equal","isMain":false,"metricName":"# contigs (>= 10000 bp)"},{"values":[0,0,0,0,0,0,0,0,0,0,0,0,0,2],"quality":"Equal","isMain":false,"metricName":"# contigs (>= 25000 bp)"},{"values":[0,0,0,0,0,0,0,0,0,0,0,0,0,0],"quality":"Equal","isMain":false,"metricName":"# contigs (>= 50000 bp)"},{"values":[8110,2344,3247,2988,4753,12816,6471,585,5701,4142,2422,4391,1126,33030],"quality":"More is better","isMain":true,"metricName":"Largest contig"},{"values":[23697,7336,22483,12420,21362,84762,26355,585,28840,10464,15339,21953,3295,84020],"quality":"More is better","isMain":true,"metricName":"Total length"},{"values":[10450,3941,11657,5494,11690,77064,23935,0,21484,9841,11916,14130,1126,84020],"quality":"More is better","isMain":true,"metricName":"Total length (>= 1000 bp)"},{"values":[8110,0,0,0,0,55472,6471,0,5701,0,0,0,0,84020],"quality":"More is better","isMain":false,"metricName":"Total length (>= 5000 bp)"},{"values":[0,0,0,0,0,25023,0,0,0,0,0,0,0,84020],"quality":"More is better","isMain":true,"metricName":"Total length (>= 10000 bp)"},{"values":[0,0,0,0,0,0,0,0,0,0,0,0,0,61593],"quality":"More is better","isMain":false,"metricName":"Total length (>= 25000 bp)"},{"values":[0,0,0,0,0,0,0,0,0,0,0,0,0,0],"quality":"More is better","isMain":true,"metricName":"Total length (>= 50000 bp)"},{"values":[975,1597,1155,999,1032,7963,2755,585,1498,3611,1655,1097,858,28563],"quality":"More is better","isMain":false,"metricName":"N50"},{"values":[618,714,676,680,761,2764,1427,585,922,1048,1048,801,664,11268],"quality":"More is better","isMain":false,"metricName":"N75"},{"values":[4,2,7,4,6,4,3,1,6,2,4,5,2,2],"quality":"Less is better","isMain":false,"metricName":"L50"},{"values":[12,4,14,8,11,9,7,1,12,3,7,10,3,3],"quality":"Less is better","isMain":false,"metricName":"L75"},{"values":["34.39","31.95","34.48","33.68","34.66","34.19","33.98","34.53","34.22","33.68","33.45","33.69","29.98","34.88"],"quality":"Equal","isMain":false,"metricName":"GC (%)"}]],["Predicted genes",[]],["Similarity statistics",[{"values":[0,0,0,0,0,0,0,0,0,0,0,0,0,0],"quality":"Equal","isMain":false,"metricName":"# similar correct contigs"},{"values":[0,0,0,0,0,0,0,0,0,0,0,0,0,0],"quality":"Equal","isMain":false,"metricName":"# similar misassembled blocks"}]],["Reference statistics",[{"values":[44342,44342,44342,44342,44342,44342,44342,44342,44342,44342,44342,44342,44342,44342],"quality":"Equal","isMain":false,"metricName":"Reference length"},{"values":[1,1,1,1,1,1,1,1,1,1,1,1,1,1],"quality":"Equal","isMain":false,"metricName":"Reference fragments"},{"values":["34.33","34.33","34.33","34.33","34.33","34.33","34.33","34.33","34.33","34.33","34.33","34.33","34.33","34.33"],"quality":"Equal","isMain":false,"metricName":"Reference GC (%)"}]]],"referenceName":"gi\_189427122\_ref\_NC","date":"18 July 2018, Wednesday, 16:39:08","order":[0,1,2,3,4,5,6,7,8,9,10,11,12,13],"assembliesNames":["ABySS\_127","ABySS\_63","CLC","IDBA\_UD","MEGAHIT","MIRA","Ray\_Meta","SOAPdenovo2","SPAdes","SPAdes\_meta","SPAdes\_sc","SPAdes\_sc\_careful","Velvet","Geneious"]},{"assembliesWithNs":null,"minContig":500,"report":[["Genome statistics",[{"values":["96.195","94.259","99.909","99.896","99.838","99.070","0.423","97.987","99.961","99.916","99.961","99.961","99.842","79.030"],"quality":"More is better","isMain":true,"metricName":"Genome fraction (%)"},{"values":["1.002","1.002","1.004","1.000","1.000","1.012","1.000","1.017","1.001","1.001","1.000","1.000","1.001","1.000"],"quality":"Less is better","isMain":true,"metricName":"Duplication ratio"},{"values":[22045,14417,102511,223527,223469,56382,714,33592,223529,223529,223529,223529,68813,56382],"quality":"More is better","isMain":true,"metricName":"Largest alignment"},{"values":[305201,298999,317618,316365,316160,317560,1341,312101,316714,316571,316563,316563,316410,250268],"quality":"More is better","isMain":true,"metricName":"Total aligned length"},{"values":[4744,3463,52319,223527,223469,19133,null,16330,223529,223529,223529,223529,35961,25631],"quality":"More is better","isMain":false,"metricName":"NG50"},{"values":[2977,1987,45382,62154,82500,14292,null,8382,72201,72201,82608,82608,27358,6997],"quality":"More is better","isMain":false,"metricName":"NG75"},{"values":[5221,3895,52319,223527,223469,19133,714,16090,223529,223529,223529,223529,35961,44854],"quality":"More is better","isMain":false,"metricName":"NA50"},{"values":[3213,2051,45382,62154,82500,14292,627,8251,72201,72201,82608,82608,27358,20388],"quality":"More is better","isMain":false,"metricName":"NA75"},{"values":[4744,3463,52319,223527,223469,19133,null,16090,223529,223529,223529,223529,35961,25631],"quality":"More is better","isMain":true,"metricName":"NGA50"},{"values":[2977,1987,45382,62154,82500,14292,null,8251,72201,72201,82608,82608,27358,6997],"quality":"More is better","isMain":false,"metricName":"NGA75"},{"values":[20,26,3,1,1,4,null,7,1,1,1,1,4,4],"quality":"Less is better","isMain":false,"metricName":"LG50"},{"values":[41,58,4,2,2,9,null,14,2,2,2,2,6,9],"quality":"Less is better","isMain":false,"metricName":"LG75"},{"values":[18,24,3,1,1,4,1,7,1,1,1,1,4,3],"quality":"Less is better","isMain":false,"metricName":"LA50"},{"values":[38,51,4,2,2,9,2,14,2,2,2,2,6,5],"quality":"Less is better","isMain":false,"metricName":"LA75"},{"values":[20,26,3,1,1,4,null,7,1,1,1,1,4,4],"quality":"Less is better","isMain":true,"metricName":"LGA50"},{"values":[41,58,4,2,2,9,null,14,2,2,2,2,6,9],"quality":"Less is better","isMain":false,"metricName":"LGA75"}]],["Misassemblies",[{"values":[0,0,0,0,0,0,0,0,0,0,0,0,0,0],"quality":"Less is better","isMain":true,"metricName":"# misassemblies"},{"values":[0,0,0,0,0,0,0,0,0,0,0,0,0,0],"quality":"Less is better","isMain":false,"metricName":" # relocations"},{"values":[0,0,0,0,0,0,0,0,0,0,0,0,0,0],"quality":"Less is better","isMain":false,"metricName":" # translocations"},{"values":[0,0,0,0,0,0,0,0,0,0,0,0,0,0],"quality":"Less is better","isMain":false,"metricName":" # inversions"},{"values":[0,0,0,0,0,0,0,0,0,0,0,0,0,0],"quality":"Less is better","isMain":false,"metricName":"# misassembled contigs"},{"values":[0,0,0,0,0,0,0,0,0,0,0,0,0,0],"quality":"Less is better","isMain":true,"metricName":"Misassembled contigs length"},{"values":[0,0,0,0,0,0,0,1,0,0,0,0,0,0],"quality":"Less is better","isMain":false,"metricName":"# local misassemblies"},{"values":[0,0,0,0,0,0,0,0,0,0,0,0,0,0],"quality":"Less is better","isMain":false,"metricName":"# unaligned mis. contigs"}]],["Unaligned",[{"values":[0,0,0,0,0,0,0,0,0,0,0,0,0,0],"quality":"Less is better","isMain":false,"metricName":"# fully unaligned contigs"},{"values":[0,0,0,0,0,0,0,0,0,0,0,0,0,0],"quality":"Less is better","isMain":false,"metricName":"Fully unaligned length"},{"values":[0,0,0,0,0,0,0,1,0,0,0,0,0,0],"quality":"Less is better","isMain":false,"metricName":"# partially unaligned contigs"},{"values":[0,0,0,0,0,0,0,1279,0,0,0,0,0,0],"quality":"Less is better","isMain":false,"metricName":"Partially unaligned length"}]],["Mismatches",[{"values":[25,6,16,11,9,5,0,9,37,34,23,14,12,2],"quality":"Less is better","isMain":false,"metricName":"# mismatches"},{"values":[0,0,0,0,0,0,0,67,0,0,0,0,0,0],"quality":"Less is better","isMain":false,"metricName":"# indels"},{"values":[0,0,0,0,0,0,0,3270,0,0,0,0,0,0],"quality":"Less is better","isMain":false,"metricName":"Indels length"},{"values":["8.21","2.01","5.06","3.48","2.85","1.59","0.00","2.90","11.69","10.75","7.27","4.42","3.80","0.80"],"quality":"Less is better","isMain":true,"metricName":"# mismatches per 100 kbp"},{"values":["0.00","0.00","0.00","0.00","0.00","0.00","0.00","21.59","0.00","0.00","0.00","0.00","0.00","0.00"],"quality":"Less is better","isMain":true,"metricName":"# indels per 100 kbp"},{"values":[0,0,0,0,0,0,0,0,0,0,0,0,0,0],"quality":"Less is better","isMain":false,"metricName":" # indels (<= 5 bp)"},{"values":[0,0,0,0,0,0,0,67,0,0,0,0,0,0],"quality":"Less is better","isMain":false,"metricName":" # indels (> 5 bp)"},{"values":[0,0,0,0,0,65,0,5942,0,0,0,0,0,53],"quality":"Less is better","isMain":false,"metricName":"# N's"},{"values":["0.00","0.00","0.00","0.00","0.00","20.47","0.00","1875.75","0.00","0.00","0.00","0.00","0.00","21.18"],"quality":"Less is better","isMain":true,"metricName":"# N's per 100 kbp"}]],["Statistics without reference",[{"values":[86,107,9,5,3,25,2,31,4,4,3,3,16,10],"quality":"Equal","isMain":true,"metricName":"# contigs"},{"values":[74,89,9,5,3,17,0,30,4,4,3,3,16,10],"quality":"Equal","isMain":false,"metricName":"# contigs (>= 1000 bp)"},{"values":[19,14,7,5,3,16,0,22,4,4,3,3,12,10],"quality":"Equal","isMain":false,"metricName":"# contigs (>= 5000 bp)"},{"values":[3,2,7,4,3,10,0,11,4,4,3,3,8,6],"quality":"Equal","isMain":false,"metricName":"# contigs (>= 10000 bp)"},{"values":[0,0,5,2,2,3,0,3,2,2,2,2,6,4],"quality":"Equal","isMain":false,"metricName":"# contigs (>= 25000 bp)"},{"values":[0,0,3,2,2,2,0,0,2,2,2,2,1,2],"quality":"Equal","isMain":false,"metricName":"# contigs (>= 50000 bp)"},{"values":[22045,14417,102511,223527,223469,56382,714,34042,223529,223529,223529,223529,68813,56382],"quality":"More is better","isMain":true,"metricName":"Largest contig"},{"values":[305201,298999,317618,316365,316160,317560,1341,316780,316714,316571,316563,316563,316410,250268],"quality":"More is better","isMain":true,"metricName":"Total length"},{"values":[296125,285081,317618,316365,316160,312250,0,316108,316714,316571,316563,316563,316410,250268],"quality":"More is better","isMain":true,"metricName":"Total length (>= 1000 bp)"},{"values":[157777,107572,308121,316365,316160,310960,0,293810,316714,316571,316563,316563,306438,250268],"quality":"More is better","isMain":false,"metricName":"Total length (>= 5000 bp)"},{"values":[50441,26903,308121,306460,316160,262759,0,216492,316714,316571,316563,316563,275908,217966],"quality":"More is better","isMain":true,"metricName":"Total length (>= 10000 bp)"},{"values":[0,0,283120,285681,305969,153053,0,86902,295730,295730,306137,306137,246276,178684],"quality":"More is better","isMain":false,"metricName":"Total length (>= 25000 bp)"},{"values":[0,0,207862,285681,305969,108199,0,0,295730,295730,306137,306137,68813,108199],"quality":"More is better","isMain":true,"metricName":"Total length (>= 50000 bp)"},{"values":[5221,3895,52319,223527,223469,19133,714,16330,223529,223529,223529,223529,35961,44854],"quality":"More is better","isMain":false,"metricName":"N50"},{"values":[3213,2051,45382,62154,82500,14292,627,8382,72201,72201,82608,82608,27358,20388],"quality":"More is better","isMain":false,"metricName":"N75"},{"values":[18,24,3,1,1,4,1,7,1,1,1,1,4,3],"quality":"Less is better","isMain":false,"metricName":"L50"},{"values":[38,51,4,2,2,9,2,14,2,2,2,2,6,5],"quality":"Less is better","isMain":false,"metricName":"L75"},{"values":["45.35","45.31","45.34","45.35","45.35","45.36","46.83","45.34","45.35","45.35","45.34","45.35","45.34","45.35"],"quality":"Equal","isMain":false,"metricName":"GC (%)"}]],["Predicted genes",[]],["Similarity statistics",[{"values":[0,0,0,0,0,0,0,0,0,0,0,0,0,0],"quality":"Equal","isMain":false,"metricName":"# similar correct contigs"},{"values":[0,0,0,0,0,0,0,0,0,0,0,0,0,0],"quality":"Equal","isMain":false,"metricName":"# similar misassembled blocks"}]],["Reference statistics",[{"values":[316674,316674,316674,316674,316674,316674,316674,316674,316674,316674,316674,316674,316674,316674],"quality":"Equal","isMain":false,"metricName":"Reference length"},{"values":[1,1,1,1,1,1,1,1,1,1,1,1,1,1],"quality":"Equal","isMain":false,"metricName":"Reference fragments"},{"values":["45.34","45.34","45.34","45.34","45.34","45.34","45.34","45.34","45.34","45.34","45.34","45.34","45.34","45.34"],"quality":"Equal","isMain":false,"metricName":"Reference GC (%)"}]]],"referenceName":"gi\_189490161\_ref\_NC","date":"18 July 2018, Wednesday, 16:39:27","order":[0,1,2,3,4,5,6,7,8,9,10,11,12,13],"assembliesNames":["ABySS\_127","ABySS\_63","CLC","IDBA\_UD","MEGAHIT","MIRA","Ray\_Meta","SOAPdenovo2","SPAdes","SPAdes\_meta","SPAdes\_sc","SPAdes\_sc\_careful","Velvet","Geneious"]},{"assembliesWithNs":null,"minContig":500,"report":[["Genome statistics",[{"values":["100.000","99.921","99.957","100.000","100.000","100.000","100.000","100.000","100.000","100.000","100.000","98.500"],"quality":"More is better","isMain":true,"metricName":"Genome fraction (%)"},{"values":["1.018","1.000","1.000","1.003","1.004","1.013","1.022","1.003","1.001","1.001","1.001","1.001"],"quality":"Less is better","isMain":true,"metricName":"Duplication ratio"},{"values":[39853,39105,39119,39235,39277,39626,39987,39263,39191,39191,39191,22157],"quality":"More is better","isMain":true,"metricName":"Largest alignment"},{"values":[39853,39105,39119,39235,39277,39626,39987,39263,39191,39191,39191,38579],"quality":"More is better","isMain":true,"metricName":"Total aligned length"},{"values":[39853,39105,39136,39235,39277,39626,39987,39263,39191,39191,39191,22157],"quality":"More is better","isMain":false,"metricName":"NG50"},{"values":[39853,39105,39136,39235,39277,39626,39987,39263,39191,39191,39191,16422],"quality":"More is better","isMain":false,"metricName":"NG75"},{"values":[39853,39105,39119,39235,39277,39626,39987,39263,39191,39191,39191,22157],"quality":"More is better","isMain":false,"metricName":"NA50"},{"values":[39853,39105,39119,39235,39277,39626,39987,39263,39191,39191,39191,16422],"quality":"More is better","isMain":false,"metricName":"NA75"},{"values":[39853,39105,39119,39235,39277,39626,39987,39263,39191,39191,39191,22157],"quality":"More is better","isMain":true,"metricName":"NGA50"},{"values":[39853,39105,39119,39235,39277,39626,39987,39263,39191,39191,39191,16422],"quality":"More is better","isMain":false,"metricName":"NGA75"},{"values":[1,1,1,1,1,1,1,1,1,1,1,1],"quality":"Less is better","isMain":false,"metricName":"LG50"},{"values":[1,1,1,1,1,1,1,1,1,1,1,2],"quality":"Less is better","isMain":false,"metricName":"LG75"},{"values":[1,1,1,1,1,1,1,1,1,1,1,1],"quality":"Less is better","isMain":false,"metricName":"LA50"},{"values":[1,1,1,1,1,1,1,1,1,1,1,2],"quality":"Less is better","isMain":false,"metricName":"LA75"},{"values":[1,1,1,1,1,1,1,1,1,1,1,1],"quality":"Less is better","isMain":true,"metricName":"LGA50"},{"values":[1,1,1,1,1,1,1,1,1,1,1,2],"quality":"Less is better","isMain":false,"metricName":"LGA75"}]],["Misassemblies",[{"values":[0,0,0,0,0,0,0,0,0,0,0,0],"quality":"Less is better","isMain":true,"metricName":"# misassemblies"},{"values":[0,0,0,0,0,0,0,0,0,0,0,0],"quality":"Less is better","isMain":false,"metricName":" # relocations"},{"values":[0,0,0,0,0,0,0,0,0,0,0,0],"quality":"Less is better","isMain":false,"metricName":" # translocations"},{"values":[0,0,0,0,0,0,0,0,0,0,0,0],"quality":"Less is better","isMain":false,"metricName":" # inversions"},{"values":[0,0,0,0,0,0,0,0,0,0,0,0],"quality":"Less is better","isMain":false,"metricName":"# misassembled contigs"},{"values":[0,0,0,0,0,0,0,0,0,0,0,0],"quality":"Less is better","isMain":true,"metricName":"Misassembled contigs length"},{"values":[1,0,0,1,1,1,1,1,0,0,0,0],"quality":"Less is better","isMain":false,"metricName":"# local misassemblies"},{"values":[0,0,0,0,0,0,0,0,0,0,0,0],"quality":"Less is better","isMain":false,"metricName":"# unaligned mis. contigs"}]],["Unaligned",[{"values":[0,0,0,0,0,0,0,0,0,0,0,0],"quality":"Less is better","isMain":false,"metricName":"# fully unaligned contigs"},{"values":[0,0,0,0,0,0,0,0,0,0,0,0],"quality":"Less is better","isMain":false,"metricName":"Fully unaligned length"},{"values":[0,0,0,0,0,0,0,0,0,0,0,0],"quality":"Less is better","isMain":false,"metricName":"# partially unaligned contigs"},{"values":[0,0,0,0,0,0,0,0,0,0,0,0],"quality":"Less is better","isMain":false,"metricName":"Partially unaligned length"}]],["Mismatches",[{"values":[0,0,0,0,0,0,0,0,0,0,0,0],"quality":"Less is better","isMain":false,"metricName":"# mismatches"},{"values":[0,0,0,0,0,0,0,0,1,1,1,0],"quality":"Less is better","isMain":false,"metricName":"# indels"},{"values":[0,0,0,0,0,0,0,0,55,55,55,0],"quality":"Less is better","isMain":false,"metricName":"Indels length"},{"values":["0.00","0.00","0.00","0.00","0.00","0.00","0.00","0.00","0.00","0.00","0.00","0.00"],"quality":"Less is better","isMain":true,"metricName":"# mismatches per 100 kbp"},{"values":["0.00","0.00","0.00","0.00","0.00","0.00","0.00","0.00","2.56","2.56","2.56","0.00"],"quality":"Less is better","isMain":true,"metricName":"# indels per 100 kbp"},{"values":[0,0,0,0,0,0,0,0,0,0,0,0],"quality":"Less is better","isMain":false,"metricName":" # indels (<= 5 bp)"},{"values":[0,0,0,0,0,0,0,0,1,1,1,0],"quality":"Less is better","isMain":false,"metricName":" # indels (> 5 bp)"},{"values":[0,0,0,0,0,1,0,0,0,0,0,0],"quality":"Less is better","isMain":false,"metricName":"# N's"},{"values":["0.00","0.00","0.00","0.00","0.00","2.52","0.00","0.00","0.00","0.00","0.00","0.00"],"quality":"Less is better","isMain":true,"metricName":"# N's per 100 kbp"}]],["Statistics without reference",[{"values":[1,1,1,1,1,1,1,1,1,1,1,2],"quality":"Equal","isMain":true,"metricName":"# contigs"},{"values":[1,1,1,1,1,1,1,1,1,1,1,2],"quality":"Equal","isMain":false,"metricName":"# contigs (>= 1000 bp)"},{"values":[1,1,1,1,1,1,1,1,1,1,1,2],"quality":"Equal","isMain":false,"metricName":"# contigs (>= 5000 bp)"},{"values":[1,1,1,1,1,1,1,1,1,1,1,2],"quality":"Equal","isMain":false,"metricName":"# contigs (>= 10000 bp)"},{"values":[1,1,1,1,1,1,1,1,1,1,1,0],"quality":"Equal","isMain":false,"metricName":"# contigs (>= 25000 bp)"},{"values":[0,0,0,0,0,0,0,0,0,0,0,0],"quality":"Equal","isMain":false,"metricName":"# contigs (>= 50000 bp)"},{"values":[39853,39105,39136,39235,39277,39626,39987,39263,39191,39191,39191,22157],"quality":"More is better","isMain":true,"metricName":"Largest contig"},{"values":[39853,39105,39136,39235,39277,39626,39987,39263,39191,39191,39191,38579],"quality":"More is better","isMain":true,"metricName":"Total length"},{"values":[39853,39105,39136,39235,39277,39626,39987,39263,39191,39191,39191,38579],"quality":"More is better","isMain":true,"metricName":"Total length (>= 1000 bp)"},{"values":[39853,39105,39136,39235,39277,39626,39987,39263,39191,39191,39191,38579],"quality":"More is better","isMain":false,"metricName":"Total length (>= 5000 bp)"},{"values":[39853,39105,39136,39235,39277,39626,39987,39263,39191,39191,39191,38579],"quality":"More is better","isMain":true,"metricName":"Total length (>= 10000 bp)"},{"values":[39853,39105,39136,39235,39277,39626,39987,39263,39191,39191,39191,0],"quality":"More is better","isMain":false,"metricName":"Total length (>= 25000 bp)"},{"values":[0,0,0,0,0,0,0,0,0,0,0,0],"quality":"More is better","isMain":true,"metricName":"Total length (>= 50000 bp)"},{"values":[39853,39105,39136,39235,39277,39626,39987,39263,39191,39191,39191,22157],"quality":"More is better","isMain":false,"metricName":"N50"},{"values":[39853,39105,39136,39235,39277,39626,39987,39263,39191,39191,39191,16422],"quality":"More is better","isMain":false,"metricName":"N75"},{"values":[1,1,1,1,1,1,1,1,1,1,1,1],"quality":"Less is better","isMain":false,"metricName":"L50"},{"values":[1,1,1,1,1,1,1,1,1,1,1,2],"quality":"Less is better","isMain":false,"metricName":"L75"},{"values":["40.43","40.50","40.50","40.51","40.50","40.48","40.50","40.50","40.49","40.49","40.49","40.54"],"quality":"Equal","isMain":false,"metricName":"GC (%)"}]],["Predicted genes",[]],["Similarity statistics",[{"values":[0,0,0,0,0,0,0,0,0,0,0,0],"quality":"Equal","isMain":false,"metricName":"# similar correct contigs"},{"values":[0,0,0,0,0,0,0,0,0,0,0,0],"quality":"Equal","isMain":false,"metricName":"# similar misassembled blocks"}]],["Reference statistics",[{"values":[39136,39136,39136,39136,39136,39136,39136,39136,39136,39136,39136,39136],"quality":"Equal","isMain":false,"metricName":"Reference length"},{"values":[1,1,1,1,1,1,1,1,1,1,1,1],"quality":"Equal","isMain":false,"metricName":"Reference fragments"},{"values":["40.51","40.51","40.51","40.51","40.51","40.51","40.51","40.51","40.51","40.51","40.51","40.51"],"quality":"Equal","isMain":false,"metricName":"Reference GC (%)"}]]],"referenceName":"gi\_190151415\_ref\_NC","date":"18 July 2018, Wednesday, 16:39:40","order":[0,1,2,3,4,5,6,7,8,9,10,11],"assembliesNames":["ABySS\_127","ABySS\_63","CLC","IDBA\_UD","MEGAHIT","MIRA","Ray\_Meta","SPAdes","SPAdes\_meta","SPAdes\_sc","SPAdes\_sc\_careful","Velvet"]},{"assembliesWithNs":null,"minContig":500,"report":[["Genome statistics",[{"values":["19.716","13.784","59.727","51.837","67.568","29.313","42.468","69.566","72.303","60.231","60.231","12.570","6.792"],"quality":"More is better","isMain":true,"metricName":"Genome fraction (%)"},{"values":["1.003","1.000","1.034","1.021","1.005","1.090","1.007","1.007","1.001","1.008","1.008","1.000","1.000"],"quality":"Less is better","isMain":true,"metricName":"Duplication ratio"},{"values":[1508,1207,3066,2293,2613,2384,1813,3066,3095,3065,3065,1486,2384],"quality":"More is better","isMain":true,"metricName":"Largest alignment"},{"values":[6940,4836,21479,18347,23793,11212,14899,24424,25394,21133,21133,4410,2384],"quality":"More is better","isMain":true,"metricName":"Total aligned length"},{"values":[null,null,937,659,1129,null,null,1228,1228,1129,1129,null,null],"quality":"More is better","isMain":false,"metricName":"NG50"},{"values":[747,842,1841,1253,1771,1219,981,1926,1926,1919,1919,973,2384],"quality":"More is better","isMain":false,"metricName":"NA50"},{"values":[729,631,937,810,901,927,752,1129,1129,1129,1129,641,2384],"quality":"More is better","isMain":false,"metricName":"NA75"},{"values":[null,null,920,546,1129,null,null,1228,1228,913,913,null,null],"quality":"More is better","isMain":true,"metricName":"NGA50"},{"values":[null,null,9,13,10,null,null,8,9,9,9,null,null],"quality":"Less is better","isMain":false,"metricName":"LG50"},{"values":[4,3,5,6,6,4,6,5,6,5,5,2,1],"quality":"Less is better","isMain":false,"metricName":"LA50"},{"values":[6,5,9,11,11,6,10,9,10,9,9,4,1],"quality":"Less is better","isMain":false,"metricName":"LA75"},{"values":[null,null,10,15,10,null,null,8,9,10,10,null,null],"quality":"Less is better","isMain":true,"metricName":"LGA50"}]],["Misassemblies",[{"values":[0,0,0,0,0,0,0,0,0,0,0,0,0],"quality":"Less is better","isMain":true,"metricName":"# misassemblies"},{"values":[0,0,0,0,0,0,0,0,0,0,0,0,0],"quality":"Less is better","isMain":false,"metricName":" # relocations"},{"values":[0,0,0,0,0,0,0,0,0,0,0,0,0],"quality":"Less is better","isMain":false,"metricName":" # translocations"},{"values":[0,0,0,0,0,0,0,0,0,0,0,0,0],"quality":"Less is better","isMain":false,"metricName":" # inversions"},{"values":[0,0,0,0,0,0,0,0,0,0,0,0,0],"quality":"Less is better","isMain":false,"metricName":"# misassembled contigs"},{"values":[0,0,0,0,0,0,0,0,0,0,0,0,0],"quality":"Less is better","isMain":true,"metricName":"Misassembled contigs length"},{"values":[0,0,1,1,0,1,0,1,0,1,1,0,1],"quality":"Less is better","isMain":false,"metricName":"# local misassemblies"},{"values":[0,0,0,0,0,0,0,0,0,0,0,0,0],"quality":"Less is better","isMain":false,"metricName":"# unaligned mis. contigs"}]],["Unaligned",[{"values":[0,0,0,0,0,0,0,0,0,0,0,0,0],"quality":"Less is better","isMain":false,"metricName":"# fully unaligned contigs"},{"values":[0,0,0,0,0,0,0,0,0,0,0,0,0],"quality":"Less is better","isMain":false,"metricName":"Fully unaligned length"},{"values":[0,0,1,1,0,0,0,0,0,1,1,0,0],"quality":"Less is better","isMain":false,"metricName":"# partially unaligned contigs"},{"values":[0,0,520,845,0,0,0,0,0,661,661,0,0],"quality":"Less is better","isMain":false,"metricName":"Partially unaligned length"}]],["Mismatches",[{"values":[1,0,189,84,36,3,25,176,30,165,165,0,2],"quality":"Less is better","isMain":false,"metricName":"# mismatches"},{"values":[0,0,13,3,0,1,1,9,0,9,9,0,1],"quality":"Less is better","isMain":false,"metricName":"# indels"},{"values":[0,0,14,3,0,1,46,10,0,10,10,0,1],"quality":"Less is better","isMain":false,"metricName":"Indels length"},{"values":["14.46","0.00","901.98","461.89","151.87","29.17","167.80","721.13","118.27","780.84","780.84","0.00","83.93"],"quality":"Less is better","isMain":true,"metricName":"# mismatches per 100 kbp"},{"values":["0.00","0.00","62.04","16.50","0.00","9.72","6.71","36.88","0.00","42.59","42.59","0.00","41.96"],"quality":"Less is better","isMain":true,"metricName":"# indels per 100 kbp"},{"values":[0,0,13,3,0,1,0,9,0,9,9,0,1],"quality":"Less is better","isMain":false,"metricName":" # indels (<= 5 bp)"},{"values":[0,0,0,0,0,0,1,0,0,0,0,0,0],"quality":"Less is better","isMain":false,"metricName":" # indels (> 5 bp)"},{"values":[0,0,0,0,0,19,100,0,0,0,0,0,5],"quality":"Less is better","isMain":false,"metricName":"# N's"},{"values":["0.00","0.00","0.00","0.00","0.00","169.46","666.71","0.00","0.00","0.00","0.00","0.00","209.73"],"quality":"Less is better","isMain":true,"metricName":"# N's per 100 kbp"}]],["Statistics without reference",[{"values":[8,6,15,16,18,9,15,16,18,14,14,5,1],"quality":"Equal","isMain":true,"metricName":"# contigs"},{"values":[2,1,7,9,10,5,5,10,11,9,9,1,1],"quality":"Equal","isMain":false,"metricName":"# contigs (>= 1000 bp)"},{"values":[0,0,0,0,0,0,0,0,0,0,0,0,0],"quality":"Equal","isMain":false,"metricName":"# contigs (>= 5000 bp)"},{"values":[0,0,0,0,0,0,0,0,0,0,0,0,0],"quality":"Equal","isMain":false,"metricName":"# contigs (>= 10000 bp)"},{"values":[0,0,0,0,0,0,0,0,0,0,0,0,0],"quality":"Equal","isMain":false,"metricName":"# contigs (>= 25000 bp)"},{"values":[0,0,0,0,0,0,0,0,0,0,0,0,0],"quality":"Equal","isMain":false,"metricName":"# contigs (>= 50000 bp)"},{"values":[1508,1207,3066,2293,2634,2384,1813,3066,3095,3065,3065,1486,2384],"quality":"More is better","isMain":true,"metricName":"Largest contig"},{"values":[6940,4836,22192,19413,23814,11212,14999,24588,25394,21958,21958,4410,2384],"quality":"More is better","isMain":true,"metricName":"Total length"},{"values":[2530,1207,15919,14571,17716,7697,7394,20044,20421,17995,17995,1486,2384],"quality":"More is better","isMain":true,"metricName":"Total length (>= 1000 bp)"},{"values":[0,0,0,0,0,0,0,0,0,0,0,0,0],"quality":"More is better","isMain":false,"metricName":"Total length (>= 5000 bp)"},{"values":[0,0,0,0,0,0,0,0,0,0,0,0,0],"quality":"More is better","isMain":true,"metricName":"Total length (>= 10000 bp)"},{"values":[0,0,0,0,0,0,0,0,0,0,0,0,0],"quality":"More is better","isMain":false,"metricName":"Total length (>= 25000 bp)"},{"values":[0,0,0,0,0,0,0,0,0,0,0,0,0],"quality":"More is better","isMain":true,"metricName":"Total length (>= 50000 bp)"},{"values":[747,842,2099,1655,1771,1219,981,1926,1926,1926,1926,973,2384],"quality":"More is better","isMain":false,"metricName":"N50"},{"values":[729,631,990,1124,901,927,752,1129,1129,1228,1228,641,2384],"quality":"More is better","isMain":false,"metricName":"N75"},{"values":[4,3,5,6,6,4,6,5,6,5,5,2,1],"quality":"Less is better","isMain":false,"metricName":"L50"},{"values":[6,5,8,9,11,6,10,9,10,8,8,4,1],"quality":"Less is better","isMain":false,"metricName":"L75"},{"values":["57.10","59.00","57.44","57.35","58.13","56.48","57.37","57.87","57.82","57.40","57.40","56.44","48.97"],"quality":"Equal","isMain":false,"metricName":"GC (%)"}]],["Predicted genes",[]],["Similarity statistics",[{"values":[0,0,0,0,0,0,0,0,0,0,0,0,0],"quality":"Equal","isMain":false,"metricName":"# similar correct contigs"},{"values":[0,0,0,0,0,0,0,0,0,0,0,0,0],"quality":"Equal","isMain":false,"metricName":"# similar misassembled blocks"}]],["Reference statistics",[{"values":[35083,35083,35083,35083,35083,35083,35083,35083,35083,35083,35083,35083,35083],"quality":"Equal","isMain":false,"metricName":"Reference length"},{"values":[1,1,1,1,1,1,1,1,1,1,1,1,1],"quality":"Equal","isMain":false,"metricName":"Reference fragments"},{"values":["57.10","57.10","57.10","57.10","57.10","57.10","57.10","57.10","57.10","57.10","57.10","57.10","57.10"],"quality":"Equal","isMain":false,"metricName":"Reference GC (%)"}]]],"referenceName":"gi\_190340974\_ref\_NC","date":"18 July 2018, Wednesday, 16:39:55","order":[0,1,2,3,4,5,6,7,8,9,10,11,12],"assembliesNames":["ABySS\_127","ABySS\_63","CLC","IDBA\_UD","MEGAHIT","MIRA","SOAPdenovo2","SPAdes","SPAdes\_meta","SPAdes\_sc","SPAdes\_sc\_careful","Velvet","Geneious"]},{"assembliesWithNs":null,"minContig":500,"report":[["Genome statistics",[{"values":["17.952","4.822","16.668","22.872","23.673","59.226","6.674","21.317","8.252","20.910","11.956","3.688","16.691"],"quality":"More is better","isMain":true,"metricName":"Genome fraction (%)"},{"values":["1.010","1.021","1.044","1.027","1.063","1.163","1.157","1.027","1.052","1.015","1.029","1.000","1.000"],"quality":"Less is better","isMain":true,"metricName":"Duplication ratio"},{"values":[1238,710,2196,1559,2036,2739,694,1407,1243,4010,1181,753,5771],"quality":"More is better","isMain":true,"metricName":"Largest alignment"},{"values":[9363,2489,8603,11909,12697,34547,3445,11004,4315,10794,6170,1904,8618],"quality":"More is better","isMain":true,"metricName":"Total aligned length"},{"values":[null,null,null,null,null,1537,null,null,null,null,null,null,null],"quality":"More is better","isMain":false,"metricName":"NG50"},{"values":[820,669,1292,964,1029,1613,531,629,675,1171,663,599,5771],"quality":"More is better","isMain":false,"metricName":"NA50"},{"values":[670,560,713,659,761,1236,429,544,628,663,568,552,2847],"quality":"More is better","isMain":false,"metricName":"NA75"},{"values":[null,null,null,null,null,1304,null,null,null,null,null,null,null],"quality":"More is better","isMain":true,"metricName":"NGA50"},{"values":[null,null,null,null,null,12,null,null,null,null,null,null,null],"quality":"Less is better","isMain":false,"metricName":"LG50"},{"values":[5,2,3,5,5,9,4,6,3,3,4,2,1],"quality":"Less is better","isMain":false,"metricName":"LA50"},{"values":[8,3,5,9,9,15,6,11,5,6,7,3,2],"quality":"Less is better","isMain":false,"metricName":"LA75"},{"values":[null,null,null,null,null,14,null,null,null,null,null,null,null],"quality":"Less is better","isMain":true,"metricName":"LGA50"}]],["Misassemblies",[{"values":[0,0,0,0,1,0,0,0,0,0,0,0,0],"quality":"Less is better","isMain":true,"metricName":"# misassemblies"},{"values":[0,0,0,0,1,0,0,0,0,0,0,0,0],"quality":"Less is better","isMain":false,"metricName":" # relocations"},{"values":[0,0,0,0,0,0,0,0,0,0,0,0,0],"quality":"Less is better","isMain":false,"metricName":" # translocations"},{"values":[0,0,0,0,0,0,0,0,0,0,0,0,0],"quality":"Less is better","isMain":false,"metricName":" # inversions"},{"values":[0,0,0,0,1,0,0,0,0,0,0,0,0],"quality":"Less is better","isMain":false,"metricName":"# misassembled contigs"},{"values":[0,0,0,0,3212,0,0,0,0,0,0,0,0],"quality":"Less is better","isMain":true,"metricName":"Misassembled contigs length"},{"values":[0,0,1,0,0,4,0,0,0,0,0,0,3],"quality":"Less is better","isMain":false,"metricName":"# local misassemblies"},{"values":[0,0,0,0,0,0,1,0,0,0,0,0,0],"quality":"Less is better","isMain":false,"metricName":"# unaligned mis. contigs"}]],["Unaligned",[{"values":[0,0,0,0,0,0,0,0,0,0,0,0,0],"quality":"Less is better","isMain":false,"metricName":"# fully unaligned contigs"},{"values":[0,0,0,0,0,0,0,0,0,0,0,0,0],"quality":"Less is better","isMain":false,"metricName":"Fully unaligned length"},{"values":[0,0,0,0,1,1,0,0,0,0,0,0,1],"quality":"Less is better","isMain":false,"metricName":"# partially unaligned contigs"},{"values":[0,0,0,0,993,1737,0,0,0,0,0,0,1430],"quality":"Less is better","isMain":false,"metricName":"Partially unaligned length"}]],["Mismatches",[{"values":[25,9,66,69,126,172,1,53,25,77,13,0,124],"quality":"Less is better","isMain":false,"metricName":"# mismatches"},{"values":[4,0,4,5,4,1,2,2,0,0,1,0,4],"quality":"Less is better","isMain":false,"metricName":"# indels"},{"values":[7,0,7,11,4,1,100,2,0,0,2,0,4],"quality":"Less is better","isMain":false,"metricName":"Indels length"},{"values":["269.77","361.59","767.09","584.40","1031.10","562.59","29.03","481.64","586.85","713.36","210.63","0.00","1439.18"],"quality":"Less is better","isMain":true,"metricName":"# mismatches per 100 kbp"},{"values":["43.16","0.00","46.49","42.35","32.73","3.27","58.06","18.18","0.00","0.00","16.20","0.00","46.43"],"quality":"Less is better","isMain":true,"metricName":"# indels per 100 kbp"},{"values":[4,0,4,4,4,1,0,2,0,0,1,0,4],"quality":"Less is better","isMain":false,"metricName":" # indels (<= 5 bp)"},{"values":[0,0,0,1,0,0,2,0,0,0,0,0,0],"quality":"Less is better","isMain":false,"metricName":" # indels (> 5 bp)"},{"values":[0,0,0,0,0,59,542,0,0,0,0,0,86],"quality":"Less is better","isMain":false,"metricName":"# N's"},{"values":["0.00","0.00","0.00","0.00","0.00","158.26","13594.18","0.00","0.00","0.00","0.00","0.00","855.89"],"quality":"Less is better","isMain":true,"metricName":"# N's per 100 kbp"}]],["Statistics without reference",[{"values":[12,4,8,14,12,21,7,15,6,10,9,3,2],"quality":"Equal","isMain":true,"metricName":"# contigs"},{"values":[2,0,3,4,5,20,0,3,1,3,1,0,2],"quality":"Equal","isMain":false,"metricName":"# contigs (>= 1000 bp)"},{"values":[0,0,0,0,0,0,0,0,0,0,0,0,1],"quality":"Equal","isMain":false,"metricName":"# contigs (>= 5000 bp)"},{"values":[0,0,0,0,0,0,0,0,0,0,0,0,0],"quality":"Equal","isMain":false,"metricName":"# contigs (>= 10000 bp)"},{"values":[0,0,0,0,0,0,0,0,0,0,0,0,0],"quality":"Equal","isMain":false,"metricName":"# contigs (>= 25000 bp)"},{"values":[0,0,0,0,0,0,0,0,0,0,0,0,0],"quality":"Equal","isMain":false,"metricName":"# contigs (>= 50000 bp)"},{"values":[1238,710,2578,1559,3212,4210,694,1698,1243,4010,1347,753,7201],"quality":"More is better","isMain":true,"metricName":"Largest contig"},{"values":[9363,2541,8985,12121,13987,37281,3987,11298,4481,10960,6350,1904,10048],"quality":"More is better","isMain":true,"metricName":"Total length"},{"values":[2377,0,5657,5345,8969,36350,0,4190,1243,6530,1347,0,10048],"quality":"More is better","isMain":true,"metricName":"Total length (>= 1000 bp)"},{"values":[0,0,0,0,0,0,0,0,0,0,0,0,7201],"quality":"More is better","isMain":false,"metricName":"Total length (>= 5000 bp)"},{"values":[0,0,0,0,0,0,0,0,0,0,0,0,0],"quality":"More is better","isMain":true,"metricName":"Total length (>= 10000 bp)"},{"values":[0,0,0,0,0,0,0,0,0,0,0,0,0],"quality":"More is better","isMain":false,"metricName":"Total length (>= 25000 bp)"},{"values":[0,0,0,0,0,0,0,0,0,0,0,0,0],"quality":"More is better","isMain":true,"metricName":"Total length (>= 50000 bp)"},{"values":[820,669,1292,964,1286,1762,541,635,675,1171,663,599,7201],"quality":"More is better","isMain":false,"metricName":"N50"},{"values":[670,602,713,659,853,1508,520,563,628,663,573,552,2847],"quality":"More is better","isMain":false,"metricName":"N75"},{"values":[5,2,3,5,4,8,4,5,3,3,4,2,1],"quality":"Less is better","isMain":false,"metricName":"L50"},{"values":[8,3,5,9,7,13,6,10,5,6,7,3,2],"quality":"Less is better","isMain":false,"metricName":"L75"},{"values":["62.92","62.97","62.27","63.06","63.24","63.70","62.24","62.39","62.62","62.97","62.17","62.66","63.58"],"quality":"Equal","isMain":false,"metricName":"GC (%)"}]],["Predicted genes",[]],["Similarity statistics",[{"values":[0,0,0,0,0,0,0,0,0,0,0,0,0],"quality":"Equal","isMain":false,"metricName":"# similar correct contigs"},{"values":[0,0,0,0,0,0,0,0,0,0,0,0,0],"quality":"Equal","isMain":false,"metricName":"# similar misassembled blocks"}]],["Reference statistics",[{"values":[51621,51621,51621,51621,51621,51621,51621,51621,51621,51621,51621,51621,51621],"quality":"Equal","isMain":false,"metricName":"Reference length"},{"values":[1,1,1,1,1,1,1,1,1,1,1,1,1],"quality":"Equal","isMain":false,"metricName":"Reference fragments"},{"values":["63.39","63.39","63.39","63.39","63.39","63.39","63.39","63.39","63.39","63.39","63.39","63.39","63.39"],"quality":"Equal","isMain":false,"metricName":"Reference GC (%)"}]]],"referenceName":"gi\_192823813\_ref\_NC","date":"18 July 2018, Wednesday, 16:40:09","order":[0,1,2,3,4,5,6,7,8,9,10,11,12],"assembliesNames":["ABySS\_127","ABySS\_63","CLC","IDBA\_UD","MEGAHIT","MIRA","SOAPdenovo2","SPAdes","SPAdes\_meta","SPAdes\_sc","SPAdes\_sc\_careful","Velvet","Geneious"]},{"assembliesWithNs":null,"minContig":500,"report":[["Genome statistics",[{"values":["3.192","9.926","13.614","13.399","2.957","18.219","2.692","8.211","11.233"],"quality":"More is better","isMain":true,"metricName":"Genome fraction (%)"},{"values":["1.000","1.002","1.038","1.282","1.000","1.000","1.000","1.000","1.000"],"quality":"Less is better","isMain":true,"metricName":"Duplication ratio"},{"values":[942,802,1242,3610,934,1540,802,1231,1231],"quality":"More is better","isMain":true,"metricName":"Largest alignment"},{"values":[1627,5063,6952,7697,1507,9286,1372,4185,5725],"quality":"More is better","isMain":true,"metricName":"Total aligned length"},{"values":[942,582,809,1278,934,655,802,915,915],"quality":"More is better","isMain":false,"metricName":"NA50"},{"values":[685,549,610,947,573,583,570,708,800],"quality":"More is better","isMain":false,"metricName":"NA75"},{"values":[null,null,null,null,null,null,null,null,null],"quality":"More is better","isMain":true,"metricName":"NGA50"},{"values":[1,5,4,2,1,6,1,2,3],"quality":"Less is better","isMain":false,"metricName":"LA50"},{"values":[2,7,7,4,2,10,2,4,5],"quality":"Less is better","isMain":false,"metricName":"LA75"}]],["Misassemblies",[{"values":[0,0,0,0,0,0,0,0,0],"quality":"Less is better","isMain":true,"metricName":"# misassemblies"},{"values":[0,0,0,0,0,0,0,0,0],"quality":"Less is better","isMain":false,"metricName":" # relocations"},{"values":[0,0,0,0,0,0,0,0,0],"quality":"Less is better","isMain":false,"metricName":" # translocations"},{"values":[0,0,0,0,0,0,0,0,0],"quality":"Less is better","isMain":false,"metricName":" # inversions"},{"values":[0,0,0,0,0,0,0,0,0],"quality":"Less is better","isMain":false,"metricName":"# misassembled contigs"},{"values":[0,0,0,0,0,0,0,0,0],"quality":"Less is better","isMain":true,"metricName":"Misassembled contigs length"},{"values":[0,0,0,3,0,0,0,0,0],"quality":"Less is better","isMain":false,"metricName":"# local misassemblies"},{"values":[0,0,0,0,0,0,0,0,0],"quality":"Less is better","isMain":false,"metricName":"# unaligned mis. contigs"}]],["Unaligned",[{"values":[0,0,0,0,0,0,0,0,0],"quality":"Less is better","isMain":false,"metricName":"# fully unaligned contigs"},{"values":[0,0,0,0,0,0,0,0,0],"quality":"Less is better","isMain":false,"metricName":"Fully unaligned length"},{"values":[0,1,1,0,0,1,0,0,0],"quality":"Less is better","isMain":false,"metricName":"# partially unaligned contigs"},{"values":[0,608,606,0,0,543,0,0,0],"quality":"Less is better","isMain":false,"metricName":"Partially unaligned length"}]],["Mismatches",[{"values":[6,41,92,58,3,24,0,15,32],"quality":"Less is better","isMain":false,"metricName":"# mismatches"},{"values":[0,7,7,6,0,0,0,0,0],"quality":"Less is better","isMain":false,"metricName":"# indels"},{"values":[0,8,15,13,0,0,0,0,0],"quality":"Less is better","isMain":false,"metricName":"Indels length"},{"values":["368.78","810.44","1325.84","849.32","199.07","258.45","0.00","358.42","558.95"],"quality":"Less is better","isMain":true,"metricName":"# mismatches per 100 kbp"},{"values":["0.00","138.37","100.88","87.86","0.00","0.00","0.00","0.00","0.00"],"quality":"Less is better","isMain":true,"metricName":"# indels per 100 kbp"},{"values":[0,7,6,5,0,0,0,0,0],"quality":"Less is better","isMain":false,"metricName":" # indels (<= 5 bp)"},{"values":[0,0,1,1,0,0,0,0,0],"quality":"Less is better","isMain":false,"metricName":" # indels (> 5 bp)"},{"values":[0,0,0,20,0,0,0,0,0],"quality":"Less is better","isMain":false,"metricName":"# N's"},{"values":["0.00","0.00","0.00","228.36","0.00","0.00","0.00","0.00","0.00"],"quality":"Less is better","isMain":true,"metricName":"# N's per 100 kbp"}]],["Statistics without reference",[{"values":[2,8,8,5,2,13,2,5,7],"quality":"Equal","isMain":true,"metricName":"# contigs"},{"values":[0,1,4,4,0,2,0,1,1],"quality":"Equal","isMain":false,"metricName":"# contigs (>= 1000 bp)"},{"values":[0,0,0,0,0,0,0,0,0],"quality":"Equal","isMain":false,"metricName":"# contigs (>= 5000 bp)"},{"values":[0,0,0,0,0,0,0,0,0],"quality":"Equal","isMain":false,"metricName":"# contigs (>= 10000 bp)"},{"values":[0,0,0,0,0,0,0,0,0],"quality":"Equal","isMain":false,"metricName":"# contigs (>= 25000 bp)"},{"values":[0,0,0,0,0,0,0,0,0],"quality":"Equal","isMain":false,"metricName":"# contigs (>= 50000 bp)"},{"values":[942,1336,1415,4408,934,1540,802,1231,1231],"quality":"More is better","isMain":true,"metricName":"Largest contig"},{"values":[1627,5675,7807,8758,1507,9829,1372,4187,5727],"quality":"More is better","isMain":true,"metricName":"Total length"},{"values":[0,1336,4926,7918,0,2716,0,1231,1231],"quality":"More is better","isMain":true,"metricName":"Total length (>= 1000 bp)"},{"values":[0,0,0,0,0,0,0,0,0],"quality":"More is better","isMain":false,"metricName":"Total length (>= 5000 bp)"},{"values":[0,0,0,0,0,0,0,0,0],"quality":"More is better","isMain":true,"metricName":"Total length (>= 10000 bp)"},{"values":[0,0,0,0,0,0,0,0,0],"quality":"More is better","isMain":false,"metricName":"Total length (>= 25000 bp)"},{"values":[0,0,0,0,0,0,0,0,0],"quality":"More is better","isMain":true,"metricName":"Total length (>= 50000 bp)"},{"values":[942,614,1063,4408,934,711,802,917,917],"quality":"More is better","isMain":false,"metricName":"N50"},{"values":[685,571,974,1210,573,623,570,708,800],"quality":"More is better","isMain":false,"metricName":"N75"},{"values":[1,4,4,1,1,5,1,2,3],"quality":"Less is better","isMain":false,"metricName":"L50"},{"values":[2,6,5,3,2,9,2,4,5],"quality":"Less is better","isMain":false,"metricName":"L75"},{"values":["63.92","64.60","63.33","61.47","63.97","63.89","62.76","63.98","63.93"],"quality":"Equal","isMain":false,"metricName":"GC (%)"}]],["Predicted genes",[]],["Similarity statistics",[{"values":[0,0,0,0,0,0,0,0,0],"quality":"Equal","isMain":false,"metricName":"# similar correct contigs"},{"values":[0,0,0,0,0,0,0,0,0],"quality":"Equal","isMain":false,"metricName":"# similar misassembled blocks"}]],["Reference statistics",[{"values":[50968,50968,50968,50968,50968,50968,50968,50968,50968],"quality":"Equal","isMain":false,"metricName":"Reference length"},{"values":[1,1,1,1,1,1,1,1,1],"quality":"Equal","isMain":false,"metricName":"Reference fragments"},{"values":["63.71","63.71","63.71","63.71","63.71","63.71","63.71","63.71","63.71"],"quality":"Equal","isMain":false,"metricName":"Reference GC (%)"}]]],"referenceName":"gi\_192823901\_ref\_NC","date":"18 July 2018, Wednesday, 16:40:20","order":[0,1,2,3,4,5,6,7,8],"assembliesNames":["CLC","IDBA\_UD","MEGAHIT","MIRA","SOAPdenovo2","SPAdes","SPAdes\_meta","SPAdes\_sc","SPAdes\_sc\_careful"]},{"assembliesWithNs":null,"minContig":500,"report":[["Genome statistics",[{"values":["5.578","2.660","4.355","3.963","22.723","1.486","6.399","2.559","3.750","3.750"],"quality":"More is better","isMain":true,"metricName":"Genome fraction (%)"},{"values":["1.004","1.000","1.026","1.000","1.106","1.000","1.000","1.000","1.000","1.000"],"quality":"Less is better","isMain":true,"metricName":"Duplication ratio"},{"values":[1227,795,1371,738,3603,796,1371,1371,1371,1371],"quality":"More is better","isMain":true,"metricName":"Largest alignment"},{"values":[3000,1425,2333,2123,13167,796,3428,1371,2009,2009],"quality":"More is better","isMain":true,"metricName":"Total aligned length"},{"values":[692,795,1371,660,1383,796,838,1371,1371,1371],"quality":"More is better","isMain":false,"metricName":"NA50"},{"values":[574,630,504,null,996,796,660,1371,638,638],"quality":"More is better","isMain":false,"metricName":"NA75"},{"values":[null,null,null,null,null,null,null,null,null,null],"quality":"More is better","isMain":true,"metricName":"NGA50"},{"values":[2,1,1,3,4,1,2,1,1,1],"quality":"Less is better","isMain":false,"metricName":"LA50"},{"values":[3,2,2,null,7,1,3,1,2,2],"quality":"Less is better","isMain":false,"metricName":"LA75"}]],["Misassemblies",[{"values":[0,0,0,0,0,0,0,0,0,0],"quality":"Less is better","isMain":true,"metricName":"# misassemblies"},{"values":[0,0,0,0,0,0,0,0,0,0],"quality":"Less is better","isMain":false,"metricName":" # relocations"},{"values":[0,0,0,0,0,0,0,0,0,0],"quality":"Less is better","isMain":false,"metricName":" # translocations"},{"values":[0,0,0,0,0,0,0,0,0,0],"quality":"Less is better","isMain":false,"metricName":" # inversions"},{"values":[0,0,0,0,0,0,0,0,0,0],"quality":"Less is better","isMain":false,"metricName":"# misassembled contigs"},{"values":[0,0,0,0,0,0,0,0,0,0],"quality":"Less is better","isMain":true,"metricName":"Misassembled contigs length"},{"values":[0,0,0,0,4,0,0,0,0,0],"quality":"Less is better","isMain":false,"metricName":"# local misassemblies"},{"values":[0,0,0,0,0,0,0,0,0,0],"quality":"Less is better","isMain":false,"metricName":"# unaligned mis. contigs"}]],["Unaligned",[{"values":[0,0,0,0,0,0,0,0,0,0],"quality":"Less is better","isMain":false,"metricName":"# fully unaligned contigs"},{"values":[0,0,0,0,0,0,0,0,0,0],"quality":"Less is better","isMain":false,"metricName":"Fully unaligned length"},{"values":[0,0,0,1,1,0,0,0,0,0],"quality":"Less is better","isMain":false,"metricName":"# partially unaligned contigs"},{"values":[0,0,0,1636,1362,0,0,0,0,0],"quality":"Less is better","isMain":false,"metricName":"Partially unaligned length"}]],["Mismatches",[{"values":[32,1,0,5,165,0,5,0,20,20],"quality":"Less is better","isMain":false,"metricName":"# mismatches"},{"values":[1,0,0,0,11,0,0,0,0,0],"quality":"Less is better","isMain":false,"metricName":"# indels"},{"values":[12,0,0,0,12,0,0,0,0,0],"quality":"Less is better","isMain":false,"metricName":"Indels length"},{"values":["1070.95","70.18","0.00","235.52","1355.46","0.00","145.86","0.00","995.52","995.52"],"quality":"Less is better","isMain":true,"metricName":"# mismatches per 100 kbp"},{"values":["33.47","0.00","0.00","0.00","90.36","0.00","0.00","0.00","0.00","0.00"],"quality":"Less is better","isMain":true,"metricName":"# indels per 100 kbp"},{"values":[0,0,0,0,11,0,0,0,0,0],"quality":"Less is better","isMain":false,"metricName":" # indels (<= 5 bp)"},{"values":[1,0,0,0,0,0,0,0,0,0],"quality":"Less is better","isMain":false,"metricName":" # indels (> 5 bp)"},{"values":[0,0,0,0,20,0,0,0,0,0],"quality":"Less is better","isMain":false,"metricName":"# N's"},{"values":["0.00","0.00","0.00","0.00","134.95","0.00","0.00","0.00","0.00","0.00"],"quality":"Less is better","isMain":true,"metricName":"# N's per 100 kbp"}]],["Statistics without reference",[{"values":[4,2,3,3,9,1,4,1,2,2],"quality":"Equal","isMain":true,"metricName":"# contigs"},{"values":[1,0,1,1,7,0,1,1,1,1],"quality":"Equal","isMain":false,"metricName":"# contigs (>= 1000 bp)"},{"values":[0,0,0,0,0,0,0,0,0,0],"quality":"Equal","isMain":false,"metricName":"# contigs (>= 5000 bp)"},{"values":[0,0,0,0,0,0,0,0,0,0],"quality":"Equal","isMain":false,"metricName":"# contigs (>= 10000 bp)"},{"values":[0,0,0,0,0,0,0,0,0,0],"quality":"Equal","isMain":false,"metricName":"# contigs (>= 25000 bp)"},{"values":[0,0,0,0,0,0,0,0,0,0],"quality":"Equal","isMain":false,"metricName":"# contigs (>= 50000 bp)"},{"values":[1227,795,1371,2374,4965,796,1371,1371,1371,1371],"quality":"More is better","isMain":true,"metricName":"Largest contig"},{"values":[3000,1425,2393,3759,14820,796,3428,1371,2009,2009],"quality":"More is better","isMain":true,"metricName":"Total length"},{"values":[1227,0,1371,2374,12917,0,1371,1371,1371,1371],"quality":"More is better","isMain":true,"metricName":"Total length (>= 1000 bp)"},{"values":[0,0,0,0,0,0,0,0,0,0],"quality":"More is better","isMain":false,"metricName":"Total length (>= 5000 bp)"},{"values":[0,0,0,0,0,0,0,0,0,0],"quality":"More is better","isMain":true,"metricName":"Total length (>= 10000 bp)"},{"values":[0,0,0,0,0,0,0,0,0,0],"quality":"More is better","isMain":false,"metricName":"Total length (>= 25000 bp)"},{"values":[0,0,0,0,0,0,0,0,0,0],"quality":"More is better","isMain":true,"metricName":"Total length (>= 50000 bp)"},{"values":[692,795,1371,2374,1423,796,838,1371,1371,1371],"quality":"More is better","isMain":false,"metricName":"N50"},{"values":[574,630,518,725,1210,796,660,1371,638,638],"quality":"More is better","isMain":false,"metricName":"N75"},{"values":[2,1,1,1,3,1,2,1,1,1],"quality":"Less is better","isMain":false,"metricName":"L50"},{"values":[3,2,2,2,6,1,3,1,2,2],"quality":"Less is better","isMain":false,"metricName":"L75"},{"values":["61.20","61.40","63.23","62.60","63.89","61.43","62.28","61.56","61.08","61.08"],"quality":"Equal","isMain":false,"metricName":"GC (%)"}]],["Predicted genes",[]],["Similarity statistics",[{"values":[0,0,0,0,0,0,0,0,0,0],"quality":"Equal","isMain":false,"metricName":"# similar correct contigs"},{"values":[0,0,0,0,0,0,0,0,0,0],"quality":"Equal","isMain":false,"metricName":"# similar misassembled blocks"}]],["Reference statistics",[{"values":[53572,53572,53572,53572,53572,53572,53572,53572,53572,53572],"quality":"Equal","isMain":false,"metricName":"Reference length"},{"values":[1,1,1,1,1,1,1,1,1,1],"quality":"Equal","isMain":false,"metricName":"Reference fragments"},{"values":["63.57","63.57","63.57","63.57","63.57","63.57","63.57","63.57","63.57","63.57"],"quality":"Equal","isMain":false,"metricName":"Reference GC (%)"}]]],"referenceName":"gi\_192823996\_ref\_NC","date":"18 July 2018, Wednesday, 16:40:32","order":[0,1,2,3,4,5,6,7,8,9],"assembliesNames":["ABySS\_127","CLC","IDBA\_UD","MEGAHIT","MIRA","SOAPdenovo2","SPAdes","SPAdes\_meta","SPAdes\_sc","SPAdes\_sc\_careful"]},{"assembliesWithNs":null,"minContig":500,"report":[["Genome statistics",[{"values":["100.000","100.000","100.000","100.000","100.000","100.000","100.000","100.000","100.000","100.000","100.000","70.264"],"quality":"More is better","isMain":true,"metricName":"Genome fraction (%)"},{"values":["1.002","1.001","1.021","1.002","1.003","1.009","1.009","1.002","1.001","1.001","1.001","1.004"],"quality":"Less is better","isMain":true,"metricName":"Duplication ratio"},{"values":[53018,52954,52892,52991,53033,53344,53381,53019,52947,52947,52947,6286],"quality":"More is better","isMain":true,"metricName":"Largest alignment"},{"values":[53018,52954,54003,52991,53033,53344,53381,53019,52947,52947,52947,37309],"quality":"More is better","isMain":true,"metricName":"Total aligned length"},{"values":[53018,52954,52892,52991,53033,53344,53381,53019,52947,52947,52947,1153],"quality":"More is better","isMain":false,"metricName":"NG50"},{"values":[53018,52954,52892,52991,53033,53344,53381,53019,52947,52947,52947,null],"quality":"More is better","isMain":false,"metricName":"NG75"},{"values":[53018,52954,52892,52991,53033,53344,53381,53019,52947,52947,52947,2049],"quality":"More is better","isMain":false,"metricName":"NA50"},{"values":[53018,52954,52892,52991,53033,53344,53381,53019,52947,52947,52947,1010],"quality":"More is better","isMain":false,"metricName":"NA75"},{"values":[53018,52954,52892,52991,53033,53344,53381,53019,52947,52947,52947,1153],"quality":"More is better","isMain":true,"metricName":"NGA50"},{"values":[53018,52954,52892,52991,53033,53344,53381,53019,52947,52947,52947,null],"quality":"More is better","isMain":false,"metricName":"NGA75"},{"values":[1,1,1,1,1,1,1,1,1,1,1,11],"quality":"Less is better","isMain":false,"metricName":"LG50"},{"values":[1,1,1,1,1,1,1,1,1,1,1,null],"quality":"Less is better","isMain":false,"metricName":"LG75"},{"values":[1,1,1,1,1,1,1,1,1,1,1,6],"quality":"Less is better","isMain":false,"metricName":"LA50"},{"values":[1,1,1,1,1,1,1,1,1,1,1,13],"quality":"Less is better","isMain":false,"metricName":"LA75"},{"values":[1,1,1,1,1,1,1,1,1,1,1,11],"quality":"Less is better","isMain":true,"metricName":"LGA50"},{"values":[1,1,1,1,1,1,1,1,1,1,1,null],"quality":"Less is better","isMain":false,"metricName":"LGA75"}]],["Misassemblies",[{"values":[0,0,0,0,0,0,0,0,0,0,0,0],"quality":"Less is better","isMain":true,"metricName":"# misassemblies"},{"values":[0,0,0,0,0,0,0,0,0,0,0,0],"quality":"Less is better","isMain":false,"metricName":" # relocations"},{"values":[0,0,0,0,0,0,0,0,0,0,0,0],"quality":"Less is better","isMain":false,"metricName":" # translocations"},{"values":[0,0,0,0,0,0,0,0,0,0,0,0],"quality":"Less is better","isMain":false,"metricName":" # inversions"},{"values":[0,0,0,0,0,0,0,0,0,0,0,0],"quality":"Less is better","isMain":false,"metricName":"# misassembled contigs"},{"values":[0,0,0,0,0,0,0,0,0,0,0,0],"quality":"Less is better","isMain":true,"metricName":"Misassembled contigs length"},{"values":[1,0,0,1,1,1,1,1,0,0,0,0],"quality":"Less is better","isMain":false,"metricName":"# local misassemblies"},{"values":[0,0,0,0,0,0,0,0,0,0,0,0],"quality":"Less is better","isMain":false,"metricName":"# unaligned mis. contigs"}]],["Unaligned",[{"values":[0,0,0,0,0,0,0,0,0,0,0,0],"quality":"Less is better","isMain":false,"metricName":"# fully unaligned contigs"},{"values":[0,0,0,0,0,0,0,0,0,0,0,0],"quality":"Less is better","isMain":false,"metricName":"Fully unaligned length"},{"values":[0,0,0,0,0,0,0,0,0,0,0,0],"quality":"Less is better","isMain":false,"metricName":"# partially unaligned contigs"},{"values":[0,0,0,0,0,0,0,0,0,0,0,0],"quality":"Less is better","isMain":false,"metricName":"Partially unaligned length"}]],["Mismatches",[{"values":[0,0,1,0,0,0,0,0,0,0,0,1],"quality":"Less is better","isMain":false,"metricName":"# mismatches"},{"values":[0,1,0,0,0,0,0,0,1,1,1,0],"quality":"Less is better","isMain":false,"metricName":"# indels"},{"values":[0,62,0,0,0,0,0,0,55,55,55,0],"quality":"Less is better","isMain":false,"metricName":"Indels length"},{"values":["0.00","0.00","1.89","0.00","0.00","0.00","0.00","0.00","0.00","0.00","0.00","2.69"],"quality":"Less is better","isMain":true,"metricName":"# mismatches per 100 kbp"},{"values":["0.00","1.89","0.00","0.00","0.00","0.00","0.00","0.00","1.89","1.89","1.89","0.00"],"quality":"Less is better","isMain":true,"metricName":"# indels per 100 kbp"},{"values":[0,0,0,0,0,0,0,0,0,0,0,0],"quality":"Less is better","isMain":false,"metricName":" # indels (<= 5 bp)"},{"values":[0,1,0,0,0,0,0,0,1,1,1,0],"quality":"Less is better","isMain":false,"metricName":" # indels (> 5 bp)"},{"values":[0,0,0,0,0,0,0,0,0,0,0,0],"quality":"Less is better","isMain":false,"metricName":"# N's"},{"values":["0.00","0.00","0.00","0.00","0.00","0.00","0.00","0.00","0.00","0.00","0.00","0.00"],"quality":"Less is better","isMain":true,"metricName":"# N's per 100 kbp"}]],["Statistics without reference",[{"values":[1,1,3,1,1,1,1,1,1,1,1,25],"quality":"Equal","isMain":true,"metricName":"# contigs"},{"values":[1,1,1,1,1,1,1,1,1,1,1,13],"quality":"Equal","isMain":false,"metricName":"# contigs (>= 1000 bp)"},{"values":[1,1,1,1,1,1,1,1,1,1,1,1],"quality":"Equal","isMain":false,"metricName":"# contigs (>= 5000 bp)"},{"values":[1,1,1,1,1,1,1,1,1,1,1,0],"quality":"Equal","isMain":false,"metricName":"# contigs (>= 10000 bp)"},{"values":[1,1,1,1,1,1,1,1,1,1,1,0],"quality":"Equal","isMain":false,"metricName":"# contigs (>= 25000 bp)"},{"values":[1,1,1,1,1,1,1,1,1,1,1,0],"quality":"Equal","isMain":false,"metricName":"# contigs (>= 50000 bp)"},{"values":[53018,52954,52892,52991,53033,53344,53381,53019,52947,52947,52947,6286],"quality":"More is better","isMain":true,"metricName":"Largest contig"},{"values":[53018,52954,54003,52991,53033,53344,53381,53019,52947,52947,52947,37309],"quality":"More is better","isMain":true,"metricName":"Total length"},{"values":[53018,52954,52892,52991,53033,53344,53381,53019,52947,52947,52947,28985],"quality":"More is better","isMain":true,"metricName":"Total length (>= 1000 bp)"},{"values":[53018,52954,52892,52991,53033,53344,53381,53019,52947,52947,52947,6286],"quality":"More is better","isMain":false,"metricName":"Total length (>= 5000 bp)"},{"values":[53018,52954,52892,52991,53033,53344,53381,53019,52947,52947,52947,0],"quality":"More is better","isMain":true,"metricName":"Total length (>= 10000 bp)"},{"values":[53018,52954,52892,52991,53033,53344,53381,53019,52947,52947,52947,0],"quality":"More is better","isMain":false,"metricName":"Total length (>= 25000 bp)"},{"values":[53018,52954,52892,52991,53033,53344,53381,53019,52947,52947,52947,0],"quality":"More is better","isMain":true,"metricName":"Total length (>= 50000 bp)"},{"values":[53018,52954,52892,52991,53033,53344,53381,53019,52947,52947,52947,2049],"quality":"More is better","isMain":false,"metricName":"N50"},{"values":[53018,52954,52892,52991,53033,53344,53381,53019,52947,52947,52947,1010],"quality":"More is better","isMain":false,"metricName":"N75"},{"values":[1,1,1,1,1,1,1,1,1,1,1,6],"quality":"Less is better","isMain":false,"metricName":"L50"},{"values":[1,1,1,1,1,1,1,1,1,1,1,13],"quality":"Less is better","isMain":false,"metricName":"L75"},{"values":["63.27","63.26","63.27","63.25","63.26","63.30","63.27","63.28","63.26","63.27","63.27","63.32"],"quality":"Equal","isMain":false,"metricName":"GC (%)"}]],["Predicted genes",[]],["Similarity statistics",[{"values":[0,0,0,0,0,0,0,0,0,0,0,0],"quality":"Equal","isMain":false,"metricName":"# similar correct contigs"},{"values":[0,0,0,0,0,0,0,0,0,0,0,0],"quality":"Equal","isMain":false,"metricName":"# similar misassembled blocks"}]],["Reference statistics",[{"values":[52892,52892,52892,52892,52892,52892,52892,52892,52892,52892,52892,52892],"quality":"Equal","isMain":false,"metricName":"Reference length"},{"values":[1,1,1,1,1,1,1,1,1,1,1,1],"quality":"Equal","isMain":false,"metricName":"Reference fragments"},{"values":["63.26","63.26","63.26","63.26","63.26","63.26","63.26","63.26","63.26","63.26","63.26","63.26"],"quality":"Equal","isMain":false,"metricName":"Reference GC (%)"}]]],"referenceName":"gi\_192824177\_ref\_NC","date":"18 July 2018, Wednesday, 16:40:46","order":[0,1,2,3,4,5,6,7,8,9,10,11],"assembliesNames":["ABySS\_127","ABySS\_63","CLC","IDBA\_UD","MEGAHIT","MIRA","Ray\_Meta","SPAdes","SPAdes\_meta","SPAdes\_sc","SPAdes\_sc\_careful","Velvet"]},{"assembliesWithNs":null,"minContig":500,"report":[["Genome statistics",[{"values":["56.851","73.938","70.603","100.000","97.435","78.354","100.000","83.380","95.935","50.886","100.000"],"quality":"More is better","isMain":true,"metricName":"Genome fraction (%)"},{"values":["1.004","1.011","1.028","4.349","1.035","1.028","1.006","1.015","1.013","1.002","1.353"],"quality":"Less is better","isMain":true,"metricName":"Duplication ratio"},{"values":[8067,6985,7702,28201,35339,10631,29468,7953,12354,3732,32819],"quality":"More is better","isMain":true,"metricName":"Largest alignment"},{"values":[20012,26202,25455,152378,35339,28229,35259,29654,34055,17868,47430],"quality":"More is better","isMain":true,"metricName":"Total aligned length"},{"values":[1953,3870,2683,34722,36232,8097,29468,3782,7702,509,49412],"quality":"More is better","isMain":false,"metricName":"NG50"},{"values":[null,null,null,34722,36232,1332,29468,1694,2929,null,49412],"quality":"More is better","isMain":false,"metricName":"NG75"},{"values":[3986,6917,5583,2609,35339,8097,29468,7702,7702,1289,32819],"quality":"More is better","isMain":false,"metricName":"NA50"},{"values":[3889,2430,2683,1656,35339,4196,29468,2929,2929,838,8200],"quality":"More is better","isMain":false,"metricName":"NA75"},{"values":[1953,3870,2683,28201,35339,8097,29468,3782,7702,509,32819],"quality":"More is better","isMain":true,"metricName":"NGA50"},{"values":[null,null,null,28201,35339,1169,29468,1694,2929,null,32819],"quality":"More is better","isMain":false,"metricName":"NGA75"},{"values":[4,3,4,1,1,2,1,3,2,14,1],"quality":"Less is better","isMain":false,"metricName":"LG50"},{"values":[null,null,null,1,1,5,1,6,4,null,1],"quality":"Less is better","isMain":false,"metricName":"LG75"},{"values":[2,2,2,15,1,2,1,2,2,4,1],"quality":"Less is better","isMain":false,"metricName":"LA50"},{"values":[3,4,4,34,1,3,1,4,4,8,2],"quality":"Less is better","isMain":false,"metricName":"LA75"},{"values":[4,3,4,1,1,2,1,3,2,14,1],"quality":"Less is better","isMain":true,"metricName":"LGA50"},{"values":[null,null,null,1,1,6,1,6,4,null,1],"quality":"Less is better","isMain":false,"metricName":"LGA75"}]],["Misassemblies",[{"values":[0,0,0,1,0,0,0,0,0,0,2],"quality":"Less is better","isMain":true,"metricName":"# misassemblies"},{"values":[0,0,0,1,0,0,0,0,0,0,2],"quality":"Less is better","isMain":false,"metricName":" # relocations"},{"values":[0,0,0,0,0,0,0,0,0,0,0],"quality":"Less is better","isMain":false,"metricName":" # translocations"},{"values":[0,0,0,0,0,0,0,0,0,0,0],"quality":"Less is better","isMain":false,"metricName":" # inversions"},{"values":[0,0,0,1,0,0,0,0,0,0,1],"quality":"Less is better","isMain":false,"metricName":"# misassembled contigs"},{"values":[0,0,0,34722,0,0,0,0,0,0,49412],"quality":"Less is better","isMain":true,"metricName":"Misassembled contigs length"},{"values":[0,0,0,2,1,0,0,0,0,0,2],"quality":"Less is better","isMain":false,"metricName":"# local misassemblies"},{"values":[0,0,0,0,0,0,0,0,0,0,0],"quality":"Less is better","isMain":false,"metricName":"# unaligned mis. contigs"}]],["Unaligned",[{"values":[0,0,0,0,0,0,0,0,0,0,0],"quality":"Less is better","isMain":false,"metricName":"# fully unaligned contigs"},{"values":[0,0,0,0,0,0,0,0,0,0,0],"quality":"Less is better","isMain":false,"metricName":"Fully unaligned length"},{"values":[0,0,0,1,1,1,0,0,0,0,1],"quality":"Less is better","isMain":false,"metricName":"# partially unaligned contigs"},{"values":[0,0,0,1982,893,697,0,0,0,0,1982],"quality":"Less is better","isMain":false,"metricName":"Partially unaligned length"}]],["Mismatches",[{"values":[3,0,7,57,37,0,5,0,0,10,5],"quality":"Less is better","isMain":false,"metricName":"# mismatches"},{"values":[0,0,0,1,0,0,0,0,0,0,1],"quality":"Less is better","isMain":false,"metricName":"# indels"},{"values":[0,0,0,1,0,0,0,0,0,0,1],"quality":"Less is better","isMain":false,"metricName":"Indels length"},{"values":["15.05","0.00","28.28","162.60","108.33","0.00","14.26","0.00","0.00","56.06","14.26"],"quality":"Less is better","isMain":true,"metricName":"# mismatches per 100 kbp"},{"values":["0.00","0.00","0.00","2.85","0.00","0.00","0.00","0.00","0.00","0.00","2.85"],"quality":"Less is better","isMain":true,"metricName":"# indels per 100 kbp"},{"values":[0,0,0,1,0,0,0,0,0,0,1],"quality":"Less is better","isMain":false,"metricName":" # indels (<= 5 bp)"},{"values":[0,0,0,0,0,0,0,0,0,0,0],"quality":"Less is better","isMain":false,"metricName":" # indels (> 5 bp)"},{"values":[0,0,0,114,0,697,0,0,0,0,121],"quality":"Less is better","isMain":false,"metricName":"# N's"},{"values":["0.00","0.00","0.00","73.82","0.00","2409.60","0.00","0.00","0.00","0.00","244.88"],"quality":"Less is better","isMain":true,"metricName":"# N's per 100 kbp"}]],["Statistics without reference",[{"values":[7,10,8,62,1,7,4,9,9,14,1],"quality":"Equal","isMain":true,"metricName":"# contigs"},{"values":[4,7,7,58,1,6,4,8,8,6,1],"quality":"Equal","isMain":false,"metricName":"# contigs (>= 1000 bp)"},{"values":[1,2,2,3,1,2,1,2,2,0,1],"quality":"Equal","isMain":false,"metricName":"# contigs (>= 5000 bp)"},{"values":[0,0,0,1,1,1,1,0,1,0,1],"quality":"Equal","isMain":false,"metricName":"# contigs (>= 10000 bp)"},{"values":[0,0,0,1,1,0,1,0,0,0,1],"quality":"Equal","isMain":false,"metricName":"# contigs (>= 25000 bp)"},{"values":[0,0,0,0,0,0,0,0,0,0,0],"quality":"Equal","isMain":false,"metricName":"# contigs (>= 50000 bp)"},{"values":[8067,6985,7702,34722,36232,11328,29468,7953,12354,3732,49412],"quality":"More is better","isMain":true,"metricName":"Largest contig"},{"values":[20012,26202,25455,154425,36232,28926,35259,29654,34055,17868,49412],"quality":"More is better","isMain":true,"metricName":"Total length"},{"values":[17895,24257,24476,151006,36232,27975,35259,28847,33248,12212,49412],"quality":"More is better","isMain":true,"metricName":"Total length (>= 1000 bp)"},{"values":[8067,13902,13285,45253,36232,19425,29468,15655,20056,0,49412],"quality":"More is better","isMain":false,"metricName":"Total length (>= 5000 bp)"},{"values":[0,0,0,34722,36232,11328,29468,0,12354,0,49412],"quality":"More is better","isMain":true,"metricName":"Total length (>= 10000 bp)"},{"values":[0,0,0,34722,36232,0,29468,0,0,0,49412],"quality":"More is better","isMain":false,"metricName":"Total length (>= 25000 bp)"},{"values":[0,0,0,0,0,0,0,0,0,0,0],"quality":"More is better","isMain":true,"metricName":"Total length (>= 50000 bp)"},{"values":[3986,6917,5583,2635,36232,8097,29468,7702,7702,1289,49412],"quality":"More is better","isMain":false,"metricName":"N50"},{"values":[3889,2430,2683,1670,36232,4196,29468,2929,2929,838,49412],"quality":"More is better","isMain":false,"metricName":"N75"},{"values":[2,2,2,13,1,2,1,2,2,4,1],"quality":"Less is better","isMain":false,"metricName":"L50"},{"values":[3,4,4,32,1,3,1,4,4,8,1],"quality":"Less is better","isMain":false,"metricName":"L75"},{"values":["46.64","45.00","45.24","45.50","44.88","45.46","44.90","45.26","44.96","46.22","44.96"],"quality":"Equal","isMain":false,"metricName":"GC (%)"}]],["Predicted genes",[]],["Similarity statistics",[{"values":[0,0,0,0,0,0,0,0,0,0,0],"quality":"Equal","isMain":false,"metricName":"# similar correct contigs"},{"values":[0,0,0,0,0,0,0,0,0,0,0],"quality":"Equal","isMain":false,"metricName":"# similar misassembled blocks"}]],["Reference statistics",[{"values":[35055,35055,35055,35055,35055,35055,35055,35055,35055,35055,35055],"quality":"Equal","isMain":false,"metricName":"Reference length"},{"values":[1,1,1,1,1,1,1,1,1,1,1],"quality":"Equal","isMain":false,"metricName":"Reference fragments"},{"values":["44.85","44.85","44.85","44.85","44.85","44.85","44.85","44.85","44.85","44.85","44.85"],"quality":"Equal","isMain":false,"metricName":"Reference GC (%)"}]]],"referenceName":"gi\_193506277\_ref\_NC","date":"18 July 2018, Wednesday, 16:40:59","order":[0,1,2,3,4,5,6,7,8,9,10],"assembliesNames":["CLC","IDBA\_UD","MEGAHIT","MIRA","Ray\_Meta","SPAdes","SPAdes\_meta","SPAdes\_sc","SPAdes\_sc\_careful","Velvet","Geneious"]},{"assembliesWithNs":null,"minContig":500,"report":[["Genome statistics",[{"values":["99.502","100.000","100.000","100.000","100.000","100.000","100.000","100.000"],"quality":"More is better","isMain":true,"metricName":"Genome fraction (%)"},{"values":["1.005","131.363","1.995","1.029","1.012","1.012","1.012","1.489"],"quality":"Less is better","isMain":true,"metricName":"Duplication ratio"},{"values":[4399,1643,5080,4548,4476,4476,4476,4099],"quality":"More is better","isMain":true,"metricName":"Largest alignment"},{"values":[4399,579760,8818,4548,4476,4476,4476,6582],"quality":"More is better","isMain":true,"metricName":"Total aligned length"},{"values":[4421,1577,8818,4548,4476,4476,4476,6582],"quality":"More is better","isMain":false,"metricName":"NG50"},{"values":[4421,1522,8818,4548,4476,4476,4476,6582],"quality":"More is better","isMain":false,"metricName":"NG75"},{"values":[4399,760,5080,4548,4476,4476,4476,4099],"quality":"More is better","isMain":false,"metricName":"NA50"},{"values":[4399,587,3738,4548,4476,4476,4476,2483],"quality":"More is better","isMain":false,"metricName":"NA75"},{"values":[4399,1577,5080,4548,4476,4476,4476,4099],"quality":"More is better","isMain":true,"metricName":"NGA50"},{"values":[4399,1522,5080,4548,4476,4476,4476,4099],"quality":"More is better","isMain":false,"metricName":"NGA75"},{"values":[1,2,1,1,1,1,1,1],"quality":"Less is better","isMain":false,"metricName":"LG50"},{"values":[1,3,1,1,1,1,1,1],"quality":"Less is better","isMain":false,"metricName":"LG75"},{"values":[1,312,1,1,1,1,1,1],"quality":"Less is better","isMain":false,"metricName":"LA50"},{"values":[1,528,2,1,1,1,1,2],"quality":"Less is better","isMain":false,"metricName":"LA75"},{"values":[1,2,1,1,1,1,1,1],"quality":"Less is better","isMain":true,"metricName":"LGA50"},{"values":[1,3,1,1,1,1,1,1],"quality":"Less is better","isMain":false,"metricName":"LGA75"}]],["Misassemblies",[{"values":[0,0,1,0,0,0,0,1],"quality":"Less is better","isMain":true,"metricName":"# misassemblies"},{"values":[0,0,1,0,0,0,0,1],"quality":"Less is better","isMain":false,"metricName":" # relocations"},{"values":[0,0,0,0,0,0,0,0],"quality":"Less is better","isMain":false,"metricName":" # translocations"},{"values":[0,0,0,0,0,0,0,0],"quality":"Less is better","isMain":false,"metricName":" # inversions"},{"values":[0,0,1,0,0,0,0,1],"quality":"Less is better","isMain":false,"metricName":"# misassembled contigs"},{"values":[0,0,8818,0,0,0,0,6582],"quality":"Less is better","isMain":true,"metricName":"Misassembled contigs length"},{"values":[0,0,1,1,0,0,0,0],"quality":"Less is better","isMain":false,"metricName":"# local misassemblies"},{"values":[0,0,0,0,0,0,0,0],"quality":"Less is better","isMain":false,"metricName":"# unaligned mis. contigs"}]],["Unaligned",[{"values":[0,0,0,0,0,0,0,0],"quality":"Less is better","isMain":false,"metricName":"# fully unaligned contigs"},{"values":[0,0,0,0,0,0,0,0],"quality":"Less is better","isMain":false,"metricName":"Fully unaligned length"},{"values":[0,0,0,0,0,0,0,0],"quality":"Less is better","isMain":false,"metricName":"# partially unaligned contigs"},{"values":[0,0,0,0,0,0,0,0],"quality":"Less is better","isMain":false,"metricName":"Partially unaligned length"}]],["Mismatches",[{"values":[0,10,0,0,0,0,0,0],"quality":"Less is better","isMain":false,"metricName":"# mismatches"},{"values":[0,0,0,0,1,1,1,0],"quality":"Less is better","isMain":false,"metricName":"# indels"},{"values":[0,0,0,0,55,55,55,0],"quality":"Less is better","isMain":false,"metricName":"Indels length"},{"values":["0.00","226.19","0.00","0.00","0.00","0.00","0.00","0.00"],"quality":"Less is better","isMain":true,"metricName":"# mismatches per 100 kbp"},{"values":["0.00","0.00","0.00","0.00","22.62","22.62","22.62","0.00"],"quality":"Less is better","isMain":true,"metricName":"# indels per 100 kbp"},{"values":[0,0,0,0,0,0,0,0],"quality":"Less is better","isMain":false,"metricName":" # indels (<= 5 bp)"},{"values":[0,0,0,0,1,1,1,0],"quality":"Less is better","isMain":false,"metricName":" # indels (> 5 bp)"},{"values":[0,21,0,0,0,0,0,0],"quality":"Less is better","isMain":false,"metricName":"# N's"},{"values":["0.00","3.62","0.00","0.00","0.00","0.00","0.00","0.00"],"quality":"Less is better","isMain":true,"metricName":"# N's per 100 kbp"}]],["Statistics without reference",[{"values":[1,786,1,1,1,1,1,1],"quality":"Equal","isMain":true,"metricName":"# contigs"},{"values":[1,75,1,1,1,1,1,1],"quality":"Equal","isMain":false,"metricName":"# contigs (>= 1000 bp)"},{"values":[0,0,1,0,0,0,0,1],"quality":"Equal","isMain":false,"metricName":"# contigs (>= 5000 bp)"},{"values":[0,0,0,0,0,0,0,0],"quality":"Equal","isMain":false,"metricName":"# contigs (>= 10000 bp)"},{"values":[0,0,0,0,0,0,0,0],"quality":"Equal","isMain":false,"metricName":"# contigs (>= 25000 bp)"},{"values":[0,0,0,0,0,0,0,0],"quality":"Equal","isMain":false,"metricName":"# contigs (>= 50000 bp)"},{"values":[4421,1643,8818,4548,4476,4476,4476,6582],"quality":"More is better","isMain":true,"metricName":"Largest contig"},{"values":[4421,580757,8818,4548,4476,4476,4476,6582],"quality":"More is better","isMain":true,"metricName":"Total length"},{"values":[4421,88904,8818,4548,4476,4476,4476,6582],"quality":"More is better","isMain":true,"metricName":"Total length (>= 1000 bp)"},{"values":[0,0,8818,0,0,0,0,6582],"quality":"More is better","isMain":false,"metricName":"Total length (>= 5000 bp)"},{"values":[0,0,0,0,0,0,0,0],"quality":"More is better","isMain":true,"metricName":"Total length (>= 10000 bp)"},{"values":[0,0,0,0,0,0,0,0],"quality":"More is better","isMain":false,"metricName":"Total length (>= 25000 bp)"},{"values":[0,0,0,0,0,0,0,0],"quality":"More is better","isMain":true,"metricName":"Total length (>= 50000 bp)"},{"values":[4421,761,8818,4548,4476,4476,4476,6582],"quality":"More is better","isMain":false,"metricName":"N50"},{"values":[4421,587,8818,4548,4476,4476,4476,6582],"quality":"More is better","isMain":false,"metricName":"N75"},{"values":[1,312,1,1,1,1,1,1],"quality":"Less is better","isMain":false,"metricName":"L50"},{"values":[1,527,1,1,1,1,1,1],"quality":"Less is better","isMain":false,"metricName":"L75"},{"values":["32.03","32.29","32.00","32.12","32.10","32.10","32.04","31.75"],"quality":"Equal","isMain":false,"metricName":"GC (%)"}]],["Predicted genes",[]],["Similarity statistics",[{"values":[0,0,0,0,0,0,0,0],"quality":"Equal","isMain":false,"metricName":"# similar correct contigs"},{"values":[0,0,0,0,0,0,0,0],"quality":"Equal","isMain":false,"metricName":"# similar misassembled blocks"}]],["Reference statistics",[{"values":[4421,4421,4421,4421,4421,4421,4421,4421],"quality":"Equal","isMain":false,"metricName":"Reference length"},{"values":[1,1,1,1,1,1,1,1],"quality":"Equal","isMain":false,"metricName":"Reference fragments"},{"values":["32.05","32.05","32.05","32.05","32.05","32.05","32.05","32.05"],"quality":"Equal","isMain":false,"metricName":"Reference GC (%)"}]]],"referenceName":"gi\_19387568\_ref\_NC","date":"18 July 2018, Wednesday, 16:41:11","order":[0,1,2,3,4,5,6,7],"assembliesNames":["CLC","MIRA","Ray\_Meta","SPAdes","SPAdes\_meta","SPAdes\_sc","SPAdes\_sc\_careful","Geneious"]},{"assembliesWithNs":null,"minContig":500,"report":[["Genome statistics",[{"values":["92.554","31.817","97.458","78.795","88.944","99.297","81.822","94.042","63.098","51.855","69.585","4.586","99.297"],"quality":"More is better","isMain":true,"metricName":"Genome fraction (%)"},{"values":["1.081","1.000","1.020","1.022","1.037","1.199","1.068","1.000","1.007","1.003","1.000","1.000","1.181"],"quality":"Less is better","isMain":true,"metricName":"Duplication ratio"},{"values":[8080,2030,12077,2817,3012,18037,5921,23655,13927,3079,8970,844,18037],"quality":"More is better","isMain":true,"metricName":"Largest alignment"},{"values":[29322,9324,29140,23610,27037,34892,25613,27559,18604,15245,20392,1344,34356],"quality":"More is better","isMain":true,"metricName":"Total aligned length"},{"values":[8910,null,6819,1363,2090,88168,2955,23655,6248,755,3550,null,106226],"quality":"More is better","isMain":false,"metricName":"NG50"},{"values":[8080,null,5424,677,1181,88168,1508,23655,973,null,null,null,106226],"quality":"More is better","isMain":false,"metricName":"NG75"},{"values":[5382,930,6819,1672,2186,null,4301,23655,13927,1897,3886,844,null],"quality":"More is better","isMain":false,"metricName":"NA50"},{"values":[2278,637,5424,921,1198,null,2219,23655,1119,1300,3550,500,null],"quality":"More is better","isMain":false,"metricName":"NA75"},{"values":[7001,null,6819,1363,2090,18037,2955,23655,2585,755,3550,null,18037],"quality":"More is better","isMain":true,"metricName":"NGA50"},{"values":[5061,null,5424,677,1181,13673,1508,23655,null,null,null,null,13673],"quality":"More is better","isMain":false,"metricName":"NGA75"},{"values":[2,null,2,8,6,1,4,1,2,8,3,null,1],"quality":"Less is better","isMain":false,"metricName":"LG50"},{"values":[3,null,3,17,11,1,7,1,4,null,null,null,1],"quality":"Less is better","isMain":false,"metricName":"LG75"},{"values":[3,4,2,6,5,null,3,1,1,3,2,1,null],"quality":"Less is better","isMain":false,"metricName":"LA50"},{"values":[5,7,3,11,10,null,5,1,3,6,3,2,null],"quality":"Less is better","isMain":false,"metricName":"LA75"},{"values":[2,null,2,8,6,1,4,1,2,8,3,null,1],"quality":"Less is better","isMain":true,"metricName":"LGA50"},{"values":[4,null,3,17,11,2,7,1,null,null,null,null,2],"quality":"Less is better","isMain":false,"metricName":"LGA75"}]],["Misassemblies",[{"values":[0,0,0,0,0,0,0,0,0,0,0,0,0],"quality":"Less is better","isMain":true,"metricName":"# misassemblies"},{"values":[0,0,0,0,0,0,0,0,0,0,0,0,0],"quality":"Less is better","isMain":false,"metricName":" # relocations"},{"values":[0,0,0,0,0,0,0,0,0,0,0,0,0],"quality":"Less is better","isMain":false,"metricName":" # translocations"},{"values":[0,0,0,0,0,0,0,0,0,0,0,0,0],"quality":"Less is better","isMain":false,"metricName":" # inversions"},{"values":[0,0,0,0,0,0,0,0,0,0,0,0,0],"quality":"Less is better","isMain":false,"metricName":"# misassembled contigs"},{"values":[0,0,0,0,0,0,0,0,0,0,0,0,0],"quality":"Less is better","isMain":true,"metricName":"Misassembled contigs length"},{"values":[1,0,0,0,0,0,0,0,1,0,0,0,0],"quality":"Less is better","isMain":false,"metricName":"# local misassemblies"},{"values":[0,0,0,0,0,1,0,0,0,0,0,0,1],"quality":"Less is better","isMain":false,"metricName":"# unaligned mis. contigs"}]],["Unaligned",[{"values":[0,0,0,0,0,0,0,0,0,0,0,0,0],"quality":"Less is better","isMain":false,"metricName":"# fully unaligned contigs"},{"values":[0,0,0,0,0,0,0,0,0,0,0,0,0],"quality":"Less is better","isMain":false,"metricName":"Fully unaligned length"},{"values":[2,0,0,0,0,1,0,0,1,0,0,0,1],"quality":"Less is better","isMain":false,"metricName":"# partially unaligned contigs"},{"values":[6843,0,0,0,0,53812,0,0,3663,0,0,0,71870],"quality":"Less is better","isMain":false,"metricName":"Partially unaligned length"}]],["Mismatches",[{"values":[23,0,2,0,9,25,0,6,287,56,175,0,24],"quality":"Less is better","isMain":false,"metricName":"# mismatches"},{"values":[1,0,0,0,0,0,0,0,6,9,0,0,0],"quality":"Less is better","isMain":false,"metricName":"# indels"},{"values":[1,0,0,0,0,0,0,0,11,30,0,0,0],"quality":"Less is better","isMain":false,"metricName":"Indels length"},{"values":["84.80","0.00","7.00","0.00","34.53","85.91","0.00","21.77","1552.11","368.52","858.18","0.00","82.48"],"quality":"Less is better","isMain":true,"metricName":"# mismatches per 100 kbp"},{"values":["3.69","0.00","0.00","0.00","0.00","0.00","0.00","0.00","32.45","59.23","0.00","0.00","0.00"],"quality":"Less is better","isMain":true,"metricName":"# indels per 100 kbp"},{"values":[1,0,0,0,0,0,0,0,6,7,0,0,0],"quality":"Less is better","isMain":false,"metricName":" # indels (<= 5 bp)"},{"values":[0,0,0,0,0,0,0,0,0,2,0,0,0],"quality":"Less is better","isMain":false,"metricName":" # indels (> 5 bp)"},{"values":[110,76,0,0,0,1,0,0,0,0,0,0,50],"quality":"Less is better","isMain":false,"metricName":"# N's"},{"values":["304.16","815.10","0.00","0.00","0.00","1.13","0.00","0.00","0.00","0.00","0.00","0.00","47.07"],"quality":"Less is better","isMain":true,"metricName":"# N's per 100 kbp"}]],["Statistics without reference",[{"values":[7,10,6,19,18,2,9,3,4,9,5,2,1],"quality":"Equal","isMain":true,"metricName":"# contigs"},{"values":[5,2,4,9,11,1,9,3,3,7,5,0,1],"quality":"Equal","isMain":false,"metricName":"# contigs (>= 1000 bp)"},{"values":[4,0,3,0,0,1,1,1,2,0,1,0,1],"quality":"Equal","isMain":false,"metricName":"# contigs (>= 5000 bp)"},{"values":[1,0,1,0,0,1,0,1,1,0,0,0,1],"quality":"Equal","isMain":false,"metricName":"# contigs (>= 10000 bp)"},{"values":[0,0,0,0,0,1,0,0,0,0,0,0,1],"quality":"Equal","isMain":false,"metricName":"# contigs (>= 25000 bp)"},{"values":[0,0,0,0,0,1,0,0,0,0,0,0,1],"quality":"Equal","isMain":false,"metricName":"# contigs (>= 50000 bp)"},{"values":[10316,2030,12077,2817,3012,88168,5921,23655,13936,3079,8970,844,106226],"quality":"More is better","isMain":true,"metricName":"Largest contig"},{"values":[36165,9324,29140,23610,27037,88704,25613,27559,22276,15249,20392,1344,106226],"quality":"More is better","isMain":true,"metricName":"Total length"},{"values":[34645,3525,27840,16178,22217,88168,25613,27559,21303,13939,20392,0,106226],"quality":"More is better","isMain":true,"metricName":"Total length (>= 1000 bp)"},{"values":[32367,0,24320,0,0,88168,5921,23655,20184,0,8970,0,106226],"quality":"More is better","isMain":false,"metricName":"Total length (>= 5000 bp)"},{"values":[10316,0,12077,0,0,88168,0,23655,13936,0,0,0,106226],"quality":"More is better","isMain":true,"metricName":"Total length (>= 10000 bp)"},{"values":[0,0,0,0,0,88168,0,0,0,0,0,0,106226],"quality":"More is better","isMain":false,"metricName":"Total length (>= 25000 bp)"},{"values":[0,0,0,0,0,88168,0,0,0,0,0,0,106226],"quality":"More is better","isMain":true,"metricName":"Total length (>= 50000 bp)"},{"values":[8910,930,6819,1672,2186,88168,4301,23655,13936,1897,3886,844,106226],"quality":"More is better","isMain":false,"metricName":"N50"},{"values":[8080,637,5424,921,1198,88168,2219,23655,6248,1302,3550,500,106226],"quality":"More is better","isMain":false,"metricName":"N75"},{"values":[2,4,2,6,5,1,3,1,1,3,2,1,1],"quality":"Less is better","isMain":false,"metricName":"L50"},{"values":[3,7,3,11,10,1,5,1,2,6,3,2,1],"quality":"Less is better","isMain":false,"metricName":"L75"},{"values":["35.10","34.14","34.68","34.80","34.77","34.51","34.63","34.67","34.93","33.96","34.22","38.91","34.35"],"quality":"Equal","isMain":false,"metricName":"GC (%)"}]],["Predicted genes",[]],["Similarity statistics",[{"values":[0,0,0,0,0,0,0,0,0,0,0,0,0],"quality":"Equal","isMain":false,"metricName":"# similar correct contigs"},{"values":[0,0,0,0,0,0,0,0,0,0,0,0,0],"quality":"Equal","isMain":false,"metricName":"# similar misassembled blocks"}]],["Reference statistics",[{"values":[29305,29305,29305,29305,29305,29305,29305,29305,29305,29305,29305,29305,29305],"quality":"Equal","isMain":false,"metricName":"Reference length"},{"values":[1,1,1,1,1,1,1,1,1,1,1,1,1],"quality":"Equal","isMain":false,"metricName":"Reference fragments"},{"values":["34.72","34.72","34.72","34.72","34.72","34.72","34.72","34.72","34.72","34.72","34.72","34.72","34.72"],"quality":"Equal","isMain":false,"metricName":"Reference GC (%)"}]]],"referenceName":"gi\_194097521\_ref\_NC","date":"18 July 2018, Wednesday, 16:41:25","order":[0,1,2,3,4,5,6,7,8,9,10,11,12],"assembliesNames":["ABySS\_127","ABySS\_63","CLC","IDBA\_UD","MEGAHIT","MIRA","Ray\_Meta","SPAdes","SPAdes\_meta","SPAdes\_sc","SPAdes\_sc\_careful","Velvet","Geneious"]},{"assembliesWithNs":null,"minContig":500,"report":[["Genome statistics",[{"values":["98.617","69.874","98.548","65.426","87.114","99.860","28.419","86.543","89.496","65.831","60.924","30.121","99.860"],"quality":"More is better","isMain":true,"metricName":"Genome fraction (%)"},{"values":["1.035","1.019","1.005","1.074","1.028","1.012","1.692","1.015","1.002","1.020","1.009","1.000","1.000"],"quality":"Less is better","isMain":true,"metricName":"Duplication ratio"},{"values":[6520,2679,12990,4158,9425,28839,2062,8672,13549,2421,2421,1966,39197],"quality":"More is better","isMain":true,"metricName":"Largest alignment"},{"values":[40081,27951,38884,27593,35136,39676,11267,34478,35185,26358,24120,11823,39197],"quality":"More is better","isMain":true,"metricName":"Total aligned length"},{"values":[2790,905,9078,1444,2844,28839,512,5934,11374,709,639,null,39197],"quality":"More is better","isMain":false,"metricName":"NG50"},{"values":[2250,null,6053,null,2067,9230,null,4093,8639,null,null,null,39197],"quality":"More is better","isMain":false,"metricName":"NG75"},{"values":[2790,1289,9078,2286,2983,28839,103,5934,11374,853,870,688,39197],"quality":"More is better","isMain":false,"metricName":"NA50"},{"values":[2099,878,6053,1429,2532,9230,null,4093,8639,709,677,545,39197],"quality":"More is better","isMain":false,"metricName":"NA75"},{"values":[2790,905,9078,1444,2844,28839,null,5934,11374,709,639,null,39197],"quality":"More is better","isMain":true,"metricName":"NGA50"},{"values":[2250,null,6053,null,2067,9230,null,4093,8639,null,null,null,39197],"quality":"More is better","isMain":false,"metricName":"NGA75"},{"values":[5,13,2,8,5,1,21,3,2,18,19,null,1],"quality":"Less is better","isMain":false,"metricName":"LG50"},{"values":[9,null,4,null,8,2,null,5,3,null,null,null,1],"quality":"Less is better","isMain":false,"metricName":"LG75"},{"values":[5,8,2,5,4,1,27,3,2,10,9,6,1],"quality":"Less is better","isMain":false,"metricName":"LA50"},{"values":[10,14,4,9,7,2,null,5,3,18,17,11,1],"quality":"Less is better","isMain":false,"metricName":"LA75"},{"values":[5,13,2,8,5,1,null,3,2,18,19,null,1],"quality":"Less is better","isMain":true,"metricName":"LGA50"},{"values":[9,null,4,null,8,2,null,5,3,null,null,null,1],"quality":"Less is better","isMain":false,"metricName":"LGA75"}]],["Misassemblies",[{"values":[0,0,0,0,0,0,0,0,0,0,0,0,0],"quality":"Less is better","isMain":true,"metricName":"# misassemblies"},{"values":[0,0,0,0,0,0,0,0,0,0,0,0,0],"quality":"Less is better","isMain":false,"metricName":" # relocations"},{"values":[0,0,0,0,0,0,0,0,0,0,0,0,0],"quality":"Less is better","isMain":false,"metricName":" # translocations"},{"values":[0,0,0,0,0,0,0,0,0,0,0,0,0],"quality":"Less is better","isMain":false,"metricName":" # inversions"},{"values":[0,0,0,0,0,0,0,0,0,0,0,0,0],"quality":"Less is better","isMain":false,"metricName":"# misassembled contigs"},{"values":[0,0,0,0,0,0,0,0,0,0,0,0,0],"quality":"Less is better","isMain":true,"metricName":"Misassembled contigs length"},{"values":[0,0,0,0,1,0,0,0,1,0,0,0,0],"quality":"Less is better","isMain":false,"metricName":"# local misassemblies"},{"values":[0,0,0,0,0,0,8,0,0,0,0,0,0],"quality":"Less is better","isMain":false,"metricName":"# unaligned mis. contigs"}]],["Unaligned",[{"values":[0,0,0,0,0,0,0,0,0,0,0,0,0],"quality":"Less is better","isMain":false,"metricName":"# fully unaligned contigs"},{"values":[0,0,0,0,0,0,0,0,0,0,0,0,0],"quality":"Less is better","isMain":false,"metricName":"Fully unaligned length"},{"values":[0,0,0,0,0,0,1,0,0,0,0,0,0],"quality":"Less is better","isMain":false,"metricName":"# partially unaligned contigs"},{"values":[0,0,0,0,0,0,977,0,0,0,0,0,0],"quality":"Less is better","isMain":false,"metricName":"Partially unaligned length"}]],["Mismatches",[{"values":[26,85,15,112,9,1,8,6,180,47,49,9,1],"quality":"Less is better","isMain":false,"metricName":"# mismatches"},{"values":[0,1,0,0,0,0,14,0,1,4,2,0,0],"quality":"Less is better","isMain":false,"metricName":"# indels"},{"values":[0,1,0,0,0,0,646,0,1,4,2,0,0],"quality":"Less is better","isMain":false,"metricName":"Indels length"},{"values":["67.17","309.91","38.78","436.12","26.32","2.55","71.72","17.66","512.40","181.89","204.90","76.12","2.55"],"quality":"Less is better","isMain":true,"metricName":"# mismatches per 100 kbp"},{"values":["0.00","3.65","0.00","0.00","0.00","0.00","125.50","0.00","2.85","15.48","8.36","0.00","0.00"],"quality":"Less is better","isMain":true,"metricName":"# indels per 100 kbp"},{"values":[0,1,0,0,0,0,1,0,1,4,2,0,0],"quality":"Less is better","isMain":false,"metricName":" # indels (<= 5 bp)"},{"values":[0,0,0,0,0,0,13,0,0,0,0,0,0],"quality":"Less is better","isMain":false,"metricName":" # indels (> 5 bp)"},{"values":[0,0,0,0,0,9,7106,0,0,0,0,0,16],"quality":"Less is better","isMain":false,"metricName":"# N's"},{"values":["0.00","0.00","0.00","0.00","0.00","22.68","35791.28","0.00","0.00","0.00","0.00","0.00","40.82"],"quality":"Less is better","isMain":true,"metricName":"# N's per 100 kbp"}]],["Statistics without reference",[{"values":[18,24,5,15,14,3,21,8,5,29,27,16,1],"quality":"Equal","isMain":true,"metricName":"# contigs"},{"values":[14,10,5,11,11,3,7,6,4,7,6,2,1],"quality":"Equal","isMain":false,"metricName":"# contigs (>= 1000 bp)"},{"values":[1,0,4,0,1,2,0,3,3,0,0,0,1],"quality":"Equal","isMain":false,"metricName":"# contigs (>= 5000 bp)"},{"values":[0,0,1,0,0,1,0,0,2,0,0,0,1],"quality":"Equal","isMain":false,"metricName":"# contigs (>= 10000 bp)"},{"values":[0,0,0,0,0,1,0,0,0,0,0,0,1],"quality":"Equal","isMain":false,"metricName":"# contigs (>= 25000 bp)"},{"values":[0,0,0,0,0,0,0,0,0,0,0,0,0],"quality":"Equal","isMain":false,"metricName":"# contigs (>= 50000 bp)"},{"values":[6520,2679,12990,4158,9425,28839,2326,8672,13549,2421,2421,1966,39197],"quality":"More is better","isMain":true,"metricName":"Largest contig"},{"values":[40081,27951,38884,27593,35136,39676,19854,34478,35185,26360,24122,11823,39197],"quality":"More is better","isMain":true,"metricName":"Total length"},{"values":[37573,17336,38884,24394,33413,39676,10655,33266,34669,11251,9567,3131,39197],"quality":"More is better","isMain":true,"metricName":"Total length (>= 1000 bp)"},{"values":[6520,0,34634,0,9425,38069,0,21007,33562,0,0,0,39197],"quality":"More is better","isMain":false,"metricName":"Total length (>= 5000 bp)"},{"values":[0,0,12990,0,0,28839,0,0,24923,0,0,0,39197],"quality":"More is better","isMain":true,"metricName":"Total length (>= 10000 bp)"},{"values":[0,0,0,0,0,28839,0,0,0,0,0,0,39197],"quality":"More is better","isMain":false,"metricName":"Total length (>= 25000 bp)"},{"values":[0,0,0,0,0,0,0,0,0,0,0,0,0],"quality":"More is better","isMain":true,"metricName":"Total length (>= 50000 bp)"},{"values":[2790,1289,9078,2286,2983,28839,1116,5934,11374,853,870,688,39197],"quality":"More is better","isMain":false,"metricName":"N50"},{"values":[2099,878,6053,1429,2532,9230,683,4093,8639,709,677,545,39197],"quality":"More is better","isMain":false,"metricName":"N75"},{"values":[5,8,2,5,4,1,7,3,2,10,9,6,1],"quality":"Less is better","isMain":false,"metricName":"L50"},{"values":[10,14,4,9,7,2,13,5,3,18,17,11,1],"quality":"Less is better","isMain":false,"metricName":"L75"},{"values":["49.88","49.97","49.96","49.37","49.98","49.96","49.07","49.78","50.04","49.59","49.52","49.04","49.95"],"quality":"Equal","isMain":false,"metricName":"GC (%)"}]],["Predicted genes",[]],["Similarity statistics",[{"values":[0,0,0,0,0,0,0,0,0,0,0,0,0],"quality":"Equal","isMain":false,"metricName":"# similar correct contigs"},{"values":[0,0,0,0,0,0,0,0,0,0,0,0,0],"quality":"Equal","isMain":false,"metricName":"# similar misassembled blocks"}]],["Reference statistics",[{"values":[39252,39252,39252,39252,39252,39252,39252,39252,39252,39252,39252,39252,39252],"quality":"Equal","isMain":false,"metricName":"Reference length"},{"values":[1,1,1,1,1,1,1,1,1,1,1,1,1],"quality":"Equal","isMain":false,"metricName":"Reference fragments"},{"values":["49.94","49.94","49.94","49.94","49.94","49.94","49.94","49.94","49.94","49.94","49.94","49.94","49.94"],"quality":"Equal","isMain":false,"metricName":"Reference GC (%)"}]]],"referenceName":"gi\_194100305\_ref\_NC","date":"18 July 2018, Wednesday, 16:41:39","order":[0,1,2,3,4,5,6,7,8,9,10,11,12],"assembliesNames":["ABySS\_127","ABySS\_63","CLC","IDBA\_UD","MEGAHIT","MIRA","SOAPdenovo2","SPAdes","SPAdes\_meta","SPAdes\_sc","SPAdes\_sc\_careful","Velvet","Geneious"]},{"assembliesWithNs":null,"minContig":500,"report":[["Genome statistics",[{"values":["43.395","33.627","62.728","89.838","84.885","73.440","22.430","96.025","89.699","78.046","66.692","41.567","31.353"],"quality":"More is better","isMain":true,"metricName":"Genome fraction (%)"},{"values":["1.007","1.000","1.007","1.025","1.046","1.004","1.013","1.089","1.024","1.046","1.040","1.003","1.000"],"quality":"Less is better","isMain":true,"metricName":"Duplication ratio"},{"values":[2156,1625,3995,11481,5761,4233,1014,12145,6968,4677,2172,2543,6453],"quality":"More is better","isMain":true,"metricName":"Largest alignment"},{"values":[16980,13060,24539,35671,34482,28645,8712,40334,35664,31695,26654,16179,12179],"quality":"More is better","isMain":true,"metricName":"Total aligned length"},{"values":[null,null,735,4115,3074,1758,null,10380,4063,1901,797,null,null],"quality":"More is better","isMain":false,"metricName":"NG50"},{"values":[null,null,null,2466,1400,null,null,5050,1989,703,null,null,null],"quality":"More is better","isMain":false,"metricName":"NG75"},{"values":[1405,1285,1090,4039,3074,2339,641,10105,4063,2105,1116,1356,6453],"quality":"More is better","isMain":false,"metricName":"NA50"},{"values":[730,606,763,2733,1606,1290,531,5050,2086,915,699,886,3011],"quality":"More is better","isMain":false,"metricName":"NA75"},{"values":[null,null,735,4039,3074,1758,null,10105,4063,1411,703,null,null],"quality":"More is better","isMain":true,"metricName":"NGA50"},{"values":[null,null,null,2466,1383,null,null,5050,1986,606,null,null,null],"quality":"More is better","isMain":false,"metricName":"NGA75"},{"values":[null,null,15,3,5,7,null,2,4,6,14,null,null],"quality":"Less is better","isMain":false,"metricName":"LG50"},{"values":[null,null,null,6,10,null,null,4,8,16,null,null,null],"quality":"Less is better","isMain":false,"metricName":"LG75"},{"values":[5,5,6,3,5,5,7,2,4,5,8,5,1],"quality":"Less is better","isMain":false,"metricName":"LA50"},{"values":[11,9,13,5,9,9,12,4,7,12,16,9,2],"quality":"Less is better","isMain":false,"metricName":"LA75"},{"values":[null,null,15,3,5,7,null,2,4,7,15,null,null],"quality":"Less is better","isMain":true,"metricName":"LGA50"},{"values":[null,null,null,6,11,null,null,4,8,18,null,null,null],"quality":"Less is better","isMain":false,"metricName":"LGA75"}]],["Misassemblies",[{"values":[0,0,0,0,0,0,0,0,0,0,0,0,0],"quality":"Less is better","isMain":true,"metricName":"# misassemblies"},{"values":[0,0,0,0,0,0,0,0,0,0,0,0,0],"quality":"Less is better","isMain":false,"metricName":" # relocations"},{"values":[0,0,0,0,0,0,0,0,0,0,0,0,0],"quality":"Less is better","isMain":false,"metricName":" # translocations"},{"values":[0,0,0,0,0,0,0,0,0,0,0,0,0],"quality":"Less is better","isMain":false,"metricName":" # inversions"},{"values":[0,0,0,0,0,0,0,0,0,0,0,0,0],"quality":"Less is better","isMain":false,"metricName":"# misassembled contigs"},{"values":[0,0,0,0,0,0,0,0,0,0,0,0,0],"quality":"Less is better","isMain":true,"metricName":"Misassembled contigs length"},{"values":[0,0,0,1,0,1,0,1,0,1,1,1,0],"quality":"Less is better","isMain":false,"metricName":"# local misassemblies"},{"values":[0,0,0,0,0,0,1,0,0,0,0,0,0],"quality":"Less is better","isMain":false,"metricName":"# unaligned mis. contigs"}]],["Unaligned",[{"values":[0,0,0,0,0,0,0,0,0,0,0,0,0],"quality":"Less is better","isMain":false,"metricName":"# fully unaligned contigs"},{"values":[0,0,0,0,0,0,0,0,0,0,0,0,0],"quality":"Less is better","isMain":false,"metricName":"Fully unaligned length"},{"values":[0,0,0,1,1,0,2,0,1,1,0,0,0],"quality":"Less is better","isMain":false,"metricName":"# partially unaligned contigs"},{"values":[0,0,0,692,692,0,1677,0,1147,1455,0,0,0],"quality":"Less is better","isMain":false,"metricName":"Partially unaligned length"}]],["Mismatches",[{"values":[25,31,119,323,179,17,10,371,126,168,84,60,20],"quality":"Less is better","isMain":false,"metricName":"# mismatches"},{"values":[4,4,6,9,5,1,1,16,7,9,4,2,1],"quality":"Less is better","isMain":false,"metricName":"# indels"},{"values":[5,5,7,20,15,1,50,20,11,12,4,4,1],"quality":"Less is better","isMain":false,"metricName":"Indels length"},{"values":["148.32","237.35","488.43","925.66","542.92","59.60","114.78","994.72","361.65","554.20","324.27","371.63","164.23"],"quality":"Less is better","isMain":true,"metricName":"# mismatches per 100 kbp"},{"values":["23.73","30.63","24.63","25.79","15.17","3.51","11.48","42.90","20.09","29.69","15.44","12.39","8.21"],"quality":"Less is better","isMain":true,"metricName":"# indels per 100 kbp"},{"values":[4,4,6,8,4,1,0,16,7,9,4,2,1],"quality":"Less is better","isMain":false,"metricName":" # indels (<= 5 bp)"},{"values":[0,0,0,1,1,0,1,0,0,0,0,0,0],"quality":"Less is better","isMain":false,"metricName":" # indels (> 5 bp)"},{"values":[0,0,0,0,0,70,642,0,0,1180,0,10,221],"quality":"Less is better","isMain":false,"metricName":"# N's"},{"values":["0.00","0.00","0.00","0.00","0.00","244.37","6111.38","0.00","0.00","3559.58","0.00","61.77","1814.60"],"quality":"Less is better","isMain":true,"metricName":"# N's per 100 kbp"}]],["Statistics without reference",[{"values":[17,14,23,11,15,15,14,10,13,22,26,15,3],"quality":"Equal","isMain":true,"metricName":"# contigs"},{"values":[6,5,8,10,13,12,2,7,11,8,8,5,3],"quality":"Equal","isMain":false,"metricName":"# contigs (>= 1000 bp)"},{"values":[0,0,0,2,1,0,0,4,2,1,0,0,1],"quality":"Equal","isMain":false,"metricName":"# contigs (>= 5000 bp)"},{"values":[0,0,0,1,0,0,0,2,0,0,0,0,0],"quality":"Equal","isMain":false,"metricName":"# contigs (>= 10000 bp)"},{"values":[0,0,0,0,0,0,0,0,0,0,0,0,0],"quality":"Equal","isMain":false,"metricName":"# contigs (>= 25000 bp)"},{"values":[0,0,0,0,0,0,0,0,0,0,0,0,0],"quality":"Equal","isMain":false,"metricName":"# contigs (>= 50000 bp)"},{"values":[2156,1625,3995,11481,5763,4233,1436,12145,6968,6132,2449,2543,6453],"quality":"More is better","isMain":true,"metricName":"Largest contig"},{"values":[16980,13062,24539,36454,35183,28645,10505,40609,36814,33150,26931,16189,12179],"quality":"More is better","isMain":true,"metricName":"Total length"},{"values":[9552,7576,14373,35698,33695,26147,2450,38622,35649,22790,14337,8729,12179],"quality":"More is better","isMain":true,"metricName":"Total length (>= 1000 bp)"},{"values":[0,0,0,17033,5763,0,0,32820,12205,6132,0,0,6453],"quality":"More is better","isMain":false,"metricName":"Total length (>= 5000 bp)"},{"values":[0,0,0,11481,0,0,0,22525,0,0,0,0,0],"quality":"More is better","isMain":true,"metricName":"Total length (>= 10000 bp)"},{"values":[0,0,0,0,0,0,0,0,0,0,0,0,0],"quality":"More is better","isMain":false,"metricName":"Total length (>= 25000 bp)"},{"values":[0,0,0,0,0,0,0,0,0,0,0,0,0],"quality":"More is better","isMain":true,"metricName":"Total length (>= 50000 bp)"},{"values":[1405,1285,1090,4115,3074,2339,735,10380,4063,2105,1116,1356,6453],"quality":"More is better","isMain":false,"metricName":"N50"},{"values":[730,608,763,3425,1901,1290,641,5050,2334,934,703,890,3011],"quality":"More is better","isMain":false,"metricName":"N75"},{"values":[5,5,6,3,5,5,6,2,4,5,8,5,1],"quality":"Less is better","isMain":false,"metricName":"L50"},{"values":[11,9,13,5,8,9,10,4,7,11,15,9,2],"quality":"Less is better","isMain":false,"metricName":"L75"},{"values":["48.97","48.47","48.09","48.54","48.82","48.60","46.90","48.42","48.41","48.32","47.97","48.45","49.72"],"quality":"Equal","isMain":false,"metricName":"GC (%)"}]],["Predicted genes",[]],["Similarity statistics",[{"values":[0,0,0,0,0,0,0,0,0,0,0,0,0],"quality":"Equal","isMain":false,"metricName":"# similar correct contigs"},{"values":[0,0,0,0,0,0,0,0,0,0,0,0,0],"quality":"Equal","isMain":false,"metricName":"# similar misassembled blocks"}]],["Reference statistics",[{"values":[38841,38841,38841,38841,38841,38841,38841,38841,38841,38841,38841,38841,38841],"quality":"Equal","isMain":false,"metricName":"Reference length"},{"values":[1,1,1,1,1,1,1,1,1,1,1,1,1],"quality":"Equal","isMain":false,"metricName":"Reference fragments"},{"values":["48.39","48.39","48.39","48.39","48.39","48.39","48.39","48.39","48.39","48.39","48.39","48.39","48.39"],"quality":"Equal","isMain":false,"metricName":"Reference GC (%)"}]]],"referenceName":"gi\_194100359\_ref\_NC","date":"18 July 2018, Wednesday, 16:41:53","order":[0,1,2,3,4,5,6,7,8,9,10,11,12],"assembliesNames":["ABySS\_127","ABySS\_63","CLC","IDBA\_UD","MEGAHIT","MIRA","SOAPdenovo2","SPAdes","SPAdes\_meta","SPAdes\_sc","SPAdes\_sc\_careful","Velvet","Geneious"]},{"assembliesWithNs":null,"minContig":500,"report":[["Genome statistics",[{"values":["100.000","99.562","100.000","99.441","100.000","100.000","100.000","100.000","100.000","99.491","98.963","21.744","100.000"],"quality":"More is better","isMain":true,"metricName":"Genome fraction (%)"},{"values":["1.025","1.062","1.016","1.011","1.004","1.011","1.072","1.002","1.007","1.012","1.003","1.005","1.011"],"quality":"Less is better","isMain":true,"metricName":"Duplication ratio"},{"values":[41601,14782,69904,15678,59593,70696,70620,70031,51718,29542,61570,1436,70696],"quality":"More is better","isMain":true,"metricName":"Largest alignment"},{"values":[71634,73926,70991,70263,70167,70696,74925,70031,70408,70395,69377,15282,70696],"quality":"More is better","isMain":true,"metricName":"Total aligned length"},{"values":[41601,6609,69904,8748,59593,70696,74946,70031,51718,11101,64997,null,70696],"quality":"More is better","isMain":false,"metricName":"NG50"},{"values":[23703,3392,69904,4639,59593,70696,74946,70031,13569,7902,64997,null,70696],"quality":"More is better","isMain":false,"metricName":"NG75"},{"values":[41601,6487,69904,8748,59593,70696,70620,70031,51718,11101,61570,713,70696],"quality":"More is better","isMain":false,"metricName":"NA50"},{"values":[23703,3329,69904,4639,59593,70696,70620,70031,5553,6420,61570,582,70696],"quality":"More is better","isMain":false,"metricName":"NA75"},{"values":[41601,6609,69904,8748,59593,70696,70620,70031,51718,11101,61570,null,70696],"quality":"More is better","isMain":true,"metricName":"NGA50"},{"values":[23703,3392,69904,4639,59593,70696,70620,70031,7179,6420,61570,null,70696],"quality":"More is better","isMain":false,"metricName":"NGA75"},{"values":[1,3,1,3,1,1,1,1,1,2,1,null,1],"quality":"Less is better","isMain":false,"metricName":"LG50"},{"values":[2,7,1,7,1,1,1,1,2,4,1,null,1],"quality":"Less is better","isMain":false,"metricName":"LG75"},{"values":[1,4,1,3,1,1,1,1,1,2,1,9,1],"quality":"Less is better","isMain":false,"metricName":"LA50"},{"values":[2,8,1,7,1,1,1,1,3,4,1,14,1],"quality":"Less is better","isMain":false,"metricName":"LA75"},{"values":[1,3,1,3,1,1,1,1,1,2,1,null,1],"quality":"Less is better","isMain":true,"metricName":"LGA50"},{"values":[2,7,1,7,1,1,1,1,2,4,1,null,1],"quality":"Less is better","isMain":false,"metricName":"LGA75"}]],["Misassemblies",[{"values":[0,0,0,0,0,0,1,0,0,0,0,0,0],"quality":"Less is better","isMain":true,"metricName":"# misassemblies"},{"values":[0,0,0,0,0,0,1,0,0,0,0,0,0],"quality":"Less is better","isMain":false,"metricName":" # relocations"},{"values":[0,0,0,0,0,0,0,0,0,0,0,0,0],"quality":"Less is better","isMain":false,"metricName":" # translocations"},{"values":[0,0,0,0,0,0,0,0,0,0,0,0,0],"quality":"Less is better","isMain":false,"metricName":" # inversions"},{"values":[0,0,0,0,0,0,1,0,0,0,0,0,0],"quality":"Less is better","isMain":false,"metricName":"# misassembled contigs"},{"values":[0,0,0,0,0,0,74946,0,0,0,0,0,0],"quality":"Less is better","isMain":true,"metricName":"Misassembled contigs length"},{"values":[0,0,0,0,0,1,1,1,0,0,0,0,1],"quality":"Less is better","isMain":false,"metricName":"# local misassemblies"},{"values":[0,0,0,0,0,0,0,0,0,0,0,0,0],"quality":"Less is better","isMain":false,"metricName":"# unaligned mis. contigs"}]],["Unaligned",[{"values":[0,0,0,0,0,0,0,0,0,0,0,0,0],"quality":"Less is better","isMain":false,"metricName":"# fully unaligned contigs"},{"values":[0,0,0,0,0,0,0,0,0,0,0,0,0],"quality":"Less is better","isMain":false,"metricName":"Fully unaligned length"},{"values":[0,0,0,0,0,0,0,0,1,1,1,0,0],"quality":"Less is better","isMain":false,"metricName":"# partially unaligned contigs"},{"values":[0,0,0,0,0,0,0,0,10026,1482,3427,0,0],"quality":"Less is better","isMain":false,"metricName":"Partially unaligned length"}]],["Mismatches",[{"values":[0,0,0,0,2,1,17,0,31,8,0,3,1],"quality":"Less is better","isMain":false,"metricName":"# mismatches"},{"values":[1,0,0,0,2,0,0,0,3,9,0,0,0],"quality":"Less is better","isMain":false,"metricName":"# indels"},{"values":[35,0,0,0,2,0,0,0,9,16,0,0,0],"quality":"Less is better","isMain":false,"metricName":"Indels length"},{"values":["0.00","0.00","0.00","0.00","2.86","1.43","24.32","0.00","44.35","11.50","0.00","19.74","1.43"],"quality":"Less is better","isMain":true,"metricName":"# mismatches per 100 kbp"},{"values":["1.43","0.00","0.00","0.00","2.86","0.00","0.00","0.00","4.29","12.94","0.00","0.00","0.00"],"quality":"Less is better","isMain":true,"metricName":"# indels per 100 kbp"},{"values":[0,0,0,0,2,0,0,0,3,9,0,0,0],"quality":"Less is better","isMain":false,"metricName":" # indels (<= 5 bp)"},{"values":[1,0,0,0,0,0,0,0,0,0,0,0,0],"quality":"Less is better","isMain":false,"metricName":" # indels (> 5 bp)"},{"values":[50,138,0,0,0,0,0,0,0,0,0,0,0],"quality":"Less is better","isMain":false,"metricName":"# N's"},{"values":["69.80","186.67","0.00","0.00","0.00","0.00","0.00","0.00","0.00","0.00","0.00","0.00","0.00"],"quality":"Less is better","isMain":true,"metricName":"# N's per 100 kbp"}]],["Statistics without reference",[{"values":[4,23,3,13,2,1,1,1,5,14,3,21,1],"quality":"Equal","isMain":true,"metricName":"# contigs"},{"values":[3,12,1,12,2,1,1,1,5,8,3,2,1],"quality":"Equal","isMain":false,"metricName":"# contigs (>= 1000 bp)"},{"values":[3,4,1,4,2,1,1,1,4,4,1,0,1],"quality":"Equal","isMain":false,"metricName":"# contigs (>= 5000 bp)"},{"values":[2,2,1,2,2,1,1,1,2,2,1,0,1],"quality":"Equal","isMain":false,"metricName":"# contigs (>= 10000 bp)"},{"values":[1,0,1,0,1,1,1,1,1,1,1,0,1],"quality":"Equal","isMain":false,"metricName":"# contigs (>= 25000 bp)"},{"values":[0,0,1,0,1,1,1,1,1,0,1,0,1],"quality":"Equal","isMain":false,"metricName":"# contigs (>= 50000 bp)"},{"values":[41601,14782,69904,15678,59593,70696,74946,70031,51718,29542,64997,1436,70696],"quality":"More is better","isMain":true,"metricName":"Largest contig"},{"values":[71634,73926,70991,70263,70167,70696,74946,70031,80434,71877,72804,15282,70696],"quality":"More is better","isMain":true,"metricName":"Total length"},{"values":[70734,66126,69904,69404,70167,70696,74946,70031,80434,66865,72804,2608,70696],"quality":"More is better","isMain":true,"metricName":"Total length (>= 1000 bp)"},{"values":[70734,41677,69904,41611,70167,70696,74946,70031,78019,56818,64997,0,70696],"quality":"More is better","isMain":false,"metricName":"Total length (>= 5000 bp)"},{"values":[65304,28581,69904,27270,70167,70696,74946,70031,65287,40643,64997,0,70696],"quality":"More is better","isMain":true,"metricName":"Total length (>= 10000 bp)"},{"values":[41601,0,69904,0,59593,70696,74946,70031,51718,29542,64997,0,70696],"quality":"More is better","isMain":false,"metricName":"Total length (>= 25000 bp)"},{"values":[0,0,69904,0,59593,70696,74946,70031,51718,0,64997,0,70696],"quality":"More is better","isMain":true,"metricName":"Total length (>= 50000 bp)"},{"values":[41601,6487,69904,8748,59593,70696,74946,70031,51718,11101,64997,713,70696],"quality":"More is better","isMain":false,"metricName":"N50"},{"values":[23703,3329,69904,4639,59593,70696,74946,70031,13569,7902,64997,582,70696],"quality":"More is better","isMain":false,"metricName":"N75"},{"values":[1,4,1,3,1,1,1,1,1,2,1,9,1],"quality":"Less is better","isMain":false,"metricName":"L50"},{"values":[2,8,1,7,1,1,1,1,2,4,1,14,1],"quality":"Less is better","isMain":false,"metricName":"L75"},{"values":["68.34","68.30","68.35","68.35","68.34","68.28","68.39","68.34","68.62","68.34","68.37","68.09","68.28"],"quality":"Equal","isMain":false,"metricName":"GC (%)"}]],["Predicted genes",[]],["Similarity statistics",[{"values":[0,0,0,0,0,0,0,0,0,0,0,0,0],"quality":"Equal","isMain":false,"metricName":"# similar correct contigs"},{"values":[0,0,0,0,0,0,0,0,0,0,0,0,0],"quality":"Equal","isMain":false,"metricName":"# similar misassembled blocks"}]],["Reference statistics",[{"values":[69904,69904,69904,69904,69904,69904,69904,69904,69904,69904,69904,69904,69904],"quality":"Equal","isMain":false,"metricName":"Reference length"},{"values":[1,1,1,1,1,1,1,1,1,1,1,1,1],"quality":"Equal","isMain":false,"metricName":"Reference fragments"},{"values":["68.34","68.34","68.34","68.34","68.34","68.34","68.34","68.34","68.34","68.34","68.34","68.34","68.34"],"quality":"Equal","isMain":false,"metricName":"Reference GC (%)"}]]],"referenceName":"gi\_194100514\_ref\_NC","date":"18 July 2018, Wednesday, 16:42:07","order":[0,1,2,3,4,5,6,7,8,9,10,11,12],"assembliesNames":["ABySS\_127","ABySS\_63","CLC","IDBA\_UD","MEGAHIT","MIRA","Ray\_Meta","SPAdes","SPAdes\_meta","SPAdes\_sc","SPAdes\_sc\_careful","Velvet","Geneious"]},{"assembliesWithNs":null,"minContig":500,"report":[["Genome statistics",[{"values":["38.224","27.263","87.147","83.417","86.933","59.271","77.069","90.465","92.945","91.217","91.217","35.434","10.221"],"quality":"More is better","isMain":true,"metricName":"Genome fraction (%)"},{"values":["1.000","1.000","1.021","1.001","1.002","1.048","1.005","1.003","1.006","1.005","1.005","1.000","1.000"],"quality":"Less is better","isMain":true,"metricName":"Duplication ratio"},{"values":[2649,1824,5735,5578,6086,3402,3578,6327,6427,6327,6327,4810,2898],"quality":"More is better","isMain":true,"metricName":"Largest alignment"},{"values":[26799,19114,61967,58514,61101,43487,54214,63457,65567,64086,64086,24843,7166],"quality":"More is better","isMain":true,"metricName":"Total aligned length"},{"values":[null,null,2792,2394,2771,1040,1080,3494,2941,3271,3271,null,null],"quality":"More is better","isMain":false,"metricName":"NG50"},{"values":[null,null,1262,1365,1340,null,537,1994,1389,1623,1623,null,null],"quality":"More is better","isMain":false,"metricName":"NG75"},{"values":[1026,814,2825,2782,2946,1689,1437,3383,3072,3271,3271,1750,2750],"quality":"More is better","isMain":false,"metricName":"NA50"},{"values":[721,609,1899,1584,2129,1276,969,1899,1581,1623,1623,1106,2750],"quality":"More is better","isMain":false,"metricName":"NA75"},{"values":[null,null,2792,2394,2771,1040,1080,3072,2941,2930,2930,null,null],"quality":"More is better","isMain":true,"metricName":"NGA50"},{"values":[null,null,1262,1178,1261,null,537,1581,1389,1494,1494,null,null],"quality":"More is better","isMain":false,"metricName":"NGA75"},{"values":[null,null,10,10,9,20,19,8,9,8,8,null,null],"quality":"Less is better","isMain":false,"metricName":"LG50"},{"values":[null,null,18,20,17,null,40,14,17,16,16,null,null],"quality":"Less is better","isMain":false,"metricName":"LG75"},{"values":[10,9,9,8,8,10,12,8,8,8,8,5,2],"quality":"Less is better","isMain":false,"metricName":"LA50"},{"values":[18,16,15,15,14,17,24,14,15,15,15,9,2],"quality":"Less is better","isMain":false,"metricName":"LA75"},{"values":[null,null,10,10,9,20,19,9,9,9,9,null,null],"quality":"Less is better","isMain":true,"metricName":"LGA50"},{"values":[null,null,18,21,18,null,40,16,17,17,17,null,null],"quality":"Less is better","isMain":false,"metricName":"LGA75"}]],["Misassemblies",[{"values":[0,0,0,0,0,0,0,0,0,0,0,0,0],"quality":"Less is better","isMain":true,"metricName":"# misassemblies"},{"values":[0,0,0,0,0,0,0,0,0,0,0,0,0],"quality":"Less is better","isMain":false,"metricName":" # relocations"},{"values":[0,0,0,0,0,0,0,0,0,0,0,0,0],"quality":"Less is better","isMain":false,"metricName":" # translocations"},{"values":[0,0,0,0,0,0,0,0,0,0,0,0,0],"quality":"Less is better","isMain":false,"metricName":" # inversions"},{"values":[0,0,0,0,0,0,0,0,0,0,0,0,0],"quality":"Less is better","isMain":false,"metricName":"# misassembled contigs"},{"values":[0,0,0,0,0,0,0,0,0,0,0,0,0],"quality":"Less is better","isMain":true,"metricName":"Misassembled contigs length"},{"values":[0,0,0,0,0,0,0,0,0,0,0,0,0],"quality":"Less is better","isMain":false,"metricName":"# local misassemblies"},{"values":[0,0,0,0,0,0,0,0,0,0,0,0,0],"quality":"Less is better","isMain":false,"metricName":"# unaligned mis. contigs"}]],["Unaligned",[{"values":[0,0,0,0,0,0,0,0,0,0,0,0,0],"quality":"Less is better","isMain":false,"metricName":"# fully unaligned contigs"},{"values":[0,0,0,0,0,0,0,0,0,0,0,0,0],"quality":"Less is better","isMain":false,"metricName":"Fully unaligned length"},{"values":[0,0,0,1,1,0,0,3,0,2,2,0,0],"quality":"Less is better","isMain":false,"metricName":"# partially unaligned contigs"},{"values":[0,0,0,816,812,0,0,3252,0,1785,1785,0,0],"quality":"Less is better","isMain":false,"metricName":"Partially unaligned length"}]],["Mismatches",[{"values":[3,0,162,46,72,22,35,141,61,117,96,5,6],"quality":"Less is better","isMain":false,"metricName":"# mismatches"},{"values":[0,0,6,1,2,0,1,5,0,4,4,0,0],"quality":"Less is better","isMain":false,"metricName":"# indels"},{"values":[0,0,7,2,2,0,52,7,0,5,5,0,0],"quality":"Less is better","isMain":false,"metricName":"Indels length"},{"values":["11.19","0.00","265.14","78.65","118.13","52.94","64.78","222.31","93.61","182.95","150.11","20.13","83.73"],"quality":"Less is better","isMain":true,"metricName":"# mismatches per 100 kbp"},{"values":["0.00","0.00","9.82","1.71","3.28","0.00","1.85","7.88","0.00","6.25","6.25","0.00","0.00"],"quality":"Less is better","isMain":true,"metricName":"# indels per 100 kbp"},{"values":[0,0,6,1,2,0,0,5,0,4,4,0,0],"quality":"Less is better","isMain":false,"metricName":" # indels (<= 5 bp)"},{"values":[0,0,0,0,0,0,1,0,0,0,0,0,0],"quality":"Less is better","isMain":false,"metricName":" # indels (> 5 bp)"},{"values":[0,0,0,0,0,38,113,0,0,0,0,0,8],"quality":"Less is better","isMain":false,"metricName":"# N's"},{"values":["0.00","0.00","0.00","0.00","0.00","87.23","208.00","0.00","0.00","0.00","0.00","0.00","111.64"],"quality":"Less is better","isMain":true,"metricName":"# N's per 100 kbp"}]],["Statistics without reference",[{"values":[28,24,30,26,27,30,43,26,31,28,28,15,3],"quality":"Equal","isMain":true,"metricName":"# contigs"},{"values":[10,3,18,23,20,20,23,20,20,21,21,12,3],"quality":"Equal","isMain":false,"metricName":"# contigs (>= 1000 bp)"},{"values":[0,0,1,2,2,0,0,2,2,2,2,0,0],"quality":"Equal","isMain":false,"metricName":"# contigs (>= 5000 bp)"},{"values":[0,0,0,0,0,0,0,0,0,0,0,0,0],"quality":"Equal","isMain":false,"metricName":"# contigs (>= 10000 bp)"},{"values":[0,0,0,0,0,0,0,0,0,0,0,0,0],"quality":"Equal","isMain":false,"metricName":"# contigs (>= 25000 bp)"},{"values":[0,0,0,0,0,0,0,0,0,0,0,0,0],"quality":"Equal","isMain":false,"metricName":"# contigs (>= 50000 bp)"},{"values":[2649,1824,5874,5578,6086,3402,3578,6935,6427,6935,6935,4810,2898],"quality":"More is better","isMain":true,"metricName":"Largest contig"},{"values":[26799,19114,62401,59330,61913,43563,54327,66848,65567,66030,66030,24843,7166],"quality":"More is better","isMain":true,"metricName":"Total length"},{"values":[14036,4373,53094,56586,56546,36088,39903,62125,56975,60493,60493,22560,7166],"quality":"More is better","isMain":true,"metricName":"Total length (>= 1000 bp)"},{"values":[0,0,5874,10765,11664,0,0,12809,12149,12809,12809,0,0],"quality":"More is better","isMain":false,"metricName":"Total length (>= 5000 bp)"},{"values":[0,0,0,0,0,0,0,0,0,0,0,0,0],"quality":"More is better","isMain":true,"metricName":"Total length (>= 10000 bp)"},{"values":[0,0,0,0,0,0,0,0,0,0,0,0,0],"quality":"More is better","isMain":false,"metricName":"Total length (>= 25000 bp)"},{"values":[0,0,0,0,0,0,0,0,0,0,0,0,0],"quality":"More is better","isMain":true,"metricName":"Total length (>= 50000 bp)"},{"values":[1026,814,2908,2782,2946,1689,1437,3511,3072,3271,3271,1750,2750],"quality":"More is better","isMain":false,"metricName":"N50"},{"values":[721,609,1899,1584,2129,1276,969,2394,1581,1899,1899,1106,2750],"quality":"More is better","isMain":false,"metricName":"N75"},{"values":[10,9,8,8,8,10,12,7,8,8,8,5,2],"quality":"Less is better","isMain":false,"metricName":"L50"},{"values":[18,16,15,15,14,17,24,13,15,14,14,9,2],"quality":"Less is better","isMain":false,"metricName":"L75"},{"values":["56.82","57.02","56.29","56.38","56.37","56.64","56.24","56.45","56.32","56.43","56.43","56.37","56.02"],"quality":"Equal","isMain":false,"metricName":"GC (%)"}]],["Predicted genes",[]],["Similarity statistics",[{"values":[0,0,0,0,0,0,0,0,0,0,0,0,0],"quality":"Equal","isMain":false,"metricName":"# similar correct contigs"},{"values":[0,0,0,0,0,0,0,0,0,0,0,0,0],"quality":"Equal","isMain":false,"metricName":"# similar misassembled blocks"}]],["Reference statistics",[{"values":[70110,70110,70110,70110,70110,70110,70110,70110,70110,70110,70110,70110,70110],"quality":"Equal","isMain":false,"metricName":"Reference length"},{"values":[1,1,1,1,1,1,1,1,1,1,1,1,1],"quality":"Equal","isMain":false,"metricName":"Reference fragments"},{"values":["56.29","56.29","56.29","56.29","56.29","56.29","56.29","56.29","56.29","56.29","56.29","56.29","56.29"],"quality":"Equal","isMain":false,"metricName":"Reference GC (%)"}]]],"referenceName":"gi\_194100609\_ref\_NC","date":"18 July 2018, Wednesday, 16:42:22","order":[0,1,2,3,4,5,6,7,8,9,10,11,12],"assembliesNames":["ABySS\_127","ABySS\_63","CLC","IDBA\_UD","MEGAHIT","MIRA","SOAPdenovo2","SPAdes","SPAdes\_meta","SPAdes\_sc","SPAdes\_sc\_careful","Velvet","Geneious"]},{"assembliesWithNs":null,"minContig":500,"report":[["Genome statistics",[{"values":["3.680","20.999","100.000","100.000","100.000","100.000","100.000","100.000","100.000","100.000","100.000","99.926","100.000"],"quality":"More is better","isMain":true,"metricName":"Genome fraction (%)"},{"values":["1.000","1.004","1.000","1.001","1.001","1.791","1.031","1.001","1.000","1.000","1.000","1.001","1.011"],"quality":"Less is better","isMain":true,"metricName":"Duplication ratio"},{"values":[638,2995,137947,138046,138088,84533,92782,138074,138002,138000,138002,28476,84534],"quality":"More is better","isMain":true,"metricName":"Largest alignment"},{"values":[5076,28968,137947,138046,138088,247073,142177,138074,138002,138000,138002,138028,139474],"quality":"More is better","isMain":true,"metricName":"Total aligned length"},{"values":[null,null,137947,138046,138088,139473,142177,138074,138002,138000,138002,14597,139474],"quality":"More is better","isMain":false,"metricName":"NG50"},{"values":[null,null,137947,138046,138088,139473,142177,138074,138002,138000,138002,11335,139474],"quality":"More is better","isMain":false,"metricName":"NG75"},{"values":[565,679,137947,138046,138088,54940,92782,138074,138002,138000,138002,14597,84534],"quality":"More is better","isMain":false,"metricName":"NA50"},{"values":[536,585,137947,138046,138088,1401,49395,138074,138002,138000,138002,11335,54940],"quality":"More is better","isMain":false,"metricName":"NA75"},{"values":[null,null,137947,138046,138088,84533,92782,138074,138002,138000,138002,14597,84534],"quality":"More is better","isMain":true,"metricName":"NGA50"},{"values":[null,null,137947,138046,138088,54940,49395,138074,138002,138000,138002,11335,54940],"quality":"More is better","isMain":false,"metricName":"NGA75"},{"values":[null,null,1,1,1,1,1,1,1,1,1,4,1],"quality":"Less is better","isMain":false,"metricName":"LG50"},{"values":[null,null,1,1,1,1,1,1,1,1,1,6,1],"quality":"Less is better","isMain":false,"metricName":"LG75"},{"values":[5,12,1,1,1,2,1,1,1,1,1,4,1],"quality":"Less is better","isMain":false,"metricName":"LA50"},{"values":[7,23,1,1,1,29,2,1,1,1,1,6,2],"quality":"Less is better","isMain":false,"metricName":"LA75"},{"values":[null,null,1,1,1,1,1,1,1,1,1,4,1],"quality":"Less is better","isMain":true,"metricName":"LGA50"},{"values":[null,null,1,1,1,2,2,1,1,1,1,6,2],"quality":"Less is better","isMain":false,"metricName":"LGA75"}]],["Misassemblies",[{"values":[0,0,0,0,0,1,1,0,0,0,0,0,1],"quality":"Less is better","isMain":true,"metricName":"# misassemblies"},{"values":[0,0,0,0,0,1,1,0,0,0,0,0,1],"quality":"Less is better","isMain":false,"metricName":" # relocations"},{"values":[0,0,0,0,0,0,0,0,0,0,0,0,0],"quality":"Less is better","isMain":false,"metricName":" # translocations"},{"values":[0,0,0,0,0,0,0,0,0,0,0,0,0],"quality":"Less is better","isMain":false,"metricName":" # inversions"},{"values":[0,0,0,0,0,1,1,0,0,0,0,0,1],"quality":"Less is better","isMain":false,"metricName":"# misassembled contigs"},{"values":[0,0,0,0,0,139473,142177,0,0,0,0,0,139474],"quality":"Less is better","isMain":true,"metricName":"Misassembled contigs length"},{"values":[0,0,0,1,1,0,0,1,0,0,0,0,0],"quality":"Less is better","isMain":false,"metricName":"# local misassemblies"},{"values":[0,0,0,0,0,0,0,0,0,0,0,0,0],"quality":"Less is better","isMain":false,"metricName":"# unaligned mis. contigs"}]],["Unaligned",[{"values":[0,0,0,0,0,0,0,0,0,0,0,0,0],"quality":"Less is better","isMain":false,"metricName":"# fully unaligned contigs"},{"values":[0,0,0,0,0,0,0,0,0,0,0,0,0],"quality":"Less is better","isMain":false,"metricName":"Fully unaligned length"},{"values":[0,0,0,0,0,0,0,0,0,0,0,0,0],"quality":"Less is better","isMain":false,"metricName":"# partially unaligned contigs"},{"values":[0,0,0,0,0,0,0,0,0,0,0,0,0],"quality":"Less is better","isMain":false,"metricName":"Partially unaligned length"}]],["Mismatches",[{"values":[0,0,0,0,0,15,0,0,0,0,0,59,3],"quality":"Less is better","isMain":false,"metricName":"# mismatches"},{"values":[0,0,0,0,0,1,0,0,1,1,1,3,0],"quality":"Less is better","isMain":false,"metricName":"# indels"},{"values":[0,0,0,0,0,1,0,0,55,53,55,3,0],"quality":"Less is better","isMain":false,"metricName":"Indels length"},{"values":["0.00","0.00","0.00","0.00","0.00","10.87","0.00","0.00","0.00","0.00","0.00","42.80","2.17"],"quality":"Less is better","isMain":true,"metricName":"# mismatches per 100 kbp"},{"values":["0.00","0.00","0.00","0.00","0.00","0.72","0.00","0.00","0.72","0.72","0.72","2.18","0.00"],"quality":"Less is better","isMain":true,"metricName":"# indels per 100 kbp"},{"values":[0,0,0,0,0,1,0,0,0,0,0,3,0],"quality":"Less is better","isMain":false,"metricName":" # indels (<= 5 bp)"},{"values":[0,0,0,0,0,0,0,0,1,1,1,0,0],"quality":"Less is better","isMain":false,"metricName":" # indels (> 5 bp)"},{"values":[0,345,0,0,0,118,0,0,0,0,0,0,223],"quality":"Less is better","isMain":false,"metricName":"# N's"},{"values":["0.00","1186.46","0.00","0.00","0.00","47.76","0.00","0.00","0.00","0.00","0.00","0.00","159.89"],"quality":"Less is better","isMain":true,"metricName":"# N's per 100 kbp"}]],["Statistics without reference",[{"values":[9,36,1,1,1,82,1,1,1,1,1,13,1],"quality":"Equal","isMain":true,"metricName":"# contigs"},{"values":[0,4,1,1,1,71,1,1,1,1,1,12,1],"quality":"Equal","isMain":false,"metricName":"# contigs (>= 1000 bp)"},{"values":[0,0,1,1,1,1,1,1,1,1,1,9,1],"quality":"Equal","isMain":false,"metricName":"# contigs (>= 5000 bp)"},{"values":[0,0,1,1,1,1,1,1,1,1,1,6,1],"quality":"Equal","isMain":false,"metricName":"# contigs (>= 10000 bp)"},{"values":[0,0,1,1,1,1,1,1,1,1,1,1,1],"quality":"Equal","isMain":false,"metricName":"# contigs (>= 25000 bp)"},{"values":[0,0,1,1,1,1,1,1,1,1,1,0,1],"quality":"Equal","isMain":false,"metricName":"# contigs (>= 50000 bp)"},{"values":[638,2995,137947,138046,138088,139473,142177,138074,138002,138000,138002,28476,139474],"quality":"More is better","isMain":true,"metricName":"Largest contig"},{"values":[5076,29078,137947,138046,138088,247079,142177,138074,138002,138000,138002,138028,139474],"quality":"More is better","isMain":true,"metricName":"Total length"},{"values":[0,8728,137947,138046,138088,237992,142177,138074,138002,138000,138002,137482,139474],"quality":"More is better","isMain":true,"metricName":"Total length (>= 1000 bp)"},{"values":[0,0,137947,138046,138088,139473,142177,138074,138002,138000,138002,130249,139474],"quality":"More is better","isMain":false,"metricName":"Total length (>= 5000 bp)"},{"values":[0,0,137947,138046,138088,139473,142177,138074,138002,138000,138002,107334,139474],"quality":"More is better","isMain":true,"metricName":"Total length (>= 10000 bp)"},{"values":[0,0,137947,138046,138088,139473,142177,138074,138002,138000,138002,28476,139474],"quality":"More is better","isMain":false,"metricName":"Total length (>= 25000 bp)"},{"values":[0,0,137947,138046,138088,139473,142177,138074,138002,138000,138002,0,139474],"quality":"More is better","isMain":true,"metricName":"Total length (>= 50000 bp)"},{"values":[565,679,137947,138046,138088,139473,142177,138074,138002,138000,138002,14597,139474],"quality":"More is better","isMain":false,"metricName":"N50"},{"values":[536,585,137947,138046,138088,1402,142177,138074,138002,138000,138002,11335,139474],"quality":"More is better","isMain":false,"metricName":"N75"},{"values":[5,12,1,1,1,1,1,1,1,1,1,4,1],"quality":"Less is better","isMain":false,"metricName":"L50"},{"values":[7,23,1,1,1,28,1,1,1,1,1,6,1],"quality":"Less is better","isMain":false,"metricName":"L75"},{"values":["44.27","43.60","43.56","43.56","43.56","43.58","43.54","43.56","43.57","43.57","43.57","43.55","43.54"],"quality":"Equal","isMain":false,"metricName":"GC (%)"}]],["Predicted genes",[]],["Similarity statistics",[{"values":[0,0,0,0,0,0,0,0,0,0,0,0,0],"quality":"Equal","isMain":false,"metricName":"# similar correct contigs"},{"values":[0,0,0,0,0,0,0,0,0,0,0,0,0],"quality":"Equal","isMain":false,"metricName":"# similar misassembled blocks"}]],["Reference statistics",[{"values":[137947,137947,137947,137947,137947,137947,137947,137947,137947,137947,137947,137947,137947],"quality":"Equal","isMain":false,"metricName":"Reference length"},{"values":[1,1,1,1,1,1,1,1,1,1,1,1,1],"quality":"Equal","isMain":false,"metricName":"Reference fragments"},{"values":["43.56","43.56","43.56","43.56","43.56","43.56","43.56","43.56","43.56","43.56","43.56","43.56","43.56"],"quality":"Equal","isMain":false,"metricName":"Reference GC (%)"}]]],"referenceName":"gi\_194172859\_ref\_NC","date":"18 July 2018, Wednesday, 16:42:38","order":[0,1,2,3,4,5,6,7,8,9,10,11,12],"assembliesNames":["ABySS\_127","ABySS\_63","CLC","IDBA\_UD","MEGAHIT","MIRA","Ray\_Meta","SPAdes","SPAdes\_meta","SPAdes\_sc","SPAdes\_sc\_careful","Velvet","Geneious"]},{"assembliesWithNs":null,"minContig":500,"report":[["Genome statistics",[{"values":["100.000","21.616","7.867","100.000","100.000","100.000","100.000","100.000","100.000","100.000"],"quality":"More is better","isMain":true,"metricName":"Genome fraction (%)"},{"values":["1.000","1.000","1.000","368.351","1.003","1.007","1.003","1.003","1.003","3.138"],"quality":"Less is better","isMain":true,"metricName":"Duplication ratio"},{"values":[19282,847,511,20257,19282,19409,19337,19337,19337,19282],"quality":"More is better","isMain":true,"metricName":"Largest alignment"},{"values":[19282,4168,1517,7101223,19282,19409,19337,19337,19337,60510],"quality":"More is better","isMain":true,"metricName":"Total aligned length"},{"values":[19282,null,null,25596,19331,19409,19337,19337,19337,60510],"quality":"More is better","isMain":false,"metricName":"NG50"},{"values":[19282,null,null,25596,19331,19409,19337,19337,19337,60510],"quality":"More is better","isMain":false,"metricName":"NG75"},{"values":[19282,744,506,3788,19282,19409,19337,19337,19337,19282],"quality":"More is better","isMain":false,"metricName":"NA50"},{"values":[19282,593,500,2383,19282,19409,19337,19337,19337,17449],"quality":"More is better","isMain":false,"metricName":"NA75"},{"values":[19282,null,null,20257,19282,19409,19337,19337,19337,19282],"quality":"More is better","isMain":true,"metricName":"NGA50"},{"values":[19282,null,null,20257,19282,19409,19337,19337,19337,19282],"quality":"More is better","isMain":false,"metricName":"NGA75"},{"values":[1,null,null,1,1,1,1,1,1,1],"quality":"Less is better","isMain":false,"metricName":"LG50"},{"values":[1,null,null,1,1,1,1,1,1,1],"quality":"Less is better","isMain":false,"metricName":"LG75"},{"values":[1,3,2,553,1,1,1,1,1,2],"quality":"Less is better","isMain":false,"metricName":"LA50"},{"values":[1,5,3,1151,1,1,1,1,1,3],"quality":"Less is better","isMain":false,"metricName":"LA75"},{"values":[1,null,null,1,1,1,1,1,1,1],"quality":"Less is better","isMain":true,"metricName":"LGA50"},{"values":[1,null,null,1,1,1,1,1,1,1],"quality":"Less is better","isMain":false,"metricName":"LGA75"}]],["Misassemblies",[{"values":[0,0,0,7,0,0,0,0,0,3],"quality":"Less is better","isMain":true,"metricName":"# misassemblies"},{"values":[0,0,0,7,0,0,0,0,0,3],"quality":"Less is better","isMain":false,"metricName":" # relocations"},{"values":[0,0,0,0,0,0,0,0,0,0],"quality":"Less is better","isMain":false,"metricName":" # translocations"},{"values":[0,0,0,0,0,0,0,0,0,0],"quality":"Less is better","isMain":false,"metricName":" # inversions"},{"values":[0,0,0,7,0,0,0,0,0,1],"quality":"Less is better","isMain":false,"metricName":"# misassembled contigs"},{"values":[0,0,0,160983,0,0,0,0,0,60510],"quality":"Less is better","isMain":true,"metricName":"Misassembled contigs length"},{"values":[0,0,0,2,0,1,0,0,0,0],"quality":"Less is better","isMain":false,"metricName":"# local misassemblies"},{"values":[0,0,0,0,0,0,0,0,0,0],"quality":"Less is better","isMain":false,"metricName":"# unaligned mis. contigs"}]],["Unaligned",[{"values":[0,0,0,0,0,0,0,0,0,0],"quality":"Less is better","isMain":false,"metricName":"# fully unaligned contigs"},{"values":[0,0,0,0,0,0,0,0,0,0],"quality":"Less is better","isMain":false,"metricName":"Fully unaligned length"},{"values":[0,0,0,0,0,0,0,0,0,0],"quality":"Less is better","isMain":false,"metricName":"# partially unaligned contigs"},{"values":[0,0,0,0,0,0,0,0,0,0],"quality":"Less is better","isMain":false,"metricName":"Partially unaligned length"}]],["Mismatches",[{"values":[0,0,12,74,0,0,0,0,0,66],"quality":"Less is better","isMain":false,"metricName":"# mismatches"},{"values":[0,0,0,5,0,0,1,1,1,0],"quality":"Less is better","isMain":false,"metricName":"# indels"},{"values":[0,0,0,5,0,0,55,55,55,0],"quality":"Less is better","isMain":false,"metricName":"Indels length"},{"values":["0.00","0.00","791.03","383.78","0.00","0.00","0.00","0.00","0.00","342.29"],"quality":"Less is better","isMain":true,"metricName":"# mismatches per 100 kbp"},{"values":["0.00","0.00","0.00","25.93","0.00","0.00","5.19","5.19","5.19","0.00"],"quality":"Less is better","isMain":true,"metricName":"# indels per 100 kbp"},{"values":[0,0,0,5,0,0,0,0,0,0],"quality":"Less is better","isMain":false,"metricName":" # indels (<= 5 bp)"},{"values":[0,0,0,0,0,0,1,1,1,0],"quality":"Less is better","isMain":false,"metricName":" # indels (> 5 bp)"},{"values":[0,0,0,1224,0,0,0,0,0,67],"quality":"Less is better","isMain":false,"metricName":"# N's"},{"values":["0.00","0.00","0.00","17.23","0.00","0.00","0.00","0.00","0.00","110.73"],"quality":"Less is better","isMain":true,"metricName":"# N's per 100 kbp"}]],["Statistics without reference",[{"values":[1,6,3,2113,1,1,1,1,1,1],"quality":"Equal","isMain":true,"metricName":"# contigs"},{"values":[1,0,0,2098,1,1,1,1,1,1],"quality":"Equal","isMain":false,"metricName":"# contigs (>= 1000 bp)"},{"values":[1,0,0,320,1,1,1,1,1,1],"quality":"Equal","isMain":false,"metricName":"# contigs (>= 5000 bp)"},{"values":[1,0,0,60,1,1,1,1,1,1],"quality":"Equal","isMain":false,"metricName":"# contigs (>= 10000 bp)"},{"values":[0,0,0,1,0,0,0,0,0,1],"quality":"Equal","isMain":false,"metricName":"# contigs (>= 25000 bp)"},{"values":[0,0,0,0,0,0,0,0,0,1],"quality":"Equal","isMain":false,"metricName":"# contigs (>= 50000 bp)"},{"values":[19282,847,511,25596,19331,19409,19337,19337,19337,60510],"quality":"More is better","isMain":true,"metricName":"Largest contig"},{"values":[19282,4168,1517,7102540,19331,19409,19337,19337,19337,60510],"quality":"More is better","isMain":true,"metricName":"Total length"},{"values":[19282,0,0,7089920,19331,19409,19337,19337,19337,60510],"quality":"More is better","isMain":true,"metricName":"Total length (>= 1000 bp)"},{"values":[19282,0,0,2574909,19331,19409,19337,19337,19337,60510],"quality":"More is better","isMain":false,"metricName":"Total length (>= 5000 bp)"},{"values":[19282,0,0,829764,19331,19409,19337,19337,19337,60510],"quality":"More is better","isMain":true,"metricName":"Total length (>= 10000 bp)"},{"values":[0,0,0,25596,0,0,0,0,0,60510],"quality":"More is better","isMain":false,"metricName":"Total length (>= 25000 bp)"},{"values":[0,0,0,0,0,0,0,0,0,60510],"quality":"More is better","isMain":true,"metricName":"Total length (>= 50000 bp)"},{"values":[19282,744,506,3793,19331,19409,19337,19337,19337,60510],"quality":"More is better","isMain":false,"metricName":"N50"},{"values":[19282,593,500,2384,19331,19409,19337,19337,19337,60510],"quality":"More is better","isMain":false,"metricName":"N75"},{"values":[1,3,2,545,1,1,1,1,1,1],"quality":"Less is better","isMain":false,"metricName":"L50"},{"values":[1,5,3,1143,1,1,1,1,1,1],"quality":"Less is better","isMain":false,"metricName":"L75"},{"values":["39.99","39.64","40.87","40.11","40.00","39.99","39.96","40.02","40.02","39.94"],"quality":"Equal","isMain":false,"metricName":"GC (%)"}]],["Predicted genes",[]],["Similarity statistics",[{"values":[0,0,0,0,0,0,0,0,0,0],"quality":"Equal","isMain":false,"metricName":"# similar correct contigs"},{"values":[0,0,0,0,0,0,0,0,0,0],"quality":"Equal","isMain":false,"metricName":"# similar misassembled blocks"}]],["Reference statistics",[{"values":[19282,19282,19282,19282,19282,19282,19282,19282,19282,19282],"quality":"Equal","isMain":false,"metricName":"Reference length"},{"values":[1,1,1,1,1,1,1,1,1,1],"quality":"Equal","isMain":false,"metricName":"Reference fragments"},{"values":["39.99","39.99","39.99","39.99","39.99","39.99","39.99","39.99","39.99","39.99"],"quality":"Equal","isMain":false,"metricName":"Reference GC (%)"}]]],"referenceName":"gi\_194186868\_ref\_NC","date":"18 July 2018, Wednesday, 16:43:27","order":[0,1,2,3,4,5,6,7,8,9],"assembliesNames":["CLC","IDBA\_UD","MEGAHIT","MIRA","Ray\_Meta","SPAdes","SPAdes\_meta","SPAdes\_sc","SPAdes\_sc\_careful","Geneious"]},{"assembliesWithNs":null,"minContig":500,"report":[["Genome statistics",[{"values":["82.197","80.060","89.531","90.606","93.341","98.761","99.972","91.962","100.000","79.572","79.776","71.062","99.803"],"quality":"More is better","isMain":true,"metricName":"Genome fraction (%)"},{"values":["1.067","1.032","1.019","1.031","1.049","1.182","1.000","1.060","1.001","1.011","1.016","1.003","1.275"],"quality":"Less is better","isMain":true,"metricName":"Duplication ratio"},{"values":[11251,10381,7623,9828,11160,42479,75790,16895,53324,15819,15666,4075,54602],"quality":"More is better","isMain":true,"metricName":"Largest alignment"},{"values":[66495,62430,69112,70841,74207,88435,75790,73400,75921,60874,61194,54014,96441],"quality":"More is better","isMain":true,"metricName":"Total aligned length"},{"values":[3246,4511,2452,2366,2821,42479,75790,3727,53324,4343,3801,1528,56961],"quality":"More is better","isMain":false,"metricName":"NG50"},{"values":[841,694,1083,933,1288,20963,75790,1956,22597,1194,880,null,56961],"quality":"More is better","isMain":false,"metricName":"NG75"},{"values":[4252,4574,2841,2504,2987,20963,75790,3480,53324,7214,4343,2264,54602],"quality":"More is better","isMain":false,"metricName":"NA50"},{"values":[2240,2266,1485,1078,1291,10165,75790,1956,22597,2657,2474,1165,19493],"quality":"More is better","isMain":false,"metricName":"NA75"},{"values":[3246,4511,2452,2366,2821,42479,75790,3480,53324,4343,3801,1528,54602],"quality":"More is better","isMain":true,"metricName":"NGA50"},{"values":[841,676,1083,933,1288,20963,75790,1602,22597,1194,880,null,22346],"quality":"More is better","isMain":false,"metricName":"NGA75"},{"values":[8,6,8,9,8,1,1,5,1,4,6,14,1],"quality":"Less is better","isMain":false,"metricName":"LG50"},{"values":[18,18,20,22,17,2,1,12,2,12,15,null,1],"quality":"Less is better","isMain":false,"metricName":"LG75"},{"values":[6,5,7,8,7,2,1,6,1,3,4,9,1],"quality":"Less is better","isMain":false,"metricName":"LA50"},{"values":[12,9,16,18,16,3,1,12,2,6,9,16,3],"quality":"Less is better","isMain":false,"metricName":"LA75"},{"values":[8,6,8,9,8,1,1,6,1,4,6,14,1],"quality":"Less is better","isMain":true,"metricName":"LGA50"},{"values":[18,18,20,22,17,2,1,13,2,12,15,null,2],"quality":"Less is better","isMain":false,"metricName":"LGA75"}]],["Misassemblies",[{"values":[0,0,0,0,0,0,0,0,0,0,0,0,0],"quality":"Less is better","isMain":true,"metricName":"# misassemblies"},{"values":[0,0,0,0,0,0,0,0,0,0,0,0,0],"quality":"Less is better","isMain":false,"metricName":" # relocations"},{"values":[0,0,0,0,0,0,0,0,0,0,0,0,0],"quality":"Less is better","isMain":false,"metricName":" # translocations"},{"values":[0,0,0,0,0,0,0,0,0,0,0,0,0],"quality":"Less is better","isMain":false,"metricName":" # inversions"},{"values":[0,0,0,0,0,0,0,0,0,0,0,0,0],"quality":"Less is better","isMain":false,"metricName":"# misassembled contigs"},{"values":[0,0,0,0,0,0,0,0,0,0,0,0,0],"quality":"Less is better","isMain":true,"metricName":"Misassembled contigs length"},{"values":[0,0,0,0,0,3,0,0,0,1,1,0,11],"quality":"Less is better","isMain":false,"metricName":"# local misassemblies"},{"values":[0,0,0,0,0,0,0,0,0,0,0,0,0],"quality":"Less is better","isMain":false,"metricName":"# unaligned mis. contigs"}]],["Unaligned",[{"values":[0,0,0,0,0,0,0,0,0,0,0,0,0],"quality":"Less is better","isMain":false,"metricName":"# fully unaligned contigs"},{"values":[0,0,0,0,0,0,0,0,0,0,0,0,0],"quality":"Less is better","isMain":false,"metricName":"Fully unaligned length"},{"values":[0,0,0,0,0,0,0,1,0,0,0,0,3],"quality":"Less is better","isMain":false,"metricName":"# partially unaligned contigs"},{"values":[0,0,0,0,0,0,0,937,0,0,0,0,7117],"quality":"Less is better","isMain":false,"metricName":"Partially unaligned length"}]],["Mismatches",[{"values":[0,3,26,0,37,129,2,0,41,16,7,10,174],"quality":"Less is better","isMain":false,"metricName":"# mismatches"},{"values":[1,2,0,0,0,5,0,0,0,0,0,1,16],"quality":"Less is better","isMain":false,"metricName":"# indels"},{"values":[1,3,0,0,0,5,0,0,0,0,0,1,32],"quality":"Less is better","isMain":false,"metricName":"Indels length"},{"values":["0.00","4.94","38.31","0.00","52.29","172.29","2.64","0.00","54.08","26.52","11.57","18.56","229.97"],"quality":"Less is better","isMain":true,"metricName":"# mismatches per 100 kbp"},{"values":["1.60","3.30","0.00","0.00","0.00","6.68","0.00","0.00","0.00","0.00","0.00","1.86","21.15"],"quality":"Less is better","isMain":true,"metricName":"# indels per 100 kbp"},{"values":[1,2,0,0,0,5,0,0,0,0,0,1,15],"quality":"Less is better","isMain":false,"metricName":" # indels (<= 5 bp)"},{"values":[0,0,0,0,0,0,0,0,0,0,0,0,1],"quality":"Less is better","isMain":false,"metricName":" # indels (> 5 bp)"},{"values":[28,244,0,0,0,5,0,1470,0,91,243,0,216],"quality":"Less is better","isMain":false,"metricName":"# N's"},{"values":["42.10","389.53","0.00","0.00","0.00","5.65","0.00","1963.40","0.00","149.27","395.53","0.00","208.58"],"quality":"Less is better","isMain":true,"metricName":"# N's per 100 kbp"}]],["Statistics without reference",[{"values":[33,27,36,42,37,11,1,31,2,16,21,36,3],"quality":"Equal","isMain":true,"metricName":"# contigs"},{"values":[13,12,24,18,22,9,1,18,2,13,13,17,3],"quality":"Equal","isMain":false,"metricName":"# contigs (>= 1000 bp)"},{"values":[1,3,3,1,3,4,1,3,2,3,3,0,3],"quality":"Equal","isMain":false,"metricName":"# contigs (>= 5000 bp)"},{"values":[1,1,0,0,1,3,1,1,2,2,1,0,3],"quality":"Equal","isMain":false,"metricName":"# contigs (>= 10000 bp)"},{"values":[0,0,0,0,0,1,1,0,1,0,0,0,2],"quality":"Equal","isMain":false,"metricName":"# contigs (>= 25000 bp)"},{"values":[0,0,0,0,0,0,1,0,1,0,0,0,1],"quality":"Equal","isMain":false,"metricName":"# contigs (>= 50000 bp)"},{"values":[11251,10381,7623,9828,11160,42479,75790,17832,53324,15819,15819,4075,56961],"quality":"More is better","isMain":true,"metricName":"Largest contig"},{"values":[66515,62639,69170,70841,74213,88464,75790,74870,75921,60965,61437,54014,103558],"quality":"More is better","isMain":true,"metricName":"Total length"},{"values":[52610,52498,61895,53610,63759,86598,75790,65850,75921,59028,55523,41699,103558],"quality":"More is better","isMain":true,"metricName":"Total length (>= 1000 bp)"},{"values":[11251,24765,20346,9828,24153,78723,75790,30284,75921,35837,27727,0,103558],"quality":"More is better","isMain":false,"metricName":"Total length (>= 5000 bp)"},{"values":[11251,10381,0,0,11160,73608,75790,17832,75921,28532,15819,0,103558],"quality":"More is better","isMain":true,"metricName":"Total length (>= 10000 bp)"},{"values":[0,0,0,0,0,42479,75790,0,53324,0,0,0,82963],"quality":"More is better","isMain":false,"metricName":"Total length (>= 25000 bp)"},{"values":[0,0,0,0,0,0,75790,0,53324,0,0,0,56961],"quality":"More is better","isMain":true,"metricName":"Total length (>= 50000 bp)"},{"values":[4252,4574,2841,2504,2987,20963,75790,3727,53324,7305,4343,2264,56961],"quality":"More is better","isMain":false,"metricName":"N50"},{"values":[2240,2266,1485,1078,1291,10166,75790,1956,22597,2657,2657,1165,26002],"quality":"More is better","isMain":false,"metricName":"N75"},{"values":[6,5,7,8,7,2,1,5,1,3,4,9,1],"quality":"Less is better","isMain":false,"metricName":"L50"},{"values":[12,9,16,18,16,3,1,12,2,6,8,16,2],"quality":"Less is better","isMain":false,"metricName":"L75"},{"values":["63.03","63.09","62.90","62.97","63.06","62.92","62.91","62.91","62.92","63.17","63.10","62.97","63.07"],"quality":"Equal","isMain":false,"metricName":"GC (%)"}]],["Predicted genes",[]],["Similarity statistics",[{"values":[0,0,0,0,0,0,0,0,0,0,0,0,0],"quality":"Equal","isMain":false,"metricName":"# similar correct contigs"},{"values":[0,0,0,0,0,0,0,0,0,0,0,0,0],"quality":"Equal","isMain":false,"metricName":"# similar misassembled blocks"}]],["Reference statistics",[{"values":[75811,75811,75811,75811,75811,75811,75811,75811,75811,75811,75811,75811,75811],"quality":"Equal","isMain":false,"metricName":"Reference length"},{"values":[1,1,1,1,1,1,1,1,1,1,1,1,1],"quality":"Equal","isMain":false,"metricName":"Reference fragments"},{"values":["62.92","62.92","62.92","62.92","62.92","62.92","62.92","62.92","62.92","62.92","62.92","62.92","62.92"],"quality":"Equal","isMain":false,"metricName":"Reference GC (%)"}]]],"referenceName":"gi\_194303053\_ref\_NC","date":"18 July 2018, Wednesday, 16:43:43","order":[0,1,2,3,4,5,6,7,8,9,10,11,12],"assembliesNames":["ABySS\_127","ABySS\_63","CLC","IDBA\_UD","MEGAHIT","MIRA","Ray\_Meta","SPAdes","SPAdes\_meta","SPAdes\_sc","SPAdes\_sc\_careful","Velvet","Geneious"]},{"assembliesWithNs":null,"minContig":500,"report":[["Genome statistics",[{"values":["0.764","8.559","0.809","3.456","7.467","13.169"],"quality":"More is better","isMain":true,"metricName":"Genome fraction (%)"},{"values":["1.000","1.000","1.000","1.000","1.011","1.006"],"quality":"Less is better","isMain":true,"metricName":"Duplication ratio"},{"values":[520,2952,551,930,930,1732],"quality":"More is better","isMain":true,"metricName":"Largest alignment"},{"values":[520,5828,551,2353,5139,9025],"quality":"More is better","isMain":true,"metricName":"Total aligned length"},{"values":[520,2952,551,915,822,896],"quality":"More is better","isMain":false,"metricName":"NA50"},{"values":[520,2876,551,915,707,853],"quality":"More is better","isMain":false,"metricName":"NA75"},{"values":[null,null,null,null,null,null],"quality":"More is better","isMain":true,"metricName":"NGA50"},{"values":[1,1,1,2,3,4],"quality":"Less is better","isMain":false,"metricName":"LA50"},{"values":[1,2,1,2,5,6],"quality":"Less is better","isMain":false,"metricName":"LA75"}]],["Misassemblies",[{"values":[0,0,0,0,0,0],"quality":"Less is better","isMain":true,"metricName":"# misassemblies"},{"values":[0,0,0,0,0,0],"quality":"Less is better","isMain":false,"metricName":" # relocations"},{"values":[0,0,0,0,0,0],"quality":"Less is better","isMain":false,"metricName":" # translocations"},{"values":[0,0,0,0,0,0],"quality":"Less is better","isMain":false,"metricName":" # inversions"},{"values":[0,0,0,0,0,0],"quality":"Less is better","isMain":false,"metricName":"# misassembled contigs"},{"values":[0,0,0,0,0,0],"quality":"Less is better","isMain":true,"metricName":"Misassembled contigs length"},{"values":[0,0,0,0,0,0],"quality":"Less is better","isMain":false,"metricName":"# local misassemblies"},{"values":[0,0,0,0,0,0],"quality":"Less is better","isMain":false,"metricName":"# unaligned mis. contigs"}]],["Unaligned",[{"values":[0,0,0,0,0,0],"quality":"Less is better","isMain":false,"metricName":"# fully unaligned contigs"},{"values":[0,0,0,0,0,0],"quality":"Less is better","isMain":false,"metricName":"Fully unaligned length"},{"values":[0,0,0,0,0,0],"quality":"Less is better","isMain":false,"metricName":"# partially unaligned contigs"},{"values":[0,0,0,0,0,0],"quality":"Less is better","isMain":false,"metricName":"Partially unaligned length"}]],["Mismatches",[{"values":[0,0,2,0,0,1],"quality":"Less is better","isMain":false,"metricName":"# mismatches"},{"values":[0,0,0,0,0,1],"quality":"Less is better","isMain":false,"metricName":"# indels"},{"values":[0,0,0,0,0,1],"quality":"Less is better","isMain":false,"metricName":"Indels length"},{"values":["0.00","0.00","362.98","0.00","0.00","11.15"],"quality":"Less is better","isMain":true,"metricName":"# mismatches per 100 kbp"},{"values":["0.00","0.00","0.00","0.00","0.00","11.15"],"quality":"Less is better","isMain":true,"metricName":"# indels per 100 kbp"},{"values":[0,0,0,0,0,1],"quality":"Less is better","isMain":false,"metricName":" # indels (<= 5 bp)"},{"values":[0,0,0,0,0,0],"quality":"Less is better","isMain":false,"metricName":" # indels (> 5 bp)"},{"values":[0,0,0,0,0,0],"quality":"Less is better","isMain":false,"metricName":"# N's"},{"values":["0.00","0.00","0.00","0.00","0.00","0.00"],"quality":"Less is better","isMain":true,"metricName":"# N's per 100 kbp"}]],["Statistics without reference",[{"values":[1,2,1,3,7,9],"quality":"Equal","isMain":true,"metricName":"# contigs"},{"values":[0,2,0,0,0,3],"quality":"Equal","isMain":false,"metricName":"# contigs (>= 1000 bp)"},{"values":[0,0,0,0,0,0],"quality":"Equal","isMain":false,"metricName":"# contigs (>= 5000 bp)"},{"values":[0,0,0,0,0,0],"quality":"Equal","isMain":false,"metricName":"# contigs (>= 10000 bp)"},{"values":[0,0,0,0,0,0],"quality":"Equal","isMain":false,"metricName":"# contigs (>= 25000 bp)"},{"values":[0,0,0,0,0,0],"quality":"Equal","isMain":false,"metricName":"# contigs (>= 50000 bp)"},{"values":[520,2952,551,930,930,1732],"quality":"More is better","isMain":true,"metricName":"Largest contig"},{"values":[520,5828,551,2353,5139,9025],"quality":"More is better","isMain":true,"metricName":"Total length"},{"values":[0,5828,0,0,0,4325],"quality":"More is better","isMain":true,"metricName":"Total length (>= 1000 bp)"},{"values":[0,0,0,0,0,0],"quality":"More is better","isMain":false,"metricName":"Total length (>= 5000 bp)"},{"values":[0,0,0,0,0,0],"quality":"More is better","isMain":true,"metricName":"Total length (>= 10000 bp)"},{"values":[0,0,0,0,0,0],"quality":"More is better","isMain":false,"metricName":"Total length (>= 25000 bp)"},{"values":[0,0,0,0,0,0],"quality":"More is better","isMain":true,"metricName":"Total length (>= 50000 bp)"},{"values":[520,2952,551,915,822,896],"quality":"More is better","isMain":false,"metricName":"N50"},{"values":[520,2876,551,915,707,853],"quality":"More is better","isMain":false,"metricName":"N75"},{"values":[1,1,1,2,3,4],"quality":"Less is better","isMain":false,"metricName":"L50"},{"values":[1,2,1,2,5,6],"quality":"Less is better","isMain":false,"metricName":"L75"},{"values":["68.65","68.63","69.15","66.94","67.68","68.00"],"quality":"Equal","isMain":false,"metricName":"GC (%)"}]],["Predicted genes",[]],["Similarity statistics",[{"values":[0,0,0,0,0,0],"quality":"Equal","isMain":false,"metricName":"# similar correct contigs"},{"values":[0,0,0,0,0,0],"quality":"Equal","isMain":false,"metricName":"# similar misassembled blocks"}]],["Reference statistics",[{"values":[68090,68090,68090,68090,68090,68090],"quality":"Equal","isMain":false,"metricName":"Reference length"},{"values":[1,1,1,1,1,1],"quality":"Equal","isMain":false,"metricName":"Reference fragments"},{"values":["67.57","67.57","67.57","67.57","67.57","67.57"],"quality":"Equal","isMain":false,"metricName":"Reference GC (%)"}]]],"referenceName":"gi\_194303197\_ref\_NC","date":"18 July 2018, Wednesday, 16:43:52","order":[0,1,2,3,4,5],"assembliesNames":["ABySS\_127","IDBA\_UD","SPAdes","SPAdes\_sc","SPAdes\_sc\_careful","Velvet"]},{"assembliesWithNs":null,"minContig":500,"report":[["Genome statistics",[{"values":["39.605","21.999","48.994","30.868","41.721","78.895","17.284","1.985","39.155","7.873","32.958","32.775","13.003","94.901"],"quality":"More is better","isMain":true,"metricName":"Genome fraction (%)"},{"values":["1.021","1.007","1.008","1.012","1.035","1.185","1.023","1.000","1.003","1.000","1.030","1.030","1.003","1.448"],"quality":"Less is better","isMain":true,"metricName":"Duplication ratio"},{"values":[4243,4256,5990,4016,6373,9161,1220,650,7041,3920,5760,6084,1765,49566],"quality":"More is better","isMain":true,"metricName":"Largest alignment"},{"values":[30849,16912,37631,23827,32934,70863,13490,1515,29973,6008,25781,25641,9951,104527],"quality":"More is better","isMain":true,"metricName":"Total aligned length"},{"values":[null,null,null,null,null,2636,null,null,null,null,null,null,null,56961],"quality":"More is better","isMain":false,"metricName":"NG50"},{"values":[null,null,null,null,null,1312,null,null,null,null,null,null,null,20595],"quality":"More is better","isMain":false,"metricName":"NG75"},{"values":[1406,1471,3939,1193,1399,2835,922,null,2437,3920,3855,3087,887,18887],"quality":"More is better","isMain":false,"metricName":"NA50"},{"values":[787,727,1350,864,875,1894,697,null,1295,1481,1632,1774,690,9094],"quality":"More is better","isMain":false,"metricName":"NA75"},{"values":[null,null,null,null,null,2575,null,null,null,null,null,null,null,49566],"quality":"More is better","isMain":true,"metricName":"NGA50"},{"values":[null,null,null,null,null,1312,null,null,null,null,null,null,null,18887],"quality":"More is better","isMain":false,"metricName":"NGA75"},{"values":[null,null,null,null,null,9,null,null,null,null,null,null,null,1],"quality":"Less is better","isMain":false,"metricName":"LG50"},{"values":[null,null,null,null,null,18,null,null,null,null,null,null,null,2],"quality":"Less is better","isMain":false,"metricName":"LG75"},{"values":[7,4,4,6,7,8,7,null,4,1,3,3,4,2],"quality":"Less is better","isMain":false,"metricName":"LA50"},{"values":[14,8,9,11,14,16,11,null,8,2,6,6,7,4],"quality":"Less is better","isMain":false,"metricName":"LA75"},{"values":[null,null,null,null,null,9,null,null,null,null,null,null,null,1],"quality":"Less is better","isMain":true,"metricName":"LGA50"},{"values":[null,null,null,null,null,18,null,null,null,null,null,null,null,2],"quality":"Less is better","isMain":false,"metricName":"LGA75"}]],["Misassemblies",[{"values":[0,0,0,0,0,0,0,0,0,0,0,0,0,0],"quality":"Less is better","isMain":true,"metricName":"# misassemblies"},{"values":[0,0,0,0,0,0,0,0,0,0,0,0,0,0],"quality":"Less is better","isMain":false,"metricName":" # relocations"},{"values":[0,0,0,0,0,0,0,0,0,0,0,0,0,0],"quality":"Less is better","isMain":false,"metricName":" # translocations"},{"values":[0,0,0,0,0,0,0,0,0,0,0,0,0,0],"quality":"Less is better","isMain":false,"metricName":" # inversions"},{"values":[0,0,0,0,0,0,0,0,0,0,0,0,0,0],"quality":"Less is better","isMain":false,"metricName":"# misassembled contigs"},{"values":[0,0,0,0,0,0,0,0,0,0,0,0,0,0],"quality":"Less is better","isMain":true,"metricName":"Misassembled contigs length"},{"values":[0,0,0,0,0,1,0,0,0,0,1,1,0,17],"quality":"Less is better","isMain":false,"metricName":"# local misassemblies"},{"values":[0,0,0,0,0,0,0,0,0,0,0,0,0,0],"quality":"Less is better","isMain":false,"metricName":"# unaligned mis. contigs"}]],["Unaligned",[{"values":[0,0,0,0,0,0,0,0,0,0,0,0,0,0],"quality":"Less is better","isMain":false,"metricName":"# fully unaligned contigs"},{"values":[0,0,0,0,0,0,0,0,0,0,0,0,0,0],"quality":"Less is better","isMain":false,"metricName":"Fully unaligned length"},{"values":[0,0,0,0,0,0,0,1,0,0,0,0,0,4],"quality":"Less is better","isMain":false,"metricName":"# partially unaligned contigs"},{"values":[0,0,0,0,0,0,0,1645,0,0,0,0,0,11222],"quality":"Less is better","isMain":false,"metricName":"Partially unaligned length"}]],["Mismatches",[{"values":[12,1,28,0,7,136,1,0,22,5,167,122,2,320],"quality":"Less is better","isMain":false,"metricName":"# mismatches"},{"values":[0,0,0,0,0,20,0,0,0,0,9,9,0,30],"quality":"Less is better","isMain":false,"metricName":"# indels"},{"values":[0,0,0,0,0,52,0,0,0,0,14,14,0,58],"quality":"Less is better","isMain":false,"metricName":"Indels length"},{"values":["39.70","5.96","74.89","0.00","21.99","225.89","7.58","0.00","73.63","83.22","663.99","487.79","20.16","441.86"],"quality":"Less is better","isMain":true,"metricName":"# mismatches per 100 kbp"},{"values":["0.00","0.00","0.00","0.00","0.00","33.22","0.00","0.00","0.00","0.00","35.78","35.98","0.00","41.42"],"quality":"Less is better","isMain":true,"metricName":"# indels per 100 kbp"},{"values":[0,0,0,0,0,18,0,0,0,0,9,9,0,29],"quality":"Less is better","isMain":false,"metricName":" # indels (<= 5 bp)"},{"values":[0,0,0,0,0,2,0,0,0,0,0,0,0,1],"quality":"Less is better","isMain":false,"metricName":" # indels (> 5 bp)"},{"values":[0,0,0,0,0,10,0,448,0,0,0,0,0,502],"quality":"Less is better","isMain":false,"metricName":"# N's"},{"values":["0.00","0.00","0.00","0.00","0.00","14.02","0.00","14177.22","0.00","0.00","0.00","0.00","0.00","432.56"],"quality":"Less is better","isMain":true,"metricName":"# N's per 100 kbp"}]],["Statistics without reference",[{"values":[26,15,19,20,26,35,16,3,18,3,12,12,11,7],"quality":"Equal","isMain":true,"metricName":"# contigs"},{"values":[10,6,13,10,11,22,3,1,9,2,9,9,3,7],"quality":"Equal","isMain":false,"metricName":"# contigs (>= 1000 bp)"},{"values":[0,0,2,0,1,1,0,0,1,0,1,1,0,5],"quality":"Equal","isMain":false,"metricName":"# contigs (>= 5000 bp)"},{"values":[0,0,0,0,0,0,0,0,0,0,0,0,0,3],"quality":"Equal","isMain":false,"metricName":"# contigs (>= 10000 bp)"},{"values":[0,0,0,0,0,0,0,0,0,0,0,0,0,1],"quality":"Equal","isMain":false,"metricName":"# contigs (>= 25000 bp)"},{"values":[0,0,0,0,0,0,0,0,0,0,0,0,0,1],"quality":"Equal","isMain":false,"metricName":"# contigs (>= 50000 bp)"},{"values":[4243,4256,5990,4016,6373,9161,1220,1931,7041,3920,5760,6084,1765,56961],"quality":"More is better","isMain":true,"metricName":"Largest contig"},{"values":[30869,16912,37689,23827,32937,71345,13490,3160,29973,6010,25899,25759,9951,116053],"quality":"More is better","isMain":true,"metricName":"Total length"},{"values":[20453,11277,33784,17026,22256,62123,3520,1931,24047,5401,24145,23983,4440,116053],"quality":"More is better","isMain":true,"metricName":"Total length (>= 1000 bp)"},{"values":[0,0,11271,0,6373,9161,0,0,7041,0,5760,6084,0,108219],"quality":"More is better","isMain":false,"metricName":"Total length (>= 5000 bp)"},{"values":[0,0,0,0,0,0,0,0,0,0,0,0,0,90327],"quality":"More is better","isMain":true,"metricName":"Total length (>= 10000 bp)"},{"values":[0,0,0,0,0,0,0,0,0,0,0,0,0,56961],"quality":"More is better","isMain":false,"metricName":"Total length (>= 25000 bp)"},{"values":[0,0,0,0,0,0,0,0,0,0,0,0,0,56961],"quality":"More is better","isMain":true,"metricName":"Total length (>= 50000 bp)"},{"values":[1406,1471,3939,1193,1399,2835,922,1931,2437,3920,3855,3087,887,20595],"quality":"More is better","isMain":false,"metricName":"N50"},{"values":[787,727,1350,864,875,1946,697,650,1295,1481,1632,1852,690,12771],"quality":"More is better","isMain":false,"metricName":"N75"},{"values":[7,4,4,6,7,8,7,1,4,1,3,3,4,2],"quality":"Less is better","isMain":false,"metricName":"L50"},{"values":[14,8,9,11,14,16,11,2,8,2,6,6,7,3],"quality":"Less is better","isMain":false,"metricName":"L75"},{"values":["62.17","62.15","62.82","62.27","62.44","62.63","61.95","58.00","62.55","64.69","62.31","62.35","61.98","63.01"],"quality":"Equal","isMain":false,"metricName":"GC (%)"}]],["Predicted genes",[]],["Similarity statistics",[{"values":[0,0,0,0,0,0,0,0,0,0,0,0,0,0],"quality":"Equal","isMain":false,"metricName":"# similar correct contigs"},{"values":[0,0,0,0,0,0,0,0,0,0,0,0,0,0],"quality":"Equal","isMain":false,"metricName":"# similar misassembled blocks"}]],["Reference statistics",[{"values":[76312,76312,76312,76312,76312,76312,76312,76312,76312,76312,76312,76312,76312,76312],"quality":"Equal","isMain":false,"metricName":"Reference length"},{"values":[1,1,1,1,1,1,1,1,1,1,1,1,1,1],"quality":"Equal","isMain":false,"metricName":"Reference fragments"},{"values":["62.84","62.84","62.84","62.84","62.84","62.84","62.84","62.84","62.84","62.84","62.84","62.84","62.84","62.84"],"quality":"Equal","isMain":false,"metricName":"Reference GC (%)"}]]],"referenceName":"gi\_194303296\_ref\_NC","date":"18 July 2018, Wednesday, 16:44:08","order":[0,1,2,3,4,5,6,7,8,9,10,11,12,13],"assembliesNames":["ABySS\_127","ABySS\_63","CLC","IDBA\_UD","MEGAHIT","MIRA","Ray\_Meta","SOAPdenovo2","SPAdes","SPAdes\_meta","SPAdes\_sc","SPAdes\_sc\_careful","Velvet","Geneious"]},{"assembliesWithNs":null,"minContig":500,"report":[["Genome statistics",[{"values":["41.776","23.696","59.721","69.836","71.061","62.793","59.357","75.581","55.662","72.560","76.299","39.257","5.012"],"quality":"More is better","isMain":true,"metricName":"Genome fraction (%)"},{"values":["1.000","1.000","1.007","1.000","1.002","1.038","1.003","1.000","1.003","1.002","1.004","1.000","1.000"],"quality":"Less is better","isMain":true,"metricName":"Duplication ratio"},{"values":[1441,1322,4955,8129,4951,4524,4953,8129,8129,8129,8129,8033,1858],"quality":"More is better","isMain":true,"metricName":"Largest alignment"},{"values":[15488,8785,22282,25900,26396,24160,22068,28030,20645,26931,28372,14554,1858],"quality":"More is better","isMain":true,"metricName":"Total aligned length"},{"values":[null,null,712,1619,1550,939,1297,2004,1619,4445,4445,null,null],"quality":"More is better","isMain":false,"metricName":"NG50"},{"values":[null,null,null,null,null,null,null,753,null,1137,1310,null,null],"quality":"More is better","isMain":false,"metricName":"NG75"},{"values":[963,808,2004,2156,1769,1493,1652,2983,4445,1892,1802,8033,1858],"quality":"More is better","isMain":false,"metricName":"NA50"},{"values":[647,691,775,1619,1265,1104,1004,1819,1810,775,836,1434,1858],"quality":"More is better","isMain":false,"metricName":"NA75"},{"values":[null,null,712,1619,1550,939,762,2004,1619,1619,1619,null,null],"quality":"More is better","isMain":true,"metricName":"NGA50"},{"values":[null,null,null,null,null,null,null,753,null,null,660,null,null],"quality":"More is better","isMain":false,"metricName":"NGA75"},{"values":[null,null,10,6,8,11,8,5,6,3,3,null,null],"quality":"Less is better","isMain":false,"metricName":"LG50"},{"values":[null,null,null,null,null,null,null,11,null,8,8,null,null],"quality":"Less is better","isMain":false,"metricName":"LG75"},{"values":[7,4,4,3,5,5,5,3,2,4,5,1,1],"quality":"Less is better","isMain":false,"metricName":"LA50"},{"values":[12,7,8,6,9,10,9,6,4,11,11,3,1],"quality":"Less is better","isMain":false,"metricName":"LA75"},{"values":[null,null,10,6,8,11,10,5,6,6,6,null,null],"quality":"Less is better","isMain":true,"metricName":"LGA50"},{"values":[null,null,null,null,null,null,null,11,null,null,14,null,null],"quality":"Less is better","isMain":false,"metricName":"LGA75"}]],["Misassemblies",[{"values":[0,0,0,0,0,0,0,0,0,0,0,0,0],"quality":"Less is better","isMain":true,"metricName":"# misassemblies"},{"values":[0,0,0,0,0,0,0,0,0,0,0,0,0],"quality":"Less is better","isMain":false,"metricName":" # relocations"},{"values":[0,0,0,0,0,0,0,0,0,0,0,0,0],"quality":"Less is better","isMain":false,"metricName":" # translocations"},{"values":[0,0,0,0,0,0,0,0,0,0,0,0,0],"quality":"Less is better","isMain":false,"metricName":" # inversions"},{"values":[0,0,0,0,0,0,0,0,0,0,0,0,0],"quality":"Less is better","isMain":false,"metricName":"# misassembled contigs"},{"values":[0,0,0,0,0,0,0,0,0,0,0,0,0],"quality":"Less is better","isMain":true,"metricName":"Misassembled contigs length"},{"values":[0,0,0,0,0,0,0,0,0,0,0,0,0],"quality":"Less is better","isMain":false,"metricName":"# local misassemblies"},{"values":[0,0,0,0,0,0,1,0,0,0,0,0,0],"quality":"Less is better","isMain":false,"metricName":"# unaligned mis. contigs"}]],["Unaligned",[{"values":[0,0,0,0,0,0,0,0,0,0,0,0,0],"quality":"Less is better","isMain":false,"metricName":"# fully unaligned contigs"},{"values":[0,0,0,0,0,0,0,0,0,0,0,0,0],"quality":"Less is better","isMain":false,"metricName":"Fully unaligned length"},{"values":[0,0,0,0,0,0,1,0,0,1,1,0,0],"quality":"Less is better","isMain":false,"metricName":"# partially unaligned contigs"},{"values":[0,0,0,0,0,0,2268,0,0,5469,5469,0,0],"quality":"Less is better","isMain":false,"metricName":"Partially unaligned length"}]],["Mismatches",[{"values":[3,2,12,10,7,2,17,34,6,27,25,6,0],"quality":"Less is better","isMain":false,"metricName":"# mismatches"},{"values":[0,0,0,0,0,0,0,4,0,4,6,0,0],"quality":"Less is better","isMain":false,"metricName":"# indels"},{"values":[0,0,0,0,0,0,0,4,0,4,6,0,0],"quality":"Less is better","isMain":false,"metricName":"Indels length"},{"values":["19.37","22.77","54.20","38.62","26.57","8.59","77.25","121.34","29.08","100.37","88.38","41.23","0.00"],"quality":"Less is better","isMain":true,"metricName":"# mismatches per 100 kbp"},{"values":["0.00","0.00","0.00","0.00","0.00","0.00","0.00","14.28","0.00","14.87","21.21","0.00","0.00"],"quality":"Less is better","isMain":true,"metricName":"# indels per 100 kbp"},{"values":[0,0,0,0,0,0,0,4,0,4,6,0,0],"quality":"Less is better","isMain":false,"metricName":" # indels (<= 5 bp)"},{"values":[0,0,0,0,0,0,0,0,0,0,0,0,0],"quality":"Less is better","isMain":false,"metricName":" # indels (> 5 bp)"},{"values":[0,0,0,0,0,24,668,0,0,0,0,0,3],"quality":"Less is better","isMain":false,"metricName":"# N's"},{"values":["0.00","0.00","0.00","0.00","0.00","99.34","2744.12","0.00","0.00","0.00","0.00","0.00","161.46"],"quality":"Less is better","isMain":true,"metricName":"# N's per 100 kbp"}]],["Statistics without reference",[{"values":[18,10,15,13,16,18,14,11,7,14,15,6,1],"quality":"Equal","isMain":true,"metricName":"# contigs"},{"values":[5,3,7,8,11,10,10,9,6,9,10,5,1],"quality":"Equal","isMain":false,"metricName":"# contigs (>= 1000 bp)"},{"values":[0,0,0,1,0,0,0,1,1,2,2,1,0],"quality":"Equal","isMain":false,"metricName":"# contigs (>= 5000 bp)"},{"values":[0,0,0,0,0,0,0,0,0,0,0,0,0],"quality":"Equal","isMain":false,"metricName":"# contigs (>= 10000 bp)"},{"values":[0,0,0,0,0,0,0,0,0,0,0,0,0],"quality":"Equal","isMain":false,"metricName":"# contigs (>= 25000 bp)"},{"values":[0,0,0,0,0,0,0,0,0,0,0,0,0],"quality":"Equal","isMain":false,"metricName":"# contigs (>= 50000 bp)"},{"values":[1441,1322,4955,8129,4951,4524,4953,8129,8129,8129,8129,8033,1858],"quality":"More is better","isMain":true,"metricName":"Largest contig"},{"values":[15488,8785,22288,25900,26396,24160,24343,28030,20699,32422,33863,14554,1858],"quality":"More is better","isMain":true,"metricName":"Total length"},{"values":[6327,3908,16695,22295,22772,18434,21569,26441,19863,28855,30296,13696,1858],"quality":"More is better","isMain":true,"metricName":"Total length (>= 1000 bp)"},{"values":[0,0,0,8129,0,0,0,8129,8129,15490,15490,8033,0],"quality":"More is better","isMain":false,"metricName":"Total length (>= 5000 bp)"},{"values":[0,0,0,0,0,0,0,0,0,0,0,0,0],"quality":"More is better","isMain":true,"metricName":"Total length (>= 10000 bp)"},{"values":[0,0,0,0,0,0,0,0,0,0,0,0,0],"quality":"More is better","isMain":false,"metricName":"Total length (>= 25000 bp)"},{"values":[0,0,0,0,0,0,0,0,0,0,0,0,0],"quality":"More is better","isMain":true,"metricName":"Total length (>= 50000 bp)"},{"values":[963,808,2004,2156,1769,1493,2350,2983,4445,4445,4445,8033,1858],"quality":"More is better","isMain":false,"metricName":"N50"},{"values":[647,691,775,1619,1265,1104,1297,1819,1864,1619,1441,1434,1858],"quality":"More is better","isMain":false,"metricName":"N75"},{"values":[7,4,4,3,5,5,4,3,2,3,3,1,1],"quality":"Less is better","isMain":false,"metricName":"L50"},{"values":[12,7,8,6,9,10,8,6,4,6,7,3,1],"quality":"Less is better","isMain":false,"metricName":"L75"},{"values":["51.78","50.24","52.15","51.89","51.93","51.88","50.97","51.73","52.07","49.48","49.43","52.62","51.91"],"quality":"Equal","isMain":false,"metricName":"GC (%)"}]],["Predicted genes",[]],["Similarity statistics",[{"values":[0,0,0,0,0,0,0,0,0,0,0,0,0],"quality":"Equal","isMain":false,"metricName":"# similar correct contigs"},{"values":[0,0,0,0,0,0,0,0,0,0,0,0,0],"quality":"Equal","isMain":false,"metricName":"# similar misassembled blocks"}]],["Reference statistics",[{"values":[37074,37074,37074,37074,37074,37074,37074,37074,37074,37074,37074,37074,37074],"quality":"Equal","isMain":false,"metricName":"Reference length"},{"values":[1,1,1,1,1,1,1,1,1,1,1,1,1],"quality":"Equal","isMain":false,"metricName":"Reference fragments"},{"values":["50.77","50.77","50.77","50.77","50.77","50.77","50.77","50.77","50.77","50.77","50.77","50.77","50.77"],"quality":"Equal","isMain":false,"metricName":"Reference GC (%)"}]]],"referenceName":"gi\_19548987\_ref\_NC","date":"18 July 2018, Wednesday, 16:44:22","order":[0,1,2,3,4,5,6,7,8,9,10,11,12],"assembliesNames":["ABySS\_127","ABySS\_63","CLC","IDBA\_UD","MEGAHIT","MIRA","SOAPdenovo2","SPAdes","SPAdes\_meta","SPAdes\_sc","SPAdes\_sc\_careful","Velvet","Geneious"]},{"assembliesWithNs":null,"minContig":500,"report":[["Genome statistics",[{"values":["100.000","100.000","100.000","100.000","85.449","55.661","100.000","100.000","100.000","100.000","100.000"],"quality":"More is better","isMain":true,"metricName":"Genome fraction (%)"},{"values":["1.000","1.002","1.002","1.007","1.008","1.001","1.002","1.001","1.001","1.001","1.000"],"quality":"Less is better","isMain":true,"metricName":"Duplication ratio"},{"values":[60195,60294,60336,60610,15548,1816,60322,60250,60250,60250,60222],"quality":"More is better","isMain":true,"metricName":"Largest alignment"},{"values":[60195,60294,60336,60610,51840,33523,60322,60250,60250,60250,60222],"quality":"More is better","isMain":true,"metricName":"Total aligned length"},{"values":[60195,60294,60336,60610,9575,560,60322,60250,60250,60250,60222],"quality":"More is better","isMain":false,"metricName":"NG50"},{"values":[60195,60294,60336,60610,6168,null,60322,60250,60250,60250,60222],"quality":"More is better","isMain":false,"metricName":"NG75"},{"values":[60195,60294,60336,60610,12459,787,60322,60250,60250,60250,60222],"quality":"More is better","isMain":false,"metricName":"NA50"},{"values":[60195,60294,60336,60610,7493,645,60322,60250,60250,60250,60222],"quality":"More is better","isMain":false,"metricName":"NA75"},{"values":[60195,60294,60336,60610,9575,560,60322,60250,60250,60250,60222],"quality":"More is better","isMain":true,"metricName":"NGA50"},{"values":[60195,60294,60336,60610,6168,null,60322,60250,60250,60250,60222],"quality":"More is better","isMain":false,"metricName":"NGA75"},{"values":[1,1,1,1,3,35,1,1,1,1,1],"quality":"Less is better","isMain":false,"metricName":"LG50"},{"values":[1,1,1,1,5,null,1,1,1,1,1],"quality":"Less is better","isMain":false,"metricName":"LG75"},{"values":[1,1,1,1,2,15,1,1,1,1,1],"quality":"Less is better","isMain":false,"metricName":"LA50"},{"values":[1,1,1,1,4,27,1,1,1,1,1],"quality":"Less is better","isMain":false,"metricName":"LA75"},{"values":[1,1,1,1,3,35,1,1,1,1,1],"quality":"Less is better","isMain":true,"metricName":"LGA50"},{"values":[1,1,1,1,5,null,1,1,1,1,1],"quality":"Less is better","isMain":false,"metricName":"LGA75"}]],["Misassemblies",[{"values":[0,0,0,0,0,0,0,0,0,0,0],"quality":"Less is better","isMain":true,"metricName":"# misassemblies"},{"values":[0,0,0,0,0,0,0,0,0,0,0],"quality":"Less is better","isMain":false,"metricName":" # relocations"},{"values":[0,0,0,0,0,0,0,0,0,0,0],"quality":"Less is better","isMain":false,"metricName":" # translocations"},{"values":[0,0,0,0,0,0,0,0,0,0,0],"quality":"Less is better","isMain":false,"metricName":" # inversions"},{"values":[0,0,0,0,0,0,0,0,0,0,0],"quality":"Less is better","isMain":false,"metricName":"# misassembled contigs"},{"values":[0,0,0,0,0,0,0,0,0,0,0],"quality":"Less is better","isMain":true,"metricName":"Misassembled contigs length"},{"values":[0,1,1,1,0,0,1,0,0,0,0],"quality":"Less is better","isMain":false,"metricName":"# local misassemblies"},{"values":[0,0,0,0,0,0,0,0,0,0,0],"quality":"Less is better","isMain":false,"metricName":"# unaligned mis. contigs"}]],["Unaligned",[{"values":[0,0,0,0,0,0,0,0,0,0,0],"quality":"Less is better","isMain":false,"metricName":"# fully unaligned contigs"},{"values":[0,0,0,0,0,0,0,0,0,0,0],"quality":"Less is better","isMain":false,"metricName":"Fully unaligned length"},{"values":[0,0,0,0,0,0,0,0,0,0,0],"quality":"Less is better","isMain":false,"metricName":"# partially unaligned contigs"},{"values":[0,0,0,0,0,0,0,0,0,0,0],"quality":"Less is better","isMain":false,"metricName":"Partially unaligned length"}]],["Mismatches",[{"values":[0,0,0,0,0,0,0,0,0,0,0],"quality":"Less is better","isMain":false,"metricName":"# mismatches"},{"values":[0,0,0,0,0,0,0,1,1,1,1],"quality":"Less is better","isMain":false,"metricName":"# indels"},{"values":[0,0,0,0,0,0,0,55,55,55,27],"quality":"Less is better","isMain":false,"metricName":"Indels length"},{"values":["0.00","0.00","0.00","0.00","0.00","0.00","0.00","0.00","0.00","0.00","0.00"],"quality":"Less is better","isMain":true,"metricName":"# mismatches per 100 kbp"},{"values":["0.00","0.00","0.00","0.00","0.00","0.00","0.00","1.66","1.66","1.66","1.66"],"quality":"Less is better","isMain":true,"metricName":"# indels per 100 kbp"},{"values":[0,0,0,0,0,0,0,0,0,0,0],"quality":"Less is better","isMain":false,"metricName":" # indels (<= 5 bp)"},{"values":[0,0,0,0,0,0,0,1,1,1,1],"quality":"Less is better","isMain":false,"metricName":" # indels (> 5 bp)"},{"values":[0,0,0,5,0,0,0,0,0,0,0],"quality":"Less is better","isMain":false,"metricName":"# N's"},{"values":["0.00","0.00","0.00","8.25","0.00","0.00","0.00","0.00","0.00","0.00","0.00"],"quality":"Less is better","isMain":true,"metricName":"# N's per 100 kbp"}]],["Statistics without reference",[{"values":[1,1,1,1,6,41,1,1,1,1,1],"quality":"Equal","isMain":true,"metricName":"# contigs"},{"values":[1,1,1,1,5,10,1,1,1,1,1],"quality":"Equal","isMain":false,"metricName":"# contigs (>= 1000 bp)"},{"values":[1,1,1,1,5,0,1,1,1,1,1],"quality":"Equal","isMain":false,"metricName":"# contigs (>= 5000 bp)"},{"values":[1,1,1,1,2,0,1,1,1,1,1],"quality":"Equal","isMain":false,"metricName":"# contigs (>= 10000 bp)"},{"values":[1,1,1,1,0,0,1,1,1,1,1],"quality":"Equal","isMain":false,"metricName":"# contigs (>= 25000 bp)"},{"values":[1,1,1,1,0,0,1,1,1,1,1],"quality":"Equal","isMain":false,"metricName":"# contigs (>= 50000 bp)"},{"values":[60195,60294,60336,60610,15548,1816,60322,60250,60250,60250,60222],"quality":"More is better","isMain":true,"metricName":"Largest contig"},{"values":[60195,60294,60336,60610,51840,33523,60322,60250,60250,60250,60222],"quality":"More is better","isMain":true,"metricName":"Total length"},{"values":[60195,60294,60336,60610,51243,13137,60322,60250,60250,60250,60222],"quality":"More is better","isMain":true,"metricName":"Total length (>= 1000 bp)"},{"values":[60195,60294,60336,60610,51243,0,60322,60250,60250,60250,60222],"quality":"More is better","isMain":false,"metricName":"Total length (>= 5000 bp)"},{"values":[60195,60294,60336,60610,28007,0,60322,60250,60250,60250,60222],"quality":"More is better","isMain":true,"metricName":"Total length (>= 10000 bp)"},{"values":[60195,60294,60336,60610,0,0,60322,60250,60250,60250,60222],"quality":"More is better","isMain":false,"metricName":"Total length (>= 25000 bp)"},{"values":[60195,60294,60336,60610,0,0,60322,60250,60250,60250,60222],"quality":"More is better","isMain":true,"metricName":"Total length (>= 50000 bp)"},{"values":[60195,60294,60336,60610,12459,787,60322,60250,60250,60250,60222],"quality":"More is better","isMain":false,"metricName":"N50"},{"values":[60195,60294,60336,60610,7493,645,60322,60250,60250,60250,60222],"quality":"More is better","isMain":false,"metricName":"N75"},{"values":[1,1,1,1,2,15,1,1,1,1,1],"quality":"Less is better","isMain":false,"metricName":"L50"},{"values":[1,1,1,1,4,27,1,1,1,1,1],"quality":"Less is better","isMain":false,"metricName":"L75"},{"values":["58.95","58.96","58.97","58.98","59.07","59.26","58.94","58.96","58.95","58.95","58.95"],"quality":"Equal","isMain":false,"metricName":"GC (%)"}]],["Predicted genes",[]],["Similarity statistics",[{"values":[0,0,0,0,0,0,0,0,0,0,0],"quality":"Equal","isMain":false,"metricName":"# similar correct contigs"},{"values":[0,0,0,0,0,0,0,0,0,0,0],"quality":"Equal","isMain":false,"metricName":"# similar misassembled blocks"}]],["Reference statistics",[{"values":[60195,60195,60195,60195,60195,60195,60195,60195,60195,60195,60195],"quality":"Equal","isMain":false,"metricName":"Reference length"},{"values":[1,1,1,1,1,1,1,1,1,1,1],"quality":"Equal","isMain":false,"metricName":"Reference fragments"},{"values":["58.95","58.95","58.95","58.95","58.95","58.95","58.95","58.95","58.95","58.95","58.95"],"quality":"Equal","isMain":false,"metricName":"Reference GC (%)"}]]],"referenceName":"gi\_195546530\_ref\_NC","date":"18 July 2018, Wednesday, 16:44:35","order":[0,1,2,3,4,5,6,7,8,9,10],"assembliesNames":["CLC","IDBA\_UD","MEGAHIT","MIRA","Ray\_Meta","SOAPdenovo2","SPAdes","SPAdes\_meta","SPAdes\_sc","SPAdes\_sc\_careful","Velvet"]},{"assembliesWithNs":null,"minContig":500,"report":[["Genome statistics",[{"values":["5.072","1.194","40.809","42.292","51.405","28.181","21.601","55.278","58.004","56.223","53.102","6.390","3.422"],"quality":"More is better","isMain":true,"metricName":"Genome fraction (%)"},{"values":["1.028","1.002","1.041","1.006","1.001","1.051","1.000","1.006","1.007","1.017","1.016","1.000","1.000"],"quality":"Less is better","isMain":true,"metricName":"Duplication ratio"},{"values":[603,514,2549,2966,3419,1559,1715,3600,2766,2698,2698,1137,1470],"quality":"More is better","isMain":true,"metricName":"Largest alignment"},{"values":[2241,514,18251,18268,22110,12727,9280,23797,25025,24425,23029,2745,1470],"quality":"More is better","isMain":true,"metricName":"Total aligned length"},{"values":[null,null,null,null,535,null,null,654,714,709,543,null,null],"quality":"More is better","isMain":false,"metricName":"NG50"},{"values":[556,514,1102,1085,1144,1058,848,1518,1102,1102,1047,886,1470],"quality":"More is better","isMain":false,"metricName":"NA50"},{"values":[550,514,728,900,848,778,642,918,841,654,730,722,1470],"quality":"More is better","isMain":false,"metricName":"NA75"},{"values":[null,null,null,null,535,null,null,654,709,589,535,null,null],"quality":"More is better","isMain":true,"metricName":"NGA50"},{"values":[null,null,null,null,19,null,null,14,17,16,20,null,null],"quality":"Less is better","isMain":false,"metricName":"LG50"},{"values":[2,1,6,6,7,5,4,5,8,9,8,2,1],"quality":"Less is better","isMain":false,"metricName":"LA50"},{"values":[3,1,11,11,12,9,7,10,14,17,14,3,1],"quality":"Less is better","isMain":false,"metricName":"LA75"},{"values":[null,null,null,null,19,null,null,14,18,19,21,null,null],"quality":"Less is better","isMain":true,"metricName":"LGA50"}]],["Misassemblies",[{"values":[0,0,0,0,0,0,0,0,0,0,0,0,0],"quality":"Less is better","isMain":true,"metricName":"# misassemblies"},{"values":[0,0,0,0,0,0,0,0,0,0,0,0,0],"quality":"Less is better","isMain":false,"metricName":" # relocations"},{"values":[0,0,0,0,0,0,0,0,0,0,0,0,0],"quality":"Less is better","isMain":false,"metricName":" # translocations"},{"values":[0,0,0,0,0,0,0,0,0,0,0,0,0],"quality":"Less is better","isMain":false,"metricName":" # inversions"},{"values":[0,0,0,0,0,0,0,0,0,0,0,0,0],"quality":"Less is better","isMain":false,"metricName":"# misassembled contigs"},{"values":[0,0,0,0,0,0,0,0,0,0,0,0,0],"quality":"Less is better","isMain":true,"metricName":"Misassembled contigs length"},{"values":[1,0,0,0,1,0,0,0,0,0,0,0,0],"quality":"Less is better","isMain":false,"metricName":"# local misassemblies"},{"values":[0,0,0,0,0,0,0,0,0,0,0,0,0],"quality":"Less is better","isMain":false,"metricName":"# unaligned mis. contigs"}]],["Unaligned",[{"values":[0,0,0,0,0,0,0,0,0,0,0,0,0],"quality":"Less is better","isMain":false,"metricName":"# fully unaligned contigs"},{"values":[0,0,0,0,0,0,0,0,0,0,0,0,0],"quality":"Less is better","isMain":false,"metricName":"Fully unaligned length"},{"values":[0,0,0,1,0,0,0,0,0,1,0,0,0],"quality":"Less is better","isMain":false,"metricName":"# partially unaligned contigs"},{"values":[0,0,0,2334,0,0,0,0,0,2063,0,0,0],"quality":"Less is better","isMain":false,"metricName":"Partially unaligned length"}]],["Mismatches",[{"values":[1,0,249,171,113,20,76,284,156,297,246,3,0],"quality":"Less is better","isMain":false,"metricName":"# mismatches"},{"values":[0,1,2,2,4,0,0,3,0,0,0,0,0],"quality":"Less is better","isMain":false,"metricName":"# indels"},{"values":[0,1,2,2,6,0,0,4,0,0,0,0,0],"quality":"Less is better","isMain":false,"metricName":"Indels length"},{"values":["45.89","0.00","1420.26","941.16","511.68","165.19","818.97","1195.89","626.03","1229.61","1078.33","109.29","0.00"],"quality":"Less is better","isMain":true,"metricName":"# mismatches per 100 kbp"},{"values":["0.00","194.93","11.41","11.01","18.11","0.00","0.00","12.63","0.00","0.00","0.00","0.00","0.00"],"quality":"Less is better","isMain":true,"metricName":"# indels per 100 kbp"},{"values":[0,1,2,2,4,0,0,3,0,0,0,0,0],"quality":"Less is better","isMain":false,"metricName":" # indels (<= 5 bp)"},{"values":[0,0,0,0,0,0,0,0,0,0,0,0,0],"quality":"Less is better","isMain":false,"metricName":" # indels (> 5 bp)"},{"values":[0,0,0,0,0,65,0,0,0,304,0,0,40],"quality":"Less is better","isMain":false,"metricName":"# N's"},{"values":["0.00","0.00","0.00","0.00","0.00","510.73","0.00","0.00","0.00","1141.27","0.00","0.00","2721.09"],"quality":"Less is better","isMain":true,"metricName":"# N's per 100 kbp"}]],["Statistics without reference",[{"values":[4,1,18,15,20,13,11,18,23,24,23,3,1],"quality":"Equal","isMain":true,"metricName":"# contigs"},{"values":[0,0,7,8,10,6,3,9,11,11,10,1,1],"quality":"Equal","isMain":false,"metricName":"# contigs (>= 1000 bp)"},{"values":[0,0,0,0,0,0,0,0,0,0,0,0,0],"quality":"Equal","isMain":false,"metricName":"# contigs (>= 5000 bp)"},{"values":[0,0,0,0,0,0,0,0,0,0,0,0,0],"quality":"Equal","isMain":false,"metricName":"# contigs (>= 10000 bp)"},{"values":[0,0,0,0,0,0,0,0,0,0,0,0,0],"quality":"Equal","isMain":false,"metricName":"# contigs (>= 25000 bp)"},{"values":[0,0,0,0,0,0,0,0,0,0,0,0,0],"quality":"Equal","isMain":false,"metricName":"# contigs (>= 50000 bp)"},{"values":[603,514,2549,3247,3419,1559,1715,3600,2766,3225,2784,1137,1470],"quality":"More is better","isMain":true,"metricName":"Largest contig"},{"values":[2241,514,18251,20610,22110,12727,9280,23890,25088,26637,23180,2745,1470],"quality":"More is better","isMain":true,"metricName":"Total length"},{"values":[0,0,10476,15062,15016,7837,3939,17742,16650,17869,14455,1137,1470],"quality":"More is better","isMain":true,"metricName":"Total length (>= 1000 bp)"},{"values":[0,0,0,0,0,0,0,0,0,0,0,0,0],"quality":"More is better","isMain":false,"metricName":"Total length (>= 5000 bp)"},{"values":[0,0,0,0,0,0,0,0,0,0,0,0,0],"quality":"More is better","isMain":true,"metricName":"Total length (>= 10000 bp)"},{"values":[0,0,0,0,0,0,0,0,0,0,0,0,0],"quality":"More is better","isMain":false,"metricName":"Total length (>= 25000 bp)"},{"values":[0,0,0,0,0,0,0,0,0,0,0,0,0],"quality":"More is better","isMain":true,"metricName":"Total length (>= 50000 bp)"},{"values":[556,514,1102,1622,1144,1058,848,1526,1102,1193,1047,886,1470],"quality":"More is better","isMain":false,"metricName":"N50"},{"values":[550,514,728,949,848,778,642,918,841,841,730,722,1470],"quality":"More is better","isMain":false,"metricName":"N75"},{"values":[2,1,6,5,7,5,4,5,8,7,8,2,1],"quality":"Less is better","isMain":false,"metricName":"L50"},{"values":[3,1,11,9,12,9,7,10,14,14,14,3,1],"quality":"Less is better","isMain":false,"metricName":"L75"},{"values":["64.39","67.12","62.56","62.29","62.42","62.48","61.83","62.47","62.39","62.43","62.51","61.49","62.45"],"quality":"Equal","isMain":false,"metricName":"GC (%)"}]],["Predicted genes",[]],["Similarity statistics",[{"values":[0,0,0,0,0,0,0,0,0,0,0,0,0],"quality":"Equal","isMain":false,"metricName":"# similar correct contigs"},{"values":[0,0,0,0,0,0,0,0,0,0,0,0,0],"quality":"Equal","isMain":false,"metricName":"# similar misassembled blocks"}]],["Reference statistics",[{"values":[42961,42961,42961,42961,42961,42961,42961,42961,42961,42961,42961,42961,42961],"quality":"Equal","isMain":false,"metricName":"Reference length"},{"values":[1,1,1,1,1,1,1,1,1,1,1,1,1],"quality":"Equal","isMain":false,"metricName":"Reference fragments"},{"values":["62.18","62.18","62.18","62.18","62.18","62.18","62.18","62.18","62.18","62.18","62.18","62.18","62.18"],"quality":"Equal","isMain":false,"metricName":"Reference GC (%)"}]]],"referenceName":"gi\_195546701\_ref\_NC","date":"18 July 2018, Wednesday, 16:44:50","order":[0,1,2,3,4,5,6,7,8,9,10,11,12],"assembliesNames":["ABySS\_127","ABySS\_63","CLC","IDBA\_UD","MEGAHIT","MIRA","SOAPdenovo2","SPAdes","SPAdes\_meta","SPAdes\_sc","SPAdes\_sc\_careful","Velvet","Geneious"]},{"assembliesWithNs":null,"minContig":500,"report":[["Genome statistics",[{"values":["99.997","87.131","99.325","95.581","96.999","100.000","100.000","100.000","100.000","92.786","95.139","44.322","100.000"],"quality":"More is better","isMain":true,"metricName":"Genome fraction (%)"},{"values":["1.075","1.016","1.000","1.046","1.025","1.341","1.028","1.003","1.543","1.013","1.010","1.001","1.213"],"quality":"Less is better","isMain":true,"metricName":"Duplication ratio"},{"values":[14448,6705,39719,6396,11374,40883,27316,40116,38987,7633,14540,2877,40883],"quality":"More is better","isMain":true,"metricName":"Largest alignment"},{"values":[42979,35404,39719,39989,39756,53607,41101,40116,61713,37599,38430,17747,48504],"quality":"More is better","isMain":true,"metricName":"Total aligned length"},{"values":[61019,5176,39719,2063,4455,84086,41101,40116,42293,6308,7964,null,84086],"quality":"More is better","isMain":false,"metricName":"NG50"},{"values":[61019,2608,39719,944,1825,84086,41101,40116,42293,1476,5260,null,84086],"quality":"More is better","isMain":false,"metricName":"NG75"},{"values":[11714,5719,39719,2063,4455,2562,27316,40116,1266,6308,7964,995,null],"quality":"More is better","isMain":false,"metricName":"NA50"},{"values":[null,2608,39719,944,3301,null,13785,40116,null,2728,5260,716,null],"quality":"More is better","isMain":false,"metricName":"NA75"},{"values":[11915,5176,39719,2063,4455,40883,27316,40116,38987,6308,7964,null,40883],"quality":"More is better","isMain":true,"metricName":"NGA50"},{"values":[11714,1268,39719,944,1825,40883,13785,40116,38987,1476,5260,null,40883],"quality":"More is better","isMain":false,"metricName":"NGA75"},{"values":[1,4,1,5,3,1,1,1,1,3,2,null,1],"quality":"Less is better","isMain":false,"metricName":"LG50"},{"values":[1,6,1,13,6,1,1,1,1,7,4,null,1],"quality":"Less is better","isMain":false,"metricName":"LG75"},{"values":[3,3,1,5,3,3,1,1,7,3,2,6,null],"quality":"Less is better","isMain":false,"metricName":"LA50"},{"values":[null,6,1,13,5,null,2,1,null,6,4,12,null],"quality":"Less is better","isMain":false,"metricName":"LA75"},{"values":[2,4,1,5,3,1,1,1,1,3,2,null,1],"quality":"Less is better","isMain":true,"metricName":"LGA50"},{"values":[3,7,1,13,6,1,2,1,1,7,4,null,1],"quality":"Less is better","isMain":false,"metricName":"LGA75"}]],["Misassemblies",[{"values":[3,1,0,0,0,2,1,0,0,0,0,0,2],"quality":"Less is better","isMain":true,"metricName":"# misassemblies"},{"values":[3,1,0,0,0,2,1,0,0,0,0,0,2],"quality":"Less is better","isMain":false,"metricName":" # relocations"},{"values":[0,0,0,0,0,0,0,0,0,0,0,0,0],"quality":"Less is better","isMain":false,"metricName":" # translocations"},{"values":[0,0,0,0,0,0,0,0,0,0,0,0,0],"quality":"Less is better","isMain":false,"metricName":" # inversions"},{"values":[1,1,0,0,0,1,1,0,0,0,0,0,1],"quality":"Less is better","isMain":false,"metricName":"# misassembled contigs"},{"values":[61019,4939,0,0,0,84086,41101,0,0,0,0,0,84086],"quality":"Less is better","isMain":true,"metricName":"Misassembled contigs length"},{"values":[2,0,0,0,0,1,0,1,0,0,0,0,1],"quality":"Less is better","isMain":false,"metricName":"# local misassemblies"},{"values":[0,0,0,0,0,0,0,0,2,0,0,0,1],"quality":"Less is better","isMain":false,"metricName":"# unaligned mis. contigs"}]],["Unaligned",[{"values":[0,0,0,0,0,0,0,0,0,0,0,0,0],"quality":"Less is better","isMain":false,"metricName":"# fully unaligned contigs"},{"values":[0,0,0,0,0,0,0,0,0,0,0,0,0],"quality":"Less is better","isMain":false,"metricName":"Fully unaligned length"},{"values":[1,1,0,0,0,2,0,0,2,0,0,0,2],"quality":"Less is better","isMain":false,"metricName":"# partially unaligned contigs"},{"values":[21039,1407,0,0,0,42444,0,0,57509,0,0,0,73731],"quality":"Less is better","isMain":false,"metricName":"Partially unaligned length"}]],["Mismatches",[{"values":[9,9,0,0,1,162,0,0,232,1,0,0,180],"quality":"Less is better","isMain":false,"metricName":"# mismatches"},{"values":[3,4,0,0,0,6,0,0,14,0,0,0,6],"quality":"Less is better","isMain":false,"metricName":"# indels"},{"values":[48,53,0,0,0,6,0,0,30,0,0,0,6],"quality":"Less is better","isMain":false,"metricName":"Indels length"},{"values":["22.51","25.83","0.00","0.00","2.58","405.11","0.00","0.00","580.16","2.70","0.00","0.00","450.12"],"quality":"Less is better","isMain":true,"metricName":"# mismatches per 100 kbp"},{"values":["7.50","11.48","0.00","0.00","0.00","15.00","0.00","0.00","35.01","0.00","0.00","0.00","15.00"],"quality":"Less is better","isMain":true,"metricName":"# indels per 100 kbp"},{"values":[2,3,0,0,0,6,0,0,13,0,0,0,6],"quality":"Less is better","isMain":false,"metricName":" # indels (<= 5 bp)"},{"values":[1,1,0,0,0,0,0,0,1,0,0,0,0],"quality":"Less is better","isMain":false,"metricName":" # indels (> 5 bp)"},{"values":[202,104,0,0,0,3,0,0,0,0,0,0,114],"quality":"Less is better","isMain":false,"metricName":"# N's"},{"values":["315.54","282.52","0.00","0.00","0.00","3.12","0.00","0.00","0.00","0.00","0.00","0.00","93.26"],"quality":"Less is better","isMain":true,"metricName":"# N's per 100 kbp"}]],["Statistics without reference",[{"values":[2,13,1,26,15,6,1,1,4,14,9,18,2],"quality":"Equal","isMain":true,"metricName":"# contigs"},{"values":[2,7,1,12,9,4,1,1,4,10,6,5,2],"quality":"Equal","isMain":false,"metricName":"# contigs (>= 1000 bp)"},{"values":[1,4,1,2,2,1,1,1,3,3,4,0,2],"quality":"Equal","isMain":false,"metricName":"# contigs (>= 5000 bp)"},{"values":[1,0,1,0,1,1,1,1,3,0,1,0,2],"quality":"Equal","isMain":false,"metricName":"# contigs (>= 10000 bp)"},{"values":[1,0,1,0,0,1,1,1,3,0,0,0,2],"quality":"Equal","isMain":false,"metricName":"# contigs (>= 25000 bp)"},{"values":[1,0,0,0,0,1,0,0,0,0,0,0,1],"quality":"Equal","isMain":false,"metricName":"# contigs (>= 50000 bp)"},{"values":[61019,6705,39719,6396,11374,84086,41101,40116,42293,7633,14540,2877,84086],"quality":"More is better","isMain":true,"metricName":"Largest contig"},{"values":[64018,36811,39719,39989,39759,96052,41101,40116,119222,37599,38430,17747,122234],"quality":"More is better","isMain":true,"metricName":"Total length"},{"values":[64018,32479,39719,29701,35626,94576,41101,40116,119222,34714,36398,8217,122234],"quality":"More is better","isMain":true,"metricName":"Total length (>= 1000 bp)"},{"values":[61019,23664,39719,11688,17976,84086,41101,40116,118110,20500,34072,0,122234],"quality":"More is better","isMain":false,"metricName":"Total length (>= 5000 bp)"},{"values":[61019,0,39719,0,11374,84086,41101,40116,118110,0,14540,0,122234],"quality":"More is better","isMain":true,"metricName":"Total length (>= 10000 bp)"},{"values":[61019,0,39719,0,0,84086,41101,40116,118110,0,0,0,122234],"quality":"More is better","isMain":false,"metricName":"Total length (>= 25000 bp)"},{"values":[61019,0,0,0,0,84086,0,0,0,0,0,0,84086],"quality":"More is better","isMain":true,"metricName":"Total length (>= 50000 bp)"},{"values":[61019,5719,39719,2063,4455,84086,41101,40116,38987,6308,7964,995,84086],"quality":"More is better","isMain":false,"metricName":"N50"},{"values":[61019,4939,39719,944,3301,84086,41101,40116,36830,2728,5260,716,38148],"quality":"More is better","isMain":false,"metricName":"N75"},{"values":[1,3,1,5,3,1,1,1,2,3,2,6,1],"quality":"Less is better","isMain":false,"metricName":"L50"},{"values":[1,5,1,13,5,1,1,1,3,6,4,12,2],"quality":"Less is better","isMain":false,"metricName":"L75"},{"values":["45.03","45.40","45.55","45.46","45.56","45.10","45.44","45.48","44.90","45.50","45.46","45.73","44.88"],"quality":"Equal","isMain":false,"metricName":"GC (%)"}]],["Predicted genes",[]],["Similarity statistics",[{"values":[0,0,0,0,0,0,0,0,0,0,0,0,0],"quality":"Equal","isMain":false,"metricName":"# similar correct contigs"},{"values":[0,0,0,0,0,0,0,0,0,0,0,0,0],"quality":"Equal","isMain":false,"metricName":"# similar misassembled blocks"}]],["Reference statistics",[{"values":[39989,39989,39989,39989,39989,39989,39989,39989,39989,39989,39989,39989,39989],"quality":"Equal","isMain":false,"metricName":"Reference length"},{"values":[1,1,1,1,1,1,1,1,1,1,1,1,1],"quality":"Equal","isMain":false,"metricName":"Reference fragments"},{"values":["45.49","45.49","45.49","45.49","45.49","45.49","45.49","45.49","45.49","45.49","45.49","45.49","45.49"],"quality":"Equal","isMain":false,"metricName":"Reference GC (%)"}]]],"referenceName":"gi\_195661200\_ref\_NC","date":"18 July 2018, Wednesday, 16:45:05","order":[0,1,2,3,4,5,6,7,8,9,10,11,12],"assembliesNames":["ABySS\_127","ABySS\_63","CLC","IDBA\_UD","MEGAHIT","MIRA","Ray\_Meta","SPAdes","SPAdes\_meta","SPAdes\_sc","SPAdes\_sc\_careful","Velvet","Geneious"]},{"assembliesWithNs":null,"minContig":500,"report":[["Genome statistics",[{"values":["100.000","100.000","100.000","100.000","100.000","10.360","100.000","100.000","100.000","100.000","99.772"],"quality":"More is better","isMain":true,"metricName":"Genome fraction (%)"},{"values":["1.000","1.002","1.003","1.009","1.006","1.000","1.003","1.001","1.001","1.001","1.000"],"quality":"Less is better","isMain":true,"metricName":"Duplication ratio"},{"values":[47453,47552,47594,47858,47632,814,47580,47508,47508,47508,47345],"quality":"More is better","isMain":true,"metricName":"Largest alignment"},{"values":[47453,47552,47594,47858,47632,4916,47580,47508,47508,47508,47345],"quality":"More is better","isMain":true,"metricName":"Total aligned length"},{"values":[47453,47552,47594,47858,47740,null,47580,47508,47508,47508,47345],"quality":"More is better","isMain":false,"metricName":"NG50"},{"values":[47453,47552,47594,47858,47740,null,47580,47508,47508,47508,47345],"quality":"More is better","isMain":false,"metricName":"NG75"},{"values":[47453,47552,47594,47858,47632,598,47580,47508,47508,47508,47345],"quality":"More is better","isMain":false,"metricName":"NA50"},{"values":[47453,47552,47594,47858,47632,548,47580,47508,47508,47508,47345],"quality":"More is better","isMain":false,"metricName":"NA75"},{"values":[47453,47552,47594,47858,47632,null,47580,47508,47508,47508,47345],"quality":"More is better","isMain":true,"metricName":"NGA50"},{"values":[47453,47552,47594,47858,47632,null,47580,47508,47508,47508,47345],"quality":"More is better","isMain":false,"metricName":"NGA75"},{"values":[1,1,1,1,1,null,1,1,1,1,1],"quality":"Less is better","isMain":false,"metricName":"LG50"},{"values":[1,1,1,1,1,null,1,1,1,1,1],"quality":"Less is better","isMain":false,"metricName":"LG75"},{"values":[1,1,1,1,1,4,1,1,1,1,1],"quality":"Less is better","isMain":false,"metricName":"LA50"},{"values":[1,1,1,1,1,6,1,1,1,1,1],"quality":"Less is better","isMain":false,"metricName":"LA75"},{"values":[1,1,1,1,1,null,1,1,1,1,1],"quality":"Less is better","isMain":true,"metricName":"LGA50"},{"values":[1,1,1,1,1,null,1,1,1,1,1],"quality":"Less is better","isMain":false,"metricName":"LGA75"}]],["Misassemblies",[{"values":[0,0,0,0,0,0,0,0,0,0,0],"quality":"Less is better","isMain":true,"metricName":"# misassemblies"},{"values":[0,0,0,0,0,0,0,0,0,0,0],"quality":"Less is better","isMain":false,"metricName":" # relocations"},{"values":[0,0,0,0,0,0,0,0,0,0,0],"quality":"Less is better","isMain":false,"metricName":" # translocations"},{"values":[0,0,0,0,0,0,0,0,0,0,0],"quality":"Less is better","isMain":false,"metricName":" # inversions"},{"values":[0,0,0,0,0,0,0,0,0,0,0],"quality":"Less is better","isMain":false,"metricName":"# misassembled contigs"},{"values":[0,0,0,0,0,0,0,0,0,0,0],"quality":"Less is better","isMain":true,"metricName":"Misassembled contigs length"},{"values":[0,1,1,1,1,0,1,0,0,0,0],"quality":"Less is better","isMain":false,"metricName":"# local misassemblies"},{"values":[0,0,0,0,0,0,0,0,0,0,0],"quality":"Less is better","isMain":false,"metricName":"# unaligned mis. contigs"}]],["Unaligned",[{"values":[0,0,0,0,0,0,0,0,0,0,0],"quality":"Less is better","isMain":false,"metricName":"# fully unaligned contigs"},{"values":[0,0,0,0,0,0,0,0,0,0,0],"quality":"Less is better","isMain":false,"metricName":"Fully unaligned length"},{"values":[0,0,0,0,0,0,0,0,0,0,0],"quality":"Less is better","isMain":false,"metricName":"# partially unaligned contigs"},{"values":[0,0,0,0,0,0,0,0,0,0,0],"quality":"Less is better","isMain":false,"metricName":"Partially unaligned length"}]],["Mismatches",[{"values":[0,0,0,0,8,2,0,0,0,0,0],"quality":"Less is better","isMain":false,"metricName":"# mismatches"},{"values":[0,0,0,0,0,0,0,1,1,1,0],"quality":"Less is better","isMain":false,"metricName":"# indels"},{"values":[0,0,0,0,0,0,0,55,55,55,0],"quality":"Less is better","isMain":false,"metricName":"Indels length"},{"values":["0.00","0.00","0.00","0.00","16.86","40.68","0.00","0.00","0.00","0.00","0.00"],"quality":"Less is better","isMain":true,"metricName":"# mismatches per 100 kbp"},{"values":["0.00","0.00","0.00","0.00","0.00","0.00","0.00","2.11","2.11","2.11","0.00"],"quality":"Less is better","isMain":true,"metricName":"# indels per 100 kbp"},{"values":[0,0,0,0,0,0,0,0,0,0,0],"quality":"Less is better","isMain":false,"metricName":" # indels (<= 5 bp)"},{"values":[0,0,0,0,0,0,0,1,1,1,0],"quality":"Less is better","isMain":false,"metricName":" # indels (> 5 bp)"},{"values":[0,0,0,1,0,0,0,0,0,0,0],"quality":"Less is better","isMain":false,"metricName":"# N's"},{"values":["0.00","0.00","0.00","2.09","0.00","0.00","0.00","0.00","0.00","0.00","0.00"],"quality":"Less is better","isMain":true,"metricName":"# N's per 100 kbp"}]],["Statistics without reference",[{"values":[1,1,1,1,1,8,1,1,1,1,1],"quality":"Equal","isMain":true,"metricName":"# contigs"},{"values":[1,1,1,1,1,0,1,1,1,1,1],"quality":"Equal","isMain":false,"metricName":"# contigs (>= 1000 bp)"},{"values":[1,1,1,1,1,0,1,1,1,1,1],"quality":"Equal","isMain":false,"metricName":"# contigs (>= 5000 bp)"},{"values":[1,1,1,1,1,0,1,1,1,1,1],"quality":"Equal","isMain":false,"metricName":"# contigs (>= 10000 bp)"},{"values":[1,1,1,1,1,0,1,1,1,1,1],"quality":"Equal","isMain":false,"metricName":"# contigs (>= 25000 bp)"},{"values":[0,0,0,0,0,0,0,0,0,0,0],"quality":"Equal","isMain":false,"metricName":"# contigs (>= 50000 bp)"},{"values":[47453,47552,47594,47858,47740,814,47580,47508,47508,47508,47345],"quality":"More is better","isMain":true,"metricName":"Largest contig"},{"values":[47453,47552,47594,47858,47740,4916,47580,47508,47508,47508,47345],"quality":"More is better","isMain":true,"metricName":"Total length"},{"values":[47453,47552,47594,47858,47740,0,47580,47508,47508,47508,47345],"quality":"More is better","isMain":true,"metricName":"Total length (>= 1000 bp)"},{"values":[47453,47552,47594,47858,47740,0,47580,47508,47508,47508,47345],"quality":"More is better","isMain":false,"metricName":"Total length (>= 5000 bp)"},{"values":[47453,47552,47594,47858,47740,0,47580,47508,47508,47508,47345],"quality":"More is better","isMain":true,"metricName":"Total length (>= 10000 bp)"},{"values":[47453,47552,47594,47858,47740,0,47580,47508,47508,47508,47345],"quality":"More is better","isMain":false,"metricName":"Total length (>= 25000 bp)"},{"values":[0,0,0,0,0,0,0,0,0,0,0],"quality":"More is better","isMain":true,"metricName":"Total length (>= 50000 bp)"},{"values":[47453,47552,47594,47858,47740,598,47580,47508,47508,47508,47345],"quality":"More is better","isMain":false,"metricName":"N50"},{"values":[47453,47552,47594,47858,47740,548,47580,47508,47508,47508,47345],"quality":"More is better","isMain":false,"metricName":"N75"},{"values":[1,1,1,1,1,4,1,1,1,1,1],"quality":"Less is better","isMain":false,"metricName":"L50"},{"values":[1,1,1,1,1,6,1,1,1,1,1],"quality":"Less is better","isMain":false,"metricName":"L75"},{"values":["46.10","46.10","46.10","46.05","46.29","45.46","46.09","46.10","46.10","46.10","46.04"],"quality":"Equal","isMain":false,"metricName":"GC (%)"}]],["Predicted genes",[]],["Similarity statistics",[{"values":[0,0,0,0,0,0,0,0,0,0,0],"quality":"Equal","isMain":false,"metricName":"# similar correct contigs"},{"values":[0,0,0,0,0,0,0,0,0,0,0],"quality":"Equal","isMain":false,"metricName":"# similar misassembled blocks"}]],["Reference statistics",[{"values":[47453,47453,47453,47453,47453,47453,47453,47453,47453,47453,47453],"quality":"Equal","isMain":false,"metricName":"Reference length"},{"values":[1,1,1,1,1,1,1,1,1,1,1],"quality":"Equal","isMain":false,"metricName":"Reference fragments"},{"values":["46.10","46.10","46.10","46.10","46.10","46.10","46.10","46.10","46.10","46.10","46.10"],"quality":"Equal","isMain":false,"metricName":"Reference GC (%)"}]]],"referenceName":"gi\_197085614\_ref\_NC","date":"18 July 2018, Wednesday, 16:45:18","order":[0,1,2,3,4,5,6,7,8,9,10],"assembliesNames":["CLC","IDBA\_UD","MEGAHIT","MIRA","Ray\_Meta","SOAPdenovo2","SPAdes","SPAdes\_meta","SPAdes\_sc","SPAdes\_sc\_careful","Velvet"]},{"assembliesWithNs":null,"minContig":500,"report":[["Genome statistics",[{"values":["33.820","21.236","32.325","45.346","43.254","55.885","3.696","12.731","38.147","34.094","31.889","23.158","55.708"],"quality":"More is better","isMain":true,"metricName":"Genome fraction (%)"},{"values":["1.028","1.001","1.028","1.016","1.055","1.079","1.451","1.000","1.042","1.011","1.013","1.003","1.001"],"quality":"Less is better","isMain":true,"metricName":"Duplication ratio"},{"values":[1976,1577,2295,2430,2403,8777,748,1672,5438,2774,2630,2309,13703],"quality":"More is better","isMain":true,"metricName":"Largest alignment"},{"values":[22386,13680,21419,29603,29384,38854,2381,8202,25371,22213,20681,14958,35920],"quality":"More is better","isMain":true,"metricName":"Total aligned length"},{"values":[null,null,null,null,null,1221,null,null,null,null,null,null,18311],"quality":"More is better","isMain":false,"metricName":"NG50"},{"values":[null,null,null,null,null,null,null,null,null,null,null,null,13577],"quality":"More is better","isMain":false,"metricName":"NG75"},{"values":[972,893,1049,1813,1043,2415,338,1023,3289,1250,1097,1014,5336],"quality":"More is better","isMain":false,"metricName":"NA50"},{"values":[821,608,813,893,748,1476,null,701,1490,923,752,791,null],"quality":"More is better","isMain":false,"metricName":"NA75"},{"values":[null,null,null,null,null,1221,null,null,null,null,null,null,5336],"quality":"More is better","isMain":true,"metricName":"NGA50"},{"values":[null,null,null,null,null,13,null,null,null,null,null,null,2],"quality":"Less is better","isMain":false,"metricName":"LG50"},{"values":[null,null,null,null,null,null,null,null,null,null,null,null,3],"quality":"Less is better","isMain":false,"metricName":"LG75"},{"values":[9,7,7,7,10,5,4,3,4,6,6,5,4],"quality":"Less is better","isMain":false,"metricName":"LA50"},{"values":[15,11,14,13,18,11,null,6,7,11,12,9,null],"quality":"Less is better","isMain":false,"metricName":"LA75"},{"values":[null,null,null,null,null,13,null,null,null,null,null,null,4],"quality":"Less is better","isMain":true,"metricName":"LGA50"}]],["Misassemblies",[{"values":[0,0,0,0,0,0,0,0,0,0,0,0,1],"quality":"Less is better","isMain":true,"metricName":"# misassemblies"},{"values":[0,0,0,0,0,0,0,0,0,0,0,0,1],"quality":"Less is better","isMain":false,"metricName":" # relocations"},{"values":[0,0,0,0,0,0,0,0,0,0,0,0,0],"quality":"Less is better","isMain":false,"metricName":" # translocations"},{"values":[0,0,0,0,0,0,0,0,0,0,0,0,0],"quality":"Less is better","isMain":false,"metricName":" # inversions"},{"values":[0,0,0,0,0,0,0,0,0,0,0,0,1],"quality":"Less is better","isMain":false,"metricName":"# misassembled contigs"},{"values":[0,0,0,0,0,0,0,0,0,0,0,0,20089],"quality":"Less is better","isMain":true,"metricName":"Misassembled contigs length"},{"values":[0,0,0,0,0,0,0,0,0,0,0,0,0],"quality":"Less is better","isMain":false,"metricName":"# local misassemblies"},{"values":[0,0,0,0,0,0,0,0,0,0,0,0,0],"quality":"Less is better","isMain":false,"metricName":"# unaligned mis. contigs"}]],["Unaligned",[{"values":[0,0,0,0,0,0,0,0,0,0,0,0,0],"quality":"Less is better","isMain":false,"metricName":"# fully unaligned contigs"},{"values":[0,0,0,0,0,0,0,0,0,0,0,0,0],"quality":"Less is better","isMain":false,"metricName":"Fully unaligned length"},{"values":[0,0,0,0,0,0,0,0,1,0,0,0,4],"quality":"Less is better","isMain":false,"metricName":"# partially unaligned contigs"},{"values":[0,0,0,0,0,0,0,0,4965,0,0,0,26596],"quality":"Less is better","isMain":false,"metricName":"Partially unaligned length"}]],["Mismatches",[{"values":[32,39,71,242,217,311,3,34,366,149,115,155,490],"quality":"Less is better","isMain":false,"metricName":"# mismatches"},{"values":[0,2,1,6,3,1,2,0,6,1,2,1,25],"quality":"Less is better","isMain":false,"metricName":"# indels"},{"values":[0,2,1,8,8,6,113,0,6,1,2,1,47],"quality":"Less is better","isMain":false,"metricName":"Indels length"},{"values":["146.86","285.05","340.92","828.34","778.70","863.77","126.00","414.53","1489.20","678.32","559.75","1038.87","1365.24"],"quality":"Less is better","isMain":true,"metricName":"# mismatches per 100 kbp"},{"values":["0.00","14.62","4.80","20.54","10.77","2.78","84.00","0.00","24.41","4.55","9.73","6.70","69.66"],"quality":"Less is better","isMain":true,"metricName":"# indels per 100 kbp"},{"values":[0,2,1,6,2,0,0,0,6,1,2,1,23],"quality":"Less is better","isMain":false,"metricName":" # indels (<= 5 bp)"},{"values":[0,0,0,0,1,1,2,0,0,0,0,0,2],"quality":"Less is better","isMain":false,"metricName":" # indels (> 5 bp)"},{"values":[0,0,0,0,0,108,434,0,236,0,131,0,1533],"quality":"Less is better","isMain":false,"metricName":"# N's"},{"values":["0.00","0.00","0.00","0.00","0.00","277.96","12561.51","0.00","771.65","0.00","629.35","0.00","2452.17"],"quality":"Less is better","isMain":true,"metricName":"# N's per 100 kbp"}]],["Statistics without reference",[{"values":[23,17,21,23,29,19,6,9,11,19,20,15,4],"quality":"Equal","isMain":true,"metricName":"# contigs"},{"values":[8,3,7,11,11,16,0,3,7,10,8,5,4],"quality":"Equal","isMain":false,"metricName":"# contigs (>= 1000 bp)"},{"values":[0,0,0,0,0,1,0,0,1,0,0,0,4],"quality":"Equal","isMain":false,"metricName":"# contigs (>= 5000 bp)"},{"values":[0,0,0,0,0,0,0,0,1,0,0,0,4],"quality":"Equal","isMain":false,"metricName":"# contigs (>= 10000 bp)"},{"values":[0,0,0,0,0,0,0,0,0,0,0,0,0],"quality":"Equal","isMain":false,"metricName":"# contigs (>= 25000 bp)"},{"values":[0,0,0,0,0,0,0,0,0,0,0,0,0],"quality":"Equal","isMain":false,"metricName":"# contigs (>= 50000 bp)"},{"values":[1976,1577,2295,2430,2403,8777,748,1672,10403,2774,2761,2309,20089],"quality":"More is better","isMain":true,"metricName":"Largest contig"},{"values":[22390,13699,21419,29673,29386,38854,3455,8202,30584,22213,20815,14958,62516],"quality":"More is better","isMain":true,"metricName":"Total length"},{"values":[11167,3853,10776,20761,16263,36286,0,4296,28253,16083,12772,8109,62516],"quality":"More is better","isMain":true,"metricName":"Total length (>= 1000 bp)"},{"values":[0,0,0,0,0,8777,0,0,10403,0,0,0,62516],"quality":"More is better","isMain":false,"metricName":"Total length (>= 5000 bp)"},{"values":[0,0,0,0,0,0,0,0,10403,0,0,0,62516],"quality":"More is better","isMain":true,"metricName":"Total length (>= 10000 bp)"},{"values":[0,0,0,0,0,0,0,0,0,0,0,0,0],"quality":"More is better","isMain":false,"metricName":"Total length (>= 25000 bp)"},{"values":[0,0,0,0,0,0,0,0,0,0,0,0,0],"quality":"More is better","isMain":true,"metricName":"Total length (>= 50000 bp)"},{"values":[972,894,1049,1823,1043,2415,562,1023,4349,1250,1097,1014,18311],"quality":"More is better","isMain":false,"metricName":"N50"},{"values":[821,608,813,949,748,1476,512,701,2154,923,755,791,13577],"quality":"More is better","isMain":false,"metricName":"N75"},{"values":[9,7,7,7,10,5,3,3,3,6,6,5,2],"quality":"Less is better","isMain":false,"metricName":"L50"},{"values":[15,11,14,13,18,11,5,6,5,11,12,9,3],"quality":"Less is better","isMain":false,"metricName":"L75"},{"values":["55.52","55.08","54.98","55.79","55.74","55.68","56.47","56.47","55.84","55.79","55.67","55.54","56.23"],"quality":"Equal","isMain":false,"metricName":"GC (%)"}]],["Predicted genes",[]],["Similarity statistics",[{"values":[0,0,0,0,0,0,0,0,0,0,0,0,0],"quality":"Equal","isMain":false,"metricName":"# similar correct contigs"},{"values":[0,0,0,0,0,0,0,0,0,0,0,0,0],"quality":"Equal","isMain":false,"metricName":"# similar misassembled blocks"}]],["Reference statistics",[{"values":[64427,64427,64427,64427,64427,64427,64427,64427,64427,64427,64427,64427,64427],"quality":"Equal","isMain":false,"metricName":"Reference length"},{"values":[1,1,1,1,1,1,1,1,1,1,1,1,1],"quality":"Equal","isMain":false,"metricName":"Reference fragments"},{"values":["55.47","55.47","55.47","55.47","55.47","55.47","55.47","55.47","55.47","55.47","55.47","55.47","55.47"],"quality":"Equal","isMain":false,"metricName":"Reference GC (%)"}]]],"referenceName":"gi\_197261329\_ref\_NC","date":"18 July 2018, Wednesday, 16:45:33","order":[0,1,2,3,4,5,6,7,8,9,10,11,12],"assembliesNames":["ABySS\_127","ABySS\_63","CLC","IDBA\_UD","MEGAHIT","MIRA","SOAPdenovo2","SPAdes","SPAdes\_meta","SPAdes\_sc","SPAdes\_sc\_careful","Velvet","Geneious"]},{"assembliesWithNs":null,"minContig":500,"report":[["Genome statistics",[{"values":["100.000","100.000","99.279","100.000","99.998","100.000","100.000","0.386","100.000","99.748","99.748","99.748","98.207"],"quality":"More is better","isMain":true,"metricName":"Genome fraction (%)"},{"values":["1.324","1.375","1.026","1.001","1.003","1.007","1.001","2.439","1.002","1.000","1.000","1.000","1.000"],"quality":"Less is better","isMain":true,"metricName":"Duplication ratio"},{"values":[53103,53103,52721,53161,35977,53462,53166,130,53228,52967,52967,52970,24141],"quality":"More is better","isMain":true,"metricName":"Largest alignment"},{"values":[70304,73013,53866,53161,53244,53462,53166,205,53228,52967,52967,52970,52154],"quality":"More is better","isMain":true,"metricName":"Total aligned length"},{"values":[70304,73013,52952,53161,35977,53462,53166,null,53228,52967,52967,52970,16056],"quality":"More is better","isMain":false,"metricName":"NG50"},{"values":[70304,73013,52952,53161,17267,53462,53166,null,53228,52967,52967,52970,16056],"quality":"More is better","isMain":false,"metricName":"NG75"},{"values":[53103,53103,52721,53161,35977,53462,53166,null,53228,52967,52967,52970,16056],"quality":"More is better","isMain":false,"metricName":"NA50"},{"values":[53103,16490,52721,53161,17267,53462,53166,null,53228,52967,52967,52970,16056],"quality":"More is better","isMain":false,"metricName":"NA75"},{"values":[53103,53103,52721,53161,35977,53462,53166,null,53228,52967,52967,52970,16056],"quality":"More is better","isMain":true,"metricName":"NGA50"},{"values":[53103,53103,52721,53161,17267,53462,53166,null,53228,52967,52967,52970,16056],"quality":"More is better","isMain":false,"metricName":"NGA75"},{"values":[1,1,1,1,1,1,1,null,1,1,1,1,2],"quality":"Less is better","isMain":false,"metricName":"LG50"},{"values":[1,1,1,1,2,1,1,null,1,1,1,1,2],"quality":"Less is better","isMain":false,"metricName":"LG75"},{"values":[1,1,1,1,1,1,1,null,1,1,1,1,2],"quality":"Less is better","isMain":false,"metricName":"LA50"},{"values":[1,2,1,1,2,1,1,null,1,1,1,1,2],"quality":"Less is better","isMain":false,"metricName":"LA75"},{"values":[1,1,1,1,1,1,1,null,1,1,1,1,2],"quality":"Less is better","isMain":true,"metricName":"LGA50"},{"values":[1,1,1,1,2,1,1,null,1,1,1,1,2],"quality":"Less is better","isMain":false,"metricName":"LGA75"}]],["Misassemblies",[{"values":[2,2,0,0,0,0,0,0,0,0,0,0,0],"quality":"Less is better","isMain":true,"metricName":"# misassemblies"},{"values":[2,2,0,0,0,0,0,0,0,0,0,0,0],"quality":"Less is better","isMain":false,"metricName":" # relocations"},{"values":[0,0,0,0,0,0,0,0,0,0,0,0,0],"quality":"Less is better","isMain":false,"metricName":" # translocations"},{"values":[0,0,0,0,0,0,0,0,0,0,0,0,0],"quality":"Less is better","isMain":false,"metricName":" # inversions"},{"values":[1,1,0,0,0,0,0,0,0,0,0,0,0],"quality":"Less is better","isMain":false,"metricName":"# misassembled contigs"},{"values":[70304,73013,0,0,0,0,0,0,0,0,0,0,0],"quality":"Less is better","isMain":true,"metricName":"Misassembled contigs length"},{"values":[0,0,0,0,0,1,0,0,1,0,0,0,0],"quality":"Less is better","isMain":false,"metricName":"# local misassemblies"},{"values":[0,0,0,0,0,0,0,1,0,0,0,0,0],"quality":"Less is better","isMain":false,"metricName":"# unaligned mis. contigs"}]],["Unaligned",[{"values":[0,0,0,0,0,0,0,0,0,0,0,0,0],"quality":"Less is better","isMain":false,"metricName":"# fully unaligned contigs"},{"values":[0,0,0,0,0,0,0,0,0,0,0,0,0],"quality":"Less is better","isMain":false,"metricName":"Fully unaligned length"},{"values":[0,0,0,0,0,0,0,0,0,0,0,0,0],"quality":"Less is better","isMain":false,"metricName":"# partially unaligned contigs"},{"values":[0,0,0,0,0,0,0,0,0,0,0,0,0],"quality":"Less is better","isMain":false,"metricName":"Partially unaligned length"}]],["Mismatches",[{"values":[0,0,2,0,0,0,0,1,0,0,0,0,1],"quality":"Less is better","isMain":false,"metricName":"# mismatches"},{"values":[1,1,0,1,0,0,1,0,1,1,1,0,1],"quality":"Less is better","isMain":false,"metricName":"# indels"},{"values":[1,1,0,57,0,0,62,0,3,3,3,0,1],"quality":"Less is better","isMain":false,"metricName":"Indels length"},{"values":["0.00","0.00","3.79","0.00","0.00","0.00","0.00","487.80","0.00","0.00","0.00","0.00","1.92"],"quality":"Less is better","isMain":true,"metricName":"# mismatches per 100 kbp"},{"values":["1.88","1.88","0.00","1.88","0.00","0.00","1.88","0.00","1.88","1.89","1.89","0.00","1.92"],"quality":"Less is better","isMain":true,"metricName":"# indels per 100 kbp"},{"values":[1,1,0,0,0,0,0,0,1,1,1,0,1],"quality":"Less is better","isMain":false,"metricName":" # indels (<= 5 bp)"},{"values":[0,0,0,1,0,0,1,0,0,0,0,0,0],"quality":"Less is better","isMain":false,"metricName":" # indels (> 5 bp)"},{"values":[41,82,1,0,0,4,0,294,0,0,0,2,0],"quality":"Less is better","isMain":false,"metricName":"# N's"},{"values":["58.32","112.31","1.85","0.00","0.00","7.48","0.00","58800.00","0.00","0.00","0.00","3.78","0.00"],"quality":"Less is better","isMain":true,"metricName":"# N's per 100 kbp"}]],["Statistics without reference",[{"values":[1,1,2,1,2,1,1,1,1,1,1,1,5],"quality":"Equal","isMain":true,"metricName":"# contigs"},{"values":[1,1,2,1,2,1,1,0,1,1,1,1,4],"quality":"Equal","isMain":false,"metricName":"# contigs (>= 1000 bp)"},{"values":[1,1,1,1,2,1,1,0,1,1,1,1,3],"quality":"Equal","isMain":false,"metricName":"# contigs (>= 5000 bp)"},{"values":[1,1,1,1,2,1,1,0,1,1,1,1,2],"quality":"Equal","isMain":false,"metricName":"# contigs (>= 10000 bp)"},{"values":[1,1,1,1,1,1,1,0,1,1,1,1,0],"quality":"Equal","isMain":false,"metricName":"# contigs (>= 25000 bp)"},{"values":[1,1,1,1,0,1,1,0,1,1,1,1,0],"quality":"Equal","isMain":false,"metricName":"# contigs (>= 50000 bp)"},{"values":[70304,73013,52952,53161,35977,53462,53166,500,53228,52967,52967,52970,24141],"quality":"More is better","isMain":true,"metricName":"Largest contig"},{"values":[70304,73013,54097,53161,53244,53462,53166,500,53228,52967,52967,52970,52154],"quality":"More is better","isMain":true,"metricName":"Total length"},{"values":[70304,73013,54097,53161,53244,53462,53166,0,53228,52967,52967,52970,51318],"quality":"More is better","isMain":true,"metricName":"Total length (>= 1000 bp)"},{"values":[70304,73013,52952,53161,53244,53462,53166,0,53228,52967,52967,52970,48658],"quality":"More is better","isMain":false,"metricName":"Total length (>= 5000 bp)"},{"values":[70304,73013,52952,53161,53244,53462,53166,0,53228,52967,52967,52970,40197],"quality":"More is better","isMain":true,"metricName":"Total length (>= 10000 bp)"},{"values":[70304,73013,52952,53161,35977,53462,53166,0,53228,52967,52967,52970,0],"quality":"More is better","isMain":false,"metricName":"Total length (>= 25000 bp)"},{"values":[70304,73013,52952,53161,0,53462,53166,0,53228,52967,52967,52970,0],"quality":"More is better","isMain":true,"metricName":"Total length (>= 50000 bp)"},{"values":[70304,73013,52952,53161,35977,53462,53166,500,53228,52967,52967,52970,16056],"quality":"More is better","isMain":false,"metricName":"N50"},{"values":[70304,73013,52952,53161,17267,53462,53166,500,53228,52967,52967,52970,16056],"quality":"More is better","isMain":false,"metricName":"N75"},{"values":[1,1,1,1,1,1,1,1,1,1,1,1,2],"quality":"Less is better","isMain":false,"metricName":"L50"},{"values":[1,1,1,1,2,1,1,1,1,1,1,1,2],"quality":"Less is better","isMain":false,"metricName":"L75"},{"values":["36.07","36.05","36.37","36.43","36.43","36.41","36.43","41.26","36.42","36.42","36.42","36.41","36.46"],"quality":"Equal","isMain":false,"metricName":"GC (%)"}]],["Predicted genes",[]],["Similarity statistics",[{"values":[0,0,0,0,0,0,0,0,0,0,0,0,0],"quality":"Equal","isMain":false,"metricName":"# similar correct contigs"},{"values":[0,0,0,0,0,0,0,0,0,0,0,0,0],"quality":"Equal","isMain":false,"metricName":"# similar misassembled blocks"}]],["Reference statistics",[{"values":[53104,53104,53104,53104,53104,53104,53104,53104,53104,53104,53104,53104,53104],"quality":"Equal","isMain":false,"metricName":"Reference length"},{"values":[1,1,1,1,1,1,1,1,1,1,1,1,1],"quality":"Equal","isMain":false,"metricName":"Reference fragments"},{"values":["36.42","36.42","36.42","36.42","36.42","36.42","36.42","36.42","36.42","36.42","36.42","36.42","36.42"],"quality":"Equal","isMain":false,"metricName":"Reference GC (%)"}]]],"referenceName":"gi\_197261511\_ref\_NC","date":"18 July 2018, Wednesday, 16:45:49","order":[0,1,2,3,4,5,6,7,8,9,10,11,12],"assembliesNames":["ABySS\_127","ABySS\_63","CLC","IDBA\_UD","MEGAHIT","MIRA","Ray\_Meta","SOAPdenovo2","SPAdes","SPAdes\_meta","SPAdes\_sc","SPAdes\_sc\_careful","Velvet"]},{"assembliesWithNs":null,"minContig":500,"report":[["Genome statistics",[{"values":["55.511","44.813","89.415","89.115","89.583","77.372","82.662","94.536","92.232","91.330","91.337","65.077","30.907"],"quality":"More is better","isMain":true,"metricName":"Genome fraction (%)"},{"values":["1.001","1.001","1.012","1.002","1.002","1.043","1.008","1.004","1.003","1.002","1.003","1.001","1.003"],"quality":"Less is better","isMain":true,"metricName":"Duplication ratio"},{"values":[2513,2574,19619,18477,18392,4273,6561,18477,19619,19619,19619,18308,6964],"quality":"More is better","isMain":true,"metricName":"Largest alignment"},{"values":[85961,69393,139863,138121,138821,124403,128009,146549,143015,141453,141542,100709,47690],"quality":"More is better","isMain":true,"metricName":"Total aligned length"},{"values":[632,null,4161,4215,4205,1524,2401,5448,5184,5325,5325,2223,null],"quality":"More is better","isMain":false,"metricName":"NG50"},{"values":[null,null,1681,1682,1970,846,1065,2609,2357,2357,2165,null,null],"quality":"More is better","isMain":false,"metricName":"NG75"},{"values":[1072,992,5041,5102,4951,1890,3226,5325,5325,5408,5408,4215,4526],"quality":"More is better","isMain":false,"metricName":"NA50"},{"values":[784,694,2300,2165,2420,1287,1769,2609,2460,2733,2733,2353,3357],"quality":"More is better","isMain":false,"metricName":"NA75"},{"values":[613,null,4161,4215,4205,1476,2401,5258,5184,5325,5325,2186,null],"quality":"More is better","isMain":true,"metricName":"NGA50"},{"values":[null,null,1681,1682,1970,830,1065,2459,2357,2165,2165,null,null],"quality":"More is better","isMain":false,"metricName":"NGA75"},{"values":[71,null,11,11,11,31,20,9,11,10,10,16,null],"quality":"Less is better","isMain":false,"metricName":"LG50"},{"values":[null,null,25,27,25,64,44,19,22,21,22,null,null],"quality":"Less is better","isMain":false,"metricName":"LG75"},{"values":[30,27,9,9,10,23,15,9,9,9,9,8,5],"quality":"Less is better","isMain":false,"metricName":"LA50"},{"values":[53,49,19,20,20,44,29,19,19,18,18,15,8],"quality":"Less is better","isMain":false,"metricName":"LA75"},{"values":[72,null,11,11,11,32,20,10,11,10,10,16,null],"quality":"Less is better","isMain":true,"metricName":"LGA50"},{"values":[null,null,25,27,25,66,44,21,22,22,22,null,null],"quality":"Less is better","isMain":false,"metricName":"LGA75"}]],["Misassemblies",[{"values":[1,0,0,0,0,2,0,2,0,0,0,1,0],"quality":"Less is better","isMain":true,"metricName":"# misassemblies"},{"values":[1,0,0,0,0,2,0,2,0,0,0,1,0],"quality":"Less is better","isMain":false,"metricName":" # relocations"},{"values":[0,0,0,0,0,0,0,0,0,0,0,0,0],"quality":"Less is better","isMain":false,"metricName":" # translocations"},{"values":[0,0,0,0,0,0,0,0,0,0,0,0,0],"quality":"Less is better","isMain":false,"metricName":" # inversions"},{"values":[1,0,0,0,0,2,0,1,0,0,0,1,0],"quality":"Less is better","isMain":false,"metricName":"# misassembled contigs"},{"values":[1839,0,0,0,0,6181,0,9415,0,0,0,2223,0],"quality":"Less is better","isMain":true,"metricName":"Misassembled contigs length"},{"values":[0,0,0,0,1,2,0,2,2,2,2,1,1],"quality":"Less is better","isMain":false,"metricName":"# local misassemblies"},{"values":[0,0,0,0,0,0,0,0,0,0,0,0,0],"quality":"Less is better","isMain":false,"metricName":"# unaligned mis. contigs"}]],["Unaligned",[{"values":[0,0,0,0,0,0,0,0,0,0,0,0,0],"quality":"Less is better","isMain":false,"metricName":"# fully unaligned contigs"},{"values":[0,0,0,0,0,0,0,0,0,0,0,0,0],"quality":"Less is better","isMain":false,"metricName":"Fully unaligned length"},{"values":[0,0,0,0,0,0,0,0,0,0,0,0,0],"quality":"Less is better","isMain":false,"metricName":"# partially unaligned contigs"},{"values":[0,0,0,0,0,0,0,0,0,0,0,0,0],"quality":"Less is better","isMain":false,"metricName":"Partially unaligned length"}]],["Mismatches",[{"values":[31,8,85,39,77,100,48,156,102,72,69,51,37],"quality":"Less is better","isMain":false,"metricName":"# mismatches"},{"values":[1,0,5,1,7,15,10,12,7,5,6,5,17],"quality":"Less is better","isMain":false,"metricName":"# indels"},{"values":[1,0,36,51,15,106,488,35,28,110,43,51,127],"quality":"Less is better","isMain":false,"metricName":"Indels length"},{"values":["36.11","11.54","61.47","28.30","55.58","83.58","37.55","106.71","71.51","50.98","48.85","50.68","77.41"],"quality":"Less is better","isMain":true,"metricName":"# mismatches per 100 kbp"},{"values":["1.16","0.00","3.62","0.73","5.05","12.54","7.82","8.21","4.91","3.54","4.25","4.97","35.57"],"quality":"Less is better","isMain":true,"metricName":"# indels per 100 kbp"},{"values":[1,0,2,0,6,11,0,9,4,1,2,2,14],"quality":"Less is better","isMain":false,"metricName":" # indels (<= 5 bp)"},{"values":[0,0,3,1,1,4,10,3,3,4,4,3,3],"quality":"Less is better","isMain":false,"metricName":" # indels (> 5 bp)"},{"values":[0,0,0,0,0,126,1027,0,0,10,0,0,99],"quality":"Less is better","isMain":false,"metricName":"# N's"},{"values":["0.00","0.00","0.00","0.00","0.00","100.96","796.70","0.00","0.00","7.06","0.00","0.00","206.47"],"quality":"Less is better","isMain":true,"metricName":"# N's per 100 kbp"}]],["Statistics without reference",[{"values":[86,78,50,48,45,76,61,40,41,41,42,35,13],"quality":"Equal","isMain":true,"metricName":"# contigs"},{"values":[34,26,34,36,36,56,45,34,37,35,36,27,13],"quality":"Equal","isMain":false,"metricName":"# contigs (>= 1000 bp)"},{"values":[0,0,9,9,9,0,4,11,11,11,10,4,3],"quality":"Equal","isMain":false,"metricName":"# contigs (>= 5000 bp)"},{"values":[0,0,1,1,1,0,0,1,1,1,1,1,0],"quality":"Equal","isMain":false,"metricName":"# contigs (>= 10000 bp)"},{"values":[0,0,0,0,0,0,0,0,0,0,0,0,0],"quality":"Equal","isMain":false,"metricName":"# contigs (>= 25000 bp)"},{"values":[0,0,0,0,0,0,0,0,0,0,0,0,0],"quality":"Equal","isMain":false,"metricName":"# contigs (>= 50000 bp)"},{"values":[2513,2574,19619,18477,18392,4611,6657,18477,19619,19619,19619,18308,6964],"quality":"More is better","isMain":true,"metricName":"Largest contig"},{"values":[85961,69393,139899,138137,138846,124799,128907,146741,143018,141570,141649,100709,47949],"quality":"More is better","isMain":true,"metricName":"Total length"},{"values":[48119,33763,129299,129436,132817,109099,117418,142604,140295,137660,137739,94829,47949],"quality":"More is better","isMain":true,"metricName":"Total length (>= 1000 bp)"},{"values":[0,0,72285,69984,68996,0,23451,88126,82373,83880,78646,36100,17450],"quality":"More is better","isMain":false,"metricName":"Total length (>= 5000 bp)"},{"values":[0,0,19619,18477,18392,0,0,18477,19619,19619,19619,18308,0],"quality":"More is better","isMain":true,"metricName":"Total length (>= 10000 bp)"},{"values":[0,0,0,0,0,0,0,0,0,0,0,0,0],"quality":"More is better","isMain":false,"metricName":"Total length (>= 25000 bp)"},{"values":[0,0,0,0,0,0,0,0,0,0,0,0,0],"quality":"More is better","isMain":true,"metricName":"Total length (>= 50000 bp)"},{"values":[1072,992,5041,5102,4951,1890,3226,5448,5325,5408,5408,4215,4526],"quality":"More is better","isMain":false,"metricName":"N50"},{"values":[784,694,2300,2165,2421,1288,1769,3298,2460,2840,2840,2353,3357],"quality":"More is better","isMain":false,"metricName":"N75"},{"values":[30,27,9,9,10,23,15,9,9,9,9,8,5],"quality":"Less is better","isMain":false,"metricName":"L50"},{"values":[53,49,19,20,20,43,29,17,19,18,18,15,8],"quality":"Less is better","isMain":false,"metricName":"L75"},{"values":["51.84","51.99","52.14","52.04","51.93","51.89","52.42","51.92","52.06","52.10","52.10","52.32","51.25"],"quality":"Equal","isMain":false,"metricName":"GC (%)"}]],["Predicted genes",[]],["Similarity statistics",[{"values":[0,0,0,0,0,0,0,0,0,0,0,0,0],"quality":"Equal","isMain":false,"metricName":"# similar correct contigs"},{"values":[0,0,0,0,0,0,0,0,0,0,0,0,0],"quality":"Equal","isMain":false,"metricName":"# similar misassembled blocks"}]],["Reference statistics",[{"values":[154641,154641,154641,154641,154641,154641,154641,154641,154641,154641,154641,154641,154641],"quality":"Equal","isMain":false,"metricName":"Reference length"},{"values":[1,1,1,1,1,1,1,1,1,1,1,1,1],"quality":"Equal","isMain":false,"metricName":"Reference fragments"},{"values":["51.80","51.80","51.80","51.80","51.80","51.80","51.80","51.80","51.80","51.80","51.80","51.80","51.80"],"quality":"Equal","isMain":false,"metricName":"Reference GC (%)"}]]],"referenceName":"gi\_197322349\_ref\_NC","date":"18 July 2018, Wednesday, 16:46:06","order":[0,1,2,3,4,5,6,7,8,9,10,11,12],"assembliesNames":["ABySS\_127","ABySS\_63","CLC","IDBA\_UD","MEGAHIT","MIRA","SOAPdenovo2","SPAdes","SPAdes\_meta","SPAdes\_sc","SPAdes\_sc\_careful","Velvet","Geneious"]},{"assembliesWithNs":null,"minContig":500,"report":[["Genome statistics",[{"values":["100.000","97.729","97.812","97.801","97.832","99.877","98.057","0.366","97.822","100.000","97.729","97.729","97.090","97.982"],"quality":"More is better","isMain":true,"metricName":"Genome fraction (%)"},{"values":["1.000","1.327","1.000","1.001","1.001","1.004","1.004","1.000","1.001","1.001","1.000","1.000","1.001","1.002"],"quality":"Less is better","isMain":true,"metricName":"Duplication ratio"},{"values":[137619,133371,131412,131566,131650,134842,123751,503,131622,137713,126434,126434,73849,134842],"quality":"More is better","isMain":true,"metricName":"Largest alignment"},{"values":[137619,178421,134630,134692,134776,137662,135411,503,134748,137713,134533,134533,133717,134842],"quality":"More is better","isMain":true,"metricName":"Total aligned length"},{"values":[137620,178421,131412,131566,131650,135157,123766,null,131622,137813,134533,134533,73849,135157],"quality":"More is better","isMain":false,"metricName":"NG50"},{"values":[137620,178421,131412,131566,131650,135157,123766,null,131622,137813,134533,134533,31881,135157],"quality":"More is better","isMain":false,"metricName":"NG75"},{"values":[137619,133371,131412,131566,131650,134842,123751,503,131622,137713,126434,126434,73849,134842],"quality":"More is better","isMain":false,"metricName":"NA50"},{"values":[137619,45050,131412,131566,131650,134842,123751,503,131622,137713,126434,126434,31881,134842],"quality":"More is better","isMain":false,"metricName":"NA75"},{"values":[137619,133371,131412,131566,131650,134842,123751,null,131622,137713,126434,126434,73849,134842],"quality":"More is better","isMain":true,"metricName":"NGA50"},{"values":[137619,133371,131412,131566,131650,134842,123751,null,131622,137713,126434,126434,31881,134842],"quality":"More is better","isMain":false,"metricName":"NGA75"},{"values":[1,1,1,1,1,1,1,null,1,1,1,1,1,1],"quality":"Less is better","isMain":false,"metricName":"LG50"},{"values":[1,1,1,1,1,1,1,null,1,1,1,1,2,1],"quality":"Less is better","isMain":false,"metricName":"LG75"},{"values":[1,1,1,1,1,1,1,1,1,1,1,1,1,1],"quality":"Less is better","isMain":false,"metricName":"LA50"},{"values":[1,2,1,1,1,1,1,1,1,1,1,1,2,1],"quality":"Less is better","isMain":false,"metricName":"LA75"},{"values":[1,1,1,1,1,1,1,null,1,1,1,1,1,1],"quality":"Less is better","isMain":true,"metricName":"LGA50"},{"values":[1,1,1,1,1,1,1,null,1,1,1,1,2,1],"quality":"Less is better","isMain":false,"metricName":"LGA75"}]],["Misassemblies",[{"values":[0,1,0,0,0,0,0,0,0,0,1,1,0,0],"quality":"Less is better","isMain":true,"metricName":"# misassemblies"},{"values":[0,1,0,0,0,0,0,0,0,0,1,1,0,0],"quality":"Less is better","isMain":false,"metricName":" # relocations"},{"values":[0,0,0,0,0,0,0,0,0,0,0,0,0,0],"quality":"Less is better","isMain":false,"metricName":" # translocations"},{"values":[0,0,0,0,0,0,0,0,0,0,0,0,0,0],"quality":"Less is better","isMain":false,"metricName":" # inversions"},{"values":[0,1,0,0,0,0,0,0,0,0,1,1,0,0],"quality":"Less is better","isMain":false,"metricName":"# misassembled contigs"},{"values":[0,178421,0,0,0,0,0,0,0,0,134533,134533,0,0],"quality":"Less is better","isMain":true,"metricName":"Misassembled contigs length"},{"values":[0,0,0,0,0,0,0,0,0,1,0,0,0,0],"quality":"Less is better","isMain":false,"metricName":"# local misassemblies"},{"values":[0,0,0,0,0,0,0,0,0,0,0,0,0,0],"quality":"Less is better","isMain":false,"metricName":"# unaligned mis. contigs"}]],["Unaligned",[{"values":[0,0,0,0,0,0,0,0,0,0,0,0,0,0],"quality":"Less is better","isMain":false,"metricName":"# fully unaligned contigs"},{"values":[0,0,0,0,0,0,0,0,0,0,0,0,0,0],"quality":"Less is better","isMain":false,"metricName":"Fully unaligned length"},{"values":[0,0,0,0,0,0,0,0,0,0,0,0,0,0],"quality":"Less is better","isMain":false,"metricName":"# partially unaligned contigs"},{"values":[0,0,0,0,0,0,0,0,0,0,0,0,0,0],"quality":"Less is better","isMain":false,"metricName":"Partially unaligned length"}]],["Mismatches",[{"values":[0,0,1,0,0,0,0,4,0,0,0,0,2,0],"quality":"Less is better","isMain":false,"metricName":"# mismatches"},{"values":[0,2,0,0,0,0,3,0,0,0,0,0,0,0],"quality":"Less is better","isMain":false,"metricName":"# indels"},{"values":[0,13,0,0,0,0,63,0,0,0,0,0,0,0],"quality":"Less is better","isMain":false,"metricName":"Indels length"},{"values":["0.00","0.00","0.74","0.00","0.00","0.00","0.00","795.23","0.00","0.00","0.00","0.00","1.50","0.00"],"quality":"Less is better","isMain":true,"metricName":"# mismatches per 100 kbp"},{"values":["0.00","1.49","0.00","0.00","0.00","0.00","2.22","0.00","0.00","0.00","0.00","0.00","0.00","0.00"],"quality":"Less is better","isMain":true,"metricName":"# indels per 100 kbp"},{"values":[0,1,0,0,0,0,1,0,0,0,0,0,0,0],"quality":"Less is better","isMain":false,"metricName":" # indels (<= 5 bp)"},{"values":[0,1,0,0,0,0,2,0,0,0,0,0,0,0],"quality":"Less is better","isMain":false,"metricName":" # indels (> 5 bp)"},{"values":[0,0,0,0,0,0,0,0,0,100,0,0,0,0],"quality":"Less is better","isMain":false,"metricName":"# N's"},{"values":["0.00","0.00","0.00","0.00","0.00","0.00","0.00","0.00","0.00","72.56","0.00","0.00","0.00","0.00"],"quality":"Less is better","isMain":true,"metricName":"# N's per 100 kbp"}]],["Statistics without reference",[{"values":[1,1,2,2,2,2,3,1,2,1,1,1,9,1],"quality":"Equal","isMain":true,"metricName":"# contigs"},{"values":[1,1,2,2,2,2,3,0,2,1,1,1,8,1],"quality":"Equal","isMain":false,"metricName":"# contigs (>= 1000 bp)"},{"values":[1,1,1,1,1,1,2,0,1,1,1,1,4,1],"quality":"Equal","isMain":false,"metricName":"# contigs (>= 5000 bp)"},{"values":[1,1,1,1,1,1,1,0,1,1,1,1,3,1],"quality":"Equal","isMain":false,"metricName":"# contigs (>= 10000 bp)"},{"values":[1,1,1,1,1,1,1,0,1,1,1,1,2,1],"quality":"Equal","isMain":false,"metricName":"# contigs (>= 25000 bp)"},{"values":[1,1,1,1,1,1,1,0,1,1,1,1,1,1],"quality":"Equal","isMain":false,"metricName":"# contigs (>= 50000 bp)"},{"values":[137620,178421,131412,131566,131650,135157,123766,503,131622,137813,134533,134533,73849,135157],"quality":"More is better","isMain":true,"metricName":"Largest contig"},{"values":[137620,178421,134631,134692,134776,137977,135426,503,134748,137813,134533,134533,133718,135157],"quality":"More is better","isMain":true,"metricName":"Total length"},{"values":[137620,178421,134631,134692,134776,137977,135426,0,134748,137813,134533,134533,133004,135157],"quality":"More is better","isMain":true,"metricName":"Total length (>= 1000 bp)"},{"values":[137620,178421,131412,131566,131650,135157,132300,0,131622,137813,134533,134533,122690,135157],"quality":"More is better","isMain":false,"metricName":"Total length (>= 5000 bp)"},{"values":[137620,178421,131412,131566,131650,135157,123766,0,131622,137813,134533,134533,117109,135157],"quality":"More is better","isMain":true,"metricName":"Total length (>= 10000 bp)"},{"values":[137620,178421,131412,131566,131650,135157,123766,0,131622,137813,134533,134533,105730,135157],"quality":"More is better","isMain":false,"metricName":"Total length (>= 25000 bp)"},{"values":[137620,178421,131412,131566,131650,135157,123766,0,131622,137813,134533,134533,73849,135157],"quality":"More is better","isMain":true,"metricName":"Total length (>= 50000 bp)"},{"values":[137620,178421,131412,131566,131650,135157,123766,503,131622,137813,134533,134533,73849,135157],"quality":"More is better","isMain":false,"metricName":"N50"},{"values":[137620,178421,131412,131566,131650,135157,123766,503,131622,137813,134533,134533,31881,135157],"quality":"More is better","isMain":false,"metricName":"N75"},{"values":[1,1,1,1,1,1,1,1,1,1,1,1,1,1],"quality":"Less is better","isMain":false,"metricName":"L50"},{"values":[1,1,1,1,1,1,1,1,1,1,1,1,2,1],"quality":"Less is better","isMain":false,"metricName":"L75"},{"values":["35.93","35.93","35.98","35.99","35.98","35.92","35.98","35.79","35.98","35.93","35.98","35.98","36.01","35.96"],"quality":"Equal","isMain":false,"metricName":"GC (%)"}]],["Predicted genes",[]],["Similarity statistics",[{"values":[0,0,0,0,0,0,0,0,0,0,0,0,0,0],"quality":"Equal","isMain":false,"metricName":"# similar correct contigs"},{"values":[0,0,0,0,0,0,0,0,0,0,0,0,0,0],"quality":"Equal","isMain":false,"metricName":"# similar misassembled blocks"}]],["Reference statistics",[{"values":[137619,137619,137619,137619,137619,137619,137619,137619,137619,137619,137619,137619,137619,137619],"quality":"Equal","isMain":false,"metricName":"Reference length"},{"values":[1,1,1,1,1,1,1,1,1,1,1,1,1,1],"quality":"Equal","isMain":false,"metricName":"Reference fragments"},{"values":["35.93","35.93","35.93","35.93","35.93","35.93","35.93","35.93","35.93","35.93","35.93","35.93","35.93","35.93"],"quality":"Equal","isMain":false,"metricName":"Reference GC (%)"}]]],"referenceName":"gi\_197322500\_ref\_NC","date":"18 July 2018, Wednesday, 16:46:25","order":[0,1,2,3,4,5,6,7,8,9,10,11,12,13],"assembliesNames":["ABySS\_127","ABySS\_63","CLC","IDBA\_UD","MEGAHIT","MIRA","Ray\_Meta","SOAPdenovo2","SPAdes","SPAdes\_meta","SPAdes\_sc","SPAdes\_sc\_careful","Velvet","Geneious"]},{"assembliesWithNs":null,"minContig":500,"report":[["Genome statistics",[{"values":["99.998","98.779","98.593","98.951","99.146","100.000","82.110","74.709","99.081","98.746","98.746","98.746","98.630"],"quality":"More is better","isMain":true,"metricName":"Genome fraction (%)"},{"values":["1.000","1.000","1.000","1.000","1.000","1.017","1.022","1.005","1.000","1.000","1.000","1.000","1.000"],"quality":"Less is better","isMain":true,"metricName":"Duplication ratio"},{"values":[43078,42553,42473,42627,42711,42982,7249,2961,42683,42539,42539,42539,42489],"quality":"More is better","isMain":true,"metricName":"Largest alignment"},{"values":[43078,42553,42473,42627,42711,43783,36134,32203,42683,42539,42539,42539,42489],"quality":"More is better","isMain":true,"metricName":"Total aligned length"},{"values":[43079,42553,42473,42627,42711,42982,3320,1199,42683,42539,42539,42539,42489],"quality":"More is better","isMain":false,"metricName":"NG50"},{"values":[43079,42553,42473,42627,42711,42982,1305,587,42683,42539,42539,42539,42489],"quality":"More is better","isMain":false,"metricName":"NG75"},{"values":[43078,42553,42473,42627,42711,42982,3375,1550,42683,42539,42539,42539,42489],"quality":"More is better","isMain":false,"metricName":"NA50"},{"values":[43078,42553,42473,42627,42711,42982,1706,816,42683,42539,42539,42539,42489],"quality":"More is better","isMain":false,"metricName":"NA75"},{"values":[43078,42553,42473,42627,42711,42982,3320,1041,42683,42539,42539,42539,42489],"quality":"More is better","isMain":true,"metricName":"NGA50"},{"values":[43078,42553,42473,42627,42711,42982,1305,null,42683,42539,42539,42539,42489],"quality":"More is better","isMain":false,"metricName":"NGA75"},{"values":[1,1,1,1,1,1,5,11,1,1,1,1,1],"quality":"Less is better","isMain":false,"metricName":"LG50"},{"values":[1,1,1,1,1,1,11,24,1,1,1,1,1],"quality":"Less is better","isMain":false,"metricName":"LG75"},{"values":[1,1,1,1,1,1,4,8,1,1,1,1,1],"quality":"Less is better","isMain":false,"metricName":"LA50"},{"values":[1,1,1,1,1,1,8,16,1,1,1,1,1],"quality":"Less is better","isMain":false,"metricName":"LA75"},{"values":[1,1,1,1,1,1,5,12,1,1,1,1,1],"quality":"Less is better","isMain":true,"metricName":"LGA50"},{"values":[1,1,1,1,1,1,11,null,1,1,1,1,1],"quality":"Less is better","isMain":false,"metricName":"LGA75"}]],["Misassemblies",[{"values":[0,0,0,0,0,0,0,0,0,0,0,0,0],"quality":"Less is better","isMain":true,"metricName":"# misassemblies"},{"values":[0,0,0,0,0,0,0,0,0,0,0,0,0],"quality":"Less is better","isMain":false,"metricName":" # relocations"},{"values":[0,0,0,0,0,0,0,0,0,0,0,0,0],"quality":"Less is better","isMain":false,"metricName":" # translocations"},{"values":[0,0,0,0,0,0,0,0,0,0,0,0,0],"quality":"Less is better","isMain":false,"metricName":" # inversions"},{"values":[0,0,0,0,0,0,0,0,0,0,0,0,0],"quality":"Less is better","isMain":false,"metricName":"# misassembled contigs"},{"values":[0,0,0,0,0,0,0,0,0,0,0,0,0],"quality":"Less is better","isMain":true,"metricName":"Misassembled contigs length"},{"values":[0,0,0,0,0,1,0,0,0,0,0,0,0],"quality":"Less is better","isMain":false,"metricName":"# local misassemblies"},{"values":[0,0,0,0,0,0,0,0,0,0,0,0,0],"quality":"Less is better","isMain":false,"metricName":"# unaligned mis. contigs"}]],["Unaligned",[{"values":[0,0,0,0,0,0,0,0,0,0,0,0,0],"quality":"Less is better","isMain":false,"metricName":"# fully unaligned contigs"},{"values":[0,0,0,0,0,0,0,0,0,0,0,0,0],"quality":"Less is better","isMain":false,"metricName":"Fully unaligned length"},{"values":[0,0,0,0,0,0,0,1,0,0,0,0,0],"quality":"Less is better","isMain":false,"metricName":"# partially unaligned contigs"},{"values":[0,0,0,0,0,0,0,1428,0,0,0,0,0],"quality":"Less is better","isMain":false,"metricName":"Partially unaligned length"}]],["Mismatches",[{"values":[0,0,0,0,0,0,0,0,0,0,0,0,0],"quality":"Less is better","isMain":false,"metricName":"# mismatches"},{"values":[0,0,0,0,0,0,0,1,0,0,0,0,0],"quality":"Less is better","isMain":false,"metricName":"# indels"},{"values":[0,0,0,0,0,0,0,45,0,0,0,0,0],"quality":"Less is better","isMain":false,"metricName":"Indels length"},{"values":["0.00","0.00","0.00","0.00","0.00","0.00","0.00","0.00","0.00","0.00","0.00","0.00","0.00"],"quality":"Less is better","isMain":true,"metricName":"# mismatches per 100 kbp"},{"values":["0.00","0.00","0.00","0.00","0.00","0.00","0.00","3.11","0.00","0.00","0.00","0.00","0.00"],"quality":"Less is better","isMain":true,"metricName":"# indels per 100 kbp"},{"values":[0,0,0,0,0,0,0,0,0,0,0,0,0],"quality":"Less is better","isMain":false,"metricName":" # indels (<= 5 bp)"},{"values":[0,0,0,0,0,0,0,1,0,0,0,0,0],"quality":"Less is better","isMain":false,"metricName":" # indels (> 5 bp)"},{"values":[0,0,0,0,0,1,0,296,0,0,0,0,0],"quality":"Less is better","isMain":false,"metricName":"# N's"},{"values":["0.00","0.00","0.00","0.00","0.00","2.28","0.00","876.80","0.00","0.00","0.00","0.00","0.00"],"quality":"Less is better","isMain":true,"metricName":"# N's per 100 kbp"}]],["Statistics without reference",[{"values":[1,1,1,1,1,2,15,26,1,1,1,1,1],"quality":"Equal","isMain":true,"metricName":"# contigs"},{"values":[1,1,1,1,1,1,12,12,1,1,1,1,1],"quality":"Equal","isMain":false,"metricName":"# contigs (>= 1000 bp)"},{"values":[1,1,1,1,1,1,1,0,1,1,1,1,1],"quality":"Equal","isMain":false,"metricName":"# contigs (>= 5000 bp)"},{"values":[1,1,1,1,1,1,0,0,1,1,1,1,1],"quality":"Equal","isMain":false,"metricName":"# contigs (>= 10000 bp)"},{"values":[1,1,1,1,1,1,0,0,1,1,1,1,1],"quality":"Equal","isMain":false,"metricName":"# contigs (>= 25000 bp)"},{"values":[0,0,0,0,0,0,0,0,0,0,0,0,0],"quality":"Equal","isMain":false,"metricName":"# contigs (>= 50000 bp)"},{"values":[43079,42553,42473,42627,42711,42982,7249,2961,42683,42539,42539,42539,42489],"quality":"More is better","isMain":true,"metricName":"Largest contig"},{"values":[43079,42553,42473,42627,42711,43818,36134,33759,42683,42539,42539,42539,42489],"quality":"More is better","isMain":true,"metricName":"Total length"},{"values":[43079,42553,42473,42627,42711,42982,33669,23695,42683,42539,42539,42539,42489],"quality":"More is better","isMain":true,"metricName":"Total length (>= 1000 bp)"},{"values":[43079,42553,42473,42627,42711,42982,7249,0,42683,42539,42539,42539,42489],"quality":"More is better","isMain":false,"metricName":"Total length (>= 5000 bp)"},{"values":[43079,42553,42473,42627,42711,42982,0,0,42683,42539,42539,42539,42489],"quality":"More is better","isMain":true,"metricName":"Total length (>= 10000 bp)"},{"values":[43079,42553,42473,42627,42711,42982,0,0,42683,42539,42539,42539,42489],"quality":"More is better","isMain":false,"metricName":"Total length (>= 25000 bp)"},{"values":[0,0,0,0,0,0,0,0,0,0,0,0,0],"quality":"More is better","isMain":true,"metricName":"Total length (>= 50000 bp)"},{"values":[43079,42553,42473,42627,42711,42982,3375,1822,42683,42539,42539,42539,42489],"quality":"More is better","isMain":false,"metricName":"N50"},{"values":[43079,42553,42473,42627,42711,42982,1706,940,42683,42539,42539,42539,42489],"quality":"More is better","isMain":false,"metricName":"N75"},{"values":[1,1,1,1,1,1,4,7,1,1,1,1,1],"quality":"Less is better","isMain":false,"metricName":"L50"},{"values":[1,1,1,1,1,1,8,14,1,1,1,1,1],"quality":"Less is better","isMain":false,"metricName":"L75"},{"values":["61.74","61.71","61.69","61.71","61.71","61.76","61.49","61.75","61.71","61.71","61.71","61.71","61.69"],"quality":"Equal","isMain":false,"metricName":"GC (%)"}]],["Predicted genes",[]],["Similarity statistics",[{"values":[1,1,1,1,1,0,0,0,1,1,1,1,1],"quality":"Equal","isMain":false,"metricName":"# similar correct contigs"},{"values":[0,0,0,0,0,0,0,0,0,0,0,0,0],"quality":"Equal","isMain":false,"metricName":"# similar misassembled blocks"}]],["Reference statistics",[{"values":[43079,43079,43079,43079,43079,43079,43079,43079,43079,43079,43079,43079,43079],"quality":"Equal","isMain":false,"metricName":"Reference length"},{"values":[1,1,1,1,1,1,1,1,1,1,1,1,1],"quality":"Equal","isMain":false,"metricName":"Reference fragments"},{"values":["61.74","61.74","61.74","61.74","61.74","61.74","61.74","61.74","61.74","61.74","61.74","61.74","61.74"],"quality":"Equal","isMain":false,"metricName":"Reference GC (%)"}]]],"referenceName":"gi\_197935853\_ref\_NC","date":"18 July 2018, Wednesday, 16:46:39","order":[0,1,2,3,4,5,6,7,8,9,10,11,12],"assembliesNames":["ABySS\_127","ABySS\_63","CLC","IDBA\_UD","MEGAHIT","MIRA","Ray\_Meta","SOAPdenovo2","SPAdes","SPAdes\_meta","SPAdes\_sc","SPAdes\_sc\_careful","Velvet"]},{"assembliesWithNs":null,"minContig":500,"report":[["Genome statistics",[{"values":["7.097","2.025","67.993","60.967","63.244","9.082","58.302","71.609","72.223","71.564","71.564","1.440"],"quality":"More is better","isMain":true,"metricName":"Genome fraction (%)"},{"values":["1.000","1.000","1.002","1.001","1.001","1.000","1.001","1.002","1.003","1.003","1.003","1.000"],"quality":"Less is better","isMain":true,"metricName":"Duplication ratio"},{"values":[873,762,2554,2400,2352,978,2545,2554,2554,2554,2554,542],"quality":"More is better","isMain":true,"metricName":"Largest alignment"},{"values":[2671,762,25645,22945,23823,3418,21969,27000,27258,26983,26983,542],"quality":"More is better","isMain":true,"metricName":"Total aligned length"},{"values":[null,null,815,841,714,null,589,1114,1023,1114,1114,null],"quality":"More is better","isMain":false,"metricName":"NG50"},{"values":[777,762,1114,1108,961,928,1108,1150,1147,1150,1150,542],"quality":"More is better","isMain":false,"metricName":"NA50"},{"values":[521,762,766,846,716,807,667,1001,939,1001,1001,542],"quality":"More is better","isMain":false,"metricName":"NA75"},{"values":[null,null,815,841,714,null,589,1108,1023,1108,1108,null],"quality":"More is better","isMain":true,"metricName":"NGA50"},{"values":[null,null,15,16,17,null,19,14,14,14,14,null],"quality":"Less is better","isMain":false,"metricName":"LG50"},{"values":[2,1,9,8,9,2,8,9,9,9,9,1],"quality":"Less is better","isMain":false,"metricName":"LA50"},{"values":[3,1,16,14,16,3,15,16,16,16,16,1],"quality":"Less is better","isMain":false,"metricName":"LA75"},{"values":[null,null,15,16,17,null,19,14,14,14,14,null],"quality":"Less is better","isMain":true,"metricName":"LGA50"}]],["Misassemblies",[{"values":[0,0,0,0,0,0,0,0,0,0,0,0],"quality":"Less is better","isMain":true,"metricName":"# misassemblies"},{"values":[0,0,0,0,0,0,0,0,0,0,0,0],"quality":"Less is better","isMain":false,"metricName":" # relocations"},{"values":[0,0,0,0,0,0,0,0,0,0,0,0],"quality":"Less is better","isMain":false,"metricName":" # translocations"},{"values":[0,0,0,0,0,0,0,0,0,0,0,0],"quality":"Less is better","isMain":false,"metricName":" # inversions"},{"values":[0,0,0,0,0,0,0,0,0,0,0,0],"quality":"Less is better","isMain":false,"metricName":"# misassembled contigs"},{"values":[0,0,0,0,0,0,0,0,0,0,0,0],"quality":"Less is better","isMain":true,"metricName":"Misassembled contigs length"},{"values":[0,0,0,0,0,0,0,0,0,0,0,0],"quality":"Less is better","isMain":false,"metricName":"# local misassemblies"},{"values":[0,0,0,0,0,0,0,0,0,0,0,0],"quality":"Less is better","isMain":false,"metricName":"# unaligned mis. contigs"}]],["Unaligned",[{"values":[0,0,0,0,0,0,0,0,0,0,0,0],"quality":"Less is better","isMain":false,"metricName":"# fully unaligned contigs"},{"values":[0,0,0,0,0,0,0,0,0,0,0,0],"quality":"Less is better","isMain":false,"metricName":"Fully unaligned length"},{"values":[0,0,0,0,0,0,0,1,0,1,1,0],"quality":"Less is better","isMain":false,"metricName":"# partially unaligned contigs"},{"values":[0,0,0,0,0,0,0,773,0,773,773,0],"quality":"Less is better","isMain":false,"metricName":"Partially unaligned length"}]],["Mismatches",[{"values":[0,0,35,23,21,0,20,41,43,37,33,0],"quality":"Less is better","isMain":false,"metricName":"# mismatches"},{"values":[0,0,0,0,0,0,0,0,0,0,0,0],"quality":"Less is better","isMain":false,"metricName":"# indels"},{"values":[0,0,0,0,0,0,0,0,0,0,0,0],"quality":"Less is better","isMain":false,"metricName":"Indels length"},{"values":["0.00","0.00","136.78","100.24","88.23","0.00","91.15","152.13","158.20","137.38","122.53","0.00"],"quality":"Less is better","isMain":true,"metricName":"# mismatches per 100 kbp"},{"values":["0.00","0.00","0.00","0.00","0.00","0.00","0.00","0.00","0.00","0.00","0.00","0.00"],"quality":"Less is better","isMain":true,"metricName":"# indels per 100 kbp"},{"values":[0,0,0,0,0,0,0,0,0,0,0,0],"quality":"Less is better","isMain":false,"metricName":" # indels (<= 5 bp)"},{"values":[0,0,0,0,0,0,0,0,0,0,0,0],"quality":"Less is better","isMain":false,"metricName":" # indels (> 5 bp)"},{"values":[0,0,0,0,0,3,0,0,0,0,0,0],"quality":"Less is better","isMain":false,"metricName":"# N's"},{"values":["0.00","0.00","0.00","0.00","0.00","87.77","0.00","0.00","0.00","0.00","0.00","0.00"],"quality":"Less is better","isMain":true,"metricName":"# N's per 100 kbp"}]],["Statistics without reference",[{"values":[4,1,25,22,25,4,24,24,25,24,24,1],"quality":"Equal","isMain":true,"metricName":"# contigs"},{"values":[0,0,11,9,7,0,9,17,15,17,17,0],"quality":"Equal","isMain":false,"metricName":"# contigs (>= 1000 bp)"},{"values":[0,0,0,0,0,0,0,0,0,0,0,0],"quality":"Equal","isMain":false,"metricName":"# contigs (>= 5000 bp)"},{"values":[0,0,0,0,0,0,0,0,0,0,0,0],"quality":"Equal","isMain":false,"metricName":"# contigs (>= 10000 bp)"},{"values":[0,0,0,0,0,0,0,0,0,0,0,0],"quality":"Equal","isMain":false,"metricName":"# contigs (>= 25000 bp)"},{"values":[0,0,0,0,0,0,0,0,0,0,0,0],"quality":"Equal","isMain":false,"metricName":"# contigs (>= 50000 bp)"},{"values":[873,762,2554,2400,2352,978,2545,2554,2554,2554,2554,542],"quality":"More is better","isMain":true,"metricName":"Largest contig"},{"values":[2671,762,25645,22977,23823,3418,21969,27773,27258,27777,27777,542],"quality":"More is better","isMain":true,"metricName":"Total length"},{"values":[0,0,15621,12762,10375,0,12072,22938,20210,22942,22942,0],"quality":"More is better","isMain":true,"metricName":"Total length (>= 1000 bp)"},{"values":[0,0,0,0,0,0,0,0,0,0,0,0],"quality":"More is better","isMain":false,"metricName":"Total length (>= 5000 bp)"},{"values":[0,0,0,0,0,0,0,0,0,0,0,0],"quality":"More is better","isMain":true,"metricName":"Total length (>= 10000 bp)"},{"values":[0,0,0,0,0,0,0,0,0,0,0,0],"quality":"More is better","isMain":false,"metricName":"Total length (>= 25000 bp)"},{"values":[0,0,0,0,0,0,0,0,0,0,0,0],"quality":"More is better","isMain":true,"metricName":"Total length (>= 50000 bp)"},{"values":[777,762,1114,1111,961,928,1108,1225,1147,1225,1225,542],"quality":"More is better","isMain":false,"metricName":"N50"},{"values":[521,762,766,846,716,807,667,1108,939,1108,1108,542],"quality":"More is better","isMain":false,"metricName":"N75"},{"values":[2,1,9,8,9,2,8,9,9,9,9,1],"quality":"Less is better","isMain":false,"metricName":"L50"},{"values":[3,1,16,14,16,3,15,15,16,15,15,1],"quality":"Less is better","isMain":false,"metricName":"L75"},{"values":["64.43","62.73","62.92","62.96","62.99","62.99","63.06","62.97","62.70","62.97","62.96","62.73"],"quality":"Equal","isMain":false,"metricName":"GC (%)"}]],["Predicted genes",[]],["Similarity statistics",[{"values":[0,0,0,0,0,0,0,0,0,0,0,0],"quality":"Equal","isMain":false,"metricName":"# similar correct contigs"},{"values":[0,0,0,0,0,0,0,0,0,0,0,0],"quality":"Equal","isMain":false,"metricName":"# similar misassembled blocks"}]],["Reference statistics",[{"values":[37635,37635,37635,37635,37635,37635,37635,37635,37635,37635,37635,37635],"quality":"Equal","isMain":false,"metricName":"Reference length"},{"values":[1,1,1,1,1,1,1,1,1,1,1,1],"quality":"Equal","isMain":false,"metricName":"Reference fragments"},{"values":["62.87","62.87","62.87","62.87","62.87","62.87","62.87","62.87","62.87","62.87","62.87","62.87"],"quality":"Equal","isMain":false,"metricName":"Reference GC (%)"}]]],"referenceName":"gi\_198449262\_ref\_NC","date":"18 July 2018, Wednesday, 16:46:54","order":[0,1,2,3,4,5,6,7,8,9,10,11],"assembliesNames":["ABySS\_127","ABySS\_63","CLC","IDBA\_UD","MEGAHIT","MIRA","SOAPdenovo2","SPAdes","SPAdes\_meta","SPAdes\_sc","SPAdes\_sc\_careful","Velvet"]},{"assembliesWithNs":null,"minContig":500,"report":[["Genome statistics",[{"values":["87.244","79.822","99.399","99.334","98.661","93.114","0.505","96.578","99.834","99.834","99.466","99.466","96.277","44.501"],"quality":"More is better","isMain":true,"metricName":"Genome fraction (%)"},{"values":["1.001","1.002","1.017","1.000","1.000","1.047","1.000","1.013","1.000","1.000","1.000","1.000","1.001","1.000"],"quality":"Less is better","isMain":true,"metricName":"Duplication ratio"},{"values":[5150,4373,22963,27212,27178,12651,562,13258,27341,27341,26391,26391,17217,12651],"quality":"More is better","isMain":true,"metricName":"Largest alignment"},{"values":[97255,89032,112573,110634,109870,108572,562,107780,111210,111210,110800,110800,107240,49557],"quality":"More is better","isMain":true,"metricName":"Total aligned length"},{"values":[1522,1324,7780,13361,19888,4415,null,6284,19951,19951,19951,19951,11445,null],"quality":"More is better","isMain":false,"metricName":"NG50"},{"values":[807,639,6697,7315,6908,3218,null,3916,7315,7315,7315,7315,5169,null],"quality":"More is better","isMain":false,"metricName":"NG75"},{"values":[2008,1736,7780,13361,19888,4415,562,6284,19951,19951,19951,19951,11381,7015],"quality":"More is better","isMain":false,"metricName":"NA50"},{"values":[1030,1049,6697,7315,6908,3218,562,3916,7315,7315,7315,7315,5169,6412],"quality":"More is better","isMain":false,"metricName":"NA75"},{"values":[1522,1324,7780,13361,19888,4415,null,6284,19951,19951,19951,19951,11381,null],"quality":"More is better","isMain":true,"metricName":"NGA50"},{"values":[807,639,6697,7315,6908,3218,null,3916,7315,7315,7315,7315,5169,null],"quality":"More is better","isMain":false,"metricName":"NGA75"},{"values":[19,25,4,3,3,7,null,7,3,3,3,3,4,null],"quality":"Less is better","isMain":false,"metricName":"LG50"},{"values":[44,55,8,6,5,14,null,12,5,5,5,5,8,null],"quality":"Less is better","isMain":false,"metricName":"LG75"},{"values":[15,17,4,3,3,7,1,7,3,3,3,3,4,3],"quality":"Less is better","isMain":false,"metricName":"LA50"},{"values":[32,34,8,6,5,14,1,12,5,5,5,5,8,5],"quality":"Less is better","isMain":false,"metricName":"LA75"},{"values":[19,25,4,3,3,7,null,7,3,3,3,3,4,null],"quality":"Less is better","isMain":true,"metricName":"LGA50"},{"values":[44,55,8,6,5,14,null,12,5,5,5,5,8,null],"quality":"Less is better","isMain":false,"metricName":"LGA75"}]],["Misassemblies",[{"values":[0,0,0,0,0,0,0,0,0,0,0,0,0,0],"quality":"Less is better","isMain":true,"metricName":"# misassemblies"},{"values":[0,0,0,0,0,0,0,0,0,0,0,0,0,0],"quality":"Less is better","isMain":false,"metricName":" # relocations"},{"values":[0,0,0,0,0,0,0,0,0,0,0,0,0,0],"quality":"Less is better","isMain":false,"metricName":" # translocations"},{"values":[0,0,0,0,0,0,0,0,0,0,0,0,0,0],"quality":"Less is better","isMain":false,"metricName":" # inversions"},{"values":[0,0,0,0,0,0,0,0,0,0,0,0,0,0],"quality":"Less is better","isMain":false,"metricName":"# misassembled contigs"},{"values":[0,0,0,0,0,0,0,0,0,0,0,0,0,0],"quality":"Less is better","isMain":true,"metricName":"Misassembled contigs length"},{"values":[0,0,0,0,1,0,0,0,0,0,1,1,1,0],"quality":"Less is better","isMain":false,"metricName":"# local misassemblies"},{"values":[0,0,0,0,0,0,0,0,0,0,0,0,0,0],"quality":"Less is better","isMain":false,"metricName":"# unaligned mis. contigs"}]],["Unaligned",[{"values":[0,0,0,0,0,0,0,0,0,0,0,0,0,0],"quality":"Less is better","isMain":false,"metricName":"# fully unaligned contigs"},{"values":[0,0,0,0,0,0,0,0,0,0,0,0,0,0],"quality":"Less is better","isMain":false,"metricName":"Fully unaligned length"},{"values":[0,0,0,0,0,0,0,0,0,0,0,0,0,1],"quality":"Less is better","isMain":false,"metricName":"# partially unaligned contigs"},{"values":[0,0,0,0,0,0,0,0,0,0,0,0,0,5233],"quality":"Less is better","isMain":false,"metricName":"Partially unaligned length"}]],["Mismatches",[{"values":[9,10,17,12,13,8,0,12,36,24,22,13,28,6],"quality":"Less is better","isMain":false,"metricName":"# mismatches"},{"values":[0,0,0,0,1,0,0,14,0,0,0,0,3,0],"quality":"Less is better","isMain":false,"metricName":"# indels"},{"values":[0,0,0,0,1,0,0,693,0,0,0,0,49,0],"quality":"Less is better","isMain":false,"metricName":"Indels length"},{"values":["9.26","11.25","15.36","10.85","11.83","7.72","0.00","11.16","32.38","21.59","19.86","11.74","26.12","12.11"],"quality":"Less is better","isMain":true,"metricName":"# mismatches per 100 kbp"},{"values":["0.00","0.00","0.00","0.00","0.91","0.00","0.00","13.02","0.00","0.00","0.00","0.00","2.80","0.00"],"quality":"Less is better","isMain":true,"metricName":"# indels per 100 kbp"},{"values":[0,0,0,0,1,0,0,0,0,0,0,0,2,0],"quality":"Less is better","isMain":false,"metricName":" # indels (<= 5 bp)"},{"values":[0,0,0,0,0,0,0,14,0,0,0,0,1,0],"quality":"Less is better","isMain":false,"metricName":" # indels (> 5 bp)"},{"values":[0,0,0,0,0,56,0,1380,0,0,10,10,0,26],"quality":"Less is better","isMain":false,"metricName":"# N's"},{"values":["0.00","0.00","0.00","0.00","0.00","51.58","0.00","1266.38","0.00","0.00","9.02","9.02","0.00","47.45"],"quality":"Less is better","isMain":true,"metricName":"# N's per 100 kbp"}]],["Statistics without reference",[{"values":[64,64,19,11,11,32,1,23,10,10,11,11,17,7],"quality":"Equal","isMain":true,"metricName":"# contigs"},{"values":[32,34,17,10,10,21,0,20,9,9,9,9,15,7],"quality":"Equal","isMain":false,"metricName":"# contigs (>= 1000 bp)"},{"values":[1,0,8,8,6,5,0,10,7,7,7,7,8,6],"quality":"Equal","isMain":false,"metricName":"# contigs (>= 5000 bp)"},{"values":[0,0,3,4,3,1,0,2,3,3,3,3,4,1],"quality":"Equal","isMain":false,"metricName":"# contigs (>= 10000 bp)"},{"values":[0,0,0,1,2,0,0,0,2,2,2,2,0,0],"quality":"Equal","isMain":false,"metricName":"# contigs (>= 25000 bp)"},{"values":[0,0,0,0,0,0,0,0,0,0,0,0,0,0],"quality":"Equal","isMain":false,"metricName":"# contigs (>= 50000 bp)"},{"values":[5150,4373,22963,27212,27178,12651,562,13466,27341,27341,26401,26401,17217,12651],"quality":"More is better","isMain":true,"metricName":"Largest contig"},{"values":[97255,89032,112573,110634,109870,108572,562,108972,111210,111210,110810,110810,107304,54790],"quality":"More is better","isMain":true,"metricName":"Total length"},{"values":[73456,67600,110690,109665,108994,100934,0,106349,110241,110241,109301,109301,106030,54790],"quality":"More is better","isMain":true,"metricName":"Total length (>= 1000 bp)"},{"values":[5150,0,86192,101840,93571,49361,0,77836,102086,102086,101146,101146,84576,50705],"quality":"More is better","isMain":false,"metricName":"Total length (>= 5000 bp)"},{"values":[0,0,50538,72991,72931,12651,0,25469,73226,73226,72286,72286,57758,12651],"quality":"More is better","isMain":true,"metricName":"Total length (>= 10000 bp)"},{"values":[0,0,0,27212,53043,0,0,0,53275,53275,52335,52335,0,0],"quality":"More is better","isMain":false,"metricName":"Total length (>= 25000 bp)"},{"values":[0,0,0,0,0,0,0,0,0,0,0,0,0,0],"quality":"More is better","isMain":true,"metricName":"Total length (>= 50000 bp)"},{"values":[2008,1736,7780,13361,19888,4415,562,6284,19951,19951,19951,19951,11445,8097],"quality":"More is better","isMain":false,"metricName":"N50"},{"values":[1030,1049,6697,7315,6908,3218,562,4577,7315,7315,7315,7315,5169,6534],"quality":"More is better","isMain":false,"metricName":"N75"},{"values":[15,17,4,3,3,7,1,7,3,3,3,3,4,3],"quality":"Less is better","isMain":false,"metricName":"L50"},{"values":[32,34,8,6,5,14,1,11,5,5,5,5,8,5],"quality":"Less is better","isMain":false,"metricName":"L75"},{"values":["54.83","54.92","54.81","54.79","54.80","54.80","54.98","54.79","54.79","54.79","54.75","54.75","54.70","53.62"],"quality":"Equal","isMain":false,"metricName":"GC (%)"}]],["Predicted genes",[]],["Similarity statistics",[{"values":[0,0,0,0,0,0,0,0,0,0,0,0,0,0],"quality":"Equal","isMain":false,"metricName":"# similar correct contigs"},{"values":[0,0,0,0,0,0,0,0,0,0,0,0,0,0],"quality":"Equal","isMain":false,"metricName":"# similar misassembled blocks"}]],["Reference statistics",[{"values":[111362,111362,111362,111362,111362,111362,111362,111362,111362,111362,111362,111362,111362,111362],"quality":"Equal","isMain":false,"metricName":"Reference length"},{"values":[1,1,1,1,1,1,1,1,1,1,1,1,1,1],"quality":"Equal","isMain":false,"metricName":"Reference fragments"},{"values":["54.78","54.78","54.78","54.78","54.78","54.78","54.78","54.78","54.78","54.78","54.78","54.78","54.78","54.78"],"quality":"Equal","isMain":false,"metricName":"Reference GC (%)"}]]],"referenceName":"gi\_19881405\_ref\_NC","date":"18 July 2018, Wednesday, 16:47:12","order":[0,1,2,3,4,5,6,7,8,9,10,11,12,13],"assembliesNames":["ABySS\_127","ABySS\_63","CLC","IDBA\_UD","MEGAHIT","MIRA","Ray\_Meta","SOAPdenovo2","SPAdes","SPAdes\_meta","SPAdes\_sc","SPAdes\_sc\_careful","Velvet","Geneious"]},{"assembliesWithNs":null,"minContig":500,"report":[["Genome statistics",[{"values":["99.951","86.848","1.798","100.000","100.000","100.000","100.000","100.000","100.000","22.073","100.000"],"quality":"More is better","isMain":true,"metricName":"Genome fraction (%)"},{"values":["1.000","1.030","1.000","217.119","1.108","1.003","1.001","1.001","1.001","1.000","1.789"],"quality":"Less is better","isMain":true,"metricName":"Duplication ratio"},{"values":[44907,3289,808,37856,45985,44929,44984,44984,44984,968,40292],"quality":"More is better","isMain":true,"metricName":"Largest alignment"},{"values":[44907,40208,808,9753683,49742,44929,44984,44984,44984,9917,80361],"quality":"More is better","isMain":true,"metricName":"Total aligned length"},{"values":[44929,1692,null,37856,49784,45056,44984,44984,44984,null,74102],"quality":"More is better","isMain":false,"metricName":"NG50"},{"values":[44929,970,null,37856,49784,45056,44984,44984,44984,null,74102],"quality":"More is better","isMain":false,"metricName":"NG75"},{"values":[44907,1692,808,3550,45985,44929,44984,44984,44984,651,40292],"quality":"More is better","isMain":false,"metricName":"NA50"},{"values":[44907,1359,808,2356,45985,44929,44984,44984,44984,560,32584],"quality":"More is better","isMain":false,"metricName":"NA75"},{"values":[44907,1692,null,37856,45985,44929,44984,44984,44984,null,40292],"quality":"More is better","isMain":true,"metricName":"NGA50"},{"values":[44907,970,null,37856,45985,44929,44984,44984,44984,null,40292],"quality":"More is better","isMain":false,"metricName":"NGA75"},{"values":[1,10,null,1,1,1,1,1,1,null,1],"quality":"Less is better","isMain":false,"metricName":"LG50"},{"values":[1,18,null,1,1,1,1,1,1,null,1],"quality":"Less is better","isMain":false,"metricName":"LG75"},{"values":[1,9,1,790,1,1,1,1,1,7,1],"quality":"Less is better","isMain":false,"metricName":"LA50"},{"values":[1,15,1,1641,1,1,1,1,1,11,2],"quality":"Less is better","isMain":false,"metricName":"LA75"},{"values":[1,10,null,1,1,1,1,1,1,null,1],"quality":"Less is better","isMain":true,"metricName":"LGA50"},{"values":[1,18,null,1,1,1,1,1,1,null,1],"quality":"Less is better","isMain":false,"metricName":"LGA75"}]],["Misassemblies",[{"values":[0,0,0,6,1,0,0,0,0,0,3],"quality":"Less is better","isMain":true,"metricName":"# misassemblies"},{"values":[0,0,0,6,1,0,0,0,0,0,3],"quality":"Less is better","isMain":false,"metricName":" # relocations"},{"values":[0,0,0,0,0,0,0,0,0,0,0],"quality":"Less is better","isMain":false,"metricName":" # translocations"},{"values":[0,0,0,0,0,0,0,0,0,0,0],"quality":"Less is better","isMain":false,"metricName":" # inversions"},{"values":[0,0,0,6,1,0,0,0,0,0,2],"quality":"Less is better","isMain":false,"metricName":"# misassembled contigs"},{"values":[0,0,0,20564,49784,0,0,0,0,0,80367],"quality":"Less is better","isMain":true,"metricName":"Misassembled contigs length"},{"values":[0,0,0,0,1,0,0,0,0,0,0],"quality":"Less is better","isMain":false,"metricName":"# local misassemblies"},{"values":[0,0,0,0,0,0,0,0,0,0,0],"quality":"Less is better","isMain":false,"metricName":"# unaligned mis. contigs"}]],["Unaligned",[{"values":[0,0,0,0,0,0,0,0,0,0,0],"quality":"Less is better","isMain":false,"metricName":"# fully unaligned contigs"},{"values":[0,0,0,0,0,0,0,0,0,0,0],"quality":"Less is better","isMain":false,"metricName":"Fully unaligned length"},{"values":[0,0,0,0,0,0,0,0,0,0,0],"quality":"Less is better","isMain":false,"metricName":"# partially unaligned contigs"},{"values":[0,0,0,0,0,0,0,0,0,0,0],"quality":"Less is better","isMain":false,"metricName":"Partially unaligned length"}]],["Mismatches",[{"values":[0,0,2,178,0,0,0,0,0,3,6],"quality":"Less is better","isMain":false,"metricName":"# mismatches"},{"values":[0,0,0,4,1,0,1,1,1,0,0],"quality":"Less is better","isMain":false,"metricName":"# indels"},{"values":[0,0,0,4,65,0,55,55,55,0,0],"quality":"Less is better","isMain":false,"metricName":"Indels length"},{"values":["0.00","0.00","247.52","396.18","0.00","0.00","0.00","0.00","0.00","30.25","13.35"],"quality":"Less is better","isMain":true,"metricName":"# mismatches per 100 kbp"},{"values":["0.00","0.00","0.00","8.90","2.23","0.00","2.23","2.23","2.23","0.00","0.00"],"quality":"Less is better","isMain":true,"metricName":"# indels per 100 kbp"},{"values":[0,0,0,4,0,0,0,0,0,0,0],"quality":"Less is better","isMain":false,"metricName":" # indels (<= 5 bp)"},{"values":[0,0,0,0,1,0,1,1,1,0,0],"quality":"Less is better","isMain":false,"metricName":" # indels (> 5 bp)"},{"values":[0,0,0,4722,0,0,0,0,0,0,105],"quality":"Less is better","isMain":false,"metricName":"# N's"},{"values":["0.00","0.00","0.00","48.41","0.00","0.00","0.00","0.00","0.00","0.00","130.65"],"quality":"Less is better","isMain":true,"metricName":"# N's per 100 kbp"}]],["Statistics without reference",[{"values":[1,26,1,2983,1,1,1,1,1,15,2],"quality":"Equal","isMain":true,"metricName":"# contigs"},{"values":[1,17,0,2965,1,1,1,1,1,0,2],"quality":"Equal","isMain":false,"metricName":"# contigs (>= 1000 bp)"},{"values":[1,0,0,415,1,1,1,1,1,0,2],"quality":"Equal","isMain":false,"metricName":"# contigs (>= 5000 bp)"},{"values":[1,0,0,68,1,1,1,1,1,0,1],"quality":"Equal","isMain":false,"metricName":"# contigs (>= 10000 bp)"},{"values":[1,0,0,2,1,1,1,1,1,0,1],"quality":"Equal","isMain":false,"metricName":"# contigs (>= 25000 bp)"},{"values":[0,0,0,0,0,0,0,0,0,0,1],"quality":"Equal","isMain":false,"metricName":"# contigs (>= 50000 bp)"},{"values":[44929,3289,808,37856,49784,45056,44984,44984,44984,968,74102],"quality":"More is better","isMain":true,"metricName":"Largest contig"},{"values":[44929,40208,808,9754953,49784,45056,44984,44984,44984,9917,80367],"quality":"More is better","isMain":true,"metricName":"Total length"},{"values":[44929,33089,0,9740553,49784,45056,44984,44984,44984,0,80367],"quality":"More is better","isMain":true,"metricName":"Total length (>= 1000 bp)"},{"values":[44929,0,0,3304559,49784,45056,44984,44984,44984,0,80367],"quality":"More is better","isMain":false,"metricName":"Total length (>= 5000 bp)"},{"values":[44929,0,0,949763,49784,45056,44984,44984,44984,0,74102],"quality":"More is better","isMain":true,"metricName":"Total length (>= 10000 bp)"},{"values":[44929,0,0,64474,49784,45056,44984,44984,44984,0,74102],"quality":"More is better","isMain":false,"metricName":"Total length (>= 25000 bp)"},{"values":[0,0,0,0,0,0,0,0,0,0,74102],"quality":"More is better","isMain":true,"metricName":"Total length (>= 50000 bp)"},{"values":[44929,1692,808,3550,49784,45056,44984,44984,44984,651,74102],"quality":"More is better","isMain":false,"metricName":"N50"},{"values":[44929,1359,808,2359,49784,45056,44984,44984,44984,560,74102],"quality":"More is better","isMain":false,"metricName":"N75"},{"values":[1,9,1,790,1,1,1,1,1,7,1],"quality":"Less is better","isMain":false,"metricName":"L50"},{"values":[1,15,1,1640,1,1,1,1,1,11,1],"quality":"Less is better","isMain":false,"metricName":"L75"},{"values":["38.63","38.60","32.43","38.66","38.30","38.63","38.63","38.62","38.62","38.64","38.53"],"quality":"Equal","isMain":false,"metricName":"GC (%)"}]],["Predicted genes",[]],["Similarity statistics",[{"values":[0,0,0,0,0,0,0,0,0,0,0],"quality":"Equal","isMain":false,"metricName":"# similar correct contigs"},{"values":[0,0,0,0,0,0,0,0,0,0,0],"quality":"Equal","isMain":false,"metricName":"# similar misassembled blocks"}]],["Reference statistics",[{"values":[44929,44929,44929,44929,44929,44929,44929,44929,44929,44929,44929],"quality":"Equal","isMain":false,"metricName":"Reference length"},{"values":[1,1,1,1,1,1,1,1,1,1,1],"quality":"Equal","isMain":false,"metricName":"Reference fragments"},{"values":["38.63","38.63","38.63","38.63","38.63","38.63","38.63","38.63","38.63","38.63","38.63"],"quality":"Equal","isMain":false,"metricName":"Reference GC (%)"}]]],"referenceName":"gi\_200003956\_ref\_NC","date":"18 July 2018, Wednesday, 16:47:53","order":[0,1,2,3,4,5,6,7,8,9,10],"assembliesNames":["CLC","IDBA\_UD","MEGAHIT","MIRA","Ray\_Meta","SPAdes","SPAdes\_meta","SPAdes\_sc","SPAdes\_sc\_careful","Velvet","Geneious"]},{"assembliesWithNs":null,"minContig":500,"report":[["Genome statistics",[{"values":["27.088","18.225","79.909","77.237","80.339","51.199","67.357","84.916","86.969","86.390","85.572","19.926","9.567"],"quality":"More is better","isMain":true,"metricName":"Genome fraction (%)"},{"values":["1.002","1.000","1.025","1.013","1.011","1.014","1.009","1.010","1.009","1.011","1.008","1.000","1.000"],"quality":"Less is better","isMain":true,"metricName":"Duplication ratio"},{"values":[1810,2155,6695,6522,6443,4133,6695,6695,6695,6695,6695,2357,3915],"quality":"More is better","isMain":true,"metricName":"Largest alignment"},{"values":[65434,43957,194838,187125,195257,124931,163034,205858,211167,209916,207301,48039,23065],"quality":"More is better","isMain":true,"metricName":"Total aligned length"},{"values":[null,null,1728,1840,1973,602,1046,2392,2206,2356,2268,null,null],"quality":"More is better","isMain":false,"metricName":"NG50"},{"values":[null,null,864,841,851,null,null,1305,1162,1250,1226,null,null],"quality":"More is better","isMain":false,"metricName":"NG75"},{"values":[752,754,1798,2083,2103,1395,1304,2368,2356,2365,2356,1430,2624],"quality":"More is better","isMain":false,"metricName":"NA50"},{"values":[629,590,1129,1321,1320,974,1006,1304,1531,1328,1328,942,null],"quality":"More is better","isMain":false,"metricName":"NA75"},{"values":[null,null,1594,1642,1958,602,1046,2254,2168,2206,2168,null,null],"quality":"More is better","isMain":true,"metricName":"NGA50"},{"values":[null,null,733,660,839,null,null,1121,1149,1149,1119,null,null],"quality":"More is better","isMain":false,"metricName":"NGA75"},{"values":[null,null,43,42,41,91,73,31,37,35,35,null,null],"quality":"Less is better","isMain":false,"metricName":"LG50"},{"values":[null,null,91,88,88,null,null,63,74,70,71,null,null],"quality":"Less is better","isMain":false,"metricName":"LG75"},{"values":[31,22,36,33,32,31,41,32,33,32,32,14,5],"quality":"Less is better","isMain":false,"metricName":"LA50"},{"values":[54,39,71,63,60,58,76,63,60,61,60,24,null],"quality":"Less is better","isMain":false,"metricName":"LA75"},{"values":[null,null,47,45,42,91,73,37,39,37,38,null,null],"quality":"Less is better","isMain":true,"metricName":"LGA50"},{"values":[null,null,101,98,91,null,null,75,77,75,77,null,null],"quality":"Less is better","isMain":false,"metricName":"LGA75"}]],["Misassemblies",[{"values":[0,0,0,0,0,0,0,1,1,0,0,0,0],"quality":"Less is better","isMain":true,"metricName":"# misassemblies"},{"values":[0,0,0,0,0,0,0,1,1,0,0,0,0],"quality":"Less is better","isMain":false,"metricName":" # relocations"},{"values":[0,0,0,0,0,0,0,0,0,0,0,0,0],"quality":"Less is better","isMain":false,"metricName":" # translocations"},{"values":[0,0,0,0,0,0,0,0,0,0,0,0,0],"quality":"Less is better","isMain":false,"metricName":" # inversions"},{"values":[0,0,0,0,0,0,0,1,1,0,0,0,0],"quality":"Less is better","isMain":false,"metricName":"# misassembled contigs"},{"values":[0,0,0,0,0,0,0,8523,8523,0,0,0,0],"quality":"Less is better","isMain":true,"metricName":"Misassembled contigs length"},{"values":[0,0,3,0,0,0,0,1,0,1,1,0,0],"quality":"Less is better","isMain":false,"metricName":"# local misassemblies"},{"values":[0,0,0,0,0,0,0,0,0,0,0,0,0],"quality":"Less is better","isMain":false,"metricName":"# unaligned mis. contigs"}]],["Unaligned",[{"values":[0,0,0,0,0,0,0,0,0,0,0,0,0],"quality":"Less is better","isMain":false,"metricName":"# fully unaligned contigs"},{"values":[0,0,0,0,0,0,0,0,0,0,0,0,0],"quality":"Less is better","isMain":false,"metricName":"Fully unaligned length"},{"values":[0,0,6,6,2,0,0,10,1,6,6,0,2],"quality":"Less is better","isMain":false,"metricName":"# partially unaligned contigs"},{"values":[0,0,6490,7235,2159,0,0,13811,1564,6225,6225,0,8239],"quality":"Less is better","isMain":false,"metricName":"Partially unaligned length"}]],["Mismatches",[{"values":[11,3,591,394,442,53,91,599,243,498,345,11,196],"quality":"Less is better","isMain":false,"metricName":"# mismatches"},{"values":[0,0,38,14,14,3,11,29,3,12,5,0,13],"quality":"Less is better","isMain":false,"metricName":"# indels"},{"values":[0,0,57,24,19,6,558,62,6,17,8,0,18],"quality":"Less is better","isMain":false,"metricName":"Indels length"},{"values":["16.84","6.83","306.77","211.59","228.20","42.94","56.04","292.59","115.90","239.11","167.23","22.90","849.77"],"quality":"Less is better","isMain":true,"metricName":"# mismatches per 100 kbp"},{"values":["0.00","0.00","19.72","7.52","7.23","2.43","6.77","14.17","1.43","5.76","2.42","0.00","56.36"],"quality":"Less is better","isMain":true,"metricName":"# indels per 100 kbp"},{"values":[0,0,37,14,14,3,0,29,3,12,5,0,13],"quality":"Less is better","isMain":false,"metricName":" # indels (<= 5 bp)"},{"values":[0,0,1,0,0,0,11,0,0,0,0,0,0],"quality":"Less is better","isMain":false,"metricName":" # indels (> 5 bp)"},{"values":[0,0,0,0,0,133,839,0,0,0,0,0,48],"quality":"Less is better","isMain":false,"metricName":"# N's"},{"values":["0.00","0.00","0.00","0.00","0.00","106.28","511.93","0.00","0.00","0.00","0.00","0.00","153.34"],"quality":"Less is better","isMain":true,"metricName":"# N's per 100 kbp"}]],["Statistics without reference",[{"values":[82,59,125,109,113,99,132,105,113,110,109,40,9],"quality":"Equal","isMain":true,"metricName":"# contigs"},{"values":[15,8,82,80,76,53,76,83,85,85,84,21,9],"quality":"Equal","isMain":false,"metricName":"# contigs (>= 1000 bp)"},{"values":[0,0,4,2,2,0,1,5,2,3,3,0,1],"quality":"Equal","isMain":false,"metricName":"# contigs (>= 5000 bp)"},{"values":[0,0,0,0,0,0,0,0,0,0,0,0,0],"quality":"Equal","isMain":false,"metricName":"# contigs (>= 10000 bp)"},{"values":[0,0,0,0,0,0,0,0,0,0,0,0,0],"quality":"Equal","isMain":false,"metricName":"# contigs (>= 25000 bp)"},{"values":[0,0,0,0,0,0,0,0,0,0,0,0,0],"quality":"Equal","isMain":false,"metricName":"# contigs (>= 50000 bp)"},{"values":[1810,2155,6695,6522,6443,4338,6695,8523,8523,6695,6695,2357,9058],"quality":"More is better","isMain":true,"metricName":"Largest contig"},{"values":[65434,43957,203996,195861,198062,125136,163889,220615,213140,216691,214076,48042,31304],"quality":"More is better","isMain":true,"metricName":"Total length"},{"values":[19589,9868,172735,174337,170269,89907,124128,204533,193043,197980,195365,33957,31304],"quality":"More is better","isMain":true,"metricName":"Total length (>= 1000 bp)"},{"values":[0,0,23011,11625,11460,0,6695,35565,15218,18902,18902,0,9058],"quality":"More is better","isMain":false,"metricName":"Total length (>= 5000 bp)"},{"values":[0,0,0,0,0,0,0,0,0,0,0,0,0],"quality":"More is better","isMain":true,"metricName":"Total length (>= 10000 bp)"},{"values":[0,0,0,0,0,0,0,0,0,0,0,0,0],"quality":"More is better","isMain":false,"metricName":"Total length (>= 25000 bp)"},{"values":[0,0,0,0,0,0,0,0,0,0,0,0,0],"quality":"More is better","isMain":true,"metricName":"Total length (>= 50000 bp)"},{"values":[752,754,2100,2213,2119,1395,1321,2610,2368,2495,2495,1430,3616],"quality":"More is better","isMain":false,"metricName":"N50"},{"values":[629,590,1297,1451,1363,974,1010,1690,1594,1594,1594,942,2624],"quality":"More is better","isMain":false,"metricName":"N75"},{"values":[31,22,33,31,31,31,41,27,31,30,29,14,3],"quality":"Less is better","isMain":false,"metricName":"L50"},{"values":[54,39,65,58,58,58,75,53,58,57,56,24,6],"quality":"Less is better","isMain":false,"metricName":"L75"},{"values":["61.32","61.78","61.06","61.54","61.62","61.55","61.52","61.26","61.61","61.47","61.42","62.01","64.59"],"quality":"Equal","isMain":false,"metricName":"GC (%)"}]],["Predicted genes",[]],["Similarity statistics",[{"values":[0,0,0,0,0,0,0,0,0,0,0,0,0],"quality":"Equal","isMain":false,"metricName":"# similar correct contigs"},{"values":[0,0,0,0,0,0,0,0,0,0,0,0,0],"quality":"Equal","isMain":false,"metricName":"# similar misassembled blocks"}]],["Reference statistics",[{"values":[241087,241087,241087,241087,241087,241087,241087,241087,241087,241087,241087,241087,241087],"quality":"Equal","isMain":false,"metricName":"Reference length"},{"values":[1,1,1,1,1,1,1,1,1,1,1,1,1],"quality":"Equal","isMain":false,"metricName":"Reference fragments"},{"values":["61.70","61.70","61.70","61.70","61.70","61.70","61.70","61.70","61.70","61.70","61.70","61.70","61.70"],"quality":"Equal","isMain":false,"metricName":"Reference GC (%)"}]]],"referenceName":"gi\_20026600\_ref\_NC","date":"18 July 2018, Wednesday, 16:48:11","order":[0,1,2,3,4,5,6,7,8,9,10,11,12],"assembliesNames":["ABySS\_127","ABySS\_63","CLC","IDBA\_UD","MEGAHIT","MIRA","SOAPdenovo2","SPAdes","SPAdes\_meta","SPAdes\_sc","SPAdes\_sc\_careful","Velvet","Geneious"]},{"assembliesWithNs":null,"minContig":500,"report":[["Genome statistics",[{"values":["100.000","100.000","100.000","100.000","100.000","100.000","100.000","100.000","100.000","100.000","100.000","100.000"],"quality":"More is better","isMain":true,"metricName":"Genome fraction (%)"},{"values":["1.010","1.002","1.000","1.003","1.004","1.024","1.001","1.004","1.002","1.002","1.002","1.001"],"quality":"Less is better","isMain":true,"metricName":"Duplication ratio"},{"values":[33845,33569,33507,33606,33648,34301,33556,33634,33562,33562,33562,33537],"quality":"More is better","isMain":true,"metricName":"Largest alignment"},{"values":[33845,33569,33507,33606,33648,34301,33556,33634,33562,33562,33562,33537],"quality":"More is better","isMain":true,"metricName":"Total aligned length"},{"values":[33845,33569,33507,33606,33648,34301,33556,33634,33562,33562,33562,33537],"quality":"More is better","isMain":false,"metricName":"NG50"},{"values":[33845,33569,33507,33606,33648,34301,33556,33634,33562,33562,33562,33537],"quality":"More is better","isMain":false,"metricName":"NG75"},{"values":[33845,33569,33507,33606,33648,34301,33556,33634,33562,33562,33562,33537],"quality":"More is better","isMain":false,"metricName":"NA50"},{"values":[33845,33569,33507,33606,33648,34301,33556,33634,33562,33562,33562,33537],"quality":"More is better","isMain":false,"metricName":"NA75"},{"values":[33845,33569,33507,33606,33648,34301,33556,33634,33562,33562,33562,33537],"quality":"More is better","isMain":true,"metricName":"NGA50"},{"values":[33845,33569,33507,33606,33648,34301,33556,33634,33562,33562,33562,33537],"quality":"More is better","isMain":false,"metricName":"NGA75"},{"values":[1,1,1,1,1,1,1,1,1,1,1,1],"quality":"Less is better","isMain":false,"metricName":"LG50"},{"values":[1,1,1,1,1,1,1,1,1,1,1,1],"quality":"Less is better","isMain":false,"metricName":"LG75"},{"values":[1,1,1,1,1,1,1,1,1,1,1,1],"quality":"Less is better","isMain":false,"metricName":"LA50"},{"values":[1,1,1,1,1,1,1,1,1,1,1,1],"quality":"Less is better","isMain":false,"metricName":"LA75"},{"values":[1,1,1,1,1,1,1,1,1,1,1,1],"quality":"Less is better","isMain":true,"metricName":"LGA50"},{"values":[1,1,1,1,1,1,1,1,1,1,1,1],"quality":"Less is better","isMain":false,"metricName":"LGA75"}]],["Misassemblies",[{"values":[0,0,0,0,0,0,0,0,0,0,0,0],"quality":"Less is better","isMain":true,"metricName":"# misassemblies"},{"values":[0,0,0,0,0,0,0,0,0,0,0,0],"quality":"Less is better","isMain":false,"metricName":" # relocations"},{"values":[0,0,0,0,0,0,0,0,0,0,0,0],"quality":"Less is better","isMain":false,"metricName":" # translocations"},{"values":[0,0,0,0,0,0,0,0,0,0,0,0],"quality":"Less is better","isMain":false,"metricName":" # inversions"},{"values":[0,0,0,0,0,0,0,0,0,0,0,0],"quality":"Less is better","isMain":false,"metricName":"# misassembled contigs"},{"values":[0,0,0,0,0,0,0,0,0,0,0,0],"quality":"Less is better","isMain":true,"metricName":"Misassembled contigs length"},{"values":[1,0,0,1,1,1,0,1,0,0,0,0],"quality":"Less is better","isMain":false,"metricName":"# local misassemblies"},{"values":[0,0,0,0,0,0,0,0,0,0,0,0],"quality":"Less is better","isMain":false,"metricName":"# unaligned mis. contigs"}]],["Unaligned",[{"values":[0,0,0,0,0,0,0,0,0,0,0,0],"quality":"Less is better","isMain":false,"metricName":"# fully unaligned contigs"},{"values":[0,0,0,0,0,0,0,0,0,0,0,0],"quality":"Less is better","isMain":false,"metricName":"Fully unaligned length"},{"values":[0,0,0,0,0,0,0,0,0,0,0,0],"quality":"Less is better","isMain":false,"metricName":"# partially unaligned contigs"},{"values":[0,0,0,0,0,0,0,0,0,0,0,0],"quality":"Less is better","isMain":false,"metricName":"Partially unaligned length"}]],["Mismatches",[{"values":[0,0,0,0,0,0,0,0,0,0,0,3],"quality":"Less is better","isMain":false,"metricName":"# mismatches"},{"values":[0,1,0,0,0,0,1,0,1,1,1,1],"quality":"Less is better","isMain":false,"metricName":"# indels"},{"values":[0,62,0,0,0,0,49,0,55,55,55,30],"quality":"Less is better","isMain":false,"metricName":"Indels length"},{"values":["0.00","0.00","0.00","0.00","0.00","0.00","0.00","0.00","0.00","0.00","0.00","8.95"],"quality":"Less is better","isMain":true,"metricName":"# mismatches per 100 kbp"},{"values":["0.00","2.98","0.00","0.00","0.00","0.00","2.98","0.00","2.98","2.98","2.98","2.98"],"quality":"Less is better","isMain":true,"metricName":"# indels per 100 kbp"},{"values":[0,0,0,0,0,0,0,0,0,0,0,0],"quality":"Less is better","isMain":false,"metricName":" # indels (<= 5 bp)"},{"values":[0,1,0,0,0,0,1,0,1,1,1,1],"quality":"Less is better","isMain":false,"metricName":" # indels (> 5 bp)"},{"values":[0,29,0,0,0,0,0,0,0,0,0,0],"quality":"Less is better","isMain":false,"metricName":"# N's"},{"values":["0.00","86.39","0.00","0.00","0.00","0.00","0.00","0.00","0.00","0.00","0.00","0.00"],"quality":"Less is better","isMain":true,"metricName":"# N's per 100 kbp"}]],["Statistics without reference",[{"values":[1,1,1,1,1,1,1,1,1,1,1,1],"quality":"Equal","isMain":true,"metricName":"# contigs"},{"values":[1,1,1,1,1,1,1,1,1,1,1,1],"quality":"Equal","isMain":false,"metricName":"# contigs (>= 1000 bp)"},{"values":[1,1,1,1,1,1,1,1,1,1,1,1],"quality":"Equal","isMain":false,"metricName":"# contigs (>= 5000 bp)"},{"values":[1,1,1,1,1,1,1,1,1,1,1,1],"quality":"Equal","isMain":false,"metricName":"# contigs (>= 10000 bp)"},{"values":[1,1,1,1,1,1,1,1,1,1,1,1],"quality":"Equal","isMain":false,"metricName":"# contigs (>= 25000 bp)"},{"values":[0,0,0,0,0,0,0,0,0,0,0,0],"quality":"Equal","isMain":false,"metricName":"# contigs (>= 50000 bp)"},{"values":[33845,33569,33507,33606,33648,34301,33556,33634,33562,33562,33562,33537],"quality":"More is better","isMain":true,"metricName":"Largest contig"},{"values":[33845,33569,33507,33606,33648,34301,33556,33634,33562,33562,33562,33537],"quality":"More is better","isMain":true,"metricName":"Total length"},{"values":[33845,33569,33507,33606,33648,34301,33556,33634,33562,33562,33562,33537],"quality":"More is better","isMain":true,"metricName":"Total length (>= 1000 bp)"},{"values":[33845,33569,33507,33606,33648,34301,33556,33634,33562,33562,33562,33537],"quality":"More is better","isMain":false,"metricName":"Total length (>= 5000 bp)"},{"values":[33845,33569,33507,33606,33648,34301,33556,33634,33562,33562,33562,33537],"quality":"More is better","isMain":true,"metricName":"Total length (>= 10000 bp)"},{"values":[33845,33569,33507,33606,33648,34301,33556,33634,33562,33562,33562,33537],"quality":"More is better","isMain":false,"metricName":"Total length (>= 25000 bp)"},{"values":[0,0,0,0,0,0,0,0,0,0,0,0],"quality":"More is better","isMain":true,"metricName":"Total length (>= 50000 bp)"},{"values":[33845,33569,33507,33606,33648,34301,33556,33634,33562,33562,33562,33537],"quality":"More is better","isMain":false,"metricName":"N50"},{"values":[33845,33569,33507,33606,33648,34301,33556,33634,33562,33562,33562,33537],"quality":"More is better","isMain":false,"metricName":"N75"},{"values":[1,1,1,1,1,1,1,1,1,1,1,1],"quality":"Less is better","isMain":false,"metricName":"L50"},{"values":[1,1,1,1,1,1,1,1,1,1,1,1],"quality":"Less is better","isMain":false,"metricName":"L75"},{"values":["28.41","28.38","28.38","28.37","28.37","28.48","28.39","28.41","28.37","28.38","28.38","28.38"],"quality":"Equal","isMain":false,"metricName":"GC (%)"}]],["Predicted genes",[]],["Similarity statistics",[{"values":[0,0,0,0,0,0,0,0,0,0,0,0],"quality":"Equal","isMain":false,"metricName":"# similar correct contigs"},{"values":[0,0,0,0,0,0,0,0,0,0,0,0],"quality":"Equal","isMain":false,"metricName":"# similar misassembled blocks"}]],["Reference statistics",[{"values":[33507,33507,33507,33507,33507,33507,33507,33507,33507,33507,33507,33507],"quality":"Equal","isMain":false,"metricName":"Reference length"},{"values":[1,1,1,1,1,1,1,1,1,1,1,1],"quality":"Equal","isMain":false,"metricName":"Reference fragments"},{"values":["28.38","28.38","28.38","28.38","28.38","28.38","28.38","28.38","28.38","28.38","28.38","28.38"],"quality":"Equal","isMain":false,"metricName":"Reference GC (%)"}]]],"referenceName":"gi\_20065963\_ref\_NC","date":"18 July 2018, Wednesday, 16:48:24","order":[0,1,2,3,4,5,6,7,8,9,10,11],"assembliesNames":["ABySS\_127","ABySS\_63","CLC","IDBA\_UD","MEGAHIT","MIRA","Ray\_Meta","SPAdes","SPAdes\_meta","SPAdes\_sc","SPAdes\_sc\_careful","Velvet"]},{"assembliesWithNs":null,"minContig":500,"report":[["Genome statistics",[{"values":["97.001","95.451","92.181","99.624","99.747","98.843","0.738","36.356","99.832","99.932","96.259","95.566","92.990","94.391"],"quality":"More is better","isMain":true,"metricName":"Genome fraction (%)"},{"values":["1.002","1.006","1.006","1.001","1.006","1.016","1.000","1.171","1.000","1.006","1.017","1.017","1.005","1.002"],"quality":"Less is better","isMain":true,"metricName":"Duplication ratio"},{"values":[15131,9262,16103,35875,35067,31244,609,3130,117672,24531,8313,8313,10209,48131],"quality":"More is better","isMain":true,"metricName":"Largest alignment"},{"values":[150757,148892,143698,154699,155575,155660,1144,56889,154800,155954,150855,149708,144922,146439],"quality":"More is better","isMain":true,"metricName":"Total aligned length"},{"values":[6259,3852,11869,24484,26979,17481,null,598,117672,12434,4089,3888,3264,17503],"quality":"More is better","isMain":false,"metricName":"NG50"},{"values":[2887,1994,4923,15412,18973,13681,null,null,117672,8328,2911,2409,1968,14913],"quality":"More is better","isMain":false,"metricName":"NG75"},{"values":[6938,3864,4772,24484,26979,17481,609,500,117672,12393,4075,3888,3457,17503],"quality":"More is better","isMain":false,"metricName":"NA50"},{"values":[3082,2132,1923,15412,18937,13681,535,null,117672,8328,3114,2549,2276,13691],"quality":"More is better","isMain":false,"metricName":"NA75"},{"values":[6259,3852,5114,24484,26979,17481,null,null,117672,12393,4075,3805,3264,17503],"quality":"More is better","isMain":true,"metricName":"NGA50"},{"values":[2887,1994,3091,15412,18937,13681,null,null,117672,8328,2911,2393,1968,13691],"quality":"More is better","isMain":false,"metricName":"NGA75"},{"values":[8,16,4,3,3,4,null,66,1,5,14,15,15,3],"quality":"Less is better","isMain":false,"metricName":"LG50"},{"values":[17,29,9,5,5,6,null,null,1,9,25,27,29,5],"quality":"Less is better","isMain":false,"metricName":"LG75"},{"values":[7,15,10,3,3,4,1,48,1,5,14,14,13,3],"quality":"Less is better","isMain":false,"metricName":"LA50"},{"values":[16,27,23,5,5,6,2,null,1,9,24,26,26,5],"quality":"Less is better","isMain":false,"metricName":"LA75"},{"values":[8,16,8,3,3,4,null,null,1,5,14,15,15,3],"quality":"Less is better","isMain":true,"metricName":"LGA50"},{"values":[17,29,17,5,5,6,null,null,1,9,25,27,29,5],"quality":"Less is better","isMain":false,"metricName":"LGA75"}]],["Misassemblies",[{"values":[0,0,0,0,0,0,0,0,0,0,0,0,0,0],"quality":"Less is better","isMain":true,"metricName":"# misassemblies"},{"values":[0,0,0,0,0,0,0,0,0,0,0,0,0,0],"quality":"Less is better","isMain":false,"metricName":" # relocations"},{"values":[0,0,0,0,0,0,0,0,0,0,0,0,0,0],"quality":"Less is better","isMain":false,"metricName":" # translocations"},{"values":[0,0,0,0,0,0,0,0,0,0,0,0,0,0],"quality":"Less is better","isMain":false,"metricName":" # inversions"},{"values":[0,0,0,0,0,0,0,0,0,0,0,0,0,0],"quality":"Less is better","isMain":false,"metricName":"# misassembled contigs"},{"values":[0,0,0,0,0,0,0,0,0,0,0,0,0,0],"quality":"Less is better","isMain":true,"metricName":"Misassembled contigs length"},{"values":[0,0,0,1,0,0,0,1,0,0,0,0,1,0],"quality":"Less is better","isMain":false,"metricName":"# local misassemblies"},{"values":[0,0,0,0,0,0,0,12,0,0,0,0,0,0],"quality":"Less is better","isMain":false,"metricName":"# unaligned mis. contigs"}]],["Unaligned",[{"values":[0,0,0,0,0,0,0,0,0,0,0,0,0,0],"quality":"Less is better","isMain":false,"metricName":"# fully unaligned contigs"},{"values":[0,0,0,0,0,0,0,0,0,0,0,0,0,0],"quality":"Less is better","isMain":false,"metricName":"Fully unaligned length"},{"values":[0,0,2,1,0,0,0,22,0,1,0,0,0,3],"quality":"Less is better","isMain":false,"metricName":"# partially unaligned contigs"},{"values":[0,0,30288,1290,0,0,0,21279,0,576,0,0,0,5810],"quality":"Less is better","isMain":false,"metricName":"Partially unaligned length"}]],["Mismatches",[{"values":[2,3,70,22,5,9,0,0,44,80,60,35,57,37],"quality":"Less is better","isMain":false,"metricName":"# mismatches"},{"values":[0,0,3,0,0,0,0,52,2,2,2,2,8,35],"quality":"Less is better","isMain":false,"metricName":"# indels"},{"values":[0,0,4,0,0,0,0,2518,2,2,2,2,14,78],"quality":"Less is better","isMain":false,"metricName":"Indels length"},{"values":["1.33","2.03","48.97","14.24","3.23","5.87","0.00","0.00","28.42","51.63","40.20","23.62","39.53","25.28"],"quality":"Less is better","isMain":true,"metricName":"# mismatches per 100 kbp"},{"values":["0.00","0.00","2.10","0.00","0.00","0.00","0.00","92.24","1.29","1.29","1.34","1.35","5.55","23.91"],"quality":"Less is better","isMain":true,"metricName":"# indels per 100 kbp"},{"values":[0,0,3,0,0,0,0,0,2,2,2,2,7,33],"quality":"Less is better","isMain":false,"metricName":" # indels (<= 5 bp)"},{"values":[0,0,0,0,0,0,0,52,0,0,0,0,1,2],"quality":"Less is better","isMain":false,"metricName":" # indels (> 5 bp)"},{"values":[0,0,4,0,0,41,0,16646,0,576,223,222,29,2466],"quality":"Less is better","isMain":false,"metricName":"# N's"},{"values":["0.00","0.00","2.30","0.00","0.00","26.34","0.00","19071.95","0.00","367.98","146.93","147.35","20.01","1618.16"],"quality":"Less is better","isMain":true,"metricName":"# N's per 100 kbp"}]],["Statistics without reference",[{"values":[36,56,36,8,9,14,2,83,2,20,50,54,52,9],"quality":"Equal","isMain":true,"metricName":"# contigs"},{"values":[32,46,30,8,9,12,0,31,2,18,41,45,45,9],"quality":"Equal","isMain":false,"metricName":"# contigs (>= 1000 bp)"},{"values":[11,6,8,7,7,9,0,0,2,10,10,9,7,8],"quality":"Equal","isMain":false,"metricName":"# contigs (>= 5000 bp)"},{"values":[3,0,4,6,5,7,0,0,2,7,0,0,1,6],"quality":"Equal","isMain":false,"metricName":"# contigs (>= 10000 bp)"},{"values":[0,0,1,2,3,1,0,0,2,0,0,0,0,1],"quality":"Equal","isMain":false,"metricName":"# contigs (>= 25000 bp)"},{"values":[0,0,0,0,0,0,0,0,1,0,0,0,0,0],"quality":"Equal","isMain":false,"metricName":"# contigs (>= 50000 bp)"},{"values":[15131,9262,43092,35875,35067,31244,609,3130,117672,24531,8313,8313,10209,48131],"quality":"More is better","isMain":true,"metricName":"Largest contig"},{"values":[150757,148928,174047,155989,155614,155660,1144,87280,154800,156530,151771,150657,144951,152395],"quality":"More is better","isMain":true,"metricName":"Total length"},{"values":[147187,141183,170250,155989,155614,154135,0,50270,154800,155208,145071,143918,139721,152395],"quality":"More is better","isMain":true,"metricName":"Total length (>= 1000 bp)"},{"values":[98746,39817,112277,153258,147942,144270,0,0,154800,126995,62003,55472,49140,148369],"quality":"More is better","isMain":false,"metricName":"Total length (>= 5000 bp)"},{"values":[41825,0,85465,143456,131181,131700,0,0,154800,103112,0,0,10209,132752],"quality":"More is better","isMain":true,"metricName":"Total length (>= 10000 bp)"},{"values":[0,0,43092,70467,92513,31244,0,0,154800,0,0,0,0,48131],"quality":"More is better","isMain":false,"metricName":"Total length (>= 25000 bp)"},{"values":[0,0,0,0,0,0,0,0,117672,0,0,0,0,0],"quality":"More is better","isMain":true,"metricName":"Total length (>= 50000 bp)"},{"values":[6938,3864,8613,24484,26979,17481,609,1127,117672,12434,4089,3907,3486,17503],"quality":"More is better","isMain":false,"metricName":"N50"},{"values":[3082,2132,4187,15412,18973,13681,535,804,117672,8328,3114,2549,2276,14913],"quality":"More is better","isMain":false,"metricName":"N75"},{"values":[7,15,5,3,3,4,1,25,1,5,14,14,13,3],"quality":"Less is better","isMain":false,"metricName":"L50"},{"values":[16,27,12,5,5,6,2,49,1,9,24,26,26,5],"quality":"Less is better","isMain":false,"metricName":"L75"},{"values":["41.69","41.79","41.21","41.73","41.69","41.65","44.14","41.24","41.68","41.66","41.67","41.67","41.81","41.59"],"quality":"Equal","isMain":false,"metricName":"GC (%)"}]],["Predicted genes",[]],["Similarity statistics",[{"values":[0,0,0,0,0,0,0,0,0,0,0,0,0,0],"quality":"Equal","isMain":false,"metricName":"# similar correct contigs"},{"values":[0,0,0,0,0,0,0,0,0,0,0,0,0,0],"quality":"Equal","isMain":false,"metricName":"# similar misassembled blocks"}]],["Reference statistics",[{"values":[155060,155060,155060,155060,155060,155060,155060,155060,155060,155060,155060,155060,155060,155060],"quality":"Equal","isMain":false,"metricName":"Reference length"},{"values":[1,1,1,1,1,1,1,1,1,1,1,1,1,1],"quality":"Equal","isMain":false,"metricName":"Reference fragments"},{"values":["41.68","41.68","41.68","41.68","41.68","41.68","41.68","41.68","41.68","41.68","41.68","41.68","41.68","41.68"],"quality":"Equal","isMain":false,"metricName":"Reference GC (%)"}]]],"referenceName":"gi\_20069879\_ref\_NC","date":"18 July 2018, Wednesday, 16:48:43","order":[0,1,2,3,4,5,6,7,8,9,10,11,12,13],"assembliesNames":["ABySS\_127","ABySS\_63","CLC","IDBA\_UD","MEGAHIT","MIRA","Ray\_Meta","SOAPdenovo2","SPAdes","SPAdes\_meta","SPAdes\_sc","SPAdes\_sc\_careful","Velvet","Geneious"]},{"assembliesWithNs":null,"minContig":500,"report":[["Genome statistics",[{"values":["99.784","99.251","99.366","99.580","99.738","99.941","17.611","79.783","100.000","100.000","99.501","99.428","98.587","99.749"],"quality":"More is better","isMain":true,"metricName":"Genome fraction (%)"},{"values":["1.001","1.000","1.001","1.001","1.001","1.005","1.001","1.016","1.001","1.001","1.000","1.000","1.001","1.001"],"quality":"Less is better","isMain":true,"metricName":"Duplication ratio"},{"values":[97511,48489,68530,97323,91393,229827,1172,45336,230375,230187,136062,136054,62635,229827],"quality":"More is better","isMain":true,"metricName":"Largest alignment"},{"values":[229806,228519,228919,229339,229795,231118,40558,186199,230375,230187,228885,228877,227095,229827],"quality":"More is better","isMain":true,"metricName":"Total aligned length"},{"values":[48617,38900,60356,76598,58010,229827,null,23655,230475,230308,136072,136054,33731,229827],"quality":"More is better","isMain":false,"metricName":"NG50"},{"values":[47008,23490,57520,76598,48647,229827,null,15362,230475,230308,92059,92059,14717,229827],"quality":"More is better","isMain":false,"metricName":"NG75"},{"values":[48617,38900,60356,76598,58010,229827,691,15856,230375,230187,136062,136054,33731,229827],"quality":"More is better","isMain":false,"metricName":"NA50"},{"values":[47008,23490,57520,76598,48647,229827,584,2730,230375,230187,92059,92059,14717,229827],"quality":"More is better","isMain":false,"metricName":"NA75"},{"values":[48617,38900,60356,76598,58010,229827,null,15856,230375,230187,136062,136054,33731,229827],"quality":"More is better","isMain":true,"metricName":"NGA50"},{"values":[47008,23490,57520,76598,48647,229827,null,4941,230375,230187,92059,92059,14717,229827],"quality":"More is better","isMain":false,"metricName":"NGA75"},{"values":[2,3,2,2,2,1,null,4,1,1,1,1,3,1],"quality":"Less is better","isMain":false,"metricName":"LG50"},{"values":[3,5,3,2,3,1,null,7,1,1,2,2,5,1],"quality":"Less is better","isMain":false,"metricName":"LG75"},{"values":[2,3,2,2,2,1,24,5,1,1,1,1,3,1],"quality":"Less is better","isMain":false,"metricName":"LA50"},{"values":[3,5,3,2,3,1,40,11,1,1,2,2,5,1],"quality":"Less is better","isMain":false,"metricName":"LA75"},{"values":[2,3,2,2,2,1,null,5,1,1,1,1,3,1],"quality":"Less is better","isMain":true,"metricName":"LGA50"},{"values":[3,5,3,2,3,1,null,10,1,1,2,2,5,1],"quality":"Less is better","isMain":false,"metricName":"LGA75"}]],["Misassemblies",[{"values":[0,0,0,0,0,0,0,0,0,0,0,0,0,0],"quality":"Less is better","isMain":true,"metricName":"# misassemblies"},{"values":[0,0,0,0,0,0,0,0,0,0,0,0,0,0],"quality":"Less is better","isMain":false,"metricName":" # relocations"},{"values":[0,0,0,0,0,0,0,0,0,0,0,0,0,0],"quality":"Less is better","isMain":false,"metricName":" # translocations"},{"values":[0,0,0,0,0,0,0,0,0,0,0,0,0,0],"quality":"Less is better","isMain":false,"metricName":" # inversions"},{"values":[0,0,0,0,0,0,0,0,0,0,0,0,0,0],"quality":"Less is better","isMain":false,"metricName":"# misassembled contigs"},{"values":[0,0,0,0,0,0,0,0,0,0,0,0,0,0],"quality":"Less is better","isMain":true,"metricName":"Misassembled contigs length"},{"values":[0,0,1,0,1,1,0,10,1,1,3,3,0,1],"quality":"Less is better","isMain":false,"metricName":"# local misassemblies"},{"values":[0,0,0,0,0,0,0,3,0,0,0,0,0,0],"quality":"Less is better","isMain":false,"metricName":"# unaligned mis. contigs"}]],["Unaligned",[{"values":[0,0,0,0,0,0,0,0,0,0,0,0,0,0],"quality":"Less is better","isMain":false,"metricName":"# fully unaligned contigs"},{"values":[0,0,0,0,0,0,0,0,0,0,0,0,0,0],"quality":"Less is better","isMain":false,"metricName":"Fully unaligned length"},{"values":[0,0,1,0,0,0,0,12,0,0,0,0,0,0],"quality":"Less is better","isMain":false,"metricName":"# partially unaligned contigs"},{"values":[0,0,749,0,0,0,0,50361,0,0,0,0,0,0],"quality":"Less is better","isMain":false,"metricName":"Partially unaligned length"}]],["Mismatches",[{"values":[6,0,2,0,3,5,0,2,1,1,0,2,0,5],"quality":"Less is better","isMain":false,"metricName":"# mismatches"},{"values":[0,0,0,1,0,5,0,116,0,1,2,4,0,5],"quality":"Less is better","isMain":false,"metricName":"# indels"},{"values":[0,0,0,25,0,145,0,5597,0,70,131,125,0,145],"quality":"Less is better","isMain":false,"metricName":"Indels length"},{"values":["2.61","0.00","0.87","0.00","1.31","2.17","0.00","1.09","0.43","0.43","0.00","0.87","0.00","2.18"],"quality":"Less is better","isMain":true,"metricName":"# mismatches per 100 kbp"},{"values":["0.00","0.00","0.00","0.44","0.00","2.17","0.00","63.18","0.00","0.43","0.87","1.75","0.00","2.18"],"quality":"Less is better","isMain":true,"metricName":"# indels per 100 kbp"},{"values":[0,0,0,0,0,3,0,0,0,0,0,1,0,3],"quality":"Less is better","isMain":false,"metricName":" # indels (<= 5 bp)"},{"values":[0,0,0,1,0,2,0,116,0,1,2,3,0,2],"quality":"Less is better","isMain":false,"metricName":" # indels (> 5 bp)"},{"values":[0,0,0,0,0,0,0,17706,100,100,20,0,0,75],"quality":"Less is better","isMain":false,"metricName":"# N's"},{"values":["0.00","0.00","0.00","0.00","0.00","0.00","0.00","7473.98","43.39","43.42","8.74","0.00","0.00","32.63"],"quality":"Less is better","isMain":true,"metricName":"# N's per 100 kbp"}]],["Statistics without reference",[{"values":[7,11,6,6,6,3,58,15,1,1,3,3,21,1],"quality":"Equal","isMain":true,"metricName":"# contigs"},{"values":[6,9,5,5,5,1,4,14,1,1,2,2,14,1],"quality":"Equal","isMain":false,"metricName":"# contigs (>= 1000 bp)"},{"values":[6,9,5,4,5,1,0,12,1,1,2,2,8,1],"quality":"Equal","isMain":false,"metricName":"# contigs (>= 5000 bp)"},{"values":[5,7,4,3,5,1,0,8,1,1,2,2,5,1],"quality":"Equal","isMain":false,"metricName":"# contigs (>= 10000 bp)"},{"values":[3,4,4,3,3,1,0,3,1,1,2,2,4,1],"quality":"Equal","isMain":false,"metricName":"# contigs (>= 25000 bp)"},{"values":[1,0,3,2,2,1,0,0,1,1,2,2,1,1],"quality":"Equal","isMain":false,"metricName":"# contigs (>= 50000 bp)"},{"values":[97511,48489,68530,97323,91393,229827,1172,48474,230475,230308,136072,136054,62635,229827],"quality":"More is better","isMain":true,"metricName":"Largest contig"},{"values":[229806,228519,229668,229339,229795,231118,40558,236902,230475,230308,228895,228877,227095,229827],"quality":"More is better","isMain":true,"metricName":"Total length"},{"values":[229301,226996,228744,228834,229290,229827,4499,236325,230475,230308,228131,228113,222626,229827],"quality":"More is better","isMain":true,"metricName":"Total length (>= 1000 bp)"},{"values":[229301,226996,228744,227677,229290,229827,0,229886,230475,230308,228131,228113,203852,229827],"quality":"More is better","isMain":false,"metricName":"Total length (>= 5000 bp)"},{"values":[223906,216597,223143,222484,229290,229827,0,201273,230475,230308,228131,228113,183123,229827],"quality":"More is better","isMain":true,"metricName":"Total length (>= 10000 bp)"},{"values":[193136,165966,223143,222484,198050,229827,0,109118,230475,230308,228131,228113,168406,229827],"quality":"More is better","isMain":false,"metricName":"Total length (>= 25000 bp)"},{"values":[97511,0,186406,173921,149403,229827,0,0,230475,230308,228131,228113,62635,229827],"quality":"More is better","isMain":true,"metricName":"Total length (>= 50000 bp)"},{"values":[48617,38900,60356,76598,58010,229827,691,23655,230475,230308,136072,136054,33731,229827],"quality":"More is better","isMain":false,"metricName":"N50"},{"values":[47008,23490,57520,76598,48647,229827,584,15362,230475,230308,92059,92059,14717,229827],"quality":"More is better","isMain":false,"metricName":"N75"},{"values":[2,3,2,2,2,1,24,4,1,1,1,1,3,1],"quality":"Less is better","isMain":false,"metricName":"L50"},{"values":[3,5,3,2,3,1,40,7,1,1,2,2,5,1],"quality":"Less is better","isMain":false,"metricName":"L75"},{"values":["60.99","61.00","61.01","61.00","60.99","61.03","59.51","61.02","61.01","61.01","60.99","60.99","60.97","60.96"],"quality":"Equal","isMain":false,"metricName":"GC (%)"}]],["Predicted genes",[]],["Similarity statistics",[{"values":[0,0,0,0,0,0,0,0,0,0,0,0,0,0],"quality":"Equal","isMain":false,"metricName":"# similar correct contigs"},{"values":[0,0,0,0,0,0,0,0,0,0,0,0,0,0],"quality":"Equal","isMain":false,"metricName":"# similar misassembled blocks"}]],["Reference statistics",[{"values":[230138,230138,230138,230138,230138,230138,230138,230138,230138,230138,230138,230138,230138,230138],"quality":"Equal","isMain":false,"metricName":"Reference length"},{"values":[1,1,1,1,1,1,1,1,1,1,1,1,1,1],"quality":"Equal","isMain":false,"metricName":"Reference fragments"},{"values":["61.01","61.01","61.01","61.01","61.01","61.01","61.01","61.01","61.01","61.01","61.01","61.01","61.01","61.01"],"quality":"Equal","isMain":false,"metricName":"Reference GC (%)"}]]],"referenceName":"gi\_20198505\_ref\_NC","date":"18 July 2018, Wednesday, 16:49:01","order":[0,1,2,3,4,5,6,7,8,9,10,11,12,13],"assembliesNames":["ABySS\_127","ABySS\_63","CLC","IDBA\_UD","MEGAHIT","MIRA","Ray\_Meta","SOAPdenovo2","SPAdes","SPAdes\_meta","SPAdes\_sc","SPAdes\_sc\_careful","Velvet","Geneious"]},{"assembliesWithNs":null,"minContig":500,"report":[["Genome statistics",[{"values":["100.000","100.000","99.841","100.000","100.000","82.519","66.584","100.000","99.993","99.993","99.993","99.251"],"quality":"More is better","isMain":true,"metricName":"Genome fraction (%)"},{"values":["1.000","1.000","1.000","1.001","1.002","1.063","1.030","1.001","1.000","1.000","1.000","1.002"],"quality":"Less is better","isMain":true,"metricName":"Duplication ratio"},{"values":[164604,164602,164341,164743,164886,5699,3785,164729,164590,164590,164590,42860],"quality":"More is better","isMain":true,"metricName":"Largest alignment"},{"values":[164604,164602,164341,164743,164886,144341,110104,164729,164590,164590,164590,163637],"quality":"More is better","isMain":true,"metricName":"Total aligned length"},{"values":[164604,164602,164341,164743,164886,1397,1004,164729,164590,164590,164590,16208],"quality":"More is better","isMain":false,"metricName":"NG50"},{"values":[164604,164602,164341,164743,164886,971,null,164729,164590,164590,164590,6798],"quality":"More is better","isMain":false,"metricName":"NG75"},{"values":[164604,164602,164341,164743,164886,1529,1344,164729,164590,164590,164590,16481],"quality":"More is better","isMain":false,"metricName":"NA50"},{"values":[164604,164602,164341,164743,164886,1129,750,164729,164590,164590,164590,6798],"quality":"More is better","isMain":false,"metricName":"NA75"},{"values":[164604,164602,164341,164743,164886,1397,819,164729,164590,164590,164590,16208],"quality":"More is better","isMain":true,"metricName":"NGA50"},{"values":[164604,164602,164341,164743,164886,971,null,164729,164590,164590,164590,6798],"quality":"More is better","isMain":false,"metricName":"NGA75"},{"values":[1,1,1,1,1,34,46,1,1,1,1,4],"quality":"Less is better","isMain":false,"metricName":"LG50"},{"values":[1,1,1,1,1,68,null,1,1,1,1,7],"quality":"Less is better","isMain":false,"metricName":"LG75"},{"values":[1,1,1,1,1,27,30,1,1,1,1,3],"quality":"Less is better","isMain":false,"metricName":"LA50"},{"values":[1,1,1,1,1,54,59,1,1,1,1,7],"quality":"Less is better","isMain":false,"metricName":"LA75"},{"values":[1,1,1,1,1,34,52,1,1,1,1,4],"quality":"Less is better","isMain":true,"metricName":"LGA50"},{"values":[1,1,1,1,1,68,null,1,1,1,1,7],"quality":"Less is better","isMain":false,"metricName":"LGA75"}]],["Misassemblies",[{"values":[0,0,0,0,0,0,0,0,0,0,0,0],"quality":"Less is better","isMain":true,"metricName":"# misassemblies"},{"values":[0,0,0,0,0,0,0,0,0,0,0,0],"quality":"Less is better","isMain":false,"metricName":" # relocations"},{"values":[0,0,0,0,0,0,0,0,0,0,0,0],"quality":"Less is better","isMain":false,"metricName":" # translocations"},{"values":[0,0,0,0,0,0,0,0,0,0,0,0],"quality":"Less is better","isMain":false,"metricName":" # inversions"},{"values":[0,0,0,0,0,0,0,0,0,0,0,0],"quality":"Less is better","isMain":false,"metricName":"# misassembled contigs"},{"values":[0,0,0,0,0,0,0,0,0,0,0,0],"quality":"Less is better","isMain":true,"metricName":"Misassembled contigs length"},{"values":[0,0,0,1,1,0,0,1,0,0,0,0],"quality":"Less is better","isMain":false,"metricName":"# local misassemblies"},{"values":[0,0,0,0,0,0,0,0,0,0,0,0],"quality":"Less is better","isMain":false,"metricName":"# unaligned mis. contigs"}]],["Unaligned",[{"values":[0,0,0,0,0,0,0,0,0,0,0,0],"quality":"Less is better","isMain":false,"metricName":"# fully unaligned contigs"},{"values":[0,0,0,0,0,0,0,0,0,0,0,0],"quality":"Less is better","isMain":false,"metricName":"Fully unaligned length"},{"values":[0,0,0,0,0,0,4,0,0,0,0,0],"quality":"Less is better","isMain":false,"metricName":"# partially unaligned contigs"},{"values":[0,0,0,0,0,0,4288,0,0,0,0,0],"quality":"Less is better","isMain":false,"metricName":"Partially unaligned length"}]],["Mismatches",[{"values":[0,0,0,0,0,3,4,0,0,0,0,0],"quality":"Less is better","isMain":false,"metricName":"# mismatches"},{"values":[1,0,0,0,0,0,30,0,0,0,0,0],"quality":"Less is better","isMain":false,"metricName":"# indels"},{"values":[2,0,0,0,0,0,1413,0,0,0,0,0],"quality":"Less is better","isMain":false,"metricName":"Indels length"},{"values":["0.00","0.00","0.00","0.00","0.00","2.21","3.65","0.00","0.00","0.00","0.00","0.00"],"quality":"Less is better","isMain":true,"metricName":"# mismatches per 100 kbp"},{"values":["0.61","0.00","0.00","0.00","0.00","0.00","27.37","0.00","0.00","0.00","0.00","0.00"],"quality":"Less is better","isMain":true,"metricName":"# indels per 100 kbp"},{"values":[1,0,0,0,0,0,0,0,0,0,0,0],"quality":"Less is better","isMain":false,"metricName":" # indels (<= 5 bp)"},{"values":[0,0,0,0,0,0,30,0,0,0,0,0],"quality":"Less is better","isMain":false,"metricName":" # indels (> 5 bp)"},{"values":[0,0,0,0,0,0,4081,0,0,0,0,0],"quality":"Less is better","isMain":false,"metricName":"# N's"},{"values":["0.00","0.00","0.00","0.00","0.00","0.00","3481.49","0.00","0.00","0.00","0.00","0.00"],"quality":"Less is better","isMain":true,"metricName":"# N's per 100 kbp"}]],["Statistics without reference",[{"values":[1,1,1,1,1,97,96,1,1,1,1,18],"quality":"Equal","isMain":true,"metricName":"# contigs"},{"values":[1,1,1,1,1,66,46,1,1,1,1,17],"quality":"Equal","isMain":false,"metricName":"# contigs (>= 1000 bp)"},{"values":[1,1,1,1,1,3,0,1,1,1,1,9],"quality":"Equal","isMain":false,"metricName":"# contigs (>= 5000 bp)"},{"values":[1,1,1,1,1,0,0,1,1,1,1,6],"quality":"Equal","isMain":false,"metricName":"# contigs (>= 10000 bp)"},{"values":[1,1,1,1,1,0,0,1,1,1,1,1],"quality":"Equal","isMain":false,"metricName":"# contigs (>= 25000 bp)"},{"values":[1,1,1,1,1,0,0,1,1,1,1,0],"quality":"Equal","isMain":false,"metricName":"# contigs (>= 50000 bp)"},{"values":[164604,164602,164341,164743,164886,5699,3906,164729,164590,164590,164590,42860],"quality":"More is better","isMain":true,"metricName":"Largest contig"},{"values":[164604,164602,164341,164743,164886,144348,117220,164729,164590,164590,164590,163637],"quality":"More is better","isMain":true,"metricName":"Total length"},{"values":[164604,164602,164341,164743,164886,121615,82386,164729,164590,164590,164590,162964],"quality":"More is better","isMain":true,"metricName":"Total length (>= 1000 bp)"},{"values":[164604,164602,164341,164743,164886,16385,0,164729,164590,164590,164590,141148],"quality":"More is better","isMain":false,"metricName":"Total length (>= 5000 bp)"},{"values":[164604,164602,164341,164743,164886,0,0,164729,164590,164590,164590,122693],"quality":"More is better","isMain":true,"metricName":"Total length (>= 10000 bp)"},{"values":[164604,164602,164341,164743,164886,0,0,164729,164590,164590,164590,42860],"quality":"More is better","isMain":false,"metricName":"Total length (>= 25000 bp)"},{"values":[164604,164602,164341,164743,164886,0,0,164729,164590,164590,164590,0],"quality":"More is better","isMain":true,"metricName":"Total length (>= 50000 bp)"},{"values":[164604,164602,164341,164743,164886,1529,1505,164729,164590,164590,164590,16481],"quality":"More is better","isMain":false,"metricName":"N50"},{"values":[164604,164602,164341,164743,164886,1129,916,164729,164590,164590,164590,6798],"quality":"More is better","isMain":false,"metricName":"N75"},{"values":[1,1,1,1,1,27,28,1,1,1,1,3],"quality":"Less is better","isMain":false,"metricName":"L50"},{"values":[1,1,1,1,1,54,52,1,1,1,1,7],"quality":"Less is better","isMain":false,"metricName":"L75"},{"values":["65.44","65.44","65.44","65.44","65.43","65.40","65.41","65.43","65.44","65.44","65.44","65.44"],"quality":"Equal","isMain":false,"metricName":"GC (%)"}]],["Predicted genes",[]],["Similarity statistics",[{"values":[0,0,0,0,0,0,0,0,0,0,0,0],"quality":"Equal","isMain":false,"metricName":"# similar correct contigs"},{"values":[0,0,0,0,0,0,0,0,0,0,0,0],"quality":"Equal","isMain":false,"metricName":"# similar misassembled blocks"}]],["Reference statistics",[{"values":[164602,164602,164602,164602,164602,164602,164602,164602,164602,164602,164602,164602],"quality":"Equal","isMain":false,"metricName":"Reference length"},{"values":[1,1,1,1,1,1,1,1,1,1,1,1],"quality":"Equal","isMain":false,"metricName":"Reference fragments"},{"values":["65.44","65.44","65.44","65.44","65.44","65.44","65.44","65.44","65.44","65.44","65.44","65.44"],"quality":"Equal","isMain":false,"metricName":"Reference GC (%)"}]]],"referenceName":"gi\_203454602\_ref\_NC","date":"18 July 2018, Wednesday, 16:49:16","order":[0,1,2,3,4,5,6,7,8,9,10,11],"assembliesNames":["ABySS\_63","CLC","IDBA\_UD","MEGAHIT","MIRA","Ray\_Meta","SOAPdenovo2","SPAdes","SPAdes\_meta","SPAdes\_sc","SPAdes\_sc\_careful","Velvet"]},{"assembliesWithNs":null,"minContig":500,"report":[["Genome statistics",[{"values":["6.756","3.168","6.334","4.259","9.419","98.286","2.565","2.344","8.335","0.807","0.902","10.981","1.582","98.423"],"quality":"More is better","isMain":true,"metricName":"Genome fraction (%)"},{"values":["1.030","1.000","1.005","1.000","1.000","1.279","1.108","1.116","1.076","1.000","1.000","1.000","1.000","1.256"],"quality":"Less is better","isMain":true,"metricName":"Duplication ratio"},{"values":[1615,2335,1748,1330,2610,60324,1246,693,3566,631,757,7547,707,61366],"quality":"More is better","isMain":true,"metricName":"Largest alignment"},{"values":[10705,4876,9791,6555,14495,193265,4374,3607,13335,1242,1388,16902,2435,190171],"quality":"More is better","isMain":true,"metricName":"Total aligned length"},{"values":[null,null,null,null,null,158195,null,null,null,null,null,null,null,160724],"quality":"More is better","isMain":false,"metricName":"NG50"},{"values":[null,null,null,null,null,158195,null,null,null,null,null,null,null,160724],"quality":"More is better","isMain":false,"metricName":"NG75"},{"values":[914,771,788,863,982,43981,968,656,1759,631,757,7071,637,45468],"quality":"More is better","isMain":false,"metricName":"NA50"},{"values":[675,767,571,631,743,5698,682,562,1035,611,631,7071,573,10136],"quality":"More is better","isMain":false,"metricName":"NA75"},{"values":[null,null,null,null,null,43981,null,null,null,null,null,null,null,45468],"quality":"More is better","isMain":true,"metricName":"NGA50"},{"values":[null,null,null,null,null,43161,null,null,null,null,null,null,null,43161],"quality":"More is better","isMain":false,"metricName":"NGA75"},{"values":[null,null,null,null,null,1,null,null,null,null,null,null,null,1],"quality":"Less is better","isMain":false,"metricName":"LG50"},{"values":[null,null,null,null,null,1,null,null,null,null,null,null,null,1],"quality":"Less is better","isMain":false,"metricName":"LG75"},{"values":[5,2,5,4,5,2,2,3,3,1,1,2,2,2],"quality":"Less is better","isMain":false,"metricName":"LA50"},{"values":[9,3,8,6,10,4,4,5,5,2,2,2,3,4],"quality":"Less is better","isMain":false,"metricName":"LA75"},{"values":[null,null,null,null,null,2,null,null,null,null,null,null,null,2],"quality":"Less is better","isMain":true,"metricName":"LGA50"},{"values":[null,null,null,null,null,3,null,null,null,null,null,null,null,3],"quality":"Less is better","isMain":false,"metricName":"LGA75"}]],["Misassemblies",[{"values":[0,0,0,0,0,2,0,0,0,0,0,0,0,2],"quality":"Less is better","isMain":true,"metricName":"# misassemblies"},{"values":[0,0,0,0,0,2,0,0,0,0,0,0,0,2],"quality":"Less is better","isMain":false,"metricName":" # relocations"},{"values":[0,0,0,0,0,0,0,0,0,0,0,0,0,0],"quality":"Less is better","isMain":false,"metricName":" # translocations"},{"values":[0,0,0,0,0,0,0,0,0,0,0,0,0,0],"quality":"Less is better","isMain":false,"metricName":" # inversions"},{"values":[0,0,0,0,0,1,0,0,0,0,0,0,0,1],"quality":"Less is better","isMain":false,"metricName":"# misassembled contigs"},{"values":[0,0,0,0,0,158195,0,0,0,0,0,0,0,160724],"quality":"Less is better","isMain":true,"metricName":"Misassembled contigs length"},{"values":[0,0,0,0,0,19,0,0,0,0,0,0,0,21],"quality":"Less is better","isMain":false,"metricName":"# local misassemblies"},{"values":[0,0,0,0,0,0,0,0,0,0,0,0,0,0],"quality":"Less is better","isMain":false,"metricName":"# unaligned mis. contigs"}]],["Unaligned",[{"values":[0,0,0,0,0,0,0,0,0,0,0,0,0,0],"quality":"Less is better","isMain":false,"metricName":"# fully unaligned contigs"},{"values":[0,0,0,0,0,0,0,0,0,0,0,0,0,0],"quality":"Less is better","isMain":false,"metricName":"Fully unaligned length"},{"values":[0,0,0,0,0,1,0,0,0,0,0,0,0,2],"quality":"Less is better","isMain":false,"metricName":"# partially unaligned contigs"},{"values":[0,0,0,0,0,10729,0,0,0,0,0,0,0,12821],"quality":"Less is better","isMain":false,"metricName":"Partially unaligned length"}]],["Mismatches",[{"values":[0,0,10,2,12,1195,14,3,35,2,0,103,1,804],"quality":"Less is better","isMain":false,"metricName":"# mismatches"},{"values":[0,0,0,0,0,66,0,2,0,0,0,9,0,58],"quality":"Less is better","isMain":false,"metricName":"# indels"},{"values":[0,0,0,0,0,292,0,86,0,0,0,15,0,270],"quality":"Less is better","isMain":false,"metricName":"Indels length"},{"values":["0.00","0.00","102.60","30.51","82.79","790.05","354.61","83.17","272.86","161.03","0.00","609.50","41.07","530.81"],"quality":"Less is better","isMain":true,"metricName":"# mismatches per 100 kbp"},{"values":["0.00","0.00","0.00","0.00","0.00","43.63","0.00","55.45","0.00","0.00","0.00","53.26","0.00","38.29"],"quality":"Less is better","isMain":true,"metricName":"# indels per 100 kbp"},{"values":[0,0,0,0,0,56,0,0,0,0,0,8,0,49],"quality":"Less is better","isMain":false,"metricName":" # indels (<= 5 bp)"},{"values":[0,0,0,0,0,10,0,2,0,0,0,1,0,9],"quality":"Less is better","isMain":false,"metricName":" # indels (> 5 bp)"},{"values":[0,0,0,0,0,40,0,307,467,0,0,0,0,655],"quality":"Less is better","isMain":false,"metricName":"# N's"},{"values":["0.00","0.00","0.00","0.00","0.00","19.59","0.00","7627.33","3383.57","0.00","0.00","0.00","0.00","322.45"],"quality":"Less is better","isMain":true,"metricName":"# N's per 100 kbp"}]],["Statistics without reference",[{"values":[13,5,12,8,15,29,5,6,8,2,2,5,4,9],"quality":"Equal","isMain":true,"metricName":"# contigs"},{"values":[2,1,3,1,4,19,1,0,5,0,0,2,0,8],"quality":"Equal","isMain":false,"metricName":"# contigs (>= 1000 bp)"},{"values":[0,0,0,0,0,2,0,0,0,0,0,2,0,4],"quality":"Equal","isMain":false,"metricName":"# contigs (>= 5000 bp)"},{"values":[0,0,0,0,0,1,0,0,0,0,0,0,0,2],"quality":"Equal","isMain":false,"metricName":"# contigs (>= 10000 bp)"},{"values":[0,0,0,0,0,1,0,0,0,0,0,0,0,1],"quality":"Equal","isMain":false,"metricName":"# contigs (>= 25000 bp)"},{"values":[0,0,0,0,0,1,0,0,0,0,0,0,0,1],"quality":"Equal","isMain":false,"metricName":"# contigs (>= 50000 bp)"},{"values":[1615,2335,1748,1330,2610,158195,1246,988,3570,631,757,7547,707,160724],"quality":"More is better","isMain":true,"metricName":"Largest contig"},{"values":[10705,4876,9791,6555,14495,204147,4374,4025,13802,1242,1388,16902,2435,203133],"quality":"More is better","isMain":true,"metricName":"Total length"},{"values":[2726,2335,3939,1330,6592,195909,1246,0,11547,0,0,14618,0,202186],"quality":"More is better","isMain":true,"metricName":"Total length (>= 1000 bp)"},{"values":[0,0,0,0,0,163905,0,0,0,0,0,14618,0,188852],"quality":"More is better","isMain":false,"metricName":"Total length (>= 5000 bp)"},{"values":[0,0,0,0,0,158195,0,0,0,0,0,0,0,172952],"quality":"More is better","isMain":true,"metricName":"Total length (>= 10000 bp)"},{"values":[0,0,0,0,0,158195,0,0,0,0,0,0,0,160724],"quality":"More is better","isMain":false,"metricName":"Total length (>= 25000 bp)"},{"values":[0,0,0,0,0,158195,0,0,0,0,0,0,0,160724],"quality":"More is better","isMain":true,"metricName":"Total length (>= 50000 bp)"},{"values":[914,771,788,863,982,158195,968,689,3566,631,757,7071,637,160724],"quality":"More is better","isMain":false,"metricName":"N50"},{"values":[675,767,571,631,743,158195,682,562,1617,611,631,7071,573,160724],"quality":"More is better","isMain":false,"metricName":"N75"},{"values":[5,2,5,4,5,1,2,3,2,1,1,2,2,1],"quality":"Less is better","isMain":false,"metricName":"L50"},{"values":[9,3,8,6,10,1,4,5,4,2,2,2,3,1],"quality":"Less is better","isMain":false,"metricName":"L75"},{"values":["64.26","65.57","63.84","63.07","64.07","64.60","65.39","62.96","64.02","59.90","59.87","64.29","62.59","64.52"],"quality":"Equal","isMain":false,"metricName":"GC (%)"}]],["Predicted genes",[]],["Similarity statistics",[{"values":[0,0,0,0,0,0,0,0,0,0,0,0,0,0],"quality":"Equal","isMain":false,"metricName":"# similar correct contigs"},{"values":[0,0,0,0,0,0,0,0,0,0,0,0,0,0],"quality":"Equal","isMain":false,"metricName":"# similar misassembled blocks"}]],["Reference statistics",[{"values":[153894,153894,153894,153894,153894,153894,153894,153894,153894,153894,153894,153894,153894,153894],"quality":"Equal","isMain":false,"metricName":"Reference length"},{"values":[1,1,1,1,1,1,1,1,1,1,1,1,1,1],"quality":"Equal","isMain":false,"metricName":"Reference fragments"},{"values":["64.74","64.74","64.74","64.74","64.74","64.74","64.74","64.74","64.74","64.74","64.74","64.74","64.74","64.74"],"quality":"Equal","isMain":false,"metricName":"Reference GC (%)"}]]],"referenceName":"gi\_203457352\_ref\_NC","date":"18 July 2018, Wednesday, 16:49:35","order":[0,1,2,3,4,5,6,7,8,9,10,11,12,13],"assembliesNames":["ABySS\_127","ABySS\_63","CLC","IDBA\_UD","MEGAHIT","MIRA","Ray\_Meta","SOAPdenovo2","SPAdes","SPAdes\_meta","SPAdes\_sc","SPAdes\_sc\_careful","Velvet","Geneious"]},{"assembliesWithNs":null,"minContig":500,"report":[["Genome statistics",[{"values":["3.936","1.052","4.250","5.837","2.464","9.286","2.134","0.551","0.892","0.356","1.089","0.477","0.936"],"quality":"More is better","isMain":true,"metricName":"Genome fraction (%)"},{"values":["1.000","1.000","1.000","1.000","1.000","1.102","0.992","1.330","1.000","1.000","0.999","1.000","1.000"],"quality":"Less is better","isMain":true,"metricName":"Duplication ratio"},{"values":[3179,835,1278,5391,849,3568,1114,614,748,549,998,735,868],"quality":"More is better","isMain":true,"metricName":"Largest alignment"},{"values":[6062,1621,6546,8990,3795,15110,3260,849,1374,549,1677,735,1442],"quality":"More is better","isMain":true,"metricName":"Total aligned length"},{"values":[3179,835,1052,5391,602,1308,1081,614,748,549,998,735,868],"quality":"More is better","isMain":false,"metricName":"NA50"},{"values":[714,786,785,1401,522,851,1065,235,626,549,679,735,574],"quality":"More is better","isMain":false,"metricName":"NA75"},{"values":[null,null,null,null,null,null,null,null,null,null,null,null,null],"quality":"More is better","isMain":true,"metricName":"NGA50"},{"values":[1,1,3,1,3,4,2,1,1,1,1,1,1],"quality":"Less is better","isMain":false,"metricName":"LA50"},{"values":[3,2,5,2,5,7,3,2,2,1,2,1,2],"quality":"Less is better","isMain":false,"metricName":"LA75"}]],["Misassemblies",[{"values":[0,0,0,0,0,0,0,0,0,0,0,0,0],"quality":"Less is better","isMain":true,"metricName":"# misassemblies"},{"values":[0,0,0,0,0,0,0,0,0,0,0,0,0],"quality":"Less is better","isMain":false,"metricName":" # relocations"},{"values":[0,0,0,0,0,0,0,0,0,0,0,0,0],"quality":"Less is better","isMain":false,"metricName":" # translocations"},{"values":[0,0,0,0,0,0,0,0,0,0,0,0,0],"quality":"Less is better","isMain":false,"metricName":" # inversions"},{"values":[0,0,0,0,0,0,0,0,0,0,0,0,0],"quality":"Less is better","isMain":false,"metricName":"# misassembled contigs"},{"values":[0,0,0,0,0,0,0,0,0,0,0,0,0],"quality":"Less is better","isMain":true,"metricName":"Misassembled contigs length"},{"values":[0,0,0,0,0,2,0,0,0,0,0,0,0],"quality":"Less is better","isMain":false,"metricName":"# local misassemblies"},{"values":[0,0,0,0,0,0,0,0,0,0,0,0,0],"quality":"Less is better","isMain":false,"metricName":"# unaligned mis. contigs"}]],["Unaligned",[{"values":[0,0,0,0,0,0,0,0,0,0,0,0,0],"quality":"Less is better","isMain":false,"metricName":"# fully unaligned contigs"},{"values":[0,0,0,0,0,0,0,0,0,0,0,0,0],"quality":"Less is better","isMain":false,"metricName":"Fully unaligned length"},{"values":[0,0,0,0,0,0,0,0,0,0,0,0,0],"quality":"Less is better","isMain":false,"metricName":"# partially unaligned contigs"},{"values":[0,0,0,0,0,0,0,0,0,0,0,0,0],"quality":"Less is better","isMain":false,"metricName":"Partially unaligned length"}]],["Mismatches",[{"values":[5,0,18,57,3,83,16,0,3,0,8,0,0],"quality":"Less is better","isMain":false,"metricName":"# mismatches"},{"values":[0,0,0,0,0,0,3,0,0,0,1,0,0],"quality":"Less is better","isMain":false,"metricName":"# indels"},{"values":[0,0,0,0,0,0,27,0,0,0,1,0,0],"quality":"Less is better","isMain":false,"metricName":"Indels length"},{"values":["82.48","0.00","274.98","634.04","79.05","580.34","486.77","0.00","218.34","0.00","476.76","0.00","0.00"],"quality":"Less is better","isMain":true,"metricName":"# mismatches per 100 kbp"},{"values":["0.00","0.00","0.00","0.00","0.00","0.00","91.27","0.00","0.00","0.00","59.59","0.00","0.00"],"quality":"Less is better","isMain":true,"metricName":"# indels per 100 kbp"},{"values":[0,0,0,0,0,0,2,0,0,0,1,0,0],"quality":"Less is better","isMain":false,"metricName":" # indels (<= 5 bp)"},{"values":[0,0,0,0,0,0,1,0,0,0,0,0,0],"quality":"Less is better","isMain":false,"metricName":" # indels (> 5 bp)"},{"values":[0,0,0,0,0,14,0,220,0,0,0,0,0],"quality":"Less is better","isMain":false,"metricName":"# N's"},{"values":["0.00","0.00","0.00","0.00","0.00","88.84","0.00","19486.27","0.00","0.00","0.00","0.00","0.00"],"quality":"Less is better","isMain":true,"metricName":"# N's per 100 kbp"}]],["Statistics without reference",[{"values":[5,2,7,5,6,12,3,2,2,1,2,1,2],"quality":"Equal","isMain":true,"metricName":"# contigs"},{"values":[2,0,4,2,0,4,3,0,0,0,0,0,0],"quality":"Equal","isMain":false,"metricName":"# contigs (>= 1000 bp)"},{"values":[0,0,0,1,0,0,0,0,0,0,0,0,0],"quality":"Equal","isMain":false,"metricName":"# contigs (>= 5000 bp)"},{"values":[0,0,0,0,0,0,0,0,0,0,0,0,0],"quality":"Equal","isMain":false,"metricName":"# contigs (>= 10000 bp)"},{"values":[0,0,0,0,0,0,0,0,0,0,0,0,0],"quality":"Equal","isMain":false,"metricName":"# contigs (>= 25000 bp)"},{"values":[0,0,0,0,0,0,0,0,0,0,0,0,0],"quality":"Equal","isMain":false,"metricName":"# contigs (>= 50000 bp)"},{"values":[3179,835,1278,5391,849,4217,1114,614,748,549,998,735,868],"quality":"More is better","isMain":true,"metricName":"Largest contig"},{"values":[6062,1621,6546,8990,3795,15759,3260,1129,1374,549,1677,735,1442],"quality":"More is better","isMain":true,"metricName":"Total length"},{"values":[4229,0,4446,6792,0,9731,3260,0,0,0,0,0,0],"quality":"More is better","isMain":true,"metricName":"Total length (>= 1000 bp)"},{"values":[0,0,0,5391,0,0,0,0,0,0,0,0,0],"quality":"More is better","isMain":false,"metricName":"Total length (>= 5000 bp)"},{"values":[0,0,0,0,0,0,0,0,0,0,0,0,0],"quality":"More is better","isMain":true,"metricName":"Total length (>= 10000 bp)"},{"values":[0,0,0,0,0,0,0,0,0,0,0,0,0],"quality":"More is better","isMain":false,"metricName":"Total length (>= 25000 bp)"},{"values":[0,0,0,0,0,0,0,0,0,0,0,0,0],"quality":"More is better","isMain":true,"metricName":"Total length (>= 50000 bp)"},{"values":[3179,835,1052,5391,602,1907,1081,614,748,549,998,735,868],"quality":"More is better","isMain":false,"metricName":"N50"},{"values":[714,786,785,1401,522,851,1065,515,626,549,679,735,574],"quality":"More is better","isMain":false,"metricName":"N75"},{"values":[1,1,3,1,3,3,2,1,1,1,1,1,1],"quality":"Less is better","isMain":false,"metricName":"L50"},{"values":[3,2,5,2,5,7,3,2,2,1,2,1,2],"quality":"Less is better","isMain":false,"metricName":"L75"},{"values":["65.09","65.02","63.84","64.95","63.79","64.43","65.09","61.17","66.45","65.21","62.97","59.18","63.38"],"quality":"Equal","isMain":false,"metricName":"GC (%)"}]],["Predicted genes",[]],["Similarity statistics",[{"values":[0,0,0,0,0,0,0,0,0,0,0,0,0],"quality":"Equal","isMain":false,"metricName":"# similar correct contigs"},{"values":[0,0,0,0,0,0,0,0,0,0,0,0,0],"quality":"Equal","isMain":false,"metricName":"# similar misassembled blocks"}]],["Reference statistics",[{"values":[154017,154017,154017,154017,154017,154017,154017,154017,154017,154017,154017,154017,154017],"quality":"Equal","isMain":false,"metricName":"Reference length"},{"values":[1,1,1,1,1,1,1,1,1,1,1,1,1],"quality":"Equal","isMain":false,"metricName":"Reference fragments"},{"values":["64.83","64.83","64.83","64.83","64.83","64.83","64.83","64.83","64.83","64.83","64.83","64.83","64.83"],"quality":"Equal","isMain":false,"metricName":"Reference GC (%)"}]]],"referenceName":"gi\_203458877\_ref\_NC","date":"18 July 2018, Wednesday, 16:49:53","order":[0,1,2,3,4,5,6,7,8,9,10,11,12],"assembliesNames":["ABySS\_127","ABySS\_63","CLC","IDBA\_UD","MEGAHIT","MIRA","Ray\_Meta","SOAPdenovo2","SPAdes","SPAdes\_meta","SPAdes\_sc","SPAdes\_sc\_careful","Velvet"]},{"assembliesWithNs":null,"minContig":500,"report":[["Genome statistics",[{"values":["2.225","2.365","2.956","2.250","7.548","9.775","2.625","0.423","6.527","2.706","9.461","10.348","4.356","2.638"],"quality":"More is better","isMain":true,"metricName":"Genome fraction (%)"},{"values":["1.000","1.009","1.000","1.051","1.001","1.000","1.000","1.000","1.133","1.000","1.036","1.045","1.000","1.000"],"quality":"Less is better","isMain":true,"metricName":"Duplication ratio"},{"values":[1114,975,1079,1087,3504,2493,1648,549,2798,1650,10304,10099,3416,4087],"quality":"More is better","isMain":true,"metricName":"Largest alignment"},{"values":[3446,3696,4579,3585,11697,15142,4067,655,11454,4192,14655,16024,6748,4087],"quality":"More is better","isMain":true,"metricName":"Total aligned length"},{"values":[916,743,1037,819,1639,1855,1518,null,1030,1342,10304,10099,3416,4087],"quality":"More is better","isMain":false,"metricName":"NA50"},{"values":[743,690,848,566,1183,1211,1518,null,602,1200,1650,1342,838,4087],"quality":"More is better","isMain":false,"metricName":"NA75"},{"values":[null,null,null,null,null,null,null,null,null,null,null,null,null,null],"quality":"More is better","isMain":true,"metricName":"NGA50"},{"values":[2,3,3,2,3,4,2,null,4,2,1,1,1,1],"quality":"Less is better","isMain":false,"metricName":"LA50"},{"values":[3,4,4,4,5,7,2,null,8,3,2,3,3,1],"quality":"Less is better","isMain":false,"metricName":"LA75"}]],["Misassemblies",[{"values":[0,0,0,0,0,0,0,0,2,0,0,0,0,0],"quality":"Less is better","isMain":true,"metricName":"# misassemblies"},{"values":[0,0,0,0,0,0,0,0,2,0,0,0,0,0],"quality":"Less is better","isMain":false,"metricName":" # relocations"},{"values":[0,0,0,0,0,0,0,0,0,0,0,0,0,0],"quality":"Less is better","isMain":false,"metricName":" # translocations"},{"values":[0,0,0,0,0,0,0,0,0,0,0,0,0,0],"quality":"Less is better","isMain":false,"metricName":" # inversions"},{"values":[0,0,0,0,0,0,0,0,1,0,0,0,0,0],"quality":"Less is better","isMain":false,"metricName":"# misassembled contigs"},{"values":[0,0,0,0,0,0,0,0,5357,0,0,0,0,0],"quality":"Less is better","isMain":true,"metricName":"Misassembled contigs length"},{"values":[0,0,0,0,0,0,0,0,1,0,1,1,0,0],"quality":"Less is better","isMain":false,"metricName":"# local misassemblies"},{"values":[0,0,0,0,0,0,0,0,0,0,0,0,0,0],"quality":"Less is better","isMain":false,"metricName":"# unaligned mis. contigs"}]],["Unaligned",[{"values":[0,0,0,0,0,0,0,0,0,0,0,0,0,0],"quality":"Less is better","isMain":false,"metricName":"# fully unaligned contigs"},{"values":[0,0,0,0,0,0,0,0,0,0,0,0,0,0],"quality":"Less is better","isMain":false,"metricName":"Fully unaligned length"},{"values":[0,0,0,0,0,0,0,1,1,0,0,0,0,0],"quality":"Less is better","isMain":false,"metricName":"# partially unaligned contigs"},{"values":[0,0,0,0,0,0,0,773,1845,0,0,0,0,0],"quality":"Less is better","isMain":false,"metricName":"Partially unaligned length"}]],["Mismatches",[{"values":[0,0,8,3,21,8,8,0,34,3,90,65,1,3],"quality":"Less is better","isMain":false,"metricName":"# mismatches"},{"values":[0,0,0,0,0,0,0,0,0,0,0,2,0,0],"quality":"Less is better","isMain":false,"metricName":"# indels"},{"values":[0,0,0,0,0,0,0,0,0,0,0,5,0,0],"quality":"Less is better","isMain":false,"metricName":"Indels length"},{"values":["0.00","0.00","174.71","86.06","179.61","52.83","196.71","0.00","336.30","71.56","614.12","405.52","14.82","73.40"],"quality":"Less is better","isMain":true,"metricName":"# mismatches per 100 kbp"},{"values":["0.00","0.00","0.00","0.00","0.00","0.00","0.00","0.00","0.00","0.00","0.00","12.48","0.00","0.00"],"quality":"Less is better","isMain":true,"metricName":"# indels per 100 kbp"},{"values":[0,0,0,0,0,0,0,0,0,0,0,2,0,0],"quality":"Less is better","isMain":false,"metricName":" # indels (<= 5 bp)"},{"values":[0,0,0,0,0,0,0,0,0,0,0,0,0,0],"quality":"Less is better","isMain":false,"metricName":" # indels (> 5 bp)"},{"values":[0,0,0,0,0,10,0,174,0,0,529,724,0,12],"quality":"Less is better","isMain":false,"metricName":"# N's"},{"values":["0.00","0.00","0.00","0.00","0.00","66.04","0.00","12184.87","0.00","0.00","3483.93","4321.61","0.00","293.61"],"quality":"Less is better","isMain":true,"metricName":"# N's per 100 kbp"}]],["Statistics without reference",[{"values":[4,5,5,5,7,10,3,2,8,3,5,7,5,1],"quality":"Equal","isMain":true,"metricName":"# contigs"},{"values":[1,0,3,1,6,8,2,0,4,3,3,4,2,1],"quality":"Equal","isMain":false,"metricName":"# contigs (>= 1000 bp)"},{"values":[0,0,0,0,0,0,0,0,1,0,1,1,0,0],"quality":"Equal","isMain":false,"metricName":"# contigs (>= 5000 bp)"},{"values":[0,0,0,0,0,0,0,0,0,0,1,1,0,0],"quality":"Equal","isMain":false,"metricName":"# contigs (>= 10000 bp)"},{"values":[0,0,0,0,0,0,0,0,0,0,0,0,0,0],"quality":"Equal","isMain":false,"metricName":"# contigs (>= 25000 bp)"},{"values":[0,0,0,0,0,0,0,0,0,0,0,0,0,0],"quality":"Equal","isMain":false,"metricName":"# contigs (>= 50000 bp)"},{"values":[1114,975,1079,1087,3504,2493,1648,879,5357,1650,10833,10828,3416,4087],"quality":"More is better","isMain":true,"metricName":"Largest contig"},{"values":[3446,3696,4579,3663,11698,15142,4067,1428,13300,4192,15184,16753,6748,4087],"quality":"More is better","isMain":true,"metricName":"Total length"},{"values":[1114,0,3155,1087,10815,13413,3166,0,10357,4192,13825,14863,4750,4087],"quality":"More is better","isMain":true,"metricName":"Total length (>= 1000 bp)"},{"values":[0,0,0,0,0,0,0,0,5357,0,10833,10828,0,0],"quality":"More is better","isMain":false,"metricName":"Total length (>= 5000 bp)"},{"values":[0,0,0,0,0,0,0,0,0,0,10833,10828,0,0],"quality":"More is better","isMain":true,"metricName":"Total length (>= 10000 bp)"},{"values":[0,0,0,0,0,0,0,0,0,0,0,0,0,0],"quality":"More is better","isMain":false,"metricName":"Total length (>= 25000 bp)"},{"values":[0,0,0,0,0,0,0,0,0,0,0,0,0,0],"quality":"More is better","isMain":true,"metricName":"Total length (>= 50000 bp)"},{"values":[916,743,1037,819,1639,1855,1518,879,2798,1342,10833,10828,3416,4087],"quality":"More is better","isMain":false,"metricName":"N50"},{"values":[743,690,848,566,1183,1211,1518,549,1030,1200,1650,1342,838,4087],"quality":"More is better","isMain":false,"metricName":"N75"},{"values":[2,3,3,2,3,4,2,1,2,2,1,1,1,1],"quality":"Less is better","isMain":false,"metricName":"L50"},{"values":[3,4,4,4,5,7,2,2,4,3,2,3,3,1],"quality":"Less is better","isMain":false,"metricName":"L75"},{"values":["65.03","64.69","65.10","64.05","64.27","65.05","64.72","58.85","63.65","64.00","64.19","64.46","65.04","65.77"],"quality":"Equal","isMain":false,"metricName":"GC (%)"}]],["Predicted genes",[]],["Similarity statistics",[{"values":[0,0,0,0,0,0,0,0,0,0,0,0,0,0],"quality":"Equal","isMain":false,"metricName":"# similar correct contigs"},{"values":[0,0,0,0,0,0,0,0,0,0,0,0,0,0],"quality":"Equal","isMain":false,"metricName":"# similar misassembled blocks"}]],["Reference statistics",[{"values":[154906,154906,154906,154906,154906,154906,154906,154906,154906,154906,154906,154906,154906,154906],"quality":"Equal","isMain":false,"metricName":"Reference length"},{"values":[1,1,1,1,1,1,1,1,1,1,1,1,1,1],"quality":"Equal","isMain":false,"metricName":"Reference fragments"},{"values":["64.78","64.78","64.78","64.78","64.78","64.78","64.78","64.78","64.78","64.78","64.78","64.78","64.78","64.78"],"quality":"Equal","isMain":false,"metricName":"Reference GC (%)"}]]],"referenceName":"gi\_203460520\_ref\_NC","date":"18 July 2018, Wednesday, 16:50:09","order":[0,1,2,3,4,5,6,7,8,9,10,11,12,13],"assembliesNames":["ABySS\_127","ABySS\_63","CLC","IDBA\_UD","MEGAHIT","MIRA","Ray\_Meta","SOAPdenovo2","SPAdes","SPAdes\_meta","SPAdes\_sc","SPAdes\_sc\_careful","Velvet","Geneious"]},{"assembliesWithNs":null,"minContig":500,"report":[["Genome statistics",[{"values":["93.233","59.650","88.534","86.948","96.553","100.000","94.316","5.276","99.772","100.000","88.338","94.970","55.524","100.000"],"quality":"More is better","isMain":true,"metricName":"Genome fraction (%)"},{"values":["1.034","1.003","1.026","1.044","1.083","1.043","1.062","1.000","1.050","1.003","1.020","1.018","1.001","1.141"],"quality":"Less is better","isMain":true,"metricName":"Duplication ratio"},{"values":[10481,2842,4859,4455,5225,42533,6940,804,13886,32616,8264,6012,2644,42534],"quality":"More is better","isMain":true,"metricName":"Largest alignment"},{"values":[47687,29581,44930,44857,51072,51261,49545,2611,51406,49601,44600,47810,27506,56150],"quality":"More is better","isMain":true,"metricName":"Total aligned length"},{"values":[4170,666,1984,1249,1789,51598,3644,null,12877,32671,2615,2891,610,55540],"quality":"More is better","isMain":false,"metricName":"NG50"},{"values":[1632,null,1375,733,1470,51598,1586,null,2893,11275,1070,1480,null,55540],"quality":"More is better","isMain":false,"metricName":"NG75"},{"values":[4170,1462,2082,1397,1777,42533,2585,665,12877,32616,4281,3178,992,42534],"quality":"More is better","isMain":false,"metricName":"NA50"},{"values":[1768,734,1463,841,1429,42533,1377,609,2488,11275,1366,1685,689,12669],"quality":"More is better","isMain":false,"metricName":"NA75"},{"values":[4170,666,1984,1249,1777,42533,3518,null,12877,32616,2615,2891,610,42534],"quality":"More is better","isMain":true,"metricName":"NGA50"},{"values":[1632,null,1375,733,1466,42533,1393,null,2893,11275,1070,1480,null,42534],"quality":"More is better","isMain":false,"metricName":"NGA75"},{"values":[4,20,8,12,9,1,5,null,2,1,5,6,25,1],"quality":"Less is better","isMain":false,"metricName":"LG50"},{"values":[10,null,15,25,17,1,10,null,5,2,13,11,null,1],"quality":"Less is better","isMain":false,"metricName":"LG75"},{"values":[4,8,7,10,10,1,6,2,2,1,4,5,10,1],"quality":"Less is better","isMain":false,"metricName":"LA50"},{"values":[9,16,13,21,19,1,13,3,6,2,9,10,18,2],"quality":"Less is better","isMain":false,"metricName":"LA75"},{"values":[4,20,8,12,10,1,5,null,2,1,5,6,25,1],"quality":"Less is better","isMain":true,"metricName":"LGA50"},{"values":[10,null,15,25,17,1,12,null,5,2,13,11,null,1],"quality":"Less is better","isMain":false,"metricName":"LGA75"}]],["Misassemblies",[{"values":[0,0,0,0,0,1,0,0,0,0,0,0,0,1],"quality":"Less is better","isMain":true,"metricName":"# misassemblies"},{"values":[0,0,0,0,0,1,0,0,0,0,0,0,0,1],"quality":"Less is better","isMain":false,"metricName":" # relocations"},{"values":[0,0,0,0,0,0,0,0,0,0,0,0,0,0],"quality":"Less is better","isMain":false,"metricName":" # translocations"},{"values":[0,0,0,0,0,0,0,0,0,0,0,0,0,0],"quality":"Less is better","isMain":false,"metricName":" # inversions"},{"values":[0,0,0,0,0,1,0,0,0,0,0,0,0,1],"quality":"Less is better","isMain":false,"metricName":"# misassembled contigs"},{"values":[0,0,0,0,0,51598,0,0,0,0,0,0,0,55540],"quality":"Less is better","isMain":true,"metricName":"Misassembled contigs length"},{"values":[0,0,0,0,1,1,0,0,0,0,0,0,0,1],"quality":"Less is better","isMain":false,"metricName":"# local misassemblies"},{"values":[0,0,0,0,0,0,0,0,0,0,0,0,0,0],"quality":"Less is better","isMain":false,"metricName":"# unaligned mis. contigs"}]],["Unaligned",[{"values":[0,0,0,0,0,0,0,0,0,0,0,0,0,0],"quality":"Less is better","isMain":false,"metricName":"# fully unaligned contigs"},{"values":[0,0,0,0,0,0,0,0,0,0,0,0,0,0],"quality":"Less is better","isMain":false,"metricName":"Fully unaligned length"},{"values":[0,0,0,0,0,0,1,0,1,0,0,0,0,1],"quality":"Less is better","isMain":false,"metricName":"# partially unaligned contigs"},{"values":[0,0,0,0,0,0,3018,0,984,0,0,0,0,4470],"quality":"Less is better","isMain":false,"metricName":"Partially unaligned length"}]],["Mismatches",[{"values":[0,0,20,0,71,12,23,0,0,101,35,25,19,111],"quality":"Less is better","isMain":false,"metricName":"# mismatches"},{"values":[0,0,0,0,2,0,0,0,0,2,2,1,0,5],"quality":"Less is better","isMain":false,"metricName":"# indels"},{"values":[0,0,0,0,9,0,0,0,0,4,4,1,0,5],"quality":"Less is better","isMain":false,"metricName":"Indels length"},{"values":["0.00","0.00","45.65","0.00","148.59","24.25","49.28","0.00","0.00","204.09","80.06","53.19","69.15","224.30"],"quality":"Less is better","isMain":true,"metricName":"# mismatches per 100 kbp"},{"values":["0.00","0.00","0.00","0.00","4.19","0.00","0.00","0.00","0.00","4.04","4.57","2.13","0.00","10.10"],"quality":"Less is better","isMain":true,"metricName":"# indels per 100 kbp"},{"values":[0,0,0,0,1,0,0,0,0,2,2,1,0,5],"quality":"Less is better","isMain":false,"metricName":" # indels (<= 5 bp)"},{"values":[0,0,0,0,1,0,0,0,0,0,0,0,0,0],"quality":"Less is better","isMain":false,"metricName":" # indels (> 5 bp)"},{"values":[0,0,0,0,0,2,0,0,1399,0,0,0,0,361],"quality":"Less is better","isMain":false,"metricName":"# N's"},{"values":["0.00","0.00","0.00","0.00","0.00","3.88","0.00","0.00","2649.37","0.00","0.00","0.00","0.00","592.22"],"quality":"Less is better","isMain":true,"metricName":"# N's per 100 kbp"}]],["Statistics without reference",[{"values":[22,27,24,38,30,1,24,4,14,3,22,22,29,2],"quality":"Equal","isMain":true,"metricName":"# contigs"},{"values":[11,9,19,16,25,1,18,0,10,3,14,15,9,2],"quality":"Equal","isMain":false,"metricName":"# contigs (>= 1000 bp)"},{"values":[3,0,0,0,1,1,2,0,3,3,3,3,0,2],"quality":"Equal","isMain":false,"metricName":"# contigs (>= 5000 bp)"},{"values":[1,0,0,0,0,1,0,0,2,2,0,0,0,1],"quality":"Equal","isMain":false,"metricName":"# contigs (>= 10000 bp)"},{"values":[0,0,0,0,0,1,0,0,0,1,0,0,0,1],"quality":"Equal","isMain":false,"metricName":"# contigs (>= 25000 bp)"},{"values":[0,0,0,0,0,1,0,0,0,0,0,0,0,1],"quality":"Equal","isMain":false,"metricName":"# contigs (>= 50000 bp)"},{"values":[10481,2865,4859,4515,5225,51598,6940,804,14870,32671,8264,6012,2644,55540],"quality":"More is better","isMain":true,"metricName":"Largest contig"},{"values":[47707,29604,44938,44917,51756,51598,52589,2611,52805,49656,44600,47822,27506,60957],"quality":"More is better","isMain":true,"metricName":"Total length"},{"values":[39538,16919,42193,29656,47861,51598,47710,0,49975,49656,39206,42703,13494,60957],"quality":"More is better","isMain":true,"metricName":"Total length (>= 1000 bp)"},{"values":[21610,0,0,0,5225,51598,12291,0,33159,49656,18732,16475,0,60957],"quality":"More is better","isMain":false,"metricName":"Total length (>= 5000 bp)"},{"values":[10481,0,0,0,0,51598,0,0,27747,43946,0,0,0,55540],"quality":"More is better","isMain":true,"metricName":"Total length (>= 10000 bp)"},{"values":[0,0,0,0,0,51598,0,0,0,32671,0,0,0,55540],"quality":"More is better","isMain":false,"metricName":"Total length (>= 25000 bp)"},{"values":[0,0,0,0,0,51598,0,0,0,0,0,0,0,55540],"quality":"More is better","isMain":true,"metricName":"Total length (>= 50000 bp)"},{"values":[4170,1462,2085,1397,1777,51598,3518,665,12877,32671,4281,3178,992,55540],"quality":"More is better","isMain":false,"metricName":"N50"},{"values":[1768,734,1466,841,1441,51598,1424,609,2893,11275,1366,1685,689,55540],"quality":"More is better","isMain":false,"metricName":"N75"},{"values":[4,8,7,10,10,1,6,2,2,1,4,5,10,1],"quality":"Less is better","isMain":false,"metricName":"L50"},{"values":[9,16,13,21,18,1,12,3,5,2,9,10,18,1],"quality":"Less is better","isMain":false,"metricName":"L75"},{"values":["63.79","64.18","63.75","64.12","63.86","63.83","63.67","65.68","63.80","63.87","63.82","63.88","64.03","63.91"],"quality":"Equal","isMain":false,"metricName":"GC (%)"}]],["Predicted genes",[]],["Similarity statistics",[{"values":[0,0,0,0,0,0,0,0,0,0,0,0,0,0],"quality":"Equal","isMain":false,"metricName":"# similar correct contigs"},{"values":[0,0,0,0,0,0,0,0,0,0,0,0,0,0],"quality":"Equal","isMain":false,"metricName":"# similar misassembled blocks"}]],["Reference statistics",[{"values":[49487,49487,49487,49487,49487,49487,49487,49487,49487,49487,49487,49487,49487,49487],"quality":"Equal","isMain":false,"metricName":"Reference length"},{"values":[1,1,1,1,1,1,1,1,1,1,1,1,1,1],"quality":"Equal","isMain":false,"metricName":"Reference fragments"},{"values":["63.84","63.84","63.84","63.84","63.84","63.84","63.84","63.84","63.84","63.84","63.84","63.84","63.84","63.84"],"quality":"Equal","isMain":false,"metricName":"Reference GC (%)"}]]],"referenceName":"gi\_203463359\_ref\_NC","date":"18 July 2018, Wednesday, 16:50:30","order":[0,1,2,3,4,5,6,7,8,9,10,11,12,13],"assembliesNames":["ABySS\_127","ABySS\_63","CLC","IDBA\_UD","MEGAHIT","MIRA","Ray\_Meta","SOAPdenovo2","SPAdes","SPAdes\_meta","SPAdes\_sc","SPAdes\_sc\_careful","Velvet","Geneious"]},{"assembliesWithNs":null,"minContig":500,"report":[["Genome statistics",[{"values":["32.877","24.555","24.738","25.028","31.055","37.173","21.637","1.328","43.134","3.565","7.290","33.140","23.469","8.234"],"quality":"More is better","isMain":true,"metricName":"Genome fraction (%)"},{"values":["1.012","1.009","1.009","1.027","1.028","1.081","1.021","1.267","1.023","0.999","1.001","1.003","1.001","0.999"],"quality":"Less is better","isMain":true,"metricName":"Duplication ratio"},{"values":[6468,4693,5591,4102,3800,7444,5982,999,12009,1168,3016,37157,4315,5363],"quality":"More is better","isMain":true,"metricName":"Largest alignment"},{"values":[51711,38478,38782,39949,49595,62407,34291,2063,68060,5536,11336,51487,36504,12778],"quality":"More is better","isMain":true,"metricName":"Total aligned length"},{"values":[3059,2543,1360,1617,1279,2038,1966,606,5492,874,1302,37157,1621,4723],"quality":"More is better","isMain":false,"metricName":"NA50"},{"values":[959,918,1079,840,914,1306,1353,458,1106,721,889,3016,1073,4723],"quality":"More is better","isMain":false,"metricName":"NA75"},{"values":[null,null,null,null,null,null,null,null,null,null,null,null,null,null],"quality":"More is better","isMain":true,"metricName":"NGA50"},{"values":[6,6,9,8,14,8,5,2,4,3,3,1,8,2],"quality":"Less is better","isMain":false,"metricName":"LA50"},{"values":[14,12,17,18,25,18,10,3,12,5,6,2,15,2],"quality":"Less is better","isMain":false,"metricName":"LA75"}]],["Misassemblies",[{"values":[0,0,0,0,0,0,0,0,0,0,0,0,0,0],"quality":"Less is better","isMain":true,"metricName":"# misassemblies"},{"values":[0,0,0,0,0,0,0,0,0,0,0,0,0,0],"quality":"Less is better","isMain":false,"metricName":" # relocations"},{"values":[0,0,0,0,0,0,0,0,0,0,0,0,0,0],"quality":"Less is better","isMain":false,"metricName":" # translocations"},{"values":[0,0,0,0,0,0,0,0,0,0,0,0,0,0],"quality":"Less is better","isMain":false,"metricName":" # inversions"},{"values":[0,0,0,0,0,0,0,0,0,0,0,0,0,0],"quality":"Less is better","isMain":false,"metricName":"# misassembled contigs"},{"values":[0,0,0,0,0,0,0,0,0,0,0,0,0,0],"quality":"Less is better","isMain":true,"metricName":"Misassembled contigs length"},{"values":[0,0,0,0,0,0,0,0,0,0,0,0,0,1],"quality":"Less is better","isMain":false,"metricName":"# local misassemblies"},{"values":[0,0,0,0,0,0,0,0,0,0,0,0,0,0],"quality":"Less is better","isMain":false,"metricName":"# unaligned mis. contigs"}]],["Unaligned",[{"values":[0,0,0,0,0,0,0,0,0,0,0,0,0,0],"quality":"Less is better","isMain":false,"metricName":"# fully unaligned contigs"},{"values":[0,0,0,0,0,0,0,0,0,0,0,0,0,0],"quality":"Less is better","isMain":false,"metricName":"Fully unaligned length"},{"values":[0,0,0,0,0,0,0,0,0,0,0,0,0,0],"quality":"Less is better","isMain":false,"metricName":"# partially unaligned contigs"},{"values":[0,0,0,0,0,0,0,0,0,0,0,0,0,0],"quality":"Less is better","isMain":false,"metricName":"Partially unaligned length"}]],["Mismatches",[{"values":[5,4,9,19,58,376,39,0,90,2,44,222,23,115],"quality":"Less is better","isMain":false,"metricName":"# mismatches"},{"values":[0,0,1,0,0,30,0,2,7,1,3,12,0,20],"quality":"Less is better","isMain":false,"metricName":"# indels"},{"values":[0,0,12,0,0,94,0,96,110,3,9,29,0,33],"quality":"Less is better","isMain":false,"metricName":"Indels length"},{"values":["9.79","10.48","23.42","48.86","120.20","651.01","116.01","0.00","134.29","36.11","388.45","431.15","63.07","898.93"],"quality":"Less is better","isMain":true,"metricName":"# mismatches per 100 kbp"},{"values":["0.00","0.00","2.60","0.00","0.00","51.94","0.00","96.95","10.44","18.05","26.49","23.31","0.00","156.34"],"quality":"Less is better","isMain":true,"metricName":"# indels per 100 kbp"},{"values":[0,0,0,0,0,25,0,0,3,1,3,11,0,20],"quality":"Less is better","isMain":false,"metricName":" # indels (<= 5 bp)"},{"values":[0,0,1,0,0,5,0,2,4,0,0,1,0,0],"quality":"Less is better","isMain":false,"metricName":" # indels (> 5 bp)"},{"values":[0,1,0,0,0,13,0,437,477,0,0,162,0,4],"quality":"Less is better","isMain":false,"metricName":"# N's"},{"values":["0.00","2.60","0.00","0.00","0.00","20.83","0.00","16717.67","695.97","0.00","0.00","313.64","0.00","31.29"],"quality":"Less is better","isMain":true,"metricName":"# N's per 100 kbp"}]],["Statistics without reference",[{"values":[32,26,30,32,42,36,19,3,33,7,9,12,28,3],"quality":"Equal","isMain":true,"metricName":"# contigs"},{"values":[13,11,17,12,23,22,13,1,18,1,5,7,16,3],"quality":"Equal","isMain":false,"metricName":"# contigs (>= 1000 bp)"},{"values":[2,0,1,0,0,1,1,0,4,0,0,1,0,1],"quality":"Equal","isMain":false,"metricName":"# contigs (>= 5000 bp)"},{"values":[0,0,0,0,0,0,0,0,2,0,0,1,0,0],"quality":"Equal","isMain":false,"metricName":"# contigs (>= 10000 bp)"},{"values":[0,0,0,0,0,0,0,0,0,0,0,1,0,0],"quality":"Equal","isMain":false,"metricName":"# contigs (>= 25000 bp)"},{"values":[0,0,0,0,0,0,0,0,0,0,0,0,0,0],"quality":"Equal","isMain":false,"metricName":"# contigs (>= 50000 bp)"},{"values":[6468,4693,5591,4102,3800,7444,5982,1104,12009,1168,3016,37319,4315,5363],"quality":"More is better","isMain":true,"metricName":"Largest contig"},{"values":[51711,38478,38782,39949,49595,62407,34313,2614,68537,5536,11336,51652,36504,12782],"quality":"More is better","isMain":true,"metricName":"Total length"},{"values":[38436,28080,29777,25004,36140,52528,29574,1104,58457,1168,8275,47894,28918,12782],"quality":"More is better","isMain":true,"metricName":"Total length (>= 1000 bp)"},{"values":[11960,0,5591,0,0,7444,5982,0,38473,0,0,37319,0,5363],"quality":"More is better","isMain":false,"metricName":"Total length (>= 5000 bp)"},{"values":[0,0,0,0,0,0,0,0,23905,0,0,37319,0,0],"quality":"More is better","isMain":true,"metricName":"Total length (>= 10000 bp)"},{"values":[0,0,0,0,0,0,0,0,0,0,0,37319,0,0],"quality":"More is better","isMain":false,"metricName":"Total length (>= 25000 bp)"},{"values":[0,0,0,0,0,0,0,0,0,0,0,0,0,0],"quality":"More is better","isMain":true,"metricName":"Total length (>= 50000 bp)"},{"values":[3059,2543,1360,1617,1279,2038,1966,831,5492,874,1302,37319,1621,4723],"quality":"More is better","isMain":false,"metricName":"N50"},{"values":[959,918,1079,840,914,1306,1353,679,1106,721,889,3016,1073,4723],"quality":"More is better","isMain":false,"metricName":"N75"},{"values":[6,6,9,8,14,8,5,2,4,3,3,1,8,2],"quality":"Less is better","isMain":false,"metricName":"L50"},{"values":[14,12,17,18,25,18,10,3,12,5,6,2,15,2],"quality":"Less is better","isMain":false,"metricName":"L75"},{"values":["64.14","64.61","64.22","64.42","64.30","64.29","65.52","63.76","64.54","63.33","64.67","65.25","64.70","64.78"],"quality":"Equal","isMain":false,"metricName":"GC (%)"}]],["Predicted genes",[]],["Similarity statistics",[{"values":[0,0,0,0,0,0,0,0,0,0,0,0,0,0],"quality":"Equal","isMain":false,"metricName":"# similar correct contigs"},{"values":[0,0,0,0,0,0,0,0,0,0,0,0,0,0],"quality":"Equal","isMain":false,"metricName":"# similar misassembled blocks"}]],["Reference statistics",[{"values":[155372,155372,155372,155372,155372,155372,155372,155372,155372,155372,155372,155372,155372,155372],"quality":"Equal","isMain":false,"metricName":"Reference length"},{"values":[1,1,1,1,1,1,1,1,1,1,1,1,1,1],"quality":"Equal","isMain":false,"metricName":"Reference fragments"},{"values":["64.72","64.72","64.72","64.72","64.72","64.72","64.72","64.72","64.72","64.72","64.72","64.72","64.72","64.72"],"quality":"Equal","isMain":false,"metricName":"Reference GC (%)"}]]],"referenceName":"gi\_204305660\_ref\_NC","date":"18 July 2018, Wednesday, 16:50:47","order":[0,1,2,3,4,5,6,7,8,9,10,11,12,13],"assembliesNames":["ABySS\_127","ABySS\_63","CLC","IDBA\_UD","MEGAHIT","MIRA","Ray\_Meta","SOAPdenovo2","SPAdes","SPAdes\_meta","SPAdes\_sc","SPAdes\_sc\_careful","Velvet","Geneious"]},{"assembliesWithNs":null,"minContig":500,"report":[["Genome statistics",[{"values":["100.000","49.562","100.000","69.310","100.000","100.000","19.992","100.000","38.285","47.451","54.892","20.739"],"quality":"More is better","isMain":true,"metricName":"Genome fraction (%)"},{"values":["1.002","1.015","1.000","1.000","1.021","1.034","1.876","1.020","1.031","1.025","1.022","1.000"],"quality":"Less is better","isMain":true,"metricName":"Duplication ratio"},{"values":[7785,1581,7768,2173,6502,8031,580,7923,912,912,912,980],"quality":"More is better","isMain":true,"metricName":"Largest alignment"},{"values":[7785,3907,7768,5384,7931,8031,1553,7923,3066,3778,4356,1611],"quality":"More is better","isMain":true,"metricName":"Total aligned length"},{"values":[7785,674,7768,1175,6502,8031,784,7923,null,null,534,null],"quality":"More is better","isMain":false,"metricName":"NG50"},{"values":[7785,null,7768,null,6502,8031,null,7923,null,null,null,null],"quality":"More is better","isMain":false,"metricName":"NG75"},{"values":[7785,926,7768,1418,6502,8031,null,7923,819,801,801,980],"quality":"More is better","isMain":false,"metricName":"NA50"},{"values":[7785,726,7768,1175,6502,8031,null,7923,801,712,578,631],"quality":"More is better","isMain":false,"metricName":"NA75"},{"values":[7785,674,7768,1175,6502,8031,null,7923,null,null,534,null],"quality":"More is better","isMain":true,"metricName":"NGA50"},{"values":[7785,null,7768,null,6502,8031,null,7923,null,null,null,null],"quality":"More is better","isMain":false,"metricName":"NGA75"},{"values":[1,4,1,3,1,1,4,1,null,null,6,null],"quality":"Less is better","isMain":false,"metricName":"LG50"},{"values":[1,null,1,null,1,1,null,1,null,null,null,null],"quality":"Less is better","isMain":false,"metricName":"LG75"},{"values":[1,2,1,2,1,1,null,1,2,3,3,1],"quality":"Less is better","isMain":false,"metricName":"LA50"},{"values":[1,3,1,3,1,1,null,1,3,4,5,2],"quality":"Less is better","isMain":false,"metricName":"LA75"},{"values":[1,4,1,3,1,1,null,1,null,null,6,null],"quality":"Less is better","isMain":true,"metricName":"LGA50"},{"values":[1,null,1,null,1,1,null,1,null,null,null,null],"quality":"Less is better","isMain":false,"metricName":"LGA75"}]],["Misassemblies",[{"values":[0,0,0,0,0,0,0,0,0,0,0,0],"quality":"Less is better","isMain":true,"metricName":"# misassemblies"},{"values":[0,0,0,0,0,0,0,0,0,0,0,0],"quality":"Less is better","isMain":false,"metricName":" # relocations"},{"values":[0,0,0,0,0,0,0,0,0,0,0,0],"quality":"Less is better","isMain":false,"metricName":" # translocations"},{"values":[0,0,0,0,0,0,0,0,0,0,0,0],"quality":"Less is better","isMain":false,"metricName":" # inversions"},{"values":[0,0,0,0,0,0,0,0,0,0,0,0],"quality":"Less is better","isMain":false,"metricName":"# misassembled contigs"},{"values":[0,0,0,0,0,0,0,0,0,0,0,0],"quality":"Less is better","isMain":true,"metricName":"Misassembled contigs length"},{"values":[0,0,0,0,0,1,0,1,0,0,0,0],"quality":"Less is better","isMain":false,"metricName":"# local misassemblies"},{"values":[0,0,0,0,0,0,1,0,0,0,0,0],"quality":"Less is better","isMain":false,"metricName":"# unaligned mis. contigs"}]],["Unaligned",[{"values":[0,0,0,0,0,0,0,0,0,0,0,0],"quality":"Less is better","isMain":false,"metricName":"# fully unaligned contigs"},{"values":[0,0,0,0,0,0,0,0,0,0,0,0],"quality":"Less is better","isMain":false,"metricName":"Fully unaligned length"},{"values":[0,0,0,0,0,0,1,0,0,0,0,0],"quality":"Less is better","isMain":false,"metricName":"# partially unaligned contigs"},{"values":[0,0,0,0,0,0,1540,0,0,0,0,0],"quality":"Less is better","isMain":false,"metricName":"Partially unaligned length"}]],["Mismatches",[{"values":[0,0,1,5,2,0,0,1,0,0,12,2],"quality":"Less is better","isMain":false,"metricName":"# mismatches"},{"values":[1,0,0,0,0,0,5,0,0,0,0,0],"quality":"Less is better","isMain":false,"metricName":"# indels"},{"values":[17,0,0,0,0,0,250,0,0,0,0,0],"quality":"Less is better","isMain":false,"metricName":"Indels length"},{"values":["0.00","0.00","12.87","92.87","25.75","0.00","0.00","12.87","0.00","0.00","281.43","124.15"],"quality":"Less is better","isMain":true,"metricName":"# mismatches per 100 kbp"},{"values":["12.87","0.00","0.00","0.00","0.00","0.00","321.96","0.00","0.00","0.00","0.00","0.00"],"quality":"Less is better","isMain":true,"metricName":"# indels per 100 kbp"},{"values":[0,0,0,0,0,0,0,0,0,0,0,0],"quality":"Less is better","isMain":false,"metricName":" # indels (<= 5 bp)"},{"values":[1,0,0,0,0,0,5,0,0,0,0,0],"quality":"Less is better","isMain":false,"metricName":" # indels (> 5 bp)"},{"values":[0,0,0,0,0,1,2262,0,0,0,0,0],"quality":"Less is better","isMain":false,"metricName":"# N's"},{"values":["0.00","0.00","0.00","0.00","0.00","12.45","50797.22","0.00","0.00","0.00","0.00","0.00"],"quality":"Less is better","isMain":true,"metricName":"# N's per 100 kbp"}]],["Statistics without reference",[{"values":[1,4,1,4,2,1,4,1,4,5,6,2],"quality":"Equal","isMain":true,"metricName":"# contigs"},{"values":[1,1,1,3,2,1,2,1,0,0,0,0],"quality":"Equal","isMain":false,"metricName":"# contigs (>= 1000 bp)"},{"values":[1,0,1,0,1,1,0,1,0,0,0,0],"quality":"Equal","isMain":false,"metricName":"# contigs (>= 5000 bp)"},{"values":[0,0,0,0,0,0,0,0,0,0,0,0],"quality":"Equal","isMain":false,"metricName":"# contigs (>= 10000 bp)"},{"values":[0,0,0,0,0,0,0,0,0,0,0,0],"quality":"Equal","isMain":false,"metricName":"# contigs (>= 25000 bp)"},{"values":[0,0,0,0,0,0,0,0,0,0,0,0],"quality":"Equal","isMain":false,"metricName":"# contigs (>= 50000 bp)"},{"values":[7785,1581,7768,2173,6502,8031,1663,7923,912,912,912,980],"quality":"More is better","isMain":true,"metricName":"Largest contig"},{"values":[7785,3907,7768,5384,7931,8031,4453,7923,3066,3778,4356,1611],"quality":"More is better","isMain":true,"metricName":"Total length"},{"values":[7785,1581,7768,4766,7931,8031,2773,7923,0,0,0,0],"quality":"More is better","isMain":true,"metricName":"Total length (>= 1000 bp)"},{"values":[7785,0,7768,0,6502,8031,0,7923,0,0,0,0],"quality":"More is better","isMain":false,"metricName":"Total length (>= 5000 bp)"},{"values":[0,0,0,0,0,0,0,0,0,0,0,0],"quality":"More is better","isMain":true,"metricName":"Total length (>= 10000 bp)"},{"values":[0,0,0,0,0,0,0,0,0,0,0,0],"quality":"More is better","isMain":false,"metricName":"Total length (>= 25000 bp)"},{"values":[0,0,0,0,0,0,0,0,0,0,0,0],"quality":"More is better","isMain":true,"metricName":"Total length (>= 50000 bp)"},{"values":[7785,926,7768,1418,6502,8031,1110,7923,819,801,801,980],"quality":"More is better","isMain":false,"metricName":"N50"},{"values":[7785,726,7768,1175,6502,8031,896,7923,801,712,578,631],"quality":"More is better","isMain":false,"metricName":"N75"},{"values":[1,2,1,2,1,1,2,1,2,3,3,1],"quality":"Less is better","isMain":false,"metricName":"L50"},{"values":[1,3,1,3,1,1,3,1,3,4,5,2],"quality":"Less is better","isMain":false,"metricName":"L75"},{"values":["23.15","23.39","23.15","24.13","23.12","23.08","25.70","23.14","24.79","24.83","24.95","25.95"],"quality":"Equal","isMain":false,"metricName":"GC (%)"}]],["Predicted genes",[]],["Similarity statistics",[{"values":[0,0,0,0,0,0,0,0,0,0,0,0],"quality":"Equal","isMain":false,"metricName":"# similar correct contigs"},{"values":[0,0,0,0,0,0,0,0,0,0,0,0],"quality":"Equal","isMain":false,"metricName":"# similar misassembled blocks"}]],["Reference statistics",[{"values":[7768,7768,7768,7768,7768,7768,7768,7768,7768,7768,7768,7768],"quality":"Equal","isMain":false,"metricName":"Reference length"},{"values":[1,1,1,1,1,1,1,1,1,1,1,1],"quality":"Equal","isMain":false,"metricName":"Reference fragments"},{"values":["23.16","23.16","23.16","23.16","23.16","23.16","23.16","23.16","23.16","23.16","23.16","23.16"],"quality":"Equal","isMain":false,"metricName":"Reference GC (%)"}]]],"referenceName":"gi\_20514675\_ref\_NC","date":"18 July 2018, Wednesday, 16:51:03","order":[0,1,2,3,4,5,6,7,8,9,10,11],"assembliesNames":["ABySS\_127","ABySS\_63","CLC","IDBA\_UD","MEGAHIT","MIRA","SOAPdenovo2","SPAdes","SPAdes\_meta","SPAdes\_sc","SPAdes\_sc\_careful","Velvet"]},{"assembliesWithNs":null,"minContig":500,"report":[["Genome statistics",[{"values":["100.000","97.573","100.000","99.479","99.658","100.000","100.000","99.598","100.000","97.917","97.917","88.903","100.000"],"quality":"More is better","isMain":true,"metricName":"Genome fraction (%)"},{"values":["1.009","1.000","1.023","1.005","1.005","1.013","1.010","1.000","1.002","1.002","1.002","1.005","1.006"],"quality":"Less is better","isMain":true,"metricName":"Duplication ratio"},{"values":[47498,42227,47057,41549,43484,46746,47507,46868,45143,41461,41461,12167,47352],"quality":"More is better","isMain":true,"metricName":"Largest alignment"},{"values":[47498,45915,48157,47024,47141,47651,47507,46868,47167,46187,46187,42037,47352],"quality":"More is better","isMain":true,"metricName":"Total aligned length"},{"values":[47498,42227,47057,41549,43484,46746,48696,46868,45143,41461,41461,3607,47352],"quality":"More is better","isMain":false,"metricName":"NG50"},{"values":[47498,42227,47057,41549,43484,46746,48696,46868,45143,41461,41461,1433,47352],"quality":"More is better","isMain":false,"metricName":"NG75"},{"values":[47498,42227,47057,41549,43484,46746,47507,46868,45143,41461,41461,3607,47352],"quality":"More is better","isMain":false,"metricName":"NA50"},{"values":[47498,42227,47057,41549,43484,46746,47507,46868,45143,41461,41461,2075,47352],"quality":"More is better","isMain":false,"metricName":"NA75"},{"values":[47498,42227,47057,41549,43484,46746,47507,46868,45143,41461,41461,3607,47352],"quality":"More is better","isMain":true,"metricName":"NGA50"},{"values":[47498,42227,47057,41549,43484,46746,47507,46868,45143,41461,41461,1433,47352],"quality":"More is better","isMain":false,"metricName":"NGA75"},{"values":[1,1,1,1,1,1,1,1,1,1,1,4,1],"quality":"Less is better","isMain":false,"metricName":"LG50"},{"values":[1,1,1,1,1,1,1,1,1,1,1,10,1],"quality":"Less is better","isMain":false,"metricName":"LG75"},{"values":[1,1,1,1,1,1,1,1,1,1,1,4,1],"quality":"Less is better","isMain":false,"metricName":"LA50"},{"values":[1,1,1,1,1,1,1,1,1,1,1,8,1],"quality":"Less is better","isMain":false,"metricName":"LA75"},{"values":[1,1,1,1,1,1,1,1,1,1,1,4,1],"quality":"Less is better","isMain":true,"metricName":"LGA50"},{"values":[1,1,1,1,1,1,1,1,1,1,1,10,1],"quality":"Less is better","isMain":false,"metricName":"LGA75"}]],["Misassemblies",[{"values":[0,0,0,0,0,0,0,0,0,0,0,0,0],"quality":"Less is better","isMain":true,"metricName":"# misassemblies"},{"values":[0,0,0,0,0,0,0,0,0,0,0,0,0],"quality":"Less is better","isMain":false,"metricName":" # relocations"},{"values":[0,0,0,0,0,0,0,0,0,0,0,0,0],"quality":"Less is better","isMain":false,"metricName":" # translocations"},{"values":[0,0,0,0,0,0,0,0,0,0,0,0,0],"quality":"Less is better","isMain":false,"metricName":" # inversions"},{"values":[0,0,0,0,0,0,0,0,0,0,0,0,0],"quality":"Less is better","isMain":false,"metricName":"# misassembled contigs"},{"values":[0,0,0,0,0,0,0,0,0,0,0,0,0],"quality":"Less is better","isMain":true,"metricName":"Misassembled contigs length"},{"values":[1,0,0,0,0,0,1,0,0,0,0,0,1],"quality":"Less is better","isMain":false,"metricName":"# local misassemblies"},{"values":[0,0,0,0,0,0,0,0,0,0,0,0,0],"quality":"Less is better","isMain":false,"metricName":"# unaligned mis. contigs"}]],["Unaligned",[{"values":[0,0,0,0,0,0,0,0,0,0,0,0,0],"quality":"Less is better","isMain":false,"metricName":"# fully unaligned contigs"},{"values":[0,0,0,0,0,0,0,0,0,0,0,0,0],"quality":"Less is better","isMain":false,"metricName":"Fully unaligned length"},{"values":[0,0,0,0,0,0,1,0,0,0,0,0,0],"quality":"Less is better","isMain":false,"metricName":"# partially unaligned contigs"},{"values":[0,0,0,0,0,0,1189,0,0,0,0,0,0],"quality":"Less is better","isMain":false,"metricName":"Partially unaligned length"}]],["Mismatches",[{"values":[0,0,0,0,0,0,0,0,0,0,0,0,0],"quality":"Less is better","isMain":false,"metricName":"# mismatches"},{"values":[0,0,0,0,0,0,0,0,0,0,0,0,0],"quality":"Less is better","isMain":false,"metricName":"# indels"},{"values":[0,0,0,0,0,0,0,0,0,0,0,0,0],"quality":"Less is better","isMain":false,"metricName":"Indels length"},{"values":["0.00","0.00","0.00","0.00","0.00","0.00","0.00","0.00","0.00","0.00","0.00","0.00","0.00"],"quality":"Less is better","isMain":true,"metricName":"# mismatches per 100 kbp"},{"values":["0.00","0.00","0.00","0.00","0.00","0.00","0.00","0.00","0.00","0.00","0.00","0.00","0.00"],"quality":"Less is better","isMain":true,"metricName":"# indels per 100 kbp"},{"values":[0,0,0,0,0,0,0,0,0,0,0,0,0],"quality":"Less is better","isMain":false,"metricName":" # indels (<= 5 bp)"},{"values":[0,0,0,0,0,0,0,0,0,0,0,0,0],"quality":"Less is better","isMain":false,"metricName":" # indels (> 5 bp)"},{"values":[0,1,0,0,0,0,0,0,0,0,0,0,0],"quality":"Less is better","isMain":false,"metricName":"# N's"},{"values":["0.00","2.18","0.00","0.00","0.00","0.00","0.00","0.00","0.00","0.00","0.00","0.00","0.00"],"quality":"Less is better","isMain":true,"metricName":"# N's per 100 kbp"}]],["Statistics without reference",[{"values":[1,2,3,4,3,2,1,1,2,3,3,18,1],"quality":"Equal","isMain":true,"metricName":"# contigs"},{"values":[1,2,1,3,2,1,1,1,2,2,2,11,1],"quality":"Equal","isMain":false,"metricName":"# contigs (>= 1000 bp)"},{"values":[1,1,1,1,1,1,1,1,1,1,1,1,1],"quality":"Equal","isMain":false,"metricName":"# contigs (>= 5000 bp)"},{"values":[1,1,1,1,1,1,1,1,1,1,1,1,1],"quality":"Equal","isMain":false,"metricName":"# contigs (>= 10000 bp)"},{"values":[1,1,1,1,1,1,1,1,1,1,1,0,1],"quality":"Equal","isMain":false,"metricName":"# contigs (>= 25000 bp)"},{"values":[0,0,0,0,0,0,0,0,0,0,0,0,0],"quality":"Equal","isMain":false,"metricName":"# contigs (>= 50000 bp)"},{"values":[47498,42227,47057,41549,43484,46746,48696,46868,45143,41461,41461,12167,47352],"quality":"More is better","isMain":true,"metricName":"Largest contig"},{"values":[47498,45915,48157,47024,47141,47651,48696,46868,47167,46187,46187,42037,47352],"quality":"More is better","isMain":true,"metricName":"Total length"},{"values":[47498,45915,47057,46250,46419,46746,48696,46868,47167,45299,45479,37344,47352],"quality":"More is better","isMain":true,"metricName":"Total length (>= 1000 bp)"},{"values":[47498,42227,47057,41549,43484,46746,48696,46868,45143,41461,41461,12167,47352],"quality":"More is better","isMain":false,"metricName":"Total length (>= 5000 bp)"},{"values":[47498,42227,47057,41549,43484,46746,48696,46868,45143,41461,41461,12167,47352],"quality":"More is better","isMain":true,"metricName":"Total length (>= 10000 bp)"},{"values":[47498,42227,47057,41549,43484,46746,48696,46868,45143,41461,41461,0,47352],"quality":"More is better","isMain":false,"metricName":"Total length (>= 25000 bp)"},{"values":[0,0,0,0,0,0,0,0,0,0,0,0,0],"quality":"More is better","isMain":true,"metricName":"Total length (>= 50000 bp)"},{"values":[47498,42227,47057,41549,43484,46746,48696,46868,45143,41461,41461,3607,47352],"quality":"More is better","isMain":false,"metricName":"N50"},{"values":[47498,42227,47057,41549,43484,46746,48696,46868,45143,41461,41461,2075,47352],"quality":"More is better","isMain":false,"metricName":"N75"},{"values":[1,1,1,1,1,1,1,1,1,1,1,4,1],"quality":"Less is better","isMain":false,"metricName":"L50"},{"values":[1,1,1,1,1,1,1,1,1,1,1,8,1],"quality":"Less is better","isMain":false,"metricName":"L75"},{"values":["66.68","66.87","66.78","66.80","66.81","66.81","66.82","66.80","66.77","66.88","66.88","67.12","66.79"],"quality":"Equal","isMain":false,"metricName":"GC (%)"}]],["Predicted genes",[]],["Similarity statistics",[{"values":[0,0,0,0,0,0,0,0,0,0,0,0,0],"quality":"Equal","isMain":false,"metricName":"# similar correct contigs"},{"values":[0,0,0,0,0,0,0,0,0,0,0,0,0],"quality":"Equal","isMain":false,"metricName":"# similar misassembled blocks"}]],["Reference statistics",[{"values":[47057,47057,47057,47057,47057,47057,47057,47057,47057,47057,47057,47057,47057],"quality":"Equal","isMain":false,"metricName":"Reference length"},{"values":[1,1,1,1,1,1,1,1,1,1,1,1,1],"quality":"Equal","isMain":false,"metricName":"Reference fragments"},{"values":["66.77","66.77","66.77","66.77","66.77","66.77","66.77","66.77","66.77","66.77","66.77","66.77","66.77"],"quality":"Equal","isMain":false,"metricName":"Reference GC (%)"}]]],"referenceName":"gi\_206599544\_ref\_NC","date":"18 July 2018, Wednesday, 16:51:19","order":[0,1,2,3,4,5,6,7,8,9,10,11,12],"assembliesNames":["ABySS\_127","ABySS\_63","CLC","IDBA\_UD","MEGAHIT","MIRA","Ray\_Meta","SPAdes","SPAdes\_meta","SPAdes\_sc","SPAdes\_sc\_careful","Velvet","Geneious"]},{"assembliesWithNs":null,"minContig":500,"report":[["Genome statistics",[{"values":["90.651","86.880","98.958","99.761","100.000","98.366","77.763","100.000","100.000","99.199","96.693","95.558","74.713"],"quality":"More is better","isMain":true,"metricName":"Genome fraction (%)"},{"values":["1.007","1.007","1.011","1.000","1.019","1.033","1.032","1.002","1.005","1.004","1.009","1.000","1.000"],"quality":"Less is better","isMain":true,"metricName":"Duplication ratio"},{"values":[11443,6749,19257,48375,25152,26141,6733,64934,64912,42827,25287,29280,48410],"quality":"More is better","isMain":true,"metricName":"Largest alignment"},{"values":[59147,56691,64819,64652,66029,65834,50682,64934,64912,64333,62765,61928,48410],"quality":"More is better","isMain":true,"metricName":"Total aligned length"},{"values":[2417,1842,15490,48375,10836,22385,2385,64934,65101,42827,16811,17043,48410],"quality":"More is better","isMain":false,"metricName":"NG50"},{"values":[1419,1057,9485,16277,8724,10302,1586,64934,65101,21750,16228,9204,null],"quality":"More is better","isMain":false,"metricName":"NG75"},{"values":[2519,2161,15490,48375,10836,22385,2105,64934,64912,42827,16697,17043,48410],"quality":"More is better","isMain":false,"metricName":"NA50"},{"values":[1579,1325,9485,16277,8724,10302,1505,64934,64912,21506,15985,9204,48410],"quality":"More is better","isMain":false,"metricName":"NA75"},{"values":[2417,1842,15490,48375,10836,22385,1821,64934,64912,42827,16697,17043,48410],"quality":"More is better","isMain":true,"metricName":"NGA50"},{"values":[1419,1057,9485,16277,8724,10302,822,64934,64912,21506,15985,9204,null],"quality":"More is better","isMain":false,"metricName":"NGA75"},{"values":[7,11,2,1,2,2,8,1,1,1,2,2,1],"quality":"Less is better","isMain":false,"metricName":"LG50"},{"values":[16,23,4,2,4,3,17,1,1,2,3,3,null],"quality":"Less is better","isMain":false,"metricName":"LG75"},{"values":[6,9,2,1,2,2,8,1,1,1,2,2,1],"quality":"Less is better","isMain":false,"metricName":"LA50"},{"values":[13,18,4,2,4,3,16,1,1,2,3,3,1],"quality":"Less is better","isMain":false,"metricName":"LA75"},{"values":[7,11,2,1,2,2,10,1,1,1,2,2,1],"quality":"Less is better","isMain":true,"metricName":"LGA50"},{"values":[16,23,4,2,4,3,21,1,1,2,3,3,null],"quality":"Less is better","isMain":false,"metricName":"LGA75"}]],["Misassemblies",[{"values":[0,0,0,0,0,0,0,0,0,0,0,0,0],"quality":"Less is better","isMain":true,"metricName":"# misassemblies"},{"values":[0,0,0,0,0,0,0,0,0,0,0,0,0],"quality":"Less is better","isMain":false,"metricName":" # relocations"},{"values":[0,0,0,0,0,0,0,0,0,0,0,0,0],"quality":"Less is better","isMain":false,"metricName":" # translocations"},{"values":[0,0,0,0,0,0,0,0,0,0,0,0,0],"quality":"Less is better","isMain":false,"metricName":" # inversions"},{"values":[0,0,0,0,0,0,0,0,0,0,0,0,0],"quality":"Less is better","isMain":false,"metricName":"# misassembled contigs"},{"values":[0,0,0,0,0,0,0,0,0,0,0,0,0],"quality":"Less is better","isMain":true,"metricName":"Misassembled contigs length"},{"values":[0,0,0,0,0,0,0,1,1,0,0,0,0],"quality":"Less is better","isMain":false,"metricName":"# local misassemblies"},{"values":[0,0,0,0,0,0,2,0,0,0,0,0,0],"quality":"Less is better","isMain":false,"metricName":"# unaligned mis. contigs"}]],["Unaligned",[{"values":[0,0,0,0,0,0,0,0,0,0,0,0,0],"quality":"Less is better","isMain":false,"metricName":"# fully unaligned contigs"},{"values":[0,0,0,0,0,0,0,0,0,0,0,0,0],"quality":"Less is better","isMain":false,"metricName":"Fully unaligned length"},{"values":[0,0,0,0,0,0,2,0,0,0,1,0,0],"quality":"Less is better","isMain":false,"metricName":"# partially unaligned contigs"},{"values":[0,0,0,0,0,0,4542,0,0,0,1104,0,0],"quality":"Less is better","isMain":false,"metricName":"Partially unaligned length"}]],["Mismatches",[{"values":[19,20,46,31,71,34,28,44,37,41,35,37,19],"quality":"Less is better","isMain":false,"metricName":"# mismatches"},{"values":[0,0,0,0,0,1,16,0,1,1,1,0,1],"quality":"Less is better","isMain":false,"metricName":"# indels"},{"values":[0,0,0,0,0,9,767,0,9,9,9,0,9],"quality":"Less is better","isMain":false,"metricName":"Indels length"},{"values":["32.34","35.52","71.73","47.95","109.56","53.34","55.56","67.89","57.09","63.78","55.85","59.75","39.24"],"quality":"Less is better","isMain":true,"metricName":"# mismatches per 100 kbp"},{"values":["0.00","0.00","0.00","0.00","0.00","1.57","31.75","0.00","1.54","1.56","1.60","0.00","2.07"],"quality":"Less is better","isMain":true,"metricName":"# indels per 100 kbp"},{"values":[0,0,0,0,0,0,0,0,0,0,0,0,0],"quality":"Less is better","isMain":false,"metricName":" # indels (<= 5 bp)"},{"values":[0,0,0,0,0,1,16,0,1,1,1,0,1],"quality":"Less is better","isMain":false,"metricName":" # indels (> 5 bp)"},{"values":[0,0,0,0,0,56,2535,0,0,244,1576,0,36],"quality":"Less is better","isMain":false,"metricName":"# N's"},{"values":["0.00","0.00","0.00","0.00","0.00","85.06","4483.71","0.00","0.00","377.84","2449.45","0.00","74.36"],"quality":"Less is better","isMain":true,"metricName":"# N's per 100 kbp"}]],["Statistics without reference",[{"values":[25,33,8,2,8,7,23,1,1,2,5,5,1],"quality":"Equal","isMain":true,"metricName":"# contigs"},{"values":[20,24,6,2,6,4,22,1,1,2,4,5,1],"quality":"Equal","isMain":false,"metricName":"# contigs (>= 1000 bp)"},{"values":[2,1,5,2,5,4,4,1,1,2,3,3,1],"quality":"Equal","isMain":false,"metricName":"# contigs (>= 5000 bp)"},{"values":[1,0,3,2,3,3,0,1,1,2,3,2,1],"quality":"Equal","isMain":false,"metricName":"# contigs (>= 10000 bp)"},{"values":[0,0,0,1,1,1,0,1,1,1,1,1,1],"quality":"Equal","isMain":false,"metricName":"# contigs (>= 25000 bp)"},{"values":[0,0,0,0,0,0,0,1,1,0,0,0,0],"quality":"Equal","isMain":false,"metricName":"# contigs (>= 50000 bp)"},{"values":[11443,6749,19257,48375,25152,26141,6837,64934,65101,42827,26391,29280,48410],"quality":"More is better","isMain":true,"metricName":"Largest contig"},{"values":[59150,56691,64819,64652,66029,65834,56538,64934,65101,64577,64341,61928,48410],"quality":"More is better","isMain":true,"metricName":"Total length"},{"values":[55055,50189,63682,64652,64494,63855,55798,64934,65101,64577,63638,61928,48410],"quality":"More is better","isMain":true,"metricName":"Total length (>= 1000 bp)"},{"values":[16927,6749,62452,64652,60488,63855,22865,64934,65101,64577,59430,55527,48410],"quality":"More is better","isMain":false,"metricName":"Total length (>= 5000 bp)"},{"values":[11443,0,46486,64652,46500,58828,0,64934,65101,64577,59430,46323,48410],"quality":"More is better","isMain":true,"metricName":"Total length (>= 10000 bp)"},{"values":[0,0,0,48375,25152,26141,0,64934,65101,42827,26391,29280,48410],"quality":"More is better","isMain":false,"metricName":"Total length (>= 25000 bp)"},{"values":[0,0,0,0,0,0,0,64934,65101,0,0,0,0],"quality":"More is better","isMain":true,"metricName":"Total length (>= 50000 bp)"},{"values":[2519,2161,15490,48375,10836,22385,2609,64934,65101,42827,16811,17043,48410],"quality":"More is better","isMain":false,"metricName":"N50"},{"values":[1582,1325,9485,16277,8724,10302,1698,64934,65101,21750,16228,9204,48410],"quality":"More is better","isMain":false,"metricName":"N75"},{"values":[6,9,2,1,2,2,6,1,1,1,2,2,1],"quality":"Less is better","isMain":false,"metricName":"L50"},{"values":[13,18,4,2,4,3,13,1,1,2,3,3,1],"quality":"Less is better","isMain":false,"metricName":"L75"},{"values":["59.52","59.54","59.57","59.55","59.57","59.53","59.65","59.57","59.55","59.51","59.49","59.54","59.62"],"quality":"Equal","isMain":false,"metricName":"GC (%)"}]],["Predicted genes",[]],["Similarity statistics",[{"values":[0,0,0,0,0,0,0,0,0,0,0,0,0],"quality":"Equal","isMain":false,"metricName":"# similar correct contigs"},{"values":[0,0,0,0,0,0,0,0,0,0,0,0,0],"quality":"Equal","isMain":false,"metricName":"# similar misassembled blocks"}]],["Reference statistics",[{"values":[64807,64807,64807,64807,64807,64807,64807,64807,64807,64807,64807,64807,64807],"quality":"Equal","isMain":false,"metricName":"Reference length"},{"values":[1,1,1,1,1,1,1,1,1,1,1,1,1],"quality":"Equal","isMain":false,"metricName":"Reference fragments"},{"values":["59.55","59.55","59.55","59.55","59.55","59.55","59.55","59.55","59.55","59.55","59.55","59.55","59.55"],"quality":"Equal","isMain":false,"metricName":"Reference GC (%)"}]]],"referenceName":"gi\_206599706\_ref\_NC","date":"18 July 2018, Wednesday, 16:51:35","order":[0,1,2,3,4,5,6,7,8,9,10,11,12],"assembliesNames":["ABySS\_127","ABySS\_63","CLC","IDBA\_UD","MEGAHIT","MIRA","SOAPdenovo2","SPAdes","SPAdes\_meta","SPAdes\_sc","SPAdes\_sc\_careful","Velvet","Geneious"]},{"assembliesWithNs":null,"minContig":500,"report":[["Genome statistics",[{"values":["3.088","1.724","31.667","24.210","32.695","10.181","25.245","34.169","36.647","35.454","35.454"],"quality":"More is better","isMain":true,"metricName":"Genome fraction (%)"},{"values":["1.000","1.000","1.040","1.024","1.013","1.000","1.002","1.015","1.000","1.017","1.017"],"quality":"Less is better","isMain":true,"metricName":"Duplication ratio"},{"values":[903,673,1696,1775,1583,1609,1477,1925,1925,1925,1925],"quality":"More is better","isMain":true,"metricName":"Largest alignment"},{"values":[2129,1189,22299,16693,22554,7020,17439,23573,25279,24469,24469],"quality":"More is better","isMain":true,"metricName":"Total aligned length"},{"values":[722,673,745,840,788,1145,839,800,1070,836,836],"quality":"More is better","isMain":false,"metricName":"NA50"},{"values":[722,516,566,599,603,819,671,null,833,514,514],"quality":"More is better","isMain":false,"metricName":"NA75"},{"values":[null,null,null,null,null,null,null,null,null,null,null],"quality":"More is better","isMain":true,"metricName":"NGA50"},{"values":[2,1,13,8,12,3,9,15,10,13,13],"quality":"Less is better","isMain":false,"metricName":"LA50"},{"values":[2,2,23,16,21,5,14,null,17,24,24],"quality":"Less is better","isMain":false,"metricName":"LA75"}]],["Misassemblies",[{"values":[0,0,0,0,0,0,0,0,0,0,0],"quality":"Less is better","isMain":true,"metricName":"# misassemblies"},{"values":[0,0,0,0,0,0,0,0,0,0,0],"quality":"Less is better","isMain":false,"metricName":" # relocations"},{"values":[0,0,0,0,0,0,0,0,0,0,0],"quality":"Less is better","isMain":false,"metricName":" # translocations"},{"values":[0,0,0,0,0,0,0,0,0,0,0],"quality":"Less is better","isMain":false,"metricName":" # inversions"},{"values":[0,0,0,0,0,0,0,0,0,0,0],"quality":"Less is better","isMain":false,"metricName":"# misassembled contigs"},{"values":[0,0,0,0,0,0,0,0,0,0,0],"quality":"Less is better","isMain":true,"metricName":"Misassembled contigs length"},{"values":[0,0,0,0,0,0,0,0,0,1,1],"quality":"Less is better","isMain":false,"metricName":"# local misassemblies"},{"values":[0,0,0,0,0,0,0,1,0,0,0],"quality":"Less is better","isMain":false,"metricName":"# unaligned mis. contigs"}]],["Unaligned",[{"values":[0,0,0,0,0,0,0,0,0,0,0],"quality":"Less is better","isMain":false,"metricName":"# fully unaligned contigs"},{"values":[0,0,0,0,0,0,0,0,0,0,0],"quality":"Less is better","isMain":false,"metricName":"Fully unaligned length"},{"values":[0,0,1,1,1,0,0,2,0,1,1],"quality":"Less is better","isMain":false,"metricName":"# partially unaligned contigs"},{"values":[0,0,3395,3152,3381,0,0,9240,0,6099,6099],"quality":"Less is better","isMain":false,"metricName":"Partially unaligned length"}]],["Mismatches",[{"values":[0,0,52,130,59,2,16,82,34,73,69],"quality":"Less is better","isMain":false,"metricName":"# mismatches"},{"values":[0,0,0,8,1,0,0,8,0,4,4],"quality":"Less is better","isMain":false,"metricName":"# indels"},{"values":[0,0,0,8,3,0,0,8,0,4,4],"quality":"Less is better","isMain":false,"metricName":"Indels length"},{"values":["0.00","0.00","238.15","778.77","261.71","28.49","91.92","348.05","134.55","298.62","282.25"],"quality":"Less is better","isMain":true,"metricName":"# mismatches per 100 kbp"},{"values":["0.00","0.00","0.00","47.92","4.44","0.00","0.00","33.96","0.00","16.36","16.36"],"quality":"Less is better","isMain":true,"metricName":"# indels per 100 kbp"},{"values":[0,0,0,8,1,0,0,8,0,4,4],"quality":"Less is better","isMain":false,"metricName":" # indels (<= 5 bp)"},{"values":[0,0,0,0,0,0,0,0,0,0,0],"quality":"Less is better","isMain":false,"metricName":" # indels (> 5 bp)"},{"values":[0,0,0,0,0,16,0,0,0,0,0],"quality":"Less is better","isMain":false,"metricName":"# N's"},{"values":["0.00","0.00","0.00","0.00","0.00","227.92","0.00","0.00","0.00","0.00","0.00"],"quality":"Less is better","isMain":true,"metricName":"# N's per 100 kbp"}]],["Statistics without reference",[{"values":[3,2,28,18,26,7,21,24,26,26,26],"quality":"Equal","isMain":true,"metricName":"# contigs"},{"values":[0,0,7,6,7,3,4,12,12,11,11],"quality":"Equal","isMain":false,"metricName":"# contigs (>= 1000 bp)"},{"values":[0,0,0,0,0,0,0,1,0,1,1],"quality":"Equal","isMain":false,"metricName":"# contigs (>= 5000 bp)"},{"values":[0,0,0,0,0,0,0,0,0,0,0],"quality":"Equal","isMain":false,"metricName":"# contigs (>= 10000 bp)"},{"values":[0,0,0,0,0,0,0,0,0,0,0],"quality":"Equal","isMain":false,"metricName":"# contigs (>= 25000 bp)"},{"values":[0,0,0,0,0,0,0,0,0,0,0],"quality":"Equal","isMain":false,"metricName":"# contigs (>= 50000 bp)"},{"values":[903,673,3702,4199,4651,1609,1477,6935,1925,6935,6935],"quality":"More is better","isMain":true,"metricName":"Largest contig"},{"values":[2129,1189,26094,20245,26226,7020,17439,33164,25279,30968,30968],"quality":"More is better","isMain":true,"metricName":"Total length"},{"values":[0,0,11909,12100,12761,3930,4948,25091,15180,20766,20766],"quality":"More is better","isMain":true,"metricName":"Total length (>= 1000 bp)"},{"values":[0,0,0,0,0,0,0,6935,0,6935,6935],"quality":"More is better","isMain":false,"metricName":"Total length (>= 5000 bp)"},{"values":[0,0,0,0,0,0,0,0,0,0,0],"quality":"More is better","isMain":true,"metricName":"Total length (>= 10000 bp)"},{"values":[0,0,0,0,0,0,0,0,0,0,0],"quality":"More is better","isMain":false,"metricName":"Total length (>= 25000 bp)"},{"values":[0,0,0,0,0,0,0,0,0,0,0],"quality":"More is better","isMain":true,"metricName":"Total length (>= 50000 bp)"},{"values":[722,673,840,1327,991,1145,839,1477,1070,1226,1226],"quality":"More is better","isMain":false,"metricName":"N50"},{"values":[722,516,706,683,741,819,671,1006,833,833,833],"quality":"More is better","isMain":false,"metricName":"N75"},{"values":[2,1,9,5,8,3,9,5,10,7,7],"quality":"Less is better","isMain":false,"metricName":"L50"},{"values":[2,2,17,10,16,5,14,12,17,14,14],"quality":"Less is better","isMain":false,"metricName":"L75"},{"values":["56.60","56.52","57.58","57.14","57.57","58.20","57.18","57.53","57.57","57.68","57.69"],"quality":"Equal","isMain":false,"metricName":"GC (%)"}]],["Predicted genes",[]],["Similarity statistics",[{"values":[0,0,0,0,0,0,0,0,0,0,0],"quality":"Equal","isMain":false,"metricName":"# similar correct contigs"},{"values":[0,0,0,0,0,0,0,0,0,0,0],"quality":"Equal","isMain":false,"metricName":"# similar misassembled blocks"}]],["Reference statistics",[{"values":[68952,68952,68952,68952,68952,68952,68952,68952,68952,68952,68952],"quality":"Equal","isMain":false,"metricName":"Reference length"},{"values":[1,1,1,1,1,1,1,1,1,1,1],"quality":"Equal","isMain":false,"metricName":"Reference fragments"},{"values":["57.34","57.34","57.34","57.34","57.34","57.34","57.34","57.34","57.34","57.34","57.34"],"quality":"Equal","isMain":false,"metricName":"Reference GC (%)"}]]],"referenceName":"gi\_206599983\_ref\_NC","date":"18 July 2018, Wednesday, 16:51:49","order":[0,1,2,3,4,5,6,7,8,9,10],"assembliesNames":["ABySS\_127","ABySS\_63","CLC","IDBA\_UD","MEGAHIT","MIRA","SOAPdenovo2","SPAdes","SPAdes\_meta","SPAdes\_sc","SPAdes\_sc\_careful"]},{"assembliesWithNs":null,"minContig":500,"report":[["Genome statistics",[{"values":["83.012","46.357","84.314","74.919","86.334","95.607","64.496","10.063","92.609","74.342","73.194","71.747","28.012","100.000"],"quality":"More is better","isMain":true,"metricName":"Genome fraction (%)"},{"values":["1.035","1.005","1.011","1.030","1.077","1.317","1.106","1.083","1.028","1.005","1.008","1.014","1.002","1.189"],"quality":"Less is better","isMain":true,"metricName":"Duplication ratio"},{"values":[5636,4869,6839,4407,4462,23512,4952,1410,9373,9092,5925,9066,4005,35744],"quality":"More is better","isMain":true,"metricName":"Largest alignment"},{"values":[50289,27288,49897,45163,54093,73311,41527,5892,55646,43751,43186,42617,16432,69619],"quality":"More is better","isMain":true,"metricName":"Total aligned length"},{"values":[3287,null,2570,1027,1469,10721,1288,null,3447,5483,1645,1339,null,92535],"quality":"More is better","isMain":false,"metricName":"NG50"},{"values":[795,null,1233,575,732,8788,569,null,2035,null,null,null,null,92535],"quality":"More is better","isMain":false,"metricName":"NG75"},{"values":[2951,1759,3862,1495,1672,8118,1346,904,5527,8221,2387,3191,1155,null],"quality":"More is better","isMain":false,"metricName":"NA50"},{"values":[775,850,1591,936,825,3628,1006,594,1984,3589,1630,1216,754,null],"quality":"More is better","isMain":false,"metricName":"NA75"},{"values":[2661,null,2570,1027,1448,8788,1152,null,3447,5483,1645,1339,null,35744],"quality":"More is better","isMain":true,"metricName":"NGA50"},{"values":[702,null,1233,575,732,7339,null,null,1984,null,null,null,null,13346],"quality":"More is better","isMain":false,"metricName":"NGA75"},{"values":[6,null,7,16,13,2,13,null,5,4,11,9,null,1],"quality":"Less is better","isMain":false,"metricName":"LG50"},{"values":[15,null,15,35,27,4,29,null,10,null,null,null,null,1],"quality":"Less is better","isMain":false,"metricName":"LG75"},{"values":[7,5,5,10,11,3,10,3,4,3,7,4,4,null],"quality":"Less is better","isMain":false,"metricName":"LA50"},{"values":[16,11,11,20,23,7,19,6,10,5,12,11,9,null],"quality":"Less is better","isMain":false,"metricName":"LA75"},{"values":[8,null,7,16,13,2,16,null,5,4,11,9,null,1],"quality":"Less is better","isMain":true,"metricName":"LGA50"},{"values":[20,null,15,35,27,4,null,null,10,null,null,null,null,2],"quality":"Less is better","isMain":false,"metricName":"LGA75"}]],["Misassemblies",[{"values":[0,0,0,0,0,0,0,0,0,0,0,0,0,0],"quality":"Less is better","isMain":true,"metricName":"# misassemblies"},{"values":[0,0,0,0,0,0,0,0,0,0,0,0,0,0],"quality":"Less is better","isMain":false,"metricName":" # relocations"},{"values":[0,0,0,0,0,0,0,0,0,0,0,0,0,0],"quality":"Less is better","isMain":false,"metricName":" # translocations"},{"values":[0,0,0,0,0,0,0,0,0,0,0,0,0,0],"quality":"Less is better","isMain":false,"metricName":" # inversions"},{"values":[0,0,0,0,0,0,0,0,0,0,0,0,0,0],"quality":"Less is better","isMain":false,"metricName":"# misassembled contigs"},{"values":[0,0,0,0,0,0,0,0,0,0,0,0,0,0],"quality":"Less is better","isMain":true,"metricName":"Misassembled contigs length"},{"values":[0,0,0,0,0,0,0,0,0,0,0,0,0,0],"quality":"Less is better","isMain":false,"metricName":"# local misassemblies"},{"values":[0,0,0,0,0,0,1,0,0,0,0,0,0,1],"quality":"Less is better","isMain":false,"metricName":"# unaligned mis. contigs"}]],["Unaligned",[{"values":[0,0,0,0,0,0,0,0,0,0,0,0,0,0],"quality":"Less is better","isMain":false,"metricName":"# fully unaligned contigs"},{"values":[0,0,0,0,0,0,0,0,0,0,0,0,0,0],"quality":"Less is better","isMain":false,"metricName":"Fully unaligned length"},{"values":[1,0,0,0,0,2,1,0,1,0,0,0,0,3],"quality":"Less is better","isMain":false,"metricName":"# partially unaligned contigs"},{"values":[3687,0,0,0,0,6144,2628,0,1861,0,0,0,0,94727],"quality":"Less is better","isMain":false,"metricName":"Partially unaligned length"}]],["Mismatches",[{"values":[16,0,28,2,67,271,103,0,21,264,139,81,6,233],"quality":"Less is better","isMain":false,"metricName":"# mismatches"},{"values":[0,0,2,0,0,3,3,4,4,3,2,1,0,13],"quality":"Less is better","isMain":false,"metricName":"# indels"},{"values":[0,0,2,0,0,7,7,202,6,5,4,3,0,25],"quality":"Less is better","isMain":false,"metricName":"Indels length"},{"values":["32.92","0.00","56.72","4.56","132.54","484.08","272.74","0.00","38.73","606.48","324.33","192.81","36.58","397.92"],"quality":"Less is better","isMain":true,"metricName":"# mismatches per 100 kbp"},{"values":["0.00","0.00","4.05","0.00","0.00","5.36","7.94","67.89","7.38","6.89","4.67","2.38","0.00","22.20"],"quality":"Less is better","isMain":true,"metricName":"# indels per 100 kbp"},{"values":[0,0,2,0,0,3,3,0,4,3,2,1,0,12],"quality":"Less is better","isMain":false,"metricName":" # indels (<= 5 bp)"},{"values":[0,0,0,0,0,0,0,4,0,0,0,0,0,1],"quality":"Less is better","isMain":false,"metricName":" # indels (> 5 bp)"},{"values":[51,0,0,0,0,16,0,489,0,0,0,0,0,1081],"quality":"Less is better","isMain":false,"metricName":"# N's"},{"values":["94.49","0.00","0.00","0.00","0.00","20.04","0.00","7663.38","0.00","0.00","0.00","0.00","0.00","657.76"],"quality":"Less is better","isMain":true,"metricName":"# N's per 100 kbp"}]],["Statistics without reference",[{"values":[31,21,23,37,43,14,29,7,20,11,22,24,16,3],"quality":"Equal","isMain":true,"metricName":"# contigs"},{"values":[13,6,16,17,19,13,19,2,16,9,17,13,5,3],"quality":"Equal","isMain":false,"metricName":"# contigs (>= 1000 bp)"},{"values":[3,0,2,0,0,5,0,0,4,4,1,2,0,3],"quality":"Equal","isMain":false,"metricName":"# contigs (>= 5000 bp)"},{"values":[0,0,0,0,0,2,0,0,0,0,0,0,0,3],"quality":"Equal","isMain":false,"metricName":"# contigs (>= 10000 bp)"},{"values":[0,0,0,0,0,0,0,0,0,0,0,0,0,3],"quality":"Equal","isMain":false,"metricName":"# contigs (>= 25000 bp)"},{"values":[0,0,0,0,0,0,0,0,0,0,0,0,0,1],"quality":"Equal","isMain":false,"metricName":"# contigs (>= 50000 bp)"},{"values":[7507,4869,6839,4407,4462,23899,4952,1410,9373,9092,5925,9066,4005,92535],"quality":"More is better","isMain":true,"metricName":"Largest contig"},{"values":[53976,27288,49897,45163,54432,79851,44382,6381,57590,43751,43186,42617,16432,164346],"quality":"More is better","isMain":true,"metricName":"Total length"},{"values":[42264,16280,45233,31289,38093,78952,36942,2735,54855,42246,39220,34630,9348,164346],"quality":"More is better","isMain":true,"metricName":"Total length (>= 1000 bp)"},{"values":[18345,0,13012,0,0,59982,0,0,29265,31655,5925,15079,0,164346],"quality":"More is better","isMain":false,"metricName":"Total length (>= 5000 bp)"},{"values":[0,0,0,0,0,34620,0,0,0,0,0,0,0,164346],"quality":"More is better","isMain":true,"metricName":"Total length (>= 10000 bp)"},{"values":[0,0,0,0,0,0,0,0,0,0,0,0,0,164346],"quality":"More is better","isMain":false,"metricName":"Total length (>= 25000 bp)"},{"values":[0,0,0,0,0,0,0,0,0,0,0,0,0,92535],"quality":"More is better","isMain":true,"metricName":"Total length (>= 50000 bp)"},{"values":[3287,1759,3862,1495,1672,9235,1500,904,5527,8221,2387,3191,1155,92535],"quality":"More is better","isMain":false,"metricName":"N50"},{"values":[1249,850,1591,936,825,7339,1152,667,2035,3589,1630,1216,754,39285],"quality":"More is better","isMain":false,"metricName":"N75"},{"values":[6,5,5,10,11,3,8,3,4,3,7,4,4,1],"quality":"Less is better","isMain":false,"metricName":"L50"},{"values":[12,11,11,20,23,5,16,5,10,5,12,11,9,2],"quality":"Less is better","isMain":false,"metricName":"L75"},{"values":["60.87","61.21","61.24","61.27","61.33","61.15","61.02","62.39","61.25","61.26","61.22","61.34","60.98","61.18"],"quality":"Equal","isMain":false,"metricName":"GC (%)"}]],["Predicted genes",[]],["Similarity statistics",[{"values":[0,0,0,0,0,0,0,0,0,0,0,0,0,0],"quality":"Equal","isMain":false,"metricName":"# similar correct contigs"},{"values":[0,0,0,0,0,0,0,0,0,0,0,0,0,0],"quality":"Equal","isMain":false,"metricName":"# similar misassembled blocks"}]],["Reference statistics",[{"values":[58554,58554,58554,58554,58554,58554,58554,58554,58554,58554,58554,58554,58554,58554],"quality":"Equal","isMain":false,"metricName":"Reference length"},{"values":[1,1,1,1,1,1,1,1,1,1,1,1,1,1],"quality":"Equal","isMain":false,"metricName":"Reference fragments"},{"values":["61.26","61.26","61.26","61.26","61.26","61.26","61.26","61.26","61.26","61.26","61.26","61.26","61.26","61.26"],"quality":"Equal","isMain":false,"metricName":"Reference GC (%)"}]]],"referenceName":"gi\_206600079\_ref\_NC","date":"18 July 2018, Wednesday, 16:52:07","order":[0,1,2,3,4,5,6,7,8,9,10,11,12,13],"assembliesNames":["ABySS\_127","ABySS\_63","CLC","IDBA\_UD","MEGAHIT","MIRA","Ray\_Meta","SOAPdenovo2","SPAdes","SPAdes\_meta","SPAdes\_sc","SPAdes\_sc\_careful","Velvet","Geneious"]},{"assembliesWithNs":null,"minContig":500,"report":[["Genome statistics",[{"values":["89.822","54.683","89.650","76.095","85.431","94.969","88.057","11.276","90.601","85.720","77.321","82.458","35.698","96.439"],"quality":"More is better","isMain":true,"metricName":"Genome fraction (%)"},{"values":["1.096","1.004","1.031","1.013","1.046","1.409","1.060","1.000","1.021","1.001","1.016","1.009","1.001","1.365"],"quality":"Less is better","isMain":true,"metricName":"Duplication ratio"},{"values":[13404,7486,12864,7747,9407,13544,13868,991,15468,28214,6875,12514,5494,16710],"quality":"More is better","isMain":true,"metricName":"Largest alignment"},{"values":[57671,32156,54111,45136,52371,77946,54223,6605,54173,50260,46005,48739,20934,77109],"quality":"More is better","isMain":true,"metricName":"Total aligned length"},{"values":[4491,645,2378,1261,2210,9235,7433,null,7976,10052,1585,3776,null,92535],"quality":"More is better","isMain":false,"metricName":"NG50"},{"values":[2849,null,1225,569,1122,7273,4281,null,2995,6238,667,848,null,92535],"quality":"More is better","isMain":false,"metricName":"NG75"},{"values":[3530,2781,2378,2005,2244,2599,7433,619,7976,28214,2973,4555,1822,null],"quality":"More is better","isMain":false,"metricName":"NA50"},{"values":[2496,954,1306,1070,1574,597,4032,542,3409,6238,1187,2609,849,null],"quality":"More is better","isMain":false,"metricName":"NA75"},{"values":[3530,645,2378,1261,2210,5101,7433,null,7976,7933,1585,3776,null,13716],"quality":"More is better","isMain":true,"metricName":"NGA50"},{"values":[2496,null,1225,569,1122,3165,3836,null,2995,5349,667,848,null,6880],"quality":"More is better","isMain":false,"metricName":"NGA75"},{"values":[4,16,6,12,8,3,3,null,3,2,8,5,null,1],"quality":"Less is better","isMain":false,"metricName":"LG50"},{"values":[9,null,14,29,17,4,6,null,6,3,22,12,null,1],"quality":"Less is better","isMain":false,"metricName":"LG75"},{"values":[5,4,5,7,7,10,3,4,3,1,5,4,4,null],"quality":"Less is better","isMain":false,"metricName":"LA50"},{"values":[10,9,12,16,13,30,6,7,5,3,12,7,8,null],"quality":"Less is better","isMain":false,"metricName":"LA75"},{"values":[5,16,6,12,8,4,3,null,3,2,8,5,null,2],"quality":"Less is better","isMain":true,"metricName":"LGA50"},{"values":[10,null,14,29,17,8,7,null,6,4,22,12,null,4],"quality":"Less is better","isMain":false,"metricName":"LGA75"}]],["Misassemblies",[{"values":[2,0,0,0,0,0,0,0,0,0,0,0,0,0],"quality":"Less is better","isMain":true,"metricName":"# misassemblies"},{"values":[2,0,0,0,0,0,0,0,0,0,0,0,0,0],"quality":"Less is better","isMain":false,"metricName":" # relocations"},{"values":[0,0,0,0,0,0,0,0,0,0,0,0,0,0],"quality":"Less is better","isMain":false,"metricName":" # translocations"},{"values":[0,0,0,0,0,0,0,0,0,0,0,0,0,0],"quality":"Less is better","isMain":false,"metricName":" # inversions"},{"values":[1,0,0,0,0,0,0,0,0,0,0,0,0,0],"quality":"Less is better","isMain":false,"metricName":"# misassembled contigs"},{"values":[7507,0,0,0,0,0,0,0,0,0,0,0,0,0],"quality":"Less is better","isMain":true,"metricName":"Misassembled contigs length"},{"values":[0,0,0,0,0,3,0,0,0,0,0,0,0,3],"quality":"Less is better","isMain":false,"metricName":"# local misassemblies"},{"values":[0,0,0,0,0,2,0,0,0,0,0,0,0,2],"quality":"Less is better","isMain":false,"metricName":"# unaligned mis. contigs"}]],["Unaligned",[{"values":[0,0,0,0,0,0,0,0,0,0,0,0,0,0],"quality":"Less is better","isMain":false,"metricName":"# fully unaligned contigs"},{"values":[0,0,0,0,0,0,0,0,0,0,0,0,0,0],"quality":"Less is better","isMain":false,"metricName":"Fully unaligned length"},{"values":[1,0,0,0,0,7,0,0,1,1,0,0,0,9],"quality":"Less is better","isMain":false,"metricName":"# partially unaligned contigs"},{"values":[966,0,0,0,0,23531,0,0,1380,2119,0,0,0,132866],"quality":"Less is better","isMain":false,"metricName":"Partially unaligned length"}]],["Mismatches",[{"values":[153,2,5,1,84,459,149,0,17,166,71,39,7,443],"quality":"Less is better","isMain":false,"metricName":"# mismatches"},{"values":[6,0,0,0,3,34,11,0,0,5,1,1,1,42],"quality":"Less is better","isMain":false,"metricName":"# indels"},{"values":[9,0,0,0,3,125,17,0,0,12,3,3,1,136],"quality":"Less is better","isMain":false,"metricName":"Indels length"},{"values":["290.79","6.24","9.52","2.24","167.85","825.08","288.86","0.00","32.03","330.59","156.76","80.74","33.48","784.18"],"quality":"Less is better","isMain":true,"metricName":"# mismatches per 100 kbp"},{"values":["11.40","0.00","0.00","0.00","5.99","61.12","21.33","0.00","0.00","9.96","2.21","2.07","4.78","74.35"],"quality":"Less is better","isMain":true,"metricName":"# indels per 100 kbp"},{"values":[6,0,0,0,3,26,11,0,0,5,1,1,1,34],"quality":"Less is better","isMain":false,"metricName":" # indels (<= 5 bp)"},{"values":[0,0,0,0,0,8,0,0,0,0,0,0,0,8],"quality":"Less is better","isMain":false,"metricName":" # indels (> 5 bp)"},{"values":[51,108,0,0,0,45,0,0,0,0,0,0,0,1430],"quality":"Less is better","isMain":false,"metricName":"# N's"},{"values":["86.98","335.86","0.00","0.00","0.00","44.16","0.00","0.00","0.00","0.00","0.00","0.00","0.00","681.03"],"quality":"Less is better","isMain":true,"metricName":"# N's per 100 kbp"}]],["Statistics without reference",[{"values":[21,21,25,31,27,31,10,10,15,6,25,18,16,9],"quality":"Equal","isMain":true,"metricName":"# contigs"},{"values":[12,8,18,16,18,22,9,0,11,6,15,11,5,9],"quality":"Equal","isMain":false,"metricName":"# contigs (>= 1000 bp)"},{"values":[2,1,2,1,1,6,3,0,4,4,2,2,1,8],"quality":"Equal","isMain":false,"metricName":"# contigs (>= 5000 bp)"},{"values":[1,0,1,0,0,2,1,0,1,2,0,1,0,4],"quality":"Equal","isMain":false,"metricName":"# contigs (>= 10000 bp)"},{"values":[0,0,0,0,0,0,0,0,0,1,0,0,0,3],"quality":"Equal","isMain":false,"metricName":"# contigs (>= 25000 bp)"},{"values":[0,0,0,0,0,0,0,0,0,0,0,0,0,1],"quality":"Equal","isMain":false,"metricName":"# contigs (>= 50000 bp)"},{"values":[13404,7486,12864,7747,9407,18536,13868,991,16848,28214,6875,12514,5494,92535],"quality":"More is better","isMain":true,"metricName":"Largest contig"},{"values":[58637,32156,54168,45136,52371,101906,54688,6605,55553,52379,46008,48739,20934,209975],"quality":"More is better","isMain":true,"metricName":"Total length"},{"values":[51512,23266,48656,34408,45679,95710,54071,0,52938,52379,38761,43755,13166,209975],"quality":"More is better","isMain":true,"metricName":"Total length (>= 1000 bp)"},{"values":[20911,7486,19204,7747,9407,58873,30960,0,40344,49853,12224,17863,5494,205201],"quality":"More is better","isMain":false,"metricName":"Total length (>= 5000 bp)"},{"values":[13404,0,12864,0,0,29257,13868,0,16848,38266,0,12514,0,175087],"quality":"More is better","isMain":true,"metricName":"Total length (>= 10000 bp)"},{"values":[0,0,0,0,0,0,0,0,0,28214,0,0,0,164346],"quality":"More is better","isMain":false,"metricName":"Total length (>= 25000 bp)"},{"values":[0,0,0,0,0,0,0,0,0,0,0,0,0,92535],"quality":"More is better","isMain":true,"metricName":"Total length (>= 50000 bp)"},{"values":[4491,2781,2378,2005,2244,7210,7433,619,7976,28214,2973,4555,1822,39285],"quality":"More is better","isMain":false,"metricName":"N50"},{"values":[2849,954,1306,1070,1574,2896,4281,542,3409,6238,1187,2609,849,32526],"quality":"More is better","isMain":false,"metricName":"N75"},{"values":[4,4,5,7,7,5,3,4,3,1,5,4,4,2],"quality":"Less is better","isMain":false,"metricName":"L50"},{"values":[9,9,12,16,13,12,6,7,5,3,12,7,8,3],"quality":"Less is better","isMain":false,"metricName":"L75"},{"values":["60.96","60.91","61.11","61.08","61.08","61.46","61.27","59.39","61.03","61.14","61.13","61.02","60.70","61.24"],"quality":"Equal","isMain":false,"metricName":"GC (%)"}]],["Predicted genes",[]],["Similarity statistics",[{"values":[0,0,0,0,0,0,0,0,0,0,0,0,0,0],"quality":"Equal","isMain":false,"metricName":"# similar correct contigs"},{"values":[0,0,0,0,0,0,0,0,0,0,0,0,0,0],"quality":"Equal","isMain":false,"metricName":"# similar misassembled blocks"}]],["Reference statistics",[{"values":[58578,58578,58578,58578,58578,58578,58578,58578,58578,58578,58578,58578,58578,58578],"quality":"Equal","isMain":false,"metricName":"Reference length"},{"values":[1,1,1,1,1,1,1,1,1,1,1,1,1,1],"quality":"Equal","isMain":false,"metricName":"Reference fragments"},{"values":["61.22","61.22","61.22","61.22","61.22","61.22","61.22","61.22","61.22","61.22","61.22","61.22","61.22","61.22"],"quality":"Equal","isMain":false,"metricName":"Reference GC (%)"}]]],"referenceName":"gi\_206600181\_ref\_NC","date":"18 July 2018, Wednesday, 16:52:24","order":[0,1,2,3,4,5,6,7,8,9,10,11,12,13],"assembliesNames":["ABySS\_127","ABySS\_63","CLC","IDBA\_UD","MEGAHIT","MIRA","Ray\_Meta","SOAPdenovo2","SPAdes","SPAdes\_meta","SPAdes\_sc","SPAdes\_sc\_careful","Velvet","Geneious"]},{"assembliesWithNs":null,"minContig":500,"report":[["Genome statistics",[{"values":["99.919","100.000","100.000","100.000","100.000","100.000","100.000","99.879","99.879","99.879","98.482"],"quality":"More is better","isMain":true,"metricName":"Genome fraction (%)"},{"values":["1.000","1.000","1.001","1.004","1.011","1.010","1.004","1.000","1.000","1.000","1.001"],"quality":"Less is better","isMain":true,"metricName":"Duplication ratio"},{"values":[35609,35638,35683,35779,36038,35983,35765,35595,35595,35595,15329],"quality":"More is better","isMain":true,"metricName":"Largest alignment"},{"values":[35609,35638,35683,35779,36038,35983,35765,35595,35595,35595,35141],"quality":"More is better","isMain":true,"metricName":"Total aligned length"},{"values":[35609,35638,35683,35779,36038,35983,35765,35595,35595,35595,11755],"quality":"More is better","isMain":false,"metricName":"NG50"},{"values":[35609,35638,35683,35779,36038,35983,35765,35595,35595,35595,11755],"quality":"More is better","isMain":false,"metricName":"NG75"},{"values":[35609,35638,35683,35779,36038,35983,35765,35595,35595,35595,11755],"quality":"More is better","isMain":false,"metricName":"NA50"},{"values":[35609,35638,35683,35779,36038,35983,35765,35595,35595,35595,11755],"quality":"More is better","isMain":false,"metricName":"NA75"},{"values":[35609,35638,35683,35779,36038,35983,35765,35595,35595,35595,11755],"quality":"More is better","isMain":true,"metricName":"NGA50"},{"values":[35609,35638,35683,35779,36038,35983,35765,35595,35595,35595,11755],"quality":"More is better","isMain":false,"metricName":"NGA75"},{"values":[1,1,1,1,1,1,1,1,1,1,2],"quality":"Less is better","isMain":false,"metricName":"LG50"},{"values":[1,1,1,1,1,1,1,1,1,1,2],"quality":"Less is better","isMain":false,"metricName":"LG75"},{"values":[1,1,1,1,1,1,1,1,1,1,2],"quality":"Less is better","isMain":false,"metricName":"LA50"},{"values":[1,1,1,1,1,1,1,1,1,1,2],"quality":"Less is better","isMain":false,"metricName":"LA75"},{"values":[1,1,1,1,1,1,1,1,1,1,2],"quality":"Less is better","isMain":true,"metricName":"LGA50"},{"values":[1,1,1,1,1,1,1,1,1,1,2],"quality":"Less is better","isMain":false,"metricName":"LGA75"}]],["Misassemblies",[{"values":[0,0,0,0,0,0,0,0,0,0,0],"quality":"Less is better","isMain":true,"metricName":"# misassemblies"},{"values":[0,0,0,0,0,0,0,0,0,0,0],"quality":"Less is better","isMain":false,"metricName":" # relocations"},{"values":[0,0,0,0,0,0,0,0,0,0,0],"quality":"Less is better","isMain":false,"metricName":" # translocations"},{"values":[0,0,0,0,0,0,0,0,0,0,0],"quality":"Less is better","isMain":false,"metricName":" # inversions"},{"values":[0,0,0,0,0,0,0,0,0,0,0],"quality":"Less is better","isMain":false,"metricName":"# misassembled contigs"},{"values":[0,0,0,0,0,0,0,0,0,0,0],"quality":"Less is better","isMain":true,"metricName":"Misassembled contigs length"},{"values":[0,0,0,1,1,3,1,0,0,0,0],"quality":"Less is better","isMain":false,"metricName":"# local misassemblies"},{"values":[0,0,0,0,0,0,0,0,0,0,0],"quality":"Less is better","isMain":false,"metricName":"# unaligned mis. contigs"}]],["Unaligned",[{"values":[0,0,0,0,0,0,0,0,0,0,0],"quality":"Less is better","isMain":false,"metricName":"# fully unaligned contigs"},{"values":[0,0,0,0,0,0,0,0,0,0,0],"quality":"Less is better","isMain":false,"metricName":"Fully unaligned length"},{"values":[0,0,0,0,0,0,0,0,0,0,0],"quality":"Less is better","isMain":false,"metricName":"# partially unaligned contigs"},{"values":[0,0,0,0,0,0,0,0,0,0,0],"quality":"Less is better","isMain":false,"metricName":"Partially unaligned length"}]],["Mismatches",[{"values":[0,0,0,0,0,0,0,0,0,0,0],"quality":"Less is better","isMain":false,"metricName":"# mismatches"},{"values":[0,0,1,0,0,0,0,0,0,0,0],"quality":"Less is better","isMain":false,"metricName":"# indels"},{"values":[0,0,45,0,0,0,0,0,0,0,0],"quality":"Less is better","isMain":false,"metricName":"Indels length"},{"values":["0.00","0.00","0.00","0.00","0.00","0.00","0.00","0.00","0.00","0.00","0.00"],"quality":"Less is better","isMain":true,"metricName":"# mismatches per 100 kbp"},{"values":["0.00","0.00","2.81","0.00","0.00","0.00","0.00","0.00","0.00","0.00","0.00"],"quality":"Less is better","isMain":true,"metricName":"# indels per 100 kbp"},{"values":[0,0,0,0,0,0,0,0,0,0,0],"quality":"Less is better","isMain":false,"metricName":" # indels (<= 5 bp)"},{"values":[0,0,1,0,0,0,0,0,0,0,0],"quality":"Less is better","isMain":false,"metricName":" # indels (> 5 bp)"},{"values":[0,0,0,0,0,0,0,0,0,0,0],"quality":"Less is better","isMain":false,"metricName":"# N's"},{"values":["0.00","0.00","0.00","0.00","0.00","0.00","0.00","0.00","0.00","0.00","0.00"],"quality":"Less is better","isMain":true,"metricName":"# N's per 100 kbp"}]],["Statistics without reference",[{"values":[1,1,1,1,1,1,1,1,1,1,5],"quality":"Equal","isMain":true,"metricName":"# contigs"},{"values":[1,1,1,1,1,1,1,1,1,1,4],"quality":"Equal","isMain":false,"metricName":"# contigs (>= 1000 bp)"},{"values":[1,1,1,1,1,1,1,1,1,1,3],"quality":"Equal","isMain":false,"metricName":"# contigs (>= 5000 bp)"},{"values":[1,1,1,1,1,1,1,1,1,1,2],"quality":"Equal","isMain":false,"metricName":"# contigs (>= 10000 bp)"},{"values":[1,1,1,1,1,1,1,1,1,1,0],"quality":"Equal","isMain":false,"metricName":"# contigs (>= 25000 bp)"},{"values":[0,0,0,0,0,0,0,0,0,0,0],"quality":"Equal","isMain":false,"metricName":"# contigs (>= 50000 bp)"},{"values":[35609,35638,35683,35779,36038,35983,35765,35595,35595,35595,15329],"quality":"More is better","isMain":true,"metricName":"Largest contig"},{"values":[35609,35638,35683,35779,36038,35983,35765,35595,35595,35595,35141],"quality":"More is better","isMain":true,"metricName":"Total length"},{"values":[35609,35638,35683,35779,36038,35983,35765,35595,35595,35595,34567],"quality":"More is better","isMain":true,"metricName":"Total length (>= 1000 bp)"},{"values":[35609,35638,35683,35779,36038,35983,35765,35595,35595,35595,32961],"quality":"More is better","isMain":false,"metricName":"Total length (>= 5000 bp)"},{"values":[35609,35638,35683,35779,36038,35983,35765,35595,35595,35595,27084],"quality":"More is better","isMain":true,"metricName":"Total length (>= 10000 bp)"},{"values":[35609,35638,35683,35779,36038,35983,35765,35595,35595,35595,0],"quality":"More is better","isMain":false,"metricName":"Total length (>= 25000 bp)"},{"values":[0,0,0,0,0,0,0,0,0,0,0],"quality":"More is better","isMain":true,"metricName":"Total length (>= 50000 bp)"},{"values":[35609,35638,35683,35779,36038,35983,35765,35595,35595,35595,11755],"quality":"More is better","isMain":false,"metricName":"N50"},{"values":[35609,35638,35683,35779,36038,35983,35765,35595,35595,35595,11755],"quality":"More is better","isMain":false,"metricName":"N75"},{"values":[1,1,1,1,1,1,1,1,1,1,2],"quality":"Less is better","isMain":false,"metricName":"L50"},{"values":[1,1,1,1,1,1,1,1,1,1,2],"quality":"Less is better","isMain":false,"metricName":"L75"},{"values":["39.27","39.26","39.25","39.24","39.25","39.20","39.28","39.27","39.27","39.27","39.34"],"quality":"Equal","isMain":false,"metricName":"GC (%)"}]],["Predicted genes",[]],["Similarity statistics",[{"values":[2,2,2,0,0,0,0,2,2,2,0],"quality":"Equal","isMain":false,"metricName":"# similar correct contigs"},{"values":[0,0,0,0,0,0,0,0,0,0,0],"quality":"Equal","isMain":false,"metricName":"# similar misassembled blocks"}]],["Reference statistics",[{"values":[35638,35638,35638,35638,35638,35638,35638,35638,35638,35638,35638],"quality":"Equal","isMain":false,"metricName":"Reference length"},{"values":[1,1,1,1,1,1,1,1,1,1,1],"quality":"Equal","isMain":false,"metricName":"Reference fragments"},{"values":["39.26","39.26","39.26","39.26","39.26","39.26","39.26","39.26","39.26","39.26","39.26"],"quality":"Equal","isMain":false,"metricName":"Reference GC (%)"}]]],"referenceName":"gi\_207270774\_ref\_NC","date":"18 July 2018, Wednesday, 16:52:38","order":[0,1,2,3,4,5,6,7,8,9,10],"assembliesNames":["ABySS\_63","CLC","IDBA\_UD","MEGAHIT","MIRA","Ray\_Meta","SPAdes","SPAdes\_meta","SPAdes\_sc","SPAdes\_sc\_careful","Velvet"]},{"assembliesWithNs":null,"minContig":500,"report":[["Genome statistics",[{"values":["1.571","100.000","100.000","100.000","100.000","100.000","100.000","100.000","100.000","100.000","99.102","100.000"],"quality":"More is better","isMain":true,"metricName":"Genome fraction (%)"},{"values":["1.000","1.000","1.003","1.004","3.171","1.002","1.003","1.001","1.001","1.001","1.001","1.071"],"quality":"Less is better","isMain":true,"metricName":"Duplication ratio"},{"values":[609,38753,38852,38894,39592,38815,38880,38808,38808,38808,15558,28769],"quality":"More is better","isMain":true,"metricName":"Largest alignment"},{"values":[609,38753,38852,38894,122874,38815,38880,38808,38808,38808,38435,41510],"quality":"More is better","isMain":true,"metricName":"Total aligned length"},{"values":[null,38753,38852,38894,39592,38815,38880,38808,38808,38808,9932,41510],"quality":"More is better","isMain":false,"metricName":"NG50"},{"values":[null,38753,38852,38894,39592,38815,38880,38808,38808,38808,5832,41510],"quality":"More is better","isMain":false,"metricName":"NG75"},{"values":[609,38753,38852,38894,1729,38815,38880,38808,38808,38808,9932,28769],"quality":"More is better","isMain":false,"metricName":"NA50"},{"values":[609,38753,38852,38894,1351,38815,38880,38808,38808,38808,5832,12741],"quality":"More is better","isMain":false,"metricName":"NA75"},{"values":[null,38753,38852,38894,39592,38815,38880,38808,38808,38808,9932,28769],"quality":"More is better","isMain":true,"metricName":"NGA50"},{"values":[null,38753,38852,38894,39592,38815,38880,38808,38808,38808,5832,12741],"quality":"More is better","isMain":false,"metricName":"NGA75"},{"values":[null,1,1,1,1,1,1,1,1,1,2,1],"quality":"Less is better","isMain":false,"metricName":"LG50"},{"values":[null,1,1,1,1,1,1,1,1,1,3,1],"quality":"Less is better","isMain":false,"metricName":"LG75"},{"values":[1,1,1,1,11,1,1,1,1,1,2,1],"quality":"Less is better","isMain":false,"metricName":"LA50"},{"values":[1,1,1,1,30,1,1,1,1,1,3,2],"quality":"Less is better","isMain":false,"metricName":"LA75"},{"values":[null,1,1,1,1,1,1,1,1,1,2,1],"quality":"Less is better","isMain":true,"metricName":"LGA50"},{"values":[null,1,1,1,1,1,1,1,1,1,3,2],"quality":"Less is better","isMain":false,"metricName":"LGA75"}]],["Misassemblies",[{"values":[0,0,0,0,0,0,0,0,0,0,0,1],"quality":"Less is better","isMain":true,"metricName":"# misassemblies"},{"values":[0,0,0,0,0,0,0,0,0,0,0,1],"quality":"Less is better","isMain":false,"metricName":" # relocations"},{"values":[0,0,0,0,0,0,0,0,0,0,0,0],"quality":"Less is better","isMain":false,"metricName":" # translocations"},{"values":[0,0,0,0,0,0,0,0,0,0,0,0],"quality":"Less is better","isMain":false,"metricName":" # inversions"},{"values":[0,0,0,0,0,0,0,0,0,0,0,1],"quality":"Less is better","isMain":false,"metricName":"# misassembled contigs"},{"values":[0,0,0,0,0,0,0,0,0,0,0,41510],"quality":"Less is better","isMain":true,"metricName":"Misassembled contigs length"},{"values":[0,0,1,1,1,0,1,0,0,0,0,0],"quality":"Less is better","isMain":false,"metricName":"# local misassemblies"},{"values":[0,0,0,0,0,0,0,0,0,0,0,0],"quality":"Less is better","isMain":false,"metricName":"# unaligned mis. contigs"}]],["Unaligned",[{"values":[0,0,0,0,0,0,0,0,0,0,0,0],"quality":"Less is better","isMain":false,"metricName":"# fully unaligned contigs"},{"values":[0,0,0,0,0,0,0,0,0,0,0,0],"quality":"Less is better","isMain":false,"metricName":"Fully unaligned length"},{"values":[0,0,0,0,0,0,0,0,0,0,0,0],"quality":"Less is better","isMain":false,"metricName":"# partially unaligned contigs"},{"values":[0,0,0,0,0,0,0,0,0,0,0,0],"quality":"Less is better","isMain":false,"metricName":"Partially unaligned length"}]],["Mismatches",[{"values":[0,0,0,0,8,0,0,0,0,0,20,0],"quality":"Less is better","isMain":false,"metricName":"# mismatches"},{"values":[0,0,0,0,0,1,0,1,1,1,2,0],"quality":"Less is better","isMain":false,"metricName":"# indels"},{"values":[0,0,0,0,0,62,0,55,55,55,2,0],"quality":"Less is better","isMain":false,"metricName":"Indels length"},{"values":["0.00","0.00","0.00","0.00","20.64","0.00","0.00","0.00","0.00","0.00","52.08","0.00"],"quality":"Less is better","isMain":true,"metricName":"# mismatches per 100 kbp"},{"values":["0.00","0.00","0.00","0.00","0.00","2.58","0.00","2.58","2.58","2.58","5.21","0.00"],"quality":"Less is better","isMain":true,"metricName":"# indels per 100 kbp"},{"values":[0,0,0,0,0,0,0,0,0,0,2,0],"quality":"Less is better","isMain":false,"metricName":" # indels (<= 5 bp)"},{"values":[0,0,0,0,0,1,0,1,1,1,0,0],"quality":"Less is better","isMain":false,"metricName":" # indels (> 5 bp)"},{"values":[0,0,0,0,84,0,0,0,0,0,0,73],"quality":"Less is better","isMain":false,"metricName":"# N's"},{"values":["0.00","0.00","0.00","0.00","68.36","0.00","0.00","0.00","0.00","0.00","0.00","175.86"],"quality":"Less is better","isMain":true,"metricName":"# N's per 100 kbp"}]],["Statistics without reference",[{"values":[1,1,1,1,57,1,1,1,1,1,5,1],"quality":"Equal","isMain":true,"metricName":"# contigs"},{"values":[0,1,1,1,50,1,1,1,1,1,5,1],"quality":"Equal","isMain":false,"metricName":"# contigs (>= 1000 bp)"},{"values":[0,1,1,1,1,1,1,1,1,1,4,1],"quality":"Equal","isMain":false,"metricName":"# contigs (>= 5000 bp)"},{"values":[0,1,1,1,1,1,1,1,1,1,1,1],"quality":"Equal","isMain":false,"metricName":"# contigs (>= 10000 bp)"},{"values":[0,1,1,1,1,1,1,1,1,1,0,1],"quality":"Equal","isMain":false,"metricName":"# contigs (>= 25000 bp)"},{"values":[0,0,0,0,0,0,0,0,0,0,0,0],"quality":"Equal","isMain":false,"metricName":"# contigs (>= 50000 bp)"},{"values":[609,38753,38852,38894,39592,38815,38880,38808,38808,38808,15558,41510],"quality":"More is better","isMain":true,"metricName":"Largest contig"},{"values":[609,38753,38852,38894,122874,38815,38880,38808,38808,38808,38435,41510],"quality":"More is better","isMain":true,"metricName":"Total length"},{"values":[0,38753,38852,38894,116475,38815,38880,38808,38808,38808,38435,41510],"quality":"More is better","isMain":true,"metricName":"Total length (>= 1000 bp)"},{"values":[0,38753,38852,38894,39592,38815,38880,38808,38808,38808,36771,41510],"quality":"More is better","isMain":false,"metricName":"Total length (>= 5000 bp)"},{"values":[0,38753,38852,38894,39592,38815,38880,38808,38808,38808,15558,41510],"quality":"More is better","isMain":true,"metricName":"Total length (>= 10000 bp)"},{"values":[0,38753,38852,38894,39592,38815,38880,38808,38808,38808,0,41510],"quality":"More is better","isMain":false,"metricName":"Total length (>= 25000 bp)"},{"values":[0,0,0,0,0,0,0,0,0,0,0,0],"quality":"More is better","isMain":true,"metricName":"Total length (>= 50000 bp)"},{"values":[609,38753,38852,38894,1729,38815,38880,38808,38808,38808,9932,41510],"quality":"More is better","isMain":false,"metricName":"N50"},{"values":[609,38753,38852,38894,1351,38815,38880,38808,38808,38808,5832,41510],"quality":"More is better","isMain":false,"metricName":"N75"},{"values":[1,1,1,1,11,1,1,1,1,1,2,1],"quality":"Less is better","isMain":false,"metricName":"L50"},{"values":[1,1,1,1,30,1,1,1,1,1,3,1],"quality":"Less is better","isMain":false,"metricName":"L75"},{"values":["30.71","30.42","30.39","30.41","30.32","30.41","30.45","30.39","30.42","30.42","30.49","30.20"],"quality":"Equal","isMain":false,"metricName":"GC (%)"}]],["Predicted genes",[]],["Similarity statistics",[{"values":[0,0,0,0,0,0,0,0,0,0,0,0],"quality":"Equal","isMain":false,"metricName":"# similar correct contigs"},{"values":[0,0,0,0,0,0,0,0,0,0,0,0],"quality":"Equal","isMain":false,"metricName":"# similar misassembled blocks"}]],["Reference statistics",[{"values":[38753,38753,38753,38753,38753,38753,38753,38753,38753,38753,38753,38753],"quality":"Equal","isMain":false,"metricName":"Reference length"},{"values":[1,1,1,1,1,1,1,1,1,1,1,1],"quality":"Equal","isMain":false,"metricName":"Reference fragments"},{"values":["30.42","30.42","30.42","30.42","30.42","30.42","30.42","30.42","30.42","30.42","30.42","30.42"],"quality":"Equal","isMain":false,"metricName":"Reference GC (%)"}]]],"referenceName":"gi\_208429855\_ref\_NC","date":"18 July 2018, Wednesday, 16:52:53","order":[0,1,2,3,4,5,6,7,8,9,10,11],"assembliesNames":["ABySS\_63","CLC","IDBA\_UD","MEGAHIT","MIRA","Ray\_Meta","SPAdes","SPAdes\_meta","SPAdes\_sc","SPAdes\_sc\_careful","Velvet","Geneious"]},{"assembliesWithNs":null,"minContig":500,"report":[["Genome statistics",[{"values":["50.667","32.052","89.076","89.441","90.943","73.183","84.173","92.946","93.480","93.329","93.329","36.082","20.233"],"quality":"More is better","isMain":true,"metricName":"Genome fraction (%)"},{"values":["1.003","1.000","1.009","1.006","1.004","1.019","1.009","1.002","1.000","1.002","1.002","1.001","1.017"],"quality":"Less is better","isMain":true,"metricName":"Duplication ratio"},{"values":[2112,1925,7631,6354,7207,4239,6611,8349,8721,7983,7983,3764,5453],"quality":"More is better","isMain":true,"metricName":"Largest alignment"},{"values":[60634,38252,107289,107012,108953,89027,100711,111104,111585,111561,111561,43087,24147],"quality":"More is better","isMain":true,"metricName":"Total aligned length"},{"values":[515,null,4266,2767,3390,1608,2052,4812,4812,4266,4266,null,null],"quality":"More is better","isMain":false,"metricName":"NG50"},{"values":[null,null,1465,1734,1902,null,1005,2677,2352,2052,2052,null,null],"quality":"More is better","isMain":false,"metricName":"NG75"},{"values":[1248,957,4550,3155,4108,1961,2462,5147,5147,5147,5147,2031,4113],"quality":"More is better","isMain":false,"metricName":"NA50"},{"values":[787,653,2052,2027,2629,1419,1704,2677,2462,2352,2352,1093,3700],"quality":"More is better","isMain":false,"metricName":"NA75"},{"values":[515,null,3284,2767,3351,1560,2047,4266,4266,4266,4266,null,null],"quality":"More is better","isMain":true,"metricName":"NGA50"},{"values":[null,null,1465,1734,1902,null,1005,2462,2220,2052,2052,null,null],"quality":"More is better","isMain":false,"metricName":"NGA75"},{"values":[60,null,11,15,12,25,17,10,10,10,10,null,null],"quality":"Less is better","isMain":false,"metricName":"LG50"},{"values":[null,null,24,28,23,null,36,18,19,21,21,null,null],"quality":"Less is better","isMain":false,"metricName":"LG75"},{"values":[20,14,10,13,10,18,13,9,9,9,9,8,3],"quality":"Less is better","isMain":false,"metricName":"LA50"},{"values":[36,26,19,24,19,31,26,17,18,19,19,15,5],"quality":"Less is better","isMain":false,"metricName":"LA75"},{"values":[60,null,12,15,12,26,17,10,10,10,10,null,null],"quality":"Less is better","isMain":true,"metricName":"LGA50"},{"values":[null,null,24,28,23,null,36,19,20,21,21,null,null],"quality":"Less is better","isMain":false,"metricName":"LGA75"}]],["Misassemblies",[{"values":[0,1,1,0,0,3,0,1,2,1,1,1,1],"quality":"Less is better","isMain":true,"metricName":"# misassemblies"},{"values":[0,1,1,0,0,3,0,1,2,1,1,1,1],"quality":"Less is better","isMain":false,"metricName":" # relocations"},{"values":[0,0,0,0,0,0,0,0,0,0,0,0,0],"quality":"Less is better","isMain":false,"metricName":" # translocations"},{"values":[0,0,0,0,0,0,0,0,0,0,0,0,0],"quality":"Less is better","isMain":false,"metricName":" # inversions"},{"values":[0,1,1,0,0,3,0,1,2,1,1,1,1],"quality":"Less is better","isMain":false,"metricName":"# misassembled contigs"},{"values":[0,1032,6563,0,0,10219,0,4812,9032,729,729,598,6254],"quality":"Less is better","isMain":true,"metricName":"Misassembled contigs length"},{"values":[1,0,0,0,0,2,0,1,0,2,2,1,1],"quality":"Less is better","isMain":false,"metricName":"# local misassemblies"},{"values":[0,0,0,0,0,0,0,0,0,0,0,0,0],"quality":"Less is better","isMain":false,"metricName":"# unaligned mis. contigs"}]],["Unaligned",[{"values":[0,0,0,0,0,0,0,0,0,0,0,0,0],"quality":"Less is better","isMain":false,"metricName":"# fully unaligned contigs"},{"values":[0,0,0,0,0,0,0,0,0,0,0,0,0],"quality":"Less is better","isMain":false,"metricName":"Fully unaligned length"},{"values":[0,0,0,0,0,0,0,0,0,0,0,0,0],"quality":"Less is better","isMain":false,"metricName":"# partially unaligned contigs"},{"values":[0,0,0,0,0,0,0,0,0,0,0,0,0],"quality":"Less is better","isMain":false,"metricName":"Partially unaligned length"}]],["Mismatches",[{"values":[15,17,109,38,37,73,33,98,99,53,46,14,53],"quality":"Less is better","isMain":false,"metricName":"# mismatches"},{"values":[0,0,2,0,0,0,7,3,3,1,1,0,0],"quality":"Less is better","isMain":false,"metricName":"# indels"},{"values":[0,0,2,0,0,0,325,3,3,1,1,0,0],"quality":"Less is better","isMain":false,"metricName":"Indels length"},{"values":["24.81","44.44","102.53","35.60","34.09","83.58","32.85","88.35","88.74","47.58","41.30","32.51","219.49"],"quality":"Less is better","isMain":true,"metricName":"# mismatches per 100 kbp"},{"values":["0.00","0.00","1.88","0.00","0.00","0.00","6.97","2.70","2.69","0.90","0.90","0.00","0.00"],"quality":"Less is better","isMain":true,"metricName":"# indels per 100 kbp"},{"values":[0,0,2,0,0,0,0,3,3,1,1,0,0],"quality":"Less is better","isMain":false,"metricName":" # indels (<= 5 bp)"},{"values":[0,0,0,0,0,0,7,0,0,0,0,0,0],"quality":"Less is better","isMain":false,"metricName":" # indels (> 5 bp)"},{"values":[0,0,0,0,0,70,674,0,0,0,0,0,52],"quality":"Less is better","isMain":false,"metricName":"# N's"},{"values":["0.00","0.00","0.00","0.00","0.00","78.63","665.12","0.00","0.00","0.00","0.00","0.00","211.76"],"quality":"Less is better","isMain":true,"metricName":"# N's per 100 kbp"}]],["Statistics without reference",[{"values":[61,42,44,44,39,52,52,33,33,40,40,29,6],"quality":"Equal","isMain":true,"metricName":"# contigs"},{"values":[24,14,29,36,33,40,36,29,29,32,32,17,6],"quality":"Equal","isMain":false,"metricName":"# contigs (>= 1000 bp)"},{"values":[0,0,6,2,7,0,2,9,9,9,9,0,1],"quality":"Equal","isMain":false,"metricName":"# contigs (>= 5000 bp)"},{"values":[0,0,0,0,0,0,0,0,0,0,0,0,0],"quality":"Equal","isMain":false,"metricName":"# contigs (>= 10000 bp)"},{"values":[0,0,0,0,0,0,0,0,0,0,0,0,0],"quality":"Equal","isMain":false,"metricName":"# contigs (>= 25000 bp)"},{"values":[0,0,0,0,0,0,0,0,0,0,0,0,0],"quality":"Equal","isMain":false,"metricName":"# contigs (>= 50000 bp)"},{"values":[2112,1925,7631,6354,7207,4697,6803,8349,8721,7983,7983,3764,6254],"quality":"More is better","isMain":true,"metricName":"Largest contig"},{"values":[60651,38252,107289,107358,108992,89027,101335,111108,111586,111564,111564,43087,24556],"quality":"More is better","isMain":true,"metricName":"Total length"},{"values":[34902,19899,96322,101417,104656,79759,90044,108513,108531,105904,105904,34559,24556],"quality":"More is better","isMain":true,"metricName":"Total length (>= 1000 bp)"},{"values":[0,0,37184,11684,41593,0,12551,58189,56593,55861,55861,0,6254],"quality":"More is better","isMain":false,"metricName":"Total length (>= 5000 bp)"},{"values":[0,0,0,0,0,0,0,0,0,0,0,0,0],"quality":"More is better","isMain":true,"metricName":"Total length (>= 10000 bp)"},{"values":[0,0,0,0,0,0,0,0,0,0,0,0,0],"quality":"More is better","isMain":false,"metricName":"Total length (>= 25000 bp)"},{"values":[0,0,0,0,0,0,0,0,0,0,0,0,0],"quality":"More is better","isMain":true,"metricName":"Total length (>= 50000 bp)"},{"values":[1248,1012,4550,3218,4108,1967,2462,5147,5147,5147,5147,2031,4156],"quality":"More is better","isMain":false,"metricName":"N50"},{"values":[787,653,2052,2179,2629,1420,1705,3155,3155,2352,2352,1093,3919],"quality":"More is better","isMain":false,"metricName":"N75"},{"values":[20,14,10,13,10,17,13,9,9,9,9,8,3],"quality":"Less is better","isMain":false,"metricName":"L50"},{"values":[36,26,19,23,19,30,26,16,16,19,19,15,4],"quality":"Less is better","isMain":false,"metricName":"L75"},{"values":["49.58","49.03","49.49","49.44","49.64","49.57","49.25","49.62","49.59","49.56","49.56","49.73","50.78"],"quality":"Equal","isMain":false,"metricName":"GC (%)"}]],["Predicted genes",[]],["Similarity statistics",[{"values":[0,0,0,0,0,0,0,0,0,0,0,0,0],"quality":"Equal","isMain":false,"metricName":"# similar correct contigs"},{"values":[0,0,0,0,0,0,0,0,0,0,0,0,0],"quality":"Equal","isMain":false,"metricName":"# similar misassembled blocks"}]],["Reference statistics",[{"values":[119343,119343,119343,119343,119343,119343,119343,119343,119343,119343,119343,119343,119343],"quality":"Equal","isMain":false,"metricName":"Reference length"},{"values":[1,1,1,1,1,1,1,1,1,1,1,1,1],"quality":"Equal","isMain":false,"metricName":"Reference fragments"},{"values":["49.66","49.66","49.66","49.66","49.66","49.66","49.66","49.66","49.66","49.66","49.66","49.66","49.66"],"quality":"Equal","isMain":false,"metricName":"Reference GC (%)"}]]],"referenceName":"gi\_208436764\_ref\_NC","date":"18 July 2018, Wednesday, 16:53:11","order":[0,1,2,3,4,5,6,7,8,9,10,11,12],"assembliesNames":["ABySS\_127","ABySS\_63","CLC","IDBA\_UD","MEGAHIT","MIRA","SOAPdenovo2","SPAdes","SPAdes\_meta","SPAdes\_sc","SPAdes\_sc\_careful","Velvet","Geneious"]},{"assembliesWithNs":null,"minContig":500,"report":[["Genome statistics",[{"values":["11.158","6.954","51.007","52.070","63.230","20.429","44.321","67.121","68.441","65.256","65.119","2.572","2.685"],"quality":"More is better","isMain":true,"metricName":"Genome fraction (%)"},{"values":["1.001","1.000","1.017","1.031","1.004","1.027","1.003","1.014","1.006","1.016","1.016","1.000","1.000"],"quality":"Less is better","isMain":true,"metricName":"Duplication ratio"},{"values":[1369,1566,2733,2967,3697,2005,2700,3697,3398,3398,3398,1483,1892],"quality":"More is better","isMain":true,"metricName":"Largest alignment"},{"values":[17328,10787,79704,81045,98123,32548,68896,104300,106355,101375,101163,3989,4165],"quality":"More is better","isMain":true,"metricName":"Total aligned length"},{"values":[null,null,579,627,765,null,null,1057,917,1005,1005,null,null],"quality":"More is better","isMain":false,"metricName":"NG50"},{"values":[726,721,1059,1083,1177,1113,921,1339,1348,1426,1364,763,null],"quality":"More is better","isMain":false,"metricName":"NA50"},{"values":[624,544,693,738,794,742,699,857,879,917,898,549,null],"quality":"More is better","isMain":false,"metricName":"NA75"},{"values":[null,null,506,570,757,null,null,905,905,917,917,null,null],"quality":"More is better","isMain":true,"metricName":"NGA50"},{"values":[null,null,70,68,57,null,null,44,48,45,44,null,null],"quality":"Less is better","isMain":false,"metricName":"LG50"},{"values":[10,6,26,24,28,11,25,28,28,26,27,2,null],"quality":"Less is better","isMain":false,"metricName":"LA50"},{"values":[17,10,51,49,53,21,46,53,52,48,50,4,null],"quality":"Less is better","isMain":false,"metricName":"LA75"},{"values":[null,null,76,72,58,null,null,48,49,48,48,null,null],"quality":"Less is better","isMain":true,"metricName":"LGA50"}]],["Misassemblies",[{"values":[0,0,0,0,0,0,0,0,0,0,0,0,0],"quality":"Less is better","isMain":true,"metricName":"# misassemblies"},{"values":[0,0,0,0,0,0,0,0,0,0,0,0,0],"quality":"Less is better","isMain":false,"metricName":" # relocations"},{"values":[0,0,0,0,0,0,0,0,0,0,0,0,0],"quality":"Less is better","isMain":false,"metricName":" # translocations"},{"values":[0,0,0,0,0,0,0,0,0,0,0,0,0],"quality":"Less is better","isMain":false,"metricName":" # inversions"},{"values":[0,0,0,0,0,0,0,0,0,0,0,0,0],"quality":"Less is better","isMain":false,"metricName":"# misassembled contigs"},{"values":[0,0,0,0,0,0,0,0,0,0,0,0,0],"quality":"Less is better","isMain":true,"metricName":"Misassembled contigs length"},{"values":[0,0,0,0,0,0,0,1,0,0,0,0,0],"quality":"Less is better","isMain":false,"metricName":"# local misassemblies"},{"values":[0,0,0,0,0,0,0,0,0,0,0,0,0],"quality":"Less is better","isMain":false,"metricName":"# unaligned mis. contigs"}]],["Unaligned",[{"values":[0,0,0,0,0,0,0,0,0,0,0,0,0],"quality":"Less is better","isMain":false,"metricName":"# fully unaligned contigs"},{"values":[0,0,0,0,0,0,0,0,0,0,0,0,0],"quality":"Less is better","isMain":false,"metricName":"Fully unaligned length"},{"values":[0,0,3,0,0,0,0,5,0,2,3,0,1],"quality":"Less is better","isMain":false,"metricName":"# partially unaligned contigs"},{"values":[0,0,2738,0,0,0,0,4006,0,1074,2687,0,5891],"quality":"Less is better","isMain":false,"metricName":"Partially unaligned length"}]],["Mismatches",[{"values":[2,1,216,251,113,9,69,274,147,264,255,2,1],"quality":"Less is better","isMain":false,"metricName":"# mismatches"},{"values":[0,0,4,8,2,0,1,7,3,7,7,0,0],"quality":"Less is better","isMain":false,"metricName":"# indels"},{"values":[0,0,4,12,6,0,53,12,36,40,40,0,0],"quality":"Less is better","isMain":false,"metricName":"Indels length"},{"values":["11.56","9.27","272.99","310.75","115.21","28.40","100.36","263.16","138.46","260.80","252.44","50.14","24.01"],"quality":"Less is better","isMain":true,"metricName":"# mismatches per 100 kbp"},{"values":["0.00","0.00","5.06","9.90","2.04","0.00","1.45","6.72","2.83","6.92","6.93","0.00","0.00"],"quality":"Less is better","isMain":true,"metricName":"# indels per 100 kbp"},{"values":[0,0,4,8,2,0,0,7,2,6,6,0,0],"quality":"Less is better","isMain":false,"metricName":" # indels (<= 5 bp)"},{"values":[0,0,0,0,0,0,1,0,1,1,1,0,0],"quality":"Less is better","isMain":false,"metricName":" # indels (> 5 bp)"},{"values":[0,0,0,0,0,49,59,0,0,0,0,0,15],"quality":"Less is better","isMain":false,"metricName":"# N's"},{"values":["0.00","0.00","0.00","0.00","0.00","150.55","85.56","0.00","0.00","0.00","0.00","0.00","149.16"],"quality":"Less is better","isMain":true,"metricName":"# N's per 100 kbp"}]],["Statistics without reference",[{"values":[24,15,80,78,90,33,75,84,88,80,80,5,3],"quality":"Equal","isMain":true,"metricName":"# contigs"},{"values":[1,2,32,28,39,12,20,47,45,45,45,1,3],"quality":"Equal","isMain":false,"metricName":"# contigs (>= 1000 bp)"},{"values":[0,0,0,0,0,0,0,0,0,0,0,0,1],"quality":"Equal","isMain":false,"metricName":"# contigs (>= 5000 bp)"},{"values":[0,0,0,0,0,0,0,0,0,0,0,0,0],"quality":"Equal","isMain":false,"metricName":"# contigs (>= 10000 bp)"},{"values":[0,0,0,0,0,0,0,0,0,0,0,0,0],"quality":"Equal","isMain":false,"metricName":"# contigs (>= 25000 bp)"},{"values":[0,0,0,0,0,0,0,0,0,0,0,0,0],"quality":"Equal","isMain":false,"metricName":"# contigs (>= 50000 bp)"},{"values":[1369,1566,2733,3006,3697,2005,2700,3697,3753,3753,3753,1483,6506],"quality":"More is better","isMain":true,"metricName":"Largest contig"},{"values":[17328,10787,83232,83277,98486,32548,68961,109599,106843,103915,105316,3989,10056],"quality":"More is better","isMain":true,"metricName":"Total length"},{"values":[1369,2586,49996,46997,62389,17576,30008,81611,74942,77701,79102,1483,10056],"quality":"More is better","isMain":true,"metricName":"Total length (>= 1000 bp)"},{"values":[0,0,0,0,0,0,0,0,0,0,0,0,6506],"quality":"More is better","isMain":false,"metricName":"Total length (>= 5000 bp)"},{"values":[0,0,0,0,0,0,0,0,0,0,0,0,0],"quality":"More is better","isMain":true,"metricName":"Total length (>= 10000 bp)"},{"values":[0,0,0,0,0,0,0,0,0,0,0,0,0],"quality":"More is better","isMain":false,"metricName":"Total length (>= 25000 bp)"},{"values":[0,0,0,0,0,0,0,0,0,0,0,0,0],"quality":"More is better","isMain":true,"metricName":"Total length (>= 50000 bp)"},{"values":[726,721,1157,1144,1177,1113,921,1533,1364,1526,1533,763,6506],"quality":"More is better","isMain":false,"metricName":"N50"},{"values":[624,544,726,772,817,742,699,951,896,951,1005,549,1892],"quality":"More is better","isMain":false,"metricName":"N75"},{"values":[10,6,25,24,28,11,25,26,27,25,24,2,1],"quality":"Less is better","isMain":false,"metricName":"L50"},{"values":[17,10,47,47,53,21,46,48,51,46,45,4,2],"quality":"Less is better","isMain":false,"metricName":"L75"},{"values":["46.34","46.68","47.92","48.14","48.72","47.49","47.75","48.03","48.53","48.38","48.42","48.23","50.86"],"quality":"Equal","isMain":false,"metricName":"GC (%)"}]],["Predicted genes",[]],["Similarity statistics",[{"values":[0,0,0,0,0,0,0,0,0,0,0,0,0],"quality":"Equal","isMain":false,"metricName":"# similar correct contigs"},{"values":[0,0,0,0,0,0,0,0,0,0,0,0,0],"quality":"Equal","isMain":false,"metricName":"# similar misassembled blocks"}]],["Reference statistics",[{"values":[155122,155122,155122,155122,155122,155122,155122,155122,155122,155122,155122,155122,155122],"quality":"Equal","isMain":false,"metricName":"Reference length"},{"values":[1,1,1,1,1,1,1,1,1,1,1,1,1],"quality":"Equal","isMain":false,"metricName":"Reference fragments"},{"values":["48.57","48.57","48.57","48.57","48.57","48.57","48.57","48.57","48.57","48.57","48.57","48.57","48.57"],"quality":"Equal","isMain":false,"metricName":"Reference GC (%)"}]]],"referenceName":"gi\_209170884\_ref\_NC","date":"18 July 2018, Wednesday, 16:53:29","order":[0,1,2,3,4,5,6,7,8,9,10,11,12],"assembliesNames":["ABySS\_127","ABySS\_63","CLC","IDBA\_UD","MEGAHIT","MIRA","SOAPdenovo2","SPAdes","SPAdes\_meta","SPAdes\_sc","SPAdes\_sc\_careful","Velvet","Geneious"]},{"assembliesWithNs":null,"minContig":500,"report":[["Genome statistics",[{"values":["25.998","13.259","51.886","23.979","37.909","95.245","24.988","53.308","13.641","31.986","31.551","17.464","100.000"],"quality":"More is better","isMain":true,"metricName":"Genome fraction (%)"},{"values":["1.002","1.000","1.028","1.008","1.042","3.682","1.040","1.038","1.017","1.044","1.015","1.007","2.411"],"quality":"Less is better","isMain":true,"metricName":"Duplication ratio"},{"values":[3735,3925,3813,2460,4870,37622,4103,3274,2565,2460,2416,2050,37621],"quality":"More is better","isMain":true,"metricName":"Largest alignment"},{"values":[12337,6278,25242,11448,18697,165972,12304,26196,6568,15363,15156,8267,113841],"quality":"More is better","isMain":true,"metricName":"Total aligned length"},{"values":[null,null,539,null,null,73503,null,600,null,null,null,null,92482],"quality":"More is better","isMain":false,"metricName":"NG50"},{"values":[null,null,null,null,null,73503,null,null,null,null,null,null,92482],"quality":"More is better","isMain":false,"metricName":"NG75"},{"values":[1642,3925,1312,1483,1107,10973,2452,1426,1293,1474,1265,1162,7904],"quality":"More is better","isMain":false,"metricName":"NA50"},{"values":[578,2353,827,727,773,1537,1146,903,1114,1216,841,896,null],"quality":"More is better","isMain":false,"metricName":"NA75"},{"values":[null,null,539,null,null,37622,null,600,null,null,null,null,37621],"quality":"More is better","isMain":true,"metricName":"NGA50"},{"values":[null,null,null,null,null,37622,null,null,null,null,null,null,37621],"quality":"More is better","isMain":false,"metricName":"NGA75"},{"values":[null,null,18,null,null,1,null,17,null,null,null,null,1],"quality":"Less is better","isMain":false,"metricName":"LG50"},{"values":[null,null,null,null,null,1,null,null,null,null,null,null,1],"quality":"Less is better","isMain":false,"metricName":"LG75"},{"values":[3,1,6,3,4,5,2,6,2,4,5,3,5],"quality":"Less is better","isMain":false,"metricName":"LA50"},{"values":[6,2,11,6,9,20,4,12,3,7,9,5,null],"quality":"Less is better","isMain":false,"metricName":"LA75"},{"values":[null,null,18,null,null,1,null,17,null,null,null,null,1],"quality":"Less is better","isMain":true,"metricName":"LGA50"},{"values":[null,null,null,null,null,1,null,null,null,null,null,null,1],"quality":"Less is better","isMain":false,"metricName":"LGA75"}]],["Misassemblies",[{"values":[0,0,0,0,0,3,0,0,0,0,0,0,4],"quality":"Less is better","isMain":true,"metricName":"# misassemblies"},{"values":[0,0,0,0,0,3,0,0,0,0,0,0,4],"quality":"Less is better","isMain":false,"metricName":" # relocations"},{"values":[0,0,0,0,0,0,0,0,0,0,0,0,0],"quality":"Less is better","isMain":false,"metricName":" # translocations"},{"values":[0,0,0,0,0,0,0,0,0,0,0,0,0],"quality":"Less is better","isMain":false,"metricName":" # inversions"},{"values":[0,0,0,0,0,1,0,0,0,0,0,0,2],"quality":"Less is better","isMain":false,"metricName":"# misassembled contigs"},{"values":[0,0,0,0,0,73503,0,0,0,0,0,0,85746],"quality":"Less is better","isMain":true,"metricName":"Misassembled contigs length"},{"values":[0,0,0,0,0,3,0,0,0,0,0,0,3],"quality":"Less is better","isMain":false,"metricName":"# local misassemblies"},{"values":[0,0,0,0,0,0,0,0,0,0,0,0,1],"quality":"Less is better","isMain":false,"metricName":"# unaligned mis. contigs"}]],["Unaligned",[{"values":[0,0,0,0,0,0,0,0,0,0,0,0,0],"quality":"Less is better","isMain":false,"metricName":"# fully unaligned contigs"},{"values":[0,0,0,0,0,0,0,0,0,0,0,0,0],"quality":"Less is better","isMain":false,"metricName":"Fully unaligned length"},{"values":[0,0,0,0,0,3,0,0,0,0,1,0,4],"quality":"Less is better","isMain":false,"metricName":"# partially unaligned contigs"},{"values":[0,0,0,0,0,22255,0,0,0,0,538,0,83507],"quality":"Less is better","isMain":false,"metricName":"Partially unaligned length"}]],["Mismatches",[{"values":[0,0,29,0,22,426,4,28,4,27,46,10,400],"quality":"Less is better","isMain":false,"metricName":"# mismatches"},{"values":[0,1,2,0,0,6,0,0,0,0,1,1,11],"quality":"Less is better","isMain":false,"metricName":"# indels"},{"values":[0,1,2,0,0,6,0,0,0,0,1,1,11],"quality":"Less is better","isMain":false,"metricName":"Indels length"},{"values":["0.00","0.00","118.06","0.00","122.58","944.76","33.81","110.95","61.94","178.30","307.96","120.95","844.92"],"quality":"Less is better","isMain":true,"metricName":"# mismatches per 100 kbp"},{"values":["0.00","15.93","8.14","0.00","0.00","13.31","0.00","0.00","0.00","0.00","6.69","12.09","23.24"],"quality":"Less is better","isMain":true,"metricName":"# indels per 100 kbp"},{"values":[0,1,2,0,0,6,0,0,0,0,1,1,11],"quality":"Less is better","isMain":false,"metricName":" # indels (<= 5 bp)"},{"values":[0,0,0,0,0,0,0,0,0,0,0,0,0],"quality":"Less is better","isMain":false,"metricName":" # indels (> 5 bp)"},{"values":[0,7,0,0,0,68,0,0,0,0,0,0,1041],"quality":"Less is better","isMain":false,"metricName":"# N's"},{"values":["0.00","111.50","0.00","0.00","0.00","36.11","0.00","0.00","0.00","0.00","0.00","0.00","526.66"],"quality":"Less is better","isMain":true,"metricName":"# N's per 100 kbp"}]],["Statistics without reference",[{"values":[11,2,21,11,16,38,7,21,5,10,14,8,5],"quality":"Equal","isMain":true,"metricName":"# contigs"},{"values":[3,2,9,4,5,32,5,9,3,9,7,3,5],"quality":"Equal","isMain":false,"metricName":"# contigs (>= 1000 bp)"},{"values":[0,0,0,0,0,3,0,0,0,0,0,0,4],"quality":"Equal","isMain":false,"metricName":"# contigs (>= 5000 bp)"},{"values":[0,0,0,0,0,3,0,0,0,0,0,0,4],"quality":"Equal","isMain":false,"metricName":"# contigs (>= 10000 bp)"},{"values":[0,0,0,0,0,2,0,0,0,0,0,0,2],"quality":"Equal","isMain":false,"metricName":"# contigs (>= 25000 bp)"},{"values":[0,0,0,0,0,1,0,0,0,0,0,0,2],"quality":"Equal","isMain":false,"metricName":"# contigs (>= 50000 bp)"},{"values":[3735,3925,3813,2460,4870,73503,4103,3274,2565,2460,2416,2050,92482],"quality":"More is better","isMain":true,"metricName":"Largest contig"},{"values":[12337,6278,25246,11448,18700,188299,12304,26196,6568,15814,15695,8330,197661],"quality":"More is better","isMain":true,"metricName":"Total length"},{"values":[7793,6278,17268,7261,10852,183588,10775,17347,4972,15016,10851,4433,197661],"quality":"More is better","isMain":true,"metricName":"Total length (>= 1000 bp)"},{"values":[0,0,0,0,0,132861,0,0,0,0,0,0,195450],"quality":"More is better","isMain":false,"metricName":"Total length (>= 5000 bp)"},{"values":[0,0,0,0,0,132861,0,0,0,0,0,0,195450],"quality":"More is better","isMain":true,"metricName":"Total length (>= 10000 bp)"},{"values":[0,0,0,0,0,119942,0,0,0,0,0,0,165986],"quality":"More is better","isMain":false,"metricName":"Total length (>= 25000 bp)"},{"values":[0,0,0,0,0,73503,0,0,0,0,0,0,165986],"quality":"More is better","isMain":true,"metricName":"Total length (>= 50000 bp)"},{"values":[1642,3925,1312,1483,1107,46439,2452,1426,1293,1799,1265,1162,73504],"quality":"More is better","isMain":false,"metricName":"N50"},{"values":[578,2353,827,727,773,2712,1146,903,1114,1216,842,959,73504],"quality":"More is better","isMain":false,"metricName":"N75"},{"values":[3,1,6,3,4,2,2,6,2,4,5,3,2],"quality":"Less is better","isMain":false,"metricName":"L50"},{"values":[6,2,11,6,9,6,4,12,3,7,9,5,2],"quality":"Less is better","isMain":false,"metricName":"L75"},{"values":["32.35","30.12","33.55","31.06","33.08","33.38","31.28","32.68","30.15","32.37","32.14","30.41","33.48"],"quality":"Equal","isMain":false,"metricName":"GC (%)"}]],["Predicted genes",[]],["Similarity statistics",[{"values":[0,0,0,0,0,0,0,0,0,0,0,0,0],"quality":"Equal","isMain":false,"metricName":"# similar correct contigs"},{"values":[0,0,0,0,0,0,0,0,0,0,0,0,0],"quality":"Equal","isMain":false,"metricName":"# similar misassembled blocks"}]],["Reference statistics",[{"values":[47342,47342,47342,47342,47342,47342,47342,47342,47342,47342,47342,47342,47342],"quality":"Equal","isMain":false,"metricName":"Reference length"},{"values":[1,1,1,1,1,1,1,1,1,1,1,1,1],"quality":"Equal","isMain":false,"metricName":"Reference fragments"},{"values":["33.04","33.04","33.04","33.04","33.04","33.04","33.04","33.04","33.04","33.04","33.04","33.04","33.04"],"quality":"Equal","isMain":false,"metricName":"Reference GC (%)"}]]],"referenceName":"gi\_209363552\_ref\_NC","date":"18 July 2018, Wednesday, 16:53:46","order":[0,1,2,3,4,5,6,7,8,9,10,11,12],"assembliesNames":["ABySS\_127","ABySS\_63","CLC","IDBA\_UD","MEGAHIT","MIRA","Ray\_Meta","SPAdes","SPAdes\_meta","SPAdes\_sc","SPAdes\_sc\_careful","Velvet","Geneious"]},{"assembliesWithNs":null,"minContig":500,"report":[["Genome statistics",[{"values":["20.362","6.365","15.325","28.867","44.067","65.631","15.280","15.549","13.857","18.723","15.846","8.919","80.563"],"quality":"More is better","isMain":true,"metricName":"Genome fraction (%)"},{"values":["1.000","1.000","1.000","1.008","1.049","1.322","1.049","1.000","1.000","1.007","1.004","1.000","1.311"],"quality":"Less is better","isMain":true,"metricName":"Duplication ratio"},{"values":[2335,1278,4619,2906,8746,22305,2237,2120,2149,2120,2120,2326,13493],"quality":"More is better","isMain":true,"metricName":"Largest alignment"},{"values":[11176,3494,8413,15962,25365,47619,8499,8536,7607,10349,8715,4896,58264],"quality":"More is better","isMain":true,"metricName":"Total aligned length"},{"values":[null,null,null,null,null,28556,null,null,null,null,null,null,179797],"quality":"More is better","isMain":false,"metricName":"NG50"},{"values":[null,null,null,null,null,1724,null,null,null,null,null,null,179797],"quality":"More is better","isMain":false,"metricName":"NG75"},{"values":[777,791,4619,1271,2104,2220,1477,1777,1719,951,951,1456,null],"quality":"More is better","isMain":false,"metricName":"NA50"},{"values":[628,714,1903,668,898,1153,1422,748,585,776,772,1456,null],"quality":"More is better","isMain":false,"metricName":"NA75"},{"values":[null,null,null,null,null,2220,null,null,null,null,null,null,5121],"quality":"More is better","isMain":true,"metricName":"NGA50"},{"values":[null,null,null,null,null,1153,null,null,null,null,null,null,3979],"quality":"More is better","isMain":false,"metricName":"NGA75"},{"values":[null,null,null,null,null,1,null,null,null,null,null,null,1],"quality":"Less is better","isMain":false,"metricName":"LG50"},{"values":[null,null,null,null,null,7,null,null,null,null,null,null,1],"quality":"Less is better","isMain":false,"metricName":"LG75"},{"values":[5,2,1,5,3,3,3,3,3,4,3,2,null],"quality":"Less is better","isMain":false,"metricName":"LA50"},{"values":[9,3,2,11,8,11,4,4,5,7,6,2,null],"quality":"Less is better","isMain":false,"metricName":"LA75"},{"values":[null,null,null,null,null,3,null,null,null,null,null,null,3],"quality":"Less is better","isMain":true,"metricName":"LGA50"},{"values":[null,null,null,null,null,11,null,null,null,null,null,null,6],"quality":"Less is better","isMain":false,"metricName":"LGA75"}]],["Misassemblies",[{"values":[0,0,0,0,0,0,0,0,0,0,0,0,1],"quality":"Less is better","isMain":true,"metricName":"# misassemblies"},{"values":[0,0,0,0,0,0,0,0,0,0,0,0,1],"quality":"Less is better","isMain":false,"metricName":" # relocations"},{"values":[0,0,0,0,0,0,0,0,0,0,0,0,0],"quality":"Less is better","isMain":false,"metricName":" # translocations"},{"values":[0,0,0,0,0,0,0,0,0,0,0,0,0],"quality":"Less is better","isMain":false,"metricName":" # inversions"},{"values":[0,0,0,0,0,0,0,0,0,0,0,0,1],"quality":"Less is better","isMain":false,"metricName":"# misassembled contigs"},{"values":[0,0,0,0,0,0,0,0,0,0,0,0,16912],"quality":"Less is better","isMain":true,"metricName":"Misassembled contigs length"},{"values":[0,0,0,0,0,3,0,0,0,0,0,0,1],"quality":"Less is better","isMain":false,"metricName":"# local misassemblies"},{"values":[0,0,0,0,0,0,0,0,0,0,0,0,3],"quality":"Less is better","isMain":false,"metricName":"# unaligned mis. contigs"}]],["Unaligned",[{"values":[0,0,0,0,0,0,0,0,0,0,0,0,0],"quality":"Less is better","isMain":false,"metricName":"# fully unaligned contigs"},{"values":[0,0,0,0,0,0,0,0,0,0,0,0,0],"quality":"Less is better","isMain":false,"metricName":"Fully unaligned length"},{"values":[0,0,0,1,0,1,0,0,1,0,0,0,4],"quality":"Less is better","isMain":false,"metricName":"# partially unaligned contigs"},{"values":[0,0,0,3064,0,6251,0,0,2375,0,0,0,308370],"quality":"Less is better","isMain":false,"metricName":"Partially unaligned length"}]],["Mismatches",[{"values":[6,1,26,18,84,448,16,19,70,45,35,1,406],"quality":"Less is better","isMain":false,"metricName":"# mismatches"},{"values":[2,0,0,2,7,26,4,0,0,0,0,0,30],"quality":"Less is better","isMain":false,"metricName":"# indels"},{"values":[2,0,0,2,9,37,206,0,0,0,0,0,40],"quality":"Less is better","isMain":false,"metricName":"Indels length"},{"values":["53.68","28.62","309.05","113.59","347.24","1243.44","190.75","222.59","920.21","437.83","402.35","20.42","918.01"],"quality":"Less is better","isMain":true,"metricName":"# mismatches per 100 kbp"},{"values":["17.89","0.00","0.00","12.62","28.94","72.16","47.69","0.00","0.00","0.00","0.00","0.00","67.83"],"quality":"Less is better","isMain":true,"metricName":"# indels per 100 kbp"},{"values":[2,0,0,2,7,26,0,0,0,0,0,0,29],"quality":"Less is better","isMain":false,"metricName":" # indels (<= 5 bp)"},{"values":[0,0,0,0,0,0,4,0,0,0,0,0,1],"quality":"Less is better","isMain":false,"metricName":" # indels (> 5 bp)"},{"values":[0,0,1,0,0,36,408,0,0,0,0,0,1291],"quality":"Less is better","isMain":false,"metricName":"# N's"},{"values":["0.00","0.00","11.89","0.00","0.00","66.83","4638.47","0.00","0.00","0.00","0.00","0.00","352.38"],"quality":"Less is better","isMain":true,"metricName":"# N's per 100 kbp"}]],["Statistics without reference",[{"values":[13,4,5,14,17,18,7,7,5,11,9,4,5],"quality":"Equal","isMain":true,"metricName":"# contigs"},{"values":[2,1,2,6,7,13,4,3,4,3,2,2,5],"quality":"Equal","isMain":false,"metricName":"# contigs (>= 1000 bp)"},{"values":[0,0,0,0,1,1,0,0,0,0,0,0,4],"quality":"Equal","isMain":false,"metricName":"# contigs (>= 5000 bp)"},{"values":[0,0,0,0,0,1,0,0,0,0,0,0,4],"quality":"Equal","isMain":false,"metricName":"# contigs (>= 10000 bp)"},{"values":[0,0,0,0,0,1,0,0,0,0,0,0,3],"quality":"Equal","isMain":false,"metricName":"# contigs (>= 25000 bp)"},{"values":[0,0,0,0,0,0,0,0,0,0,0,0,3],"quality":"Equal","isMain":false,"metricName":"# contigs (>= 50000 bp)"},{"values":[2335,1278,4619,3351,8746,28556,2237,2120,4524,2120,2120,2326,179797],"quality":"More is better","isMain":true,"metricName":"Largest contig"},{"values":[11176,3494,8413,19032,25365,53872,8796,8536,9982,10349,8735,4896,366366],"quality":"More is better","isMain":true,"metricName":"Total length"},{"values":[3698,1278,6522,13272,18616,49868,6919,5981,9397,4779,3701,3782,366366],"quality":"More is better","isMain":true,"metricName":"Total length (>= 1000 bp)"},{"values":[0,0,0,0,8746,28556,0,0,0,0,0,0,362681],"quality":"More is better","isMain":false,"metricName":"Total length (>= 5000 bp)"},{"values":[0,0,0,0,0,28556,0,0,0,0,0,0,362681],"quality":"More is better","isMain":true,"metricName":"Total length (>= 10000 bp)"},{"values":[0,0,0,0,0,28556,0,0,0,0,0,0,345769],"quality":"More is better","isMain":false,"metricName":"Total length (>= 25000 bp)"},{"values":[0,0,0,0,0,0,0,0,0,0,0,0,345769],"quality":"More is better","isMain":true,"metricName":"Total length (>= 50000 bp)"},{"values":[777,791,4619,1760,2104,28556,1477,1777,2120,951,951,1456,107051],"quality":"More is better","isMain":false,"metricName":"N50"},{"values":[628,714,1903,815,898,1724,1422,748,1719,776,776,1456,107051],"quality":"More is better","isMain":false,"metricName":"N75"},{"values":[5,2,1,4,3,1,3,3,2,4,3,2,2],"quality":"Less is better","isMain":false,"metricName":"L50"},{"values":[9,3,2,8,8,7,4,4,3,7,6,2,2],"quality":"Less is better","isMain":false,"metricName":"L75"},{"values":["49.54","46.88","47.54","49.06","50.97","53.25","45.99","48.24","49.82","50.64","49.52","46.63","50.06"],"quality":"Equal","isMain":false,"metricName":"GC (%)"}]],["Predicted genes",[]],["Similarity statistics",[{"values":[0,0,0,0,0,0,0,0,0,0,0,0,0],"quality":"Equal","isMain":false,"metricName":"# similar correct contigs"},{"values":[0,0,0,0,0,0,0,0,0,0,0,0,0],"quality":"Equal","isMain":false,"metricName":"# similar misassembled blocks"}]],["Reference statistics",[{"values":[54896,54896,54896,54896,54896,54896,54896,54896,54896,54896,54896,54896,54896],"quality":"Equal","isMain":false,"metricName":"Reference length"},{"values":[1,1,1,1,1,1,1,1,1,1,1,1,1],"quality":"Equal","isMain":false,"metricName":"Reference fragments"},{"values":["51.12","51.12","51.12","51.12","51.12","51.12","51.12","51.12","51.12","51.12","51.12","51.12","51.12"],"quality":"Equal","isMain":false,"metricName":"Reference GC (%)"}]]],"referenceName":"gi\_209427726\_ref\_NC","date":"18 July 2018, Wednesday, 16:54:03","order":[0,1,2,3,4,5,6,7,8,9,10,11,12],"assembliesNames":["ABySS\_127","ABySS\_63","CLC","IDBA\_UD","MEGAHIT","MIRA","SOAPdenovo2","SPAdes","SPAdes\_meta","SPAdes\_sc","SPAdes\_sc\_careful","Velvet","Geneious"]},{"assembliesWithNs":null,"minContig":500,"report":[["Genome statistics",[{"values":["79.600","63.945","90.014","77.626","70.526","99.902","81.742","5.975","92.135","83.013","83.626","84.606","41.605","90.049"],"quality":"More is better","isMain":true,"metricName":"Genome fraction (%)"},{"values":["1.007","1.029","1.011","1.024","1.030","1.287","1.029","1.034","1.028","1.002","1.018","1.025","1.000","1.064"],"quality":"Less is better","isMain":true,"metricName":"Duplication ratio"},{"values":[8534,7801,8560,7080,7622,26620,9930,918,20619,34662,10418,8874,4338,17107],"quality":"More is better","isMain":true,"metricName":"Largest alignment"},{"values":[49821,40893,56187,49404,45023,79410,52297,3736,58849,51643,52628,53872,25856,59398],"quality":"More is better","isMain":true,"metricName":"Total aligned length"},{"values":[3696,2603,2537,1663,1887,12615,3873,null,57853,34662,3901,1929,null,107051],"quality":"More is better","isMain":false,"metricName":"NG50"},{"values":[756,null,1664,615,null,7889,1257,null,57853,2550,1246,849,null,107051],"quality":"More is better","isMain":false,"metricName":"NG75"},{"values":[3763,3366,3338,2063,2767,7889,5686,622,1377,34662,7805,2748,1565,null],"quality":"More is better","isMain":false,"metricName":"NA50"},{"values":[2230,869,1987,1181,1271,2751,2104,550,null,10738,1930,1116,1048,null],"quality":"More is better","isMain":false,"metricName":"NA75"},{"values":[3696,1639,2516,1663,1887,12615,3873,null,5738,34662,3901,1929,null,15321],"quality":"More is better","isMain":true,"metricName":"NGA50"},{"values":[756,null,1664,615,null,7889,1257,null,2311,2550,1246,849,null,7263],"quality":"More is better","isMain":false,"metricName":"NGA75"},{"values":[6,6,7,10,9,2,5,null,1,1,4,7,null,1],"quality":"Less is better","isMain":false,"metricName":"LG50"},{"values":[16,null,14,26,null,3,12,null,1,3,12,20,null,1],"quality":"Less is better","isMain":false,"metricName":"LG75"},{"values":[5,5,6,6,5,3,4,3,10,1,3,5,5,null],"quality":"Less is better","isMain":false,"metricName":"LA50"},{"values":[8,10,12,14,11,7,8,5,null,2,8,13,10,null],"quality":"Less is better","isMain":false,"metricName":"LA75"},{"values":[6,8,7,10,9,2,5,null,3,1,4,7,null,2],"quality":"Less is better","isMain":true,"metricName":"LGA50"},{"values":[16,null,14,26,null,3,12,null,7,3,12,20,null,4],"quality":"Less is better","isMain":false,"metricName":"LGA75"}]],["Misassemblies",[{"values":[0,0,0,0,0,1,0,0,0,0,0,0,0,1],"quality":"Less is better","isMain":true,"metricName":"# misassemblies"},{"values":[0,0,0,0,0,1,0,0,0,0,0,0,0,1],"quality":"Less is better","isMain":false,"metricName":" # relocations"},{"values":[0,0,0,0,0,0,0,0,0,0,0,0,0,0],"quality":"Less is better","isMain":false,"metricName":" # translocations"},{"values":[0,0,0,0,0,0,0,0,0,0,0,0,0,0],"quality":"Less is better","isMain":false,"metricName":" # inversions"},{"values":[0,0,0,0,0,1,0,0,0,0,0,0,0,1],"quality":"Less is better","isMain":false,"metricName":"# misassembled contigs"},{"values":[0,0,0,0,0,3576,0,0,0,0,0,0,0,7676],"quality":"Less is better","isMain":true,"metricName":"Misassembled contigs length"},{"values":[0,0,0,0,0,2,0,0,0,1,0,0,0,3],"quality":"Less is better","isMain":false,"metricName":"# local misassemblies"},{"values":[0,0,0,0,0,0,0,0,1,0,0,0,0,1],"quality":"Less is better","isMain":false,"metricName":"# unaligned mis. contigs"}]],["Unaligned",[{"values":[0,0,0,0,0,0,0,0,0,0,0,0,0,0],"quality":"Less is better","isMain":false,"metricName":"# fully unaligned contigs"},{"values":[0,0,0,0,0,0,0,0,0,0,0,0,0,0],"quality":"Less is better","isMain":false,"metricName":"Fully unaligned length"},{"values":[0,1,0,0,0,2,0,0,1,2,0,0,0,4],"quality":"Less is better","isMain":false,"metricName":"# partially unaligned contigs"},{"values":[0,3935,0,0,0,3342,0,0,45958,2018,0,0,0,89239],"quality":"Less is better","isMain":false,"metricName":"Partially unaligned length"}]],["Mismatches",[{"values":[1,6,26,3,165,131,22,0,7,111,82,22,3,129],"quality":"Less is better","isMain":false,"metricName":"# mismatches"},{"values":[0,0,1,1,10,5,3,1,0,8,4,3,0,23],"quality":"Less is better","isMain":false,"metricName":"# indels"},{"values":[0,0,1,2,18,5,32,48,0,10,5,4,0,31],"quality":"Less is better","isMain":false,"metricName":"Indels length"},{"values":["2.02","15.10","46.48","6.22","376.45","211.00","43.31","0.00","12.23","215.16","157.78","41.84","11.60","230.51"],"quality":"Less is better","isMain":true,"metricName":"# mismatches per 100 kbp"},{"values":["0.00","0.00","1.79","2.07","22.82","8.05","5.91","26.93","0.00","15.51","7.70","5.71","0.00","41.10"],"quality":"Less is better","isMain":true,"metricName":"# indels per 100 kbp"},{"values":[0,0,1,1,9,5,2,0,0,8,4,3,0,23],"quality":"Less is better","isMain":false,"metricName":" # indels (<= 5 bp)"},{"values":[0,0,0,0,1,0,1,1,0,0,0,0,0,0],"quality":"Less is better","isMain":false,"metricName":" # indels (> 5 bp)"},{"values":[0,127,0,0,0,15,0,103,0,0,265,0,0,706],"quality":"Less is better","isMain":false,"metricName":"# N's"},{"values":["0.00","283.31","0.00","0.00","0.00","18.01","0.00","2682.99","0.00","0.00","501.01","0.00","0.00","474.49"],"quality":"Less is better","isMain":true,"metricName":"# N's per 100 kbp"}]],["Statistics without reference",[{"values":[20,20,23,30,23,20,17,6,15,7,20,30,19,5],"quality":"Equal","isMain":true,"metricName":"# contigs"},{"values":[11,9,17,16,15,13,14,0,12,6,12,14,11,5],"quality":"Equal","isMain":false,"metricName":"# contigs (>= 1000 bp)"},{"values":[3,3,2,2,1,4,4,0,3,2,3,2,0,4],"quality":"Equal","isMain":false,"metricName":"# contigs (>= 5000 bp)"},{"values":[0,0,0,0,0,2,0,0,2,2,1,0,0,3],"quality":"Equal","isMain":false,"metricName":"# contigs (>= 10000 bp)"},{"values":[0,0,0,0,0,1,0,0,1,1,0,0,0,1],"quality":"Equal","isMain":false,"metricName":"# contigs (>= 25000 bp)"},{"values":[0,0,0,0,0,0,0,0,1,0,0,0,0,1],"quality":"Equal","isMain":false,"metricName":"# contigs (>= 50000 bp)"},{"values":[8534,7801,8560,7086,7622,28556,9930,918,57853,34662,10418,8874,4338,107051],"quality":"More is better","isMain":true,"metricName":"Largest contig"},{"values":[49821,44828,56541,49410,45158,83266,52297,3839,104807,53700,52893,53872,25859,148792],"quality":"More is better","isMain":true,"metricName":"Total length"},{"values":[42871,37289,51913,39510,39178,77625,50051,0,102556,52931,47014,41902,20654,148792],"quality":"More is better","isMain":true,"metricName":"Total length (>= 1000 bp)"},{"values":[20541,22012,15013,13847,7622,54515,28666,0,84210,46010,27549,16871,0,143823],"quality":"More is better","isMain":false,"metricName":"Total length (>= 5000 bp)"},{"values":[0,0,0,0,0,41171,0,0,78472,46010,10418,0,0,136147],"quality":"More is better","isMain":true,"metricName":"Total length (>= 10000 bp)"},{"values":[0,0,0,0,0,28556,0,0,57853,34662,0,0,0,107051],"quality":"More is better","isMain":false,"metricName":"Total length (>= 25000 bp)"},{"values":[0,0,0,0,0,0,0,0,57853,0,0,0,0,107051],"quality":"More is better","isMain":true,"metricName":"Total length (>= 50000 bp)"},{"values":[3763,3836,3338,2063,2767,7889,5686,622,57853,34662,7805,2748,1568,107051],"quality":"More is better","isMain":false,"metricName":"N50"},{"values":[2230,2399,1987,1181,1271,2955,2104,550,5738,11348,1930,1116,1048,16912],"quality":"More is better","isMain":false,"metricName":"N75"},{"values":[5,4,6,6,5,3,4,3,1,1,3,5,5,1],"quality":"Less is better","isMain":false,"metricName":"L50"},{"values":[8,7,12,14,11,7,8,5,3,2,8,13,10,2],"quality":"Less is better","isMain":false,"metricName":"L75"},{"values":["51.28","50.73","51.07","51.24","50.87","50.80","51.14","48.10","50.28","51.24","51.03","51.04","51.68","49.05"],"quality":"Equal","isMain":false,"metricName":"GC (%)"}]],["Predicted genes",[]],["Similarity statistics",[{"values":[0,0,0,0,0,0,0,0,0,0,0,0,0,0],"quality":"Equal","isMain":false,"metricName":"# similar correct contigs"},{"values":[0,0,0,0,0,0,0,0,0,0,0,0,0,0],"quality":"Equal","isMain":false,"metricName":"# similar misassembled blocks"}]],["Reference statistics",[{"values":[62147,62147,62147,62147,62147,62147,62147,62147,62147,62147,62147,62147,62147,62147],"quality":"Equal","isMain":false,"metricName":"Reference length"},{"values":[1,1,1,1,1,1,1,1,1,1,1,1,1,1],"quality":"Equal","isMain":false,"metricName":"Reference fragments"},{"values":["50.91","50.91","50.91","50.91","50.91","50.91","50.91","50.91","50.91","50.91","50.91","50.91","50.91","50.91"],"quality":"Equal","isMain":false,"metricName":"Reference GC (%)"}]]],"referenceName":"gi\_209447126\_ref\_NC","date":"18 July 2018, Wednesday, 16:54:21","order":[0,1,2,3,4,5,6,7,8,9,10,11,12,13],"assembliesNames":["ABySS\_127","ABySS\_63","CLC","IDBA\_UD","MEGAHIT","MIRA","Ray\_Meta","SOAPdenovo2","SPAdes","SPAdes\_meta","SPAdes\_sc","SPAdes\_sc\_careful","Velvet","Geneious"]},{"assembliesWithNs":null,"minContig":500,"report":[["Genome statistics",[{"values":["100.000","97.113","100.000","98.912","100.000","100.000","76.071","50.715","100.000","97.529","96.787","96.787","89.177"],"quality":"More is better","isMain":true,"metricName":"Genome fraction (%)"},{"values":["1.003","1.011","1.000","1.004","1.002","1.007","1.047","1.041","1.003","1.000","1.002","1.000","1.003"],"quality":"Less is better","isMain":true,"metricName":"Duplication ratio"},{"values":[46996,42309,46872,41924,46983,47220,2923,6151,46998,45714,41836,43751,18945],"quality":"More is better","isMain":true,"metricName":"Largest alignment"},{"values":[46996,46040,46872,46557,46983,47220,37318,24058,46998,45714,45476,45366,41931],"quality":"More is better","isMain":true,"metricName":"Total aligned length"},{"values":[46996,42309,46872,41924,46983,47220,1107,2227,46998,45714,41836,43751,6033],"quality":"More is better","isMain":false,"metricName":"NG50"},{"values":[46996,42309,46872,41924,46983,47220,593,null,46998,45714,41836,43751,1130],"quality":"More is better","isMain":false,"metricName":"NG75"},{"values":[46996,42309,46872,41924,46983,47220,1236,2227,46998,45714,41836,43751,6033],"quality":"More is better","isMain":false,"metricName":"NA50"},{"values":[46996,42309,46872,41924,46983,47220,1020,null,46998,45714,41836,43751,2544],"quality":"More is better","isMain":false,"metricName":"NA75"},{"values":[46996,42309,46872,41924,46983,47220,1107,477,46998,45714,41836,43751,6033],"quality":"More is better","isMain":true,"metricName":"NGA50"},{"values":[46996,42309,46872,41924,46983,47220,593,null,46998,45714,41836,43751,1130],"quality":"More is better","isMain":false,"metricName":"NGA75"},{"values":[1,1,1,1,1,1,15,5,1,1,1,1,2],"quality":"Less is better","isMain":false,"metricName":"LG50"},{"values":[1,1,1,1,1,1,29,null,1,1,1,1,6],"quality":"Less is better","isMain":false,"metricName":"LG75"},{"values":[1,1,1,1,1,1,11,4,1,1,1,1,2],"quality":"Less is better","isMain":false,"metricName":"LA50"},{"values":[1,1,1,1,1,1,19,null,1,1,1,1,4],"quality":"Less is better","isMain":false,"metricName":"LA75"},{"values":[1,1,1,1,1,1,15,13,1,1,1,1,2],"quality":"Less is better","isMain":true,"metricName":"LGA50"},{"values":[1,1,1,1,1,1,29,null,1,1,1,1,6],"quality":"Less is better","isMain":false,"metricName":"LGA75"}]],["Misassemblies",[{"values":[0,0,0,0,0,0,0,0,0,0,0,0,0],"quality":"Less is better","isMain":true,"metricName":"# misassemblies"},{"values":[0,0,0,0,0,0,0,0,0,0,0,0,0],"quality":"Less is better","isMain":false,"metricName":" # relocations"},{"values":[0,0,0,0,0,0,0,0,0,0,0,0,0],"quality":"Less is better","isMain":false,"metricName":" # translocations"},{"values":[0,0,0,0,0,0,0,0,0,0,0,0,0],"quality":"Less is better","isMain":false,"metricName":" # inversions"},{"values":[0,0,0,0,0,0,0,0,0,0,0,0,0],"quality":"Less is better","isMain":false,"metricName":"# misassembled contigs"},{"values":[0,0,0,0,0,0,0,0,0,0,0,0,0],"quality":"Less is better","isMain":true,"metricName":"Misassembled contigs length"},{"values":[1,0,0,0,1,1,0,0,1,0,0,0,0],"quality":"Less is better","isMain":false,"metricName":"# local misassemblies"},{"values":[0,0,0,0,0,0,0,1,0,0,0,0,0],"quality":"Less is better","isMain":false,"metricName":"# unaligned mis. contigs"}]],["Unaligned",[{"values":[0,0,0,0,0,0,0,0,0,0,0,0,0],"quality":"Less is better","isMain":false,"metricName":"# fully unaligned contigs"},{"values":[0,0,0,0,0,0,0,0,0,0,0,0,0],"quality":"Less is better","isMain":false,"metricName":"Fully unaligned length"},{"values":[0,0,0,0,0,0,0,4,0,0,0,0,0],"quality":"Less is better","isMain":false,"metricName":"# partially unaligned contigs"},{"values":[0,0,0,0,0,0,0,7541,0,0,0,0,0],"quality":"Less is better","isMain":false,"metricName":"Partially unaligned length"}]],["Mismatches",[{"values":[0,0,0,0,0,1,1,0,0,0,0,0,0],"quality":"Less is better","isMain":false,"metricName":"# mismatches"},{"values":[0,0,0,0,0,0,0,16,0,0,0,0,0],"quality":"Less is better","isMain":false,"metricName":"# indels"},{"values":[0,0,0,0,0,0,0,757,0,0,0,0,0],"quality":"Less is better","isMain":false,"metricName":"Indels length"},{"values":["0.00","0.00","0.00","0.00","0.00","2.13","2.80","0.00","0.00","0.00","0.00","0.00","0.00"],"quality":"Less is better","isMain":true,"metricName":"# mismatches per 100 kbp"},{"values":["0.00","0.00","0.00","0.00","0.00","0.00","0.00","67.31","0.00","0.00","0.00","0.00","0.00"],"quality":"Less is better","isMain":true,"metricName":"# indels per 100 kbp"},{"values":[0,0,0,0,0,0,0,0,0,0,0,0,0],"quality":"Less is better","isMain":false,"metricName":" # indels (<= 5 bp)"},{"values":[0,0,0,0,0,0,0,16,0,0,0,0,0],"quality":"Less is better","isMain":false,"metricName":" # indels (> 5 bp)"},{"values":[0,0,0,0,0,0,0,3023,0,0,0,0,0],"quality":"Less is better","isMain":false,"metricName":"# N's"},{"values":["0.00","0.00","0.00","0.00","0.00","0.00","0.00","9363.48","0.00","0.00","0.00","0.00","0.00"],"quality":"Less is better","isMain":true,"metricName":"# N's per 100 kbp"}]],["Statistics without reference",[{"values":[1,3,1,4,1,1,32,13,1,1,4,2,14],"quality":"Equal","isMain":true,"metricName":"# contigs"},{"values":[1,2,1,2,1,1,19,7,1,1,3,2,7],"quality":"Equal","isMain":false,"metricName":"# contigs (>= 1000 bp)"},{"values":[1,1,1,1,1,1,0,2,1,1,1,1,2],"quality":"Equal","isMain":false,"metricName":"# contigs (>= 5000 bp)"},{"values":[1,1,1,1,1,1,0,0,1,1,1,1,1],"quality":"Equal","isMain":false,"metricName":"# contigs (>= 10000 bp)"},{"values":[1,1,1,1,1,1,0,0,1,1,1,1,0],"quality":"Equal","isMain":false,"metricName":"# contigs (>= 25000 bp)"},{"values":[0,0,0,0,0,0,0,0,0,0,0,0,0],"quality":"Equal","isMain":false,"metricName":"# contigs (>= 50000 bp)"},{"values":[46996,42309,46872,41924,46983,47220,2923,7857,46998,45714,41836,43751,18945],"quality":"More is better","isMain":true,"metricName":"Largest contig"},{"values":[46996,46040,46872,46557,46983,47220,37318,32285,46998,45714,45476,45366,41931],"quality":"More is better","isMain":true,"metricName":"Total length"},{"values":[46996,45181,46872,44830,46983,47220,28019,27510,46998,45714,44955,45366,36406],"quality":"More is better","isMain":true,"metricName":"Total length (>= 1000 bp)"},{"values":[46996,42309,46872,41924,46983,47220,0,13758,46998,45714,41836,43751,24978],"quality":"More is better","isMain":false,"metricName":"Total length (>= 5000 bp)"},{"values":[46996,42309,46872,41924,46983,47220,0,0,46998,45714,41836,43751,18945],"quality":"More is better","isMain":true,"metricName":"Total length (>= 10000 bp)"},{"values":[46996,42309,46872,41924,46983,47220,0,0,46998,45714,41836,43751,0],"quality":"More is better","isMain":false,"metricName":"Total length (>= 25000 bp)"},{"values":[0,0,0,0,0,0,0,0,0,0,0,0,0],"quality":"More is better","isMain":true,"metricName":"Total length (>= 50000 bp)"},{"values":[46996,42309,46872,41924,46983,47220,1236,4774,46998,45714,41836,43751,6033],"quality":"More is better","isMain":false,"metricName":"N50"},{"values":[46996,42309,46872,41924,46983,47220,1020,2227,46998,45714,41836,43751,2544],"quality":"More is better","isMain":false,"metricName":"N75"},{"values":[1,1,1,1,1,1,11,3,1,1,1,1,2],"quality":"Less is better","isMain":false,"metricName":"L50"},{"values":[1,1,1,1,1,1,19,5,1,1,1,1,4],"quality":"Less is better","isMain":false,"metricName":"L75"},{"values":["56.28","56.25","56.26","56.24","56.25","56.22","56.20","55.68","56.28","56.21","56.19","56.17","55.84"],"quality":"Equal","isMain":false,"metricName":"GC (%)"}]],["Predicted genes",[]],["Similarity statistics",[{"values":[0,0,0,0,0,0,0,0,0,0,0,0,0],"quality":"Equal","isMain":false,"metricName":"# similar correct contigs"},{"values":[0,0,0,0,0,0,0,0,0,0,0,0,0],"quality":"Equal","isMain":false,"metricName":"# similar misassembled blocks"}]],["Reference statistics",[{"values":[46872,46872,46872,46872,46872,46872,46872,46872,46872,46872,46872,46872,46872],"quality":"Equal","isMain":false,"metricName":"Reference length"},{"values":[1,1,1,1,1,1,1,1,1,1,1,1,1],"quality":"Equal","isMain":false,"metricName":"Reference fragments"},{"values":["56.26","56.26","56.26","56.26","56.26","56.26","56.26","56.26","56.26","56.26","56.26","56.26","56.26"],"quality":"Equal","isMain":false,"metricName":"Reference GC (%)"}]]],"referenceName":"gi\_209552419\_ref\_NC","date":"18 July 2018, Wednesday, 16:54:37","order":[0,1,2,3,4,5,6,7,8,9,10,11,12],"assembliesNames":["ABySS\_127","ABySS\_63","CLC","IDBA\_UD","MEGAHIT","MIRA","Ray\_Meta","SOAPdenovo2","SPAdes","SPAdes\_meta","SPAdes\_sc","SPAdes\_sc\_careful","Velvet"]},{"assembliesWithNs":null,"minContig":500,"report":[["Genome statistics",[{"values":["83.894","80.522","81.881","79.281","89.042","100.000","86.234","82.058","62.902","75.618","77.451","68.095","99.006"],"quality":"More is better","isMain":true,"metricName":"Genome fraction (%)"},{"values":["1.000","1.002","1.002","1.004","1.006","1.737","1.009","1.000","1.000","1.000","1.000","1.000","1.600"],"quality":"Less is better","isMain":true,"metricName":"Duplication ratio"},{"values":[29240,27413,27375,23699,27471,31151,28180,27585,22485,27055,27055,21834,24172],"quality":"More is better","isMain":true,"metricName":"Largest alignment"},{"values":[42727,41072,41779,40528,45630,88253,44301,41792,32045,38512,39446,34681,80657],"quality":"More is better","isMain":true,"metricName":"Total aligned length"},{"values":[29240,27413,27375,4685,27471,116371,28180,27585,28279,27055,27055,4141,213470],"quality":"More is better","isMain":false,"metricName":"NG50"},{"values":[4313,6293,3796,1332,4034,116371,4503,4006,null,1055,3280,null,213470],"quality":"More is better","isMain":false,"metricName":"NG75"},{"values":[29240,27413,27375,23699,27471,null,28180,27585,22485,27055,27055,21834,null],"quality":"More is better","isMain":false,"metricName":"NA50"},{"values":[8182,4251,7529,3707,4485,null,8347,7832,5823,4362,7960,3084,null],"quality":"More is better","isMain":false,"metricName":"NA75"},{"values":[29240,27413,27375,4685,27471,31151,28180,27585,5823,27055,27055,4141,17483],"quality":"More is better","isMain":true,"metricName":"NGA50"},{"values":[4313,1633,3796,1332,4034,7771,4503,4006,null,1055,3280,null,17483],"quality":"More is better","isMain":false,"metricName":"NGA75"},{"values":[1,1,1,2,1,1,1,1,1,1,1,2,1],"quality":"Less is better","isMain":false,"metricName":"LG50"},{"values":[3,3,3,6,4,1,3,3,null,6,3,null,1],"quality":"Less is better","isMain":false,"metricName":"LG75"},{"values":[1,1,1,1,1,null,1,1,1,1,1,1,null],"quality":"Less is better","isMain":false,"metricName":"LA50"},{"values":[2,3,2,3,3,null,2,2,2,2,2,3,null],"quality":"Less is better","isMain":false,"metricName":"LA75"},{"values":[1,1,1,2,1,1,1,1,2,1,1,2,2],"quality":"Less is better","isMain":true,"metricName":"LGA50"},{"values":[3,4,3,6,4,2,3,3,null,6,3,null,2],"quality":"Less is better","isMain":false,"metricName":"LGA75"}]],["Misassemblies",[{"values":[0,0,0,0,0,3,0,0,1,0,0,0,0],"quality":"Less is better","isMain":true,"metricName":"# misassemblies"},{"values":[0,0,0,0,0,3,0,0,1,0,0,0,0],"quality":"Less is better","isMain":false,"metricName":" # relocations"},{"values":[0,0,0,0,0,0,0,0,0,0,0,0,0],"quality":"Less is better","isMain":false,"metricName":" # translocations"},{"values":[0,0,0,0,0,0,0,0,0,0,0,0,0],"quality":"Less is better","isMain":false,"metricName":" # inversions"},{"values":[0,0,0,0,0,1,0,0,1,0,0,0,0],"quality":"Less is better","isMain":false,"metricName":"# misassembled contigs"},{"values":[0,0,0,0,0,62391,0,0,28279,0,0,0,0],"quality":"Less is better","isMain":true,"metricName":"Misassembled contigs length"},{"values":[0,0,0,0,0,2,0,0,0,0,0,0,0],"quality":"Less is better","isMain":false,"metricName":"# local misassemblies"},{"values":[0,0,0,0,0,1,0,0,0,0,0,0,1],"quality":"Less is better","isMain":false,"metricName":"# unaligned mis. contigs"}]],["Unaligned",[{"values":[0,0,0,0,0,0,0,0,0,0,0,0,0],"quality":"Less is better","isMain":false,"metricName":"# fully unaligned contigs"},{"values":[0,0,0,0,0,0,0,0,0,0,0,0,0],"quality":"Less is better","isMain":false,"metricName":"Fully unaligned length"},{"values":[0,1,0,0,0,3,0,0,1,0,0,0,1],"quality":"Less is better","isMain":false,"metricName":"# partially unaligned contigs"},{"values":[0,4568,0,0,0,133637,0,0,2057,0,0,0,132813],"quality":"Less is better","isMain":false,"metricName":"Partially unaligned length"}]],["Mismatches",[{"values":[0,27,1,0,1,89,0,0,55,5,49,1,232],"quality":"Less is better","isMain":false,"metricName":"# mismatches"},{"values":[0,0,0,0,0,11,0,0,11,0,0,0,16],"quality":"Less is better","isMain":false,"metricName":"# indels"},{"values":[0,0,0,0,0,13,0,0,19,0,0,0,23],"quality":"Less is better","isMain":false,"metricName":"Indels length"},{"values":["0.00","65.84","2.40","0.00","2.21","174.75","0.00","0.00","171.68","12.98","124.22","2.88","460.10"],"quality":"Less is better","isMain":true,"metricName":"# mismatches per 100 kbp"},{"values":["0.00","0.00","0.00","0.00","0.00","21.60","0.00","0.00","34.34","0.00","0.00","0.00","31.73"],"quality":"Less is better","isMain":true,"metricName":"# indels per 100 kbp"},{"values":[0,0,0,0,0,11,0,0,10,0,0,0,16],"quality":"Less is better","isMain":false,"metricName":" # indels (<= 5 bp)"},{"values":[0,0,0,0,0,0,0,0,1,0,0,0,0],"quality":"Less is better","isMain":false,"metricName":" # indels (> 5 bp)"},{"values":[1,323,0,0,0,28,0,0,0,0,0,0,210],"quality":"Less is better","isMain":false,"metricName":"# N's"},{"values":["2.34","707.71","0.00","0.00","0.00","12.61","0.00","0.00","0.00","0.00","0.00","0.00","98.37"],"quality":"Less is better","isMain":true,"metricName":"# N's per 100 kbp"}]],["Statistics without reference",[{"values":[4,6,6,9,8,8,5,5,2,6,4,9,1],"quality":"Equal","isMain":true,"metricName":"# contigs"},{"values":[3,4,5,6,7,8,5,5,2,6,4,5,1],"quality":"Equal","isMain":false,"metricName":"# contigs (>= 1000 bp)"},{"values":[2,3,2,1,1,3,2,2,2,1,2,1,1],"quality":"Equal","isMain":false,"metricName":"# contigs (>= 5000 bp)"},{"values":[1,1,1,1,1,3,1,1,1,1,1,1,1],"quality":"Equal","isMain":false,"metricName":"# contigs (>= 10000 bp)"},{"values":[1,1,1,0,1,3,1,1,1,1,1,0,1],"quality":"Equal","isMain":false,"metricName":"# contigs (>= 25000 bp)"},{"values":[0,0,0,0,0,2,0,0,0,0,0,0,1],"quality":"Equal","isMain":false,"metricName":"# contigs (>= 50000 bp)"},{"values":[29240,27413,27375,23699,27471,116371,28180,27585,28279,27055,27055,21834,213470],"quality":"More is better","isMain":true,"metricName":"Largest contig"},{"values":[42727,45640,41779,40528,45630,222098,44301,41792,34102,38512,39446,34681,213470],"quality":"More is better","isMain":true,"metricName":"Total length"},{"values":[41735,44158,40965,38292,44729,222098,44301,41792,34102,38512,39446,31758,213470],"quality":"More is better","isMain":true,"metricName":"Total length (>= 1000 bp)"},{"values":[37422,42525,34904,23699,27471,211022,36527,35417,34102,27055,35015,21834,213470],"quality":"More is better","isMain":false,"metricName":"Total length (>= 5000 bp)"},{"values":[29240,27413,27375,23699,27471,211022,28180,27585,28279,27055,27055,21834,213470],"quality":"More is better","isMain":true,"metricName":"Total length (>= 10000 bp)"},{"values":[29240,27413,27375,0,27471,211022,28180,27585,28279,27055,27055,0,213470],"quality":"More is better","isMain":false,"metricName":"Total length (>= 25000 bp)"},{"values":[0,0,0,0,0,178762,0,0,0,0,0,0,213470],"quality":"More is better","isMain":true,"metricName":"Total length (>= 50000 bp)"},{"values":[29240,27413,27375,23699,27471,116371,28180,27585,28279,27055,27055,21834,213470],"quality":"More is better","isMain":false,"metricName":"N50"},{"values":[8182,8819,7529,3707,4485,62391,8347,7832,28279,4362,7960,3084,213470],"quality":"More is better","isMain":false,"metricName":"N75"},{"values":[1,1,1,1,1,1,1,1,1,1,1,1,1],"quality":"Less is better","isMain":false,"metricName":"L50"},{"values":[2,2,2,3,3,2,2,2,1,2,2,3,1],"quality":"Less is better","isMain":false,"metricName":"L75"},{"values":["29.78","29.72","29.64","29.73","29.29","28.83","29.77","29.69","29.60","29.72","29.71","29.75","28.86"],"quality":"Equal","isMain":false,"metricName":"GC (%)"}]],["Predicted genes",[]],["Similarity statistics",[{"values":[1,1,1,1,1,0,1,1,0,1,1,1,0],"quality":"Equal","isMain":false,"metricName":"# similar correct contigs"},{"values":[0,0,0,0,0,0,0,0,0,0,0,0,0],"quality":"Equal","isMain":false,"metricName":"# similar misassembled blocks"}]],["Reference statistics",[{"values":[50930,50930,50930,50930,50930,50930,50930,50930,50930,50930,50930,50930,50930],"quality":"Equal","isMain":false,"metricName":"Reference length"},{"values":[1,1,1,1,1,1,1,1,1,1,1,1,1],"quality":"Equal","isMain":false,"metricName":"Reference fragments"},{"values":["29.33","29.33","29.33","29.33","29.33","29.33","29.33","29.33","29.33","29.33","29.33","29.33","29.33"],"quality":"Equal","isMain":false,"metricName":"Reference GC (%)"}]]],"referenceName":"gi\_209901237\_ref\_NC","date":"18 July 2018, Wednesday, 16:54:55","order":[0,1,2,3,4,5,6,7,8,9,10,11,12],"assembliesNames":["ABySS\_127","ABySS\_63","CLC","IDBA\_UD","MEGAHIT","MIRA","Ray\_Meta","SPAdes","SPAdes\_meta","SPAdes\_sc","SPAdes\_sc\_careful","Velvet","Geneious"]},{"assembliesWithNs":null,"minContig":500,"report":[["Genome statistics",[{"values":["99.754","100.000","100.000","100.000","100.000","100.000","100.000","100.000","100.000","98.466","100.000"],"quality":"More is better","isMain":true,"metricName":"Genome fraction (%)"},{"values":["1.002","1.011","1.016","1.075","1.005","1.014","1.006","1.006","1.006","1.000","1.037"],"quality":"Less is better","isMain":true,"metricName":"Duplication ratio"},{"values":[8907,9028,9070,8890,8978,9056,8984,8984,8984,8792,9262],"quality":"More is better","isMain":true,"metricName":"Largest alignment"},{"values":[8907,9028,9070,9599,8978,9056,8984,8984,8984,8792,9262],"quality":"More is better","isMain":true,"metricName":"Total aligned length"},{"values":[8929,9028,9070,8890,8978,9056,8984,8984,8984,8792,9262],"quality":"More is better","isMain":false,"metricName":"NG50"},{"values":[8929,9028,9070,8890,8978,9056,8984,8984,8984,8792,9262],"quality":"More is better","isMain":false,"metricName":"NG75"},{"values":[8907,9028,9070,8890,8978,9056,8984,8984,8984,8792,9262],"quality":"More is better","isMain":false,"metricName":"NA50"},{"values":[8907,9028,9070,8890,8978,9056,8984,8984,8984,8792,9262],"quality":"More is better","isMain":false,"metricName":"NA75"},{"values":[8907,9028,9070,8890,8978,9056,8984,8984,8984,8792,9262],"quality":"More is better","isMain":true,"metricName":"NGA50"},{"values":[8907,9028,9070,8890,8978,9056,8984,8984,8984,8792,9262],"quality":"More is better","isMain":false,"metricName":"NGA75"},{"values":[1,1,1,1,1,1,1,1,1,1,1],"quality":"Less is better","isMain":false,"metricName":"LG50"},{"values":[1,1,1,1,1,1,1,1,1,1,1],"quality":"Less is better","isMain":false,"metricName":"LG75"},{"values":[1,1,1,1,1,1,1,1,1,1,1],"quality":"Less is better","isMain":false,"metricName":"LA50"},{"values":[1,1,1,1,1,1,1,1,1,1,1],"quality":"Less is better","isMain":false,"metricName":"LA75"},{"values":[1,1,1,1,1,1,1,1,1,1,1],"quality":"Less is better","isMain":true,"metricName":"LGA50"},{"values":[1,1,1,1,1,1,1,1,1,1,1],"quality":"Less is better","isMain":false,"metricName":"LGA75"}]],["Misassemblies",[{"values":[0,0,0,0,0,0,0,0,0,0,0],"quality":"Less is better","isMain":true,"metricName":"# misassemblies"},{"values":[0,0,0,0,0,0,0,0,0,0,0],"quality":"Less is better","isMain":false,"metricName":" # relocations"},{"values":[0,0,0,0,0,0,0,0,0,0,0],"quality":"Less is better","isMain":false,"metricName":" # translocations"},{"values":[0,0,0,0,0,0,0,0,0,0,0],"quality":"Less is better","isMain":false,"metricName":" # inversions"},{"values":[0,0,0,0,0,0,0,0,0,0,0],"quality":"Less is better","isMain":false,"metricName":"# misassembled contigs"},{"values":[0,0,0,0,0,0,0,0,0,0,0],"quality":"Less is better","isMain":true,"metricName":"Misassembled contigs length"},{"values":[0,1,1,0,0,1,0,0,0,0,1],"quality":"Less is better","isMain":false,"metricName":"# local misassemblies"},{"values":[0,0,0,0,0,0,0,0,0,0,0],"quality":"Less is better","isMain":false,"metricName":"# unaligned mis. contigs"}]],["Unaligned",[{"values":[0,0,0,0,0,0,0,0,0,0,0],"quality":"Less is better","isMain":false,"metricName":"# fully unaligned contigs"},{"values":[0,0,0,0,0,0,0,0,0,0,0],"quality":"Less is better","isMain":false,"metricName":"Fully unaligned length"},{"values":[0,0,0,0,0,0,0,0,0,0,0],"quality":"Less is better","isMain":false,"metricName":"# partially unaligned contigs"},{"values":[0,0,0,0,0,0,0,0,0,0,0],"quality":"Less is better","isMain":false,"metricName":"Partially unaligned length"}]],["Mismatches",[{"values":[0,0,0,0,0,0,0,0,0,0,0],"quality":"Less is better","isMain":false,"metricName":"# mismatches"},{"values":[0,0,0,0,1,0,1,1,1,0,0],"quality":"Less is better","isMain":false,"metricName":"# indels"},{"values":[0,0,0,0,49,0,55,55,55,0,0],"quality":"Less is better","isMain":false,"metricName":"Indels length"},{"values":["0.00","0.00","0.00","0.00","0.00","0.00","0.00","0.00","0.00","0.00","0.00"],"quality":"Less is better","isMain":true,"metricName":"# mismatches per 100 kbp"},{"values":["0.00","0.00","0.00","0.00","11.20","0.00","11.20","11.20","11.20","0.00","0.00"],"quality":"Less is better","isMain":true,"metricName":"# indels per 100 kbp"},{"values":[0,0,0,0,0,0,0,0,0,0,0],"quality":"Less is better","isMain":false,"metricName":" # indels (<= 5 bp)"},{"values":[0,0,0,0,1,0,1,1,1,0,0],"quality":"Less is better","isMain":false,"metricName":" # indels (> 5 bp)"},{"values":[0,0,0,0,0,0,0,0,0,0,0],"quality":"Less is better","isMain":false,"metricName":"# N's"},{"values":["0.00","0.00","0.00","0.00","0.00","0.00","0.00","0.00","0.00","0.00","0.00"],"quality":"Less is better","isMain":true,"metricName":"# N's per 100 kbp"}]],["Statistics without reference",[{"values":[1,1,1,2,1,1,1,1,1,1,1],"quality":"Equal","isMain":true,"metricName":"# contigs"},{"values":[1,1,1,1,1,1,1,1,1,1,1],"quality":"Equal","isMain":false,"metricName":"# contigs (>= 1000 bp)"},{"values":[1,1,1,1,1,1,1,1,1,1,1],"quality":"Equal","isMain":false,"metricName":"# contigs (>= 5000 bp)"},{"values":[0,0,0,0,0,0,0,0,0,0,0],"quality":"Equal","isMain":false,"metricName":"# contigs (>= 10000 bp)"},{"values":[0,0,0,0,0,0,0,0,0,0,0],"quality":"Equal","isMain":false,"metricName":"# contigs (>= 25000 bp)"},{"values":[0,0,0,0,0,0,0,0,0,0,0],"quality":"Equal","isMain":false,"metricName":"# contigs (>= 50000 bp)"},{"values":[8929,9028,9070,8890,8978,9056,8984,8984,8984,8792,9262],"quality":"More is better","isMain":true,"metricName":"Largest contig"},{"values":[8929,9028,9070,9599,8978,9056,8984,8984,8984,8792,9262],"quality":"More is better","isMain":true,"metricName":"Total length"},{"values":[8929,9028,9070,8890,8978,9056,8984,8984,8984,8792,9262],"quality":"More is better","isMain":true,"metricName":"Total length (>= 1000 bp)"},{"values":[8929,9028,9070,8890,8978,9056,8984,8984,8984,8792,9262],"quality":"More is better","isMain":false,"metricName":"Total length (>= 5000 bp)"},{"values":[0,0,0,0,0,0,0,0,0,0,0],"quality":"More is better","isMain":true,"metricName":"Total length (>= 10000 bp)"},{"values":[0,0,0,0,0,0,0,0,0,0,0],"quality":"More is better","isMain":false,"metricName":"Total length (>= 25000 bp)"},{"values":[0,0,0,0,0,0,0,0,0,0,0],"quality":"More is better","isMain":true,"metricName":"Total length (>= 50000 bp)"},{"values":[8929,9028,9070,8890,8978,9056,8984,8984,8984,8792,9262],"quality":"More is better","isMain":false,"metricName":"N50"},{"values":[8929,9028,9070,8890,8978,9056,8984,8984,8984,8792,9262],"quality":"More is better","isMain":false,"metricName":"N75"},{"values":[1,1,1,1,1,1,1,1,1,1,1],"quality":"Less is better","isMain":false,"metricName":"L50"},{"values":[1,1,1,1,1,1,1,1,1,1,1],"quality":"Less is better","isMain":false,"metricName":"L75"},{"values":["59.68","59.70","59.61","59.96","59.67","59.60","59.64","59.64","59.64","59.60","59.74"],"quality":"Equal","isMain":false,"metricName":"GC (%)"}]],["Predicted genes",[]],["Similarity statistics",[{"values":[0,0,0,0,0,0,0,0,0,0,0],"quality":"Equal","isMain":false,"metricName":"# similar correct contigs"},{"values":[0,0,0,0,0,0,0,0,0,0,0],"quality":"Equal","isMain":false,"metricName":"# similar misassembled blocks"}]],["Reference statistics",[{"values":[8929,8929,8929,8929,8929,8929,8929,8929,8929,8929,8929],"quality":"Equal","isMain":false,"metricName":"Reference length"},{"values":[1,1,1,1,1,1,1,1,1,1,1],"quality":"Equal","isMain":false,"metricName":"Reference fragments"},{"values":["59.65","59.65","59.65","59.65","59.65","59.65","59.65","59.65","59.65","59.65","59.65"],"quality":"Equal","isMain":false,"metricName":"Reference GC (%)"}]]],"referenceName":"gi\_209901313\_ref\_NC","date":"18 July 2018, Wednesday, 16:55:08","order":[0,1,2,3,4,5,6,7,8,9,10],"assembliesNames":["CLC","IDBA\_UD","MEGAHIT","MIRA","Ray\_Meta","SPAdes","SPAdes\_meta","SPAdes\_sc","SPAdes\_sc\_careful","Velvet","Geneious"]},{"assembliesWithNs":null,"minContig":500,"report":[["Genome statistics",[{"values":["100.000","99.949","100.000","100.000","100.000","99.985","99.952","0.390","100.000","99.943","99.943","99.943","99.897"],"quality":"More is better","isMain":true,"metricName":"Genome fraction (%)"},{"values":["1.918","1.000","1.000","1.001","1.001","1.000","1.001","1.000","1.001","1.000","1.000","1.000","1.001"],"quality":"Less is better","isMain":true,"metricName":"Duplication ratio"},{"values":[132562,132494,132562,132562,132703,132542,132499,517,132689,132487,132487,132487,66983],"quality":"More is better","isMain":true,"metricName":"Largest alignment"},{"values":[254220,132494,132562,132562,132703,132542,132499,517,132689,132487,132487,132487,132470],"quality":"More is better","isMain":true,"metricName":"Total aligned length"},{"values":[254220,132558,132562,132661,132703,132542,132568,null,132689,132544,132544,132544,66983],"quality":"More is better","isMain":false,"metricName":"NG50"},{"values":[254220,132558,132562,132661,132703,132542,132568,null,132689,132544,132544,132544,51368],"quality":"More is better","isMain":false,"metricName":"NG75"},{"values":[132562,132494,132562,132562,132703,132542,132499,517,132689,132487,132487,132487,66983],"quality":"More is better","isMain":false,"metricName":"NA50"},{"values":[62971,132494,132562,132562,132703,132542,132499,517,132689,132487,132487,132487,51368],"quality":"More is better","isMain":false,"metricName":"NA75"},{"values":[132562,132494,132562,132562,132703,132542,132499,null,132689,132487,132487,132487,66983],"quality":"More is better","isMain":true,"metricName":"NGA50"},{"values":[132562,132494,132562,132562,132703,132542,132499,null,132689,132487,132487,132487,51368],"quality":"More is better","isMain":false,"metricName":"NGA75"},{"values":[1,1,1,1,1,1,1,null,1,1,1,1,1],"quality":"Less is better","isMain":false,"metricName":"LG50"},{"values":[1,1,1,1,1,1,1,null,1,1,1,1,2],"quality":"Less is better","isMain":false,"metricName":"LG75"},{"values":[1,1,1,1,1,1,1,1,1,1,1,1,1],"quality":"Less is better","isMain":false,"metricName":"LA50"},{"values":[2,1,1,1,1,1,1,1,1,1,1,1,2],"quality":"Less is better","isMain":false,"metricName":"LA75"},{"values":[1,1,1,1,1,1,1,null,1,1,1,1,1],"quality":"Less is better","isMain":true,"metricName":"LGA50"},{"values":[1,1,1,1,1,1,1,null,1,1,1,1,2],"quality":"Less is better","isMain":false,"metricName":"LGA75"}]],["Misassemblies",[{"values":[2,0,0,0,0,0,0,0,0,0,0,0,0],"quality":"Less is better","isMain":true,"metricName":"# misassemblies"},{"values":[2,0,0,0,0,0,0,0,0,0,0,0,0],"quality":"Less is better","isMain":false,"metricName":" # relocations"},{"values":[0,0,0,0,0,0,0,0,0,0,0,0,0],"quality":"Less is better","isMain":false,"metricName":" # translocations"},{"values":[0,0,0,0,0,0,0,0,0,0,0,0,0],"quality":"Less is better","isMain":false,"metricName":" # inversions"},{"values":[1,0,0,0,0,0,0,0,0,0,0,0,0],"quality":"Less is better","isMain":false,"metricName":"# misassembled contigs"},{"values":[254220,0,0,0,0,0,0,0,0,0,0,0,0],"quality":"Less is better","isMain":true,"metricName":"Misassembled contigs length"},{"values":[0,0,0,0,1,0,0,0,1,0,0,0,0],"quality":"Less is better","isMain":false,"metricName":"# local misassemblies"},{"values":[0,0,0,0,0,0,0,0,0,0,0,0,0],"quality":"Less is better","isMain":false,"metricName":"# unaligned mis. contigs"}]],["Unaligned",[{"values":[0,0,0,0,0,0,0,0,0,0,0,0,0],"quality":"Less is better","isMain":false,"metricName":"# fully unaligned contigs"},{"values":[0,0,0,0,0,0,0,0,0,0,0,0,0],"quality":"Less is better","isMain":false,"metricName":"Fully unaligned length"},{"values":[0,0,0,0,0,0,0,0,0,0,0,0,0],"quality":"Less is better","isMain":false,"metricName":"# partially unaligned contigs"},{"values":[0,0,0,0,0,0,0,0,0,0,0,0,0],"quality":"Less is better","isMain":false,"metricName":"Partially unaligned length"}]],["Mismatches",[{"values":[0,0,2,0,0,0,0,0,0,0,0,0,0],"quality":"Less is better","isMain":false,"metricName":"# mismatches"},{"values":[0,0,0,0,0,0,0,0,0,0,0,0,0],"quality":"Less is better","isMain":false,"metricName":"# indels"},{"values":[0,0,0,0,0,0,0,0,0,0,0,0,0],"quality":"Less is better","isMain":false,"metricName":"Indels length"},{"values":["0.00","0.00","1.51","0.00","0.00","0.00","0.00","0.00","0.00","0.00","0.00","0.00","0.00"],"quality":"Less is better","isMain":true,"metricName":"# mismatches per 100 kbp"},{"values":["0.00","0.00","0.00","0.00","0.00","0.00","0.00","0.00","0.00","0.00","0.00","0.00","0.00"],"quality":"Less is better","isMain":true,"metricName":"# indels per 100 kbp"},{"values":[0,0,0,0,0,0,0,0,0,0,0,0,0],"quality":"Less is better","isMain":false,"metricName":" # indels (<= 5 bp)"},{"values":[0,0,0,0,0,0,0,0,0,0,0,0,0],"quality":"Less is better","isMain":false,"metricName":" # indels (> 5 bp)"},{"values":[0,0,0,0,0,1,0,0,0,0,0,0,0],"quality":"Less is better","isMain":false,"metricName":"# N's"},{"values":["0.00","0.00","0.00","0.00","0.00","0.75","0.00","0.00","0.00","0.00","0.00","0.00","0.00"],"quality":"Less is better","isMain":true,"metricName":"# N's per 100 kbp"}]],["Statistics without reference",[{"values":[1,1,1,1,1,1,1,1,1,1,1,1,4],"quality":"Equal","isMain":true,"metricName":"# contigs"},{"values":[1,1,1,1,1,1,1,0,1,1,1,1,4],"quality":"Equal","isMain":false,"metricName":"# contigs (>= 1000 bp)"},{"values":[1,1,1,1,1,1,1,0,1,1,1,1,3],"quality":"Equal","isMain":false,"metricName":"# contigs (>= 5000 bp)"},{"values":[1,1,1,1,1,1,1,0,1,1,1,1,3],"quality":"Equal","isMain":false,"metricName":"# contigs (>= 10000 bp)"},{"values":[1,1,1,1,1,1,1,0,1,1,1,1,2],"quality":"Equal","isMain":false,"metricName":"# contigs (>= 25000 bp)"},{"values":[1,1,1,1,1,1,1,0,1,1,1,1,2],"quality":"Equal","isMain":false,"metricName":"# contigs (>= 50000 bp)"},{"values":[254220,132558,132562,132661,132703,132542,132568,517,132689,132544,132544,132544,66983],"quality":"More is better","isMain":true,"metricName":"Largest contig"},{"values":[254220,132558,132562,132661,132703,132542,132568,517,132689,132544,132544,132544,132502],"quality":"More is better","isMain":true,"metricName":"Total length"},{"values":[254220,132558,132562,132661,132703,132542,132568,0,132689,132544,132544,132544,132502],"quality":"More is better","isMain":true,"metricName":"Total length (>= 1000 bp)"},{"values":[254220,132558,132562,132661,132703,132542,132568,0,132689,132544,132544,132544,129760],"quality":"More is better","isMain":false,"metricName":"Total length (>= 5000 bp)"},{"values":[254220,132558,132562,132661,132703,132542,132568,0,132689,132544,132544,132544,129760],"quality":"More is better","isMain":true,"metricName":"Total length (>= 10000 bp)"},{"values":[254220,132558,132562,132661,132703,132542,132568,0,132689,132544,132544,132544,118351],"quality":"More is better","isMain":false,"metricName":"Total length (>= 25000 bp)"},{"values":[254220,132558,132562,132661,132703,132542,132568,0,132689,132544,132544,132544,118351],"quality":"More is better","isMain":true,"metricName":"Total length (>= 50000 bp)"},{"values":[254220,132558,132562,132661,132703,132542,132568,517,132689,132544,132544,132544,66983],"quality":"More is better","isMain":false,"metricName":"N50"},{"values":[254220,132558,132562,132661,132703,132542,132568,517,132689,132544,132544,132544,51368],"quality":"More is better","isMain":false,"metricName":"N75"},{"values":[1,1,1,1,1,1,1,1,1,1,1,1,1],"quality":"Less is better","isMain":false,"metricName":"L50"},{"values":[1,1,1,1,1,1,1,1,1,1,1,1,2],"quality":"Less is better","isMain":false,"metricName":"L75"},{"values":["39.97","39.96","39.97","39.97","39.96","39.97","39.97","39.46","39.96","39.96","39.96","39.96","39.97"],"quality":"Equal","isMain":false,"metricName":"GC (%)"}]],["Predicted genes",[]],["Similarity statistics",[{"values":[0,0,0,0,0,0,0,0,0,0,0,0,0],"quality":"Equal","isMain":false,"metricName":"# similar correct contigs"},{"values":[0,0,0,0,0,0,0,0,0,0,0,0,0],"quality":"Equal","isMain":false,"metricName":"# similar misassembled blocks"}]],["Reference statistics",[{"values":[132562,132562,132562,132562,132562,132562,132562,132562,132562,132562,132562,132562,132562],"quality":"Equal","isMain":false,"metricName":"Reference length"},{"values":[1,1,1,1,1,1,1,1,1,1,1,1,1],"quality":"Equal","isMain":false,"metricName":"Reference fragments"},{"values":["39.96","39.96","39.96","39.96","39.96","39.96","39.96","39.96","39.96","39.96","39.96","39.96","39.96"],"quality":"Equal","isMain":false,"metricName":"Reference GC (%)"}]]],"referenceName":"gi\_209972829\_ref\_NC","date":"18 July 2018, Wednesday, 16:55:27","order":[0,1,2,3,4,5,6,7,8,9,10,11,12],"assembliesNames":["ABySS\_127","ABySS\_63","CLC","IDBA\_UD","MEGAHIT","MIRA","Ray\_Meta","SOAPdenovo2","SPAdes","SPAdes\_meta","SPAdes\_sc","SPAdes\_sc\_careful","Velvet"]},{"assembliesWithNs":null,"minContig":500,"report":[["Genome statistics",[{"values":["99.944","100.000","100.000","100.000","100.000","100.000","100.000","100.000","100.000","100.000"],"quality":"More is better","isMain":true,"metricName":"Genome fraction (%)"},{"values":["1.001","1.004","1.010","1.030","1.003","1.009","1.004","1.004","1.004","1.004"],"quality":"Less is better","isMain":true,"metricName":"Duplication ratio"},{"values":[14390,14457,14539,14827,14443,14525,14453,14453,14453,9027],"quality":"More is better","isMain":true,"metricName":"Largest alignment"},{"values":[14390,14457,14539,14827,14443,14525,14453,14453,14453,14427],"quality":"More is better","isMain":true,"metricName":"Total aligned length"},{"values":[14398,14457,14539,14827,14443,14525,14453,14453,14453,9038],"quality":"More is better","isMain":false,"metricName":"NG50"},{"values":[14398,14457,14539,14827,14443,14525,14453,14453,14453,4713],"quality":"More is better","isMain":false,"metricName":"NG75"},{"values":[14390,14457,14539,14827,14443,14525,14453,14453,14453,9027],"quality":"More is better","isMain":false,"metricName":"NA50"},{"values":[14390,14457,14539,14827,14443,14525,14453,14453,14453,4713],"quality":"More is better","isMain":false,"metricName":"NA75"},{"values":[14390,14457,14539,14827,14443,14525,14453,14453,14453,9027],"quality":"More is better","isMain":true,"metricName":"NGA50"},{"values":[14390,14457,14539,14827,14443,14525,14453,14453,14453,4713],"quality":"More is better","isMain":false,"metricName":"NGA75"},{"values":[1,1,1,1,1,1,1,1,1,1],"quality":"Less is better","isMain":false,"metricName":"LG50"},{"values":[1,1,1,1,1,1,1,1,1,2],"quality":"Less is better","isMain":false,"metricName":"LG75"},{"values":[1,1,1,1,1,1,1,1,1,1],"quality":"Less is better","isMain":false,"metricName":"LA50"},{"values":[1,1,1,1,1,1,1,1,1,2],"quality":"Less is better","isMain":false,"metricName":"LA75"},{"values":[1,1,1,1,1,1,1,1,1,1],"quality":"Less is better","isMain":true,"metricName":"LGA50"},{"values":[1,1,1,1,1,1,1,1,1,2],"quality":"Less is better","isMain":false,"metricName":"LGA75"}]],["Misassemblies",[{"values":[0,0,0,0,0,0,0,0,0,0],"quality":"Less is better","isMain":true,"metricName":"# misassemblies"},{"values":[0,0,0,0,0,0,0,0,0,0],"quality":"Less is better","isMain":false,"metricName":" # relocations"},{"values":[0,0,0,0,0,0,0,0,0,0],"quality":"Less is better","isMain":false,"metricName":" # translocations"},{"values":[0,0,0,0,0,0,0,0,0,0],"quality":"Less is better","isMain":false,"metricName":" # inversions"},{"values":[0,0,0,0,0,0,0,0,0,0],"quality":"Less is better","isMain":false,"metricName":"# misassembled contigs"},{"values":[0,0,0,0,0,0,0,0,0,0],"quality":"Less is better","isMain":true,"metricName":"Misassembled contigs length"},{"values":[0,0,1,1,0,1,0,0,0,0],"quality":"Less is better","isMain":false,"metricName":"# local misassemblies"},{"values":[0,0,0,0,0,0,0,0,0,0],"quality":"Less is better","isMain":false,"metricName":"# unaligned mis. contigs"}]],["Unaligned",[{"values":[0,0,0,0,0,0,0,0,0,0],"quality":"Less is better","isMain":false,"metricName":"# fully unaligned contigs"},{"values":[0,0,0,0,0,0,0,0,0,0],"quality":"Less is better","isMain":false,"metricName":"Fully unaligned length"},{"values":[0,0,0,0,0,0,0,0,0,0],"quality":"Less is better","isMain":false,"metricName":"# partially unaligned contigs"},{"values":[0,0,0,0,0,0,0,0,0,0],"quality":"Less is better","isMain":false,"metricName":"Partially unaligned length"}]],["Mismatches",[{"values":[3,0,0,0,0,0,0,0,0,0],"quality":"Less is better","isMain":false,"metricName":"# mismatches"},{"values":[0,1,0,0,1,0,1,1,1,0],"quality":"Less is better","isMain":false,"metricName":"# indels"},{"values":[0,59,0,0,45,0,55,55,55,0],"quality":"Less is better","isMain":false,"metricName":"Indels length"},{"values":["20.85","0.00","0.00","0.00","0.00","0.00","0.00","0.00","0.00","0.00"],"quality":"Less is better","isMain":true,"metricName":"# mismatches per 100 kbp"},{"values":["0.00","6.95","0.00","0.00","6.95","0.00","6.95","6.95","6.95","0.00"],"quality":"Less is better","isMain":true,"metricName":"# indels per 100 kbp"},{"values":[0,0,0,0,0,0,0,0,0,0],"quality":"Less is better","isMain":false,"metricName":" # indels (<= 5 bp)"},{"values":[0,1,0,0,1,0,1,1,1,0],"quality":"Less is better","isMain":false,"metricName":" # indels (> 5 bp)"},{"values":[0,0,0,1,0,0,0,0,0,0],"quality":"Less is better","isMain":false,"metricName":"# N's"},{"values":["0.00","0.00","0.00","6.74","0.00","0.00","0.00","0.00","0.00","0.00"],"quality":"Less is better","isMain":true,"metricName":"# N's per 100 kbp"}]],["Statistics without reference",[{"values":[1,1,1,1,1,1,1,1,1,3],"quality":"Equal","isMain":true,"metricName":"# contigs"},{"values":[1,1,1,1,1,1,1,1,1,2],"quality":"Equal","isMain":false,"metricName":"# contigs (>= 1000 bp)"},{"values":[1,1,1,1,1,1,1,1,1,1],"quality":"Equal","isMain":false,"metricName":"# contigs (>= 5000 bp)"},{"values":[1,1,1,1,1,1,1,1,1,0],"quality":"Equal","isMain":false,"metricName":"# contigs (>= 10000 bp)"},{"values":[0,0,0,0,0,0,0,0,0,0],"quality":"Equal","isMain":false,"metricName":"# contigs (>= 25000 bp)"},{"values":[0,0,0,0,0,0,0,0,0,0],"quality":"Equal","isMain":false,"metricName":"# contigs (>= 50000 bp)"},{"values":[14398,14457,14539,14827,14443,14525,14453,14453,14453,9038],"quality":"More is better","isMain":true,"metricName":"Largest contig"},{"values":[14398,14457,14539,14827,14443,14525,14453,14453,14453,14449],"quality":"More is better","isMain":true,"metricName":"Total length"},{"values":[14398,14457,14539,14827,14443,14525,14453,14453,14453,13751],"quality":"More is better","isMain":true,"metricName":"Total length (>= 1000 bp)"},{"values":[14398,14457,14539,14827,14443,14525,14453,14453,14453,9038],"quality":"More is better","isMain":false,"metricName":"Total length (>= 5000 bp)"},{"values":[14398,14457,14539,14827,14443,14525,14453,14453,14453,0],"quality":"More is better","isMain":true,"metricName":"Total length (>= 10000 bp)"},{"values":[0,0,0,0,0,0,0,0,0,0],"quality":"More is better","isMain":false,"metricName":"Total length (>= 25000 bp)"},{"values":[0,0,0,0,0,0,0,0,0,0],"quality":"More is better","isMain":true,"metricName":"Total length (>= 50000 bp)"},{"values":[14398,14457,14539,14827,14443,14525,14453,14453,14453,9038],"quality":"More is better","isMain":false,"metricName":"N50"},{"values":[14398,14457,14539,14827,14443,14525,14453,14453,14453,4713],"quality":"More is better","isMain":false,"metricName":"N75"},{"values":[1,1,1,1,1,1,1,1,1,1],"quality":"Less is better","isMain":false,"metricName":"L50"},{"values":[1,1,1,1,1,1,1,1,1,2],"quality":"Less is better","isMain":false,"metricName":"L75"},{"values":["38.65","38.69","38.72","38.85","38.69","38.65","38.66","38.66","38.66","38.67"],"quality":"Equal","isMain":false,"metricName":"GC (%)"}]],["Predicted genes",[]],["Similarity statistics",[{"values":[0,0,0,0,0,0,0,0,0,0],"quality":"Equal","isMain":false,"metricName":"# similar correct contigs"},{"values":[0,0,0,0,0,0,0,0,0,0],"quality":"Equal","isMain":false,"metricName":"# similar misassembled blocks"}]],["Reference statistics",[{"values":[14398,14398,14398,14398,14398,14398,14398,14398,14398,14398],"quality":"Equal","isMain":false,"metricName":"Reference length"},{"values":[1,1,1,1,1,1,1,1,1,1],"quality":"Equal","isMain":false,"metricName":"Reference fragments"},{"values":["38.65","38.65","38.65","38.65","38.65","38.65","38.65","38.65","38.65","38.65"],"quality":"Equal","isMain":false,"metricName":"Reference GC (%)"}]]],"referenceName":"gi\_211956446\_ref\_NC","date":"18 July 2018, Wednesday, 16:55:40","order":[0,1,2,3,4,5,6,7,8,9],"assembliesNames":["CLC","IDBA\_UD","MEGAHIT","MIRA","Ray\_Meta","SPAdes","SPAdes\_meta","SPAdes\_sc","SPAdes\_sc\_careful","Velvet"]},{"assembliesWithNs":null,"minContig":500,"report":[["Genome statistics",[{"values":["97.040","96.917","96.940","96.744","83.024","100.000","99.074","4.788","91.763","99.681","97.647","97.110","77.066","100.000"],"quality":"More is better","isMain":true,"metricName":"Genome fraction (%)"},{"values":["1.020","1.006","1.025","1.026","1.034","1.354","1.110","1.000","1.049","1.018","1.019","1.015","1.001","1.465"],"quality":"Less is better","isMain":true,"metricName":"Duplication ratio"},{"values":[12841,9222,9877,6428,6987,14765,12065,747,7488,11878,11896,11003,5902,44253],"quality":"More is better","isMain":true,"metricName":"Largest alignment"},{"values":[39443,38886,39631,39559,34207,53675,43840,1909,38361,40180,39387,39155,30754,58392],"quality":"More is better","isMain":true,"metricName":"Total aligned length"},{"values":[8564,6847,4469,3468,2641,13307,11577,null,4246,10106,6100,6100,1601,67670],"quality":"More is better","isMain":false,"metricName":"NG50"},{"values":[5057,4784,2578,1538,879,7969,7763,null,1533,6100,5131,4784,521,67670],"quality":"More is better","isMain":false,"metricName":"NG75"},{"values":[8564,6847,4469,3468,3022,12995,11577,654,4246,9596,6100,6100,2490,44253],"quality":"More is better","isMain":false,"metricName":"NA50"},{"values":[5057,5787,2578,1538,1736,5998,6643,508,1663,6100,5131,3039,1177,12231],"quality":"More is better","isMain":false,"metricName":"NA75"},{"values":[8564,6847,4469,3468,2641,12995,11577,null,4246,9596,6100,6100,1601,44253],"quality":"More is better","isMain":true,"metricName":"NGA50"},{"values":[5057,4784,2578,1538,879,7969,7763,null,1533,6100,5131,3039,521,44253],"quality":"More is better","isMain":false,"metricName":"NGA75"},{"values":[2,3,3,5,5,2,2,null,4,2,3,3,6,1],"quality":"Less is better","isMain":false,"metricName":"LG50"},{"values":[4,5,7,9,11,3,3,null,8,4,4,4,17,1],"quality":"Less is better","isMain":false,"metricName":"LG75"},{"values":[2,3,3,5,4,2,2,2,4,2,3,3,4,1],"quality":"Less is better","isMain":false,"metricName":"LA50"},{"values":[4,4,7,9,8,4,4,3,7,4,4,5,8,2],"quality":"Less is better","isMain":false,"metricName":"LA75"},{"values":[2,3,3,5,5,2,2,null,4,2,3,3,6,1],"quality":"Less is better","isMain":true,"metricName":"LGA50"},{"values":[4,5,7,9,11,3,3,null,8,4,4,5,17,1],"quality":"Less is better","isMain":false,"metricName":"LGA75"}]],["Misassemblies",[{"values":[0,0,0,0,0,0,0,0,0,0,0,0,0,2],"quality":"Less is better","isMain":true,"metricName":"# misassemblies"},{"values":[0,0,0,0,0,0,0,0,0,0,0,0,0,2],"quality":"Less is better","isMain":false,"metricName":" # relocations"},{"values":[0,0,0,0,0,0,0,0,0,0,0,0,0,0],"quality":"Less is better","isMain":false,"metricName":" # translocations"},{"values":[0,0,0,0,0,0,0,0,0,0,0,0,0,0],"quality":"Less is better","isMain":false,"metricName":" # inversions"},{"values":[0,0,0,0,0,0,0,0,0,0,0,0,0,1],"quality":"Less is better","isMain":false,"metricName":"# misassembled contigs"},{"values":[0,0,0,0,0,0,0,0,0,0,0,0,0,67670],"quality":"Less is better","isMain":true,"metricName":"Misassembled contigs length"},{"values":[0,0,0,0,0,1,0,0,0,1,1,0,0,2],"quality":"Less is better","isMain":false,"metricName":"# local misassemblies"},{"values":[0,0,0,0,0,0,0,0,0,0,0,0,0,0],"quality":"Less is better","isMain":false,"metricName":"# unaligned mis. contigs"}]],["Unaligned",[{"values":[0,0,0,0,0,0,0,0,0,0,0,0,0,0],"quality":"Less is better","isMain":false,"metricName":"# fully unaligned contigs"},{"values":[0,0,0,0,0,0,0,0,0,0,0,0,0,0],"quality":"Less is better","isMain":false,"metricName":"Fully unaligned length"},{"values":[0,0,0,0,0,1,0,0,0,1,0,1,0,1],"quality":"Less is better","isMain":false,"metricName":"# partially unaligned contigs"},{"values":[0,0,0,0,0,532,0,0,0,510,0,1002,0,9278],"quality":"Less is better","isMain":false,"metricName":"Partially unaligned length"}]],["Mismatches",[{"values":[0,0,5,0,47,94,36,0,0,1,0,2,1,35],"quality":"Less is better","isMain":false,"metricName":"# mismatches"},{"values":[0,0,0,0,1,3,0,0,0,0,1,0,0,2],"quality":"Less is better","isMain":false,"metricName":"# indels"},{"values":[0,0,0,0,1,18,0,0,0,0,18,0,0,17],"quality":"Less is better","isMain":false,"metricName":"Indels length"},{"values":["0.00","0.00","12.94","0.00","142.00","235.78","91.14","0.00","0.00","2.52","0.00","5.17","3.25","87.79"],"quality":"Less is better","isMain":true,"metricName":"# mismatches per 100 kbp"},{"values":["0.00","0.00","0.00","0.00","3.02","7.53","0.00","0.00","0.00","0.00","2.57","0.00","0.00","5.02"],"quality":"Less is better","isMain":true,"metricName":"# indels per 100 kbp"},{"values":[0,0,0,0,1,1,0,0,0,0,0,0,0,0],"quality":"Less is better","isMain":false,"metricName":" # indels (<= 5 bp)"},{"values":[0,0,0,0,0,2,0,0,0,0,1,0,0,2],"quality":"Less is better","isMain":false,"metricName":" # indels (> 5 bp)"},{"values":[0,39,0,0,0,1,0,0,0,778,291,1142,0,147],"quality":"Less is better","isMain":false,"metricName":"# N's"},{"values":["0.00","100.29","0.00","0.00","0.00","1.83","0.00","0.00","0.00","1899.51","733.83","2833.82","0.00","217.23"],"quality":"Less is better","isMain":true,"metricName":"# N's per 100 kbp"}]],["Statistics without reference",[{"values":[10,11,15,19,16,9,5,3,17,7,12,12,18,1],"quality":"Equal","isMain":true,"metricName":"# contigs"},{"values":[5,6,11,13,10,8,5,0,12,6,6,7,9,1],"quality":"Equal","isMain":false,"metricName":"# contigs (>= 1000 bp)"},{"values":[4,4,2,1,1,5,5,0,2,4,4,3,1,1],"quality":"Equal","isMain":false,"metricName":"# contigs (>= 5000 bp)"},{"values":[1,0,0,0,0,2,2,0,0,2,1,1,0,1],"quality":"Equal","isMain":false,"metricName":"# contigs (>= 10000 bp)"},{"values":[0,0,0,0,0,0,0,0,0,0,0,0,0,1],"quality":"Equal","isMain":false,"metricName":"# contigs (>= 25000 bp)"},{"values":[0,0,0,0,0,0,0,0,0,0,0,0,0,1],"quality":"Equal","isMain":false,"metricName":"# contigs (>= 50000 bp)"},{"values":[12841,9222,9877,6428,6987,14765,12065,747,7488,12146,12164,12005,5902,67670],"quality":"More is better","isMain":true,"metricName":"Largest contig"},{"values":[39443,38886,39631,39559,34210,54521,43840,1909,38361,40958,39655,40299,30756,67670],"quality":"More is better","isMain":true,"metricName":"Total length"},{"values":[35498,35692,36917,35287,29368,53565,43840,0,34865,40338,35225,36947,24614,67670],"quality":"More is better","isMain":true,"metricName":"Total length (>= 1000 bp)"},{"values":[32564,29731,16081,6428,6987,47814,43840,0,12545,35966,31009,25719,5902,67670],"quality":"More is better","isMain":false,"metricName":"Total length (>= 5000 bp)"},{"values":[12841,0,0,0,0,28072,23642,0,0,22252,12164,12005,0,67670],"quality":"More is better","isMain":true,"metricName":"Total length (>= 10000 bp)"},{"values":[0,0,0,0,0,0,0,0,0,0,0,0,0,67670],"quality":"More is better","isMain":false,"metricName":"Total length (>= 25000 bp)"},{"values":[0,0,0,0,0,0,0,0,0,0,0,0,0,67670],"quality":"More is better","isMain":true,"metricName":"Total length (>= 50000 bp)"},{"values":[8564,6847,4469,3468,3022,13307,11577,654,4246,10106,6100,6100,2490,67670],"quality":"More is better","isMain":false,"metricName":"N50"},{"values":[5057,5787,2578,1538,1736,5998,6643,508,1663,6100,5131,4784,1177,67670],"quality":"More is better","isMain":false,"metricName":"N75"},{"values":[2,3,3,5,4,2,2,2,4,2,3,3,4,1],"quality":"Less is better","isMain":false,"metricName":"L50"},{"values":[4,4,7,9,8,4,4,3,7,4,4,4,8,1],"quality":"Less is better","isMain":false,"metricName":"L75"},{"values":["42.84","42.80","42.84","42.95","42.72","43.26","43.17","37.77","42.76","42.91","42.90","42.97","42.39","43.11"],"quality":"Equal","isMain":false,"metricName":"GC (%)"}]],["Predicted genes",[]],["Similarity statistics",[{"values":[0,0,0,0,0,0,0,0,0,0,0,0,0,0],"quality":"Equal","isMain":false,"metricName":"# similar correct contigs"},{"values":[0,0,0,0,0,0,0,0,0,0,0,0,0,0],"quality":"Equal","isMain":false,"metricName":"# similar misassembled blocks"}]],["Reference statistics",[{"values":[39867,39867,39867,39867,39867,39867,39867,39867,39867,39867,39867,39867,39867,39867],"quality":"Equal","isMain":false,"metricName":"Reference length"},{"values":[1,1,1,1,1,1,1,1,1,1,1,1,1,1],"quality":"Equal","isMain":false,"metricName":"Reference fragments"},{"values":["42.91","42.91","42.91","42.91","42.91","42.91","42.91","42.91","42.91","42.91","42.91","42.91","42.91","42.91"],"quality":"Equal","isMain":false,"metricName":"Reference GC (%)"}]]],"referenceName":"gi\_212499705\_ref\_NC","date":"18 July 2018, Wednesday, 16:55:58","order":[0,1,2,3,4,5,6,7,8,9,10,11,12,13],"assembliesNames":["ABySS\_127","ABySS\_63","CLC","IDBA\_UD","MEGAHIT","MIRA","Ray\_Meta","SOAPdenovo2","SPAdes","SPAdes\_meta","SPAdes\_sc","SPAdes\_sc\_careful","Velvet","Geneious"]},{"assembliesWithNs":null,"minContig":500,"report":[["Genome statistics",[{"values":["20.290","10.059","18.112","15.837","21.982","22.221","22.221","22.221"],"quality":"More is better","isMain":true,"metricName":"Genome fraction (%)"},{"values":["1.000","1.000","1.000","1.000","1.000","1.000","1.000","1.000"],"quality":"Less is better","isMain":true,"metricName":"Duplication ratio"},{"values":[2108,1355,1659,1442,2108,2108,2108,2108],"quality":"More is better","isMain":true,"metricName":"Largest alignment"},{"values":[25893,12837,23114,20211,28053,28357,28357,28357],"quality":"More is better","isMain":true,"metricName":"Total aligned length"},{"values":[877,830,698,832,819,832,832,832],"quality":"More is better","isMain":false,"metricName":"NA50"},{"values":[698,613,552,563,665,677,677,677],"quality":"More is better","isMain":false,"metricName":"NA75"},{"values":[null,null,null,null,null,null,null,null],"quality":"More is better","isMain":true,"metricName":"NGA50"},{"values":[11,6,13,10,14,13,13,13],"quality":"Less is better","isMain":false,"metricName":"LA50"},{"values":[20,11,22,18,24,22,22,22],"quality":"Less is better","isMain":false,"metricName":"LA75"}]],["Misassemblies",[{"values":[0,0,0,0,0,0,0,0],"quality":"Less is better","isMain":true,"metricName":"# misassemblies"},{"values":[0,0,0,0,0,0,0,0],"quality":"Less is better","isMain":false,"metricName":" # relocations"},{"values":[0,0,0,0,0,0,0,0],"quality":"Less is better","isMain":false,"metricName":" # translocations"},{"values":[0,0,0,0,0,0,0,0],"quality":"Less is better","isMain":false,"metricName":" # inversions"},{"values":[0,0,0,0,0,0,0,0],"quality":"Less is better","isMain":false,"metricName":"# misassembled contigs"},{"values":[0,0,0,0,0,0,0,0],"quality":"Less is better","isMain":true,"metricName":"Misassembled contigs length"},{"values":[0,0,0,0,0,0,0,0],"quality":"Less is better","isMain":false,"metricName":"# local misassemblies"},{"values":[0,0,0,0,0,0,0,0],"quality":"Less is better","isMain":false,"metricName":"# unaligned mis. contigs"}]],["Unaligned",[{"values":[0,0,0,0,0,0,0,0],"quality":"Less is better","isMain":false,"metricName":"# fully unaligned contigs"},{"values":[0,0,0,0,0,0,0,0],"quality":"Less is better","isMain":false,"metricName":"Fully unaligned length"},{"values":[0,0,0,0,1,0,0,0],"quality":"Less is better","isMain":false,"metricName":"# partially unaligned contigs"},{"values":[0,0,0,0,1671,0,0,0],"quality":"Less is better","isMain":false,"metricName":"Partially unaligned length"}]],["Mismatches",[{"values":[56,26,38,37,64,56,51,49],"quality":"Less is better","isMain":false,"metricName":"# mismatches"},{"values":[0,0,0,0,1,0,0,0],"quality":"Less is better","isMain":false,"metricName":"# indels"},{"values":[0,0,0,0,1,0,0,0],"quality":"Less is better","isMain":false,"metricName":"Indels length"},{"values":["216.27","202.54","164.40","183.07","228.15","197.48","179.85","172.80"],"quality":"Less is better","isMain":true,"metricName":"# mismatches per 100 kbp"},{"values":["0.00","0.00","0.00","0.00","3.56","0.00","0.00","0.00"],"quality":"Less is better","isMain":true,"metricName":"# indels per 100 kbp"},{"values":[0,0,0,0,1,0,0,0],"quality":"Less is better","isMain":false,"metricName":" # indels (<= 5 bp)"},{"values":[0,0,0,0,0,0,0,0],"quality":"Less is better","isMain":false,"metricName":" # indels (> 5 bp)"},{"values":[0,0,0,0,0,0,0,0],"quality":"Less is better","isMain":false,"metricName":"# N's"},{"values":["0.00","0.00","0.00","0.00","0.00","0.00","0.00","0.00"],"quality":"Less is better","isMain":true,"metricName":"# N's per 100 kbp"}]],["Statistics without reference",[{"values":[30,16,32,27,34,34,34,34],"quality":"Equal","isMain":true,"metricName":"# contigs"},{"values":[7,4,3,4,8,7,7,7],"quality":"Equal","isMain":false,"metricName":"# contigs (>= 1000 bp)"},{"values":[0,0,0,0,0,0,0,0],"quality":"Equal","isMain":false,"metricName":"# contigs (>= 5000 bp)"},{"values":[0,0,0,0,0,0,0,0],"quality":"Equal","isMain":false,"metricName":"# contigs (>= 10000 bp)"},{"values":[0,0,0,0,0,0,0,0],"quality":"Equal","isMain":false,"metricName":"# contigs (>= 25000 bp)"},{"values":[0,0,0,0,0,0,0,0],"quality":"Equal","isMain":false,"metricName":"# contigs (>= 50000 bp)"},{"values":[2108,1355,1659,1442,2108,2108,2108,2108],"quality":"More is better","isMain":true,"metricName":"Largest contig"},{"values":[25893,12842,23124,20211,29724,28357,28357,28357],"quality":"More is better","isMain":true,"metricName":"Total length"},{"values":[9504,4846,4202,4906,11377,9504,9504,9504],"quality":"More is better","isMain":true,"metricName":"Total length (>= 1000 bp)"},{"values":[0,0,0,0,0,0,0,0],"quality":"More is better","isMain":false,"metricName":"Total length (>= 5000 bp)"},{"values":[0,0,0,0,0,0,0,0],"quality":"More is better","isMain":true,"metricName":"Total length (>= 10000 bp)"},{"values":[0,0,0,0,0,0,0,0],"quality":"More is better","isMain":false,"metricName":"Total length (>= 25000 bp)"},{"values":[0,0,0,0,0,0,0,0],"quality":"More is better","isMain":true,"metricName":"Total length (>= 50000 bp)"},{"values":[877,830,698,832,877,832,832,832],"quality":"More is better","isMain":false,"metricName":"N50"},{"values":[698,613,552,563,678,677,677,677],"quality":"More is better","isMain":false,"metricName":"N75"},{"values":[11,6,13,10,12,13,13,13],"quality":"Less is better","isMain":false,"metricName":"L50"},{"values":[20,11,22,18,22,22,22,22],"quality":"Less is better","isMain":false,"metricName":"L75"},{"values":["41.70","41.52","41.53","41.62","41.34","41.66","41.65","41.65"],"quality":"Equal","isMain":false,"metricName":"GC (%)"}]],["Predicted genes",[]],["Similarity statistics",[{"values":[0,0,0,0,0,0,0,0],"quality":"Equal","isMain":false,"metricName":"# similar correct contigs"},{"values":[0,0,0,0,0,0,0,0],"quality":"Equal","isMain":false,"metricName":"# similar misassembled blocks"}]],["Reference statistics",[{"values":[127615,127615,127615,127615,127615,127615,127615,127615],"quality":"Equal","isMain":false,"metricName":"Reference length"},{"values":[1,1,1,1,1,1,1,1],"quality":"Equal","isMain":false,"metricName":"Reference fragments"},{"values":["41.63","41.63","41.63","41.63","41.63","41.63","41.63","41.63"],"quality":"Equal","isMain":false,"metricName":"Reference GC (%)"}]]],"referenceName":"gi\_213159268\_ref\_NC","date":"18 July 2018, Wednesday, 16:56:11","order":[0,1,2,3,4,5,6,7],"assembliesNames":["CLC","IDBA\_UD","MEGAHIT","SOAPdenovo2","SPAdes","SPAdes\_meta","SPAdes\_sc","SPAdes\_sc\_careful"]},{"assembliesWithNs":null,"minContig":500,"report":[["Genome statistics",[{"values":["100.000","100.000","100.000","100.000","100.000","26.604","100.000","100.000","100.000","100.000","100.000"],"quality":"More is better","isMain":true,"metricName":"Genome fraction (%)"},{"values":["1.000","1.002","1.003","1.007","1.011","1.000","1.003","1.001","1.001","1.001","1.001"],"quality":"Less is better","isMain":true,"metricName":"Duplication ratio"},{"values":[40287,40386,40428,40573,40717,1033,40414,40342,40342,40342,40317],"quality":"More is better","isMain":true,"metricName":"Largest alignment"},{"values":[40287,40386,40428,40573,40717,10718,40414,40342,40342,40342,40317],"quality":"More is better","isMain":true,"metricName":"Total aligned length"},{"values":[40287,40386,40428,40573,40717,null,40414,40342,40342,40342,40317],"quality":"More is better","isMain":false,"metricName":"NG50"},{"values":[40287,40386,40428,40573,40717,null,40414,40342,40342,40342,40317],"quality":"More is better","isMain":false,"metricName":"NG75"},{"values":[40287,40386,40428,40573,40717,611,40414,40342,40342,40342,40317],"quality":"More is better","isMain":false,"metricName":"NA50"},{"values":[40287,40386,40428,40573,40717,543,40414,40342,40342,40342,40317],"quality":"More is better","isMain":false,"metricName":"NA75"},{"values":[40287,40386,40428,40573,40717,null,40414,40342,40342,40342,40317],"quality":"More is better","isMain":true,"metricName":"NGA50"},{"values":[40287,40386,40428,40573,40717,null,40414,40342,40342,40342,40317],"quality":"More is better","isMain":false,"metricName":"NGA75"},{"values":[1,1,1,1,1,null,1,1,1,1,1],"quality":"Less is better","isMain":false,"metricName":"LG50"},{"values":[1,1,1,1,1,null,1,1,1,1,1],"quality":"Less is better","isMain":false,"metricName":"LG75"},{"values":[1,1,1,1,1,7,1,1,1,1,1],"quality":"Less is better","isMain":false,"metricName":"LA50"},{"values":[1,1,1,1,1,11,1,1,1,1,1],"quality":"Less is better","isMain":false,"metricName":"LA75"},{"values":[1,1,1,1,1,null,1,1,1,1,1],"quality":"Less is better","isMain":true,"metricName":"LGA50"},{"values":[1,1,1,1,1,null,1,1,1,1,1],"quality":"Less is better","isMain":false,"metricName":"LGA75"}]],["Misassemblies",[{"values":[0,0,0,0,0,0,0,0,0,0,0],"quality":"Less is better","isMain":true,"metricName":"# misassemblies"},{"values":[0,0,0,0,0,0,0,0,0,0,0],"quality":"Less is better","isMain":false,"metricName":" # relocations"},{"values":[0,0,0,0,0,0,0,0,0,0,0],"quality":"Less is better","isMain":false,"metricName":" # translocations"},{"values":[0,0,0,0,0,0,0,0,0,0,0],"quality":"Less is better","isMain":false,"metricName":" # inversions"},{"values":[0,0,0,0,0,0,0,0,0,0,0],"quality":"Less is better","isMain":false,"metricName":"# misassembled contigs"},{"values":[0,0,0,0,0,0,0,0,0,0,0],"quality":"Less is better","isMain":true,"metricName":"Misassembled contigs length"},{"values":[0,1,1,1,1,0,1,0,0,0,0],"quality":"Less is better","isMain":false,"metricName":"# local misassemblies"},{"values":[0,0,0,0,0,0,0,0,0,0,0],"quality":"Less is better","isMain":false,"metricName":"# unaligned mis. contigs"}]],["Unaligned",[{"values":[0,0,0,0,0,0,0,0,0,0,0],"quality":"Less is better","isMain":false,"metricName":"# fully unaligned contigs"},{"values":[0,0,0,0,0,0,0,0,0,0,0],"quality":"Less is better","isMain":false,"metricName":"Fully unaligned length"},{"values":[0,0,0,0,0,0,0,0,0,0,0],"quality":"Less is better","isMain":false,"metricName":"# partially unaligned contigs"},{"values":[0,0,0,0,0,0,0,0,0,0,0],"quality":"Less is better","isMain":false,"metricName":"Partially unaligned length"}]],["Mismatches",[{"values":[0,0,0,0,0,2,0,0,0,0,0],"quality":"Less is better","isMain":false,"metricName":"# mismatches"},{"values":[0,0,0,0,0,0,0,1,1,1,1],"quality":"Less is better","isMain":false,"metricName":"# indels"},{"values":[0,0,0,0,0,0,0,55,55,55,30],"quality":"Less is better","isMain":false,"metricName":"Indels length"},{"values":["0.00","0.00","0.00","0.00","0.00","18.66","0.00","0.00","0.00","0.00","0.00"],"quality":"Less is better","isMain":true,"metricName":"# mismatches per 100 kbp"},{"values":["0.00","0.00","0.00","0.00","0.00","0.00","0.00","2.48","2.48","2.48","2.48"],"quality":"Less is better","isMain":true,"metricName":"# indels per 100 kbp"},{"values":[0,0,0,0,0,0,0,0,0,0,0],"quality":"Less is better","isMain":false,"metricName":" # indels (<= 5 bp)"},{"values":[0,0,0,0,0,0,0,1,1,1,1],"quality":"Less is better","isMain":false,"metricName":" # indels (> 5 bp)"},{"values":[0,0,0,0,0,0,0,0,0,0,0],"quality":"Less is better","isMain":false,"metricName":"# N's"},{"values":["0.00","0.00","0.00","0.00","0.00","0.00","0.00","0.00","0.00","0.00","0.00"],"quality":"Less is better","isMain":true,"metricName":"# N's per 100 kbp"}]],["Statistics without reference",[{"values":[1,1,1,1,1,16,1,1,1,1,1],"quality":"Equal","isMain":true,"metricName":"# contigs"},{"values":[1,1,1,1,1,2,1,1,1,1,1],"quality":"Equal","isMain":false,"metricName":"# contigs (>= 1000 bp)"},{"values":[1,1,1,1,1,0,1,1,1,1,1],"quality":"Equal","isMain":false,"metricName":"# contigs (>= 5000 bp)"},{"values":[1,1,1,1,1,0,1,1,1,1,1],"quality":"Equal","isMain":false,"metricName":"# contigs (>= 10000 bp)"},{"values":[1,1,1,1,1,0,1,1,1,1,1],"quality":"Equal","isMain":false,"metricName":"# contigs (>= 25000 bp)"},{"values":[0,0,0,0,0,0,0,0,0,0,0],"quality":"Equal","isMain":false,"metricName":"# contigs (>= 50000 bp)"},{"values":[40287,40386,40428,40573,40717,1033,40414,40342,40342,40342,40317],"quality":"More is better","isMain":true,"metricName":"Largest contig"},{"values":[40287,40386,40428,40573,40717,10718,40414,40342,40342,40342,40317],"quality":"More is better","isMain":true,"metricName":"Total length"},{"values":[40287,40386,40428,40573,40717,2048,40414,40342,40342,40342,40317],"quality":"More is better","isMain":true,"metricName":"Total length (>= 1000 bp)"},{"values":[40287,40386,40428,40573,40717,0,40414,40342,40342,40342,40317],"quality":"More is better","isMain":false,"metricName":"Total length (>= 5000 bp)"},{"values":[40287,40386,40428,40573,40717,0,40414,40342,40342,40342,40317],"quality":"More is better","isMain":true,"metricName":"Total length (>= 10000 bp)"},{"values":[40287,40386,40428,40573,40717,0,40414,40342,40342,40342,40317],"quality":"More is better","isMain":false,"metricName":"Total length (>= 25000 bp)"},{"values":[0,0,0,0,0,0,0,0,0,0,0],"quality":"More is better","isMain":true,"metricName":"Total length (>= 50000 bp)"},{"values":[40287,40386,40428,40573,40717,611,40414,40342,40342,40342,40317],"quality":"More is better","isMain":false,"metricName":"N50"},{"values":[40287,40386,40428,40573,40717,543,40414,40342,40342,40342,40317],"quality":"More is better","isMain":false,"metricName":"N75"},{"values":[1,1,1,1,1,7,1,1,1,1,1],"quality":"Less is better","isMain":false,"metricName":"L50"},{"values":[1,1,1,1,1,11,1,1,1,1,1],"quality":"Less is better","isMain":false,"metricName":"L75"},{"values":["63.69","63.68","63.68","63.67","63.70","63.75","63.70","63.69","63.69","63.69","63.70"],"quality":"Equal","isMain":false,"metricName":"GC (%)"}]],["Predicted genes",[]],["Similarity statistics",[{"values":[0,0,0,0,0,0,0,0,0,0,0],"quality":"Equal","isMain":false,"metricName":"# similar correct contigs"},{"values":[0,0,0,0,0,0,0,0,0,0,0],"quality":"Equal","isMain":false,"metricName":"# similar misassembled blocks"}]],["Reference statistics",[{"values":[40287,40287,40287,40287,40287,40287,40287,40287,40287,40287,40287],"quality":"Equal","isMain":false,"metricName":"Reference length"},{"values":[1,1,1,1,1,1,1,1,1,1,1],"quality":"Equal","isMain":false,"metricName":"Reference fragments"},{"values":["63.69","63.69","63.69","63.69","63.69","63.69","63.69","63.69","63.69","63.69","63.69"],"quality":"Equal","isMain":false,"metricName":"Reference GC (%)"}]]],"referenceName":"gi\_213163898\_ref\_NC","date":"18 July 2018, Wednesday, 16:56:26","order":[0,1,2,3,4,5,6,7,8,9,10],"assembliesNames":["CLC","IDBA\_UD","MEGAHIT","MIRA","Ray\_Meta","SOAPdenovo2","SPAdes","SPAdes\_meta","SPAdes\_sc","SPAdes\_sc\_careful","Velvet"]},{"assembliesWithNs":null,"minContig":500,"report":[["Genome statistics",[{"values":["0.755","12.715","6.439","14.477","1.182","8.869","17.501","17.705","17.233","17.856"],"quality":"More is better","isMain":true,"metricName":"Genome fraction (%)"},{"values":["1.000","1.000","1.079","1.007","1.000","1.000","1.020","1.001","1.034","1.036"],"quality":"Less is better","isMain":true,"metricName":"Duplication ratio"},{"values":[598,1840,1071,1189,994,1289,1840,1840,1840,1840],"quality":"More is better","isMain":true,"metricName":"Largest alignment"},{"values":[1129,19022,9635,21657,1768,13268,26181,26486,25781,26716],"quality":"More is better","isMain":true,"metricName":"Total aligned length"},{"values":[598,793,660,745,994,706,793,805,798,798],"quality":"More is better","isMain":false,"metricName":"NA50"},{"values":[531,650,544,644,774,576,635,695,658,658],"quality":"More is better","isMain":false,"metricName":"NA75"},{"values":[null,null,null,null,null,null,null,null,null,null],"quality":"More is better","isMain":true,"metricName":"NGA50"},{"values":[1,10,6,13,1,8,15,13,14,15],"quality":"Less is better","isMain":false,"metricName":"LA50"},{"values":[2,18,11,21,2,13,25,22,24,25],"quality":"Less is better","isMain":false,"metricName":"LA75"}]],["Misassemblies",[{"values":[0,0,0,0,0,0,0,0,0,0],"quality":"Less is better","isMain":true,"metricName":"# misassemblies"},{"values":[0,0,0,0,0,0,0,0,0,0],"quality":"Less is better","isMain":false,"metricName":" # relocations"},{"values":[0,0,0,0,0,0,0,0,0,0],"quality":"Less is better","isMain":false,"metricName":" # translocations"},{"values":[0,0,0,0,0,0,0,0,0,0],"quality":"Less is better","isMain":false,"metricName":" # inversions"},{"values":[0,0,0,0,0,0,0,0,0,0],"quality":"Less is better","isMain":false,"metricName":"# misassembled contigs"},{"values":[0,0,0,0,0,0,0,0,0,0],"quality":"Less is better","isMain":true,"metricName":"Misassembled contigs length"},{"values":[0,0,0,0,0,0,0,0,0,0],"quality":"Less is better","isMain":false,"metricName":"# local misassemblies"},{"values":[0,0,0,0,0,0,0,0,0,0],"quality":"Less is better","isMain":false,"metricName":"# unaligned mis. contigs"}]],["Unaligned",[{"values":[0,0,0,0,0,0,0,0,0,0],"quality":"Less is better","isMain":false,"metricName":"# fully unaligned contigs"},{"values":[0,0,0,0,0,0,0,0,0,0],"quality":"Less is better","isMain":false,"metricName":"Fully unaligned length"},{"values":[0,1,0,0,0,0,3,0,2,2],"quality":"Less is better","isMain":false,"metricName":"# partially unaligned contigs"},{"values":[0,2071,0,0,0,0,2816,0,2006,2006],"quality":"Less is better","isMain":false,"metricName":"Partially unaligned length"}]],["Mismatches",[{"values":[0,41,31,61,0,20,100,47,81,117],"quality":"Less is better","isMain":false,"metricName":"# mismatches"},{"values":[0,0,3,0,0,0,4,0,4,11],"quality":"Less is better","isMain":false,"metricName":"# indels"},{"values":[0,0,3,0,0,0,4,0,4,11],"quality":"Less is better","isMain":false,"metricName":"Indels length"},{"values":["0.00","215.54","321.84","281.66","0.00","150.74","381.96","177.45","314.18","437.99"],"quality":"Less is better","isMain":true,"metricName":"# mismatches per 100 kbp"},{"values":["0.00","0.00","31.15","0.00","0.00","0.00","15.28","0.00","15.52","41.18"],"quality":"Less is better","isMain":true,"metricName":"# indels per 100 kbp"},{"values":[0,0,3,0,0,0,4,0,4,11],"quality":"Less is better","isMain":false,"metricName":" # indels (<= 5 bp)"},{"values":[0,0,0,0,0,0,0,0,0,0],"quality":"Less is better","isMain":false,"metricName":" # indels (> 5 bp)"},{"values":[0,0,0,0,6,0,0,0,0,0],"quality":"Less is better","isMain":false,"metricName":"# N's"},{"values":["0.00","0.00","0.00","0.00","339.37","0.00","0.00","0.00","0.00","0.00"],"quality":"Less is better","isMain":true,"metricName":"# N's per 100 kbp"}]],["Statistics without reference",[{"values":[2,23,15,30,2,19,33,32,33,34],"quality":"Equal","isMain":true,"metricName":"# contigs"},{"values":[0,6,2,1,0,1,8,5,7,8],"quality":"Equal","isMain":false,"metricName":"# contigs (>= 1000 bp)"},{"values":[0,0,0,0,0,0,0,0,0,0],"quality":"Equal","isMain":false,"metricName":"# contigs (>= 5000 bp)"},{"values":[0,0,0,0,0,0,0,0,0,0],"quality":"Equal","isMain":false,"metricName":"# contigs (>= 10000 bp)"},{"values":[0,0,0,0,0,0,0,0,0,0],"quality":"Equal","isMain":false,"metricName":"# contigs (>= 25000 bp)"},{"values":[0,0,0,0,0,0,0,0,0,0],"quality":"Equal","isMain":false,"metricName":"# contigs (>= 50000 bp)"},{"values":[598,2532,1071,1189,994,1289,1840,1840,1840,1840],"quality":"More is better","isMain":true,"metricName":"Largest contig"},{"values":[1129,21093,10397,21812,1768,13268,29510,26513,28675,29679],"quality":"More is better","isMain":true,"metricName":"Total length"},{"values":[0,9211,2114,1189,0,1289,11314,6679,9593,10596],"quality":"More is better","isMain":true,"metricName":"Total length (>= 1000 bp)"},{"values":[0,0,0,0,0,0,0,0,0,0],"quality":"More is better","isMain":false,"metricName":"Total length (>= 5000 bp)"},{"values":[0,0,0,0,0,0,0,0,0,0],"quality":"More is better","isMain":true,"metricName":"Total length (>= 10000 bp)"},{"values":[0,0,0,0,0,0,0,0,0,0],"quality":"More is better","isMain":false,"metricName":"Total length (>= 25000 bp)"},{"values":[0,0,0,0,0,0,0,0,0,0],"quality":"More is better","isMain":true,"metricName":"Total length (>= 50000 bp)"},{"values":[598,852,667,745,994,706,822,805,818,822],"quality":"More is better","isMain":false,"metricName":"N50"},{"values":[531,695,560,650,774,576,721,695,703,703],"quality":"More is better","isMain":false,"metricName":"N75"},{"values":[1,8,6,13,1,8,12,13,13,13],"quality":"Less is better","isMain":false,"metricName":"L50"},{"values":[2,15,11,21,2,13,22,22,22,23],"quality":"Less is better","isMain":false,"metricName":"L75"},{"values":["28.26","25.71","24.92","25.03","23.84","24.62","25.76","24.98","25.22","25.30"],"quality":"Equal","isMain":false,"metricName":"GC (%)"}]],["Predicted genes",[]],["Similarity statistics",[{"values":[0,0,0,0,0,0,0,0,0,0],"quality":"Equal","isMain":false,"metricName":"# similar correct contigs"},{"values":[0,0,0,0,0,0,0,0,0,0],"quality":"Equal","isMain":false,"metricName":"# similar misassembled blocks"}]],["Reference statistics",[{"values":[149599,149599,149599,149599,149599,149599,149599,149599,149599,149599],"quality":"Equal","isMain":false,"metricName":"Reference length"},{"values":[1,1,1,1,1,1,1,1,1,1],"quality":"Equal","isMain":false,"metricName":"Reference fragments"},{"values":["25.31","25.31","25.31","25.31","25.31","25.31","25.31","25.31","25.31","25.31"],"quality":"Equal","isMain":false,"metricName":"Reference GC (%)"}]]],"referenceName":"gi\_21426072\_ref\_NC","date":"18 July 2018, Wednesday, 16:56:39","order":[0,1,2,3,4,5,6,7,8,9],"assembliesNames":["ABySS\_127","CLC","IDBA\_UD","MEGAHIT","MIRA","SOAPdenovo2","SPAdes","SPAdes\_meta","SPAdes\_sc","SPAdes\_sc\_careful"]},{"assembliesWithNs":null,"minContig":500,"report":[["Genome statistics",[{"values":["40.166","17.817","65.049","36.691","57.183","100.000","88.082","75.382","97.089","55.584","69.839","21.496","100.000"],"quality":"More is better","isMain":true,"metricName":"Genome fraction (%)"},{"values":["1.024","1.000","1.036","1.018","1.042","2.573","1.117","1.033","1.022","1.033","1.020","1.002","1.780"],"quality":"Less is better","isMain":true,"metricName":"Duplication ratio"},{"values":[3311,3790,4452,2656,2660,37293,13994,6278,25239,6515,6955,2275,37293],"quality":"More is better","isMain":true,"metricName":"Largest alignment"},{"values":[18645,8079,29970,16934,27009,115943,44622,35197,44245,26045,32314,9763,80505],"quality":"More is better","isMain":true,"metricName":"Total aligned length"},{"values":[null,null,1011,null,924,73503,11878,2019,25967,1014,1292,null,73504],"quality":"More is better","isMain":false,"metricName":"NG50"},{"values":[null,null,null,null,null,73503,2579,788,16063,null,null,null,73504],"quality":"More is better","isMain":false,"metricName":"NG75"},{"values":[1798,2181,1808,1186,1508,4210,11878,2592,25239,3435,2051,1390,6013],"quality":"More is better","isMain":false,"metricName":"NA50"},{"values":[789,2108,1011,690,1008,1625,7883,1592,16063,1347,1006,1188,null],"quality":"More is better","isMain":false,"metricName":"NA75"},{"values":[null,null,1011,null,924,37293,11878,2019,25239,1014,1023,null,37293],"quality":"More is better","isMain":true,"metricName":"NGA50"},{"values":[null,null,null,null,null,37293,2579,688,16063,null,null,null,37293],"quality":"More is better","isMain":false,"metricName":"NGA75"},{"values":[null,null,11,null,15,1,2,8,1,8,8,null,1],"quality":"Less is better","isMain":false,"metricName":"LG50"},{"values":[null,null,null,null,null,1,4,16,2,null,null,null,1],"quality":"Less is better","isMain":false,"metricName":"LG75"},{"values":[4,2,6,5,7,3,2,5,1,3,5,3,3],"quality":"Less is better","isMain":false,"metricName":"LA50"},{"values":[10,3,11,10,13,17,3,10,2,6,11,5,null],"quality":"Less is better","isMain":false,"metricName":"LA75"},{"values":[null,null,11,null,15,1,2,8,1,8,9,null,1],"quality":"Less is better","isMain":true,"metricName":"LGA50"},{"values":[null,null,null,null,null,1,4,17,2,null,null,null,1],"quality":"Less is better","isMain":false,"metricName":"LGA75"}]],["Misassemblies",[{"values":[0,0,0,0,0,3,0,0,0,0,0,0,1],"quality":"Less is better","isMain":true,"metricName":"# misassemblies"},{"values":[0,0,0,0,0,3,0,0,0,0,0,0,1],"quality":"Less is better","isMain":false,"metricName":" # relocations"},{"values":[0,0,0,0,0,0,0,0,0,0,0,0,0],"quality":"Less is better","isMain":false,"metricName":" # translocations"},{"values":[0,0,0,0,0,0,0,0,0,0,0,0,0],"quality":"Less is better","isMain":false,"metricName":" # inversions"},{"values":[0,0,0,0,0,3,0,0,0,0,0,0,1],"quality":"Less is better","isMain":false,"metricName":"# misassembled contigs"},{"values":[0,0,0,0,0,87814,0,0,0,0,0,0,73504],"quality":"Less is better","isMain":true,"metricName":"Misassembled contigs length"},{"values":[1,0,0,0,0,1,0,0,0,0,0,0,1],"quality":"Less is better","isMain":false,"metricName":"# local misassemblies"},{"values":[0,0,0,0,0,0,0,0,0,0,0,0,1],"quality":"Less is better","isMain":false,"metricName":"# unaligned mis. contigs"}]],["Unaligned",[{"values":[0,0,0,0,0,0,0,0,0,0,0,0,0],"quality":"Less is better","isMain":false,"metricName":"# fully unaligned contigs"},{"values":[0,0,0,0,0,0,0,0,0,0,0,0,0],"quality":"Less is better","isMain":false,"metricName":"Fully unaligned length"},{"values":[1,0,0,0,0,3,0,0,0,0,1,0,5],"quality":"Less is better","isMain":false,"metricName":"# partially unaligned contigs"},{"values":[2092,0,0,0,0,15627,0,0,0,0,940,0,44089],"quality":"Less is better","isMain":false,"metricName":"Partially unaligned length"}]],["Mismatches",[{"values":[7,0,20,0,31,131,125,9,328,198,50,17,114],"quality":"Less is better","isMain":false,"metricName":"# mismatches"},{"values":[2,0,0,0,0,2,3,0,2,1,2,0,3],"quality":"Less is better","isMain":false,"metricName":"# indels"},{"values":[2,0,0,0,0,2,4,0,4,16,19,0,3],"quality":"Less is better","isMain":false,"metricName":"Indels length"},{"values":["38.43","0.00","67.81","0.00","119.56","288.90","312.97","26.33","745.05","785.59","157.89","174.41","251.41"],"quality":"Less is better","isMain":true,"metricName":"# mismatches per 100 kbp"},{"values":["10.98","0.00","0.00","0.00","0.00","4.41","7.51","0.00","4.54","3.97","6.32","0.00","6.62"],"quality":"Less is better","isMain":true,"metricName":"# indels per 100 kbp"},{"values":[2,0,0,0,0,2,3,0,2,0,1,0,3],"quality":"Less is better","isMain":false,"metricName":" # indels (<= 5 bp)"},{"values":[0,0,0,0,0,0,0,0,0,1,1,0,0],"quality":"Less is better","isMain":false,"metricName":" # indels (> 5 bp)"},{"values":[51,105,0,0,0,40,0,125,728,0,937,0,574],"quality":"Less is better","isMain":false,"metricName":"# N's"},{"values":["245.94","1299.67","0.00","0.00","0.00","30.24","0.00","353.89","1618.75","0.00","2817.71","0.00","460.00"],"quality":"Less is better","isMain":true,"metricName":"# N's per 100 kbp"}]],["Statistics without reference",[{"values":[14,3,21,16,21,29,9,18,5,13,21,8,6],"quality":"Equal","isMain":true,"metricName":"# contigs"},{"values":[5,3,11,6,13,25,9,13,3,8,11,5,6],"quality":"Equal","isMain":false,"metricName":"# contigs (>= 1000 bp)"},{"values":[1,0,0,0,0,3,3,1,2,2,1,0,3],"quality":"Equal","isMain":false,"metricName":"# contigs (>= 5000 bp)"},{"values":[0,0,0,0,0,1,2,0,2,0,0,0,3],"quality":"Equal","isMain":false,"metricName":"# contigs (>= 10000 bp)"},{"values":[0,0,0,0,0,1,0,0,1,0,0,0,2],"quality":"Equal","isMain":false,"metricName":"# contigs (>= 25000 bp)"},{"values":[0,0,0,0,0,1,0,0,0,0,0,0,1],"quality":"Equal","isMain":false,"metricName":"# contigs (>= 50000 bp)"},{"values":[5403,3790,4452,2656,2660,73503,13994,6278,25967,6515,7895,2277,73504],"quality":"More is better","isMain":true,"metricName":"Largest contig"},{"values":[20737,8079,30565,16934,27009,132285,44622,35322,44973,26045,33254,9765,124782],"quality":"More is better","isMain":true,"metricName":"Total length"},{"values":[14176,8079,23456,10302,20977,129372,44622,31545,43719,22797,26295,7852,124782],"quality":"More is better","isMain":true,"metricName":"Total length (>= 1000 bp)"},{"values":[5403,0,0,0,0,87814,33755,6278,42030,11642,7895,0,114341],"quality":"More is better","isMain":false,"metricName":"Total length (>= 5000 bp)"},{"values":[0,0,0,0,0,73503,25872,0,42030,0,0,0,114341],"quality":"More is better","isMain":true,"metricName":"Total length (>= 10000 bp)"},{"values":[0,0,0,0,0,73503,0,0,25967,0,0,0,102099],"quality":"More is better","isMain":false,"metricName":"Total length (>= 25000 bp)"},{"values":[0,0,0,0,0,73503,0,0,0,0,0,0,73504],"quality":"More is better","isMain":true,"metricName":"Total length (>= 50000 bp)"},{"values":[2190,2181,1808,1186,1508,73503,11878,2712,25967,3435,2437,1390,73504],"quality":"More is better","isMain":false,"metricName":"N50"},{"values":[836,2108,1011,690,1008,2605,7883,1592,16063,1347,1014,1188,28595],"quality":"More is better","isMain":false,"metricName":"N75"},{"values":[3,2,6,5,7,1,2,5,1,3,4,3,1],"quality":"Less is better","isMain":false,"metricName":"L50"},{"values":[7,3,11,10,13,7,3,10,2,6,10,5,2],"quality":"Less is better","isMain":false,"metricName":"L75"},{"values":["33.05","30.89","33.28","32.23","32.59","33.26","33.21","33.11","33.32","32.15","33.07","32.78","33.25"],"quality":"Equal","isMain":false,"metricName":"GC (%)"}]],["Predicted genes",[]],["Similarity statistics",[{"values":[0,0,0,0,0,0,0,0,0,0,0,0,0],"quality":"Equal","isMain":false,"metricName":"# similar correct contigs"},{"values":[0,0,0,0,0,0,0,0,0,0,0,0,0],"quality":"Equal","isMain":false,"metricName":"# similar misassembled blocks"}]],["Reference statistics",[{"values":[45344,45344,45344,45344,45344,45344,45344,45344,45344,45344,45344,45344,45344],"quality":"Equal","isMain":false,"metricName":"Reference length"},{"values":[1,1,1,1,1,1,1,1,1,1,1,1,1],"quality":"Equal","isMain":false,"metricName":"Reference fragments"},{"values":["33.25","33.25","33.25","33.25","33.25","33.25","33.25","33.25","33.25","33.25","33.25","33.25","33.25"],"quality":"Equal","isMain":false,"metricName":"Reference GC (%)"}]]],"referenceName":"gi\_215401108\_ref\_NC","date":"18 July 2018, Wednesday, 16:56:57","order":[0,1,2,3,4,5,6,7,8,9,10,11,12],"assembliesNames":["ABySS\_127","ABySS\_63","CLC","IDBA\_UD","MEGAHIT","MIRA","Ray\_Meta","SPAdes","SPAdes\_meta","SPAdes\_sc","SPAdes\_sc\_careful","Velvet","Geneious"]},{"assembliesWithNs":null,"minContig":500,"report":[["Genome statistics",[{"values":["84.452","26.718","71.206","62.545","80.428","94.787","12.686","4.235","87.892","29.293","53.986","53.374","8.670","96.833"],"quality":"More is better","isMain":true,"metricName":"Genome fraction (%)"},{"values":["1.016","1.000","1.016","1.021","1.026","1.302","1.030","1.527","1.003","1.044","1.014","1.012","1.000","1.308"],"quality":"Less is better","isMain":true,"metricName":"Duplication ratio"},{"values":[5712,2178,3773,4609,3413,9917,1052,686,10974,6379,3817,3817,1370,17887],"quality":"More is better","isMain":true,"metricName":"Largest alignment"},{"values":[36474,11362,30722,27155,35085,52474,5559,1801,37504,12456,23266,22967,3687,53744],"quality":"More is better","isMain":true,"metricName":"Total aligned length"},{"values":[3116,null,1644,673,1078,28212,null,null,5243,null,783,697,null,42152],"quality":"More is better","isMain":false,"metricName":"NG50"},{"values":[701,null,null,null,657,10133,null,null,4217,null,null,null,null,42152],"quality":"More is better","isMain":false,"metricName":"NG75"},{"values":[3910,1224,3105,1561,1379,1846,632,null,8373,2981,1632,1864,658,null],"quality":"More is better","isMain":false,"metricName":"NA50"},{"values":[986,669,1209,775,960,null,540,null,4300,1357,1250,868,547,null],"quality":"More is better","isMain":false,"metricName":"NA75"},{"values":[3116,null,1644,673,1078,5151,null,null,5243,null,783,697,null,14978],"quality":"More is better","isMain":true,"metricName":"NGA50"},{"values":[701,null,null,null,657,4694,null,null,4217,null,null,null,null,14978],"quality":"More is better","isMain":false,"metricName":"NGA75"},{"values":[5,null,8,14,11,1,null,null,3,null,13,13,null,1],"quality":"Less is better","isMain":false,"metricName":"LG50"},{"values":[15,null,null,null,23,2,null,null,5,null,null,null,null,1],"quality":"Less is better","isMain":false,"metricName":"LG75"},{"values":[4,4,5,5,7,10,4,null,2,2,5,4,2,null],"quality":"Less is better","isMain":false,"metricName":"LA50"},{"values":[10,8,9,12,15,null,6,null,4,3,9,8,4,null],"quality":"Less is better","isMain":false,"metricName":"LA75"},{"values":[5,null,8,14,11,3,null,null,3,null,13,13,null,2],"quality":"Less is better","isMain":true,"metricName":"LGA50"},{"values":[15,null,null,null,23,5,null,null,5,null,null,null,null,2],"quality":"Less is better","isMain":false,"metricName":"LGA75"}]],["Misassemblies",[{"values":[0,0,0,0,0,0,0,0,0,0,0,0,0,0],"quality":"Less is better","isMain":true,"metricName":"# misassemblies"},{"values":[0,0,0,0,0,0,0,0,0,0,0,0,0,0],"quality":"Less is better","isMain":false,"metricName":" # relocations"},{"values":[0,0,0,0,0,0,0,0,0,0,0,0,0,0],"quality":"Less is better","isMain":false,"metricName":" # translocations"},{"values":[0,0,0,0,0,0,0,0,0,0,0,0,0,0],"quality":"Less is better","isMain":false,"metricName":" # inversions"},{"values":[0,0,0,0,0,0,0,0,0,0,0,0,0,0],"quality":"Less is better","isMain":false,"metricName":"# misassembled contigs"},{"values":[0,0,0,0,0,0,0,0,0,0,0,0,0,0],"quality":"Less is better","isMain":true,"metricName":"Misassembled contigs length"},{"values":[0,0,0,0,0,0,0,0,0,1,0,0,0,1],"quality":"Less is better","isMain":false,"metricName":"# local misassemblies"},{"values":[0,0,0,0,0,2,0,3,0,0,0,0,0,4],"quality":"Less is better","isMain":false,"metricName":"# unaligned mis. contigs"}]],["Unaligned",[{"values":[0,0,0,0,0,0,0,0,0,0,0,0,0,0],"quality":"Less is better","isMain":false,"metricName":"# fully unaligned contigs"},{"values":[0,0,0,0,0,0,0,0,0,0,0,0,0,0],"quality":"Less is better","isMain":false,"metricName":"Fully unaligned length"},{"values":[0,0,0,0,0,7,0,3,0,0,0,0,0,6],"quality":"Less is better","isMain":false,"metricName":"# partially unaligned contigs"},{"values":[0,0,0,0,0,38953,0,2440,0,0,0,0,0,92797],"quality":"Less is better","isMain":false,"metricName":"Partially unaligned length"}]],["Mismatches",[{"values":[1,8,61,4,29,221,5,0,16,107,226,185,0,263],"quality":"Less is better","isMain":false,"metricName":"# mismatches"},{"values":[0,0,0,0,0,7,0,0,0,1,4,4,0,11],"quality":"Less is better","isMain":false,"metricName":"# indels"},{"values":[0,0,0,0,0,7,0,0,0,1,8,8,0,13],"quality":"Less is better","isMain":false,"metricName":"Indels length"},{"values":["2.78","70.41","201.45","15.04","84.79","548.26","92.68","0.00","42.81","858.95","984.41","815.05","0.00","638.68"],"quality":"Less is better","isMain":true,"metricName":"# mismatches per 100 kbp"},{"values":["0.00","0.00","0.00","0.00","0.00","17.37","0.00","0.00","0.00","8.03","17.42","17.62","0.00","26.71"],"quality":"Less is better","isMain":true,"metricName":"# indels per 100 kbp"},{"values":[0,0,0,0,0,7,0,0,0,1,4,4,0,11],"quality":"Less is better","isMain":false,"metricName":" # indels (<= 5 bp)"},{"values":[0,0,0,0,0,0,0,0,0,0,0,0,0,0],"quality":"Less is better","isMain":false,"metricName":" # indels (> 5 bp)"},{"values":[0,0,0,0,0,24,0,1390,0,0,0,0,0,1221],"quality":"Less is better","isMain":false,"metricName":"# N's"},{"values":["0.00","0.00","0.00","0.00","0.00","26.25","0.00","26782.27","0.00","0.00","0.00","0.00","0.00","832.53"],"quality":"Less is better","isMain":true,"metricName":"# N's per 100 kbp"}]],["Statistics without reference",[{"values":[22,12,19,23,28,16,8,6,9,5,16,15,5,7],"quality":"Equal","isMain":true,"metricName":"# contigs"},{"values":[9,4,10,7,14,13,2,2,6,4,10,7,1,7],"quality":"Equal","isMain":false,"metricName":"# contigs (>= 1000 bp)"},{"values":[1,0,0,0,0,7,0,0,3,1,0,0,0,6],"quality":"Equal","isMain":false,"metricName":"# contigs (>= 5000 bp)"},{"values":[0,0,0,0,0,2,0,0,1,0,0,0,0,5],"quality":"Equal","isMain":false,"metricName":"# contigs (>= 10000 bp)"},{"values":[0,0,0,0,0,1,0,0,0,0,0,0,0,2],"quality":"Equal","isMain":false,"metricName":"# contigs (>= 25000 bp)"},{"values":[0,0,0,0,0,0,0,0,0,0,0,0,0,0],"quality":"Equal","isMain":false,"metricName":"# contigs (>= 50000 bp)"},{"values":[5712,2178,3773,4609,3413,28212,1052,1452,10974,6387,3817,3817,1370,42152],"quality":"More is better","isMain":true,"metricName":"Largest contig"},{"values":[36474,11362,30780,27155,35085,91435,5559,5190,37504,13006,23269,22970,3687,146662],"quality":"More is better","isMain":true,"metricName":"Total length"},{"values":[27157,6158,24605,16146,25455,89179,2097,2603,35227,12430,18803,17074,1370,146662],"quality":"More is better","isMain":true,"metricName":"Total length (>= 1000 bp)"},{"values":[5712,0,0,0,0,73589,0,0,24590,6387,0,0,0,144807],"quality":"More is better","isMain":false,"metricName":"Total length (>= 5000 bp)"},{"values":[0,0,0,0,0,38345,0,0,10974,0,0,0,0,138673],"quality":"More is better","isMain":true,"metricName":"Total length (>= 10000 bp)"},{"values":[0,0,0,0,0,28212,0,0,0,0,0,0,0,81999],"quality":"More is better","isMain":false,"metricName":"Total length (>= 25000 bp)"},{"values":[0,0,0,0,0,0,0,0,0,0,0,0,0,0],"quality":"More is better","isMain":true,"metricName":"Total length (>= 50000 bp)"},{"values":[3910,1224,3105,1561,1379,9917,632,1151,8373,3471,1632,1864,658,39847],"quality":"More is better","isMain":false,"metricName":"N50"},{"values":[986,669,1209,775,960,5158,540,686,4300,3471,1250,868,547,18008],"quality":"More is better","isMain":false,"metricName":"N75"},{"values":[4,4,5,5,7,3,4,2,2,2,5,4,2,2],"quality":"Less is better","isMain":false,"metricName":"L50"},{"values":[10,8,9,12,15,7,6,4,4,2,9,8,4,4],"quality":"Less is better","isMain":false,"metricName":"L75"},{"values":["34.83","35.14","34.98","34.88","34.67","34.88","31.89","34.97","34.79","34.26","34.84","34.84","33.60","34.72"],"quality":"Equal","isMain":false,"metricName":"GC (%)"}]],["Predicted genes",[]],["Similarity statistics",[{"values":[0,0,0,0,0,0,0,0,0,0,0,0,0,0],"quality":"Equal","isMain":false,"metricName":"# similar correct contigs"},{"values":[0,0,0,0,0,0,0,0,0,0,0,0,0,0],"quality":"Equal","isMain":false,"metricName":"# similar misassembled blocks"}]],["Reference statistics",[{"values":[42526,42526,42526,42526,42526,42526,42526,42526,42526,42526,42526,42526,42526,42526],"quality":"Equal","isMain":false,"metricName":"Reference length"},{"values":[1,1,1,1,1,1,1,1,1,1,1,1,1,1],"quality":"Equal","isMain":false,"metricName":"Reference fragments"},{"values":["34.91","34.91","34.91","34.91","34.91","34.91","34.91","34.91","34.91","34.91","34.91","34.91","34.91","34.91"],"quality":"Equal","isMain":false,"metricName":"Reference GC (%)"}]]],"referenceName":"gi\_215401171\_ref\_NC","date":"18 July 2018, Wednesday, 16:57:16","order":[0,1,2,3,4,5,6,7,8,9,10,11,12,13],"assembliesNames":["ABySS\_127","ABySS\_63","CLC","IDBA\_UD","MEGAHIT","MIRA","Ray\_Meta","SOAPdenovo2","SPAdes","SPAdes\_meta","SPAdes\_sc","SPAdes\_sc\_careful","Velvet","Geneious"]},{"assembliesWithNs":null,"minContig":500,"report":[["Genome statistics",[{"values":["1.020","2.821","4.194","11.279","1.702","0.931","13.351","11.603","11.827","11.018"],"quality":"More is better","isMain":true,"metricName":"Genome fraction (%)"},{"values":["1.000","1.306","1.000","1.007","1.000","1.341","1.019","1.000","1.022","1.024"],"quality":"Less is better","isMain":true,"metricName":"Duplication ratio"},{"values":[550,1097,920,1665,1129,740,1879,2109,1879,1879],"quality":"More is better","isMain":true,"metricName":"Largest alignment"},{"values":[1573,4765,6467,17394,2624,1435,20593,17887,18288,17042],"quality":"More is better","isMain":true,"metricName":"Total aligned length"},{"values":[513,707,626,922,807,158,1135,1173,966,900],"quality":"More is better","isMain":false,"metricName":"NA50"},{"values":[510,298,542,660,688,null,593,614,593,593],"quality":"More is better","isMain":false,"metricName":"NA75"},{"values":[null,null,null,null,null,null,null,null,null,null],"quality":"More is better","isMain":true,"metricName":"NGA50"},{"values":[2,4,5,7,2,3,8,6,7,7],"quality":"Less is better","isMain":false,"metricName":"LA50"},{"values":[3,6,7,13,3,null,15,13,14,13],"quality":"Less is better","isMain":false,"metricName":"LA75"}]],["Misassemblies",[{"values":[0,0,0,0,0,0,1,0,0,0],"quality":"Less is better","isMain":true,"metricName":"# misassemblies"},{"values":[0,0,0,0,0,0,1,0,0,0],"quality":"Less is better","isMain":false,"metricName":" # relocations"},{"values":[0,0,0,0,0,0,0,0,0,0],"quality":"Less is better","isMain":false,"metricName":" # translocations"},{"values":[0,0,0,0,0,0,0,0,0,0],"quality":"Less is better","isMain":false,"metricName":" # inversions"},{"values":[0,0,0,0,0,0,1,0,0,0],"quality":"Less is better","isMain":false,"metricName":"# misassembled contigs"},{"values":[0,0,0,0,0,0,1408,0,0,0],"quality":"Less is better","isMain":true,"metricName":"Misassembled contigs length"},{"values":[0,0,0,0,0,0,0,0,0,0],"quality":"Less is better","isMain":false,"metricName":"# local misassemblies"},{"values":[0,0,0,0,0,0,0,0,0,0],"quality":"Less is better","isMain":false,"metricName":"# unaligned mis. contigs"}]],["Unaligned",[{"values":[0,0,0,0,0,0,0,0,0,0],"quality":"Less is better","isMain":false,"metricName":"# fully unaligned contigs"},{"values":[0,0,0,0,0,0,0,0,0,0],"quality":"Less is better","isMain":false,"metricName":"Fully unaligned length"},{"values":[0,0,0,0,0,1,1,1,0,0],"quality":"Less is better","isMain":false,"metricName":"# partially unaligned contigs"},{"values":[0,0,0,0,0,731,1468,1119,0,0],"quality":"Less is better","isMain":false,"metricName":"Partially unaligned length"}]],["Mismatches",[{"values":[0,50,7,84,2,5,83,99,81,54],"quality":"Less is better","isMain":false,"metricName":"# mismatches"},{"values":[0,0,0,1,0,1,1,10,3,2],"quality":"Less is better","isMain":false,"metricName":"# indels"},{"values":[0,0,0,1,0,52,6,18,8,7],"quality":"Less is better","isMain":false,"metricName":"Indels length"},{"values":["0.00","1149.43","108.24","483.01","76.22","348.43","403.17","553.35","444.15","317.83"],"quality":"Less is better","isMain":true,"metricName":"# mismatches per 100 kbp"},{"values":["0.00","0.00","0.00","5.75","0.00","69.69","4.86","55.89","16.45","11.77"],"quality":"Less is better","isMain":true,"metricName":"# indels per 100 kbp"},{"values":[0,0,0,1,0,0,0,9,2,1],"quality":"Less is better","isMain":false,"metricName":" # indels (<= 5 bp)"},{"values":[0,0,0,0,0,1,1,1,1,1],"quality":"Less is better","isMain":false,"metricName":" # indels (> 5 bp)"},{"values":[0,0,0,0,7,590,0,0,0,0],"quality":"Less is better","isMain":false,"metricName":"# N's"},{"values":["0.00","0.00","0.00","0.00","266.77","22222.22","0.00","0.00","0.00","0.00"],"quality":"Less is better","isMain":true,"metricName":"# N's per 100 kbp"}]],["Statistics without reference",[{"values":[3,7,10,20,3,4,21,19,21,20],"quality":"Equal","isMain":true,"metricName":"# contigs"},{"values":[0,2,0,6,1,0,10,8,6,5],"quality":"Equal","isMain":false,"metricName":"# contigs (>= 1000 bp)"},{"values":[0,0,0,0,0,0,0,0,0,0],"quality":"Equal","isMain":false,"metricName":"# contigs (>= 5000 bp)"},{"values":[0,0,0,0,0,0,0,0,0,0],"quality":"Equal","isMain":false,"metricName":"# contigs (>= 10000 bp)"},{"values":[0,0,0,0,0,0,0,0,0,0],"quality":"Equal","isMain":false,"metricName":"# contigs (>= 25000 bp)"},{"values":[0,0,0,0,0,0,0,0,0,0],"quality":"Equal","isMain":false,"metricName":"# contigs (>= 50000 bp)"},{"values":[550,1162,920,1665,1129,818,2192,2118,1880,1880],"quality":"More is better","isMain":true,"metricName":"Largest contig"},{"values":[1573,5682,6468,17511,2624,2655,22440,19016,18639,17393],"quality":"More is better","isMain":true,"metricName":"Total length"},{"values":[0,2173,0,8311,1129,0,15709,11981,8676,7485],"quality":"More is better","isMain":true,"metricName":"Total length (>= 1000 bp)"},{"values":[0,0,0,0,0,0,0,0,0,0],"quality":"More is better","isMain":false,"metricName":"Total length (>= 5000 bp)"},{"values":[0,0,0,0,0,0,0,0,0,0],"quality":"More is better","isMain":true,"metricName":"Total length (>= 10000 bp)"},{"values":[0,0,0,0,0,0,0,0,0,0],"quality":"More is better","isMain":false,"metricName":"Total length (>= 25000 bp)"},{"values":[0,0,0,0,0,0,0,0,0,0],"quality":"More is better","isMain":true,"metricName":"Total length (>= 50000 bp)"},{"values":[513,730,626,922,807,741,1408,1223,966,900],"quality":"More is better","isMain":false,"metricName":"N50"},{"values":[510,704,542,660,688,589,779,692,665,614],"quality":"More is better","isMain":false,"metricName":"N75"},{"values":[2,3,5,7,2,2,7,6,7,7],"quality":"Less is better","isMain":false,"metricName":"L50"},{"values":[3,5,7,13,3,3,12,11,13,13],"quality":"Less is better","isMain":false,"metricName":"L75"},{"values":["40.43","38.74","36.43","38.92","41.04","44.55","40.04","39.24","39.60","39.91"],"quality":"Equal","isMain":false,"metricName":"GC (%)"}]],["Predicted genes",[]],["Similarity statistics",[{"values":[0,0,0,0,0,0,0,0,0,0],"quality":"Equal","isMain":false,"metricName":"# similar correct contigs"},{"values":[0,0,0,0,0,0,0,0,0,0],"quality":"Equal","isMain":false,"metricName":"# similar misassembled blocks"}]],["Reference statistics",[{"values":[154196,154196,154196,154196,154196,154196,154196,154196,154196,154196],"quality":"Equal","isMain":false,"metricName":"Reference length"},{"values":[1,1,1,1,1,1,1,1,1,1],"quality":"Equal","isMain":false,"metricName":"Reference fragments"},{"values":["40.07","40.07","40.07","40.07","40.07","40.07","40.07","40.07","40.07","40.07"],"quality":"Equal","isMain":false,"metricName":"Reference GC (%)"}]]],"referenceName":"gi\_215401232\_ref\_NC","date":"18 July 2018, Wednesday, 16:57:29","order":[0,1,2,3,4,5,6,7,8,9],"assembliesNames":["ABySS\_127","CLC","IDBA\_UD","MEGAHIT","MIRA","SOAPdenovo2","SPAdes","SPAdes\_meta","SPAdes\_sc","SPAdes\_sc\_careful"]},{"assembliesWithNs":null,"minContig":500,"report":[["Genome statistics",[{"values":["6.507","1.733","8.703","2.876","9.803","10.630","10.775","10.775"],"quality":"More is better","isMain":true,"metricName":"Genome fraction (%)"},{"values":["1.108","0.999","1.049","1.000","1.000","1.017","1.048","1.048"],"quality":"Less is better","isMain":true,"metricName":"Duplication ratio"},{"values":[858,587,1083,852,868,928,1287,1287],"quality":"More is better","isMain":true,"metricName":"Largest alignment"},{"values":[9985,2574,13091,4274,14575,15797,16015,16015],"quality":"More is better","isMain":true,"metricName":"Total aligned length"},{"values":[708,null,662,742,562,673,579,579],"quality":"More is better","isMain":false,"metricName":"NA50"},{"values":[null,null,541,612,null,515,null,null],"quality":"More is better","isMain":false,"metricName":"NA75"},{"values":[null,null,null,null,null,null,null,null],"quality":"More is better","isMain":true,"metricName":"NGA50"},{"values":[9,null,10,3,15,13,17,17],"quality":"Less is better","isMain":false,"metricName":"LA50"},{"values":[null,null,16,5,null,21,null,null],"quality":"Less is better","isMain":false,"metricName":"LA75"}]],["Misassemblies",[{"values":[0,0,0,0,0,0,0,0],"quality":"Less is better","isMain":true,"metricName":"# misassemblies"},{"values":[0,0,0,0,0,0,0,0],"quality":"Less is better","isMain":false,"metricName":" # relocations"},{"values":[0,0,0,0,0,0,0,0],"quality":"Less is better","isMain":false,"metricName":" # translocations"},{"values":[0,0,0,0,0,0,0,0],"quality":"Less is better","isMain":false,"metricName":" # inversions"},{"values":[0,0,0,0,0,0,0,0],"quality":"Less is better","isMain":false,"metricName":"# misassembled contigs"},{"values":[0,0,0,0,0,0,0,0],"quality":"Less is better","isMain":true,"metricName":"Misassembled contigs length"},{"values":[0,0,1,0,0,0,1,1],"quality":"Less is better","isMain":false,"metricName":"# local misassemblies"},{"values":[0,0,0,0,0,0,0,0],"quality":"Less is better","isMain":false,"metricName":"# unaligned mis. contigs"}]],["Unaligned",[{"values":[0,0,0,0,0,0,0,0],"quality":"Less is better","isMain":false,"metricName":"# fully unaligned contigs"},{"values":[0,0,0,0,0,0,0,0],"quality":"Less is better","isMain":false,"metricName":"Fully unaligned length"},{"values":[4,3,1,0,7,1,6,6],"quality":"Less is better","isMain":false,"metricName":"# partially unaligned contigs"},{"values":[3131,2651,574,0,6872,3283,8553,8553],"quality":"Less is better","isMain":false,"metricName":"Partially unaligned length"}]],["Mismatches",[{"values":[74,21,57,5,49,55,91,88],"quality":"Less is better","isMain":false,"metricName":"# mismatches"},{"values":[9,1,10,0,2,2,6,6],"quality":"Less is better","isMain":false,"metricName":"# indels"},{"values":[12,2,17,0,4,5,12,12],"quality":"Less is better","isMain":false,"metricName":"Indels length"},{"values":["765.17","815.22","440.66","116.99","336.28","348.10","568.22","549.48"],"quality":"Less is better","isMain":true,"metricName":"# mismatches per 100 kbp"},{"values":["93.06","38.82","77.31","0.00","13.73","12.66","37.46","37.46"],"quality":"Less is better","isMain":true,"metricName":"# indels per 100 kbp"},{"values":[9,1,10,0,2,2,6,6],"quality":"Less is better","isMain":false,"metricName":" # indels (<= 5 bp)"},{"values":[0,0,0,0,0,0,0,0],"quality":"Less is better","isMain":false,"metricName":" # indels (> 5 bp)"},{"values":[0,0,0,0,0,0,0,0],"quality":"Less is better","isMain":false,"metricName":"# N's"},{"values":["0.00","0.00","0.00","0.00","0.00","0.00","0.00","0.00"],"quality":"Less is better","isMain":true,"metricName":"# N's per 100 kbp"}]],["Statistics without reference",[{"values":[16,5,20,6,23,23,24,24],"quality":"Equal","isMain":true,"metricName":"# contigs"},{"values":[4,3,2,0,7,1,7,7],"quality":"Equal","isMain":false,"metricName":"# contigs (>= 1000 bp)"},{"values":[0,0,0,0,0,0,0,0],"quality":"Equal","isMain":false,"metricName":"# contigs (>= 5000 bp)"},{"values":[0,0,0,0,0,0,0,0],"quality":"Equal","isMain":false,"metricName":"# contigs (>= 10000 bp)"},{"values":[0,0,0,0,0,0,0,0],"quality":"Equal","isMain":false,"metricName":"# contigs (>= 25000 bp)"},{"values":[0,0,0,0,0,0,0,0],"quality":"Equal","isMain":false,"metricName":"# contigs (>= 50000 bp)"},{"values":[1419,1788,1161,852,1945,3753,3753,3753],"quality":"More is better","isMain":true,"metricName":"Largest contig"},{"values":[13844,5225,14142,4274,21447,19357,25331,25331],"quality":"More is better","isMain":true,"metricName":"Total length"},{"values":[5371,4103,2302,0,10289,3753,13059,13059],"quality":"More is better","isMain":true,"metricName":"Total length (>= 1000 bp)"},{"values":[0,0,0,0,0,0,0,0],"quality":"More is better","isMain":false,"metricName":"Total length (>= 5000 bp)"},{"values":[0,0,0,0,0,0,0,0],"quality":"More is better","isMain":true,"metricName":"Total length (>= 10000 bp)"},{"values":[0,0,0,0,0,0,0,0],"quality":"More is better","isMain":false,"metricName":"Total length (>= 25000 bp)"},{"values":[0,0,0,0,0,0,0,0],"quality":"More is better","isMain":true,"metricName":"Total length (>= 50000 bp)"},{"values":[852,1224,700,742,868,808,1245,1245],"quality":"More is better","isMain":false,"metricName":"N50"},{"values":[708,1091,642,612,708,662,742,742],"quality":"More is better","isMain":false,"metricName":"N75"},{"values":[6,2,9,3,8,8,7,7],"quality":"Less is better","isMain":false,"metricName":"L50"},{"values":[11,3,14,5,15,15,15,15],"quality":"Less is better","isMain":false,"metricName":"L75"},{"values":["44.91","43.94","45.80","45.60","44.95","45.40","44.94","44.94"],"quality":"Equal","isMain":false,"metricName":"GC (%)"}]],["Predicted genes",[]],["Similarity statistics",[{"values":[0,0,0,0,0,0,0,0],"quality":"Equal","isMain":false,"metricName":"# similar correct contigs"},{"values":[0,0,0,0,0,0,0,0],"quality":"Equal","isMain":false,"metricName":"# similar misassembled blocks"}]],["Reference statistics",[{"values":[148634,148634,148634,148634,148634,148634,148634,148634],"quality":"Equal","isMain":false,"metricName":"Reference length"},{"values":[1,1,1,1,1,1,1,1],"quality":"Equal","isMain":false,"metricName":"Reference fragments"},{"values":["45.00","45.00","45.00","45.00","45.00","45.00","45.00","45.00"],"quality":"Equal","isMain":false,"metricName":"Reference GC (%)"}]]],"referenceName":"gi\_215401395\_ref\_NC","date":"18 July 2018, Wednesday, 16:57:41","order":[0,1,2,3,4,5,6,7],"assembliesNames":["CLC","IDBA\_UD","MEGAHIT","SOAPdenovo2","SPAdes","SPAdes\_meta","SPAdes\_sc","SPAdes\_sc\_careful"]},{"assembliesWithNs":null,"minContig":500,"report":[["Genome statistics",[{"values":["63.690","41.155","62.077","46.401","70.863","76.354","22.443","54.285","29.887","42.559","38.742","22.063","46.249"],"quality":"More is better","isMain":true,"metricName":"Genome fraction (%)"},{"values":["1.028","1.012","1.046","1.001","1.054","1.069","1.365","1.029","1.000","1.010","1.016","1.004","1.120"],"quality":"Less is better","isMain":true,"metricName":"Duplication ratio"},{"values":[1511,2702,3215,4111,2379,8637,2096,6155,2200,3637,2128,2225,8637],"quality":"More is better","isMain":true,"metricName":"Largest alignment"},{"values":[24157,15366,23948,17115,27493,30115,8301,20323,11024,15862,14509,8168,19098],"quality":"More is better","isMain":true,"metricName":"Total aligned length"},{"values":[738,null,952,null,1269,2774,null,1441,null,null,null,null,5321],"quality":"More is better","isMain":false,"metricName":"NG50"},{"values":[null,null,null,null,null,1613,null,null,null,null,null,null,null],"quality":"More is better","isMain":false,"metricName":"NG75"},{"values":[991,1288,1650,1380,1394,3335,null,2366,2113,1299,1002,1006,4721],"quality":"More is better","isMain":false,"metricName":"NA50"},{"values":[738,589,952,938,1030,1684,null,1375,1485,958,754,922,4480],"quality":"More is better","isMain":false,"metricName":"NA75"},{"values":[738,null,952,null,1174,2774,null,923,null,null,null,null,1260],"quality":"More is better","isMain":true,"metricName":"NGA50"},{"values":[null,null,null,null,null,1076,null,null,null,null,null,null,null],"quality":"More is better","isMain":false,"metricName":"NGA75"},{"values":[18,null,11,null,11,4,null,6,null,null,null,null,3],"quality":"Less is better","isMain":false,"metricName":"LG50"},{"values":[null,null,null,null,null,8,null,null,null,null,null,null,null],"quality":"Less is better","isMain":false,"metricName":"LG75"},{"values":[10,4,6,4,8,3,null,3,3,4,5,3,2],"quality":"Less is better","isMain":false,"metricName":"LA50"},{"values":[18,9,11,7,14,6,null,6,5,7,10,5,3],"quality":"Less is better","isMain":false,"metricName":"LA75"},{"values":[18,null,11,null,12,4,null,8,null,null,null,null,4],"quality":"Less is better","isMain":true,"metricName":"LGA50"},{"values":[null,null,null,null,null,9,null,null,null,null,null,null,null],"quality":"Less is better","isMain":false,"metricName":"LGA75"}]],["Misassemblies",[{"values":[0,0,0,0,0,0,0,0,0,0,0,0,0],"quality":"Less is better","isMain":true,"metricName":"# misassemblies"},{"values":[0,0,0,0,0,0,0,0,0,0,0,0,0],"quality":"Less is better","isMain":false,"metricName":" # relocations"},{"values":[0,0,0,0,0,0,0,0,0,0,0,0,0],"quality":"Less is better","isMain":false,"metricName":" # translocations"},{"values":[0,0,0,0,0,0,0,0,0,0,0,0,0],"quality":"Less is better","isMain":false,"metricName":" # inversions"},{"values":[0,0,0,0,0,0,0,0,0,0,0,0,0],"quality":"Less is better","isMain":false,"metricName":"# misassembled contigs"},{"values":[0,0,0,0,0,0,0,0,0,0,0,0,0],"quality":"Less is better","isMain":true,"metricName":"Misassembled contigs length"},{"values":[0,0,0,0,0,1,0,0,0,0,0,0,1],"quality":"Less is better","isMain":false,"metricName":"# local misassemblies"},{"values":[0,0,0,0,0,0,4,0,0,0,0,0,0],"quality":"Less is better","isMain":false,"metricName":"# unaligned mis. contigs"}]],["Unaligned",[{"values":[0,0,0,0,0,0,0,0,0,0,0,0,0],"quality":"Less is better","isMain":false,"metricName":"# fully unaligned contigs"},{"values":[0,0,0,0,0,0,0,0,0,0,0,0,0],"quality":"Less is better","isMain":false,"metricName":"Fully unaligned length"},{"values":[0,0,0,0,0,2,4,1,0,0,0,0,3],"quality":"Less is better","isMain":false,"metricName":"# partially unaligned contigs"},{"values":[0,0,0,0,0,2218,5556,1528,0,0,0,0,3059],"quality":"Less is better","isMain":false,"metricName":"Partially unaligned length"}]],["Mismatches",[{"values":[13,60,60,30,52,49,14,137,46,49,54,21,28],"quality":"Less is better","isMain":false,"metricName":"# mismatches"},{"values":[0,0,2,0,1,0,3,7,2,0,0,0,0],"quality":"Less is better","isMain":false,"metricName":"# indels"},{"values":[0,0,2,0,1,0,165,7,2,0,0,0,0],"quality":"Less is better","isMain":false,"metricName":"Indels length"},{"values":["55.34","395.26","262.04","175.28","198.94","173.99","169.12","684.21","417.27","312.14","377.89","258.05","164.14"],"quality":"Less is better","isMain":true,"metricName":"# mismatches per 100 kbp"},{"values":["0.00","0.00","8.73","0.00","3.83","0.00","36.24","34.96","18.14","0.00","0.00","0.00","0.00"],"quality":"Less is better","isMain":true,"metricName":"# indels per 100 kbp"},{"values":[0,0,2,0,1,0,0,7,2,0,0,0,0],"quality":"Less is better","isMain":false,"metricName":" # indels (<= 5 bp)"},{"values":[0,0,0,0,0,0,3,0,0,0,0,0,0],"quality":"Less is better","isMain":false,"metricName":" # indels (> 5 bp)"},{"values":[0,0,0,0,0,25,3719,277,0,0,0,0,39],"quality":"Less is better","isMain":false,"metricName":"# N's"},{"values":["0.00","0.00","0.00","0.00","0.00","77.32","22062.05","1251.81","0.00","0.00","0.00","0.00","176.02"],"quality":"Less is better","isMain":true,"metricName":"# N's per 100 kbp"}]],["Statistics without reference",[{"values":[27,15,18,13,22,12,14,10,7,13,15,8,4],"quality":"Equal","isMain":true,"metricName":"# contigs"},{"values":[8,4,9,6,14,10,5,7,6,6,5,3,4],"quality":"Equal","isMain":false,"metricName":"# contigs (>= 1000 bp)"},{"values":[0,0,0,0,0,2,0,1,0,0,0,0,3],"quality":"Equal","isMain":false,"metricName":"# contigs (>= 5000 bp)"},{"values":[0,0,0,0,0,0,0,0,0,0,0,0,0],"quality":"Equal","isMain":false,"metricName":"# contigs (>= 10000 bp)"},{"values":[0,0,0,0,0,0,0,0,0,0,0,0,0],"quality":"Equal","isMain":false,"metricName":"# contigs (>= 25000 bp)"},{"values":[0,0,0,0,0,0,0,0,0,0,0,0,0],"quality":"Equal","isMain":false,"metricName":"# contigs (>= 50000 bp)"},{"values":[1511,2702,3215,4111,2379,8637,3245,6155,2200,3637,2128,2225,8637],"quality":"More is better","isMain":true,"metricName":"Largest contig"},{"values":[24157,15366,23954,17128,27545,32333,16857,22128,11024,15862,14512,8168,22157],"quality":"More is better","isMain":true,"metricName":"Total length"},{"values":[10113,7937,16942,11967,21728,31145,10112,19838,10486,11066,7352,4377,22157],"quality":"More is better","isMain":true,"metricName":"Total length (>= 1000 bp)"},{"values":[0,0,0,0,0,14039,0,6155,0,0,0,0,19666],"quality":"More is better","isMain":false,"metricName":"Total length (>= 5000 bp)"},{"values":[0,0,0,0,0,0,0,0,0,0,0,0,0],"quality":"More is better","isMain":true,"metricName":"Total length (>= 10000 bp)"},{"values":[0,0,0,0,0,0,0,0,0,0,0,0,0],"quality":"More is better","isMain":false,"metricName":"Total length (>= 25000 bp)"},{"values":[0,0,0,0,0,0,0,0,0,0,0,0,0],"quality":"More is better","isMain":true,"metricName":"Total length (>= 50000 bp)"},{"values":[991,1288,1650,1380,1407,4322,1735,2903,2113,1299,1002,1006,5708],"quality":"More is better","isMain":false,"metricName":"N50"},{"values":[738,589,952,938,1036,1685,868,1646,1485,958,754,922,5321],"quality":"More is better","isMain":false,"metricName":"N75"},{"values":[10,4,6,4,8,3,4,3,3,4,5,3,2],"quality":"Less is better","isMain":false,"metricName":"L50"},{"values":[18,9,11,7,13,6,8,5,5,7,10,5,3],"quality":"Less is better","isMain":false,"metricName":"L75"},{"values":["64.83","65.64","64.14","63.49","63.95","63.95","63.94","63.87","64.13","63.82","64.26","64.47","63.44"],"quality":"Equal","isMain":false,"metricName":"GC (%)"}]],["Predicted genes",[]],["Similarity statistics",[{"values":[0,0,0,0,0,0,0,0,0,0,0,0,0],"quality":"Equal","isMain":false,"metricName":"# similar correct contigs"},{"values":[0,0,0,0,0,0,0,0,0,0,0,0,0],"quality":"Equal","isMain":false,"metricName":"# similar misassembled blocks"}]],["Reference statistics",[{"values":[36885,36885,36885,36885,36885,36885,36885,36885,36885,36885,36885,36885,36885],"quality":"Equal","isMain":false,"metricName":"Reference length"},{"values":[1,1,1,1,1,1,1,1,1,1,1,1,1],"quality":"Equal","isMain":false,"metricName":"Reference fragments"},{"values":["64.51","64.51","64.51","64.51","64.51","64.51","64.51","64.51","64.51","64.51","64.51","64.51","64.51"],"quality":"Equal","isMain":false,"metricName":"Reference GC (%)"}]]],"referenceName":"gi\_215479924\_ref\_NC","date":"18 July 2018, Wednesday, 16:57:58","order":[0,1,2,3,4,5,6,7,8,9,10,11,12],"assembliesNames":["ABySS\_127","ABySS\_63","CLC","IDBA\_UD","MEGAHIT","MIRA","SOAPdenovo2","SPAdes","SPAdes\_meta","SPAdes\_sc","SPAdes\_sc\_careful","Velvet","Geneious"]},{"assembliesWithNs":null,"minContig":500,"report":[["Genome statistics",[{"values":["70.337","44.729","60.311","71.817","66.196","98.332","39.960","85.376","64.504","73.949","54.832","50.066","82.193"],"quality":"More is better","isMain":true,"metricName":"Genome fraction (%)"},{"values":["1.023","1.006","1.001","1.025","1.017","1.338","1.129","1.038","1.008","1.010","1.006","1.003","1.234"],"quality":"Less is better","isMain":true,"metricName":"Duplication ratio"},{"values":[2542,2008,9968,5709,10888,13228,4233,12369,10090,5561,4583,4936,13228],"quality":"More is better","isMain":true,"metricName":"Largest alignment"},{"values":[26369,16447,22109,26884,24632,48195,14638,32475,23795,27364,20195,18394,37158],"quality":"More is better","isMain":true,"metricName":"Total aligned length"},{"values":[891,null,1062,1730,1987,7407,null,11399,1566,1505,639,502,19751],"quality":"More is better","isMain":false,"metricName":"NG50"},{"values":[null,null,null,null,null,5717,null,11399,null,null,null,null,10568],"quality":"More is better","isMain":false,"metricName":"NG75"},{"values":[1020,1127,2477,2176,3616,5717,1164,11399,4057,4114,3261,1824,7407],"quality":"More is better","isMain":false,"metricName":"NA50"},{"values":[807,758,1439,1466,1398,4070,620,2712,1537,1057,927,1131,null],"quality":"More is better","isMain":false,"metricName":"NA75"},{"values":[891,null,1062,1730,1961,7407,null,11399,1537,1503,639,502,9750],"quality":"More is better","isMain":true,"metricName":"NGA50"},{"values":[null,null,null,null,null,4805,null,2712,null,null,null,null,7407],"quality":"More is better","isMain":false,"metricName":"NGA75"},{"values":[15,null,6,6,4,2,null,2,4,5,10,11,1],"quality":"Less is better","isMain":false,"metricName":"LG50"},{"values":[null,null,null,null,null,3,null,2,null,null,null,null,2],"quality":"Less is better","isMain":false,"metricName":"LG75"},{"values":[10,6,2,4,2,3,4,2,2,3,3,3,3],"quality":"Less is better","isMain":false,"metricName":"LA50"},{"values":[17,10,5,8,5,6,9,4,4,7,6,6,null],"quality":"Less is better","isMain":false,"metricName":"LA75"},{"values":[15,null,6,6,4,2,null,2,4,5,10,11,2],"quality":"Less is better","isMain":true,"metricName":"LGA50"},{"values":[null,null,null,null,null,4,null,4,null,null,null,null,3],"quality":"Less is better","isMain":false,"metricName":"LGA75"}]],["Misassemblies",[{"values":[0,0,0,0,0,0,0,0,0,0,0,0,0],"quality":"Less is better","isMain":true,"metricName":"# misassemblies"},{"values":[0,0,0,0,0,0,0,0,0,0,0,0,0],"quality":"Less is better","isMain":false,"metricName":" # relocations"},{"values":[0,0,0,0,0,0,0,0,0,0,0,0,0],"quality":"Less is better","isMain":false,"metricName":" # translocations"},{"values":[0,0,0,0,0,0,0,0,0,0,0,0,0],"quality":"Less is better","isMain":false,"metricName":" # inversions"},{"values":[0,0,0,0,0,0,0,0,0,0,0,0,0],"quality":"Less is better","isMain":false,"metricName":"# misassembled contigs"},{"values":[0,0,0,0,0,0,0,0,0,0,0,0,0],"quality":"Less is better","isMain":true,"metricName":"Misassembled contigs length"},{"values":[0,0,0,0,0,1,1,1,0,0,0,1,1],"quality":"Less is better","isMain":false,"metricName":"# local misassemblies"},{"values":[0,0,0,0,0,0,2,0,0,0,0,0,0],"quality":"Less is better","isMain":false,"metricName":"# unaligned mis. contigs"}]],["Unaligned",[{"values":[0,0,0,0,0,0,0,0,0,0,0,0,0],"quality":"Less is better","isMain":false,"metricName":"# fully unaligned contigs"},{"values":[0,0,0,0,0,0,0,0,0,0,0,0,0],"quality":"Less is better","isMain":false,"metricName":"Fully unaligned length"},{"values":[0,0,0,0,0,1,1,1,0,0,0,0,2],"quality":"Less is better","isMain":false,"metricName":"# partially unaligned contigs"},{"values":[0,0,0,0,0,2745,1216,4265,0,0,0,0,16333],"quality":"Less is better","isMain":false,"metricName":"Partially unaligned length"}]],["Mismatches",[{"values":[11,1,0,46,40,123,4,123,107,105,24,18,71],"quality":"Less is better","isMain":false,"metricName":"# mismatches"},{"values":[0,0,0,2,0,0,6,2,4,6,0,0,0],"quality":"Less is better","isMain":false,"metricName":"# indels"},{"values":[0,0,0,2,0,0,288,2,5,7,0,0,0],"quality":"Less is better","isMain":false,"metricName":"Indels length"},{"values":["42.69","6.10","0.00","174.85","164.96","341.47","27.33","393.29","452.83","387.61","119.49","98.15","235.81"],"quality":"Less is better","isMain":true,"metricName":"# mismatches per 100 kbp"},{"values":["0.00","0.00","0.00","7.60","0.00","0.00","40.99","6.39","16.93","22.15","0.00","0.00","0.00"],"quality":"Less is better","isMain":true,"metricName":"# indels per 100 kbp"},{"values":[0,0,0,2,0,0,0,2,4,6,0,0,0],"quality":"Less is better","isMain":false,"metricName":" # indels (<= 5 bp)"},{"values":[0,0,0,0,0,0,6,0,0,0,0,0,0],"quality":"Less is better","isMain":false,"metricName":" # indels (> 5 bp)"},{"values":[0,0,0,0,0,25,1945,0,0,0,0,10,429],"quality":"Less is better","isMain":false,"metricName":"# N's"},{"values":["0.00","0.00","0.00","0.00","0.00","49.08","10962.69","0.00","0.00","0.00","0.00","54.34","802.00"],"quality":"Less is better","isMain":true,"metricName":"# N's per 100 kbp"}]],["Statistics without reference",[{"values":[27,17,11,15,11,11,12,8,9,15,13,11,5],"quality":"Equal","isMain":true,"metricName":"# contigs"},{"values":[10,6,6,9,6,9,5,4,6,7,5,7,5],"quality":"Equal","isMain":false,"metricName":"# contigs (>= 1000 bp)"},{"values":[0,0,1,1,1,3,0,2,1,1,0,0,5],"quality":"Equal","isMain":false,"metricName":"# contigs (>= 5000 bp)"},{"values":[0,0,0,0,1,1,0,2,1,0,0,0,2],"quality":"Equal","isMain":false,"metricName":"# contigs (>= 10000 bp)"},{"values":[0,0,0,0,0,0,0,0,0,0,0,0,0],"quality":"Equal","isMain":false,"metricName":"# contigs (>= 25000 bp)"},{"values":[0,0,0,0,0,0,0,0,0,0,0,0,0],"quality":"Equal","isMain":false,"metricName":"# contigs (>= 50000 bp)"},{"values":[2542,2008,9968,5709,10888,15973,4380,16634,10090,5561,4583,4936,19751],"quality":"More is better","isMain":true,"metricName":"Largest contig"},{"values":[26369,16483,22109,26970,24658,50942,17742,36742,23824,27366,20197,18404,53491],"quality":"More is better","isMain":true,"metricName":"Total length"},{"values":[13817,9205,18749,22547,21005,49157,12288,34242,21439,21370,14520,15543,53491],"quality":"More is better","isMain":true,"metricName":"Total length (>= 1000 bp)"},{"values":[0,0,9968,5709,10888,29097,0,28033,10090,5561,0,0,53491],"quality":"More is better","isMain":false,"metricName":"Total length (>= 5000 bp)"},{"values":[0,0,0,0,10888,15973,0,28033,10090,0,0,0,30319],"quality":"More is better","isMain":true,"metricName":"Total length (>= 10000 bp)"},{"values":[0,0,0,0,0,0,0,0,0,0,0,0,0],"quality":"More is better","isMain":false,"metricName":"Total length (>= 25000 bp)"},{"values":[0,0,0,0,0,0,0,0,0,0,0,0,0],"quality":"More is better","isMain":true,"metricName":"Total length (>= 50000 bp)"},{"values":[1020,1127,2477,2176,3616,5717,1748,11399,4057,4114,3261,1824,10568],"quality":"More is better","isMain":false,"metricName":"N50"},{"values":[807,794,1439,1466,1398,4793,918,11399,1566,1057,927,1131,7407],"quality":"More is better","isMain":false,"metricName":"N75"},{"values":[10,6,2,4,2,3,3,2,2,3,3,3,2],"quality":"Less is better","isMain":false,"metricName":"L50"},{"values":[17,10,5,8,5,5,7,2,4,7,6,6,4],"quality":"Less is better","isMain":false,"metricName":"L75"},{"values":["64.53","63.87","64.15","64.03","64.16","64.46","63.80","64.20","64.03","64.12","63.95","63.96","64.38"],"quality":"Equal","isMain":false,"metricName":"GC (%)"}]],["Predicted genes",[]],["Similarity statistics",[{"values":[0,0,0,0,0,0,0,0,0,0,0,0,0],"quality":"Equal","isMain":false,"metricName":"# similar correct contigs"},{"values":[0,0,0,0,0,0,0,0,0,0,0,0,0],"quality":"Equal","isMain":false,"metricName":"# similar misassembled blocks"}]],["Reference statistics",[{"values":[36632,36632,36632,36632,36632,36632,36632,36632,36632,36632,36632,36632,36632],"quality":"Equal","isMain":false,"metricName":"Reference length"},{"values":[1,1,1,1,1,1,1,1,1,1,1,1,1],"quality":"Equal","isMain":false,"metricName":"Reference fragments"},{"values":["64.33","64.33","64.33","64.33","64.33","64.33","64.33","64.33","64.33","64.33","64.33","64.33","64.33"],"quality":"Equal","isMain":false,"metricName":"Reference GC (%)"}]]],"referenceName":"gi\_215479976\_ref\_NC","date":"18 July 2018, Wednesday, 16:58:14","order":[0,1,2,3,4,5,6,7,8,9,10,11,12],"assembliesNames":["ABySS\_127","ABySS\_63","CLC","IDBA\_UD","MEGAHIT","MIRA","SOAPdenovo2","SPAdes","SPAdes\_meta","SPAdes\_sc","SPAdes\_sc\_careful","Velvet","Geneious"]},{"assembliesWithNs":null,"minContig":500,"report":[["Genome statistics",[{"values":["23.826","13.465","75.781","74.248","79.214","43.584","73.792","86.216","85.908","86.046","86.046","9.873"],"quality":"More is better","isMain":true,"metricName":"Genome fraction (%)"},{"values":["1.001","1.000","1.017","1.007","1.003","1.000","1.004","1.000","1.001","1.002","1.002","0.997"],"quality":"Less is better","isMain":true,"metricName":"Duplication ratio"},{"values":[1589,1210,3995,5045,4387,3682,4424,5341,5341,5341,5341,2062],"quality":"More is better","isMain":true,"metricName":"Largest alignment"},{"values":[28436,16053,91840,88512,94471,51960,88057,102829,102566,102558,102564,11728],"quality":"More is better","isMain":true,"metricName":"Total aligned length"},{"values":[null,null,1288,1468,1598,null,1072,2020,1915,1985,1985,null],"quality":"More is better","isMain":false,"metricName":"NG50"},{"values":[null,null,562,null,645,null,null,1105,1069,1091,1091,null],"quality":"More is better","isMain":false,"metricName":"NG75"},{"values":[796,750,1599,1775,1998,1312,1468,2259,2159,2310,2316,900],"quality":"More is better","isMain":false,"metricName":"NA50"},{"values":[654,656,1035,1231,1209,952,889,1509,1382,1509,1509,628],"quality":"More is better","isMain":false,"metricName":"NA75"},{"values":[null,null,1288,1468,1598,null,1030,2020,1915,1985,1985,null],"quality":"More is better","isMain":true,"metricName":"NGA50"},{"values":[null,null,562,null,630,null,null,1105,1069,1091,1091,null],"quality":"More is better","isMain":false,"metricName":"NGA75"},{"values":[null,null,27,26,24,null,31,19,20,19,19,null],"quality":"Less is better","isMain":false,"metricName":"LG50"},{"values":[null,null,63,null,51,null,null,38,41,39,39,null],"quality":"Less is better","isMain":false,"metricName":"LG75"},{"values":[15,9,18,17,17,15,19,15,16,15,15,5],"quality":"Less is better","isMain":false,"metricName":"LA50"},{"values":[24,15,35,32,32,26,38,29,31,29,29,9],"quality":"Less is better","isMain":false,"metricName":"LA75"},{"values":[null,null,27,26,24,null,31,19,20,19,19,null],"quality":"Less is better","isMain":true,"metricName":"LGA50"},{"values":[null,null,63,null,52,null,null,38,41,39,39,null],"quality":"Less is better","isMain":false,"metricName":"LGA75"}]],["Misassemblies",[{"values":[0,0,0,0,0,0,0,0,0,0,0,1],"quality":"Less is better","isMain":true,"metricName":"# misassemblies"},{"values":[0,0,0,0,0,0,0,0,0,0,0,1],"quality":"Less is better","isMain":false,"metricName":" # relocations"},{"values":[0,0,0,0,0,0,0,0,0,0,0,0],"quality":"Less is better","isMain":false,"metricName":" # translocations"},{"values":[0,0,0,0,0,0,0,0,0,0,0,0],"quality":"Less is better","isMain":false,"metricName":" # inversions"},{"values":[0,0,0,0,0,0,0,0,0,0,0,1],"quality":"Less is better","isMain":false,"metricName":"# misassembled contigs"},{"values":[0,0,0,0,0,0,0,0,0,0,0,2325],"quality":"Less is better","isMain":true,"metricName":"Misassembled contigs length"},{"values":[0,0,0,0,1,0,1,0,1,0,0,0],"quality":"Less is better","isMain":false,"metricName":"# local misassemblies"},{"values":[0,0,0,0,0,0,0,0,0,0,0,0],"quality":"Less is better","isMain":false,"metricName":"# unaligned mis. contigs"}]],["Unaligned",[{"values":[0,0,0,0,0,0,0,0,0,0,0,0],"quality":"Less is better","isMain":false,"metricName":"# fully unaligned contigs"},{"values":[0,0,0,0,0,0,0,0,0,0,0,0],"quality":"Less is better","isMain":false,"metricName":"Fully unaligned length"},{"values":[0,0,0,0,0,0,1,0,0,0,0,0],"quality":"Less is better","isMain":false,"metricName":"# partially unaligned contigs"},{"values":[0,0,0,0,0,0,624,0,0,0,0,0],"quality":"Less is better","isMain":false,"metricName":"Partially unaligned length"}]],["Mismatches",[{"values":[3,2,99,79,72,14,69,109,94,96,81,6],"quality":"Less is better","isMain":false,"metricName":"# mismatches"},{"values":[0,0,4,3,2,0,5,0,4,5,4,1],"quality":"Less is better","isMain":false,"metricName":"# indels"},{"values":[0,0,48,80,45,0,158,0,104,110,104,42],"quality":"Less is better","isMain":false,"metricName":"Indels length"},{"values":["10.56","12.46","109.58","89.25","76.24","26.94","78.43","106.05","91.78","93.58","78.96","50.98"],"quality":"Less is better","isMain":true,"metricName":"# mismatches per 100 kbp"},{"values":["0.00","0.00","4.43","3.39","2.12","0.00","5.68","0.00","3.91","4.87","3.90","8.50"],"quality":"Less is better","isMain":true,"metricName":"# indels per 100 kbp"},{"values":[0,0,3,1,1,0,1,0,1,1,1,0],"quality":"Less is better","isMain":false,"metricName":" # indels (<= 5 bp)"},{"values":[0,0,1,2,1,0,4,0,3,4,3,1],"quality":"Less is better","isMain":false,"metricName":" # indels (> 5 bp)"},{"values":[0,0,0,0,0,50,274,0,0,0,0,10],"quality":"Less is better","isMain":false,"metricName":"# N's"},{"values":["0.00","0.00","0.00","0.00","0.00","96.23","308.14","0.00","0.00","0.00","0.00","85.19"],"quality":"Less is better","isMain":true,"metricName":"# N's per 100 kbp"}]],["Statistics without reference",[{"values":[36,21,67,56,60,43,70,55,58,55,55,13],"quality":"Equal","isMain":true,"metricName":"# contigs"},{"values":[6,3,36,41,41,25,35,42,42,43,43,3],"quality":"Equal","isMain":false,"metricName":"# contigs (>= 1000 bp)"},{"values":[0,0,0,1,0,0,0,2,2,2,2,0],"quality":"Equal","isMain":false,"metricName":"# contigs (>= 5000 bp)"},{"values":[0,0,0,0,0,0,0,0,0,0,0,0],"quality":"Equal","isMain":false,"metricName":"# contigs (>= 10000 bp)"},{"values":[0,0,0,0,0,0,0,0,0,0,0,0],"quality":"Equal","isMain":false,"metricName":"# contigs (>= 25000 bp)"},{"values":[0,0,0,0,0,0,0,0,0,0,0,0],"quality":"Equal","isMain":false,"metricName":"# contigs (>= 50000 bp)"},{"values":[1589,1210,3995,5045,4387,3682,4424,5341,5341,5341,5341,2325],"quality":"More is better","isMain":true,"metricName":"Largest contig"},{"values":[28436,16053,91897,89143,94726,51960,88920,102829,102566,102767,102773,11738],"quality":"More is better","isMain":true,"metricName":"Total length"},{"values":[7444,3436,70128,78039,81797,38381,64663,93734,90956,94575,94581,5013],"quality":"More is better","isMain":true,"metricName":"Total length (>= 1000 bp)"},{"values":[0,0,0,5045,0,0,0,10408,10408,10408,10408,0],"quality":"More is better","isMain":false,"metricName":"Total length (>= 5000 bp)"},{"values":[0,0,0,0,0,0,0,0,0,0,0,0],"quality":"More is better","isMain":true,"metricName":"Total length (>= 10000 bp)"},{"values":[0,0,0,0,0,0,0,0,0,0,0,0],"quality":"More is better","isMain":false,"metricName":"Total length (>= 25000 bp)"},{"values":[0,0,0,0,0,0,0,0,0,0,0,0],"quality":"More is better","isMain":true,"metricName":"Total length (>= 50000 bp)"},{"values":[796,750,1599,1775,1998,1312,1468,2259,2159,2310,2316,929],"quality":"More is better","isMain":false,"metricName":"N50"},{"values":[654,656,1035,1242,1209,952,937,1509,1382,1509,1509,681],"quality":"More is better","isMain":false,"metricName":"N75"},{"values":[15,9,18,17,17,15,19,15,16,15,15,4],"quality":"Less is better","isMain":false,"metricName":"L50"},{"values":[24,15,35,31,32,26,38,29,31,29,29,8],"quality":"Less is better","isMain":false,"metricName":"L75"},{"values":["35.03","35.89","35.68","35.62","35.56","34.74","35.98","35.52","35.58","35.55","35.55","34.91"],"quality":"Equal","isMain":false,"metricName":"GC (%)"}]],["Predicted genes",[]],["Similarity statistics",[{"values":[0,0,0,0,0,0,0,0,0,0,0,0],"quality":"Equal","isMain":false,"metricName":"# similar correct contigs"},{"values":[0,0,0,0,0,0,0,0,0,0,0,0],"quality":"Equal","isMain":false,"metricName":"# similar misassembled blocks"}]],["Reference statistics",[{"values":[119217,119217,119217,119217,119217,119217,119217,119217,119217,119217,119217,119217],"quality":"Equal","isMain":false,"metricName":"Reference length"},{"values":[1,1,1,1,1,1,1,1,1,1,1,1],"quality":"Equal","isMain":false,"metricName":"Reference fragments"},{"values":["35.70","35.70","35.70","35.70","35.70","35.70","35.70","35.70","35.70","35.70","35.70","35.70"],"quality":"Equal","isMain":false,"metricName":"Reference GC (%)"}]]],"referenceName":"gi\_21686665\_ref\_NC","date":"18 July 2018, Wednesday, 16:58:31","order":[0,1,2,3,4,5,6,7,8,9,10,11],"assembliesNames":["ABySS\_127","ABySS\_63","CLC","IDBA\_UD","MEGAHIT","MIRA","SOAPdenovo2","SPAdes","SPAdes\_meta","SPAdes\_sc","SPAdes\_sc\_careful","Velvet"]},{"assembliesWithNs":null,"minContig":500,"report":[["Genome statistics",[{"values":["35.382","24.821","69.296","77.707","77.296","54.424","26.728","80.539","78.385","76.984","70.928","29.076","24.116"],"quality":"More is better","isMain":true,"metricName":"Genome fraction (%)"},{"values":["1.004","1.002","1.056","1.020","1.043","1.043","1.080","1.020","1.017","1.025","1.029","1.000","1.000"],"quality":"Less is better","isMain":true,"metricName":"Duplication ratio"},{"values":[7160,3925,6471,11488,6768,9445,1315,10122,8813,4633,3820,4895,12642],"quality":"More is better","isMain":true,"metricName":"Largest alignment"},{"values":[52706,36895,108373,117276,119272,83770,39813,121592,118015,116663,107679,43150,35780],"quality":"More is better","isMain":true,"metricName":"Total aligned length"},{"values":[null,null,1150,2088,1955,787,null,3858,2038,1723,1209,null,null],"quality":"More is better","isMain":false,"metricName":"NG50"},{"values":[null,null,null,829,819,null,null,1519,862,624,null,null,null],"quality":"More is better","isMain":false,"metricName":"NG75"},{"values":[809,922,1638,2612,2269,1620,686,4627,2338,2188,1754,1295,5407],"quality":"More is better","isMain":false,"metricName":"NA50"},{"values":[639,644,960,1718,1477,1018,541,2605,1601,1359,954,764,2642],"quality":"More is better","isMain":false,"metricName":"NA75"},{"values":[null,null,1144,2000,1955,776,null,3845,2038,1654,1209,null,null],"quality":"More is better","isMain":true,"metricName":"NGA50"},{"values":[null,null,null,807,805,null,null,1519,862,619,null,null,null],"quality":"More is better","isMain":false,"metricName":"NGA75"},{"values":[null,null,34,20,25,46,null,12,24,29,35,null,null],"quality":"Less is better","isMain":false,"metricName":"LG50"},{"values":[null,null,null,48,53,null,null,27,51,63,null,null,null],"quality":"Less is better","isMain":false,"metricName":"LG75"},{"values":[17,14,22,14,18,18,25,9,17,21,22,10,2],"quality":"Less is better","isMain":false,"metricName":"LA50"},{"values":[35,25,43,28,34,34,43,18,32,39,42,21,5],"quality":"Less is better","isMain":false,"metricName":"LA75"},{"values":[null,null,36,20,25,47,null,12,24,30,35,null,null],"quality":"Less is better","isMain":true,"metricName":"LGA50"},{"values":[null,null,null,49,54,null,null,27,51,65,null,null,null],"quality":"Less is better","isMain":false,"metricName":"LGA75"}]],["Misassemblies",[{"values":[0,0,0,0,0,0,0,0,0,0,0,0,0],"quality":"Less is better","isMain":true,"metricName":"# misassemblies"},{"values":[0,0,0,0,0,0,0,0,0,0,0,0,0],"quality":"Less is better","isMain":false,"metricName":" # relocations"},{"values":[0,0,0,0,0,0,0,0,0,0,0,0,0],"quality":"Less is better","isMain":false,"metricName":" # translocations"},{"values":[0,0,0,0,0,0,0,0,0,0,0,0,0],"quality":"Less is better","isMain":false,"metricName":" # inversions"},{"values":[0,0,0,0,0,0,0,0,0,0,0,0,0],"quality":"Less is better","isMain":false,"metricName":"# misassembled contigs"},{"values":[0,0,0,0,0,0,0,0,0,0,0,0,0],"quality":"Less is better","isMain":true,"metricName":"Misassembled contigs length"},{"values":[0,0,1,0,0,0,0,3,0,0,0,0,0],"quality":"Less is better","isMain":false,"metricName":"# local misassemblies"},{"values":[0,0,0,0,0,0,1,0,0,0,0,0,0],"quality":"Less is better","isMain":false,"metricName":"# unaligned mis. contigs"}]],["Unaligned",[{"values":[0,0,0,0,0,0,0,0,0,0,0,0,0],"quality":"Less is better","isMain":false,"metricName":"# fully unaligned contigs"},{"values":[0,0,0,0,0,0,0,0,0,0,0,0,0],"quality":"Less is better","isMain":false,"metricName":"Fully unaligned length"},{"values":[0,0,1,0,0,0,0,0,0,1,1,0,0],"quality":"Less is better","isMain":false,"metricName":"# partially unaligned contigs"},{"values":[0,0,1446,0,0,0,0,0,0,633,633,0,0],"quality":"Less is better","isMain":false,"metricName":"Partially unaligned length"}]],["Mismatches",[{"values":[16,25,835,854,610,150,54,1343,552,937,772,137,73],"quality":"Less is better","isMain":false,"metricName":"# mismatches"},{"values":[4,3,36,26,12,5,21,75,13,33,29,9,6],"quality":"Less is better","isMain":false,"metricName":"# indels"},{"values":[50,16,68,119,53,47,971,208,57,132,129,24,20],"quality":"Less is better","isMain":false,"metricName":"Indels length"},{"values":["30.48","67.88","812.14","740.72","531.89","185.76","136.17","1123.89","474.63","820.33","733.59","317.57","204.02"],"quality":"Less is better","isMain":true,"metricName":"# mismatches per 100 kbp"},{"values":["7.62","8.15","35.01","22.55","10.46","6.19","52.96","62.76","11.18","28.89","27.56","20.86","16.77"],"quality":"Less is better","isMain":true,"metricName":"# indels per 100 kbp"},{"values":[2,2,35,21,11,3,0,69,11,29,26,8,5],"quality":"Less is better","isMain":false,"metricName":" # indels (<= 5 bp)"},{"values":[2,1,1,5,1,2,21,6,2,4,3,1,1],"quality":"Less is better","isMain":false,"metricName":" # indels (> 5 bp)"},{"values":[0,0,0,0,0,209,3170,0,192,417,533,0,217],"quality":"Less is better","isMain":false,"metricName":"# N's"},{"values":["0.00","0.00","0.00","0.00","0.00","248.11","7398.24","0.00","162.41","354.25","489.27","0.00","606.48"],"quality":"Less is better","isMain":true,"metricName":"# N's per 100 kbp"}]],["Statistics without reference",[{"values":[57,41,79,57,65,61,58,37,60,74,78,38,10],"quality":"Equal","isMain":true,"metricName":"# contigs"},{"values":[13,10,41,37,44,35,7,31,44,44,42,16,10],"quality":"Equal","isMain":false,"metricName":"# contigs (>= 1000 bp)"},{"values":[1,0,1,3,2,1,0,7,2,0,0,0,2],"quality":"Equal","isMain":false,"metricName":"# contigs (>= 5000 bp)"},{"values":[0,0,0,1,0,0,0,1,0,0,0,0,1],"quality":"Equal","isMain":false,"metricName":"# contigs (>= 10000 bp)"},{"values":[0,0,0,0,0,0,0,0,0,0,0,0,0],"quality":"Equal","isMain":false,"metricName":"# contigs (>= 25000 bp)"},{"values":[0,0,0,0,0,0,0,0,0,0,0,0,0],"quality":"Equal","isMain":false,"metricName":"# contigs (>= 50000 bp)"},{"values":[7160,3925,6471,11488,6768,9445,1432,10298,9005,4633,3820,4895,12642],"quality":"More is better","isMain":true,"metricName":"Largest contig"},{"values":[52706,36895,110055,117633,119610,84237,42848,121940,118221,117715,108937,43150,35780],"quality":"More is better","isMain":true,"metricName":"Total length"},{"values":[23736,15283,82295,101632,103239,64975,8654,117137,105385,96466,83071,28540,35780],"quality":"More is better","isMain":true,"metricName":"Total length (>= 1000 bp)"},{"values":[7160,0,6471,23979,12065,9445,0,52171,15467,0,0,0,18049],"quality":"More is better","isMain":false,"metricName":"Total length (>= 5000 bp)"},{"values":[0,0,0,11488,0,0,0,10298,0,0,0,0,12642],"quality":"More is better","isMain":true,"metricName":"Total length (>= 10000 bp)"},{"values":[0,0,0,0,0,0,0,0,0,0,0,0,0],"quality":"More is better","isMain":false,"metricName":"Total length (>= 25000 bp)"},{"values":[0,0,0,0,0,0,0,0,0,0,0,0,0],"quality":"More is better","isMain":true,"metricName":"Total length (>= 50000 bp)"},{"values":[809,922,1642,2612,2269,1620,735,4627,2338,2188,1761,1295,5407],"quality":"More is better","isMain":false,"metricName":"N50"},{"values":[639,644,989,1718,1477,1030,619,2605,1601,1375,1078,764,2642],"quality":"More is better","isMain":false,"metricName":"N75"},{"values":[17,14,21,14,18,18,23,9,17,21,21,10,2],"quality":"Less is better","isMain":false,"metricName":"L50"},{"values":[35,25,42,28,34,34,39,18,32,38,41,21,5],"quality":"Less is better","isMain":false,"metricName":"L75"},{"values":["56.63","56.79","55.29","55.20","55.22","55.40","55.09","55.34","55.29","55.02","55.25","56.22","57.52"],"quality":"Equal","isMain":false,"metricName":"GC (%)"}]],["Predicted genes",[]],["Similarity statistics",[{"values":[0,0,0,0,0,0,0,0,0,0,0,0,0],"quality":"Equal","isMain":false,"metricName":"# similar correct contigs"},{"values":[0,0,0,0,0,0,0,0,0,0,0,0,0],"quality":"Equal","isMain":false,"metricName":"# similar misassembled blocks"}]],["Reference statistics",[{"values":[148371,148371,148371,148371,148371,148371,148371,148371,148371,148371,148371,148371,148371],"quality":"Equal","isMain":false,"metricName":"Reference length"},{"values":[1,1,1,1,1,1,1,1,1,1,1,1,1],"quality":"Equal","isMain":false,"metricName":"Reference fragments"},{"values":["56.10","56.10","56.10","56.10","56.10","56.10","56.10","56.10","56.10","56.10","56.10","56.10","56.10"],"quality":"Equal","isMain":false,"metricName":"Reference GC (%)"}]]],"referenceName":"gi\_216905852\_ref\_NC","date":"18 July 2018, Wednesday, 16:58:51","order":[0,1,2,3,4,5,6,7,8,9,10,11,12],"assembliesNames":["ABySS\_127","ABySS\_63","CLC","IDBA\_UD","MEGAHIT","MIRA","SOAPdenovo2","SPAdes","SPAdes\_meta","SPAdes\_sc","SPAdes\_sc\_careful","Velvet","Geneious"]},{"assembliesWithNs":null,"minContig":500,"report":[["Genome statistics",[{"values":["100.000","100.000","100.000","100.000","100.000","100.000","100.000","100.000","100.000","100.000","100.000"],"quality":"More is better","isMain":true,"metricName":"Genome fraction (%)"},{"values":["1.003","1.000","1.003","1.004","1.016","1.001","1.003","1.001","1.001","1.001","1.001"],"quality":"Less is better","isMain":true,"metricName":"Duplication ratio"},{"values":[37582,37456,37555,37597,38057,37505,37583,37511,37511,37511,37486],"quality":"More is better","isMain":true,"metricName":"Largest alignment"},{"values":[37582,37456,37555,37597,38057,37505,37583,37511,37511,37511,37486],"quality":"More is better","isMain":true,"metricName":"Total aligned length"},{"values":[37582,37456,37555,37597,38057,37505,37583,37511,37511,37511,37486],"quality":"More is better","isMain":false,"metricName":"NG50"},{"values":[37582
