## Supplementary material for "Choice of assembly software has a critical impact on virome characterisation"

### QUAST

**Quality Assessment Tool for Genome Assemblies**
by CAB

Loading...

Aligned to
""

Combined reference
Estimated reference size:
 bp
|
 bp
|
 references
|
 fragments
|
 % G+C
|
 chromosomes
  
 reads
|
 mapped
|
 properly paired
  
 genes
|
 operons

Unfortunately, JavaScript in your
browser is disabled or is not supported.
We need JavaScript to build report and plots.

Worst
Median
Best

Show heatmap

|  |
| --- |
| Contigs are ordered from largest (contig #1) to smallest.  Contigs are broken into nonoverlapping 100 bp windows. Plot shows numbers of windows for each GC percentage. |

[{"assembliesWithNs":["ABySS\_63","ABySS\_127","SOAPdenovo2","SPAdes\_sc","Geneious"],"minContig":500,"report":[["Genome statistics",[{"values":["59.042","49.157","99.719","99.645","99.735","84.597","99.680","99.646","20.487","99.737","99.737","99.625","99.713","84.597","80.944","99.741"],"quality":"More is better","isMain":true,"metricName":"Genome fraction (%)"},{"values":["1.005","1.004","1.063","1.003","1.002","1.001","1.070","1.014","1.019","1.002","1.001","1.002","1.001","1.001","1.082","1.072"],"quality":"Less is better","isMain":true,"metricName":"Duplication ratio"},{"values":[76649,76666,129415,128851,129556,129478,128970,129048,36281,129492,129470,128748,129470,129478,48827,129441],"quality":"More is better","isMain":true,"metricName":"Largest alignment"},{"values":[362058,301060,626708,609282,610345,516986,650157,616906,124681,609876,609689,609027,609283,516986,534402,652197],"quality":"More is better","isMain":true,"metricName":"Total aligned length"},{"values":[null,null,null,71548,71590,36833,39164,69825,null,70642,70598,null,70598,36833,null,null],"quality":"More is better","isMain":false,"metricName":"NA50"},{"values":[null,null,null,39290,38545,22872,25697,37514,null,38545,38521,null,38521,22872,null,null],"quality":"More is better","isMain":false,"metricName":"NA75"},{"values":[null,null,null,4,4,5,5,4,null,4,4,null,4,5,null,null],"quality":"Less is better","isMain":false,"metricName":"LA50"},{"values":[null,null,null,7,7,9,9,7,null,7,7,null,7,9,null,null],"quality":"Less is better","isMain":false,"metricName":"LA75"}]],["Misassemblies",[{"values":[1,0,0,0,0,0,0,4,1,0,0,0,0,0,0,3],"quality":"Less is better","isMain":true,"metricName":"# misassemblies"},{"values":[1,0,0,0,0,0,0,4,1,0,0,0,0,0,0,3],"quality":"Less is better","isMain":false,"metricName":" # relocations"},{"values":[0,0,0,0,0,0,0,0,0,0,0,0,0,0,0,0],"quality":"Less is better","isMain":false,"metricName":" # translocations"},{"values":[0,0,0,0,0,0,0,0,0,0,0,0,0,0,0,0],"quality":"Less is better","isMain":false,"metricName":" # inversions"},{"values":[0,0,0,0,0,0,0,0,0,0,0,0,0,0,0,0],"quality":"Less is better","isMain":false,"metricName":" # interspecies translocations"},{"values":[1,0,0,0,0,0,0,4,1,0,0,0,0,0,0,2],"quality":"Less is better","isMain":false,"metricName":"# misassembled contigs"},{"values":[55287,0,0,0,0,0,0,290262,76440,0,0,0,0,0,0,209554],"quality":"Less is better","isMain":true,"metricName":"Misassembled contigs length"},{"values":[0,0,0,0,0,0,0,0,1,0,0,0,0,0,0,0],"quality":"Less is better","isMain":false,"metricName":"# possibly misassembled contigs"},{"values":[0,0,0,0,0,0,0,0,4,0,0,0,0,0,0,0],"quality":"Less is better","isMain":false,"metricName":" # possible misassemblies"},{"values":[0,1,3,9,9,2,4,4,2,3,3,3,3,2,2,6],"quality":"Less is better","isMain":false,"metricName":"# local misassemblies"},{"values":[0,0,5,0,0,0,0,0,16,0,0,0,0,0,0,0],"quality":"Less is better","isMain":false,"metricName":"# unaligned mis. contigs"}]],["Unaligned",[{"values":[52,50,1143,0,0,0,0,0,2,0,0,1513,0,0,4969,53],"quality":"Less is better","isMain":false,"metricName":"# fully unaligned contigs"},{"values":[4523550,4539144,5327286,0,0,0,0,0,1453,0,0,5699396,0,0,3125238,4491700],"quality":"Less is better","isMain":false,"metricName":"Fully unaligned length"},{"values":[0,0,97,0,0,0,0,0,14,0,0,0,0,0,0,0],"quality":"Less is better","isMain":false,"metricName":"# partially unaligned contigs"},{"values":[0,0,72903,0,0,0,0,0,343704,0,0,0,0,0,0,0],"quality":"Less is better","isMain":false,"metricName":"Partially unaligned length"}]],["Mismatches",[{"values":[27,17,27,27,27,2,29,29,73,27,27,39,28,2,19,27],"quality":"Less is better","isMain":false,"metricName":"# mismatches"},{"values":[22,12,22,18,19,25,21,21,78,23,28,27,24,25,16,27],"quality":"Less is better","isMain":false,"metricName":"# indels"},{"values":[267,39,45,18,19,86,21,50,962,403,422,443,249,86,106,102],"quality":"Less is better","isMain":false,"metricName":"Indels length"},{"values":["7.49","5.66","4.43","4.44","4.43","0.39","4.77","4.77","58.36","4.43","4.43","6.41","4.60","0.39","3.84","4.43"],"quality":"Less is better","isMain":true,"metricName":"# mismatches per 100 kbp"},{"values":["6.10","4.00","3.61","2.96","3.12","4.84","3.45","3.45","62.36","3.78","4.60","4.44","3.94","4.84","3.24","4.43"],"quality":"Less is better","isMain":true,"metricName":"# indels per 100 kbp"},{"values":[16,10,21,18,19,24,21,20,28,18,20,19,19,24,14,26],"quality":"Less is better","isMain":false,"metricName":" # indels (<= 5 bp)"},{"values":[6,2,1,0,0,1,0,1,50,5,8,8,5,1,2,1],"quality":"Less is better","isMain":false,"metricName":" # indels (> 5 bp)"},{"values":[379,300,0,0,0,0,3,0,96628,0,0,2315,0,0,0,1367],"quality":"Less is better","isMain":false,"metricName":"# N's"},{"values":["7.76","6.20","0.00","0.00","0.00","0.00","0.46","0.00","20446.95","0.00","0.00","36.69","0.00","0.00","0.00","26.57"],"quality":"Less is better","isMain":true,"metricName":"# N's per 100 kbp"}]],["Statistics without reference",[{"values":[61,56,1299,12,13,26,89,12,23,14,14,1527,15,26,5385,65],"quality":"Equal","isMain":true,"metricName":"# contigs"},{"values":[329,156,3330,12,17,26,198,12,37,15,17,10459,20,26,36928,71],"quality":"Equal","isMain":false,"metricName":"# contigs (>= 0 bp)"},{"values":[55,46,242,12,12,23,46,12,17,13,13,333,13,23,126,60],"quality":"Equal","isMain":false,"metricName":"# contigs (>= 1000 bp)"},{"values":[43,37,47,12,12,18,15,12,10,12,12,39,11,18,9,47],"quality":"Equal","isMain":false,"metricName":"# contigs (>= 5000 bp)"},{"values":[36,33,42,10,10,13,11,10,7,10,10,33,10,13,8,40],"quality":"Equal","isMain":false,"metricName":"# contigs (>= 10000 bp)"},{"values":[32,29,38,10,10,8,9,10,7,10,10,29,10,8,5,34],"quality":"Equal","isMain":false,"metricName":"# contigs (>= 25000 bp)"},{"values":[24,23,27,5,5,1,3,5,4,5,5,22,5,1,0,29],"quality":"Equal","isMain":false,"metricName":"# contigs (>= 50000 bp)"},{"values":[606039,850094,517843,129514,129556,129478,129652,131360,111231,129492,129470,961438,129470,129478,48827,570751],"quality":"More is better","isMain":true,"metricName":"Largest contig"},{"values":[4885659,4840330,6047398,609973,610384,517099,651026,616996,472579,610018,609790,6309145,609358,517099,3659913,5144531],"quality":"More is better","isMain":true,"metricName":"Total length"},{"values":[4925478,4864782,6793591,609973,611244,517099,683649,616996,476008,610105,609993,8733808,609775,517099,15002577,5146971],"quality":"More is better","isMain":false,"metricName":"Total length (>= 0 bp)"},{"values":[4881764,4833798,5344924,609973,609604,514825,622939,616996,468398,609218,609034,5498689,607975,514825,412440,5140797],"quality":"More is better","isMain":true,"metricName":"Total length (>= 1000 bp)"},{"values":[4848452,4809805,5072171,609973,609604,500287,552295,616996,450690,607171,607031,5078327,601163,500287,269150,5110525],"quality":"More is better","isMain":false,"metricName":"Total length (>= 5000 bp)"},{"values":[4793602,4779566,5038605,598394,597997,469142,528785,605111,431582,595614,595506,5038964,595029,469142,262975,5062855],"quality":"More is better","isMain":true,"metricName":"Total length (>= 10000 bp)"},{"values":[4727943,4712281,4975828,598394,597997,383456,498500,605111,431582,595614,595506,4976556,595029,383456,203753,4961380],"quality":"More is better","isMain":false,"metricName":"Total length (>= 25000 bp)"},{"values":[4401634,4447961,4535797,404576,404786,129478,278058,407905,321696,402496,402519,4686157,402079,129478,0,4759938],"quality":"More is better","isMain":true,"metricName":"Total length (>= 50000 bp)"},{"values":[232572,246998,147644,71548,71590,36833,39183,72105,67075,70642,70598,234413,70598,36833,633,197323],"quality":"More is better","isMain":false,"metricName":"N50"},{"values":[108259,113859,52891,39290,38545,22872,25697,40281,43295,38545,38521,49081,38521,22872,551,107191],"quality":"More is better","isMain":false,"metricName":"N75"},{"values":[8,6,11,4,4,5,5,4,3,4,4,8,4,5,2078,8],"quality":"Less is better","isMain":false,"metricName":"L50"},{"values":[16,13,27,7,7,9,9,7,5,7,7,23,7,9,3626,17],"quality":"Less is better","isMain":false,"metricName":"L75"}]],["Predicted genes",[]],["Similarity statistics",[{"values":[2,2,1,1,1,1,1,2,0,2,2,2,2,1,1,2],"quality":"Equal","isMain":false,"metricName":"# similar correct contigs"},{"values":[0,0,0,0,0,0,0,0,0,0,0,0,0,0,0,0],"quality":"Equal","isMain":false,"metricName":"# similar misassembled blocks"}]],["Reference statistics",[{"values":[610540,610540,610540,610540,610540,610540,610540,610540,610540,610540,610540,610540,610540,610540,610540,610540],"quality":"Equal","isMain":false,"metricName":"Reference length"},{"values":[12,12,12,12,12,12,12,12,12,12,12,12,12,12,12,12],"quality":"Equal","isMain":false,"metricName":"Reference fragments"}]]],"referenceName":"combined\_reference","date":"18 July 2018, Wednesday, 16:00:53","order":[0,1,2,3,4,5,6,7,8,9,10,11,12,13,14,15],"assembliesNames":["ABySS\_63","ABySS\_127","CLC","IDBA\_UD","MEGAHIT","metaVelvet","MIRA","Ray\_Meta","SOAPdenovo2","SPAdes","SPAdes\_meta","SPAdes\_sc","SPAdes\_sc\_careful","Velvet","vicuna","Geneious"]},{"assembliesWithNs":null,"minContig":500,"report":[["Genome statistics",[{"values":["99.978","100.000","99.873","100.000","100.000","100.000","99.849","91.570","100.000","100.000","100.000","99.860","75.336","100.000"],"quality":"More is better","isMain":true,"metricName":"Genome fraction (%)"},{"values":["1.000","1.002","1.053","1.001","1.002","1.097","1.001","0.999","1.001","1.003","1.004","1.000","1.185","1.005"],"quality":"Less is better","isMain":true,"metricName":"Duplication ratio"},{"values":[76649,76666,52891,76765,76807,76666,76550,36281,76743,76925,76993,76559,1579,76666],"quality":"More is better","isMain":true,"metricName":"Largest alignment"},{"values":[76649,76666,80629,76765,76807,84045,76550,70114,76743,76925,76993,76559,68410,76666],"quality":"More is better","isMain":true,"metricName":"Total aligned length"},{"values":[76649,76792,52891,76765,76807,76720,76618,76440,76743,76925,76993,76559,659,77087],"quality":"More is better","isMain":false,"metricName":"NG50"},{"values":[76649,76792,27738,76765,76807,76720,76618,76440,76743,76925,76993,76559,550,77087],"quality":"More is better","isMain":false,"metricName":"NG75"},{"values":[76649,76666,52891,76765,76807,76666,76550,33833,76743,76925,76993,76559,676,76666],"quality":"More is better","isMain":false,"metricName":"NA50"},{"values":[76649,76666,27738,76765,76807,76666,76550,33833,76743,76925,76993,76559,586,76666],"quality":"More is better","isMain":false,"metricName":"NA75"},{"values":[76649,76666,52891,76765,76807,76666,76550,33833,76743,76925,76993,76559,659,76666],"quality":"More is better","isMain":true,"metricName":"NGA50"},{"values":[76649,76666,27738,76765,76807,76666,76550,33833,76743,76925,76993,76559,547,76666],"quality":"More is better","isMain":false,"metricName":"NGA75"},{"values":[1,1,1,1,1,1,1,1,1,1,1,1,46,1],"quality":"Less is better","isMain":false,"metricName":"LG50"},{"values":[1,1,2,1,1,1,1,1,1,1,1,1,77,1],"quality":"Less is better","isMain":false,"metricName":"LG75"},{"values":[1,1,1,1,1,1,1,2,1,1,1,1,40,1],"quality":"Less is better","isMain":false,"metricName":"LA50"},{"values":[1,1,2,1,1,1,1,2,1,1,1,1,67,1],"quality":"Less is better","isMain":false,"metricName":"LA75"},{"values":[1,1,1,1,1,1,1,2,1,1,1,1,46,1],"quality":"Less is better","isMain":true,"metricName":"LGA50"},{"values":[1,1,2,1,1,1,1,2,1,1,1,1,78,1],"quality":"Less is better","isMain":false,"metricName":"LGA75"}]],["Misassemblies",[{"values":[0,0,0,0,0,0,0,1,0,0,0,0,0,0],"quality":"Less is better","isMain":true,"metricName":"# misassemblies"},{"values":[0,0,0,0,0,0,0,1,0,0,0,0,0,0],"quality":"Less is better","isMain":false,"metricName":" # relocations"},{"values":[0,0,0,0,0,0,0,0,0,0,0,0,0,0],"quality":"Less is better","isMain":false,"metricName":" # translocations"},{"values":[0,0,0,0,0,0,0,0,0,0,0,0,0,0],"quality":"Less is better","isMain":false,"metricName":" # inversions"},{"values":[0,0,0,0,0,0,0,1,0,0,0,0,0,0],"quality":"Less is better","isMain":false,"metricName":"# misassembled contigs"},{"values":[0,0,0,0,0,0,0,76440,0,0,0,0,0,0],"quality":"Less is better","isMain":true,"metricName":"Misassembled contigs length"},{"values":[0,0,0,1,1,0,0,1,0,0,0,0,0,0],"quality":"Less is better","isMain":false,"metricName":"# local misassemblies"},{"values":[0,0,0,0,0,0,0,0,0,0,0,0,0,0],"quality":"Less is better","isMain":false,"metricName":"# unaligned mis. contigs"}]],["Unaligned",[{"values":[0,0,0,0,0,0,0,0,0,0,0,0,0,0],"quality":"Less is better","isMain":false,"metricName":"# fully unaligned contigs"},{"values":[0,0,0,0,0,0,0,0,0,0,0,0,0,0],"quality":"Less is better","isMain":false,"metricName":"Fully unaligned length"},{"values":[0,0,0,0,0,0,0,1,0,0,0,0,0,0],"quality":"Less is better","isMain":false,"metricName":"# partially unaligned contigs"},{"values":[0,0,0,0,0,0,0,6326,0,0,0,0,0,0],"quality":"Less is better","isMain":false,"metricName":"Partially unaligned length"}]],["Mismatches",[{"values":[1,1,1,1,1,1,1,35,1,1,13,1,1,1],"quality":"Less is better","isMain":false,"metricName":"# mismatches"},{"values":[0,0,0,0,0,0,0,44,1,3,4,1,1,0],"quality":"Less is better","isMain":false,"metricName":"# indels"},{"values":[0,0,0,0,0,0,0,553,77,204,272,68,68,0],"quality":"Less is better","isMain":false,"metricName":"Indels length"},{"values":["1.30","1.30","1.31","1.30","1.30","1.30","1.31","49.86","1.30","1.30","16.96","1.31","1.73","1.30"],"quality":"Less is better","isMain":true,"metricName":"# mismatches per 100 kbp"},{"values":["0.00","0.00","0.00","0.00","0.00","0.00","0.00","62.68","1.30","3.91","5.22","1.31","1.73","0.00"],"quality":"Less is better","isMain":true,"metricName":"# indels per 100 kbp"},{"values":[0,0,0,0,0,0,0,14,0,0,0,0,0,0],"quality":"Less is better","isMain":false,"metricName":" # indels (<= 5 bp)"},{"values":[0,0,0,0,0,0,0,30,1,3,4,1,1,0],"quality":"Less is better","isMain":false,"metricName":" # indels (> 5 bp)"},{"values":[0,0,0,0,0,1,0,1824,0,0,0,0,0,0],"quality":"Less is better","isMain":false,"metricName":"# N's"},{"values":["0.00","0.00","0.00","0.00","0.00","1.19","0.00","2386.19","0.00","0.00","0.00","0.00","0.00","0.00"],"quality":"Less is better","isMain":true,"metricName":"# N's per 100 kbp"}]],["Statistics without reference",[{"values":[1,1,2,1,1,4,1,1,1,1,1,1,98,1],"quality":"Equal","isMain":true,"metricName":"# contigs"},{"values":[1,1,2,1,1,4,1,1,1,1,1,1,10,1],"quality":"Equal","isMain":false,"metricName":"# contigs (>= 1000 bp)"},{"values":[1,1,2,1,1,1,1,1,1,1,1,1,0,1],"quality":"Equal","isMain":false,"metricName":"# contigs (>= 5000 bp)"},{"values":[1,1,2,1,1,1,1,1,1,1,1,1,0,1],"quality":"Equal","isMain":false,"metricName":"# contigs (>= 10000 bp)"},{"values":[1,1,2,1,1,1,1,1,1,1,1,1,0,1],"quality":"Equal","isMain":false,"metricName":"# contigs (>= 25000 bp)"},{"values":[1,1,1,1,1,1,1,1,1,1,1,1,0,1],"quality":"Equal","isMain":false,"metricName":"# contigs (>= 50000 bp)"},{"values":[76649,76792,52891,76765,76807,76720,76618,76440,76743,76925,76993,76559,1579,77087],"quality":"More is better","isMain":true,"metricName":"Largest contig"},{"values":[76649,76792,80629,76765,76807,84099,76618,76440,76743,76925,76993,76559,68443,77087],"quality":"More is better","isMain":true,"metricName":"Total length"},{"values":[76649,76792,80629,76765,76807,84099,76618,76440,76743,76925,76993,76559,11938,77087],"quality":"More is better","isMain":true,"metricName":"Total length (>= 1000 bp)"},{"values":[76649,76792,80629,76765,76807,76720,76618,76440,76743,76925,76993,76559,0,77087],"quality":"More is better","isMain":false,"metricName":"Total length (>= 5000 bp)"},{"values":[76649,76792,80629,76765,76807,76720,76618,76440,76743,76925,76993,76559,0,77087],"quality":"More is better","isMain":true,"metricName":"Total length (>= 10000 bp)"},{"values":[76649,76792,80629,76765,76807,76720,76618,76440,76743,76925,76993,76559,0,77087],"quality":"More is better","isMain":false,"metricName":"Total length (>= 25000 bp)"},{"values":[76649,76792,52891,76765,76807,76720,76618,76440,76743,76925,76993,76559,0,77087],"quality":"More is better","isMain":true,"metricName":"Total length (>= 50000 bp)"},{"values":[76649,76792,52891,76765,76807,76720,76618,76440,76743,76925,76993,76559,676,77087],"quality":"More is better","isMain":false,"metricName":"N50"},{"values":[76649,76792,27738,76765,76807,76720,76618,76440,76743,76925,76993,76559,586,77087],"quality":"More is better","isMain":false,"metricName":"N75"},{"values":[1,1,1,1,1,1,1,1,1,1,1,1,40,1],"quality":"Less is better","isMain":false,"metricName":"L50"},{"values":[1,1,2,1,1,1,1,1,1,1,1,1,67,1],"quality":"Less is better","isMain":false,"metricName":"L75"},{"values":["30.22","30.23","30.18","30.22","30.22","30.13","30.22","30.22","30.22","30.23","30.26","30.22","29.92","30.25"],"quality":"Equal","isMain":false,"metricName":"GC (%)"}]],["Predicted genes",[]],["Similarity statistics",[{"values":[1,1,0,0,0,1,1,0,1,1,1,1,0,1],"quality":"Equal","isMain":false,"metricName":"# similar correct contigs"},{"values":[0,0,0,0,0,0,0,0,0,0,0,0,0,0],"quality":"Equal","isMain":false,"metricName":"# similar misassembled blocks"}]],["Reference statistics",[{"values":[76666,76666,76666,76666,76666,76666,76666,76666,76666,76666,76666,76666,76666,76666],"quality":"Equal","isMain":false,"metricName":"Reference length"},{"values":[1,1,1,1,1,1,1,1,1,1,1,1,1,1],"quality":"Equal","isMain":false,"metricName":"Reference fragments"},{"values":["30.22","30.22","30.22","30.22","30.22","30.22","30.22","30.22","30.22","30.22","30.22","30.22","30.22","30.22"],"quality":"Equal","isMain":false,"metricName":"Reference GC (%)"}]]],"referenceName":"KC821625","date":"18 July 2018, Wednesday, 16:01:02","order":[0,1,2,3,4,5,6,7,8,9,10,11,12,13],"assembliesNames":["ABySS\_63","ABySS\_127","CLC","IDBA\_UD","MEGAHIT","MIRA","Ray\_Meta","SOAPdenovo2","SPAdes","SPAdes\_meta","SPAdes\_sc","SPAdes\_sc\_careful","vicuna","Geneious"]},{"assembliesWithNs":null,"minContig":500,"report":[["Genome statistics",[{"values":["100.000","100.000","100.000","100.000","99.343","100.000","100.000","5.088","100.000","100.000","100.000","100.000","99.343","99.920","100.000"],"quality":"More is better","isMain":true,"metricName":"Genome fraction (%)"},{"values":["1.024","1.000","1.002","1.003","1.004","1.168","1.015","1.555","1.001","1.001","1.001","1.001","1.004","1.002","1.002"],"quality":"Less is better","isMain":true,"metricName":"Duplication ratio"},{"values":[44392,54016,54115,54157,35739,35913,54799,214,54093,54041,54042,54042,35739,43063,54016],"quality":"More is better","isMain":true,"metricName":"Largest alignment"},{"values":[55287,54016,54115,54157,53817,63047,54799,2748,54093,54041,54042,54042,53817,53973,54016],"quality":"More is better","isMain":true,"metricName":"Total aligned length"},{"values":[55287,54016,54115,54157,35769,35913,54799,null,54093,54041,54042,54042,35769,43162,54124],"quality":"More is better","isMain":false,"metricName":"NG50"},{"values":[55287,54016,54115,54157,8710,5119,54799,null,54093,54041,54042,54042,8710,43162,54124],"quality":"More is better","isMain":false,"metricName":"NG75"},{"values":[44392,54016,54115,54157,35739,35913,54799,null,54093,54041,54042,54042,35739,43063,54016],"quality":"More is better","isMain":false,"metricName":"NA50"},{"values":[44392,54016,54115,54157,8710,2313,54799,null,54093,54041,54042,54042,8710,43063,54016],"quality":"More is better","isMain":false,"metricName":"NA75"},{"values":[44392,54016,54115,54157,35739,35913,54799,null,54093,54041,54042,54042,35739,43063,54016],"quality":"More is better","isMain":true,"metricName":"NGA50"},{"values":[44392,54016,54115,54157,8710,5119,54799,null,54093,54041,54042,54042,8710,43063,54016],"quality":"More is better","isMain":false,"metricName":"NGA75"},{"values":[1,1,1,1,1,1,1,null,1,1,1,1,1,1,1],"quality":"Less is better","isMain":false,"metricName":"LG50"},{"values":[1,1,1,1,2,2,1,null,1,1,1,1,2,1,1],"quality":"Less is better","isMain":false,"metricName":"LG75"},{"values":[1,1,1,1,1,1,1,null,1,1,1,1,1,1,1],"quality":"Less is better","isMain":false,"metricName":"LA50"},{"values":[1,1,1,1,2,5,1,null,1,1,1,1,2,1,1],"quality":"Less is better","isMain":false,"metricName":"LA75"},{"values":[1,1,1,1,1,1,1,null,1,1,1,1,1,1,1],"quality":"Less is better","isMain":true,"metricName":"LGA50"},{"values":[1,1,1,1,2,2,1,null,1,1,1,1,2,1,1],"quality":"Less is better","isMain":false,"metricName":"LGA75"}]],["Misassemblies",[{"values":[1,0,0,0,0,0,0,0,0,0,0,0,0,0,0],"quality":"Less is better","isMain":true,"metricName":"# misassemblies"},{"values":[1,0,0,0,0,0,0,0,0,0,0,0,0,0,0],"quality":"Less is better","isMain":false,"metricName":" # relocations"},{"values":[0,0,0,0,0,0,0,0,0,0,0,0,0,0,0],"quality":"Less is better","isMain":false,"metricName":" # translocations"},{"values":[0,0,0,0,0,0,0,0,0,0,0,0,0,0,0],"quality":"Less is better","isMain":false,"metricName":" # inversions"},{"values":[1,0,0,0,0,0,0,0,0,0,0,0,0,0,0],"quality":"Less is better","isMain":false,"metricName":"# misassembled contigs"},{"values":[55287,0,0,0,0,0,0,0,0,0,0,0,0,0,0],"quality":"Less is better","isMain":true,"metricName":"Misassembled contigs length"},{"values":[0,0,1,1,0,0,1,0,0,0,0,0,0,0,0],"quality":"Less is better","isMain":false,"metricName":"# local misassemblies"},{"values":[0,0,0,0,0,0,0,8,0,0,0,0,0,0,0],"quality":"Less is better","isMain":false,"metricName":"# unaligned mis. contigs"}]],["Unaligned",[{"values":[0,0,0,0,0,0,0,0,0,0,0,0,0,0,0],"quality":"Less is better","isMain":false,"metricName":"# fully unaligned contigs"},{"values":[0,0,0,0,0,0,0,0,0,0,0,0,0,0,0],"quality":"Less is better","isMain":false,"metricName":"Fully unaligned length"},{"values":[0,0,0,0,0,0,0,6,0,0,0,0,0,0,0],"quality":"Less is better","isMain":false,"metricName":"# partially unaligned contigs"},{"values":[0,0,0,0,0,0,0,14181,0,0,0,0,0,0,0],"quality":"Less is better","isMain":false,"metricName":"Partially unaligned length"}]],["Mismatches",[{"values":[0,0,0,0,0,0,1,1,0,0,0,1,0,0,0],"quality":"Less is better","isMain":false,"metricName":"# mismatches"},{"values":[5,6,3,4,7,5,5,0,5,7,6,6,7,4,4],"quality":"Less is better","isMain":false,"metricName":"# indels"},{"values":[22,6,3,4,7,5,34,0,81,67,66,66,7,4,4],"quality":"Less is better","isMain":false,"metricName":"Indels length"},{"values":["0.00","0.00","0.00","0.00","0.00","0.00","1.85","36.39","0.00","0.00","0.00","1.85","0.00","0.00","0.00"],"quality":"Less is better","isMain":true,"metricName":"# mismatches per 100 kbp"},{"values":["9.26","11.11","5.55","7.41","13.05","9.26","9.26","0.00","9.26","12.96","11.11","11.11","13.05","7.41","7.41"],"quality":"Less is better","isMain":true,"metricName":"# indels per 100 kbp"},{"values":[4,6,3,4,7,5,4,0,4,5,4,4,7,4,4],"quality":"Less is better","isMain":false,"metricName":" # indels (<= 5 bp)"},{"values":[1,0,0,0,0,0,1,0,1,2,2,2,0,0,0],"quality":"Less is better","isMain":false,"metricName":" # indels (> 5 bp)"},{"values":[0,0,0,0,0,0,0,8165,0,0,0,0,0,0,3],"quality":"Less is better","isMain":false,"metricName":"# N's"},{"values":["0.00","0.00","0.00","0.00","0.00","0.00","0.00","44247.55","0.00","0.00","0.00","0.00","0.00","0.00","5.54"],"quality":"Less is better","isMain":true,"metricName":"# N's per 100 kbp"}]],["Statistics without reference",[{"values":[1,1,1,1,4,22,1,8,1,1,1,1,4,2,1],"quality":"Equal","isMain":true,"metricName":"# contigs"},{"values":[1,1,1,1,4,9,1,7,1,1,1,1,4,2,1],"quality":"Equal","isMain":false,"metricName":"# contigs (>= 1000 bp)"},{"values":[1,1,1,1,3,2,1,0,1,1,1,1,3,2,1],"quality":"Equal","isMain":false,"metricName":"# contigs (>= 5000 bp)"},{"values":[1,1,1,1,1,1,1,0,1,1,1,1,1,2,1],"quality":"Equal","isMain":false,"metricName":"# contigs (>= 10000 bp)"},{"values":[1,1,1,1,1,1,1,0,1,1,1,1,1,1,1],"quality":"Equal","isMain":false,"metricName":"# contigs (>= 25000 bp)"},{"values":[1,1,1,1,0,0,1,0,1,1,1,1,0,0,1],"quality":"Equal","isMain":false,"metricName":"# contigs (>= 50000 bp)"},{"values":[55287,54016,54115,54157,35769,35913,54799,4974,54093,54041,54042,54042,35769,43162,54124],"quality":"More is better","isMain":true,"metricName":"Largest contig"},{"values":[55287,54016,54115,54157,53847,63083,54799,18453,54093,54041,54042,54042,53847,54081,54124],"quality":"More is better","isMain":true,"metricName":"Total length"},{"values":[55287,54016,54115,54157,53847,54245,54799,17708,54093,54041,54042,54042,53847,54081,54124],"quality":"More is better","isMain":true,"metricName":"Total length (>= 1000 bp)"},{"values":[55287,54016,54115,54157,49799,41032,54799,0,54093,54041,54042,54042,49799,54081,54124],"quality":"More is better","isMain":false,"metricName":"Total length (>= 5000 bp)"},{"values":[55287,54016,54115,54157,35769,35913,54799,0,54093,54041,54042,54042,35769,54081,54124],"quality":"More is better","isMain":true,"metricName":"Total length (>= 10000 bp)"},{"values":[55287,54016,54115,54157,35769,35913,54799,0,54093,54041,54042,54042,35769,43162,54124],"quality":"More is better","isMain":false,"metricName":"Total length (>= 25000 bp)"},{"values":[55287,54016,54115,54157,0,0,54799,0,54093,54041,54042,54042,0,0,54124],"quality":"More is better","isMain":true,"metricName":"Total length (>= 50000 bp)"},{"values":[55287,54016,54115,54157,35769,35913,54799,2437,54093,54041,54042,54042,35769,43162,54124],"quality":"More is better","isMain":false,"metricName":"N50"},{"values":[55287,54016,54115,54157,8710,2322,54799,1587,54093,54041,54042,54042,8710,43162,54124],"quality":"More is better","isMain":false,"metricName":"N75"},{"values":[1,1,1,1,1,1,1,3,1,1,1,1,1,1,1],"quality":"Less is better","isMain":false,"metricName":"L50"},{"values":[1,1,1,1,2,5,1,5,1,1,1,1,2,1,1],"quality":"Less is better","isMain":false,"metricName":"L75"},{"values":["33.43","33.47","33.46","33.48","33.45","33.23","33.45","33.27","33.46","33.47","33.47","33.47","33.45","33.48","33.48"],"quality":"Equal","isMain":false,"metricName":"GC (%)"}]],["Predicted genes",[]],["Similarity statistics",[{"values":[0,0,0,0,0,0,0,0,0,0,0,0,0,0,0],"quality":"Equal","isMain":false,"metricName":"# similar correct contigs"},{"values":[0,0,0,0,0,0,0,0,0,0,0,0,0,0,0],"quality":"Equal","isMain":false,"metricName":"# similar misassembled blocks"}]],["Reference statistics",[{"values":[54012,54012,54012,54012,54012,54012,54012,54012,54012,54012,54012,54012,54012,54012,54012],"quality":"Equal","isMain":false,"metricName":"Reference length"},{"values":[1,1,1,1,1,1,1,1,1,1,1,1,1,1,1],"quality":"Equal","isMain":false,"metricName":"Reference fragments"},{"values":["33.48","33.48","33.48","33.48","33.48","33.48","33.48","33.48","33.48","33.48","33.48","33.48","33.48","33.48","33.48"],"quality":"Equal","isMain":false,"metricName":"Reference GC (%)"}]]],"referenceName":"KC821629","date":"18 July 2018, Wednesday, 16:01:12","order":[0,1,2,3,4,5,6,7,8,9,10,11,12,13,14],"assembliesNames":["ABySS\_63","CLC","IDBA\_UD","MEGAHIT","metaVelvet","MIRA","Ray\_Meta","SOAPdenovo2","SPAdes","SPAdes\_meta","SPAdes\_sc","SPAdes\_sc\_careful","Velvet","vicuna","Geneious"]},{"assembliesWithNs":null,"minContig":500,"report":[["Genome statistics",[{"values":["98.812","98.812","98.812","98.979","98.812","98.812","12.056","98.812","98.812","98.812","98.812","98.979","98.812","98.812"],"quality":"More is better","isMain":true,"metricName":"Genome fraction (%)"},{"values":["1.000","1.003","1.004","1.000","1.006","1.020","1.000","1.002","1.002","1.002","1.002","1.000","1.006","1.010"],"quality":"Less is better","isMain":true,"metricName":"Duplication ratio"},{"values":[36771,36870,36912,36833,36990,37514,251,36848,36826,36826,36826,36833,36998,37131],"quality":"More is better","isMain":true,"metricName":"Largest alignment"},{"values":[36771,36870,36912,36833,36990,37514,4486,36848,36826,36826,36826,36833,36998,37131],"quality":"More is better","isMain":true,"metricName":"Total aligned length"},{"values":[36771,36870,36912,36833,36990,37514,30106,36848,36826,36826,36826,36833,36998,37131],"quality":"More is better","isMain":false,"metricName":"NG50"},{"values":[36771,36870,36912,36833,36990,37514,30106,36848,36826,36826,36826,36833,36998,37131],"quality":"More is better","isMain":false,"metricName":"NG75"},{"values":[36771,36870,36912,36833,36990,37514,null,36848,36826,36826,36826,36833,36998,37131],"quality":"More is better","isMain":false,"metricName":"NA50"},{"values":[36771,36870,36912,36833,36990,37514,null,36848,36826,36826,36826,36833,36998,37131],"quality":"More is better","isMain":false,"metricName":"NA75"},{"values":[36771,36870,36912,36833,36990,37514,null,36848,36826,36826,36826,36833,36998,37131],"quality":"More is better","isMain":true,"metricName":"NGA50"},{"values":[36771,36870,36912,36833,36990,37514,null,36848,36826,36826,36826,36833,36998,37131],"quality":"More is better","isMain":false,"metricName":"NGA75"},{"values":[1,1,1,1,1,1,1,1,1,1,1,1,1,1],"quality":"Less is better","isMain":false,"metricName":"LG50"},{"values":[1,1,1,1,1,1,1,1,1,1,1,1,1,1],"quality":"Less is better","isMain":false,"metricName":"LG75"},{"values":[1,1,1,1,1,1,null,1,1,1,1,1,1,1],"quality":"Less is better","isMain":false,"metricName":"LA50"},{"values":[1,1,1,1,1,1,null,1,1,1,1,1,1,1],"quality":"Less is better","isMain":false,"metricName":"LA75"},{"values":[1,1,1,1,1,1,null,1,1,1,1,1,1,1],"quality":"Less is better","isMain":true,"metricName":"LGA50"},{"values":[1,1,1,1,1,1,null,1,1,1,1,1,1,1],"quality":"Less is better","isMain":false,"metricName":"LGA75"}]],["Misassemblies",[{"values":[0,0,0,0,0,0,0,0,0,0,0,0,0,0],"quality":"Less is better","isMain":true,"metricName":"# misassemblies"},{"values":[0,0,0,0,0,0,0,0,0,0,0,0,0,0],"quality":"Less is better","isMain":false,"metricName":" # relocations"},{"values":[0,0,0,0,0,0,0,0,0,0,0,0,0,0],"quality":"Less is better","isMain":false,"metricName":" # translocations"},{"values":[0,0,0,0,0,0,0,0,0,0,0,0,0,0],"quality":"Less is better","isMain":false,"metricName":" # inversions"},{"values":[0,0,0,0,0,0,0,0,0,0,0,0,0,0],"quality":"Less is better","isMain":false,"metricName":"# misassembled contigs"},{"values":[0,0,0,0,0,0,0,0,0,0,0,0,0,0],"quality":"Less is better","isMain":true,"metricName":"Misassembled contigs length"},{"values":[1,1,1,0,1,1,0,1,1,1,1,0,1,1],"quality":"Less is better","isMain":false,"metricName":"# local misassemblies"},{"values":[0,0,0,0,0,0,1,0,0,0,0,0,0,0],"quality":"Less is better","isMain":false,"metricName":"# unaligned mis. contigs"}]],["Unaligned",[{"values":[0,0,0,0,0,0,0,0,0,0,0,0,0,0],"quality":"Less is better","isMain":false,"metricName":"# fully unaligned contigs"},{"values":[0,0,0,0,0,0,0,0,0,0,0,0,0,0],"quality":"Less is better","isMain":false,"metricName":"Fully unaligned length"},{"values":[0,0,0,0,0,0,1,0,0,0,0,0,0,0],"quality":"Less is better","isMain":false,"metricName":"# partially unaligned contigs"},{"values":[0,0,0,0,0,0,25620,0,0,0,0,0,0,0],"quality":"Less is better","isMain":false,"metricName":"Partially unaligned length"}]],["Mismatches",[{"values":[1,1,1,1,1,1,2,1,1,1,1,1,1,1],"quality":"Less is better","isMain":false,"metricName":"# mismatches"},{"values":[2,2,2,2,2,2,0,1,2,2,2,2,2,2],"quality":"Less is better","isMain":false,"metricName":"# indels"},{"values":[2,2,2,2,2,2,0,1,2,2,2,2,2,2],"quality":"Less is better","isMain":false,"metricName":"Indels length"},{"values":["2.72","2.72","2.72","2.72","2.72","2.72","44.58","2.72","2.72","2.72","2.72","2.72","2.72","2.72"],"quality":"Less is better","isMain":true,"metricName":"# mismatches per 100 kbp"},{"values":["5.44","5.44","5.44","5.43","5.44","5.44","0.00","2.72","5.44","5.44","5.44","5.43","5.44","5.44"],"quality":"Less is better","isMain":true,"metricName":"# indels per 100 kbp"},{"values":[2,2,2,2,2,2,0,1,2,2,2,2,2,2],"quality":"Less is better","isMain":false,"metricName":" # indels (<= 5 bp)"},{"values":[0,0,0,0,0,0,0,0,0,0,0,0,0,0],"quality":"Less is better","isMain":false,"metricName":" # indels (> 5 bp)"},{"values":[0,0,0,0,0,0,10076,0,0,0,0,0,0,1],"quality":"Less is better","isMain":false,"metricName":"# N's"},{"values":["0.00","0.00","0.00","0.00","0.00","0.00","33468.41","0.00","0.00","0.00","0.00","0.00","0.00","2.69"],"quality":"Less is better","isMain":true,"metricName":"# N's per 100 kbp"}]],["Statistics without reference",[{"values":[1,1,1,1,1,1,1,1,1,1,1,1,1,1],"quality":"Equal","isMain":true,"metricName":"# contigs"},{"values":[1,1,1,1,1,1,1,1,1,1,1,1,1,1],"quality":"Equal","isMain":false,"metricName":"# contigs (>= 1000 bp)"},{"values":[1,1,1,1,1,1,1,1,1,1,1,1,1,1],"quality":"Equal","isMain":false,"metricName":"# contigs (>= 5000 bp)"},{"values":[1,1,1,1,1,1,1,1,1,1,1,1,1,1],"quality":"Equal","isMain":false,"metricName":"# contigs (>= 10000 bp)"},{"values":[1,1,1,1,1,1,1,1,1,1,1,1,1,1],"quality":"Equal","isMain":false,"metricName":"# contigs (>= 25000 bp)"},{"values":[0,0,0,0,0,0,0,0,0,0,0,0,0,0],"quality":"Equal","isMain":false,"metricName":"# contigs (>= 50000 bp)"},{"values":[36771,36870,36912,36833,36990,37514,30106,36848,36826,36826,36826,36833,36998,37131],"quality":"More is better","isMain":true,"metricName":"Largest contig"},{"values":[36771,36870,36912,36833,36990,37514,30106,36848,36826,36826,36826,36833,36998,37131],"quality":"More is better","isMain":true,"metricName":"Total length"},{"values":[36771,36870,36912,36833,36990,37514,30106,36848,36826,36826,36826,36833,36998,37131],"quality":"More is better","isMain":true,"metricName":"Total length (>= 1000 bp)"},{"values":[36771,36870,36912,36833,36990,37514,30106,36848,36826,36826,36826,36833,36998,37131],"quality":"More is better","isMain":false,"metricName":"Total length (>= 5000 bp)"},{"values":[36771,36870,36912,36833,36990,37514,30106,36848,36826,36826,36826,36833,36998,37131],"quality":"More is better","isMain":true,"metricName":"Total length (>= 10000 bp)"},{"values":[36771,36870,36912,36833,36990,37514,30106,36848,36826,36826,36826,36833,36998,37131],"quality":"More is better","isMain":false,"metricName":"Total length (>= 25000 bp)"},{"values":[0,0,0,0,0,0,0,0,0,0,0,0,0,0],"quality":"More is better","isMain":true,"metricName":"Total length (>= 50000 bp)"},{"values":[36771,36870,36912,36833,36990,37514,30106,36848,36826,36826,36826,36833,36998,37131],"quality":"More is better","isMain":false,"metricName":"N50"},{"values":[36771,36870,36912,36833,36990,37514,30106,36848,36826,36826,36826,36833,36998,37131],"quality":"More is better","isMain":false,"metricName":"N75"},{"values":[1,1,1,1,1,1,1,1,1,1,1,1,1,1],"quality":"Less is better","isMain":false,"metricName":"L50"},{"values":[1,1,1,1,1,1,1,1,1,1,1,1,1,1],"quality":"Less is better","isMain":false,"metricName":"L75"},{"values":["40.51","40.52","40.51","40.52","40.51","40.50","40.42","40.50","40.51","40.48","40.51","40.52","40.51","40.52"],"quality":"Equal","isMain":false,"metricName":"GC (%)"}]],["Predicted genes",[]],["Similarity statistics",[{"values":[0,0,0,0,0,0,0,0,0,0,0,0,0,0],"quality":"Equal","isMain":false,"metricName":"# similar correct contigs"},{"values":[0,0,0,0,0,0,0,0,0,0,0,0,0,0],"quality":"Equal","isMain":false,"metricName":"# similar misassembled blocks"}]],["Reference statistics",[{"values":[37211,37211,37211,37211,37211,37211,37211,37211,37211,37211,37211,37211,37211,37211],"quality":"Equal","isMain":false,"metricName":"Reference length"},{"values":[1,1,1,1,1,1,1,1,1,1,1,1,1,1],"quality":"Equal","isMain":false,"metricName":"Reference fragments"},{"values":["40.53","40.53","40.53","40.53","40.53","40.53","40.53","40.53","40.53","40.53","40.53","40.53","40.53","40.53"],"quality":"Equal","isMain":false,"metricName":"Reference GC (%)"}]]],"referenceName":"KF302033","date":"18 July 2018, Wednesday, 16:01:21","order":[0,1,2,3,4,5,6,7,8,9,10,11,12,13],"assembliesNames":["CLC","IDBA\_UD","MEGAHIT","metaVelvet","MIRA","Ray\_Meta","SOAPdenovo2","SPAdes","SPAdes\_meta","SPAdes\_sc","SPAdes\_sc\_careful","Velvet","vicuna","Geneious"]},{"assembliesWithNs":null,"minContig":500,"report":[["Genome statistics",[{"values":["99.981","99.546","99.981","99.981","99.816","99.981","12.445","99.981","99.981","99.466","99.981","99.981","91.183","100.000"],"quality":"More is better","isMain":true,"metricName":"Genome fraction (%)"},{"values":["1.000","1.005","1.001","1.000","1.009","1.015","0.984","1.001","1.000","1.006","1.000","1.000","1.083","1.184"],"quality":"Less is better","isMain":true,"metricName":"Duplication ratio"},{"values":[129415,128851,129556,129478,128970,129048,977,129492,129470,128748,129470,129478,48827,129441],"quality":"More is better","isMain":true,"metricName":"Largest alignment"},{"values":[129415,128851,129556,129478,129662,131360,16037,129492,129470,128748,129470,129478,127741,153292],"quality":"More is better","isMain":true,"metricName":"Total aligned length"},{"values":[129415,129514,129556,129478,129652,131360,111231,129492,129470,129470,129470,129478,24884,153292],"quality":"More is better","isMain":false,"metricName":"NG50"},{"values":[129415,129514,129556,129478,129652,131360,111231,129492,129470,129470,129470,129478,690,153292],"quality":"More is better","isMain":false,"metricName":"NG75"},{"values":[129415,128851,129556,129478,128970,129048,null,129492,129470,128748,129470,129478,24884,129441],"quality":"More is better","isMain":false,"metricName":"NA50"},{"values":[129415,128851,129556,129478,128970,129048,null,129492,129470,128748,129470,129478,694,129441],"quality":"More is better","isMain":false,"metricName":"NA75"},{"values":[129415,128851,129556,129478,128970,129048,null,129492,129470,128748,129470,129478,24884,129441],"quality":"More is better","isMain":true,"metricName":"NGA50"},{"values":[129415,128851,129556,129478,128970,129048,null,129492,129470,128748,129470,129478,690,129441],"quality":"More is better","isMain":false,"metricName":"NGA75"},{"values":[1,1,1,1,1,1,1,1,1,1,1,1,2,1],"quality":"Less is better","isMain":false,"metricName":"LG50"},{"values":[1,1,1,1,1,1,1,1,1,1,1,1,17,1],"quality":"Less is better","isMain":false,"metricName":"LG75"},{"values":[1,1,1,1,1,1,null,1,1,1,1,1,2,1],"quality":"Less is better","isMain":false,"metricName":"LA50"},{"values":[1,1,1,1,1,1,null,1,1,1,1,1,16,1],"quality":"Less is better","isMain":false,"metricName":"LA75"},{"values":[1,1,1,1,1,1,null,1,1,1,1,1,2,1],"quality":"Less is better","isMain":true,"metricName":"LGA50"},{"values":[1,1,1,1,1,1,null,1,1,1,1,1,17,1],"quality":"Less is better","isMain":false,"metricName":"LGA75"}]],["Misassemblies",[{"values":[0,0,0,0,0,1,0,0,0,0,0,0,0,2],"quality":"Less is better","isMain":true,"metricName":"# misassemblies"},{"values":[0,0,0,0,0,1,0,0,0,0,0,0,0,2],"quality":"Less is better","isMain":false,"metricName":" # relocations"},{"values":[0,0,0,0,0,0,0,0,0,0,0,0,0,0],"quality":"Less is better","isMain":false,"metricName":" # translocations"},{"values":[0,0,0,0,0,0,0,0,0,0,0,0,0,0],"quality":"Less is better","isMain":false,"metricName":" # inversions"},{"values":[0,0,0,0,0,1,0,0,0,0,0,0,0,1],"quality":"Less is better","isMain":false,"metricName":"# misassembled contigs"},{"values":[0,0,0,0,0,131360,0,0,0,0,0,0,0,153292],"quality":"Less is better","isMain":true,"metricName":"Misassembled contigs length"},{"values":[0,0,1,0,0,0,0,0,0,0,0,0,0,0],"quality":"Less is better","isMain":false,"metricName":"# local misassemblies"},{"values":[0,0,0,0,0,0,1,0,0,0,0,0,0,0],"quality":"Less is better","isMain":false,"metricName":"# unaligned mis. contigs"}]],["Unaligned",[{"values":[0,0,0,0,0,0,0,0,0,0,0,0,0,0],"quality":"Less is better","isMain":false,"metricName":"# fully unaligned contigs"},{"values":[0,0,0,0,0,0,0,0,0,0,0,0,0,0],"quality":"Less is better","isMain":false,"metricName":"Fully unaligned length"},{"values":[0,0,0,0,0,0,1,0,0,0,0,0,0,0],"quality":"Less is better","isMain":false,"metricName":"# partially unaligned contigs"},{"values":[0,0,0,0,0,0,95386,0,0,0,0,0,0,0],"quality":"Less is better","isMain":false,"metricName":"Partially unaligned length"}]],["Mismatches",[{"values":[0,0,0,0,0,0,5,0,0,0,0,0,1,0],"quality":"Less is better","isMain":false,"metricName":"# mismatches"},{"values":[1,0,0,2,0,0,10,1,1,0,1,2,1,1],"quality":"Less is better","isMain":false,"metricName":"# indels"},{"values":[24,0,0,63,0,0,78,77,55,0,55,63,24,2],"quality":"Less is better","isMain":false,"metricName":"Indels length"},{"values":["0.00","0.00","0.00","0.00","0.00","0.00","31.04","0.00","0.00","0.00","0.00","0.00","0.85","0.00"],"quality":"Less is better","isMain":true,"metricName":"# mismatches per 100 kbp"},{"values":["0.77","0.00","0.00","1.55","0.00","0.00","62.08","0.77","0.77","0.00","0.77","1.55","0.85","0.77"],"quality":"Less is better","isMain":true,"metricName":"# indels per 100 kbp"},{"values":[0,0,0,1,0,0,7,0,0,0,0,1,0,1],"quality":"Less is better","isMain":false,"metricName":" # indels (<= 5 bp)"},{"values":[1,0,0,1,0,0,3,1,1,0,1,1,1,0],"quality":"Less is better","isMain":false,"metricName":" # indels (> 5 bp)"},{"values":[0,0,0,0,0,0,35487,0,0,0,0,0,0,57],"quality":"Less is better","isMain":false,"metricName":"# N's"},{"values":["0.00","0.00","0.00","0.00","0.00","0.00","31903.88","0.00","0.00","0.00","0.00","0.00","0.00","37.18"],"quality":"Less is better","isMain":true,"metricName":"# N's per 100 kbp"}]],["Statistics without reference",[{"values":[1,1,1,1,2,1,1,1,1,1,1,1,74,1],"quality":"Equal","isMain":true,"metricName":"# contigs"},{"values":[1,1,1,1,1,1,1,1,1,1,1,1,10,1],"quality":"Equal","isMain":false,"metricName":"# contigs (>= 1000 bp)"},{"values":[1,1,1,1,1,1,1,1,1,1,1,1,3,1],"quality":"Equal","isMain":false,"metricName":"# contigs (>= 5000 bp)"},{"values":[1,1,1,1,1,1,1,1,1,1,1,1,2,1],"quality":"Equal","isMain":false,"metricName":"# contigs (>= 10000 bp)"},{"values":[1,1,1,1,1,1,1,1,1,1,1,1,1,1],"quality":"Equal","isMain":false,"metricName":"# contigs (>= 25000 bp)"},{"values":[1,1,1,1,1,1,1,1,1,1,1,1,0,1],"quality":"Equal","isMain":false,"metricName":"# contigs (>= 50000 bp)"},{"values":[129415,129514,129556,129478,129652,131360,111231,129492,129470,129470,129470,129478,48827,153292],"quality":"More is better","isMain":true,"metricName":"Largest contig"},{"values":[129415,129514,129556,129478,130344,131360,111231,129492,129470,129470,129470,129478,127774,153292],"quality":"More is better","isMain":true,"metricName":"Total length"},{"values":[129415,129514,129556,129478,129652,131360,111231,129492,129470,129470,129470,129478,91928,153292],"quality":"More is better","isMain":true,"metricName":"Total length (>= 1000 bp)"},{"values":[129415,129514,129556,129478,129652,131360,111231,129492,129470,129470,129470,129478,79886,153292],"quality":"More is better","isMain":false,"metricName":"Total length (>= 5000 bp)"},{"values":[129415,129514,129556,129478,129652,131360,111231,129492,129470,129470,129470,129478,73711,153292],"quality":"More is better","isMain":true,"metricName":"Total length (>= 10000 bp)"},{"values":[129415,129514,129556,129478,129652,131360,111231,129492,129470,129470,129470,129478,48827,153292],"quality":"More is better","isMain":false,"metricName":"Total length (>= 25000 bp)"},{"values":[129415,129514,129556,129478,129652,131360,111231,129492,129470,129470,129470,129478,0,153292],"quality":"More is better","isMain":true,"metricName":"Total length (>= 50000 bp)"},{"values":[129415,129514,129556,129478,129652,131360,111231,129492,129470,129470,129470,129478,24884,153292],"quality":"More is better","isMain":false,"metricName":"N50"},{"values":[129415,129514,129556,129478,129652,131360,111231,129492,129470,129470,129470,129478,694,153292],"quality":"More is better","isMain":false,"metricName":"N75"},{"values":[1,1,1,1,1,1,1,1,1,1,1,1,2,1],"quality":"Less is better","isMain":false,"metricName":"L50"},{"values":[1,1,1,1,1,1,1,1,1,1,1,1,16,1],"quality":"Less is better","isMain":false,"metricName":"L75"},{"values":["35.73","35.73","35.73","35.73","35.71","35.65","35.75","35.73","35.74","35.74","35.73","35.73","35.77","35.44"],"quality":"Equal","isMain":false,"metricName":"GC (%)"}]],["Predicted genes",[]],["Similarity statistics",[{"values":[0,0,0,0,0,0,0,0,0,0,0,0,0,0],"quality":"Equal","isMain":false,"metricName":"# similar correct contigs"},{"values":[0,0,0,0,0,0,0,0,0,0,0,0,0,0],"quality":"Equal","isMain":false,"metricName":"# similar misassembled blocks"}]],["Reference statistics",[{"values":[129439,129439,129439,129439,129439,129439,129439,129439,129439,129439,129439,129439,129439,129439],"quality":"Equal","isMain":false,"metricName":"Reference length"},{"values":[1,1,1,1,1,1,1,1,1,1,1,1,1,1],"quality":"Equal","isMain":false,"metricName":"Reference fragments"},{"values":["35.73","35.73","35.73","35.73","35.73","35.73","35.73","35.73","35.73","35.73","35.73","35.73","35.73","35.73"],"quality":"Equal","isMain":false,"metricName":"Reference GC (%)"}]]],"referenceName":"KF302034","date":"18 July 2018, Wednesday, 16:01:37","order":[0,1,2,3,4,5,6,7,8,9,10,11,12,13],"assembliesNames":["CLC","IDBA\_UD","MEGAHIT","metaVelvet","MIRA","Ray\_Meta","SOAPdenovo2","SPAdes","SPAdes\_meta","SPAdes\_sc","SPAdes\_sc\_careful","Velvet","vicuna","Geneious"]},{"assembliesWithNs":null,"minContig":500,"report":[["Genome statistics",[{"values":["99.558","99.558","99.584","99.567","99.558","99.558","99.558","99.558","3.733","99.558","99.558","99.558","99.556","99.558","99.558","99.558"],"quality":"More is better","isMain":true,"metricName":"Genome fraction (%)"},{"values":["1.000","1.000","1.974","1.000","1.001","1.000","1.344","1.000","1.703","1.003","1.002","1.000","1.001","1.000","1.001","1.002"],"quality":"Less is better","isMain":true,"metricName":"Duplication ratio"},{"values":[35176,35176,35176,35179,35176,35177,7023,35176,350,35176,35176,35176,35175,35177,35176,35176],"quality":"More is better","isMain":true,"metricName":"Largest alignment"},{"values":[35176,35176,48992,35179,35176,35177,47238,35176,1311,35176,35176,35176,35175,35177,35176,35176],"quality":"More is better","isMain":true,"metricName":"Total aligned length"},{"values":[35177,35176,35384,35181,35193,35177,3869,35176,null,35292,35251,35176,35214,35177,35214,35227],"quality":"More is better","isMain":false,"metricName":"NG50"},{"values":[35177,35176,35384,35181,35193,35177,2597,35176,null,35292,35251,35176,35214,35177,35214,35227],"quality":"More is better","isMain":false,"metricName":"NG75"},{"values":[35176,35176,null,35179,35176,35177,2770,35176,null,35176,35176,35176,35175,35177,35176,35176],"quality":"More is better","isMain":false,"metricName":"NA50"},{"values":[35176,35176,null,35179,35176,35177,1045,35176,null,35176,35176,35176,35175,35177,35176,35176],"quality":"More is better","isMain":false,"metricName":"NA75"},{"values":[35176,35176,35176,35179,35176,35177,3869,35176,null,35176,35176,35176,35175,35177,35176,35176],"quality":"More is better","isMain":true,"metricName":"NGA50"},{"values":[35176,35176,35176,35179,35176,35177,2597,35176,null,35176,35176,35176,35175,35177,35176,35176],"quality":"More is better","isMain":false,"metricName":"NGA75"},{"values":[1,1,1,1,1,1,4,1,null,1,1,1,1,1,1,1],"quality":"Less is better","isMain":false,"metricName":"LG50"},{"values":[1,1,1,1,1,1,7,1,null,1,1,1,1,1,1,1],"quality":"Less is better","isMain":false,"metricName":"LG75"},{"values":[1,1,null,1,1,1,6,1,null,1,1,1,1,1,1,1],"quality":"Less is better","isMain":false,"metricName":"LA50"},{"values":[1,1,null,1,1,1,13,1,null,1,1,1,1,1,1,1],"quality":"Less is better","isMain":false,"metricName":"LA75"},{"values":[1,1,1,1,1,1,4,1,null,1,1,1,1,1,1,1],"quality":"Less is better","isMain":true,"metricName":"LGA50"},{"values":[1,1,1,1,1,1,7,1,null,1,1,1,1,1,1,1],"quality":"Less is better","isMain":false,"metricName":"LGA75"}]],["Misassemblies",[{"values":[0,0,0,0,0,0,0,0,0,0,0,0,0,0,0,0],"quality":"Less is better","isMain":true,"metricName":"# misassemblies"},{"values":[0,0,0,0,0,0,0,0,0,0,0,0,0,0,0,0],"quality":"Less is better","isMain":false,"metricName":" # relocations"},{"values":[0,0,0,0,0,0,0,0,0,0,0,0,0,0,0,0],"quality":"Less is better","isMain":false,"metricName":" # translocations"},{"values":[0,0,0,0,0,0,0,0,0,0,0,0,0,0,0,0],"quality":"Less is better","isMain":false,"metricName":" # inversions"},{"values":[0,0,0,0,0,0,0,0,0,0,0,0,0,0,0,0],"quality":"Less is better","isMain":false,"metricName":"# misassembled contigs"},{"values":[0,0,0,0,0,0,0,0,0,0,0,0,0,0,0,0],"quality":"Less is better","isMain":true,"metricName":"Misassembled contigs length"},{"values":[0,0,0,0,0,0,0,0,1,0,0,0,0,0,0,0],"quality":"Less is better","isMain":false,"metricName":"# local misassemblies"},{"values":[0,0,5,0,0,0,0,0,2,0,0,0,0,0,0,0],"quality":"Less is better","isMain":false,"metricName":"# unaligned mis. contigs"}]],["Unaligned",[{"values":[0,0,0,0,0,0,0,0,0,0,0,0,0,0,0,0],"quality":"Less is better","isMain":false,"metricName":"# fully unaligned contigs"},{"values":[0,0,0,0,0,0,0,0,0,0,0,0,0,0,0,0],"quality":"Less is better","isMain":false,"metricName":"Fully unaligned length"},{"values":[0,0,97,0,0,0,0,0,1,0,0,0,0,0,0,0],"quality":"Less is better","isMain":false,"metricName":"# partially unaligned contigs"},{"values":[0,0,72903,0,0,0,0,0,6864,0,0,0,0,0,0,0],"quality":"Less is better","isMain":false,"metricName":"Partially unaligned length"}]],["Mismatches",[{"values":[0,0,0,0,0,0,1,0,3,0,0,0,0,0,0,0],"quality":"Less is better","isMain":false,"metricName":"# mismatches"},{"values":[2,2,2,2,2,3,3,2,2,2,2,2,2,3,2,2],"quality":"Less is better","isMain":false,"metricName":"# indels"},{"values":[2,2,2,2,2,3,3,2,8,2,2,2,2,3,2,2],"quality":"Less is better","isMain":false,"metricName":"Indels length"},{"values":["0.00","0.00","0.00","0.00","0.00","0.00","2.84","0.00","227.45","0.00","0.00","0.00","0.00","0.00","0.00","0.00"],"quality":"Less is better","isMain":true,"metricName":"# mismatches per 100 kbp"},{"values":["5.69","5.69","5.68","5.69","5.69","8.53","8.53","5.69","151.63","5.69","5.69","5.69","5.69","8.53","5.69","5.69"],"quality":"Less is better","isMain":true,"metricName":"# indels per 100 kbp"},{"values":[2,2,2,2,2,3,3,2,1,2,2,2,2,3,2,2],"quality":"Less is better","isMain":false,"metricName":" # indels (<= 5 bp)"},{"values":[0,0,0,0,0,0,0,0,1,0,0,0,0,0,0,0],"quality":"Less is better","isMain":false,"metricName":" # indels (> 5 bp)"},{"values":[0,0,0,0,0,0,1,0,4361,0,0,0,0,0,0,22],"quality":"Less is better","isMain":false,"metricName":"# N's"},{"values":["0.00","0.00","0.00","0.00","0.00","0.00","2.12","0.00","47870.47","0.00","0.00","0.00","0.00","0.00","0.00","62.45"],"quality":"Less is better","isMain":true,"metricName":"# N's per 100 kbp"}]],["Statistics without reference",[{"values":[1,1,144,1,1,1,29,1,3,1,1,1,1,1,1,1],"quality":"Equal","isMain":true,"metricName":"# contigs"},{"values":[1,1,21,1,1,1,14,1,1,1,1,1,1,1,1,1],"quality":"Equal","isMain":false,"metricName":"# contigs (>= 1000 bp)"},{"values":[1,1,1,1,1,1,1,1,1,1,1,1,1,1,1,1],"quality":"Equal","isMain":false,"metricName":"# contigs (>= 5000 bp)"},{"values":[1,1,1,1,1,1,0,1,0,1,1,1,1,1,1,1],"quality":"Equal","isMain":false,"metricName":"# contigs (>= 10000 bp)"},{"values":[1,1,1,1,1,1,0,1,0,1,1,1,1,1,1,1],"quality":"Equal","isMain":false,"metricName":"# contigs (>= 25000 bp)"},{"values":[0,0,0,0,0,0,0,0,0,0,0,0,0,0,0,0],"quality":"Equal","isMain":false,"metricName":"# contigs (>= 50000 bp)"},{"values":[35177,35176,35384,35181,35193,35177,7023,35176,7679,35292,35251,35176,35214,35177,35214,35227],"quality":"More is better","isMain":true,"metricName":"Largest contig"},{"values":[35177,35176,142352,35181,35193,35177,47270,35176,9110,35292,35251,35176,35214,35177,35214,35227],"quality":"More is better","isMain":true,"metricName":"Total length"},{"values":[35177,35176,60999,35181,35193,35177,37000,35176,7679,35292,35251,35176,35214,35177,35214,35227],"quality":"More is better","isMain":true,"metricName":"Total length (>= 1000 bp)"},{"values":[35177,35176,35384,35181,35193,35177,7023,35176,7679,35292,35251,35176,35214,35177,35214,35227],"quality":"More is better","isMain":false,"metricName":"Total length (>= 5000 bp)"},{"values":[35177,35176,35384,35181,35193,35177,0,35176,0,35292,35251,35176,35214,35177,35214,35227],"quality":"More is better","isMain":true,"metricName":"Total length (>= 10000 bp)"},{"values":[35177,35176,35384,35181,35193,35177,0,35176,0,35292,35251,35176,35214,35177,35214,35227],"quality":"More is better","isMain":false,"metricName":"Total length (>= 25000 bp)"},{"values":[0,0,0,0,0,0,0,0,0,0,0,0,0,0,0,0],"quality":"More is better","isMain":true,"metricName":"Total length (>= 50000 bp)"},{"values":[35177,35176,896,35181,35193,35177,2770,35176,7679,35292,35251,35176,35214,35177,35214,35227],"quality":"More is better","isMain":false,"metricName":"N50"},{"values":[35177,35176,638,35181,35193,35177,1045,35176,7679,35292,35251,35176,35214,35177,35214,35227],"quality":"More is better","isMain":false,"metricName":"N75"},{"values":[1,1,32,1,1,1,6,1,1,1,1,1,1,1,1,1],"quality":"Less is better","isMain":false,"metricName":"L50"},{"values":[1,1,81,1,1,1,13,1,1,1,1,1,1,1,1,1],"quality":"Less is better","isMain":false,"metricName":"L75"},{"values":["44.91","44.91","40.43","44.91","44.90","44.91","44.62","44.91","46.39","44.88","44.91","44.91","44.91","44.91","44.88","44.88"],"quality":"Equal","isMain":false,"metricName":"GC (%)"}]],["Predicted genes",[]],["Similarity statistics",[{"values":[1,1,1,1,1,1,0,1,0,1,1,1,1,1,1,1],"quality":"Equal","isMain":false,"metricName":"# similar correct contigs"},{"values":[0,0,0,0,0,0,0,0,0,0,0,0,0,0,0,0],"quality":"Equal","isMain":false,"metricName":"# similar misassembled blocks"}]],["Reference statistics",[{"values":[35330,35330,35330,35330,35330,35330,35330,35330,35330,35330,35330,35330,35330,35330,35330,35330],"quality":"Equal","isMain":false,"metricName":"Reference length"},{"values":[1,1,1,1,1,1,1,1,1,1,1,1,1,1,1,1],"quality":"Equal","isMain":false,"metricName":"Reference fragments"},{"values":["44.87","44.87","44.87","44.87","44.87","44.87","44.87","44.87","44.87","44.87","44.87","44.87","44.87","44.87","44.87","44.87"],"quality":"Equal","isMain":false,"metricName":"Reference GC (%)"}]]],"referenceName":"KF302035","date":"18 July 2018, Wednesday, 16:01:47","order":[0,1,2,3,4,5,6,7,8,9,10,11,12,13,14,15],"assembliesNames":["ABySS\_63","ABySS\_127","CLC","IDBA\_UD","MEGAHIT","metaVelvet","MIRA","Ray\_Meta","SOAPdenovo2","SPAdes","SPAdes\_meta","SPAdes\_sc","SPAdes\_sc\_careful","Velvet","vicuna","Geneious"]},{"assembliesWithNs":null,"minContig":500,"report":[["Genome statistics",[{"values":["98.686","98.744","98.744","89.743","98.744","98.744","6.318","98.744","98.744","98.744","98.744","89.743","54.824","98.744"],"quality":"More is better","isMain":true,"metricName":"Genome fraction (%)"},{"values":["1.001","1.003","1.004","1.000","1.007","1.024","0.981","1.002","1.001","1.001","1.001","1.000","1.216","1.010"],"quality":"Less is better","isMain":true,"metricName":"Duplication ratio"},{"values":[37706,37827,37869,20798,37981,38637,933,37805,37783,37783,37783,20798,851,38107],"quality":"More is better","isMain":true,"metricName":"Largest alignment"},{"values":[37706,37827,37869,34297,37981,38637,2368,37805,37783,37783,37783,34297,25452,38107],"quality":"More is better","isMain":true,"metricName":"Total aligned length"},{"values":[37728,37827,37869,20798,37981,38637,36485,37805,37783,37783,37783,20798,510,38107],"quality":"More is better","isMain":false,"metricName":"NG50"},{"values":[37728,37827,37869,4056,37981,38637,36485,37805,37783,37783,37783,4056,null,38107],"quality":"More is better","isMain":false,"metricName":"NG75"},{"values":[37706,37827,37869,20798,37981,38637,null,37805,37783,37783,37783,20798,535,38107],"quality":"More is better","isMain":false,"metricName":"NA50"},{"values":[37706,37827,37869,6359,37981,38637,null,37805,37783,37783,37783,6359,510,38107],"quality":"More is better","isMain":false,"metricName":"NA75"},{"values":[37706,37827,37869,20798,37981,38637,null,37805,37783,37783,37783,20798,510,38107],"quality":"More is better","isMain":true,"metricName":"NGA50"},{"values":[37706,37827,37869,4056,37981,38637,null,37805,37783,37783,37783,4056,null,38107],"quality":"More is better","isMain":false,"metricName":"NGA75"},{"values":[1,1,1,1,1,1,1,1,1,1,1,1,33,1],"quality":"Less is better","isMain":false,"metricName":"LG50"},{"values":[1,1,1,3,1,1,1,1,1,1,1,3,null,1],"quality":"Less is better","isMain":false,"metricName":"LG75"},{"values":[1,1,1,1,1,1,null,1,1,1,1,1,21,1],"quality":"Less is better","isMain":false,"metricName":"LA50"},{"values":[1,1,1,2,1,1,null,1,1,1,1,2,33,1],"quality":"Less is better","isMain":false,"metricName":"LA75"},{"values":[1,1,1,1,1,1,null,1,1,1,1,1,33,1],"quality":"Less is better","isMain":true,"metricName":"LGA50"},{"values":[1,1,1,3,1,1,null,1,1,1,1,3,null,1],"quality":"Less is better","isMain":false,"metricName":"LGA75"}]],["Misassemblies",[{"values":[0,0,0,0,0,0,0,0,0,0,0,0,0,0],"quality":"Less is better","isMain":true,"metricName":"# misassemblies"},{"values":[0,0,0,0,0,0,0,0,0,0,0,0,0,0],"quality":"Less is better","isMain":false,"metricName":" # relocations"},{"values":[0,0,0,0,0,0,0,0,0,0,0,0,0,0],"quality":"Less is better","isMain":false,"metricName":" # translocations"},{"values":[0,0,0,0,0,0,0,0,0,0,0,0,0,0],"quality":"Less is better","isMain":false,"metricName":" # inversions"},{"values":[0,0,0,0,0,0,0,0,0,0,0,0,0,0],"quality":"Less is better","isMain":false,"metricName":"# misassembled contigs"},{"values":[0,0,0,0,0,0,0,0,0,0,0,0,0,0],"quality":"Less is better","isMain":true,"metricName":"Misassembled contigs length"},{"values":[1,1,1,1,1,1,0,1,1,1,1,1,0,1],"quality":"Less is better","isMain":false,"metricName":"# local misassemblies"},{"values":[0,0,0,0,0,0,1,0,0,0,0,0,0,0],"quality":"Less is better","isMain":false,"metricName":"# unaligned mis. contigs"}]],["Unaligned",[{"values":[0,0,0,0,0,0,0,0,0,0,0,0,0,0],"quality":"Less is better","isMain":false,"metricName":"# fully unaligned contigs"},{"values":[0,0,0,0,0,0,0,0,0,0,0,0,0,0],"quality":"Less is better","isMain":false,"metricName":"Fully unaligned length"},{"values":[0,0,0,0,0,0,1,0,0,0,0,0,0,0],"quality":"Less is better","isMain":false,"metricName":"# partially unaligned contigs"},{"values":[0,0,0,0,0,0,34117,0,0,0,0,0,0,0],"quality":"Less is better","isMain":false,"metricName":"Partially unaligned length"}]],["Mismatches",[{"values":[0,0,0,0,0,0,0,0,0,0,0,0,1,0],"quality":"Less is better","isMain":false,"metricName":"# mismatches"},{"values":[0,0,0,0,0,0,2,0,0,0,0,0,0,0],"quality":"Less is better","isMain":false,"metricName":"# indels"},{"values":[0,0,0,0,0,0,46,0,0,0,0,0,0,0],"quality":"Less is better","isMain":false,"metricName":"Indels length"},{"values":["0.00","0.00","0.00","0.00","0.00","0.00","0.00","0.00","0.00","0.00","0.00","0.00","4.77","0.00"],"quality":"Less is better","isMain":true,"metricName":"# mismatches per 100 kbp"},{"values":["0.00","0.00","0.00","0.00","0.00","0.00","82.85","0.00","0.00","0.00","0.00","0.00","0.00","0.00"],"quality":"Less is better","isMain":true,"metricName":"# indels per 100 kbp"},{"values":[0,0,0,0,0,0,0,0,0,0,0,0,0,0],"quality":"Less is better","isMain":false,"metricName":" # indels (<= 5 bp)"},{"values":[0,0,0,0,0,0,2,0,0,0,0,0,0,0],"quality":"Less is better","isMain":false,"metricName":" # indels (> 5 bp)"},{"values":[0,0,0,0,0,0,3043,0,0,0,0,0,0,0],"quality":"Less is better","isMain":false,"metricName":"# N's"},{"values":["0.00","0.00","0.00","0.00","0.00","0.00","8340.41","0.00","0.00","0.00","0.00","0.00","0.00","0.00"],"quality":"Less is better","isMain":true,"metricName":"# N's per 100 kbp"}]],["Statistics without reference",[{"values":[1,1,1,5,1,1,1,1,1,1,1,5,45,1],"quality":"Equal","isMain":true,"metricName":"# contigs"},{"values":[1,1,1,4,1,1,1,1,1,1,1,4,0,1],"quality":"Equal","isMain":false,"metricName":"# contigs (>= 1000 bp)"},{"values":[1,1,1,2,1,1,1,1,1,1,1,2,0,1],"quality":"Equal","isMain":false,"metricName":"# contigs (>= 5000 bp)"},{"values":[1,1,1,1,1,1,1,1,1,1,1,1,0,1],"quality":"Equal","isMain":false,"metricName":"# contigs (>= 10000 bp)"},{"values":[1,1,1,0,1,1,1,1,1,1,1,0,0,1],"quality":"Equal","isMain":false,"metricName":"# contigs (>= 25000 bp)"},{"values":[0,0,0,0,0,0,0,0,0,0,0,0,0,0],"quality":"Equal","isMain":false,"metricName":"# contigs (>= 50000 bp)"},{"values":[37728,37827,37869,20798,37981,38637,36485,37805,37783,37783,37783,20798,851,38107],"quality":"More is better","isMain":true,"metricName":"Largest contig"},{"values":[37728,37827,37869,34297,37981,38637,36485,37805,37783,37783,37783,34297,25469,38107],"quality":"More is better","isMain":true,"metricName":"Total length"},{"values":[37728,37827,37869,33546,37981,38637,36485,37805,37783,37783,37783,33546,0,38107],"quality":"More is better","isMain":true,"metricName":"Total length (>= 1000 bp)"},{"values":[37728,37827,37869,27157,37981,38637,36485,37805,37783,37783,37783,27157,0,38107],"quality":"More is better","isMain":false,"metricName":"Total length (>= 5000 bp)"},{"values":[37728,37827,37869,20798,37981,38637,36485,37805,37783,37783,37783,20798,0,38107],"quality":"More is better","isMain":true,"metricName":"Total length (>= 10000 bp)"},{"values":[37728,37827,37869,0,37981,38637,36485,37805,37783,37783,37783,0,0,38107],"quality":"More is better","isMain":false,"metricName":"Total length (>= 25000 bp)"},{"values":[0,0,0,0,0,0,0,0,0,0,0,0,0,0],"quality":"More is better","isMain":true,"metricName":"Total length (>= 50000 bp)"},{"values":[37728,37827,37869,20798,37981,38637,36485,37805,37783,37783,37783,20798,535,38107],"quality":"More is better","isMain":false,"metricName":"N50"},{"values":[37728,37827,37869,6359,37981,38637,36485,37805,37783,37783,37783,6359,510,38107],"quality":"More is better","isMain":false,"metricName":"N75"},{"values":[1,1,1,1,1,1,1,1,1,1,1,1,21,1],"quality":"Less is better","isMain":false,"metricName":"L50"},{"values":[1,1,1,2,1,1,1,1,1,1,1,2,33,1],"quality":"Less is better","isMain":false,"metricName":"L75"},{"values":["40.21","40.22","40.21","40.16","40.22","40.15","40.19","40.21","40.21","40.20","40.21","40.16","39.81","40.22"],"quality":"Equal","isMain":false,"metricName":"GC (%)"}]],["Predicted genes",[]],["Similarity statistics",[{"values":[0,0,0,0,0,0,0,0,0,0,0,0,0,0],"quality":"Equal","isMain":false,"metricName":"# similar correct contigs"},{"values":[0,0,0,0,0,0,0,0,0,0,0,0,0,0],"quality":"Equal","isMain":false,"metricName":"# similar misassembled blocks"}]],["Reference statistics",[{"values":[38208,38208,38208,38208,38208,38208,38208,38208,38208,38208,38208,38208,38208,38208],"quality":"Equal","isMain":false,"metricName":"Reference length"},{"values":[1,1,1,1,1,1,1,1,1,1,1,1,1,1],"quality":"Equal","isMain":false,"metricName":"Reference fragments"},{"values":["40.23","40.23","40.23","40.23","40.23","40.23","40.23","40.23","40.23","40.23","40.23","40.23","40.23","40.23"],"quality":"Equal","isMain":false,"metricName":"Reference GC (%)"}]]],"referenceName":"KF302036","date":"18 July 2018, Wednesday, 16:01:55","order":[0,1,2,3,4,5,6,7,8,9,10,11,12,13],"assembliesNames":["CLC","IDBA\_UD","MEGAHIT","metaVelvet","MIRA","Ray\_Meta","SOAPdenovo2","SPAdes","SPAdes\_meta","SPAdes\_sc","SPAdes\_sc\_careful","Velvet","vicuna","Geneious"]},{"assembliesWithNs":null,"minContig":500,"report":[["Genome statistics",[{"values":["98.925","98.925","98.925","98.925","98.925","98.925","6.615","98.925","98.925","98.925","98.925","98.925","51.054","98.925"],"quality":"More is better","isMain":true,"metricName":"Genome fraction (%)"},{"values":["1.000","1.002","1.003","1.000","1.003","1.024","0.977","1.002","1.001","1.001","1.001","1.000","1.160","1.006"],"quality":"Less is better","isMain":true,"metricName":"Duplication ratio"},{"values":[44551,44650,44692,44558,44678,26064,1075,44628,44606,44606,44606,44558,839,44829],"quality":"More is better","isMain":true,"metricName":"Largest alignment"},{"values":[44551,44650,44692,44558,44678,45598,2910,44628,44606,44606,44606,44558,26656,44829],"quality":"More is better","isMain":true,"metricName":"Total aligned length"},{"values":[44551,44650,44692,44558,44678,45598,43295,44628,44606,44606,44606,44558,507,44829],"quality":"More is better","isMain":false,"metricName":"NG50"},{"values":[44551,44650,44692,44558,44678,45598,43295,44628,44606,44606,44606,44558,null,44829],"quality":"More is better","isMain":false,"metricName":"NG75"},{"values":[44551,44650,44692,44558,44678,26064,null,44628,44606,44606,44606,44558,520,44829],"quality":"More is better","isMain":false,"metricName":"NA50"},{"values":[44551,44650,44692,44558,44678,19534,null,44628,44606,44606,44606,44558,510,44829],"quality":"More is better","isMain":false,"metricName":"NA75"},{"values":[44551,44650,44692,44558,44678,26064,null,44628,44606,44606,44606,44558,507,44829],"quality":"More is better","isMain":true,"metricName":"NGA50"},{"values":[44551,44650,44692,44558,44678,19534,null,44628,44606,44606,44606,44558,null,44829],"quality":"More is better","isMain":false,"metricName":"NGA75"},{"values":[1,1,1,1,1,1,1,1,1,1,1,1,41,1],"quality":"Less is better","isMain":false,"metricName":"LG50"},{"values":[1,1,1,1,1,1,1,1,1,1,1,1,null,1],"quality":"Less is better","isMain":false,"metricName":"LG75"},{"values":[1,1,1,1,1,1,null,1,1,1,1,1,23,1],"quality":"Less is better","isMain":false,"metricName":"LA50"},{"values":[1,1,1,1,1,2,null,1,1,1,1,1,36,1],"quality":"Less is better","isMain":false,"metricName":"LA75"},{"values":[1,1,1,1,1,1,null,1,1,1,1,1,41,1],"quality":"Less is better","isMain":true,"metricName":"LGA50"},{"values":[1,1,1,1,1,2,null,1,1,1,1,1,null,1],"quality":"Less is better","isMain":false,"metricName":"LGA75"}]],["Misassemblies",[{"values":[0,0,0,0,0,1,0,0,0,0,0,0,0,0],"quality":"Less is better","isMain":true,"metricName":"# misassemblies"},{"values":[0,0,0,0,0,1,0,0,0,0,0,0,0,0],"quality":"Less is better","isMain":false,"metricName":" # relocations"},{"values":[0,0,0,0,0,0,0,0,0,0,0,0,0,0],"quality":"Less is better","isMain":false,"metricName":" # translocations"},{"values":[0,0,0,0,0,0,0,0,0,0,0,0,0,0],"quality":"Less is better","isMain":false,"metricName":" # inversions"},{"values":[0,0,0,0,0,1,0,0,0,0,0,0,0,0],"quality":"Less is better","isMain":false,"metricName":"# misassembled contigs"},{"values":[0,0,0,0,0,45598,0,0,0,0,0,0,0,0],"quality":"Less is better","isMain":true,"metricName":"Misassembled contigs length"},{"values":[1,1,1,1,1,0,0,1,1,1,1,1,0,1],"quality":"Less is better","isMain":false,"metricName":"# local misassemblies"},{"values":[0,0,0,0,0,0,1,0,0,0,0,0,0,0],"quality":"Less is better","isMain":false,"metricName":"# unaligned mis. contigs"}]],["Unaligned",[{"values":[0,0,0,0,0,0,0,0,0,0,0,0,0,0],"quality":"Less is better","isMain":false,"metricName":"# fully unaligned contigs"},{"values":[0,0,0,0,0,0,0,0,0,0,0,0,0,0],"quality":"Less is better","isMain":false,"metricName":"Fully unaligned length"},{"values":[0,0,0,0,0,0,1,0,0,0,0,0,0,0],"quality":"Less is better","isMain":false,"metricName":"# partially unaligned contigs"},{"values":[0,0,0,0,0,0,40385,0,0,0,0,0,0,0],"quality":"Less is better","isMain":false,"metricName":"Partially unaligned length"}]],["Mismatches",[{"values":[0,0,0,0,0,0,0,0,0,0,0,0,1,0],"quality":"Less is better","isMain":false,"metricName":"# mismatches"},{"values":[0,0,0,0,0,0,3,0,0,0,0,0,0,0],"quality":"Less is better","isMain":false,"metricName":"# indels"},{"values":[0,0,0,0,0,0,69,0,0,0,0,0,0,0],"quality":"Less is better","isMain":false,"metricName":"Indels length"},{"values":["0.00","0.00","0.00","0.00","0.00","0.00","0.00","0.00","0.00","0.00","0.00","0.00","4.35","0.00"],"quality":"Less is better","isMain":true,"metricName":"# mismatches per 100 kbp"},{"values":["0.00","0.00","0.00","0.00","0.00","0.00","100.70","0.00","0.00","0.00","0.00","0.00","0.00","0.00"],"quality":"Less is better","isMain":true,"metricName":"# indels per 100 kbp"},{"values":[0,0,0,0,0,0,0,0,0,0,0,0,0,0],"quality":"Less is better","isMain":false,"metricName":" # indels (<= 5 bp)"},{"values":[0,0,0,0,0,0,3,0,0,0,0,0,0,0],"quality":"Less is better","isMain":false,"metricName":" # indels (> 5 bp)"},{"values":[0,0,0,0,0,0,4715,0,0,0,0,0,0,0],"quality":"Less is better","isMain":false,"metricName":"# N's"},{"values":["0.00","0.00","0.00","0.00","0.00","0.00","10890.40","0.00","0.00","0.00","0.00","0.00","0.00","0.00"],"quality":"Less is better","isMain":true,"metricName":"# N's per 100 kbp"}]],["Statistics without reference",[{"values":[1,1,1,1,1,1,1,1,1,1,1,1,49,1],"quality":"Equal","isMain":true,"metricName":"# contigs"},{"values":[1,1,1,1,1,1,1,1,1,1,1,1,0,1],"quality":"Equal","isMain":false,"metricName":"# contigs (>= 1000 bp)"},{"values":[1,1,1,1,1,1,1,1,1,1,1,1,0,1],"quality":"Equal","isMain":false,"metricName":"# contigs (>= 5000 bp)"},{"values":[1,1,1,1,1,1,1,1,1,1,1,1,0,1],"quality":"Equal","isMain":false,"metricName":"# contigs (>= 10000 bp)"},{"values":[1,1,1,1,1,1,1,1,1,1,1,1,0,1],"quality":"Equal","isMain":false,"metricName":"# contigs (>= 25000 bp)"},{"values":[0,0,0,0,0,0,0,0,0,0,0,0,0,0],"quality":"Equal","isMain":false,"metricName":"# contigs (>= 50000 bp)"},{"values":[44551,44650,44692,44558,44678,45598,43295,44628,44606,44606,44606,44558,839,44829],"quality":"More is better","isMain":true,"metricName":"Largest contig"},{"values":[44551,44650,44692,44558,44678,45598,43295,44628,44606,44606,44606,44558,26669,44829],"quality":"More is better","isMain":true,"metricName":"Total length"},{"values":[44551,44650,44692,44558,44678,45598,43295,44628,44606,44606,44606,44558,0,44829],"quality":"More is better","isMain":true,"metricName":"Total length (>= 1000 bp)"},{"values":[44551,44650,44692,44558,44678,45598,43295,44628,44606,44606,44606,44558,0,44829],"quality":"More is better","isMain":false,"metricName":"Total length (>= 5000 bp)"},{"values":[44551,44650,44692,44558,44678,45598,43295,44628,44606,44606,44606,44558,0,44829],"quality":"More is better","isMain":true,"metricName":"Total length (>= 10000 bp)"},{"values":[44551,44650,44692,44558,44678,45598,43295,44628,44606,44606,44606,44558,0,44829],"quality":"More is better","isMain":false,"metricName":"Total length (>= 25000 bp)"},{"values":[0,0,0,0,0,0,0,0,0,0,0,0,0,0],"quality":"More is better","isMain":true,"metricName":"Total length (>= 50000 bp)"},{"values":[44551,44650,44692,44558,44678,45598,43295,44628,44606,44606,44606,44558,521,44829],"quality":"More is better","isMain":false,"metricName":"N50"},{"values":[44551,44650,44692,44558,44678,45598,43295,44628,44606,44606,44606,44558,510,44829],"quality":"More is better","isMain":false,"metricName":"N75"},{"values":[1,1,1,1,1,1,1,1,1,1,1,1,23,1],"quality":"Less is better","isMain":false,"metricName":"L50"},{"values":[1,1,1,1,1,1,1,1,1,1,1,1,36,1],"quality":"Less is better","isMain":false,"metricName":"L75"},{"values":["44.65","44.67","44.68","44.66","44.69","44.64","44.64","44.67","44.66","44.66","44.67","44.66","44.41","44.72"],"quality":"Equal","isMain":false,"metricName":"GC (%)"}]],["Predicted genes",[]],["Similarity statistics",[{"values":[0,0,0,0,0,0,0,0,0,0,0,0,0,0],"quality":"Equal","isMain":false,"metricName":"# similar correct contigs"},{"values":[0,0,0,0,0,0,0,0,0,0,0,0,0,0],"quality":"Equal","isMain":false,"metricName":"# similar misassembled blocks"}]],["Reference statistics",[{"values":[45035,45035,45035,45035,45035,45035,45035,45035,45035,45035,45035,45035,45035,45035],"quality":"Equal","isMain":false,"metricName":"Reference length"},{"values":[1,1,1,1,1,1,1,1,1,1,1,1,1,1],"quality":"Equal","isMain":false,"metricName":"Reference fragments"},{"values":["44.67","44.67","44.67","44.67","44.67","44.67","44.67","44.67","44.67","44.67","44.67","44.67","44.67","44.67"],"quality":"Equal","isMain":false,"metricName":"Reference GC (%)"}]]],"referenceName":"KF302037","date":"18 July 2018, Wednesday, 16:02:04","order":[0,1,2,3,4,5,6,7,8,9,10,11,12,13],"assembliesNames":["CLC","IDBA\_UD","MEGAHIT","metaVelvet","MIRA","Ray\_Meta","SOAPdenovo2","SPAdes","SPAdes\_meta","SPAdes\_sc","SPAdes\_sc\_careful","Velvet","vicuna","Geneious"]},{"assembliesWithNs":null,"minContig":500,"report":[["Genome statistics",[{"values":["100.000","100.000","100.000","100.000","100.000","100.000","99.984","100.000","100.000","100.000","100.000","72.187","100.000"],"quality":"More is better","isMain":true,"metricName":"Genome fraction (%)"},{"values":["1.010","1.000","1.017","1.023","1.008","1.069","0.991","1.013","1.008","1.008","1.008","1.182","1.049"],"quality":"Less is better","isMain":true,"metricName":"Duplication ratio"},{"values":[6148,6089,6188,6230,6089,6508,6034,6166,6134,6134,6134,560,6385],"quality":"More is better","isMain":true,"metricName":"Largest alignment"},{"values":[6148,6089,6188,6230,6089,6508,6034,6166,6134,6134,6134,5190,6385],"quality":"More is better","isMain":true,"metricName":"Total aligned length"},{"values":[6148,6089,6188,6230,6135,6508,6034,6166,6134,6134,6134,515,6385],"quality":"More is better","isMain":false,"metricName":"NG50"},{"values":[6148,6089,6188,6230,6135,6508,6034,6166,6134,6134,6134,505,6385],"quality":"More is better","isMain":false,"metricName":"NG75"},{"values":[6148,6089,6188,6230,6089,6508,6034,6166,6134,6134,6134,516,6385],"quality":"More is better","isMain":false,"metricName":"NA50"},{"values":[6148,6089,6188,6230,6089,6508,6034,6166,6134,6134,6134,506,6385],"quality":"More is better","isMain":false,"metricName":"NA75"},{"values":[6148,6089,6188,6230,6089,6508,6034,6166,6134,6134,6134,515,6385],"quality":"More is better","isMain":true,"metricName":"NGA50"},{"values":[6148,6089,6188,6230,6089,6508,6034,6166,6134,6134,6134,505,6385],"quality":"More is better","isMain":false,"metricName":"NGA75"},{"values":[1,1,1,1,1,1,1,1,1,1,1,6,1],"quality":"Less is better","isMain":false,"metricName":"LG50"},{"values":[1,1,1,1,1,1,1,1,1,1,1,9,1],"quality":"Less is better","isMain":false,"metricName":"LG75"},{"values":[1,1,1,1,1,1,1,1,1,1,1,5,1],"quality":"Less is better","isMain":false,"metricName":"LA50"},{"values":[1,1,1,1,1,1,1,1,1,1,1,8,1],"quality":"Less is better","isMain":false,"metricName":"LA75"},{"values":[1,1,1,1,1,1,1,1,1,1,1,6,1],"quality":"Less is better","isMain":true,"metricName":"LGA50"},{"values":[1,1,1,1,1,1,1,1,1,1,1,9,1],"quality":"Less is better","isMain":false,"metricName":"LGA75"}]],["Misassemblies",[{"values":[0,0,0,0,0,0,0,0,0,0,0,0,0],"quality":"Less is better","isMain":true,"metricName":"# misassemblies"},{"values":[0,0,0,0,0,0,0,0,0,0,0,0,0],"quality":"Less is better","isMain":false,"metricName":" # relocations"},{"values":[0,0,0,0,0,0,0,0,0,0,0,0,0],"quality":"Less is better","isMain":false,"metricName":" # translocations"},{"values":[0,0,0,0,0,0,0,0,0,0,0,0,0],"quality":"Less is better","isMain":false,"metricName":" # inversions"},{"values":[0,0,0,0,0,0,0,0,0,0,0,0,0],"quality":"Less is better","isMain":false,"metricName":"# misassembled contigs"},{"values":[0,0,0,0,0,0,0,0,0,0,0,0,0],"quality":"Less is better","isMain":true,"metricName":"Misassembled contigs length"},{"values":[0,0,1,1,0,1,0,0,0,0,0,0,1],"quality":"Less is better","isMain":false,"metricName":"# local misassemblies"},{"values":[0,0,0,0,0,0,0,0,0,0,0,0,0],"quality":"Less is better","isMain":false,"metricName":"# unaligned mis. contigs"}]],["Unaligned",[{"values":[0,0,0,0,0,0,0,0,0,0,0,0,0],"quality":"Less is better","isMain":false,"metricName":"# fully unaligned contigs"},{"values":[0,0,0,0,0,0,0,0,0,0,0,0,0],"quality":"Less is better","isMain":false,"metricName":"Fully unaligned length"},{"values":[0,0,0,0,0,0,0,0,0,0,0,0,0],"quality":"Less is better","isMain":false,"metricName":"# partially unaligned contigs"},{"values":[0,0,0,0,0,0,0,0,0,0,0,0,0],"quality":"Less is better","isMain":false,"metricName":"Partially unaligned length"}]],["Mismatches",[{"values":[9,9,9,9,9,9,9,9,9,9,9,7,9],"quality":"Less is better","isMain":false,"metricName":"# mismatches"},{"values":[3,2,2,2,2,2,5,3,3,3,3,0,3],"quality":"Less is better","isMain":false,"metricName":"# indels"},{"values":[61,2,2,2,2,2,56,79,47,47,47,0,77],"quality":"Less is better","isMain":false,"metricName":"Indels length"},{"values":["147.86","147.86","147.86","147.86","147.86","147.86","147.88","147.86","147.86","147.86","147.86","159.31","147.86"],"quality":"Less is better","isMain":true,"metricName":"# mismatches per 100 kbp"},{"values":["49.29","32.86","32.86","32.86","32.86","32.86","82.16","49.29","49.29","49.29","49.29","0.00","49.29"],"quality":"Less is better","isMain":true,"metricName":"# indels per 100 kbp"},{"values":[2,2,2,2,2,2,2,2,2,2,2,0,2],"quality":"Less is better","isMain":false,"metricName":" # indels (<= 5 bp)"},{"values":[1,0,0,0,0,0,3,1,1,1,1,0,1],"quality":"Less is better","isMain":false,"metricName":" # indels (> 5 bp)"},{"values":[0,0,0,0,0,0,42,0,0,0,0,0,0],"quality":"Less is better","isMain":false,"metricName":"# N's"},{"values":["0.00","0.00","0.00","0.00","0.00","0.00","696.06","0.00","0.00","0.00","0.00","0.00","0.00"],"quality":"Less is better","isMain":true,"metricName":"# N's per 100 kbp"}]],["Statistics without reference",[{"values":[1,1,1,1,1,1,1,1,1,1,1,10,1],"quality":"Equal","isMain":true,"metricName":"# contigs"},{"values":[1,1,1,1,1,1,1,1,1,1,1,0,1],"quality":"Equal","isMain":false,"metricName":"# contigs (>= 1000 bp)"},{"values":[1,1,1,1,1,1,1,1,1,1,1,0,1],"quality":"Equal","isMain":false,"metricName":"# contigs (>= 5000 bp)"},{"values":[0,0,0,0,0,0,0,0,0,0,0,0,0],"quality":"Equal","isMain":false,"metricName":"# contigs (>= 10000 bp)"},{"values":[0,0,0,0,0,0,0,0,0,0,0,0,0],"quality":"Equal","isMain":false,"metricName":"# contigs (>= 25000 bp)"},{"values":[0,0,0,0,0,0,0,0,0,0,0,0,0],"quality":"Equal","isMain":false,"metricName":"# contigs (>= 50000 bp)"},{"values":[6148,6089,6188,6230,6135,6508,6034,6166,6134,6134,6134,562,6385],"quality":"More is better","isMain":true,"metricName":"Largest contig"},{"values":[6148,6089,6188,6230,6135,6508,6034,6166,6134,6134,6134,5192,6385],"quality":"More is better","isMain":true,"metricName":"Total length"},{"values":[6148,6089,6188,6230,6135,6508,6034,6166,6134,6134,6134,0,6385],"quality":"More is better","isMain":true,"metricName":"Total length (>= 1000 bp)"},{"values":[6148,6089,6188,6230,6135,6508,6034,6166,6134,6134,6134,0,6385],"quality":"More is better","isMain":false,"metricName":"Total length (>= 5000 bp)"},{"values":[0,0,0,0,0,0,0,0,0,0,0,0,0],"quality":"More is better","isMain":true,"metricName":"Total length (>= 10000 bp)"},{"values":[0,0,0,0,0,0,0,0,0,0,0,0,0],"quality":"More is better","isMain":false,"metricName":"Total length (>= 25000 bp)"},{"values":[0,0,0,0,0,0,0,0,0,0,0,0,0],"quality":"More is better","isMain":true,"metricName":"Total length (>= 50000 bp)"},{"values":[6148,6089,6188,6230,6135,6508,6034,6166,6134,6134,6134,516,6385],"quality":"More is better","isMain":false,"metricName":"N50"},{"values":[6148,6089,6188,6230,6135,6508,6034,6166,6134,6134,6134,506,6385],"quality":"More is better","isMain":false,"metricName":"N75"},{"values":[1,1,1,1,1,1,1,1,1,1,1,5,1],"quality":"Less is better","isMain":false,"metricName":"L50"},{"values":[1,1,1,1,1,1,1,1,1,1,1,8,1],"quality":"Less is better","isMain":false,"metricName":"L75"},{"values":["45.22","45.21","45.41","45.35","45.12","44.99","45.18","45.31","45.19","45.19","45.19","47.03","44.96"],"quality":"Equal","isMain":false,"metricName":"GC (%)"}]],["Predicted genes",[]],["Similarity statistics",[{"values":[0,0,0,0,0,0,0,0,0,0,0,0,0],"quality":"Equal","isMain":false,"metricName":"# similar correct contigs"},{"values":[0,0,0,0,0,0,0,0,0,0,0,0,0],"quality":"Equal","isMain":false,"metricName":"# similar misassembled blocks"}]],["Reference statistics",[{"values":[6087,6087,6087,6087,6087,6087,6087,6087,6087,6087,6087,6087,6087],"quality":"Equal","isMain":false,"metricName":"Reference length"},{"values":[1,1,1,1,1,1,1,1,1,1,1,1,1],"quality":"Equal","isMain":false,"metricName":"Reference fragments"},{"values":["45.18","45.18","45.18","45.18","45.18","45.18","45.18","45.18","45.18","45.18","45.18","45.18","45.18"],"quality":"Equal","isMain":false,"metricName":"Reference GC (%)"}]]],"referenceName":"NC\_001330","date":"18 July 2018, Wednesday, 16:02:11","order":[0,1,2,3,4,5,6,7,8,9,10,11,12],"assembliesNames":["ABySS\_63","CLC","IDBA\_UD","MEGAHIT","MIRA","Ray\_Meta","SOAPdenovo2","SPAdes","SPAdes\_meta","SPAdes\_sc","SPAdes\_sc\_careful","vicuna","Geneious"]},{"assembliesWithNs":null,"minContig":500,"report":[["Genome statistics",[{"values":["100.000","95.395","100.000","99.610","99.424","97.159","99.424","99.424","99.610","99.610","100.000","99.610","32.882","99.610"],"quality":"More is better","isMain":true,"metricName":"Genome fraction (%)"},{"values":["1.011","1.004","1.000","1.005","1.004","1.000","1.004","1.007","1.005","1.005","1.008","1.013","1.006","1.007"],"quality":"Less is better","isMain":true,"metricName":"Duplication ratio"},{"values":[5445,5158,5386,5365,5355,5233,5355,5373,5365,5365,5431,4783,764,5365],"quality":"More is better","isMain":true,"metricName":"Largest alignment"},{"values":[5445,5158,5386,5365,5355,5233,5355,5373,5365,5365,5431,5410,1771,5365],"quality":"More is better","isMain":true,"metricName":"Total aligned length"},{"values":[5445,5158,5386,5391,5377,5233,5377,5395,5391,5391,5431,4809,null,5402],"quality":"More is better","isMain":false,"metricName":"NG50"},{"values":[5445,5158,5386,5391,5377,5233,5377,5395,5391,5391,5431,4809,null,5402],"quality":"More is better","isMain":false,"metricName":"NG75"},{"values":[5445,5158,5386,5365,5355,5233,5355,5373,5365,5365,5431,4783,506,5365],"quality":"More is better","isMain":false,"metricName":"NA50"},{"values":[5445,5158,5386,5365,5355,5233,5355,5373,5365,5365,5431,4783,501,5365],"quality":"More is better","isMain":false,"metricName":"NA75"},{"values":[5445,5158,5386,5365,5355,5233,5355,5373,5365,5365,5431,4783,null,5365],"quality":"More is better","isMain":true,"metricName":"NGA50"},{"values":[5445,5158,5386,5365,5355,5233,5355,5373,5365,5365,5431,4783,null,5365],"quality":"More is better","isMain":false,"metricName":"NGA75"},{"values":[1,1,1,1,1,1,1,1,1,1,1,1,null,1],"quality":"Less is better","isMain":false,"metricName":"LG50"},{"values":[1,1,1,1,1,1,1,1,1,1,1,1,null,1],"quality":"Less is better","isMain":false,"metricName":"LG75"},{"values":[1,1,1,1,1,1,1,1,1,1,1,1,2,1],"quality":"Less is better","isMain":false,"metricName":"LA50"},{"values":[1,1,1,1,1,1,1,1,1,1,1,1,3,1],"quality":"Less is better","isMain":false,"metricName":"LA75"},{"values":[1,1,1,1,1,1,1,1,1,1,1,1,null,1],"quality":"Less is better","isMain":true,"metricName":"LGA50"},{"values":[1,1,1,1,1,1,1,1,1,1,1,1,null,1],"quality":"Less is better","isMain":false,"metricName":"LGA75"}]],["Misassemblies",[{"values":[0,0,0,0,0,0,0,0,0,0,0,0,0,0],"quality":"Less is better","isMain":true,"metricName":"# misassemblies"},{"values":[0,0,0,0,0,0,0,0,0,0,0,0,0,0],"quality":"Less is better","isMain":false,"metricName":" # relocations"},{"values":[0,0,0,0,0,0,0,0,0,0,0,0,0,0],"quality":"Less is better","isMain":false,"metricName":" # translocations"},{"values":[0,0,0,0,0,0,0,0,0,0,0,0,0,0],"quality":"Less is better","isMain":false,"metricName":" # inversions"},{"values":[0,0,0,0,0,0,0,0,0,0,0,0,0,0],"quality":"Less is better","isMain":false,"metricName":"# misassembled contigs"},{"values":[0,0,0,0,0,0,0,0,0,0,0,0,0,0],"quality":"Less is better","isMain":true,"metricName":"Misassembled contigs length"},{"values":[0,0,0,0,0,0,0,0,0,0,0,0,0,0],"quality":"Less is better","isMain":false,"metricName":"# local misassemblies"},{"values":[0,0,0,0,0,0,0,0,0,0,0,0,0,0],"quality":"Less is better","isMain":false,"metricName":"# unaligned mis. contigs"}]],["Unaligned",[{"values":[0,0,0,0,0,0,0,0,0,0,0,0,0,0],"quality":"Less is better","isMain":false,"metricName":"# fully unaligned contigs"},{"values":[0,0,0,0,0,0,0,0,0,0,0,0,0,0],"quality":"Less is better","isMain":false,"metricName":"Fully unaligned length"},{"values":[0,0,0,0,0,0,0,0,0,0,0,0,0,0],"quality":"Less is better","isMain":false,"metricName":"# partially unaligned contigs"},{"values":[0,0,0,0,0,0,0,0,0,0,0,0,0,0],"quality":"Less is better","isMain":false,"metricName":"Partially unaligned length"}]],["Mismatches",[{"values":[16,16,16,16,16,16,16,16,16,16,16,16,6,16],"quality":"Less is better","isMain":false,"metricName":"# mismatches"},{"values":[1,1,0,0,0,0,0,4,0,0,1,0,0,0],"quality":"Less is better","isMain":false,"metricName":"# indels"},{"values":[59,20,0,0,0,0,0,46,0,0,45,0,0,0],"quality":"Less is better","isMain":false,"metricName":"Indels length"},{"values":["297.07","311.41","297.07","298.23","298.79","305.75","298.79","298.79","298.23","298.23","297.07","298.23","338.79","298.23"],"quality":"Less is better","isMain":true,"metricName":"# mismatches per 100 kbp"},{"values":["18.57","19.46","0.00","0.00","0.00","0.00","0.00","74.70","0.00","0.00","18.57","0.00","0.00","0.00"],"quality":"Less is better","isMain":true,"metricName":"# indels per 100 kbp"},{"values":[0,0,0,0,0,0,0,1,0,0,0,0,0,0],"quality":"Less is better","isMain":false,"metricName":" # indels (<= 5 bp)"},{"values":[1,1,0,0,0,0,0,3,0,0,1,0,0,0],"quality":"Less is better","isMain":false,"metricName":" # indels (> 5 bp)"},{"values":[0,71,0,0,0,0,0,161,0,0,0,0,0,0],"quality":"Less is better","isMain":false,"metricName":"# N's"},{"values":["0.00","1376.50","0.00","0.00","0.00","0.00","0.00","2984.24","0.00","0.00","0.00","0.00","0.00","0.00"],"quality":"Less is better","isMain":true,"metricName":"# N's per 100 kbp"}]],["Statistics without reference",[{"values":[1,1,1,1,1,1,1,1,1,1,1,2,3,1],"quality":"Equal","isMain":true,"metricName":"# contigs"},{"values":[1,1,1,1,1,1,1,1,1,1,1,1,0,1],"quality":"Equal","isMain":false,"metricName":"# contigs (>= 1000 bp)"},{"values":[1,1,1,1,1,1,1,1,1,1,1,0,0,1],"quality":"Equal","isMain":false,"metricName":"# contigs (>= 5000 bp)"},{"values":[0,0,0,0,0,0,0,0,0,0,0,0,0,0],"quality":"Equal","isMain":false,"metricName":"# contigs (>= 10000 bp)"},{"values":[0,0,0,0,0,0,0,0,0,0,0,0,0,0],"quality":"Equal","isMain":false,"metricName":"# contigs (>= 25000 bp)"},{"values":[0,0,0,0,0,0,0,0,0,0,0,0,0,0],"quality":"Equal","isMain":false,"metricName":"# contigs (>= 50000 bp)"},{"values":[5445,5158,5386,5391,5377,5233,5377,5395,5391,5391,5431,4809,764,5402],"quality":"More is better","isMain":true,"metricName":"Largest contig"},{"values":[5445,5158,5386,5391,5377,5233,5377,5395,5391,5391,5431,5436,1782,5402],"quality":"More is better","isMain":true,"metricName":"Total length"},{"values":[5445,5158,5386,5391,5377,5233,5377,5395,5391,5391,5431,4809,0,5402],"quality":"More is better","isMain":true,"metricName":"Total length (>= 1000 bp)"},{"values":[5445,5158,5386,5391,5377,5233,5377,5395,5391,5391,5431,0,0,5402],"quality":"More is better","isMain":false,"metricName":"Total length (>= 5000 bp)"},{"values":[0,0,0,0,0,0,0,0,0,0,0,0,0,0],"quality":"More is better","isMain":true,"metricName":"Total length (>= 10000 bp)"},{"values":[0,0,0,0,0,0,0,0,0,0,0,0,0,0],"quality":"More is better","isMain":false,"metricName":"Total length (>= 25000 bp)"},{"values":[0,0,0,0,0,0,0,0,0,0,0,0,0,0],"quality":"More is better","isMain":true,"metricName":"Total length (>= 50000 bp)"},{"values":[5445,5158,5386,5391,5377,5233,5377,5395,5391,5391,5431,4809,512,5402],"quality":"More is better","isMain":false,"metricName":"N50"},{"values":[5445,5158,5386,5391,5377,5233,5377,5395,5391,5391,5431,4809,506,5402],"quality":"More is better","isMain":false,"metricName":"N75"},{"values":[1,1,1,1,1,1,1,1,1,1,1,1,2,1],"quality":"Less is better","isMain":false,"metricName":"L50"},{"values":[1,1,1,1,1,1,1,1,1,1,1,1,3,1],"quality":"Less is better","isMain":false,"metricName":"L75"},{"values":["44.70","44.64","44.69","44.70","44.71","44.89","44.71","44.75","44.70","44.70","44.67","44.68","46.07","44.67"],"quality":"Equal","isMain":false,"metricName":"GC (%)"}]],["Predicted genes",[]],["Similarity statistics",[{"values":[0,0,0,0,0,0,0,0,0,0,0,0,0,0],"quality":"Equal","isMain":false,"metricName":"# similar correct contigs"},{"values":[0,0,0,0,0,0,0,0,0,0,0,0,0,0],"quality":"Equal","isMain":false,"metricName":"# similar misassembled blocks"}]],["Reference statistics",[{"values":[5386,5386,5386,5386,5386,5386,5386,5386,5386,5386,5386,5386,5386,5386],"quality":"Equal","isMain":false,"metricName":"Reference length"},{"values":[1,1,1,1,1,1,1,1,1,1,1,1,1,1],"quality":"Equal","isMain":false,"metricName":"Reference fragments"},{"values":["44.76","44.76","44.76","44.76","44.76","44.76","44.76","44.76","44.76","44.76","44.76","44.76","44.76","44.76"],"quality":"Equal","isMain":false,"metricName":"Reference GC (%)"}]]],"referenceName":"NC\_001422","date":"18 July 2018, Wednesday, 16:02:19","order":[0,1,2,3,4,5,6,7,8,9,10,11,12,13],"assembliesNames":["ABySS\_63","ABySS\_127","CLC","IDBA\_UD","MEGAHIT","MIRA","Ray\_Meta","SOAPdenovo2","SPAdes","SPAdes\_meta","SPAdes\_sc","SPAdes\_sc\_careful","vicuna","Geneious"]},{"assembliesWithNs":null,"minContig":500,"report":[["Genome statistics",[{"values":["100.000","100.000","100.000","100.000","100.000","99.546","100.000","100.000","0.255","100.000","100.000","100.000","100.000","99.546","100.000","100.000"],"quality":"More is better","isMain":true,"metricName":"Genome fraction (%)"},{"values":["1.004","1.023","1.000","1.003","1.003","1.004","1.093","1.028","5.520","1.004","1.002","1.002","1.002","1.004","1.009","1.436"],"quality":"Less is better","isMain":true,"metricName":"Duplication ratio"},{"values":[38543,40089,39191,39290,38545,22872,39164,23054,100,38545,38521,38521,38521,22872,39552,39166],"quality":"More is better","isMain":true,"metricName":"Largest alignment"},{"values":[39313,40089,39191,39290,39325,39128,42829,40281,100,39345,39277,39277,39277,39128,39552,56245],"quality":"More is better","isMain":true,"metricName":"Total aligned length"},{"values":[38593,40089,39191,39290,38545,22872,39183,40281,null,38545,38521,38521,38521,22872,39552,56262],"quality":"More is better","isMain":false,"metricName":"NG50"},{"values":[38593,40089,39191,39290,38545,15512,39183,40281,null,38545,38521,38521,38521,15512,39552,56262],"quality":"More is better","isMain":false,"metricName":"NG75"},{"values":[38543,40089,39191,39290,38545,22872,39164,23054,null,38545,38521,38521,38521,22872,39552,39166],"quality":"More is better","isMain":false,"metricName":"NA50"},{"values":[38543,40089,39191,39290,38545,15486,39164,17227,null,38545,38521,38521,38521,15486,39552,17079],"quality":"More is better","isMain":false,"metricName":"NA75"},{"values":[38543,40089,39191,39290,38545,22872,39164,23054,null,38545,38521,38521,38521,22872,39552,39166],"quality":"More is better","isMain":true,"metricName":"NGA50"},{"values":[38543,40089,39191,39290,38545,15486,39164,17227,null,38545,38521,38521,38521,15486,39552,39166],"quality":"More is better","isMain":false,"metricName":"NGA75"},{"values":[1,1,1,1,1,1,1,1,null,1,1,1,1,1,1,1],"quality":"Less is better","isMain":false,"metricName":"LG50"},{"values":[1,1,1,1,1,2,1,1,null,1,1,1,1,2,1,1],"quality":"Less is better","isMain":false,"metricName":"LG75"},{"values":[1,1,1,1,1,1,1,1,null,1,1,1,1,1,1,1],"quality":"Less is better","isMain":false,"metricName":"LA50"},{"values":[1,1,1,1,1,2,1,2,null,1,1,1,1,2,1,2],"quality":"Less is better","isMain":false,"metricName":"LA75"},{"values":[1,1,1,1,1,1,1,1,null,1,1,1,1,1,1,1],"quality":"Less is better","isMain":true,"metricName":"LGA50"},{"values":[1,1,1,1,1,2,1,2,null,1,1,1,1,2,1,1],"quality":"Less is better","isMain":false,"metricName":"LGA75"}]],["Misassemblies",[{"values":[0,0,0,0,0,0,0,1,0,0,0,0,0,0,0,1],"quality":"Less is better","isMain":true,"metricName":"# misassemblies"},{"values":[0,0,0,0,0,0,0,1,0,0,0,0,0,0,0,1],"quality":"Less is better","isMain":false,"metricName":" # relocations"},{"values":[0,0,0,0,0,0,0,0,0,0,0,0,0,0,0,0],"quality":"Less is better","isMain":false,"metricName":" # translocations"},{"values":[0,0,0,0,0,0,0,0,0,0,0,0,0,0,0,0],"quality":"Less is better","isMain":false,"metricName":" # inversions"},{"values":[0,0,0,0,0,0,0,1,0,0,0,0,0,0,0,1],"quality":"Less is better","isMain":false,"metricName":"# misassembled contigs"},{"values":[0,0,0,0,0,0,0,40281,0,0,0,0,0,0,0,56262],"quality":"Less is better","isMain":true,"metricName":"Misassembled contigs length"},{"values":[0,1,0,1,0,0,0,0,0,0,0,0,0,0,1,0],"quality":"Less is better","isMain":false,"metricName":"# local misassemblies"},{"values":[0,0,0,0,0,0,0,0,0,0,0,0,0,0,0,0],"quality":"Less is better","isMain":false,"metricName":"# unaligned mis. contigs"}]],["Unaligned",[{"values":[0,0,0,0,0,0,0,0,0,0,0,0,0,0,0,0],"quality":"Less is better","isMain":false,"metricName":"# fully unaligned contigs"},{"values":[0,0,0,0,0,0,0,0,0,0,0,0,0,0,0,0],"quality":"Less is better","isMain":false,"metricName":"Fully unaligned length"},{"values":[0,0,0,0,0,0,0,0,0,0,0,0,0,0,0,0],"quality":"Less is better","isMain":false,"metricName":"# partially unaligned contigs"},{"values":[0,0,0,0,0,0,0,0,0,0,0,0,0,0,0,0],"quality":"Less is better","isMain":false,"metricName":"Partially unaligned length"}]],["Mismatches",[{"values":[0,0,0,0,0,0,0,0,1,0,0,0,0,0,0,0],"quality":"Less is better","isMain":false,"metricName":"# mismatches"},{"values":[2,2,2,2,2,5,2,2,0,2,2,2,2,5,2,8],"quality":"Less is better","isMain":false,"metricName":"# indels"},{"values":[60,2,2,2,2,5,2,2,0,2,2,2,2,5,2,8],"quality":"Less is better","isMain":false,"metricName":"Indels length"},{"values":["0.00","0.00","0.00","0.00","0.00","0.00","0.00","0.00","1000.00","0.00","0.00","0.00","0.00","0.00","0.00","0.00"],"quality":"Less is better","isMain":true,"metricName":"# mismatches per 100 kbp"},{"values":["5.10","5.10","5.10","5.10","5.10","12.82","5.10","5.10","0.00","5.10","5.10","5.10","5.10","12.82","5.10","20.41"],"quality":"Less is better","isMain":true,"metricName":"# indels per 100 kbp"},{"values":[1,2,2,2,2,5,2,2,0,2,2,2,2,5,2,8],"quality":"Less is better","isMain":false,"metricName":" # indels (<= 5 bp)"},{"values":[1,0,0,0,0,0,0,0,0,0,0,0,0,0,0,0],"quality":"Less is better","isMain":false,"metricName":" # indels (> 5 bp)"},{"values":[50,0,0,0,0,0,0,0,310,0,0,0,0,0,0,249],"quality":"Less is better","isMain":false,"metricName":"# N's"},{"values":["127.02","0.00","0.00","0.00","0.00","0.00","0.00","0.00","56159.42","0.00","0.00","0.00","0.00","0.00","0.00","442.57"],"quality":"Less is better","isMain":true,"metricName":"# N's per 100 kbp"}]],["Statistics without reference",[{"values":[2,1,1,1,2,3,6,1,1,2,2,2,2,3,1,1],"quality":"Equal","isMain":true,"metricName":"# contigs"},{"values":[1,1,1,1,1,2,2,1,0,1,1,1,1,2,1,1],"quality":"Equal","isMain":false,"metricName":"# contigs (>= 1000 bp)"},{"values":[1,1,1,1,1,2,1,1,0,1,1,1,1,2,1,1],"quality":"Equal","isMain":false,"metricName":"# contigs (>= 5000 bp)"},{"values":[1,1,1,1,1,2,1,1,0,1,1,1,1,2,1,1],"quality":"Equal","isMain":false,"metricName":"# contigs (>= 10000 bp)"},{"values":[1,1,1,1,1,0,1,1,0,1,1,1,1,0,1,1],"quality":"Equal","isMain":false,"metricName":"# contigs (>= 25000 bp)"},{"values":[0,0,0,0,0,0,0,0,0,0,0,0,0,0,0,1],"quality":"Equal","isMain":false,"metricName":"# contigs (>= 50000 bp)"},{"values":[38593,40089,39191,39290,38545,22872,39183,40281,552,38545,38521,38521,38521,22872,39552,56262],"quality":"More is better","isMain":true,"metricName":"Largest contig"},{"values":[39363,40089,39191,39290,39325,39154,42848,40281,552,39345,39277,39277,39277,39154,39552,56262],"quality":"More is better","isMain":true,"metricName":"Total length"},{"values":[38593,40089,39191,39290,38545,38384,40234,40281,0,38545,38521,38521,38521,38384,39552,56262],"quality":"More is better","isMain":true,"metricName":"Total length (>= 1000 bp)"},{"values":[38593,40089,39191,39290,38545,38384,39183,40281,0,38545,38521,38521,38521,38384,39552,56262],"quality":"More is better","isMain":false,"metricName":"Total length (>= 5000 bp)"},{"values":[38593,40089,39191,39290,38545,38384,39183,40281,0,38545,38521,38521,38521,38384,39552,56262],"quality":"More is better","isMain":true,"metricName":"Total length (>= 10000 bp)"},{"values":[38593,40089,39191,39290,38545,0,39183,40281,0,38545,38521,38521,38521,0,39552,56262],"quality":"More is better","isMain":false,"metricName":"Total length (>= 25000 bp)"},{"values":[0,0,0,0,0,0,0,0,0,0,0,0,0,0,0,56262],"quality":"More is better","isMain":true,"metricName":"Total length (>= 50000 bp)"},{"values":[38593,40089,39191,39290,38545,22872,39183,40281,552,38545,38521,38521,38521,22872,39552,56262],"quality":"More is better","isMain":false,"metricName":"N50"},{"values":[38593,40089,39191,39290,38545,15512,39183,40281,552,38545,38521,38521,38521,15512,39552,56262],"quality":"More is better","isMain":false,"metricName":"N75"},{"values":[1,1,1,1,1,1,1,1,1,1,1,1,1,1,1,1],"quality":"Less is better","isMain":false,"metricName":"L50"},{"values":[1,1,1,1,1,2,1,1,1,1,1,1,1,2,1,1],"quality":"Less is better","isMain":false,"metricName":"L75"},{"values":["36.50","36.43","36.51","36.50","36.50","36.55","36.48","36.65","36.78","36.50","36.50","36.50","36.50","36.55","36.52","35.87"],"quality":"Equal","isMain":false,"metricName":"GC (%)"}]],["Predicted genes",[]],["Similarity statistics",[{"values":[0,0,0,0,0,0,0,0,0,0,0,0,0,0,0,0],"quality":"Equal","isMain":false,"metricName":"# similar correct contigs"},{"values":[0,0,0,0,0,0,0,0,0,0,0,0,0,0,0,0],"quality":"Equal","isMain":false,"metricName":"# similar misassembled blocks"}]],["Reference statistics",[{"values":[39189,39189,39189,39189,39189,39189,39189,39189,39189,39189,39189,39189,39189,39189,39189,39189],"quality":"Equal","isMain":false,"metricName":"Reference length"},{"values":[1,1,1,1,1,1,1,1,1,1,1,1,1,1,1,1],"quality":"Equal","isMain":false,"metricName":"Reference fragments"},{"values":["36.52","36.52","36.52","36.52","36.52","36.52","36.52","36.52","36.52","36.52","36.52","36.52","36.52","36.52","36.52","36.52"],"quality":"Equal","isMain":false,"metricName":"Reference GC (%)"}]]],"referenceName":"NC\_021790","date":"18 July 2018, Wednesday, 16:02:29","order":[0,1,2,3,4,5,6,7,8,9,10,11,12,13,14,15],"assembliesNames":["ABySS\_63","ABySS\_127","CLC","IDBA\_UD","MEGAHIT","metaVelvet","MIRA","Ray\_Meta","SOAPdenovo2","SPAdes","SPAdes\_meta","SPAdes\_sc","SPAdes\_sc\_careful","Velvet","vicuna","Geneious"]},{"assembliesWithNs":null,"minContig":500,"report":[["Genome statistics",[{"values":["100.000","100.000","100.000","100.000","100.000","99.484","100.000","100.000","7.693","100.000","100.000","99.945","99.945","99.484","72.612","100.000"],"quality":"More is better","isMain":true,"metricName":"Genome fraction (%)"},{"values":["1.001","1.000","1.000","1.001","1.002","1.000","1.012","1.022","0.989","1.001","1.001","1.000","1.000","1.000","1.142","1.009"],"quality":"Less is better","isMain":true,"metricName":"Duplication ratio"},{"values":[71499,71457,71449,71548,71590,38880,71686,69825,888,71526,71485,71410,71410,38880,2444,72069],"quality":"More is better","isMain":true,"metricName":"Largest alignment"},{"values":[71499,71457,71449,71548,71590,71097,72282,73023,5437,71526,71485,71410,71410,71097,59248,72069],"quality":"More is better","isMain":true,"metricName":"Total aligned length"},{"values":[71499,71457,71449,71548,71590,38881,71686,73023,66950,71526,71485,71410,71410,38881,572,72069],"quality":"More is better","isMain":false,"metricName":"NG50"},{"values":[71499,71457,71449,71548,71590,12817,71686,73023,66950,71526,71485,71410,71410,12817,508,72069],"quality":"More is better","isMain":false,"metricName":"NG75"},{"values":[71499,71457,71449,71548,71590,38880,71686,69825,null,71526,71485,71410,71410,38880,617,72069],"quality":"More is better","isMain":false,"metricName":"NA50"},{"values":[71499,71457,71449,71548,71590,12817,71686,69825,null,71526,71485,71410,71410,12817,523,72069],"quality":"More is better","isMain":false,"metricName":"NA75"},{"values":[71499,71457,71449,71548,71590,38880,71686,69825,null,71526,71485,71410,71410,38880,572,72069],"quality":"More is better","isMain":true,"metricName":"NGA50"},{"values":[71499,71457,71449,71548,71590,12817,71686,69825,null,71526,71485,71410,71410,12817,508,72069],"quality":"More is better","isMain":false,"metricName":"NGA75"},{"values":[1,1,1,1,1,1,1,1,1,1,1,1,1,1,41,1],"quality":"Less is better","isMain":false,"metricName":"LG50"},{"values":[1,1,1,1,1,3,1,1,1,1,1,1,1,3,75,1],"quality":"Less is better","isMain":false,"metricName":"LG75"},{"values":[1,1,1,1,1,1,1,1,null,1,1,1,1,1,31,1],"quality":"Less is better","isMain":false,"metricName":"LA50"},{"values":[1,1,1,1,1,3,1,1,null,1,1,1,1,3,57,1],"quality":"Less is better","isMain":false,"metricName":"LA75"},{"values":[1,1,1,1,1,1,1,1,null,1,1,1,1,1,41,1],"quality":"Less is better","isMain":true,"metricName":"LGA50"},{"values":[1,1,1,1,1,3,1,1,null,1,1,1,1,3,75,1],"quality":"Less is better","isMain":false,"metricName":"LGA75"}]],["Misassemblies",[{"values":[0,0,0,0,0,0,0,1,0,0,0,0,0,0,0,0],"quality":"Less is better","isMain":true,"metricName":"# misassemblies"},{"values":[0,0,0,0,0,0,0,1,0,0,0,0,0,0,0,0],"quality":"Less is better","isMain":false,"metricName":" # relocations"},{"values":[0,0,0,0,0,0,0,0,0,0,0,0,0,0,0,0],"quality":"Less is better","isMain":false,"metricName":" # translocations"},{"values":[0,0,0,0,0,0,0,0,0,0,0,0,0,0,0,0],"quality":"Less is better","isMain":false,"metricName":" # inversions"},{"values":[0,0,0,0,0,0,0,1,0,0,0,0,0,0,0,0],"quality":"Less is better","isMain":false,"metricName":"# misassembled contigs"},{"values":[0,0,0,0,0,0,0,73023,0,0,0,0,0,0,0,0],"quality":"Less is better","isMain":true,"metricName":"Misassembled contigs length"},{"values":[0,0,0,1,1,0,1,0,0,0,0,0,0,0,0,1],"quality":"Less is better","isMain":false,"metricName":"# local misassemblies"},{"values":[0,0,0,0,0,0,0,0,1,0,0,0,0,0,0,0],"quality":"Less is better","isMain":false,"metricName":"# unaligned mis. contigs"}]],["Unaligned",[{"values":[0,0,0,0,0,0,0,0,0,0,0,0,0,0,0,0],"quality":"Less is better","isMain":false,"metricName":"# fully unaligned contigs"},{"values":[0,0,0,0,0,0,0,0,0,0,0,0,0,0,0,0],"quality":"Less is better","isMain":false,"metricName":"Fully unaligned length"},{"values":[0,0,0,0,0,0,0,0,1,0,0,0,0,0,0,0],"quality":"Less is better","isMain":false,"metricName":"# partially unaligned contigs"},{"values":[0,0,0,0,0,0,0,0,61513,0,0,0,0,0,0,0],"quality":"Less is better","isMain":false,"metricName":"Partially unaligned length"}]],["Mismatches",[{"values":[0,0,0,0,0,0,0,0,0,0,0,0,0,0,1,0],"quality":"Less is better","isMain":false,"metricName":"# mismatches"},{"values":[7,6,6,6,6,4,5,6,5,7,7,6,6,4,3,6],"quality":"Less is better","isMain":false,"metricName":"# indels"},{"values":[56,14,6,6,6,4,5,6,81,83,42,6,6,4,3,6],"quality":"Less is better","isMain":false,"metricName":"Indels length"},{"values":["0.00","0.00","0.00","0.00","0.00","0.00","0.00","0.00","0.00","0.00","0.00","0.00","0.00","0.00","1.93","0.00"],"quality":"Less is better","isMain":true,"metricName":"# mismatches per 100 kbp"},{"values":["9.80","8.40","8.40","8.40","8.40","5.63","7.00","8.40","90.98","9.80","9.80","8.40","8.40","5.63","5.78","8.40"],"quality":"Less is better","isMain":true,"metricName":"# indels per 100 kbp"},{"values":[6,5,6,6,6,4,5,6,1,6,6,6,6,4,3,6],"quality":"Less is better","isMain":false,"metricName":" # indels (<= 5 bp)"},{"values":[1,1,0,0,0,0,0,0,4,1,1,0,0,0,0,0],"quality":"Less is better","isMain":false,"metricName":" # indels (> 5 bp)"},{"values":[0,0,0,0,0,0,0,0,10926,0,0,0,0,0,0,0],"quality":"Less is better","isMain":false,"metricName":"# N's"},{"values":["0.00","0.00","0.00","0.00","0.00","0.00","0.00","0.00","16319.64","0.00","0.00","0.00","0.00","0.00","0.00","0.00"],"quality":"Less is better","isMain":true,"metricName":"# N's per 100 kbp"}]],["Statistics without reference",[{"values":[1,1,1,1,1,4,2,1,1,1,1,1,1,4,86,1],"quality":"Equal","isMain":true,"metricName":"# contigs"},{"values":[1,1,1,1,1,4,1,1,1,1,1,1,1,4,7,1],"quality":"Equal","isMain":false,"metricName":"# contigs (>= 1000 bp)"},{"values":[1,1,1,1,1,4,1,1,1,1,1,1,1,4,0,1],"quality":"Equal","isMain":false,"metricName":"# contigs (>= 5000 bp)"},{"values":[1,1,1,1,1,3,1,1,1,1,1,1,1,3,0,1],"quality":"Equal","isMain":false,"metricName":"# contigs (>= 10000 bp)"},{"values":[1,1,1,1,1,1,1,1,1,1,1,1,1,1,0,1],"quality":"Equal","isMain":false,"metricName":"# contigs (>= 25000 bp)"},{"values":[1,1,1,1,1,0,1,1,1,1,1,1,1,0,0,1],"quality":"Equal","isMain":false,"metricName":"# contigs (>= 50000 bp)"},{"values":[71499,71457,71449,71548,71590,38881,71686,73023,66950,71526,71485,71410,71410,38881,2444,72069],"quality":"More is better","isMain":true,"metricName":"Largest contig"},{"values":[71499,71457,71449,71548,71590,71100,72282,73023,66950,71526,71485,71410,71410,71100,59259,72069],"quality":"More is better","isMain":true,"metricName":"Total length"},{"values":[71499,71457,71449,71548,71590,71100,71686,73023,66950,71526,71485,71410,71410,71100,12431,72069],"quality":"More is better","isMain":true,"metricName":"Total length (>= 1000 bp)"},{"values":[71499,71457,71449,71548,71590,71100,71686,73023,66950,71526,71485,71410,71410,71100,0,72069],"quality":"More is better","isMain":false,"metricName":"Total length (>= 5000 bp)"},{"values":[71499,71457,71449,71548,71590,65385,71686,73023,66950,71526,71485,71410,71410,65385,0,72069],"quality":"More is better","isMain":true,"metricName":"Total length (>= 10000 bp)"},{"values":[71499,71457,71449,71548,71590,38881,71686,73023,66950,71526,71485,71410,71410,38881,0,72069],"quality":"More is better","isMain":false,"metricName":"Total length (>= 25000 bp)"},{"values":[71499,71457,71449,71548,71590,0,71686,73023,66950,71526,71485,71410,71410,0,0,72069],"quality":"More is better","isMain":true,"metricName":"Total length (>= 50000 bp)"},{"values":[71499,71457,71449,71548,71590,38881,71686,73023,66950,71526,71485,71410,71410,38881,617,72069],"quality":"More is better","isMain":false,"metricName":"N50"},{"values":[71499,71457,71449,71548,71590,12817,71686,73023,66950,71526,71485,71410,71410,12817,523,72069],"quality":"More is better","isMain":false,"metricName":"N75"},{"values":[1,1,1,1,1,1,1,1,1,1,1,1,1,1,31,1],"quality":"Less is better","isMain":false,"metricName":"L50"},{"values":[1,1,1,1,1,3,1,1,1,1,1,1,1,3,57,1],"quality":"Less is better","isMain":false,"metricName":"L75"},{"values":["32.88","32.88","32.88","32.88","32.89","32.88","32.96","32.80","32.81","32.89","32.88","32.88","32.88","32.88","32.54","32.91"],"quality":"Equal","isMain":false,"metricName":"GC (%)"}]],["Predicted genes",[]],["Similarity statistics",[{"values":[0,0,0,0,0,0,0,0,0,0,0,0,0,0,0,0],"quality":"Equal","isMain":false,"metricName":"# similar correct contigs"},{"values":[0,0,0,0,0,0,0,0,0,0,0,0,0,0,0,0],"quality":"Equal","isMain":false,"metricName":"# similar misassembled blocks"}]],["Reference statistics",[{"values":[71443,71443,71443,71443,71443,71443,71443,71443,71443,71443,71443,71443,71443,71443,71443,71443],"quality":"Equal","isMain":false,"metricName":"Reference length"},{"values":[1,1,1,1,1,1,1,1,1,1,1,1,1,1,1,1],"quality":"Equal","isMain":false,"metricName":"Reference fragments"},{"values":["32.88","32.88","32.88","32.88","32.88","32.88","32.88","32.88","32.88","32.88","32.88","32.88","32.88","32.88","32.88","32.88"],"quality":"Equal","isMain":false,"metricName":"Reference GC (%)"}]]],"referenceName":"NC\_021794","date":"18 July 2018, Wednesday, 16:02:39","order":[0,1,2,3,4,5,6,7,8,9,10,11,12,13,14,15],"assembliesNames":["ABySS\_63","ABySS\_127","CLC","IDBA\_UD","MEGAHIT","metaVelvet","MIRA","Ray\_Meta","SOAPdenovo2","SPAdes","SPAdes\_meta","SPAdes\_sc","SPAdes\_sc\_careful","Velvet","vicuna","Geneious"]},{"assembliesWithNs":null,"minContig":500,"report":[["Genome statistics",[{"values":["100.000","99.971","99.970","100.000","100.000","99.946","100.000","99.409","10.732","100.000","100.000","100.000","100.000","99.946","70.768","100.000"],"quality":"More is better","isMain":true,"metricName":"Genome fraction (%)"},{"values":["1.000","1.000","1.000","1.001","1.002","1.002","1.104","1.000","0.997","1.002","1.001","1.001","1.001","1.002","1.057","1.005"],"quality":"Less is better","isMain":true,"metricName":"Duplication ratio"},{"values":[72541,72514,72513,72634,72676,36945,25697,72105,784,70642,70598,70598,70598,36945,23419,72916],"quality":"More is better","isMain":true,"metricName":"Largest alignment"},{"values":[72541,72514,72513,72634,72676,72601,80083,72105,7763,72689,72601,72601,72591,72601,54235,72916],"quality":"More is better","isMain":true,"metricName":"Total aligned length"},{"values":[72541,72514,72535,72634,72676,36945,17449,72105,67075,70642,70598,70598,70598,36945,656,72916],"quality":"More is better","isMain":false,"metricName":"NG50"},{"values":[72541,72514,72535,72634,72676,25815,12836,72105,67075,70642,70598,70598,70598,25815,null,72916],"quality":"More is better","isMain":false,"metricName":"NG75"},{"values":[72541,72514,72513,72634,72676,36945,17449,72105,null,70642,70598,70598,70598,36945,1875,72916],"quality":"More is better","isMain":false,"metricName":"NA50"},{"values":[72541,72514,72513,72634,72676,25815,4642,72105,null,70642,70598,70598,70598,25815,549,72916],"quality":"More is better","isMain":false,"metricName":"NA75"},{"values":[72541,72514,72513,72634,72676,36945,17449,72105,null,70642,70598,70598,70598,36945,656,72916],"quality":"More is better","isMain":true,"metricName":"NGA50"},{"values":[72541,72514,72513,72634,72676,25815,12836,72105,null,70642,70598,70598,70598,25815,null,72916],"quality":"More is better","isMain":false,"metricName":"NGA75"},{"values":[1,1,1,1,1,1,2,1,1,1,1,1,1,1,13,1],"quality":"Less is better","isMain":false,"metricName":"LG50"},{"values":[1,1,1,1,1,2,3,1,1,1,1,1,1,2,null,1],"quality":"Less is better","isMain":false,"metricName":"LG75"},{"values":[1,1,1,1,1,1,2,1,null,1,1,1,1,1,3,1],"quality":"Less is better","isMain":false,"metricName":"LA50"},{"values":[1,1,1,1,1,2,4,1,null,1,1,1,1,2,20,1],"quality":"Less is better","isMain":false,"metricName":"LA75"},{"values":[1,1,1,1,1,1,2,1,null,1,1,1,1,1,13,1],"quality":"Less is better","isMain":true,"metricName":"LGA50"},{"values":[1,1,1,1,1,2,3,1,null,1,1,1,1,2,null,1],"quality":"Less is better","isMain":false,"metricName":"LGA75"}]],["Misassemblies",[{"values":[0,0,0,0,0,0,0,0,0,0,0,0,0,0,0,0],"quality":"Less is better","isMain":true,"metricName":"# misassemblies"},{"values":[0,0,0,0,0,0,0,0,0,0,0,0,0,0,0,0],"quality":"Less is better","isMain":false,"metricName":" # relocations"},{"values":[0,0,0,0,0,0,0,0,0,0,0,0,0,0,0,0],"quality":"Less is better","isMain":false,"metricName":" # translocations"},{"values":[0,0,0,0,0,0,0,0,0,0,0,0,0,0,0,0],"quality":"Less is better","isMain":false,"metricName":" # inversions"},{"values":[0,0,0,0,0,0,0,0,0,0,0,0,0,0,0,0],"quality":"Less is better","isMain":false,"metricName":"# misassembled contigs"},{"values":[0,0,0,0,0,0,0,0,0,0,0,0,0,0,0,0],"quality":"Less is better","isMain":true,"metricName":"Misassembled contigs length"},{"values":[0,0,0,1,1,0,0,0,0,0,0,0,0,0,0,1],"quality":"Less is better","isMain":false,"metricName":"# local misassemblies"},{"values":[0,0,0,0,0,0,0,0,1,0,0,0,0,0,0,0],"quality":"Less is better","isMain":false,"metricName":"# unaligned mis. contigs"}]],["Unaligned",[{"values":[0,0,0,0,0,0,0,0,0,0,0,0,0,0,0,0],"quality":"Less is better","isMain":false,"metricName":"# fully unaligned contigs"},{"values":[0,0,0,0,0,0,0,0,0,0,0,0,0,0,0,0],"quality":"Less is better","isMain":false,"metricName":"Fully unaligned length"},{"values":[0,0,0,0,0,0,0,0,1,0,0,0,0,0,0,0],"quality":"Less is better","isMain":false,"metricName":"# partially unaligned contigs"},{"values":[0,0,0,0,0,0,0,0,59312,0,0,0,0,0,0,0],"quality":"Less is better","isMain":false,"metricName":"Partially unaligned length"}]],["Mismatches",[{"values":[1,0,0,0,0,1,1,1,1,0,0,0,0,1,0,0],"quality":"Less is better","isMain":false,"metricName":"# mismatches"},{"values":[2,1,1,1,1,2,2,2,3,1,1,1,1,2,1,1],"quality":"Less is better","isMain":false,"metricName":"# indels"},{"values":[7,1,1,1,1,2,2,2,25,1,1,1,1,2,1,1],"quality":"Less is better","isMain":false,"metricName":"Indels length"},{"values":["1.38","0.00","0.00","0.00","0.00","1.38","1.38","1.39","12.85","0.00","0.00","0.00","0.00","1.38","0.00","0.00"],"quality":"Less is better","isMain":true,"metricName":"# mismatches per 100 kbp"},{"values":["2.76","1.38","1.38","1.38","1.38","2.76","2.76","2.77","38.54","1.38","1.38","1.38","1.38","2.76","1.95","1.38"],"quality":"Less is better","isMain":true,"metricName":"# indels per 100 kbp"},{"values":[1,1,1,1,1,2,2,2,2,1,1,1,1,2,1,1],"quality":"Less is better","isMain":false,"metricName":" # indels (<= 5 bp)"},{"values":[1,0,0,0,0,0,0,0,1,0,0,0,0,0,0,0],"quality":"Less is better","isMain":false,"metricName":" # indels (> 5 bp)"},{"values":[0,0,0,0,0,0,1,0,16810,0,0,0,0,0,0,2],"quality":"Less is better","isMain":false,"metricName":"# N's"},{"values":["0.00","0.00","0.00","0.00","0.00","0.00","1.25","0.00","25061.50","0.00","0.00","0.00","0.00","0.00","0.00","2.74"],"quality":"Less is better","isMain":true,"metricName":"# N's per 100 kbp"}]],["Statistics without reference",[{"values":[1,1,1,1,1,6,19,1,1,2,2,2,2,6,46,1],"quality":"Equal","isMain":true,"metricName":"# contigs"},{"values":[1,1,1,1,1,5,10,1,1,2,2,2,2,5,6,1],"quality":"Equal","isMain":false,"metricName":"# contigs (>= 1000 bp)"},{"values":[1,1,1,1,1,3,3,1,1,1,1,1,1,3,1,1],"quality":"Equal","isMain":false,"metricName":"# contigs (>= 5000 bp)"},{"values":[1,1,1,1,1,2,3,1,1,1,1,1,1,2,1,1],"quality":"Equal","isMain":false,"metricName":"# contigs (>= 10000 bp)"},{"values":[1,1,1,1,1,2,1,1,1,1,1,1,1,2,0,1],"quality":"Equal","isMain":false,"metricName":"# contigs (>= 25000 bp)"},{"values":[1,1,1,1,1,0,0,1,1,1,1,1,1,0,0,1],"quality":"Equal","isMain":false,"metricName":"# contigs (>= 50000 bp)"},{"values":[72541,72514,72535,72634,72676,36945,25697,72105,67075,70642,70598,70598,70598,36945,23419,72916],"quality":"More is better","isMain":true,"metricName":"Largest contig"},{"values":[72541,72514,72535,72634,72676,72655,80083,72105,67075,72689,72601,72601,72601,72655,54242,72916],"quality":"More is better","isMain":true,"metricName":"Total length"},{"values":[72541,72514,72535,72634,72676,71902,75006,72105,67075,72689,72601,72601,72601,71902,31615,72916],"quality":"More is better","isMain":true,"metricName":"Total length (>= 1000 bp)"},{"values":[72541,72514,72535,72634,72676,67801,55982,72105,67075,70642,70598,70598,70598,67801,23419,72916],"quality":"More is better","isMain":false,"metricName":"Total length (>= 5000 bp)"},{"values":[72541,72514,72535,72634,72676,62760,55982,72105,67075,70642,70598,70598,70598,62760,23419,72916],"quality":"More is better","isMain":true,"metricName":"Total length (>= 10000 bp)"},{"values":[72541,72514,72535,72634,72676,62760,25697,72105,67075,70642,70598,70598,70598,62760,0,72916],"quality":"More is better","isMain":false,"metricName":"Total length (>= 25000 bp)"},{"values":[72541,72514,72535,72634,72676,0,0,72105,67075,70642,70598,70598,70598,0,0,72916],"quality":"More is better","isMain":true,"metricName":"Total length (>= 50000 bp)"},{"values":[72541,72514,72535,72634,72676,36945,17449,72105,67075,70642,70598,70598,70598,36945,1875,72916],"quality":"More is better","isMain":false,"metricName":"N50"},{"values":[72541,72514,72535,72634,72676,25815,4642,72105,67075,70642,70598,70598,70598,25815,549,72916],"quality":"More is better","isMain":false,"metricName":"N75"},{"values":[1,1,1,1,1,1,2,1,1,1,1,1,1,1,3,1],"quality":"Less is better","isMain":false,"metricName":"L50"},{"values":[1,1,1,1,1,2,4,1,1,1,1,1,1,2,20,1],"quality":"Less is better","isMain":false,"metricName":"L75"},{"values":["38.05","38.05","38.05","38.05","38.06","38.04","37.89","38.06","37.85","38.06","38.06","38.06","38.05","38.04","37.82","38.06"],"quality":"Equal","isMain":false,"metricName":"GC (%)"}]],["Predicted genes",[]],["Similarity statistics",[{"values":[0,0,0,0,0,0,0,0,0,0,0,0,0,0,0,0],"quality":"Equal","isMain":false,"metricName":"# similar correct contigs"},{"values":[0,0,0,0,0,0,0,0,0,0,0,0,0,0,0,0],"quality":"Equal","isMain":false,"metricName":"# similar misassembled blocks"}]],["Reference statistics",[{"values":[72534,72534,72534,72534,72534,72534,72534,72534,72534,72534,72534,72534,72534,72534,72534,72534],"quality":"Equal","isMain":false,"metricName":"Reference length"},{"values":[1,1,1,1,1,1,1,1,1,1,1,1,1,1,1,1],"quality":"Equal","isMain":false,"metricName":"Reference fragments"},{"values":["38.06","38.06","38.06","38.06","38.06","38.06","38.06","38.06","38.06","38.06","38.06","38.06","38.06","38.06","38.06","38.06"],"quality":"Equal","isMain":false,"metricName":"Reference GC (%)"}]]],"referenceName":"NC\_021796","date":"18 July 2018, Wednesday, 16:02:49","order":[0,1,2,3,4,5,6,7,8,9,10,11,12,13,14,15],"assembliesNames":["ABySS\_63","ABySS\_127","CLC","IDBA\_UD","MEGAHIT","metaVelvet","MIRA","Ray\_Meta","SOAPdenovo2","SPAdes","SPAdes\_meta","SPAdes\_sc","SPAdes\_sc\_careful","Velvet","vicuna","Geneious"]},{"assembliesWithNs":null,"minContig":500,"report":[["Genome statistics",[]],["Misassemblies",[]],["Unaligned",[]],["Mismatches",[{"values":[329,229,0,708,2315,0,1033],"quality":"Less is better","isMain":false,"metricName":"# N's"},{"values":["7.27","5.05","0.00","48726.77","40.62","0.00","23.00"],"quality":"Less is better","isMain":true,"metricName":"# N's per 100 kbp"}]],["Statistics without reference",[{"values":[52,50,1143,2,1513,4969,53],"quality":"Equal","isMain":true,"metricName":"# contigs"},{"values":[47,40,209,0,320,88,48],"quality":"Equal","isMain":false,"metricName":"# contigs (>= 1000 bp)"},{"values":[35,31,34,0,27,0,35],"quality":"Equal","isMain":false,"metricName":"# contigs (>= 5000 bp)"},{"values":[30,28,31,0,23,0,30],"quality":"Equal","isMain":false,"metricName":"# contigs (>= 10000 bp)"},{"values":[26,24,27,0,19,0,24],"quality":"Equal","isMain":false,"metricName":"# contigs (>= 25000 bp)"},{"values":[20,20,22,0,17,0,23],"quality":"Equal","isMain":false,"metricName":"# contigs (>= 50000 bp)"},{"values":[606039,850094,517843,759,961438,1713,570751],"quality":"More is better","isMain":true,"metricName":"Largest contig"},{"values":[4523550,4539144,5327286,1453,5699396,3125238,4491700],"quality":"More is better","isMain":true,"metricName":"Total length"},{"values":[4520425,4532612,4706165,0,4889696,98683,4487966],"quality":"More is better","isMain":true,"metricName":"Total length (>= 1000 bp)"},{"values":[4487113,4508619,4459027,0,4471337,0,4457694],"quality":"More is better","isMain":false,"metricName":"Total length (>= 5000 bp)"},{"values":[4443856,4483538,4436936,0,4443539,0,4421811],"quality":"More is better","isMain":true,"metricName":"Total length (>= 10000 bp)"},{"values":[4378197,4416253,4374159,0,4381131,0,4320336],"quality":"More is better","isMain":false,"metricName":"Total length (>= 25000 bp)"},{"values":[4125658,4227198,4155491,0,4283644,0,4274188],"quality":"More is better","isMain":true,"metricName":"Total length (>= 50000 bp)"},{"values":[242354,246998,183441,759,254168,622,270337],"quality":"More is better","isMain":false,"metricName":"N50"},{"values":[128038,147853,56152,694,53841,549,120025],"quality":"More is better","isMain":false,"metricName":"N75"},{"values":[7,6,9,1,7,2130,6],"quality":"Less is better","isMain":false,"metricName":"L50"},{"values":[13,11,20,2,17,3466,13],"quality":"Less is better","isMain":false,"metricName":"L75"},{"values":["34.63","34.65","35.37","34.90","35.62","34.29","34.60"],"quality":"Equal","isMain":false,"metricName":"GC (%)"}]],["Predicted genes",[]],["Similarity statistics",[]],["Reference statistics",[]]],"referenceName":"","date":"18 July 2018, Wednesday, 16:02:54","order":[0,1,2,3,4,5,6],"assembliesNames":["ABySS\_63","ABySS\_127","CLC","SOAPdenovo2","SPAdes\_sc","vicuna","Geneious"]}]

{}

{}

{}

{}

{}

{}

{}

{}

{}

{}

{}

{}

[{"refnames":["KF302033","NC\_001330","NC\_001422","NC\_021790","KC821629","KF302037","KF302036","KF302034","NC\_021796","NC\_021794","KC821625","KF302035"],"coord\_y":[[null,1,1,2,1,null,null,null,1,1,1,1],[null,null,1,1,null,null,null,null,1,1,1,1],[1,1,1,1,1,1,1,1,1,1,2,144],[1,1,1,1,1,1,1,1,1,1,1,1],[1,1,1,2,1,1,1,1,1,1,1,1],[1,null,null,3,4,1,5,1,6,4,null,1],[1,1,1,6,22,1,1,2,19,2,4,29],[1,1,1,1,1,1,1,1,1,1,1,1],[1,1,1,1,8,1,1,1,1,1,1,3],[1,1,1,2,1,1,1,1,2,1,1,1],[1,1,1,2,1,1,1,1,2,1,1,1],[1,1,1,2,1,1,1,1,2,1,1,1],[1,1,2,2,1,1,1,1,2,1,1,1],[1,null,null,3,4,1,5,1,6,4,null,1],[1,10,3,1,2,49,45,74,46,86,98,1],[1,1,1,1,1,1,1,1,1,1,1,1]],"coord\_x":[[0.475,1.475,2.475,3.475,4.475,5.475,6.475,7.475,8.475,9.475,10.475,11.475],[0.5449999999999999,1.545,2.545,3.545,4.545,5.545,6.545,7.545,8.545,9.545,10.545,11.545],[0.615,1.615,2.615,3.615,4.615,5.615,6.615,7.615,8.615,9.615,10.615,11.615],[0.6849999999999999,1.685,2.685,3.685,4.685,5.685,6.685,7.685,8.685,9.685,10.685,11.685],[0.755,1.755,2.755,3.755,4.755,5.755,6.755,7.755,8.755,9.755,10.755,11.755],[0.825,1.825,2.825,3.825,4.825,5.825,6.825,7.825,8.825,9.825,10.825,11.825],[0.895,1.895,2.895,3.895,4.895,5.895,6.895,7.895,8.895,9.895,10.895,11.895],[0.965,1.965,2.965,3.965,4.965,5.965,6.965,7.965,8.965,9.965,10.965,11.965],[1.035,2.035,3.035,4.035,5.035,6.035,7.035,8.035,9.035,10.035,11.035,12.035],[1.105,2.105,3.105,4.105,5.105,6.105,7.105,8.105,9.105,10.105,11.105,12.105],[1.175,2.175,3.175,4.175,5.175,6.175,7.175,8.175,9.175,10.175,11.175,12.175],[1.245,2.245,3.245,4.245,5.245,6.245,7.245,8.245,9.245,10.245,11.245,12.245],[1.315,2.315,3.315,4.315,5.315,6.315,7.315,8.315,9.315,10.315,11.315,12.315],[1.385,2.385,3.385,4.385,5.385,6.385,7.385,8.385,9.385,10.385,11.385,12.385],[1.455,2.455,3.455,4.455,5.455,6.455,7.455,8.455,9.455,10.455,11.455,12.455],[1.525,2.525,3.525,4.525,5.525,6.525,7.525,8.525,9.525,10.525,11.525,12.525]],"filenames":["ABySS\_63","ABySS\_127","CLC","IDBA\_UD","MEGAHIT","metaVelvet","MIRA","Ray\_Meta","SOAPdenovo2","SPAdes","SPAdes\_meta","SPAdes\_sc","SPAdes\_sc\_careful","Velvet","vicuna","Geneious"]},
{"refnames":["KF302034","NC\_021794","KC821625","NC\_021796","KC821629","NC\_021790","KF302037","KF302035","KF302033","KF302036","NC\_001330","NC\_001422"],"coord\_y":[[null,71499,76649,72541,44392,38543,null,35176,null,null,6148,5445],[null,71457,76666,72514,null,40089,null,35176,null,null,null,5158],[129415,71449,52891,72513,54016,39191,44551,35176,36771,37706,6089,5386],[128851,71548,76765,72634,54115,39290,44650,35179,36870,37827,6188,5365],[129556,71590,76807,72676,54157,38545,44692,35176,36912,37869,6230,5355],[129478,38880,null,36945,35739,22872,44558,35177,36833,20798,null,null],[128970,71686,76666,25697,35913,39164,44678,7023,36990,37981,6089,5233],[129048,69825,76550,72105,54799,23054,26064,35176,37514,38637,6508,5355],[977,888,36281,784,214,100,1075,350,251,933,6034,5373],[129492,71526,76743,70642,54093,38545,44628,35176,36848,37805,6166,5365],[129470,71485,76925,70598,54041,38521,44606,35176,36826,37783,6134,5365],[128748,71410,76993,70598,54042,38521,44606,35176,36826,37783,6134,5431],[129470,71410,76559,70598,54042,38521,44606,35175,36826,37783,6134,4783],[129478,38880,null,36945,35739,22872,44558,35177,36833,20798,null,null],[48827,2444,1579,23419,43063,39552,839,35176,36998,851,560,764],[129441,72069,76666,72916,54016,39166,44829,35176,37131,38107,6385,5365]],"coord\_x":[[0.475,1.475,2.475,3.475,4.475,5.475,6.475,7.475,8.475,9.475,10.475,11.475],[0.5449999999999999,1.545,2.545,3.545,4.545,5.545,6.545,7.545,8.545,9.545,10.545,11.545],[0.615,1.615,2.615,3.615,4.615,5.615,6.615,7.615,8.615,9.615,10.615,11.615],[0.6849999999999999,1.685,2.685,3.685,4.685,5.685,6.685,7.685,8.685,9.685,10.685,11.685],[0.755,1.755,2.755,3.755,4.755,5.755,6.755,7.755,8.755,9.755,10.755,11.755],[0.825,1.825,2.825,3.825,4.825,5.825,6.825,7.825,8.825,9.825,10.825,11.825],[0.895,1.895,2.895,3.895,4.895,5.895,6.895,7.895,8.895,9.895,10.895,11.895],[0.965,1.965,2.965,3.965,4.965,5.965,6.965,7.965,8.965,9.965,10.965,11.965],[1.035,2.035,3.035,4.035,5.035,6.035,7.035,8.035,9.035,10.035,11.035,12.035],[1.105,2.105,3.105,4.105,5.105,6.105,7.105,8.105,9.105,10.105,11.105,12.105],[1.175,2.175,3.175,4.175,5.175,6.175,7.175,8.175,9.175,10.175,11.175,12.175],[1.245,2.245,3.245,4.245,5.245,6.245,7.245,8.245,9.245,10.245,11.245,12.245],[1.315,2.315,3.315,4.315,5.315,6.315,7.315,8.315,9.315,10.315,11.315,12.315],[1.385,2.385,3.385,4.385,5.385,6.385,7.385,8.385,9.385,10.385,11.385,12.385],[1.455,2.455,3.455,4.455,5.455,6.455,7.455,8.455,9.455,10.455,11.455,12.455],[1.525,2.525,3.525,4.525,5.525,6.525,7.525,8.525,9.525,10.525,11.525,12.525]],"filenames":["ABySS\_63","ABySS\_127","CLC","IDBA\_UD","MEGAHIT","metaVelvet","MIRA","Ray\_Meta","SOAPdenovo2","SPAdes","SPAdes\_meta","SPAdes\_sc","SPAdes\_sc\_careful","Velvet","vicuna","Geneious"]},
{"refnames":["KF302034","NC\_021796","KC821625","NC\_021794","KC821629","NC\_021790","KF302037","KF302035","KF302033","KF302036","NC\_001330","NC\_001422"],"coord\_y":[[null,72541,76649,71499,55287,39313,null,35176,null,null,6148,5445],[null,72514,76666,71457,null,40089,null,35176,null,null,null,5158],[129415,72513,80629,71449,54016,39191,44551,48992,36771,37706,6089,5386],[128851,72634,76765,71548,54115,39290,44650,35179,36870,37827,6188,5365],[129556,72676,76807,71590,54157,39325,44692,35176,36912,37869,6230,5355],[129478,72601,null,71097,53817,39128,44558,35177,36833,34297,null,null],[129662,80083,84045,72282,63047,42829,44678,47238,36990,37981,6089,5233],[131360,72105,76550,73023,54799,40281,45598,35176,37514,38637,6508,5355],[16037,7763,70114,5437,2748,100,2910,1311,4486,2368,6034,5373],[129492,72689,76743,71526,54093,39345,44628,35176,36848,37805,6166,5365],[129470,72601,76925,71485,54041,39277,44606,35176,36826,37783,6134,5365],[128748,72601,76993,71410,54042,39277,44606,35176,36826,37783,6134,5431],[129470,72591,76559,71410,54042,39277,44606,35175,36826,37783,6134,5410],[129478,72601,null,71097,53817,39128,44558,35177,36833,34297,null,null],[127741,54235,68410,59248,53973,39552,26656,35176,36998,25452,5190,1771],[153292,72916,76666,72069,54016,56245,44829,35176,37131,38107,6385,5365]],"coord\_x":[[0.475,1.475,2.475,3.475,4.475,5.475,6.475,7.475,8.475,9.475,10.475,11.475],[0.5449999999999999,1.545,2.545,3.545,4.545,5.545,6.545,7.545,8.545,9.545,10.545,11.545],[0.615,1.615,2.615,3.615,4.615,5.615,6.615,7.615,8.615,9.615,10.615,11.615],[0.6849999999999999,1.685,2.685,3.685,4.685,5.685,6.685,7.685,8.685,9.685,10.685,11.685],[0.755,1.755,2.755,3.755,4.755,5.755,6.755,7.755,8.755,9.755,10.755,11.755],[0.825,1.825,2.825,3.825,4.825,5.825,6.825,7.825,8.825,9.825,10.825,11.825],[0.895,1.895,2.895,3.895,4.895,5.895,6.895,7.895,8.895,9.895,10.895,11.895],[0.965,1.965,2.965,3.965,4.965,5.965,6.965,7.965,8.965,9.965,10.965,11.965],[1.035,2.035,3.035,4.035,5.035,6.035,7.035,8.035,9.035,10.035,11.035,12.035],[1.105,2.105,3.105,4.105,5.105,6.105,7.105,8.105,9.105,10.105,11.105,12.105],[1.175,2.175,3.175,4.175,5.175,6.175,7.175,8.175,9.175,10.175,11.175,12.175],[1.245,2.245,3.245,4.245,5.245,6.245,7.245,8.245,9.245,10.245,11.245,12.245],[1.315,2.315,3.315,4.315,5.315,6.315,7.315,8.315,9.315,10.315,11.315,12.315],[1.385,2.385,3.385,4.385,5.385,6.385,7.385,8.385,9.385,10.385,11.385,12.385],[1.455,2.455,3.455,4.455,5.455,6.455,7.455,8.455,9.455,10.455,11.455,12.455],[1.525,2.525,3.525,4.525,5.525,6.525,7.525,8.525,9.525,10.525,11.525,12.525]],"filenames":["ABySS\_63","ABySS\_127","CLC","IDBA\_UD","MEGAHIT","metaVelvet","MIRA","Ray\_Meta","SOAPdenovo2","SPAdes","SPAdes\_meta","SPAdes\_sc","SPAdes\_sc\_careful","Velvet","vicuna","Geneious"]},
{"refnames":["KC821625","KC821629","KF302033","KF302034","KF302035","KF302036","KF302037","NC\_001330","NC\_001422","NC\_021790","NC\_021794","NC\_021796"],"coord\_y":[[0,1,null,null,0,null,null,0,0,0,0,0],[0,null,null,null,0,null,null,null,0,0,0,0],[0,0,0,0,0,0,0,0,0,0,0,0],[0,0,0,0,0,0,0,0,0,0,0,0],[0,0,0,0,0,0,0,0,0,0,0,0],[null,0,0,0,0,0,0,null,null,0,0,0],[0,0,0,0,0,0,0,0,0,0,0,0],[0,0,0,1,0,0,1,0,0,1,1,0],[1,0,0,0,0,0,0,0,0,0,0,0],[0,0,0,0,0,0,0,0,0,0,0,0],[0,0,0,0,0,0,0,0,0,0,0,0],[0,0,0,0,0,0,0,0,0,0,0,0],[0,0,0,0,0,0,0,0,0,0,0,0],[null,0,0,0,0,0,0,null,null,0,0,0],[0,0,0,0,0,0,0,0,0,0,0,0],[0,0,0,2,0,0,0,0,0,1,0,0]],"coord\_x":[[0.475,1.475,2.475,3.475,4.475,5.475,6.475,7.475,8.475,9.475,10.475,11.475],[0.5449999999999999,1.545,2.545,3.545,4.545,5.545,6.545,7.545,8.545,9.545,10.545,11.545],[0.615,1.615,2.615,3.615,4.615,5.615,6.615,7.615,8.615,9.615,10.615,11.615],[0.6849999999999999,1.685,2.685,3.685,4.685,5.685,6.685,7.685,8.685,9.685,10.685,11.685],[0.755,1.755,2.755,3.755,4.755,5.755,6.755,7.755,8.755,9.755,10.755,11.755],[0.825,1.825,2.825,3.825,4.825,5.825,6.825,7.825,8.825,9.825,10.825,11.825],[0.895,1.895,2.895,3.895,4.895,5.895,6.895,7.895,8.895,9.895,10.895,11.895],[0.965,1.965,2.965,3.965,4.965,5.965,6.965,7.965,8.965,9.965,10.965,11.965],[1.035,2.035,3.035,4.035,5.035,6.035,7.035,8.035,9.035,10.035,11.035,12.035],[1.105,2.105,3.105,4.105,5.105,6.105,7.105,8.105,9.105,10.105,11.105,12.105],[1.175,2.175,3.175,4.175,5.175,6.175,7.175,8.175,9.175,10.175,11.175,12.175],[1.245,2.245,3.245,4.245,5.245,6.245,7.245,8.245,9.245,10.245,11.245,12.245],[1.315,2.315,3.315,4.315,5.315,6.315,7.315,8.315,9.315,10.315,11.315,12.315],[1.385,2.385,3.385,4.385,5.385,6.385,7.385,8.385,9.385,10.385,11.385,12.385],[1.455,2.455,3.455,4.455,5.455,6.455,7.455,8.455,9.455,10.455,11.455,12.455],[1.525,2.525,3.525,4.525,5.525,6.525,7.525,8.525,9.525,10.525,11.525,12.525]],"filenames":["ABySS\_63","ABySS\_127","CLC","IDBA\_UD","MEGAHIT","metaVelvet","MIRA","Ray\_Meta","SOAPdenovo2","SPAdes","SPAdes\_meta","SPAdes\_sc","SPAdes\_sc\_careful","Velvet","vicuna","Geneious"]},
{"refnames":["KF302033","KF302035","KF302036","NC\_001330","NC\_001422","NC\_021796","KF302037","KC821629","NC\_021794","KC821625","NC\_021790","KF302034"],"coord\_y":[[null,0,null,0,0,0,null,55287,0,0,0,null],[null,0,null,null,0,0,null,null,0,0,0,null],[0,0,0,0,0,0,0,0,0,0,0,0],[0,0,0,0,0,0,0,0,0,0,0,0],[0,0,0,0,0,0,0,0,0,0,0,0],[0,0,0,null,null,0,0,0,0,null,0,0],[0,0,0,0,0,0,0,0,0,0,0,0],[0,0,0,0,0,0,45598,0,73023,0,40281,131360],[0,0,0,0,0,0,0,0,0,76440,0,0],[0,0,0,0,0,0,0,0,0,0,0,0],[0,0,0,0,0,0,0,0,0,0,0,0],[0,0,0,0,0,0,0,0,0,0,0,0],[0,0,0,0,0,0,0,0,0,0,0,0],[0,0,0,null,null,0,0,0,0,null,0,0],[0,0,0,0,0,0,0,0,0,0,0,0],[0,0,0,0,0,0,0,0,0,0,56262,153292]],"coord\_x":[[0.475,1.475,2.475,3.475,4.475,5.475,6.475,7.475,8.475,9.475,10.475,11.475],[0.5449999999999999,1.545,2.545,3.545,4.545,5.545,6.545,7.545,8.545,9.545,10.545,11.545],[0.615,1.615,2.615,3.615,4.615,5.615,6.615,7.615,8.615,9.615,10.615,11.615],[0.6849999999999999,1.685,2.685,3.685,4.685,5.685,6.685,7.685,8.685,9.685,10.685,11.685],[0.755,1.755,2.755,3.755,4.755,5.755,6.755,7.755,8.755,9.755,10.755,11.755],[0.825,1.825,2.825,3.825,4.825,5.825,6.825,7.825,8.825,9.825,10.825,11.825],[0.895,1.895,2.895,3.895,4.895,5.895,6.895,7.895,8.895,9.895,10.895,11.895],[0.965,1.965,2.965,3.965,4.965,5.965,6.965,7.965,8.965,9.965,10.965,11.965],[1.035,2.035,3.035,4.035,5.035,6.035,7.035,8.035,9.035,10.035,11.035,12.035],[1.105,2.105,3.105,4.105,5.105,6.105,7.105,8.105,9.105,10.105,11.105,12.105],[1.175,2.175,3.175,4.175,5.175,6.175,7.175,8.175,9.175,10.175,11.175,12.175],[1.245,2.245,3.245,4.245,5.245,6.245,7.245,8.245,9.245,10.245,11.245,12.245],[1.315,2.315,3.315,4.315,5.315,6.315,7.315,8.315,9.315,10.315,11.315,12.315],[1.385,2.385,3.385,4.385,5.385,6.385,7.385,8.385,9.385,10.385,11.385,12.385],[1.455,2.455,3.455,4.455,5.455,6.455,7.455,8.455,9.455,10.455,11.455,12.455],[1.525,2.525,3.525,4.525,5.525,6.525,7.525,8.525,9.525,10.525,11.525,12.525]],"filenames":["ABySS\_63","ABySS\_127","CLC","IDBA\_UD","MEGAHIT","metaVelvet","MIRA","Ray\_Meta","SOAPdenovo2","SPAdes","SPAdes\_meta","SPAdes\_sc","SPAdes\_sc\_careful","Velvet","vicuna","Geneious"]},
{"refnames":["NC\_021794","KF302037","KF302036","NC\_021796","KF302034","KC821629","KF302033","KC821625","KF302035","NC\_021790","NC\_001330","NC\_001422"],"coord\_y":[[0.0,null,null,1.38,null,0.0,null,1.3,0.0,0.0,147.86,297.07],[0.0,null,null,0.0,null,null,null,1.3,0.0,0.0,null,311.41],[0.0,0.0,0.0,0.0,0.0,0.0,2.72,1.31,0.0,0.0,147.86,297.07],[0.0,0.0,0.0,0.0,0.0,0.0,2.72,1.3,0.0,0.0,147.86,298.23],[0.0,0.0,0.0,0.0,0.0,0.0,2.72,1.3,0.0,0.0,147.86,298.79],[0.0,0.0,0.0,1.38,0.0,0.0,2.72,null,0.0,0.0,null,null],[0.0,0.0,0.0,1.38,0.0,0.0,2.72,1.3,2.84,0.0,147.86,305.75],[0.0,0.0,0.0,1.39,0.0,1.85,2.72,1.31,0.0,0.0,147.86,298.79],[0.0,0.0,0.0,12.85,31.04,36.39,44.58,49.86,227.45,1000.0,147.88,298.79],[0.0,0.0,0.0,0.0,0.0,0.0,2.72,1.3,0.0,0.0,147.86,298.23],[0.0,0.0,0.0,0.0,0.0,0.0,2.72,1.3,0.0,0.0,147.86,298.23],[0.0,0.0,0.0,0.0,0.0,0.0,2.72,16.96,0.0,0.0,147.86,297.07],[0.0,0.0,0.0,0.0,0.0,1.85,2.72,1.31,0.0,0.0,147.86,298.23],[0.0,0.0,0.0,1.38,0.0,0.0,2.72,null,0.0,0.0,null,null],[1.93,4.35,4.77,0.0,0.85,0.0,2.72,1.73,0.0,0.0,159.31,338.79],[0.0,0.0,0.0,0.0,0.0,0.0,2.72,1.3,0.0,0.0,147.86,298.23]],"coord\_x":[[0.475,1.475,2.475,3.475,4.475,5.475,6.475,7.475,8.475,9.475,10.475,11.475],[0.5449999999999999,1.545,2.545,3.545,4.545,5.545,6.545,7.545,8.545,9.545,10.545,11.545],[0.615,1.615,2.615,3.615,4.615,5.615,6.615,7.615,8.615,9.615,10.615,11.615],[0.6849999999999999,1.685,2.685,3.685,4.685,5.685,6.685,7.685,8.685,9.685,10.685,11.685],[0.755,1.755,2.755,3.755,4.755,5.755,6.755,7.755,8.755,9.755,10.755,11.755],[0.825,1.825,2.825,3.825,4.825,5.825,6.825,7.825,8.825,9.825,10.825,11.825],[0.895,1.895,2.895,3.895,4.895,5.895,6.895,7.895,8.895,9.895,10.895,11.895],[0.965,1.965,2.965,3.965,4.965,5.965,6.965,7.965,8.965,9.965,10.965,11.965],[1.035,2.035,3.035,4.035,5.035,6.035,7.035,8.035,9.035,10.035,11.035,12.035],[1.105,2.105,3.105,4.105,5.105,6.105,7.105,8.105,9.105,10.105,11.105,12.105],[1.175,2.175,3.175,4.175,5.175,6.175,7.175,8.175,9.175,10.175,11.175,12.175],[1.245,2.245,3.245,4.245,5.245,6.245,7.245,8.245,9.245,10.245,11.245,12.245],[1.315,2.315,3.315,4.315,5.315,6.315,7.315,8.315,9.315,10.315,11.315,12.315],[1.385,2.385,3.385,4.385,5.385,6.385,7.385,8.385,9.385,10.385,11.385,12.385],[1.455,2.455,3.455,4.455,5.455,6.455,7.455,8.455,9.455,10.455,11.455,12.455],[1.525,2.525,3.525,4.525,5.525,6.525,7.525,8.525,9.525,10.525,11.525,12.525]],"filenames":["ABySS\_63","ABySS\_127","CLC","IDBA\_UD","MEGAHIT","metaVelvet","MIRA","Ray\_Meta","SOAPdenovo2","SPAdes","SPAdes\_meta","SPAdes\_sc","SPAdes\_sc\_careful","Velvet","vicuna","Geneious"]},
{"refnames":["NC\_021796","KF302033","KF302034","KC821625","KF302036","KF302037","NC\_021790","NC\_001422","KC821629","NC\_021794","KF302035","NC\_001330"],"coord\_y":[[2.76,null,null,0.0,null,null,5.1,18.57,9.26,9.8,5.69,49.29],[1.38,null,null,0.0,null,null,5.1,19.46,null,8.4,5.69,null],[1.38,5.44,0.77,0.0,0.0,0.0,5.1,0.0,11.11,8.4,5.68,32.86],[1.38,5.44,0.0,0.0,0.0,0.0,5.1,0.0,5.55,8.4,5.69,32.86],[1.38,5.44,0.0,0.0,0.0,0.0,5.1,0.0,7.41,8.4,5.69,32.86],[2.76,5.43,1.55,null,0.0,0.0,12.82,null,13.05,5.63,8.53,null],[2.76,5.44,0.0,0.0,0.0,0.0,5.1,0.0,9.26,7.0,8.53,32.86],[2.77,5.44,0.0,0.0,0.0,0.0,5.1,0.0,9.26,8.4,5.69,32.86],[38.54,0.0,62.08,62.68,82.85,100.7,0.0,74.7,0.0,90.98,151.63,82.16],[1.38,2.72,0.77,1.3,0.0,0.0,5.1,0.0,9.26,9.8,5.69,49.29],[1.38,5.44,0.77,3.91,0.0,0.0,5.1,0.0,12.96,9.8,5.69,49.29],[1.38,5.44,0.0,5.22,0.0,0.0,5.1,18.57,11.11,8.4,5.69,49.29],[1.38,5.44,0.77,1.31,0.0,0.0,5.1,0.0,11.11,8.4,5.69,49.29],[2.76,5.43,1.55,null,0.0,0.0,12.82,null,13.05,5.63,8.53,null],[1.95,5.44,0.85,1.73,0.0,0.0,5.1,0.0,7.41,5.78,5.69,0.0],[1.38,5.44,0.77,0.0,0.0,0.0,20.41,0.0,7.41,8.4,5.69,49.29]],"coord\_x":[[0.475,1.475,2.475,3.475,4.475,5.475,6.475,7.475,8.475,9.475,10.475,11.475],[0.5449999999999999,1.545,2.545,3.545,4.545,5.545,6.545,7.545,8.545,9.545,10.545,11.545],[0.615,1.615,2.615,3.615,4.615,5.615,6.615,7.615,8.615,9.615,10.615,11.615],[0.6849999999999999,1.685,2.685,3.685,4.685,5.685,6.685,7.685,8.685,9.685,10.685,11.685],[0.755,1.755,2.755,3.755,4.755,5.755,6.755,7.755,8.755,9.755,10.755,11.755],[0.825,1.825,2.825,3.825,4.825,5.825,6.825,7.825,8.825,9.825,10.825,11.825],[0.895,1.895,2.895,3.895,4.895,5.895,6.895,7.895,8.895,9.895,10.895,11.895],[0.965,1.965,2.965,3.965,4.965,5.965,6.965,7.965,8.965,9.965,10.965,11.965],[1.035,2.035,3.035,4.035,5.035,6.035,7.035,8.035,9.035,10.035,11.035,12.035],[1.105,2.105,3.105,4.105,5.105,6.105,7.105,8.105,9.105,10.105,11.105,12.105],[1.175,2.175,3.175,4.175,5.175,6.175,7.175,8.175,9.175,10.175,11.175,12.175],[1.245,2.245,3.245,4.245,5.245,6.245,7.245,8.245,9.245,10.245,11.245,12.245],[1.315,2.315,3.315,4.315,5.315,6.315,7.315,8.315,9.315,10.315,11.315,12.315],[1.385,2.385,3.385,4.385,5.385,6.385,7.385,8.385,9.385,10.385,11.385,12.385],[1.455,2.455,3.455,4.455,5.455,6.455,7.455,8.455,9.455,10.455,11.455,12.455],[1.525,2.525,3.525,4.525,5.525,6.525,7.525,8.525,9.525,10.525,11.525,12.525]],"filenames":["ABySS\_63","ABySS\_127","CLC","IDBA\_UD","MEGAHIT","metaVelvet","MIRA","Ray\_Meta","SOAPdenovo2","SPAdes","SPAdes\_meta","SPAdes\_sc","SPAdes\_sc\_careful","Velvet","vicuna","Geneious"]},
{"refnames":["NC\_001330","KC821625","NC\_001422","KF302036","KF302037","NC\_021794","NC\_021796","KF302034","KF302033","KC821629","KF302035","NC\_021790"],"coord\_y":[[0.0,0.0,0.0,null,null,0.0,0.0,null,null,0.0,0.0,127.02],[null,0.0,1376.5,null,null,0.0,0.0,null,null,null,0.0,0.0],[0.0,0.0,0.0,0.0,0.0,0.0,0.0,0.0,0.0,0.0,0.0,0.0],[0.0,0.0,0.0,0.0,0.0,0.0,0.0,0.0,0.0,0.0,0.0,0.0],[0.0,0.0,0.0,0.0,0.0,0.0,0.0,0.0,0.0,0.0,0.0,0.0],[null,null,null,0.0,0.0,0.0,0.0,0.0,0.0,0.0,0.0,0.0],[0.0,1.19,0.0,0.0,0.0,0.0,1.25,0.0,0.0,0.0,2.12,0.0],[0.0,0.0,0.0,0.0,0.0,0.0,0.0,0.0,0.0,0.0,0.0,0.0],[696.06,2386.19,2984.24,8340.41,10890.4,16319.64,25061.5,31903.88,33468.41,44247.55,47870.47,56159.42],[0.0,0.0,0.0,0.0,0.0,0.0,0.0,0.0,0.0,0.0,0.0,0.0],[0.0,0.0,0.0,0.0,0.0,0.0,0.0,0.0,0.0,0.0,0.0,0.0],[0.0,0.0,0.0,0.0,0.0,0.0,0.0,0.0,0.0,0.0,0.0,0.0],[0.0,0.0,0.0,0.0,0.0,0.0,0.0,0.0,0.0,0.0,0.0,0.0],[null,null,null,0.0,0.0,0.0,0.0,0.0,0.0,0.0,0.0,0.0],[0.0,0.0,0.0,0.0,0.0,0.0,0.0,0.0,0.0,0.0,0.0,0.0],[0.0,0.0,0.0,0.0,0.0,0.0,2.74,37.18,2.69,5.54,62.45,442.57]],"coord\_x":[[0.475,1.475,2.475,3.475,4.475,5.475,6.475,7.475,8.475,9.475,10.475,11.475],[0.5449999999999999,1.545,2.545,3.545,4.545,5.545,6.545,7.545,8.545,9.545,10.545,11.545],[0.615,1.615,2.615,3.615,4.615,5.615,6.615,7.615,8.615,9.615,10.615,11.615],[0.6849999999999999,1.685,2.685,3.685,4.685,5.685,6.685,7.685,8.685,9.685,10.685,11.685],[0.755,1.755,2.755,3.755,4.755,5.755,6.755,7.755,8.755,9.755,10.755,11.755],[0.825,1.825,2.825,3.825,4.825,5.825,6.825,7.825,8.825,9.825,10.825,11.825],[0.895,1.895,2.895,3.895,4.895,5.895,6.895,7.895,8.895,9.895,10.895,11.895],[0.965,1.965,2.965,3.965,4.965,5.965,6.965,7.965,8.965,9.965,10.965,11.965],[1.035,2.035,3.035,4.035,5.035,6.035,7.035,8.035,9.035,10.035,11.035,12.035],[1.105,2.105,3.105,4.105,5.105,6.105,7.105,8.105,9.105,10.105,11.105,12.105],[1.175,2.175,3.175,4.175,5.175,6.175,7.175,8.175,9.175,10.175,11.175,12.175],[1.245,2.245,3.245,4.245,5.245,6.245,7.245,8.245,9.245,10.245,11.245,12.245],[1.315,2.315,3.315,4.315,5.315,6.315,7.315,8.315,9.315,10.315,11.315,12.315],[1.385,2.385,3.385,4.385,5.385,6.385,7.385,8.385,9.385,10.385,11.385,12.385],[1.455,2.455,3.455,4.455,5.455,6.455,7.455,8.455,9.455,10.455,11.455,12.455],[1.525,2.525,3.525,4.525,5.525,6.525,7.525,8.525,9.525,10.525,11.525,12.525]],"filenames":["ABySS\_63","ABySS\_127","CLC","IDBA\_UD","MEGAHIT","metaVelvet","MIRA","Ray\_Meta","SOAPdenovo2","SPAdes","SPAdes\_meta","SPAdes\_sc","SPAdes\_sc\_careful","Velvet","vicuna","Geneious"]},
{"refnames":["NC\_021790","KF302035","NC\_021796","NC\_021794","KC821629","KC821625","NC\_001422","KF302034","KF302033","NC\_001330","KF302037","KF302036"],"coord\_y":[[100.0,99.558,100.0,100.0,100.0,99.978,100.0,null,null,100.0,null,null],[100.0,99.558,99.971,100.0,null,100.0,95.395,null,null,null,null,null],[100.0,99.584,99.97,100.0,100.0,99.873,100.0,99.981,98.812,100.0,98.925,98.686],[100.0,99.567,100.0,100.0,100.0,100.0,99.61,99.546,98.812,100.0,98.925,98.744],[100.0,99.558,100.0,100.0,100.0,100.0,99.424,99.981,98.812,100.0,98.925,98.744],[99.546,99.558,99.946,99.484,99.343,null,null,99.981,98.979,null,98.925,89.743],[100.0,99.558,100.0,100.0,100.0,100.0,97.159,99.816,98.812,100.0,98.925,98.744],[100.0,99.558,99.409,100.0,100.0,99.849,99.424,99.981,98.812,100.0,98.925,98.744],[0.255,3.733,10.732,7.693,5.088,91.57,99.424,12.445,12.056,99.984,6.615,6.318],[100.0,99.558,100.0,100.0,100.0,100.0,99.61,99.981,98.812,100.0,98.925,98.744],[100.0,99.558,100.0,100.0,100.0,100.0,99.61,99.981,98.812,100.0,98.925,98.744],[100.0,99.558,100.0,99.945,100.0,100.0,100.0,99.466,98.812,100.0,98.925,98.744],[100.0,99.556,100.0,99.945,100.0,99.86,99.61,99.981,98.812,100.0,98.925,98.744],[99.546,99.558,99.946,99.484,99.343,null,null,99.981,98.979,null,98.925,89.743],[100.0,99.558,70.768,72.612,99.92,75.336,32.882,91.183,98.812,72.187,51.054,54.824],[100.0,99.558,100.0,100.0,100.0,100.0,99.61,100.0,98.812,100.0,98.925,98.744]],"coord\_x":[[0.475,1.475,2.475,3.475,4.475,5.475,6.475,7.475,8.475,9.475,10.475,11.475],[0.5449999999999999,1.545,2.545,3.545,4.545,5.545,6.545,7.545,8.545,9.545,10.545,11.545],[0.615,1.615,2.615,3.615,4.615,5.615,6.615,7.615,8.615,9.615,10.615,11.615],[0.6849999999999999,1.685,2.685,3.685,4.685,5.685,6.685,7.685,8.685,9.685,10.685,11.685],[0.755,1.755,2.755,3.755,4.755,5.755,6.755,7.755,8.755,9.755,10.755,11.755],[0.825,1.825,2.825,3.825,4.825,5.825,6.825,7.825,8.825,9.825,10.825,11.825],[0.895,1.895,2.895,3.895,4.895,5.895,6.895,7.895,8.895,9.895,10.895,11.895],[0.965,1.965,2.965,3.965,4.965,5.965,6.965,7.965,8.965,9.965,10.965,11.965],[1.035,2.035,3.035,4.035,5.035,6.035,7.035,8.035,9.035,10.035,11.035,12.035],[1.105,2.105,3.105,4.105,5.105,6.105,7.105,8.105,9.105,10.105,11.105,12.105],[1.175,2.175,3.175,4.175,5.175,6.175,7.175,8.175,9.175,10.175,11.175,12.175],[1.245,2.245,3.245,4.245,5.245,6.245,7.245,8.245,9.245,10.245,11.245,12.245],[1.315,2.315,3.315,4.315,5.315,6.315,7.315,8.315,9.315,10.315,11.315,12.315],[1.385,2.385,3.385,4.385,5.385,6.385,7.385,8.385,9.385,10.385,11.385,12.385],[1.455,2.455,3.455,4.455,5.455,6.455,7.455,8.455,9.455,10.455,11.455,12.455],[1.525,2.525,3.525,4.525,5.525,6.525,7.525,8.525,9.525,10.525,11.525,12.525]],"filenames":["ABySS\_63","ABySS\_127","CLC","IDBA\_UD","MEGAHIT","metaVelvet","MIRA","Ray\_Meta","SOAPdenovo2","SPAdes","SPAdes\_meta","SPAdes\_sc","SPAdes\_sc\_careful","Velvet","vicuna","Geneious"]},
{"refnames":["NC\_001330","KF302033","NC\_001422","KF302037","KF302036","KF302034","KC821625","KC821629","NC\_021796","NC\_021794","KF302035","NC\_021790"],"coord\_y":[[1.01,null,1.011,null,null,null,1.0,1.024,1.0,1.001,1.0,1.004],[null,null,1.004,null,null,null,1.002,null,1.0,1.0,1.0,1.023],[1.0,1.0,1.0,1.0,1.001,1.0,1.053,1.0,1.0,1.0,1.974,1.0],[1.017,1.003,1.005,1.002,1.003,1.005,1.001,1.002,1.001,1.001,1.0,1.003],[1.023,1.004,1.004,1.003,1.004,1.001,1.002,1.003,1.002,1.002,1.001,1.003],[null,1.0,null,1.0,1.0,1.0,null,1.004,1.002,1.0,1.0,1.004],[1.008,1.006,1.0,1.003,1.007,1.009,1.097,1.168,1.104,1.012,1.344,1.093],[1.069,1.02,1.004,1.024,1.024,1.015,1.001,1.015,1.0,1.022,1.0,1.028],[0.991,1.0,1.007,0.977,0.981,0.984,0.999,1.555,0.997,0.989,1.703,5.52],[1.013,1.002,1.005,1.002,1.002,1.001,1.001,1.001,1.002,1.001,1.003,1.004],[1.008,1.002,1.005,1.001,1.001,1.0,1.003,1.001,1.001,1.001,1.002,1.002],[1.008,1.002,1.008,1.001,1.001,1.006,1.004,1.001,1.001,1.0,1.0,1.002],[1.008,1.002,1.013,1.001,1.001,1.0,1.0,1.001,1.001,1.0,1.001,1.002],[null,1.0,null,1.0,1.0,1.0,null,1.004,1.002,1.0,1.0,1.004],[1.182,1.006,1.006,1.16,1.216,1.083,1.185,1.002,1.057,1.142,1.001,1.009],[1.049,1.01,1.007,1.006,1.01,1.184,1.005,1.002,1.005,1.009,1.002,1.436]],"coord\_x":[[0.475,1.475,2.475,3.475,4.475,5.475,6.475,7.475,8.475,9.475,10.475,11.475],[0.5449999999999999,1.545,2.545,3.545,4.545,5.545,6.545,7.545,8.545,9.545,10.545,11.545],[0.615,1.615,2.615,3.615,4.615,5.615,6.615,7.615,8.615,9.615,10.615,11.615],[0.6849999999999999,1.685,2.685,3.685,4.685,5.685,6.685,7.685,8.685,9.685,10.685,11.685],[0.755,1.755,2.755,3.755,4.755,5.755,6.755,7.755,8.755,9.755,10.755,11.755],[0.825,1.825,2.825,3.825,4.825,5.825,6.825,7.825,8.825,9.825,10.825,11.825],[0.895,1.895,2.895,3.895,4.895,5.895,6.895,7.895,8.895,9.895,10.895,11.895],[0.965,1.965,2.965,3.965,4.965,5.965,6.965,7.965,8.965,9.965,10.965,11.965],[1.035,2.035,3.035,4.035,5.035,6.035,7.035,8.035,9.035,10.035,11.035,12.035],[1.105,2.105,3.105,4.105,5.105,6.105,7.105,8.105,9.105,10.105,11.105,12.105],[1.175,2.175,3.175,4.175,5.175,6.175,7.175,8.175,9.175,10.175,11.175,12.175],[1.245,2.245,3.245,4.245,5.245,6.245,7.245,8.245,9.245,10.245,11.245,12.245],[1.315,2.315,3.315,4.315,5.315,6.315,7.315,8.315,9.315,10.315,11.315,12.315],[1.385,2.385,3.385,4.385,5.385,6.385,7.385,8.385,9.385,10.385,11.385,12.385],[1.455,2.455,3.455,4.455,5.455,6.455,7.455,8.455,9.455,10.455,11.455,12.455],[1.525,2.525,3.525,4.525,5.525,6.525,7.525,8.525,9.525,10.525,11.525,12.525]],"filenames":["ABySS\_63","ABySS\_127","CLC","IDBA\_UD","MEGAHIT","metaVelvet","MIRA","Ray\_Meta","SOAPdenovo2","SPAdes","SPAdes\_meta","SPAdes\_sc","SPAdes\_sc\_careful","Velvet","vicuna","Geneious"]},
{"refnames":["KF302034","NC\_021794","KC821625","NC\_021796","KC821629","NC\_021790","KF302037","KF302035","KF302033","KF302036","NC\_001330","NC\_001422"],"coord\_y":[[null,71499,76649,72541,44392,38543,null,35176,null,null,6148,5445],[null,71457,76666,72514,null,40089,null,35176,null,null,null,5158],[129415,71449,52891,72513,54016,39191,44551,35176,36771,37706,6089,5386],[128851,71548,76765,72634,54115,39290,44650,35179,36870,37827,6188,5365],[129556,71590,76807,72676,54157,38545,44692,35176,36912,37869,6230,5355],[129478,38880,null,36945,35739,22872,44558,35177,36833,20798,null,null],[128970,71686,76666,17449,35913,39164,44678,3869,36990,37981,6089,5233],[129048,69825,76550,72105,54799,23054,26064,35176,37514,38637,6508,5355],[null,null,33833,null,null,null,null,null,null,null,6034,5373],[129492,71526,76743,70642,54093,38545,44628,35176,36848,37805,6166,5365],[129470,71485,76925,70598,54041,38521,44606,35176,36826,37783,6134,5365],[128748,71410,76993,70598,54042,38521,44606,35176,36826,37783,6134,5431],[129470,71410,76559,70598,54042,38521,44606,35175,36826,37783,6134,4783],[129478,38880,null,36945,35739,22872,44558,35177,36833,20798,null,null],[24884,572,659,656,43063,39552,507,35176,36998,510,515,null],[129441,72069,76666,72916,54016,39166,44829,35176,37131,38107,6385,5365]],"coord\_x":[[0.475,1.475,2.475,3.475,4.475,5.475,6.475,7.475,8.475,9.475,10.475,11.475],[0.5449999999999999,1.545,2.545,3.545,4.545,5.545,6.545,7.545,8.545,9.545,10.545,11.545],[0.615,1.615,2.615,3.615,4.615,5.615,6.615,7.615,8.615,9.615,10.615,11.615],[0.6849999999999999,1.685,2.685,3.685,4.685,5.685,6.685,7.685,8.685,9.685,10.685,11.685],[0.755,1.755,2.755,3.755,4.755,5.755,6.755,7.755,8.755,9.755,10.755,11.755],[0.825,1.825,2.825,3.825,4.825,5.825,6.825,7.825,8.825,9.825,10.825,11.825],[0.895,1.895,2.895,3.895,4.895,5.895,6.895,7.895,8.895,9.895,10.895,11.895],[0.965,1.965,2.965,3.965,4.965,5.965,6.965,7.965,8.965,9.965,10.965,11.965],[1.035,2.035,3.035,4.035,5.035,6.035,7.035,8.035,9.035,10.035,11.035,12.035],[1.105,2.105,3.105,4.105,5.105,6.105,7.105,8.105,9.105,10.105,11.105,12.105],[1.175,2.175,3.175,4.175,5.175,6.175,7.175,8.175,9.175,10.175,11.175,12.175],[1.245,2.245,3.245,4.245,5.245,6.245,7.245,8.245,9.245,10.245,11.245,12.245],[1.315,2.315,3.315,4.315,5.315,6.315,7.315,8.315,9.315,10.315,11.315,12.315],[1.385,2.385,3.385,4.385,5.385,6.385,7.385,8.385,9.385,10.385,11.385,12.385],[1.455,2.455,3.455,4.455,5.455,6.455,7.455,8.455,9.455,10.455,11.455,12.455],[1.525,2.525,3.525,4.525,5.525,6.525,7.525,8.525,9.525,10.525,11.525,12.525]],"filenames":["ABySS\_63","ABySS\_127","CLC","IDBA\_UD","MEGAHIT","metaVelvet","MIRA","Ray\_Meta","SOAPdenovo2","SPAdes","SPAdes\_meta","SPAdes\_sc","SPAdes\_sc\_careful","Velvet","vicuna","Geneious"]}]

{"refnames":["KC821625","KC821629","KF302033","KF302034","KF302035","KF302036","KF302037","NC\_001330","NC\_001422","NC\_021790","NC\_021794","NC\_021796"],"coord\_y":[[0.0,0.0,0.0,1.0,0.0,0.0,0,0,0,0,0,0,0.0,0.0,0.0,0,0,0,0,0,0,0.0,0.0,0.0,0.0,0.0,0.0,0.0,0.0,0.0,0.0,0.0,0.0,0.0,0.0,0.0],[0.0,0.0,0.0,0,0,0,0,0,0,0,0,0,0.0,0.0,0.0,0,0,0,0,0,0,0,0,0,0.0,0.0,0.0,0.0,0.0,0.0,0.0,0.0,0.0,0.0,0.0,0.0],[0.0,0.0,0.0,0.0,0.0,0.0,0.0,0.0,0.0,0.0,0.0,0.0,0.0,0.0,0.0,0.0,0.0,0.0,0.0,0.0,0.0,0.0,0.0,0.0,0.0,0.0,0.0,0.0,0.0,0.0,0.0,0.0,0.0,0.0,0.0,0.0],[0.0,0.0,0.0,0.0,0.0,0.0,0.0,0.0,0.0,0.0,0.0,0.0,0.0,0.0,0.0,0.0,0.0,0.0,0.0,0.0,0.0,0.0,0.0,0.0,0.0,0.0,0.0,0.0,0.0,0.0,0.0,0.0,0.0,0.0,0.0,0.0],[0.0,0.0,0.0,0.0,0.0,0.0,0.0,0.0,0.0,0.0,0.0,0.0,0.0,0.0,0.0,0.0,0.0,0.0,0.0,0.0,0.0,0.0,0.0,0.0,0.0,0.0,0.0,0.0,0.0,0.0,0.0,0.0,0.0,0.0,0.0,0.0],[0,0,0,0.0,0.0,0.0,0.0,0.0,0.0,0.0,0.0,0.0,0.0,0.0,0.0,0.0,0.0,0.0,0.0,0.0,0.0,0,0,0,0,0,0,0.0,0.0,0.0,0.0,0.0,0.0,0.0,0.0,0.0],[0.0,0.0,0.0,0.0,0.0,0.0,0.0,0.0,0.0,0.0,0.0,0.0,0.0,0.0,0.0,0.0,0.0,0.0,0.0,0.0,0.0,0.0,0.0,0.0,0.0,0.0,0.0,0.0,0.0,0.0,0.0,0.0,0.0,0.0,0.0,0.0],[0.0,0.0,0.0,0.0,0.0,0.0,0.0,0.0,0.0,1.0,0.0,0.0,0.0,0.0,0.0,0.0,0.0,0.0,1.0,0.0,0.0,0.0,0.0,0.0,0.0,0.0,0.0,1.0,0.0,0.0,1.0,0.0,0.0,0.0,0.0,0.0],[1.0,0.0,0.0,0.0,0.0,0.0,0.0,0.0,0.0,0.0,0.0,0.0,0.0,0.0,0.0,0.0,0.0,0.0,0.0,0.0,0.0,0.0,0.0,0.0,0.0,0.0,0.0,0.0,0.0,0.0,0.0,0.0,0.0,0.0,0.0,0.0],[0.0,0.0,0.0,0.0,0.0,0.0,0.0,0.0,0.0,0.0,0.0,0.0,0.0,0.0,0.0,0.0,0.0,0.0,0.0,0.0,0.0,0.0,0.0,0.0,0.0,0.0,0.0,0.0,0.0,0.0,0.0,0.0,0.0,0.0,0.0,0.0],[0.0,0.0,0.0,0.0,0.0,0.0,0.0,0.0,0.0,0.0,0.0,0.0,0.0,0.0,0.0,0.0,0.0,0.0,0.0,0.0,0.0,0.0,0.0,0.0,0.0,0.0,0.0,0.0,0.0,0.0,0.0,0.0,0.0,0.0,0.0,0.0],[0.0,0.0,0.0,0.0,0.0,0.0,0.0,0.0,0.0,0.0,0.0,0.0,0.0,0.0,0.0,0.0,0.0,0.0,0.0,0.0,0.0,0.0,0.0,0.0,0.0,0.0,0.0,0.0,0.0,0.0,0.0,0.0,0.0,0.0,0.0,0.0],[0.0,0.0,0.0,0.0,0.0,0.0,0.0,0.0,0.0,0.0,0.0,0.0,0.0,0.0,0.0,0.0,0.0,0.0,0.0,0.0,0.0,0.0,0.0,0.0,0.0,0.0,0.0,0.0,0.0,0.0,0.0,0.0,0.0,0.0,0.0,0.0],[0,0,0,0.0,0.0,0.0,0.0,0.0,0.0,0.0,0.0,0.0,0.0,0.0,0.0,0.0,0.0,0.0,0.0,0.0,0.0,0,0,0,0,0,0,0.0,0.0,0.0,0.0,0.0,0.0,0.0,0.0,0.0],[0.0,0.0,0.0,0.0,0.0,0.0,0.0,0.0,0.0,0.0,0.0,0.0,0.0,0.0,0.0,0.0,0.0,0.0,0.0,0.0,0.0,0.0,0.0,0.0,0.0,0.0,0.0,0.0,0.0,0.0,0.0,0.0,0.0,0.0,0.0,0.0],[0.0,0.0,0.0,0.0,0.0,0.0,0.0,0.0,0.0,2.0,0.0,0.0,0.0,0.0,0.0,0.0,0.0,0.0,0.0,0.0,0.0,0.0,0.0,0.0,0.0,0.0,0.0,1.0,0.0,0.0,0.0,0.0,0.0,0.0,0.0,0.0]],"coord\_x":[[1,1,1,2,2,2,3,3,3,4,4,4,5,5,5,6,6,6,7,7,7,8,8,8,9,9,9,10,10,10,11,11,11,12,12,12],[1,1,1,2,2,2,3,3,3,4,4,4,5,5,5,6,6,6,7,7,7,8,8,8,9,9,9,10,10,10,11,11,11,12,12,12],[1,1,1,2,2,2,3,3,3,4,4,4,5,5,5,6,6,6,7,7,7,8,8,8,9,9,9,10,10,10,11,11,11,12,12,12],[1,1,1,2,2,2,3,3,3,4,4,4,5,5,5,6,6,6,7,7,7,8,8,8,9,9,9,10,10,10,11,11,11,12,12,12],[1,1,1,2,2,2,3,3,3,4,4,4,5,5,5,6,6,6,7,7,7,8,8,8,9,9,9,10,10,10,11,11,11,12,12,12],[1,1,1,2,2,2,3,3,3,4,4,4,5,5,5,6,6,6,7,7,7,8,8,8,9,9,9,10,10,10,11,11,11,12,12,12],[1,1,1,2,2,2,3,3,3,4,4,4,5,5,5,6,6,6,7,7,7,8,8,8,9,9,9,10,10,10,11,11,11,12,12,12],[1,1,1,2,2,2,3,3,3,4,4,4,5,5,5,6,6,6,7,7,7,8,8,8,9,9,9,10,10,10,11,11,11,12,12,12],[1,1,1,2,2,2,3,3,3,4,4,4,5,5,5,6,6,6,7,7,7,8,8,8,9,9,9,10,10,10,11,11,11,12,12,12],[1,1,1,2,2,2,3,3,3,4,4,4,5,5,5,6,6,6,7,7,7,8,8,8,9,9,9,10,10,10,11,11,11,12,12,12],[1,1,1,2,2,2,3,3,3,4,4,4,5,5,5,6,6,6,7,7,7,8,8,8,9,9,9,10,10,10,11,11,11,12,12,12],[1,1,1,2,2,2,3,3,3,4,4,4,5,5,5,6,6,6,7,7,7,8,8,8,9,9,9,10,10,10,11,11,11,12,12,12],[1,1,1,2,2,2,3,3,3,4,4,4,5,5,5,6,6,6,7,7,7,8,8,8,9,9,9,10,10,10,11,11,11,12,12,12],[1,1,1,2,2,2,3,3,3,4,4,4,5,5,5,6,6,6,7,7,7,8,8,8,9,9,9,10,10,10,11,11,11,12,12,12],[1,1,1,2,2,2,3,3,3,4,4,4,5,5,5,6,6,6,7,7,7,8,8,8,9,9,9,10,10,10,11,11,11,12,12,12],[1,1,1,2,2,2,3,3,3,4,4,4,5,5,5,6,6,6,7,7,7,8,8,8,9,9,9,10,10,10,11,11,11,12,12,12]],"filenames":["ABySS\_63","ABySS\_127","CLC","IDBA\_UD","MEGAHIT","metaVelvet","MIRA","Ray\_Meta","SOAPdenovo2","SPAdes","SPAdes\_meta","SPAdes\_sc","SPAdes\_sc\_careful","Velvet","vicuna","Geneious"]}

{}

{}

{"links\_names":["View in Icarus contig browser"],"links":["icarus.html"]}

{
"# contigs" : "is the total number of contigs in the assembly.",
"Largest contig" : "is the length of the longest contig in the assembly.",
"Total length" : "is the total number of bases in the assembly.",
"Reference length" : "is the total number of bases in the reference.",
"# contigs (>= 0 bp)" : "is the total number of contigs in the assembly that have size greater than or equal to 0 bp.",
"Total length (>= 0 bp)" : "is the total number of bases in the contigs having size greater than or equal to 0 bp.",
"N50" : "is the contig length such that using longer or equal length contigs produces half (50%) of the bases of the assembly. Usually there is no value that produces exactly 50%, so the technical definition is the maximum length x such that using contigs of length at least x accounts for at least 50% of the total assembly length.",
"NG50" : "is the contig length such that using longer or equal length contigs produces half (50%) of the bases of the reference genome. This metric is computed only if a reference genome is provided.",
"N75" : "is the contig length such that using longer or equal length contigs produces 75% of the bases of the assembly. Usually there is no value that produces exactly 75%, so the technical definition is the maximum length x such that using contigs of length at least x accounts for at least 75% of the total assembly length.",
"NG75" : "is the contig length such that using longer or equal length contigs produces 75% of the bases of the reference genome. This metric is computed only if a reference genome is provided.",
"L50" : "is the minimum number of contigs that produce half (50%) of the bases of the assembly. In other words, it's the number of contigs of length at least N50.",
"LG50" : "is the minimum number of contigs that produce half (50%) of the bases of the reference genome. In other words, it's the number of contigs of length at least NG50. This metric is computed only if a reference genome is provided.",
"L75" : "is the minimum number of contigs that produce 75% of the bases of the assembly. In other words, it's the number of contigs of length at least N75.",
"LG75" : "is the minimum number of contigs that produce 75% of the bases of the reference genome. In other words, it's the number of contigs of length at least NG75. This metric is computed only if a reference genome is provided.",
"NA50" : "is N50 where the lengths of aligned blocks are counted instead of contig lengths. I.e., if a contig has a misassembly with respect to the reference, the contig is broken into smaller pieces. This metric is computed only if a reference genome is provided.",
"NGA50" : "is NG50 where the lengths of aligned blocks are counted instead of contig lengths. I.e., if a contig has a misassembly with respect to the reference, the contig is broken into smaller pieces. This metric is computed only if a reference genome is provided.",
"NA75" : "is N75 where the lengths of aligned blocks are counted instead of contig lengths. I.e., if a contig has a misassembly with respect to the reference, the contig is broken into smaller pieces. This metric is computed only if a reference genome is provided.",
"NGA75" : "is NG75 where the lengths of aligned blocks are counted instead of contig lengths. I.e., if a contig has a misassembly with respect to the reference, the contig is broken into smaller pieces. This metric is computed only if a reference genome is provided.",
"LA50" : "is L50 where aligned blocks are counted instead of contigs. I.e., if a contig has a misassembly with respect to the reference, the contig is broken into smaller pieces.",
"LGA50" : "is LG50 where aligned blocks are counted instead of contigs. I.e., if a contig has a misassembly with respect to the reference, the contig is broken into smaller pieces.",
"LA75" : "is L75 where aligned blocks are counted instead of contigs. I.e., if a contig has a misassembly with respect to the reference, the contig is broken into smaller pieces.",
"LGA75" : "is LG75 where aligned blocks are counted instead of contigs. I.e., if a contig has a misassembly with respect to the reference, the contig is broken into smaller pieces.",
"Average %IDY" : "is the average of alignment identity percent (Nucmer measure of alignment accuracy) among all contigs.",
"# misassemblies" : "is the number of positions in the assembled contigs where the left flanking sequence aligns over 1 kbp away from the right flanking sequence on the reference (*relocation*) or they overlap on more than 1 kbp (*relocation*) or flanking sequences align on different strands (*inversion*) or different chromosomes (*translocation*).",
"# misassembled contigs" : "is the number of contigs that contain misassembly events.",
"Misassembled contigs length" : "is the number of total bases contained in all contigs that have one or more misassemblies.",
"# relocations" : "is the number of relocation events among all misassembly events. Relocation is a misassembly where the left flanking sequence aligns over 1 kbp away from the right flanking sequence on the reference, or they overlap by more than 1 kbp and both flanking sequences align on the same chromosome.",
"# translocations" : "is the number of translocation events among all misassembly events. Translocation is a misassembly where the flanking sequences align on different chromosomes.",
"# interspecies translocations" : "is the number of interspecies translocation events among all misassembly events. Interspecies translocation is a misassembly where the flanking sequences align on different references (--meta only).",
"# inversions" : "is the number of inversion events among all misassembly events. Inversion is a misassembly where it is not a *relocation* and the flanking sequences align on opposite strands of the same chromosome.",
"# local misassemblies" : "is the number of local misassemblies. We define a local misassembly breakpoint as a breakpoint that satisfies these conditions:

1. Two or more distinct alignments cover the breakpoint.
2. The gap between left and right flanking sequences is less than 1 kbp.
3. The left and right flanking sequences both are on the same strand of the same chromosome of the reference genome.
",
"# scaffold gap size misassemblies" : "is the number of scaffold gap size misassemblies. We define scaffold gap size misassembly as a breakpoint where the flanking sequences combined in scaffold on the wrong distance. These misassemblies are not included in the total number of misassemblies. ",
"# possibly misassembled contigs": "is the number of contigs that contain large unaligned fragment (default min length is 500 bp) and thus could possibly contain interspecies translocation with unknown reference.",
"# possible misassemblies" : "is the number of putative interspecies translocations in possibly misassembled contigs if each large unaligned fragment is supposed to be a fragment of unknown reference.",
"# structural variations" : "is the number of misassemblies matched with structural variations.",
"# unaligned mis. contigs" : "is the number of contigs that have the number of unaligned bases more than 50% of contig length and a misassembly event in their aligned fragment. Note that such misassemblies are not counted in *# misassemblies* and other *misassemblies* statistics.",
"# fully unaligned contigs" : "is the number of contigs that have no alignment to the reference sequence.",
"Fully unaligned length" : "is the total number of bases contained in all fully unaligned contigs.",
"# partially unaligned contigs" : "is the number of contigs that have at least one alignment to the reference sequence but also have at least one unaligned fragment of length ≥ *unaligned-part-size threshold*.",
"Partially unaligned length" : "is the total number of unaligned bases in all partially unaligned contigs.",
"# ambiguous contigs" : "is the number of contigs that have reference alignments of equal quality in multiple locations on the reference.",
"Ambiguous contigs length" : "is the total number of bases contained in all ambiguous contigs.",
"Genome fraction (%)" : "is the total number of aligned bases in the reference, divided by the genome size. A base in the reference genome is counted as aligned if there is at least one contig with at least one alignment to this base. Contigs from repeat regions may map to multiple places, and thus may be counted multiple times in this quantity.",
"GC (%)" : "is the total number of G and C nucleotides in the assembly, divided by the total length of the assembly.",
"Reference GC (%)" : "is the total number of G and C nucleotides in the reference, divided by the total length of the reference.",
"# mismatches per 100 kbp" : "is the average number of mismatches per 100000 aligned bases.",
"# mismatches" : "is the number of mismatches in all aligned bases.",
"# indels per 100 kbp" : "is the average number of indels per 100000 aligned bases.",
"# indels" : "is the number of indels in all aligned bases",
"# indels (<= 5 bp)" : "is the number of indels of length less than or equal to 5 bp",
"# indels (> 5 bp)" : "is the number of indels of length greater than 5 bp",
"Indels length" : "is the number of total bases contained in all indels",
"# genes" : "is the number of genes in the assembly (complete and partial), based on a user-provided annotated list of gene positions in the reference genome. A gene counts as 'partially covered' if the assembly contains at least 100 bp of this gene but not the whole gene.",
"# operons" : "is the number of operons in the assembly (complete and partial), based on a user-provided annotated list of operon positions in the reference genome. An operon counts as 'partially covered' if the assembly contains at least 100 bp of this operon but not the whole operon.",
"# predicted genes (unique)" : "is the number of unique genes in the assembly found by a gene prediction tool.",
"# predicted genes (>= 0 bp)" : "is the number of found genes having length greater than or equal to 0 bp.",
"Cumulative length" : "plot shows the growth of assembly contig lengths. On the x-axis, contigs are ordered from largest (contig #1) to smallest. The y-axis gives the size of the x largest contigs in the assembly.",
"Nx" : "plot shows the Nx metric value as x varies from 0 to 100. Nx is the minimum contig length **y** such that using contigs of length at least **y** accounts for at least x% of the total assembly length.",
"NGx" : "plot shows the NGx metric value as x varies from 0 to 100. NGx is the minimum contig length **y** such that using contigs of length at least **y** accounts for at least x% of the bases of the reference genome. This metric is computed only if a reference genome is provided.",
"NAx" : "plot shows the NAx metric value as x varies from 0 to 100. NAx is computed similarly to Nx, but based on lengths of aligned blocks instead of contig lengths. Contigs are broken into aligned blocks at misassembly breakpoints. NAx is the minimum block length **y** such that using blocks of length at least **y** accounts for at least x% of the bases of the assembly. This metric is computed only if a reference genome is provided.",
"NGAx" : "plot shows the NGAx metric value as x varies from 0 to 100.NGAx is computed similarly to NGx, but based on lengths of aligned blocks instead of contig lengths. Contigs are broken at misassembly breakpoints. NGAx is the minimum block length **y** such that using blocks of length at least **y** accounts for at least x% of the bases of the reference genome. This metric is computed only if a reference genome is provided.",
"GC content" : "plot shows the distribution of GC percentage among the contigs, i.e., the total number of bases in contigs with such GC content. Typically, the distribution is approximately Gaussian. However, for some genomes it is not Gaussian. For assembly projects with contaminants, the GC distribution of the contaminants often differs from the reference genome and may give a superposition of multiple curves with different peaks.",
"Duplication ratio" : "is the total number of aligned bases in the assembly (i.e. *Total length* - *Fully unaligned length* - *Partially unaligned length*), divided by the total number of aligned bases in the reference (see the **Genome fraction (%)** metric). If the assembly contains many contigs that cover the same regions of the reference, its *Duplication ratio* may be much larger than 1. This may occur due to overestimating repeat multiplicities and due to small overlaps between contigs, among other reasons.",
"Largest alignment" : "is the length of the largest continuous alignment in the assembly. This metric is always equal to the *Largest contig* metric but it can be smaller if the largest contig of the assembly contains a misassembly event.",
"Total aligned length" : "is the total number of aligned bases in the assembly.",
"Avg contig read support" : "is the average coverage of contigs that have large unique alignments to the reference. Read coverage is extracted from contig names (SPAdes/Velvet naming scheme only).",
"# N's" : "is the total number of uncalled bases (N's) in the assembly.",
"# N's per 100 kbp" : "is the average number of uncalled bases (N's) per 100000 assembly bases.",
"# similar correct contigs" : "is the number of correct contigs similar among > 50% assemblies (see Icarus for visualization).",
"# similar misassembled blocks" : "is the number of misassembled blocks similar among > 50% assemblies (see Icarus for visualization)."
}
