## Supplementary material for "Choice of assembly software has a critical impact on virome characterisation"

### QUAST

**Quality Assessment Tool for Genome Assemblies**
by CAB

Loading...

Aligned to
""

Combined reference
Estimated reference size:
 bp
|
 bp
|
 references
|
 fragments
|
 % G+C
|
 chromosomes
  
 reads
|
 mapped
|
 properly paired
  
 genes
|
 operons

Unfortunately, JavaScript in your
browser is disabled or is not supported.
We need JavaScript to build report and plots.

Worst
Median
Best

Show heatmap

|  |
| --- |
| Contigs are ordered from largest (contig #1) to smallest.  Contigs are broken into nonoverlapping 100 bp windows. Plot shows numbers of windows for each GC percentage. |

[{"assembliesWithNs":["ABySS\_63","ABySS\_127","SOAPdenovo2","SPAdes\_sc","Geneious"],"minContig":500,"report":[["Genome statistics",[{"values":["65.446","46.446","99.706","99.740","99.357","82.414","98.194","99.684","30.720","99.649","99.730","99.738","99.730","82.414","99.740","99.740"],"quality":"More is better","isMain":true,"metricName":"Genome fraction (%)"},{"values":["1.002","1.015","1.021","1.002","1.002","1.002","1.046","1.046","0.997","1.003","1.001","1.001","1.001","1.002","1.006","1.225"],"quality":"Less is better","isMain":true,"metricName":"Duplication ratio"},{"values":[76649,76666,129415,129514,129556,129477,128822,129072,73921,128853,129470,129470,129470,129477,129712,128822],"quality":"More is better","isMain":true,"metricName":"Largest alignment"},{"values":[400020,287650,612755,609963,608100,503771,621785,636072,186293,609185,609469,609484,609459,503771,612080,745091],"quality":"More is better","isMain":true,"metricName":"Total aligned length"},{"values":[null,null,null,71548,71588,36833,null,71664,null,70642,70598,null,70598,36833,54016,null],"quality":"More is better","isMain":false,"metricName":"NA50"},{"values":[null,null,null,39290,38545,23050,null,38215,null,39268,38521,null,38521,23050,39191,null],"quality":"More is better","isMain":false,"metricName":"NA75"},{"values":[null,null,null,4,4,4,null,4,null,4,4,null,4,4,4,null],"quality":"Less is better","isMain":false,"metricName":"LA50"},{"values":[null,null,null,7,7,8,null,7,null,7,7,null,7,8,7,null],"quality":"Less is better","isMain":false,"metricName":"LA75"}]],["Misassemblies",[{"values":[0,2,0,0,0,0,1,6,0,0,0,0,0,0,2,11],"quality":"Less is better","isMain":true,"metricName":"# misassemblies"},{"values":[0,2,0,0,0,0,1,6,0,0,0,0,0,0,0,9],"quality":"Less is better","isMain":false,"metricName":" # relocations"},{"values":[0,0,0,0,0,0,0,0,0,0,0,0,0,0,0,0],"quality":"Less is better","isMain":false,"metricName":" # translocations"},{"values":[0,0,0,0,0,0,0,0,0,0,0,0,0,0,0,0],"quality":"Less is better","isMain":false,"metricName":" # inversions"},{"values":[0,0,0,0,0,0,0,0,0,0,0,0,0,0,2,2],"quality":"Less is better","isMain":false,"metricName":" # interspecies translocations"},{"values":[0,1,0,0,0,0,1,4,0,0,0,0,0,0,1,9],"quality":"Less is better","isMain":false,"metricName":"# misassembled contigs"},{"values":[0,10015,0,0,0,0,521,127096,0,0,0,0,0,0,148165,647195],"quality":"Less is better","isMain":true,"metricName":"Misassembled contigs length"},{"values":[0,0,0,0,0,0,0,0,3,0,0,0,0,0,0,0],"quality":"Less is better","isMain":false,"metricName":"# possibly misassembled contigs"},{"values":[0,0,0,0,0,0,0,0,4,0,0,0,0,0,0,0],"quality":"Less is better","isMain":false,"metricName":" # possible misassemblies"},{"values":[1,0,3,10,8,3,7,4,5,3,3,3,3,3,7,4],"quality":"Less is better","isMain":false,"metricName":"# local misassemblies"},{"values":[0,0,1,0,0,0,2,0,15,0,0,0,0,0,0,0],"quality":"Less is better","isMain":false,"metricName":"# unaligned mis. contigs"}]],["Unaligned",[{"values":[60,132,450,0,0,0,94,0,2,0,0,593,0,0,0,14],"quality":"Less is better","isMain":false,"metricName":"# fully unaligned contigs"},{"values":[4534725,4510461,4733009,0,0,0,4546615,0,4899,0,0,4858632,0,0,0,3487048],"quality":"Less is better","isMain":false,"metricName":"Fully unaligned length"},{"values":[0,0,24,0,0,0,3,0,22,0,0,0,0,0,0,2],"quality":"Less is better","isMain":false,"metricName":"# partially unaligned contigs"},{"values":[0,0,15071,0,0,0,1758,0,320088,0,0,0,0,0,0,2489],"quality":"Less is better","isMain":false,"metricName":"Partially unaligned length"}]],["Mismatches",[{"values":[10,10,29,27,21,19,35,28,41,27,27,27,28,19,63,52],"quality":"Less is better","isMain":false,"metricName":"# mismatches"},{"values":[23,11,20,18,19,32,21,23,158,22,26,25,26,32,19,21],"quality":"Less is better","isMain":false,"metricName":"# indels"},{"values":[498,66,43,18,83,94,190,291,2821,419,330,262,330,94,19,156],"quality":"Less is better","isMain":false,"metricName":"Indels length"},{"values":["2.50","3.53","4.76","4.43","3.46","3.78","5.84","4.60","21.86","4.44","4.43","4.43","4.60","3.78","10.35","8.54"],"quality":"Less is better","isMain":true,"metricName":"# mismatches per 100 kbp"},{"values":["5.76","3.88","3.29","2.96","3.13","6.36","3.50","3.78","84.24","3.62","4.27","4.11","4.27","6.36","3.12","3.45"],"quality":"Less is better","isMain":true,"metricName":"# indels per 100 kbp"},{"values":[13,10,19,18,18,31,18,19,23,16,19,19,19,31,19,19],"quality":"Less is better","isMain":false,"metricName":" # indels (<= 5 bp)"},{"values":[10,1,1,0,1,1,3,4,135,6,7,6,7,1,0,2],"quality":"Less is better","isMain":false,"metricName":" # indels (> 5 bp)"},{"values":[496,6556,0,0,0,0,34,0,63405,0,0,2480,0,0,0,589],"quality":"Less is better","isMain":false,"metricName":"# N's"},{"values":["10.05","136.63","0.00","0.00","0.00","0.00","0.66","0.00","12382.36","0.00","0.00","45.35","0.00","0.00","0.00","13.91"],"quality":"Less is better","isMain":true,"metricName":"# N's per 100 kbp"}]],["Statistics without reference",[{"values":[69,139,505,12,14,24,157,13,27,13,14,607,14,24,15,30],"quality":"Equal","isMain":true,"metricName":"# contigs"},{"values":[5688,3865,1705,16,314,24,4272,13,35,13,19,7281,19,24,20,42],"quality":"Equal","isMain":false,"metricName":"# contigs (>= 0 bp)"},{"values":[63,129,93,12,12,21,83,13,21,13,13,94,13,21,15,27],"quality":"Equal","isMain":false,"metricName":"# contigs (>= 1000 bp)"},{"values":[48,109,59,12,11,17,52,12,10,12,12,41,12,17,13,24],"quality":"Equal","isMain":false,"metricName":"# contigs (>= 5000 bp)"},{"values":[42,87,50,10,10,11,48,12,9,10,10,34,10,11,10,22],"quality":"Equal","isMain":false,"metricName":"# contigs (>= 10000 bp)"},{"values":[37,53,40,10,10,7,43,10,8,10,10,31,10,7,8,21],"quality":"Equal","isMain":false,"metricName":"# contigs (>= 25000 bp)"},{"values":[25,29,28,5,5,2,27,5,3,5,5,22,5,2,4,17],"quality":"Equal","isMain":false,"metricName":"# contigs (>= 50000 bp)"},{"values":[606442,316102,517832,129514,129556,129477,431858,129851,121445,129492,129470,849779,129470,129477,148165,670062],"quality":"More is better","isMain":true,"metricName":"Largest contig"},{"values":[4934995,4798321,5369684,610062,608106,503971,5175361,636894,512059,609947,609539,5468166,609539,503971,612442,4235338],"quality":"More is better","isMain":true,"metricName":"Total length"},{"values":[5485388,5402268,5805229,610651,692677,503971,6413442,636894,513400,609947,609881,7308437,609881,503971,613743,4239476],"quality":"More is better","isMain":false,"metricName":"Total length (>= 0 bp)"},{"values":[4931100,4791720,5111614,610062,606277,501343,5129676,636894,507818,609947,608783,5137988,608783,501343,612442,4233741],"quality":"More is better","isMain":true,"metricName":"Total length (>= 1000 bp)"},{"values":[4891940,4730755,5057608,610062,604134,491430,5073804,634296,482630,607900,606780,5061582,606780,491430,605319,4226067],"quality":"More is better","isMain":false,"metricName":"Total length (>= 5000 bp)"},{"values":[4842101,4567742,4998041,598389,597980,452189,5042100,634296,474818,596271,595215,5016803,595215,452189,587735,4212107],"quality":"More is better","isMain":true,"metricName":"Total length (>= 10000 bp)"},{"values":[4764925,3986544,4836905,598389,597980,375908,4968286,602931,455231,596271,595215,4965314,595215,375908,559516,4192782],"quality":"More is better","isMain":false,"metricName":"Total length (>= 25000 bp)"},{"values":[4279678,3124173,4348368,404576,404786,195020,4332172,406415,266922,402472,402229,4584455,402229,195020,404640,4037202],"quality":"More is better","isMain":true,"metricName":"Total length (>= 50000 bp)"},{"values":[191791,75742,133924,71548,71590,36833,166085,71664,68168,70642,70598,254188,70598,36833,72708,370813],"quality":"More is better","isMain":false,"metricName":"N50"},{"values":[100856,35789,71449,39290,38545,23076,82889,40498,37358,39268,38521,99830,38521,23076,39243,189216],"quality":"More is better","isMain":false,"metricName":"N75"},{"values":[7,18,10,4,4,4,10,4,3,4,4,7,4,4,3,4],"quality":"Less is better","isMain":false,"metricName":"L50"},{"values":[17,41,23,7,7,8,20,7,6,7,7,16,7,8,6,8],"quality":"Less is better","isMain":false,"metricName":"L75"}]],["Predicted genes",[]],["Similarity statistics",[{"values":[2,2,1,1,2,2,0,1,0,2,2,2,2,2,0,1],"quality":"Equal","isMain":false,"metricName":"# similar correct contigs"},{"values":[1,0,0,0,1,1,1,0,0,1,1,0,1,1,0,1],"quality":"Equal","isMain":false,"metricName":"# similar misassembled blocks"}]],["Reference statistics",[{"values":[610540,610540,610540,610540,610540,610540,610540,610540,610540,610540,610540,610540,610540,610540,610540,610540],"quality":"Equal","isMain":false,"metricName":"Reference length"},{"values":[12,12,12,12,12,12,12,12,12,12,12,12,12,12,12,12],"quality":"Equal","isMain":false,"metricName":"Reference fragments"}]]],"referenceName":"combined\_reference","date":"18 July 2018, Wednesday, 16:08:04","order":[0,1,2,3,4,5,6,7,8,9,10,11,12,13,14,15],"assembliesNames":["ABySS\_63","ABySS\_127","CLC","IDBA\_UD","MEGAHIT","MetaVelvet","MIRA","Ray\_Meta","SOAPdenovo2","SPAdes","SPAdes\_meta","SPAdes\_sc","SPAdes\_sc\_careful","Velvet","VICUNA","Geneious"]},{"assembliesWithNs":null,"minContig":500,"report":[["Genome statistics",[{"values":["99.978","100.000","99.871","100.000","100.000","100.000","100.000","96.204","100.000","99.911","99.983","99.911","100.000","100.000"],"quality":"More is better","isMain":true,"metricName":"Genome fraction (%)"},{"values":["1.000","1.002","1.000","1.001","1.002","1.084","1.014","1.002","1.001","1.000","1.001","1.000","1.005","1.023"],"quality":"Less is better","isMain":true,"metricName":"Duplication ratio"},{"values":[76649,76666,76567,76666,76807,35010,77710,73921,76666,76634,76653,76634,77070,71811],"quality":"More is better","isMain":true,"metricName":"Largest alignment"},{"values":[76649,76666,76567,76666,76807,83080,77710,73921,76666,76634,76653,76634,77070,78464],"quality":"More is better","isMain":true,"metricName":"Total aligned length"},{"values":[76649,76792,76567,76765,76807,30919,77715,77309,76719,76634,76703,76634,148165,189216],"quality":"More is better","isMain":false,"metricName":"NG50"},{"values":[76649,76792,76567,76765,76807,30919,77715,77309,76719,76634,76703,76634,148165,189216],"quality":"More is better","isMain":false,"metricName":"NG75"},{"values":[76649,76666,76567,76666,76807,30919,77710,73921,76666,76634,76653,76634,77070,null],"quality":"More is better","isMain":false,"metricName":"NA50"},{"values":[76649,76666,76567,76666,76807,30919,77710,73921,76666,76634,76653,76634,null,null],"quality":"More is better","isMain":false,"metricName":"NA75"},{"values":[76649,76666,76567,76666,76807,30919,77710,73921,76666,76634,76653,76634,77070,71811],"quality":"More is better","isMain":true,"metricName":"NGA50"},{"values":[76649,76666,76567,76666,76807,30919,77710,73921,76666,76634,76653,76634,77070,71811],"quality":"More is better","isMain":false,"metricName":"NGA75"},{"values":[1,1,1,1,1,2,1,1,1,1,1,1,1,1],"quality":"Less is better","isMain":false,"metricName":"LG50"},{"values":[1,1,1,1,1,2,1,1,1,1,1,1,1,1],"quality":"Less is better","isMain":false,"metricName":"LG75"},{"values":[1,1,1,1,1,2,1,1,1,1,1,1,1,null],"quality":"Less is better","isMain":false,"metricName":"LA50"},{"values":[1,1,1,1,1,2,1,1,1,1,1,1,null,null],"quality":"Less is better","isMain":false,"metricName":"LA75"},{"values":[1,1,1,1,1,2,1,1,1,1,1,1,1,1],"quality":"Less is better","isMain":true,"metricName":"LGA50"},{"values":[1,1,1,1,1,2,1,1,1,1,1,1,1,1],"quality":"Less is better","isMain":false,"metricName":"LGA75"}]],["Misassemblies",[{"values":[0,0,0,0,0,0,0,0,0,0,0,0,0,0],"quality":"Less is better","isMain":true,"metricName":"# misassemblies"},{"values":[0,0,0,0,0,0,0,0,0,0,0,0,0,0],"quality":"Less is better","isMain":false,"metricName":" # relocations"},{"values":[0,0,0,0,0,0,0,0,0,0,0,0,0,0],"quality":"Less is better","isMain":false,"metricName":" # translocations"},{"values":[0,0,0,0,0,0,0,0,0,0,0,0,0,0],"quality":"Less is better","isMain":false,"metricName":" # inversions"},{"values":[0,0,0,0,0,0,0,0,0,0,0,0,0,0],"quality":"Less is better","isMain":false,"metricName":"# misassembled contigs"},{"values":[0,0,0,0,0,0,0,0,0,0,0,0,0,0],"quality":"Less is better","isMain":true,"metricName":"Misassembled contigs length"},{"values":[0,0,0,0,1,0,0,2,0,0,0,0,1,0],"quality":"Less is better","isMain":false,"metricName":"# local misassemblies"},{"values":[0,0,0,0,0,0,0,0,0,0,0,0,0,1],"quality":"Less is better","isMain":false,"metricName":"# unaligned mis. contigs"}]],["Unaligned",[{"values":[0,0,0,0,0,0,0,0,0,0,0,0,0,0],"quality":"Less is better","isMain":false,"metricName":"# fully unaligned contigs"},{"values":[0,0,0,0,0,0,0,0,0,0,0,0,0,0],"quality":"Less is better","isMain":false,"metricName":"Fully unaligned length"},{"values":[0,0,0,0,0,0,0,1,0,0,0,0,1,1],"quality":"Less is better","isMain":false,"metricName":"# partially unaligned contigs"},{"values":[0,0,0,0,0,0,0,3388,0,0,0,0,71095,112146],"quality":"Less is better","isMain":false,"metricName":"Partially unaligned length"}]],["Mismatches",[{"values":[1,1,1,1,1,1,1,0,1,1,1,1,26,26],"quality":"Less is better","isMain":false,"metricName":"# mismatches"},{"values":[0,0,0,0,0,2,4,20,0,1,0,1,0,2],"quality":"Less is better","isMain":false,"metricName":"# indels"},{"values":[0,0,0,0,0,136,272,367,0,68,0,68,0,136],"quality":"Less is better","isMain":false,"metricName":"Indels length"},{"values":["1.30","1.30","1.31","1.30","1.30","1.30","1.30","0.00","1.30","1.31","1.30","1.31","33.91","33.91"],"quality":"Less is better","isMain":true,"metricName":"# mismatches per 100 kbp"},{"values":["0.00","0.00","0.00","0.00","0.00","2.61","5.22","27.12","0.00","1.31","0.00","1.31","0.00","2.61"],"quality":"Less is better","isMain":true,"metricName":"# indels per 100 kbp"},{"values":[0,0,0,0,0,0,0,4,0,0,0,0,0,0],"quality":"Less is better","isMain":false,"metricName":" # indels (<= 5 bp)"},{"values":[0,0,0,0,0,2,4,16,0,1,0,1,0,2],"quality":"Less is better","isMain":false,"metricName":" # indels (> 5 bp)"},{"values":[0,0,0,0,0,0,0,1270,0,0,0,0,0,1],"quality":"Less is better","isMain":false,"metricName":"# N's"},{"values":["0.00","0.00","0.00","0.00","0.00","0.00","0.00","1642.76","0.00","0.00","0.00","0.00","0.00","0.52"],"quality":"Less is better","isMain":true,"metricName":"# N's per 100 kbp"}]],["Statistics without reference",[{"values":[1,1,1,1,1,8,1,1,1,1,1,1,1,2],"quality":"Equal","isMain":true,"metricName":"# contigs"},{"values":[1,1,1,1,1,7,1,1,1,1,1,1,1,2],"quality":"Equal","isMain":false,"metricName":"# contigs (>= 1000 bp)"},{"values":[1,1,1,1,1,3,1,1,1,1,1,1,1,1],"quality":"Equal","isMain":false,"metricName":"# contigs (>= 5000 bp)"},{"values":[1,1,1,1,1,2,1,1,1,1,1,1,1,1],"quality":"Equal","isMain":false,"metricName":"# contigs (>= 10000 bp)"},{"values":[1,1,1,1,1,2,1,1,1,1,1,1,1,1],"quality":"Equal","isMain":false,"metricName":"# contigs (>= 25000 bp)"},{"values":[1,1,1,1,1,0,1,1,1,1,1,1,1,1],"quality":"Equal","isMain":false,"metricName":"# contigs (>= 50000 bp)"},{"values":[76649,76792,76567,76765,76807,35010,77715,77309,76719,76634,76703,76634,148165,189216],"quality":"More is better","isMain":true,"metricName":"Largest contig"},{"values":[76649,76792,76567,76765,76807,83080,77715,77309,76719,76634,76703,76634,148165,190610],"quality":"More is better","isMain":true,"metricName":"Total length"},{"values":[76649,76792,76567,76765,76807,82466,77715,77309,76719,76634,76703,76634,148165,190610],"quality":"More is better","isMain":true,"metricName":"Total length (>= 1000 bp)"},{"values":[76649,76792,76567,76765,76807,72979,77715,77309,76719,76634,76703,76634,148165,189216],"quality":"More is better","isMain":false,"metricName":"Total length (>= 5000 bp)"},{"values":[76649,76792,76567,76765,76807,65929,77715,77309,76719,76634,76703,76634,148165,189216],"quality":"More is better","isMain":true,"metricName":"Total length (>= 10000 bp)"},{"values":[76649,76792,76567,76765,76807,65929,77715,77309,76719,76634,76703,76634,148165,189216],"quality":"More is better","isMain":false,"metricName":"Total length (>= 25000 bp)"},{"values":[76649,76792,76567,76765,76807,0,77715,77309,76719,76634,76703,76634,148165,189216],"quality":"More is better","isMain":true,"metricName":"Total length (>= 50000 bp)"},{"values":[76649,76792,76567,76765,76807,30919,77715,77309,76719,76634,76703,76634,148165,189216],"quality":"More is better","isMain":false,"metricName":"N50"},{"values":[76649,76792,76567,76765,76807,30919,77715,77309,76719,76634,76703,76634,148165,189216],"quality":"More is better","isMain":false,"metricName":"N75"},{"values":[1,1,1,1,1,2,1,1,1,1,1,1,1,1],"quality":"Less is better","isMain":false,"metricName":"L50"},{"values":[1,1,1,1,1,2,1,1,1,1,1,1,1,1],"quality":"Less is better","isMain":false,"metricName":"L75"},{"values":["30.22","30.23","30.22","30.23","30.21","30.12","30.23","30.25","30.23","30.22","30.23","30.22","31.51","32.01"],"quality":"Equal","isMain":false,"metricName":"GC (%)"}]],["Predicted genes",[]],["Similarity statistics",[{"values":[0,0,0,0,0,0,0,0,0,0,0,0,0,0],"quality":"Equal","isMain":false,"metricName":"# similar correct contigs"},{"values":[0,0,0,0,0,0,0,0,0,0,0,0,0,0],"quality":"Equal","isMain":false,"metricName":"# similar misassembled blocks"}]],["Reference statistics",[{"values":[76666,76666,76666,76666,76666,76666,76666,76666,76666,76666,76666,76666,76666,76666],"quality":"Equal","isMain":false,"metricName":"Reference length"},{"values":[1,1,1,1,1,1,1,1,1,1,1,1,1,1],"quality":"Equal","isMain":false,"metricName":"Reference fragments"},{"values":["30.22","30.22","30.22","30.22","30.22","30.22","30.22","30.22","30.22","30.22","30.22","30.22","30.22","30.22"],"quality":"Equal","isMain":false,"metricName":"Reference GC (%)"}]]],"referenceName":"KC821625","date":"18 July 2018, Wednesday, 16:08:14","order":[0,1,2,3,4,5,6,7,8,9,10,11,12,13],"assembliesNames":["ABySS\_63","ABySS\_127","CLC","IDBA\_UD","MEGAHIT","MIRA","Ray\_Meta","SOAPdenovo2","SPAdes","SPAdes\_meta","SPAdes\_sc","SPAdes\_sc\_careful","VICUNA","Geneious"]},{"assembliesWithNs":null,"minContig":500,"report":[["Genome statistics",[{"values":["99.880","99.880","99.961","100.000","100.000","99.583","97.328","100.000","3.484","100.000","100.000","100.000","100.000","99.583","100.000","100.000"],"quality":"More is better","isMain":true,"metricName":"Genome fraction (%)"},{"values":["1.005","1.001","1.000","1.002","1.003","1.003","1.009","1.023","1.111","1.001","1.001","1.001","1.001","1.003","1.001","1.006"],"quality":"Less is better","isMain":true,"metricName":"Duplication ratio"},{"values":[54004,53956,53995,54115,54155,35740,48400,53943,332,54093,54042,54042,54042,35740,54016,54016],"quality":"More is better","isMain":true,"metricName":"Largest alignment"},{"values":[54004,53956,53995,54115,54155,53946,53062,55233,1881,54093,54042,54042,54042,53946,54016,54016],"quality":"More is better","isMain":true,"metricName":"Total aligned length"},{"values":[54204,54006,54016,54115,54157,35767,48400,55233,null,54093,54042,54042,54042,35767,54055,54310],"quality":"More is better","isMain":false,"metricName":"NG50"},{"values":[54204,54006,54016,54115,54157,8710,48400,55233,null,54093,54042,54042,54042,8710,54055,54310],"quality":"More is better","isMain":false,"metricName":"NG75"},{"values":[54004,53956,53995,54115,54155,35740,48400,53943,null,54093,54042,54042,54042,35740,54016,54016],"quality":"More is better","isMain":false,"metricName":"NA50"},{"values":[54004,53956,53995,54115,54155,8710,48400,53943,null,54093,54042,54042,54042,8710,54016,54016],"quality":"More is better","isMain":false,"metricName":"NA75"},{"values":[54004,53956,53995,54115,54155,35740,48400,53943,null,54093,54042,54042,54042,35740,54016,54016],"quality":"More is better","isMain":true,"metricName":"NGA50"},{"values":[54004,53956,53995,54115,54155,8710,48400,53943,null,54093,54042,54042,54042,8710,54016,54016],"quality":"More is better","isMain":false,"metricName":"NGA75"},{"values":[1,1,1,1,1,1,1,1,null,1,1,1,1,1,1,1],"quality":"Less is better","isMain":false,"metricName":"LG50"},{"values":[1,1,1,1,1,2,1,1,null,1,1,1,1,2,1,1],"quality":"Less is better","isMain":false,"metricName":"LG75"},{"values":[1,1,1,1,1,1,1,1,null,1,1,1,1,1,1,1],"quality":"Less is better","isMain":false,"metricName":"LA50"},{"values":[1,1,1,1,1,2,1,1,null,1,1,1,1,2,1,1],"quality":"Less is better","isMain":false,"metricName":"LA75"},{"values":[1,1,1,1,1,1,1,1,null,1,1,1,1,1,1,1],"quality":"Less is better","isMain":true,"metricName":"LGA50"},{"values":[1,1,1,1,1,2,1,1,null,1,1,1,1,2,1,1],"quality":"Less is better","isMain":false,"metricName":"LGA75"}]],["Misassemblies",[{"values":[0,0,0,0,0,0,0,1,0,0,0,0,0,0,0,0],"quality":"Less is better","isMain":true,"metricName":"# misassemblies"},{"values":[0,0,0,0,0,0,0,1,0,0,0,0,0,0,0,0],"quality":"Less is better","isMain":false,"metricName":" # relocations"},{"values":[0,0,0,0,0,0,0,0,0,0,0,0,0,0,0,0],"quality":"Less is better","isMain":false,"metricName":" # translocations"},{"values":[0,0,0,0,0,0,0,0,0,0,0,0,0,0,0,0],"quality":"Less is better","isMain":false,"metricName":" # inversions"},{"values":[0,0,0,0,0,0,0,1,0,0,0,0,0,0,0,0],"quality":"Less is better","isMain":false,"metricName":"# misassembled contigs"},{"values":[0,0,0,0,0,0,0,55233,0,0,0,0,0,0,0,0],"quality":"Less is better","isMain":true,"metricName":"Misassembled contigs length"},{"values":[0,0,0,1,1,0,0,0,0,0,0,0,0,0,0,0],"quality":"Less is better","isMain":false,"metricName":"# local misassemblies"},{"values":[0,0,0,0,0,0,0,0,3,0,0,0,0,0,0,0],"quality":"Less is better","isMain":false,"metricName":"# unaligned mis. contigs"}]],["Unaligned",[{"values":[0,0,0,0,0,0,0,0,0,0,0,0,0,0,0,0],"quality":"Less is better","isMain":false,"metricName":"# fully unaligned contigs"},{"values":[0,0,0,0,0,0,0,0,0,0,0,0,0,0,0,0],"quality":"Less is better","isMain":false,"metricName":"Fully unaligned length"},{"values":[0,0,0,0,0,0,0,0,7,0,0,0,0,0,0,0],"quality":"Less is better","isMain":false,"metricName":"# partially unaligned contigs"},{"values":[0,0,0,0,0,0,0,0,8203,0,0,0,0,0,0,0],"quality":"Less is better","isMain":false,"metricName":"Partially unaligned length"}]],["Mismatches",[{"values":[0,0,0,0,0,0,2,0,1,0,0,0,1,0,0,0],"quality":"Less is better","isMain":false,"metricName":"# mismatches"},{"values":[7,4,4,3,4,8,6,4,1,5,6,6,6,8,4,4],"quality":"Less is better","isMain":false,"metricName":"# indels"},{"values":[293,59,4,3,4,8,41,4,1,98,66,66,66,8,4,4],"quality":"Less is better","isMain":false,"metricName":"Indels length"},{"values":["0.00","0.00","0.00","0.00","0.00","0.00","3.80","0.00","53.13","0.00","0.00","0.00","1.85","0.00","0.00","0.00"],"quality":"Less is better","isMain":true,"metricName":"# mismatches per 100 kbp"},{"values":["12.98","7.41","7.41","5.55","7.41","14.87","11.41","7.41","53.13","9.26","11.11","11.11","11.11","14.87","7.41","7.41"],"quality":"Less is better","isMain":true,"metricName":"# indels per 100 kbp"},{"values":[1,3,4,3,4,8,5,4,1,3,4,4,4,8,4,4],"quality":"Less is better","isMain":false,"metricName":" # indels (<= 5 bp)"},{"values":[6,1,0,0,0,0,1,0,0,2,2,2,2,0,0,0],"quality":"Less is better","isMain":false,"metricName":" # indels (> 5 bp)"},{"values":[250,50,0,0,0,0,1,0,4578,0,0,0,0,0,0,43],"quality":"Less is better","isMain":false,"metricName":"# N's"},{"values":["461.22","92.58","0.00","0.00","0.00","0.00","1.88","0.00","44472.51","0.00","0.00","0.00","0.00","0.00","0.00","79.18"],"quality":"Less is better","isMain":true,"metricName":"# N's per 100 kbp"}]],["Statistics without reference",[{"values":[1,1,1,1,1,4,5,1,8,1,1,1,1,4,1,1],"quality":"Equal","isMain":true,"metricName":"# contigs"},{"values":[1,1,1,1,1,4,3,1,4,1,1,1,1,4,1,1],"quality":"Equal","isMain":false,"metricName":"# contigs (>= 1000 bp)"},{"values":[1,1,1,1,1,3,1,1,0,1,1,1,1,3,1,1],"quality":"Equal","isMain":false,"metricName":"# contigs (>= 5000 bp)"},{"values":[1,1,1,1,1,1,1,1,0,1,1,1,1,1,1,1],"quality":"Equal","isMain":false,"metricName":"# contigs (>= 10000 bp)"},{"values":[1,1,1,1,1,1,1,1,0,1,1,1,1,1,1,1],"quality":"Equal","isMain":false,"metricName":"# contigs (>= 25000 bp)"},{"values":[1,1,1,1,1,0,0,1,0,1,1,1,1,0,1,1],"quality":"Equal","isMain":false,"metricName":"# contigs (>= 50000 bp)"},{"values":[54204,54006,54016,54115,54157,35767,48400,55233,2281,54093,54042,54042,54042,35767,54055,54310],"quality":"More is better","isMain":true,"metricName":"Largest contig"},{"values":[54204,54006,54016,54115,54157,53973,53062,55233,10294,54093,54042,54042,54042,53973,54055,54310],"quality":"More is better","isMain":true,"metricName":"Total length"},{"values":[54204,54006,54016,54115,54157,53973,51842,55233,7403,54093,54042,54042,54042,53973,54055,54310],"quality":"More is better","isMain":true,"metricName":"Total length (>= 1000 bp)"},{"values":[54204,54006,54016,54115,54157,49796,48400,55233,0,54093,54042,54042,54042,49796,54055,54310],"quality":"More is better","isMain":false,"metricName":"Total length (>= 5000 bp)"},{"values":[54204,54006,54016,54115,54157,35767,48400,55233,0,54093,54042,54042,54042,35767,54055,54310],"quality":"More is better","isMain":true,"metricName":"Total length (>= 10000 bp)"},{"values":[54204,54006,54016,54115,54157,35767,48400,55233,0,54093,54042,54042,54042,35767,54055,54310],"quality":"More is better","isMain":false,"metricName":"Total length (>= 25000 bp)"},{"values":[54204,54006,54016,54115,54157,0,0,55233,0,54093,54042,54042,54042,0,54055,54310],"quality":"More is better","isMain":true,"metricName":"Total length (>= 50000 bp)"},{"values":[54204,54006,54016,54115,54157,35767,48400,55233,1923,54093,54042,54042,54042,35767,54055,54310],"quality":"More is better","isMain":false,"metricName":"N50"},{"values":[54204,54006,54016,54115,54157,8710,48400,55233,873,54093,54042,54042,54042,8710,54055,54310],"quality":"More is better","isMain":false,"metricName":"N75"},{"values":[1,1,1,1,1,1,1,1,3,1,1,1,1,1,1,1],"quality":"Less is better","isMain":false,"metricName":"L50"},{"values":[1,1,1,1,1,2,1,1,5,1,1,1,1,2,1,1],"quality":"Less is better","isMain":false,"metricName":"L75"},{"values":["33.47","33.48","33.47","33.48","33.46","33.44","33.53","33.41","32.89","33.46","33.47","33.47","33.47","33.44","33.46","33.47"],"quality":"Equal","isMain":false,"metricName":"GC (%)"}]],["Predicted genes",[]],["Similarity statistics",[{"values":[0,0,0,0,0,0,0,0,0,0,0,0,0,0,0,0],"quality":"Equal","isMain":false,"metricName":"# similar correct contigs"},{"values":[0,0,0,0,0,0,0,0,0,0,0,0,0,0,0,0],"quality":"Equal","isMain":false,"metricName":"# similar misassembled blocks"}]],["Reference statistics",[{"values":[54012,54012,54012,54012,54012,54012,54012,54012,54012,54012,54012,54012,54012,54012,54012,54012],"quality":"Equal","isMain":false,"metricName":"Reference length"},{"values":[1,1,1,1,1,1,1,1,1,1,1,1,1,1,1,1],"quality":"Equal","isMain":false,"metricName":"Reference fragments"},{"values":["33.48","33.48","33.48","33.48","33.48","33.48","33.48","33.48","33.48","33.48","33.48","33.48","33.48","33.48","33.48","33.48"],"quality":"Equal","isMain":false,"metricName":"Reference GC (%)"}]]],"referenceName":"KC821629","date":"18 July 2018, Wednesday, 16:08:25","order":[0,1,2,3,4,5,6,7,8,9,10,11,12,13,14,15],"assembliesNames":["ABySS\_63","ABySS\_127","CLC","IDBA\_UD","MEGAHIT","MetaVelvet","MIRA","Ray\_Meta","SOAPdenovo2","SPAdes","SPAdes\_meta","SPAdes\_sc","SPAdes\_sc\_careful","Velvet","VICUNA","Geneious"]},{"assembliesWithNs":null,"minContig":500,"report":[["Genome statistics",[{"values":["98.812","98.812","98.812","98.812","98.812","98.812","7.087","98.812","98.812","98.812","98.812","98.812","98.812","98.812"],"quality":"More is better","isMain":true,"metricName":"Genome fraction (%)"},{"values":["1.000","1.003","1.004","1.002","1.004","1.021","0.979","1.002","1.002","1.002","1.002","1.002","1.008","1.089"],"quality":"Less is better","isMain":true,"metricName":"Duplication ratio"},{"values":[36771,36870,36910,36833,36911,37544,710,36848,36826,36826,36826,36833,37057,35127],"quality":"More is better","isMain":true,"metricName":"Largest alignment"},{"values":[36771,36870,36910,36833,36911,37544,2581,36848,36826,36826,36826,36833,37057,40038],"quality":"More is better","isMain":true,"metricName":"Total aligned length"},{"values":[36771,36870,36912,36833,36911,37544,34341,36848,36826,36826,36826,36833,37057,40038],"quality":"More is better","isMain":false,"metricName":"NG50"},{"values":[36771,36870,36912,36833,36911,37544,34341,36848,36826,36826,36826,36833,37057,40038],"quality":"More is better","isMain":false,"metricName":"NG75"},{"values":[36771,36870,36910,36833,36911,37544,null,36848,36826,36826,36826,36833,37057,35127],"quality":"More is better","isMain":false,"metricName":"NA50"},{"values":[36771,36870,36910,36833,36911,37544,null,36848,36826,36826,36826,36833,37057,35127],"quality":"More is better","isMain":false,"metricName":"NA75"},{"values":[36771,36870,36910,36833,36911,37544,null,36848,36826,36826,36826,36833,37057,35127],"quality":"More is better","isMain":true,"metricName":"NGA50"},{"values":[36771,36870,36910,36833,36911,37544,null,36848,36826,36826,36826,36833,37057,35127],"quality":"More is better","isMain":false,"metricName":"NGA75"},{"values":[1,1,1,1,1,1,1,1,1,1,1,1,1,1],"quality":"Less is better","isMain":false,"metricName":"LG50"},{"values":[1,1,1,1,1,1,1,1,1,1,1,1,1,1],"quality":"Less is better","isMain":false,"metricName":"LG75"},{"values":[1,1,1,1,1,1,null,1,1,1,1,1,1,1],"quality":"Less is better","isMain":false,"metricName":"LA50"},{"values":[1,1,1,1,1,1,null,1,1,1,1,1,1,1],"quality":"Less is better","isMain":false,"metricName":"LA75"},{"values":[1,1,1,1,1,1,null,1,1,1,1,1,1,1],"quality":"Less is better","isMain":true,"metricName":"LGA50"},{"values":[1,1,1,1,1,1,null,1,1,1,1,1,1,1],"quality":"Less is better","isMain":false,"metricName":"LGA75"}]],["Misassemblies",[{"values":[0,0,0,0,0,0,0,0,0,0,0,0,0,1],"quality":"Less is better","isMain":true,"metricName":"# misassemblies"},{"values":[0,0,0,0,0,0,0,0,0,0,0,0,0,1],"quality":"Less is better","isMain":false,"metricName":" # relocations"},{"values":[0,0,0,0,0,0,0,0,0,0,0,0,0,0],"quality":"Less is better","isMain":false,"metricName":" # translocations"},{"values":[0,0,0,0,0,0,0,0,0,0,0,0,0,0],"quality":"Less is better","isMain":false,"metricName":" # inversions"},{"values":[0,0,0,0,0,0,0,0,0,0,0,0,0,1],"quality":"Less is better","isMain":false,"metricName":"# misassembled contigs"},{"values":[0,0,0,0,0,0,0,0,0,0,0,0,0,40038],"quality":"Less is better","isMain":true,"metricName":"Misassembled contigs length"},{"values":[1,1,1,1,1,1,0,1,1,1,1,1,1,0],"quality":"Less is better","isMain":false,"metricName":"# local misassemblies"},{"values":[0,0,0,0,0,0,1,0,0,0,0,0,0,0],"quality":"Less is better","isMain":false,"metricName":"# unaligned mis. contigs"}]],["Unaligned",[{"values":[0,0,0,0,0,0,0,0,0,0,0,0,0,0],"quality":"Less is better","isMain":false,"metricName":"# fully unaligned contigs"},{"values":[0,0,0,0,0,0,0,0,0,0,0,0,0,0],"quality":"Less is better","isMain":false,"metricName":"Fully unaligned length"},{"values":[0,0,0,0,0,0,1,0,0,0,0,0,0,0],"quality":"Less is better","isMain":false,"metricName":"# partially unaligned contigs"},{"values":[0,0,0,0,0,0,31760,0,0,0,0,0,0,0],"quality":"Less is better","isMain":false,"metricName":"Partially unaligned length"}]],["Mismatches",[{"values":[1,1,1,1,1,1,1,1,1,1,1,1,1,1],"quality":"Less is better","isMain":false,"metricName":"# mismatches"},{"values":[2,2,2,2,2,2,2,1,2,2,2,2,2,2],"quality":"Less is better","isMain":false,"metricName":"# indels"},{"values":[2,2,2,2,2,2,56,1,2,2,2,2,2,2],"quality":"Less is better","isMain":false,"metricName":"Indels length"},{"values":["2.72","2.72","2.72","2.72","2.72","2.72","37.92","2.72","2.72","2.72","2.72","2.72","2.72","2.72"],"quality":"Less is better","isMain":true,"metricName":"# mismatches per 100 kbp"},{"values":["5.44","5.44","5.44","5.44","5.44","5.44","75.84","2.72","5.44","5.44","5.44","5.44","5.44","5.44"],"quality":"Less is better","isMain":true,"metricName":"# indels per 100 kbp"},{"values":[2,2,2,2,2,2,0,1,2,2,2,2,2,2],"quality":"Less is better","isMain":false,"metricName":" # indels (<= 5 bp)"},{"values":[0,0,0,0,0,0,2,0,0,0,0,0,0,0],"quality":"Less is better","isMain":false,"metricName":" # indels (> 5 bp)"},{"values":[0,0,0,0,0,0,4722,0,0,0,0,0,0,0],"quality":"Less is better","isMain":false,"metricName":"# N's"},{"values":["0.00","0.00","0.00","0.00","0.00","0.00","13750.33","0.00","0.00","0.00","0.00","0.00","0.00","0.00"],"quality":"Less is better","isMain":true,"metricName":"# N's per 100 kbp"}]],["Statistics without reference",[{"values":[1,1,1,1,1,1,1,1,1,1,1,1,1,1],"quality":"Equal","isMain":true,"metricName":"# contigs"},{"values":[1,1,1,1,1,1,1,1,1,1,1,1,1,1],"quality":"Equal","isMain":false,"metricName":"# contigs (>= 1000 bp)"},{"values":[1,1,1,1,1,1,1,1,1,1,1,1,1,1],"quality":"Equal","isMain":false,"metricName":"# contigs (>= 5000 bp)"},{"values":[1,1,1,1,1,1,1,1,1,1,1,1,1,1],"quality":"Equal","isMain":false,"metricName":"# contigs (>= 10000 bp)"},{"values":[1,1,1,1,1,1,1,1,1,1,1,1,1,1],"quality":"Equal","isMain":false,"metricName":"# contigs (>= 25000 bp)"},{"values":[0,0,0,0,0,0,0,0,0,0,0,0,0,0],"quality":"Equal","isMain":false,"metricName":"# contigs (>= 50000 bp)"},{"values":[36771,36870,36912,36833,36911,37544,34341,36848,36826,36826,36826,36833,37057,40038],"quality":"More is better","isMain":true,"metricName":"Largest contig"},{"values":[36771,36870,36912,36833,36911,37544,34341,36848,36826,36826,36826,36833,37057,40038],"quality":"More is better","isMain":true,"metricName":"Total length"},{"values":[36771,36870,36912,36833,36911,37544,34341,36848,36826,36826,36826,36833,37057,40038],"quality":"More is better","isMain":true,"metricName":"Total length (>= 1000 bp)"},{"values":[36771,36870,36912,36833,36911,37544,34341,36848,36826,36826,36826,36833,37057,40038],"quality":"More is better","isMain":false,"metricName":"Total length (>= 5000 bp)"},{"values":[36771,36870,36912,36833,36911,37544,34341,36848,36826,36826,36826,36833,37057,40038],"quality":"More is better","isMain":true,"metricName":"Total length (>= 10000 bp)"},{"values":[36771,36870,36912,36833,36911,37544,34341,36848,36826,36826,36826,36833,37057,40038],"quality":"More is better","isMain":false,"metricName":"Total length (>= 25000 bp)"},{"values":[0,0,0,0,0,0,0,0,0,0,0,0,0,0],"quality":"More is better","isMain":true,"metricName":"Total length (>= 50000 bp)"},{"values":[36771,36870,36912,36833,36911,37544,34341,36848,36826,36826,36826,36833,37057,40038],"quality":"More is better","isMain":false,"metricName":"N50"},{"values":[36771,36870,36912,36833,36911,37544,34341,36848,36826,36826,36826,36833,37057,40038],"quality":"More is better","isMain":false,"metricName":"N75"},{"values":[1,1,1,1,1,1,1,1,1,1,1,1,1,1],"quality":"Less is better","isMain":false,"metricName":"L50"},{"values":[1,1,1,1,1,1,1,1,1,1,1,1,1,1],"quality":"Less is better","isMain":false,"metricName":"L75"},{"values":["40.51","40.51","40.49","40.51","40.51","40.54","40.45","40.50","40.51","40.52","40.51","40.51","40.50","40.65"],"quality":"Equal","isMain":false,"metricName":"GC (%)"}]],["Predicted genes",[]],["Similarity statistics",[{"values":[0,0,0,0,0,0,0,0,0,0,0,0,0,0],"quality":"Equal","isMain":false,"metricName":"# similar correct contigs"},{"values":[0,0,0,0,0,0,0,0,0,0,0,0,0,0],"quality":"Equal","isMain":false,"metricName":"# similar misassembled blocks"}]],["Reference statistics",[{"values":[37211,37211,37211,37211,37211,37211,37211,37211,37211,37211,37211,37211,37211,37211],"quality":"Equal","isMain":false,"metricName":"Reference length"},{"values":[1,1,1,1,1,1,1,1,1,1,1,1,1,1],"quality":"Equal","isMain":false,"metricName":"Reference fragments"},{"values":["40.53","40.53","40.53","40.53","40.53","40.53","40.53","40.53","40.53","40.53","40.53","40.53","40.53","40.53"],"quality":"Equal","isMain":false,"metricName":"Reference GC (%)"}]]],"referenceName":"KF302033","date":"18 July 2018, Wednesday, 16:08:34","order":[0,1,2,3,4,5,6,7,8,9,10,11,12,13],"assembliesNames":["CLC","IDBA\_UD","MEGAHIT","MetaVelvet","MIRA","Ray\_Meta","SOAPdenovo2","SPAdes","SPAdes\_meta","SPAdes\_sc","SPAdes\_sc\_careful","Velvet","VICUNA","Geneious"]},{"assembliesWithNs":null,"minContig":500,"report":[["Genome statistics",[{"values":["99.981","99.981","99.981","99.981","99.523","99.716","8.668","99.547","99.981","99.981","99.981","99.981","99.981","99.981"],"quality":"More is better","isMain":true,"metricName":"Genome fraction (%)"},{"values":["1.000","1.001","1.001","1.000","1.006","1.006","0.989","1.005","1.000","1.000","1.000","1.000","1.002","1.021"],"quality":"Less is better","isMain":true,"metricName":"Duplication ratio"},{"values":[129415,129514,129556,129477,128822,129072,872,128853,129470,129470,129470,129477,129712,128822],"quality":"More is better","isMain":true,"metricName":"Largest alignment"},{"values":[129415,129514,129556,129477,128822,129072,11123,128853,129470,129470,129470,129477,129712,132154],"quality":"More is better","isMain":true,"metricName":"Total aligned length"},{"values":[129415,129514,129556,129477,129589,129851,121445,129492,129470,129470,129470,129477,129712,132154],"quality":"More is better","isMain":false,"metricName":"NG50"},{"values":[129415,129514,129556,129477,129589,129851,121445,129492,129470,129470,129470,129477,129712,132154],"quality":"More is better","isMain":false,"metricName":"NG75"},{"values":[129415,129514,129556,129477,128822,129072,null,128853,129470,129470,129470,129477,129712,128822],"quality":"More is better","isMain":false,"metricName":"NA50"},{"values":[129415,129514,129556,129477,128822,129072,null,128853,129470,129470,129470,129477,129712,128822],"quality":"More is better","isMain":false,"metricName":"NA75"},{"values":[129415,129514,129556,129477,128822,129072,null,128853,129470,129470,129470,129477,129712,128822],"quality":"More is better","isMain":true,"metricName":"NGA50"},{"values":[129415,129514,129556,129477,128822,129072,null,128853,129470,129470,129470,129477,129712,128822],"quality":"More is better","isMain":false,"metricName":"NGA75"},{"values":[1,1,1,1,1,1,1,1,1,1,1,1,1,1],"quality":"Less is better","isMain":false,"metricName":"LG50"},{"values":[1,1,1,1,1,1,1,1,1,1,1,1,1,1],"quality":"Less is better","isMain":false,"metricName":"LG75"},{"values":[1,1,1,1,1,1,null,1,1,1,1,1,1,1],"quality":"Less is better","isMain":false,"metricName":"LA50"},{"values":[1,1,1,1,1,1,null,1,1,1,1,1,1,1],"quality":"Less is better","isMain":false,"metricName":"LA75"},{"values":[1,1,1,1,1,1,null,1,1,1,1,1,1,1],"quality":"Less is better","isMain":true,"metricName":"LGA50"},{"values":[1,1,1,1,1,1,null,1,1,1,1,1,1,1],"quality":"Less is better","isMain":false,"metricName":"LGA75"}]],["Misassemblies",[{"values":[0,0,0,0,0,0,0,0,0,0,0,0,0,1],"quality":"Less is better","isMain":true,"metricName":"# misassemblies"},{"values":[0,0,0,0,0,0,0,0,0,0,0,0,0,1],"quality":"Less is better","isMain":false,"metricName":" # relocations"},{"values":[0,0,0,0,0,0,0,0,0,0,0,0,0,0],"quality":"Less is better","isMain":false,"metricName":" # translocations"},{"values":[0,0,0,0,0,0,0,0,0,0,0,0,0,0],"quality":"Less is better","isMain":false,"metricName":" # inversions"},{"values":[0,0,0,0,0,0,0,0,0,0,0,0,0,1],"quality":"Less is better","isMain":false,"metricName":"# misassembled contigs"},{"values":[0,0,0,0,0,0,0,0,0,0,0,0,0,132154],"quality":"Less is better","isMain":true,"metricName":"Misassembled contigs length"},{"values":[0,1,1,0,0,0,0,0,0,0,0,0,1,0],"quality":"Less is better","isMain":false,"metricName":"# local misassemblies"},{"values":[0,0,0,0,0,0,1,0,0,0,0,0,0,0],"quality":"Less is better","isMain":false,"metricName":"# unaligned mis. contigs"}]],["Unaligned",[{"values":[0,0,0,0,0,0,0,0,0,0,0,0,0,0],"quality":"Less is better","isMain":false,"metricName":"# fully unaligned contigs"},{"values":[0,0,0,0,0,0,0,0,0,0,0,0,0,0],"quality":"Less is better","isMain":false,"metricName":"Fully unaligned length"},{"values":[0,0,0,0,0,0,1,0,0,0,0,0,0,0],"quality":"Less is better","isMain":false,"metricName":"# partially unaligned contigs"},{"values":[0,0,0,0,0,0,110346,0,0,0,0,0,0,0],"quality":"Less is better","isMain":false,"metricName":"Partially unaligned length"}]],["Mismatches",[{"values":[0,0,0,0,0,0,1,0,0,0,0,0,0,0],"quality":"Less is better","isMain":false,"metricName":"# mismatches"},{"values":[1,0,0,1,0,0,8,0,1,1,1,1,0,0],"quality":"Less is better","isMain":false,"metricName":"# indels"},{"values":[24,0,0,62,0,0,141,0,55,55,55,62,0,0],"quality":"Less is better","isMain":false,"metricName":"Indels length"},{"values":["0.00","0.00","0.00","0.00","0.00","0.00","8.91","0.00","0.00","0.00","0.00","0.00","0.00","0.00"],"quality":"Less is better","isMain":true,"metricName":"# mismatches per 100 kbp"},{"values":["0.77","0.00","0.00","0.77","0.00","0.00","71.30","0.00","0.77","0.77","0.77","0.77","0.00","0.00"],"quality":"Less is better","isMain":true,"metricName":"# indels per 100 kbp"},{"values":[0,0,0,0,0,0,1,0,0,0,0,0,0,0],"quality":"Less is better","isMain":false,"metricName":" # indels (<= 5 bp)"},{"values":[1,0,0,1,0,0,7,0,1,1,1,1,0,0],"quality":"Less is better","isMain":false,"metricName":" # indels (> 5 bp)"},{"values":[0,0,0,0,0,0,22057,0,0,0,0,0,0,0],"quality":"Less is better","isMain":false,"metricName":"# N's"},{"values":["0.00","0.00","0.00","0.00","0.00","0.00","18162.13","0.00","0.00","0.00","0.00","0.00","0.00","0.00"],"quality":"Less is better","isMain":true,"metricName":"# N's per 100 kbp"}]],["Statistics without reference",[{"values":[1,1,1,1,1,1,1,1,1,1,1,1,1,1],"quality":"Equal","isMain":true,"metricName":"# contigs"},{"values":[1,1,1,1,1,1,1,1,1,1,1,1,1,1],"quality":"Equal","isMain":false,"metricName":"# contigs (>= 1000 bp)"},{"values":[1,1,1,1,1,1,1,1,1,1,1,1,1,1],"quality":"Equal","isMain":false,"metricName":"# contigs (>= 5000 bp)"},{"values":[1,1,1,1,1,1,1,1,1,1,1,1,1,1],"quality":"Equal","isMain":false,"metricName":"# contigs (>= 10000 bp)"},{"values":[1,1,1,1,1,1,1,1,1,1,1,1,1,1],"quality":"Equal","isMain":false,"metricName":"# contigs (>= 25000 bp)"},{"values":[1,1,1,1,1,1,1,1,1,1,1,1,1,1],"quality":"Equal","isMain":false,"metricName":"# contigs (>= 50000 bp)"},{"values":[129415,129514,129556,129477,129589,129851,121445,129492,129470,129470,129470,129477,129712,132154],"quality":"More is better","isMain":true,"metricName":"Largest contig"},{"values":[129415,129514,129556,129477,129589,129851,121445,129492,129470,129470,129470,129477,129712,132154],"quality":"More is better","isMain":true,"metricName":"Total length"},{"values":[129415,129514,129556,129477,129589,129851,121445,129492,129470,129470,129470,129477,129712,132154],"quality":"More is better","isMain":true,"metricName":"Total length (>= 1000 bp)"},{"values":[129415,129514,129556,129477,129589,129851,121445,129492,129470,129470,129470,129477,129712,132154],"quality":"More is better","isMain":false,"metricName":"Total length (>= 5000 bp)"},{"values":[129415,129514,129556,129477,129589,129851,121445,129492,129470,129470,129470,129477,129712,132154],"quality":"More is better","isMain":true,"metricName":"Total length (>= 10000 bp)"},{"values":[129415,129514,129556,129477,129589,129851,121445,129492,129470,129470,129470,129477,129712,132154],"quality":"More is better","isMain":false,"metricName":"Total length (>= 25000 bp)"},{"values":[129415,129514,129556,129477,129589,129851,121445,129492,129470,129470,129470,129477,129712,132154],"quality":"More is better","isMain":true,"metricName":"Total length (>= 50000 bp)"},{"values":[129415,129514,129556,129477,129589,129851,121445,129492,129470,129470,129470,129477,129712,132154],"quality":"More is better","isMain":false,"metricName":"N50"},{"values":[129415,129514,129556,129477,129589,129851,121445,129492,129470,129470,129470,129477,129712,132154],"quality":"More is better","isMain":false,"metricName":"N75"},{"values":[1,1,1,1,1,1,1,1,1,1,1,1,1,1],"quality":"Less is better","isMain":false,"metricName":"L50"},{"values":[1,1,1,1,1,1,1,1,1,1,1,1,1,1],"quality":"Less is better","isMain":false,"metricName":"L75"},{"values":["35.73","35.73","35.73","35.73","35.73","35.72","35.58","35.73","35.73","35.73","35.73","35.73","35.74","35.70"],"quality":"Equal","isMain":false,"metricName":"GC (%)"}]],["Predicted genes",[]],["Similarity statistics",[{"values":[0,0,0,0,0,0,0,0,0,0,0,0,0,0],"quality":"Equal","isMain":false,"metricName":"# similar correct contigs"},{"values":[0,0,0,0,0,0,0,0,0,0,0,0,0,0],"quality":"Equal","isMain":false,"metricName":"# similar misassembled blocks"}]],["Reference statistics",[{"values":[129439,129439,129439,129439,129439,129439,129439,129439,129439,129439,129439,129439,129439,129439],"quality":"Equal","isMain":false,"metricName":"Reference length"},{"values":[1,1,1,1,1,1,1,1,1,1,1,1,1,1],"quality":"Equal","isMain":false,"metricName":"Reference fragments"},{"values":["35.73","35.73","35.73","35.73","35.73","35.73","35.73","35.73","35.73","35.73","35.73","35.73","35.73","35.73"],"quality":"Equal","isMain":false,"metricName":"Reference GC (%)"}]]],"referenceName":"KF302034","date":"18 July 2018, Wednesday, 16:08:44","order":[0,1,2,3,4,5,6,7,8,9,10,11,12,13],"assembliesNames":["CLC","IDBA\_UD","MEGAHIT","MetaVelvet","MIRA","Ray\_Meta","SOAPdenovo2","SPAdes","SPAdes\_meta","SPAdes\_sc","SPAdes\_sc\_careful","Velvet","VICUNA","Geneious"]},{"assembliesWithNs":null,"minContig":500,"report":[["Genome statistics",[{"values":["99.558","99.558","99.590","99.558","99.558","99.561","99.578","99.561","15.092","99.570","99.570","99.558","99.570","99.561","99.558","99.561"],"quality":"More is better","isMain":true,"metricName":"Genome fraction (%)"},{"values":["1.000","1.001","1.362","1.000","1.000","1.000","1.177","1.000","1.001","1.002","1.002","1.000","1.002","1.000","1.027","1.020"],"quality":"Less is better","isMain":true,"metricName":"Duplication ratio"},{"values":[35176,35176,35176,35176,35176,35177,27926,35177,284,35180,35180,35176,35180,35177,15625,35176],"quality":"More is better","isMain":true,"metricName":"Largest alignment"},{"values":[35176,35176,39179,35176,35176,35177,36988,35177,5336,35180,35180,35176,35180,35177,36088,35615],"quality":"More is better","isMain":true,"metricName":"Total aligned length"},{"values":[35176,35210,35304,35176,35176,35178,28423,35181,28742,35250,35250,35176,35250,35178,12574,36100],"quality":"More is better","isMain":false,"metricName":"NG50"},{"values":[35176,35210,35304,35176,35176,35178,28423,35181,28742,35250,35250,35176,35250,35178,12574,36100],"quality":"More is better","isMain":false,"metricName":"NG75"},{"values":[35176,35176,35176,35176,35176,35177,27926,35177,null,35180,35180,35176,35180,35177,12556,35176],"quality":"More is better","isMain":false,"metricName":"NA50"},{"values":[35176,35176,null,35176,35176,35177,7315,35177,null,35180,35180,35176,35180,35177,12556,35176],"quality":"More is better","isMain":false,"metricName":"NA75"},{"values":[35176,35176,35176,35176,35176,35177,27926,35177,null,35180,35180,35176,35180,35177,12556,35176],"quality":"More is better","isMain":true,"metricName":"NGA50"},{"values":[35176,35176,35176,35176,35176,35177,27926,35177,null,35180,35180,35176,35180,35177,12556,35176],"quality":"More is better","isMain":false,"metricName":"NGA75"},{"values":[1,1,1,1,1,1,1,1,1,1,1,1,1,1,2,1],"quality":"Less is better","isMain":false,"metricName":"LG50"},{"values":[1,1,1,1,1,1,1,1,1,1,1,1,1,1,2,1],"quality":"Less is better","isMain":false,"metricName":"LG75"},{"values":[1,1,1,1,1,1,1,1,null,1,1,1,1,1,2,1],"quality":"Less is better","isMain":false,"metricName":"LA50"},{"values":[1,1,null,1,1,1,2,1,null,1,1,1,1,1,2,1],"quality":"Less is better","isMain":false,"metricName":"LA75"},{"values":[1,1,1,1,1,1,1,1,null,1,1,1,1,1,2,1],"quality":"Less is better","isMain":true,"metricName":"LGA50"},{"values":[1,1,1,1,1,1,1,1,null,1,1,1,1,1,2,1],"quality":"Less is better","isMain":false,"metricName":"LGA75"}]],["Misassemblies",[{"values":[0,0,0,0,0,0,0,0,0,0,0,0,0,0,0,0],"quality":"Less is better","isMain":true,"metricName":"# misassemblies"},{"values":[0,0,0,0,0,0,0,0,0,0,0,0,0,0,0,0],"quality":"Less is better","isMain":false,"metricName":" # relocations"},{"values":[0,0,0,0,0,0,0,0,0,0,0,0,0,0,0,0],"quality":"Less is better","isMain":false,"metricName":" # translocations"},{"values":[0,0,0,0,0,0,0,0,0,0,0,0,0,0,0,0],"quality":"Less is better","isMain":false,"metricName":" # inversions"},{"values":[0,0,0,0,0,0,0,0,0,0,0,0,0,0,0,0],"quality":"Less is better","isMain":false,"metricName":"# misassembled contigs"},{"values":[0,0,0,0,0,0,0,0,0,0,0,0,0,0,0,0],"quality":"Less is better","isMain":true,"metricName":"Misassembled contigs length"},{"values":[0,0,0,0,0,0,1,0,0,0,0,0,0,0,0,1],"quality":"Less is better","isMain":false,"metricName":"# local misassemblies"},{"values":[0,0,1,0,0,0,2,0,1,0,0,0,0,0,0,0],"quality":"Less is better","isMain":false,"metricName":"# unaligned mis. contigs"}]],["Unaligned",[{"values":[0,0,0,0,0,0,0,0,0,0,0,0,0,0,0,0],"quality":"Less is better","isMain":false,"metricName":"# fully unaligned contigs"},{"values":[0,0,0,0,0,0,0,0,0,0,0,0,0,0,0,0],"quality":"Less is better","isMain":false,"metricName":"Fully unaligned length"},{"values":[0,0,24,0,0,0,3,0,1,0,0,0,0,0,0,2],"quality":"Less is better","isMain":false,"metricName":"# partially unaligned contigs"},{"values":[0,0,15071,0,0,0,1758,0,23406,0,0,0,0,0,0,2489],"quality":"Less is better","isMain":false,"metricName":"Partially unaligned length"}]],["Mismatches",[{"values":[0,0,1,0,0,0,1,0,2,0,0,0,0,0,0,0],"quality":"Less is better","isMain":false,"metricName":"# mismatches"},{"values":[2,2,2,2,2,2,2,2,3,2,2,2,2,2,2,2],"quality":"Less is better","isMain":false,"metricName":"# indels"},{"values":[2,2,2,2,2,2,2,2,6,2,2,2,2,2,2,2],"quality":"Less is better","isMain":false,"metricName":"Indels length"},{"values":["0.00","0.00","2.84","0.00","0.00","0.00","2.84","0.00","37.51","0.00","0.00","0.00","0.00","0.00","0.00","0.00"],"quality":"Less is better","isMain":true,"metricName":"# mismatches per 100 kbp"},{"values":["5.69","5.69","5.68","5.69","5.69","5.69","5.68","5.69","56.26","5.69","5.69","5.69","5.69","5.69","5.69","5.69"],"quality":"Less is better","isMain":true,"metricName":"# indels per 100 kbp"},{"values":[2,2,2,2,2,2,2,2,3,2,2,2,2,2,2,2],"quality":"Less is better","isMain":false,"metricName":" # indels (<= 5 bp)"},{"values":[0,0,0,0,0,0,0,0,0,0,0,0,0,0,0,0],"quality":"Less is better","isMain":false,"metricName":" # indels (> 5 bp)"},{"values":[0,0,0,0,0,0,7,0,8275,0,0,0,0,0,0,60],"quality":"Less is better","isMain":false,"metricName":"# N's"},{"values":["0.00","0.00","0.00","0.00","0.00","0.00","16.22","0.00","28790.62","0.00","0.00","0.00","0.00","0.00","0.00","156.37"],"quality":"Less is better","isMain":true,"metricName":"# N's per 100 kbp"}]],["Statistics without reference",[{"values":[1,1,44,1,1,1,14,1,1,1,1,1,1,1,4,3],"quality":"Equal","isMain":true,"metricName":"# contigs"},{"values":[1,1,1,1,1,1,2,1,1,1,1,1,1,1,4,2],"quality":"Equal","isMain":false,"metricName":"# contigs (>= 1000 bp)"},{"values":[1,1,1,1,1,1,2,1,1,1,1,1,1,1,3,1],"quality":"Equal","isMain":false,"metricName":"# contigs (>= 5000 bp)"},{"values":[1,1,1,1,1,1,1,1,1,1,1,1,1,1,2,1],"quality":"Equal","isMain":false,"metricName":"# contigs (>= 10000 bp)"},{"values":[1,1,1,1,1,1,1,1,1,1,1,1,1,1,0,1],"quality":"Equal","isMain":false,"metricName":"# contigs (>= 25000 bp)"},{"values":[0,0,0,0,0,0,0,0,0,0,0,0,0,0,0,0],"quality":"Equal","isMain":false,"metricName":"# contigs (>= 50000 bp)"},{"values":[35176,35210,35304,35176,35176,35178,28423,35181,28742,35250,35250,35176,35250,35178,15645,36100],"quality":"More is better","isMain":true,"metricName":"Largest contig"},{"values":[35176,35210,62977,35176,35176,35178,43155,35181,28742,35250,35250,35176,35250,35178,36126,38371],"quality":"More is better","isMain":true,"metricName":"Total length"},{"values":[35176,35210,35304,35176,35176,35178,36158,35181,28742,35250,35250,35176,35250,35178,36126,37827],"quality":"More is better","isMain":true,"metricName":"Total length (>= 1000 bp)"},{"values":[35176,35210,35304,35176,35176,35178,36158,35181,28742,35250,35250,35176,35250,35178,33586,36100],"quality":"More is better","isMain":false,"metricName":"Total length (>= 5000 bp)"},{"values":[35176,35210,35304,35176,35176,35178,28423,35181,28742,35250,35250,35176,35250,35178,28219,36100],"quality":"More is better","isMain":true,"metricName":"Total length (>= 10000 bp)"},{"values":[35176,35210,35304,35176,35176,35178,28423,35181,28742,35250,35250,35176,35250,35178,0,36100],"quality":"More is better","isMain":false,"metricName":"Total length (>= 25000 bp)"},{"values":[0,0,0,0,0,0,0,0,0,0,0,0,0,0,0,0],"quality":"More is better","isMain":true,"metricName":"Total length (>= 50000 bp)"},{"values":[35176,35210,35304,35176,35176,35178,28423,35181,28742,35250,35250,35176,35250,35178,12574,36100],"quality":"More is better","isMain":false,"metricName":"N50"},{"values":[35176,35210,636,35176,35176,35178,7735,35181,28742,35250,35250,35176,35250,35178,12574,36100],"quality":"More is better","isMain":false,"metricName":"N75"},{"values":[1,1,1,1,1,1,1,1,1,1,1,1,1,1,2,1],"quality":"Less is better","isMain":false,"metricName":"L50"},{"values":[1,1,17,1,1,1,2,1,1,1,1,1,1,1,2,1],"quality":"Less is better","isMain":false,"metricName":"L75"},{"values":["44.91","44.89","42.47","44.91","44.91","44.91","43.79","44.92","44.73","44.90","44.90","44.91","44.90","44.91","44.94","44.41"],"quality":"Equal","isMain":false,"metricName":"GC (%)"}]],["Predicted genes",[]],["Similarity statistics",[{"values":[1,1,1,1,1,1,0,1,0,1,1,1,1,1,0,1],"quality":"Equal","isMain":false,"metricName":"# similar correct contigs"},{"values":[0,0,0,0,0,0,0,0,0,0,0,0,0,0,0,0],"quality":"Equal","isMain":false,"metricName":"# similar misassembled blocks"}]],["Reference statistics",[{"values":[35330,35330,35330,35330,35330,35330,35330,35330,35330,35330,35330,35330,35330,35330,35330,35330],"quality":"Equal","isMain":false,"metricName":"Reference length"},{"values":[1,1,1,1,1,1,1,1,1,1,1,1,1,1,1,1],"quality":"Equal","isMain":false,"metricName":"Reference fragments"},{"values":["44.87","44.87","44.87","44.87","44.87","44.87","44.87","44.87","44.87","44.87","44.87","44.87","44.87","44.87","44.87","44.87"],"quality":"Equal","isMain":false,"metricName":"Reference GC (%)"}]]],"referenceName":"KF302035","date":"18 July 2018, Wednesday, 16:08:54","order":[0,1,2,3,4,5,6,7,8,9,10,11,12,13,14,15],"assembliesNames":["ABySS\_63","ABySS\_127","CLC","IDBA\_UD","MEGAHIT","MetaVelvet","MIRA","Ray\_Meta","SOAPdenovo2","SPAdes","SPAdes\_meta","SPAdes\_sc","SPAdes\_sc\_careful","Velvet","VICUNA","Geneious"]},{"assembliesWithNs":null,"minContig":500,"report":[["Genome statistics",[{"values":["98.699","98.744","98.744","37.186","98.744","98.744","90.018","98.744","98.744","98.744","98.744","37.186","98.744","98.744"],"quality":"More is better","isMain":true,"metricName":"Genome fraction (%)"},{"values":["1.000","1.003","1.004","1.000","1.004","1.013","0.987","1.002","1.001","1.001","1.001","1.000","1.007","1.359"],"quality":"Less is better","isMain":true,"metricName":"Duplication ratio"},{"values":[37711,37827,37869,7580,37863,38215,33963,37805,37783,37783,37783,7580,38007,29512],"quality":"More is better","isMain":true,"metricName":"Largest alignment"},{"values":[37711,37827,37869,14208,37863,38215,33963,37805,37783,37783,37783,14208,38007,51269],"quality":"More is better","isMain":true,"metricName":"Total aligned length"},{"values":[37728,37827,37869,null,37863,38215,37358,37805,37783,37783,37783,null,38007,51269],"quality":"More is better","isMain":false,"metricName":"NG50"},{"values":[37728,37827,37869,null,37863,38215,37358,37805,37783,37783,37783,null,38007,51269],"quality":"More is better","isMain":false,"metricName":"NG75"},{"values":[37711,37827,37869,7580,37863,38215,33963,37805,37783,37783,37783,7580,38007,29512],"quality":"More is better","isMain":false,"metricName":"NA50"},{"values":[37711,37827,37869,6628,37863,38215,33963,37805,37783,37783,37783,6628,38007,21757],"quality":"More is better","isMain":false,"metricName":"NA75"},{"values":[37711,37827,37869,null,37863,38215,33963,37805,37783,37783,37783,null,38007,29512],"quality":"More is better","isMain":true,"metricName":"NGA50"},{"values":[37711,37827,37869,null,37863,38215,33963,37805,37783,37783,37783,null,38007,29512],"quality":"More is better","isMain":false,"metricName":"NGA75"},{"values":[1,1,1,null,1,1,1,1,1,1,1,null,1,1],"quality":"Less is better","isMain":false,"metricName":"LG50"},{"values":[1,1,1,null,1,1,1,1,1,1,1,null,1,1],"quality":"Less is better","isMain":false,"metricName":"LG75"},{"values":[1,1,1,1,1,1,1,1,1,1,1,1,1,1],"quality":"Less is better","isMain":false,"metricName":"LA50"},{"values":[1,1,1,2,1,1,1,1,1,1,1,2,1,2],"quality":"Less is better","isMain":false,"metricName":"LA75"},{"values":[1,1,1,null,1,1,1,1,1,1,1,null,1,1],"quality":"Less is better","isMain":true,"metricName":"LGA50"},{"values":[1,1,1,null,1,1,1,1,1,1,1,null,1,1],"quality":"Less is better","isMain":false,"metricName":"LGA75"}]],["Misassemblies",[{"values":[0,0,0,0,0,0,0,0,0,0,0,0,0,1],"quality":"Less is better","isMain":true,"metricName":"# misassemblies"},{"values":[0,0,0,0,0,0,0,0,0,0,0,0,0,1],"quality":"Less is better","isMain":false,"metricName":" # relocations"},{"values":[0,0,0,0,0,0,0,0,0,0,0,0,0,0],"quality":"Less is better","isMain":false,"metricName":" # translocations"},{"values":[0,0,0,0,0,0,0,0,0,0,0,0,0,0],"quality":"Less is better","isMain":false,"metricName":" # inversions"},{"values":[0,0,0,0,0,0,0,0,0,0,0,0,0,1],"quality":"Less is better","isMain":false,"metricName":"# misassembled contigs"},{"values":[0,0,0,0,0,0,0,0,0,0,0,0,0,51269],"quality":"Less is better","isMain":true,"metricName":"Misassembled contigs length"},{"values":[1,1,1,1,1,1,1,1,1,1,1,1,1,0],"quality":"Less is better","isMain":false,"metricName":"# local misassemblies"},{"values":[0,0,0,0,0,0,0,0,0,0,0,0,0,0],"quality":"Less is better","isMain":false,"metricName":"# unaligned mis. contigs"}]],["Unaligned",[{"values":[0,0,0,0,0,0,0,0,0,0,0,0,0,0],"quality":"Less is better","isMain":false,"metricName":"# fully unaligned contigs"},{"values":[0,0,0,0,0,0,0,0,0,0,0,0,0,0],"quality":"Less is better","isMain":false,"metricName":"Fully unaligned length"},{"values":[0,0,0,0,0,0,1,0,0,0,0,0,0,0],"quality":"Less is better","isMain":false,"metricName":"# partially unaligned contigs"},{"values":[0,0,0,0,0,0,3395,0,0,0,0,0,0,0],"quality":"Less is better","isMain":false,"metricName":"Partially unaligned length"}]],["Mismatches",[{"values":[1,0,0,0,0,0,0,0,0,0,0,0,0,0],"quality":"Less is better","isMain":false,"metricName":"# mismatches"},{"values":[0,0,0,0,0,0,47,0,0,0,0,0,0,0],"quality":"Less is better","isMain":false,"metricName":"# indels"},{"values":[0,0,0,0,0,0,779,0,0,0,0,0,0,0],"quality":"Less is better","isMain":false,"metricName":"Indels length"},{"values":["2.65","0.00","0.00","0.00","0.00","0.00","0.00","0.00","0.00","0.00","0.00","0.00","0.00","0.00"],"quality":"Less is better","isMain":true,"metricName":"# mismatches per 100 kbp"},{"values":["0.00","0.00","0.00","0.00","0.00","0.00","136.65","0.00","0.00","0.00","0.00","0.00","0.00","0.00"],"quality":"Less is better","isMain":true,"metricName":"# indels per 100 kbp"},{"values":[0,0,0,0,0,0,5,0,0,0,0,0,0,0],"quality":"Less is better","isMain":false,"metricName":" # indels (<= 5 bp)"},{"values":[0,0,0,0,0,0,42,0,0,0,0,0,0,0],"quality":"Less is better","isMain":false,"metricName":" # indels (> 5 bp)"},{"values":[0,0,0,0,0,0,1010,0,0,0,0,0,0,0],"quality":"Less is better","isMain":false,"metricName":"# N's"},{"values":["0.00","0.00","0.00","0.00","0.00","0.00","2703.57","0.00","0.00","0.00","0.00","0.00","0.00","0.00"],"quality":"Less is better","isMain":true,"metricName":"# N's per 100 kbp"}]],["Statistics without reference",[{"values":[1,1,1,2,1,1,1,1,1,1,1,2,1,1],"quality":"Equal","isMain":true,"metricName":"# contigs"},{"values":[1,1,1,2,1,1,1,1,1,1,1,2,1,1],"quality":"Equal","isMain":false,"metricName":"# contigs (>= 1000 bp)"},{"values":[1,1,1,2,1,1,1,1,1,1,1,2,1,1],"quality":"Equal","isMain":false,"metricName":"# contigs (>= 5000 bp)"},{"values":[1,1,1,0,1,1,1,1,1,1,1,0,1,1],"quality":"Equal","isMain":false,"metricName":"# contigs (>= 10000 bp)"},{"values":[1,1,1,0,1,1,1,1,1,1,1,0,1,1],"quality":"Equal","isMain":false,"metricName":"# contigs (>= 25000 bp)"},{"values":[0,0,0,0,0,0,0,0,0,0,0,0,0,1],"quality":"Equal","isMain":false,"metricName":"# contigs (>= 50000 bp)"},{"values":[37728,37827,37869,7580,37863,38215,37358,37805,37783,37783,37783,7580,38007,51269],"quality":"More is better","isMain":true,"metricName":"Largest contig"},{"values":[37728,37827,37869,14208,37863,38215,37358,37805,37783,37783,37783,14208,38007,51269],"quality":"More is better","isMain":true,"metricName":"Total length"},{"values":[37728,37827,37869,14208,37863,38215,37358,37805,37783,37783,37783,14208,38007,51269],"quality":"More is better","isMain":true,"metricName":"Total length (>= 1000 bp)"},{"values":[37728,37827,37869,14208,37863,38215,37358,37805,37783,37783,37783,14208,38007,51269],"quality":"More is better","isMain":false,"metricName":"Total length (>= 5000 bp)"},{"values":[37728,37827,37869,0,37863,38215,37358,37805,37783,37783,37783,0,38007,51269],"quality":"More is better","isMain":true,"metricName":"Total length (>= 10000 bp)"},{"values":[37728,37827,37869,0,37863,38215,37358,37805,37783,37783,37783,0,38007,51269],"quality":"More is better","isMain":false,"metricName":"Total length (>= 25000 bp)"},{"values":[0,0,0,0,0,0,0,0,0,0,0,0,0,51269],"quality":"More is better","isMain":true,"metricName":"Total length (>= 50000 bp)"},{"values":[37728,37827,37869,7580,37863,38215,37358,37805,37783,37783,37783,7580,38007,51269],"quality":"More is better","isMain":false,"metricName":"N50"},{"values":[37728,37827,37869,6628,37863,38215,37358,37805,37783,37783,37783,6628,38007,51269],"quality":"More is better","isMain":false,"metricName":"N75"},{"values":[1,1,1,1,1,1,1,1,1,1,1,1,1,1],"quality":"Less is better","isMain":false,"metricName":"L50"},{"values":[1,1,1,2,1,1,1,1,1,1,1,2,1,1],"quality":"Less is better","isMain":false,"metricName":"L75"},{"values":["40.22","40.22","40.21","39.47","40.21","40.19","40.24","40.20","40.19","40.24","40.19","39.47","40.19","40.10"],"quality":"Equal","isMain":false,"metricName":"GC (%)"}]],["Predicted genes",[]],["Similarity statistics",[{"values":[0,0,0,0,0,0,0,0,0,0,0,0,0,0],"quality":"Equal","isMain":false,"metricName":"# similar correct contigs"},{"values":[0,0,0,0,0,0,0,0,0,0,0,0,0,0],"quality":"Equal","isMain":false,"metricName":"# similar misassembled blocks"}]],["Reference statistics",[{"values":[38208,38208,38208,38208,38208,38208,38208,38208,38208,38208,38208,38208,38208,38208],"quality":"Equal","isMain":false,"metricName":"Reference length"},{"values":[1,1,1,1,1,1,1,1,1,1,1,1,1,1],"quality":"Equal","isMain":false,"metricName":"Reference fragments"},{"values":["40.23","40.23","40.23","40.23","40.23","40.23","40.23","40.23","40.23","40.23","40.23","40.23","40.23","40.23"],"quality":"Equal","isMain":false,"metricName":"Reference GC (%)"}]]],"referenceName":"KF302036","date":"18 July 2018, Wednesday, 16:09:03","order":[0,1,2,3,4,5,6,7,8,9,10,11,12,13],"assembliesNames":["CLC","IDBA\_UD","MEGAHIT","MetaVelvet","MIRA","Ray\_Meta","SOAPdenovo2","SPAdes","SPAdes\_meta","SPAdes\_sc","SPAdes\_sc\_careful","Velvet","VICUNA","Geneious"]},{"assembliesWithNs":null,"minContig":500,"report":[["Genome statistics",[{"values":["98.925","98.925","98.925","98.925","98.925","98.925","98.925","87.396","98.925","98.925","98.925","98.925","98.925","98.925","98.925"],"quality":"More is better","isMain":true,"metricName":"Genome fraction (%)"},{"values":["1.003","1.000","1.002","1.003","1.001","1.004","1.012","0.986","1.002","1.001","1.001","1.001","1.001","1.013","1.198"],"quality":"Less is better","isMain":true,"metricName":"Duplication ratio"},{"values":[44671,44551,44650,44692,44613,44737,45078,38805,44628,44606,44606,44606,44613,40569,29763],"quality":"More is better","isMain":true,"metricName":"Largest alignment"},{"values":[44671,44551,44650,44692,44613,44737,45078,38805,44628,44606,44606,44606,44613,45152,53369],"quality":"More is better","isMain":true,"metricName":"Total aligned length"},{"values":[44671,44551,44650,44692,44613,44737,45078,44105,44628,44606,44606,44606,44613,40569,53369],"quality":"More is better","isMain":false,"metricName":"NG50"},{"values":[44671,44551,44650,44692,44613,44737,45078,44105,44628,44606,44606,44606,44613,40569,53369],"quality":"More is better","isMain":false,"metricName":"NG75"},{"values":[44671,44551,44650,44692,44613,44737,45078,38805,44628,44606,44606,44606,44613,40569,29763],"quality":"More is better","isMain":false,"metricName":"NA50"},{"values":[44671,44551,44650,44692,44613,44737,45078,38805,44628,44606,44606,44606,44613,40569,23606],"quality":"More is better","isMain":false,"metricName":"NA75"},{"values":[44671,44551,44650,44692,44613,44737,45078,38805,44628,44606,44606,44606,44613,40569,29763],"quality":"More is better","isMain":true,"metricName":"NGA50"},{"values":[44671,44551,44650,44692,44613,44737,45078,38805,44628,44606,44606,44606,44613,40569,23606],"quality":"More is better","isMain":false,"metricName":"NGA75"},{"values":[1,1,1,1,1,1,1,1,1,1,1,1,1,1,1],"quality":"Less is better","isMain":false,"metricName":"LG50"},{"values":[1,1,1,1,1,1,1,1,1,1,1,1,1,1,1],"quality":"Less is better","isMain":false,"metricName":"LG75"},{"values":[1,1,1,1,1,1,1,1,1,1,1,1,1,1,1],"quality":"Less is better","isMain":false,"metricName":"LA50"},{"values":[1,1,1,1,1,1,1,1,1,1,1,1,1,1,2],"quality":"Less is better","isMain":false,"metricName":"LA75"},{"values":[1,1,1,1,1,1,1,1,1,1,1,1,1,1,1],"quality":"Less is better","isMain":true,"metricName":"LGA50"},{"values":[1,1,1,1,1,1,1,1,1,1,1,1,1,1,2],"quality":"Less is better","isMain":false,"metricName":"LGA75"}]],["Misassemblies",[{"values":[0,0,0,0,0,0,0,0,0,0,0,0,0,0,1],"quality":"Less is better","isMain":true,"metricName":"# misassemblies"},{"values":[0,0,0,0,0,0,0,0,0,0,0,0,0,0,1],"quality":"Less is better","isMain":false,"metricName":" # relocations"},{"values":[0,0,0,0,0,0,0,0,0,0,0,0,0,0,0],"quality":"Less is better","isMain":false,"metricName":" # translocations"},{"values":[0,0,0,0,0,0,0,0,0,0,0,0,0,0,0],"quality":"Less is better","isMain":false,"metricName":" # inversions"},{"values":[0,0,0,0,0,0,0,0,0,0,0,0,0,0,1],"quality":"Less is better","isMain":false,"metricName":"# misassembled contigs"},{"values":[0,0,0,0,0,0,0,0,0,0,0,0,0,0,53369],"quality":"Less is better","isMain":true,"metricName":"Misassembled contigs length"},{"values":[1,1,1,1,1,1,1,2,1,1,1,1,1,1,0],"quality":"Less is better","isMain":false,"metricName":"# local misassemblies"},{"values":[0,0,0,0,0,0,0,0,0,0,0,0,0,0,0],"quality":"Less is better","isMain":false,"metricName":"# unaligned mis. contigs"}]],["Unaligned",[{"values":[0,0,0,0,0,0,0,0,0,0,0,0,0,0,0],"quality":"Less is better","isMain":false,"metricName":"# fully unaligned contigs"},{"values":[0,0,0,0,0,0,0,0,0,0,0,0,0,0,0],"quality":"Less is better","isMain":false,"metricName":"Fully unaligned length"},{"values":[0,0,0,0,0,0,0,1,0,0,0,0,0,0,0],"quality":"Less is better","isMain":false,"metricName":"# partially unaligned contigs"},{"values":[0,0,0,0,0,0,0,5300,0,0,0,0,0,0,0],"quality":"Less is better","isMain":false,"metricName":"Partially unaligned length"}]],["Mismatches",[{"values":[0,0,0,0,0,0,0,31,0,0,0,0,0,0,0],"quality":"Less is better","isMain":false,"metricName":"# mismatches"},{"values":[0,0,0,0,0,0,0,59,0,0,0,0,0,0,0],"quality":"Less is better","isMain":false,"metricName":"# indels"},{"values":[0,0,0,0,0,0,0,1167,0,0,0,0,0,0,0],"quality":"Less is better","isMain":false,"metricName":"Indels length"},{"values":["0.00","0.00","0.00","0.00","0.00","0.00","0.00","78.76","0.00","0.00","0.00","0.00","0.00","0.00","0.00"],"quality":"Less is better","isMain":true,"metricName":"# mismatches per 100 kbp"},{"values":["0.00","0.00","0.00","0.00","0.00","0.00","0.00","149.90","0.00","0.00","0.00","0.00","0.00","0.00","0.00"],"quality":"Less is better","isMain":true,"metricName":"# indels per 100 kbp"},{"values":[0,0,0,0,0,0,0,6,0,0,0,0,0,0,0],"quality":"Less is better","isMain":false,"metricName":" # indels (<= 5 bp)"},{"values":[0,0,0,0,0,0,0,53,0,0,0,0,0,0,0],"quality":"Less is better","isMain":false,"metricName":" # indels (> 5 bp)"},{"values":[0,0,0,0,0,0,0,1430,0,0,0,0,0,0,0],"quality":"Less is better","isMain":false,"metricName":"# N's"},{"values":["0.00","0.00","0.00","0.00","0.00","0.00","0.00","3242.26","0.00","0.00","0.00","0.00","0.00","0.00","0.00"],"quality":"Less is better","isMain":true,"metricName":"# N's per 100 kbp"}]],["Statistics without reference",[{"values":[1,1,1,1,1,1,1,1,1,1,1,1,1,2,1],"quality":"Equal","isMain":true,"metricName":"# contigs"},{"values":[1,1,1,1,1,1,1,1,1,1,1,1,1,2,1],"quality":"Equal","isMain":false,"metricName":"# contigs (>= 1000 bp)"},{"values":[1,1,1,1,1,1,1,1,1,1,1,1,1,1,1],"quality":"Equal","isMain":false,"metricName":"# contigs (>= 5000 bp)"},{"values":[1,1,1,1,1,1,1,1,1,1,1,1,1,1,1],"quality":"Equal","isMain":false,"metricName":"# contigs (>= 10000 bp)"},{"values":[1,1,1,1,1,1,1,1,1,1,1,1,1,1,1],"quality":"Equal","isMain":false,"metricName":"# contigs (>= 25000 bp)"},{"values":[0,0,0,0,0,0,0,0,0,0,0,0,0,0,1],"quality":"Equal","isMain":false,"metricName":"# contigs (>= 50000 bp)"},{"values":[44671,44551,44650,44692,44613,44737,45078,44105,44628,44606,44606,44606,44613,40569,53369],"quality":"More is better","isMain":true,"metricName":"Largest contig"},{"values":[44671,44551,44650,44692,44613,44737,45078,44105,44628,44606,44606,44606,44613,45152,53369],"quality":"More is better","isMain":true,"metricName":"Total length"},{"values":[44671,44551,44650,44692,44613,44737,45078,44105,44628,44606,44606,44606,44613,45152,53369],"quality":"More is better","isMain":true,"metricName":"Total length (>= 1000 bp)"},{"values":[44671,44551,44650,44692,44613,44737,45078,44105,44628,44606,44606,44606,44613,40569,53369],"quality":"More is better","isMain":false,"metricName":"Total length (>= 5000 bp)"},{"values":[44671,44551,44650,44692,44613,44737,45078,44105,44628,44606,44606,44606,44613,40569,53369],"quality":"More is better","isMain":true,"metricName":"Total length (>= 10000 bp)"},{"values":[44671,44551,44650,44692,44613,44737,45078,44105,44628,44606,44606,44606,44613,40569,53369],"quality":"More is better","isMain":false,"metricName":"Total length (>= 25000 bp)"},{"values":[0,0,0,0,0,0,0,0,0,0,0,0,0,0,53369],"quality":"More is better","isMain":true,"metricName":"Total length (>= 50000 bp)"},{"values":[44671,44551,44650,44692,44613,44737,45078,44105,44628,44606,44606,44606,44613,40569,53369],"quality":"More is better","isMain":false,"metricName":"N50"},{"values":[44671,44551,44650,44692,44613,44737,45078,44105,44628,44606,44606,44606,44613,40569,53369],"quality":"More is better","isMain":false,"metricName":"N75"},{"values":[1,1,1,1,1,1,1,1,1,1,1,1,1,1,1],"quality":"Less is better","isMain":false,"metricName":"L50"},{"values":[1,1,1,1,1,1,1,1,1,1,1,1,1,1,1],"quality":"Less is better","isMain":false,"metricName":"L75"},{"values":["44.69","44.65","44.65","44.62","44.68","44.70","44.65","44.66","44.67","44.67","44.66","44.67","44.68","44.71","44.70"],"quality":"Equal","isMain":false,"metricName":"GC (%)"}]],["Predicted genes",[]],["Similarity statistics",[{"values":[0,0,0,0,0,0,0,0,0,0,0,0,0,0,0],"quality":"Equal","isMain":false,"metricName":"# similar correct contigs"},{"values":[2,0,0,2,2,2,0,0,2,2,0,2,2,0,1],"quality":"Equal","isMain":false,"metricName":"# similar misassembled blocks"}]],["Reference statistics",[{"values":[45035,45035,45035,45035,45035,45035,45035,45035,45035,45035,45035,45035,45035,45035,45035],"quality":"Equal","isMain":false,"metricName":"Reference length"},{"values":[1,1,1,1,1,1,1,1,1,1,1,1,1,1,1],"quality":"Equal","isMain":false,"metricName":"Reference fragments"},{"values":["44.67","44.67","44.67","44.67","44.67","44.67","44.67","44.67","44.67","44.67","44.67","44.67","44.67","44.67","44.67"],"quality":"Equal","isMain":false,"metricName":"Reference GC (%)"}]]],"referenceName":"KF302037","date":"18 July 2018, Wednesday, 16:09:12","order":[0,1,2,3,4,5,6,7,8,9,10,11,12,13,14],"assembliesNames":["ABySS\_63","CLC","IDBA\_UD","MEGAHIT","MetaVelvet","MIRA","Ray\_Meta","SOAPdenovo2","SPAdes","SPAdes\_meta","SPAdes\_sc","SPAdes\_sc\_careful","Velvet","VICUNA","Geneious"]},{"assembliesWithNs":null,"minContig":500,"report":[["Genome statistics",[{"values":["100.000","100.000","100.000","100.000","100.000","89.207","100.000","100.000","100.000","100.000","100.000","89.207","100.000","100.000"],"quality":"More is better","isMain":true,"metricName":"Genome fraction (%)"},{"values":["1.009","1.645","1.000","1.017","1.011","1.012","1.878","1.013","1.008","1.008","1.008","1.012","1.071","1.071"],"quality":"Less is better","isMain":true,"metricName":"Duplication ratio"},{"values":[6141,6089,6089,6188,6154,2581,5716,6166,6134,6134,6134,2581,6508,6508],"quality":"More is better","isMain":true,"metricName":"Largest alignment"},{"values":[6141,10015,6089,6188,6154,5494,11429,6166,6134,6134,6134,5494,6508,6508],"quality":"More is better","isMain":true,"metricName":"Total aligned length"},{"values":[6141,10015,6089,6188,6154,2009,11429,6166,6134,6134,6134,2009,6518,6518],"quality":"More is better","isMain":false,"metricName":"NG50"},{"values":[6141,10015,6089,6188,6154,2009,11429,6166,6134,6134,6134,2009,6518,6518],"quality":"More is better","isMain":false,"metricName":"NG75"},{"values":[6141,6089,6089,6188,6154,2009,5716,6166,6134,6134,6134,2009,6508,6508],"quality":"More is better","isMain":false,"metricName":"NA50"},{"values":[6141,2766,6089,6188,6154,2009,5713,6166,6134,6134,6134,2009,6508,6508],"quality":"More is better","isMain":false,"metricName":"NA75"},{"values":[6141,6089,6089,6188,6154,2009,5716,6166,6134,6134,6134,2009,6508,6508],"quality":"More is better","isMain":true,"metricName":"NGA50"},{"values":[6141,6089,6089,6188,6154,2009,5716,6166,6134,6134,6134,2009,6508,6508],"quality":"More is better","isMain":false,"metricName":"NGA75"},{"values":[1,1,1,1,1,2,1,1,1,1,1,2,1,1],"quality":"Less is better","isMain":false,"metricName":"LG50"},{"values":[1,1,1,1,1,2,1,1,1,1,1,2,1,1],"quality":"Less is better","isMain":false,"metricName":"LG75"},{"values":[1,1,1,1,1,2,1,1,1,1,1,2,1,1],"quality":"Less is better","isMain":false,"metricName":"LA50"},{"values":[1,2,1,1,1,2,2,1,1,1,1,2,1,1],"quality":"Less is better","isMain":false,"metricName":"LA75"},{"values":[1,1,1,1,1,2,1,1,1,1,1,2,1,1],"quality":"Less is better","isMain":true,"metricName":"LGA50"},{"values":[1,1,1,1,1,2,1,1,1,1,1,2,1,1],"quality":"Less is better","isMain":false,"metricName":"LGA75"}]],["Misassemblies",[{"values":[0,2,0,0,0,0,1,0,0,0,0,0,0,0],"quality":"Less is better","isMain":true,"metricName":"# misassemblies"},{"values":[0,2,0,0,0,0,1,0,0,0,0,0,0,0],"quality":"Less is better","isMain":false,"metricName":" # relocations"},{"values":[0,0,0,0,0,0,0,0,0,0,0,0,0,0],"quality":"Less is better","isMain":false,"metricName":" # translocations"},{"values":[0,0,0,0,0,0,0,0,0,0,0,0,0,0],"quality":"Less is better","isMain":false,"metricName":" # inversions"},{"values":[0,1,0,0,0,0,1,0,0,0,0,0,0,0],"quality":"Less is better","isMain":false,"metricName":"# misassembled contigs"},{"values":[0,10015,0,0,0,0,11429,0,0,0,0,0,0,0],"quality":"Less is better","isMain":true,"metricName":"Misassembled contigs length"},{"values":[0,0,0,1,0,0,0,0,0,0,0,0,1,1],"quality":"Less is better","isMain":false,"metricName":"# local misassemblies"},{"values":[0,0,0,0,0,0,0,0,0,0,0,0,0,0],"quality":"Less is better","isMain":false,"metricName":"# unaligned mis. contigs"}]],["Unaligned",[{"values":[0,0,0,0,0,0,0,0,0,0,0,0,0,0],"quality":"Less is better","isMain":false,"metricName":"# fully unaligned contigs"},{"values":[0,0,0,0,0,0,0,0,0,0,0,0,0,0],"quality":"Less is better","isMain":false,"metricName":"Fully unaligned length"},{"values":[0,0,0,0,0,0,0,0,0,0,0,0,0,0],"quality":"Less is better","isMain":false,"metricName":"# partially unaligned contigs"},{"values":[0,0,0,0,0,0,0,0,0,0,0,0,0,0],"quality":"Less is better","isMain":false,"metricName":"Partially unaligned length"}]],["Mismatches",[{"values":[9,9,9,9,9,10,9,9,9,9,9,10,9,9],"quality":"Less is better","isMain":false,"metricName":"# mismatches"},{"values":[3,2,2,2,3,2,2,3,3,3,3,2,2,2],"quality":"Less is better","isMain":false,"metricName":"# indels"},{"values":[54,2,2,2,67,2,2,79,47,47,47,2,2,2],"quality":"Less is better","isMain":false,"metricName":"Indels length"},{"values":["147.86","147.86","147.86","147.86","147.86","184.16","147.86","147.86","147.86","147.86","147.86","184.16","147.86","147.86"],"quality":"Less is better","isMain":true,"metricName":"# mismatches per 100 kbp"},{"values":["49.29","32.86","32.86","32.86","49.29","36.83","32.86","49.29","49.29","49.29","49.29","36.83","32.86","32.86"],"quality":"Less is better","isMain":true,"metricName":"# indels per 100 kbp"},{"values":[2,2,2,2,2,2,2,2,2,2,2,2,2,2],"quality":"Less is better","isMain":false,"metricName":" # indels (<= 5 bp)"},{"values":[1,0,0,0,1,0,0,1,1,1,1,0,0,0],"quality":"Less is better","isMain":false,"metricName":" # indels (> 5 bp)"},{"values":[0,0,0,0,0,0,0,0,0,0,0,0,0,12],"quality":"Less is better","isMain":false,"metricName":"# N's"},{"values":["0.00","0.00","0.00","0.00","0.00","0.00","0.00","0.00","0.00","0.00","0.00","0.00","0.00","184.11"],"quality":"Less is better","isMain":true,"metricName":"# N's per 100 kbp"}]],["Statistics without reference",[{"values":[1,1,1,1,1,3,1,1,1,1,1,3,1,1],"quality":"Equal","isMain":true,"metricName":"# contigs"},{"values":[1,1,1,1,1,2,1,1,1,1,1,2,1,1],"quality":"Equal","isMain":false,"metricName":"# contigs (>= 1000 bp)"},{"values":[1,1,1,1,1,0,1,1,1,1,1,0,1,1],"quality":"Equal","isMain":false,"metricName":"# contigs (>= 5000 bp)"},{"values":[0,1,0,0,0,0,1,0,0,0,0,0,0,0],"quality":"Equal","isMain":false,"metricName":"# contigs (>= 10000 bp)"},{"values":[0,0,0,0,0,0,0,0,0,0,0,0,0,0],"quality":"Equal","isMain":false,"metricName":"# contigs (>= 25000 bp)"},{"values":[0,0,0,0,0,0,0,0,0,0,0,0,0,0],"quality":"Equal","isMain":false,"metricName":"# contigs (>= 50000 bp)"},{"values":[6141,10015,6089,6188,6154,2581,11429,6166,6134,6134,6134,2581,6518,6518],"quality":"More is better","isMain":true,"metricName":"Largest contig"},{"values":[6141,10015,6089,6188,6154,5494,11429,6166,6134,6134,6134,5494,6518,6518],"quality":"More is better","isMain":true,"metricName":"Total length"},{"values":[6141,10015,6089,6188,6154,4590,11429,6166,6134,6134,6134,4590,6518,6518],"quality":"More is better","isMain":true,"metricName":"Total length (>= 1000 bp)"},{"values":[6141,10015,6089,6188,6154,0,11429,6166,6134,6134,6134,0,6518,6518],"quality":"More is better","isMain":false,"metricName":"Total length (>= 5000 bp)"},{"values":[0,10015,0,0,0,0,11429,0,0,0,0,0,0,0],"quality":"More is better","isMain":true,"metricName":"Total length (>= 10000 bp)"},{"values":[0,0,0,0,0,0,0,0,0,0,0,0,0,0],"quality":"More is better","isMain":false,"metricName":"Total length (>= 25000 bp)"},{"values":[0,0,0,0,0,0,0,0,0,0,0,0,0,0],"quality":"More is better","isMain":true,"metricName":"Total length (>= 50000 bp)"},{"values":[6141,10015,6089,6188,6154,2009,11429,6166,6134,6134,6134,2009,6518,6518],"quality":"More is better","isMain":false,"metricName":"N50"},{"values":[6141,10015,6089,6188,6154,2009,11429,6166,6134,6134,6134,2009,6518,6518],"quality":"More is better","isMain":false,"metricName":"N75"},{"values":[1,1,1,1,1,2,1,1,1,1,1,2,1,1],"quality":"Less is better","isMain":false,"metricName":"L50"},{"values":[1,1,1,1,1,2,1,1,1,1,1,2,1,1],"quality":"Less is better","isMain":false,"metricName":"L75"},{"values":["45.19","45.12","45.21","45.27","45.22","45.54","45.25","45.22","45.19","45.19","45.19","45.54","45.04","45.07"],"quality":"Equal","isMain":false,"metricName":"GC (%)"}]],["Predicted genes",[]],["Similarity statistics",[{"values":[0,0,0,0,0,0,0,0,0,0,0,0,0,0],"quality":"Equal","isMain":false,"metricName":"# similar correct contigs"},{"values":[0,0,0,0,0,0,0,0,0,0,0,0,0,0],"quality":"Equal","isMain":false,"metricName":"# similar misassembled blocks"}]],["Reference statistics",[{"values":[6087,6087,6087,6087,6087,6087,6087,6087,6087,6087,6087,6087,6087,6087],"quality":"Equal","isMain":false,"metricName":"Reference length"},{"values":[1,1,1,1,1,1,1,1,1,1,1,1,1,1],"quality":"Equal","isMain":false,"metricName":"Reference fragments"},{"values":["45.18","45.18","45.18","45.18","45.18","45.18","45.18","45.18","45.18","45.18","45.18","45.18","45.18","45.18"],"quality":"Equal","isMain":false,"metricName":"Reference GC (%)"}]]],"referenceName":"NC\_001330","date":"18 July 2018, Wednesday, 16:09:22","order":[0,1,2,3,4,5,6,7,8,9,10,11,12,13],"assembliesNames":["ABySS\_63","ABySS\_127","CLC","IDBA\_UD","MEGAHIT","MetaVelvet","Ray\_Meta","SPAdes","SPAdes\_meta","SPAdes\_sc","SPAdes\_sc\_careful","Velvet","VICUNA","Geneious"]},{"assembliesWithNs":null,"minContig":500,"report":[["Genome statistics",[{"values":["99.610","100.000","56.517","17.267","75.455","100.000","100.000","100.000","100.000","100.000","17.267","100.000","100.000"],"quality":"More is better","isMain":true,"metricName":"Genome fraction (%)"},{"values":["1.004","1.018","1.000","1.026","3.539","3.701","1.014","1.008","1.008","1.008","1.026","1.058","1.577"],"quality":"Less is better","isMain":true,"metricName":"Duplication ratio"},{"values":[5365,5485,2143,939,661,5386,5463,5431,5431,5431,939,5699,3775],"quality":"More is better","isMain":true,"metricName":"Largest alignment"},{"values":[5365,5485,3044,939,14373,19908,5463,5431,5431,5431,939,5699,8415],"quality":"More is better","isMain":true,"metricName":"Total aligned length"},{"values":[5386,5485,901,null,593,19936,5463,5431,5431,5431,null,5699,7442],"quality":"More is better","isMain":false,"metricName":"NG50"},{"values":[5386,5485,null,null,592,19936,5463,5431,5431,5431,null,5699,7442],"quality":"More is better","isMain":false,"metricName":"NG75"},{"values":[5365,5485,2143,939,534,5386,5463,5431,5431,5431,939,5699,3652],"quality":"More is better","isMain":false,"metricName":"NA50"},{"values":[5365,5485,901,939,514,5386,5463,5431,5431,5431,939,5699,3652],"quality":"More is better","isMain":false,"metricName":"NA75"},{"values":[5365,5485,901,null,593,5386,5463,5431,5431,5431,null,5699,3775],"quality":"More is better","isMain":true,"metricName":"NGA50"},{"values":[5365,5485,null,null,592,5386,5463,5431,5431,5431,null,5699,3652],"quality":"More is better","isMain":false,"metricName":"NGA75"},{"values":[1,1,2,null,5,1,1,1,1,1,null,1,1],"quality":"Less is better","isMain":false,"metricName":"LG50"},{"values":[1,1,null,null,7,1,1,1,1,1,null,1,1],"quality":"Less is better","isMain":false,"metricName":"LG75"},{"values":[1,1,1,1,13,2,1,1,1,1,1,1,2],"quality":"Less is better","isMain":false,"metricName":"LA50"},{"values":[1,1,2,1,19,3,1,1,1,1,1,1,2],"quality":"Less is better","isMain":false,"metricName":"LA75"},{"values":[1,1,2,null,5,1,1,1,1,1,null,1,1],"quality":"Less is better","isMain":true,"metricName":"LGA50"},{"values":[1,1,null,null,7,1,1,1,1,1,null,1,2],"quality":"Less is better","isMain":false,"metricName":"LGA75"}]],["Misassemblies",[{"values":[0,0,0,0,1,3,0,0,0,0,0,0,2],"quality":"Less is better","isMain":true,"metricName":"# misassemblies"},{"values":[0,0,0,0,1,3,0,0,0,0,0,0,2],"quality":"Less is better","isMain":false,"metricName":" # relocations"},{"values":[0,0,0,0,0,0,0,0,0,0,0,0,0],"quality":"Less is better","isMain":false,"metricName":" # translocations"},{"values":[0,0,0,0,0,0,0,0,0,0,0,0,0],"quality":"Less is better","isMain":false,"metricName":" # inversions"},{"values":[0,0,0,0,1,1,0,0,0,0,0,0,2],"quality":"Less is better","isMain":false,"metricName":"# misassembled contigs"},{"values":[0,0,0,0,521,19936,0,0,0,0,0,0,7993],"quality":"Less is better","isMain":true,"metricName":"Misassembled contigs length"},{"values":[0,1,0,0,0,0,0,0,0,0,0,1,1],"quality":"Less is better","isMain":false,"metricName":"# local misassemblies"},{"values":[0,0,0,0,0,0,0,0,0,0,0,0,0],"quality":"Less is better","isMain":false,"metricName":"# unaligned mis. contigs"}]],["Unaligned",[{"values":[0,0,0,0,0,0,0,0,0,0,0,0,0],"quality":"Less is better","isMain":false,"metricName":"# fully unaligned contigs"},{"values":[0,0,0,0,0,0,0,0,0,0,0,0,0],"quality":"Less is better","isMain":false,"metricName":"Fully unaligned length"},{"values":[0,0,0,0,0,0,0,0,0,0,0,0,0],"quality":"Less is better","isMain":false,"metricName":"# partially unaligned contigs"},{"values":[0,0,0,0,0,0,0,0,0,0,0,0,0],"quality":"Less is better","isMain":false,"metricName":"Partially unaligned length"}]],["Mismatches",[{"values":[16,16,10,8,29,17,16,16,16,16,8,16,16],"quality":"Less is better","isMain":false,"metricName":"# mismatches"},{"values":[0,0,0,8,0,0,1,1,1,1,8,0,0],"quality":"Less is better","isMain":false,"metricName":"# indels"},{"values":[0,0,0,9,0,0,77,45,45,45,9,0,0],"quality":"Less is better","isMain":false,"metricName":"Indels length"},{"values":["298.23","297.07","328.52","860.22","713.58","315.63","297.07","297.07","297.07","297.07","860.22","297.07","297.07"],"quality":"Less is better","isMain":true,"metricName":"# mismatches per 100 kbp"},{"values":["0.00","0.00","0.00","860.22","0.00","0.00","18.57","18.57","18.57","18.57","860.22","0.00","0.00"],"quality":"Less is better","isMain":true,"metricName":"# indels per 100 kbp"},{"values":[0,0,0,8,0,0,0,0,0,0,8,0,0],"quality":"Less is better","isMain":false,"metricName":" # indels (<= 5 bp)"},{"values":[0,0,0,0,0,0,1,1,1,1,0,0,0],"quality":"Less is better","isMain":false,"metricName":" # indels (> 5 bp)"},{"values":[0,0,0,0,14,0,0,0,0,0,0,0,13],"quality":"Less is better","isMain":false,"metricName":"# N's"},{"values":["0.00","0.00","0.00","0.00","97.34","0.00","0.00","0.00","0.00","0.00","0.00","0.00","153.03"],"quality":"Less is better","isMain":true,"metricName":"# N's per 100 kbp"}]],["Statistics without reference",[{"values":[1,1,2,1,26,1,1,1,1,1,1,1,3],"quality":"Equal","isMain":true,"metricName":"# contigs"},{"values":[1,1,1,0,0,1,1,1,1,1,0,1,1],"quality":"Equal","isMain":false,"metricName":"# contigs (>= 1000 bp)"},{"values":[1,1,0,0,0,1,1,1,1,1,0,1,1],"quality":"Equal","isMain":false,"metricName":"# contigs (>= 5000 bp)"},{"values":[0,0,0,0,0,1,0,0,0,0,0,0,0],"quality":"Equal","isMain":false,"metricName":"# contigs (>= 10000 bp)"},{"values":[0,0,0,0,0,0,0,0,0,0,0,0,0],"quality":"Equal","isMain":false,"metricName":"# contigs (>= 25000 bp)"},{"values":[0,0,0,0,0,0,0,0,0,0,0,0,0],"quality":"Equal","isMain":false,"metricName":"# contigs (>= 50000 bp)"},{"values":[5386,5485,2143,954,661,19936,5463,5431,5431,5431,954,5699,7442],"quality":"More is better","isMain":true,"metricName":"Largest contig"},{"values":[5386,5485,3044,954,14382,19936,5463,5431,5431,5431,954,5699,8495],"quality":"More is better","isMain":true,"metricName":"Total length"},{"values":[5386,5485,2143,0,0,19936,5463,5431,5431,5431,0,5699,7442],"quality":"More is better","isMain":true,"metricName":"Total length (>= 1000 bp)"},{"values":[5386,5485,0,0,0,19936,5463,5431,5431,5431,0,5699,7442],"quality":"More is better","isMain":false,"metricName":"Total length (>= 5000 bp)"},{"values":[0,0,0,0,0,19936,0,0,0,0,0,0,0],"quality":"More is better","isMain":true,"metricName":"Total length (>= 10000 bp)"},{"values":[0,0,0,0,0,0,0,0,0,0,0,0,0],"quality":"More is better","isMain":false,"metricName":"Total length (>= 25000 bp)"},{"values":[0,0,0,0,0,0,0,0,0,0,0,0,0],"quality":"More is better","isMain":true,"metricName":"Total length (>= 50000 bp)"},{"values":[5386,5485,2143,954,534,19936,5463,5431,5431,5431,954,5699,7442],"quality":"More is better","isMain":false,"metricName":"N50"},{"values":[5386,5485,901,954,519,19936,5463,5431,5431,5431,954,5699,7442],"quality":"More is better","isMain":false,"metricName":"N75"},{"values":[1,1,1,1,13,1,1,1,1,1,1,1,1],"quality":"Less is better","isMain":false,"metricName":"L50"},{"values":[1,1,2,1,19,1,1,1,1,1,1,1,1],"quality":"Less is better","isMain":false,"metricName":"L75"},{"values":["44.67","44.76","44.28","46.54","45.62","44.65","44.70","44.67","44.67","44.67","46.54","44.67","44.61"],"quality":"Equal","isMain":false,"metricName":"GC (%)"}]],["Predicted genes",[]],["Similarity statistics",[{"values":[0,0,0,0,0,0,0,0,0,0,0,0,0],"quality":"Equal","isMain":false,"metricName":"# similar correct contigs"},{"values":[0,0,0,0,0,0,0,0,0,0,0,0,0],"quality":"Equal","isMain":false,"metricName":"# similar misassembled blocks"}]],["Reference statistics",[{"values":[5386,5386,5386,5386,5386,5386,5386,5386,5386,5386,5386,5386,5386],"quality":"Equal","isMain":false,"metricName":"Reference length"},{"values":[1,1,1,1,1,1,1,1,1,1,1,1,1],"quality":"Equal","isMain":false,"metricName":"Reference fragments"},{"values":["44.76","44.76","44.76","44.76","44.76","44.76","44.76","44.76","44.76","44.76","44.76","44.76","44.76"],"quality":"Equal","isMain":false,"metricName":"Reference GC (%)"}]]],"referenceName":"NC\_001422","date":"18 July 2018, Wednesday, 16:09:40","order":[0,1,2,3,4,5,6,7,8,9,10,11,12],"assembliesNames":["CLC","IDBA\_UD","MEGAHIT","MetaVelvet","MIRA","Ray\_Meta","SPAdes","SPAdes\_meta","SPAdes\_sc","SPAdes\_sc\_careful","Velvet","VICUNA","Geneious"]},{"assembliesWithNs":null,"minContig":500,"report":[["Genome statistics",[{"values":["100.000","100.000","99.946","100.000","100.000","100.000","100.000","100.000","9.156","100.000","100.000","100.000","100.000","100.000","100.000","100.000"],"quality":"More is better","isMain":true,"metricName":"Genome fraction (%)"},{"values":["1.004","1.003","1.001","1.003","1.007","1.004","1.007","1.033","1.165","1.002","1.002","1.002","1.002","1.004","1.001","1.584"],"quality":"Less is better","isMain":true,"metricName":"Duplication ratio"},{"values":[38543,38545,39170,39290,38545,23050,39447,23054,167,39268,38521,38521,38521,23050,39191,39192],"quality":"More is better","isMain":true,"metricName":"Largest alignment"},{"values":[39313,39325,39170,39290,39473,39306,39447,40498,3587,39268,39277,39277,39277,39306,39191,62047],"quality":"More is better","isMain":true,"metricName":"Total aligned length"},{"values":[38593,38545,39191,39290,38545,23076,39447,40498,743,39268,38521,38521,38521,23076,39243,62073],"quality":"More is better","isMain":false,"metricName":"NG50"},{"values":[38593,38545,39191,39290,38545,15512,39447,40498,null,39268,38521,38521,38521,15512,39243,62073],"quality":"More is better","isMain":false,"metricName":"NG75"},{"values":[38543,38545,39170,39290,38545,23050,39447,23054,null,39268,38521,38521,38521,23050,39191,39192],"quality":"More is better","isMain":false,"metricName":"NA50"},{"values":[38543,38545,39170,39290,38545,15486,39447,17444,null,39268,38521,38521,38521,15486,39191,22855],"quality":"More is better","isMain":false,"metricName":"NA75"},{"values":[38543,38545,39170,39290,38545,23050,39447,23054,null,39268,38521,38521,38521,23050,39191,39192],"quality":"More is better","isMain":true,"metricName":"NGA50"},{"values":[38543,38545,39170,39290,38545,15486,39447,17444,null,39268,38521,38521,38521,15486,39191,39192],"quality":"More is better","isMain":false,"metricName":"NGA75"},{"values":[1,1,1,1,1,1,1,1,6,1,1,1,1,1,1,1],"quality":"Less is better","isMain":false,"metricName":"LG50"},{"values":[1,1,1,1,1,2,1,1,null,1,1,1,1,2,1,1],"quality":"Less is better","isMain":false,"metricName":"LG75"},{"values":[1,1,1,1,1,1,1,1,null,1,1,1,1,1,1,1],"quality":"Less is better","isMain":false,"metricName":"LA50"},{"values":[1,1,1,1,1,2,1,2,null,1,1,1,1,2,1,2],"quality":"Less is better","isMain":false,"metricName":"LA75"},{"values":[1,1,1,1,1,1,1,1,null,1,1,1,1,1,1,1],"quality":"Less is better","isMain":true,"metricName":"LGA50"},{"values":[1,1,1,1,1,2,1,2,null,1,1,1,1,2,1,1],"quality":"Less is better","isMain":false,"metricName":"LGA75"}]],["Misassemblies",[{"values":[0,0,0,0,0,0,0,1,0,0,0,0,0,0,0,1],"quality":"Less is better","isMain":true,"metricName":"# misassemblies"},{"values":[0,0,0,0,0,0,0,1,0,0,0,0,0,0,0,1],"quality":"Less is better","isMain":false,"metricName":" # relocations"},{"values":[0,0,0,0,0,0,0,0,0,0,0,0,0,0,0,0],"quality":"Less is better","isMain":false,"metricName":" # translocations"},{"values":[0,0,0,0,0,0,0,0,0,0,0,0,0,0,0,0],"quality":"Less is better","isMain":false,"metricName":" # inversions"},{"values":[0,0,0,0,0,0,0,1,0,0,0,0,0,0,0,1],"quality":"Less is better","isMain":false,"metricName":"# misassembled contigs"},{"values":[0,0,0,0,0,0,0,40498,0,0,0,0,0,0,0,62073],"quality":"Less is better","isMain":true,"metricName":"Misassembled contigs length"},{"values":[0,0,0,1,0,0,1,0,0,0,0,0,0,0,0,0],"quality":"Less is better","isMain":false,"metricName":"# local misassemblies"},{"values":[0,0,0,0,0,0,0,0,6,0,0,0,0,0,0,0],"quality":"Less is better","isMain":false,"metricName":"# unaligned mis. contigs"}]],["Unaligned",[{"values":[0,0,0,0,0,0,0,0,0,0,0,0,0,0,0,0],"quality":"Less is better","isMain":false,"metricName":"# fully unaligned contigs"},{"values":[0,0,0,0,0,0,0,0,0,0,0,0,0,0,0,0],"quality":"Less is better","isMain":false,"metricName":"Fully unaligned length"},{"values":[0,0,0,0,0,0,0,0,5,0,0,0,0,0,0,0],"quality":"Less is better","isMain":false,"metricName":"# partially unaligned contigs"},{"values":[0,0,0,0,0,0,0,0,15572,0,0,0,0,0,0,0],"quality":"Less is better","isMain":false,"metricName":"Partially unaligned length"}]],["Mismatches",[{"values":[0,0,0,0,0,0,1,0,4,0,0,0,0,0,0,0],"quality":"Less is better","isMain":false,"metricName":"# mismatches"},{"values":[2,2,2,2,2,3,2,2,2,3,2,2,2,3,2,2],"quality":"Less is better","isMain":false,"metricName":"# indels"},{"values":[60,2,2,2,2,3,2,2,5,79,2,2,2,3,2,3],"quality":"Less is better","isMain":false,"metricName":"Indels length"},{"values":["0.00","0.00","0.00","0.00","0.00","0.00","2.55","0.00","111.48","0.00","0.00","0.00","0.00","0.00","0.00","0.00"],"quality":"Less is better","isMain":true,"metricName":"# mismatches per 100 kbp"},{"values":["5.10","5.10","5.11","5.10","5.10","7.66","5.10","5.10","55.74","7.66","5.10","5.10","5.10","7.66","5.10","5.10"],"quality":"Less is better","isMain":true,"metricName":"# indels per 100 kbp"},{"values":[1,2,2,2,2,3,2,2,2,2,2,2,2,3,2,2],"quality":"Less is better","isMain":false,"metricName":" # indels (<= 5 bp)"},{"values":[1,0,0,0,0,0,0,0,0,1,0,0,0,0,0,0],"quality":"Less is better","isMain":false,"metricName":" # indels (> 5 bp)"},{"values":[50,0,0,0,0,0,0,0,8183,0,0,0,0,0,0,6],"quality":"Less is better","isMain":false,"metricName":"# N's"},{"values":["127.02","0.00","0.00","0.00","0.00","0.00","0.00","0.00","41428.72","0.00","0.00","0.00","0.00","0.00","0.00","9.67"],"quality":"Less is better","isMain":true,"metricName":"# N's per 100 kbp"}]],["Statistics without reference",[{"values":[2,2,1,1,2,3,1,1,6,1,2,2,2,3,1,1],"quality":"Equal","isMain":true,"metricName":"# contigs"},{"values":[1,1,1,1,1,2,1,1,5,1,1,1,1,2,1,1],"quality":"Equal","isMain":false,"metricName":"# contigs (>= 1000 bp)"},{"values":[1,1,1,1,1,2,1,1,1,1,1,1,1,2,1,1],"quality":"Equal","isMain":false,"metricName":"# contigs (>= 5000 bp)"},{"values":[1,1,1,1,1,2,1,1,0,1,1,1,1,2,1,1],"quality":"Equal","isMain":false,"metricName":"# contigs (>= 10000 bp)"},{"values":[1,1,1,1,1,0,1,1,0,1,1,1,1,0,1,1],"quality":"Equal","isMain":false,"metricName":"# contigs (>= 25000 bp)"},{"values":[0,0,0,0,0,0,0,0,0,0,0,0,0,0,0,1],"quality":"Equal","isMain":false,"metricName":"# contigs (>= 50000 bp)"},{"values":[38593,38545,39191,39290,38545,23076,39447,40498,7812,39268,38521,38521,38521,23076,39243,62073],"quality":"More is better","isMain":true,"metricName":"Largest contig"},{"values":[39363,39325,39191,39290,39473,39358,39447,40498,19752,39268,39277,39277,39277,39358,39243,62073],"quality":"More is better","isMain":true,"metricName":"Total length"},{"values":[38593,38545,39191,39290,38545,38588,39447,40498,19009,39268,38521,38521,38521,38588,39243,62073],"quality":"More is better","isMain":true,"metricName":"Total length (>= 1000 bp)"},{"values":[38593,38545,39191,39290,38545,38588,39447,40498,7812,39268,38521,38521,38521,38588,39243,62073],"quality":"More is better","isMain":false,"metricName":"Total length (>= 5000 bp)"},{"values":[38593,38545,39191,39290,38545,38588,39447,40498,0,39268,38521,38521,38521,38588,39243,62073],"quality":"More is better","isMain":true,"metricName":"Total length (>= 10000 bp)"},{"values":[38593,38545,39191,39290,38545,0,39447,40498,0,39268,38521,38521,38521,0,39243,62073],"quality":"More is better","isMain":false,"metricName":"Total length (>= 25000 bp)"},{"values":[0,0,0,0,0,0,0,0,0,0,0,0,0,0,0,62073],"quality":"More is better","isMain":true,"metricName":"Total length (>= 50000 bp)"},{"values":[38593,38545,39191,39290,38545,23076,39447,40498,4231,39268,38521,38521,38521,23076,39243,62073],"quality":"More is better","isMain":false,"metricName":"N50"},{"values":[38593,38545,39191,39290,38545,15512,39447,40498,3216,39268,38521,38521,38521,15512,39243,62073],"quality":"More is better","isMain":false,"metricName":"N75"},{"values":[1,1,1,1,1,1,1,1,2,1,1,1,1,1,1,1],"quality":"Less is better","isMain":false,"metricName":"L50"},{"values":[1,1,1,1,1,2,1,1,3,1,1,1,1,2,1,1],"quality":"Less is better","isMain":false,"metricName":"L75"},{"values":["36.50","36.50","36.52","36.50","36.47","36.49","36.57","36.69","34.89","36.50","36.50","36.50","36.50","36.49","36.50","37.18"],"quality":"Equal","isMain":false,"metricName":"GC (%)"}]],["Predicted genes",[]],["Similarity statistics",[{"values":[1,1,0,0,1,1,0,0,0,1,1,1,1,1,0,0],"quality":"Equal","isMain":false,"metricName":"# similar correct contigs"},{"values":[0,0,0,0,0,0,0,0,0,0,0,0,0,0,0,0],"quality":"Equal","isMain":false,"metricName":"# similar misassembled blocks"}]],["Reference statistics",[{"values":[39189,39189,39189,39189,39189,39189,39189,39189,39189,39189,39189,39189,39189,39189,39189,39189],"quality":"Equal","isMain":false,"metricName":"Reference length"},{"values":[1,1,1,1,1,1,1,1,1,1,1,1,1,1,1,1],"quality":"Equal","isMain":false,"metricName":"Reference fragments"},{"values":["36.52","36.52","36.52","36.52","36.52","36.52","36.52","36.52","36.52","36.52","36.52","36.52","36.52","36.52","36.52","36.52"],"quality":"Equal","isMain":false,"metricName":"Reference GC (%)"}]]],"referenceName":"NC\_021790","date":"18 July 2018, Wednesday, 16:09:50","order":[0,1,2,3,4,5,6,7,8,9,10,11,12,13,14,15],"assembliesNames":["ABySS\_63","ABySS\_127","CLC","IDBA\_UD","MEGAHIT","MetaVelvet","MIRA","Ray\_Meta","SOAPdenovo2","SPAdes","SPAdes\_meta","SPAdes\_sc","SPAdes\_sc\_careful","Velvet","VICUNA","Geneious"]},{"assembliesWithNs":null,"minContig":500,"report":[["Genome statistics",[{"values":["100.000","99.971","100.000","100.000","100.000","100.000","100.000","5.406","100.000","100.000","100.000","100.000","100.000","100.000","100.000"],"quality":"More is better","isMain":true,"metricName":"Genome fraction (%)"},{"values":["1.001","1.000","1.001","1.002","1.001","1.002","1.007","0.988","1.001","1.001","1.001","1.001","1.001","1.006","1.581"],"quality":"Less is better","isMain":true,"metricName":"Duplication ratio"},{"values":[71499,71428,71548,71588,65519,71593,71952,610,71526,71485,71485,71485,65519,53925,53925],"quality":"More is better","isMain":true,"metricName":"Largest alignment"},{"values":[71499,71428,71548,71588,71457,71593,71952,3814,71526,71485,71485,71485,71457,71853,112928],"quality":"More is better","isMain":true,"metricName":"Total aligned length"},{"values":[71499,71449,71548,71590,65543,71593,71952,68168,71526,71485,71485,71485,65543,148165,189216],"quality":"More is better","isMain":false,"metricName":"NG50"},{"values":[71499,71449,71548,71590,65543,71593,71952,68168,71526,71485,71485,71485,65543,148165,189216],"quality":"More is better","isMain":false,"metricName":"NG75"},{"values":[71499,71428,71548,71588,65519,71593,71952,null,71526,71485,71485,71485,65519,null,41075],"quality":"More is better","isMain":false,"metricName":"NA50"},{"values":[71499,71428,71548,71588,65519,71593,71952,null,71526,71485,71485,71485,65519,null,null],"quality":"More is better","isMain":false,"metricName":"NA75"},{"values":[71499,71428,71548,71588,65519,71593,71952,null,71526,71485,71485,71485,65519,53925,53925],"quality":"More is better","isMain":true,"metricName":"NGA50"},{"values":[71499,71428,71548,71588,65519,71593,71952,null,71526,71485,71485,71485,65519,53925,53925],"quality":"More is better","isMain":false,"metricName":"NGA75"},{"values":[1,1,1,1,1,1,1,1,1,1,1,1,1,1,1],"quality":"Less is better","isMain":false,"metricName":"LG50"},{"values":[1,1,1,1,1,1,1,1,1,1,1,1,1,1,1],"quality":"Less is better","isMain":false,"metricName":"LG75"},{"values":[1,1,1,1,1,1,1,null,1,1,1,1,1,null,2],"quality":"Less is better","isMain":false,"metricName":"LA50"},{"values":[1,1,1,1,1,1,1,null,1,1,1,1,1,null,null],"quality":"Less is better","isMain":false,"metricName":"LA75"},{"values":[1,1,1,1,1,1,1,null,1,1,1,1,1,1,1],"quality":"Less is better","isMain":true,"metricName":"LGA50"},{"values":[1,1,1,1,1,1,1,null,1,1,1,1,1,1,1],"quality":"Less is better","isMain":false,"metricName":"LGA75"}]],["Misassemblies",[{"values":[0,0,0,0,0,0,0,0,0,0,0,0,0,0,2],"quality":"Less is better","isMain":true,"metricName":"# misassemblies"},{"values":[0,0,0,0,0,0,0,0,0,0,0,0,0,0,2],"quality":"Less is better","isMain":false,"metricName":" # relocations"},{"values":[0,0,0,0,0,0,0,0,0,0,0,0,0,0,0],"quality":"Less is better","isMain":false,"metricName":" # translocations"},{"values":[0,0,0,0,0,0,0,0,0,0,0,0,0,0,0],"quality":"Less is better","isMain":false,"metricName":" # inversions"},{"values":[0,0,0,0,0,0,0,0,0,0,0,0,0,0,1],"quality":"Less is better","isMain":false,"metricName":"# misassembled contigs"},{"values":[0,0,0,0,0,0,0,0,0,0,0,0,0,0,189216],"quality":"Less is better","isMain":true,"metricName":"Misassembled contigs length"},{"values":[0,0,1,1,0,1,1,0,0,0,0,0,0,0,0],"quality":"Less is better","isMain":false,"metricName":"# local misassemblies"},{"values":[0,0,0,0,0,0,0,1,0,0,0,0,0,1,0],"quality":"Less is better","isMain":false,"metricName":"# unaligned mis. contigs"}]],["Unaligned",[{"values":[0,0,0,0,0,0,0,0,0,0,0,0,0,0,0],"quality":"Less is better","isMain":false,"metricName":"# fully unaligned contigs"},{"values":[0,0,0,0,0,0,0,0,0,0,0,0,0,0,0],"quality":"Less is better","isMain":false,"metricName":"Fully unaligned length"},{"values":[0,0,0,0,0,0,0,1,0,0,0,0,0,1,1],"quality":"Less is better","isMain":false,"metricName":"# partially unaligned contigs"},{"values":[0,0,0,0,0,0,0,64961,0,0,0,0,0,76312,76288],"quality":"Less is better","isMain":false,"metricName":"Partially unaligned length"}]],["Mismatches",[{"values":[0,0,0,0,0,0,0,0,0,0,0,0,0,5,5],"quality":"Less is better","isMain":false,"metricName":"# mismatches"},{"values":[7,6,6,5,5,6,6,3,6,7,7,7,5,6,6],"quality":"Less is better","isMain":false,"metricName":"# indels"},{"values":[56,6,6,5,5,6,6,48,82,42,42,42,5,6,6],"quality":"Less is better","isMain":false,"metricName":"Indels length"},{"values":["0.00","0.00","0.00","0.00","0.00","0.00","0.00","0.00","0.00","0.00","0.00","0.00","0.00","7.00","7.00"],"quality":"Less is better","isMain":true,"metricName":"# mismatches per 100 kbp"},{"values":["9.80","8.40","8.40","7.00","7.00","8.40","8.40","77.68","8.40","9.80","9.80","9.80","7.00","8.40","8.40"],"quality":"Less is better","isMain":true,"metricName":"# indels per 100 kbp"},{"values":[6,6,6,5,5,6,6,0,5,6,6,6,5,6,6],"quality":"Less is better","isMain":false,"metricName":" # indels (<= 5 bp)"},{"values":[1,0,0,0,0,0,0,3,1,1,1,1,0,0,0],"quality":"Less is better","isMain":false,"metricName":" # indels (> 5 bp)"},{"values":[0,0,0,0,0,0,0,4544,0,0,0,0,0,0,1],"quality":"Less is better","isMain":false,"metricName":"# N's"},{"values":["0.00","0.00","0.00","0.00","0.00","0.00","0.00","6607.05","0.00","0.00","0.00","0.00","0.00","0.00","0.53"],"quality":"Less is better","isMain":true,"metricName":"# N's per 100 kbp"}]],["Statistics without reference",[{"values":[1,1,1,1,2,1,1,2,1,1,1,1,2,1,1],"quality":"Equal","isMain":true,"metricName":"# contigs"},{"values":[1,1,1,1,2,1,1,1,1,1,1,1,2,1,1],"quality":"Equal","isMain":false,"metricName":"# contigs (>= 1000 bp)"},{"values":[1,1,1,1,2,1,1,1,1,1,1,1,2,1,1],"quality":"Equal","isMain":false,"metricName":"# contigs (>= 5000 bp)"},{"values":[1,1,1,1,1,1,1,1,1,1,1,1,1,1,1],"quality":"Equal","isMain":false,"metricName":"# contigs (>= 10000 bp)"},{"values":[1,1,1,1,1,1,1,1,1,1,1,1,1,1,1],"quality":"Equal","isMain":false,"metricName":"# contigs (>= 25000 bp)"},{"values":[1,1,1,1,1,1,1,1,1,1,1,1,1,1,1],"quality":"Equal","isMain":false,"metricName":"# contigs (>= 50000 bp)"},{"values":[71499,71449,71548,71590,65543,71593,71952,68168,71526,71485,71485,71485,65543,148165,189216],"quality":"More is better","isMain":true,"metricName":"Largest contig"},{"values":[71499,71449,71548,71590,71507,71593,71952,68775,71526,71485,71485,71485,71507,148165,189216],"quality":"More is better","isMain":true,"metricName":"Total length"},{"values":[71499,71449,71548,71590,71507,71593,71952,68168,71526,71485,71485,71485,71507,148165,189216],"quality":"More is better","isMain":true,"metricName":"Total length (>= 1000 bp)"},{"values":[71499,71449,71548,71590,71507,71593,71952,68168,71526,71485,71485,71485,71507,148165,189216],"quality":"More is better","isMain":false,"metricName":"Total length (>= 5000 bp)"},{"values":[71499,71449,71548,71590,65543,71593,71952,68168,71526,71485,71485,71485,65543,148165,189216],"quality":"More is better","isMain":true,"metricName":"Total length (>= 10000 bp)"},{"values":[71499,71449,71548,71590,65543,71593,71952,68168,71526,71485,71485,71485,65543,148165,189216],"quality":"More is better","isMain":false,"metricName":"Total length (>= 25000 bp)"},{"values":[71499,71449,71548,71590,65543,71593,71952,68168,71526,71485,71485,71485,65543,148165,189216],"quality":"More is better","isMain":true,"metricName":"Total length (>= 50000 bp)"},{"values":[71499,71449,71548,71590,65543,71593,71952,68168,71526,71485,71485,71485,65543,148165,189216],"quality":"More is better","isMain":false,"metricName":"N50"},{"values":[71499,71449,71548,71590,65543,71593,71952,68168,71526,71485,71485,71485,65543,148165,189216],"quality":"More is better","isMain":false,"metricName":"N75"},{"values":[1,1,1,1,1,1,1,1,1,1,1,1,1,1,1],"quality":"Less is better","isMain":false,"metricName":"L50"},{"values":[1,1,1,1,1,1,1,1,1,1,1,1,1,1,1],"quality":"Less is better","isMain":false,"metricName":"L75"},{"values":["32.88","32.88","32.87","32.88","32.88","32.88","32.86","32.82","32.88","32.88","32.88","32.88","32.88","31.51","32.02"],"quality":"Equal","isMain":false,"metricName":"GC (%)"}]],["Predicted genes",[]],["Similarity statistics",[{"values":[0,0,0,0,0,0,0,0,0,0,0,0,0,0,0],"quality":"Equal","isMain":false,"metricName":"# similar correct contigs"},{"values":[0,0,0,0,0,0,0,0,0,0,0,0,0,0,0],"quality":"Equal","isMain":false,"metricName":"# similar misassembled blocks"}]],["Reference statistics",[{"values":[71443,71443,71443,71443,71443,71443,71443,71443,71443,71443,71443,71443,71443,71443,71443],"quality":"Equal","isMain":false,"metricName":"Reference length"},{"values":[1,1,1,1,1,1,1,1,1,1,1,1,1,1,1],"quality":"Equal","isMain":false,"metricName":"Reference fragments"},{"values":["32.88","32.88","32.88","32.88","32.88","32.88","32.88","32.88","32.88","32.88","32.88","32.88","32.88","32.88","32.88"],"quality":"Equal","isMain":false,"metricName":"Reference GC (%)"}]]],"referenceName":"NC\_021794","date":"18 July 2018, Wednesday, 16:10:01","order":[0,1,2,3,4,5,6,7,8,9,10,11,12,13,14],"assembliesNames":["ABySS\_63","CLC","IDBA\_UD","MEGAHIT","MetaVelvet","MIRA","Ray\_Meta","SOAPdenovo2","SPAdes","SPAdes\_meta","SPAdes\_sc","SPAdes\_sc\_careful","Velvet","VICUNA","Geneious"]},{"assembliesWithNs":null,"minContig":500,"report":[["Genome statistics",[{"values":["100.000","99.968","99.971","100.000","100.000","99.640","100.000","100.000","15.892","100.000","100.000","100.000","100.000","99.640","100.000","100.000"],"quality":"More is better","isMain":true,"metricName":"Genome fraction (%)"},{"values":["1.000","1.000","1.000","1.001","1.002","1.001","1.033","1.024","0.979","1.002","1.001","1.001","1.001","1.001","1.002","1.531"],"quality":"Less is better","isMain":true,"metricName":"Duplication ratio"},{"values":[72567,72512,72514,72634,72676,28469,72799,71664,3041,70642,70598,70598,70598,28469,72535,72680],"quality":"More is better","isMain":true,"metricName":"Largest alignment"},{"values":[72567,72512,72514,72634,72676,72321,74909,74256,11282,72689,72601,72601,72591,72321,72535,111076],"quality":"More is better","isMain":true,"metricName":"Total aligned length"},{"values":[72567,72512,72535,72634,72676,21286,72806,71664,43763,70642,70598,70598,70598,21286,72708,111083],"quality":"More is better","isMain":false,"metricName":"NG50"},{"values":[72567,72512,72535,72634,72676,16407,72806,71664,19587,70642,70598,70598,70598,16407,72708,111083],"quality":"More is better","isMain":false,"metricName":"NG75"},{"values":[72567,72512,72514,72634,72676,21285,72799,71664,null,70642,70598,70598,70598,21285,72535,72680],"quality":"More is better","isMain":false,"metricName":"NA50"},{"values":[72567,72512,72514,72634,72676,16407,72799,71664,null,70642,70598,70598,70598,16407,72535,38396],"quality":"More is better","isMain":false,"metricName":"NA75"},{"values":[72567,72512,72514,72634,72676,21285,72799,71664,null,70642,70598,70598,70598,21285,72535,72680],"quality":"More is better","isMain":true,"metricName":"NGA50"},{"values":[72567,72512,72514,72634,72676,16407,72799,71664,null,70642,70598,70598,70598,16407,72535,72680],"quality":"More is better","isMain":false,"metricName":"NGA75"},{"values":[1,1,1,1,1,2,1,1,1,1,1,1,1,2,1,1],"quality":"Less is better","isMain":false,"metricName":"LG50"},{"values":[1,1,1,1,1,3,1,1,2,1,1,1,1,3,1,1],"quality":"Less is better","isMain":false,"metricName":"LG75"},{"values":[1,1,1,1,1,2,1,1,null,1,1,1,1,2,1,1],"quality":"Less is better","isMain":false,"metricName":"LA50"},{"values":[1,1,1,1,1,3,1,1,null,1,1,1,1,3,1,2],"quality":"Less is better","isMain":false,"metricName":"LA75"},{"values":[1,1,1,1,1,2,1,1,null,1,1,1,1,2,1,1],"quality":"Less is better","isMain":true,"metricName":"LGA50"},{"values":[1,1,1,1,1,3,1,1,null,1,1,1,1,3,1,1],"quality":"Less is better","isMain":false,"metricName":"LGA75"}]],["Misassemblies",[{"values":[0,0,0,0,0,0,0,0,0,0,0,0,0,0,0,1],"quality":"Less is better","isMain":true,"metricName":"# misassemblies"},{"values":[0,0,0,0,0,0,0,0,0,0,0,0,0,0,0,1],"quality":"Less is better","isMain":false,"metricName":" # relocations"},{"values":[0,0,0,0,0,0,0,0,0,0,0,0,0,0,0,0],"quality":"Less is better","isMain":false,"metricName":" # translocations"},{"values":[0,0,0,0,0,0,0,0,0,0,0,0,0,0,0,0],"quality":"Less is better","isMain":false,"metricName":" # inversions"},{"values":[0,0,0,0,0,0,0,0,0,0,0,0,0,0,0,1],"quality":"Less is better","isMain":false,"metricName":"# misassembled contigs"},{"values":[0,0,0,0,0,0,0,0,0,0,0,0,0,0,0,111083],"quality":"Less is better","isMain":true,"metricName":"Misassembled contigs length"},{"values":[0,0,0,1,1,0,1,0,0,0,0,0,0,0,0,1],"quality":"Less is better","isMain":false,"metricName":"# local misassemblies"},{"values":[0,0,0,0,0,0,0,0,2,0,0,0,0,0,0,0],"quality":"Less is better","isMain":false,"metricName":"# unaligned mis. contigs"}]],["Unaligned",[{"values":[0,0,0,0,0,0,0,0,0,0,0,0,0,0,0,0],"quality":"Less is better","isMain":false,"metricName":"# fully unaligned contigs"},{"values":[0,0,0,0,0,0,0,0,0,0,0,0,0,0,0,0],"quality":"Less is better","isMain":false,"metricName":"Fully unaligned length"},{"values":[0,0,0,0,0,0,0,0,3,0,0,0,0,0,0,0],"quality":"Less is better","isMain":false,"metricName":"# partially unaligned contigs"},{"values":[0,0,0,0,0,0,0,0,53757,0,0,0,0,0,0,0],"quality":"Less is better","isMain":false,"metricName":"Partially unaligned length"}]],["Mismatches",[{"values":[0,0,0,0,0,0,0,0,1,0,0,0,0,0,0,0],"quality":"Less is better","isMain":false,"metricName":"# mismatches"},{"values":[2,1,1,1,1,1,1,1,13,1,1,1,1,1,1,1],"quality":"Less is better","isMain":false,"metricName":"# indels"},{"values":[33,1,1,1,1,1,1,1,251,1,1,1,1,1,1,1],"quality":"Less is better","isMain":false,"metricName":"Indels length"},{"values":["0.00","0.00","0.00","0.00","0.00","0.00","0.00","0.00","8.68","0.00","0.00","0.00","0.00","0.00","0.00","0.00"],"quality":"Less is better","isMain":true,"metricName":"# mismatches per 100 kbp"},{"values":["2.76","1.38","1.38","1.38","1.38","1.38","1.38","1.38","112.78","1.38","1.38","1.38","1.38","1.38","1.38","1.38"],"quality":"Less is better","isMain":true,"metricName":"# indels per 100 kbp"},{"values":[1,1,1,1,1,1,1,1,1,1,1,1,1,1,1,1],"quality":"Less is better","isMain":false,"metricName":" # indels (<= 5 bp)"},{"values":[1,0,0,0,0,0,0,0,12,0,0,0,0,0,0,0],"quality":"Less is better","isMain":false,"metricName":" # indels (> 5 bp)"},{"values":[0,0,0,0,0,0,2,0,5886,0,0,0,0,0,0,25],"quality":"Less is better","isMain":false,"metricName":"# N's"},{"values":["0.00","0.00","0.00","0.00","0.00","0.00","2.67","0.00","9049.95","0.00","0.00","0.00","0.00","0.00","0.00","22.51"],"quality":"Less is better","isMain":true,"metricName":"# N's per 100 kbp"}]],["Statistics without reference",[{"values":[1,1,1,1,1,5,4,2,3,2,2,2,2,5,1,1],"quality":"Equal","isMain":true,"metricName":"# contigs"},{"values":[1,1,1,1,1,5,2,2,3,2,2,2,2,5,1,1],"quality":"Equal","isMain":false,"metricName":"# contigs (>= 1000 bp)"},{"values":[1,1,1,1,1,4,1,1,2,1,1,1,1,4,1,1],"quality":"Equal","isMain":false,"metricName":"# contigs (>= 5000 bp)"},{"values":[1,1,1,1,1,3,1,1,2,1,1,1,1,3,1,1],"quality":"Equal","isMain":false,"metricName":"# contigs (>= 10000 bp)"},{"values":[1,1,1,1,1,1,1,1,1,1,1,1,1,1,1,1],"quality":"Equal","isMain":false,"metricName":"# contigs (>= 25000 bp)"},{"values":[1,1,1,1,1,0,1,1,0,1,1,1,1,0,1,1],"quality":"Equal","isMain":false,"metricName":"# contigs (>= 50000 bp)"},{"values":[72567,72512,72535,72634,72676,28497,72806,71664,43763,70642,70598,70598,70598,28497,72708,111083],"quality":"More is better","isMain":true,"metricName":"Largest contig"},{"values":[72567,72512,72535,72634,72676,72376,74927,74262,65039,72689,72601,72601,72601,72376,72708,111083],"quality":"More is better","isMain":true,"metricName":"Total length"},{"values":[72567,72512,72535,72634,72676,72376,73872,74262,65039,72689,72601,72601,72601,72376,72708,111083],"quality":"More is better","isMain":true,"metricName":"Total length (>= 1000 bp)"},{"values":[72567,72512,72535,72634,72676,71230,72806,71664,63350,70642,70598,70598,70598,71230,72708,111083],"quality":"More is better","isMain":false,"metricName":"Total length (>= 5000 bp)"},{"values":[72567,72512,72535,72634,72676,66190,72806,71664,63350,70642,70598,70598,70598,66190,72708,111083],"quality":"More is better","isMain":true,"metricName":"Total length (>= 10000 bp)"},{"values":[72567,72512,72535,72634,72676,28497,72806,71664,43763,70642,70598,70598,70598,28497,72708,111083],"quality":"More is better","isMain":false,"metricName":"Total length (>= 25000 bp)"},{"values":[72567,72512,72535,72634,72676,0,72806,71664,0,70642,70598,70598,70598,0,72708,111083],"quality":"More is better","isMain":true,"metricName":"Total length (>= 50000 bp)"},{"values":[72567,72512,72535,72634,72676,21286,72806,71664,43763,70642,70598,70598,70598,21286,72708,111083],"quality":"More is better","isMain":false,"metricName":"N50"},{"values":[72567,72512,72535,72634,72676,16407,72806,71664,19587,70642,70598,70598,70598,16407,72708,111083],"quality":"More is better","isMain":false,"metricName":"N75"},{"values":[1,1,1,1,1,2,1,1,1,1,1,1,1,2,1,1],"quality":"Less is better","isMain":false,"metricName":"L50"},{"values":[1,1,1,1,1,3,1,1,2,1,1,1,1,3,1,1],"quality":"Less is better","isMain":false,"metricName":"L75"},{"values":["38.06","38.05","38.06","38.05","38.05","38.07","37.99","37.97","38.09","38.06","38.06","38.06","38.05","38.07","38.06","38.24"],"quality":"Equal","isMain":false,"metricName":"GC (%)"}]],["Predicted genes",[]],["Similarity statistics",[{"values":[0,0,0,0,0,0,0,0,0,0,0,0,0,0,0,0],"quality":"Equal","isMain":false,"metricName":"# similar correct contigs"},{"values":[0,0,0,0,0,0,0,0,0,0,0,0,0,0,0,0],"quality":"Equal","isMain":false,"metricName":"# similar misassembled blocks"}]],["Reference statistics",[{"values":[72534,72534,72534,72534,72534,72534,72534,72534,72534,72534,72534,72534,72534,72534,72534,72534],"quality":"Equal","isMain":false,"metricName":"Reference length"},{"values":[1,1,1,1,1,1,1,1,1,1,1,1,1,1,1,1],"quality":"Equal","isMain":false,"metricName":"Reference fragments"},{"values":["38.06","38.06","38.06","38.06","38.06","38.06","38.06","38.06","38.06","38.06","38.06","38.06","38.06","38.06","38.06","38.06"],"quality":"Equal","isMain":false,"metricName":"Reference GC (%)"}]]],"referenceName":"NC\_021796","date":"18 July 2018, Wednesday, 16:10:12","order":[0,1,2,3,4,5,6,7,8,9,10,11,12,13,14,15],"assembliesNames":["ABySS\_63","ABySS\_127","CLC","IDBA\_UD","MEGAHIT","MetaVelvet","MIRA","Ray\_Meta","SOAPdenovo2","SPAdes","SPAdes\_meta","SPAdes\_sc","SPAdes\_sc\_careful","Velvet","VICUNA","Geneious"]},{"assembliesWithNs":null,"minContig":500,"report":[["Genome statistics",[]],["Misassemblies",[]],["Unaligned",[]],["Mismatches",[{"values":[196,6506,0,10,1450,2480,429],"quality":"Less is better","isMain":false,"metricName":"# N's"},{"values":["4.32","144.24","0.00","0.22","29597.88","51.04","12.30"],"quality":"Less is better","isMain":true,"metricName":"# N's per 100 kbp"}]],["Statistics without reference",[{"values":[60,132,450,94,2,593,14],"quality":"Equal","isMain":true,"metricName":"# contigs"},{"values":[55,123,81,63,2,81,14],"quality":"Equal","isMain":false,"metricName":"# contigs (>= 1000 bp)"},{"values":[40,103,47,39,0,29,13],"quality":"Equal","isMain":false,"metricName":"# contigs (>= 5000 bp)"},{"values":[35,81,40,37,0,24,13],"quality":"Equal","isMain":false,"metricName":"# contigs (>= 10000 bp)"},{"values":[30,48,30,32,0,21,12],"quality":"Equal","isMain":false,"metricName":"# contigs (>= 25000 bp)"},{"values":[21,26,23,24,0,17,10],"quality":"Equal","isMain":false,"metricName":"# contigs (>= 50000 bp)"},{"values":[606442,316102,517832,431858,3844,849779,670062],"quality":"More is better","isMain":true,"metricName":"Largest contig"},{"values":[4534725,4510461,4733009,4546615,4899,4858632,3487048],"quality":"More is better","isMain":true,"metricName":"Total length"},{"values":[4531600,4504640,4502612,4525198,4899,4529210,3487048],"quality":"More is better","isMain":true,"metricName":"Total length (>= 1000 bp)"},{"values":[4492440,4443675,4448606,4483321,0,4454807,3482495],"quality":"More is better","isMain":false,"metricName":"Total length (>= 5000 bp)"},{"values":[4448742,4280662,4400514,4466402,0,4421593,3482495],"quality":"More is better","isMain":true,"metricName":"Total length (>= 10000 bp)"},{"values":[4371566,3709479,4239378,4392588,0,4370104,3463170],"quality":"More is better","isMain":false,"metricName":"Total length (>= 25000 bp)"},{"values":[4004759,2920863,3944386,4058184,0,4182157,3383728],"quality":"More is better","isMain":true,"metricName":"Total length (>= 50000 bp)"},{"values":[242354,79531,152917,234860,3844,288931,448073],"quality":"More is better","isMain":false,"metricName":"N50"},{"values":[105301,35789,88464,111768,3844,119210,245657],"quality":"More is better","isMain":false,"metricName":"N75"},{"values":[6,16,8,8,1,6,3],"quality":"Less is better","isMain":false,"metricName":"L50"},{"values":[14,37,18,16,1,12,6],"quality":"Less is better","isMain":false,"metricName":"L75"},{"values":["34.64","34.60","34.85","34.66","35.84","34.97","34.64"],"quality":"Equal","isMain":false,"metricName":"GC (%)"}]],["Predicted genes",[]],["Similarity statistics",[]],["Reference statistics",[]]],"referenceName":"","date":"18 July 2018, Wednesday, 16:10:19","order":[0,1,2,3,4,5,6],"assembliesNames":["ABySS\_63","ABySS\_127","CLC","MIRA","SOAPdenovo2","SPAdes\_sc","Geneious"]}]

{}

{}

{}

{}

{}

{}

{}

{}

{}

{}

{}

{}

[{"refnames":["KF302033","KF302034","KC821625","KF302036","KF302037","NC\_001330","NC\_021790","NC\_021794","KC821629","NC\_001422","NC\_021796","KF302035"],"coord\_y":[[null,null,1,null,1,1,2,1,1,null,1,1],[null,null,1,null,null,1,2,null,1,null,1,1],[1,1,1,1,1,1,1,1,1,1,1,44],[1,1,1,1,1,1,1,1,1,1,1,1],[1,1,1,1,1,1,2,1,1,2,1,1],[1,1,null,2,1,3,3,2,4,1,5,1],[1,1,8,1,1,null,1,1,5,26,4,14],[1,1,1,1,1,1,1,1,1,1,2,1],[1,1,1,1,1,null,6,2,8,null,3,1],[1,1,1,1,1,1,1,1,1,1,2,1],[1,1,1,1,1,1,2,1,1,1,2,1],[1,1,1,1,1,1,2,1,1,1,2,1],[1,1,1,1,1,1,2,1,1,1,2,1],[1,1,null,2,1,3,3,2,4,1,5,1],[1,1,1,1,2,1,1,1,1,1,1,4],[1,1,2,1,1,1,1,1,1,3,1,3]],"coord\_x":[[0.475,1.475,2.475,3.475,4.475,5.475,6.475,7.475,8.475,9.475,10.475,11.475],[0.5449999999999999,1.545,2.545,3.545,4.545,5.545,6.545,7.545,8.545,9.545,10.545,11.545],[0.615,1.615,2.615,3.615,4.615,5.615,6.615,7.615,8.615,9.615,10.615,11.615],[0.6849999999999999,1.685,2.685,3.685,4.685,5.685,6.685,7.685,8.685,9.685,10.685,11.685],[0.755,1.755,2.755,3.755,4.755,5.755,6.755,7.755,8.755,9.755,10.755,11.755],[0.825,1.825,2.825,3.825,4.825,5.825,6.825,7.825,8.825,9.825,10.825,11.825],[0.895,1.895,2.895,3.895,4.895,5.895,6.895,7.895,8.895,9.895,10.895,11.895],[0.965,1.965,2.965,3.965,4.965,5.965,6.965,7.965,8.965,9.965,10.965,11.965],[1.035,2.035,3.035,4.035,5.035,6.035,7.035,8.035,9.035,10.035,11.035,12.035],[1.105,2.105,3.105,4.105,5.105,6.105,7.105,8.105,9.105,10.105,11.105,12.105],[1.175,2.175,3.175,4.175,5.175,6.175,7.175,8.175,9.175,10.175,11.175,12.175],[1.245,2.245,3.245,4.245,5.245,6.245,7.245,8.245,9.245,10.245,11.245,12.245],[1.315,2.315,3.315,4.315,5.315,6.315,7.315,8.315,9.315,10.315,11.315,12.315],[1.385,2.385,3.385,4.385,5.385,6.385,7.385,8.385,9.385,10.385,11.385,12.385],[1.455,2.455,3.455,4.455,5.455,6.455,7.455,8.455,9.455,10.455,11.455,12.455],[1.525,2.525,3.525,4.525,5.525,6.525,7.525,8.525,9.525,10.525,11.525,12.525]],"filenames":["ABySS\_63","ABySS\_127","CLC","IDBA\_UD","MEGAHIT","MetaVelvet","MIRA","Ray\_Meta","SOAPdenovo2","SPAdes","SPAdes\_meta","SPAdes\_sc","SPAdes\_sc\_careful","Velvet","VICUNA","Geneious"]},
{"refnames":["KF302034","KC821625","NC\_021796","NC\_021794","KC821629","KF302037","NC\_021790","KF302035","KF302033","KF302036","NC\_001330","NC\_001422"],"coord\_y":[[null,76649,72567,71499,54004,44671,38543,35176,null,null,6141,null],[null,76666,72512,null,53956,null,38545,35176,null,null,6089,null],[129415,76567,72514,71428,53995,44551,39170,35176,36771,37711,6089,5365],[129514,76666,72634,71548,54115,44650,39290,35176,36870,37827,6188,5485],[129556,76807,72676,71588,54155,44692,38545,35176,36910,37869,6154,2143],[129477,null,28469,65519,35740,44613,23050,35177,36833,7580,2581,939],[128822,35010,72799,71593,48400,44737,39447,27926,36911,37863,null,661],[129072,77710,71664,71952,53943,45078,23054,35177,37544,38215,5716,5386],[872,73921,3041,610,332,38805,167,284,710,33963,null,null],[128853,76666,70642,71526,54093,44628,39268,35180,36848,37805,6166,5463],[129470,76634,70598,71485,54042,44606,38521,35180,36826,37783,6134,5431],[129470,76653,70598,71485,54042,44606,38521,35176,36826,37783,6134,5431],[129470,76634,70598,71485,54042,44606,38521,35180,36826,37783,6134,5431],[129477,null,28469,65519,35740,44613,23050,35177,36833,7580,2581,939],[129712,77070,72535,53925,54016,40569,39191,15625,37057,38007,6508,5699],[128822,71811,72680,53925,54016,29763,39192,35176,35127,29512,6508,3775]],"coord\_x":[[0.475,1.475,2.475,3.475,4.475,5.475,6.475,7.475,8.475,9.475,10.475,11.475],[0.5449999999999999,1.545,2.545,3.545,4.545,5.545,6.545,7.545,8.545,9.545,10.545,11.545],[0.615,1.615,2.615,3.615,4.615,5.615,6.615,7.615,8.615,9.615,10.615,11.615],[0.6849999999999999,1.685,2.685,3.685,4.685,5.685,6.685,7.685,8.685,9.685,10.685,11.685],[0.755,1.755,2.755,3.755,4.755,5.755,6.755,7.755,8.755,9.755,10.755,11.755],[0.825,1.825,2.825,3.825,4.825,5.825,6.825,7.825,8.825,9.825,10.825,11.825],[0.895,1.895,2.895,3.895,4.895,5.895,6.895,7.895,8.895,9.895,10.895,11.895],[0.965,1.965,2.965,3.965,4.965,5.965,6.965,7.965,8.965,9.965,10.965,11.965],[1.035,2.035,3.035,4.035,5.035,6.035,7.035,8.035,9.035,10.035,11.035,12.035],[1.105,2.105,3.105,4.105,5.105,6.105,7.105,8.105,9.105,10.105,11.105,12.105],[1.175,2.175,3.175,4.175,5.175,6.175,7.175,8.175,9.175,10.175,11.175,12.175],[1.245,2.245,3.245,4.245,5.245,6.245,7.245,8.245,9.245,10.245,11.245,12.245],[1.315,2.315,3.315,4.315,5.315,6.315,7.315,8.315,9.315,10.315,11.315,12.315],[1.385,2.385,3.385,4.385,5.385,6.385,7.385,8.385,9.385,10.385,11.385,12.385],[1.455,2.455,3.455,4.455,5.455,6.455,7.455,8.455,9.455,10.455,11.455,12.455],[1.525,2.525,3.525,4.525,5.525,6.525,7.525,8.525,9.525,10.525,11.525,12.525]],"filenames":["ABySS\_63","ABySS\_127","CLC","IDBA\_UD","MEGAHIT","MetaVelvet","MIRA","Ray\_Meta","SOAPdenovo2","SPAdes","SPAdes\_meta","SPAdes\_sc","SPAdes\_sc\_careful","Velvet","VICUNA","Geneious"]},
{"refnames":["KF302034","NC\_021796","KC821625","NC\_021794","KC821629","KF302037","NC\_021790","KF302035","KF302036","KF302033","NC\_001330","NC\_001422"],"coord\_y":[[null,72567,76649,71499,54004,44671,39313,35176,null,null,6141,null],[null,72512,76666,null,53956,null,39325,35176,null,null,10015,null],[129415,72514,76567,71428,53995,44551,39170,39179,37711,36771,6089,5365],[129514,72634,76666,71548,54115,44650,39290,35176,37827,36870,6188,5485],[129556,72676,76807,71588,54155,44692,39473,35176,37869,36910,6154,3044],[129477,72321,null,71457,53946,44613,39306,35177,14208,36833,5494,939],[128822,74909,83080,71593,53062,44737,39447,36988,37863,36911,null,14373],[129072,74256,77710,71952,55233,45078,40498,35177,38215,37544,11429,19908],[11123,11282,73921,3814,1881,38805,3587,5336,33963,2581,null,null],[128853,72689,76666,71526,54093,44628,39268,35180,37805,36848,6166,5463],[129470,72601,76634,71485,54042,44606,39277,35180,37783,36826,6134,5431],[129470,72601,76653,71485,54042,44606,39277,35176,37783,36826,6134,5431],[129470,72591,76634,71485,54042,44606,39277,35180,37783,36826,6134,5431],[129477,72321,null,71457,53946,44613,39306,35177,14208,36833,5494,939],[129712,72535,77070,71853,54016,45152,39191,36088,38007,37057,6508,5699],[132154,111076,78464,112928,54016,53369,62047,35615,51269,40038,6508,8415]],"coord\_x":[[0.475,1.475,2.475,3.475,4.475,5.475,6.475,7.475,8.475,9.475,10.475,11.475],[0.5449999999999999,1.545,2.545,3.545,4.545,5.545,6.545,7.545,8.545,9.545,10.545,11.545],[0.615,1.615,2.615,3.615,4.615,5.615,6.615,7.615,8.615,9.615,10.615,11.615],[0.6849999999999999,1.685,2.685,3.685,4.685,5.685,6.685,7.685,8.685,9.685,10.685,11.685],[0.755,1.755,2.755,3.755,4.755,5.755,6.755,7.755,8.755,9.755,10.755,11.755],[0.825,1.825,2.825,3.825,4.825,5.825,6.825,7.825,8.825,9.825,10.825,11.825],[0.895,1.895,2.895,3.895,4.895,5.895,6.895,7.895,8.895,9.895,10.895,11.895],[0.965,1.965,2.965,3.965,4.965,5.965,6.965,7.965,8.965,9.965,10.965,11.965],[1.035,2.035,3.035,4.035,5.035,6.035,7.035,8.035,9.035,10.035,11.035,12.035],[1.105,2.105,3.105,4.105,5.105,6.105,7.105,8.105,9.105,10.105,11.105,12.105],[1.175,2.175,3.175,4.175,5.175,6.175,7.175,8.175,9.175,10.175,11.175,12.175],[1.245,2.245,3.245,4.245,5.245,6.245,7.245,8.245,9.245,10.245,11.245,12.245],[1.315,2.315,3.315,4.315,5.315,6.315,7.315,8.315,9.315,10.315,11.315,12.315],[1.385,2.385,3.385,4.385,5.385,6.385,7.385,8.385,9.385,10.385,11.385,12.385],[1.455,2.455,3.455,4.455,5.455,6.455,7.455,8.455,9.455,10.455,11.455,12.455],[1.525,2.525,3.525,4.525,5.525,6.525,7.525,8.525,9.525,10.525,11.525,12.525]],"filenames":["ABySS\_63","ABySS\_127","CLC","IDBA\_UD","MEGAHIT","MetaVelvet","MIRA","Ray\_Meta","SOAPdenovo2","SPAdes","SPAdes\_meta","SPAdes\_sc","SPAdes\_sc\_careful","Velvet","VICUNA","Geneious"]},
{"refnames":["KC821625","KC821629","KF302033","KF302034","KF302035","KF302036","KF302037","NC\_001330","NC\_001422","NC\_021790","NC\_021794","NC\_021796"],"coord\_y":[[0,0,null,null,0,null,0,0,null,0,0,0],[0,0,null,null,0,null,null,2,null,0,null,0],[0,0,0,0,0,0,0,0,0,0,0,0],[0,0,0,0,0,0,0,0,0,0,0,0],[0,0,0,0,0,0,0,0,0,0,0,0],[null,0,0,0,0,0,0,0,0,0,0,0],[0,0,0,0,0,0,0,null,1,0,0,0],[0,1,0,0,0,0,0,1,3,1,0,0],[0,0,0,0,0,0,0,null,null,0,0,0],[0,0,0,0,0,0,0,0,0,0,0,0],[0,0,0,0,0,0,0,0,0,0,0,0],[0,0,0,0,0,0,0,0,0,0,0,0],[0,0,0,0,0,0,0,0,0,0,0,0],[null,0,0,0,0,0,0,0,0,0,0,0],[0,0,0,0,0,0,0,0,0,0,0,0],[0,0,1,1,0,1,1,0,2,1,2,1]],"coord\_x":[[0.475,1.475,2.475,3.475,4.475,5.475,6.475,7.475,8.475,9.475,10.475,11.475],[0.5449999999999999,1.545,2.545,3.545,4.545,5.545,6.545,7.545,8.545,9.545,10.545,11.545],[0.615,1.615,2.615,3.615,4.615,5.615,6.615,7.615,8.615,9.615,10.615,11.615],[0.6849999999999999,1.685,2.685,3.685,4.685,5.685,6.685,7.685,8.685,9.685,10.685,11.685],[0.755,1.755,2.755,3.755,4.755,5.755,6.755,7.755,8.755,9.755,10.755,11.755],[0.825,1.825,2.825,3.825,4.825,5.825,6.825,7.825,8.825,9.825,10.825,11.825],[0.895,1.895,2.895,3.895,4.895,5.895,6.895,7.895,8.895,9.895,10.895,11.895],[0.965,1.965,2.965,3.965,4.965,5.965,6.965,7.965,8.965,9.965,10.965,11.965],[1.035,2.035,3.035,4.035,5.035,6.035,7.035,8.035,9.035,10.035,11.035,12.035],[1.105,2.105,3.105,4.105,5.105,6.105,7.105,8.105,9.105,10.105,11.105,12.105],[1.175,2.175,3.175,4.175,5.175,6.175,7.175,8.175,9.175,10.175,11.175,12.175],[1.245,2.245,3.245,4.245,5.245,6.245,7.245,8.245,9.245,10.245,11.245,12.245],[1.315,2.315,3.315,4.315,5.315,6.315,7.315,8.315,9.315,10.315,11.315,12.315],[1.385,2.385,3.385,4.385,5.385,6.385,7.385,8.385,9.385,10.385,11.385,12.385],[1.455,2.455,3.455,4.455,5.455,6.455,7.455,8.455,9.455,10.455,11.455,12.455],[1.525,2.525,3.525,4.525,5.525,6.525,7.525,8.525,9.525,10.525,11.525,12.525]],"filenames":["ABySS\_63","ABySS\_127","CLC","IDBA\_UD","MEGAHIT","MetaVelvet","MIRA","Ray\_Meta","SOAPdenovo2","SPAdes","SPAdes\_meta","SPAdes\_sc","SPAdes\_sc\_careful","Velvet","VICUNA","Geneious"]},
{"refnames":["KC821625","KF302035","NC\_001330","NC\_001422","KF302033","KF302036","KF302037","KC821629","NC\_021790","NC\_021796","KF302034","NC\_021794"],"coord\_y":[[0,0,0,null,null,null,0,0,0,0,null,0],[0,0,10015,null,null,null,null,0,0,0,null,null],[0,0,0,0,0,0,0,0,0,0,0,0],[0,0,0,0,0,0,0,0,0,0,0,0],[0,0,0,0,0,0,0,0,0,0,0,0],[null,0,0,0,0,0,0,0,0,0,0,0],[0,0,null,521,0,0,0,0,0,0,0,0],[0,0,11429,19936,0,0,0,55233,40498,0,0,0],[0,0,null,null,0,0,0,0,0,0,0,0],[0,0,0,0,0,0,0,0,0,0,0,0],[0,0,0,0,0,0,0,0,0,0,0,0],[0,0,0,0,0,0,0,0,0,0,0,0],[0,0,0,0,0,0,0,0,0,0,0,0],[null,0,0,0,0,0,0,0,0,0,0,0],[0,0,0,0,0,0,0,0,0,0,0,0],[0,0,0,7993,40038,51269,53369,0,62073,111083,132154,189216]],"coord\_x":[[0.475,1.475,2.475,3.475,4.475,5.475,6.475,7.475,8.475,9.475,10.475,11.475],[0.5449999999999999,1.545,2.545,3.545,4.545,5.545,6.545,7.545,8.545,9.545,10.545,11.545],[0.615,1.615,2.615,3.615,4.615,5.615,6.615,7.615,8.615,9.615,10.615,11.615],[0.6849999999999999,1.685,2.685,3.685,4.685,5.685,6.685,7.685,8.685,9.685,10.685,11.685],[0.755,1.755,2.755,3.755,4.755,5.755,6.755,7.755,8.755,9.755,10.755,11.755],[0.825,1.825,2.825,3.825,4.825,5.825,6.825,7.825,8.825,9.825,10.825,11.825],[0.895,1.895,2.895,3.895,4.895,5.895,6.895,7.895,8.895,9.895,10.895,11.895],[0.965,1.965,2.965,3.965,4.965,5.965,6.965,7.965,8.965,9.965,10.965,11.965],[1.035,2.035,3.035,4.035,5.035,6.035,7.035,8.035,9.035,10.035,11.035,12.035],[1.105,2.105,3.105,4.105,5.105,6.105,7.105,8.105,9.105,10.105,11.105,12.105],[1.175,2.175,3.175,4.175,5.175,6.175,7.175,8.175,9.175,10.175,11.175,12.175],[1.245,2.245,3.245,4.245,5.245,6.245,7.245,8.245,9.245,10.245,11.245,12.245],[1.315,2.315,3.315,4.315,5.315,6.315,7.315,8.315,9.315,10.315,11.315,12.315],[1.385,2.385,3.385,4.385,5.385,6.385,7.385,8.385,9.385,10.385,11.385,12.385],[1.455,2.455,3.455,4.455,5.455,6.455,7.455,8.455,9.455,10.455,11.455,12.455],[1.525,2.525,3.525,4.525,5.525,6.525,7.525,8.525,9.525,10.525,11.525,12.525]],"filenames":["ABySS\_63","ABySS\_127","CLC","IDBA\_UD","MEGAHIT","MetaVelvet","MIRA","Ray\_Meta","SOAPdenovo2","SPAdes","SPAdes\_meta","SPAdes\_sc","SPAdes\_sc\_careful","Velvet","VICUNA","Geneious"]},
{"refnames":["KF302036","NC\_021796","KF302034","NC\_021794","KF302035","KC821629","KF302033","KF302037","KC821625","NC\_021790","NC\_001330","NC\_001422"],"coord\_y":[[null,0.0,null,0.0,0.0,0.0,null,0.0,1.3,0.0,147.86,null],[null,0.0,null,null,0.0,0.0,null,null,1.3,0.0,147.86,null],[2.65,0.0,0.0,0.0,2.84,0.0,2.72,0.0,1.31,0.0,147.86,298.23],[0.0,0.0,0.0,0.0,0.0,0.0,2.72,0.0,1.3,0.0,147.86,297.07],[0.0,0.0,0.0,0.0,0.0,0.0,2.72,0.0,1.3,0.0,147.86,328.52],[0.0,0.0,0.0,0.0,0.0,0.0,2.72,0.0,null,0.0,184.16,860.22],[0.0,0.0,0.0,0.0,2.84,3.8,2.72,0.0,1.3,2.55,null,713.58],[0.0,0.0,0.0,0.0,0.0,0.0,2.72,0.0,1.3,0.0,147.86,315.63],[0.0,8.68,8.91,0.0,37.51,53.13,37.92,78.76,0.0,111.48,null,null],[0.0,0.0,0.0,0.0,0.0,0.0,2.72,0.0,1.3,0.0,147.86,297.07],[0.0,0.0,0.0,0.0,0.0,0.0,2.72,0.0,1.31,0.0,147.86,297.07],[0.0,0.0,0.0,0.0,0.0,0.0,2.72,0.0,1.3,0.0,147.86,297.07],[0.0,0.0,0.0,0.0,0.0,1.85,2.72,0.0,1.31,0.0,147.86,297.07],[0.0,0.0,0.0,0.0,0.0,0.0,2.72,0.0,null,0.0,184.16,860.22],[0.0,0.0,0.0,7.0,0.0,0.0,2.72,0.0,33.91,0.0,147.86,297.07],[0.0,0.0,0.0,7.0,0.0,0.0,2.72,0.0,33.91,0.0,147.86,297.07]],"coord\_x":[[0.475,1.475,2.475,3.475,4.475,5.475,6.475,7.475,8.475,9.475,10.475,11.475],[0.5449999999999999,1.545,2.545,3.545,4.545,5.545,6.545,7.545,8.545,9.545,10.545,11.545],[0.615,1.615,2.615,3.615,4.615,5.615,6.615,7.615,8.615,9.615,10.615,11.615],[0.6849999999999999,1.685,2.685,3.685,4.685,5.685,6.685,7.685,8.685,9.685,10.685,11.685],[0.755,1.755,2.755,3.755,4.755,5.755,6.755,7.755,8.755,9.755,10.755,11.755],[0.825,1.825,2.825,3.825,4.825,5.825,6.825,7.825,8.825,9.825,10.825,11.825],[0.895,1.895,2.895,3.895,4.895,5.895,6.895,7.895,8.895,9.895,10.895,11.895],[0.965,1.965,2.965,3.965,4.965,5.965,6.965,7.965,8.965,9.965,10.965,11.965],[1.035,2.035,3.035,4.035,5.035,6.035,7.035,8.035,9.035,10.035,11.035,12.035],[1.105,2.105,3.105,4.105,5.105,6.105,7.105,8.105,9.105,10.105,11.105,12.105],[1.175,2.175,3.175,4.175,5.175,6.175,7.175,8.175,9.175,10.175,11.175,12.175],[1.245,2.245,3.245,4.245,5.245,6.245,7.245,8.245,9.245,10.245,11.245,12.245],[1.315,2.315,3.315,4.315,5.315,6.315,7.315,8.315,9.315,10.315,11.315,12.315],[1.385,2.385,3.385,4.385,5.385,6.385,7.385,8.385,9.385,10.385,11.385,12.385],[1.455,2.455,3.455,4.455,5.455,6.455,7.455,8.455,9.455,10.455,11.455,12.455],[1.525,2.525,3.525,4.525,5.525,6.525,7.525,8.525,9.525,10.525,11.525,12.525]],"filenames":["ABySS\_63","ABySS\_127","CLC","IDBA\_UD","MEGAHIT","MetaVelvet","MIRA","Ray\_Meta","SOAPdenovo2","SPAdes","SPAdes\_meta","SPAdes\_sc","SPAdes\_sc\_careful","Velvet","VICUNA","Geneious"]},
{"refnames":["KC821625","KF302034","NC\_021796","KF302036","NC\_021790","KF302035","KF302033","KF302037","NC\_021794","KC821629","NC\_001330","NC\_001422"],"coord\_y":[[0.0,null,2.76,null,5.1,5.69,null,0.0,9.8,12.98,49.29,null],[0.0,null,1.38,null,5.1,5.69,null,null,null,7.41,32.86,null],[0.0,0.77,1.38,0.0,5.11,5.68,5.44,0.0,8.4,7.41,32.86,0.0],[0.0,0.0,1.38,0.0,5.1,5.69,5.44,0.0,8.4,5.55,32.86,0.0],[0.0,0.0,1.38,0.0,5.1,5.69,5.44,0.0,7.0,7.41,49.29,0.0],[null,0.77,1.38,0.0,7.66,5.69,5.44,0.0,7.0,14.87,36.83,860.22],[2.61,0.0,1.38,0.0,5.1,5.68,5.44,0.0,8.4,11.41,null,0.0],[5.22,0.0,1.38,0.0,5.1,5.69,5.44,0.0,8.4,7.41,32.86,0.0],[27.12,71.3,112.78,136.65,55.74,56.26,75.84,149.9,77.68,53.13,null,null],[0.0,0.0,1.38,0.0,7.66,5.69,2.72,0.0,8.4,9.26,49.29,18.57],[1.31,0.77,1.38,0.0,5.1,5.69,5.44,0.0,9.8,11.11,49.29,18.57],[0.0,0.77,1.38,0.0,5.1,5.69,5.44,0.0,9.8,11.11,49.29,18.57],[1.31,0.77,1.38,0.0,5.1,5.69,5.44,0.0,9.8,11.11,49.29,18.57],[null,0.77,1.38,0.0,7.66,5.69,5.44,0.0,7.0,14.87,36.83,860.22],[0.0,0.0,1.38,0.0,5.1,5.69,5.44,0.0,8.4,7.41,32.86,0.0],[2.61,0.0,1.38,0.0,5.1,5.69,5.44,0.0,8.4,7.41,32.86,0.0]],"coord\_x":[[0.475,1.475,2.475,3.475,4.475,5.475,6.475,7.475,8.475,9.475,10.475,11.475],[0.5449999999999999,1.545,2.545,3.545,4.545,5.545,6.545,7.545,8.545,9.545,10.545,11.545],[0.615,1.615,2.615,3.615,4.615,5.615,6.615,7.615,8.615,9.615,10.615,11.615],[0.6849999999999999,1.685,2.685,3.685,4.685,5.685,6.685,7.685,8.685,9.685,10.685,11.685],[0.755,1.755,2.755,3.755,4.755,5.755,6.755,7.755,8.755,9.755,10.755,11.755],[0.825,1.825,2.825,3.825,4.825,5.825,6.825,7.825,8.825,9.825,10.825,11.825],[0.895,1.895,2.895,3.895,4.895,5.895,6.895,7.895,8.895,9.895,10.895,11.895],[0.965,1.965,2.965,3.965,4.965,5.965,6.965,7.965,8.965,9.965,10.965,11.965],[1.035,2.035,3.035,4.035,5.035,6.035,7.035,8.035,9.035,10.035,11.035,12.035],[1.105,2.105,3.105,4.105,5.105,6.105,7.105,8.105,9.105,10.105,11.105,12.105],[1.175,2.175,3.175,4.175,5.175,6.175,7.175,8.175,9.175,10.175,11.175,12.175],[1.245,2.245,3.245,4.245,5.245,6.245,7.245,8.245,9.245,10.245,11.245,12.245],[1.315,2.315,3.315,4.315,5.315,6.315,7.315,8.315,9.315,10.315,11.315,12.315],[1.385,2.385,3.385,4.385,5.385,6.385,7.385,8.385,9.385,10.385,11.385,12.385],[1.455,2.455,3.455,4.455,5.455,6.455,7.455,8.455,9.455,10.455,11.455,12.455],[1.525,2.525,3.525,4.525,5.525,6.525,7.525,8.525,9.525,10.525,11.525,12.525]],"filenames":["ABySS\_63","ABySS\_127","CLC","IDBA\_UD","MEGAHIT","MetaVelvet","MIRA","Ray\_Meta","SOAPdenovo2","SPAdes","SPAdes\_meta","SPAdes\_sc","SPAdes\_sc\_careful","Velvet","VICUNA","Geneious"]},
{"refnames":["NC\_001330","NC\_001422","KC821625","KF302036","KF302037","NC\_021794","NC\_021796","KF302033","KF302034","KF302035","NC\_021790","KC821629"],"coord\_y":[[0.0,null,0.0,null,0.0,0.0,0.0,null,null,0.0,127.02,461.22],[0.0,null,0.0,null,null,null,0.0,null,null,0.0,0.0,92.58],[0.0,0.0,0.0,0.0,0.0,0.0,0.0,0.0,0.0,0.0,0.0,0.0],[0.0,0.0,0.0,0.0,0.0,0.0,0.0,0.0,0.0,0.0,0.0,0.0],[0.0,0.0,0.0,0.0,0.0,0.0,0.0,0.0,0.0,0.0,0.0,0.0],[0.0,0.0,null,0.0,0.0,0.0,0.0,0.0,0.0,0.0,0.0,0.0],[null,97.34,0.0,0.0,0.0,0.0,2.67,0.0,0.0,16.22,0.0,1.88],[0.0,0.0,0.0,0.0,0.0,0.0,0.0,0.0,0.0,0.0,0.0,0.0],[null,null,1642.76,2703.57,3242.26,6607.05,9049.95,13750.33,18162.13,28790.62,41428.72,44472.51],[0.0,0.0,0.0,0.0,0.0,0.0,0.0,0.0,0.0,0.0,0.0,0.0],[0.0,0.0,0.0,0.0,0.0,0.0,0.0,0.0,0.0,0.0,0.0,0.0],[0.0,0.0,0.0,0.0,0.0,0.0,0.0,0.0,0.0,0.0,0.0,0.0],[0.0,0.0,0.0,0.0,0.0,0.0,0.0,0.0,0.0,0.0,0.0,0.0],[0.0,0.0,null,0.0,0.0,0.0,0.0,0.0,0.0,0.0,0.0,0.0],[0.0,0.0,0.0,0.0,0.0,0.0,0.0,0.0,0.0,0.0,0.0,0.0],[184.11,153.03,0.52,0.0,0.0,0.53,22.51,0.0,0.0,156.37,9.67,79.18]],"coord\_x":[[0.475,1.475,2.475,3.475,4.475,5.475,6.475,7.475,8.475,9.475,10.475,11.475],[0.5449999999999999,1.545,2.545,3.545,4.545,5.545,6.545,7.545,8.545,9.545,10.545,11.545],[0.615,1.615,2.615,3.615,4.615,5.615,6.615,7.615,8.615,9.615,10.615,11.615],[0.6849999999999999,1.685,2.685,3.685,4.685,5.685,6.685,7.685,8.685,9.685,10.685,11.685],[0.755,1.755,2.755,3.755,4.755,5.755,6.755,7.755,8.755,9.755,10.755,11.755],[0.825,1.825,2.825,3.825,4.825,5.825,6.825,7.825,8.825,9.825,10.825,11.825],[0.895,1.895,2.895,3.895,4.895,5.895,6.895,7.895,8.895,9.895,10.895,11.895],[0.965,1.965,2.965,3.965,4.965,5.965,6.965,7.965,8.965,9.965,10.965,11.965],[1.035,2.035,3.035,4.035,5.035,6.035,7.035,8.035,9.035,10.035,11.035,12.035],[1.105,2.105,3.105,4.105,5.105,6.105,7.105,8.105,9.105,10.105,11.105,12.105],[1.175,2.175,3.175,4.175,5.175,6.175,7.175,8.175,9.175,10.175,11.175,12.175],[1.245,2.245,3.245,4.245,5.245,6.245,7.245,8.245,9.245,10.245,11.245,12.245],[1.315,2.315,3.315,4.315,5.315,6.315,7.315,8.315,9.315,10.315,11.315,12.315],[1.385,2.385,3.385,4.385,5.385,6.385,7.385,8.385,9.385,10.385,11.385,12.385],[1.455,2.455,3.455,4.455,5.455,6.455,7.455,8.455,9.455,10.455,11.455,12.455],[1.525,2.525,3.525,4.525,5.525,6.525,7.525,8.525,9.525,10.525,11.525,12.525]],"filenames":["ABySS\_63","ABySS\_127","CLC","IDBA\_UD","MEGAHIT","MetaVelvet","MIRA","Ray\_Meta","SOAPdenovo2","SPAdes","SPAdes\_meta","SPAdes\_sc","SPAdes\_sc\_careful","Velvet","VICUNA","Geneious"]},
{"refnames":["NC\_021796","NC\_021790","KF302035","KC821629","KF302037","NC\_021794","KC821625","NC\_001330","KF302034","KF302033","KF302036","NC\_001422"],"coord\_y":[[100.0,100.0,99.558,99.88,98.925,100.0,99.978,100.0,null,null,null,null],[99.968,100.0,99.558,99.88,null,null,100.0,100.0,null,null,null,null],[99.971,99.946,99.59,99.961,98.925,99.971,99.871,100.0,99.981,98.812,98.699,99.61],[100.0,100.0,99.558,100.0,98.925,100.0,100.0,100.0,99.981,98.812,98.744,100.0],[100.0,100.0,99.558,100.0,98.925,100.0,100.0,100.0,99.981,98.812,98.744,56.517],[99.64,100.0,99.561,99.583,98.925,100.0,null,89.207,99.981,98.812,37.186,17.267],[100.0,100.0,99.578,97.328,98.925,100.0,100.0,null,99.523,98.812,98.744,75.455],[100.0,100.0,99.561,100.0,98.925,100.0,100.0,100.0,99.716,98.812,98.744,100.0],[15.892,9.156,15.092,3.484,87.396,5.406,96.204,null,8.668,7.087,90.018,null],[100.0,100.0,99.57,100.0,98.925,100.0,100.0,100.0,99.547,98.812,98.744,100.0],[100.0,100.0,99.57,100.0,98.925,100.0,99.911,100.0,99.981,98.812,98.744,100.0],[100.0,100.0,99.558,100.0,98.925,100.0,99.983,100.0,99.981,98.812,98.744,100.0],[100.0,100.0,99.57,100.0,98.925,100.0,99.911,100.0,99.981,98.812,98.744,100.0],[99.64,100.0,99.561,99.583,98.925,100.0,null,89.207,99.981,98.812,37.186,17.267],[100.0,100.0,99.558,100.0,98.925,100.0,100.0,100.0,99.981,98.812,98.744,100.0],[100.0,100.0,99.561,100.0,98.925,100.0,100.0,100.0,99.981,98.812,98.744,100.0]],"coord\_x":[[0.475,1.475,2.475,3.475,4.475,5.475,6.475,7.475,8.475,9.475,10.475,11.475],[0.5449999999999999,1.545,2.545,3.545,4.545,5.545,6.545,7.545,8.545,9.545,10.545,11.545],[0.615,1.615,2.615,3.615,4.615,5.615,6.615,7.615,8.615,9.615,10.615,11.615],[0.6849999999999999,1.685,2.685,3.685,4.685,5.685,6.685,7.685,8.685,9.685,10.685,11.685],[0.755,1.755,2.755,3.755,4.755,5.755,6.755,7.755,8.755,9.755,10.755,11.755],[0.825,1.825,2.825,3.825,4.825,5.825,6.825,7.825,8.825,9.825,10.825,11.825],[0.895,1.895,2.895,3.895,4.895,5.895,6.895,7.895,8.895,9.895,10.895,11.895],[0.965,1.965,2.965,3.965,4.965,5.965,6.965,7.965,8.965,9.965,10.965,11.965],[1.035,2.035,3.035,4.035,5.035,6.035,7.035,8.035,9.035,10.035,11.035,12.035],[1.105,2.105,3.105,4.105,5.105,6.105,7.105,8.105,9.105,10.105,11.105,12.105],[1.175,2.175,3.175,4.175,5.175,6.175,7.175,8.175,9.175,10.175,11.175,12.175],[1.245,2.245,3.245,4.245,5.245,6.245,7.245,8.245,9.245,10.245,11.245,12.245],[1.315,2.315,3.315,4.315,5.315,6.315,7.315,8.315,9.315,10.315,11.315,12.315],[1.385,2.385,3.385,4.385,5.385,6.385,7.385,8.385,9.385,10.385,11.385,12.385],[1.455,2.455,3.455,4.455,5.455,6.455,7.455,8.455,9.455,10.455,11.455,12.455],[1.525,2.525,3.525,4.525,5.525,6.525,7.525,8.525,9.525,10.525,11.525,12.525]],"filenames":["ABySS\_63","ABySS\_127","CLC","IDBA\_UD","MEGAHIT","MetaVelvet","MIRA","Ray\_Meta","SOAPdenovo2","SPAdes","SPAdes\_meta","SPAdes\_sc","SPAdes\_sc\_careful","Velvet","VICUNA","Geneious"]},
{"refnames":["KF302034","KF302033","KC821625","KF302036","KF302037","NC\_021794","NC\_001330","KC821629","NC\_021796","KF302035","NC\_021790","NC\_001422"],"coord\_y":[[null,null,1.0,null,1.003,1.001,1.009,1.005,1.0,1.0,1.004,null],[null,null,1.002,null,null,null,1.645,1.001,1.0,1.001,1.003,null],[1.0,1.0,1.0,1.0,1.0,1.0,1.0,1.0,1.0,1.362,1.001,1.004],[1.001,1.003,1.001,1.003,1.002,1.001,1.017,1.002,1.001,1.0,1.003,1.018],[1.001,1.004,1.002,1.004,1.003,1.002,1.011,1.003,1.002,1.0,1.007,1.0],[1.0,1.002,null,1.0,1.001,1.001,1.012,1.003,1.001,1.0,1.004,1.026],[1.006,1.004,1.084,1.004,1.004,1.002,null,1.009,1.033,1.177,1.007,3.539],[1.006,1.021,1.014,1.013,1.012,1.007,1.878,1.023,1.024,1.0,1.033,3.701],[0.989,0.979,1.002,0.987,0.986,0.988,null,1.111,0.979,1.001,1.165,null],[1.005,1.002,1.001,1.002,1.002,1.001,1.013,1.001,1.002,1.002,1.002,1.014],[1.0,1.002,1.0,1.001,1.001,1.001,1.008,1.001,1.001,1.002,1.002,1.008],[1.0,1.002,1.001,1.001,1.001,1.001,1.008,1.001,1.001,1.0,1.002,1.008],[1.0,1.002,1.0,1.001,1.001,1.001,1.008,1.001,1.001,1.002,1.002,1.008],[1.0,1.002,null,1.0,1.001,1.001,1.012,1.003,1.001,1.0,1.004,1.026],[1.002,1.008,1.005,1.007,1.013,1.006,1.071,1.001,1.002,1.027,1.001,1.058],[1.021,1.089,1.023,1.359,1.198,1.581,1.071,1.006,1.531,1.02,1.584,1.577]],"coord\_x":[[0.475,1.475,2.475,3.475,4.475,5.475,6.475,7.475,8.475,9.475,10.475,11.475],[0.5449999999999999,1.545,2.545,3.545,4.545,5.545,6.545,7.545,8.545,9.545,10.545,11.545],[0.615,1.615,2.615,3.615,4.615,5.615,6.615,7.615,8.615,9.615,10.615,11.615],[0.6849999999999999,1.685,2.685,3.685,4.685,5.685,6.685,7.685,8.685,9.685,10.685,11.685],[0.755,1.755,2.755,3.755,4.755,5.755,6.755,7.755,8.755,9.755,10.755,11.755],[0.825,1.825,2.825,3.825,4.825,5.825,6.825,7.825,8.825,9.825,10.825,11.825],[0.895,1.895,2.895,3.895,4.895,5.895,6.895,7.895,8.895,9.895,10.895,11.895],[0.965,1.965,2.965,3.965,4.965,5.965,6.965,7.965,8.965,9.965,10.965,11.965],[1.035,2.035,3.035,4.035,5.035,6.035,7.035,8.035,9.035,10.035,11.035,12.035],[1.105,2.105,3.105,4.105,5.105,6.105,7.105,8.105,9.105,10.105,11.105,12.105],[1.175,2.175,3.175,4.175,5.175,6.175,7.175,8.175,9.175,10.175,11.175,12.175],[1.245,2.245,3.245,4.245,5.245,6.245,7.245,8.245,9.245,10.245,11.245,12.245],[1.315,2.315,3.315,4.315,5.315,6.315,7.315,8.315,9.315,10.315,11.315,12.315],[1.385,2.385,3.385,4.385,5.385,6.385,7.385,8.385,9.385,10.385,11.385,12.385],[1.455,2.455,3.455,4.455,5.455,6.455,7.455,8.455,9.455,10.455,11.455,12.455],[1.525,2.525,3.525,4.525,5.525,6.525,7.525,8.525,9.525,10.525,11.525,12.525]],"filenames":["ABySS\_63","ABySS\_127","CLC","IDBA\_UD","MEGAHIT","MetaVelvet","MIRA","Ray\_Meta","SOAPdenovo2","SPAdes","SPAdes\_meta","SPAdes\_sc","SPAdes\_sc\_careful","Velvet","VICUNA","Geneious"]},
{"refnames":["KF302034","KC821625","NC\_021796","NC\_021794","KC821629","KF302037","NC\_021790","KF302035","KF302033","KF302036","NC\_001330","NC\_001422"],"coord\_y":[[null,76649,72567,71499,54004,44671,38543,35176,null,null,6141,null],[null,76666,72512,null,53956,null,38545,35176,null,null,6089,null],[129415,76567,72514,71428,53995,44551,39170,35176,36771,37711,6089,5365],[129514,76666,72634,71548,54115,44650,39290,35176,36870,37827,6188,5485],[129556,76807,72676,71588,54155,44692,38545,35176,36910,37869,6154,901],[129477,null,21285,65519,35740,44613,23050,35177,36833,null,2009,null],[128822,30919,72799,71593,48400,44737,39447,27926,36911,37863,null,593],[129072,77710,71664,71952,53943,45078,23054,35177,37544,38215,5716,5386],[null,73921,null,null,null,38805,null,null,null,33963,null,null],[128853,76666,70642,71526,54093,44628,39268,35180,36848,37805,6166,5463],[129470,76634,70598,71485,54042,44606,38521,35180,36826,37783,6134,5431],[129470,76653,70598,71485,54042,44606,38521,35176,36826,37783,6134,5431],[129470,76634,70598,71485,54042,44606,38521,35180,36826,37783,6134,5431],[129477,null,21285,65519,35740,44613,23050,35177,36833,null,2009,null],[129712,77070,72535,53925,54016,40569,39191,12556,37057,38007,6508,5699],[128822,71811,72680,53925,54016,29763,39192,35176,35127,29512,6508,3775]],"coord\_x":[[0.475,1.475,2.475,3.475,4.475,5.475,6.475,7.475,8.475,9.475,10.475,11.475],[0.5449999999999999,1.545,2.545,3.545,4.545,5.545,6.545,7.545,8.545,9.545,10.545,11.545],[0.615,1.615,2.615,3.615,4.615,5.615,6.615,7.615,8.615,9.615,10.615,11.615],[0.6849999999999999,1.685,2.685,3.685,4.685,5.685,6.685,7.685,8.685,9.685,10.685,11.685],[0.755,1.755,2.755,3.755,4.755,5.755,6.755,7.755,8.755,9.755,10.755,11.755],[0.825,1.825,2.825,3.825,4.825,5.825,6.825,7.825,8.825,9.825,10.825,11.825],[0.895,1.895,2.895,3.895,4.895,5.895,6.895,7.895,8.895,9.895,10.895,11.895],[0.965,1.965,2.965,3.965,4.965,5.965,6.965,7.965,8.965,9.965,10.965,11.965],[1.035,2.035,3.035,4.035,5.035,6.035,7.035,8.035,9.035,10.035,11.035,12.035],[1.105,2.105,3.105,4.105,5.105,6.105,7.105,8.105,9.105,10.105,11.105,12.105],[1.175,2.175,3.175,4.175,5.175,6.175,7.175,8.175,9.175,10.175,11.175,12.175],[1.245,2.245,3.245,4.245,5.245,6.245,7.245,8.245,9.245,10.245,11.245,12.245],[1.315,2.315,3.315,4.315,5.315,6.315,7.315,8.315,9.315,10.315,11.315,12.315],[1.385,2.385,3.385,4.385,5.385,6.385,7.385,8.385,9.385,10.385,11.385,12.385],[1.455,2.455,3.455,4.455,5.455,6.455,7.455,8.455,9.455,10.455,11.455,12.455],[1.525,2.525,3.525,4.525,5.525,6.525,7.525,8.525,9.525,10.525,11.525,12.525]],"filenames":["ABySS\_63","ABySS\_127","CLC","IDBA\_UD","MEGAHIT","MetaVelvet","MIRA","Ray\_Meta","SOAPdenovo2","SPAdes","SPAdes\_meta","SPAdes\_sc","SPAdes\_sc\_careful","Velvet","VICUNA","Geneious"]}]

{"refnames":["KC821625","KC821629","KF302033","KF302034","KF302035","KF302036","KF302037","NC\_001330","NC\_001422","NC\_021790","NC\_021794","NC\_021796"],"coord\_y":[[0.0,0.0,0.0,0.0,0.0,0.0,0,0,0,0,0,0,0.0,0.0,0.0,0,0,0,0.0,0.0,0.0,0.0,0.0,0.0,0,0,0,0.0,0.0,0.0,0.0,0.0,0.0,0.0,0.0,0.0],[0.0,0.0,0.0,0.0,0.0,0.0,0,0,0,0,0,0,0.0,0.0,0.0,0,0,0,0,0,0,2.0,0.0,0.0,0,0,0,0.0,0.0,0.0,0,0,0,0.0,0.0,0.0],[0.0,0.0,0.0,0.0,0.0,0.0,0.0,0.0,0.0,0.0,0.0,0.0,0.0,0.0,0.0,0.0,0.0,0.0,0.0,0.0,0.0,0.0,0.0,0.0,0.0,0.0,0.0,0.0,0.0,0.0,0.0,0.0,0.0,0.0,0.0,0.0],[0.0,0.0,0.0,0.0,0.0,0.0,0.0,0.0,0.0,0.0,0.0,0.0,0.0,0.0,0.0,0.0,0.0,0.0,0.0,0.0,0.0,0.0,0.0,0.0,0.0,0.0,0.0,0.0,0.0,0.0,0.0,0.0,0.0,0.0,0.0,0.0],[0.0,0.0,0.0,0.0,0.0,0.0,0.0,0.0,0.0,0.0,0.0,0.0,0.0,0.0,0.0,0.0,0.0,0.0,0.0,0.0,0.0,0.0,0.0,0.0,0.0,0.0,0.0,0.0,0.0,0.0,0.0,0.0,0.0,0.0,0.0,0.0],[0,0,0,0.0,0.0,0.0,0.0,0.0,0.0,0.0,0.0,0.0,0.0,0.0,0.0,0.0,0.0,0.0,0.0,0.0,0.0,0.0,0.0,0.0,0.0,0.0,0.0,0.0,0.0,0.0,0.0,0.0,0.0,0.0,0.0,0.0],[0.0,0.0,0.0,0.0,0.0,0.0,0.0,0.0,0.0,0.0,0.0,0.0,0.0,0.0,0.0,0.0,0.0,0.0,0.0,0.0,0.0,0,0,0,1.0,0.0,0.0,0.0,0.0,0.0,0.0,0.0,0.0,0.0,0.0,0.0],[0.0,0.0,0.0,1.0,0.0,0.0,0.0,0.0,0.0,0.0,0.0,0.0,0.0,0.0,0.0,0.0,0.0,0.0,0.0,0.0,0.0,1.0,0.0,0.0,3.0,0.0,0.0,1.0,0.0,0.0,0.0,0.0,0.0,0.0,0.0,0.0],[0.0,0.0,0.0,0.0,0.0,0.0,0.0,0.0,0.0,0.0,0.0,0.0,0.0,0.0,0.0,0.0,0.0,0.0,0.0,0.0,0.0,0,0,0,0,0,0,0.0,0.0,0.0,0.0,0.0,0.0,0.0,0.0,0.0],[0.0,0.0,0.0,0.0,0.0,0.0,0.0,0.0,0.0,0.0,0.0,0.0,0.0,0.0,0.0,0.0,0.0,0.0,0.0,0.0,0.0,0.0,0.0,0.0,0.0,0.0,0.0,0.0,0.0,0.0,0.0,0.0,0.0,0.0,0.0,0.0],[0.0,0.0,0.0,0.0,0.0,0.0,0.0,0.0,0.0,0.0,0.0,0.0,0.0,0.0,0.0,0.0,0.0,0.0,0.0,0.0,0.0,0.0,0.0,0.0,0.0,0.0,0.0,0.0,0.0,0.0,0.0,0.0,0.0,0.0,0.0,0.0],[0.0,0.0,0.0,0.0,0.0,0.0,0.0,0.0,0.0,0.0,0.0,0.0,0.0,0.0,0.0,0.0,0.0,0.0,0.0,0.0,0.0,0.0,0.0,0.0,0.0,0.0,0.0,0.0,0.0,0.0,0.0,0.0,0.0,0.0,0.0,0.0],[0.0,0.0,0.0,0.0,0.0,0.0,0.0,0.0,0.0,0.0,0.0,0.0,0.0,0.0,0.0,0.0,0.0,0.0,0.0,0.0,0.0,0.0,0.0,0.0,0.0,0.0,0.0,0.0,0.0,0.0,0.0,0.0,0.0,0.0,0.0,0.0],[0,0,0,0.0,0.0,0.0,0.0,0.0,0.0,0.0,0.0,0.0,0.0,0.0,0.0,0.0,0.0,0.0,0.0,0.0,0.0,0.0,0.0,0.0,0.0,0.0,0.0,0.0,0.0,0.0,0.0,0.0,0.0,0.0,0.0,0.0],[0.0,0.0,0.0,0.0,0.0,0.0,0.0,0.0,0.0,0.0,0.0,0.0,0.0,0.0,0.0,0.0,0.0,0.0,0.0,0.0,0.0,0.0,0.0,0.0,0.0,0.0,0.0,0.0,0.0,0.0,0.0,0.0,0.0,0.0,0.0,0.0],[0.0,0.0,0.0,0.0,0.0,0.0,1.0,0.0,0.0,1.0,0.0,0.0,0.0,0.0,0.0,1.0,0.0,0.0,1.0,0.0,0.0,0.0,0.0,0.0,2.0,0.0,0.0,1.0,0.0,0.0,2.0,0.0,0.0,1.0,0.0,0.0]],"coord\_x":[[1,1,1,2,2,2,3,3,3,4,4,4,5,5,5,6,6,6,7,7,7,8,8,8,9,9,9,10,10,10,11,11,11,12,12,12],[1,1,1,2,2,2,3,3,3,4,4,4,5,5,5,6,6,6,7,7,7,8,8,8,9,9,9,10,10,10,11,11,11,12,12,12],[1,1,1,2,2,2,3,3,3,4,4,4,5,5,5,6,6,6,7,7,7,8,8,8,9,9,9,10,10,10,11,11,11,12,12,12],[1,1,1,2,2,2,3,3,3,4,4,4,5,5,5,6,6,6,7,7,7,8,8,8,9,9,9,10,10,10,11,11,11,12,12,12],[1,1,1,2,2,2,3,3,3,4,4,4,5,5,5,6,6,6,7,7,7,8,8,8,9,9,9,10,10,10,11,11,11,12,12,12],[1,1,1,2,2,2,3,3,3,4,4,4,5,5,5,6,6,6,7,7,7,8,8,8,9,9,9,10,10,10,11,11,11,12,12,12],[1,1,1,2,2,2,3,3,3,4,4,4,5,5,5,6,6,6,7,7,7,8,8,8,9,9,9,10,10,10,11,11,11,12,12,12],[1,1,1,2,2,2,3,3,3,4,4,4,5,5,5,6,6,6,7,7,7,8,8,8,9,9,9,10,10,10,11,11,11,12,12,12],[1,1,1,2,2,2,3,3,3,4,4,4,5,5,5,6,6,6,7,7,7,8,8,8,9,9,9,10,10,10,11,11,11,12,12,12],[1,1,1,2,2,2,3,3,3,4,4,4,5,5,5,6,6,6,7,7,7,8,8,8,9,9,9,10,10,10,11,11,11,12,12,12],[1,1,1,2,2,2,3,3,3,4,4,4,5,5,5,6,6,6,7,7,7,8,8,8,9,9,9,10,10,10,11,11,11,12,12,12],[1,1,1,2,2,2,3,3,3,4,4,4,5,5,5,6,6,6,7,7,7,8,8,8,9,9,9,10,10,10,11,11,11,12,12,12],[1,1,1,2,2,2,3,3,3,4,4,4,5,5,5,6,6,6,7,7,7,8,8,8,9,9,9,10,10,10,11,11,11,12,12,12],[1,1,1,2,2,2,3,3,3,4,4,4,5,5,5,6,6,6,7,7,7,8,8,8,9,9,9,10,10,10,11,11,11,12,12,12],[1,1,1,2,2,2,3,3,3,4,4,4,5,5,5,6,6,6,7,7,7,8,8,8,9,9,9,10,10,10,11,11,11,12,12,12],[1,1,1,2,2,2,3,3,3,4,4,4,5,5,5,6,6,6,7,7,7,8,8,8,9,9,9,10,10,10,11,11,11,12,12,12]],"filenames":["ABySS\_63","ABySS\_127","CLC","IDBA\_UD","MEGAHIT","MetaVelvet","MIRA","Ray\_Meta","SOAPdenovo2","SPAdes","SPAdes\_meta","SPAdes\_sc","SPAdes\_sc\_careful","Velvet","VICUNA","Geneious"]}
