## Supplementary material for "Choice of assembly software has a critical impact on virome characterisation"

### QUAST

**Quality Assessment Tool for Genome Assemblies**
by CAB

Loading...

Aligned to
""

Combined reference
Estimated reference size:
 bp
|
 bp
|
 references
|
 fragments
|
 % G+C
|
 chromosomes
  
 reads
|
 mapped
|
 properly paired
  
 genes
|
 operons

Unfortunately, JavaScript in your
browser is disabled or is not supported.
We need JavaScript to build report and plots.

Worst
Median
Best

Show heatmap

|  |
| --- |
| Contigs are ordered from largest (contig #1) to smallest.  Contigs are broken into nonoverlapping 100 bp windows. Plot shows numbers of windows for each GC percentage. |

{"assembliesWithNs":["Geneious"],"minContig":500,"report":[["Genome statistics",[{"values":["99.527","97.174","99.589","8.487","99.589","96.525","99.743","93.799","94.737","99.589","99.155"],"quality":"More is better","isMain":true,"metricName":"Genome fraction (%)"},{"values":["1.000","1.006","1.004","1.151","1.323","1.000","1.000","1.001","1.001","1.027","1.020"],"quality":"Less is better","isMain":true,"metricName":"Duplication ratio"},{"values":[30946,16305,31106,1517,38103,16092,31013,16048,16196,31552,31461],"quality":"More is better","isMain":true,"metricName":"Largest alignment"},{"values":[30946,30401,31106,2900,40923,30012,31013,29219,29511,31552,31461],"quality":"More is better","isMain":true,"metricName":"Total aligned length"},{"values":[30965,16305,31106,null,40990,16092,31020,16048,16196,31829,32871],"quality":"More is better","isMain":false,"metricName":"NG50"},{"values":[30965,5625,31106,null,40990,10831,31020,11640,11970,31829,32871],"quality":"More is better","isMain":false,"metricName":"NG75"},{"values":[30946,16305,31106,1123,38103,16092,31013,10553,5582,31552,31461],"quality":"More is better","isMain":false,"metricName":"NA50"},{"values":[30946,5625,31106,1123,38103,10831,31013,null,null,31552,31461],"quality":"More is better","isMain":false,"metricName":"NA75"},{"values":[30946,16305,31106,null,38103,16092,31013,16048,16196,31552,31461],"quality":"More is better","isMain":true,"metricName":"NGA50"},{"values":[30946,5625,31106,null,38103,10831,31013,10553,5027,31552,31461],"quality":"More is better","isMain":false,"metricName":"NGA75"},{"values":[1,1,1,null,1,1,1,1,1,1,1],"quality":"Less is better","isMain":false,"metricName":"LG50"},{"values":[1,3,1,null,1,2,1,2,2,1,1],"quality":"Less is better","isMain":false,"metricName":"LG75"},{"values":[1,1,1,2,1,1,1,2,2,1,1],"quality":"Less is better","isMain":false,"metricName":"LA50"},{"values":[1,3,1,2,1,2,1,null,null,1,1],"quality":"Less is better","isMain":false,"metricName":"LA75"},{"values":[1,1,1,null,1,1,1,1,1,1,1],"quality":"Less is better","isMain":true,"metricName":"LGA50"},{"values":[1,3,1,null,1,2,1,2,3,1,1],"quality":"Less is better","isMain":false,"metricName":"LGA75"}]],["Misassemblies",[{"values":[0,0,0,1,3,0,0,0,0,0,0],"quality":"Less is better","isMain":true,"metricName":"# misassemblies"},{"values":[0,0,0,0,2,0,0,0,0,0,0],"quality":"Less is better","isMain":false,"metricName":" # relocations"},{"values":[0,0,0,0,0,0,0,0,0,0,0],"quality":"Less is better","isMain":false,"metricName":" # translocations"},{"values":[0,0,0,1,1,0,0,0,0,0,0],"quality":"Less is better","isMain":false,"metricName":" # inversions"},{"values":[0,0,0,1,1,0,0,0,0,0,0],"quality":"Less is better","isMain":false,"metricName":"# misassembled contigs"},{"values":[0,0,0,1481,40990,0,0,0,0,0,0],"quality":"Less is better","isMain":true,"metricName":"Misassembled contigs length"},{"values":[1,1,1,1,1,1,0,1,1,2,2],"quality":"Less is better","isMain":false,"metricName":"# local misassemblies"},{"values":[0,0,0,0,0,0,0,0,0,0,0],"quality":"Less is better","isMain":false,"metricName":"# unaligned mis. contigs"}]],["Unaligned",[{"values":[0,0,0,0,0,0,0,0,0,0,0],"quality":"Less is better","isMain":false,"metricName":"# fully unaligned contigs"},{"values":[0,0,0,0,0,0,0,0,0,0,0],"quality":"Less is better","isMain":false,"metricName":"Fully unaligned length"},{"values":[0,0,0,0,0,0,0,2,2,0,1],"quality":"Less is better","isMain":false,"metricName":"# partially unaligned contigs"},{"values":[0,0,0,0,0,0,0,11110,11632,0,1410],"quality":"Less is better","isMain":false,"metricName":"Partially unaligned length"}]],["Mismatches",[{"values":[27,27,27,0,5,25,27,28,28,21,153],"quality":"Less is better","isMain":false,"metricName":"# mismatches"},{"values":[10,10,10,0,1,10,10,12,12,4,40],"quality":"Less is better","isMain":false,"metricName":"# indels"},{"values":[17,17,17,0,1,17,17,19,19,5,51],"quality":"Less is better","isMain":false,"metricName":"Indels length"},{"values":["87.21","89.32","87.16","0.00","16.14","83.26","87.02","95.97","95.02","67.79","496.06"],"quality":"Less is better","isMain":true,"metricName":"# mismatches per 100 kbp"},{"values":["32.30","33.08","32.28","0.00","3.23","33.31","32.23","41.13","40.72","12.91","129.69"],"quality":"Less is better","isMain":true,"metricName":"# indels per 100 kbp"},{"values":[10,10,10,0,1,10,10,12,12,4,40],"quality":"Less is better","isMain":false,"metricName":" # indels (<= 5 bp)"},{"values":[0,0,0,0,0,0,0,0,0,0,0],"quality":"Less is better","isMain":false,"metricName":" # indels (> 5 bp)"},{"values":[0,0,0,0,0,0,0,0,0,0,778],"quality":"Less is better","isMain":false,"metricName":"# N's"},{"values":["0.00","0.00","0.00","0.00","0.00","0.00","0.00","0.00","0.00","0.00","2366.83"],"quality":"Less is better","isMain":true,"metricName":"# N's per 100 kbp"}]],["Statistics without reference",[{"values":[1,5,1,2,1,3,1,4,5,1,1],"quality":"Equal","isMain":true,"metricName":"# contigs"},{"values":[1,5,1,2,1,3,1,4,5,1,1],"quality":"Equal","isMain":false,"metricName":"# contigs (>= 0 bp)"},{"values":[1,5,1,2,1,3,1,4,5,1,1],"quality":"Equal","isMain":false,"metricName":"# contigs (>= 1000 bp)"},{"values":[1,3,1,0,1,2,1,3,4,1,1],"quality":"Equal","isMain":false,"metricName":"# contigs (>= 5000 bp)"},{"values":[1,1,1,0,1,2,1,3,2,1,1],"quality":"Equal","isMain":false,"metricName":"# contigs (>= 10000 bp)"},{"values":[1,0,1,0,1,0,1,0,0,1,1],"quality":"Equal","isMain":false,"metricName":"# contigs (>= 25000 bp)"},{"values":[0,0,0,0,0,0,0,0,0,0,0],"quality":"Equal","isMain":false,"metricName":"# contigs (>= 50000 bp)"},{"values":[30965,16305,31106,1557,40990,16092,31020,16048,16196,31829,32871],"quality":"More is better","isMain":true,"metricName":"Largest contig"},{"values":[30965,30401,31106,3038,40990,30012,31020,40329,41143,31829,32871],"quality":"More is better","isMain":true,"metricName":"Total length"},{"values":[30965,30401,31106,3038,40990,30012,31020,40329,41143,31829,32871],"quality":"More is better","isMain":false,"metricName":"Total length (>= 0 bp)"},{"values":[30965,30401,31106,3038,40990,30012,31020,40329,41143,31829,32871],"quality":"More is better","isMain":true,"metricName":"Total length (>= 1000 bp)"},{"values":[30965,27558,31106,0,40990,26923,31020,39135,39861,31829,32871],"quality":"More is better","isMain":false,"metricName":"Total length (>= 5000 bp)"},{"values":[30965,16305,31106,0,40990,26923,31020,39135,28166,31829,32871],"quality":"More is better","isMain":true,"metricName":"Total length (>= 10000 bp)"},{"values":[30965,0,31106,0,40990,0,31020,0,0,31829,32871],"quality":"More is better","isMain":false,"metricName":"Total length (>= 25000 bp)"},{"values":[0,0,0,0,0,0,0,0,0,0,0],"quality":"More is better","isMain":true,"metricName":"Total length (>= 50000 bp)"},{"values":[30965,16305,31106,1557,40990,16092,31020,11640,11970,31829,32871],"quality":"More is better","isMain":false,"metricName":"N50"},{"values":[30965,5625,31106,1481,40990,10831,31020,11447,6113,31829,32871],"quality":"More is better","isMain":false,"metricName":"N75"},{"values":[1,1,1,1,1,1,1,2,2,1,1],"quality":"Less is better","isMain":false,"metricName":"L50"},{"values":[1,3,1,2,1,2,1,3,3,1,1],"quality":"Less is better","isMain":false,"metricName":"L75"},{"values":["36.18","36.21","36.19","36.21","35.80","36.20","36.19","35.57","35.57","36.10","35.73"],"quality":"Equal","isMain":false,"metricName":"GC (%)"}]],["Predicted genes",[]],["Similarity statistics",[{"values":[0,0,0,0,0,0,0,0,0,0,0],"quality":"Equal","isMain":false,"metricName":"# similar correct contigs"},{"values":[0,0,0,0,0,0,0,0,0,0,0],"quality":"Equal","isMain":false,"metricName":"# similar misassembled blocks"}]],["Reference statistics",[{"values":[31106,31106,31106,31106,31106,31106,31106,31106,31106,31106,31106],"quality":"Equal","isMain":false,"metricName":"Reference length"},{"values":[1,1,1,1,1,1,1,1,1,1,1],"quality":"Equal","isMain":false,"metricName":"Reference fragments"},{"values":["36.21","36.21","36.21","36.21","36.21","36.21","36.21","36.21","36.21","36.21","36.21"],"quality":"Equal","isMain":false,"metricName":"Reference GC (%)"}]]],"referenceName":"Q33\_phage","date":"05 July 2018, Thursday, 15:30:41","order":[0,1,2,3,4,5,6,7,8,9,10],"assembliesNames":["CLC","IDBA\_UD","MEGAHIT","MIRA","Ray\_Meta","SPAdes","SPAdes\_meta","SPAdes\_sc","SPAdes\_sc\_careful","VICUNA","Geneious"]}

{{ qualities }}

{{ mainMetrics }}

{"lists\_of\_lengths":[[30965],[16305,5628,5625,1581,1262],[31106],[1557,1481],[40990],[16092,10831,3089],[31020],[16048,11640,11447,1194],[16196,11970,6113,5582,1282],[31829],[32871]],"filenames":["CLC","IDBA\_UD","MEGAHIT","MIRA","Ray\_Meta","SPAdes","SPAdes\_meta","SPAdes\_sc","SPAdes\_sc\_careful","VICUNA","Geneious"]}

{"assemblies\_lengths":[30965,30401,31106,3038,40990,30012,31020,40329,41143,31829,32871],"filenames":["CLC","IDBA\_UD","MEGAHIT","MIRA","Ray\_Meta","SPAdes","SPAdes\_meta","SPAdes\_sc","SPAdes\_sc\_careful","VICUNA","Geneious"]}

{"reflen":[31106]}

{"tickX":1}

{"coord\_y":[[30965,30965,30965,0.0],[16305,16305,16305,5628,5628,5625,5625,1581,1581,1262,1262,0.0],[31106,31106,31106,0.0],[1557,1557,1557,1481,1481,0.0],[40990,40990,40990,0.0],[16092,16092,16092,10831,10831,3089,3089,0.0],[31020,31020,31020,0.0],[16048,16048,16048,11640,11640,11447,11447,1194,1194,0.0],[16196,16196,16196,11970,11970,6113,6113,5582,5582,1282,1282,0.0],[31829,31829,31829,0.0],[32871,32871,32871,0.0]],"coord\_x":[[0.0,1e-10,100.0,100.0000000001],[0.0,1e-10,53.633104174204796,53.6331041743048,72.14565310351634,72.14565310361634,90.64833393638366,90.64833393648367,95.84882076247492,95.84882076257492,100.0,100.0000000001],[0.0,1e-10,100.0,100.0000000001],[0.0,1e-10,51.25082290980909,51.25082290990909,100.0,100.0000000001],[0.0,1e-10,100.0,100.0000000001],[0.0,1e-10,53.61855257896841,53.618552579068414,89.70745035319206,89.70745035329206,100.0,100.0000000001],[0.0,1e-10,100.0,100.0000000001],[0.0,1e-10,39.792705001363785,39.79270500146379,68.65531007463612,68.65531007473612,97.03935133526743,97.03935133536743,100.0,100.0000000001],[0.0,1e-10,39.36514109326009,39.365141093360094,68.45879007364557,68.45879007374558,83.31672459470627,83.31672459480627,96.8840385970882,96.88403859718821,100.0,100.0000000001],[0.0,1e-10,100.0,100.0000000001],[0.0,1e-10,100.0,100.0000000001]],"filenames":["CLC","IDBA\_UD","MEGAHIT","MIRA","Ray\_Meta","SPAdes","SPAdes\_meta","SPAdes\_sc","SPAdes\_sc\_careful","VICUNA","Geneious"]}

{"coord\_y":[[30965,30965,30965,0.0],[16305,16305,16305,5628,5628,5625,5625,1581,1581,1262,1262,0.0],[31106,31106,31106,0.0],[1557,1557,1557,1481,1481,0.0],[40990,40990,40990,0.0],[16092,16092,16092,10831,10831,3089,3089,0.0],[31020,31020,31020,0.0],[16048,16048,16048,11640,11640,11447,11447,1194,1194,0.0],[16196,16196,16196,11970,11970,6113,6113,5582,5582,1282,1282,0.0],[31829,31829,31829,0.0],[32871,32871,32871,0.0]],"coord\_x":[[0.0,1e-10,99.54671124541889,99.54671124551889],[0.0,1e-10,52.41754002443258,52.417540024532585,70.51051244132965,70.51051244142965,88.59384041663988,88.59384041673988,93.6764611329004,93.6764611330004,97.73355622709445,97.73355622719446],[0.0,1e-10,100.0,100.0000000001],[0.0,1e-10,5.005465183565872,5.005465183665872,9.766604513598663,9.766604513698663],[0.0,1e-10,131.77522021474957,131.77522021484955],[0.0,1e-10,51.732784671767504,51.732784671867506,86.55243361409374,86.55243361419375,96.48299363466855,96.48299363476855],[0.0,1e-10,99.72352600784414,99.72352600794414],[0.0,1e-10,51.5913328618273,51.5913328619273,89.01176621873593,89.01176621883593,125.81174050022504,125.81174050032504,129.65022825178423,129.65022825188422],[0.0,1e-10,52.06712531344435,52.06712531354435,90.54844724490452,90.54844724500452,110.2006043850061,110.2006043851061,128.1456953642384,128.1456953643384,132.267086735678,132.26708673577798],[0.0,1e-10,102.32431042242654,102.32431042252654],[0.0,1e-10,105.67414646691957,105.67414646701957]],"filenames":["CLC","IDBA\_UD","MEGAHIT","MIRA","Ray\_Meta","SPAdes","SPAdes\_meta","SPAdes\_sc","SPAdes\_sc\_careful","VICUNA","Geneious"]}

{"coord\_y":[[30946,30946,30946,0.0],[16305,16305,16305,5628,5628,5625,5625,1581,1581,1262,1262,0.0],[31106,31106,31106,0.0],[1517,1517,1517,1123,1123,260,260,0.0],[38103,38103,38103,1089,1089,947,947,784,784,0.0],[16092,16092,16092,10831,10831,3089,3089,0.0],[31013,31013,31013,0.0],[16048,16048,16048,10553,10553,1424,1424,1194,1194,0.0],[16196,16196,16196,5582,5582,5027,5027,1424,1424,1282,1282,0.0],[31552,31552,31552,0.0],[31461,31461,31461,0.0]],"coord\_x":[[0.0,1e-10,99.93864040045213,99.93864040055213],[0.0,1e-10,53.633104174204796,53.6331041743048,72.14565310351634,72.14565310361634,90.64833393638366,90.64833393648367,95.84882076247492,95.84882076257492,100.0,100.0000000001],[0.0,1e-10,100.0,100.0000000001],[0.0,1e-10,49.934167215273206,49.93416721537321,86.899275839368,86.899275839468,95.45753785385122,95.45753785395122],[0.0,1e-10,92.95681873627714,92.95681873637714,95.6135642839717,95.6135642840717,97.92388387411563,97.92388387421563,99.83654549890217,99.83654549900217],[0.0,1e-10,53.61855257896841,53.618552579068414,89.70745035319206,89.70745035329206,100.0,100.0000000001],[0.0,1e-10,99.97743391360413,99.97743391370413],[0.0,1e-10,39.792705001363785,39.79270500146379,65.95997917131592,65.95997917141592,69.49093704282278,69.49093704292278,72.45158570755535,72.45158570765535],[0.0,1e-10,39.36514109326009,39.365141093360094,52.93245509564203,52.93245509574203,65.15081544855747,65.15081544865747,68.6119145419634,68.6119145420634,71.7278759448752,71.7278759449752],[0.0,1e-10,99.12972446511043,99.12972446521043],[0.0,1e-10,95.71050470019166,95.71050470029166]],"filenames":["CLC","IDBA\_UD","MEGAHIT","MIRA","Ray\_Meta","SPAdes","SPAdes\_meta","SPAdes\_sc","SPAdes\_sc\_careful","VICUNA","Geneious"]}

{"coord\_y":[[30946,30946,30946,0.0],[16305,16305,16305,5628,5628,5625,5625,1581,1581,1262,1262,0.0],[31106,31106,31106,0.0],[1517,1517,1517,1123,1123,260,260,0.0],[38103,38103,38103,1089,1089,947,947,784,784,0.0],[16092,16092,16092,10831,10831,3089,3089,0.0],[31013,31013,31013,0.0],[16048,16048,16048,10553,10553,1424,1424,1194,1194,0.0],[16196,16196,16196,5582,5582,5027,5027,1424,1424,1282,1282,0.0],[31552,31552,31552,0.0],[31461,31461,31461,0.0]],"coord\_x":[[0.0,1e-10,99.48562978203562,99.48562978213562],[0.0,1e-10,52.41754002443258,52.417540024532585,70.51051244132965,70.51051244142965,88.59384041663988,88.59384041673988,93.6764611329004,93.6764611330004,97.73355622709445,97.73355622719446],[0.0,1e-10,100.0,100.0000000001],[0.0,1e-10,4.876872629074777,4.876872629174777,8.487108596412268,8.487108596512268,9.322960200604385,9.322960200704385],[0.0,1e-10,122.49405259435478,122.49405259445479,125.99498489037485,125.99498489047485,129.0394136179515,129.0394136180515,131.55982768597698,131.55982768607697],[0.0,1e-10,51.732784671767504,51.732784671867506,86.55243361409374,86.55243361419375,96.48299363466855,96.48299363476855],[0.0,1e-10,99.7010223108082,99.7010223109082],[0.0,1e-10,51.5913328618273,51.5913328619273,85.51726355044043,85.51726355054043,90.09515849032341,90.09515849042342,93.9336462418826,93.9336462419826],[0.0,1e-10,52.06712531344435,52.06712531354435,70.01221629267665,70.01221629277666,86.17308557834501,86.17308557844501,90.75098051822799,90.75098051832799,94.87237188966759,94.87237188976759],[0.0,1e-10,101.4338069825757,101.4338069826757],[0.0,1e-10,101.14125892110847,101.14125892120848]],"filenames":["CLC","IDBA\_UD","MEGAHIT","MIRA","Ray\_Meta","SPAdes","SPAdes\_meta","SPAdes\_sc","SPAdes\_sc\_careful","VICUNA","Geneious"]}

{{ genesInContigs }}

{{ operonsInContigs }}

[{{ num\_contigs }},
{{ Largest\_alignment }},
{{ Total\_aligned\_length }},
{{ num\_misassemblies }},
{{ Misassembled\_contigs\_length }},
{{ num\_mismatches\_per\_100\_kbp }},
{{ num\_indels\_per\_100\_kbp }},
{{ num\_N's\_per\_100\_kbp }},
{{ Genome\_fraction }},
{{ Duplication\_ratio }},
{{ NGA50 }}]

{{ allMisassemblies }}

{{ krona }}

{"list\_of\_GC\_distributions":[[[0.0,1.0,2.0,3.0,4.0,5.0,6.0,7.0,8.0,9.0,10.0,11.0,12.0,13.0,14.0,15.0,16.0,17.0,18.0,19.0,20.0,21.0,22.0,23.0,24.0,25.0,26.0,27.0,28.0,29.0,30.0,31.0,32.0,33.0,34.0,35.0,36.0,37.0,38.0,39.0,40.0,41.0,42.0,43.0,44.0,45.0,46.0,47.0,48.0,49.0,50.0,51.0,52.0,53.0,54.0,55.0,56.0,57.0,58.0,59.0,60.0,61.0,62.0,63.0,64.0,65.0,66.0,67.0,68.0,69.0,70.0,71.0,72.0,73.0,74.0,75.0,76.0,77.0,78.0,79.0,80.0,81.0,82.0,83.0,84.0,85.0,86.0,87.0,88.0,89.0,90.0,91.0,92.0,93.0,94.0,95.0,96.0,97.0,98.0,99.0,100.0],[0,0,0,0,0,0,0,0,0,0,0,0,0,0,0,0,0,1,0,1,1,0,1,1,1,3,1,4,6,8,17,9,15,19,31,22,26,18,21,28,22,9,9,11,6,3,4,4,3,1,1,2,0,1,0,0,0,0,0,0,0,0,0,0,0,0,0,0,0,0,0,0,0,0,0,0,0,0,0,0,0,0,0,0,0,0,0,0,0,0,0,0,0,0,0,0,0,0,0,0,0]],[[0.0,1.0,2.0,3.0,4.0,5.0,6.0,7.0,8.0,9.0,10.0,11.0,12.0,13.0,14.0,15.0,16.0,17.0,18.0,19.0,20.0,21.0,22.0,23.0,24.0,25.0,26.0,27.0,28.0,29.0,30.0,31.0,32.0,33.0,34.0,35.0,36.0,37.0,38.0,39.0,40.0,41.0,42.0,43.0,44.0,45.0,46.0,47.0,48.0,49.0,50.0,51.0,52.0,53.0,54.0,55.0,56.0,57.0,58.0,59.0,60.0,61.0,62.0,63.0,64.0,65.0,66.0,67.0,68.0,69.0,70.0,71.0,72.0,73.0,74.0,75.0,76.0,77.0,78.0,79.0,80.0,81.0,82.0,83.0,84.0,85.0,86.0,87.0,88.0,89.0,90.0,91.0,92.0,93.0,94.0,95.0,96.0,97.0,98.0,99.0,100.0],[0,0,0,0,0,0,0,0,0,0,0,0,0,0,0,1,0,0,0,0,1,0,1,2,1,3,2,7,2,11,6,8,21,25,29,23,18,26,21,16,17,11,14,10,10,6,4,2,1,1,1,0,0,3,0,0,0,0,0,0,0,0,0,0,0,0,0,0,0,0,0,0,0,0,0,0,0,0,0,0,0,0,0,0,0,0,0,0,0,0,0,0,0,0,0,0,0,0,0,0,0]],[[0.0,1.0,2.0,3.0,4.0,5.0,6.0,7.0,8.0,9.0,10.0,11.0,12.0,13.0,14.0,15.0,16.0,17.0,18.0,19.0,20.0,21.0,22.0,23.0,24.0,25.0,26.0,27.0,28.0,29.0,30.0,31.0,32.0,33.0,34.0,35.0,36.0,37.0,38.0,39.0,40.0,41.0,42.0,43.0,44.0,45.0,46.0,47.0,48.0,49.0,50.0,51.0,52.0,53.0,54.0,55.0,56.0,57.0,58.0,59.0,60.0,61.0,62.0,63.0,64.0,65.0,66.0,67.0,68.0,69.0,70.0,71.0,72.0,73.0,74.0,75.0,76.0,77.0,78.0,79.0,80.0,81.0,82.0,83.0,84.0,85.0,86.0,87.0,88.0,89.0,90.0,91.0,92.0,93.0,94.0,95.0,96.0,97.0,98.0,99.0,100.0],[0,0,0,0,0,0,0,0,0,0,0,0,0,0,0,0,1,0,1,0,1,0,1,3,1,1,2,3,6,8,15,14,19,12,27,27,31,17,20,19,13,12,15,14,9,7,5,1,1,2,1,1,0,1,0,0,0,0,0,0,0,0,0,0,0,0,0,0,0,0,0,0,0,0,0,0,0,0,0,0,0,0,0,0,0,0,0,0,0,0,0,0,0,0,0,0,0,0,0,0,0]],[[0.0,1.0,2.0,3.0,4.0,5.0,6.0,7.0,8.0,9.0,10.0,11.0,12.0,13.0,14.0,15.0,16.0,17.0,18.0,19.0,20.0,21.0,22.0,23.0,24.0,25.0,26.0,27.0,28.0,29.0,30.0,31.0,32.0,33.0,34.0,35.0,36.0,37.0,38.0,39.0,40.0,41.0,42.0,43.0,44.0,45.0,46.0,47.0,48.0,49.0,50.0,51.0,52.0,53.0,54.0,55.0,56.0,57.0,58.0,59.0,60.0,61.0,62.0,63.0,64.0,65.0,66.0,67.0,68.0,69.0,70.0,71.0,72.0,73.0,74.0,75.0,76.0,77.0,78.0,79.0,80.0,81.0,82.0,83.0,84.0,85.0,86.0,87.0,88.0,89.0,90.0,91.0,92.0,93.0,94.0,95.0,96.0,97.0,98.0,99.0,100.0],[0,0,0,0,0,0,0,0,0,0,0,0,0,0,0,0,0,0,0,0,0,0,0,0,0,0,0,0,1,0,0,4,1,4,2,3,1,4,2,3,3,0,1,0,1,0,1,0,0,0,0,0,0,0,0,0,0,0,0,0,0,0,0,0,0,0,0,0,0,0,0,0,0,0,0,0,0,0,0,0,0,0,0,0,0,0,0,0,0,0,0,0,0,0,0,0,0,0,0,0,0]],[[0.0,1.0,2.0,3.0,4.0,5.0,6.0,7.0,8.0,9.0,10.0,11.0,12.0,13.0,14.0,15.0,16.0,17.0,18.0,19.0,20.0,21.0,22.0,23.0,24.0,25.0,26.0,27.0,28.0,29.0,30.0,31.0,32.0,33.0,34.0,35.0,36.0,37.0,38.0,39.0,40.0,41.0,42.0,43.0,44.0,45.0,46.0,47.0,48.0,49.0,50.0,51.0,52.0,53.0,54.0,55.0,56.0,57.0,58.0,59.0,60.0,61.0,62.0,63.0,64.0,65.0,66.0,67.0,68.0,69.0,70.0,71.0,72.0,73.0,74.0,75.0,76.0,77.0,78.0,79.0,80.0,81.0,82.0,83.0,84.0,85.0,86.0,87.0,88.0,89.0,90.0,91.0,92.0,93.0,94.0,95.0,96.0,97.0,98.0,99.0,100.0],[0,0,0,0,0,0,0,0,0,0,0,0,0,0,0,1,0,0,1,1,0,0,1,3,3,7,5,4,14,7,10,18,36,23,43,25,28,32,24,25,21,13,16,13,10,12,4,1,4,0,1,0,2,1,1,0,0,0,0,0,0,0,0,0,0,0,0,0,0,0,0,0,0,0,0,0,0,0,0,0,0,0,0,0,0,0,0,0,0,0,0,0,0,0,0,0,0,0,0,0,0]],[[0.0,1.0,2.0,3.0,4.0,5.0,6.0,7.0,8.0,9.0,10.0,11.0,12.0,13.0,14.0,15.0,16.0,17.0,18.0,19.0,20.0,21.0,22.0,23.0,24.0,25.0,26.0,27.0,28.0,29.0,30.0,31.0,32.0,33.0,34.0,35.0,36.0,37.0,38.0,39.0,40.0,41.0,42.0,43.0,44.0,45.0,46.0,47.0,48.0,49.0,50.0,51.0,52.0,53.0,54.0,55.0,56.0,57.0,58.0,59.0,60.0,61.0,62.0,63.0,64.0,65.0,66.0,67.0,68.0,69.0,70.0,71.0,72.0,73.0,74.0,75.0,76.0,77.0,78.0,79.0,80.0,81.0,82.0,83.0,84.0,85.0,86.0,87.0,88.0,89.0,90.0,91.0,92.0,93.0,94.0,95.0,96.0,97.0,98.0,99.0,100.0],[0,0,0,0,0,0,0,0,0,0,0,0,0,0,0,0,0,0,0,0,1,1,0,1,1,2,3,0,8,9,15,13,21,22,18,21,25,17,25,19,13,18,11,10,15,0,3,1,3,0,3,0,0,0,1,0,0,0,0,0,0,0,0,0,0,0,0,0,0,0,0,0,0,0,0,0,0,0,0,0,0,0,0,0,0,0,0,0,0,0,0,0,0,0,0,0,0,0,0,0,0]],[[0.0,1.0,2.0,3.0,4.0,5.0,6.0,7.0,8.0,9.0,10.0,11.0,12.0,13.0,14.0,15.0,16.0,17.0,18.0,19.0,20.0,21.0,22.0,23.0,24.0,25.0,26.0,27.0,28.0,29.0,30.0,31.0,32.0,33.0,34.0,35.0,36.0,37.0,38.0,39.0,40.0,41.0,42.0,43.0,44.0,45.0,46.0,47.0,48.0,49.0,50.0,51.0,52.0,53.0,54.0,55.0,56.0,57.0,58.0,59.0,60.0,61.0,62.0,63.0,64.0,65.0,66.0,67.0,68.0,69.0,70.0,71.0,72.0,73.0,74.0,75.0,76.0,77.0,78.0,79.0,80.0,81.0,82.0,83.0,84.0,85.0,86.0,87.0,88.0,89.0,90.0,91.0,92.0,93.0,94.0,95.0,96.0,97.0,98.0,99.0,100.0],[0,0,0,0,0,0,0,0,0,0,0,0,0,0,0,0,0,1,1,1,0,1,0,0,2,2,2,2,7,12,14,17,20,16,17,24,22,23,17,21,21,17,14,11,8,3,4,3,3,2,0,1,1,0,0,0,0,0,0,0,0,0,0,0,0,0,0,0,0,0,0,0,0,0,0,0,0,0,0,0,0,0,0,0,0,0,0,0,0,0,0,0,0,0,0,0,0,0,0,0,0]],[[0.0,1.0,2.0,3.0,4.0,5.0,6.0,7.0,8.0,9.0,10.0,11.0,12.0,13.0,14.0,15.0,16.0,17.0,18.0,19.0,20.0,21.0,22.0,23.0,24.0,25.0,26.0,27.0,28.0,29.0,30.0,31.0,32.0,33.0,34.0,35.0,36.0,37.0,38.0,39.0,40.0,41.0,42.0,43.0,44.0,45.0,46.0,47.0,48.0,49.0,50.0,51.0,52.0,53.0,54.0,55.0,56.0,57.0,58.0,59.0,60.0,61.0,62.0,63.0,64.0,65.0,66.0,67.0,68.0,69.0,70.0,71.0,72.0,73.0,74.0,75.0,76.0,77.0,78.0,79.0,80.0,81.0,82.0,83.0,84.0,85.0,86.0,87.0,88.0,89.0,90.0,91.0,92.0,93.0,94.0,95.0,96.0,97.0,98.0,99.0,100.0],[0,0,0,0,0,0,0,0,0,0,0,0,0,0,0,0,0,0,0,1,4,1,2,1,1,7,6,3,6,14,19,20,30,20,32,34,31,22,29,25,24,23,13,6,6,7,5,5,2,1,0,0,0,1,1,0,0,0,0,0,0,0,0,0,0,0,0,0,0,0,0,0,0,0,0,0,0,0,0,0,0,0,0,0,0,0,0,0,0,0,0,0,0,0,0,0,0,0,0,0,0]],[[0.0,1.0,2.0,3.0,4.0,5.0,6.0,7.0,8.0,9.0,10.0,11.0,12.0,13.0,14.0,15.0,16.0,17.0,18.0,19.0,20.0,21.0,22.0,23.0,24.0,25.0,26.0,27.0,28.0,29.0,30.0,31.0,32.0,33.0,34.0,35.0,36.0,37.0,38.0,39.0,40.0,41.0,42.0,43.0,44.0,45.0,46.0,47.0,48.0,49.0,50.0,51.0,52.0,53.0,54.0,55.0,56.0,57.0,58.0,59.0,60.0,61.0,62.0,63.0,64.0,65.0,66.0,67.0,68.0,69.0,70.0,71.0,72.0,73.0,74.0,75.0,76.0,77.0,78.0,79.0,80.0,81.0,82.0,83.0,84.0,85.0,86.0,87.0,88.0,89.0,90.0,91.0,92.0,93.0,94.0,95.0,96.0,97.0,98.0,99.0,100.0],[0,0,0,0,0,0,0,0,0,0,0,0,0,0,0,0,0,1,0,0,2,2,4,2,3,1,3,6,13,7,24,16,23,28,39,33,27,32,29,27,21,18,10,14,6,9,4,2,2,0,1,0,2,1,0,0,0,0,0,0,0,0,0,0,0,0,0,0,0,0,0,0,0,0,0,0,0,0,0,0,0,0,0,0,0,0,0,0,0,0,0,0,0,0,0,0,0,0,0,0,0]],[[0.0,1.0,2.0,3.0,4.0,5.0,6.0,7.0,8.0,9.0,10.0,11.0,12.0,13.0,14.0,15.0,16.0,17.0,18.0,19.0,20.0,21.0,22.0,23.0,24.0,25.0,26.0,27.0,28.0,29.0,30.0,31.0,32.0,33.0,34.0,35.0,36.0,37.0,38.0,39.0,40.0,41.0,42.0,43.0,44.0,45.0,46.0,47.0,48.0,49.0,50.0,51.0,52.0,53.0,54.0,55.0,56.0,57.0,58.0,59.0,60.0,61.0,62.0,63.0,64.0,65.0,66.0,67.0,68.0,69.0,70.0,71.0,72.0,73.0,74.0,75.0,76.0,77.0,78.0,79.0,80.0,81.0,82.0,83.0,84.0,85.0,86.0,87.0,88.0,89.0,90.0,91.0,92.0,93.0,94.0,95.0,96.0,97.0,98.0,99.0,100.0],[0,0,0,0,0,0,0,0,0,0,0,0,0,0,0,0,0,1,0,1,2,0,0,0,0,3,2,3,8,5,13,12,23,22,35,23,26,17,21,26,20,10,8,12,6,5,3,3,2,1,1,3,1,0,0,0,0,0,0,0,0,0,0,0,0,0,0,0,0,0,0,0,0,0,0,0,0,0,0,0,0,0,0,0,0,0,0,0,0,0,0,0,0,0,0,0,0,0,0,0,0]],[[0.0,1.0,2.0,3.0,4.0,5.0,6.0,7.0,8.0,9.0,10.0,11.0,12.0,13.0,14.0,15.0,16.0,17.0,18.0,19.0,20.0,21.0,22.0,23.0,24.0,25.0,26.0,27.0,28.0,29.0,30.0,31.0,32.0,33.0,34.0,35.0,36.0,37.0,38.0,39.0,40.0,41.0,42.0,43.0,44.0,45.0,46.0,47.0,48.0,49.0,50.0,51.0,52.0,53.0,54.0,55.0,56.0,57.0,58.0,59.0,60.0,61.0,62.0,63.0,64.0,65.0,66.0,67.0,68.0,69.0,70.0,71.0,72.0,73.0,74.0,75.0,76.0,77.0,78.0,79.0,80.0,81.0,82.0,83.0,84.0,85.0,86.0,87.0,88.0,89.0,90.0,91.0,92.0,93.0,94.0,95.0,96.0,97.0,98.0,99.0,100.0],[0,0,0,0,0,0,0,0,0,0,0,0,0,0,0,0,0,2,0,1,1,0,1,2,1,3,4,5,7,15,9,13,18,27,34,24,26,16,24,21,15,9,14,11,11,6,1,1,0,2,1,2,0,1,0,0,0,0,0,0,0,0,0,0,0,0,0,0,0,0,0,0,0,0,0,0,0,0,0,0,0,0,0,0,0,0,0,0,0,0,0,0,0,0,0,0,0,0,0,0,0]],[[0.0,1.0,2.0,3.0,4.0,5.0,6.0,7.0,8.0,9.0,10.0,11.0,12.0,13.0,14.0,15.0,16.0,17.0,18.0,19.0,20.0,21.0,22.0,23.0,24.0,25.0,26.0,27.0,28.0,29.0,30.0,31.0,32.0,33.0,34.0,35.0,36.0,37.0,38.0,39.0,40.0,41.0,42.0,43.0,44.0,45.0,46.0,47.0,48.0,49.0,50.0,51.0,52.0,53.0,54.0,55.0,56.0,57.0,58.0,59.0,60.0,61.0,62.0,63.0,64.0,65.0,66.0,67.0,68.0,69.0,70.0,71.0,72.0,73.0,74.0,75.0,76.0,77.0,78.0,79.0,80.0,81.0,82.0,83.0,84.0,85.0,86.0,87.0,88.0,89.0,90.0,91.0,92.0,93.0,94.0,95.0,96.0,97.0,98.0,99.0,100.0],[0,0,0,0,0,0,0,0,0,0,0,0,0,0,0,0,0,0,0,1,1,1,0,0,1,3,5,2,4,8,10,21,14,19,31,22,25,20,20,18,21,17,10,11,6,8,6,4,0,0,0,0,0,1,1,0,0,0,0,0,0,0,0,0,0,0,0,0,0,0,0,0,0,0,0,0,0,0,0,0,0,0,0,0,0,0,0,0,0,0,0,0,0,0,0,0,0,0,0,0,0]]],"reference\_index":11,"lists\_of\_gc\_info":null,"filenames":["CLC","IDBA\_UD","MEGAHIT","MIRA","Ray\_Meta","SPAdes","SPAdes\_meta","SPAdes\_sc","SPAdes\_sc\_careful","VICUNA","Geneious"]}
